# Elucidation of Japanese pepper (*Zanthoxylum piperitum* De Candolle) domestication using RAD-Seq

**DOI:** 10.1101/2020.12.29.424752

**Authors:** Maddumage Dona Ginushika Priyadarshani Premarathne, Nami Fukutome, Kazuaki Yamasaki, Fumiyo Hayakawa, Atsushi J. Nagano, Hisataka Mizuno, Nobuo Ibaragi, Yukio Nagano

**Author notes:** Correspondence to Y. N. These authors contributed equally to this work (Nami Fukutome, Yukio Nagano).

## Abstract

Japanese pepper, *Zanthoxylum piperitum*, is native to Japan and has four well-known lineages (Asakura, Takahara, Budou, and Arima), which are named after their production area or morphology. Restriction-site associated DNA sequencing (RAD-Seq) was used to analyse 93 accessions from various areas, including these four lineages. Single nucleotide variant analysis was used to classify the plants into eight groups: the Asakura and Arima lineages each had two groups, the Takahara and Budou lineages each had one group, and two additional groups were present. In one Asakura group and two Arima groups, the plants were present in agricultural fields and mountains, thus representing the early stage of domestication of the Japanese pepper. The second Asakura lineage group was closely related to plants present in various areas, and this represents the second stage of domestication of this plant because, after early domestication, genetically related lineages with desirable traits spread to the periphery. These results demonstrate that domestication of Japanese pepper is ongoing. In addition, this study shows that spineless plants are polyphyletic, despite the spineless lineage being considered a subspecies of Japanese pepper.

## Introduction

*Zanthoxylum* species belongs to the family Rutaceae, commonly known as the citrus family. Many *Zanthoxylum* species are distributed worldwide and are used for culinary and medicinal purposes. For example, the Āyurvedic traditional medicine of India and Sri Lanka uses *Z. tetraspermum* and *Z. rhetsa* (“*Katu Keena*”)^1^. Sichuan pepper is an essential ingredient in the cuisines of the Sichuan province of China and contains several species of *Zanthoxylum*, including *Z. simulans*. *Z. simulans* is used in traditional Chinese medicine for its stomachic, analgesic, and anthelmintic properties^2^.

*Zanthoxylum piperitum* is native to Japan (except for Ryukyu) and southern Korea. In Japan, it is mainly used for culinary purposes. In English, it is known as Japanese pepper; in Japanese, it is called sanshō. This plant is considered to be highly medicinal in various uses^3–10^. Although *Zanthoxylum piperitum* is widely consumed and distributed throughout Japan, lineages have been formed based on the production area and morphology of this plant. The Asakura, Arima, and Takahara lineages are classified based on the place of production, while the Budou lineage is classified based on its morphology.

The Asakura lineage is believed to have originated in the Asakura district of Yabu City in central Hyōgo Prefecture. A typical feature of this lineage is the absence of spines. For this reason, the spineless plant is called the Asakura lineage in Japan. This spineless plant has been identified as a subspecies of *Z. piperitum* and named *Z. piperitum* (L.) DC forma *inerme* (Makino) Makino^11^. However, there is no evidence that all spineless plants are either from Asakura or are monophyletic. In Yabu city, *Zanthoxylum piperitum* grow as wild plants in mountainous areas and as cultivated plants in agricultural fields and gardens. The Arima lineage, also known as the Rokko lineage, is another Japanese pepper lineage found in Kobe City in southern Hyōgo Prefecture. The flowers of these plants are typically used as ingredients in Arima cuisine.

The Takahara lineage is grown in Takayama City in northern Gifu Prefecture. These plants are grown at high altitudes 800 m above sea level, in contrast to the other lineages, which are cultivated on the plains. These plants bloom in mid-May, whereas other lineages bloom in late April. The wild Takahara lineage is difficult to find in the mountains of this region.

The Budou lineage (Budou means “grape” in Japanese) is categorised based on its morphology. This lineage is cultivated mainly in Kainan City, Wakayama Prefecture. It bears clusters of fruits that resemble grapes. According to oral tradition, it appeared as a chance seedling in a garden about 200 years ago and has been cultivated ever since. Therefore, the wild lineage is probably absent.

Many studies have investigated the genetic relationships among *Zanthoxylum* species^12–23^. There are no reports that compare the intraspecific diversity of *Zanthoxylum piperitum* in detail. However, three important studies have examined the DNA sequences of *Zanthoxylum piperitum* growing in Korea. One is the complete sequencing of the chloroplast genome^12^, and the others involved the development of markers to identify *Zanthoxylum piperitum*^13, 14^. Many other studies have used the genetic information for *Zanthoxylum piperitum* as a reference for comparison^15–20, 22, 23^. There are two important studies on the intraspecific genetic diversity of *Zanthoxylum* species, which focus on: (1) the relationship and diversity between wild and cultivated plants^22^ and (2) the genetic structure of cultivated plants^23^. A previous study^22^ compared wild plants to cultivated plants using DNA sequence data and detected genetic similarities between cultivated and wild *Z. armatum*. Although this finding may be relevant to the domestication of *Z. armatum*, the study only used a small number of markers and was unable to examine the species domestication details.

Studies on intraspecific genetic diversity of *Zanthoxylum* species have been based on the analysis of a small number of markers, such as sequence-related amplified polymorphism markers^22^ and simple sequence repeat (SSR) markers^23^. Compared to genetic studies using a small number of markers, high-throughput sequencing can generate many single nucleotide variants (SNVs) that are suitable for analysing intraspecific differences^24, 25^. Restriction site-associated DNA sequencing (RAD-Seq)^26^ is a high-throughput sequencing method. Initially, RAD-Seq cleaves individual genomes using a selected site-specific restriction enzyme. Then, DNA fragments adjacent to the restriction site are sequenced using a high-throughput sequencer. Double-digest RAD-Seq (ddRAD-Seq)^27^ is a variation of RAD-Seq that uses two restriction enzymes to cleave the genome. These methods allow for reduced representation of individual genomes. Moreover, these methods do not require prior knowledge of the genome sequence^27^. Our group has previously used the original RAD-Seq and ddRAD-Seq methods to analyse citrus^28–30^, loquat^31^, and firefly^32^.

The four lineages are well known, and *Zanthoxylum piperitum* is cultivated all over Japan, except for Ryukyu. Some lineages are present in both the mountains and agricultural fields. Therefore, the domestication of Japanese pepper may be an ongoing process. In the present study, ddRAD-Seq was used to elucidate the intraspecific genetic diversity and occurrence of domestication in Japanese pepper.

## Results

### Variant detection by *de novo* mapping of RAD-Seq data

Double digest RAD-Seq generated more than 11.9 gigabases of data for a total of 235.2 million raw single-end 51-bp reads. Quality-based filtering resulted in an average of 2.5 million reads (maximum of 4.3 million and a minimum of 0.4 million) among 93 samples (Supplementary Table 1). The Stacks program built loci *de novo* with an average coverage depth of 27.32 times (Supplementary Table 2). We performed the following analysis using data from 4,334 variant sites.

### Correlation of genetic similarity with geographic locations

Based on the principal component analysis (PCA) (Figure 1) and geographical information (Figure 2, Table 1), 93 samples were classified into seven groups. PC1 divided group A from the other groups (Figure 1a); PC2 divided groups D and E from the other groups (Figure 1a); PC3 (Figure 1b) identified groups C and F and confirmed the presence of group E; PC4 identified group B (Figure 1c) and confirmed the presence of groups C and F; PC5 reconfirmed the presence of groups B and F (Figure 1d); and PC6 identified group G (Figure 1e). Thus, probably because the genetic relationships among plants were complex, it was necessary to use PC1 through PC6 for classification.

**Figure 1.**
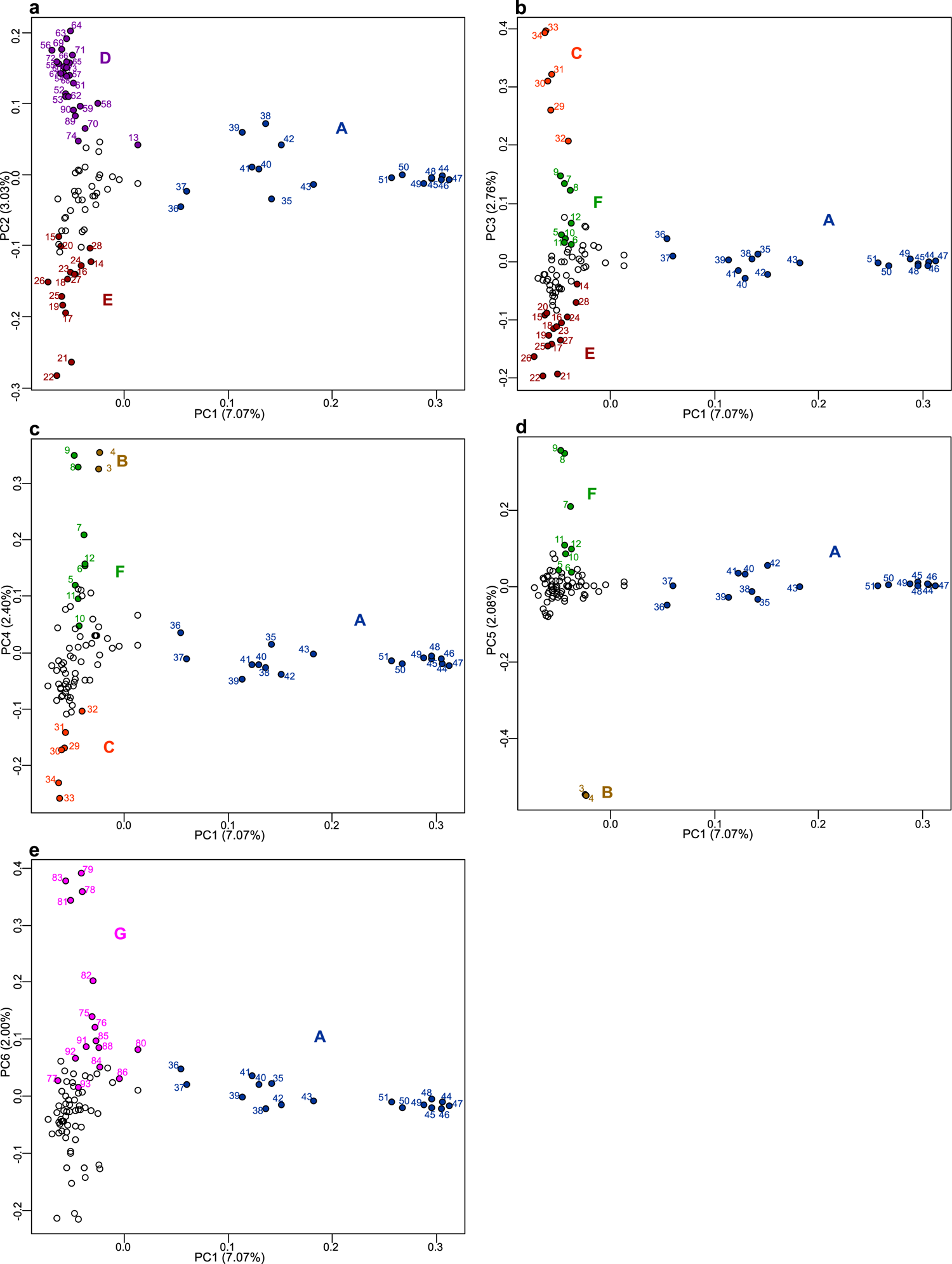
Principal component (PC) analysis of *Zanthoxylum* accessions used in the study. The six major component data are shown by the five two-dimensional data sets: (**a**) first and second PCA axes, (**b**) first and third PCA axes, (**c**) first and fourth PCA axes, (**d**) first and fifth PCA axes, and (**e**) first and sixth PCA axes. Group A of the Asakura lineage is shown in blue; group B of the Budou lineage is shown in brown; group C of the Arima lineage is shown in orange; group D of the Asakura lineage is shown in violet; group E of the Takahara lineage is shown in maroon; group F of the Arima lineage is shown in green; group G is shown in violet. The letters indicating the sample number are also shown in the same colour. The contribution rate of each principal component is shown in parentheses. Figure was generated using R software (version 3.6.2)^44^.

**Figure 2.**
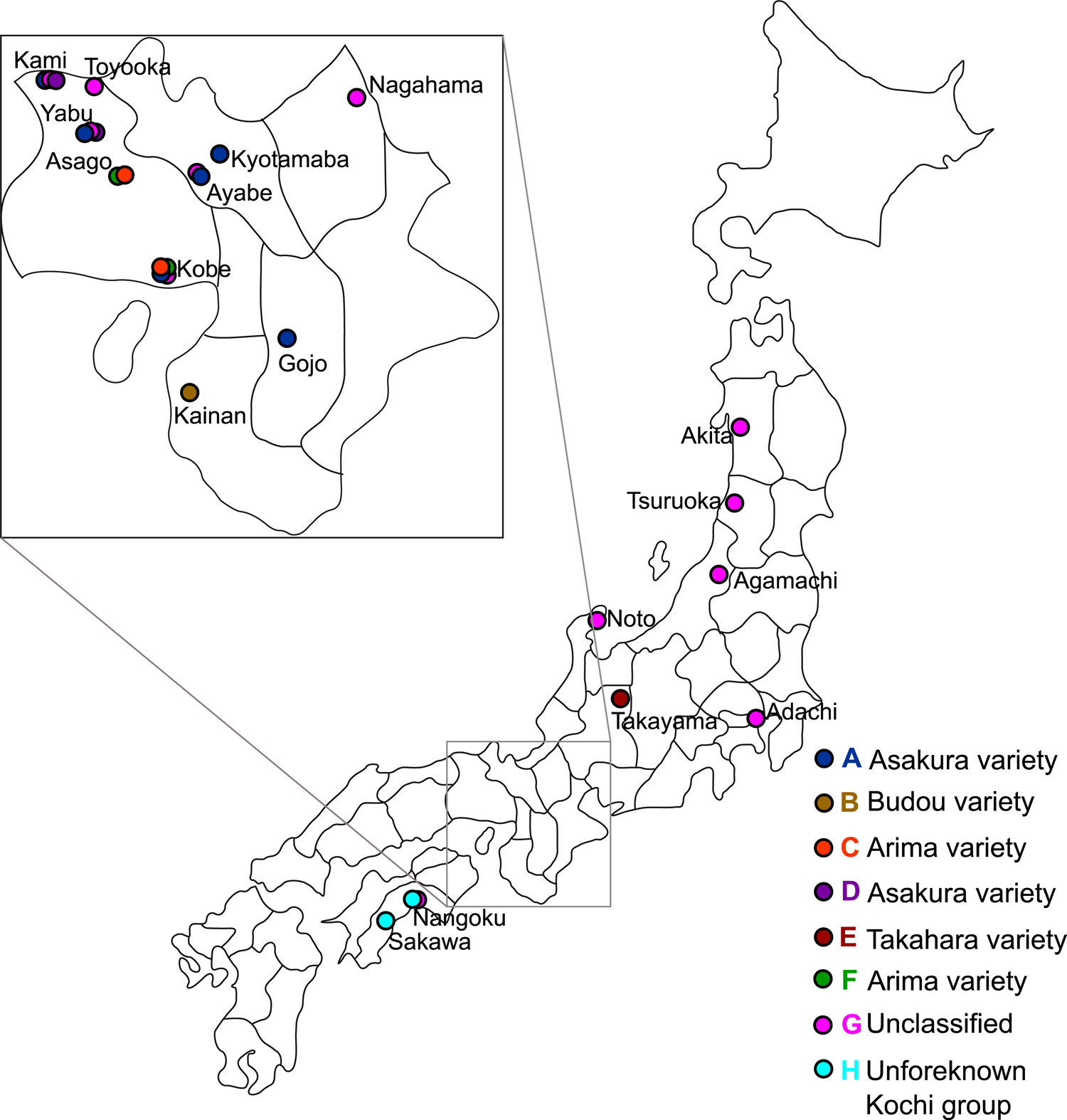
Sampling points and locations of the lineages. The colour scheme is the same as in Figure 1.

**Table 1.** Information on *Zanthoxylum piperitum* accessions used in the study.

| Plant Number | Prefecture | City/Town | Spiny/Spineless | Information | Group | Lineage |
| --- | --- | --- | --- | --- | --- | --- |
| 1 | Kochi | Sakawa | Spiny | Agricultural field, cultivated | H | Unforeknown<br>Kochi group |
| 2 | Kochi | Nangoku | Spiny | Mountain, wild tree | H | Unforeknown<br>Kochi group |
| 3 | Wakayama | Kainan | Spiny | Agricultural field, cultivated, Nagamine agricultural cooperative | B | Budou |
| 4 | Wakayama | Kainan | Spiny | Agricultural field, cultivated, Nagamine agricultural cooperative | B | Budou |
| 5 | Hyogo | Kobe | Spiny | Garden, cultivated | F | Arima |
| 6 | Hyogo | Kobe | Spiny | Garden, cultivated | F | Arima |
| 7 | Hyogo | Asago | Spiny | Agricultural field in fruit tree research institute for preservation purpose, cultivated, originally transplanted from Mt. Inariyama, agricultural field | F | Arima |
| 8 | Hyogo | Asago | Spiny | Agricultural field in fruit tree research institute for preservation purpose, cultivated, originally transplanted from Mt. Inariyama, agricultural field, grafted from the same tree as plant 9 | F | Arima |
| 9 | Hyogo | Asago | Spiny | Agricultural field in fruit tree research institute for preservation purpose, cultivated, originally transplanted from Mt. Inariyama, agricultural field, grafted from the same tree as plant 8 | F | Arima |
| 10 | Hyogo | Kobe | Spiny | Garden, cultivated | F | Arima |
| 11 | Hyogo | Kobe | Spiny | Garden, cultivated | F | Arima |
| 12 | Hyogo | Kobe | Spiny | Garden, cultivated | F | Arima |
| 13 | Hyogo | Yabu | Spineless | Garden, cultivated | D | Asakura |
| 14 | Gifu | Takayama | Spiny | Garden, cultivated | E | Takahara |
| 15 | Gifu | Takayama | Spiny | Agricultural field, cultivated | E | Takahara |
| 16 | Gifu | Takayama | Spineless | Agricultural field, cultivated | E | Takahara |
| 17 | Gifu | Takayama | Spineless | Agricultural field, cultivated | E | Takahara |
| 18 | Gifu | Takayama | Spineless | Agricultural field, cultivated | E | Takahara |
| 19 | Gifu | Takayama | Spineless | Agricultural field, cultivated | E | Takahara |
| 20 | Gifu | Takayama | Spineless | Garden, cultivated, used as mother tree for grafting | E | Takahara |
| 21 | Gifu | Takayama | Spineless | Agricultural field, cultivated, used as mother tree for grafting | E | Takahara |
| 22 | Gifu | Takayama | Spineless | Agricultural field, cultivated | E | Takahara |
| 23 | Gifu | Takayama | Spineless | Agricultural field, cultivated | E | Takahara |
| 24 | Gifu | Takayama | Spineless | Garden, cultivated | E | Takahara |
| 25 | Gifu | Takayama | Spineless | Agricultural field, cultivated, about 30 years old | E | Takahara |
| 26 | Gifu | Takayama | Spineless | Agricultural field, cultivated, about 25-26 years old | E | Takahara |
| 27 | Gifu | Takayama | Spiny | Agricultural field, cultivated | E | Takahara |
| 28 | Gifu | Takayama | Spiny | Agricultural field, cultivated | E | Takahara |
| 29 | Hyogo | Kobe | Spiny | Garden, cultivated, originally transplanted from Mt. Rokko | C | Arima |
| 30 | Hyogo | Kobe | Spiny | Garden, cultivated, originally transplanted from Mt. Rokko | C | Arima |
| 31 | Hyogo | Asago | Spiny | Agricultural field in fruit tree research institute for preservation purpose, cultivated, originally transplanted from Yubunedani of Mt. Rokko | C | Arima |
| 32 | Hyogo | Asago | Spiny | Agricultural field in fruit tree research institute for preservation purpose, cultivated, originally transplanted from Yubunedani of Mt. Rokko | C | Arima |
| 33 | Hyogo | Asago | Spiny | Agricultural field in fruit tree research institute for preservation purpose, cultivated, originally transplanted from Yubunedani of Mt. Rokko, grafted from the same tree as plant 34 | C | Arima |
| 34 | Hyogo | Asago | Spiny | Agricultural field in fruit tree research institute for preservation purpose, cultivated, originally transplanted from Yubunedani of Mt. Rokko, grafted from the same tree as plant 33 | C | Arima |
| 35 | Hyogo | Yabu | Spineless | Agricultural field, cultivated, near the Konryuji temple | A | Asakura |
| 36 | Kyoto | Kyōtamba | Spineless | Agricultural field, cultivated | A | Asakura |
| 37 | Hyogo | Kobe | Spineless | Agricultural field, cultivated, purchased as nursery stock of high-yielding Asakura sanshō | A | Asakura |
| 38 | Hyogo | Yabu | Spineless | Agricultural field, cultivated, near the Konryuji temple | A | Asakura |
| 39 | Hyogo | Yabu | Spineless | Agricultural field, cultivated, near the Konryuji temple | A | Asakura |
| 40 | Kyoto | Kyōtamba | Spineless | Agricultural field, cultivated | A | Asakura |
| 41 | Kyoto | Kyōtamba | Spineless | Agricultural field, cultivated | A | Asakura |
| 42 | Hyogo | Yabu | Spineless | Agricultural field, cultivated, used as high-yielding mother tree for grafting | A | Asakura |
| 43 | Kyoto | Ayabe | Spineless | Agricultural field, cultivated | A | Asakura |
| 44 | Nara | Gojō | Spineless | Agricultural field, cultivated | A | Asakura |
| 45 | Kyoto | Kyōtamba | Spineless | Agricultural field, cultivated | A | Asakura |
| 46 | Hyogo | Yabu | Spineless | Agricultural field, cultivated | A | Asakura |
| 47 | Hyogo | Kobe | Spineless | Agricultural field, cultivated, purchased as nursery stock of Asakura sanshō | A | Asakura |
| 48 | Hyogo | Yabu | Spineless | Agricultural field, cultivated, used as mother tree for grafting | A | Asakura |
| 49 | Nara | Gojō | Spineless | Agricultural field, cultivated | A | Asakura |
| 50 | Nara | Gojō | Spineless | Agricultural field, cultivated | A | Asakura |
| 51 | Hyogo | Kami | Spineless | Agricultural field, cultivated | A | Asakura |
| 52 | Hyogo | Yabu | Spineless | Mountain, wild tree | D | Asakura |
| 53 | Hyogo | Yabu | Spineless | Mountain, wild tree, near the Konryuji temple | D | Asakura |
| 54 | Hyogo | Yabu | Spineless | Mountain, wild tree, near the Konryuji temple | D | Asakura |
| 55 | Hyogo | Yabu | Spiny | Agricultural field, cultivated, near the Konryuji temple | D | Asakura |
| 56 | Hyogo | Yabu | Spineless | Mountain, wild tree | D | Asakura |
| 57 | Hyogo | Yabu | Spineless | Mountain, wild tree | D | Asakura |
| 58 | Hyogo | Yabu | Spineless in<br>2017, spiny in<br>2020 | Agricultural field, cultivated, 100 years old tree | D | Asakura |
| 59 | Hyogo | Yabu | Spineless | Mountain, wild tree | D | Asakura |
| 60 | Hyogo | Yabu | Spineless | Mountain, wild tree, near the Konryuji temple | D | Asakura |
| 61 | Hyogo | Yabu | Spiny | Possible abandoned tree near an abandoned house | D | Asakura |
| 62 | Hyogo | Yabu | Spineless | Mountain, wild tree | D | Asakura |
| 63 | Hyogo | Yabu | Spineless | Mountain, wild tree | D | Asakura |
| 64 | Hyogo | Yabu | Spiny | Mountain, wild tree | D | Asakura |
| 65 | Hyogo | Yabu | Spineless | Mountain, wild tree | D | Asakura |
| 66 | Hyogo | Yabu | Spiny | Mountain, wild tree, near the Konryuji temple | D | Asakura |
| 67 | Hyogo | Yabu | Spiny | Mountain, wild tree | D | Asakura |
| 68 | Hyogo | Yabu | Spiny | Mountain, wild tree | D | Asakura |
| 69 | Hyogo | Yabu | Spineless | Mountain, wild tree | D | Asakura |
| 70 | Hyogo | Yabu | Spineless | Mountain, wild tree | D | Asakura |
| 71 | Hyogo | Yabu | Spineless | Mountain, wild tree | D | Asakura |
| 72 | Hyogo | Yabu | Spiny | Mountain, wild tree | D | Asakura |
| 73 | Hyogo | Yabu | Spiny | Mountain, wild tree | D | Asakura |
| 74 | Hyogo | Kami | Spineless | Garden, spontaneously grown tree | D | Asakura |
| 75 | Ishikawa | Noto | Spineless | Mountain, wild tree | G | Unclassified |
| 76 | Yamagata | Tsuruoka | Spiny | Garden, spontaneously grown tree | G | Unclassified |
| 77 | Shiga | Nagahama | Spineless | Agricultural field, cultivated | G | Unclassified |
| 78 | Niigata | Agamachi | Spineless | Mountain, wild tree | G | Unclassified |
| 79 | Niigata | Agamachi | Spineless | Mountain, wild tree | G | Unclassified |
| 80 | Hyogo | Kami | Spineless | Agricultural field, cultivated | G | Unclassified |
| 81 | Yamagata | Tsuruoka | Spineless | Garden, spontaneous grown tree, growing adjacent to plant<br>83 | G | Unclassified |
| 82 | Akita | Akita | Spiny | Transplanted from mountain, wild tree | G | Unclassified |
| 83 | Yamagata | Tsuruoka | Spineless | Garden, spontaneous grown tree, growing adjacent to plant<br>81 | G | Unclassified |
| 84 | Tokyo | Adachi | Spiny | Garden, cultivated from the seed of an old tree | G | Unclassified |
| 85 | Ishikawa | Noto | Spineless | Mountain, wild tree | G | Unclassified |
| 86 | Ishikawa | Noto | Spineless | Mountain, wild tree | G | Unclassified |
| 87 | Kochi | Sakawa | Spiny | Agricultural field, cultivated | H | Unforeknown<br>Kochi group |
| 88 | Kyoto | Ayabe | Spineless | Agricultural field, cultivated | G | Unclassified |
| 89 | Hyogo | Yabu | Spineless | Garden, cultivated, near the Konryuji temple | D | Asakura |
| 90 | Hyogo | Yabu | Spineless | Mountain, wild tree, near the Konryuji temple, located in the<br>recorded place of the origin of Asakura lineage | D | Asakura |
| 91 | Hyogo | Toyooka | Spiny | Mountain, wild tree | G | Unclassified |
| 92 | Kochi | Nangoku | Spiny | Agricultural field, cultivated | G | Unclassified |
| 93 | Hyogo | Kobe | Spineless | Garden, spontaneously grown tree | G | Unclassified |

Group A may be an Asakura lineage group because the area across which the group A plants were distributed coincided with the production area for the Asakura lineage in Yabu City, central Hyogo Prefecture (Figure 2, Table 1). All plants in group A were spineless. Group A was widely distributed and cultivated in other areas, including Kobe and Kami in Hyogo Prefecture, Ayabe, and Kyōtamba in Kyoto Prefecture, and Gojō in Nara Prefecture (Table 1). Interestingly, plants 42, 43, 44, 45, 46, 47, 48, 49, 50, and 51 formed tight cluster, and are from Yabu, Ayabe, Gojō, Kyōtamba, Yabu, Kobe, Yabu, Gojō, Gojō, and Kami, respectively. The other plants in group A (35, 36, 37, 38, 39, 40, and 41) were from Yabu, Kyōtamba, Kobe, Yabu, Yabu, Kyōtamba, and Kyōtamba, respectively.

The analysis identified group D as the second Asakura lineage group. Group D plants were from Yabu City only (Figure 1a, Figure 2, Table 1), except for plant 74, which was from Kami. Unlike group A, which had more cultivated plants, group D contained many wild plants found in mountainous areas. Most, but not all, of the group D plants were spineless. Plant 58 was 100 years old and was growing in an agricultural field. When we observed plant 58 in 2017, this plant was spineless. As of 2020, this plant was near death, and its spines were visible.

The analysis identified group E as the Takahara lineage. Group E (Figure 1a) was an isolated group with plants only from Takayama city, located in the highland region of northern Gifu Prefecture (Figure 2, Table 1). This group consisted of only cultivated plants. Most of the group E plants were spineless (Table 1).

The analysis identified group C as the first group of the Arima lineage. Group C plants were from Kobe and Asago cities in southern Hyogo Prefecture (Figure 2, Table 1), where the Arima lineage was produced. There is an agricultural field in Asago City for the preservation of the Arima lineage. All plants in group C were spiny and transplanted from Mt. Rokko (Table 1).

The analysis identified group F as the second Arima lineage group. All plants in group F (Table 1) were from the Arima lineage production area in Asago and Kobe (Figure 2). Group F contained spiny plants. Three plants cultivated at the Fruit Tree Research Institute came from Mt. Inariyama, located near Arima. PCA did not clearly separate group F from the other plants.

The analysis identified group B as the Budou lineage. Group B was from Kainan city in Wakayama Prefecture (Figure 2), where the Budou lineage is produced (Table 1). Group B is the grape-like Budou lineage, which has a different phenotype than the other lineages. The plants in group B were all cultivated and had spines.

Group G was designated as an unclassified lineage. This plant group included plants from different locations across Japan (Table 1), including Adachi, Akita, Agamachi, Ayabe, Kami, Kobe, Nagahama, Nangoku, Noto, Toyooka, and Tsuruoka. The plants in group G were both cultivated and wild (Table 1).

### Tight cluster and additional groups revealed by cluster analysis

Cluster analysis was performed using a pairwise distance matrix. The cluster analysis identified the same seven groups that were identified in the PCA analysis and showed that plants 42, 43, 44, 45, 46, 47, 48, 49, 50, and 51 formed tight cluster.

In addition, cluster analysis identified an eighth group (Figure 3), named group H. Group H (Figure 3) contained the plants from Kochi Prefecture (Figure 2, Table 1). This was named “unforeknown Kochi group”. The plants in this group were spiny. Plants 1 and 87 were cultivated, and plant 2 was wild. These plants are used as ingredients to make rice cakes in this region (Table 1). The other plant (number 92) from Kochi Prefecture did not belong to group H.

**Figure 3.**
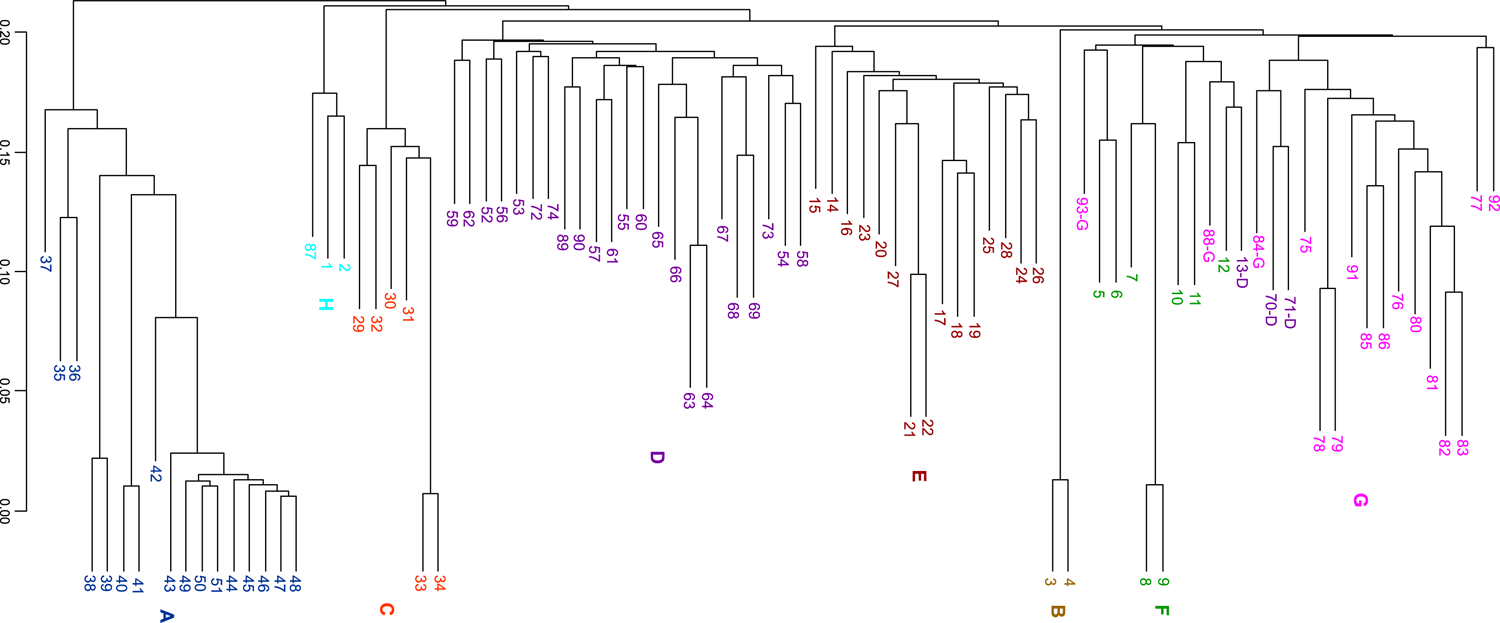
Cluster analysis of *Zanthoxylum* accessions. The new group, H group, is shown in light blue. The colour scheme is the same as in Figure 1, with an additional colour. Figure was generated using R software (version 3.6.2)^44^.

### Regional groups and their exceptions elucidated through phylogenetic analyses

To further establish the above classifications, we performed phylogenetic analyses using the maximum likelihood (ML) method (Figure 4) and a method based on the coalescent model (Figure 5). Both phylogenetic trees classified plants into eight groups. The phylogenetic trees (Figures 4 and 5) also showed a tight cluster in group A, containing plants 42, 43, 44, 45, 46, 47, 48, 49, 50, and 51, present as a subclade within clade A with a high bootstrap value. The branch lengths of the ML tree (Figure 4) indicated that these nine plants had a high degree of genetic similarity.

**Figure 4.**
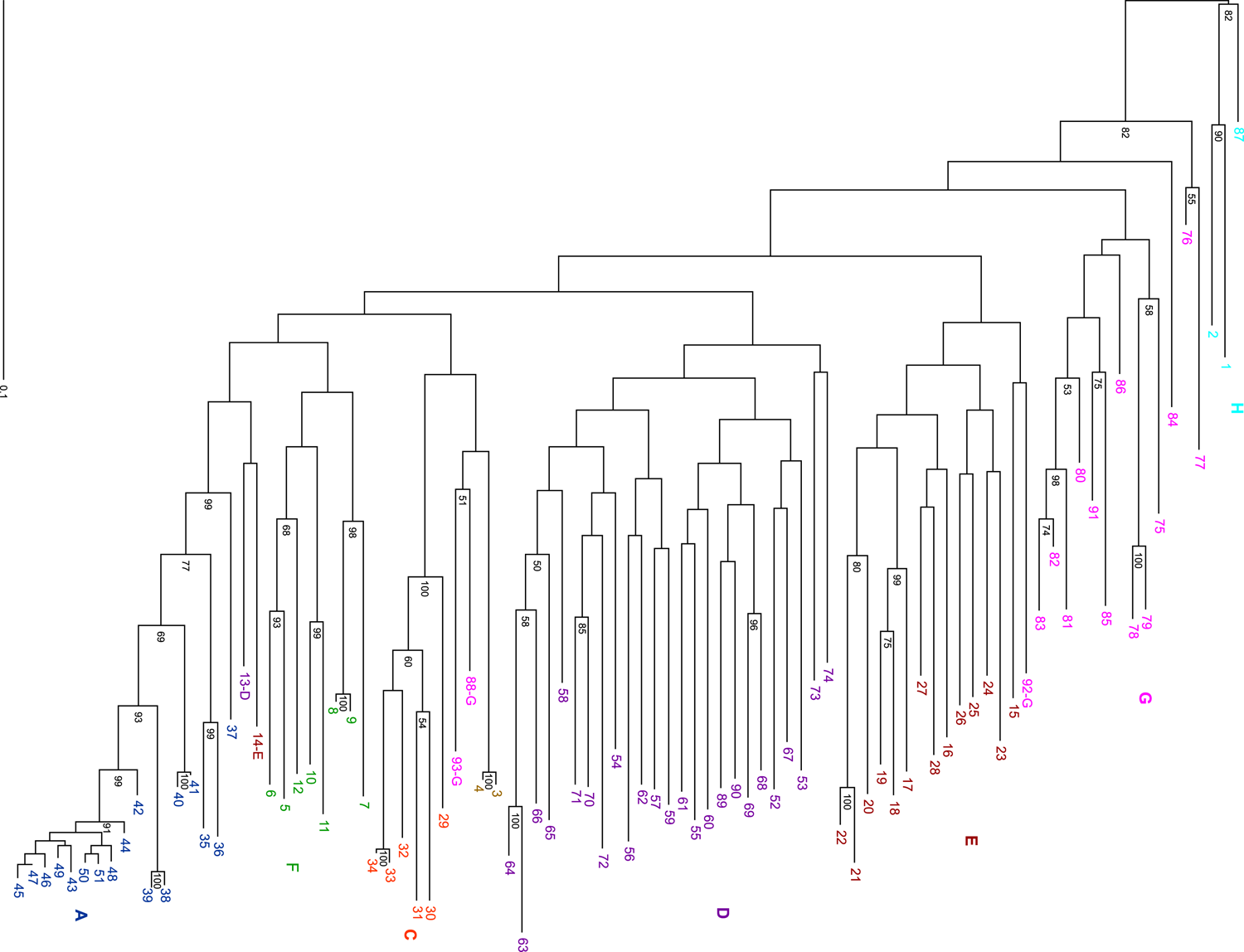
Maximum likelihood phylogenetic analysis of *Zanthoxylum* accessions. The colour scheme is the same as in Figure 3. The numbers at the nodes indicate bootstrap values (% over 1,000 replicates). The nodes with ≥50% bootstrap value are labeled. Figure was generated using Dendroscope (version 3.6.3)^48^ and TreeView X (version 0.5.0, https://treeview-x.en.softonic.com/)

**Figure 5.**
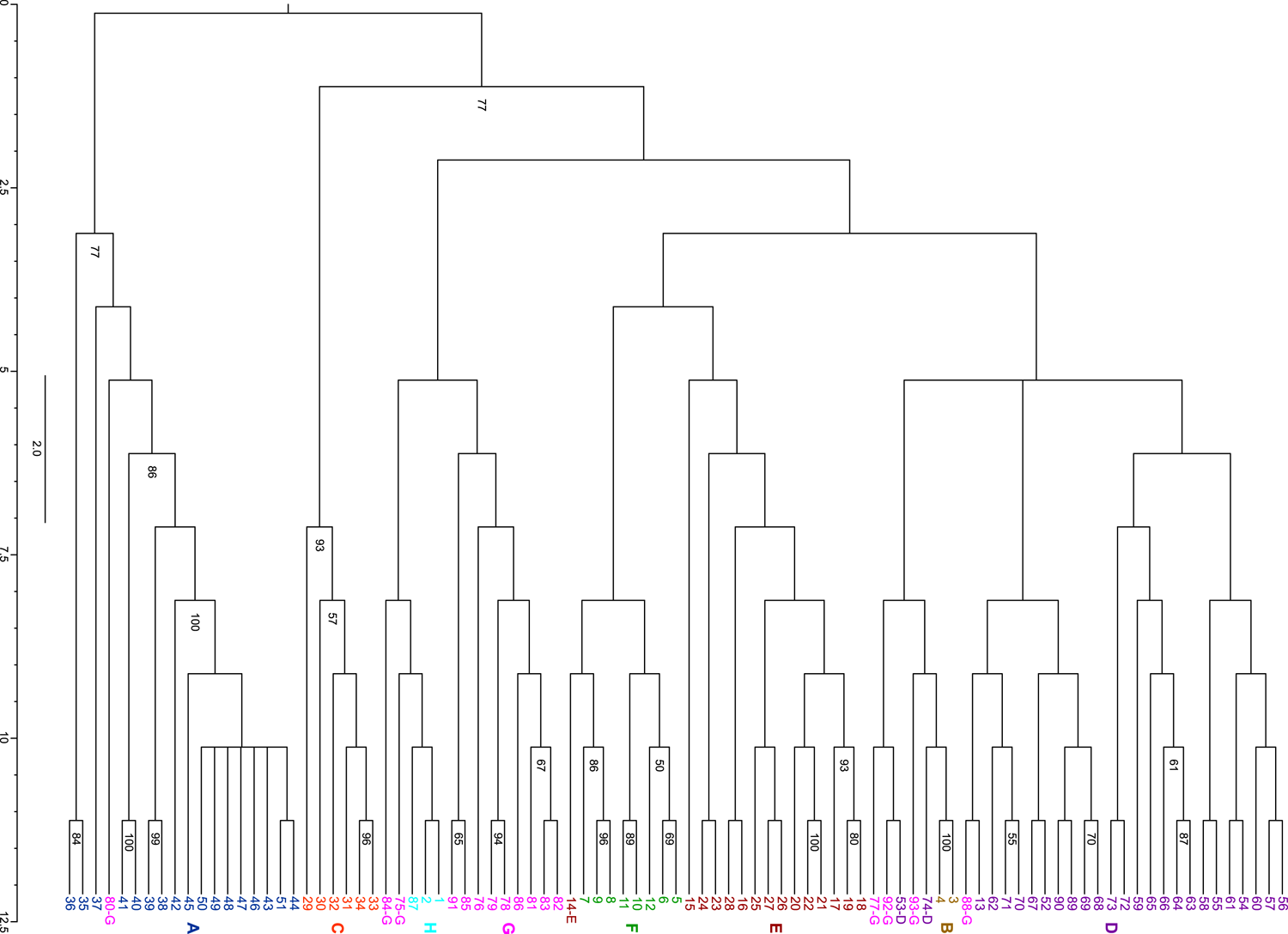
Coalescent model phylogenetic tree of *Zanthoxylum* accessions. The colour scheme is the same as in Figure 3. The numbers at the nodes indicate bootstrap values (% over 1,000 replicates). The nodes with ≥50% bootstrap value are labeled. Figure was generated using FigTree (version 1.4.4, http://tree.bio.ed.ac.uk/).

There were some differences in these eight groups between the two phylogenetic trees. Two trees separated plant 14 from the other plants in group E. The ML tree separated plant 13 from other plants in group D. Indeed, in PCA, these two plants were closer to the plants in the other groups (Figure 1a and 1b). The discrepancies between the two phylogenetic analyses, including these ones, are related to the low bootstrap values.

The cluster analysis and both phylogenetic analyses showed that group G had two regional groups in common: (1) plants 78 and 79, and (2) plants 81, 82, and 83. Plants 78 and 79 in the first group were from Agamachi, Niigata Prefecture, and were wild spineless plants. In the second group, plant 81 was from Yamagata Prefecture, plant 82 was from Akita Prefecture, and plant 83 was from Tsuruoka, Yamagata Prefecture, which are adjacent to each other. However, plant 76 from Tsuruoka, Yamagata Prefecture, did not belong to this regional group.

In both phylogenetic trees, group G plants 88, 92, and 93 were isolated from the other group G plants. In group G in the coalescent tree (Figure 5), plants 77 and 80 were separated from other plants in the same group. This observation suggests that group G may not be a single isolated group. Thus, the grouping of group G is ambiguous, probably because plants in various locations belonged to group G.

### Admixture history of the eight groups and the tight cluster within group A

Admixture analysis was used to estimate ancestral history and calculated cross-validation (CV) errors to estimate possible values of *K* (number of ancestral populations) = 1–12 (Supplementary Figure 1). *K* = 2 was the most likely value because the error value of the cross-validation was minimal. We presented the results of the admixture analysis from *K* = 2 to *K* = 7 (Figure 6).

**Figure 6.**
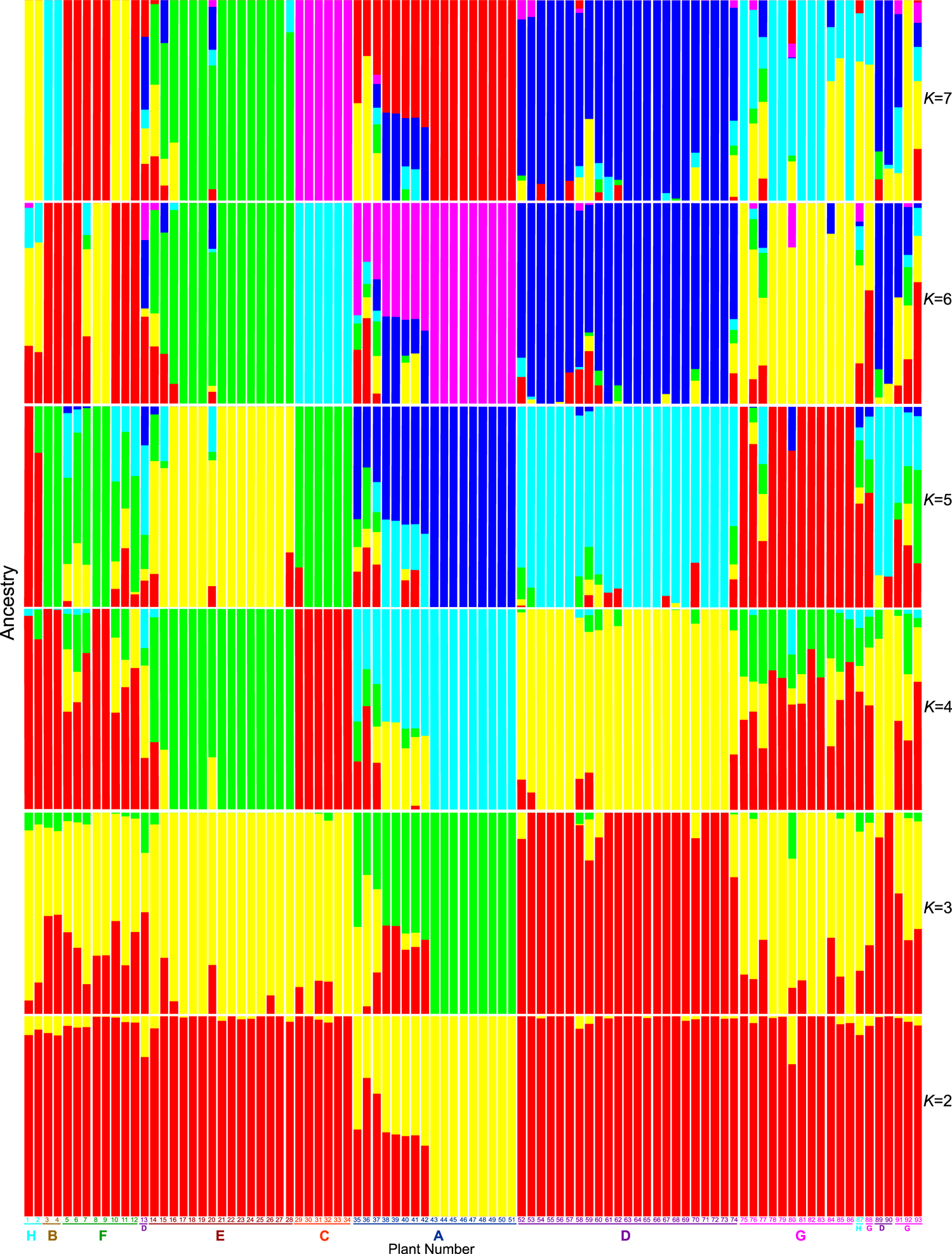
*K* = 2-7 admixture plots in admixture analysis of *Zanthoxylum* accessions used in this study. The horizontal axis shows the group names and the sample numbers. The colour scheme for the group names and the sample numbers is the same as in Figure 3. The colour for the genetic cluster is different from that used to denote the group names and sample numbers. Figure was generated using R software (version 3.6.2)^44^.

In the case of *K* = 2 (Figure 6), except for plant 42, the tight cluster within group A (plants 43, 44, 45, 46, 47, 48, 49, 50, and 51) formed a single population, shown in yellow. The red colour corresponds to plants other than those in group A. The remaining group A plants were admixed groups of the two populations. Thus, at *K* = 2, group A members were clearly separated from the other plants.

The analysis of *K* = 3–7 separated the six groups A, B, C, D, E, and H (Figure 6). However, the analysis did not separate group F, which is consistent with the results of PCA, which also did not clearly separate group F from other plants.

The admixture analysis also showed that the grouping of group G was ambiguous, and group G may not be a single isolated group. Reflecting this observation, the pattern of colours in group G was not consistent across several *K* values. Thus, the plants in group G plants may be admixtures of a variety of groups.

### Conservation of heterozygosity

Asexual reproduction was confirmed by checking for heterozygosity conservation using pairwise alignments. Plants 8 and 9 of group F were grafted from the same tree (Table 1). Between these two plants, 2288 of 3699 sites were conserved heterozygous sites (61.9%) (Supplementary Fig. 2). Similarly, plants 33 and 34 of group C were grafted from the same tree (Table 1). Between these two plants, 1947 of the 3097 sites were conserved heterozygous sites (62.9%) (Supplementary Fig. 3). Grafting is a common method of propagating the Budou lineage, namely, group B plants. Between plants 3 and 4 of group B, 2198 of 3879 sites were conserved heterozygous sites (56.7%) (Supplementary Fig. 4). Thus, this degree of heterozygosity conservation was found among asexually reproduced plants when analysing the data using *de novo* mapping.

Between plants 40 and 41 of group A, growing nearby, 2140 of 3414 sites were conserved heterozygous sites (62.7%) (Supplementary Fig. 5); between plants 38 and 39 of group A, growing nearby, 1700 of 3502 sites were conserved heterozygous sites (48.5%) (Supplementary Fig. 6). Thus, plant pairs “40 and 41” and “38 and 39” were identified as asexually propagated plants, although no records have been kept regarding their propagation. Based on the analysis of these five pairs, we observed a conservation of heterozygosity of approximately 50% or above between asexually propagated plants when the RAD-Seq data were analysed *de novo*.

We examined the conservation of heterozygosity for the tight cluster in group A (plants 42, 43, 44, 45, 46, 47, 48, 49, 50, and 51). The degree of conservation ranged from 50.4% to 68.9% (Supplementary Fig. 7, Supplementary Table 3). By contrast, the plants from the tight clusters within group A shared lower levels of conservation (from 29.5% to 36.5%) with the other group A plants (plants, 35, 38, and 40) (Supplementary Fig. 7, Supplementary Table 3). Therefore, the members of the tight cluster in group A were asexually propagated plants derived from the same tree.

### Statistical interpretations of the classifications

In our statistical analysis (Table 2 and 3, Supplementary Table 4, 5, and 6), we removed the plants (13 and 14) for which we obtained uncertain results and the plants in group G that caused the ambiguity. Although groups C and F belonged to the Arima lineages, the Fst value between them was high (0.110, Table 2). Thus, the difference in the pairwise Fst values between these two groups may indicate two places of origin.

**Table 2.** Fst values for each pair of groups for A, B, C, D, E, F, H.

| Group | B | F | E | C | A | D |
| --- | --- | --- | --- | --- | --- | --- |
| H | 0.307 | 0.120 | 0.0847 | 0.165 | 0.1080 | 0.0490 |
| B |  | 0.130 | 0.0950 | 0.191 | 0.125 | 0.0529 |
| F |  |  | 0.0689 | 0.110 | 0.103 | 0.0457 |
| E |  |  |  | 0.0874 | 0.0994 | 0.0464 |
| C |  |  |  |  | 0.126 | 0.0572 |
| A |  |  |  |  |  | 0.0803 |

Similarly, the pairwise Fst value between groups A and D of the Asakura lineage was high (0.0803, Table 2). The reason for the higher Fst value may be the presence of a tight cluster in group A. Therefore, we performed a second statistical analysis by modifying the grouping. In the new grouping, we designated the tight cluster in group A as A2. We denoted the plants that remained in group A as A1. As expected, the pairwise Fst values were high between group A2 and each of the other groups (Table 3). Table 3 shows that group A1 was genetically closer to group D (0.0539). Thus, the high genetic distance between groups A and D in Table 2 was due to the tight cluster within group A.

**Table 3.** Fst values for each pair of groups for A1, A2, B, C, D, E, F, H.

| Group | B | F | E | C | A1 | A2 | D |
| --- | --- | --- | --- | --- | --- | --- | --- |
| H | 0.268 | 0.109 | 0.0760 | 0.156 | 0.132 | 0.196 | 0.0464 |
| B |  | 0.127 | 0.0950 | 0.189 | 0.159 | 0.226 | 0.0531 |
| F |  |  | 0.0697 | 0.111 | 0.100 | 0.154 | 0.0454 |
| E |  |  |  | 0.0878 | 0.0807 | 0.132 | 0.0464 |
| C |  |  |  |  | 0.134 | 0.195 | 0.0582 |
| A1 |  |  |  |  |  | 0.0990 | 0.0539 |
| A2 |  |  |  |  |  |  | 0.0993 |

We performed tests for f3-statistics and f4-statistics (Supplementary Tables 5 and 6, respectively). Negative values of f3-statistics with A1 as the outgroup (Supplementary Table 5) showed that group A1 is an admixture population of group A2 and the remaining plants. This result is in good agreement with the results of the admixture analysis (Figure 6). The f4-statistics test (Supplementary Table 6) also supports this result with high or low Z-scores. As for source populations other than A2, groups C and E were more likely than group D. Although groups C and F belonged to the Arima lineages, positive values of f3-statistics with A2 as the outgroup showed that group C may have more gene flow from groups E, D, and H than from group F.

## Discussion

Our study classified *Zanthoxylum piperitum* into several groups based on DNA sequences and geographic information. Each group showed interesting characteristics, some of which may have been related to domestication. The domestication of *Zanthoxylum piperitum* can be described in two steps. The first step is the cultivation of wild plants in agricultural fields, in which case the wild and cultivated plants are genetically similar. In the second step, the lineages with desirable traits spread from their place of origin to the surrounding areas, in which case the genetic diversity of individuals within the lineages should be low. Grafting is a means of spreading crops such as fruit trees.

Group D plants reflect the first step of domestication, because the cultivated plants probably originated in the Yabu Mountains. Group D comprised both wild plants in mountainous areas and cultivated plants in agricultural fields (Table 1).

The spineless Asakura lineage (*Z. piperitum* (L.) DC forma *inerme* (Makino) Makino), first described 90 years ago^11^, may be placed in group D. This 90-year-old documentation^11^, based on a much older book, states that the origin of the Asakura lineage was on a cliff near the Konryuji temple. Plant 90, belonging to group D, grows in a place similar to the place of origin and is spineless. In addition, spineless wild plants 53, 54, 60, and 69 were collected near the Konryuji temple.

Group A plants reflect the second domestication step because they include lineages with desirable traits and appear to have been spread. Group A contained a tight cluster comprising plants 42, 43, 44, 45, 46, 47, 48, 49, 50, and 51. This cluster showed minimal genetic diversity (Supplementary Table 4). Furthermore, the amount of conservation of heterozygosity showed that these plants were propagated by grafting (Supplementary Fig. 7, Supplementary Table 3). These plants were also found to be widely cultivated in many cities (Yabu, Ayabe, Gojo, Kyotamba, Kobe, and Kami), indicating that they have been spread. Plants 42 and 48 are highly productive trees in Yabu City, which supplies scions for grafting. Thus, the plants in this tight cluster exhibited a variety of desirable traits.

There are two potential places of origin for group A: Yabu City, where the Asakura district is located and another distant location. The following reasons may explain the first possibility: (1) many of the group A plants grow in Yabu City (plants 35, 38, 39, 42, 46, and 48); (2) a 90-year-old document^11^ describes the spineless plants originating from Asakura District, Yabu City; and (3) all of the group A plants are spineless. Interestingly, we collected plants 35, 38, and 39 near the Konryuji temple, a potential place of origin for the Asakura lineage. Therefore, we do not exclude the possibility that the spineless Asakura lineage described 90 years ago^11^ may represent group A plants. However, plants 35, 38, and 39 were cultivated. Thus, a serious problem for the first possibility is the absence of wild plants in group A in the mountains of Yabu City.

The second possibility is that the group A plants are from another distant location; that is, the plants in the tight cluster of group A of unknown origin hybridised with the group D plants to form the rest of the group A plants. This possibility is consistent with the results of admixture analysis for *K* = 2. However, it is worth noting that the results of the admixture analysis may have been artificial since the second step of domestication formed the ancestral population containing the plants in the tight cluster. In addition, the f3-statistics test results (Supplementary Table 5) do not necessarily indicate that the source population of group A is group D. If the second possibility is true, the origin of the plants in the tight cluster remains unknown, although the origin of the rest of the group A plants can be explained by hybridisation.

In groups C and F of the Arima lineage, we observed the first domestication step based on documentation. According to the records, the group F plants (plants 7, 8, and 9) were transplanted from Mt. Inariyama to Asago city (Table 1). Group C plants (plants 29, 30, 31, 32, 33, and 34) were transplanted from Mt. Rokko to Asago and Kobe cities. The other group F plants (plants 5, 6, 10, 11, and 12) should have the same origin. According to the PCA results, the group F plants (plants 5, 6, 10, 11, and 12) were more closely related to the plants in the other groups.

We did not find any records describing the origins of the group E plants. In addition, we could not find wild representatives of this group in the mountains. If the group E plants have adaptations to high altitudes, the mountainous areas near Takayama city may be where group E originated. Spineless plants are present in group E (plants 16, 17, 18,19, 20, 21, 22, 23, 24, 25, and 26) and are genetically different from the plants in groups A and D in Yabu City, where the Asakura district is located.

In addition to these observations, the results for group G also suggest that domestication is underway in many areas. Ongoing domestication is associated with the cultivation of wild lineages on nearby agricultural lands and gardens. Group G was genetically different from the other plant groups undergoing domestication. Therefore, these plants may have been domesticated at their respective locations.

Recent reports have stated that the domestication of other crop species, such as olives in Morocco^33^ and dragon fruits/pitaya in Mexico^34^, is also ongoing. Both olives and dragon fruits undergo two common steps to become domesticated: first, the *in situ* conservation of wild plants, and second, the introduction of wild plants into agricultural fields. These plants were selected as the desirable plants for cultivation. The ongoing domestication of Japanese pepper is not entirely similar to that of olives and dragon fruit. In the first step of olive domestication, farmers continue to disturb the natural habitat of the olive when selecting wild populations^33^. For Japanese pepper, wild plants are harvested from their natural habitats and transplanted into agricultural fields or gardens while leaving other wild plants intact in the mountains. The second step of the domestication of olives and dragon fruits occurs through the continuous introduction of wild plants into agricultural fields to further select for preferred traits^33, 34^. In Japanese pepper, wild plants are not frequently reintroduced into cultivated plant sites.

On the other hand, the ongoing domestication of Japanese pepper is similar to that of olives and dragon fruit in which domestication syndrome has been observed. Domestication syndrome changes the genomic, physiological, and morphological characteristics of cultivated plants compared to those of wild plants^35^. The genomic characteristics of the plants in the tight cluster in group A were different from those of the wild plants. Similarly, the morphological characteristics of the plants in group B were different from those of wild plants.

Spineless plants are known as the Asakura lineage and are classified as a single subspecies, *Z. piperitum* (L.) DC forma *inerme* (Makino) Makino^11^. However, it is inappropriate to refer to the spineless plant as the Asakura lineage or *Z. piperitum* (L.) DC forma *inerme* (Makino) Makino. One Asakura lineage group (group A) contained a tight cluster of spineless plants of unknown origin. Although the second Asakura lineage group (group D) had a plant growing in a recorded place of origin^11^, it contained some spiny plants. In addition, the spineless trait is not unique to the Asakura lineage, as some of the plants in the Takahara lineage (group E) are also spineless. Some group G plants were spineless. Importantly, these spineless plants were from many locations. Therefore, our study does not support the notion that the spineless lineage is either from Asakura or monophyletic. Our results indicate that the spineless lineage is not a single subspecies.

Some Fis values were negative (Supplementary Table 4). Notably, the tight cluster plants in group A (group A2) had the highest negative Fis value. Therefore, we speculated that some of these plants had been asexually reproduced by grafting, as asexual reproduction may cause negative Fis values^36^. Pairwise alignments support this possibility (Supplementary Fig. 7 and Supplementary Table 3).

In this study, we elucidated the genetic diversity of Japanese pepper, while tracing the process of domestication. Our results demonstrate that the conservation of plant genetic resources is an important agricultural practice. Several cultivated lineages of Japanese pepper were found to be closely related to plants growing in the mountains, in contrast to other crops. On the other hand, several cultivated lineages, groups A, B, and E, had no close relatives growing in the mountains, which suggests that wild plants of this group no longer exist. Therefore, there is a need to conserve both plants growing in the mountains *in situ* and plants growing in agricultural fields or preservation facilities *ex situ*. The results presented in this study will be useful for developing a plan for the conservation and breeding of Japanese peppers.

## Methods

### Plant materials

Ninety-three *Zanthoxylum piperitum* samples collected from various parts of Japan were used (Table 1). Photographs of the materials were taken, and the leaves were stored in a freezer (‒80°C). At each location, where possible, plants growing in the mountains and agricultural fields were collected. Any lineages of Budou or Takahara that grew in the mountains were not found. All methods involving plants were carried out in accordance with relevant guidelines and regulations. Plants were collected with permission from their owners.

### DNA extraction and double-digest restriction site amplified DNA sequencing (ddRAD-Seq)

DNA extraction and purification were performed using previously described methods^30^. The library for ddRAD-Seq was prepared based on the original protocol^27^ with some modifications^37^. The first restriction site near the primer binding site, which reads the single-ended DNA sequence, was *Bgl*II. The second restriction site, adjacent to the binding site for reading the index sequence, was *Eco*RI. The library was sequenced with 51-bp single-end reads at Macrogen (Seoul, Korea) on two lanes of an Illumina HiSeq2000 (Illumina, San Diego, CA, USA).

### Processing and quality control of ddRAD-Seq data

The Stacks package^38^ was used to analyse the ddRAD-Seq reads. The process_shortreads program of the Stacks package (version 2.4) was used to clean reads using -c (clean data, remove any read with an uncalled base), -q (discard reads with low-quality scores), and -r (rescue barcodes) options. The cutadapt program (version 2.6)^39^ removed adaptor sequences from reads using the -b (remove adapter anywhere, the sequence of an adapter that may be ligated to the 5ʹ or 3ʹ end) and -e 0.1 (maximum allowed error rate by keeping the default value 0.1 = 10%) options. Any low-quality bases in the single-end reads were trimmed using the trimmomatic program (version 0.39)^40^ with the following settings LEADING:19 (cut bases off the start of a read, if below a threshold quality), TRAILING:19 (cut bases off the end of a read, if below a threshold quality), AVGQUAL:20 (drop the read if the average quality is below a specified level), MINLEN:51 (drop the read if it is below a specified length), and SLIDINGWINDOW:20:20 (perform sliding window trimming, clipping once the average quality within the window falls below a threshold).

### *De novo* mapping and variant calling

The denovo_map.pl of the Stacks package (version 2.5), a wrapper script for ustacks, cstacks, sstacks, tsv2bam, and gstacks, was used to map reads *de novo* without using a reference genome. The options for denovo_map.pl were -M 4 (number of mismatches allowed between stacks within individuals [for ustacks]), -n 4 (number of mismatches allowed between stacks between individuals [for cstacks]), and -m 3 (number of identical reads required to initiate a new putative allele [for ustacks]). Genotyping data were created to specify all the samples under analysis by assigning one individual per population. After performing denovo_map.pl, the populations program of the Stacks package (version 2.5) created the vcf (variant call format)^41^, plink^42^, phylip, and treemix files using the -R 0.5 (minimum percentage of individuals across populations required to process a locus), --write-single-snp (restrict data analysis to only the first SNP per locus), --min-maf 0.05 (minimum minor allele count required to process a SNP), --vcf, --plink, --phylip-var-all --treemix options. The populations program was also used to create pairwise alignments between the two individuals. In this case, the -R option was set to one.

### Principal component analysis (PCA) and cluster analysis

Principal component analysis was performed based on the vcf file generated by the populations program. The SNPRelate program^43^ in the R software environment (version 3.6.2)^44^ converted the vcf file to a gds (genomic data structure) file, drew the PCA diagrams, and calculated the contribution ratios for each principal component. This step used only bi-allelic loci. The SNPRelate program plotted the dendrogram by considering the identity by state (IBS) pairwise distances. Images of the results were generated using the basic functions of the R software environment.

### Admixture analysis

The plink program (plink 2, version 1.90p)^42^ was used to create the input files for the admixture program. The admixture program (version 1.3)^45^ was used to determine the admixture history and the cross-validation (CV) error for the hypothetical runs from *K* (number of ancestral populations) = 1–12. The CV error plot was drawn using the received log data to determine the optimal *K* value. R software (version 3.6.2) was used to draw admixture plots using the Q estimate files created by the admixture programs. Images of the results were generated using the basic functions of the R software environment.

### Maximum likelihood phylogenetic tree analysis

ModelTest-NG was used to select the model^46^. The maximum likelihood (ML) phylogenetic tree was created using the raxmlHPC-PTHREADS-SSE3 program (version 8.2.12)^47^. The phylip file created by the populations program was used as the input file. The parameters for raxml were -f a (rapid bootstrap analysis and search for the best-scoring ML tree in one program run), -x 12345 (an integer number [random seed] and turn on rapid bootstrapping), -p 12345 (a random number seed for parsimony inferences), -N 1000 (bootstrap value), and -m GTRGAMMAX (model of binary [morphological], nucleotide, multi-state, or amino acid substitution). Group H was used as the root. Image of the phylogenetic tree was generated using Dendroscope (version 3.6.3)^48^ and TreeView X (version 0.5.0, https://treeview-x.en.softonic.com/).

### Phylogenetic tree analysis under the coalescent model

Phylogenetic tree analysis was performed using SVDquartets^49^ integrated into PAUP (version 4.0a) software^50^. The input file for the PAUP was created by converting a phylip file containing all sites to the nexus file format. The parameters used for SVDquartets were ‘quartet evaluation’, ‘evaluate all possible quartets’, ‘tree inference’, ‘select trees’ ‘using QFM quartet assembly’, ‘tree model’, ‘multi-species coalescent’, ‘write quartets file’ ‘QMC format’, ‘handling of ambiguities’ and ‘distribute’. The number of bootstrap analyses was 1,000 replicates. Group A was used as the root. Image of the phylogenetic tree was generated using FigTree (version 1.4.4, http://tree.bio.ed.ac.uk/).

### Statistical analysis

Groups separated by PCA and cluster analysis were assigned as separate populations in the population map data required for the Stacks package. The denovo_map.pl was re-performed after modifying the data for the population map. The populations program was run with the options -R 0.5, --write-single-snp, --min-maf 0.05, --fstats (SNP and haplotype-based F statistics), and --Fst_correction (a correction to be applied to Fst values with p-value [default p-value to keep an Fst measurement: 0.05]). The program treemix (version 1.12)^51^ inferred gene flow using the f3- and f4-statistics with the option -k 500 (standard errors in blocks of 500 SNPs).

## Acknowledgments

We are grateful to those who helped us collect samples from various locations. We thank Ms. Yuki Sano, Mr. Atushi Shirakawa, and Mr. Kazuma Kubo for preparing the DNA samples. This work was supported by the Urakami Foundation for Food and Food Culture Promotion.

## Author contributions

N.F., K.Y., F.H., and Y.N. designed the study. N.F., H.M., N.I., and Y.N. collected the samples. Y.N. performed DNA extraction. M.D.G.P.P. and Y.N. performed the bioinformatic analysis. A.J.N. performed the RAD-Seq analysis. M.D.G.P.P., N.F., K.Y., F.H.H.M., N.I., and Y.N. analysed the data. M.D.G.P.P. and Y.N. wrote the manuscript.

## Data availability

Sequences are available at the DNA Data Bank of Japan Sequence Read Archive (https://www.ddbj.nig.ac.jp/dra/index-e.html; Accession no. DRA011046).

