## Supplementary information for "Elucidation of Japanese pepper (*Zanthoxylum piperitum* De Candolle) domestication using RAD-Seq"

[7] Gifu Agriculture and Forestry Office

[8] Association for Conservation of Asakura Sanshō

[#]These authors contributed equally to this work

**Supplementary Table 1**. Number of reads for each sample.

| Plant Number | Number of reads |
|---|---|
| 1 | 492,671 |
| 2 | 1,020,522 |
| 3 | 2,415,554 |
| 4 | 2,355,924 |
| 5 | 3,693,650 |
| 6 | 4,164,827 |
| 7 | 3,422,496 |
| 8 | 2,772,637 |
| 9 | 3,171,379 |
| 10 | 2,440,405 |
| 11 | 1,943,956 |
| 12 | 1,807,098 |
| 13 | 2,128,142 |
| 14 | 2,635,067 |
| 15 | 2,802,228 |
| 16 | 3,229,276 |
| 17 | 2,564,887 |
| 18 | 3,563,144 |
| 19 | 3,372,967 |
| 20 | 2,710,040 |
| 21 | 3,862,417 |
| 22 | 3,422,944 |
| 23 | 3,453,792 |
| 24 | 3,312,854 |
| 25 | 2,829,563 |
| 26 | 3,107,223 |
| 27 | 1,449,052 |
| 28 | 2,211,696 |
| 29 | 4,053,452 |
| 30 | 3,324,127 |
| 31 | 3,878,846 |
| 32 | 1,599,156 |
| 33 | 4,035,473 |
| 34 | 2,114,298 |
| 35 | 3,053,713 |
| 36 | 2,889,502 |
| 37 | 1,724,458 |
| 38 | 3,824,334 |
| 39 | 1,555,847 |
| 40 | 3,627,469 |
| 41 | 2,364,229 |
| 42 | 1,962,361 |
| 43 | 1,161,984 |
| 44 | 2,956,102 |
| 45 | 3,184,775 |
| 46 | 3,517,124 |
| 47 | 2,917,710 |
| 48 | 1,960,905 |
| 49 | 2,128,278 |
| 50 | 1,500,949 |

| | |
|---|---|
| 51 | 1,827,290 |
| 52 | 2,368,973 |
| 53 | 1,858,291 |
| 54 | 3,405,094 |
| 55 | 3,524,157 |
| 56 | 2,544,278 |
| 57 | 2,611,042 |
| 58 | 2,726,379 |
| 59 | 3,071,209 |
| 60 | 4,002,421 |
| 61 | 3,209,430 |
| 62 | 2,592,742 |
| 63 | 3,827,753 |
| 64 | 3,010,996 |
| 65 | 3,226,056 |
| 66 | 4,378,973 |
| 67 | 2,407,660 |
| 68 | 2,256,885 |
| 69 | 2,081,604 |
| 70 | 1,104,256 |
| 71 | 3,356,050 |
| 72 | 2,439,638 |
| 73 | 2,376,232 |
| 74 | 2,534,822 |
| 75 | 1,135,946 |
| 76 | 1,304,899 |
| 77 | 2,427,326 |
| 78 | 2,043,761 |
| 79 | 2,032,929 |
| 80 | 1,001,683 |
| 81 | 2,411,373 |
| 82 | 831,665 |
| 83 | 1,568,828 |
| 84 | 703,535 |
| 85 | 1,855,774 |
| 86 | 530,151 |
| 87 | 1,336,823 |
| 88 | 1,837,433 |
| 89 | 2,661,439 |
| 90 | 1,815,876 |
| 91 | 1,268,454 |
| 92 | 3,136,579 |
| 93 | 2,870,434 |
| Total | 235,208,642 |
| Average reads | 2,529,125 |

**Supplementary Table 2.** Depths of coverage for processed samples.

| Plant Number | Depth of coverage |
|---|---|
| 1 | 24.34x |
| 2 | 24.73x |
| 3 | 25.88x |
| 4 | 26.25x |
| 5 | 28.62x |
| 6 | 28.64x |
| 7 | 29.41x |
| 8 | 26.09x |
| 9 | 28.60x |
| 10 | 26.44x |
| 11 | 27.96x |
| 12 | 26.76x |
| 13 | 28.99x |
| 14 | 28.71x |
| 15 | 28.45x |
| 16 | 28.58x |
| 17 | 28.09x |
| 18 | 29.55x |
| 19 | 30.17x |
| 20 | 27.86x |
| 21 | 30.28x |
| 22 | 28.57x |
| 23 | 29.14x |
| 24 | 29.35x |
| 25 | 28.79x |
| 26 | 28.90x |
| 27 | 28.31x |
| 28 | 28.51x |
| 29 | 28.03x |
| 30 | 27.77x |
| 31 | 28.05x |
| 32 | 27.73x |
| 33 | 29.46x |
| 34 | 28.60x |
| 35 | 26.25x |
| 36 | 27.75x |
| 37 | 27.40x |
| 38 | 28.30x |
| 39 | 23.34x |
| 40 | 28.92x |
| 41 | 28.94x |
| 42 | 28.17x |
| 43 | 22.95x |
| 44 | 26.68x |
| 45 | 28.05x |
| 46 | 29.57x |
| 47 | 29.69x |
| 48 | 27.16x |
| 49 | 26.13x |
| 50 | 27.08x |
| 51 | 24.57x |
| 52 | 26.14x |
| 53 | 25.39x |

| | |
|---|---|
| 54 | 27.14x |
| 55 | 26.86x |
| 56 | 24.35x |
| 57 | 24.24x |
| 58 | 29.66x |
| 59 | 28.69x |
| 60 | 27.61x |
| 61 | 26.99x |
| 62 | 26.76x |
| 63 | 28.86x |
| 64 | 27.45x |
| 65 | 28.11x |
| 66 | 30.15x |
| 67 | 28.28x |
| 68 | 28.11x |
| 69 | 28.80x |
| 70 | 26.70x |
| 71 | 29.54x |
| 72 | 26.98x |
| 73 | 29.29x |
| 74 | 23.22x |
| 75 | 20.60x |
| 76 | 26.06x |
| 77 | 27.74x |
| 78 | 20.97x |
| 79 | 24.68x |
| 80 | 28.15x |
| 81 | 29.73x |
| 82 | 27.13x |
| 83 | 26.95x |
| 84 | 25.55x |
| 85 | 22.23x |
| 86 | 24.57x |
| 87 | 27.43x |
| 88 | 27.23x |
| 89 | 29.49x |
| 90 | 25.79x |
| 91 | 27.84x |
| 92 | 24.64x |
| 93 | 27.77x |
| Average | 27.32x |

**Supplementary Table 3.** The degrees of conservation of heterozygosity between each pair of the plants 35, 38, 40, 42, 43, 44, 45, 46, 47, 48, 49, 50, and 51.

| | 42 | 43 | 44 | 45 | 46 | 47 | 48 | 49 | 50 | 51 |
|---|---|---|---|---|---|---|---|---|---|---|
| 35 | 35.8 | 29.5 | 30.9 | 32.3 | 32.0 | 34.4 | 32.3 | 31.6 | 33.0 | 32.3 |
| 38 | 35.6 | 30.5 | 32.5 | 32.1 | 34.1 | 36.5 | 32.9 | 33.0 | 34.5 | 33.5 |
| 40 | 34.7 | 29.5 | 33.1 | 31.9 | 34.3 | 33.9 | 33.4 | 33.2 | 34.7 | 34.9 |
| 42 | | 50.4 | 50.9 | 50.4 | 52.6 | 55.9 | 50.6 | 52.9 | 55.0 | 53.0 |
| 43 | | | 52.9 | 51.0 | 51.9 | 56.9 | 52.6 | 54.9 | 61.9 | 59.3 |
| 44 | | | | 55.5 | 59.5 | 62.9 | 57.1 | 56.2 | 60.4 | 57.8 |
| 45 | | | | | 62.1 | 63.9 | 60.9 | 59.0 | 59.2 | 55.5 |
| 46 | | | | | | 68.9 | 62.3 | 62.3 | 58.8 | 58.6 |
| 47 | | | | | | | 66.4 | 66.1 | 66.7 | 62.9 |
| 48 | | | | | | | | 60.1 | 61.7 | 58.9 |
| 49 | | | | | | | | | 63.9 | 61.5 |
| 50 | | | | | | | | | | 67.6 |

**Supplementary Table 4.** Summary statistics for each group for all positions (variant and fixed).

| Population | Sites | Variant Sites | Poly Sites | %Poly Loci | Obs Het | Obs Homo | Exp Hete | Exp Homo | Nucleotide diversity | Inbreeding coefficient (Fis) |
|---|---|---|---|---|---|---|---|---|---|---|
| H | 242383 | 2287 | 751 | 0.310 | 0.00265 | 0.99735 | 0.00147 | 0.99853 | 0.00256 | -0.000130 |
| B | 402530 | 4085 | 1378 | 0.342 | 0.00312 | 0.99688 | 0.00165 | 0.99835 | 0.00258 | -0.000810 |
| F | 463469 | 4967 | 3598 | 0.776 | 0.00322 | 0.99678 | 0.00289 | 0.99711 | 0.00323 | 0.000200 |
| E | 464236 | 4973 | 4095 | 0.882 | 0.00319 | 0.99681 | 0.00303 | 0.99697 | 0.00322 | 0.000540 |
| C | 460079 | 4923 | 2973 | 0.646 | 0.00320 | 0.99680 | 0.00253 | 0.99747 | 0.00298 | -0.000330 |
| A1 | 463672 | 4964 | 3433 | 0.740 | 0.00347 | 0.99653 | 0.00277 | 0.99723 | 0.00312 | -0.000540 |
| A2 | 450414 | 4802 | 2142 | 0.476 | 0.00362 | 0.99638 | 0.00209 | 0.99791 | 0.00235 | -0.00232 |
| D | 464287 | 4973 | 4593 | 0.989 | 0.00324 | 0.99676 | 0.00321 | 0.99679 | 0.00331 | 0.000840 |

Poly – Polymorphic, Obs Hete – Observed Heterozygosity, Obs Homo – Observed Homozygosity, Exp Hete – Expected Heterozygosity, Exp Homo – Expected Homozygosity.

**Supplementary Table 5.** F3-statistics. The table was sorted by the values of the f3-statistic.

| Outgroup; X, Y | F3-statistic | Standard error | Z-score |
|---|---|---|---|
| A1;C,A2 | -0.00942 | 0.00118 | -7.97 |
| A1;E,A2 | -0.00931 | 0.00133 | -6.99 |
| A1;A2,D | -0.00877 | 0.00133 | -6.61 |
| A1;H,A2 | -0.00782 | 0.00184 | -4.25 |
| A1;B,A2 | -0.00738 | 0.00169 | -4.37 |
| A1;F,A2 | -0.00682 | 0.00134 | -5.09 |
| D;B,E | 0.00452 | 0.00121 | 3.75 |
| D;C,A2 | 0.00467 | 0.00120 | 3.88 |
| D;C,A1 | 0.00532 | 0.00140 | 3.79 |
| D;F,E | 0.00552 | 0.000834 | 6.62 |
| D;E,A2 | 0.00553 | 0.00106 | 5.20 |
| F;C,A2 | 0.00572 | 0.00127 | 4.50 |
| D;B,C | 0.00576 | 0.00182 | 3.16 |
| D;F,A1 | 0.00595 | 0.00152 | 3.92 |
| D;E,A1 | 0.00606 | 0.00119 | 5.10 |
| D;B,A1 | 0.00647 | 0.000929 | 6.97 |
| D;B,F | 0.00654 | 0.00108 | 6.08 |
| D;F,C | 0.00683 | 0.00105 | 6.54 |
| D;E,C | 0.00717 | 0.00112 | 6.40 |
| F;H,D | 0.00728 | 0.000853 | 8.53 |
| D;B,A2 | 0.00786 | 0.00128 | 6.14 |
| F;A2,D | 0.00788 | 0.000818 | 9.63 |
| F;E,A2 | 0.00790 | 0.000926 | 8.53 |
| D;F,A2 | 0.00790 | 0.00109 | 7.27 |
| F;B,C | 0.00817 | 0.00200 | 4.08 |
| F;B,E | 0.00825 | 0.00186 | 4.43 |
| F;C,A1 | 0.00832 | 0.000860 | 9.68 |
| D;H,A1 | 0.00838 | 0.00213 | 3.93 |
| D;H,E | 0.00848 | 0.00113 | 7.52 |
| D;H,F | 0.00850 | 0.000411 | 20.7 |
| F;H,A2 | 0.00871 | 0.00145 | 6.00 |
| D;H,C | 0.00881 | 0.000746 | 11.8 |
| F;C,D | 0.00895 | 0.000650 | 13.8 |
| E;H,D | 0.00903 | 0.00104 | 8.68 |
| F;B,A2 | 0.00921 | 0.00163 | 5.66 |
| F;B,D | 0.00924 | 0.00152 | 6.09 |
| F;H,C | 0.00926 | 0.000822 | 11.3 |
| D;H,A2 | 0.00933 | 0.00157 | 5.95 |
| E;C,A2 | 0.00949 | 0.00105 | 9.00 |
| E;C,A1 | 0.00960 | 0.00139 | 6.89 |
| F;H,A1 | 0.00971 | 0.00152 | 6.40 |
| D;H,B | 0.00977 | 0.00208 | 4.70 |
| F;B,A1 | 0.00977 | 0.00158 | 6.17 |
| F;A1,D | 0.00984 | 0.00106 | 9.31 |
| F;H,E | 0.0102 | 0.00107 | 9.59 |
| F;E,D | 0.0103 | 0.00137 | 7.50 |
| E;C,D | 0.0103 | 0.00105 | 9.83 |
| F;E,A1 | 0.0104 | 0.00106 | 9.80 |
| F;H,B | 0.0105 | 0.00295 | 3.56 |
| F;E,C | 0.0106 | 0.000824 | 12.9 |

| | | | |
|---|---|---|---|
| E;H,C | 0.0107 | 0.00180 | 5.95 |
| E;H,A1 | 0.0113 | 0.00171 | 6.63 |
| E;A1,D | 0.0115 | 0.000938 | 12.2 |
| E;B,C | 0.0116 | 0.00193 | 6.00 |
| E;F,C | 0.0117 | 0.00178 | 6.57 |
| E;F,A1 | 0.0119 | 0.00133 | 8.93 |
| E;A2,D | 0.0120 | 0.000528 | 22.7 |
| E;F,D | 0.0120 | 0.000434 | 27.6 |
| E;H,F | 0.0120 | 0.00141 | 8.53 |
| E;H,A2 | 0.0128 | 0.00136 | 9.44 |
| E;B,D | 0.0130 | 0.00120 | 10.8 |
| E;B,A1 | 0.0134 | 0.00129 | 10.4 |
| E;B,F | 0.0140 | 0.000979 | 14.3 |
| E;H,B | 0.0143 | 0.00293 | 4.86 |
| E;F,A2 | 0.0144 | 0.00120 | 12.0 |
| E;B,A2 | 0.0153 | 0.00141 | 10.9 |
| A1;H,D | 0.0177 | 0.00184 | 9.64 |
| H;B,E | 0.0178 | 0.00297 | 6.01 |
| H;C,A2 | 0.0181 | 0.00201 | 8.99 |
| A1;B,E | 0.0181 | 0.00139 | 13.0 |
| H;B,C | 0.0187 | 0.00345 | 5.42 |
| H;E,A2 | 0.0193 | 0.00348 | 5.53 |
| A1;F,E | 0.0196 | 0.00111 | 17.6 |
| A1;B,D | 0.0196 | 0.00190 | 10.4 |
| H;C,A1 | 0.0197 | 0.00294 | 6.68 |
| H;B,F | 0.0198 | 0.00380 | 5.21 |
| H;B,A1 | 0.0199 | 0.00350 | 5.68 |
| A1;E,D | 0.0200 | 0.00139 | 14.4 |
| A1;B,C | 0.0201 | 0.00245 | 8.19 |
| H;F,E | 0.0201 | 0.00264 | 7.61 |
| A1;H,E | 0.0201 | 0.00148 | 13.6 |
| A1;F,D | 0.0201 | 0.00141 | 14.2 |
| A1;B,F | 0.0202 | 0.00173 | 11.7 |
| A1;H,F | 0.0203 | 0.00245 | 8.28 |
| H;B,A2 | 0.0203 | 0.00316 | 6.44 |
| H;F,A1 | 0.0206 | 0.00337 | 6.11 |
| H;E,A1 | 0.0207 | 0.00360 | 5.77 |
| A1;C,D | 0.0208 | 0.00130 | 16.0 |
| A1;H,B | 0.0210 | 0.00351 | 5.99 |
| H;F,C | 0.0211 | 0.00202 | 10.4 |
| A1;H,C | 0.0212 | 0.00227 | 9.34 |
| H;E,C | 0.0214 | 0.00224 | 9.57 |
| H;F,A2 | 0.0216 | 0.00298 | 7.26 |
| A1;F,C | 0.0217 | 0.00146 | 14.8 |
| H;B,D | 0.0218 | 0.00333 | 6.53 |
| A1;E,C | 0.0219 | 0.00112 | 19.6 |
| H;A2,D | 0.0222 | 0.00337 | 6.58 |
| H;C,D | 0.0227 | 0.00241 | 9.44 |
| H;F,D | 0.0230 | 0.00265 | 8.69 |
| H;E,D | 0.0231 | 0.00240 | 9.62 |
| H;A1,D | 0.0232 | 0.00415 | 5.58 |
| C;H,D | 0.0232 | 0.00202 | 11.5 |
| C;F,E | 0.0236 | 0.00160 | 14.7 |
| C;B,E | 0.0236 | 0.000759 | 31.2 |

| | | | |
|---|---|---|---|
| C;H,E | 0.0246 | 0.00143 | 17.2 |
| C;E,D | 0.0249 | 0.00136 | 18.3 |
| C;H,F | 0.0249 | 0.00158 | 15.8 |
| C;F,D | 0.0252 | 0.00172 | 14.7 |
| C;E,A1 | 0.0256 | 0.00154 | 16.6 |
| C;E,A2 | 0.0257 | 0.00211 | 12.2 |
| C;F,A1 | 0.0259 | 0.00210 | 12.3 |
| C;B,F | 0.0260 | 0.00226 | 11.5 |
| C;B,D | 0.0263 | 0.00118 | 22.2 |
| C;H,A1 | 0.0263 | 0.00220 | 11.9 |
| C;A1,D | 0.0267 | 0.00184 | 14.5 |
| C;H,B | 0.0272 | 0.00276 | 9.89 |
| C;A2,D | 0.0274 | 0.00171 | 16.1 |
| C;B,A1 | 0.0275 | 0.00124 | 22.2 |
| C;H,A2 | 0.0279 | 0.00277 | 10.1 |
| C;F,A2 | 0.0285 | 0.00210 | 13.5 |
| C;B,A2 | 0.0295 | 0.00213 | 13.9 |
| A2;F,A1 | 0.0325 | 0.00145 | 22.4 |
| A2;B,A1 | 0.0331 | 0.00151 | 21.8 |
| A2;H,A1 | 0.0335 | 0.00144 | 23.2 |
| A2;A1,D | 0.0345 | 0.00139 | 24.8 |
| D;A1,A2 | 0.0349 | 0.00193 | 18.1 |
| A2;E,A1 | 0.0350 | 0.00153 | 22.9 |
| A2;C,A1 | 0.0351 | 0.00113 | 31.1 |
| B;H,D | 0.0364 | 0.00273 | 13.3 |
| F;A1,A2 | 0.0368 | 0.000855 | 43.1 |
| B;C,A2 | 0.0372 | 0.00242 | 15.3 |
| B;H,A2 | 0.0378 | 0.00365 | 10.4 |
| B;H,A1 | 0.0383 | 0.00371 | 10.3 |
| B;A2,D | 0.0383 | 0.00247 | 15.5 |
| B;H,F | 0.0383 | 0.00301 | 12.7 |
| B;F,A1 | 0.0391 | 0.00245 | 15.9 |
| B;C,A1 | 0.0392 | 0.00174 | 22.6 |
| B;E,A2 | 0.0393 | 0.00310 | 12.7 |
| B;H,C | 0.0394 | 0.00322 | 12.2 |
| B;F,D | 0.0396 | 0.00223 | 17.8 |
| B;F,A2 | 0.0396 | 0.00362 | 10.9 |
| B;A1,D | 0.0397 | 0.00120 | 33.2 |
| B;H,E | 0.0403 | 0.00290 | 13.9 |
| B;C,D | 0.0404 | 0.00190 | 21.2 |
| B;F,E | 0.0406 | 0.00236 | 17.2 |
| B;F,C | 0.0407 | 0.00270 | 15.1 |
| E;A1,A2 | 0.0408 | 0.00139 | 29.4 |
| B;E,A1 | 0.0412 | 0.00174 | 23.7 |
| B;E,D | 0.0416 | 0.00201 | 20.7 |
| B;E,C | 0.0430 | 0.00219 | 19.7 |
| H;A1,A2 | 0.0487 | 0.00423 | 11.5 |
| C;A1,A2 | 0.0569 | 0.00284 | 20.0 |
| A2;H,D | 0.0600 | 0.00166 | 36.1 |
| A2;B,F | 0.0601 | 0.00239 | 25.1 |
| A2;B,E | 0.0605 | 0.00295 | 20.5 |
| A2;H,F | 0.0606 | 0.00112 | 54.2 |
| A2;F,E | 0.0614 | 0.00219 | 27.9 |
| A2;F,D | 0.0614 | 0.00216 | 28.4 |

| | | | |
|---|---|---|---|
| A2;B,D | 0.0615 | 0.00270 | 22.8 |
| A2;H,B | 0.0619 | 0.00221 | 27.9 |
| A2;B,C | 0.0626 | 0.00389 | 16.1 |
| A2;H,E | 0.0630 | 0.00144 | 43.8 |
| A2;F,C | 0.0636 | 0.00203 | 31.4 |
| A2;E,D | 0.0638 | 0.00226 | 28.2 |
| A2;H,C | 0.0641 | 0.00192 | 33.4 |
| A2;C,D | 0.0647 | 0.00225 | 28.7 |
| A2;E,C | 0.0663 | 0.00241 | 27.5 |
| B;A1,A2 | 0.0667 | 0.00288 | 23.1 |

**Supplementary Table 6.** F4-statistics. The table was sorted by the values of the Z-score.

| Outgroup, X; Y, Z | F4-statistic | Standard error | Z-score |
|---|---|---|---|
| F,A2;A1,D | -0.0289 | 0.000912 | -31.7 |
| F,A1;A2,D | -0.0270 | 0.00108 | -25.0 |
| C,A2;A1,D | -0.0296 | 0.00128 | -23.2 |
| E,A2;A1,D | -0.0288 | 0.00125 | -23.0 |
| B,A2;A1,D | -0.0284 | 0.00145 | -19.6 |
| E,A1;A2,D | -0.0293 | 0.00153 | -19.2 |
| C,A1;A2,D | -0.0302 | 0.00164 | -18.4 |
| H,A2;A1,D | -0.0265 | 0.00165 | -16.1 |
| H,A1;A2,D | -0.0255 | 0.00161 | -15.9 |
| B,A1;A2,D | -0.0270 | 0.00244 | -11.1 |
| F,C;E,A2 | -0.00488 | 0.000675 | -7.24 |
| F,E;A1,A2 | -0.00249 | 0.000408 | -6.09 |
| B,C;E,A1 | -0.00380 | 0.00100 | -3.81 |
| F,D;A1,A2 | -0.00195 | 0.000557 | -3.51 |
| F,C;A1,A2 | -0.00260 | 0.000781 | -3.32 |
| B,C;E,A2 | -0.00584 | 0.00182 | -3.21 |
| F,C;E,A1 | -0.00228 | 0.000718 | -3.18 |
| F,A1;E,C | -0.00206 | 0.000756 | -2.73 |
| B,D;E,A2 | -0.00334 | 0.00129 | -2.59 |
| B,C;E,D | -0.00265 | 0.00104 | -2.55 |
| F,C;E,D | -0.00165 | 0.000656 | -2.52 |
| H,A2;F,C | -0.00354 | 0.00148 | -2.39 |
| B,C;F,A2 | -0.00348 | 0.00146 | -2.38 |
| H,C;E,A2 | -0.00334 | 0.00148 | -2.25 |
| F,E;C,A2 | -0.00271 | 0.00123 | -2.20 |
| F,D;E,A2 | -0.00238 | 0.00110 | -2.17 |
| H,C;F,A2 | -0.00299 | 0.00146 | -2.04 |
| B,C;A1,A2 | -0.00203 | 0.00107 | -1.90 |
| F,D;C,A2 | -0.00107 | 0.00062 | -1.73 |
| H,A2;F,E | -0.00235 | 0.00137 | -1.72 |
| B,D;E,A1 | -0.00195 | 0.00120 | -1.62 |
| B,A1;E,C | -0.00198 | 0.00122 | -1.62 |
| H,C;E,A1 | -0.00175 | 0.00115 | -1.52 |
| F,A2;E,C | -0.00217 | 0.00144 | -1.51 |
| B,E;C,A2 | -0.00375 | 0.00290 | -1.29 |
| H,B;F,E | -0.00200 | 0.00161 | -1.24 |
| H,A2;B,C | -0.00224 | 0.00191 | -1.18 |
| B,A1;E,D | -0.00154 | 0.00133 | -1.16 |
| H,C;A1,A2 | -0.00160 | 0.00141 | -1.13 |
| H,A1;E,C | -0.00108 | 0.000978 | -1.10 |
| B,E;A1,A2 | -0.00192 | 0.00178 | -1.08 |
| B,A2;E,C | -0.00209 | 0.00196 | -1.07 |
| F,D;E,C | -0.00131 | 0.00124 | -1.06 |
| B,E;C,A1 | -0.00182 | 0.00176 | -1.03 |
| B,F;E,A1 | -0.00152 | 0.00147 | -1.03 |
| B,D;F,A2 | -0.00132 | 0.00129 | -1.03 |
| B,C;F,A1 | -0.00145 | 0.00144 | -1.01 |
| B,D;C,A2 | -0.00210 | 0.00222 | -0.950 |
| B,D;E,C | -0.00123 | 0.00131 | -0.945 |
| B,F;E,D | -0.000993 | 0.00108 | -0.923 |

| | | | |
|---|---|---|---|
| B,A2;F,D | -0.00136 | 0.00149 | -0.911 |
| H,E;A1,A2 | -0.00149 | 0.00169 | -0.878 |
| B,A2;F,C | -0.00245 | 0.00284 | -0.862 |
| H,E;C,A2 | -0.00215 | 0.00253 | -0.851 |
| H,C;F,A1 | -0.00139 | 0.00168 | -0.825 |
| H,A1;F,C | -0.000941 | 0.00116 | -0.809 |
| B,E;F,A2 | -0.00131 | 0.00164 | -0.801 |
| B,F;C,A1 | -0.00160 | 0.00208 | -0.769 |
| B,E;C,D | -0.00141 | 0.00188 | -0.750 |
| B,D;A1,A2 | -0.00139 | 0.00187 | -0.745 |
| B,A2;E,D | -0.00101 | 0.00136 | -0.742 |
| B,F;C,D | -0.00107 | 0.00155 | -0.692 |
| H,A2;E,C | -0.00119 | 0.00183 | -0.649 |
| F,A1;E,D | -0.000548 | 0.000892 | -0.615 |
| H,D;E,A2 | -0.000845 | 0.00148 | -0.572 |
| H,D;A1,A2 | -0.000952 | 0.00194 | -0.491 |
| H,B;F,C | -0.00109 | 0.00227 | -0.480 |
| H,D;F,C | -0.000311 | 0.000657 | -0.474 |
| B,F;E,A2 | -0.000957 | 0.00210 | -0.457 |
| B,D;C,A1 | -0.000715 | 0.00157 | -0.455 |
| H,D;F,A2 | -0.000825 | 0.00187 | -0.441 |
| B,F;C,A2 | -0.00104 | 0.00244 | -0.425 |
| H,A2;B,E | -0.00105 | 0.00251 | -0.417 |
| H,E;F,A2 | -0.000811 | 0.00208 | -0.390 |
| H,D;E,C | -0.000331 | 0.000912 | -0.363 |
| H,E;C,A1 | -0.000668 | 0.00187 | -0.357 |
| H,D;C,A2 | -0.000514 | 0.00178 | -0.289 |
| F,D;E,A1 | -0.000429 | 0.00152 | -0.282 |
| H,F;C,A1 | -0.000447 | 0.00175 | -0.256 |
| F,E;C,D | -0.000339 | 0.00148 | -0.230 |
| H,C;B,A2 | -0.000648 | 0.00322 | -0.201 |
| B,A2;F,E | -0.000352 | 0.00178 | -0.198 |
| B,C;F,D | -0.000292 | 0.00159 | -0.184 |
| F,E;C,A1 | -0.000221 | 0.00122 | -0.181 |
| F,E;A1,D | -0.000119 | 0.00105 | -0.114 |
| E,C;A1,A2 | -0.000110 | 0.00104 | -0.106 |
| H,A1;B,C | -0.000205 | 0.00237 | -0.0864 |
| F,A2;E,D | -0.0000137 | 0.000565 | -0.0243 |
| B,F;A2,D | -0.0000360 | 0.00191 | -0.0189 |
| H,D;F,E | 0.0000199 | 0.00121 | 0.0164 |
| H,B;F,A1 | 0.0000609 | 0.00218 | 0.0279 |
| H,D;E,A1 | 0.000107 | 0.00224 | 0.0478 |
| B,D;F,A1 | 0.0000663 | 0.00129 | 0.0514 |
| H,D;F,A1 | 0.000127 | 0.00216 | 0.0588 |
| B,A1;F,C | 0.000151 | 0.00250 | 0.0604 |
| B,F;E,C | 0.0000805 | 0.00119 | 0.0677 |
| H,F;B,E | 0.000260 | 0.00323 | 0.0805 |
| H,A1;F,E | 0.000139 | 0.00146 | 0.0950 |
| H,D;B,A2 | 0.000438 | 0.00286 | 0.153 |
| H,D;C,A1 | 0.000438 | 0.00210 | 0.208 |
| H,F;B,A1 | 0.000797 | 0.00369 | 0.216 |
| H,B;F,A2 | 0.000498 | 0.00210 | 0.238 |
| H,C;B,A1 | 0.000948 | 0.00362 | 0.262 |
| B,A1;C,D | 0.000442 | 0.00160 | 0.276 |

| | | | |
|---|---|---|---|
| H,F;C,A2 | 0.000553 | 0.00179 | 0.309 |
| H,A1;B,F | 0.000736 | 0.00229 | 0.321 |
| H,A1;B,E | 0.000875 | 0.00260 | 0.336 |
| B,A1;F,D | 0.000593 | 0.00173 | 0.343 |
| H,D;B,A1 | 0.00139 | 0.00392 | 0.355 |
| B,F;A1,A2 | 0.000563 | 0.00154 | 0.367 |
| H,D;B,C | 0.000952 | 0.00256 | 0.372 |
| B,D;F,C | 0.000781 | 0.00206 | 0.379 |
| H,C;F,E | 0.000359 | 0.000930 | 0.387 |
| H,F;B,C | 0.00124 | 0.00316 | 0.394 |
| H,E;F,A1 | 0.000675 | 0.00169 | 0.400 |
| H,B;C,A1 | 0.00115 | 0.00282 | 0.408 |
| B,E;A1,D | 0.000408 | 0.00100 | 0.409 |
| H,E;B,A2 | 0.00145 | 0.00320 | 0.452 |
| H,D;B,E | 0.00128 | 0.00274 | 0.469 |
| H,A2;F,D | 0.000605 | 0.00124 | 0.486 |
| H,B;A1,A2 | 0.000437 | 0.00087 | 0.504 |
| H,F;B,A2 | 0.00180 | 0.00342 | 0.526 |
| H,F;E,A1 | 0.000537 | 0.00102 | 0.528 |
| H,B;C,A2 | 0.00159 | 0.00301 | 0.528 |
| H,D;B,F | 0.00126 | 0.00232 | 0.544 |
| B,E;F,A1 | 0.000615 | 0.00106 | 0.578 |
| B,A2;C,D | 0.00109 | 0.00177 | 0.615 |
| H,A2;B,F | 0.00130 | 0.00207 | 0.628 |
| H,F;A1,A2 | 0.00100 | 0.00158 | 0.633 |
| E,C;A2,D | 0.000859 | 0.00136 | 0.633 |
| B,F;A1,D | 0.000527 | 0.000780 | 0.676 |
| C,D;A1,A2 | 0.000644 | 0.000921 | 0.700 |
| H,E;B,A1 | 0.00293 | 0.00366 | 0.801 |
| H,B;E,C | 0.000904 | 0.00112 | 0.807 |
| E,D;A1,A2 | 0.000535 | 0.000602 | 0.888 |
| F,C;A1,D | 0.000630 | 0.000690 | 0.913 |
| E,C;A1,D | 0.000749 | 0.000809 | 0.926 |
| H,C;B,E | 0.00270 | 0.00289 | 0.933 |
| H,F;E,C | 0.000984 | 0.00105 | 0.935 |
| H,E;B,F | 0.00226 | 0.00236 | 0.956 |
| H,F;A2,D | 0.00143 | 0.00144 | 0.992 |
| H,C;B,F | 0.00234 | 0.00234 | 1.00 |
| H,A2;B,D | 0.00190 | 0.00191 | 1.00 |
| H,F;E,A2 | 0.00154 | 0.00147 | 1.05 |
| H,B;E,A1 | 0.00206 | 0.00196 | 1.05 |
| F,D;C,A1 | 0.000886 | 0.000842 | 1.05 |
| E,D;C,A1 | 0.00111 | 0.00104 | 1.06 |
| B,E;F,C | 0.00244 | 0.00228 | 1.07 |
| H,B;A2,D | 0.00147 | 0.00133 | 1.10 |
| H,F;B,D | 0.00323 | 0.00282 | 1.14 |
| B,E;F,D | 0.00102 | 0.000890 | 1.15 |
| B,A1;F,E | 0.00213 | 0.00184 | 1.16 |
| E,A1;C,D | 0.00186 | 0.00159 | 1.17 |
| B,C;F,E | 0.00236 | 0.00201 | 1.17 |
| H,A1;B,D | 0.00329 | 0.00277 | 1.19 |
| H,B;E,A2 | 0.00249 | 0.00208 | 1.20 |
| B,E;A2,D | 0.00233 | 0.00185 | 1.26 |
| H,E;F,C | 0.00134 | 0.00103 | 1.30 |

| | | | |
|---|---|---|---|
| H,E;C,D | 0.00164 | 0.00123 | 1.33 |
| H,C;E,D | 0.00131 | 0.000979 | 1.34 |
| H,C;B,D | 0.00401 | 0.00290 | 1.38 |
| H,E;B,C | 0.00360 | 0.00259 | 1.39 |
| H,B;C,D | 0.00306 | 0.00212 | 1.44 |
| H,B;A1,D | 0.00190 | 0.00131 | 1.45 |
| H,E;A1,D | 0.00231 | 0.00157 | 1.47 |
| B,C;A1,D | 0.00116 | 0.00077 | 1.51 |
| H,C;F,D | 0.00167 | 0.00110 | 1.52 |
| H,F;C,D | 0.00198 | 0.00129 | 1.54 |
| H,A1;F,D | 0.00256 | 0.00162 | 1.58 |
| E,D;C,A2 | 0.00164 | 0.00101 | 1.62 |
| H,B;F,D | 0.00196 | 0.00119 | 1.65 |
| H,A1;E,D | 0.00242 | 0.00146 | 1.66 |
| H,E;B,D | 0.00524 | 0.00258 | 2.03 |
| B,D;F,E | 0.00202 | 0.00098 | 2.05 |
| F,A2;C,D | 0.00216 | 0.00100 | 2.16 |
| B,C;A2,D | 0.00319 | 0.00147 | 2.17 |
| H,F;A1,D | 0.00243 | 0.00107 | 2.26 |
| H,A1;C,D | 0.00350 | 0.00153 | 2.29 |
| H,A2;E,D | 0.00295 | 0.00128 | 2.31 |
| F,E;A2,D | 0.00237 | 0.00100 | 2.36 |
| E,A2;C,D | 0.00250 | 0.00105 | 2.39 |
| H,E;A2,D | 0.00380 | 0.00149 | 2.55 |
| F,A1;C,D | 0.00152 | 0.00059 | 2.57 |
| H,A2;C,D | 0.00414 | 0.00149 | 2.77 |
| F,C;A2,D | 0.00323 | 0.00114 | 2.83 |
| H,C;A1,D | 0.00306 | 0.00103 | 2.98 |
| H,E;F,D | 0.00299 | 0.000992 | 3.01 |
| H,F;E,D | 0.00297 | 0.000885 | 3.35 |
| H,B;E,D | 0.00396 | 0.00113 | 3.51 |
| H,C;A2,D | 0.00466 | 0.00101 | 4.60 |
| B,A1;C,A2 | 0.0274 | 0.00303 | 9.04 |
| H,A2;B,A1 | 0.0284 | 0.00296 | 9.61 |
| B,A2;C,A1 | 0.0295 | 0.00288 | 10.2 |
| B,A1;E,A2 | 0.0255 | 0.00227 | 11.2 |
| H,A1;B,A2 | 0.0288 | 0.00253 | 11.4 |
| H,A2;C,A1 | 0.0306 | 0.00238 | 12.9 |
| B,A2;E,A1 | 0.0274 | 0.00202 | 13.5 |
| H,A2;F,A1 | 0.0271 | 0.00186 | 14.5 |
| B,A2;F,A1 | 0.0270 | 0.00160 | 16.9 |
| H,A1;C,A2 | 0.0290 | 0.00169 | 17.2 |
| H,A1;F,A2 | 0.0281 | 0.00161 | 17.5 |
| H,A2;E,A1 | 0.0294 | 0.00162 | 18.2 |
| B,A1;F,A2 | 0.0276 | 0.00142 | 19.4 |
| E,A1;C,A2 | 0.0312 | 0.00155 | 20.1 |
| H,A1;E,A2 | 0.0280 | 0.00135 | 20.7 |
| E,A2;C,A1 | 0.0313 | 0.00144 | 21.8 |
| F,A2;C,A1 | 0.0311 | 0.00138 | 22.5 |
| F,A1;C,A2 | 0.0285 | 0.00106 | 27.0 |
| F,A1;E,A2 | 0.0264 | 0.000953 | 27.7 |
| F,A2;E,A1 | 0.0289 | 0.000919 | 31.5 |

**Supplementary Figure 1**. Cross-validation (cv) error against the number of ancestors (*K*) from *K*=1-12 and *K*=1,8,9,10,11,12 admixture plots.

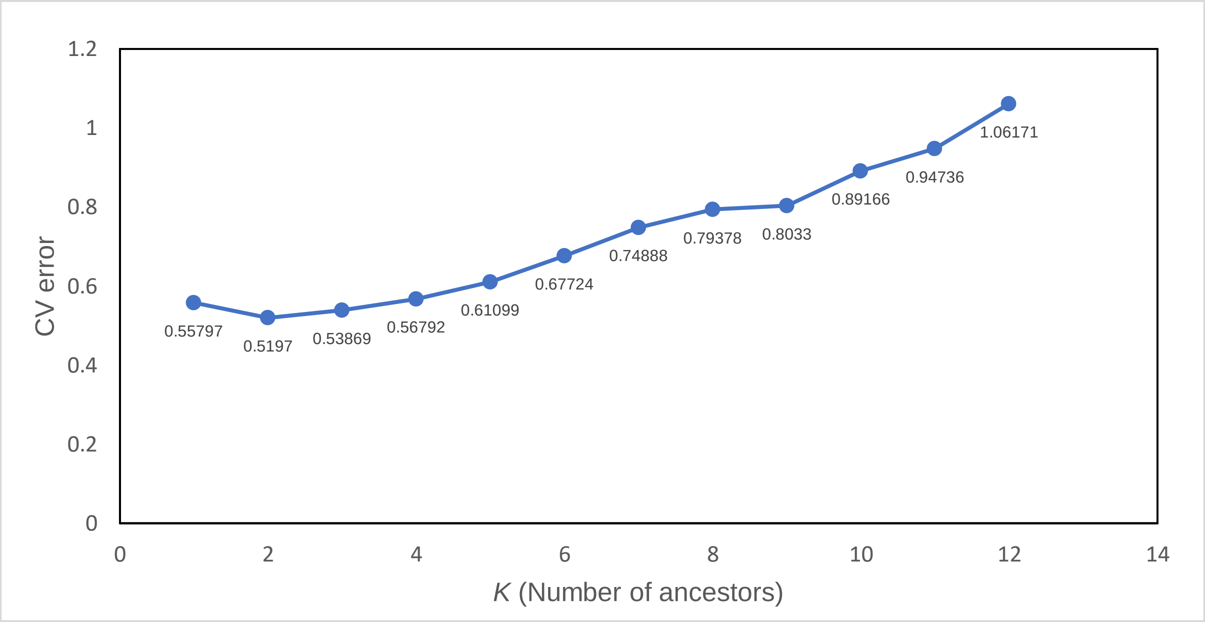


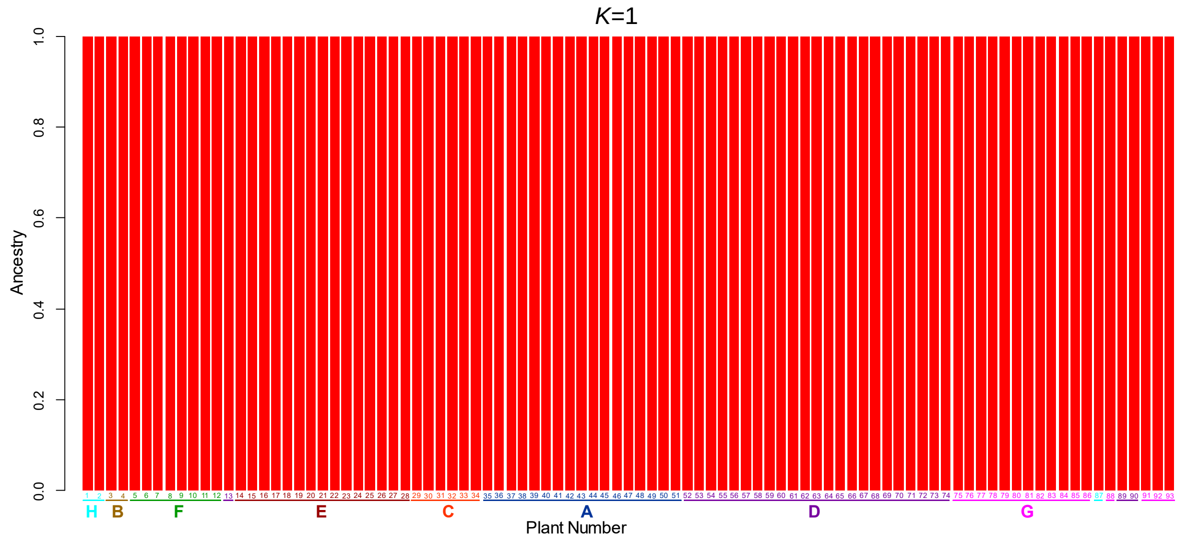


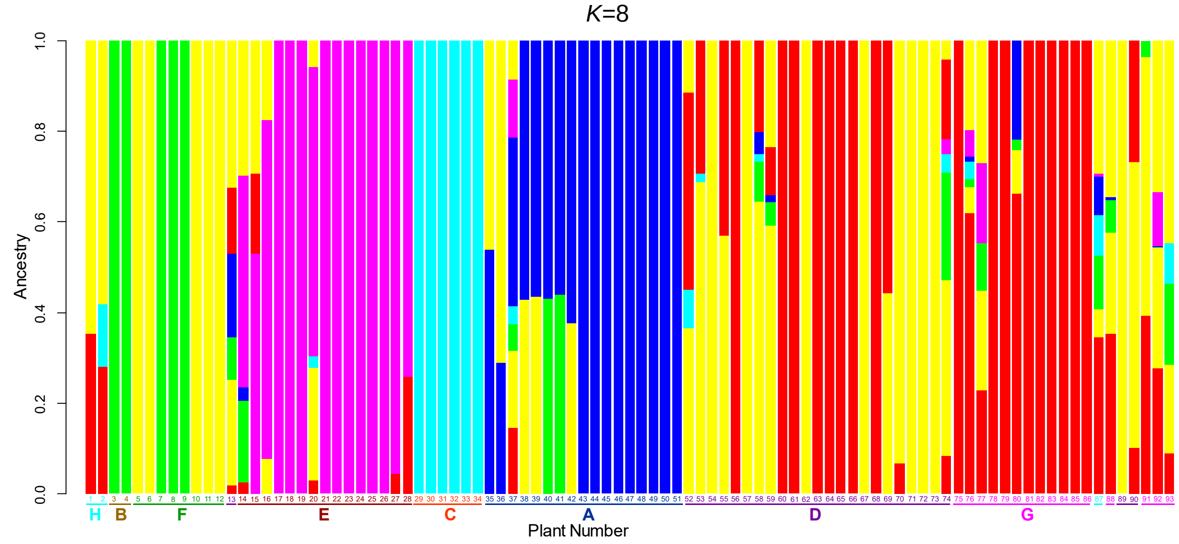

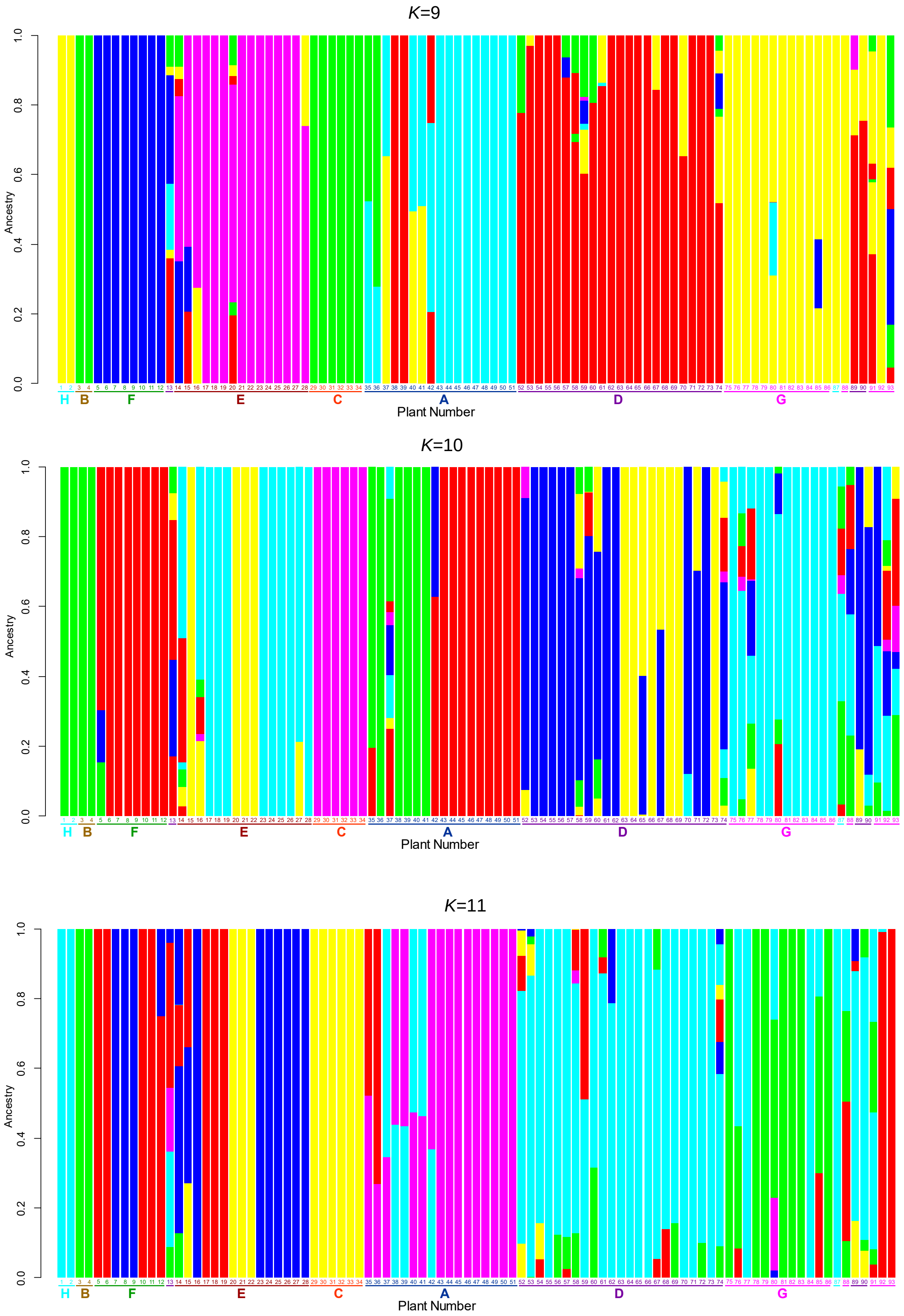
K=9
Ancestry
0.0
0.2
0.4
0.6
0.8
1.0
H
B
F
E
C
A
D
G
Plant Number
K=10
Ancestry
0.0
0.2
0.4
0.6
0.8
1.0
H
B
F
E
C
A
D
G
Plant Number
K=11
Ancestry
0.0
0.2
0.4
0.6
0.8
1.0
H
B
F
E
C
A
D
G
Plant Number

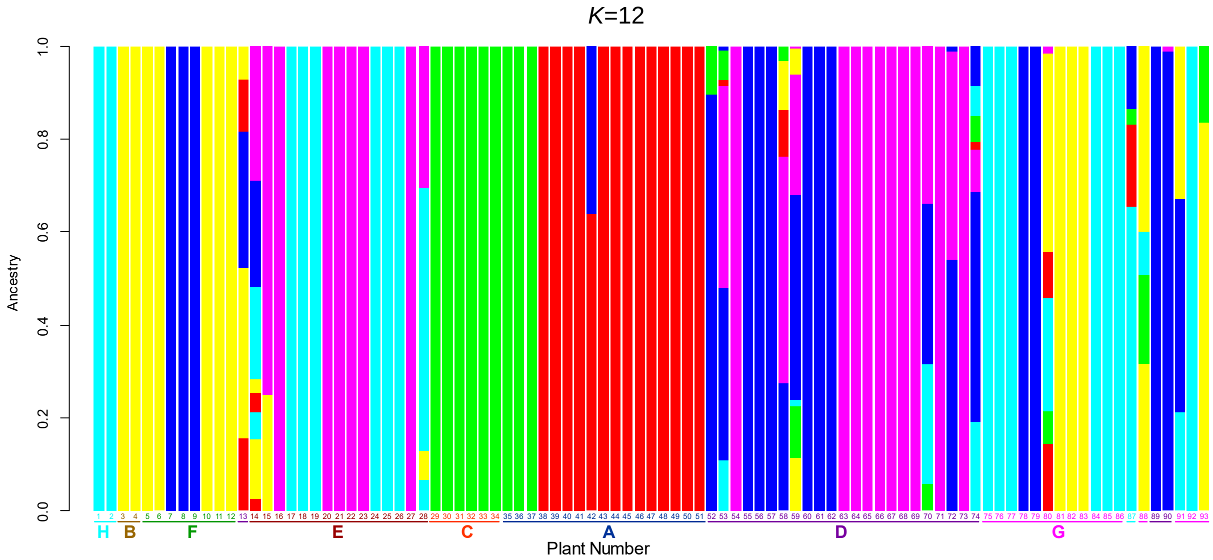

K=12
Ancestry
0.0
0.2
0.4
0.6
0.8
1.0
Plant Number
H
B
F
E
C
A
D
G

**Supplementary Figure 2.** The pairwise alignments between plants 8 and 9.

```
CLUSTAL 2.1 multiple sequence alignment (clustal format)

8               YGGCKGTWTMCYWAMYKYKSRWYCRKWKCARRRYKYRYYRMKMKSAMWRRMYYKRWKYYY
9               YCRSKRYWKMTYWRMCKYKSRWYTRKWKGGRRRTGCAYYRMKMKSGMWRRMYYKRWKYYY
                *  .*  *.* ** * ******* **** .***    ********.**************

8               RRRKSMWTGTRYYYKMRKKTTARYKYWYYKYYYWYYMRYWRWMKKKRRRMYRYRKSRRKM
9               RRRTSMWYRWRYYYKMRGKAGGRYKYWYTKYYYWYYMRYWRWMKKKRGGATRYRKSRRKM
                ***.***   ******* *: .****** ******************    *********

8               MYMWRMSYKSRRYKRYYYYYYRRYYSYWWCRYSYRYAMWRYWYRYYMYRSRWKSWTWRRM
9               MYMWRMGTKSRRYKRYYYYYYRRYCGTATYGTGYRYCMAACWYRYYMYRSRWKSAWWRRM
                ******. **************** .      .***.*   *************  ****

8               WYRYMMWRYWMRYRWRRYYRKWRYKTRYRRWMRYMSRAYYMTCRWYCTGYYYRCTRWCYR
9               WYRYMMWRYWMRYRWRRYYRKWRYTCAYRRWMRYMSGGYYMAYRWYMWKCYYGYGRAGYR
                ************************.  ********* .***: ***    **   *  **

8               WSKYWKKRTMKMYRYMRMYMRKYYGWRKMMRGKKRYYYMKYKMYYRYYYRGAYYTRGYRR
9               TCKYWKKGYMKMYRYARMYMRKTYAWRKMMRRKKRCTCCKYKAYYRYTYRTGYYGGACRR
                 .*****  ****** ****** *.****** ***    *** **** ** .**  . **

8               YRYYKYWRRYWMCGYGCYRYYMKKWWMWMYKSWYCWRTWRWRYCMYYYYKRRMMYTWRRR
9               YRYYKYWRRYWCYKCRYYRYTMKKWWMAMYKSWYMWGYWRTACYMYYYYKRRMMYYWRRR
                ***********      *** ****** ****** *  **    *********** ****

8               RMYRWRMRYKYKYRRYCRKARRWWYYMRKRMMSSYRMWKMKRRKMWYYMYRRWSRYCSWK
9               RMYRWRMRYKYKYRATARKCARWWCTARKRMMSSYRMWKMKRRKCAYCMCARWSRTYSAT
                **************  .**. ***   *****************  * *  ****  * .

8               YWWRYMRRSWRMTTTYYYRYYRWWRYYKYRCRGYTAAGWYWYCMYATSKRTYYRRWMSYC
9               YWWRYMRRSWRCCCCYYYRYYRWWRYYKYGTGRTWWTATCAYYAYMKSKRAYYRRWMSTY
                ***********    **************       :.   *  * .***:*******

8               GGGATWWYAYMYYWMMKRCAKYRYRMTYRTTKKYYRTTCKKGCCTCRRYYRYRTGMWYGK
9               SRACGWWYGYMYYWMMGGYRKYAYRMKCGGCGKYYRYCAKKKYTCTRRYYRYRCTMAYRK
                . .. ***.*******    ** ***.     ****  .**     *******  * * *

8               RSMWMYYKRKKTKRMWMCCTMYYGKYRYRYRYRRTCTYRWAYTKRWKGRWMYKRMYKKYR
9               RSMWMYYKRKKYTRMACGTCMYYRKYRYRCRCRRYSYYRATTAKRWKRAWMYKRMYKKYR
                *********** .**     *** ***** * ** . ** : :****  ***********

8               CYAKMKWRATCGRCKCYRYYYKKRKWMYRWKKCKWCGMYKKTWWGKRKKYKATTKAAMKW
9               YCWKMKWRCCYTRTKTCGYYYKGRKWMYRWKKTKWYTMCKKGATRKRKTYKMYYKGGMKW
                   *****.   * *   **** ********* **  * **    ***.**   *..***

8               YKRRRYYYTRTTYGGGRKSRYTGCWTYMKKKRKKYMKKRSMSMWKKRKRGAAKWRKYAYT
9               TKRRRYYYYRYKYRRRRKSRCGRAAKYMKKKRKGYMKKRSMCAWKKGKRATTKWRKYRCC
                 ******* * .*   ****   . .******* *******. *** **.::*****
```

```
8     CGRRSYYKWYYRKYKCRSRRGRKRAYKKRKCKKGGMRSWYRKKMMYKKKMRYTKMKAAGY
9     TAGRSYYKWYYGGYTSRSRRKRKRMYGKRKMGKARMRSWYRKKMMYKKGMRYCKMKCRKY
       . ********  *..**** *** * ***  *. ************* *** ***.  *

8     SKTTTYYYRRKMKYRMKYMKAMRRYYYKRRKAGACCYGRTYYKRGMWKKRRKARYRYWKY
9     SKCCCYYYRRKMKYAAGYMKCMARTYYKRRKRRWYYTRRYYYKRKAWTKRRGWRYRYWKY
      **   *********   ***.* * ******       * ****  *.***  *******

8     TKKWRWGYSKKKYKRRRKYMYRSYYRKRMMMRYRYRSWWYRWKMWKMYCKRKAKYKYMWY
9     KKKWRWKYSGKKYKRRGKYMYRSYYRKRMMMRYRYRSWAYRWKMWKMYSKRKRKYKYMWY
      .***** ** ****** ********************* *********.*** *******

8     YASKRRGYYWSYYKMKWMKYKYGKRAAKGMYCGTMRYYYYCRRGAGAKYAYKGGCWGMWY
9     YRSKGRKYYWSYYKMKWMKTKYAKRWTKRMYYKWMRCTTYYAATGRTKYGYKRRMWKMWY
      * ** * ************ **.** :* **   **   *    . :**.**   * ***

8     AGACGKYYKKKKKYRKWRMMKCYAYYRSRKRRYYRRWKKTAAYMWYKYAKKKTYRWRMYK
9     GSWYKGYYKGKGKYRKWRMMKMCWYYRSRKRGYYRRWKGYMRYMWYKYCKKKKYRWRMYK
      ..    *** * *********   ******* ******    ******.***.*******

8     KYRYKYYWKRMKRKRRYYKWRKAYYYRKYKTGCYRYCRYRRGKRKGRYGWYKTYWYMYKK
9     KYRYKTTAKRCKRKGGTCKWRKMYYYRKYKYRYYRYYRTRRRKRKKACKWTKYCTYMYKK
      *****   ** ***    **** *******   *** * ** ***    * *   *****

8     TRYKWTAKKGYGTCRYKKCAKRKMMMYRYTKKKYKKKWKYGRYSWKCRCYYWMMYAGKRR
9     ARYKWCWKKKYAAYRYKKYRKRKMMMYGCCKGGYGKKWKYAACSWKYRMYYWMMYMRKRR
      :****  ** *.: ****  *******   *  * *****.  *** * ******  ***

8     KYRKRKTKKKCTTYRRYCCATKAYKYRWKGTCKMAAMWCKYKMWYYSRMAKKSYRYRKCT
9     KYRKRKCTTKTGGYRRYGAGGKWYKYRWKSCTKCWCMWTKYKMWYYGRMMKKSYRYRTMY
      ****** ..*   **** .. * ******.  *  .** *******.** *******.

8     RKRMYRGRKYYWKRRMYKKWYYKKSKYRTCMKKYRRKKKYKRKKMRKKGGRRYKRYWYYK
9     RKRMCRKRKYYWKRACYKKWYYKKSKYRCTMKKYRRKKKYKRKKMRKKRRRRYKRYWYYK
      **** * *******  ************  ******************  **********

8     KWRWYRRCGGARKYKKAYTGKKKRKKAAACMRKGKRRRMKWKKMRKYSKYYKCTGYCGSR
9     KWRWYRRYKKRRKYKGCCCCKKKRKKGGTMMRKKGRAAMKWKKMRKYSKYYKYKRYSRSR
      *******    **** .   ******..: ***  *  ************** . *. **

8     GAATAWWMKGKKKAATYTCAKMKGKGKTRYAYCTARGCAMGGYYGAGWRRKGYAKTTGYC
9     ARWWCWWAKKKKKCCGTKMCKAKKKKKYRYGCYCCRAYGMATTYRRRTGRKRTRKWYKYY
      .   .** * ***..  . .* * * * **.   .*. .*.  *     **   *   *

8     RRRCTGTWMWTCRYRAAKRMWYYKWYTKKYYGWMKRMSKCYRRRKYWKMYMARRKYMMKM
9     RRRMYKCACWCTRYRRRKGAWYYKAYYKKTYTAMKRMSKTCAAGTYTKMYMTRRKYMMKM
      ***      *  ***  *  **** * ** *  ******     .* ****:********

8     KYMGRMRKYRKACKYTSTMMYRRRMRWARKKKATGKMRRKYWYRRGGCKMYKKYMKYKYM
9     KYMKRMRKYRKGGGCYSYMMYRRRMRTMRKKKTWATARRKYWYRRTKYKCTKKYMKYKYC
```

```
            *** *******.    * ********  ****: .. ********   *  ********

8           YYRRCTYRKWYCCKYWRYTYYMKYCMKYYKGGWKKRKYTGGAAAKKWKRWMWWWMMYYWR
9           YYRRTYCGKWYAYKTARYKCCMKCTMKYCGACTKKRKYCRKRRRKKWKRWMWWWMMYYWR
            ****    ***. *  **.  **  ***  .  *****      ****************

8           RRKRYKKRMCKRCWYRGRARKMKTMYKTYKTWKRKKKYKTTRRRYAKSRSYYKAMKMKYR
9           RRKRYKKGMAKGYWYRKGGRTMKKMYKYYKAWKRKKKYKYYRRGYRKSRSYYKTMKMKYR
            ******* *.*  ***  .*.**.*** **:********  ** * *******:******

8           RRWYTKCTKGSMWKRKRYKYKTTTAWRRKKYMWYKWKAYTRTWRRYRYTTKYRKRTKCRW
9           RRWCAKGAKRSMTKRKRYKYKAACGWRRKKYMWYKWKRYAACARRYGCYYKYRKRYGYRT
            *** :* :* ** ********:: .************ *:   ***    *****   *

8           SRYRRGCKYYMKGKKKTTGMSKYRGKTWMYKGGTKWGAGTWKTAKTGAAMACGTAATMGT
9           SRCGRKTGYYMKKKKKKYKMSGCGAKYWMYKKRYKAAGCCWKYRKGSGWMCMAAMRYMKK
            **  *   **** ***.  **   .* ****   * ..  **  * .. *. .:   * .

8           YYYYYWAKWKKRKYKMYGCKKKMMKRCCCTRYYWRRKYRKMKKMYYKTMTYYGYTAKRYK
9           YYYYYWGKWTKRKYKMYKMKKKMMKRTTMGRYYWRRGYRKMKKMYCTAAKYYKCYGKRYK
            ******.**.*******  *******    ****** ******** .: .**   .****

8           KKWYMKKRRMMYARKRWSGKKMKACCKTAGGMKRRYRRCYMYMRYKCKKKKKKCWWRSYY
9           KKWYMKKRRMMYMAKRWSRKKCGCTTGYMRKMKRRYRRYYMYMRYKYKKTKKKYWWRSYY
            ************  **** **   .       ******* ******* **.*** ******

8           YCCTTCYYKKGGRCRYYMAWYYRYKWKKKKSMKYWKMGAGYMMSKRKTKGTAYKKWYYMG
9           YYYCCTYYKKKKRMRYYMMWYYRYKWKKKKSMKYWKMKWRYACGKRKYKKGGYKKATTCR
            *     ****  * **** ******************   *  .*** *  .***

8           CTYMYKRKKKKRWWRRRYRKYAMKYCCYRKYMWRKWWWRRKMSKGATAKYYYAGRRRKRY
9           AATMYKRKKKKRWWRRRYRKYWMKYTTYRKYMWRKWWWRRKMSKRRCCKYYYRKARRKRY
            .: ****************** ***  *****************   .****   *****

8           WWKYYYCCYYRKKKKRRRTRMKKWRKCCRKYKKKRRKRKCCKYMKRSRKYTWWKMKYRRA
9           WWGYYYTTCCAKKKKRRRYGMKKWRGTTGKYGKKRRKRKYYKYCKRSRKCAWWKMKYRGM
            ** ***     *******  *****    ** *******  ** ***** :*******

8           GGAWKGRKKARMKRRRYGGAKARKYRKCWKKAYCTGKYYYYWTKYGYAGMRYYMAKMMSM
9           SRRWKKRKKRRMKRRRYRRMKCRKYGGYWKKWYTAKKYYYYWYKYAYMTAATTAMKMMSM
            .  ** *** *******   *.***   *** * : ****** **.*        *****

8           YYCCRWRMMRRYMRWRKRKYKYGCKMKCKRYRRKRYMKKCCGMKKMMCGRYKKKKMYKKR
9           CTYTRWGMMRRYMRWRKRKYKTAYKMTTKRYRRKRYMTKMARMKKMMMRRYKKKKMYKKR
                ** ************** . **. *********.* . *****  ***********

8           MKCKMKCKKYRYKSRYYMYWKACRTKRRTGWRRRRYWRYMAGYCGATCWKWRGGTYKKYG
9           MGYKMKMKKYGYKSRYTMYWKGMRCKRRARARRRAYWRYMTRCGTRGYWKTRAAYYKKYR
            *  *** *** ***** ****. * ***:  *** *****:       ** *.. ****

8           TCYWCRMRTKATYWKYMKYRGKRRRKYKGRKGRMKYYTTTAAKSRGTGCKYMCKRRGTGW
```

```
9      WYYWYAMAAGCCTWKYMGTGKKRAGTCTKRKRRATYYACCWWTSRKCAYKYMMKRGKGAA
         **  * : .  ****    **  . . ** * .**:    .**  . *** **   .

8      YWCMMYMYYRKAMRYKYASKYGRWRCTKWKKACGWYRRGTARRKKMCYATYMRKSRRRKM
9      YAYMMYMYYRKGMRYTYWSKYRAWRCTKWKGMYRWYRGTAGRAKGMYTRKYMRKSRRRKM
       *  ********.***.* ***  **  ***    ***  :.* * *   .**********

8      MYYRKATKGGTTKKWWARKKMRRRAKSKWRYYRKYRRYYRRYWMRRMGAKAGKMRTTTGT
9      MYYRKRYKKAGGKKWACRKKCGGGWKSKARYYRKYAAYYRRYWCGGARGKWTKAGCGYTY
       *****  * .  *** .***     *** ******  ******     .*  *

8      MAGARACGMYYYYKGRRYGWYYYCAAYGASKRTCCRMRRTYWGACKWYYGRYARMAAACK
9      ARATRWYKMYYYYKKRRCAAYYYMTGYTWSKGCAMRMRRACTATMGWYYKAYWRMWRWYK
         .:*   ****** ** . *** :.*  **  . ****:  .:  ***  * **    *

8      GYTYKRRYWTCTTAKWKYAGKTKATMRAKMKYYKRRRKKKRKAYKYRCKYKYCAKYKRYM
9      RYWYKRRYWGTAWWKWKCTTKCTWAMRGGMKYYKRRRKKTRKMYKYRTTTKCYTGTKRYM
        * ******  :  *** : * . :**. **********.** **** . *  :  ****

8      YRRWKGRMRKKYAMRTCWTGWKRKRRKYMRCAAKMYWKYRRSRRYRRMRTCKWRKYYGAA
9      YRATKAGCRKKYWMRCSWCAWTGKRRKYMRMRRKMYWKYRASRRYRRMRYMKWRKCCRRR
       **  *.  **** ** .* .*. *******   ******* ********  ****

8      MCTCTYKAGTCKKRTKKYKAACCTTYRWWRRYRYYYYKMYMKWYACCYCKKYRRGYKTWK
9      MMYTYYKGTAYKKRYKKYKMRSMYKCATWRRYGCCCYGMYMKWYTYMCYKKYRRKYKCWK
       *    **. : *** ****  .  .   ****    * ******:    ***** ** **

8      RMRRKKKGAGAKRWYKKYKMGKRRRGGWRYMYKRMKYWTAWRYKRKWSKGYRCRRYYKKG
9      RMRRKKKRRRGKRATGKYKMRKRRRRRWAYMYKRMKYWCCWRYKRKWSKKCGMRRYCKTR
       *******   .**   **** ****  * ********* .*********    *** *.

8      YWKRTGCAYRSMYWKMKAKGCGMRYYKRKKWMYRKRRKWKYTYRYGRYSARKRKKYGKKY
9      YWKRWRYWYRSMYWKMKWKAARMRYYKRKKWMYRKGAKWKYYYRYKGYSMRKRKKYKKKY
       ****    ********* *.. *************  **** ***  ** ****** ***

8      YYMKKRCKYWRTGKWKMYKCRWKYGYTAYKYGRYKTKTTYGCRKKMYKCTKKKKTCTCCT
9      YYMGKRYKYWRCAGWKMYGYRWKYATCRCKYKGTGKKYYYRMRKKMYKYYKKKKKSYYMY
       *** ** **** . ****  ****.    **    .*  * ******   ****..

8      CTRWCAGKKYRAATYRMAMKKYMYTTCYCKRKKYAKRKKWSMYCATTGGKYYKCTGYATW
9      TYRWTGKKKYRWRYYRCRAKTTCYKKYCYTRKKYWKGKKWSMYTCKYRKKYYKMYRTGCT
         ** . ****   **   *.  *..   .**** * ******  ..   ****    .

8      YYKRKACAMRKMKRWCAKKAAGGCYGGSRARYKGYYCKRRKMASCYGTGKWRCTRTARSY
9      YYTGGMTGMRKMKRTYCGKGTTAACAACGRGYKKYTYKRGGCRSMYKWRKWRSAGCTRSY
       **.    .******  . *.: .. ...   ** *  **    * *   ***.:  :***

8      RMYYTRRYRYMTTWWRCCYCTKTYMMYRYRYKCTGACCCTTAACYRGKTYARYYRWTMTA
9      RCYYKGRYRYCKWWWGTATTKKCCMMYRYRYKYGKCYGYCAWWYYRKTCTCGCCGWWMYR
       * **. **** . **  .  .*  ********   .    :   ** .  .    * *
```

```
8   TCKYYYCSRTKKKGTTSYAYAYWRTMCKYYWRYRWKKYTGCGRYKYRWCRKRKWSMYKWR
9   YYKYCYYCAYTKKKYKSYTTGYWRCATGCTARTRWKKYWRMAGTKYRWYGGAKWSAYKWR
      ** * .  .**  .**: .***       * *****   .  ****    *** ****

8   CMRAGKYKKKWGRKYRYAAKRMACWKRWYGARCMYWYCGKKRKYASRMMKKRAAAKYYKR
9   MMGGAKYKKKWRGGTAYCGKRMTYTKRWYRRRMMYWYMSKKRKCGSRMMKKRGTGKYCKR
     * ..******     *..***:  ****  * **** .**** .*******.:.** **

8   KGARKGAMKKWCGWYKAYKKYSMKYMMRKCCGRRKRCCCYWYWSMWRKCATKMCMRRYMY
9   KTRRKAGCKKWSRWCKMCKKTSCTYMMRKYYKRRKRYYYCWYWSMWRKYRYKAMMRRYMY
    *  **.. ***. * *  ** * .*****   ****    ********   *  ******

8   GYMKKMYYYRKTTKWRRRWTYYRAKARRWKSRYYKKKYYRTTKYKRKSKRCRYKSKYRKK
9   RYMKKMYYYRKWYKWAGAWYYCGRKTRGWKSAYYKKKYYRYKKYKRKSKRMRYKSKYRKK
     **********  **   * *   *:* *** ******** .******** *********

8   YRRSYKRYYKYKSSWRKKYSTKKYWKKYGCSGKWMRRRMGCCRRKKKYMRTAKRMMGCWY
9   YRRSYKGYYKYKSSWRKKYSYTKYWKKCTYGAKWMRRRMKYYRRKKKYMRAGKRMAKYWC
    ****** ************* .*****   ..*******   ********:.***   *

8   MKRKKYYYYKWSSAMRSCGGAKKYTAYMSRRRTMRMKYKYRKKWWWYYMYKYRKWYGCKKY
9   MKGKKYYCKWSSTAGGTRKWKKYYRYMSRRGYMRMKYKYRKKWWWYYCCKYRKWYAYKKY
    ** **** ****:  .    ***  *****  ***************  ******. ***

8   KYRRYRKRKTTSWMMWRTGATCGKKKRRGYGWYYMKTKMRWRGYYRKYKCGYMKMYRYKK
9   TYRRYRKRKYKSWMMWRGTRWMRKKKRGRCSTYYMKWKMRWRKYYRKYGAATAKMYRYGK
    .******** .******      ****   . **** ***** *****  ..  ***** *

8   KMWARYMYRRMRRMMRRYWKMSRRWKYWKKTKRRTKTKRWKYYKTYAMACAKSWYYYWWK
9   KMWRRYMYRRMRRMMRGYWKMSRRWKYWKKKKRRAKYKRWKYYKKYWMRYRKSWYYYWWK
    *** ************ *************.***:* *******.* *   *********

8   KYYTRYRTWYYYAKYRCMMRWYKYGYKRRYKKGYKYRYKRKRKRYMYKRKRYYKKYWCAT
9   KYCYGTRYWYTYRKCRYACGWYKYRYKRRYKKKYTYGYKRGRKRYCCKRKRYYGKYWSGC
    **    * ** * * *    **** ******* *.* *** ****  ****** ***..

8   GKTCTYMKKKWKGTRAMARYYWYKYKYRKMMKRYTKWYYYRGMKTKYMTYGRYKRYWYCA
9   CKGYCTAKKKWKRKGMCMRYYWYKYKYRKMAKRCGKWYYYRKMTGKCMCYKRYKRYWYTG
     *     ***** .    ************ **  ****** *. * * * ******* .

8   CWTYYKTYSYWYSYYYRGYMYYGYMARKRCRKYMRYTTTGKKYYRRWWRYKAKYRCCAAY
9   AWGCYKKYSYWYSYYYAKCCCYRYMWGGRMRKYMRYCCCAKKTCGRWWACKWKCGTTGRY
    .*  **.*********     * **   * ******   .**   ***  * *    . *

8   WRRMCSRKMKYAKYRMRKYYMSWKRYYRKYR
9   WRRCYGAKMKYMKYRCRKYYMSWTRYYRKCG
    ***  . **** *** *******.*****
```

**Supplementary Figure 3.** The pairwise alignments between plants 33 and 34.

```
CLUSTAL 2.1 multiple sequence alignment (clustal format)

33              RRYYWYMCACKKGARTWRKYRYYYMMYWWATYYKMWRYWRWSRMKYCSYRKKKYTMMSWY
34              RRYYWYMTGYKKKCGYWRKYRTYYMMYWWWWYCGATRYWRWSRMKYTCTATKKYGMMSWY
                ******* . ** .  ***** *******  *    ********** .  .*** *****

33              ARYGRYYRYRRWYYWRWYRRMCAGYGMSCKRRRTSMRMYRRKRYYRWWMRGCRMMRRMYA
34              TRYRRYYRYRRWYYWRWYRRCGGKCCMSTKRRRYSMAMYRRKGYYRWWMRATRMMAACYG
                :** ****************  .   ** **** ** ***** *******. ***   *.

33              YYSWTKYMYYTGWKSYYRRKYMWYKGCWSTMKYRWYYYYYYRRMYRYTRTMYYMWYKRRR
34              YYSAWKYMYYAAWKSYYRRKYMWYKAAASYMKYRWYYYYYYRRMYRYCGYMYTCTYKRRR
                ***  *****:.*************.. * *****************   **   *****

33              KRKKRYTCKMRAMRYRWWYYMMMTCYTCWKRYCTGGSARRKMMYKYSYYYYYYSMYRTAR
34              TRKKRYCMTMRRMRYRWWYTMMMAAYWTWKRYTKRKSWRRKMMYKYSYYYYYYSCYGGTR
                .*****  .** ******* ***:.*  **** .  * **************** *  :*

33              YYRAATYMRTKYCMGYWKRMKYSRYMYTRRYWRKGMCMMYSCYAMKWYYYWRYWRWTKYK
34              TTGRGCTARCKYYAKYWKRMKYSRYMYWRRYWRKKMYMCCCMYCMKWYYTAGCWRTKKYK
                    .   * **   ************ ****** * *  . *.*****    ** .***

33              MTGYYKGARYRCRCSYYWKYYWRYYCMYWWYSRRYRMRRMRMYWMGGGRTYKWWRYKKRK
34              MAAYYKTRAYRAAASYYWKYYWRYYYMTATYSRRYRMRRMRMYWMRRRACYKWWRYKKRK
                *:.***   **. .*********** *   ***************     **********

33              MCCAWMMYYMRSKRYKTYKYRMYKRYMKKKMTYRKMYMWMCAAGTYTACGMSYKRRRMTK
34              MMMRAMMYYMRSKRTKAYKYRMTTRYMKKKMCYRKMYMWMYWRRYCYGAKASYKGRRMGG
                *    ********* *:***** .******* ********       ..  *** ***

33              MWAGARWTYYRYMMRRCCGYRTRRACTTAMWWSYMYWATYTAYGYKKCYGCTTYCGTGGG
34              MWRRMRAKYYRYMMRRYYKTACRRGYKWMMWWSYMYTGCTKRTRYKKTCSTYKYMRWTAA
                **   * .********      **. .  ******* .  .   ***  .  .*     ..

33              CAGCKYKRKSYYMKMKWYSYYKYMCAATTAWTTYYKARRRRSYSKRSRYKMAKYWKGGAA
34              MMRSKYKRGSTTAKMKWYSYYKYMYCCAATTACCTKRRRRRSYSKRSATKMRKYWTTARW
                   .**** *   *********** ..::: :   * **********  ** ***. .

33              YYRRMKKRYYRKYSRKWMCYCGYCARMKYACGRCAAAATYYSWWKRATGRWRYRRRWTCK
34              YYRRMKKRYYRKYSRKWMTCGACYRRMKYGTAATTTTCKYYSWWKRGGRRWRYRRRWYYK
                ******************   .   ****. .  :::..*******.  ********  *

33              MMYRWKYYKWYKKYRKKKMYYMYWGMAMYKSYYKMKGTAKATRRKYKRKWYRTTRKYYWK
34              MMYRWKYYKWYKTYRKKKMYYMYWRMMCCKSYCKMKTWGKGCRRKYKRKACAYCRKYYWK
                ************.*********** *   *** ***  .*. *******     ******

33              KMKTRWKMYGSMKWRAKGCMKKWWYKKYYWKRKKWWYMMYRYMRKKKKAYRMSKYWYKRK
34              KMKKRWKMTTGMKWRRGAYCKKWWYKKCYWKRKGWWYMMYRYMRGTTGCYRMSKYWYKRK
```

```
            ***.****  .****  .  ******* ***** ********** .. .***********

33          KKMMMKRKKMMYKKRTRYMRWTRKYYYYKKMWKKKYYYYKKGCCCAKCRRKRKTWTATGC
34          KKMMMKRGKMMYKKACRYMRWAGKYYYYKKMWKKKYYYCGKRMYYMKMGRKRKGWYRYAT
            ******* ******  *****: ***************  *     *  **** *   .

33          KRTMRSKCCCKKMYYAAAKMMKMYMKSWKKKKYKRKRWKRKYMKYKKSKMTTRRMYKRWC
34          KRWCRSKTYTKKMCYCGGKMMKMYCKSWKKKGYKRKRWKRKYMKYKKSKMGCRRMYKRWY
            **  ***   *** *...****** ****** ******************  *******

33          RRYKKGKYRYRRYRKKGRKRRAKCTYRYKKKYRRYYKRWYRWRTSYYRGGGTRWMYRWYT
34          RAYKKRGYRYRRYRKGARKRRGKGCCACKKKYRRYYGRWYRWRWSYYRRKKKRWMYRWTC
            * ***  ******** .****.*     ******** ****** ****   .******

33          YKYYYGARSYYYSYRWRRYYKKWYKRTRKGGGKAATKRKYYKAGGMKRYCKKYMMYWRRK
34          YKYYYRCRSYYYSYRTAGYYKKWYKRKRKTKAKWRYKRKYYKCCKCGRYAKKCACCARRK
            ***** .********   ********.**  .*   ******.    **.**     ***

33          RYSKYKSKYAYGWYATTGTYYYTWKCCACAYGRKWWWRKYGCTTRYYYYKWSWRMYKRWM
34          RYSKYTSKYWYRWYGGCTYTTTATGGYGAGYRRKWWWRKYRYWKRYYYYKWSWRMYKRWM
            *****.*** * **.       :    ...* ********   .****************

33          WRTYYAWYTKMYYAAYYYKKTMKRAYRKRKYYKMYMRRKAKYRYYRMYYRYKKKMYWKYR
34          WRKYYMTYCKMYYRRYYYKKKMKRTYRKRKYYKMYMRRKGGYRYYRMYYRYKTKAYWKYR
            **.**  * ****  *****.***:**************. ***********.* *****

33          YKYWKMYCTTGKKMRGCGKAAAKRAAYMRKRSAWWKKYMRKYRRACAWTCYSWYKATATC
34          YKYWKMYACCAKKARRMRKGGTKRRMCMRKRSWWWGKYMRGYRRTTMTCTYSWYKTCGCY
            *******.  .** *   *..:**   ***** ** **** ***:     *****: .

33          YSCCYRKMCACYKYYWYSYYRMKYKWYKAGGYTYKMWKKKCKGCTTAYGARRAACKMCGT
34          YSGMYRGMMMYYTYYWYSYYRMKYKWYKGAACCYKMWKKKTKAACYGYKWARGGTGMMKA
            **  ** *   *.***************...  ******* *..  .*   *..  *  :

33          CCKWKKRWRWYRKSTRKGYRCCGCKYKYMTAGRKKKYKKYRWCTATCTTCRYYRYKKKKR
34          TTKWKKRWGWYRKSCRKAYRYYRYKYGYCCMKRKKKYKKYRWGGGYYYYMRYYGYKKKKR
              ****** ***** **.**    ** *    **********  .     *** ******

33          MWRYYKWRWKTKACTWRYRKKMRKKYMYCKKKCKKYWKKRRKKKMKMYKYCTTTRTYWCK
34          MWRYYKWRWKKKWYKWRYRKKMRKTTACYKTKTKKYWKKRRKKKAKMYKYACACGATTTK
            **********.*  .*********.    *.* *********** *****. :  :   *

33          KKWWKYSMYRRRKTATTWGKRKKRYKMSRSYMMYWAGRYKMCAAGKKRAGYGAWWKMKKY
34          KKWWKYSMYRRRKAGCAWCKAKKRYKMSRSYMMYWMKRYKAYRMRKKRGACARWWGMGKY
            *************:. :* * **************  ***     ***.. . ** * **

33          YYKKWKKRKKRRWRRRCSRAATAKYTTGYWYGKAGGWYKYWYMYMRKCAARKRKKKKKYY
34          YYKTWKKRGTRRWGGGTCGTGKWKYWCRYWYATGRAWYKYWYMYMRGYRRGTGKKKKKYY
            ***.**** .***    . :.. **   ***... .**********     . *******

33          WKMKWRWWWYWKKRRRYYRYYTKRKKTRWMKGTMKKSYKRTWAKYKMRKMYKKMRRWKY
```

```
34        WKMKWRWWACWKKRRRYYRYYYKRKKKRWCTRWMKKSYKRGTRGYKMRKMKYKKMRRWKY
          ********  *********** ****.** .  *******    ****************

33        RKKAGTKRKGTRYYRYMCMKWWYKKCYYYYWYYKYKKAGTKRKYYRWKYKKMRKKKRKTK
34        RKKGAGGRKAYRYYGCCYMKWWYKKTCYYTACYKYKKMTYKRKYYRWGCKKMRKKKRKAK
          ***..  **. ***    *******  **   *****   *******  *********:*

33        KKKSYYRRGYRYGTTYMYWRSRYAYWTAGKKTMKKAWWRMMKYKKRYYYYMGMRYRKMKK
34        KGKSYYRRRYRYAGCYMYWRSRYRYWWWSTKKMKKMTWGMMKYKKRYYYYMRMRYRKMKK
          * ****** ***.  ******** **  ..*.***  * ************ ********

33        KKMMKMMKKSYRWGWRRYYRWKKKKRTMTMKKKKCRYYRWSRRKRWWTRYYYKRRYYKRR
34        KKMMKMMKKSYRWSWRRYYRWKTKGGCMGMKKKKTGTYGWSRATAWTGRYYYKRRTTKRR
          *************.********.*   * *****   * *** . *  *******  ***

33        RKKGKRYKRCAYYKKAKCCAYRKTGGCTAKKSYKMYGAKKYCTTGKYYGARRYRRGGCCT
34        RKKRGRYKRTMCYKKWKYYWYRKGTTYCRKKCYKMYKTKKYMYWKKYYATGGTAGRAYYK
          ***  ****   *** *   ***      **.**** :***    ***.:      .  .

33        YKRRKRKCKRRKRKKYTKRYTTTTTTYYTWTMKYTWYRWGWTGACTMCGMYRKYSYYKKK
34        YKAAKRKMKRRKRKKYAKRYYKYWWYTYKAACKYWWCGARWCTCMKCTTMYRKYSYYKKG
          **  *** ********:*** .     *. : ** *    *  . .   **********

33        WMRYRWRKGGRYYKGTYSYKKSRYMRYRRMCRYKWRRYMKKKTRTCTTAKYRKKMRRTGA
34        AMRYRWRGAAATCKRKYSYKKSRYMRYRRMYRTKWRRYMGKKKRYMYWWKYRKKMRRYSR
           ****** ..   * .************** * ****** **.*     ******** .

33        RKMRKGRYWKKYWSMRYSTTCCRYKYMCGCGWAGTMSRCRKKYYKRYSGCCCGKRYRCAK
34        RKMRGARCAKKYWSMRYSYYYYRYKYMAAAAWRAKMSRAGKKYYKRYSAATTAKRYRARK
          **** .*  *********    *****....* ..***. ********..  .****. *

33        KYKRKMWYYSKCSTKMTRMGRGCGRRKKKRYKKYYGCTATMTYWYKKYYYKCAGRRAYWM
34        KYKRKMWYYSKTSKKMARMRGRYARRKKKRYKKYYKTWWYCYYWYKKYYYKYRRRRRYWM
          *********** *.**:**    .***********       *********   ** ***

33        RMWKRTRKWATAGYYRKRKRSKKKKKGYAWKKTWYRRRRKYWRRYYYYMGKKKKWRSCYK
34        RMWKRKRKATGTKYYRKGKRSKKKKKKYGTKKCTCARRRKYWRRYYYYMRKKKKWRSMYK
          *****.** : : **** ******** *. **    ************* ******* **

33        AYCYRYWYMRGAYKKWTYYRYYWWYGATACYRRYYKMKRKMRRWMAMGRASMMARMMWGA
34        GCYYRCWYMRRRYKKWKYYRYYWWYTCWGMTGRYYKMKAKMRGACMMAAMSMMMRMMWRW
          .  ** ****  ****.******** . .   ****** ***    *.  *** ****

33        WKRKMYYRYKKRYYTACCYKKAYRKYTCKYYRRWMCCKRAAKMKRWRMKRRYKWWYMWWG
34        WKRKMYYATTKRYYCCTAYKKWYRKCCAKYYRRWCTTKARGKMTGWAMKRRYKWWYMWWS
          *******  .**** . .*** ***  .******   *  .**. * ************.

33        RTKKCYYARKCTGCSYRKKTACKRWRRRMYMYRYYRRRGRWCRYMYMRYWGGAGYKYKYC
34        AAKGYYCRATYYRSSYRKKWMMKRWRRRMYMYRYYRRRKRWMRYMYMRYWAATAYKYKYA
           :*  *   .   .*****   **************** ** ********..:.*****.
```

```
33    ARKYMYSGAACAAWYYKKKKCGWKMSKMRRYYRKCYKYRKWKKKRSSKWKKKSRKRKYAW
34    RRKYMYSRRMYWGTYYKKGKGKWKCSKMRRYYRKSYKYRKWKKKRSSKWKTKSRKRKYTW
       ******     . **** *  ** *********.***************.*******:*

33    AGGTTRCGCTKYKSCWKCCYYMWYRRRCAGMWKKKKWTTTCGCRCYCRMMKKGKMYRMRK
34    RRRYYRYATAGYKSTWKYYYYMWYRGRSRRMWGKKKWYWYMRMRYYYRMMKGAKMYGMRK
           * . : *** **  ****** *.  ** ****      * * **** .*** ***

33    GYKWWKWAYYKKMYWYCCGMKKSWSYMKYMMKKTAMGRYKKRKKATTTYWTKKACGRGGK
34    AYKWWKWRYCTKMYWYTMRMKKSWSYMKYMMKKWMCRRYKKRKKCCCCCACKKGTARKSK
      .****** * .*****   **************    *******.      **. .* .*

33    RRRKCYGMYWYMYKMCAAGTYCGMSYWYYCRTMYYSRKRGCGMYKMRKYYRKMWRYKRAG
34    RRRKGCKMYWYMYKMARCRYCTAASYTTCYRYAYYSRKGATAMYGMGTYYRKMARTKRCA
      ****   ********. .    . **    *  ***** . .** * .***** * **..

33    TAWRSTCAGYAATKKGTAWRRKTGTYGKYKGCYMRGRYKWMYCTRWCYKYKKKTCMRKMR
34    AGWRSWMMRCGGAKTKAWWAGGCACYTKCKKTYMRTRYKWMYTAGTTYKYKKKKMARKMR
      :.***     ..:*. : *    . * * *  *** ******  :  ******.  ****

33    KGGGTWYKSMWRWKMWKWWKRKAGYYKRKRYTRMKYMYKYRWYRYYRYKRAKCGTKTKKY
34    KKTRCWYKCMWRWGCAKWWKRKGTYCKRKRYKRAKYMYKYRWYRYYRYKACKYRYKYKKY
      *    ***.****   ******. * *****.* *************** .*   * ***

33    MMKYRSRKKKKKRRRRRYMWRYRKTTMAKYYYACTKYKKKGTCKKWSKKTGYKKRYKYRW
34    MAKYRSRKKKKKRRRRRYMWRYRGACAGGCTYRAYKYKKKACGKKWSKKKRYKKGCTYRW
      * ********************* :  .   * . *****.  ******. ***  .***

33    RRYKMKTTTKYKGWRATTRYTAAYTYWWRYRYMWRTMTWYTCKRKYTATYRMKYKKTRSM
34    RRYKMKWWYKYKKWRRYYACWWWYYYWWRYRYMWRYCGWYYYKRKYYMKYRMTYKKKRSM
      ******   *** **        * **********   **  ****  .***.***.***

33    YAKTWATGGARMMCKYYYMYAGRARKKKRKTTTTCAKWATKKRMRGWYMWMMYGGYYKKY
34    YGTGTRYSKRGAAMKYYYMYGRACRKKKRKGGACTGKWRYTKRCGKWCMWMMCAAYYKKY
      *..    .      ******.  .******  :  .**  .**   * **** ..*****

33    YYWYRKRRGYATTCKKYMAAKRRKYMATTMKKKKKMKWKKYYMKGRKMYYWRKWRRGWCK
34    YCAYRGRRACCCYMKKYCGWKRGKYMRACMKKKKKMKWKKYYMKKRKMYCWRKWRARAYK
      *  ** **. .   *** . ** *** : *************** **** *****    *

33    YKKYKGRKTGTTGATCCGGAKGTAYTKKGKYYWMKKRMMRYKKGYKRKKWKKRYMWRRYK
34    YTKYTAGKARCYARGYYATGGAGGCKKGKKYYWMKTRMMRTTGRYKAKKWKKRYMWRGCK
      *.**.. *:   .    . . . . .*  ******.**** .  ** **********  *

33    YWYYMYTTCYRWKYGMRRRRKMKTGCRKYYRWMYMRSTKRSRAAWYRTTRKYYYKKYMKS
34    YWYYMYWKYYRWKYRCRRRRKMKCRYGKYYRWMYMRSCKRCGGTTCGACRKYYYKKTAKS
      ****** . *****  *******    ********** **. .:   : *******  **

33    MYKTTRCYYYWWKMRYYSYCMGCWKTRKGAWAKRCMYKMAAKKAKKSKWKAYRKYYKYSY
34    MYKKKGYYYYWWKMRYYSYMMKYWKKGGAGTTKRTCTKMGCKKTTKSGWKMYRKYYKYST
      ***..  ************ *  **.  .. :**   **..**:.** ** ********
```

**Supplementary Figure 4.** The pairwise alignments between plants 3 and 4.

```
CLUSTAL 2.1 multiple sequence alignment (clustal format)

3               KRYMGAYACYGTAGATMGGTTGTTMRGMYMWCGYGYWYWWKYRRKRGWMSMARYYYMGKR
4               KGTCTCCTYYRWMACCAATCAACAARTMYMWSRYKYWYWWKTARKRAWMGARRCYYMAKR
                *    . : *   ..  .  :. : * ****. * ******  ***.**.  * ***.**

3               WRRKWTKRRYAARYYYMMYSMYSYYYGRRAMSWTGKCRRRMTCCYGAYWCWAWYWRWYYR
4               AAGKWKKRRYMRRYCYMMYSCYSYYYRRRGMSWWRTYRAAAYYMTTGYWYWMWCAGACTG
                   **.****  ** ***** ***** **.***  . *        .** * *

3               WKMWKRYKKYSWRYWYRKYRYMYWRMMMYRCMWWRYMTCWYRRWAYGKSWSCRRYYRRCM
4               WKMAKRYKKYSWRCWYRKYRCAYWRMMMYRYMWWRYMWTTYRRWGCAKSWSYRRYYRRAM
                *** ********* ******  ******** ******   ****. .**** ******.*

3               KSMMYRRCSKMMARWYTGKRTYMRYRKYYYRCMRYRWWYYGYKAAWWKKYYYRGWWRMWY
4               KSMMYRRYSKMCTRWYYRKAKTCRYRKYYTRYMRYRWWYYRTKGRTWKKYCYRKATACWY
                ******* *** :***  * .  ****** * ********  *.  **** **     **

3               YKKWKMAGYMKKWWWMYRRYWYYSMWMWRWRMYYTSYWRRKMYWKGCRYSRMRMMKYGSY
4               YKKWKCGAYMKKWWWMYGRYWYYSMWMWRWRMYYKSYWRAKMYWKCTAYSRMRMMKYRSY
                ***** ..********* ****************.**** *****   ********* **

3               KRRRKYRRYYYYGWYARRRYMYSMRYYRYGWYSWMRMYRYYMKMKWRCKKRMGCMWYRKY
4               KRRRKYGRYYYCTAYRARRYMYSMRCYRYRWYSWMRMYRYTCGCKTGMKKRMTYAAYRKY
                ****** ****   *  ******** *** **********    *   ****    ****

3               WYMRYYKKWCKAYYYRGAKGRRGAMRWGRRSRYKRMWATWWYYYMYWRYWRMMYTYCKWS
4               WYMRYTKKWMKRYYYRSGGRARRRMRWAARCRYKRMWWWWWYYYMYWRYWRCCTCYMKWG
                ***** *** * ****..   *  ***. *.******  ************    * **.

3               RKRRRWRYWRYYRKRKAKGAKKMYGAMWKYRWKSRWYKYWRCCCRTAYRTTTYMYRYRWY
4               AKRRRTATARCTGKRKTTRGGGCCARCTKYRWKSRWYKYTGTYSRCGYGGCGYMYRYRWY
                 ****    *   ***:. .    .   ***********    .* .*    ********

3               KYRRKGWRGKAGKTTCTGASWMYCACGGYWMRKTMTTTWRRAYGCYTTCATRGMMKRCYR
4               KYRRKAWARKMKKKYSKRMSWMYSRTATYWMRKCCGWGTGRRTKMCACYWWRACMKRYYR
                *****.*  *  *. ..  ****.  . *****       *     :    *. *** **

3               ARGKRRRYYTRRTSKCAAAYYGGWYYKTWKMWMMWSMYWYWMKAGGCTCATYWRRGTTRR
4               RRTGRRRYYKRRKSKTGGGYCRRACYKKWKMWMMWSMCACTMKTATYWMRKYWRRRYYRA
                 *  *****.**.** ...*     **.*********    **:.     .****   *

3               YRWMMYSSKGTGYRYAWAGTGYYMMAGKRKRYCYYYYCAKWKAKKRAYKAGYYATYTTAA
4               YATMMYSSKSWKCGTGWGKWRCYCMRRKRKRCAYYYYTRKWGGKKRGCKGAYYGCYACCR
                *  ******.     .*.    * *  **** .****  ** .***. *..**. *: .

3               AYTWCASGTYKWRRYSRGGGARKRKKGKKYKWMKSCCGKKTYRGACTKWTATCCGYRAGK
4               GTYTYTCSATKWGAYSRARACRKRKGKKKYKTMKSATATKWYGRMYCKWGTAAGKTRGTK
```

```
  .    :..: **  ***. ..****  **** ***. ..* *     ** ::.  *.  *

3   TWMRYATTKYRKWYYRMMRGGTKRRTYYKKWYGTGTARMRRYCAGYTYTWCSRKYTSYAK
4   YACGCRKCKYRKWYYRMMRSKKGRRWYYKKWYTATCGGMAACYWKCWCYAMSRKYWGCWT
          . ***********. . ** ****** :  . *            ****  . .

3   TGRKRYRATCKRYMKKKTKKAYGCACTYRRYRWRGKKRCAMWAATTKKACTWGKATTACT
4   GARKRYRWWMKRYMKTKYKKRCTTGTAYRRYRWRAKKRAGAWGGYCKKWMWWATCCCGAK
     .*****   *****.* **    . :*******.***.. *..  **   *...  ...

3   MTATRRKTRKYTAGCKRMKGTARGGGCMMMRMRRYGGARMRTGYWKMYRRRMMSTAGKGC
4   MKRYRRKCRKYCWRYKRMKACCRKCAYMMMRMRRYAAGAMRYSTWKMYRRRMMSWWCKRM
    *.  *** ***    ****. .*  . ********... ** . **********   *

3   YWMMRKCGATWAGKACGCWKRKRRTTCWWCYRYKARGTAWTGAYKARKMTAAKYYMYKRY
4   CTMMRKTTTYWWAKMYRYWGRKRGCWYWWMYRYKGGACWWCKMCKRRKMKMMKCCCYKRY
      ****  : * .*    * ***    ** ****. .  *    * ***.  *   ****

3   KRKKTAYKYWKCYYGATYKMGKRMGKMYYCACMYRYWMYRGTACAAKRKACCCKKKMGKW
4   TGKKCTCKYWKTTYCRCYKAAKRMKKMYYTCTMYRTWMYRAYMGGWKGKGTTTKGKMKKW
    . ** : ****  *   ** .*** **** . *** ****.   . * *.   * ** **

3   YRRMYRGYWSTCAGKAGKYAAKGTGYYRSYMYYGTCTCCTTRMRYMCGTTKYYACCGCGK
4   YRRCYAKCWSAATAKMAKYWMKKAKTYRSYMYTRWYCGTCCGCRYMATGGKYYTTTRMKK
    *** *   **:.:.* .**  * :  ******           ***.   ***:     *

3   KYRRTYTTSYRRKRKRRYCTGKYRWRGRYCAWCATCRKTRCCACAYAKMRMRMRYRRYCG
4   KYRRYTYKSYRRKRKRRYYYRGCRAATRYSMWTRYTRKCRTTGMGCRKMRMRMRYRRYYT
    ****   .**********     *   **. *    ** *  . .  ***********

3   MRKKYRYKTTWYKYKYRACYYKYYYYSAYWAKKSMKMYWWRYRGCKYKYRMKKYYKRKRC
4   MGKKYRYKKCWYKTKYARAYYKYYYYSGYWWKKSMKCYWWRYRKYGYKYRMKKYYKRKRY
    * ******. *** **  .********.** ***** ******   *************

3   TTKYWYSYCYCCTCAAGYYTTWYKKKKARYCTTMCRWKMKCTCAARAWGGCASKYKSRTS
4   CWKYWYSYYYMYCTGGKTYKKWYKKKKTGCAYWMYRWKCKYWTRCRMWRRMWGGYKSRKS
      ****** *    ..  *..******:  .  * *** *    .* *    . ****.*

3   RSMGTYWKWTTWGMWKGTGAGCRATRWRGACTTRYWRYYTGRRKKRYYGCYTKYMGYRKK
4   AGMCKYWKWYYWRMWKKKTCAGGRYRWRRGYKGATTRYYGARRKKRCTAYCKKYMCYRKK
     .* .****  * *** . ..    *** . .    *** .*****  .  .*** ****

3   WYKMMRWCMMKYRRRYMRWRMYYTKRRATTAKYCYWYKYKKTWWSMKKKGRYTAATKRKR
4   WYKMMRWGMMKYRRRYMATAMYYWKRGWYWRKYGYWYKYKKCWWSATKKCRYCTTWKRKG
    ******* *********   *** **     ** ******* *** .** ** :: ***

3   RKAKKWYKKRYKKTYKKRWAKGTYYCCCCYRWMWCATSMYYMMKMWWKKKKRRMKRSCKRK
4   RKMKKAYKKRYKKYYKGGTMKRKYYTTYSCRWMWMRYSMYYMMKMWAGKKRRMKASYKRK
    ** ** ******* **    * .**   . ****   *********  ****** * ***

3   YKYWWRSKKKWMYRGYTKWKRKGYKRYGKGAYRKACAYSRMRRYRRKWTGGWYYRSRWKR
```

```
4                 TKYWWRSGKKWMYRKYWKWKRKKYKRCTTRMCRKMMMYSRMRRYRRKWYKRWYYRSRTKA
                   ****** ****** * ***** ***  .   **    ***********   ****** *

3                 TCCRYGCKGKYWKYKYRGACCRYWTAARRTRRCGWMRSRTAKRGAGYRCTARWAAKKKTR
4                 GTYRYTAKKKYWKYKYRAMYSRTACGGRRARRMRWMRSRCGGGKWAYRYYGRWRCKKKYR
                     ** .* ********.  .*   ..**:**  ***** .    .**  .** .*** *

3                 YYRMKSYRSYRMYACGYGCKYSRCTGTSRKTCKMRYKRKRGAGTGYMRYCRKRKCTAGSA
4                 YYRCKSYRSYRMTTYATKYKTSRYASYCRKYMKMRYKRKRRRRYKYMRYMGKRGYWGSSR
                  *** ******** : .   * ** :. .**  ********     ****  **  ..*

3                 YMYYYWMRYTGGMYYYKTYRTYRCGYWRRKYKWSGKRACAAATARRCTWARGTTGCYYMC
4                 CMYYYWMAYYTKCTTYKYYRCTRTAYWGRKYKWSKKRWMWRRCGGRTCTCRRWKATYTMY
                   ****** *      ** **  * .** ****** **      . *   .*  .. * *

3                 CTYWWYRRKKKKMASGTTYGAGKYCYYGCYKKRKRYAYTGKYRWGYTSTAGWRCTMYKRT
4                 TWYWWYRRKKKKMTGACCYKWAKYYCCAYYKKRKRYGCKATYRTRCAGKRRAAMKMYKRW
                    ***********:..  *  .**   . *******. ...**   :..     .****

3                 KYKYRWYYYMMMRKRWMMKRKKSMWRMKRKMCGAAGTCYRYTTTKKYWCYYMRWKCYKKY
4                 KYKYRWYYYMMMRKRWMMKRKKSMWRMKRKCTTGCTGYCRYCACKKYWYYYMRWKTYKKY
                  ******************************   ..    ** : **** ****** ****

3                 GTSYWRKMKKRMYKRKWYKRMYKYCGKRSRYMRYTMTCTAAACCAKYRYMYRRWWSYKMS
4                 ACSTWRKMKKRMYKAGTCKRMYKYMRKRSRCMGYCACTWRWRMMWKYRYMYRRWWSYKMS
                  . * **********    ******  **** * *           ***************

3                 YYKRKKCCKKCYRYRKKKRKSKKKKMKRKYKKACMRKYGKKWYWWKRRTRKYYRRKSYRS
4                 YYKRKKTTKKYYRYRKTKRKSTKKKMTAKYKKWMARTCAGGWTTTKRRKRKTYRRKSCGS
                  ******  ** *****.****.****. ****   *. .  *   ***.** *****  *

3                 KGKKKRCYKYYRYKYMTYKKTCACKMAAAYMGGACYKSKYKGYKWMTRCMTRYCKKYMRY
4                 KTKKKRYYKYYRYKYCAYKTKYRYKAGRTYMATGTYKSKYKKCKWMCRTMCRYTGTYARY
                  * **** ******** :**..   * . :**. ****** *** * * **  .* **

3                 YSYCMTTKMYRKMAMCRRMKKTTRWYRKYRYYRCATCMGCKARYYYRMRGRWKWMRYTTG
4                 YSYYCCCKMYRKACMYRRMKKGCGWYRKTRYYRYCAAMAAKGRYYYRMRKAWKWMRYYYR
                  ***    ***** .* *****    **** **** .:.*..*.*******  ******

3                 GAYTYGKKYTAWMTGAMRYKYKRRGWSYMYMWMGTACKTRRMMRYWAGYCMKSWWKYMKK
4                 RGCKYKKGTARWMCAGMGCKYKRGTWGYAYMWCSWRMGAGRMMRYWMRTYMGSWWKYMKK
                   . .* *  : **  ..*  ****  *.* *** .    : ******    * ********

3                 RCGRTKKRKMRYRKRTKKYYRKMTYCAAWRSTCTYYKKGKRYYYYRMMKYRMYTKKYRKY
4                 RYKGGKTRGMAYRKRYKKYYRKMGYSRMWRSAACYYKTRKRYYCYRMMKYRMCYGGCAKY
                  *    *.* * **** ******* *.  ***:. ***. **** ********      **

3                 YMWGKKYRRYRMYMRYKKRKYCTAWKRYSWMRRYYWTYKSYAAMKYRKCCKRRAKKGAYT
4                 YMWKKKYRRYRCTAGTKKRKYYCWWKRYSWMRRYYWYYKSYRGMKYRKTTKRRMKKKTCC
                  *** *******     *****   ************ **** .*****  *** ** :
```

```
3    TGKYWRYWYRYYRMWKYWMMYRRWSYYWCAYRYYWYCTKMRRAKRRKGARKWWRKKYKKY
4    CKGCWRYWYRYYRMWKYWMCYAAACYYWARYGCCWYYYKMRRWKRRKRMGKWWRKKYKKC
         *************** *   .***. *   **  **** ****   *********

3    KYKRKKYGKMTTTTMYMMKYACYKYRMKWGGTGMKYYRKTGRYRWGKRCATTARKWKRKC
4    KYKRKKYKKMYYYYMYMMKCCSYKYRMKWRACAMKYYRKKKRYGWAKGSWCKMGGATRKA
     ******* **    ***** ..******* . .******. ** *.* .  .    .**.

3    GGYYYRKTCGCAGGGAKKMKTYGATCWTRYGAMTTGMSWKYRYYATRMWMASMTRWYMMR
4    AATTTAKYMRMRRRRWKKMKAYAMCYWYGYKRMWKRMSWKYRCCTCRMWMRGMWRWYMAR
     ..    *         ****:*.   *  *  * . ******  : **** .* **** *

3    MTYAYCYYTTYKTTYSAWKYYYKRRRRWYYSATGWKKRYKWKRWRKKYMRYKYRYYYGKY
4    MCYGYMCYGACKAGYSMWKYYYKRRRRWYYSRYKWTKRYKAKRWRKKCMRYGYRYYYKKY
     * *.*  * : *: ** **************   *.**** ****** *** ***** **

3    RRSRKYMYRCAGCAYAGMKYYYRYMMYRRKKRYYYKSRRWRRWKYYRMYMMYYTMYKRKY
4    RRSGGTCYAYMTTMYRRMKCCYRYMMCRRKGRTCTKSRRWRGWKYYRMYMCYYKMYKRKY
     ***    *      *  **  ***** *** *   ****** ******** **.******

3    WYRTKRCTCGTYGATGKRKYYKYWWKAAKYYRYRRWKYKKKSRRKKKMMKKKYYKARKMY
4    WYRKKRACGAYYKCCAKRKYYKTTTKMWKCCAYRRWKYKKKSRRKKKMMKKKYYKGAKMY
     ***.**.  . * . .******   *  *   ***********************. ***

3    YYKKYKKWTYKRKYKRKGAKKYGKMMRKMYYRSRTARYWKKKMCWTCARKWWSYKTCCKY
4    YYKKYKKAWYKRKYKRKACTKYATCAAKMYYGSGKRRYWKKKMYTWMMGKWWSYKCTTKY
     *******  ********...**..   **** * . *******      ******   **

3    YKYMKWWWKGKGRWARTCGYMKKKKRWWKKRMKKYMGRKRGMMCMYYYWGYMRRRGCCCM
4    YGYMKWWTKRKRRWWAYYSYMKKKKRATGGRAKKYMKRKRKMMYCTYYWKYMRRAKTTTM
     * ***** * * **    .*******    * **** *** **   *** ****     *

3    WRYKKTKYRYMYYKKKTSCKTKYYCCARRYKYYSKGYWRKGYMKTKKYKYYKRKMAGCMR
4    WRYKGKKTGTCYYKKKKCAKYGYCYAWRRYKYYSKTCARGRYMKKKKYKYYKGKMTAMMA
     **** .*    *****...*  *  . ********   *  ***.******* **:. *

3    KYTKYYKAYKWRCAKKKYWKYYYWRKKTARARYMYYSRRKSRKMRTMRYRGCYWSTCTKK
4    KYWKYYKWYKWGTGKKKYAKTYCTAKGYWRMRYCYCGRRKSRKCRYMRYRCGYWSCTCKT
     ** **** ***  .**** * *   *   * ** * .****** * ****  ***   *.

3    TTGRMMYYAGGKKYRKWKKKYYTYKWRCCAASWWKYYRKWWKYYGCACTAGRWKWRCTGC
4    YYKAAMYYGRKKKYRKWKKKTCKYKWRGYMGCAWKYYRGWWKCTRTMTCGARWKWRYKRS
          ***.  *********  .****   .. ***** ***       ..***** . .

3    KYYKYSRTATAMCKYYKRATCGYWWYRKYAYRYKARRGYRWMTAWYCAKRKKCAYARRSY
4    KYYKYSRYWYRMYKYYKRMKMRCAWYRKYWYRYKGGRAYRWMWRTYTTKRKKGCCTRRSY
     *******    * ***** .    ***** ****. *.****   * :**** . :****

3    MKTMRGRYGGRKGYYKMYRYKCWRRCMYWTSKSSAKKKTYRRYRWMRGTGAYRARKGTTT
4    AKYMRCRYRAGKRCCKMYRYGMTRRGMYWWSKSSWKKKYTRRTGWMRKYSMYGGAGKYYC
      * ** ** . *   *****   ** *** **** ***  **  ***  . * .
```

```
3    TCRSRARYSWWKKRSMYYWRTKRGRCCYYMGYGKKRYRYYKKATCCGRWYYYKKKRKTKT
4    GMGCAWRYSWWKKGGMYYWRYTRKRTTYTMKYKKKRYRYYKKWWYYRRWYCYKKKAKGKG
        .  ******* .***** .* *  * * * *********     *** **** * *

3    KYWRGGMRKRYRWRSYRYRRTTKWYWRYWKYCACYWMMWTKRYMSKKYTWYYKYRYKRYY
4    KYWRATMGGRCRWRSYRTGAWKKWYWRYWGYMRMCWMMAYKRYCSKKYKWYYKYRYKRYY
     ****. *  * ******    .******* *    ***  *** ****.***********

3    KYKYCMKRYYYCYKRYAGSMTRKYSYKMMKARKYKYMKKKKYWKWKTKRMWSRYMRKMAY
4    KYKYYAKRCTTTCKRYRRSMAGKYSYKMCKGRKYKYMKKKKYWKWKCGAMWSRYMRKMCT
     ****  **     ***  **: ****** *.***************   *********.

3    CAGTKRKSRKWKCYTAATGRYYKRMKKRRTCGYACYRYGGAAGGYMGGRGCKCTTCKYYR
4    MRRAKRKSGKWTYYYMRYRGYYTGAKKRRCYACGTTRYARCGCTYMATAATKTYYTGCCR
        :**** **. *      **.  ****  . .  **. ..  **.  . *       *

3    ATKKRKYRYRMKKKAKRGGATYWRSMKMRKRCCYAKYRYRRWRMYCYRYKMMWYYYGMGR
4    CGKTRKYRYRCKKKWKRCKCKTTRSMKMRKRYSYTTYRYRRWRMYTYRYKMMWYYYCMKR
     . *.****** *** **  ..  ******** .*:.********* ********** * *

3    YRYWWWYRTCTGWRYWKGKKTWWYKKKYCCAKKKGWYRRCRRMCACKTTRWAYKKYWKKY
4    YRYWWWYRWYWRWRYWKKKKYATCKKKYTYGKKGRWYRRMAACYRSKGCRWGYKKYWKKY
     ********    ***** **    ****  .**  ****      .*  **.********

3    RSKMGYYKWWKWYCRYKYYGRKKWKYKKYKKCTTTKYKYKYKTYKYTWKYWKTGKRKMGK
4    RSKMRYYKWWGWCTGCKYYKRKKWKYKKYKKMYWYKYKYKYKKCTTKWKYWTWRKRKMKK
     **** ***** *    *** ***********    *******. . .****.  **** *

3    KRRKYTKYRAGAYKMSRRMKRRYKKGRYAKKKRAKTKRWTAYKKWYYWYYKKCCWKKTMM
4    KRRKYKKYRTRGYKMSRRMKRRYKKARYMKGGRGKCGAAGGYKKWYYWYYKKYMWTGYCC
     *****.***: .*************.** *  *.*     .***********  *.

3    TRYKKKWRCRYMWYTTMRYKKYTYYSGGAYKGKYKARRKKKWKMKGMKKRMKWWKYSRWR
4    WRYKKKWGMRYMTCYAMRYKKTWYYSRRRYKRKYKGRAKKKWKAKKMKKRMKWWKYSRAG
      ******  ***   :*****  ***   ** ***.* ***** * ************

3    TYKYKYYRKRKKMRGKRYYRWSAGRAYKGARWYTCCTRKMRRGAAGWRGWWKATYTYRCA
4    YYKYKYYRKGKKMRAKRYYRWSWRRWYGRRRWYKMSYRKCGARGGAWRRWAKWGCCYRYR
      ******** ****.*******  * *   ***. . **    ...** * *    **

3    TAATYYKCGGCATTCYWGRTRMMWSKKKRRKKTTGTTWYYMRKCRSYKGAGCSKKMRKMY
4    YWRYYYKMRRYTAYTYTARWAMMWSKKGAGKKKWKCCWYYMRKTRSYKATATCKKCAKMY
         ***    ::  * .*  ******   **.    ****** ****.:. .**  ***

3    KYWMYYWMKMYKYRRRKGCTYKSKYKAGTCAYTKRKKTKRRKYAYWKRRKRWRYSWGACG
4    KYWMCYWMKMYKYRAGKSYYYKSKYKRRWMGCCGRKKKKRRKYRYWKRRKRWGYSWKGYK
     **** *********  *.  ******    .   ***.***** ******** *** .

3    GGSRCCMRTSTCRSARGAYMYTAAACWKKSYWYRYGGCAMYRMYYYRRSGRYKKCGSYYY
4    KRSGYMCRKSCYRSMGTCYMYYRTCTWKKSYWYRYAATGMYRMYYYRRSRGYGKYKGCTY
```

```
                   *    *.*  **   .***  :. *********.. .**********  * *  .  *

3                YKSRRYRRRGGGYMRRYRWYYGGTKKWMRRMSKMRKCYYYRWKKMCYAYGYRTKKYRKSY
4                YKSRRYRRAACRTMRRCGWYYKRKKKAMGGCSKMRKYYYYRWKKMACGYCYRKKKTGKSY
                 ******** .   ***  ***  .** *   ***** ********. .* **.**  ***

3                CTGCWYMWRMMRWYCGTAKWGMYMWKARMRR
4                YCCSWYMTRMMRWYYCCTKWKMYMWKGRMRR
                    .*** ******   :** *****.****
```

**Supplementary Figure 5.** The pairwise alignments between plants 40 and 41.

```
CLUSTAL 2.1 multiple sequence alignment (clustal format)

40              YRYYYYYSCYWKWMKRYMRYWCTKRRCYYRYGAGYRKKYRYMKYRYMRMRYRRGWSRKYM
41              YATCYYYSYYWKWMKRYMRCTTWKRAYYYRYRCTCRKKYRCCGYRYCRMRYRRAWSRKYM
                *   **** **********    **  **** .  *****   *** ******.******

40              RSWMKWYTWGYMMKWYYGGTRRTYRYRWRRYMMKWYRKYYGGTTYWRYRRYYRRYWMRYY
41              ASWMKTTCATCMMKATTTACRRKYRYRWAGTMMKWYRTYYTTAYYWRYRGYYRAYWMRYY
                 ****      ***    . **.*****   ******.**  : ***** *** ******

40              RRYMYSKMWYRTCWWTGCYMYGYWWWYWAWRKYYTWMYSKGARKYYMYRRMCWKYMMRYM
41              RRYAYSKMAYRWMWWYKMTMYTYWWWYWRWRKYYWWMYSKKRRKYYMYRRMMTKYCARYM
                *** **** **  **    ** ****** ***** *****  *********  **  ***

40              KKRWMYWRRRKYMCRKMRYSYYYKRMMRSMYRCMYKCCWSYMYWYRSKYYKRYSYYAWRK
41              KKRWMYWRRRKYMMRGMGYSYYYKRMMRSMYRYMTKYTWSYMYWYRSKCYKRYSYYCWRK
                ************* * * ************** * *  ********** *******.***

40              MRRRYWMYYRRKKWSWKTTYYRYKRYWKWKRYYKRKYMRYGWYKYSWYWKGTKWMRKYWR
41              MRRGCWMYYRRKKWCWGWWYYRYKRTTKTTRYYKRKYMRYKAYKYSWYWKAWKWMRKYTA
                ***  *********.*   ******  * .**********  ********. ******

40              YKKYMWRKRYRGKYMRWRCYMYKRRKATWASMRRSWMMKYMWKMWRARARRYKMGMYRKR
41              TKKCCTRKRYRRTYMRWGYYMYGRRKTYTWSMRRCWMMKYMWKMWRWRTRRYKMKMYRKA
                 **   ***** .****  *** ***:   ****.*********** *:***** ****

40              YYYMMRRAAKACRCMYYRRTTRGMRRRGCRRRYRYMYYMKMMKWWYMYYYYRRCTTRYRK
41              CCCMMRRTGKMYRMMYYRRKWRKMRRRTAGGACGYMYYMKMMKWWTMYYYYRRYCYRYRK
                   ****:.*  * *****. * **** .     *********** *******   ****

40              YRRMRAGAGKTAAKRGWGGRRRKYKRKAKYYRWWYRKYRYCGTYWKKAGYYSATAYKGMR
41              YRRMRTAGAGYMRKRTAKAGAGKTKATGKYYRWWYRKYRYTAGYWKKGATYSGGWCGRMR
                *****:...    **   .   * * ..************ . ****.. **.     **

40              YYKYYKWRRSKKYTYRGMTYKRKKKRYYRYTTRGMYKMKRMKKCGAGRMRKRYTTTKMKK
41              YYGYYKTRRSKKYKYRKMCTTAKKKACYRYAWRRMCKMKRMKKSATKRARGRCCAGKAKK
                ** *** ******.** *  . ***  ***: * * *******..: * * *  : * **

40              KTTCGAYYSRYKYKTTTYYMMRYRRKGGCKCCMRYKCTAWMYAYSAKTGGRRGACARRMK
41              KYCTRCTTGGYKYKWYYYYMMGTGGKRAYKYTMRYKGAWWMCTYSCKKRRRARRMMARMK
                *    .  . ****   ****    * . *  **** : ** :**.*.  *      ***

40              AWCTCWGYAYTKYATRYMYCRTGRYWKAAAMRYMKRCCTWCRYKARYTMRWWAYSRMSYK
41              GWYCATKYMYYKYTGGYAYMRWRRYWKGRWMRYMKRSMYWMRYKMRYCMRWACYGRMSYK
                .*  .  * * **:  * * *  ****.  ******.  * *** ** *** .*.*****

40              RRYKYKRTTWAKKKKKMCKTCKMRRRKYARGGTKARTKRKKKYWMRWACACKYYKKYYMYY
41              RRYGTKGGYWGKKKKKMYKCTKMAAGTYRRKTCGCGWGGKKTTAGWGYRYKYYKKYYMYC
```

```
      ***  *   *.****** *  **   .* *    .    **    *.   *********

40    SRYYARTAYKKKMYRRRKSMKGTRRRRRAGYATKKKYKYYTAYYKKMWKYKRTKRYKMAR
41    SRYYTRGMYKKTMYARRKSMKKGRRRRRGAYGCKKKYKTYYRYYKKCTKYKACKATKMRR
      ****:*  ***.** ******  *****..*. ***** *  ****  ***  *  ** *

40    MSYKMYSKRYWRKYWKRYMTKGTKKKKRTKKYRKMRKRTGRKCSKTAYKKKRYKCCMKYM
41    MSYKMYSKRYWRKYAKRYMKKKCKGGKRWTKYRKMRKRAARGYSKYWYKKKRYKSSMKYM
      ************** ****.*  *  ** .********:.*  **  *******..****

40    KYKRMRRKRRYKGAAKRYKYYRKWKKKRMKRCTTGGKTYRCKCCCCKKKKTRSKKSRKTR
41    KYKRMRRKRRYKKRWKRCGYTAKWKGKRAKATCYKTGCTRYKYSTMKKKKYRSKKSAGYR
      ************   **  *  *** ** *         * * .  **** *****   *

40    AGKMKTTTKSRYKSYWYKWSMMMYYKYYGYMYKRYYTTRKWRRCGCMWWKWWYMKRYKKY
41    CAKMKCAATSRCGSYWYKWSMMMYYKYYKYMYKRYYYCRKWRRTKTMWWKWAYMKRYKKC
      ..*** ::.**  *************** *******  *****   ***** *******

40    YYCGTCGTCGWRKWSRWSYKRMKYYYRYSKKRYKRYRRKWYRMKKKYRGCRRYKKKKMTK
41    TCGTGTAWMKWRKWSRTSYKRMKCTYRYSTKRYKRYRRKWCAATKTYRSMRRYKKKKMAG
            .   ****** ******  ****.**********   .*.**. ********:

40    GTMMYCAAAKKMMKRATCCWGTRMKRKRCCGRSRYKMWTMWWKGTTYKRWKCCTKKWRKY
41    AWAMYYRMWKKMMTGMWYYWTWAMKRKRYYRRSRYKMACCWWKRWYYKRWKMYYKKWRKY
      .  **    ****.     *   *****   ******   ***   *****   ******

40    MKACYRKWWRKMRRYKRKYKRKKYKWTKCTTCTKYCKRYKKMAMCKGYYYYRKKYRKRKK
41    MKCTYGKTTRKMRRYKRKYKRKKYKWYKYAAMWGCSKRYKKAGCYKKYYYYAKKYRKRKK
      **. * *  ***************** * ::    .***** .  * **** ********

40    SWYKRYYRSYYYRYYKWKWWYMYYRKMMWCKMWWYSRKKKKKWRYRKSRMKWSKYSKMKK
41    CWYKRYYRSTYYRYYKAGWWYMYYRKMMWYKMWWYSRKKKKKWRYRKSRMKWSKYSKAKK
      .******** ******  *********** *************************** **

40    MMMSKKKRWYRKMMRRRAATKKYKYACYCYRKYTKYYMYYWAKCKWRKRKWRMKKCMKYW
41    MMCSKKKRWYRKMMRRRMWYKKYKYGYYYYRKTCKYYMYYTGTYKWRKRKWRMTGMCTCW
      ** **************   *****. * ***  ****** .. *********.   . *

40    MWKMYKKYYMKKRWAACTYMCSWAYYCWMRRGGYSKMMWSYKRYTCAYKKMMCYYTAWMR
41    CWKMYKGYYMKKRTRCTYYMYSWMYYYWMRRKACGGACASYKACYTGYKKMAMYCYRWMG
       ***** ******  .  ** ** ** **** . .    ***    .****  *   **

40    KYRCKAKYRMKTGRACMTAGGWYRCKYRWRYKKKMRYKKKWYRTYRRRAYYKWTGGARTA
41    GYRTTGKYRMKYRGRACWWRRWYRYKYRWRYKGKMRCKKKWYRWYRRRGYYKWKRRRACG
       ** ..*****    .     *** ******* *** ****** ****.****.     .

40    TYYYKRWSWRRGMGWYMKCRKKMMRGKYGKMRYMGRRYYMYMYKKYRYWMYYYYRYYGRR
41    AYYYKRWSWRRKMAWCCKYAKKACAKKYRKMRTMKRACYAYMYKGYRYWATTYYRYTKRG
      :********** *.*  *  **    ** *** * *  * **** ****   ****  *

40    YYSYRTCAARAAMAGCRYMACAYGWWMKKYKWMRRWKKWWYGGCYGKKKKRYYYRAAKKK
```

```
41                   TCCYRWAGGACTMRSYRYMTMCYTWWAKKYKWMRRWKKWWCARTTAKKKKRYYYRCCKKK
                       .** ... .:* . ***: .* ** ************* .   .*********..***

40                   RCTMRKCCYYRYMRRRYRKKWTAWYRGKTRYKKKMAWWRKMKKRKAGCWMYYYRSRRKKK
41                   RYWCRKMYYYRYMRAATAKKWCGWYGRKKRYKKKMTWWRKMKKRKMKAWMYYYRSRRKKK
                     *   **  ******    *** .**  *.******:*********  .************

40                   YWWARYKRSMWYRMRRWKMWYKRYYMKKKRKKKKYKSRYWKRRRMYWRKRYKRYKRKWAT
41                   YWWMRYKRSMWYRMRRWKMWYKGYYMKKKRKKKKYKSRYWKRRRMYWRGGYKRYKAKWTA
                     *** ****************** *************************  ***** **::

40                   AYWRKKRKKYCRYSCWWGKKAKYKKMYKWYMYRRKMKYMYCKTTCTYYKWRYKKRWMRRK
41                   GYWRKKRKKTTAYSATTTKTMTYKKMYKWYMYRRKAKCMYSKAAGAYYKWRYTKRWMRGK
                     .********   **.   *. .************* * **.*:: :******.***** *

40                   TTKKRGGSMRYYYKKKWRGWKRYKWCGCGTYKKSKMMSRKRKAGAAGKKYKKKKKYRSMR
41                   YWKKRRRSMACTCGKGWRKAKGCKWTKMKKYKKSKMMSRKRKRRMRAKGYKKKKKYRSMR
                       ***  **     * **  *  **    .************    .* ***********

40                   KMYYKRSYRRKKKSKRYKRRKYRKMMKMYKGRKYRMMRYKKKMSKYYKCYCCRSKYYRYG
41                   KMYCKGSYRRKKKSKRYKRRKYRKMMKMYKKRKYRMMAYKKKMSKCYKMCYSRSKYCGYA
                     *** * ************************ ****** ******* **   .****  *.

40                   MKCCCWMYRKKKYRYKKKYTTTGWKYWYYRGCTGTMGSKMYYKWRYMWGRKMRYYMTAGT
41                   MKTMMAMYRKKKYRYKTGTGWCTWKYTYYRRYCAYARSKMYYKWRYMWTGGCGYYMAGTC
                     **    **********.      *** ***   .   ***********     ***:.

40                   AKARRWRGKWYRTSCKMKRRMGRCWGCMRWRYWWGTRKTRWKKKWGGGKMKYKRKYYMYY
41                   GKRRRWRKKWTRGSYGMKRRMRRGTTAMRWRYWWRKRKGRWKKKWRKTKMKYKRKYYCCC
                     .* **** ** * *  ***** *   .******* .** ******   *********

40                   CTYKYMRWRYRRWRKTCGCAKKWYRYRKRRYRRTTCTCKGSGRYRKRAMKYKKWWKCGYR
41                   TKCKYMRWRYRRWRKKMKSMKKWYRYRKRRYRRYYYYYKACAAYRKRRMKYKKWWKTTYR
                      . ************.  . *************     *... **** ********  **

40                   MYSYCGTGMRRKKCKTYWYKWKKYKKWCRWKSWGWYYWRKGGCGTATYMMRRTMKYWRMY
41                   MYSYYRYRMRAKKYKGYWYKWKKYKKWTAWKSWKWYYARKRKMTAGCYMMGRKMKYWRMY
                     ****    ** ** * ***********  **** *** **    :. *** *.*******

40                   RYRGGKKRKYKGAKYYAKKRKYYRWARATWYMTYYMYRMTCCRRMMCTWRKKSGYKRACG
41                   RYRRRGKRKYKACKYYMKKRKTYRWRRWWWYMCYCMYRMCTTRRMMTCWRKGCACKRTTA
                     ***   *****..*** **** *** *  *** * ****   ****  *** .. **: .

40                   RSMKYKTCRYYAAKMRRYYKKYWRYSYRKSGWMKYYRRTTMTRYTTGTGACWYKYKRRRR
41                   RSMKYKCYAYCRGKMRGYYKKYWRYSYRTSSAMKYYAGACCGRYCCAKKCTWYKYKRRRR
                     ******   *  .*** ***********.*. ****  :   **  .. . *********

40                   YWKKYYRYKSRKRWMRRYKTCAYYKKAYMKYYRRYWRGYKMYYMMTKKRTTCCCMYCKYR
41                   CTTGCCAYKSRKRWMRRYKWYGCCKKGYMKYYRRYWRKCKCTYCCAKKRCAMASMYYGYA
                       .    ************  .  **.**********  *  *  :*** :  ..**  *
```

```
40              YRAWYYAGCGKKKKWRRMKMGSGKMMKGGCKCCKGCCMCRKWRWRYRWKWTTTSYWRKKM
41              YRGTCCGSYRKKKKWRRMKMRCKKMMGSKYKTYKATTCTRKWRWRYRWGWKWWSYWRKKC
                **.   ..  ********** . *** .  *  *.    ********* *.  ******

40              RRMRTKGCMYMKCTKWWSKKYWYRGYKYMCWYYMCYCYGKRSSSRRYMMMYRYYKKWGCG
41              RRMRYKTTMYMKSYKWWSKKYWYGSYKYMYWYYMYTACAGRSSSRRYMMMYAYYKKWATA
                **** *  ****. ********* .**** ****   . . *********** *****. .

40              GGTTAGRYRKKTACWKGMAYKRWYRGKYMMRRTRGCGKKMKKMGWATKWRRWRYYMTYWR
41              TKAGGARYRKKYGMTKRCRYKRTYRSKYMMRRCATTKKKMKKCTAGKKWRRWRYYMKYWR
                  : ..***** .  *   *** **.******     *****    ..*********.***

40              KTAGCYTCMYWYWKKRKKYKRKKMKYSYRKKYRRWMYYYKMYRRGKKGGKYGKYYRWMMY
41              TAWAMYYMAYACWKKRKKYKRKKMKYSYRKKYRRWMYYYKMYRRRKKRRKYRKYYRWMMY
                .: . *   *  ****************************** **  ** ********

40              TKKWCKRRWKTYTMMTKRGKYYYKTACCCMWYTGWYSRRYYKYKYKWKWATCYYYYCYRY
41              YKKWYKAAWGCCCMMAKRTGCCCKKGTAACACYRTYSRRCYKYKYKWKARCACYYYMYRY
                 *** *  *    **:**     *.. ..      **** ********   . *** ***

40              CSWRRYYGKMRYRGRYYCYAWGAMTTTAKYMYYMYMSRSRMMYMYKWTAARYRCCTKRMR
41              YSWRRYTAKMRYRTGYYYYGTTGMGACGKCACCCYMSRSRCATACKWACGRYATTGKRMR
                 ***** .*****  ** *.  .* : .*     ******     **:..**    ****

40              YKMTAMYSRRSCKKRRYKKWMRRMRYGTCWCYYMYMKKSYKYWCTYKCTRRWTAAKWWRR
41              YKMWTMYSRRSMKKRRYKKWCRRAACCCTWMYYMYMKKGTTYAMCYKTCGRWKTWKWWRR
                *** :****** ******** **      * *******. .*   **   **.: *****

40              YRWMAAATYYYGTKMRWYRRWGATSRRTKCCKRKYYYYTGYRYYKMTGAWYKMYRYMRYM
41              YRWMRWRYYYYRCKARWYRRTCGWSRRYGYYKRKYYYYARYRYYKMATGWYKMYRYMRYM
                ****    ***  * *****  . ***    *******: ******: .***********

40              WKKRYYYKWCRMYKKYKMGKYYYCYRKRRYGCMMMKSMYMYRMYCARWRKKKYKKRGYRW
41              WKKRYYYGWMRMYKKYKCRGYYYYYRTRRCKYMMAKSMYMYRMYAWGTRKKKYKKRRYRW
                ******* * *******   *** **.**   ** *********.   ******** ***

40              WKRMRKWYRRKMRRMMKWYYTKGCGGMMMKCKKKKRRACAMRSRWKYKRMKSCRYYWKCC
41              WGRMRTWYRRGMRRMMKWYYKKAAAAMMMKYKKKKRRWMRMRSRWKYKRMKCSRCCWKTY
                * ***.**** *********.*....**** ******   ***********..*  **

40              GKYKKKKYTYSRYYTAGTKYRRYSCYTACAMGRRTYRSRRYKAKYGTAYWWRYRGGRMKY
41              AKTGKKGYATSRYYKRAWKYRRYGYTYRYWCRRRGYRSRRYKGKYRKRCWWRYRRTGMKY
                .*  ** *: ****. . *****.        ** *******.** .  *****   ***

40              KYYYWMAGMKRYMYKYKRYRCKTWYYYRYMKGCKKRMGRKKKYCKRRMYTYCCKRWGTMR
41              KYYYWAMAMKRYMYKYKRYRMKCWYYYRYCGRYKKGCRRKKKCGKRGMYACTTKRWTAMR
                *****  .*********** * ******    **   ****  ** **:   *** :**

40              RRRYGYGAATMCATAKRAGACRGKMYYYYRYYMRCTYAGWCGTRWRRGMYCTYCTRWMGK
41              RRGYKYTWMYCAGCTKRWRCYAAKMYYYYRYYMRTAYGAWTKKRWRRSMYYWCYWGTMRK
                ** * *     .. :**  .  .*********** :*..*  .****.**        * *
```

```
40    KAAAKYAWKWMYWYRATCMGCRKKMWGYKCGYYWKKCTWTYRAKKMAAYRWMKKRKRRYR
41    KWGTKYTWKWMYWYATGTMATATGCTTYKSKYYWKKGCWKYRMKKMRWYRWMKKRKRRYR
      * .:**:******* :  *.  .    **. *****  *.** ***  ************

40    YRYKYGKYKRYKYRTRRYWMKRMWMKMMAKYMGGGYACKWSRRKRCRRRWKRRKKKYKWT
41    YRYKTRKYKRYKYRKRRYWMKRMWCKMMMKCMKRRYMYKWSRAKRTRRRWKARKKGYTTC
      ****  ********.********* *** * *   *  **** ** ***** *** *.

40    YWRRWKYKKYTCWRKMRGGKMKSYRWAYGTYRTRYKRRRMSRYYYYWYCCACYKARYTGT
41    YWRRWKYKKYYMWRKMRRKKMKCYRWMYAYTGYRYKRRRMSRYYYYWYSMGTYKGRYYTW
      **********  *****  ***.*** *.    ***************. . **.**

40    RWYWRWKMSYGSSKKWWKWSAGCARMMYRRAYYYYWRWRGMTGCKYMYYKKRRKYKKATT
41    RWYWRWKMSCRSSKKTATACTTTRRAATGATYYYYWRWRACCATKYMYYKKRRKYKKWKY
      *********  ****  . .:   *     :********.  . ************* .

40    TAWRYYYYYYKRKSTKKKRSKKKYYKKKCKWKYRRWYRRAKRCRMCCKKYSMKRRRYKYK
41    KWWRYCTCTYKRTCKKKTRSGKKTYKKKMKWKYRRWYAARKAMRMATKKYSMKRGAYKYK
      . ***    ***...**.** ** **** ********   *  **. *******  ****

40    KWYKYYYCKKWMYYMWYGYKKYRKWWAGGRMWTMRYGKCCKTKAKKGGAMKYMWCKRGKR
41    GWYKYYYSKKWMYYMWCRCGKYRKWWTAAACAWAGYAGGYKYKWKTRAWMKYATYGRAKR
       ******.********    ******:..      *.   * * *. . ***    *.**

40    RKSRYYYWWTCTCCWAWKKKRYMMKAYCKTWYWKKKKKWYKKWAGCK
41    RKSRYYYTTAACYMWTWKKKGTMMKTYYKYWYAGKKKKWYKKAGATK
      *******  :.   *:****  ***:* * **  ******** .. *
```

**Supplementary Figure 6.** The pairwise alignments between plants 38 and 39.

```
CLUSTAL 2.1 multiple sequence alignment (clustal format)

38              YKYYRRKRKYRMAMWKMYRMYYAGTWSRKRGTAAGKMRYRYYRCCGGCCMYYKYRWRKAA
39              YKYYRRKRKYGMRAWTMYRCYYGACASAGGRACMKTCRYRYYRMTRKMTATCKYRWRKMR
                ********** *  *.*** **..  *    :.  . ******         ******  

38              RRRRKRWWSRMKMYKKYYRTKYMRYTTGGGYYMKYKYWMWRCGMGCYMRYAMYRSYRKWR
39              AGGRKRAWGRMKMYKGTCRYKYMRYAYRRKYYMKYKYWMWRYRMRYYMRYGMYRCTRKTA
                   *** *.******   * *****:    *********** *  ****.***. **  

38              MRYWRKMGRYAGARYWKYMRYSYMGTTARRCMAWMATKRRRRYMYMWCCSKYRRMTTRRY
39              MRYWRGMTRYTATRCWGYMRYSYMRCCGRRMMRACWYKRRRRYMYMWYYGKYRAMAGRRY
                ***** * **:.:* * *******   .** *     **********  .*** *: ***

38              YCAGGKYWRYWWCCYTMMTCGTKMRKRKGMMYYRKMYTTCRCWYMWRATYMAAGYYWMRW
39              YACKRTTARYWAYYYMMCGTAKMRKRGRMCTCGKMCYKTGTTCATAWWYMWGAYYWMGA
                *..  .  ***   * **   :*****  *    **  .          ** ..****  

38              WYRKWRYRKWRAYKCMGWYWRRYRYRKCWMRYYRWCMRRWWMRRRSYYYRRKKRRRRTCY
39              WYRTWRYRKWRRYKMMSWYWRRYRYRKAWMRYYRWAMRRWWMRRRSTTTRRKGAAGGKTY
                ***.******* ** *.**********.*******.**********   ***     . *

38              RYYWATWRWTWKTTAYMMMMMRKTYRRRYMGKRCRMKYKYRWMYRYGRMMYWWSRRWSRK
39              GTYWWYAGTKWKWWGYMMMMMRKYYRRRYMRKRMRMKYKYRTMYRYAGCMYWTSRRWSAG
                  **     .**  .******** ****** ** ******* ****.  *** *****  

38              WMMYAGYRRTCYRWRYRWYYYWTWWMYMRMRKKRTTCWGRSTRYRKSYAYTGGGTCAWAW
39              WMMCRKYRRCTYRTGTGWYYYWWTWACAAARKKGYAAARRSYRTGKSCGTCATRYMWTRW
                ***   ***  **    *****  *     ***  :.  ** *  ** .  .       *

38              KRWTRWCYWYRKYSWYTGGRYWWYYARYRWWMKKRYYKKKGGYKYMTYRKSWKKYYYKRR
39              KRAYRWYYWYRKCGTYAACRYAATYRAYRWWCKKRYYKKKKKYKYMKCRKCAKKYYYKRG
                **  ** ***** . *:. **   *  **** ********  ****. **. ******* 

38              GYMYYRTTWRYKYKMCMKSSKMYRWRGYRYKYWYRWKMYRYKYTARKKCAGKYKRWRYKR
39              KYMYYRYCWRYKTTCYMKSSKMYRWARYRYGYWYRAKMYRCGYKMRKKSCKKYKRWRYKR
                 *****  **** .  *********  *** **** ****  *. ***.. *********

38              YKRWMYYRKKAGGTYKKRYYMRRKSKARGGAGRAYRTYMYWAYKRYMRYYKRKKAWTWKS
39              YKRACYYRKKRRRWYTKRCYCGRGSKGRAKTKRRYRKYMYWGYGRYMRYYKRKKWWKWKS
                ***  *****    *.** *  * **.*. : * **.****.* ********** *.***

38              CKTSYKKTWCGSYTKTYRRSRWYKYKYRMSRWWSRRWKMYKSRWYYGYCKYSYMKMCATC
39              YKGGCKKKWYRSYGKGYRRSRWYTTGCAMSAWWSRRWKMYKSRWYCTTYKYSYAKCARYM
                 * . **.*  ** * *******.    ** **************    **** * .   

38              ARYATRKRRTCTRWRYGRWYGACTMMWKKTTKKKKKYGWGRKRTYRRYRYYWGCTGGCYR
39              RRYRYRKRRKSKAAGCKRWCACAGMCTKTACKTGTGCKTRAKRGYRGTRYYATTGTKMYR
```

```
      **  ****...     ** ... *  *.: *. .      ** **  ***       **

38    SYGYGTKAMCWMYKKMYYKAYKKAKYRRYKRKMWKCGKTTYYWYTACCTYKYKKGTCRYK
39    SYKCKGGTCYWCYGKAYYKMYKKMKYRRYGRTMTKTTGYYCYWYWRTMKYKYKKTCYRYK
      **     :  * * * *** *** ***** *.* *      ***    .*****   ***

38    RYKYRKKMYTKGRRRKYMTAARWKRKKYRYYRCRWSRYCCKYYRGRMGRKYKYRYWKKKS
39    RYKYRTKCTKGSRRRKYCCCGRWKGGKCGYYRYRWSRYTTKTCRARMTRKYTYRYWKGKS
      *****.*  . .*****  ..***  *  *** *****  *  *.** ***.***** **

38    MYYRRWGARTARGTAGGCACRKAGKTYKKKGKCTRAKYMRMKGKKKYWTCKRCKMYMKTG
39    MTCGRARMACCGACGAATTYRKWRKWYKKKCTTKRGKYMRMKKKKKCTATGRYKMCMKCR
      *   *     . . ... : **  * **** . .*.****** ***  :  * ** **

38    GTAKKYWKYGGSTRRYYTGWARAGTTTWMSKAMKAWYCTACKYRCMCTKRARYCGYCRKR
39    ACRKKCAKCRACARRYYGRTWGRRWYYAMSTGCGMWCMCGYKYRTMYWGRGRCTRTYRKR
      .  **  *  ..:****           **..   *   . *** *   *.*     ***

38    YKGKTCARMRKTCYRRGCTKRWGGWATGKYWYYYSYYKCGTGYRMMWRRWYYRRKWKWKA
39    YKKKGTTRMGGKYYRGAACGRWTRWCGRTYWYYYCYYKMRYAYRMMTGGTCCRRKWTWKM
      ** *  :**  . ** ..  **  *.  .*****.***   .****      ****.**

38    YGYTWRRCRRKRKYYMKTYGGGGTAYMWYRMKRRRYRWCCCGRRMRRSKYYYSKRKGGSR
39    YCTAARRTAAKRKYYMKYTRSAKYGYMWYGMTRRRYRATTTTRRMRRSKCYYSKAKTAGG
      *  : **   *******   ..  .**** *.*****     ******* **** * ..

38    YRYYCGWKYMRCCTGSYWKCYKGTGGKCRKGYCKKSRKMRWWKGCGGYAWKYMWKWKRYK
39    YRTYYAWKTAAYMKKSYWKMYKRWRRKTATTCTKTSRKMRWWKATARTRWKYMWKWKRYK
      ** * .**     . **** **    *  .   *.********. .   ***********

38    AACCYTGKYGYTTCGRMACYTTTKYGYRMRCTGAATRYRRAGAARGWKTGAGKYKKYYGT
39    RRAAYYRKTAYWGTARMGTYCWCGYRYRMRAATMGCGTGAGTTGRKTGWKRRKCKKTTAG
        ..*  * .*   .**. *    * ****.:  .     . :.*       * **  .

38    AKMCYKCGCMKAAGGAYCYYGYKRAMWCKYWRYKKARRRKCACRCCTCRRKRCKAYTGAK
39    GKMYYKMAACTRGRTRCYCCRCKRGMAYKYWGCKKTGGRKYGTGGTCTRRGRYKTTGATK
      .** **  .. . .         **.*  ***  **:  ** .      ** * *:  .:*

38    RYTTACYCKSTKRKRTWYCRGWGYKRTWKYRSKYKKWYMYYKRMWKKRRWRKRKRRGGCT
39    GTWWRTTTKSGKRKRKWYTRSAAYKAYTKCACKYKKWTAYCKRMWKKRGARKRKRGRATC
              ** ****.** *. .**   *  .*****  * *******  *****  .

38    RRYTGAGYRMRGWAGGSKWSKYRCYYYMMRKRMACGCAKKKCCWCYACTKKYRRMKKTAA
39    RRCARTCYRMRATCTASKWSKCAMYYYCMATAAWYATGKKKYYWMYRSWKKYRRMKKCCG
      ** : : ****. . .*****   *** * .    . .***  * * . ******** ..

38    ATYCAAWCGCCRRACCTGGGYTGCGYTGGWRATYRTWTGKWKRMKKGGYSTWYRYRRRYA
39    RYYAATGTATTTGGTTSYSARCYATRCYATTGCKYGCWGKKTGRMKKCRYCWTYRTGGRYR
        *.:. .     : . ..   .    .   ..*  *  *  ****  *.  **   **

38    RWRMMRGRRKKGTWRKYRRACYGYTKYMRRRYRRRYKKYKRGRYKWRWMYYKTRTACTMT
```

```
39          RWRMMRTGRGKRKWRKYRRTACATWGYMRRRYRAACTKYGGARYKAGAMYYKKRGRYAMW
            ******  * * .******:. .   *******   .**  .***   ****.*   :*

38          TRKRWKRRSYGRKGYTGTYYCGTACGGAGAARKATAKYWSSYMKYWKKGYWTWWMGKWKT
39          KRKRWGRASYCGKKTYAKYYYRYCTAAGTCWRTRYWKTASGTCKYWKKCYWWWWMKKWKK
            .**** * **  *   ..**   . ... . *.   *  *.  ***** ** *** ***.

38          CCKYRYYWWKYYRMRWKSGYKKTCKYYKYRYKYRYKMYKMRRWWMSSYGRMTKRYASRYY
39          TTTYRYYWWKYTRMRWTCKTKTWYGYYKCAYKYRTGCCGCRRWWMCCTRGMCTAYRSRYY
              .******** ****..  *.   ***  ****      *****..   * . * ****

38          WKYRKMKKYCKMCCWMGTWTKWGYKKRGRKMYYGRYTTKYKRYRATAAWKMKKTTYRKKW
39          WKYRKMKTYYGCAMWCKCTKTAKYKKGKAGMYYKACACKYTRYAGCGGWGAKKWWYRGGT
            *******.*   . *    ..  ***    ***   : **.** . ..*  **  **

38          WAMYKRRYKRYKTCCMKYYKRRKGMGKYKKKWRSYYYMYYKKKKYSKRYKGWTAMKKKYS
39          AWMYTRRYKRYKWYYMKYYKRRKCARKYKKKWRSYYYMYTKKKKYSKRYKRWYRMKKKCC
              **.*******   ********   ************* ********** *  **** .

38          KGKYYRKKWRGTKYKCAKRYKKRKTWCWYWKRCRKAAYRMKWRWYAKRYRKYGMMWTKKY
39          GKKYYRKKWRACKYKMWTAYKKRKKWYWYWKGMGKRWYRMKWGTCTGRYRKYRMMWCKKY
              ********. ***  . *****.* ****   *  *****   : ***** *** ***

38          RWRKCYAGYMAYRRWKKTKTKMAGYRYMRWYWKMTATRWYMRYRKCGRCRKSYKKWYYTT
39          GARGMCCTYMMYRRWKKYKCTCRACRYMRWYWKCAGCRTYMRYRKYRRTRKSYKTWYYWW
              *   . ** ****** * .  . ******** :. * ******  * *****.***

38          RWAWYKTYRGGYRRKKYKKYRRKRCTCCCCKCKWKTGMKYWWRKARKRRMKKYMRWWCRW
39          RWWTYKWYRRRCARKKYTKYARTRYYYMYYKYTAKYRMKYWWRKCAKRRCKKYMRWWTRW
            **  ** **    ****.** *.*      * . *  *******. *** ******* **

38          KRYKRMCRSCTMKGCTTTRMCTTKSKCACGKYYGGMCYTRRMKKYYYMRWRRMYRKCTRK
39          KRYKRCYRSYWMKTTAAWRMTCYKSKYRYRKYYRRMTCGRACKTYCTARWRRMCAKTGGK
            *****  **  **  :: **   ***    ***  *   *  *.*   *****  *   *

38          YRYMKKRRCCCTWWTKAAKTRCWCRTYTARKWAYGGKRWKWWRMAWCCSYYYGRYTTTAK
39          TRYMKKRRATTGTTYGTGGCGTTMRCTCGRKTWYKKKAAGTARAGWTYCYCCAGTCGATG
             *******.       :.      *   .**  *  *     * .*  .*  .    ::

38          RWGKATYKWYTSKRTMCTCAGMYRCYTTGMKYYYSRKMKYGCAAWGYMKKMWYYMTCCCY
39          GARKTCCKTCCCKGWCYYMMRMYRYCYWRMKYTCGGGCKYRTTGTRYCTTMWYTCKMMYY
               *:  *   .*        ***     ***  .   **  :.  * ..***  .   *

38          MYRRRTMYGMMKMKMSMKKYKKMRKMKCTKCAAGACAYCKKKCKKYYTMACTKYRSYCYK
39          MYRRRKMYRMCKAKCCAKTCKKMRKMKMYKYWMRRYRYTKTKMKKYTWAGYYKYGSYYCG
            *****.** * * * . *. *******  *       * *.* ***   .  ** **

38          YTCYRCYKCRTAGTYYYRMKKKAAAGRKYCGKKWRRKKYMRTAKWTAKCSYRRKWMKWWM
39          TYYCGMTKTAGTACYTTRMKKKTTTTRKCYKKKWRRKKYCAYRKWKWKYSYRGGTAKTWM
                   *   :. *  *****::: **   ********    **. * ***    * **
```

```
38          GKTMYCTCRMKAYRYCKSRWKRKKAKARYYRKSKKKRRKKYYKKKKGTRMRGCCGTCAKY
39          RKKMYMYYRMKTCGCYKSRWKRKKMKRRYYRKSKKTRRKKYYKKKKCCRMRRYYRYGGKT
             *.**   ***:    ******** * ********.**********  ***      .*

38          YRKYRMKWTAKKKTTSYMYYKKWYRAGTCAAGKKRYSCAYMGTYTTGTAMYTCTSRKKYT
39          CRTYRMKWCTKKGCCGCMYYKGWYRWAAAGGTKKRYSYMYCKWCCWRYTCCAYCCAKKYC
             *.***** :**   . **** *** .:... *****  *        :  :  . ***

38          RAASWWCMMACGYSMWYRKTRTRYKRSKTRRRYRCKWKKGMRYGRRYMKYWKYKGTGKWR
39          RTGSWWYMMRTAYSMWYRGCGCRYGRSKCGRRTGYGWKKCMRYKRRCCKYWKTKKKKKWR
            *:.*** **  .******    ** ***  **    *** *** **  **** * . ***

38          KRGTYGGYYYYKRARYYRRRCGCTACTCYAGCTTYGCWKGGAAGWRKKKSMYWTKRTTAY
39          KRKGYKKCYYYKRTGTYRRRTTAGGTYTYMRTWCYRYTKAATGRWRKKKCMCWKTRYCTY
            **  *   *****:  ****  . .   *     *   *..:. *****.* *..*  :*

38          CTTRMYWWKYKRKYMMRCTRRATRWKWMYYYWMWSWGYCTCYRSCCSSYRYWKYWWSKAK
39          YWYRCCTTKYKRGYMMRYWRGGCGWKWMYYYWMWSWRYMCTYRSAASSYRYWKYWWSKRK
               *    **** ****  * .  ************ *   ***..************ *

38          KKYGTATYRCCYRRKRWYRARYKGYMMASRKMAGTCYSYKKTTYMYKGKRAGMYKTTAAC
39          KKCRYTCCATAYGRGRWYRGGTTKYMMRSRKMGACTYSCGTACTMYKKGRMRMYTCCGGT
            **   :    .* * ****.  . *** ****..  **  .:  ***  *  **.  ..

38          KKYYKMKKCGTAARKKKKKGCGCRSYWRKGKKKTCAKKKYRRMKRYWKKRSCARTCYYYY
39          KKYTKCKKSRYMRRKKKKGTTTTRGTTAKRKKKAYRKKKYGGMGRYTKKGSYTRGTYYYY
            *** * **.    *****     *.   * ***:  ****  * ** ** * :*  ****

38          CRGGGCCYCYCTMARWYGACARWKKKKRMYYYWKKKKKRYRRRRAARWKRWCKMMKRYYM
39          YRKKAYYCYCYWMGATTKGYGRWKKKTRMYYYWKKKKKRCGGRRMWRWGRWAKMMKRTCM
             *  .       *.    . .*****.************   **  ** **.*****  *

38          KKAGYAKRMKYKMRKYKYAKRTGTKRRKSTGTATRTKGSKYYCCMKRWMSGCMKWKSACT
39          KKMRYMKRMKYKMAKYKCRGGYRWGGAKCYRWRYGYKRGKYYYMATAACSKYMKWKSGTA
            **  * ******* ***          *.       * .***   .   *  *****. :

38          TGYKYKGGYYYYKKKCKTWSKYKRKYSSKYWYGAGYRYKYYRKYCKKWRRKWRWCRTCRR
39          GAYKYKKRYYTTGKKTGCTGKYKAKYSSKTWYRRKTGYKYYRKYSKKWRRKWRATGATGG
             .****  **   **    .*** ***** **     *******.********   :

38          ATTKRRMYRKCRKWKCGWYRCKKKKAGAWYRYYGYYRYGCYRKYKKSWGRCKGKMRKKTG
39          TGCKGRMYRKTRKWTTAWYRTTTKGWSRTCGCCAYYRYTTTRKYKKSTCATTAKMGKKWR
            :  * ***** ***. .*** ..*  .      .****   ******    ..** **

38          RKMRRSYKRYRRTYYWYRYGTKTYATAYSYAATKYRKGRYKYKRYRRGKKYKMRKKCTWM
39          RKCARSYGRCARAYCWYRYRYKCTCYGCGYWWWGYAKARYGYGGCRGAKKYKMGKKTAAM
            **  *** *  *:* ****  *  . . .*    * *.** *   * .***** ** : *

38          RRYRKTYRYWKMRKYYKMTRYWYKYKYRGRKRYMARRATYGYTGAKTCGCRGCKKRWKYY
39          GACATGCGCAGMRKCTGACRYWYKTGYRARKRYMGAGTYYRYKRWKCTAARAYGKRWTYC
                .      ***     *****  **.*****.  : * *.  *  ..*.  ***.*
```

```
38          MWYYAATRWCAMSGRTCYMMKTYSRWAARWCYYMGRKGMCCYKYRMYWKGTTWRKKTAGA
39          CTCCRWYGTTRCSARYACACKCYGGTWMAAYTYMRAKACTGYKTRACAGRAAWRKKAGAC
                        *.* .   * *.        **  *.   ** *     ::****:...

38          KMKKAAAAAKRKCTAYKMKKKKWARRKCYRGRRRATKKCTYRYYWRKKRKKWGGTWKRRY
39          KCKKCWCRRKAKYYWTKMKKTKARGRKYYRSRRRCCKKTATAYYWRGKRKKWRTCAKGRT
            * **. .  * *    ****.*   ** **.***. ** :  **** *****    * *

38          RKKKRKYRRWKYTRRYWYSYKKMRKKCARYYKKKARRYMRCGGYKKKMYKGGCKTGCMYT
39          GKKKRKYRRWKYCRRYWYSCGKCRKTTRACTKKKGRRYMRMRRYKKGMYKKAYGYKYMYY
             *********** ******  * **.     ***.*****   *** *** .     **

38          GYKRRAKTGTKYCCYCAMWYYKCRGGYMRWKSSKKWYRAKRWRTMMWRYRKAYGYTKMRK
39          RYKRRMGARWKCYYCTCCWYYKMRKRYMRAKSSKKWYRCKRWRGACTRYGGRCATWKMRK
             ****  :  *     . **** *  *** ********.****    **    .  ****

38          KKGRGGYYYRCMMKAMKKKYWKKTGWKWKYRKRKRCTYYYRYSYYYKRGCKKKRTARKAA
39          KKRRKRYYTAAAMKMMTTKYTTKYAAKWGTAKRKRYKTTCGYSYYYGRRYKKKRYRRKMG
            ** *  **  . ** *..** .* . **   **** .    ***** *  ****  ** .

38          RMYYKYYKRRKCKRRCKTYYWKCTCCYKYYRKKYKYKKRKTKYTKRMYRYMKMKGKKKWT
39          RMYYKTCKRRKAKRGMKACCTKTCMMCKYYRKKYKYKKAGKKYKKAACACMKMKAKKKWW
            *****  ****.**  *:   *     ***********  .**.*     ****.****

38          AGKKMRRARMKMKRKMKGCGGKRRYMKATTRYKKRKKGGKKRRRAMSKAWSWMKKWRKKY
39          WRKKMGGRRMKCGGTAKASAKKRRYCKGCYRYGKRKGKRKKRRRTMSKWAGTATKWRKGT
              ***   ***   . *... **** *.  ** ***   *****:***  .  .****

38          YYRKWYAAYWRYGRYYYKSRYCWTTGTYSYKWAWATGKTTTRKMYRMTRYGRCCYYKYYR
39          TTAKWTRWCWRTAGYYYKCGYATYKSKTCCKWRWTWRKYWYGKMYGCGRYRRMYYYGYYR
               **    ** . ****. *.  ... . ** *:  *    ***   ** *  ** ***

38          RYWTTCCSARCGYYK
39          RYAYCYSGMATACCK
            **    ..   .  *
```

**Supplementary Figure 7.** The pairwise alignments between each pair of the plants 35, 38, 40, 42, 43, 44, 45, 46, 47, 48, 49, 50, and 51.

```
CLUSTAL 2.1 multiple sequence alignment (clustal format)

The pairwise alignments of two accessions 35 and 42

35              KYYYRYYRMYSWGTGTKWYTMWYYMKRGYGGRKKYYGCMKWYKMRMRYRGKMKYYRAYMA
42              GCYYRTYRMYSWRGTGTWTWAWYYMKRRTRRATKYYATMKWTKARMRCRRKMKYYRGCMG
                  *** ******    .*   ******     .***. *** * *** * ******. *.

35              CRKRRSYWMRATRKCWRYYCMYAGMWKMTYATMRRWWWRKGTRMTCCTAACRKRYYRRCA
42              GRKRRSCWCGMGGKMWRYYMACWKMWKMKYTAMRRWWTRGRYAAKAYWGGYGKRYYRRTW
                 ***** *     * ****     ****.*::***** *      ..  .. ******  

35              TSRRCTAAGKWMWYWYYYKKMYRRRRRWRRRYMYWTMWYCAAKSRKSCCTAYWYRYRYMR
42              KCAATWCWATWMWYACCCKKACRRRRAWRRRYMCWKMTCYRWKSRKSAMYRYWYGCGYMR
                ..    . ..****    **  **** ****** *.*     *****.   ***   ***

35              KGGWKAYTCCMSCAAMYYYYYYYCRWKRMRRRWCMKRRYRGTRYKSYRYCYYMRSGGWYM
42              TTRWKRYYYYCGYRWMYYYYCCCARAKRMARRWYMKRRTRRWRYKSYRYAYTMRSRAWYM
                .  ** *    .   *****   .* *** *** **** *  *******.* *** .***

35              YKYKKRRTTRRAASGKRYWRKYRYKMYTRWGCWRKRYTTTGYMRYRRAYKRYGMRWYYYY
42              YKYKTRAYYGRGGGTKRYWGKTATKCCYRWAYWRTRYCACKYMRYRRWYKRYAAAWYYTY
                ****.*    *... **** *   *   **. **.** :  ****** ****.  *** *

35              KKTRCGRYGRYMACGYTYTYYKRYRSMCMRYMCCYCCKRWWWRWMWYYYRSRGATSAYYT
42              TKYGMSRYTRYMRMTTCTCYYKRYRGCSMRYMTYCSSKRWAAAWMTCCYRSRKRWSGCCG
                .*   .** ***       ******. .****   ..***   **   ****   *.   

35              GMYKRYMYKRYRMKARYMRRYRRRGKCYYWTYMRAGYGMRKMMWMRYYKGWRWYRWRYYT
42              RMCKRYMYKRCRCKRGYMRRCRRGRGTYCWYCCGGKYKMRKMMTCACTGAARWYRWRTYY
                 * ******* * *  **** **    * *   .  * *****      . ****** * 

35              TCCATYGKWYSRRCRCMRKYGTYMYKYRCCKAKCGAAATGGCCKKCYSRAAMGMYAGGAR
42              YATWCCSGACSRRSRAMRKYSKYCYKYRTSKMKYKRRCWKCTTKKAYCGCTMAMYRRRCR
                 .    .   ***.*.****..* **** .* *     .    **.*. .:*.**   .*

35              TCCAYKYKYRYKWWRWTGKKKTRGMYCAGCKTMTACCGAKSKYGKKTKCYMRWRKYMRKR
42              YYYRYGYKYRYGWAATAAKTKWRRMYYGCMTYCYWATKCGSKYAKTYKYYMAWGTYMRGR
                    * ***** *   :.*.* * ** .  .    .  . ***.*. * ** * .*** *

35              MRRRTAKCTCGKRYAAWGAYYRRRKGCGWKKMRKKWTGAMKWACYRRRTTWKKCCGYMCK
42              MRRAYRKYWMKKRYRMWRMTYRRRGRATWGGAAKKWYRGAKWGTYGRGWCTKKTTRYCMK
                ***   *    ***  *   ****   .*    ***  . **. * *    **   *  *

35              YRYTATTAMKRWRYKRTTMRWKRWCAKAGCKCMYGRKGTGGYYRTCTCYWTKARTYKMYY
42              YRYCMWWMMGAARYKRWCMRWKRWYRKGATKSMYKRTAYKRYYRWYCMYWGGRGCYKMYY
                ***     *   ****  ******  *.. *.** *..   ***    **     *****

35              RMWYRCCCTRTKGACRMWMYKGWARYKRTCWCCWRYMYTRSRTGCMMAGWTYWMTYTCSC
```

```
42    RCATASYMYRCTTRMAMWMYKSWRRYKRYYAYMWRYMYWRGGKRYCAWAAATAMCTKYGT
      *    .   * .    *****.* ****     ***** *. .     . :  *  . .

35    CTTYMCRTTYGYWGTGATKCRTYKWKYRRAWAACTAKCCGYYYYRKYYMYWMRYGTTGKK
42    TCCYMMAAAYRYWAAKRYKSAGTKWGYRRGAWWMKMKSYKYYCCRTCYCCAMRYRWWCKK
         **  ::* **.:   *.   ** ***.    . *.  **  *. *   ***    **

35    KACCKWRTKGCAKWKRWWCWMCGCACYWYGCRYMKKKRKKGCYKKKRKYKRCMRYGGAKK
42    KRYGKARYGATTKTGRWWYTCYAMMTCTTCTRCCTKKRKKRYYKGKAGTKRYCGYKKMKK
      *   * *  . :*  ***    .        *  .*****  ** *   **   *   **

35    KTCGTTMKYWGCYYMTYYCKMKAAYTGATCAAAGRRTTCYCKGSCKTGGGKRKWYWWSTG
42    GKTTGYMKTTKYTCCYYYYTAGRGTYKRYYRRWRRRKYYCMKKSYKWKRSKRKTTWASCR
       .    **         ** .   .          **.     * * *   .***  * *

35    WRGGAGRGCCCCCTCKKWRSYGTGGYGYMYYYATGKRMYCCTATKKMRKYWWWYCYTACK
42    WRKKWRRRMMYYSYMKKTGGTTGKRCKYMYCYMKRTRMYTAYRYKKCRKYWWWYGYWGTK
      **    *     .  **  .       *** * . .*** .   ** ******* * . *

35    CTTGKGTKASYCKCKYKYKRYGCCGSTARKKRRCTYKWATAAGMKKKTTTMYAKGWRYRM
42    MYYKKRKKWCTMTTKYKYGRYATYASWMRKKRRTWYKWWKMRRMKKKACYMTWKRWRYRC
          * .* .  . **** **.  .*  *****  *** .   ****:  *  * ****

35    KYRGCAYTRGTKTTSWTATMGCGGCCAAACCTTYARTTGWYRMTMYRWRWKGRSKRATCG
42    KCAKMGCYRAWKWGGTKGAATTACAYRMMYYYYCRRYCKTTACGCCRTRWKRACGRRWTA
      *    .  *. *  . ..:   . .          *          * ***  . *   .

35    ASYGWYTTRTYTYMGMATATKMAKAAMKATGYRMRCTGRCKSMGGAAGGMYRYTWTYTTS
42    WGYAWYYYGKTCCAACTCGCGCCKRGAKTAAYRCASYRRMKGCATCRTRMYRCKWYYKKS
       .*.**   .    . : .   .* . *::.**  .  * *. . .   *** .* *..*

35    YWTTTAKYYKYACAYGGRWYKYMYMYRRCCGTCCGRWRRGTKRMARWTTMGTCCGGKKRR
42    YWWWYMKYYTCGYWTKTRWYKYMYMYAGYSRWYARRWGAACTRMRRTWCCRCTSRRKKRR
      **    ***. .     *********   .   . **  . .** *       .  ****

35    RYATWAGKRWCKGRRRYKCYWKSAKGAKKGRCRYARKKRCAWRRRTMYGGTGKRSKGWKR
42    GCRGTGAKRWMKKRRRYKYYWKCTKATKTRAYRYCRKKGMMARRGACTKRYTKRSKRWGR
           ..*** * ***** ***.:*.:*.   **.***    ** :      **** * *

35    GSYGTRKATGYRKAARKRYMRKGKKCWCCTKMYYRYYWCCAKSYMAAGGYYTCTRYAKKK
42    ASCAWGKTCKYRKGCRKRCCAGKKKYAYTKKMYTGYYWYSRKGCMRRRKCTKYYACCKKK
      .* .  *:  ***..***     **    .***  *** . *. *       .    .***

35    KKCGWKWKCGGRYKKYTYYRTTAAKAGCRCYTGRRKCKTMKTAYGKCATTKTMCKRKTAC
42    TGAATGWKTAARYKGCCYYRCCTTKMSYRYYWKGRKAKWMKYMYRKTRACKYMMGAGYRS
      . ..  ** ..***   ***  ::* . * *   **.* **  * *  : * *      .

35    TYACAYRMYKKCYRMTGYKYYMATAWMAGWWGYRRKRCKCKGKKKWTMRYYTCKCRRKMG
42    CYGTCTAMCGKMYRMYRYTYCCTATWCWSWTRYGATRYKYKAKKKWWMRYYYYKYRRKMA
       *. .  *  * ***  *.*  :::*  .*  *  .* * *.**** ****  * ****.

35    RCGCKRGGCCKKKKRKACGRMTWKTTGTAMYKMKAATTACMRMRTMRTRTAKRARYTAAK
42    RTAGKRRTTGKKKKAKRYKGCKTKKYRYWMYGMKMTYKRYMRMRYMRWRCGKGRRYYWTK
```

```
                 * .**     **** *     . *.    ** ** : .  **** ** * .*  **  :*

35               WRRWKGTWMTWRRGAKRYYYWCTWTGMKCGGMYYTTCGMWMWTYGGCYCKTTTAGRGCCA
42               AAAWKRYTCCWRRRCKRYYYWMYAWRMKYAKACCYYMRAWMWCTAKYCSKGAWMRGAATG
                    **     *** .******     ** .         ***  .  .* :    .. .

35               YRAATRTGMWTKTKWSKRTGRATGRRMTRKMMYAKSKGKYKGKYYMWTCAACAACGGGAR
42               YRWGYGKAATWKYKAGKRYKGRWKGAACRKACCGKCKRKTGRKYYMWYMGTMCWMRRRRR
                 ** .  ..   * * .**          **   .*.* *   *****  .: .      *

35               TMWGCWGAGAKRRCCKYWYACKCTRYKYGGRSRSKAYRRGRKKMCGTKRRMRAKCTAYCA
42               CMTRMWRMKMKRAGYKYWYRYKSYRYKYRRGSRSKWCAARRKKCTACKRRMRRKSCRCMR
                  *   *    **   ****  *. ****   ****     ***  . ***** *.

35               TGAGACKRYRYYKYRTGGTMKGTTYCGRTRYKKYRRWKAGGRKTTKWYAYYKTWRRMGCW
42               YRRKMYKRYRYTKCAWRCCCKSWWTYRRCAYTKYRRWKRCRRKYYKATRYYGYAGACRYW
                       ***** *       *.     * *.******    **  *   **        *

35               KYYWYYGKKRKWYATCAAYMSYAGCTTKRTCRTYAKWAYTARMWWYMSKTGTCWATATKT
42               KYYWCYKKKAKWYWKYRRYMSYWSYWKTGWTRKYRTARYWMGCATCAGKCAYSWWKMWKW
                 **** * ** *** .   ****  .  ..   *.* .  *        .*  . .* .  *

35               CGGKKKTCWCYKYKYYTKYKKTYTKCCYGTGCACKKMRAAGYTTWTWGRKCCTTRACGGK
42               MRRKGKYATTYKYKYYYKCKKCYKKMMCKCRYRMTGAACGACYKWWTRAGTYWWRRMKRK
                    * * .  ****** * ** *.*          .   ...  .*          *    *

35               RKAAGCKKKYKWYTAYMGTYGYAKCKRYCTYRYRTCGYCRYAKYKCRGWKWYRAAKYAWK
42               RKWRKYKKKYKAYWRCCCCTKYCKYKGCAGCRYRCYKYYATGTCTMARWKWYRGGKYRWG
                 **    ***** *         *.* *   .  ***    *    .. .   *****..** *

35               MTAKWYMAYYYTWTCCYCTKCYCYRATYRYGKRATATCGCRKSTATTRTGRCCMYKKKTK
42               MCRTTTAGYYYCWGTTTYCKYYYYARGYRYRTRRWMAYCMGGGCCGKRCKGAMMTKGKKK
                 *  .   .*** *       * * *   *** .*    :      . . .*   . * * *.*

35               KCSYKWRCTTATAKGTTTGMKKWCCWRARYWSTYKRMGGWKKRSGKKCCWATCTCKYGYK
42               GTSCKWRYYCRCGKCCCYRMKKAMTTGWRTASAYTGCRATKKRSAKKMYWMWTGTKYRYK
                   * ***     .*     ***       *  *:*.    . ****.**   *     ** **

35               AACSTRKAKTAYKRKKTKGAWMYKGTATKCYSSCAKTKRMKKTMYRRGAYATYTYRKWKY
42               WMMSKAKGKYWYKRKKCKTRWCTKACGATTYSSYGKYKRMKKKMYAGRWCGCCYYRGWKY
                    *. *.*  ***** *  *  *. .:. *** .* *****.**      .   ** ***

35               WYRWAKGMGGKAKKKTYYMYAAGTKAYYAGWGTAYGYATTAYRWYYAYRKYRKKGATAYR
42               WCRWTGRMKKKGKTKGYCMYMWRWKWYYRKWRYGYRYRYWRTRWCCRYRKYRKKRRYWYR
                 * **:  *  *.*.* * **     * **  *  .* *     **   ******    **

35               GRCKYWWKWYWCGYYYRTRRARWKYYMYTMARGTCAAGWCTACGAYACKKYYKKKCGRKY
42               AGYKYWWKTCTTACCCRWRRRAATYYMYKMRRKWSRRKWYCGYAGYWYKKYYKKGTKRKY
                 .  *****     .   * **   .****.* *  .   *  . ..*  ******   ***

35               KRKYAYRCYRRYCGGCGMAAWYSRCTYYMGRGYRYMATASWWKKGAAYGAGGGKWKYRCG
42               KAKYRCRYYRRYYRRYKCMWTYSRYAYYCRARYGCMRWRGTWKKRRRTRMRKRKWKYRMR
                 * **  * ****          *** :**    *  *   . ***         *****
```

```
35      TARKRTARMYCKTYCYRYMCCTRKAYSKMYYGATCRATKMMKYRGCTCTTSMKCYKKATR
42      YMRKRKRRCTMKYYYYRYAMYKRKMTSKMYYRWATARYGMMGYRATATWWSMKYCGKGCG
          ***. *   * * ***   .**  *****  :     ** **. :   ***   *.

35      GTAACAKKAATCCGTCYRAWGTGRRTYYKRGTGYGAGTMWRWYATGTYGWTACGTYGACA
42      AYRGMRKKGRYYYAKYYRTTCARRRKCCKGKCTYAGRYAWRWYGCKGYRTYTMRYYAWSR
        .  .  **.    .. **:  : **.  *    *..   ****.   *   :   *. .

35      MWASRKGGKTRRKWWATASRRSKYWKKMGYYKGCGTTTGMMYRRRAKKKCGYYCRRKYMC
42      CTRSAKRKKWRRKTTWYRSRGSKTTKKMTYYKRSRWYKCMMYRRRCTKKTKCTMRRKYMM
           * *  * ***     ** **  *** *** .   . ******..**     *****

35      YRRKMYGAMCWKRTTMGTTMYWCTTATMGKCKWTKWWSKKTATTRRGAKKGCASMGRKMT
42      YGGKCTRGMMWKRYYCSWKAYWYAYRYMRGYKWKKTWSTKAMWKRAKCKKTTGGAAGGMG
        *  *   .* ***   . . ** :   *   **.* **.*:  .*  .**  .. .  *

35      RRKRRACCACAACKYCMCYYSYYYAATTTCGTYCCCYTMMYYYRTGKMSRSWKSKARRWK
42      RRKRRCTYCTCRSGYYMYYYSYYYMWGWKYRCCYYYCYCCTCYRYAKMSRCTGSGRAATK
        *****.  . . . * * ******    .             ** .****.  *     *

35      CCATKTCRGYYRRRWSACCAWTAKCAMYRYKTMCYYYCAGYRRMATATRTKAGTYYGGAM
42      TYWGKYYGRCYRRRWSCYYWWGGKTTMCGTKYMYYYYYRTYGRAMCTYGWKGTAYYTKMC
            *     ******.   * .* :*   * * ***   * *   :   *. :**

35      GTATYWGAYKKYRAYWKKKYYKYYRTYGGKRMTYSSGKRKWRGRCMAMRAGTKKGWKMMC
42      RKRACATWYKTYRRYWKKKYYGYYRWYKRKRCYYSSKGGKWRKRMCMMRWRKKKATTMMT
         . :    **.** ******* *** *  **  ***   *** *   **  .**. .**

35      RRKGRRKKYCTRCATCCKKYGAAWAKYYWRARKTRMRGRTRTTAAGGYGYRAGTGYYRKM
42      RRKRRRKKYTGRYWYTTKKCRRMWRKYYWGRRKYRMARRKGWWMCAACKYRWRKKYYRGA
        *** *****  *     **    * ****  ** **  *.    ...  **  . ***

35      RCGKTKYKRATAAKTWRSYWYYRTKYTARYAYYWTRRTMMSRTAKTCCTGGAWKGCGCKK
42      RYRGWKTGRMYRRGYWRSYWYCGCGYCCRYTCCWKRRYMMSAKMKCTYWKRGWKRTATTT
        *    *  *      ******    * .**:  *.** *** . *      .**  . ..

35      GAMCYKCTTGKYRKCKGMKGGTGYCWTRTGRYKWRWKYACGATGTCKRRCRATRTGMARG
42      TWMTYKMYKKKYRKMKTAKCTYRTTAAAYARCGTGTGCGTRWCKYMKRRSGWKAWKCMGK
          * **  . **** *  *       :  .*       .       ***.  .

35      KAGCTCTCGWTCGTTKKKGARRCACATTTATTTCYCAACCGKGCRKAGGCKWRYKYCAWT
42      TRATAAYMRWKMRYCKKGRCAATTTGCYWMKWAYCTRRYYRKRAGTWTATKWRYKYYWWC
        . .:.   *.    **  .   : .    . :        * . .  . ******  *

35      AYYATCYKYMRGYYKCYTKKRCMYRKCKCATRGCRCGTCKYKRRKACATGKCKWSAKRYK
42      WYCWCYTTYMRKYYGMYCKKRMMYRTMGSMCRRYRATGAKYKRGKMYMWRKMKTCWTGTK
         *     .*** **  * *** ***.  .  *  *.  .**** *     * * . .  *

35      MAAYMKCCCKGTSRYAKGTAAYGKACTATYRRRWYRTYRRAWYYCARRCRSYARRTKGTG
42      MRMCAKYYMKKGGRYRTKWMRYRKWMWGCYARRWYRYCARWTCCTTGGMRSYRRGYKAYR
        *    *   *  .** .    * *   . * *****   *     :   *** *  *.
```

```
35      TRACAWGGGKMKWMKKTGGAACMRWRSMRSRRKATTTCCMRMYKMKRMRTTGYRYGARMY
42      YGCTGTATAKCGWMKKYRRGCAARWRSMRSRRKGWWKYGMRMCKAKAAGYKKYRCACGCY
          . . . .*  ****   ... **********.  .  *** * *    . ** ..  *

35      RGKAAGAAGYWYYSKCTGTAKTAAGMWMRYCTKCARKRYRTMRKTMGGSRKWKTRKGCCK
42      RTKGWARMKYWYCGKYYSWGKCCCAMWMRYMYGTGAKRCRKMRKAMCRGRGTKARKAYYK
        * *. .   *** .*  . .* ...*****    . ** *.***:*  .*  *:**.  *

35      YCCCRRYKAKRARYYRTRKYRMRYKKMYATCATGYCCRKYGATYKMMKKCTTAKKKKYTM
42      CYSMGGCKCKRRRYYRCRKYRMRYKKMCWYYRARTYYAKYRWKYKMMKKYYWMTKKKYKM
          .    *.** **** **********     :     **  .******    .****.*

35      YMKRSTCYRAGYKYCMMRGTMRYSTGRGWWAACCGRGACCARTRAGATCCAYATYYCKCT
42      YMKRSATYGGACGCYMMARKMGCCWRARWTRCTMRGRMTYRRCGGRMYTTGTGCYYYKMG
        *****: * ..    **  .*  .    *  .         *  .     . . ** *

35      WCYGKCTTATTYTGCGCGMAYTGACGYMKTKCGRRACTTGKTCKAYAGYCCGAAYCMRMM
42      WMCRGAYYWKWCWRYRGKMRTKKWMKYMKAKMKRRWYCCKKYSKWYTTYAGAGGTTAGAC
        *    .   .        *  .    ***:*  **     * .* *: *. ...

35      YRCCTCSKRTGTRRRYYCYRGYRTKRWTKTTTYTKARGYYSTYYRTTAGATTATTRTCCT
42      YAYYCASKRYRCAGRCYTYRRCRCGRTGKCAACYKWARYCSGYYGYKRRMCCMYWRYYYY
        *    .***     * * **  *  *  * ::  *   * * **  .        *

35      TCYCGYWAKYWYRWKAAGRWRRYMTACACAARTKYTTTKRRKYKTTYMKGCYRKWTTCRT
42      KYCMSCWMKYWYRTTRRAGWGRYMYRMMMRMRGKYYWYKAGGYKACYAKAYTAGTYYTGC
        .   . * ***** .  . * ***       * **   *   **: * *.

35      SRMWRGRAKMAAGTRTWAYRMYWYCCWGMKKGCRGWYASYAGGGGCATAKMRRTWYRGMK
42      GAMTGRRMKMMCSYRKWMYRMCWYMYWAAGTAYRRTCCCCTACATTGCGKARRCACRKAT
        . *   * ** .. *.* *** **  *.  .. *   .. :. .  . .* **   *  .

35      ARKRCMRCMSTTTAKWTTACYRWARCRTGTTCCKYRGGAKWGTAGMCWMAATTGTGRKRC
42      GRTGYMGGMSCCKMTTYWCSYGAGGTGYKWYYYKCRKKMKWKGGAATTCRRYGACCRKGA
        .*.  *  **  . .   ..*  .         * *   **  ..        .  ** .

35      AYSYKKKAWGCRRGMGCTCYMGKKAKYKYGTARGTCKRKYRWRRGACYGWYGSKAAATTA
42      TCSYKKKMWTTRRRCKACYCCKKKGGTKYRYWGAWYKRKYRTAATRYCRWCRSKRGTYKM
        : ***** *  **   .     **.  **    .  *****        *  ** .: .

35      ATAYMKKSWKAKKGRCCWTRYGKGKAGRTWGWRTAATKCRCRTAKGGCYYCCGTGGRYGR
42      MKMCMTKSTKGGKARSMAYGTKKKKGKAKTATRYMMWGARMGCGTTATTCTMRYRKACKG
         .  *.** *. *.*.      * *.  . . *     .*   .. .

35      MKYACCAYTSAAKKARAKYKCRTMWTCTYKTTGGKRKMGGATTTGGGGGKKYWCYGCKKY
42      AKTTYYGYYCRMKKRAGKYKTRCMWWYWYKKKACKAGCKKGWGYRKSRKKTYAYCRYKKY
         * :  .* .  **  .*** * **   **... *     .     .  *.*     ***

35      YYMWKKYMKMCGCWRYWTMWKTRYAATTKSYKMCGCSGAKRKRRGACGTGKKTAKYTGCT
42      YYCTGKYMGASSTAGCWWCWKWACGGACKSYGMYRYCAGKRKRAKWYKYKKKYMKTCAAY
        **   ***  ..    *  **   ..: *** *   ...****       **  *  ..

35      TRMKTKCYRTAYYYMKTGYCGRYTWWGTTRKKKCWKRRKYSKGRYYAGMYRKRGGCKAWA
```

```
42              YGCKKKTYGWMYTTCKAAYMKRCYTWTGYGKKKTWGRGGTCGRACYWRAYRKARRYKGWG
                   *.* *   *   *:.*  *   *    *** * *   .    *   ***    *.*.

35              CTTRCGYMAYKGTKRCYYAGYCCRGGAYKKGTRYKRWTAGTGTKRYYYAYYWATCACCAG
42              YCKGYKCMRCGRWKRMYYWATMTGATGYGKKCRYGRTKRKAKYKRCTYMYYWRYTTTTRK
                  .    *     ** ** .    . .* *  ** * .  :  **  * ***   :

35              CKRKRYTGGATYRYRKYKKMTMRRRWYRCGGWKTGAWCKCKSKAKWAYYAAGGTATMAAT
42              YKRKRCYARGCCGYGKYKKCGMRRRAYGYRRATARRWYKYKSKRTWRYYWRKRKRYCTRY
                 ****  . .   * ****  **** *     .:  * * *** .* **    .   :

35              GATGCRRSKKWYWRTGTARYRWGTAGCYYKSKTSKACCMRMATGYCSGTMYYYKCAMRTT
42              KRKSYGGSKKWTWGCRAGGCRWAKWATYYGCKCCGCGMMRARCAYMCCWMYYYKYMCACY
                  ..   **** *   :.  **.. . ** .* . .  **   .* .  *****

35              CKGTAKYYACCAYWYTRMYWTYCGKKKT
42              TKSKRGYYTTAMYACWAMCWWTYRKKKW
                 *..  **: . *    * *    ***
```

The pairwise alignments of two accessions 35 and 43

```
35              WYGYMYSYWYKYKRKRMWRYYWRTGRAMRGGCTTMTAMYACRYRKCRKMYMRAAKSRKSY
43              WYTYMYSCTCKYGGKRMWAYYWRKRGRCGKATCCMCRACRYRTATTGKMYMRWWKSRKSY
                ** ****   **  **** ****.      .   *      *  .  *****  ******

35              YRYYRKCGTRYYYRWRKYYYYYYGCARGYRCTTRWARCYKRSMGARCGWWMRRYYYYRGT
43              YRYYRKSRKGYYCRWRKYYYYYYRMGRRTGYYYRWRGYYKRGCRGGYRTWMRRYYYYGAY
                ******. . ** **********  .*      **   ***.  .    ******** .

35              TCCATTRWWYYRRATMWTAMWCYKWMYYYMRRYKWYYAWTRWKKCRWYCKCCYGCRMKTC
43              GYYRAWGWWYYRRWKCAWGMAAYTWMCYYMGGCGTTYWTGGAGKYAWCTGYYYRTGMGYY
                    :  ****** .   .* .*.** ***      *      *  *     *   *

35              WKGAYCRTTTWKYKMWYWWMKKYKRMRRAGKCRYRCAKKGGYMACCKKMKRKYTCTTYWR
43              WKAGYMAGYYWKTGCWYWWAKKTKRMRRCSKMRYRATGKKRCCGTTKKMKGGCCMAATWR
                **..*     **   **** ** *****..* ***.: *    .  ****     :: **

35              CCCTTYTTMCYTRRYYTYWMRSARAKWRGYCCAYYRYWTTYYRYYRTTKYYRYYKTTWYC
43              TTYGGTGYCMCGRRYYWYTCGGGGTKTRKCAGGTYATTWKYCAYYRGKGYYRTYKGKAYY
                            **** *   .. :* *  . . *    .*  *** . *** ** . *

35              RYWACCCKACMYYGRCGRKMSYRYCMSRWSYMKYYMYKKRMYKKGKYRWCYMYMCRYKMR
43              ACWRYTAKWMMYYSATARKMSCGTMMSRTCCCTTCMCKKRMYKKRKTRWTTMYAYRYKCG
                  *   .*  ***.  .****    ***  .  .  * ******* * **  **  ***

35              SRGCCRYKGWMRWAYRYTYSRKAWWYTMAAGKRYYYAKCTCGYACTTGGACTGRGRYCRA
43              CGKYTRYGKTCRWTYGTYCSRKWAACWMRWRTGCTCMKYYTATRTGYCKRTYCGKRYTRC
                .    **    **:*    ***     *   .     *   .             ** *.

35              GTWYATRRGTGAACCRRYACCCGKAAAKYGTAGKCACRTRCGGWGTCGRKGYRSKTYRYT
43              AYATMCRGRYRWMYYGRYCTATAKWMMKYAGGRKTGYGYGTAAAAYAKRKTCASKKYRYC
                .     *         **. . .*   **. . * .     .. . . **   **.***

35              MYYAAGGMATGATKMYYYRACAYWAYKGRWGGRMWCYYRGYGGGCASKMRKMRGYAMYYY
43              MYYTGTACRYKRKTMYYYGGGGTWGYKKATARACWTCTASYATRMGCKCAKMRAYRMYYY
                ***:. .     ..**** . . *.**   .    *    .*.   ..*  ***.* ****

35              YGAGMTCRMTAKYTYKYCRRRKKKAYRYRTAGAARKACGACGRTCKKCACRWRYCYGRMA
43              YCRRMYYAMKRKCGCKCTGGAKKTTCATACTRRMRKMMTRYKAGYKKYGYRTGCYCRGMG
                *   *   *. *   *     **.:     :   **         ** . *       *.

35              CCKATGCCYCMCKYKAYRTRARYMCRAACAWYTGGGCAAYGKSKRTAATWAACRAATCTK
43              TAKMYKYMYMMMGCKGYRWGRACCMARRMWWYYRTTYGCTKTGKRKTCYATGARGGGTYK
                 .*     * *   *.**            **     ..  ..**.:.  :..*..   *

35              GSYRCYCGATWRRCWRYTCCYWYYGATTAAMYYCKCAKKTGAKKCGAKCAYTACRCRGGY
43              AGCRYCTTTAWGGTAGCWYMYWYCRRKYRRAYYMKMCKKCAGGTMRRGSMYYCTRMGRSY
                .. *    ::*         ***   .    ** * .** .. .    . * . *   .*

35              GCRKTYCRSKGGGTWAYWSKMYKGMKYMCCKGMSATYTRRYTYAACAYARYCGCAGRYYS
43              RYRKCYTRGGKRRYAWCTCKMCKRCKYMYYKRMGRWYWRRYYCRGTTYTACARACRGCTS
```

```
                  ** * *.         .** *  ***  * *.  * ***   . :*:  . ..    *

35              YRRKGRTGTWGAAYKRTKKGGMRSMRSRRKRMYKGTAMKRYGRRWKYGCWCWRYSYKMRK
43              YGGKARYRYTARRYKGYKKRRARSMGCGGKRMCKRYRMKGTAGGTKCATWYWGCCTKMRK
                *  *.*    .  **  **   *** .  *** *   **  .   * . * *  . ****

35              GTRWCCTKYTGCCKRGRSRYMCMMYKMYTRKATAKCTCMGCCRRTRRAYGAWCRKKYYCT
43              ACRWYMKKTCAAAKGRRGGYCMAATKMCYRKGCCGYYMMRYYGGCGGGYKTAYRKGTYTC
                . **  .*  ...*  *. *     **  **. .    *        .* :  **  *

35              KGRCRGCTTTKRCTGTRKTRWCGMYGTTKYRGCMWCKRYYYASKKAGRKTMYAATWWGAK
43              KAAMRRSKYKKGYYSYGKKGATTCCRWWKYRRMMTTKRYYTRSGKGTRKCACRGYWTRRK
                *.  * .. .*   .  *.         ***  *  ****  * *. **    . *   *

35              RYCKKYSAKGYCKYRKKKRRTTGAAYWKCTSAWYWTMYKGCWRYRATCMKRCCGTCTAGG
43              RYTKKCCGKTTTKYGKKKRRAATGGYWKMWSRACAYMTTAGWRYRWYYCKRYMKCYYRKC
                ** ** ..*   ** *****:: ..***  *     * .. ****    **

35              AAKYKKAYATAYRRKWYKTWTTCCGGRYAGCCRRCKRWRRMARMRYRKCKCRYRWKYMYR
43              CMKYKKWTRWRYRRKWYKGTKYYYRAGYGAATAGYKRWGAMRRMRYRTYKMGCAAKCATA
                . ****     *******  .    . *...    ***  * *****. *     *

35              CGGMKKKYWTKYAAGMWCGWCYMCGKRRAGTMYACCYRARYMGGKGKCKGKRSTRMCYWR
43              TRTAKTTTWAKTRMRATYKTYCCMKKGRMRAMYGMSYRRGCMRKGTKMKRGRSCGMTYWR
                    *.. *:*              * *  :**. .**   *    * *  **  * ***

35              YRRKMYKGRYYWKASYYYYRTKRMACKYSWWSYYKGCKACKMGCKYKKYGTARGCCYRGY
43              TRAGMCKAGYYWKMGCCCCRGKGCMYKYSWWGTYKTTKWTKCTYGYKKYRYWGAYYYATC
                 *  * *. **** .    * *    *****. **  *  *    ****    .  *

35              GYTYTGGKMKAMSGCWKAKCCWSGKGGWRTTYTATYKTRCATAAKCWGRWKGGCKGGGGY
43              ACKCAARTAKMMSCTWKGKSMACKTRRWAWKYWGCCGARYWYMMGMTRATKTATKRKKKY
                . . :. . * **  **.*.  . .  *  .* .   :*           * . *    *

35              RYKWKRMMKKYACATCATKTKAYMYKRRWMWRGTCAYAATAYKRKMYKWKCTYYKAYRAG
43              ACKAKGMMKTYTYGWYWWKYKGTCYKGRACAARYMRYGTGGTKAGCYKAKYATTKGTGGA
                  * * ***.*: .    * *.  ** *        *.: . *   ** * :  *.  ..

35              CTKRWTAATCCGYYYRRTKKRGTAACAMMATWYCYGKWRTAAKKMYMRYKYCWAMGCTTW
43              YYKRTCRTGYTACCCRRCTKRKWTRTGMMRYAYTYRKTRCWRKKATAGYKYYTCCRMYYT
                  **   :   .   ** .**  :  .**   * * * *   **    ***  .

35              YYCCGCYRRYKRYYWYSKYAYRMYGTAYYRATCASYKWTWCGKTYTCRKKRYKYYGRMKK
43              CCYMATCGRYGRYYTCSKYRYAMCRAGTCAGAMRGTKTATYAKKYKYGGKGTKTCTRATK
                    .   ** ***  *** * *  :.   .:  . * :  .*.*.   *  *   * .*

35              ACYACTGSYRGTTACTCSMCTKRCCWRRAKATRCATGTGTTTMKAGKTSTWMKATTRCRY
43              CMCRAYRGCAAYYMTATCCYAGGYYTAARKGCGTCCACRYYWMKRAKYGCAMKMYWRMRC
                .   .  .  .    : .  :        *.   . .     ** .* .  **   * *

35              GKYGAWYRWKYRYMSACACAGARRKYYCKWARWTCCACRTSACGMYCTWCYRMRKKTYSC
43              RKYRRWYRWKYATCSCAGTGARGATCYTGTMGTYAYRTGCGTAKACAKTTTGAGKKWCCY
                 **  ******   *... ..   . *       .     .:.   ..      **  .
```

```
35          YYCTTTKGATATCMAWKRRTWYCCAAARKRKGCRCTGCCCGCGAKKYKGMACRWAKTCWK
43          CCACAGTAGCWWTACAKRGCACATTGRATGKRYRYACYYMTATCKKYKAMMMRWGKKTAG
              . : ...     . **    . :.  . *  * :     . .****.*  **.*.

35          YKYRMWCAAAMYRGRCGAYGSSTKYYTRCTKWKKKWMYKMKTKAATGCMRYAKRACKYGR
43          YKYACTMRRRCYGRGTTGYASSGKTTGAYKGWKKKWMYKMTGGMWGCACRCRGACGKYKG
            ***        *     .*.** *     . *********.      . *    . **

35          WAAKCATTRYAYYRRKYYSRYYYKCCWCTYGYMTCMYGYRKSATCWRWKAKKYKYRYMYR
43          WWRKSRYWGTRCCGGKYYSRYCCGYMTTYYKYCKACTKYRKSCCYTRTKMKKYKYGYMCG
            *  *.          ******        * * ..   ****.   * * ***** **

35          YKTCKTCCCGGAYAARAACCCGAYKKAWKAKKKRCTYRRGAMAMYYKKAKKWKYYCKMCT
43          TKCMKCAYYRRMCTGGGWYSYATCGKGWKTTKKRTGCRRKMCRMYCKTRTKWKYTSKCMY
             *  * .      :. .  . .:  *.**:.***   **    ** *. .**** .*

35          ATKTRRTCCMACMKMYKTCAGATKKKRYAAWKGTTTYCRYRTCYACCYTRTKTAMCTTAC
43          RKGYRRCYACMTMKMCKCTTAGCKKKRCGCWKRCACTMRYRYYTMYMCWGWKCCCTKCCT
             .  **  .   *** *  :.. **** ..**  :   ***          * .  . .

35          GMRRYCKKGYWYRRWKYGRCKTCTTCKYKKYYYWYAMCCCGYRTKKRGKRRYMRRSYRKR
43          KMGGTMTKKYTYRRWKYCRYKYYCCTKCKKYYYWCCCATTKCGKKKASKGACARRSYRKR
             *    .* * ****** * *     * ****** . .     .** .*    *******

35          CKRYRAGCATTRAAYRYTAMWWYMSGWTKRMKKGTAWWYRKYKRYKRGGCCCWKTTKWKT
43          MTGCGWRMCKYGGRCRYWMCATCAGRWGKRMKKRGCWWTRKCKRYKARRYATWKCCGTKW
             .      ..  .  **       . * *****  .** ** ****    . **    *

35          TRKRKKKCGKKRCRAYCCKRCWRCYATGAKTYCRYCRMAGWYMWYCTSYMYCAGCTGCCY
43          WGKRKKKTRKKRTRRYATKAGWRAYGCAGGYYTGCMACCCTYMTYYYSCMYTGATCATMT
              *****  *** * *. *  **.*. ..  *      .  ** *  * ** ..  .

35          MMTAWSTTCCAKGAYWKYWACAGRRYTMYAGRACGTTKRGMGYCYAATATTARGACCCKT
43          MAWWTCAYTTGKAGYWKTWGTRRRATCCCGCRRTTYWGRRAKYACRWKGYGGGAWSYSKK
            *    .:   .*..*** *.   *     . *      *   *.   ..  . . . .*.

35          AAGKATAKRGCCGGTGTACTGRGYRKAYCAGKRYRACGTMYRKYAYYWYYTGARSGRAKY
43          RTTKTGTGRRYTTRWRYRTYAATYRGRCTGAKGYRCGAYAYRKYWYCWCCCRRGSRGWGY
             : *: : *           .  **    ..* **. .  **** * *      *    *

35          RMRKCCRRKWWASKYKMKGMMYYTMRMYMKKTTYTYRRKGRWKRYWTKYRKTATGKTYYY
43          RMRKTMRRKTTWSKTKMKRMMYCKMRMYMKGCYCWTGAKRRWGAYWAGYGGCRYRKWYYY
            ****  ***   ** *** *** .******        * **  **: *      * ***

35          ACCYYWMKMRKMACYWRKKRAGRKCRTGAGGYMRKTKKTMRCTGGRSYWYCKATWTKYRK
43          CYYYTACGCAKMWTCARGKGCAGKTRYRRTAYMGGAKGYMRTGAARSYWYTTRWWWGCGK
            .  *      **    * * .. * *    .**  :*  **  ..***** .  *    *

35          WGYYKKATRMRMGWWYCCTKWKKTTATCKYYWGKWTYGTGYKYWWSTGYYMTMATRTRCA
43          WRYCKKGKRMRMAATYSYWTWTTYKMKMKTCAKKACTRAAYGCAAGGKTTCCCWKRCRMG
            * * **..****.  *.  .*.. . . *    *    :.*    .        .* * .
```

```
35      ATKWRCKYGGTTRGCCKTGCRMRTTKKAGTTYAYGCRTCTTTKRMGTKTRCATACGYYYC
43      RYKWRYTYTAAYGRYYGGAAGMRWCKKRRYKYWYAYGYYKKKKGCRCKCRTGATGAYYYA
          *** .* .:       .. **  **   .* *.    ...*    * * .:: .***.

35      CGMYYMAMTGKRWMWTYRTATYKTKWSCCRTCAKTKCATGRRMTRAKKKYKKKGCTCGGT
43      YKMCCMCAYAGRTCAWCGGTKCKYTAGYMGCYGKYKTRWKGAACAGKKGTGKKRSYAAAY
          *  *.  . *       :. * . .     .* *         .**  ** . ...

35      RRYRGKCYRGGCYYKRWGCGYWKRRMRAKCCGRYKYRTTCCATCCYRCGKKMRKWAGCCG
43      RRYGRKTYGTKGTYTRWKYAYAKRRMRRGTMRRYKYRWCATCCYAYGATGGAAKWGATTT
        ***  * *     *.**  .* *****     *****  . .  .* .     **..

35      GAYMYGTGGRKAGGMARATTYRARRTTTCGTAGTTCMWCAMYTAKYYTCKYRYGACRRRT
43      KGCCCAGCCGKMKACRGWYCYGWRGCGCTCKGRACYCWYRCTYRTYYWYKCGCRCAGRGC
         .   .    *  .      *  *      .. :   *      .**  *    .. *

35      GAAKRRARTWTGATAKRYKCYTTWCKRMYRYACMRYRAMGCMAKWRSWSCCTTYMTTYKK
43      RCGKGGGRCWGAGATGRTTTYCCTYKRCTACWYMRYGGMATCWKAGSAGTTCCTAAAYTG
         ..*  .* * ..:: * . *    **      *** .*.   *  * .      ::*.

35      GGCGYRATAGGACTACGYKCAKYCKYYRWWARCTRGGTRTCTMYRYTTTKKGRWTGWWWM
43      KASRYRRKMARWAGCGRTTMMTYYKCCAAAWRTWRRKKRYYYMYRYAWYGKCGAWRAAAM
         .. **  . .  . .   .  .* *      *  *  .*   ****:   *        *

35      KKMWGCRKKKRMGAYRYMKYCGGKKARCRMRCGKTYYKWYAYGKATCAAAGKGKGATGCCR
43      KKMACTGKKACTGYRCCKTTATKKTRYGCGMKKKTYTWYGTKKRYYRRWRKAKKTATAYA
        ***    **   .**  *  . **:*      *. *.**.  *       *.* :: .

35      YGTCACGARCYRYWYWWSWRTTTTYGGCCGGYRYMMYCKRYKMYCYRYCYTTAAGTCATT
43      CKAYGTAGRMCRYWTWWSWRYKKKYARMMTAYRYMCYYKACKMYTCACTCCCTTRWYGWC
          : . ..*  *** ***** ...*.    .**** * *  ***        ::   .

35      YCRKATCRYCKCGAGKTRMYAACYYKYRMYYGYTGAKRCRWAGWCGYRTKKKWGMAAYGW
43      CYRKGGTRYMTYKRRKGRMTWRMYYKTAMCTAYCACKGYRWRRTSRYRYKTKARMWWCKW
          **.  **  .    * **    ***  *  .* ..*  **   . ** *.*  *    *

35      YKMCYSCGSKCGTRRKKKCGKACARMWWGATCGMYKYRRWAKKRRCATGYATGACKGKKC
43      CGMTCCYRGKMRKRGKKKTAKRYRGCTTAMYTRATGCGGTMKGGAYWCACGCRRYKRKKY
          *  .  .*  .* ***  .*       .            *      . .    * **

35      TGACKYAKAGGTTAYYSRRTCTGACWYWGKWRYGRGGYCYRMRTCARRCTGKCMKYRRTA
43      WSMYKTWTWRACYRTCGRRKGYRWMWYARKTGTKRARTTCRAACYWRRMYTKAAKTRGCT
         .  *  .  .     .**.     **  *    *.    *     **   *. * * :

35      YCTGGGAYCRTTCARYKCKKAYTTKYYWTAKKTTCRRRRRYRKYGKYCGCCCTTTTTAAA
43      YTAASRCCYGWGAWRCTMKGRCCWKCYWWRKGYKYRGARRYGTCAKCYAATYGKCCAWRR
        * :.. .     . * . *     * **  *  . *  *** . .*  ..   .  :

35      AKCKYMKTCTCGKMKKRKGTCAGRGMSSYRAGYTAGTGYKWTYCCTAAWTACTCTAMYSK
43      MGMKCAGYTWMKKMKKGKRYYCRRRAGSTRRTCCTAGACGTCCAACRTWCMAAAAGCCGK
           *        **** *   . *  .* *    :.  .     ..  :*  .:.:.   .*

35      KYAKMWWRYTRKRRMSKRMKKSWMRRRRYKAASKKYKYCKKYARYYRRYRKKTYARAAYS
```

```
43                  KYGKATARYCRKAGMGKGMKGGAAAAGRTKTTGKKYKCSKKYRATTARYRKKKCTATGYS
                    **.*   ** **  *.* ** .     * *::.**** .***     *****. : :.**

35                  GYRRAATKGGCYCYKKGAKAAGCTYWTCKRKTCYGGKYGCGYMYKYRRRCGAATMRYSGT
43                  KCGRRRAKRKMYMTKKRCTMWRYWYTAMKAKYTYRSKCRMRCAYGCGRAMRCTKMGCCRY
                       *  :*   *  ** ..     * : * *  * .*      *   *   .:.*  .

35                  RCCACGWWC
43                  RTTTMRWTT
                    *  :  *
```

The pairwise alignments of two accessions 35 and 44

```
35              YYWGARKWKYRSKKRYYYMRWWRYTMTYACTYYYRRATYAAYYYYKYRTTYRYMCMGRAK
44              YYWRGATWKYRCGKRYYCCGTWRYWMATRAWTYYRRRCYTGYYYYKYGYWYAYCGARAMK
                *** . .****. ****    *** *:  .  ****  *:.******   * *      *

35              RRKWARMCTSMKRWATMSRRRKYWMKYMTARACCYWWRRYYGTWWMRMYKKYYYYAYYRW
44              RRKWRGAYGGCKRWWKMSRRRTYWCTCMAWGGYYCAAGGYYTCWWMRMYGKYYYYWCYGW
                ****     . *** .*****.** . *:  .       **  ****** *****  * *

35              RKSRRYMAAGMYRRRKWRWWYYACYYMRKKKSYYRYYKRTRYKYKYRAYRKCYTCCMCRY
44              RKSRGYMMCTACARAGTGWWCCGYYYMRTGKSYYRYYGGKGYKCKYRWYRKYYGTTCYRY
                **** ** .    *    **  . ****. *******  . ** *** *** *     **

35              YYMMKYMRKMKTTCYRCYWRMYYMKRYRMCTTTTMRGAKWYKMKTRAMRYMTCTCSYRKM
44              TCMMTYMRTMKYYMCGTCWRMYYMKRYRMYYWKWMRRRGTTTAKYRGCRCMATCMSYGKC
                  **.***.**       ***********   . **     . * *. * *:   ** *

35              RYRRRKWWRYCKRWYRAMYWYCCTACARRKTCTYRATWRSRGCTCAMRTTMRWKKYRYCR
44              GTAGAKTWRYYGRWYGGMCWCTYWGTTGGKGMAYARWWRGRAYYTRMAYYMAWKKYGTYA
                     * ***  *** .* *    . :  *  :*   **.*.    *   * ****

35              YCRYSKKWCYYKRMRWMSGWCYMRKAMGTCKKMATKWRTAMYYTYGYMRYRYGGTRRYRK
44              TAGYSKKAYYTKRMRTCGKWYTMRGRARWTGGCRYTTRCMATTWTAYMRYAYKRWRRCAK
                 . ****  * ****  . *  **           . *       .**** *   **  *

35              TYRYCCGCWYWYRKRRAGMYKWCGGMTTKRKCRWYYGCAYYRRTWYYRYRSRYMSYYYYR
44              YCACYYRTWCWCGKRRRCMYTAMRKMGKGRKYRWTCRMTCTGRATTCATRSRYCGTCYYR
                        * *  ***  **.    * . ** **    :   *:     **** .  ***

35              YRYRWYWYWCYRYRWRWKCTGRGCKYMWYYRMYKYYRAGATTYTYYRAYCATTTYRGYYA
44              YGTGWYWTAYCGCRWRWKYYSRAMKTATCCRCCKTTGWRMGCYCCTGTTMRKYYTRRYCW
                *   ***      *****  .*. *     *  *        *    :   .   * *

35              KCGGGYTTAMRRMYYYRWGYAKTAAGAGTACAGAATACAAAKCATCCTCACATGACGYCW
44              KAARACCCCCARCCYYGTRYRKYWGCCCYRATCTTAMYMRMKGWYYMWAMMWYRWMKYYW
                *.. .   .  *  **   * *  . .   .: :::     *      .        * *

35              YCAMWYRRRMKCKYAWRATYGKAAGWRYYWWMRMYRRYYCAGCTCCCCYTAYRYCMTGWT
44              YYRMWYRGACTAKYCTGTAYSKWRTWGYYWWAACTAGYYYGRYCTATSYYMCRYTCAATK
                *  ****   ..**.  ::*.*   * ****      ** .    . .*   **  :. .

35              ARYMMRMATKCTGWKCATTRTGRTRGTAYAGTRYGGGKAWMTAGGCRKKWRYKKGWTKAY
44              RGCACGCTYKAKRAGYRYYRGRRKRRCGYGRCRYKRAKTAMCTTATRKKARYKKSWYKWY
                       : *..       *  *.*  .*.  **  .*: * : . *** ****.* * *

35              CYKYMGYTWKYYCCYKTYYYKKCTCTRTKKCTKRRKKAATYTRYTYRMYRRWYATACKCT
44              YTTCAKCKTKYYYMCKCCCTTKSYYGRAKTMCKRRKKTTCYWRYWYRMYRRWYTCGTTYW
                  .    . ***   *    .*.   *:*.  *****:: * ** ********: . .

35              SMYRRCGWYSWKGTCYGCRWWSAWAYMKYTYAKCYKMRSAGTYKTAAAYWYARYSRRTTR
44              GACGRYRAYSTKKWYTAGAWTCWARCAKTYCMKYYKMRGRKWCKAGCCYWYGRYSRRAAR
```

```
      .   *   ** *    .  * .     *    * ****.    *:...***.*****::*

35    RYTGAWYARRTGGCTCRRRYCSARRKSRWYWTGGYWKMRRRTMGGKCGRSMTTRSRRKAG
44    RYYKWTCGARKRKMGAGRGYMSRGGKCGTYAYAATTKCRRGYCRRGYRRSMACRSRRKGS
      **     . *.    . * * *   *.  *  ..  * **        ***: *****..

35    GMYKGYRMRKRRTYAGRGYARMRYYYGARKMATWWGGYKYTGYKRGRMYCCTRRYRCKCG
44    TMCGRYAAGGAGYTWKRRTCGMRYYYKWATAGYATARCKYKRYKRRRMYTACGGCGAKYA
       *   *          *  . *****   . .   .  **. *** *** .     .* .

35    YTCMCCKTCCATRSKGAARRMMRKYCACKKCYWKGYKYTAMYGCGGCKRKRRTKAARRMY
44    YYGMTYKKMTTGRSKKRWAACMRGTMGTGTMTAKTYKYWCCYRYRRYGRKRRKKGGGACY
      *  *  *.  : ***      **   .  .   * *** . *      ****.*..   *

35    KKSYAKKGWYTWMYCYYGMKKKYWRYRKKRRKAATCKGKKCACCCGRRKKGYCRYWCGYY
44    KKSTCKKKTTCAATMYTKCKTGCWRYRKTRRGCTATKKKGGGMMYCGGKKRCTGCWYKYY
      *** .**         *   *. *****.** .:: * *  .      **     *  **

35    CKMRTMRYMAATTYSTACYMKAKRGTKAKGRKTGTCKAYCAGTRMWRKATKRRYKKCMYT
44    TKAGWAACMGGACCCYWTTMKRKRRYGMKKRKYKATKMTMGRKGCWRKWKKGGYGKYMCY
       *      *..:  .    ** **    * **  : *   . .  *** .*  * * *

35    RRRGTCAAKATCYKRRRRRRKYKGYTCKKRKRCTMYSSCKWYTRCRTRRWKMTYMGTTAC
44    RRRSYYMGKGGYYKGRRARGGTGRYCYTKAKRYKATGGYKACCRYGCRGAKCYCMRWWWA
      ***.   .*.  ** ** *     *  .* ** .  .. *   *   *  *   *    .

35    AGKYYCARAYKGYYKTCRYYAAAKYMMYTATMGRYYKACSYTTYKYRWYCYRTARGKTTG
44    GRKYYTGARYGACTKCAGTTRWRKYCCCARYCKRYYGMTGCACCKCAATGCGWWRTKGCK
      . *** .  * .  * .      **   :    ***   . :  *         * *

35    YGCTAGACCCRRKTTAWRMTMRYKYYARWKTCAAGTMGTGTTTCGSGGGAGATYKRMRYG
44    YCAAGRCMYYRRKYAGAACCCGTTCCGRWKCTGGRCCRYCCGGMRGAKKRRMATTGARTK
      * .:. .   *** :.       .  .***  ..           ..     : .  *

35    TMTCWYYYYTGAGKAGCTCASRMKRMMYRWKAAYMTRAYMYTGGATACTYAGTTGAGYST
44    ACCYTCYYCKKCTGCKMYYWGAMGAAMTGWKMGTMWACYATYRTWWMTWTMTCCATKYGC
      :     ** . .  .     . *   *  ** . *  .*               .: *.

35    ATGMGCAWGTSKAATCMTGTTYRGMMATTCRGATASGCAATCTKTATTRGYTCCTYCGCC
44    GKACRMRARWSKGGGTMWTGGCGRCATGATRRMYWSRYGGGTCKCMYWRTYYYYYTYRTT
      ...       **..  *         : : *    *  ..   *    * *

35    KRYTKWRKYYYGYTGWYRGRYMKRARGTYMGKWKTTYKMAGKRRKYYTYCGMKTACWYWR
44    KACYTWRKYYYRYYRWYRKRTCKRRAAYTMRKWGYWTKMWAGGATCYYCYKATYRTWYAG
      *   .****** *  *** *  **  .  * **    **  .   . *     .   **

35    WCYRATCAMWRMRMGGTTMCCWWKGCAYRMWWRKKYRYYCTGTWRYMCAAYMKCYYYSCY
44    WYYRMYYRCTGAGMKSYYAYTTTKATGYRMWWRKKYRCCMYSKWRYMTGGCMTAYYCCYC
      * **         * .       *. .**********    ..**** .. *..** .

35    KYAYKTTKCATAMAAMKRRTWYCTGGCRRTSWMYKYYRKRGCRAGKWYCGATCAAYSYWR
44    KCRYKWKTTGCGMCMAKGRCACTWTATRRKSWAYKYYATGRYRWRTTYTTCYSCGCSYWR
      *  ** .. . .*.  * *      . **.** **** .   *  . *  . ... ****
```

```
35      TRYRWYYRTTKYGCAGRKARRMWTGRRYRGTTGYRKACGKKGTMYRCATYKYYMKYYMAA
44      CRYGWYTAWYKYRYRRGKWAACTAARRCGTCCTYRTCGRKKRYAYRMWWYGTCAKYYMGW
         ** **    **     *     :.**      **..  **   **   *    ****.

35      YTATARSYCTAKKTKYKWYKKWWYAWSKYKYGGAWKMYMCRGCKYWWYYMMKGKYRTMYR
44      YYCWRAGYYWRKGCKYKWYKKTTCWWSKYKTAARWKATMYRRYKTTTCTAMKRGTACAYR
        * .   .*   *  *******    ***** .. **  * *  *      **      **

35      RAGWKKRAYKMMYRRYAKGKKKYYRRKMGTCYGGRKKKRMKKAARRKGGTATWMMGACKG
44      RRCWKKGCYGMMYRRYMTRKKGCTGAGMRAMYTAGTKGACKKGMRRKAAYRYACCRMTTR
        *  *** .* ****** . **      * : * . .*   **. ***..         .

35      AYRGTRWRYWKYYYWSATCYRYYWMRAKRRMKKWRARCGGAAACRKYRRRRRCTWKCKKM
44      GTRTWRWRYTKYYYWSCCYYRCTAMACKGGATKWRGGYKRCGRTGKYRRRRRYKWKYGGA
        . *  **** ******.  **   * .*   .***.    ..   ******* .**

35      RAGCCGACAATMRKKAYYMYGYRRRRTGRGKAGGYMTKYTTWTGATKTARRYTCATMTTC
44      AGATTTWMRRKCAKKWYCCCRTGRGRKRRKKMKRYCAKCYYTCSMWKWMRRYKAGCMWKY
         ..       .  ** *      * *. * *   * :*     .  *  ***... * .

35      ATWCCKCMYTAKGYYCYRTKKRKYYRYGKMYYRGRRCRWSGGTWGCCKRKGYYWWCATRC
44      RYWYYKSCYGRTAYYYYRKTKRKTCGYRKMYYRRRGAGTGATCWRYYKRKSYYWAMWGRY
          *  *. *  ..** **..***   * ***** * .  ..  *   ***.***    *

35      MTACGKGGTAYYKSMGAMTGWTAWTCYYYGSCCTTTYMCRRSCGGATTGTRTWAYSRARS
44      CYRMRKKTGRCYKSMKMCWTTKTACTYTTKGTTCCYTAMAACYRKGKCAGRCAWCSRWRS
             *      ****      .:   *   .          .   .. . *    ** **

35      GRMTKRYYKYAKTCTAMGGGKKRTGATSYYGRRCGKKTTGGAAAKMATRKCTCACRWKCY
44      RRMWTGCTTCRKAYCRMASKKGRYKGCSYTRRGYRKKCGRRWMMKCTCRKACYRYRAKYC
         ** .   .  *:   *.. * *  . **  *   **       * : **.    * *

35      GCRCGGWCMWGTAGCTKSRRYYKYYAAGGTAGCCACCCRKRGATARAMSKKWGKRRYKRW
44      RSGTAAATCWRWTTYKKSGRYYTTTTGTACWCTARYYAGKRACAGGGMSKKWTKRGCKAT
         .  ..   *  :  .** ***.  :. .    .   . **..:. .***** **  *

35      TAGGRMGGCRWYKAAKKMRCKSGCMYRCYMYGCMYYYAWKMKYAGAAMTCAKTGMTAKRY
44      WTRAACAATGTTKGRKKMRMGSAMMYGMCCCSYMYYYTAKAGCRRCGAWYMTCKMYRKAC
         : .  ..    *. ****  *. **     . ****: *     ..    .  *  *

35      YYCCKKCTGTAKAKTRTYKRMYGCWAYYTKCCKTGTTKAKATWCKGATRRCRAGCKMGWR
44      CCYMKKTCSKRKTKCGCYKAACRTAWYYWKTTTWRYKTRKWYTSKAGGARMGRRYKMRTG
            **  .. *:*   **       ** *  .   .. *   .*..  *     **

35      TTYKRYMYYSMMRGTYCTGTTWTYAGGARCATTTGAKCYCGRGSKGGCCRYYKKYYRWMA
44      CGCKRTMYYSAARKWYYCRYYAKCMRTGGYGAGCACKTYMKRRCGKKSTGYYKKYYRTCR
           ** ****  *  *      .    .  .:  ..* *  * .   .  *******

35      WYRYRRYKYTTMMWAWTATYYMYGRGCMGCRATGYYCGGKGGAGTAKGGYSGKACTGYYA
44      TCRCGRYKYKYMCTMTYRCYYMCRARTARYAGATCCYRKKKSRKAMKRRYSAKWYYRTCG
          *  ****. *       ***         .:      * .  : *  **.*      .
```

```
35          TCTTAYRKKKYTRYKYYGRMYYYGYGAMYACGCYYKGATCAAGTCCKAKKCSCMTGKYRG
44          ATCYGYGGKKTYGTKTCTRACYYSYRGMYCMRMCYKRWATRRAYAMKTKKMGYMKKKCAA
            :    .*   **      *    *   **.* .**.      **   :    . . *:** . *. *   .

35          CTCTTYSMKRCKATGTCTAGKTCGCKMRKYCGAAGTCKRKKGRCAAGAWKYGAGTTCKCA
44          TATWWYCCKGYKCCACYAGRKATAAKMRKYTAMRAKYKRKGRRAGMACWKYCCRGGYKGT
             :     *. *   *. .    :. *: ..***** .   .. ***   *.. ..*** .      * :

35          GCYMYRCKYYYTCGKTGCRMYRYSTTTGKKCACAKWWWYRYATATGKWRTKYCACGWMWW
44          RMCCTRYTTYYWMRKKRYRMYRYGAWYTKGYRMTKAAAYRYMCTYCKAAWGYYMYKTMWW
                  * . **    *.  *****.:     *     :*    ***  :   *      *       ***

35          GCTGKCCACMKCKWWGCRCRKYRRMRKKTRARKYSKRYAMMRCCTGKYWRKKGWMKCRCC
44          KTGTKYACTMGSKAACTRYGKYRACRKKKGMAGTSKRCCCCGYACKKYWRKKKAAKYGSM
                 * .. * .*     *  ***  ***.       ***  .      .  ******    *   .

35          CTYYRKKMWKAMAKRAAAAGTGTCKAYATGTSCSGRCTRYGKAGYRKYWKWWSTGCCAGR
44          YATYGTTAWGRMGKRRRRWRYAYYKGCWGTYGYSRRYGRYKKGRCRKYWKWWSCRYTRAR
             : * .. *   *.**        .   *.        . * *   ** *.   ********       .*

35          KGKRCRCCKWRSTGCGYGGYGRYMYTCWYMYAKKYWYYSMRRTYGTKYYMMKCMWKWWWM
44          KRGGMGMYKTGGCAATYTAYKRYMYYYACCCWTKYWYCGAGGYYCAKYYCMKMATKWTAM
            *          *   . .. * .* ****          .**** .       * :*** **     **   *

35          RYRKWCGMKTRKYRYRKKMASYATGCTCATSWMRRMYSWKKAASWYCKTTRACTKTKGRR
44          RYGKAYAMGYAGCACGKKMRGYGWAYYYMWGWMRRMTSWKKTTGTTTKKYACTCKGKAAA
            ** *   .*          *** .*. .       .***** ****::.    *.   .   * *.

35          YYYYKAGTATKYGACYYGYKYYARYYRTKYRACMRKKMYTCAKRAGAYRAAWKGKMGCMT
44          TCTYKRRYTAKCKRSYYKCKYYRATTACKYRTAMRKKMCYTGKRGACCRRRWKRKCRYCC
                **     ::*    .**  ***        ***:.*****     .**... *   ** *

35          YTKARRYKAYKMYRYTKCTCMKKTCGKARKYKYRMAAAGMTKGTGKCMGGRGTCTGACCR
44          CAKGAGCKMYKMYRYKGTAAMKKATSKRGKCGCGMGWMRCWKRWRKSAKRRRYMYKRYYR
             :*.     * ******.   :.***: .*   *      *.        *     *.     *          *

35          YGWWKCCGCGRRGCYYCGMYAAGTWMGGYWRYKRYCACRTTAAAWYTCCTKRKCWKMWST
44          CRWTKTTKYRAGRYCTMKMYCGAYWMKRYTRYKACYMYGCCTTMACYYYKKRKMWTCWSY
               * *                **... **   * ***           ::        .*** *. **

35          AYCGKGSTGTRTMCYYKAATCTKRAKYRMYYAAMTKCKCYWRRTWRGCYKCCYMMMCTGA
44          MCMRKKSARCRYMMCCGGTCMYGAMKTAMCTGGAAGSGMYWRRCWRRMCGYSCCMMYCAC
                 * *:   * *      .:         *   *    .. : .   **** **       .   **   ..

35          WMKGRKWWRGGTCGRKTKKKTMRYYTCKCWRKKGYGMMTYCAKWYYKMYYSGTAGKYRCG
44          WCKAGTWWRRKWYRRKYKTKKMRYYYYKYTAKKRYRAAKCMRKATTKMYYSRWWRKCATA
            * *. .***      ** *.*.****   *     ** *     .     *     *****     *    .

35          CWKTTKCKKKKAKTAGTAACAKKAKRMWTYATMKWKKRKGRYRGTGKYTRYSCYTKYYWR
44          GWTKKKTGKKKRKACAKTRYRKKMKGCWKYGKMKWGKGGRACGRKRKCGGTCMTYGYCWR
             *...*   *** *:...:    ** *   *.*..*** *          . *      .     * **

35          AGRCACGKMRAKKWGTAYYMARSYKSSTTCAAYKCGKGGWCAYKKYYYKKTWTYGCYRKY
```

```
44        RRRMRTRGAGRKKWAWRCTAGRSYKCGYKYWWYKTAKRTAMWTKGYYYKKGWATKTCRKY
            *        ***.     .****.. .   **  .*     * ***** *:    ***

35        GAGGGGCATGTCCSTACWAAAARKSYRKGTRRKYCMKKKCSTCYAKARTCYKTGTAKKAA
44        RMRKRAMMKKGAMSCWATWRRMGGGYRKRKGRKCYAGKGTSAMTTTWAYTCKCSCGKKMR
               .  .  . *  .       .*** . **    *  *:  :.     * . .**

35        GTTRRGMGAKWYYKATKMSSTMGCCCCYMGKRMTTKCGTGGAYGTGTRRKMRSGGGGTKY
44        RKYAGRCARGWCCGCGTCCGACRMYTTYMRTGCCWKTKGCTWCRKRGRRKMRSRARKKKY
           .     .  *   . . ..:      ** .    *        .  ****** .  .**

35        KAACSWCCRWAKMYARWKTRMWMCWMMKYKMYSSCKMCRKKKAMRRYRCTWCKCYACWGK
44        KWMMSWYYAWGGMYWRAKCACTMMWCAKYKCYSSYKMAGKKKGMAGYAYKWYKYCWYWKK
          *   **  *. **  * *    * *  *** *** **. ***.*  *  .* *    * *

35        YKTWGAKKTGTGCKKYYAAGGTAGTCYARCAGYTATTAAYTAATTAYYRYRKMRTCKTKT
44        CKYWARKKGKKSMKGCYRRRKKRKWTYCGSWRCCRYWGMTCGCAGRYYRYRKMRWYGCKW
           * *. **  .. *  *    .    *. .       .   ..:  ********    *

35        TCAKRKRKYKTKYKKKGRRWTTGGTAAWRCTRKCRKTAGTATATACGKYYYYCWRCGMMA
44        YYMKGKRKYKCKYKKKRAAWYYRCYGTWRYCRKYGKCCKYRYCATSAKCYYYAARYAMMM
             * ***** *****   *     .:**  **  * .    .::..* ***. * .**

35        MTKRKTTWMWKTCTAYKTATSAWRYRARWTAGKTTTCKWSTATYRTARRYRRRATGYRTA
44        ACKGGAYTCAKWTWGTTWMYGGTRCRGGWKMAKYGCTTAGCWYYGCGGRYRAARWKYAGT
            *  :    *   . .   .. * *. *. .*    . .   *  . ***     *  :

35        KTTTKGKYKKTCRKCTRKGAKGCRGCMRYKGCTKGTGCCRYRCAGYACKRWWGGYYRRYK
44        GGGKKRKTGKYMRKYYRKSWGAAARACRYKKYWKRGCGARYRYGAYGTKRWWTKTYGRTK
             .* *  *  **  **.  ..  . ***   *    .*** ..*. ****   * * *

35        GAWRWGKYYMKMKYYGGATWKRCGGYTYRAYGTGTAYSRTCCRCKWYRCRMGYMYKTTKY
44        RWTGARTYCCKCKCCKRRKAGRTTTTATRWYAATCRCSRWYTRGGTTRAGCKYCCKCAGC
                .*  * *     .  *    : * *.:    **   *    *.   *  * :

35        GKYKGCCGGRYYRKGCMYGAGGTGACTCTCTATTCMRCCYSTYWRRRCYMWMRKGGCATT
44        KKYKTTAAAATCAKKYMYRRRCCKRYYACTKWAWTCAGYYCWYWGRRSTAWMRKRRGCKC
           ***  ...    *  **         .  . :      *. ** **.  ****   ..

35        CCAGCGGTYKYRYWACATKGATRGYWTCRWCACKCRRTRYTKAAATMRMWCAYACTAAAT
44        TTGSYKKYYKYRTWRYRWGSGYGAYAYTATAGTTMGGWATCKGTGYCRCWYGYMSWWRWK
            ..    **** *     ..  .*     .. .       *.:.  * * .* .    .

35        RGSKYTTGWRTYKRGTGTTTRYMKKWGMKWYRYRTTAATAKCCACTGGTACWTTCTTTYY
44        RSSKYGKTTAYYTGRYACGGRYAKTWKMKWTGTGKCGTGTGAYRYYKKYRTWCATGKKYY
          *.*** .    *.   .   ** *.* ***    . .: : .         * :  ..**

35        CKMCAAYSAAGGGKGTGTYYKYMWTYRCCYTMRTGKMYYRTMYRGKCCKMWSKKGCGGCS
44        SGAMWRYSCGCATGTWAKCCGYMTCCGYYCYCAAAKCTYRCMYRAKGYKMTSKKAMKKYC
          .     **.. .    ..   **          :.*  ** ***.*  ** ***.    .

35        KKRSKCGSKYRRRRWKKKKWWCARYKTACACKCAAGRKCRMAMMGSCGAAGTCKGGARKK
44        KKACGMRGGYRRAATKKGGAWTTGYKYRMRMKMRRRGTYGAMMMTSYAGGAYMKSKRRGK
```

```
      ** .   . ***   **   * : **     *     .    ** * ....  *.  * *

35    KKKRYGYRMMMACACYWGAKKWRAKCGAKKYYKYAGYGGKRTGAGKKARKRRAKMCRAKK
44    KKKRYRCRAACRTRYCATWKGWRRTAKGKKCCKYGRYTAKRYTTAKGRRGRRRKCYRWKK
      *****  *           * ** .. .**  **. * .**  :.*  * ** *  * **

35    MKTRRKKKYYKTCAGAYKKYYWGYAYCCACRRYRTCGGYKYCRGKMYWYRGATMGTRKCT
44    MKYRRKKKCTKCTTAGYGGCYWKCRCMMMYRRYRYYRRTKCTRRTATATGCTCCCAAKTC
      ** *****  *  :..*   **        ****     *  * .      :   : *

35    ACMTGWGGGTRTTRYACGCGYYGARCKRTTCKYYKKMAKGGGGYYWYAMCCCGYRKACRG
44    CTMCKWCTTCAYYRYWMRTKYYSGRYKGYYYKYCGKMRKRRKRYYWCCCATTKYRKRSAS
      . *  *       **     **..* *    **  ** *    *** . .   *** . .

35    KRYKKARRRTMRRYRSYSGCTRKKRYTRAGGTWGYYYATGCCYKYTARGKYMKAYMKWTC
44    KATKKWGGGKMRRYASYSSYWRKTGYAGWARYAATCYRCRMMCKYWMGKTCAGWCMKWGA
      *  **    .**** ***.  **. *:  .   .  *      **    .     *** .

35    TTAGTTKGKAGYKYYKYGGTAACYYRGCKKMRTMTGCYARCCYTKKRWTTKWRAKRTTYM
44    WGMRYYTRKRKYKTCKYRKYGGMCCARYKKMRYMKKTTWRMYCYKKRWCCKTAMKRWWCM
            . *  **  **   ..      **** *.    *    ****  *   **   *

35    KRKKYAMKKCTGGTAYMTGYCGCKKGATRMYYRRRGGKWTTRRWYWTGWGKYGRSYKGCG
44    KRKKYWMKKYKRRWRCCCCTGCAKKTRKRMYCGARKKKWGKGGTYWGTWAKYKGGTKTTA
      ***** *** .           .**  .***   *  ** .   **  *.**  . *  .

35    KMACAGCYKKKKRTYTMRCGTCGCGYYWWYCMKWAYGAMSGCWCRCCWYRMWMKGTTGKG
44    KMRYTAMYKKKGGGCGMGYKWYAGKCYWWCMAKWMCAGMSCTWYRSMACGMTAKKCATKR
      **  :. ****     *     .   ***   **  ..**  * *.    *  *  : *

35    GYGCCGTARTAKRTAAGRRKCWRWAGCTAAWGKGGCYYRYGKWRKRSTTMMYRCCAKTCC
44    RYRTMRYRAKMKRYMMRGGGAWRWMTTGTWWCKTATTCACTKARKGCYWMMYAYYGKYMM
       *       . **       .***    : * * .      * ** .  ***   .*

35    ATTYTRCTYRKKKRAYMYYYKRGAAYKKGYKRKMCRRTYTGYCTRWAKATGYKGKKAGTG
44    RYYYKGMCCGKKKRGYCYYTKRKGTYKKATKAGCYRRACGACYCRAGKGGRYKKKKGRKK
         *.     ****.* ** ** .:***. *    **:  .   * .*.  ** **. .

35    GRGWKGTYACGMRRYWKRRRMRARRTAGWMRATKTGYAATARACWKTTSYYRSYWYRWKG
44    SRRAKKKYCYACGGCAKRRAMACGRYWTWAGRYGAKYTMYRRRSWGYGGCCRSYWYRTKT
      .*  * .*. .     *** * . * *   *     : *:   * .*   .  ****** *

35    TCAYGGGTCTYWKTATRCGSRGCYTACCTKWKGCTTYCAGGYGGGGYGTGRKCYKYWGTA
44    GYGYSRKCACYAKKRKRYTSAKYYCGYYWKWGRYYYYMTARTTRATYRGKRKMYKYWKYW
        .*.   . * *. .*  *   * .   **     * :.    . *   ** ****

35    GSWYKRGRGGCRWTAKYGWWSKYKGKGGRTCCATGSRKATGCMCRTTRYRCATGKCKKAY
44    KSWTKRAGRCARWKGGCRAAGKYKRKKRGAATWKKSRKRYRTCMGYYRCRMRYKKTKKWC
       ** **.   .**..     .*** *   :.  . ***         * *    * **

35    GCAKWMKGGTMCRKACGRGTWGAKYKKAYKYRYTCRWYKRAYRRCAYYGCKRWRYMARMR
44    RMRKTCTTAAMYGKRARGCYWCTKYKKWCTCRTWYRWYKGMYAGYWCCRAKRWGTMRRMR
         *  . .:*  * .    * :****  . *   ****  *       .***  * ***
```

```
35              KRYKRCRRWKYMYRKKCARRRYYYTGKYTTGWWKGAKGRCRYACRGKRAGRRGWYWGKKK
44              TGCKRYRRWKCATAKKYMRRRYYYYRTTCCAWWKATKRRYRCMYRKKGTTAGRACTRKKK
                .  ** ****    **  ******  .   .***.:* * *   * * :        ***

35              AAAGMKKCKTRYSWRKGRYGKKKRKRYMKRKGKCWKCMYYRGRYYWKSYMAC
44              WGTAMKKMKWRTCWGKRATRKKKATACCTAGKKYAKTMYYRAGYYWKGCMRM
                 .:.*** * * .* *    *** .   .   *  * ****. ****. *
```

The pairwise alignments of two accessions 35 and 45

```
35              YRYRWMSWYRMGCYYMRKRRSATKYMYGYYYYKYWKYRYMGMRATKRRRWAATGRYGGGA
45              TAYRWMSTTGCKAYTMRKGRSWKKYMYRYYYYKCTKYGCCTAAMCKRRRWCRYKRYAAAR
                  *****     .* *** ** .**** *****  **        *****.   **...

35              RYKRSMYMRYYYRKTKYRYRYSYRYKRCWRKCCYCAGYYKKYTRKGARAWMRYYYYYWWR
45              GYKRGCYMRYCTGGGGYGCACSYRYGGAWRKTTCMTCCYKKTWRKKMRRWMRYYYYYAAR
                 ***. ****      *    ****  .***    :  ***  **  * ********  *

35              RRWRKSRKYRKYMMCTCTMCAYTATARTCWKCTGTGCRAYKWMMRSTKRYMMTTTCKRWY
45              RRWRKSRKYRKYMMTYYACTCTYTCGRWYWKYYSYTYGTYKTMMRGWKRYMMYWKYGRWY
                **************   :  .  : .*  **  .    :** ***. *****  .  ***

35              MACYYCCKGGWRRWRKMYMMRMMYRRRYKYMYRRYAYMTKYRKMGRYRYRCKRRYMRRTR
45              MTYCCTTGRKARRWRKMYCARMMTAGATGYCCRRYMYMGGYGTMRRTGTRAKRRYMRRYR
                *:         *******  ***      *  *** **  * .* *   *.******* *

35              WRYYKRYRYTYWYWWKMYTCWYKRMAARWAGRYYWGGTRWYRRMWYYRWRRRCYMYKYRT
45              WRYYTRYGTYYATAAKAYCTAYKRMWWRWRKRYCWKRWRWYAAATCCRWRRRTCCYTYRK
                ****.**   *    * *   ****  **  ** *   ***      *****   *.**.

35              TARWMYRRYYYAYARKKWYMYYYYACWRYWYMCKWWRKWYRYKKYRWGTGCGTCGRRYYY
45              ARRWMCRRYTTRYWRKKWCMCCYYRYWRCWYMMGWWATWCAYKKCGWSWKTAATARRTCC
                : *** ***   * **** *  **  ** ***  ** .*  ***  *.  .:  .**

35              RRRYCRYRSYYYRYKRMWMWYRMKRGYRRKGYYMKMGMMKRSRRWTKAGKWYKWRTTGTG
45              AARYYACAGTYYRYKRAAAWTRMKRKTARTRYYMKMRCMKRSRRWAKWKGTTTTRCGTGR
                  **    . ******   * ****   *. *****  *******:*     . *

35              GTYYKRYRRSGCYWGWMAKYTYRWMTCRCTCKYMWRMYRRTRMYATACRYYYKRSMRTTW
45              KCYYKRYRGCATCAAWMRKCYYRTCGTGTAYKCCTRCYRGKGACGGTAGYYCKGCMRWCW
                  ****** ..   .** *  **      : *   * **  .   . :. ** * .**  *

35              ACACRRKYAYKAKKCGCRCGKRMTGGRGWGRWRWTRCATACGKGYYYYWCWCCGGAGKTA
45              WYMTGRKYWTKGKKYRTAYAKGCCAAARWRRWRWCRYCATGAKRCYYYWAAYYRRMAKYW
                     ***  *.**     .*   ..  * **** * .:: .*  ****.      .*

35              MTTGKARKKRWMWYTCTTRYKKACTTAGRKTATYRRAWTTGTTTTCKWSTTGRWRGCRWT
45              ACYAKWGGGRTCAYWTCCRTTKWYGWMTGKYRWCRGGWKYAAYGCTTAGACAAARKTGAC
                   .*    *   *    * .*       *    * .*. .:    . .: .  *

35              MKRCYKYRRMRTKYKKKKCWTGTCGCTAACATYAYGGCTRGAMWYYCTKKKCWTYYSKCC
45              CKGTYGYGRMAGKTGKKKGTGAYMAGCGTMTACMYAKYYRSWCTCYYWKKKGAKTCGKTM
                 *  * * **  *  ***   .  .  .: ::  *.   *.    *  ***  .  .*

35              YGGTACKKGYGAKTGCWKRRSKCYRRKKMACATYYACCKRCGTGCCGAGGCRAKMRKYKW
45              YRTKMYKGKTRWKCATATGGGGMCAAKKCRYRWCCRMYGRTTWRYYRTTATRWGCRKYKW
                *  .  *     * .  .  .     **           *       : . *   *****

35              YARRCTKAYMRYWSMGRYRMWMYYKCGGYKTYYYKKKGKCTGGKRKAAACWYSARKCYTA
45              YWRRTATWYMRYTCCRGYACTAYYGYAACKCCCTTKGKTMCARKRKTTCTATCTRGTYCG
```

```
                    *  **  :.  ****  .    *     **    ..  *     .*   .   .  ***::.    .:*   *  .

35        RKYWYATAGKCKSMKKGCWSWKACCYGTCTGRGAWSTTARRYWKKAYKYAKRKCAACKCS
45        RKYATTCGAKTTGAKKRYAGTKRSYYTKTAAGTGTCYKWAATAKKRCKYTGRKYWGTKTG
          ***  : ..* .. **   . * . * . :.  . . .     **  **: **  . * .

35        AKWWTKSWYWRASRYSRCATWCTRRWRTATYRGAYRWYAAYSMMGRKGGCTCYAYGWRSY
45        RTATKKGTCWGWGRYSRMTGTAARRAACGWCAKWTATCRGYSMMKRKRKMGATTCRWRSY
           .  .*.  *  .**** :  .:**   .          .**** **    . :  ****

35        RRSKGYRYKWSYGAAATGKKRYYRKTMMAAMTRSMRRRKMGKGWYRMRRMYYRYYYTKYY
45        GGCKAYGYKTSYAWRMWRKTRCYRKGCMCTAKRSMRRRKMRKTWYRMGAMYYRTYYYKTY
            .*.* ** **.     *.* ***  *.: .******** * ****  **** ** * *

35        YARTWGGRKMRSGGKKWAKYRKRKYGAAACSATCRKRMWAKRYRARYAGCKKRMWWMKKK
45        YCGGAACAKMRSRKKKWMKYGTGKYRMWMMSTCAAKRAWWKRYRWRYGTTKKACTWCKGK
          *.   .  ****  *** ** . **     *: . ** * **** **.  **   * * *

35        MYYSSAKYWKTKKMKRRRGRRKWKCCCGKKYWCYRCKYCTTTWMYYCKAKKTGGTGASKK
45        CYYCGGKYWKGKKCKAAGKAAGTKYMMRKKTTYYGYGYTCCAWMTYMKRKKGTRWCCCKG
           **...**** ** *        *    **   *   *   :** * * **     ..*

35        YYGKTKYRMRGTKCYYGGTTATTARYWYAGAYRYRKCMRMRWKKMCWYYKMKGCCRRTRY
45        YYKKCGYACAACKSCCKRYWRYWGGTWCRRRYRYRKMCGMGTKGAYAYYKMKSYMRGWAC
          ** *  *   . *.         .  *    *****   *  *    *****.  *

35        YSYWKKMAKRCCRTMKYTAGGCCTYMRAGCMWRSKATKRRYKKACGYMRKMRGWYRTAAK
45        CCTAGKMRKRYMRYCKTCGARMSWYMGTKYCWRGGWKKGGYGKGMKTCRKMRSTTRCTGK
           .   ** **  *  *   ..  . ** :   **.  .*  * *.    ****.  * :.*

35        CGKATCCGTYYWKCRKRRRTRRRCYKCGKSTRYTWTMTKGCCGYGRTCTCGCWYKRCTTT
45        GRKWGYTTCYYTKTAGRGGWARGATKYAGCARYCTKCGGTTTTTKAYYGYRGWTKGSKYC
            *      ** *   *    *  . *  . .:**  .                * * ..

35        WYTTATRRAGTRWKGGMYCMKYMTAKTYCGRCACMRAYRYAAAKKATYAARTTKGKRKYC
45        ACCCGGRGTTKGAKRKCTYCKCMYTTYYMKRMTMARRTGTGTGKKRYCGWRKCGRKAKYM
              . * :  .  *      * * :.  *  * :  *    .:.**   . *.   * **

35        TKRYYRTCCYYKYCGKCTTMGTSYTYGRTRYKYWGWGAGRKTKWYYYYMYWRRMGCACYY
45        KKRYYRYYTTTGCTAKAGCCSWSTCTKAYRYKYWKACRRRKYKWYTCCMYAGACATRTYY
          .*****         .*.   . *      *****      ** ***    **     .   **

35        WYYYRKRGKRKGGGAKRRRMMKMYRRMSYSAGCTGKRRACWWYARRTYCYKACYKRMWWY
45        WCYYRKASKRGRRRGKRRGMMKMYRACSYSWSYWRTGGWMWWYRRRCYMCKWYCKGCATC
          * **** .**     .*** ******  ***  .     .     *** ** *   *    *

35        MSKYRMTCWGTGATTGYATTAGTGTRMKAYYKACKYKYKGYKYKKMRYMRTTKWKTTKKR
45        AGKCAMYSWCGKRYAACCWYRACACRMKRTYKRMKTKCKKYKCKKMRYMRYKKWKCCKKA
           .*  * .*      :. .    . . ***  **  * * * ** ******* .***  **

35        RTRTTTTKRKYGCTCKGACCKCGGKYTATCAKGRYRRKARSRYCAGMAAKRTYYKYAYYT
45        GWRWWCYKRKYKYKMKCGTTKYRRKYWRWYRKTRYRGKWRSAYTWAMGTTGKCTKCMTTA
            *    ****  . * .  *   **     * *** * ** *  .*.:. .  *     :
```

```
35            CTYGTCTATCGGKAKRRGKRYAGKRRGGGTCCKYRMKRWYRYKCAAAKRGGKYKKKWRCG
45            YCTRYYGCGMSKKRKRAATRYCCKRGRRRYYYGTAATGWYRYKYWMMKRCCKYKKKAGTA
                     .  . * ** ..**. **           . *****     **  *****    .

35            GWKYGAKKKGYYRYYKYYSTAAAGGKCRTMCRKKGATATGYRACSKKWKRTRYRWTRGGR
45            AAKTARKKKTCYRYYTYYCCGTGTAGMGCCTGKKRMWRYRCGMYSKKWTAYRYATCRRRA
              . * . ***  ****.**. .:. .        **           ****.  **    *

35            MAGMGGCCYYKMKMCTGCYGCMWYCGTYSYTWWKMTGRKRWMKCTGAKYGGCCKYWYKRM
45            CRKCAATYYYTMKAYAATYAMMWYSKYYCTWWWKAWRRKAWMKTCCRKYKRYAKYAYKAA
                  ..  **.**  :. *. ***.  *.  ***   ** ***    **   .** **

35            YMGCTYYRCWGKCTRGYCTKYCCGTTKKKYARGRTGGGYGGCYTRYTKCRKRYKGCTYCY
45            CCRYATTGAWKKYYRSYYWTYMYRYKTKKCGGRRKRKAYSTTCGRYYKYRKRCTKTCCTC
                  :   .* *  *.*  .*    ..** .  *.  .*.     ** * *** .

35            WRCAKRYCAGKTAAAWGYYTATRRYKCGYMAWRRWKKYAGTMATTGTTCAAATSAGKARY
45            TGYRKACMRTKYWRRWAYYWRKRGTTTRTAGTARWGKYWKYCRYYRYKYWWCWGWRGRRY
                  *     *    *.**  .*  .    .  ** **          .   . .    **

35            KKTKGKGKTYCKTAAYYTYGCKTTCTCKARGTKAAAARKSRTAYMKKSTAYKTTCARCGT
45            KGKKKKKKYCMTACWTCYCAYGYYYGAKWRRYGWMRMGGGRYRCAGKSARCKKWYWATRC
              * .* * *   .:.      .      .* *          .*      **:  *.

35            ACGTAWKKKKRTRGMGGAWRWSWWRACCAASRTCGGKRMGKSKMTGGTGYWAYRKYYCKM
45            MYSCGAGGKKRYGRCACGWRWSWTGTMMGCGRYMRATGCKGGKCGTRKRCTGTGKYTYKC
                . .   ***     . .*****  :  ...*    ..    .*     .   .  **  *

35            CAWRTYKCGYYYKRTRWKKYMYSYMYSMRGTTWRYRKCKRTRGWCSTYKKCRYYKGYYKW
45            YRTGCYKTACCCKRCAATKTATSCAYSMRKWYARYRKTKRYRRWMCYYKKSGYYKKKTCGT
                   ** .   **    .*   *  ****    **** ** * * . ***. ***

35            MAWYYCYRRYKMCYKTMCWAWKMTTYYGAARGYTGCCYMGCRATCTYYTWGKGKAYSGAC
45            MRTCTTCGRYKCGYKCCYTMTGCYCYYTGTARTKRMTCARYAGAMGYYCTKKKKTYSAWY
              *       ***  **          ** .:    .         .:  **    * *:**.

35            TGYGTGACRKKRKKMYKRAGTGCTRYKCYYGCMYRYYGYKKKMYCYYRTCRCCYYYKKCC
45            YRTRWRRTGGKRKKCTTGTAACMYGTKTTCTYACGYYSYKKKMYMCCGATAAACTCKKAM
                         ****  . :.:     *        **.******    :  ..   **.

35            MTGKYRGCTCSMTGAGAGAGKRTRATTRCAACMRAKRTTAGAGTCYYYCWGRCGGGKYWWTK
45            MKKKCAATATCCARCRGCTGCRGCCRARGTMRRKRAACAGAKYCCCYWKATTKKKYWWGK
              *. *  . : . : . . .   *.  *. . ** **::.....      *       **** *

35            TAGKMWMRRWGATRRKRKYRKCCSAGYSWWSYYKAWGCGRRTRYRYCCTTACGMGGACYM
45            KGAKATCGGTSRKRRKRKYRKAACGTTSWWGTYKMWTTARRCRYRYYYYWWTRCTTRYCC
              ...*      . .********....  ***. ** *   .** ****

35            CYKMGTARTCTYWRRKYGSWMKWKYRMGCWCYRASGKGTCGKTACMYCYTWKKRCTRTCW
45            MYKMRYWGWYYTTAAKCRSWAKWKCRMCTWYCGWCRKRCYAKKRYMYYYKAKKRTCAKYT
               ***             *  ** *** **  *     . *    .*.  ** *. ***    .
```

```
35    RATAAKYCYKGKGGCYYAGRYGKTAWCGKRSGYMMKKKRCCTYTKYYYYYYYWRAKTTTR
45    RWYMMGYAYKCKTATTCRKACKGKWAYRKGCKCMMKTKAYYATCTCTCTCCTTATKCGKR
      *     *.** * .         .    * .  ***.*   :  .         :*  .*

35    KRAKYRWRMWTKKGTGAGKTRMYYARKGACGGAYTTGGYKKGKAYWKWAMGGTRYYYCRY
45    KRGKYRAAMWWKKAACWKKWRMYYGAGRWTTAGCCAKRYKKKKGYAKACCKKKRYYCYGC
      **.***  ** **.:   * ****.      ..  :  *** *.* * .   .***

35    WCRRTWSKYMGTRKRTTGAGGKSMRYRGATKTYKCAKKYAKTWTRSYWYMAAKYRAGGGT
45    AMRRYWSKCMCAGKRYWKWRTGGAGTAAGCGAYKYRGGYGGYWGRSYWYMWMKCGCKRKC
        ** *** * : **       .    ..  :**    *.  * ******  *  .

35    KYCWKCYYKYTRGYTWMMSRTKYCCTCTTCTAATGWKCGCKGCTKAYGYWMWACKCTGTR
45    GCAWKYYYKCKRRYWWMMSAGKYTAKMWWSWGMYRWKYRYTRYYKMYKTTCTRMTTKKKG
        .** *** .* * ****  ** ..   . .   **   .   * *       . . .

35    AYARYKYWGCCKKKKRGYWRWKMGTRCGRAWAKYGWWSTMCKRRAATKTGACGTYYYRCG
45    RTWAYGCWTATKKKKRRTWRWKCAAGTRGCTGGCRAAGCMYKRRRWKKWKRYKCCCCRYA
          *  * . *****  **** .:    . .     . * ***  .*         * .

35    AYYCKKWRCRKGYKTGAACATMRGCCKKGKKCGRGCMWYYTCGTTTYYYAKKWMTGWATR
45    RTCSKKWRMGKKCKAACGTGCMGRYYKKRKTARGRYMTCTKYACGAYYYCKKWMCATCCR
         .****  *  *:... . *    ** *..    *   . .  :***.**** . . *

35    YKCYTTAAKTRRCCKRCTKYTGRRKRCKMKGTAGKKGSMKKRKAYRMYYKWRASCKCWRY
45    YKTCCCTTKYGAYYKRYKKYWKRRKRMKCKKYMRKKKSMKGAKMTAMCTKTRWGAGMWRY
      **    ::*     ** .**  **** * *    ** ***  *   *  * * ..  ***

35    MMCTGAWMKRYWWRYKCGWRKTKYRKKKWMRYYTCKCRRKGAAMRYCTSKYTTGWWYYYY
45    AMYCACWCKGYWWRCKMRTGKYKYRTKKAMRYYYYKYARKRWWMRYMYCTCACTWWYYYY
       *  ..* * **** *    * ***.** ****  *  **   ***  .. :  ******

35    SGTAGCCYCCMCATKCKKKTAGSRAKACARMWKWGTYAKATMAMYKMKGTYYYYGWKMRR
45    SRWWRYTCMTCTTKGTKKGATCGRRKRYRGCWKWRKYWKGKCRATGMKRYCYTYKWKAAG
      *           :.  ** :: .* *     *** .* *..     **   * * **

35    AGAACYYCAATGCKACGGGKACGAMKCKCMTKMYCRWACCMRKTAYTAYTGTTACSRYRY
45    GRMCACTMRRYRYGRSRTCKMYRCCTMKYMWKCTMRWGAGMAKYWTYRTCAAACTSRYRY
      .  ..          .   *   . . * * *   **.. * *       .::. *****

35    CCKKCTATKACGYYWGCRMKCKRMKMRGCYTGTYTAACMTGARWCCGMRMYRYTCCCCKC
45    TTGKGCGCKWMKYYWAMRCTMKRMKAGAYYWTGTKRTTCAACRTTTKAACYATCTATMKY
         *  . *   ***. * . ****  . *    . :  :..*       *    .  *

35    RTAARRYYKTGWTARKCRWYKWKYATCCMYGWGTGSGGKGGCCCMWYRCCYCMYRCYYKC
45    RYGMRRYYKAATKRGKARWYKWKCTGSYCTAATYKSKRKATYYMMWTAATYTMYRYCYKS
      * . *****:. .  *.****** : .   .    *  *.    **  . * ***  **.

35    AYKYWASRTCTYCRCTKCKWYWTGGTRCWMYGWYYRMYAYCYAGGCCGCACKAATCTTGM
45    MCGCTTSRWYYTMRGYTGKWYWAAACRAACCRTTYRCCCYTYMRTYAATRYKMMYMYWKM
           :**     *  . ****:.. *.      **  .* *    ..   *       *

35    GCMKGTTTGACTYCASRMTKYMMCCMMRCTCCAYWGYYWCGCAYKGYKYTGGKYCCTTTG
```

```
45      SYATCCCYKRYYYAMCRMGGYMMSYMARGCTTGYWKYTWAATGYGSYKCGAATYYYCCYT
        .  .        *. .**  ***. * *    .** * *.. .* .**  ...*

35      TGWTMKRRYCAGCYRKATATGGCTTGMYYTAAATGACCRTWRAKRGTKCRYMRTYWMRRW
45      WSTCATRAYTGCMYGTRYGWRRMYYKAYYACRWKCTATRGTARTGRYKYGYMRYCWMAGA
         .   .* * .  * .  .        **:.  . :. *    .   *  ***  **

35      CCATAAKTMRCCCACGCSTTRTTMRRYCTRMRTKTMKRYTGTYGKRMYCCACTRKKKMRK
45      MYTGTWGACGTAYRMKTGYGRKKCAGCYYGARGKWMKRCKRYYKTRMYTAMYKRKKKCRK
          : :  :   .     .  *..        * * *** .  * .*** .  .**** **

35      GKCCKTTCTTMRYRYKARRMRSWKGYWGRYKYMCGYTKAMYRTGCKKGRKRRRAAYWKKG
45      RKTYKYKMCCARCGYKWAACRCWKKTATGYKYASACCGTCYRCRYKGRGKRRRGGTTTTK
         *  * .    *  **    *.**     *** ..   : **   *   ****..  ..

35      KMMYAYGMKGCKWRYRMKGTKRKGRWSYWKWGAKYKYWYKRYAGCCRRYYCKRWRRWMCA
45      KMATWTTCTATKWRYRCGRWKGGCRWSYWKWRMKTKTWYKGTGAATAGYCAKRWGAWMYC
        **      .. *****    *   *******  * * ***  ...   * .***  ** .

35      GGCGGRMKKTGCRCRYCTKGCYTTWRRWKKYWWKKKYTAYYTGKCYKKKGAKYMWAGRCR
45      ASMRRAMTKYRTRMGCTCTATCCGWRRWKKYWWKKGTCCYYYRKTTKKKATKTATWAATG
        ..    *.*   *     ..    ***********   .**  *  ***.:*    .

35      CYCMRKRGRCGTGRAYKYGSKYRMRGWKKWTTRYKKYMGARKGTCAACGGCWMCMYYGRY
45      YCYCRKGTGTTCRRGYKYRGKYRMRKWKKWWCGTTKCCRRAGRYYTGTCKYACYMYYAGY
            **       *.*** .***** ****    .*         :.       ***. *

35      TAKSYMAACYRTKAMTRYKACKGTMCYKKRYYYKKKYAYTGWTAYRYMWARSRTGWKKAG
45      CTKGCMMGAYRYKWCWRCKCGKRYAGTKGAYYYGKKYGYYRWYCYRYCTRGSAWRWKGTT
         :*. * ..** *   * *. *     *  *** ***.*  * .***    *   ** :

35      YWYRRAWWYACASCKKKWKTKMYMTCCKACYWWYYMCKKYRAGMMYRRYTCAKGKKWKYK
45      CWYRRCTTCWGRSYKKKWGCKATMWTTGCTTTTYYASKTCRRCMMYRRYATGTRKKWGYK
         ****.      * ****  *  *    .    ** .*. *  ******: .. *** **

35      KYWYRKMACMGTGYWGYGGGKKRGKCATGKGKAKKYTATMGCKGAYYKAKKATCGYYYWS
45      KCWTGKMGYMRASYWRYTAAKGARGACAAKKKMKGYYRYARTTAGYTKGKKRYMRYYYWS
        * *  **. * :.** * ..*    ..:.* * * *      ...* *.**    *****

35      YKAAKYRMKMGWRTKYMYACRRYMAACGCGMMKWCYAKAYYSKYYKKGTKKYYMCCKCAT
45      CKCMKYRCKMKTRYTYMYCYGGCCTCTTAAMMGTYTRKMYYGKTCKTCAKTTYCGTKMCA
         *. *** **  * .***.     :.  ..**     * **.*  *. :*. *   * .:

35      TTMMWWMRYMMAKGRAGKCGCRYRCKMASRKMAMYSYRCACAKKAATTKTGTCATAYRRY
45      KCAATAMAYMCGKTGRKGATTACGAKMRGGKMRMTSCAYRYMKKTTCWKKKWYGCTYAAT
        .     * ** .*     .     .** . ** * *      **::  *.   . :*

35      YKKYCRRYYYAKCGAKAGRYYRTYTAATKYRATMRCMYWCAKKKKYRKRAATKKYCCTMG
45      YKKCSGGYYCGKMRWKRKATTACCCGTCKYRGGMRMMCWTGKTGKCGKRRRAKKCTTYMK
        *** .  ** .*   *         .: ***. ** * * .*. *  **  :**    *

35      CYCYGCCYWAKKRGRYCTCMYWKKATMTYKKKCTCTAAGKCKTATYYGCGYMYKYRACCC
45      MYMYRYMCWMTKRRRCYWMMYTKKMYMKYKKKYYSYGGSTGKYRGCCRMACACGCRGYTA
```

```
         * *    * .** *    ** **  *.****  . .... *       .     *.  .

35      KMCWATACMWRYRYSRGCTGAYGWWKAKCCCCGCCCGARRYRRMGTCTACTCKACYKCAK
45      KCTWCCCTMWRCGCCRRMYKRYRWTGCKTMYYATMYSRRRYRRARAAGCGCGKTMTTACT
        *  *. . ***   .*     * *  .*    .   . *****  :. .   *:  ....

35      YCMKCYYWWKTKYYKRKMMYWTAGCYTTTKTKWWWCCGRYRYYKKGWRTWKYTYRWKCCA
45      CTCKYTCAAKKTYYKRKMCCAKCRYYAWYGWKAAAYSSRYRYYKKKAAWAGYATGTKYAC
           *     *..******   ..  *:    *    ..*******       *:  * ..

35      CMYWYKWWGCRCRKYKKKRMYTTKKYAYRWYKKRMRMYGYAYWWCTKGWMCRCMMGGTRT
45      TMCTTKAACTRYGTYKKKACCAGKKYMYATYGKRCGMYKYWCTTGYKKAAYGYACAACRA
         *   *    *  .****   : *** *  * **  ** *      *        .. *:

35      YYRKKMKKKAKKTATCTAGTCGYASKTGKARCYAAAGAGKYRYKYWWWSTGWRYYRGTAA
45      TYGTTAKKKGKGGRYYYGAYYKYRGKYKKGATCTGCKGRKYACKYWWWSCTWRCTGAYMG
         * .. ***.*      ..   * .*  *.   :.. . **  ******  **   .  .

35      GKCYYCYGTTCKRKWRSYYMYCWKKWTKKYYKRCGCTGGGWKKKMRKKWGCAMTGCCGKM
45      AKMYYMTRYWYKGKTGGYTAYYAKKWCGGYYKRYRYKTTTWKGGAAKKWKYGAKATTTGC
        .* **      * *  .*  *  ***   ****   .   **    ***  . ..

35      RYMYGYRRRTYGGKWWRGKKYMTKTTAGTCYRKCRKTAKRRRYTTTMGTATWCCAKKYMY
45      ACCCRYRRGCTRTGTWRKKKYCAKYYGAYTYRKYGGWMKRRRYKACCRGCGWYYRKKYCY
             ***       ** *** :*  ..  ***     *****.:    . *   *** *

35      TTCKGTAYYTCTGKWKACMWYRAACTARGKTAGAWSMCWMKGTAAKTAGAKRGRWAWTRK
45      GCAKKYRYYWYCAKWKCTCATAGCACGRATCCARWSMAAMKKCWMKCCRGGRSRWTWGRT
          .*   **   .***.     ... .*.. .. ***. **    * . . *.**:* *.

35      YAACCATACGTGTKGKRWRYMYKGMAGCMKTYMGWYMWGYRCSGCTTGGYMCRYRSCGYK
45      YGRYYWYRMRCACKKKGWACCCKTMMATCKAYATTYMARTGYGRACCRRTAMACACYAYK
        *.         . * * *    * * .  *:*   **     . .          . .**

35      ACWCCGTTMGYTTRATKYRTTCACRYKYTTAAKTCTCCATACTAGGCTCCTAYWYGCARK
45      CTWTYACACKTAWGRCKCAWATGTGTKYYYRGGGYCTAWKGYYRACAAYYAGCWTATCAK
        . *  . :   :    *   : .   **   .     . ..   . .:  :. * . . *

35      RKMMCCAYGCAKRWKAATMCGGATTTMYWMYGTAGTSGMYYKGTCYYTCTGKAYYSKKYM
45      AKCCMYWCSSGKRWKRRGMTSKCGCCMYWMYRCTTCSACYTTKAYYCKYGKTGCCGGKCC
         *      ...****   * . .   *****  :  *. * . : * .   ..  . *

35      GGRMMAMRRAACTGYTGGKWRGTTTGAGCGGAKSKRCACTTCMKTWAKWCWAGAMRATGC
45      KRGCCGMRARRMYRCYRTGARKWCCATTSKKMTGKRMMYYYYMKAAGGTTTWKRCRTGAS
             .**            *    .: .   ..**      **: .        *: ..

35      TCYMYSKSMRTRYKYYKMGYYGGRYRGKCCKCAASATKSYRTCTTGCKKRSKCGSKRRWK
45      CYYAYSKSMRKRYKCCGMRCYKRRYRRKGYKTCCSCCKCYRAMKWTTKKRSKMRGKAATK
          * ******.***   *  *  *** *  * ..*. *.**: .   *****  .*   *

35      RKCKKYYCYRYTWCKYYYYRMRRMMMRGMYKTAAYTTKKRGYTAMMMAMYAYWYKMYRKY
45      ATYGGYYTCGTYWMKYYYYAMGGAAMARMCGKRRYKWKKRRCKTMMCRMYRCAYKMYRKY
         .   **     * ***** *    *  *  .  *. ***  .:**  **   *******
```

```
35      RGKTAKCGMWWYCGTAKKYGCGWRMRWYSSGKMRGKKRRKMACYMYAKMARRKYTKRCAT
45      RKKWRTYAMWWYYRWRKKYKSRWRCGTYSSKKMAAGGRRKCMYYMYWKMWGRKYGKRYWY
        * *  . .****    *** . **   *** ** .  ***   *** **  *** **

35      TWKRRKKYWGATMTGKSTCWATTRTKKCGAGTCCKTWRKKYARYGYWRWRRYMRTMCRCR
45      YTKRAKGCWRMCCYKGSCYAMYWRCGGTAGRWYTKYWRKGYGRYRYWRTRRTCRKMYGMR
          ** *  *       *      *    ..    * *** *.** *** **  *.*   *

35      YKKYYCTTTTRRKYYCMKRWMSRACCTTTCCACKTCKKGMWRCARAGCTAGTWKYRMAWW
45      TKKTYTYWCKRRKCYYATRWMSRRYYWGGTTRMGYMKKKCTGSTGMRMWWCCTKYRMRWW
         ** *    .*** *  .*****             **    .:         **** **

35      RKKYTTCGAMMRYYKKYTYSYKYTKTTKCGTTAKRCGRTCKSWKAMYGCYYRKRMTAGTK
45      RKKCWYYTTMMRTCKTCKCCCKCCKWKTMKKGCKRMTAKTTSWKGACTYYYATGMYTAAK
        ***     :*** *. . . *  * ..  . .**   . .***.    ** . * :.:*

35      GCRACMSGKWYAKCYKRTTT
45      RYRGAMSRTTYCKSCKRGCY
          *..** . *.*. **
```

The pairwise alignments of two accessions 35 and 46

```
35                  YYYYRMWRYYATWAYYTYMKMRRWYYKGCCGWYWWRRMAYMMRCKWYCTKRYCRYRYRRR
46                  CCYYRMWRYYRWTGCCGYMKMRRAYCGKSYRWYWWRRMWYMMATKWCTCGRYYATRYRRR
                      ********   .   ****** *   .  ******* ***  **    **   *****

35                  RRTTCKKKTARYYYRRKWWTTKMYYSCGTKKKGTRMWYMKRMRWMYYRCWKCGTTRMYYG
46                  RRYYTKKKCGATTYRRKWWAGKAYYSYCYKKKAKRMWTMKRMRTCYTRYAGMRGGACYYA
                    **   *** .   ******: * ***   ***..*** *****  * *         **.

35                  RTAAAYGGWMTRRMTCGCCYMYRMKYYMRYMSWACYKCCRCAGARMRTTYRRRRYKWTRW
46                  RCGRRTATTMGRRAYAKMMYMYRMKYYCRCMSWGGTKTAGTWAMACGCKTAGAGCKTKGW
                    * .   .  * **  .   ******** * ***.  * .   .     .      * . *

35                  RYYAMMTTAWTGAYMRYRKYYRKRRRYRRWYYYKKKTYTCCTGTWWTCRRTCCGKYRKGY
46                  RYYRCMWYCWCATYMRYAKCTAKRRRCARWCTYKKKAYYSTCAYWWAAGRYTTRKYRKRY
                    ***  *  .* .:**** *   ****  **  ****:* .  . **:. *    **** *

35                  YYYKYYKKRMTARYKRSMCMRWAYWYYRYGMWAMMCRYKYYMTTTTGTKRWYYRRCGATM
46                  YYYKCTKKGCYRGYKRGCMCATMTACCRYRMTWMMMRYKYYMYWKWRWKRWYYRRTAWKA
                    ****  **     ***.          ** *  ** ******  .   ******* . .

35                  WMGSRRGGGARMWYCKWCKGARWYKMMRRKTKKWYASWRWMMRCGYWCWGWYWMRMMRRR
46                  AARSRRTAAGAMACYKWYGRMRWYKMMRGKCKKWYMGWRWMMAYKCWMTRWYWMRMMRRR
                       *** ... *   **    ******* * **** .*****    *   **********

35                  WCRCRRRRGCMSYTTKTYGRWYYGRMKRRMCYYYKCCKYKTGMMCKTMYYMYARYRRRWA
46                  WTATARARCACGTGKGAYRGWCTKGMKGGAMTTYKTTTYGYACMSGYACCATTGYRRRAW
                    *    * * . .  . :*  *    **      **  .*  . *.       : ****

35                  ATCAKRKRKSKTKWYKGTRYRGKYYYYRAMRRRKYRYAWYTMWYYYWCCYSRWRKRKWKR
46                  RWARKGGGKSKYKWCGRYAYGCKYYYYGRARRRTYRYWWYKCWCCCWYYTSRWRKRKTKA
                      . *   *** **     *  *****   ***.*** **. *   *   ******* *

35                  MKMRYTCMKWKMRKSKRSRKKMYRSTCAYKWGYYWWMYYTYMRCMRYWKTRRTRMRKMCA
46                  CGCRYGYCKWKMRKSKGCRTKMTRSYAWYGTTTTTTMYYYCCGYMGCAKKRRYRMRKMAT
                       **   ******** .*.** ** . *       ***     *   *.** *****.:

35                  YMRMACGACRKKARCTAGAGYGKSMRKGAYTYRACTGWKTMWCTAACCTRSYWYKRKGTG
46                  CMRMCATGTGKKWRYYWRRAYTKSAGGRMYAYRRYGTWGYMWYGMWYYKRSYWYKRKACA
                     ***.. .  ** *     .* **     *:**    *  **      .********. .

35                  GKYWRYGYYKYKAATRWRTYMRGTCYCTWKKTYYGKMWTGKRKCGRACGAMGGATCYYMM
46                  KGYWRYAYYKCGMRKRWAGTMAKYYYYWWGKYYYAKCTKKKRKMKAMMRRAAATATYYMM
                      ****.***    .**   *    *  * * **.*  . ***        ..:: ****

35                  KKCKYRKKWGGGCTTCAARAKWTGKYWWSTGMYYYYRRATGGGTCSRCCTGACGAYYRCA
46                  KKYKTRKGWKARTKYYWWGCKWKKGCAAGCKMTCTTRRWKKATGGSRYMWKRYKTCCRMR
                    ** * ** * .  .     .**.     .  *    ** . .   **       :  *

35                  TGYACYCYGCACATKYRKCGRWARCRYYAGCTKAGAAWYWAWYYTKTRCWKYSSMRGMAG
46                  YRCRSYMCRMRTGCKYGKARGWRRMGYTRRYYKMKRRTCWTWCCCGCGTTGTSSARKMRK
```

```
                    .*      . ** *.  * *  *     *      *:*          ** * *

35              SGCTCRYGRCRYGCRKCYKWRTACYKKMSKKKMMGCRYYYWMKWYYRGYRRKRYMYKTMM
46              CTTYYRYRGYGCRMRKTYKTRCCYYKKCCKKKMMKSGYYYTCKTCCRRCGRKRYMYKCMC
                .    **       ** ** * . *** .***** . ***  *   *   ******* *

35              CAWAWSYKATYYMGRGCAYMGCCWAAYMWGKKAYSGYKKGYTGKYMRCRKTTARWMYKRK
46              YWTMTSYGRAYYMRARTGTARYMTCRYMWKKKTYSACKKRTWRKYMRYGGGWRATCTTGK
                     **  :***    .      . *** **:**. **    ****          . *

35              TRYCYKTYYTMGCARYRTKKKMYCYYKTCRKKGGCCMYRGCTCTTSMTYAAGCYTYKRAT
46              YGTMYKCTCAAATCGYRWKKKMYMCYKWTAKKRRMMMCAATATWWCCAYCGCYTCCGGCC
                    **   : . . ** *****  **   **    *  . :   . :*..       .

35              GTRARCAMRAKRYGTCTYRRKACGACKKRKYRYKYYRTCGMYWGCRMYRYSTTTGGGKKW
46              ACRRRAWMRRKGYAKYCYRRKRYKRGGKRKTRTTYYRWMRCCARYRMYRYSAWYTRKKKA
                . * *. ** * *..  ****      *** * .***        ******:     **

35              WWGRKWRTWKYGCYRCCACAGGKMYKKWWGCCRKKRRMGTTKTGRKYARWKAYKRACAYG
46              AASRKAAWAGYATYRYACTWAAGMYKGAACTYGTKRACKGGKATRKYMATKGYKGMARYT
                  .**     *. ** ..  .. ***       .**     *: ***   *.**  . *

35              RKSKYGTAMMRTMMTATGYSKGWMKCRTTYKKMKKAAKKRATKYCTGTCGYATAGATGGA
46              AKSKYRWRCCGCMMKMWKYSKKAAKYRKATTTAKGRGKKRRYKCGYAYYKYWGCAGACRG
                 ****       **.   ***   * *.: .. *  .***  *   .   *  ...:  .

35              CYRYSYMRRKTKTCTAGYKCKWYYWYWWSWRGGGGTMTTAAGKKRCTASYYRYTTCKAWR
46              YCRCGYMRRTYGYYWGRCKAKTTYWTWWSWRKRRRCMYWMRAKGGMYRSYYRTYWYKGTG
                  * .****.     .  *.*  ** *****     *    .*     ****    *.

35              SKTYYMMYGTCCWKGYKYWTTKYYCTGAWKCKKMRKWTGRARGCCGKMRGKTMAGYMYRY
46              GKCYYMCYRYYYAKKYKYWWCKYCYKACWGYGGAAKWYRRGRATTTGCAAGKCTKCCCGY
                .* *** *     * ****  **  ...*      **  *.*.      . . :     *

35              RRRWRTCKWTWRGKKAGTKMKTTCGCTKTTCCRKRWRTRYTTTCTTTMTRRWCMAKCKCM
46              RRGWRCAGTCWRKKKMKYKCKYYMRYWKCGTTGKRWRKRYYKYYKACMWGGWYARKGKSC
                ** ** .   ** **   * *      *     ****.** .  .: *   *   * *.

35              YTTGRTAKGAGGYTCTTGKTAYYRYCKMYYRYATRRSCTKKGCWSMAKCGTGYRKRRKRG
46              YGCKRYRTKCARYWYKCAKGGTCGTYKMYYRYGWRRSMGKTAGWSMGKYKCRYRKRRKRS
                *   *  . .. *  . .* .     ******. ***  *.. ***.*    *******.

35              GTACRTAKTKRYKMCATACGTGTTCCKRCYTCWCCWYYKGCMYMGAWTCYYTGSCCCTTC
46              RYRYRWTKGKRTTMYWYRMRYRYCTTKRYCCTATAWYYGATCYAKMACTYTWAGYTTCCM
                    * :* ** .*            **      .*** .  *      *  ..

35              TTYMCRRSGYTGCTAGCGRYYWTATRGTTKKCCRKAWKYWYYWRRGTSKTYKKGTGKYYM
46              YYTAMAACKYYKSYWRTKRTYWCRYAACWKKTTRKMATYWYCWRRKYGKYTKTSWRTTYC
                       . *  .     * **    .  **  **  .*** ***  .*  *..  . *

35              YKCMMWKWTWMRYCKACMKAACGCRYRKAMYMWASYKSRYMGYKAAWKCGTGCYRRYYYY
46              YKMAATKWYAMRYMGTYMKWMATTACGKMMTMWRGYKGRYCRTGTTTKTKWATYAATCTY
                **    **  ***  : **  .     * * ** .**.**    :: *   . *     *
```

```
35      KTAGTATYGAAACGKYYYRWKYKRSSGTCCCAYARYYRTKYRWYYACMRYKKMYRYWWWC
46      KGRRYTACKRGCTCKYYCRTTCKRSCAAMSMWYRATTACKYRWCYTAMRYKKMCRYWWWT
        *    ::   ..  *** * . ***..: .  *      **** *:.****** *****

35      AAYRKRAACRGYCCRKKYMGCYCYMGCMCACKRGKTCTTWAKKGTCYKYKTMYWYTRRAK
46      GRCGKRRRMARTYYAKKYMKMYMYCRYCMGYKRCKCYYWWMTKSYYYKCKWMYWCAAGGK
        .   **         ****  * *     . ** *    * .*.  ** * *** :  .*

35      MYRKKKRSCCGYYTARKYKYMMATMYRGCTGACRRCYGWWAYYKAKCRATCRGCGAGGCR
46      MYRKKGRSMMSYYYRGKCGCMCGWMYRRMYKRYRATCRWTRTTGCKTRWYTRRYRTTATR
        ***** **  .**   *   * . ***      *    *     .* *   *   : . *

35      RKWAGRYYKWTKYCYCTKAYMGGRYRMWMYYKGGCTYKTKARRRRKKKCTTCKAATYWWY
46      RKARKRYYKWYKYYTYATWYMKRGYACTAYYKRKTYCKCKWRRRAGKTTGCAKTTCYAAT
        **   ***** **   :. **   *    ***     * * ***  *.   .*:: *

35      SKWAGACRTGCRCKRTTGGAGSMKGSWKYYCGWSRRYCAGCAGGKRYGMTAAAYKYKRKC
46      CGAGCTGRAKARTTRWYSRGAGAKRGTKTCTATCAATMRKYRKTTGTACCGTRCGYTRGY
        .  . : *: .* .*  . ... * . *   . .          .  .  .:   *.*

35      AKMSKWATGKTAGTAAYTKWSRRYRWYGRRTCCTRSYRWRRTSGWGTKMYYKGGMRWRSM
46      WKMGKACCRGCRKWMRYYKWGRRCATCKRRGAACRSYRWGGCCKTAGTCCYKRRARWRSM
         **.* .         * **.**     ** .. *****   .  . .  **   *****

35      RSRRKTARMYKGKYRMRKRMYYRRYKYKRYTAYRTGRWRRYKAGTGKACGTMTKKYRYKK
46      RSRRTYCGMCGRKYAMGGAMYYRGTKTKRYKCYGCRRWRRCKCTCTKCGRYAYGGYRYGK
        ****. . *   ** *   ****  * ***..*   **** *.   *.       *** *

35      YAYTWGSYYRYYKYTAYRYMWTARSRSYRAKTYRMRYKWYKKYASKKYRWAYYMMYWWYY
46      YWYWARSYTGCCTCYCYRYCTCRGSACYRGGCCAAGYKWYKKCTSKKTRWGCYMMTTTYY
        * *   **    .  .***     * .**.      ****** :*** **. ***   **

35      MCAGYRAMKWGRAGKWKCYKKYYRKMGYKMGGAAWYGYMKAGRKKCWKRAGTTKYKMKKA
46      AMRRYRMCTARRRRGWGYCKKCTAKMRYKMTKRGWTRYAKRTGKTMWGAWSYYGCTCKKM
            **  .  *   *   **   ** ***   .*  * *   *. *   .    . **

35      RMKKTATWMGRGKTAKYYYWSKATMRWCCRKSKGCRSMCGAAGGWRCTATTACRRAMMAT
46      RAKGYRYACRGKKCGGYYYWSKCCMATYYRKGTTTRGAAARGCAWRYGGCYCTGGMAMMC
        * *         * . ******. *    **..  *. .. . .**  .  .     *

35      RKKKTYYRMYCCGTGACAKKKMRAAAYYYCTCCMGGYRYKYGARRMMRSKAKKYWGKYKW
46      ATKKKYYRMYTAKCAGTCKKKCRWWMYYYGKTYAKSCGYKYKWAACMRCKGKKTATGCTA
         .**.***** .  .. .*** *   *** .    .  ***     **.*.**     .

35      MYRYSCGCRAAKYKSMKKGKMMYAGGCKAYGMRYKKMTTWRYRMGRAAKKKARSGRRRRW
46      CYRYSYRYRGGKTTSMKKKKMATGRAMKWTKCRTTTCGCWRYRCRRWWKKKWGGRARRRW
         ****   *..* .**** **  . . *    * ..   ****  *  ***  .  ****

35      STACGAAAGKRTYYYYYYCTCCSTTACAACKGAGGKKAKYYTTGRYRGGGTKYRRRGYYY
46      SWGMATGTATGKCTCTTCMAYYSKCWYRGMKSRRKGGRGYCYYTRYRRRRYGTAGRTTYY
        * . .:.:.. .       :  *.    . *.       *    ***        *  **
```

```
35          YMAAAKMRKKKMARKRCGGWGRKKGKYRYGRRYYKYRYYYAAGGKCRTTAKGGCTRKATA
46          YMWMMKCRKKKMRGKGTAAAKRKKTKYRYRRRYYTYRYYCTGTAGAGCYWGKCTKGKMWR
            **   * *****  *  ..  *** **** ****.**** :. . .        . *

35          AYGKYRMMKGKGYTARRTYKRWGGRMAGGGAGGCRYGYTAGRRKMRTATKCCMGGCMYYS
46          GCRKTGMAKKGTYAGRRGYKATRRACRKRRAATACSYKRKGRGCAAGAKMAATAAMYYC
            .  *  * *   *:.** **           ..   .*.   *   :.:* .  ..***.

35          YKMYYYKKKACCCMKRKRKKCTGGTAACCKAKTAYKRMYKGRTYYRCTKYTKGTTTYKKK
46          YKMYYYKGKCYYYMKRKAKKTCSCKRTTAGGKGWYKAACKRAATTGAYKYWTAYKKYTKG
            ******* *.   **** **  . . : . .*  **   *  :   . ** .. ..*.*

35          TRAWRCWATTGWTRRCKATRKKRWYWRCKRCASYSYKAWTGCGRRCCATAMAGAGKKMGT
46          WAMWAYWWYWKWGGGYKWKRKKGTYWRYKGATSYGTKMWKTTARRGAWCWMRKWRKKCKG
               *  *    *    * .***  *** * .:**. * *.  .** .   *    **

35          YWYGTARWMTCYYWRRKYYTYTAARMMAYMTGCKRTKCKMWKKGKWRRATAAKWAGGCYY
46          CWTRYWGWMWYYYWAAKCCCCYGTRMAMCMKCTGAKKTKMAKKRKWARWYMMGWWTATTC
             *     **  ***  *     .:**   *.    .* ** ** ** *     *  .

35          KGRYGGAWRKRRMMKAKRCCGTATTSKKKTTYRARCYCMGGWYYKAKRTTAKKKYMRKMC
46          KKACTKWARKGRMMTTKAYMRYGWYCKKKCCCRGRYCSARRAYYKRKRGCTKKKTMAGCM
            *       ** ***.:*     .  .***   *.*  .    *** **  :*** *

35          CGGACWAAYKGYKAATWKKKYWTGTKYTTYGTYCCRYGWGRTCYYKWGYGKCAYKYSRKC
46          TTAGYAGRYKRYKGTAAKKKYAYKKKYAWTRAYATRYRTCAYYCYGATYKKSMCGCSRKT
              ..  . ** **.:: ****   .**:   :*. **       *   * *.    ***

35          GRGCCKWYWRCWSYYMYKYTCYKCAGAATCGAAGCCGCCYCRRGMMTGGCYTGACTCCRG
46          RRKGGKWYWRAWGCYCCGCYTYKMRRMRWYKRRTYAATTCYAGKCCWKSYYYKRYYYTGA
             *   *****.*. *      **            ..           . *        .

35          YKCCYTSMYAGKCYCCYMMAKGGCARCGTCCAAYYGYMCMWKTYWCYKGAAYWTRWCTGT
46          YKAYYYCCYMAKGYYMTMARKRRGCRSKCTTGRCYTCCGMWKCTWMYGSGMCAYATMCAW
            **. * . * .* *   *  *   .*.    .  *    ***  * * ..        .

35          RKCCRRRTRMMCWMTRRKMGRKRAATTTCTRWRRTKRGTSCAMKCGTGGRRKGMTTTRRR
46          GGYYRRRWGCCMWMYRRTAKRKRRWKCAMKRTAGYTGRYSYWMTTAWTKRGKRMWYYGAG
                ***     ** **.  ***  . : .*    .   *  *. .   * * *

35          WTATAKKCAKTACTGTTKMRRRRYRWWYAARCKTTCMKACCTTTCRMRTMRYWRKCCYGC
46          AATGTKGYRKYRTGRKKKCAGGGCATAYGCRTKGCTMKRYTAYKYACRACRYWRKYYCCG
             :: :*   *     ..*         *..* *   **   : .   *: *****

35          RACRWRYYTYATTCTATGGTRWACGGCTYKRMYYKGTCMCCYYYRYYTYRTYAMRRRKYT
46          RGYRTRYYWYRGWSCGKKRYGAGTACTCTTGCYTTKATCTYYYYRYYKTGYYTAAGRKYY
            *. * *** *   . ..     . .    .  * . :    ******.   *:   ***

35          YTAGCGAMARYRGKKYMMMCATTTRCGAKYTGAKAACCTAYYAMRSTKTTCRKAACWSKA
46          YYGATCGAGRTRSKKCCCCMMYYCASKWKCYRTTMMGMWWCCMCRGGTACARKCRYASKG
            * ..  . .* *.**          .  *   :.          *. .: .**.   **.

35          ATCMCCYMATGYGYMTTKRRTKRTSTGCGRKYGCRYKCRTGCTAWCWYAYRRCACYAKGK
```

```
46          GGTMYYYMTGACRYCGCKRRKKRGCGAMRRKCAYGCTMRYRTGCWMWCCTRGMRTCGKTK
            .  *  **: .  *   ***.** . .  ** .   . *    .* * . *     .* *

35          YYWWAWGTTGTAKCYGRYMCMYYAYYTGTYCTKSYKGMKATTGKMRACGRYKRKGRYYKT
46          YCTWRWRYYAWRKYYRRTAYCYCGYYCKCTTCGSYKKCGRYYRKMRRTKRYKGTRRCYTA
            *  * *   .  * * *    * .**       ***       ***   *** . * *.:

35          AKATKGRGTTCSGMGRAAAAARKSATGAGMMMTARATAKYCYMKCMKKKCSTCCYKKCWW
46          CKRYKKRAKGASKMRAWWRRMGGGWYRMRMMMYRGTKTKCYCAGMCGKGTSKYMTTGYWA
            .*  * *.. .* *         .     ***   :.:*        *  *.   .  *

35          KKGTTTRRCGMRTCWRWYTGTTMSSATTCMGYRKGTGKCCKMYSKASMRKAYCCGRYYYK
46          GKSYYYAGYRMRKTTGWYYRYYCCGWCCAMRTGKKKKKYSGAYSKRSMRGWYTMRRYCCG
             *.       **.   **     ..   .*   * . * .  *** ***  *   **

35          MWGTYYTRTTCCTGYMYGAKTMYRGKCCKMKCTTWSATKKRYRRSKSKRRWKKMKKKAWK
46          MTRCYYCGYYYYWRCCTAGKYMYRRTGYKMKYKKTSCCKKRYRRSKGGRRTKKAKGGGWK
            *   **           ..* *** .  *** .. *. ********. ** ** *  .**

35          CAYRYKTCCCYAGRRYAMMKGAGRRCYRKKKRKYRYCMMMYAKAYAKKMYRKYRACKCGT
46          TTYGYKYMMYYRRARYMAMKAGAGGMYRKKKRKCRYYMMCTRGRCRKKMYRKYRRMTYAY
             :* **    *   **  **...   ******* ** **       ********  . .

35          CTKKYKRYCGGGWRRWTRKRKTWRAAKMACMAMMARTYCTKRAAGTTTGGRRYAYCCCAR
46          SYKKCGRYMKRRWRGTYRGRGYWRWRKCMYMWMMWRGCYKTRTWRWYCTTGACTCTTTGG
            . **  **    **   * *  **  *   * ** *   ..*:          :    .

35          RKKYGCMWYYCTGTCGTTTRYACCWMGCRGWRYKTTAAAAGTCATTYKAGYCRCTKKYTG
46          AKGCRYMTCTMKRYYACGAACCTTWMKAASTRYKCCTGTMRWYGWCCKMSTYRYKKKYWK
             *    *    .   .  :  .  ** . . ***  :.:    .   * .  * .***

35          CRKKCMGAGKACGGSRCKTTTKRYKAYRRMYYKKCYRMWRKKYMMMMCTGAKWMRRRWYC
46          YRKKMCKMRKGTAKSRMKCCWGAYKMTRAMYTKGMYRMWRTGCCMAMYCACKWCGRRWCM
             ***     *. . ** *     **  * ** *  *****.   * *  ..**  ***

35          KCWGRARRYKTKKWMRYYTCKCWRGTTTCMGACCMMYWCGATMTYYTSTGCCKCCGCWRC
46          KYTRRGGATKYKTAMRYYYYKYTARYCWYACGTAAATWYKWGMATCKCAAYTGYTAGWRM
            *   *.   * *. ****  *          . .   *    *:  ..:.     . **

35          CSYGKCTGARTKCKKKAKACKAARMWKTYAKKATTYTMYKMKKRKRMRMRRTTRGWGTSY
46          TSYRKGCRGGKKTKKGRKRYKTTRMWKKYRKKGKYCAATGMKKRGGMRMRACYACWRWSY
             ** *   . .* **  *  *::****.* **..  :   ****  ****     *  **

35          WYKCYRCGAYKYTACRYRRYYGCKRWRRKMARWTMTCKGCGGMKKCRYRWYMYRCKKGTT
46          WYGYTRYRRYKYWRYGYAGCCTAKRWGAKMRRTYCYYKSMRRMTKMGCRWCATATKTTAY
            **   *   ***    *     .***  ** *     *.   *.*   **     *. :

35          MMWMKCKARARRYYYTGKYKKKKGAKGRCRYCYGKRRCWYWGRAKAATCAKYSKKKAGCK
46          MMWCKYGMRCRAYYYYRTTKKKKATKRRYRCYTKKGGYACTRRGGCCCMMKYSKKTCKTG
            *** *   *.* ***  . ****.:* * *    *       *. ..   *****..

35          CWSTRCGWRYKKRKRYMKKATCWAGMYGCRYCCAKSYMTCCTSTCRATTKKTCAKTKATC
46          MWSCGYATATGKRKRCCKKRYYAGAMYATGYYSRKGCMWYAYSCYGMYCGGCATKYTWCS
```

```
       **   .    ****  **    ..**.  * . *. *  . *         .:* .  .

35    WGRYTGWYRCGGGARRYMRSCKRYRGTTKAYYYTTMGYCTTCCTARATCCTACATCCCKR
46    WKGYYRWYRYAKRMRRTCGSMKATRKGAKMYTYYWMRCYCYSTYGRMGTYARMMYYSMKR
      *  *  *** .   **   * *  *  :* * *  *     .  .*    :     . **

35    RACCRGCMWRCARAKWRRYYGGGTWKCAKWWRKKCYCRMYMKKCYYCSCYYTKATAMGYA
46    RGYMRRMCTGYMGMKWRRYYRRSYTKMRKWWRKKYYYRMYAGTAYYSCYCYCTGCCARTG
      *.  *         ******  .  *  ****** * ***  ..**..  * .. .   .

35    KRCWKTAARRTSWYGRKRMGRGCRMSGKWYAWTCRYSYWRKCARYCCKYRSSWTARTKYC
46    KRMWKARRRRKSWCRATGMAARYRMSRTTYCWYSRCSYWRKTTGCTTGYGSSACGGYGYA
      ** **:  **.**   . *.   *** . *.* .* ***** :     * **  .   *.

35    MCGCSMATGRSAKMMWTYYAYGAACSRRTWAAYGCARYKKKRMWGMWKMYAATAMYKKKK
46    AYRYCMRARRSMKMAWWYYWYRMMMSARGWGGYSYWRYKKKACTAMWKMCGGCGCYKKKK
          .* : ** ** * ** *    * * *..*.  *****   .**** .. . *****

35    MYSSCKMTKAGMKGRRGARWGCATARTAKCCWYCTTTRWAYKAAKKTAYAAKKTACCCYA
46    CYSSYKAKKGAMGTAGRWAWRYRYWGKGKYSWYYCCARWWYGRRTKGRCMWKGGRMTAYR
       *** * .*..*       *      ..* .**   :** *   .*     *     .*

35    RAKRMWYGTAAAYKYYYGACTWYYCAAYRRYRKTKYCCAACRRTCCAGGYYKYCRGTKMY
46    GTGACWYRYWRMTKTCCKGYYWCCGCRCRGYRKWGCMMWMYRRYYTCAATYKCTRAWTAT
       :   **      *    .  *   .  * ***        **   ... **  *. .

35    GATMGTRMTAGARGCCGTGTRYKKGYWYGRKCRTTCWYRMMYGYTYKYRGCAYYWYYYRK
46    CTCCSWRMCCRWAKMMCYRYRYTKKYAYRRKYGYYYAYGACTRCCCKYGATRYYWCYYRK
       :  . ** .          **.* * * **      *        ** .  *** ****

35    SKRGKRCKRYKWKAGWYAMMKARMMYMRRRYRMSYSAGCTRKTKRYRATRGYAKRKCTTK
46    SKASKRYKGCGAKGATTWMMKGGAMYMRRGYACSYSWSYWAGCTGYGWKRRYRTGKYCWK
      ** .** *    *..   ***.  ***** *  *** .     . *  .* * . *   *

35    KYYAKTRMWWYMSYRKKWGTTTTGATTYAYCMCGGRTCATTAGKYYCYCCGACRSKAGCK
46    KCCWKYGCATCAGCAKKWCGGGCRRYKCMYYMYRRRYYRKWRKKTYMTYTAGYASGGATG
      *   *       .  ***        .  * *   *   .   * *    ..  * ..

35    YGCAKKMRCTYRRTCCCYAMRCMYTWKTTKWKRKRTTKRKSWATACTCCCGGKTATAGRM
46    CRYGKKMRAATGACTATTWARMCCYWKCCKTKAKRWWKRKSWCGWYKMYYRRKWRYRTRM
         .****.:     .    *    **  * * **  *****.   .     *     **

35    RTTTCAACACCCTTAGTTKYAKGKKRMSGACTTATRCRGCAAKYYYGCGCKCGGGAGMYY
46    RWWYYMWYMTYYYWRRCYKTGKCKKGCGRGAYYCCRYGKYGCGYYYRARYGYRRRMAACC
      *                 * .* **  . ..  . *    .. *** .        .

35    TMAMTKTWMWRTTYCKYTGTARKRRAKGKTTTCTKWSKTTGCRWRYRGCTCTYTTRYRTY
46    YMMACKYTCARWWCYKCGAWMGKRGGKAKYGCTKTAGKACAMAARYGRTCYYYYCGYACC
       *   *    *    *  .   ** .*.*    .. .*: .   **      *   *

35    AATGRTTRKKYKKKCGCTAACAARGYCGGYYGCCRGAKMWAYGYYTTYATATTCCTAKRG
46    RRWKAGKAKKTGKKMRSYGTMTWGRCMRRYYKTTRSWGCTGCKYYYYTGCTWCYMCMKRR
            . **  **  . .: :       **   *.    .  **   . :      **
```

```
35                  KCMYKCTGKGYWWCYGGACTKGYKWWCSTRRSYRRMCYMYYKMCGTCYYGCKGGACKCGC
46                  KGCYKGCRGKYWWMYATGTYKKTKWAYSKRRSCAAMYCCYYKCTACTCCRAKKRRYKTKY
                    *  **     *** *. .  *  **  *.***   *   ***  .     .*    *

35                  YMWKGGAGRACTAYMRWKRYCTGYWRYTMRYAATTCYRCKSYKMGAKRGTYMYRWCRSTY
46                  YCTGKKWRGRYYRYMGTKGCYCKYWGCWAACGGACYCAYKCTKMKRKRAYTATRWMRCYY
                    *            **  *     **      ..:     *. **  **.    ** *. *

35                  MAKYAGKTTRTYMWRSKKRYYMGCCKKKKYCRMKYTRRRGWKKKCGYCYWKRRTRTRCCW
46                  CMKTWKTYYGATCWRGTGGYTCATGGKKKYYRMKCYRRRSTGKKYRCYYWKGGCGYGYST
                      *   .   :  **..  *  .   **** ***  ***.  **    ***       .

35                  YKRKKYTRYGTKCGTCTGKKTTKAWYYTRTGTRTRTRWKGGMYCMTTGGCAAMTCYRRYK
46                  TKGKGTARYRWKAAGTCRKKKKGWACYCRYSYGCRKGAKRKCTYCCKTATGWMYMYRRTK
                     * *  :**  *..    **..    * * .   *.  *       . . . *  *** *

35                  TCYTMAMYCTKKGATACAKACSYTYTTKRTCRAGAACYWWYKATGACGKATTCTCCGWAG
46                  YMYCCWCCYAKKARYGMTGCYGCCYACKAATGGRMCACWWTKRYRRYAKGCKTCGATWCR
                      *      :**.  . : . .  *: * :  .  .. ** *     .*. .   . *.

35                  AGCKRMCMCCYTAYTGWRMCGMCGCCYYKCMTGGYCWYKMRRTYRWMMKGRMKGKACYAC
46                  MKYGAAMMSYYYRTCATRCYKMMRTTYYGGMYRKYYWYKAGGACRACCKKGMKAKCATRG
                           *. *    . *   *    **  *   * ***   : *   *  **.*..

35                  ACATRMRMYRTCCCSTCAKRATYTRRCYGWTRCAGYRTKCWTTGWTRWYYMTSTYATMYC
46                  TTMAGAACTACTATSAATKRMYCYRRYCKTCRTCRCRCKATYKRACGWTCCCGACWKCTM
                    :  :        . *:.:**    **     * .  * *.  .    *    .:  .

35                  KTAGCMSGGGWTTKGGGCMW
46                  GCGATMGKRRAWAKSRKACA
                      .. *.     :*.  .
```

The pairwise alignments of two accessions 35 and 47

```
35              TGAAWMRRMYKYYYYYRKWRKSRKRGYYYRRATWMMMTSRRKWMKYMRYKSYKCRRKYRW
47              YRRRWMRRMYKYYYYYRGWRKSRKRRYYYRRWKAAMCWSRRTWMKYMRCKSYKYRRGTAW
                    ************* ******* ***** .  *  ***.****** **** **   *

35              KKSGAYYRYAAGMYKKYRKRSMGTYRYKWYCCCYCGRCMWRYKYATTKKCCWRMTKCAMY
47              KKSTMTCRTRRAAYKKYRKRSMAWYRYKWYYYYYAARYCWAYKTCAYTKYYTAAYGYRCY
                ***    *   . *********. ******   *..*  * ** .: .*          *

35              RYRCGSWGRYWWYTWKCYKCMRWYYWYKKYRKRKRYSGRWSKMYMMRRKYMCCMYRRKYR
47              RYGYAGWAGCATYGWKYYGACRWYCTCKKYGKRKRYSKAWGKMYMMAGKYMYYACARKYG
                **  ..*.    * ** * . ***   *** ******  *.*****  ***     ***

35              WMMWYRKRWMAYYTCRAGYCTYYRYYYWRTRYYYMAMTYRRKYGSYWRYRYTKARGRMMR
47              WCATYRKGAMGTYGTGWRYMWYYRYYCARKGYYCMGCCTGAGYCCYARYRYYGMRKRMCG
                *   ***  *. *     *  *****  *. ** *.       * .* ****   * **

35              TKRKMACCWKTTAGCRRRYRKTYYKMKRTRWRMKMGMRRYYKTGYYMKKYYKRMRYMRYY
47              GGRTMTAMWKWYRRYGRRCRKYTTKCKRYGTACGCTMARYYKYKTYAKKYTKRMRYMRYY
                  *.*:. **      ** **   * **        * ****   * *** *********

35              YWASYYYWRKCCMCACGRGGYTYACMMRKCYTTTSMKGYGGTRWYACYKRRTMWGYYRKY
47              YWWSCCTWRKTTMMRYARRKYCCCYMMRKYTYWKGCKKCKRWRWYGYTKRRWATRCCRKY
                ** *   ***  *   .*  *  . ****    .. *     ***.  ***      ***

35              YYRSYCTCMYYRYRGTMYSTGYKKWYYMKACCARRMCYTGAKGTYWWCKYYMRWKTYYRG
47              CCAGTMCYCYYRYGRCMTGCATGKWCTMKCYYTGGAMTCATTRYYWWAGYCCGTKATYRA
                   .     ****   * . .  ** **.  :      .:.  ***. *   *: **.

35              YYYYYMRYRWAARRSYMRRKRKAYYTYKYKAMCRWKKYMRWKKRYWTGCRKWKTRYMYSY
47              YYYYCMRYRACRGGCYMGRKRKGCCGYKCGWCTGTGKYMGTKGGCWCTYGKWKWACMCCT
                **** ****  .   .** ****.   **        ***  *   *    ***   * .

35              MKRKGTKKRKKTMAYYGKGCATTRMWRKKATKKKRKKMRGKYTRRRGWKTKKMKAATCYC
47              MKRKRYKKRKKYCMTYRKRTGYYGCWRKKWKKKTGGKMRRKCYRRRSTGWGTMKGTGYCY
                ****  *****    * *  .    **** .**.  *** *  ***.    .**.:

35              YRTRTRCWKRRKYGGSTRYMKYTAAKGRTCGRRCTCGYMYSSWYTCRTRTGTRWCGGGMY
47              CGCGYGYWKRGGTKRCARYATTWWWKKAYYRGRSKYRCMYSSACCYGCRCTKGAYKRKCT
                       ***     .:** .    *      *..   ****      *  .

35              CKTYGTCCWKYRAYYYSRTCYAKYTMAMYTATAARKRYTKYTTKMMATWWTYYAAMRYYR
47              YKYCRWMTAKYRRYYTSGYMTRKYCCWCCARYGGRKRCCGYACKMMGCATKCCTTARYYR
                 *       *** ** *     **     :  ..***   *: ***.   .  :: ****

35              RMYTCAKKKMRKGAAMATTAYCWKTGTCYMAKTCKRCCGAARRMCAGCSAKKYWGKGCKG
47              RMYYGCKKKCRKAWWMTYCTTTTKYRYGYMGKKMKGMMKRTAACYWRYCRKKTAATTGKT
                ***  .*** **.  *:  :   *    **.*.*     :       . **  ..  *

35              YKYKSYKYKRWWGCKKKRRKAAKYWKWSKGKMMYCAYGMKMWRYRMTRGAATCGYGKTCG
47              YKYKSYKYKRAARYGKKRRKGGKYWKWSKKKMATMWTKCTCWRYRCYGRMWWYAYRKCTA
```

```
     **********     *****..******* **       . ****        .* *  .

35   MRTTGTTCACKCRATAAKGRTMKKRMRRWSTGCYATWRGCTRCCGYTCAATACGKYYYYC
47   MRWCKWYYMYKTRMCWWKCGKMKKRCAAWGYRTYTYWRRACRMTAYATGCATGAKCYYYA
     **        * *    *  .****   *.   *: ** . *  .*: ..:: .* ***.

35   KCGYMTTGKARKKTWMWYTKYTMTCYKTATRAWGYARARWGKCYRKYKTTCWKWSTTGTC
47   KYAYMCYAKWGGGYTCACCTCWMWMCKWMYGGTTCRRGGWKKMCGKYKGCTWTAGACAAM
     * .**  .*          .  *   *    .    *. * *   ***   *. .:  .:

35   GWGSSRMRRCRMWATGRGGCACCTGKKMKKKARYYWCTRGAKKGCKTTYYYYAGCYKCAG
47   AAACCGMRRTGMWRWKATATTYYGRKTCKKGWGYYWYYRSWGKAAGKWCYCYMRGYKGGK
     . ... ***  **     .  :   *. **   ***  *.  *.. .  * *   ** .

35   YWCKCRWGYGARWRRSYRRTYCYMKMRRYYATYAGGACCKKRGRYRRAMRACGAACTAKC
47   YWMKMATKTRWGAGGGCAACCGCCKCGGCCCCYMKRRMYTGRKRYRRGMRRYKRWAGCKG
     ** *           .        *     . *      . * ****.**     . .*

35   GAAACTTCYKKYMYKRCKYYRTKYYRKMYWMGCRMYRYSTTTGKWWWCCGRKWRTWKYYR
47   KTRMMKCYYKKYCTKRYKTCRGTYYRKCCAMRYRMYRYSAWYRGAAAYSSRKAAWAGCYR
      :   .  ****  ** *  * .****   *  ******:        ..**      **

35   SMCATGKCACTKKYWYKAWWGCCRMGRYKKKKKKRYRKYKSKAYRYGKGYYYWYCYWTKG
47   SMTTGTKACTKKKCTTKRAACTYACKRYKKKKKKGYAGTGSKTYGYKKTCCCTYGYWYKK
     ** :  *.. .**   *         ******** *    **:* * *     * ** *

35   AWMCTRTYRKKMKMAKRATKCWATGTCGTAYSKCCAAGCYTCTGRKAGKKTATKKYWYWW
47   WWAYKRATGTTAGMGKRRYKTWGAAYYKGCCGTATTRAGYYYWKRKGRKKCCCKKYWTWW
      *  .*:  ..  *.**  * *.:.    . ... : . *    **. ** . **** **

35   STMCWRTAGRKGKRCYRYCRCKWRSTGCGGGYGYMMYGTCGKGSYWAKKYKKYAYMTACC
47   SCCYWRCMARKRGGMYRYMRYKTGGCAATTAYKYMCYACTKGTSYWMKKYTGCWCMYRYY
     *   **  .**    *** * *  . ..  .* ** *.     *** ***.    *

35   YKARYKACTWYYRYRRCKKCKRWRRKGCGGRRKKRCRYWGCYTTGWRGRWKYMYRCGTTK
47   TKRRYKGYCATYGYAGYKKAKRWGAKSMRRAATKRMGCTATCCGATAKRWKCATATKWYK
      * ***.    * *   **.***  *.     .**    .    .  ***         *

35   GYTTAYYTGKYKWKKGAKYGGRCRYACRGKRRRRRCTTATYAKYGRAYAGKKYKKRSTTT
47   RTKCCYYYRKTKWKKATKCRRRYRCMYRKKGGRRRSKAWYTRKYRRRCWRKKYKKRSCCK
       . .**  * ****.:*   * *   * *  ***..:    ** *    *******  .

35   GRKAAGYTGGWRYKKCCYMKRKGKCWCCAGMYGRYCCASYMWKMTGKRYYYYAYKTCTCK
47   KGKWWRYCARWATKKYMCCKAGKKYAYAGAMYKGYYSRGCMTKMWATGCTCTRYKAYCYT
       *   * . *  **    *   *   ...**  * . . * ** ..      **:   .

35   GGYARGGKKRYYTGTYRKKYRRMKRRTCYGKMKAAAKMTKCTKCARWRGTKRCGGWAGCG
47   TRYRASKKGRYYYKKYGKGTAGMKRRWYCRTMKWMMKCYKACKYRRAGACKGTAAATAYT
       *  . * ***  .* *    ****    .**   *  *. *  *  . *  .. :.

35   GSGRRYYKCCYRYYYAAGGKRMAATGATTGCKRGRCAACTWTKRYWGYRRTYKRWGGRMA
47   CCRGRYYTMYYRYYYTGTAGGCTRYKRKYCTKRRGTGMYCAAGGTWTYRRGYKATRAACR
      .  ***.  *****:. .   :    .   **   .    :   * *** **   .
```

```
35                GWYYRRTYKCGGGACKKMATGCMYRYMYMYYYKRKRMKKGTAYTACCKYWYKRMYGTCWK
47                KWYYRRCYKYACRGTKKARCAMMCGYCCMYYYKRKAMKKKKRTYTTAGCAYKAACRAAWK
                   ***** ** .  . **   . *  *  ******* *** .   : .   **    :.**

35                TKKKKTYRYTYGRKTTAMCTCCTATTYWYAGYTATMATTWYARWKYRYYTYAGTAAAMTA
47                WTTKKWCAYYYKGKGCTMAAYYAGAATACWATARGCRWGWCGRWKYRYYWYGAARRRMCG
                   ..**   * *  *  :*.:  :.::    . :      * .******* *..:   * .

35                GTTTMGTMYKGKCYYYTMKMCGKSRMAMCAYRGCCGGRMMRATGGKTTTTYMYRKWARTS
47                RCAYCAACTTKKMYYYKATAYTTGAMGATCTGSYMKRGCCAWYRTGWCAWCCYRKWRACG
                    :  .:  . * ***. .   .. *.   . .               :   ****   .

35                ATTAKCACTTKMAWYYMKYWAKWWAYAMGYRYKRGTWKGRYGCTRYKKCTAWYRMAACYT
47                RGARKMMYYYKCRWCYMKYAGGTTWCWMACACKRRWATRRCAYWGCTKMGCWYRCRRTYY
                    : *     *  * **** .      *.   **   . * .    .*  .***    *

35                CGRYCAKKWWTAWTACYYSRYRYMARRKRRMCKGATTGTTCKRGKGKKTKTTCCSAAARK
47                YRGCMRKKWWCRWWRACYSRYRTAGRRKGACSGKRYYRYKYKRKKRKKYGYGAMSWWRGG
                        ****  *  . *****  .***   .       . ** * **    . *

35                SKTGAGKTAWRAYMYACKKSTYAKKWRATCTGTAWKKRCGWCCWAWYKMRYCKATKMSSA
47                GKYRMAKYRTGTCMYMYKGSKCTTGWRWYTCSCGAGGRYRAMTTWWYKCGCTGTGTCCGW
                  .*   .*    : **  * *. :. **    . .   *       ***     : . ..

35                TCGYKRMKCGSKYWGCKCMSAGGSMRKRMWRYYYKYGCCYYMATYTGGCTMKYRGKCCKM
47                GYRYTGCKYRGKYWTSGSASCCASMRGRMWRYCCGCSYYYYMGCCWKRYYCKYRRKGTKM
                     *.  *  .*** . . *. .*** *****    .  ***.        *** *  **

35                SYKKCSYYKRSKKRRWKKKKKKATWRYRYKTAACYRMGRKCRMAMCGSRRYYKKKKKKRK
47                SYKKMCTCKRSKKAATKKKKGGKGKWRCGTKYTTMTAMRGTYGAMMMKSGGTYKKKKKRK
                  **** .  *****   ****  *..**   * ::   *  .    *  *   ********

35                GYGAYMYAATYWYKKMYRGGKRAKCKCCAAKWKYYYKYGCGCGWRACWMYSSGAGKRRTA
47                RCKMYMYRRACAYKKMYRATKRRKYTYGGRKWKYYCGYKSRYRWRTTTMYSSTTAKRRYG
                      ***  :  ******. ** * .  . *****  * .   **:  **** :.*** .

35                KRRKTMACYYMRARRAKKRMARRRKKYKTCAGATTKKRYYRRYAAWTAYRCRKCTKKMGW
47                GGRKCCMYCYMGWGRWKKRMWRRRKKCKCTTAGYKKKRYYRACGCTGRTRMGKYYKKCRT
                    **     **   * **** ***** *  :.. .******  ..    *  *  **

35                TTYWYCGYYYTRTRKKWYMYSYMTYSGTACMTGTAGTWMYRKCYACKRWMYKKGCRGGYK
47                CCCTYTACCCWRCATKAYMYSYMKYSKWGMAKRWRKYAMYRKTYTTKRWCYKGKSGAAYK
                      * .    *  .* ******.**  .  .      **** *: *** **  . ..**

35                GGRYYRWMGWMKWYYRGYKYRYATTCTAMMKWWGKSRKMATRGTGAYMTMAKKYSGKGYT
47                RRAYYATCKTMKTCCRRTKCGYGYWMYWMCGWTAKSRGCRCARCAGTAGCCKKYSAKRTW
                     **     **   *  *  *.     *  * .***       ..    .****.*

35                GKRKRKMYKRTRYKRYMRTYGMYCCYKKCTRKMACTGACSCMTGYRGTTCTCSMTTRYAT
47                RKGGRKCTTGYGTKGCARWYSMYYYYKKTWAKCRAYACMGMMKKCAAYYTATCCAWGCCC
                   *  **  .    *   * *.**  ***   *  . .. . *.   .   :  . :   .
```

```
35    GTARMRAKKGTCKATGMCYARSGKWKGYGCYAKYGAACSRWAAKYGMCCAYARKKRMWTM
47    ACRRMRRKKAKYKTCAMAYGRSRKWKKYRYYWKYRWMMSRWGWKCAMYYGYWRKKACTAM
      .  *** **.. *: .*.*.** *** *  * **    ***. * .*  .* ***   :*

35    WMWAAMYKMGTYYAASSCYWAKTTCYTACGRKRRRRKWWCSTKYWCWTYKKAATYMYGKK
47    WMWRRCYKCKYYTGGSSYYWMKKGMYKRYRGKAGAAGWAYCGKYWYWCYKKRRGYCCCKG
      ***   **   * ..** ** *.  *.    *     *  . *** * ***   *   *

35    TACACYAAAAKRWGTKWWTYYKYYYGRRGATTTWYYAYRYRGARTYYACGTYTACYKYYM
47    GRMRAYRTGTGRWRYGTTCCTKTCCKAARGACGWCCRYRYRCMRKCCCGRYYGGMYGTCA
          .* :.: **        *       .:  *   ****  *.  .   * . *

35    KYARYYKAGTARRSYRKTYRMRYRRRKWWTCYASTCKYGTGKWKMYYWWACMCAKAGAAY
47    KYWRYCTCRCRGASYRGCCAAGCRRRKTTKMCWSATKTAGAKWKMYTTTCTASGKTRCCC
      ** ** ..     ***       ****  .   *: * . .*****   .  ..*: ..

35    TTTACYCYMMYYKGGKKYKGYTYRGRKMTYKWMGYGGGRYWKRAGTTYKMKKAAWYATCG
47    WYKRYYYYMMYYTTRKGCKKCWTGSAKMAYKWARYKAAGYWGAWSYYCTCKKGMWCRYAK
        .  * *****.  *  *     . **:***  * .. **   .   . **. *   .

35    WTKRWYYYKTAYKTYYGGTWMARRSMGYRKRTACYRKRRGTKWRKYWWTKKTRRWYWWRK
47    TYGRTYYYKACYGKCTACAAMCGGGARYRKGCCYYGKGGRCKWAKYWWGKGKGGTYWWRK
         * ****:.* .  . : *.  .  ***  . * *    ** **** * .   *****

35    RCACYSYYKKGCRRMMGACCYKKMWGTAGCTCGYYRYWRRAYGYGSAAMKWAMSGCWWKA
47    GATMYGTYKKTTRRCMKRYMYKKMWRYWRYWYAYYRYWAARCRCRSGTAKWMMSCTWWKG
       .: *. ***  ** *    *****       .*****       *.: ** **  ***.

35    YRAKGTACCCWYRAACSGRCKWYTRKCYCMGTTATGGCGARYGKWRKRSRMMYKCTATTS
47    CGWKRKRYSMACGCTYCRAYKTYCRGACMARYWWKTATKRACKKARKGCRMMYKYYGWYC
         * .  .    .: .   * * * .       . .      * ** .*****  .  .

35    KKKKKYYKRGAAMWYKTYKRMMRKMKGACAYYMCAGYGKGKAKCTKWCRTGKYTYYYYAA
47    KKKKKYYKRTGTMWYKYTKRMMAGCKRWMGYYMYRRYCKKKGKYKTAYGCAKYWYYYTCR
      ********* .:****  ****   *   .***   * * *.* ..    .** *** .

35    GTGTTCYRTCYKAYKCYTRRRCTCRCSAACAGYYTKYWYCCMYKTTRRSRYKYKACGAAK
47    SYRWYATACYCGMCGMCKRRRGYGRASTGGRRTCGGCWYTTCCKYWRACGYKYKMYKRRK
      .    .           .***   *.*:.        **    *  * . ****     *

35    MCMTTGRAGWYMMCYGACTCCCMRCYTSMYGGRMKCYMMCTGCMTGMKGGATCCATCKTY
47    CACWWKRMSWYMAYYKRYYYMAMRYYYCCCRRGMKTYMMTCASMKAAKRRCCTTGKAKCT
       .    * .***  *      .** * .     ** ***  ..*.. *  .   ...*

35    WTYKGATGKYTTTRTRRYRMWTAAATYCRCTTGMYCGAAAATRGSKYRKWRATTKRGTKY
47    WWYGSGCKTTCCYRWGATRMWCMWRYCARYYYKAYYRWRRWKRSSKYRKTAGCYTGRYKY
      * * ..  .    *    ***      .*     *      .*.*****  .  .   **

35    MKRYRRYRKGGTRMYTGACATAAKMRKCKTACKRRAGAATGYYATKYYKAGAGCKRRMRK
47    MTRYRRYRTKAAAMYKSGMTGTWGMRKAKYRTKAGGRMCCACTRYKYYKCRCTTGARMAK
      *.******. .: **... : :  ***.*   *  .  . .    ****. .    ** *

35    TWAYTAMCCAGAATKCCYKKCCTRGMACGYCMRRCRMMKRKMWRKKCAKMWARMRMYRCY
```

```
47          WARYYRMMMWSWWWGTTYGKGYYAKMWMKYYAGGMRCCKGKMTGTKGTKCAGRAACYAYT
               *  *    .      * *     *   *     *  * **  .* :*  .*   *

35          WTCCCGKTCRARATTRCAKYCRYRRTYGGCRWTWRRWKYMTSTYACKTGTGMYSGGGGGA
47          WCTATRKCAGRRMYYRYRKYTGYGRYCRRARWAARGTKCCCGACWMGKTCTMCSKRASRC
            *  .  * .  *   *  **  * *    .**: *  *    .:     .   * *  .. .

35          YWYCKCRYRTCMTAGGYAAAKYRGTKGCATCGGATMKGTRYGRTTTGWRATRCYKYRMWY
47          CWCMGYRYRYYMYCAATGGCKCGRWTRAGGMRCTCCKSWRTKAYKYRWRWCGMYTYRMTY
             *    ***  * ... ...*     . ..     :  *. *     .  **    *.*** *

35          RRWKYAGRTCRWYYYKRGYAYYWYYAMCCCGYRKAKRGKRTGATKWYAKTMYMRRYMSYS
47          RRWKYRRRYYGWYYCKRSYRYYWCYCCATTKYRKRKASKRARGCGTTWKKMYMRRYCSYS
            *****  *   *** **.* *** *. .    *** * .**: .      *.****** ***

35          AGCTRKKRTRAYRRAYAARKCTGYRYKKKTRWYRMWWYMSKYRMTCKWGCTTGYAATTMY
47          WSYWRKTGCGWCGRGYRRRKMYSCRYKKKYGWYRCATCAGKCAMYSKWCYGAACCMKWMY
             .  **.       *.*  **  . *****  ***      .*  * .**   :. . . **

35          KRMGKYACYKWWCKYYCCGGCKKMRYMRCMMTWKTTKWTRAKRKRTTRKRKSWYATACAG
47          KRMKKYRAYKWWMKTYMARRYKKMRTARMMCYWKCCGTYAMKGTRWWGKRKCAYCGWYRR
            *** ** .**** * * .    ****  * *  **         * .*    ***. *.

35          ATCCCGGKYTAYRCKKGYYYKGCWYYCTCTGYYYKAGKATTTRACMCGGTWRRYKTTAMY
47          WKMYYRRKYWRCRTKKTYCCKRYTCTMKYCRYYYKRRKWYCARGYMARRYWRRYKCCTMY
             .      **   * ** *  *        .   ****  *    :*. *.   *****  :**

35          AKAGYWYTCRCYTGRKTCMKGTAGGSTGATRTMKCYCKKRKAATCKYCRMYYWRSCKCYW
47          TKMSYACYYRYYWKRKYMCKKYMRKSAARCRYMTYCMGGRKMTCAKTTAMCTTGGAGMYW
            :* .*     * *  **    *     *:.  * *.      ** : .*    *     ..  **

35          RRWRYRYKYWYYMMMMCTGAKWMWKYRWWRRYGRRKTKGCTKWMRYYTCKCWRRKGMRYK
47          RRWRYRYTYWYCCMAMYCACKWCWKYGWWARCRRRKYKRYKTAMRYYYYKYTARKRMRYK
            *******.***   * *   ..** *** ** *  *** *    .. ****   *    ** ****

35          MYYSRYRRCCSYGTTKCKTSRKACAKKRMWMMRYCWTYKATTYCKRWCMMYKMKGTYKKR
47          MYYSRCARMTSYRKKKTKKGRKRYRKGGCTMMRYMWKYKGKYCYKRAYCATGMKRYCKKR
            *****  *  **  ..* *..**    *    **** *.**..    **       **   ***

35          KMRMRRATGKSTATTRCRYKTRYTGCCAGRYMKMSGCAACYKGTRKKYTTMGTGRRKYRC
47          GMRMRAMYKGSCMYWRTRCTCGYYRTATARTCKMSKMCGTTKKGGKKTYWMACRRRKCAY
             ****     *    * * .  *    .:.*  ***   ..   *    **    *.  ***

35          MKRCMYTCKMACGGAGMWRYGARAMGTTMCWWGCAYRMWWRKRYYMMYWYSCYYGTTKCG
47          ATRYMYGTKMMMKKMKMCTGYKMGMMSYYATTTAAGYRMWWRKRYYCAYTCCYCYKWKTTA
             .* **  **          *     *.       ...**********  *   .   *   .. .

35          GAMYAKRCAATKSWAYTAYRKRAMAATGCRATACSGKWYRACCYSYWRYAGGTGTKYRYR
47          CCMYGKRMGGKTSWMCKTCATGRMRRWRYRWWTGSRTTYRCSMCSYWRYRRRAACKYRYR
             .**.** ....**   .:   .   *       *   : * . **..   *****    :. *****

35          RRRWTKYRCTKRWWRRGGGWGKKRKKTGAMKKMRMTTAGAYMYYRRRRGKAGMTKTAKRW
47          RRRACKYRYKKRWWRRTTTWRGGAKTYRGAKGCACYWWKGCCCYRRGRKKMKCAKWMKRW
```

```
                    ***  *** .******   *    *.  . *        .   *** * *   :*  ***

35                  RGCRYTTTCTTTRWCKKGRTRTAKGGGCYRTCRACTKWKMYYRTGMYRAATTCRCGRWSA
47                  RKYRYYKYYKACGWYKKTGGRYRTAKRYYRWYRRYYKWKMYYRCTCTAGMGAAGYRRWSG
                    *  ** .  .:  * **   *  ..   **  *   *******     .  :.   ***.

35                  KARWKTAGAAKRGYTAWTKRYKGTYGGTAKYATKWYRYMYGMGCMMAGWGCTTCYSCCTT
47                  KMRWKCCRCGKRSTWRWGKRTTRYYRKYRKTGCKWYRYMYAMATCAMKARYWCTTGTTCC
                    * *** . ..**.   * ** .  *    * . *******.*.            .

35                  YMCRYRSGAGYTGCTKTCGTATTKWKAYYSMRRTYGTMYTTCMWWMRYCTAMKCARRYRK
47                  TAMAYACACAYYKSYKKTKCRYYGTKMYYGAGGYYCACYWWMATAMRYMYRMKTGAACGT
                        * ....*  . *.        * **.    * : *      ***   ** .    .

35                  MRASRKYRSSKAASKWKYKCCKTACKKYRRYKKAYGAYYYTRCKARYYRTAATKYRTWAC
47                  MRRGGKTAGSKTTGGTKYKGAKYGTKKYAATKKMCKRYYCKGTKGATTACGTCKYRGTTA
                    ** . *  .**::.  *** .* . ***   **    ** . *.      .: ***  :.

35                  MRKKYCAAARAACTKGGCYCYMGCMKYYSYAKWKKCYGAKYKKWTACAKKMSGCTYGGYC
47                  MRKKCTGRRRRRYAKKKMYMYCRYCTCYSYGKWTKMYRWKCKKTAGMWKKMGRAYCSSCY
                    ****  .  *   :*   * *    . ***.**.* *  * ** :.  ***. .  ..

35                  ATCARTTYKYRRMRTCMGGGCTGARAAKWWYYCGGCRRGGGGCCGAGGCRKKWRYKYKTK
47                  MYYRGKYCGCGGCAWYMTKRMYKRRTWKWTYYTRRTAGRRKRYYRTTATRGKARYGYKCG
                         .          *       *: ** **             : . * * ** **

35                  GAYRRCYKGAYTGGSMKGMYYCYKTKKKKKKCTKTCYTGRRTRKTTRTCKWTCTSMKMGW
47                  KWCRRYTTATYACKCCKRAYYMCKCKKKTKTMCKCTCAARGARKKYRWTTTCYWGATARA
                       **  ..:*:  . *  **  * ***.*.  *   :.* :**. *  .    . .

35                  SWKTCYWSRRYYCMYCAKMAYKKRRKCAKMSAKGTAAKYKSYWMRRWGTTAKWYYACAGR
47                  GTKWYTTCAATYTCTACKMRCKKRRGYWKMGRKKWMRGCKGYTARRWACYWKTCCRTTKR
                    . *    .   *   ..**  ****   **. *      *.*  ***.   *    :  *

35                  AKARRSKGWSKGKKMYRMWMRWKRSMGTGCCMWMYGYCRMRRYWYRMYYRRYYTRYARYW
47                  WKRGGCKATSKAKTCCGCWARWKRSMRWRYSMWMCRYMRMRRYWYAMYYRGTTWRYCGCA
                     *   .*. **.*.    * ******    .***  * ******* ****    **.

35                  RRTYKYRMRKRCARARKYRTGGKSYATMRYRTTTYKTGATCATARACTWGKTWRSYWYWA
47                  RRYYKYAARKAATRCGKCRYRTKSYWYAGYRCKGCGAKRKMMYRRGTWAKGYWRSYWYWW
                    ** ***  ** .:*. * *   ***    ** .   :  .    *.      *******

35                  YRKGGTYGCYYWYKYKTRRTKKGSTYCTYKWKKGCTTTAAYKMWATGRKYWMKKKSWYWK
47                  CGKRRAYRYYCAYKCKKRAGKKRGYYYWYKWKKRYYKKMMYKCTRKKRKYWAKKKSWTWG
                      *  :*  *  ** *.*  ** . *  *****   ..  **   . **** ***** *

35                  KGRGRWAKYGAWWSMYMTRRATRKGSRMTATMRCCRMATGKKYKCRAGGTCRGKKYKCGR
47                  KARRRWGGCRRAAGACCCRRWKRGKSRMATCMRMYRMRYRKKCGYRCTAAYGRKGCKARG
                    *.* **.      .    ** .*  ***:: **  **   **   *. .:   *  *.
```

The pairwise alignments of two accessions 35 and 48

```
35          AYYYRCYYRMYSYRYCCKACARYGWYARMKRYYAYWRKGKYMAAATMGARKCCCYYYYKY
48          TCYYRMTYGATSYRCTTTWTRRTKWYMRMKRTCRCARTAGYCRRMKAAGRKATTYYYYKY
            : ***  *   ***   .   *  ** ****     *.. *    . ..**.  ******

35          YRGMCRMGKGARYKRCKWWWKRWYCCCAAYSACYWYYYCCKMRRKKYGCTMWRKRATWRS
48          CGRCGRARKARGYKRYGTWWGRWYYYYGTYSWYCTCYYTTGMRGGKYRYWMWRKAGAWRG
                 *  *.  ***   ** ***   .:**     **   **  **   **** .:**.

35          WRMWYMMRMRKTRGYCKWWWGRYKYYTYMRATKTAYAYYYYRRYGWKKTGSRRKWMKYMW
48          WRMAYMMAMRGYRKYMGWWAKRYKCYYCCGWWKATTRTYYYRRYKWKKWRSRRTWMKYMW
            *** *** **  * *  **  *** *      *::   ****** ***  ***.******

35          ARYRYMYYYRRGRGGAGWTMWYGKTMYAKTTKSRKSYCRTRYRYRGYRRYYRRYTKMRWR
48          TRCGTACCTARKGRTGTTGMWYRKKMTWGCGKSRKSYYRKGYAYAKYRRTTRRYYKAAWR
            :*        *    .   *** *.*     ****** *. * *  ***  *** *  **

35          YYKRTYGYRTATMGCYSCGWMATYYKRMCKRYYGGTRWYAWMRYKRRWYYRWRRCMRKKS
48          YYGGATATAWRKAAAYSTCWCTKYTKRMYKRYYKRWRTYGAAGTKRRTCCRWRRYCGKKS
            **  : .    . ..**  * :.* *** ****   * *.    ***   ****   ***

35          CGCGARWYRMYRYYYRYCCRSMRWMYMYWMTCYCYMKWKMGKKTKWRGAATRGTARACKG
48          MRYKWATTGCCGTTCRTMYRSMRWCYMYTCCYTYYMKTTCAKKYTWRRGGGAAYCRRYGK
                           *   ***** ***      *** . .** .** ..  . .*

35          TCMRMRYRMTTTRMRKKTMCKYTATYRYKYAYYMMCKRRYYYRTMMCRMRYRRGYTCKWW
48          YMMAARYRMYWKRMRKKCAMKYWRYYRCTYMCYMMYKRRTYYRACMMACGTGARYWMKTW
              *  ****  .*****   **   ** .*  *** *** ***: *        *  * *

35          RRSRYKACWWCGCRRMKKGWKCRGKCCYMRYWYTRTRYRYRRRKWKRWRKMYKGTTGYYY
48          RRSRCKWYWTYRMARMKKRWGYAKKYYYMRYACYGYACACRRGKWKRWAGCYKCCAATYY
            **** *  *     **** *    *  ****         ** *****   **  :. **

35          YYWARYRYRCYKWKSRTCTCYRWSCKWYRCGMRWYYMYKGGATTCYTGCCCGAGCGCRYS
48          YYWMRTRTAYCGAKSRATCTYRTSGGTTRAKCAWYYCCKTKTGATYAAGTYKRTAATGCS
            *** * *      ***:   ** *    *.   ***  *  : : *:.      ..   *

35          AKGYTGACTMTCSMYGMMGCTGRRCTCYMGGTCCATGCYYWGTTGGMYKGYGCATKKRAR
48          TKTYYKRYYCWYCCCRMMATCARRGCYTARRCTTGKTGYTWRYWRKCYGSYRYGCTTGWA
            :* *        .   **.  .**          ..  * *      * .*  . ..

35          WCTAGCYRAATMGRRCKKACACTRAGKSKYCTRGWRTYKRCYATCTKRCGMAWYRKMWYA
48          TYGGCMYGWRYMRGRTKKTTMSWRWSKSKYAGRATAYTTGYYTCYGTRTRCGAYRKMWYR
               .  *    *  * **:  . * .****. *.    .  *:   .*   . ******

35          RYATAGCKMRCARKKTAACACCRAGGCRRWYRCGKWRKKGAWYTKCYCTKKTWGRMWMYY
48          GYTGTRSGMRYRRKKYRGTTATATTATRRWYRYKKARKKTTWYYKYTTATGATRACTAYY
             *: : . **  ***  . :.  : . *****  * *** :** *   :. :      **

35          CKYKTAYAKKKKCKRKAATYYTACRTCRKCTTRTYCKRCTSMKTTTGGGGSWKKWSAWWY
48          YGCKCCCCGKKTMKATTTCYTACTRAARKTGCGATTTAYWGAKYCCRRRKSTKKTCWAAT
```

```
           * . . **. * .:: * :. *:.**    :  .   . *       * ** .

35      CMAYKYMRRCCAKCMSKAGKTAAYSWYWRSGRRWTYRAWYYAYRAKMKSGGCTCARRRRS
48      YMRCKYMRRYYWKTAGKRKKWMRYSWYWRGRRRWYCAWTCCRCRTGCTGRKMGACGRGRS
         *  *****   *  .*  *   ******. ***         *:  ..   ..  * **

35      YARRKACTKTTSKGWKGTYRRRRAYMKKGGMRWKRSMRSRRCAKKRRMKMTKRMRRGYCK
48      YRGGKTTGGWACKATKAKTGRRRRCCKKRRARWKRSMRSRRMGKKRRMKAYKRMAGKTMK
        *   *:    :.*. *..  ***   **   ********** .****** ***      *

35      TYMWAWWKRWMWKRTYYKYYYTCWKCAYGAGGKRYRATGKWRYGRGGGTCCRGGACKAAA
48      YTCATWWKGTMWTGKCTKCTTAYWKYRYSRRKKRYRCYKTWGYRGAAACYYGAARYKWMM
            :***  **. .  *   : **  *.   ****.  .* *  ...    ..  *

35      KTATSRKKCTCACTTRWRRCGGWCGKTKCKKKGYRRKYRYAAGGMAGCYYYRCATYWYYY
48      KYTCSRKTACYRMCCRARGTAAAAATYKYGKKTCRGKYRYTGTACWCTYYCGGGATWCYY
        * : ***..      * *  .. ... *  **  * ****:. .    **   .: * **

35      TARYKRWGGRMAGMKGGCYYRTGMATRYGWYWYGGCKMKYRYYTYGCTAASATGTCMRCM
48      AGRYKATTAACRKCKAATYYRWKCRGACSWYWYACTTAKCRTCCYTYGWRSGAACTMRMA
        :.***   .     *.. ***       .****.  . * *   *     *.:.  **

35      RMTGGGGCMWCCMYSGCMYYYWWKAGMKRTTCKYYGYATCKATTAACCYRMYGAGTKMYT
48      GACACAKMMWYYMYCCTMYYYWWKRRMKRYYYKCTACMWYGGCKRTTAYAACRRRKKMYY
           . .  **  **.  *******  ***   *  .     . . : .*      .***

35      TYKKKRATCACYYWWYATTYYWCKATYCCYGGCSKRMGTRRKTATAYYGRCCCKKCMGAY
48      KTTKKGGKTGGTCWWTRYKCCTYKGCYMTYRRMGGAARYGAKYWYRTCARATTGKGMAGT
        . .** .. .   **   .    *. *  *   .       *      .*.   * *..

35      CCCTGCTCTYCACYCWYKRMWTKTCGRRTATMKGCTTMWRGGTRGYAACGTAMTGAMRRM
48      TATARTCTCTAWMYYWYKRMWATWYRRRYRWMKRAKYMTGAAYRKTRTTRYGCAACARAC
         . :      .  * ******:.   **   ** .. * ..  *   :   . :.. *

35      RYYGATRKTRRRYKYTAAATRTKCTGMCATCYCTGKATACGGKYYYAWCYWGGGWMGGGK
48      AYYKRYRKYRRRYKCCGTWWGWKAGACTTGAYYKKKTGWTTAKYYCGWYCWKRRACATRG
         **   ** *****  .:    *. .  : .* . *:    .*** .*  *     .

35      GCATRYAATWWRYGTAGGTKRRMYTKKKKMRACYYCCTTCGTTMKGARRMCAGCRSKAYC
48      ATCCRCCGCATRYKKGAKGTRRMYYKKKKCRRYYCTYYKMKCCAKKRAACYWRYRCKGTA
        . . * ..   ** ...  .**** **** *  *    .     *         *.*. .

35      AGCYWKYKYACGRGGCMGRGAKYKAGGGARKRAKKRTMYRAMKKMWRYRMKGAATCKKKY
48      GTTTAKYKYGMRRRRYARARGTYKTAAAWGKRWTKGYATRWCTGCWRYRCKRMWWYKKGC
        .    ****.  *       ..**:...  ** .*    *  .  **** *     **

35      RWWKKTCTMCACKTAYWRGMTWYCAGTRGAAATMTATGTTTRYARRTGGSWRYWAGKRRR
48      AAWKKGYCMMGYTAGCWAACAACSMRYRKRRRCMCGYCCGGRYWGRYRASWGYAGAKRGA
          ***   * . .:. * . :  .   *     * .     **  *  .** * ..**

35      YYAGACTTKACKATCGGKMKMCMAYCRYAASGGKRAKGGRMRCGRWTWARTTGGKWRTYM
48      CYGKRYYKKTYGWWMRKKATAYCWTTGTGGGRSKGMKKRGCRYRATYWGGCYRTGARWCC
         *.    .*:       * .        ... .*  *    *     *.       *
```

```
35          STAACACTTGMCCRKYACRAAYMAATWAWWYTCCATGYGTKTCRKYAGCKWWWTTYWCGY
48          GGCGMMYYYACATRKYRYGWWYMGTGAGTTYYYGTGAYRYKGAGTCRRYKTWWYYCTTAC
            . ..     . . ***     **.:  .  *   : .*  * . .    * **     .

35          YYRKTGYMYSYSGTMAGTWAGGRCYKRTAYTKATCRGKCTCKYGRYKGKRRWYMAKMWYY
48          CCGKYRTMYSYSKWMGTYAWKKRTYKRCTYWKCWYGATYYMKTTGYKRGRATCCTGCTCC
               *   ***** *.      * *** :* *.   ..   *   **  *    :

35          RAAYYRKRYCGRMYTGKMMAAWSRKTGYYYRMGRGYMRCWGKKAAYSTGTGSKWYYGTRT
48          RRWTCGKAYYKRMYCRKMCWRTSRGAKYYYAMRARTAAMWKKKRTTGYRWRSKWYYRYGG
            *     * *  ***  **    ** : *** *       * ** : .    *****

35          RMGTTCTKYKRTYYGCMYWWRKKKMYCYYKGTTYRTCRCCYYYMKYRGACTCGATCTCTK
48          RCACYMYKTKGYTCTMACWWRKKKMYMCYKRYKTAATAAAYTCMKCAAWYWTARCTATAK
            * .    * *        ********  **  .  :  ..*  **  .    .   : :*

35          ASCKGAATRAARTCGKMRACGGTCATGGYRKMYWWTTMWMRGTGKRCSYKKAWACGCRRR
48          GCYKKGGCGTRRATAKCGRTAAKYRYKKYRKMYWWGKATCGACCKGMGTKKMWGYTTRRG
            .. * ..  : *: .*    ...     ******* .    .  *  . ** *.   **

35          YRYKKMAMGKCKCYKKTWMWKYGRKYRGYGAYGYMAKWAMCGKKRWKACGTACCCWGKKG
48          CGYKKMWMKGYGMYKKGAMWKYARKYRAYTRCRCMRKWMMYSKGAWKGYRKRYSMARKKK
              **** *     ***  ****.****.*     * ** * .*  **.  .  .   **

35          TWRTCKWYRTAACTAAKGGCYYTCTWYYRYGATGGKWRMSGRMMYKCCCTATYYKRTSTY
48          YWAKYGTYRYMMAKGWKTATTCKSKTCTACRRWTKKAGMCRGMMTTYYMYGWCTKGYCCC
             * .   **   ... * .   ...          *  *.  ** .    .   *  .

35          ATRCGAYMMGYKARGRMWYKTKTYTRMMRKGACGGARWYCACTGGYKKGGTKTYMRYYST
48          TGGYKRTCMKYKRGRRMWYKKKYTWRMMAGRWTTAGRAYYRYGKRYKKAKKKWCMRCYCC
            :       * **   *****.*   ***      ..* *      ***. .*  ** *.

35          TTCAKCRKRYTTAYTKGCKYKKGTRTCGCTCRKGCATAAACGKKWCWCGGARGMYYTMAM
48          ACTCKTGKRYCYWTYKATKCGGRCRYMKAWYGKKYGCCCTGAKKWAAYRRMAKMYYYMMA
            :  .*  ***     *. *     *   .   *  . ..: .***.       *** *

35          TTGKARKTKRTWMWTTTARTKYRRRATGWATAKTKWSTTGWCRWRTAGRYTYRRMCYTRT
48          CYAKWGGAGRYTCAWGWMGGKCRGGWKATRKWKYTAGACAAMAAGYRRATCCAGMYYCAK
              .*   : *          * *   ..  . * . .: .              * *  .

35          RGKYKKTGWGCTAAARRTATRYYMRATTTTTKTTAKGCYYYKKTAYYWRCYTKGRWKKGT
48          RAKTGKKSWAGCGTTRRYRKRYCCRGACYYWKCCMKRGYTYKKATYYWRMYYKKATKTTG
            *.*  *..*.  .::**  .**  *.:    *   *  * ***::**** * *   *.

35          GTRGYGAGWCKRYCACKGCKMCGTGGGCRRMRGGAKTGYTRYRKMRACTAKCGRKCKKCK
48          TCGKTATAAMKRYYRYKRYKCTAYKARTRRMRKRRKCKYARYRKMRWAGCKGKGGYKKYK
                 .:.  ***   *  *  .  .  ****   *  *:******  . .*     ** *

35          YRTKYYKMYWTAAGCRMYRYTTTGGKKKWWWCCGRWRKWKYYGCWWWCTGKCGCACKMYW
48          TRYTYYKCCAKMMRYRMYRYAWYTRKKKAAATGSRAAKAGCYATTWWYKKKYAACTKMCT
             * .***    .    *****:    ***     .*  *   *.  ** . * ... **
```

```
35          YWWGCYAAGYRMGKRKYKYTKRGYRKYSKAYMMRGYYYWRKKGWKGGTRYKYWKKMKAAA
48          TAACTYCGAYACKKRKYKYKKRRYAGTSKTYMCGTYTYWRKKKAKAACRTKYWTTAGRRG
                 *...*   ******.** *   **:**   * *****  *.. * ***..    .

35          MATKKGGGTCGAASCYMGTCYCCGGRKARYKKYWYWWSWRGGGGGYTGCRTAAGKGKRCY
48          MRYTGRAAYYKRWGAYCAGYYGAAKRKGGCKKYWTWWSWRKRRRKCCRYRYWRAKAGGMY
            *  .  ..     ..* .  * .. **.  **** *****         *   .*.   *

35          YSYYCCKKAWRSTGCGKGGYGRYMMYGTCWYRTCGKMTGRAGYKAGYRACTKKTMKYGRY
48          YSYYMYKKGTGGCAATKTAYKRYMCYRYYAYRCYRKCWRRGRCKCRYRCGYKKYAGYRGY
            ****  **.  . .. * .* *** *    **   *   *.  *. **.  **   *  *

35          GGYKYTGWAACCTTARSRYKKTYRMRKKYASKYRAKKMYWWYYMCCAGTKYKYRTMAKGG
48          KRTGYYRWGGYYWCRGSAYKGCCAAGKKCWSGTRGKKMTTTYYASMRRYTCGTACARKCA
                *  *..      * **      **  *  *.***   ** .    .       * .

35          CMMYRRAKGKWKYKKYYRRKMGTYKRRRGYCKRGTTGCCGKKWKRATWMGKRKGKKYATK
48          AMMYRRMTRKWGCKKYTGAKMRAYGRRRRCMGASYYKMAKKKWGARYACRKGKKKKCWKG
            .***** . **  ***   ** :* ***     .    . ***       * * **  .

35          WYYWSKKTAGYTKKKMRASKYRKYCRKAATRGSTMCRCWATRYAYRYTKATYTGWYRCTG
48          WCYWSKKACAYYKKKMACGKYRKYTGGGTCRKSCCYGYAMYRCTYRCYTCAYYRWYRTCA
            * *****:..* **** ..*****   .: * *        * :**  ..:*  ***  .

35          TATARRYMGTACACRGWKKYTTMKRRKYMKYRCKWAKRMRCARMWRYACMTAGTATCWKR
48          AGKWRRTCKKCAGTAAWKKYYWMKRRKCATCGYGTMKRMRYRRCTGYWTMWWSYTCTTKR
            :.. **   ....  .****  *****  .      **** *    *  *  . :   **

35          YRMAWKWRMTTMYGYTTKYGYYYKRYAARRKSWAGYKAGTAYYYRKRMGCRCSGKWYTGT
48          YRMRAKWRMAAAYKYWKTCRCTTKRYGGRRTSWRRCKRRWGCYYATGCRYRGSRTTYCAC
            ***  ****:: * * ..     ***..**.**   *   . ** .    * * . * .

35          RATCAWAYYWRATGRYYTSSTTTCAAKYTKYRMTGYYCRCTGGGAWKKMRKKWTGARKMR
48          RCWSGAGCYWRRYARYCWSSYKYTGRKYKKTAMCTYYMRYKTTTCWGGAAKKWYRGRGCA
            *. .. . ***  .**  ** .  . **.*  *  ** * .   .*    ***  .*

35          WCGARTGRGTACTKKTTTMWGTKMTKKGCAKRWTWTTTCTCTTMGWCAWMYTGTAAKCTG
48          TYKWGYRRRYGMAKKCACCTRYKCAKKRYMGAAKWYKYYKYACCRWYRACYGKYRGTMYK
                   *  . :** :     * :**      .* .  . :   *    *    ..

35          TGAYCYRTCGKGAYACTCKARKMYYRYMWYRGSCWMRWKRCWKRTRYKMCKKRWKKSMGC
48          YRGYYYRWYATKRYGTCAGCAKMYYRYCATARSMWMRWKRYWKAGGTTMYKKRWKKSMAT
              .* **  ..  *.  . . ******     * ****** **    .* ********.

35          MKYAKCYYYSYMCRYRSGTGCTTAKTGGCAACCRYYATRWWKMRYWKTACRCTRMRKMCA
48          CKYWKTTYTGTAMAYACAYKSYWMTKRATGCTTATTRYGATKMGCAKAGYRYYAARKMAT
             ** *  * .    * ..  .   .. . ..          **   *:. *    ***.:

35          RRACACKRATTAGMRYYRGATKTGATARWKTWRSYWYMKTGTWMRAGGKTCAAGTWGYWC
48          GGCGCYKRMYKWTAGCYGAGCGAKMYRRWGYWRSYWYMKYACACGTRKKYYMWRCWACAY
              . . **  .     * ..  :    **  ******** .    :  *      *.

35          YKYKAATTRCMCMSGSCRTTKWKTRKKYKAGGGKGGMAGMRKYYKGRGYKWRAKYGWWST
```

```
48      YKCGMRKKRMCYASKGYRWWKWKGATKYKRTRAKRKMWKMRKTCKAARYGTRGGCRAAGC
        **    ..*    * . *  ***  .***   .*  *  ***  *.  *  *.     .

35      GCARRMATCGKGMRCRKTYRRMRATGGTKARRTTYCRCRKGGTKYRGKCGRMYYCGGGAK
48      KYMRRCWKAAGKCRYGKCCRRMRRYRRYKRRGCCCYRGAKTAAKCGRKARGMCTYTTACK
           **  ...   *  *  ****     * *     *  * .:*   *.  *     ..*

35      AGAWCMTTRWRYYYTTAMGRAAKWYCRKYTGRAACCTMRYGGKACGGSTGTCMYKYKRKA
48      RRGWYAWKRTRYYYCCTMRRTCKACYRKYWKRRRYMYCRYKRKGTAKSARCYMYTCGAKM
          .*   .* ****  :* *:.*   ***  *      **  *. . *:   **.   *

35      KYCRMYYWRSCKYWRYYMMCTGACYACWMRWWRRWCCSYRWGGCRRYRKTKKWMRYYWRR
48      KTTAMCTWRSSGYWRYCCAYCACMYGYWCGWWARWMAGTGTARTARYRKYTKAMRYYTAR
        *   *  ***. ****     .. *. *  ** ** ..   .   **** .* ****  *

35      KGYMMKTKWYYGYTYYSKKTCACCGSKCMGTTCKKKAACAKRMWCKYATKGTTKRTTMYK
48      KRCAAGCKWYYRYGYYSKKYTTTYRGKTCRKKTKKGRRYGKGCTMKYGKKAAWGGCCATG
        *      **** * *****  :   .*   .. **    .*    **..*.:

35      GYWKKMMRMRKMTCGGGTKKKAYWYSKTYYKGTYKTKCMMWWWWMRYCTWKWCGMGKRRK
48      KCWGKAMRMRKAWMRRAGKTKMTAYGKYTYKCAYKAKMAATTAAARYAYWKATACSKARK
          * * *****     . *.*   *.*  ** :**:*        **. **  . .* **

35      RYRKAMSYYTCRCGSWMRRYKKAASAWKKKYRRYYKRTKYGACYYGYKCYYARYYRAATK
48      ACGKCCGYYATRYAGWMRRTKKTTGRTKKKYAATYGRAKCKRSYYKCKMCYRATTAGTCK
           *. .**: * ..**** **::.  ****   * *:*   .**  *  *     .: *

35      YRRTCMRKKCMYWCASYRTAACRAATGYCCRKGMYYMGCMGKRYSYKWKYGRKYKCTCMY
48      YRGCAMRKKMACWTGSCGCCGTRRRAATTTAKRMTCCRYCRKGTGCKWKYRRKCKYWMMY
        **  .****   * .*   .. *  :.    * *       *  . **** ** *   **

35      WCAKMYYSYCRCGGYTAYKYMMACCACATTACKGTCRYKGWWYYAKCGCGWYAKYYRWYK
48      WMWGMYYGCTRYKSYYRCGCMCCYMWYTYWGTKRKYRCTRWTYYCKTRTSWTWKTYRWYK
        *   ***.  *  .*     * .    :  . * . * . * **.*   .*  * *****

35      GTYMYRRRWRRGTKYMARYWMRRGKRKRMCGCYTTRWKYMYRACCKRRYYYTGCYWKAGS
48      ACYMYAGRWGAACGYMRRYWMRRKTRKRMTATCCGRWKCATARYYKRRYYYYRTTWKTAC
        . ***  **  .  ** ****** .**** .    ***       ******    **:..

35      KGRCRYACGKRARCMCRGRRRRKCKSAGWRGRSKCGWRSYGCTRKATCCTGKKRKRYMKK
48      TAATGCMYKKGMGACTGRRRRRKYKSCKWRKRSKMRWRSYRMCGKTCTTGCKKATACCKG
        ..       *   .    ***** **. ** ***  ****    *:     ** .   *

35      KKCWCCTCAGMGRYYKSYCTKYYYCTYCGTYRRYRTCYKTAYCRKYGGCCTTGATMGTGC
48      KKYAYTGAGAMAGYYKGCMWGCCCACYMRYYRRYRYYTKKMCTRTCARGTCWCTCCSWRT
        **     ...*. ***.       . *   *****   *.   *. .      :  .

35      TACYCCAWKYTRYCKKGYRMAACCYRRWKGARKAARTTCKWYRMMYGKMYWRRMGCAKYY
48      CCTTYYMAKYYRYTKTKYAAWGMYYRRWKCRRGRWGYYYKWYRMMCKGCYAGACRYRKYY
         .      ** ** *. *   .  *****  *       ******    *       ***

35      WYYAKKRGCRYKWGAGTKARRRMMMGKMYRGTRMSYSAGCTRKTKRYRACAYKWWRYYKR
48      WCYRKKASYRYKWAGACKWRRGMCCRGMYGRYACSYSWSYWRKKTGYGWYRCTAARCYKG
```

```
      * * ** . ****... * ** *    **     *** .  **.. *     .  * **

35    MWWYMSWGTATTKRMYAGYKCYKYKCYKCYCGCKKMRRAYRGYYYRYCYAMRCMYMTWTT
48    CATCAGWCGMAGTRMYRKTKMTKCKMYKMCYRYKKMRRGTATCCTATTYWMGMACMYWCC
           .*   : .***   *  * * **     *****.         * *    * *

35    KTAWKRGRTTRACTGKRKATKKYGKKCGKTAGTRKMRATGAKMRSRGKKYYAACYRMWKA
48    KCGTKARRWWGTTATKRKCGKKYRKKYRKWRKGRKMRTCARKMRSAAKKYYWMMYAAWKG
      * . *  *   : : ***. *** **  *    ****: . **** .****   *  **.

35    RGAGRYKKTRMWRWTAGARAATAMYKKTCMYASSCAACGGKYWRRTWTKAMKKRRGATRR
48    RKWRAYKKCACTAWATTCGGGCGCYKKWYCYRSSYMRYRRKCTRRAWYKGMKKAGRWKAA
      *    ***     *:: . .. . ***   * **      *  **:* *.***    .

35    CTKKWCYWYTTTWMTAAKKTYYYMYAGTGAGCRTKKAAKWYGTYYATTARTGAAWAYRKC
48    YKGKTYYWYCCAWMWRRKKGYCYMYWRWAGTTAWKGRCGWYRYYYRYWGGWRRGWRYRKA
       . *  ***  :**   ** * ***   ..    *  . **  **   .    .* ***.

35    KYCCRKYMRWKKCCRYWKCRWRATRYAATTSTCAGCYMTRGTKYCKTGGKAYKYRRKRKC
48    GYYTGKYMGTGGTAGCWKYRWGMWACGGACCCMTRTTMYRRYKYTKAAAKMTKYRRKGTY
       *   ***     .  ** **     ..: .  :   * *  ** *:..*  ***** .

35    YGCMRSATKRYKGYRCYYMYTRRRGWRGAATKATCYKRKRKCKYKAGARYMYYTKGRKYR
48    TTTCRGWKKGTKRYRYYYMCYRAGSTRTGGCKWGYYKGKRKYGTKGATRYATYKKKAKCR
          *. .*  * ** ***  *  . * .. *   ** ***   *..:**  *.*  * *

35    KYYWYKRCTWYYGKWYTCRKATGTRCGGGMYCKYMGTYRCAYYTCYCGCYGTKRKYTKCK
48    KYCWYKGSKATTRKACCYGKCCTKGYKRKCTYKCMRWCGMRTTYMTYATYKCKRKTCKSG
      ** *** ..    *     *.  .        * *            . *  ***  *.

35    YMYYSYYYAAGGSMRKWGRYYKYWGCTYYYRTRRYYRYMYGCCKSATMRKKKRSKSKWGC
48    YAYYSYYYCGCASMRKWKRYCGYTRTYCYYGYGRCYRYMYRGYKSCCMRKKKRSKGGAAT
      * ******.. .***** **  *     **   * *****   **. ********.  .

35    KKYKKKWKCARYRYTRRAGCCCYYRYRRMAKTCAKTKGTKKRYTMYCMMGAMMCMCAAYW
48    KKYKGGWGTTRCGYYRRGAGAMTYACGGAMKYMWGAKRWKKRCKMYYMMKMMMMCYGRCA
      ****  *  :*  * **.. .  *      *    :*  *** .** **  **   .

35    YKKYRTGGKAKCGKAKKACCATYAGCGWTRGMGRSRGKTRKTMCMACMRKKTGRYRCATA
48    YKGTGGAAKRTYAKRKKRSMGYCCKSRAYGRCKGSGRKYRTCCYMWYMRKKGAGCRYWYT
      **    ..* . .* ** . .  . .        *  * *.   *  **** .  *   :

35    ATTTCKRCKYAAYWATAGYRGYMRTKSTGGGACGYTCYKCGTKGTCWATAATYWRRGRYK
48    TYGCTKAYGCRMTWRCRSCRRCCRKKCKRAARTRCGTCTMRAGAAMWGCGGKYWRGRRCK
      :    *       *   . *   *.*.. ..       .  : .: *. ...***  * *

35    YAWGMKKKCCYMATWGTTYKRCYMYRKYRYKGKKGGRKTRGTCRKYAARKSATGACTARW
48    YRWAMKKKTMTAGCWKCCYKGTCCTRKYRYKRTKAKGKKGAGAAGYWRGGGWCRCMYRRW
      * *.****    . *   **     ****** .*.  *. . .  *    .   .   **

35    YGWKKTGGRCSGTYACARYKYTRRGMWYYKWKWGTSCSCCKRYKMYW
48    CRWKGCAARMSRKCCATAYKCCARRMWCCGAKWCYGYSMYGGYTCCT
        **  ..* * . ..: **   * **    **  . *    *.
```

The pairwise alignments of two accessions 35 and 49

```
35              RRGWWMMWWMYYYYMKKRKRKSRKSYYRYYGYWRRTRTRYSYRMAKCYWYYRYWAGGGMA
49              RAKTWCAWAMCYYYCGKRGRKSRKSYYRYYRYARGKGKAYSTGCTKMTACCRYWWRKRMW
                *   *  * * ***  ** *********** * * . . **   :*     ***    *

35              RGYRYMRYYKMSCASYTTTTSMYRYRCCCARCRYSYYYMRGWSGCTRYRKRSTRWKMYYY
49              ARYAYMRYYKMSYRCTYWKWGCCGTRTYYRRMGYSCCTMAKWSRAGRTRKRSYGWKMYYY
                  * ********  .   . .    *    *  **   *  ** . * ****  ******

35              YYYYYRMATCMRKTRRACKRRAAKRRKMATMMAKGGYRATWKTYKWWAYSWWKAACRCAM
49              YYYCTGCRYMARKWRRWYGRGTWKRRKMTACCMKRRTRRGWKYYKTTTYGWWKCMGAYGM
                ***        ** **   * : *****::   *   *  ** **  :*.***.    .*

35              GYKYKYYMRYCYKYKCTRAGTCYYCKKYKWRWYTGGYACYWYYYKYRTKGWATRARTYCY
49              RYKTKYYMRYYYGYKYCRTRWYCYYKKTGARWYYTATGMCTCYYKYGYGRAGGRMAWYSY
                 ** ****** * **  *:    * **   ***  . .    ****     . *   *.*

35              WWRGCCMMMYRSYRTTKYYYRGTMYSWYKCYWWYMMKRAWWYWRWKCGGYMTCMMYMRRY
49              WWRAYTMMMCAGTRGKGCTYGRCMTGWCTYTWWYMMKRGWWTWRWTTKRCCCTCCYMRRY
                ***.  ***  . * .   *   * .* .  *******.** ***.         *****

35              RGKWYKSRRYWKYYRRKWMKYGAYRRRKGYGCKRTKRWYRGMMRGAGKWAYYTYMRGGTR
49              GRGTTTSRRYTKYYRRTTMKYRRTAGAGKYSYGRYKRAYRKMCGRWKKWGCCGYMRKRWR
                     .**** *****. ***        *.  * ** ** *     **.   ***   *

35              KYYRRTWGTTRTGTKWRWYAWRGKRYTYMRTCCCCAYRRMWYYRWRMTACRYGYRYYKMR
49              KYTRRATRKAGWRYKWRWYRWRAKRYYCCGGYTTTRTRRMTCCRWRMYRYGTTCRCCGCR
                ** **:  .:    ***** **.***           ***   ****       *    *

35              WRMRYYKCCATKGAMMRKAYRWRYYYMKMYKWYMMRGGKYYCWRKRMGTYGTRWRWKYRW
49              WRMATTGYYRAKRRAMAKMYRWRYYCCTMCKAYMMASRKYYYWAKRMRYTAARWRWKCAA
                ***       :*   * * ******  .* * *** . *** * ***   .:*****

35              GRAATCTCTAYMAGACGCRCCAACCYACCCCYCMYWYGWYRMMCCCWGYRAKRWKARAAC
49              RGGGGYCYGWYMRRWMKMAYTGRYTCGTYTACSMCTCKTTACCYYYWKYRGKGAKMRRRY
                  ..      **         .    .   . .*            * **.*  * *

35              SYWAMWGTGTWMRYRMYYKCGYYTCACTYCRKAKATTAMYRRRKRAGKKRMMMGRAKTGC
49              SYWGMWRYACTARTRCCTTYKYYKMRYYYYRGTGGYCGATGGATRWSGGGCCMKGWKYRY
                ***.**  .   * *   .  **.    * * : .  .     .* .     *   *

35              TAGTGGRGTCGAAAARRSTAMGTAAACSMKYWMGYRTGGKRRMWAWAACRYKKKTKMKGA
49              WMTWKRRKWTTMRRRARGGGCAWTCRMSMKCAMAYRCCAKAACTTWGTTRCTKKYKAKRR
                      *         *. . .:.  ***  *.**  .*    :*.: * .** * *

35              YRGKYAKCYWGAYMWYRSYRTTAKMRTYRMRYRKKGYKRRKKWWYASKYAWKKMYWWMCK
49              YRRTCTKTYWKRYMWYASYRYWRGMRCYRMRCRKKKCKRRKKTTCWSKTGWGKMTTTASK
                ** . :* **  **** ***    ** **** ***  *****    ** .* **    .*

35              KAMKYKYRTMAYMMYRKGRMYMRKKKWKCYKKYYRTRKMAGYKYRCWKRGTTMTKKMKWG
49              KTATCKTACARYMMYRTRGCTCRKKKTGYCKKCTGAAKMWRYKTGGMWGASYYCGKKAKWR
```

```
        *: . *     *****.     ****    **   : **  **   *  .    ** **

35      KCTTATRWMMMGKGKYYYKTAYKRTYRMRWCCATTMKWRRTAARYKCKYRTGCKCTAWKK
49      GTWYRYRAMCMRKKKYYYKACYKRKYRCATTTCYYMKWRGCGCGCKYKYGGAYKYCGAKK
              * * * * *****:.***.**     .  ****  ..  * **  . *  . **

35      ACACKRRYTTGAYAKGTACKRTCKTGCTCGTTWTRTKCGAAYCTACGKYYYYGWCWGCMA
49      WYMTKGGYCCRWTGKCKGYKGGTKYRTYARYYWCRAKYTGCTYATGAKCYYYKAAARYCC
            *  *     .* .. *   *    .   * *:*  ..  :: .* ***  .    .

35      MTTKWMWGTRTGAYAKYTATGRYAWTATAGWRTKWSTRWGCRTATATRRYYKWRATGRGY
49      ACCGTCAKWRWTGTGTYWMYKGCGTKWKMAARYTAGAAAAYGYRACCRRYYKTGRWKARC
                 *  . ..*      . . . . * . .:  .    :. *****

35      CCCGYTTKRYKKGKAAACGGCTRKGAACGCMWMGGCTTGCATYYYYAKGTCYAGKGACTK
49      YAYRCGKKATGKKKGTTMRRYYRKCTWYAYCTCRKYYYATGWCYCYMKAKGYRKKAGTCT
         .    .*   * *.::     ** :  .         . .  * * *.. *  *..  .

35      YYWWTTCKTRWRYKKCAAGATCAKKKMCGTYYTATGGGAKKTRYGCCGAGGCRRKWRYKK
49      YYWTYCAGYRWGTKKTGGAMATGTKKCTACCCKCCAKRRTGARYATTATTATRRKARYKK
        ***   .  **  ** ... : ..**  .   .. .   . :**.  .: . *** ****

35      RRCYKAYMRWGRMWMYYCKTTYYKAKKCGCTCKCTGGARCAYWYSKWRKRKRAAATKWYA
49      RRYTTWYMRTRACTAYYMKCCCYKCKKARYGTTMCRRWRSMYATCGARKRKAWMMYKATT
        **  . ***      ** *   **.**.    .     *. *  .  ****     *  :

35      TAGCKCSMKGSWKRCKKCARCCYWWSARRYMTKYCMAYKRRRKCCTCGAKGTWMSTGKAW
49      CGATTYGAKRSTKRGKKTRAYYTTTCWAATMKGTTMRCGRRRKYTCAAWKRGAMGYKGRA
         .. . . * * ** **        .    *.   *   ****   .. *   *.

35      WGTYSWWRRYSRTCYRRTYRRAWYYARGGCTCYSYCACGARRSRWGAKMWMKKGGTRWRS
49      TKWYCWTAGTCGGGTRRYCARWTCCRRRKMGATSYAMMARGGCGTATKCTCKKRRKRWRS
           *.*    .    **   *     *    . **.  .   .  .:*  **   .****

35      MGKAGGMYKYRMRRRKGCKGCGKGCMWRWKRCCAYTATRYRTCAMKYYRKMYGATMKGTC
49      MKKGTRMCGYRMGAGKRMKAMKKRYCAGWKGMMWYCCCRYRYYMAKTCGTATCTCCGSWT
        * *.  *  ***   *  *.  *     **    * . ***    *   .   :   .

35      TACYCAGRKTWKRYGTTYTCKCCCAYKWYRRWKGAGRCKTCTATTCKYYKKYWRRMCYYW
49      CCTTYMKAKYWKGYRAYCWATAYSRTKWYRRWKSRRRYKYTWRYYYKYCKGTAGACTYYW
         .      * ** * :   ... .  *******.  * *       ** *       ***

35      YYTKCRKRKAMYMRRRYRMSYGCTSRKRTRACYAAMKKCYRYKARMMMWWTGGYMSKTCT
49      CYGKSAKRKWMYMRRGYACSYSYWSRTGCGWTYRGMKKMCRYKWGMMCATATACAGKWYY
         * *. *** ***** *  **.  **.     * .***  ***  **   : .  .*

35      YRMTWTTGYAAWYKRMAGKYKYKYAKYRKYKKTYYGCAKKMRGYYRYCYAMRCTTWTTWG
49      CACYTGAACCMWYKRMRKKYGCKYRKTRKCKKCYCRYGKKMRKCTATTTWARMYKWCCTR
              :. . *****  **  ** * ** ** *   .****         *  .*

35      ARYKRTTTSRGYAAAKKCTGGTAYMTGYKKKGGRYRYKYGRSMRGKAYTAYYKACSTTRR
49      MACKRWWCSRACTCWKKTGRRWRCCCCTKKKTKRYRYKYCRSMRRGWYGGYYKMMSAARR
           **   **. :. **           ***  ****** ****   * .***  *::**
```

```
35               MWAGGCRYKKCTRMWKMKWMTRKRKMGKKKYWAKKAMKRGYRRCTKRCWMMWKCYYTTWM
49               AWGTCTAYKKYCACTTMKWCKRKRKCSKKKYWGKKGMKRKYAAYKGAYACCTKYYCCAWM
                  *.    ***      .*** .**** .*****.**.*** *    .       * *  :**

35               CKAKRKKTGAACAKKWWYAATKKYYYATTYGTKRWYAYRKYYRKYSRAMGKRYYYAYYKR
49               MKRKRTKGKMWSRKGWWYTGCTGTCCRYWTAATRWCRYRKYYRKCSRWCATGCTCMTTKA
                  * **.*     . * ***:. .         .:.**  ******* **   ..       *

35               TCTTYGARYRGTTCCTGSYYTYGKRRGGGTKYRRKKGYAAAKMTAATRTAACARRCGGWG
49               AYAWYKRRYRAGKTTYKGTYYYRKRGAAACGTAGKTAYWMMKCGRTCRYGGYRRGTAAAA
                 : : *  ***. .    . * * ** ...     *..*   *    : * ..  *  .. .

35               TCKSYYAAGGKGGCCAARKCRKGAGTGRMKRGRKYKGAGCGYMGCYYRYKKRRAAGKAGM
49               ATKSTTTGTAGKCTAGMGGAGKRGAYRGMTRTRKYKRRKYAYCATTCGYKKRRWMKGRAM
                 : **  :. .    ..   . * ..   *.* ****    .* .    *****     .*

35               CMYSRGCMYYYWWKYGCAMRKYYKTKMTACCAKYYKRKMYKTYRCTYTTKKCYGCYYYYK
49               YMYCACTMYYYWWKCCYMMRKYYKKKMYTTAGKYYKAKACKATGAAYYKTKATTTYYYYG
                  **.   *******    ******.** : ..**** *  *:  .:* ..*.   ****

35               YKAGTSRTCTRCTGGTRRYMYTYACGAAGCCGYACATTTYTGMGAGGACTYTYKCYYGYG
49               CGGAWSRATCRGAAACRGCCCYYMYKRRTTAAYGYGCYCYWKMAGAKRYYYYYKAYTRCR
                   .. **:  * :.. *     *        ..*. .    *  *...    * **.*

35               SMAGATYMMYMAGGCKKCCAGTYWKGRWRWCCTCTRKCWTRATTMAGMGAACTRAAAATR
49               CMTTGAYMMTARRRSKKTTGKCTWGSRAATAYKYWAKMACGRYCMRKAKCMSWRWRRWKG
                 .*: .:***     .**  .    * .*   . .    *       *     . . *    .

35               KWRTKRMKATKGTYKYTTYRATAKTCARTGGTTATCAGCCRGCRWKYKCKYRWYKYYMGG
49               TTAYTGCTWKKACYKCYYCGTGTGCYRAKRRYYRCTTAATACYRWKYKMKYRWYKYYMRR
                 .   .  . .*. **     : :     .       :..    ***** *********

35               GRYAGCCRRYGRWRRKYMARGCGGMKKRYWCGKAGCAYTTRWKYMYRKKWWRRRRYYYTG
49               RGYGAATAGYKRWGAKYMRRSMRRMTKGCWYKTMRARYYKRWKCATAKGTAARRRYYYYR
                   *...   * **  *** *.    *.*  *   .  . * .***     *    ******

35               YWWKGAKKGRCRYCYGKRRRGRCTTAKYSYKGRSKKCGTRSRKYTCRTWRGMRGGYKAYC
49               TWWGATKKAATGCYYKKGGGRRGGKGKYSYKKRSKKMRWRSGKTWMAYTACMRATTGTTG
                  ** .:**.     * *    *   ..***** ****   ** *         **.   :

35               TTGARKKRYMKKGCWKAWMYGRYKSYMYYKATWWYTRMYCTRYKACAKKMRCGKATGCCKT
49               CGTCATACCKKKYAKGAMYAGYKGCMCYKGYATCKRMYYKACGRYMKKCRYTKGCATTKY
                    . .   **   *. **. **. * **.    .*** .       ** *  *. .  *

35               TCRKAARRMRSYKKYWKGACCMGYKYCYAWAGCKAAKMYWKAWKCGCKGKMMYYRMKKKY
49               KMRKRTAACGCYKKTATTCTGATYKYTCMWCRYGGGKATTTGAKYSYKKKMATYAMKKTY
                 . ** :     .***   . .     ***    *.    ..*    .. * . * **  * ***.*

35               MYRCTKAATYRKRGRYTKRYKGAWGCAGWGYATYKYAKYGKYWTAKKKYCGYSRGAYYMM
49               CTGAWGMWWCRKRRRCCKRCTKMARYRRARYCAYKCRKYTKYWWRKKKCTRCCGKRYTAC
                    .      *** *  ** .         *.:**  ** ***  ***      .   *
```

```
35      YYAYCYTGSYKTTCTCKAARYKKRKKTRCTATCCAARKSRTTGATRTARWRYCMTRRRWR
49      YCGTYYCKSYKYKYGMKWWRYKKGKKARTWMGAMRMGGGRACACGRYRRWRYYCYRRRWR
        * .  *  *** .   *  **** **:*    .     .*: .. *  ****   *****

35      KKSTAACWRKAKKGRRCMKYWCWAWKTTSRTCYAGGRMGATGCGSKMYKAWRYYGWTYSM
49      KGSKTGMWRKWKKRAGYMGYAMTWWKKYGRYAYRKRRMRTAKYRGKMTKMARYYAAKYGA
        * *.:. *** **    * *    **. .* .*   ** ::   .** *  ***. .*.

35      RRTKKGTRMKCMMWWMRYRKGCARAAKYKKCGCRYGRAATAMASRYKAMYSAASKWKYKY
49      GGYKTCAGMGMAATAMRYGKAYRAWMGCKKATTACKGGGCMMRGGYKRMTSTTGKTKTGY
           *. : *      *** *.       **.      ..  * . ** * *::.* *  *

35      RRYYAKCYGACYKKKYARYYRTWRKYRMRKKCYWCAKSYRRAATRGRGYCYMMKYTRKYS
49      AATYRKYCKRSCKKKYRATTACWRKYRMRKKMCTTGKSCRRRRARRRKYMYCCKYCGKYS
           * *    . ****      *********    .** **  :* * * *  **  ***

35      YGKKKYRKYKWYTRACCAMKKKGGCTCCTGTGKARYYKYKKMRCMGACYRGGTCTGARGW
49      YRKTKTRKCKTCAAGYMWMKKTRSYGYSYSAAKRGYCGCKGCRSAKTYYAKRYMYKRRRW
        * *.* ** *  : .   ***. .   . .:.*  *   *  *.  : *        * *

35      WRCKYWYCATGCTSCYTCYKCAASRACMKRCGYRKTATWAYYGYWTATAWRMMCCCYMAC
49      TRTKYWYTCCAYWCMCGTTKAGRGAMYCTAMATAKYWWAWTYKYWYRGCTRCMMTTYMWM
         * **** . .  .     *.. .    .  .  *      * **   . * *   **

35      GYCRSAKGKCAWTTGTKGKMKTRGAAKYCAYGCGYMGCGYRRMYRCAGKRTYRTRKGTAT
49      KYYGGRKKKMRACCACKAKMKKGAGGTTGTCKSTTARYRCRACYAYRSKRYCGYRKAYRC
         *  . * *     . *.***. ....  :  .       *  *   .**    **.

35      GYRTKCRKTTGGGWKGGGCAGGKCGGCACGGYGGYWWGTTKRRWYWMRYKYRRCASYYKR
49      ACGCKARKKATAAAKATAYRKRKYRRTWTTTYKKYWWAGGKGGTYWCRYKYRGATGTYKR
        .   *.**.: .. *. .    *        *  ***.  *   ** ***** .:. ***

35      TGCGRRTKAACGYMCYKKWTATYGTAYRMTCYYWRRYWYGSKTMSGCRWKCYRASKYAKC
49      YTTARRCKWRYKCCMYKKWCGCCRYWYGMWYYCWAACWCRSKACSCTAWKYCGWCKCMKS
           .** *       **** .     * *  * *   *  **: *   **    .*  *.

35      CWRYKAGTWRTTCWKGWRKTAAKCACKAGGGCYYTCTWYGRYKGKWRKRGTATCARYMMK
49      MARYKTKYWAWKTTKATRKYMMGTWTGGCTATTCKSKTTKACKTKAGKGAGGCATRYMMT
          ***:  *  .  *. **        .  .   ...     * *  * . . .:****.

35      RCCAYKKRCKGAAKYCRRKYKKGACTYKRMRKMKGRAACMYMKCGGAAAYWCMCAGYWKW
49      AYYGCKKRMGKRGKYTGRKYKGACMGTKRMAGCKAGGGMCTCGTTAGGRYAYCTGRYWKA
           . ***    .**  **** ..   ***   *. ..       ... *    . ***

35      ATYAMCCKMKTAGMMYYSKRAAGGWCYYKYGAYMWTRMGYCGGYRCTTGTKCSKKWRSKS
49      WKYCAYSGAKYWRCACYSKRCGCAAGCCGCRMYMTCRCRTYRKYRSYYKGKGSKKTRSKG
         .*.  .  *      ****.. .        **  *      **.    * *** ***.

35      RRWKYKKKWYRYKTCTGAGRRGMGAGAYKKGAAGYKTCGYRKKRCTYRACMMGAMYGKAM
49      AATKYKGGWYGYKYMYRWRARRMSRSMCKKAGGACGKMRYRKKRYKCRCYMMKMCTKGRM
           ***  ** **       * *. .  **....  .  ***** . *. **       *

35      YKMYRKTARYKCWCAKWTCAAYGAYGCGRMWWRWRCRTMMRRRKMTTTRAYYKKTCAGAT
```

```
49              YKMYRKWRATKYAGGKWKTGRCRTYKSRRCTWRWGMRCCMGGRKMCCYRMYCKKCTTAGY
                ******    *   .**. .   :* . *  ***  *  *  ***   * * **  :..

35              KRYAYCCCARCYRWKTCTTKTCTGTGRRGGGGWKKMRKCTGARGCCGKMRYMYTGGRRRG
49              KACTCTTTGGYCRTKCGCCKWTWKYTRRATTTWGGAAKMCAGRATTTGCACCCGSSGRRK
                *  :    .   * *    *      **.   *    *  ..*.          .. **

35              KTYAGGYCMTTAMRYTTTCTTTCMRWCAGAKKTKCGTAKGGYYCYRTCKATTWKYKMYYR
49              KWYMKRYMCAWMMRYYKYYKACYCGWYRRRKKYKSKYRTARYYYYRWYKMYCWKCKMYYR
                * *   *  :  *** .  .:    *    ** *.   .. ** **  *   ** *****

35              CRRTWSMKRWKRGGWATKRYKYTAKMYTRYTYYKRAMGCMTCTAAGWAWTCKAGGYYSCC
49              AGAWWSMKGWKASKWMGKRTTYYRKCTCACKTTKRGMATCWYWWRRTTACTKRKKYTGTT
                .   **** ** . *  ** .*  *     .  **.*.         :   *   * .

35              TTCTYMCTARRSGKYKTKGCAACWRKRKCCTGTMRYWRYKYKRRRKAKARKRTAGGYRGY
49              CCSYTAMWRAACKKYKKKATGCTWRKGGYYAKCMGCARCGTKAGGKGKCGKRYWRATATY
                  .        . ***.*. .. ***    :  *   *   *   *.*. **   .   *

35              TYKMATKTKATARWKWGGRSYWYMKYYWMKTRTGTCCSRTKYKTYKAWKKCTKYWCTGYR
49              YYKAGCGAKMYRRWGTAARSYWYCKYCAAKKRWAWYSSAGKYGKTTGWKKYYKCTMKKYA
                 ** .  :*   **  ..***** **   *.* .  .*  ** . ..***  *   . *

35              KGGGAGAGCYYMMKASWYRWKYRYGRGRCKGCKYGWWSKTGKKCTCATTKGRRTCTYATK
49              TKAATRRAYCTACKMSWTRTKCRTARRGYGAAGCRAAGKCKKKTATWKAGTARCYYYRYK
                . ..:  .     * ** * * * .*    ..     .*  ** :  .:   *   *  *

35              GCKWYRKCRKTCGRKGTGKTCCTTATTRCWCAYTCYTGCCATWAKTYAYRKCCAYYRRRY
49              SSKWCAKYGKAARGKKYRKCTTYCMYWRYTATTKGCYRTATCWMKCYRTAKTATYYRRRT
                ..**  *  *:.  *   *        *  .: .     .: * * *   * .:*****

35              MAMSYRTKKYYCGTTKTYYYCMKGYRCATCCCCAMWRARMAATCWYWCAYRMWWRKKCCR
49              CRMSTGWGKTYTACAKKCYYYATATATMYYSMYRCTGMGACCCTWYTMRYRMWWRKKYYR
                  **    * * . :*. **  ..      .         ..  **   ********  *

35              CGMAYKYKKCATACYRYKRKRRSWYAGTTTYYMRMSGKWAWRYRGYTGTTSSKMYTTMKW
49              YCATCKYKTTGCGMTRTTGKRRSWCGAAYKYYMAMSRTTCWRCGAYGACCSSGCCCYCKT
                   : ***. . .  * . ***** ..: .*** ** . .**  .* .  **      *

35              KYKMRYCKMCWCACRYWYMRWRTRYSAAAYRAKRGTKTTYTGAGTTCCRKAKTYGTCKAY
49              GTTCGYMKAYATGMGCWYCGWGWACCTGMTARGRRYKCCTYTGAWGTARKMKYYKATKMT
                  .  * *    .   **  *    .:.     *  *     ..   .** * * : *

35              GTCKRKMWRSKYGRYYYKMRRRYTRGKKGAATGCRKATCYYWRMTTRRRKYKSTRYTYYA
49              RKTKGTCWRGKYKGYTYGCARRCYRSGKTGGCAAAKTGYYYWRMCGARGKTKCARYCYTR
                 . * . **.**  * *   **  *. * .. .. *:  *****   * * *.:** *

35              CAKRTCGCTCTYWYTKCTGTRTRGMYCMTTRYAYKTKYCAAKYYYSKAAAGMGYKCAYKG
49              MRKAYYRSKYKYACCKYYSYGKGKCTYCYGGCWYKCKYAGGKYYTSTGTGRAAYKYRYKR
                  *    .. .*   *  .  .           ** **...*** *..:.  .**  **

35              GGAATRTKTKGCCGAGGKKAGCTGKCKTMGYWKYRYYTATATTAACAYGRRYKAGTCRYT
49              SRCRYRKKCKRYYRGTRKKRRTCATYTYMRYTKCACYKTAGCCTTMRYSRRYKMSYYRCA
```

```
                 . .  *.* *    .  **    .. . * * *   *.::.  ::  *.**** .  * :

35               GRKAACCKMRYGGKACGTGTRMCKKTCCTRKTYAKYRMYYWRSCTAATAAKKCKMMCKWM
49               TRKRRYMTCRYKRKGTAAACRMMKGCTYWAKCYMKTAMCTWRSMATGYRWKGMGAMYKWC
                  **    . **  *. .:. ** *      * * *  *  *** ::.   *    * **

35               YRWWGRGRRKTKTKWMRYYTCKCWRGCCCYMMTGWTMRYYSKYYGTRRCKKGKMACKWKT
49               YGWWSRRRRKYGKTAMRYYYYKYTARYMSCAAATWGMRYYSKCYRKRRTKKKKMRYGWKK
                 * **.* ***  .. ****  *      .   : * ****** * .** ** **   **.

35               YAATCAGYTTMKAACKKMMRMRAYRRYCRGCTAGAAWCGYYYRKRTRWKGYYMYMMGTCA
49               YRGCTTATKYMKTCTKKAAAAGGTRGTYRATCMKRRTTACCCRKACRWTRTYMYMAKWYG
                 * .  :. . **:. **     . *   *.        .   **  **.  ****    .

35               GATMTWTATAAYMKKWMCWTYCRKCYKRTGWSMSKYKGCRTGACCYKKYYYWAKWYYRYR
49               ACAMYAYWYRRYCGGTCAACTYRKTYKRCRASCCKYGKSGGAGTTYKTYYYTRKTCCRCG
                 ..:*       *     .    ** ***   * .**  .  ..  **.***  *   *

35               KCGKRYMAWAWSYSRTKTCYYMGCTARGTYKMGCCASTCCWKGKKWTYAYSGGYAYMRCR
49               GYKKAYCWTMTSYSRKGYYYYMRAGTARATKAKMYRGCYYTKKKKTATMYSARTRYMRYG
                    * *     ****.   *** . :  : *     .    * ** :  **.   ***

35               KTKKYKRTRYKYYAMRKKKCACYYKGTTCTTRMACTCYKKSCMTTGYRGCTCWSMTAGCK
49               GGKKTTGYGTKYCRARTKKYCMCYKRYWTCWACRAYMTKKGMMYKKCAATATWCCARSYK
                   ** .    **   *.** .  **         .   **. * .   . : *. : . *

35               TKCATGTAKTKGGCKAAKMRAKGCTYKAWGGYRCAKKACTAKCGAAACMKRCKYTTYTAT
49               KKYMCACGGCGATTGRWTMRRKAYKTKCAAAYRYGKKWAGCKGKTRMYCKRYTTWYCATC
                 .*   . .   .     .** *. . *. ..** .** . .*  :    ** .    ::

35               RRKCCKYYKRMYWRMYRYYTGKKAGGGRKWRTWYRWWWCGCACKWKMGCYWYKWGCCYYM
49               ARKAATYYKRCCARMYRYYARKKCAACRKAAWAYRTWWYAACTKTKMKTCTTKACTYTCM
                  **...****   ******: **... **    ** ** ... * **     *      *

35               KYRRMKRRKKRSKAYTAMRMGGTTGRKTKGMAYYGRKKWKAAKRATCMRYRAGKGYASCK
49               KYRACKRRGKASKTCYMCGMTTGCARKYKKACTYKGTKWGRGKRRYYCACGGAKKYWGAK
                 ***  *** * **:     *    .** *  . *  .**  .**       ..* * ..*

35               CRYGGCAKKKYYYWKWWSTACCAWRGGGGATTATAGGRKGKRGCCCCKKKRSGGYGYCW
49               YGYKKYGKKKYCYWKWWSWWMYRWRKRRRMKKMWGAARKRGGKMMYYKKKGGTAYKYYA
                   *   .**** ******     **     ..  ...**        *** . .* *
```

The pairwise alignments of two accessions 35 and 50

```
35              WRYYKGAYYKTTTMYKKKMWYYMKRMAACRYKGRGYRRMYYRCYYYTRRYYKWCRRCKRW
50              WRYYKAGTYKAGCAYKKKMWYTMTACWWYRYKKGRCGGCCCGGCCCYRRTTKWYGRYKRW
                *****.. **:   ******* *.     ***               **  **  * ***

35              YWMCMRMKMKYSKSACYYRRYCYRYWCYRYRKCRSCCMARWYMYMTGYKKWSYWAYYYYK
50              YTCYMACGCKYSKSGGCTARYTCGTWTYRTRKYRSAMCGRWYMYMCATGGACYAWYYYYK
                *   *    *****.    **    * ** ** **.  .****** .    .*  *****

35              GTRMAMTTKYYCWRWCTAGTGYKKYYRRTAKWMKYRTMMWYAYRSYRCYYYRYRGTMYSK
50              KWGATMGKGYYYAAWYCWTCKYKKYYRRAGTWMKYRCCMWYMCAGTGYCYYRYGRYATSK
                    :* . **   *      *******:..*****  ***   .    ****     **

35              RWYTGTRYWYMKWRRMYCWKGYMKYKRYYGGTRYAWMRYRRKWRWYMWYYRRRTTMCMCC
50              RWCGAGRTWYMKWGGATAWTACCGCKRYYKRWRYGWMRTRRGARWYMTCCRARCCCYMMT
                **  . * *****    .*..    ****   **.*** **  ****   * *    *

35              CCGYTYTMYYMGRCTMCYMTTSMAMRMRKYRMKCARWKMMRRAMKAAAGYWYYTKYKWRK
50              AGTTCTCMYYMRGGCCYTAWKGCRMRMRKYRMKYRGAGMMGRTCKGGGACTCCYKYKWRK
                .      ****         ..  *********     ** *: *....     ******

35              RWSRKRSRTTSAAYYTMKWTYKKRYAGWTWWRAGTTKRMRWKTKWWRYMYYYYYYYWKWR
50              AWGAKRSRKACRGCCGAGWYCKKRYRTTKWWRWRGGGRCRWKYKTWRYMYYYYYCYWKWR
                 *. ****.:. .     *  ****   .***     * *** * ********* *****

35              MMGAMWKWYKRRYKTCCTKWYKRTRWRYMRYMRYYYAGYRYRKSYMRRYYMCWMYCMYRR
50              MMRRCTGTTTRRYKGTTAGWTKGKGWRYMRYMRCTYTATGYRKSCCGGTYMYTATYCCAG
                **       .****   : * * . ********  *:.  ****     **

35              KRRYKSRMYWRKAYRKYAGCTKMWYWRCCGYYYRKRGTCYKYYMYSCTARGTYAGTWKMA
50              KRATKSRACTGKRTGKYGAYCKMTCWGYTACCCRGRACTTKTYMYSYKMRKWYGTYATAW
                **  ***    *   **..  **  *   .   * *.   * **** . *  *.   .

35              RCYACKYRGKYGTKCCKMYMGCRYKWKMAKGCTTYYAYTYRRYCGARYMTAGMWSYKAMG
50              RTYWYKYRRKYRWGMMGATAKSGYKWGMRKRMYYCCWTCCGRYYKCRYMCRRCTSYGRMR
                * *  *** **          . *** * *           **  .***     **  *

35              YCYMCASYWYKKYSGYCRGCMRRKTKYKRCTRYCYKRYYGMRTKCYYKTCACTCKCMYYR
50              TTCAMRGYTCKKYSATMRRYMRGGGKTTGMYGTMYKRTCTARWKMCYKATRAYAKMMCCA
                      .*  ****.  *  **   * .      ***    * *  **:  . .* *

35              TACGTTCTCTTSMTKKCYKATGRATGTKKCAGTCATGGMKMSRASMRRYYYKMWKYRTCT
50              WRYAYYTATWWCCAKKYYKGCRGCCACKGARAKYRYKKMGASRWSMRRYCCGMWKYGYYW
                   .   :   . :** **.   . . * . ..     *  ** *****   ****

35              GGYMRYRYMYKGCCKKGCTGCSRRWKKYRRRWSKKKWKCARYRYKGRTACYYYCAGGYCT
50              KRCCRYRTMYKRGYKKAMWKYCAATKKYRRRWSKGGWKTTRYGYKTRKRMYYYAGACCTC
                    *** ***   **.    .   *********  ** :** ** *.  ***...

35              RMGGCGAMMAAMYKAAMKKKKRYSWWKKYTWGAMAKWGMYTAAWGAKKTACKKGWCAWAR
50              ACRRMRMMMCMCCKWWCKKKKRYSWWKKCGWKMCGGWAMYKRRATWKTWRMTKRAGGWCR
```

```
                       **.   *   ***********  *   . *.**.     *.   .*   .*.*

35              YYKWKRKRTRRRTMAMGAKMRYAGTTKTKAYYCCARCYAAWMRTKTTCAKACACRYTKYR
50              CCGWKRGRYRRRCCMMKWGMRCGRWYKYKWCCYYRATCRMWCGCKWYYMKTTCTRYCKTR
                   *** * ***   *   ** .   * *           *   *    *: . ** * *

35              TMKWTRYKTAAKYWCCCGGMAMTGKKRTWMWTYYTTYKKTATGGRSAWGAYKRATATKTK
50              KAKWCRCKCCCKYWAYYRAMMACAGGRYTCACTCWWYKKWMYARGGGTTWYGRGKWKKYK
                . ** * * ..***.   .*   .  *         ***   .  ..   * *.. .* *

35              RKTCKWSTTGTCRWWYATGYRMKACTTKYKMTCRCGGYCTGYKMYTTAKCTYTTGYWKCR
50              RKAMTAGACAAMAAWYRWKYRMKWSGKKTGCYMGMRRYYYRYKATCAMKGATACKYWKMR
                **: . .: .:   **   **** . .*         *   **   : * : :  *** *

35              WRGAYCCRYWTYYWCTWRRSKGACKGACGTKKGAKAAWRTTYGRTTCAGATCTTTRYTRY
50              TGKRYYYRTAGTCWYKARRSGCTGKRRTACKKRRTTGTRCCCRRKGAWAGCYGYCRCCGT
                    *  *     * . ***  : *   . **  .:. *    *. . ..     *

35              SYKCGCTAWAARATCCGKWGRTAYYSRTMYYATRKTTCGRYKKKTGAKCTAARKSKTARA
50              CCTMRAYRWRWGCGTYRKWARWRTCCGCCYCTYGKYKYKRYGKKYTRKYCWRGGGKYRGT
                . .  .  *   .    **.*    .   * :  * .  ** **   *      .*   :

35              WYKKGSYTYTAAKYKKCMKCGCGKTGTAKKKCGWWCWAWMRWSYRKCGSKGTGMRGGCAY
50              TCKGSSYKCCGWKYTGYCGAKYRKCSCGKKKTRWAMTWWMRAGTGKYRGKRKRMRRTYWC
                  * .**.  . **.    .   * . .***  *    *** .  *  .* . **

35              ATWWTYYYRMYTYMCKMYGAARRMMRSTKAKYWGYGCAKKRRMYKKCYMMYTWKMCATGC
50              GYATKCCYRMYWYMTKAYKRWAACMGCKGGKTATYRYCTKRRMYKKYYCATYAKMYGCRM
                .   .  **** ** * *      * .. .*   *  ..******* *     ** .

35              CCMYAKMWRYRARMKCKYCYYYCSRRCCTRATTCAYWYYMYMCGARAGRGGGGCTGACTY
50              YSATWTCWRYRTRCKYKTTYCYASRRGYYRRYYGCCWYYCCAYKRRGAGARKSYYKRYYY
                 .  . ****:* * *  * *.***   *    . ***      *.. .  .      *

35              YGCGGYYYASYCCCCRTRMMYMMAGGATCCAGAWKYWKKKKAYAATCAGRAACCCRWAMA
50              YTAACCYCTCCATATGGAMMTMARRRCYTTGCTWKTWGKKTTTRTWYGCRWRATGRWMCM
                * ..  * :. . .    ** *    .   . :** * **.:  :  . *  .  **

35              CTRAAATRAGTTCMYRMWTATAKCCACTCGCGAWTTGTGKYRRYSYKGCRRKKTMGCKTA
50              SWRWRWKRGCCYYMYRMWKTGTGAYRMATATKMAGGACCKYAGCGTKTTRRTKYMKYKWR
                . *   .*.    *****.: : .   : .      .  **   . *  **.* *  *

35              MTYKKWKMTYCCCWRMTCWRRKYWRGACYGWYGSKWTAKSKACYRGRKMGCCWKGKGGWR
50              MYYKKAGMKTYYSWGMWYWRGTCWAKRYCRWCRSKWYMTSKGYCGARKASSMAKKTRRWA
                * ***  *.   .* *  ** . *      *  ***  .**.   .** ..  * .  *

35              MCTTACTKWRCGAKKWRWGTGAGGGCYYKRYGWASGTTAMMYKKCCATTKTCAKCAGYYK
50              MYAYRYKKTRTCTKGWRWCYRGTTATTCGACKACCKCYWMMYTKYYGAKKATTKYGKYYK
                * :   .* *  :* ***   .  .        ..    ***.*  .:.*: :* . ***

35              ARMWTYKKGTGYKRKMGACGGAAMCAGYGKKYKGTATCGTYYGYYRCGRTGRKYKKKTMY
50              RRMWWYKKAGCTKAGCATTTAGGMYRRYKKKYKKKTAYAKCCKTYRTAAWRRKCTKKYAY
                 *** ***.   *   .:  ...*   * **** .:: ..    ** .   ** .**  *
```

```
35      RYCACGKYYMKKYRGTARKYMCTYRMRKRKWWYAKKYTWKMYWWKGKYAYKGMMYKMKKK
50      RYYRYRGTCAKTYRRCRAGCMYCYRMRKRKTTCWKKTGWKMTTTKRTCRYKRMMYTMKKK
        **        *.**      *  *******    **  ***   * .  ** ***.****

35      KKCYYRRKKCYKYWYKMKKCAKAWKTATRGRCRRKTRWRYTTAYYYWSKTACKCTGRCKA
50      KGTCTGAKKMYKTWCKCKKYGKMWKYRYRRGAARKYAAGTAWMYYYWSKACYKYKKAYKW
        *      ** ** * * ** .* **   *  . **     :  ******:. * .   *

35      TAAKARGTKTTCGATCATTRACAYTKGKWYRCWRRYMCTTTYGGCYRCGCTATKCRTSYR
50      YWMKGRKYKCCMTMYYWWWRWMWCYTRKWYRYTRRTCMGCACAAYTATSTGMYKTGCGYR
           *.*  *          *     . ****  **     : ..    .    *   .**

35      CGGMGTMWKYRMMRRGATAMYKWYRRMYGKKCCATAMAKTWKAMYKAGTCAAACWRKRMG
50      TRKMSYATKYRMMRRRTCGATKTCRRMYKKTSTGCGMCKCAKRAYKRRWMWRRMAATGAR
           *.   ******* : .  *  **** *.. . .*.*  *  **          .

35      KWACYSYWRYYTRSSKGTGTTTTWYSRAAAATCKRYYYATCTGGAYAGAGKKRWRYTGAW
50      TTCYCSYWRYTWASSKACAYYWYTCSRWWWRWMTGCTCRAYCKARTRSMKKKRARYYKTA
        . .  *****   ***. .      **      .     :   .   .  *** **  :

35      GCKRGCKYRRRKAAAKTRKYKATGACARWKMWAAGGRKYAAGGKRTMCAGCRRMSKKWCA
50      ATKGRYGTAGRKWMMKGRKYKRYRRYRRAKMWTWCRGTYTGTAKGCCTWCTRGASKKWAT
        . *       **   * ****      * ***:    .*:. .*       *  ****.:

35      CKGRRYKGMWGGRRYKWYTWCYKMCGGGSRRAASCARMGTAYCMMWRYMYMYYYTAYKYG
50      TTTRRYKACWAARGYKWYGTYYTAYRRRCRRTTCTWGAACGCMCMWRYMYMYYYGTYKCC
         . ****. *..* ****   *.     .**::.    . .   ********** :**

35      CAGMRKYYMKKAWKTCTATKYRYYACCKTTAKRGCKTTTYWKRKTRMKRKGRRCGRMRTT
50      GRRMRKCCMKKCAGWYCRKGYRCYTYAKCGTKARSKWYKCAKRKYRMKAKAGGTAGMGWW
           ***  ***.      . ** *: .*  :*  .*  .  *** *** *.   . *

35      ATACARKRTAGGKSRMMRKTAAKRKTWRSYWYKRWKTGTWMRACYRTKYCTKTRRAGCMC
50      MWWMCGKRYWRTGGRMAGGATGKRGYWRSYWYKRWKYACACGTYCGCGCAKKKRRRRMCY
            . **     .**   ::.**  ********** .    :      ..*.**

35      MSRGACTTKWKKGCTTTAKMWCTGRKCKRGKKYWWKGRGGCYCKWGWGGTCKYWWSKTGA
50      ASAKGYWWTWKKRYYKKMKCTMKKRKMKAKKKTWWGARRRTCMGTKWKSGYGCAAGKCKG
         *  .   .***   .. *   . ** *  ** ** .*        * .      .*  .

35      ATRTKYTGACGTRCTAYACRAAAKYCATGGTRGKKKCGRTTYRKACAYTRYRTCGGKKYC
50      WKAAGCWKRYKCRMYCCRYRRRMGCTGCTAAARKTKARGWCTGGWMRCWRYRYYAAGKCY
         . :        *  .   *      .  .:  *.*.            ***  .. *

35      RTKYGTTTGATMKGTRKCTACTCGTYYYKGYRWYGRKTTTKYRMMYKKTGGYAKYYWYAM
50      GWTCSYACCTCCKSWRKTCCTCMTCCCYGKYRAYSRGYYYKTRMMCKGCARYRKYYWCCC
          . . :  :  *. **  .       *  ** *.*    * *** *  . * **** .

35      CCCGYRTKRGKKRKWCGRRMSYRKRKRYRAGTRYYKTATRCAMWWKYMSKRYKTAYRYKR
50      ATTKYRKKASKKRKTMRGACSYRKRTGYGWRKRCYKWMYGMRCATTCAGGACKGMTRYTR
        .   **.* .****      *****. *   .* **         .  .   *   **.*
```

```
35          MTCGYGAKCCYYKMYYKYYKYKKYYGGCKKMRCYRYYYRYCTTWKKTTKWGTYRTTAYCK
50          MWMRYRGKMYTCKAYYKYTKCKKYCRRYKKMRATAYCTATTYKWKKCCGTRACRWWGYYK
            *   * .*    * **** * ***    ****.  *      .***     : *  .* *

35          TARRKGYRMYAKCKKKRAWMWAAWRRYYYKYRTTARYAGYTCRMRWCKTMGGKGSTTRCM
50          WRARGTCRMYCKYTGTAGTMTCRWRRYCCKCGCCTRYTSYWYRMRWMTWCKRKKSACRTM
               *   ***.* . . . * . ****  *    :**:.*  **** .    * *: * *

35          CYKRKTYCTAYAAKYCRYYYWRRWYKAAMMCTGAWMKRCGRWGKTKYKKMRYYTCCWRRK
50          MYGAKCCTGGTCMKTTACTYWRRWYGRWMMYCACWCKGYSRTRKYKYTKMRYYYYYTARG
             *  *    . . *     ******   **  ..* *  .*  * **.*****     *

35          YGTMYYSKYGCKKCKKKACRCKATTTMYKMKTTRTTCTWASWKRKWAKYRRCWWKKKMRK
50          YKGMYYSKYRGKKTKKKRYGMKGKYCATGMKCGRYWMYTWCWKAGWMKYRAYWWGGGAAK
            *  ******  ** ***    *..     **  *      .**  * ***  **     *

35          WAMTTCCTARRKAGMTKYRAACCKTAKRWRRYTTATWCAKKCMCAATYTGTAKGTCTTGY
50          WGAKAYYCGGRKMKCAKYRGCMYKWMKRWRRYKGCGWYGKKSCSMMWYGKYRTRYYYYRY
            *. .:   . **   :***..  *  ******. . * .**. .   *    .      *

35          AAGYYGMSCTCGCKGAYKMYYRAKAWSKKRYWKRYKRYKCGWKRWKKMGCMKRSWTWTCY
50          RGCYTKMCYWYATKRGCKMYYRGKWWSGKRYWGGTGRTTMRTKRWKGMATCKGSTKACTT
             . *  *.   . * . *****.* ** ****    * .   **** *.  * * .

35          STYMCRYRSKKCGTTGTATKAMKAYYTCTAMWYSKTYKGTKYYMCYKTMMWMRYRAWCMK
50          GYTAMAYACTKYACAKCRYKRMKMYYAAAGCACGKYYTCAKYYCYTKAAAAMRYRRAYAK
            .     * ..* . :    * ** **:.:.   .* *. :***   *:   ****    *

35          RGGCGRKKRYRKAMRASYAAYMYYMKKSMMCYSKAASAWKGKGGGRRYYYKCYKYCTTCR
50          ATRYTRKKACGTMMRRGYWRYMYTMKKGMCMTSKTTGWTKTKARRAATYYGYCKCMWKMR
                 ***   . ** .*  *** ***.*   **::.  * *.     **   *   . *

35          YKYYARYYRTRKMRKKYGMTKACWTARAATAGYYKKKGRRMAKKCYGYTGMYWTACAMKY
50          TKYYRATTACRKMRKKCRACGGMWKRRRRAMRTCKKKRRRMMTKMYRCYRCTTAGMWMKY
             ***      ******     . *. *  :    *** *** .* *       :.  ***

35          KSGGCGYAYKYTMRGGWTTGCCRYKWWCRCRGAATGYYASYYRWKRGGKKCACKCACRAM
50          KGRSYSYRCGCYCAARACGRYYRYKWTTRTGRMCCACTRGCCGTGRAATGTMYKYRMRGM
            *. . .*       .       ****  *    . .   .     *...    *   *.*

35          GACYRKWYYGTTARCCYCCKAGYCACYCWYATACAGKWTRKKMKRWRKAAAYMWCRYMRM
50          SMYYAKWYYACYRRMTYTTGWRTTWMYYWYGWRYGRKWARKKCKGTGKGGTTCATRYAAC
            .  * ****.   *  *         * **.   . **:*** *   *..:    **

35          YYTCCCRRTRYCGCKCWTRWKTMYKSCTGWMGYRRACTAKCTRACACCMCMKRCTYYYRW
50          TYCTATRRCRCMKYKAAYGWKYCTKSAGKAMRYRGWAGCKGCGMSTAYCYCKRYKTCCAA
             *  . ** *    *.   **   **.   * **  . .*    .:.    ** .

35          WRTYYYTRTCGMYWMGCYTKWRWRTWYRWWWKCGCACKCKCCGCMKRMKKKKCCTKKKYK
50          ARGYYYWRAAACCAMRYYAKARAAWAYRTWWKYAACTKSKYYCTCTACKKKKYYKGKKYK
             * *** *:..   *  *:* *    ** *** ... *.*     .  ****  . ****

35          RKYKRYMMRYRKKGCGTRYKRKMWMAKKRATCAAAGKGTCKGYCASKKKGGYARYWYWWS
```

```
50    AGTKRYMCGCAKKKYRCRTKGTAWMGKKRRYYRRWRKAYYKKYYWGKGKKKTGRYWTWWS
         ****    **    * * . **.***       *.  * *  .* *   .*** ***

35    MWRKTMWGCAGRCWCYRYCYCCTCKKSGGYYMYCWTAGKKACGGYYARKMWAGTAGARMT
50    MWRKCCARYMARGWMCGTMTYSYMKKGTAYYMYYAGTAKKWYRKYYWRKAWGKCWRGGMK
      ****      .* *       .  **. .****   :.**    ** ** *.    . *.

35    YAYYKKGCCGCTTRMWACWCMYYKKMKYCKYWAMKAMKYRRCTKWCKYWCYWMYKMAAKK
50    YWCYKKRMYATCWACTGMWACYYKTCKYYKYWGMKGMKYAAYKGTYKYWYCWMYGMRRKT
      *  ***   .      . *. ***. ** ***.**.***   .   ***  *** *  *.

35    TAAYAAKTGCGTKKYKYYYAGYAYGTGYYWYYAGYYRRGYRKGAKYTGYKKYYYYRWYKA
50    GRRYMWKGCARWKGTKTCCWRCGTAGACTWCCRKCCGGRYRKSMKTGAYTKCYTYRWYKR
         *  *  .  *  *      . . .  *         ***. *  .*.* * *****

35    GWYMRRCKRWRRKKMARGMKKGARCRYTKYRWYMYRCKWRRRYYYTGKYKAAATAKKGAK
50    RTYMAGAGRWGAKKMRRKMTKRRAMGCGTYAACATATGWRRRYYYYRTTKRRMYWKKATT
        **  . **  *** * *.*       .*        *******  . *     **.:.

35    GRCRYACRGKRCAGCTKRKYSKYTCAWRKGRSCKKRSRKKYMKKRKGKCWMMTYKGRYCC
50    RRYRCMYGKKGMMRGGGRKYSKYCTRTRKKRSMKGRSGKKCCKKAGKKYACMWCTAGYYS
       * *     *       ******    ** ** * ** **  **   *   *  .. * .

35    ASYMMKMKKYWWTRYCYRYSYMRKGTKKRTGTCKYGTKTTRKKMWRKATRYYYTRKATCY
50    RGCMCKMKGYWWWACYCACCTMRKRYKKRYKATKTACTYYGKTCWRKWKRYYCYRKGGYY
       . * *** ***       . ***  ***  : * . .   *. *** .***  **.  *

35    WKKTRYYKKGRTCRGCTCTYGGGAGMYSSWYTRTRTRCGMYMCAWAGARRYKKYKTATYT
50    WKGARYYKKKAYYRRSKYKTARAGAMYSSWYCGCRKGYKCCMMTTWRGRRYKKYGYRYTC
      ** :*****    * .. . . ...******   *.     * :   .******

35    KTTRYRAARCTCTCWRMGGCARAARGWRMMKMMRGGTACASYTCGCGSKGRRGCCMYYKA
50    GACACARWGTGYCMAAMACATGTMAKTACCKMCRSRYGYGSYCTRTASGARRSYYCCTTT
       :              *. .: :       ** *.  . .**    .* .**.     .:

35    GCTYTCYAGAKGGWTTCACKAAMGTGTMMMRATGCAMCGRKAATCMCYMGKAGGCKKWYY
50    KMKTCACCTCKATAGKYCAGGGAKWRWCCMRWYRARCSKRKGGGTMTCCKKTTATKKATT
        .  . . .*.   . .. ..       **   .  . **..  *    *: . **

35    RYKKCCYKRGKYYCCGCATRKCKKYACKRACKKGAYCKYWSKKAMAYYCAKMSAKGAAKT
50    RYKTMYTTRRKYYYTKMCCGTMKKYCTKRTTTKRRYSKTTCKKRMRCYYWKMGRKKMRKC
      ***.   .* ***    .  . ***. **: .*  *.*  .** *  *  **. *   *

35    ATARRWAWYYCAYYCARRSKGRTWWGYKGMKRRYWMKKGGMRSMRRKMTACAKWRMRRMY
50    TCTRRWWTYCTTCTMRGGCKAGKATKCKACKRRCWCKKRRARSMRRKMYRMRGWAAGAMY
      : :***  *  :      .*. .    *. *** * **   *******     *    **

35    YRYYYAR
50    YRYTCCG
      ***  .
```

The pairwise alignments of two accessions 35 and 51

```
35              KMYAGYYRWMRTTKKCCCCRWYYACYYWRAMYYMCKKMYTTTSMYRYMGASGRGTTTCYR
51              KMYTRYYGTCGGKGKYYMYAWYYRYYYWRWMCYMTKKMTWKWGCCGTMKRCKGRWGCMCG
                ***: **     . *     ***  **** * ** ***  . .    *  .

35              RCWYCKTYYMKRRKTYMCTRWWYTMTKCYMRRRKYACTKMRWRYYRYRGYYKGMCYKKRY
51              RGWCMKYTTCKGRKGYMYYGTWYKMAKYYMRRRKYWYYKAATATTGCGATYGKAYYKKRY
                * *  *    * ** **    **.*:* *******   *         . *    *****

35              YMKRWYYRRMSRRKWMKYTWAYRWMRMYYTAGGYRWRYRGSYYYRYTYKRWYGKWYMKRR
51              TMKRWYYRRMSRRTTMKYCWWCGWMRMYCYMKAYGWRCAAGTYTRCACKRWCRKWYMKGG
                 ************. *** *   *****    .* **  .. * * : ***  *****

35              MCYYYAGGMYWTRYKCCCWMMMATCMAAMRATAKCKARTRCKKCMYYMYRRRWWYRKYKG
51              AMTYYMKRCCAGRCKTTAWCCCCGMAGCMRWYWKYKRGCGMGKTACCATARATWYRKYGK
                   **       * *  .*   .   ..**   * *      *       *  *****

35              KYKSYKRTRTTTYYAKTYGCMKWYKWWKMCCMTMRMWRYRATYRAMWKRGKWWRYMRWYY
51              TYKSYKRGGYWKTCGGATRYCGTTTTWKMYYCCMRAWRYRTGYGGMWKRRKTWRYCRWYC
                .******    .  . :       . ***    ** ****: * .**** * *** ***

35              WYYYKRTSWKWTYMRYRYGGTRKKYYKRRATYYWTWMYYTTTMRYCRTYWYYYWGYRKYY
51              WCYYGAGGWKWWYMRYAYKRWRKKTYKRRTCYYWCTMCYKAKCRTYAYCWCTYWRYRKYY
                * **   .*** **** *   *** ****: ***  * *.:. *     * **  *****

35              RKYYYYYYATKRTYWTGRYTYKCGCWGRMYYRKGRYCACYAGTTGAGGTARRYMAAGSST
51              RKTTTCTYCGGRKYACARCGCTMKYWRRMYYRTARCMRYCRTYCARAAWRRRTAGTKCGY
                **     *.  *.*  .*   .   * *****..*         . ..  **  .: ..

35              TCAATKYYMRYGYKKRKGYACTCTCRKAARKSTMGTARAKYYCYKKKCSWYYWRKMKWKK
51              KYWWCKYYMRYRCTKATACRYCYGAAGWRGGGGCKYRGTKCYYYGKKTSWYYWRKCGAGK
                .    ******  .* ..      .      .      :* * * ** *******    *

35              ARAWTCRRGKRTGTTTGTTSKGGYRACCRACATCYCCGKKKYRACTMKRYRKYYYTAMRG
51              MRTWCYAGRKRYRYYKCCAGGAKYRWYMGTTGGYYTYKTKKYRCGYAKAYRKYCTCRCGR
                 *:*     **    .  :. . **    : .  *   .****.   * ****

35              SAYCRKTYRMRYGGKKWYRRWWAWSKYAKKWKKCCCYMYYWWMRKKGAGGCCMMYRRCKG
51              GRCMGGCYRMCRRKKWYRRTTWWSKTGTKAGKTAYYMYTTTAAGKRRCATAMMYRRMTR
                .      ****   ******   *** ..*  * . ***      *   . .***** .

35              KKWKTYTYRRRTMYGKGCYKCTYAAWTTYKMTAMKWKYTATGKWTGYYYARGKAWKGYRM
51              KKWGCCWTRRGAMYRKCMYKMWYRTWYYYTCAGAKWKYYRYRKTATYYYCAKKCWKAYGM
                ***     ** :** *  **  * :*  *. :. ****    * : ***.  *.**.* *

35              ACGACGRTCKMTCAYSRAAAGTARYYYKGYWARYKMWKAYRGASATKCGWCCCKRRWCKY
51              WTAMYRGCYGAWYRYSRWCMACRRYCCGKYTRGTKMWKRYRKMSCCKMTTATMKAATTKY
                  .           *** . .  **    *    **** **  *. *   .  *    **

35              KRYYKYWYRYKYCYCCYAGGRGTMSYGRKKRKRYCMMGAMMMYRWTYKKAKCGAKRCATY
51              TRYYKYWYGYKYMCMYYRRAGKCASYRRKKRKRYYMMKMMMCTRAGYKKRKYKGKRSGYC
```

```
            .******* ***    *  .    ** ******* **  **  *  *** *  .**..

35          AGKACMWRTRRTMACMACARRRRRKKYYCGAYCACCWCYRYYRRKKYAAWYWCTTRKARY
51          CRKTTCTRCRRCCMYMWTGARRRRRKKYCARMCMMMMWYCACYGAKGCRMWYTSGYGKRGY
            . *:   * **    *  . ******* .       *    *  *    ** .   *  *

35          CKTCWCGYYYRTYYMYYSMRGTMAGTWYKCYKYRACKYGCRYKWYYRWYMAKMWYYAAYY
51          YKATTTACCCRCTYMYCCARKWMRKYACKTYKYRGGKYKSGYKWYYATCCTGMTCCRWTC
             *:   .   *  *** . *  *     * ****. ** . *****    : *

35          RYKRMMCWGCYMGRGATAYMRATTCASMAKKAYSGGKWTRRYKTYYRKMYRTYSWKMYWR
51          GYKGCCYTRYYMRARGAGTAAGAKMRGCCGKTTGARKWKRGTKWTCRTACRWYSWKCYWR
             **       **   .:.   .:.  . . *: .. **.*  *   *.  * **** ***

35          MMCYYKTCYACTYYYKTGGTTCTCWSMGATKCYATRACAYAKATCCCACGGGYAWWTYRC
51          CMMCYKATYRAYYTCGKKAYYTATWCCAGAKYYGCGWTRYGGRCYTGRTAASCCWTKYAT
             *  **: * . *   . .   : *. ..:* *.     *.        ... .* .*

35          KCRRTTATKAAKAYWWWGCKTCMCKTCKRRCKYKSKAWKKYWWGYKYSYKATGAKKRRTT
51          KYGGWYRYKWWTRYTWTKYKWYMTKKMKRAYKYKCGGWKTTATTYKYSCGCAAGKKRRAA
            *       *  . * *   *  * *. **  ***. .**.    ****  .:..****::

35          AAYAWSTCTGYAYGMKCWRYRMKMRRATCCTKTMKWKRMWYRYWYRTRYMATTSYMRYCK
51          GGYRWSCACKTWTKCTYARYRCTMGGTWYAWKCCGTGGAACGCWYGAACMGWCCTMRYMK
            ..* ** .       .  *** .*  :  . *           ** :  *.  . *** *

35          KTGGYTCGYYRRKRWRKGGKRYKKKKYRTMWMAMYTRGWKKATCYYWKAATYWWKGKRKK
51          KYRKTCMRTYRRKGWRKRSKGYKKKKYRWAAAWMCYRSTKKTGYYYWKGGCCAAGRKRKG
            *        **** *** .* *******     *  *. **:  ****..      ***

35          YRYYYWMYKRKRCTTWTKCRKATTRMTRYGCCGGCAMGTMAACYKGYYSYYKKRGRTGGK
51          TRYYYAMTKAKRYWKWCKYGKTCKGCGGCRYMTATGMRWMGRYYKAYTSYCKKAKRCKRK
             **** * * **  .* *  *: .         . .*  *.  **.* ** **  *   *

35          YYYRCKYWSRRKCCRYMYKCYKCGAATCAMWGACTYYGYYGYCSMCYGGCCCATWTYYWK
51          TYYRAKCWSRRTGGRTCCGTYKYKRRKYMMTKRYYYYATTATTCMATRRSTTGKWCCTWG
             ***.* ****.  *     **    .  *     **.  .  .*.   .  ..*   *

35          GAYGKAAATTKRYCAGCYTYKMGAAMTGRKRACTRAAATRRWRTKRGTYTTTKKYRMKAA
51          SRCRTTRTWWKATYGCMYWYGCKCCAYRRKRMGAATRWKRRTACTGRYYCCCGTCGMKWT
            .   .: :  *   .  * *   ..   ***  : :  .**   .   *    .  ** :

35          TAKACGTCTACTAKGGCCGGCARTTGTSATTRMAYRYKGRKKWYRCACCARYMCATYGKG
51          GTGWTACTYRTYGGAATTARTRRYCACSMYWRMTTRCTRATKWYRYCTATRTCYRKTRKA
             :   .      . ..  .   *  . *   **: * .  .**** . .:*    .  *.

35          AKTGRRKCACRCTAWTCCAKCAAAGGWKYRMAAYYGCYKAYKYMRYKMYAMRRTWYSWKM
51          GKYARRKTTTRTAMTYMYRKTTGMRKTKYRMGTCCRYYKTCKCMGYTMYRARGCACSWKA
            .* .*** : * :      * :.    ****.:    **: * * *.**  *    ***

35          GCRMYCACKCGYTGRYKRYRSSKYRWCTRYTTRTTKGSAYTGYAAGGCGAGTCCCGGYGW
51          RYRMYACSKTKCCAGCGRYGSSKCAAACRYYGGYAKKGWTKKYWWACAACKWMYYRKYKT
              ***...*    .   ** ***   . **    :* .  . *  . ...       *
```

```
35          RMKTCKKCYGCRACKSARCAMWCGCGRTGTYRMYTKGKWCRYTCTRGKAKGAMWYRATAC
51          ACKGYKKYCCGGGTGSRRYGMWTRTARYAAYRCTYTKKWYRYKMCRTGCKAWAATRWAWM
              *  **     .  * * .**   .* .:**   . ** **.  * .*.     * :

35          GGRMMRRAWTATCYTGWATGRKSTAYCAGAWMWAKWATTAGMATCCARYATGAGACCCCY
51          KRGCMRRWWYRYYYYKAWCTRTGGCTMRRRWMAGGTWYYMRATGTMGGTRKCMCTTSYYT
                *** *    *      *.. .     ** .        :   .   .   : .

35          RWYKYRYTRRCRWRRKYMARGCGMYRYKKTCTRYKTGYMYRTTRWKKARRYYTGKKKGRC
51          RWYKYGYKAGYRWGAKYMRRSMRMYRYTKYTGGCTGCCATAWARWGKCRRYYYRKKTRRY
            ***** *.   **  *** *.  ****.*     .       :** *.**** **.  *

35          RYCGKRGGCTYKKAGCKCGWKRSTRKKYMRKGKKCWKMYKGRYKSYMAGKRTTCTAKCAY
51          RCYKKGRKGGYKKCTTGMRWGRSCGKKCCAGKKKYAKMCTAGYKGCMRRGRCGAGCKGTY
            *   *     ***.     * **  **     **  ** .. **. *   *  . .* :*

35          KKYKYRCYYYWWYYTRMYWTAAACCRMYRYSTTTKGRYRYYKWRKMYYYYWWKYWTWYRW
51          KKYKTRYTCCAAYYWRCCAKCWMYYRMYRYSAWYKCRYRYYKAAKMYYYYWAGYWWWCAT
            **** *      ** *   ..    ******:  * ******  *******  ** *

35          CATGKKCMWRMGMGKYRWKAGYKKAYMMRMGTCYWYCYWKGMCRYGTTYKKMKTWAAAYT
51          TTGTKKTMAACKMTKYATKGTYTKGYMCGMKYTYTYGYWKKAYGYTCATTTAGWAGRGTY
             :  ** *    * **  *. *.*.**  *   * * ***    *  : ..    . .

35          GKATCAAAGGTCKGYSCCRCARYGAARTTKKYWWSMWRGGGGTGGRKGKRKGYCSYYCRS
51          KKRYYRRWRAYYKKYGAYGYTRCKRGRCCKKTWWSCWRKRRRCRARKAGGKACMGCCMGG
             *       .  * *..   :*   .*  **  *** **      .**.  *.  .    .

35          GGYMTCWKYRRRTWTKRCTWCKKMRKTGAMCCAKTGMRKGYMYGTATYRRCTACRRRKYA
51          TAYMKYAKYRRRYACGRYKWYGGAATYRGAYMRGKSMAKKCCCTAGKYRRTCGTGRRKYM
             .**.  *****    * .*     .  .     ..* *     :..***  .  ****

35          GYTMGCCTCAMGRATAKRWRTTTTWCMKCCMYTAGYTGYCYRTCTTYAGAYRCRWSKWMR
51          KYYCATGCYGCRAGWMKRWRKKACWYMKGSCYYMAYKRYYYRWYKCYGSCYRAGWSKWMR
             *  .    .   .  ****..: * ** . *  .*. * **  . *...**. ******

35          KKRWRYKCYTGWYYMGCMYKWRSAWYSTYMRRSMGGKACGTGKCGYSRAATKRTYYYACY
51          KGRWRTTYTCRWYYMRYCYTAGCMTTGYTAAACMKRKCTAAKGSRCSRWWWTGKCTCMTT
            * *** .    ****   *.  .   .     .*  *. .:  .  **   . .

35          TYGTCKGYATCGKKRMYYTATGRCCWYRYMKKAAAKTMRYRWAAAWGSKRYYYGYAAGGG
51          ATKAYKTYRGTKKKRMYTGGCRRYYWYRYMKKWMMKGMRYRAMCGWTSKRYYCRCTGTAA
            :  : * *    *****  .  *  *******   * ****  ..* *****   :. ..

35          KCRTAGCYYTCRCGRCASKKWYWKKCTGKRRATYKKRWATGRMAAYTACYKRRKGGCMRM
51          GMGCWCTYYYMRTRGYGSKKWTWKKYCTKRRRAYKKATRYRACGGCCMTYKRGKAAMMGC
                   **  *    .**** ***   *** :***       ..    *** *.. *

35          YTMYYYKYGAGTMKRYYYKAACGATKAAKRMYARYYKTKKKKMYGGMGAGKKWYYWRRSK
51          CWMYYYKCSRRWMKACCCGRRTACKGTGGAACWRYYKWTTKGCCKKCKRKTKAYCAAGGK
              ***** .   **        ... :.     **** ..*         .* *    .*
```

```
35     TKKGTYCKCMMWWMRYTAWGGGKRYRCTATGYCATSYRYTCAKCSYSYGRCACAAASKKW
51     YKTCAYTKMAATAMRYYRAATTKACGSYMYSTMTWGYGYYYMGYGTSCAAYRYMTTGKKW
        *. :* *     ***   .  *   .   .  : .* *     . * .     ::.***

35     KGGTRRYYKYAACKYSAYTAAGRYYRTRKYRMRYKKMYCAAKGYRTAARAATGGCYCYMM
51     GSKWAATYKCGCTKCGGYYRRKATTACRKYRMRYKKCCTGCGACGACGRRRARTACATCC
        .     ** .. * ..*         *********   .. .  :..*  :  . .

35     KYGKCGTGKKCTCYWCKMKKKYKRRRYGTGTTARKAGGCYKYRARTMTGRYSTGCTGACC
51     TCRKTAATKGTAAYWMKAKGKYGRAGTSWRYWRGKGAAYCGCGGRKCGRGCCWRMYKRYY
       .  * .: *  :.** * * ** *   .      *...     .*.     .

35     RRKWWCCAYWKRKTCTRTMRKRARMARKRTAGGGYMRKTKYYTRTKTMWRSYWYAGKAAY
51     RRKWTTYRCAKRKYGYACARKARRACGKRYWRRKYAGGAKYTCRKGYMWRSYWYWKKTWC
       ****      ***      **  * . **     *   :**  *.  *******  *:

35     CGKWRYGYYWKKATRKSRCKKWKCTKTATMKCRKCGKKWTYRRTYRWKGRWGTCKYGTGR
51     ATKWRYAYYAKKGKRKSAYTKWKYYKKMKMKMRKMKKTACTRRCTRWKARWSGYGCRCKA
       . ****.** **..***  .***  *. .** **  *.   **  ***.**.

35     CMATTTKYYATRCKWRRYKGTCATCYCAARKKCGRRKTRAAGTCYYAKGTWAGGMGCKCT
51     YCWKYAGCYTYRMKWRGCKRKTGCYCTRRGKKARGGGWGGCAWYCTRKRKWCARCKSTYY
          . :  *: * ***  * . .       **.      ...     * .*..   ..

35     AYYRTGACTAGYAKAGYCTYMKKTRKWRGRCGRGCRMRMYRYTCCCKMRMYWKCRWTGWT
51     RYYGYRWMGTKYRKRTTGATCKKCGKTGAAYKARTGMACTATCTATKARCCTKARWKRAY
        **      : * *    :  **  *  .       *       . * *   *.**.

35     RGCGCTYYTASGGYAYCMWTMRRTKTGTATKCRRTTYTAYGRGTTGTGRYKKTWRTRYGC
51     GRMRYWYYGGSCRTCCMCAYMRRCKCACGKKTGRYGYCWTAGAYYRYRACGGYWRCRCKY
             ** .*   .     *** * . ..*  *  *   . .          ** *

35     AAWCCCGMMTKKRTWMWTYKYKTTCGRRRWATGKTKWSTYYRYATGRMGTKGKKMTKAAA
51     GCAAYYKMACKGRYTCAWYKCKCATAGGGWWKAKYTAGAYTGYRWKRMRGKAKKCYKGTT
       .. .   *  * *     ** * : .   * ..* . .:*  *   **  *.**  *.::

35     RCGGRCTRGCTATKMRYTKCKKKRWGYGAWSRRSKCYRRTYAGGTYYRRYRGGACKKTAM
51     RMRRRTCARACGAKCGYWKGKKKATTTRWASGGGKACAAWYWKAYCYRRYRKRRAGGGCA
       *   *    . .:*  * * ***       *  .*.    *  .  *****   .   .

35     GTTTTYCCWCYKKRYRGGYKYCRTGATMTKACGRRCMRYYCKRKYKWYRRWKYARKTCTT
51     TYYCWYAMWMYKKRYRRRTKCTRWCTCCYKRMKRAMMRYYTTRTYKTCGGTGYGRKYYAA
            *. * ******   *  *  :   *   *  **** .*.**      *.**  ::

35     KWYGYYKKTGGKYYWYYAMCCCGYRTKGRGKRCKRRRMCMRYRMSYRKKRGKAMYRYKTA
51     KTYAYCKKCARKYYWCYCCATTKYRKKKASGGYKRRGMMMRYACSYRKTGRKRMCRYTWM
       * *.* ** . **** *. .   **.*  .   *** * ***  ****.  * * **.

35     TRCAMWWYMWGTAAKRMKKKYYKYKYKKYGCKKMRYYRYCYMRTWTTKWRTYTTGAYRKR
51     YGMRCATCATCGMMKRMKKKYYKTKCKKYRYKKMRYYRYTCCGYWCCKTGAYWWRGCGKR
                     ********* * ***  ********     *  *  :*   .  **

35     KATCKCKCGGTAYRGYMGTKKGATKCRCTWRCKKYACRKATMGAWKKKMRAYRMWAYWMK
```

```
51              KCGYKYKYRRWRCGTCCCCKKTGGKTAYGWRYGKYMMAKGYMRWAKKGAGRYACTGYWCK
                *.  * *            ** . *    **  **   *. *   **    *   .** *

35              KRSSCACCTATYRGYYMKCRRRRKSCYTTTWYCMAKKKTYMYKYAAAWYGTKCAGATTAY
51              KRSSYMMYKGGYGRYCMKYAGAAGSYCCCAWYMMRKTKGYCCGYGGCWYRYKSWRRYWGT
                ****    .. *  * **      *    :** * *.* *   *...**  *.     .

35              TGTGAYGGACYYARYYKWRWTKTKRRWGTGKWARGCAKSYGCKRRRYRYKCCATAGCKCY
51              YAGAMTRRCYCCRGCCKWAWGGKKGGTACCKWRGAATGGTTTKRRGCGCKGAWCWKYGMY
                 . .    .       ** *  .*   .  **  ..: .   ***    * .       *

35              KRKMWTYRMCYRKYRACYGWYAWTAYRMGCKGRKAYASKGCCWGGGGRTYTCGACWRKKY
51              KRKMWKTGMYYRKYRRYCRWCRWYMCRMCTGRAKGCWCKASMARKRRAKYCYRRYWRKGY
                *****.  * *****    *  *   **     *.  .*..       .*     *** *

35              TYARKGGCKRYGGWKRMMYKCAKCKKCKKAKMKCRWMWTCTYKKGTGYKTRMRKMGACTC
51              CYGGKTATGACTKAKGMMTTSTKYKGYKKGKMKYRAMWWYWYKGAGCTKWRMAGCRWAYM
                 *. * .       * ** ..:* *  **.*** * **   ** .   * **     .

35              TAGTTTTYCAGYKKATKYCMYYYYCYGCCGAGCKWYYRYYKWYTKCYKGWTGKMYYCKTY
51              YRRCCAYYYRRYKKGKTYYMYYYYTCATTATATKATTRYYKWYCGYTTRTYSKAYYMTCC
                     : *   ***...* *****  .  .:. *   ******    .   .* ** .

35              YYKTKCTKYYWWACYGCTCRCKRSMRKCWGGTTWTCRCTRCAGMAKYKMGTGRRCCCYKM
51              CTTCTMCKYCAACTYRYAARTTAGAGKYATRCCAWYRYKRMWAMTKCGCAAARRYYMTKM
                  . .  **   . *  :.* . .  *         * .*  .*:*   .:.**    **

35              SAKTYGRRWRAWYYACAYTCCCGTSYRRKTSKGWWYKGKMYWMRMKRSMRSRRCKMYKYR
51              GRTWTRRRWRWTCCGTTYGATATCSYGGKYCKAWTYKAKCYWMGCKRSMRSRRAKMCKYA
                . .   ****    . :* . .  **  * .*.* **.* ***  ********.** **

35              MRRYRRYYCYCGTCRYAKAGCCMYWACTRYKYRYYRTTAAKAGWYCRRCKMKRTMCYKKR
51              AGATRGCTMYYACTACCKRRTTMYTTYWRYKCACYRCCTTKMSACYRRMTCKRYMMYKGA
                    *    * .    .*    ** :  ***   **  ::* .   ** . ** * **

35              KAKYYKCKYGYMMCRYWCWAACRWKGCGCAGARKKMRYYWRGMMRWMGTAGCACGCCGTK
51              KMKTTKSGYRYMMYGYAATMWYATKRYKTCAGRKTMRYYTARAAGWMRWWRYRTAGMRKK
                * *  *. * ***  * .      *    ...**.****      **       .   .*

35              CKAKACKRMWKKTYATTMKK
51              TKRKRYGGCTKKWTGKYMKK
                 * *      **  .. ***
```

The pairwise alignments of two accessions 38 and 42

```
38              KYYYKYSRTTGCRMYMYSYRAMSRRGTRWGGRKWMRRYMYRMWYRWRYYAMWRRKCTGTA
42              KCYTKYGRCAATGMYMTSYRWMCAATYRWRRAGWMRRYMYGMWTRTAYYCCWRRKTGAWG
                * * **.* :.  *** *** *.    **    ******* ** *  **. ****  . .

38              AYGWWGYWYRKTRRWKWYKWCGRYMSRMMYRWRKTGYMARWTRATYRRAKKYYYKYYWWR
42              RCRWARYWYGKARRWKWYKWARGCCSGMMCATGKYKYMWRWWGWWYRRGKKYYYKYYWWR
                   *  *** *:********.    * **    *  ** **    ***.***********

38              SRWYAMMGRRWTTGCTMYYRMYAGTGYAAWYMKRMWYARYYSYTCAKKCTCCTGMRYKGR
42              SRWYMMMKRAAYAATCMYYRMYWTCATRWWYMGACWCWGYYSYAATGKYCMYCACATKRG
                **** ** *   :.  ******   .   ***   *   ****:.: *     .   *

38              WTAYRKTGCWRMWRWWMRSSWCYKWYYRRGCTAYYKTRSKWYGMCYRYYTRSRMKMYKTM
42              WWMTGKAAAWGMWRWTCASSWTTTWYYRRATWMYYKGASKWYRMTYGYYGGCRMTCYKCM
                *    *:..* ****   ***  .*****.   ***  **** * * **  .**. ** *

38              AATKGTCRGGYRYWYYWKGGTCWYCYRAKMKCGRKYWTRRKGYMKMYRRRYRYRRRRTTC
42              MWKTRKMAKRYRTATCWKRKYTWYMYRTKMKMARTTACRRKRYMKCTGARCAYAGGAWWY
                  .. .    **    **    ** **:*** .*.   *** ***    *  *

38              TKAYMKCGTYAWAATGARYYMKTTGTGRKYMCTGYWYMYARWRRTCCGRRSTRMGCKGAT
42              CKGYMKMKYYWTMRCATGCCMKYCKWRRKYATGAYACMTRGWRAWYYRRGCCRMATKRRY
                 *.***   *     .:   **     ***   .*  *   **     * . **. *

38              YRKWMYGTYMGSYRRYGKRRSKGRTCCASMYWGTGMGTAYYMRYCRYYRSKCAARRKKSR
42              CRGWMTKYYMACYRRCRGGRSKAGYYYMSACTRYRMTAGTYMRYTAYYRCTSRTARTKSR
                 * **   **..***    ***.     *      * :. ****  ***... : *.***

38              RKRGMKYRAWRYRYAKYRWYYSRWWMKYMRCTCYARCTTKMRRARGTKRATTGWMCGCGT
42              RKRRMGYRWAACATMGTATYYSRWWATYMGYWTCGRTAYKMRGRGRYGGWKWKWMYRTTG
                *** * **           ****** .**     .* : ***        .  **

38              TKAKCGYWTYYYTCYYYAGGAYRTAAYKATAGTMMRAYYCMKWKWKRRYMWYACWRTCAG
42              CKMTYRTWKYYYGYCYYRAAGCGARWCKTKRSAMMACCCYMKWKTKRRCAATWMTGKACK
                 * .   *.***   ** ...  :   *:. .:** .   **** ***        ...

38              GCYYKRYRYSTGTATTRRMRGSYSGKGAMYRAYCCTYTTSCRKYMRATGGSYWMYMKKTG
42              KSCTKGCRYSGRYWCYRRMRRCYSSKACMYGRCTTATCACAAKYMGWGAASYWMCCGKAA
                 .  *  ***      **** .**.*..**     :  :.. ***   ..****   *:.

38              CRKKKRRKKKYTCTTSRKRMRRWKYARRGYWYMKAAGYGKAGRCKCTGAKTKYYYKMYRY
42              AAKKKRRKGKCWTCCSRKAMRRWKYMRRRYAYMKRRRYKKGRAYKGCAGKYKTYYKMCAY
                . ****** *     *** ****** ** * ***   * *.   *  ..* * ****  *

38              GGCSGYKKKKAAYGTKTMKTKKKCGCAGKGARTTTRRRSKTYKYKRKYRARYKKRGAATC
42              KRSGRYKKKKMRCRYKKCKKKKKSRYRRKRMGWCCAGASKYYKYKRTYRRRYKKGRMCWY
                  .. *****     *. *.***.    *         ** *****.** ****   .

38              AYACCAKGTRYKRKKRKAKGTKCKGTYRTSKKRKCCCAGTAKYTCCCRYMTYRCTCARKM
42              GYCTYGTRKRYKRKKRKRKRYGMKCGCRYGKGGKYMYGAWWKYCYSSGCMWYRYYYWRGC
```

```
                     .*.  .. .******** *    *   * .*  *   ..  **  ..  * **    *

38                   AGRKAYKYKKKYRKKCARTRTTATAGTTYKKKMYCGTRKACGMTYYAYKYGTWAMYYKAC
42                   WRRGRYGCKKTYRKKYGGWAGAMKRAKWYKKKAYYCKGKTYCAKYYRYKYKGTCAYCKTT
                       *  *  **.****  .    : . .. **** *  . *:   .** ***   . * *:

38                   CSTCGRYRYSWTRAKWRTSWTAYRRATAAYWRKKWRTTCRGTCRKTTYKKTGKRTMYCRM
42                   GGCTAAYRYGTKRCKWRYSAWMYGGGARGYWRKKWRYKMRRGGATYYYKGYRKRWMCTRM
                      .  . ***. .*.*** *   *   .: .******* . *     .  **   ** *  **

38                   RRWSYKKRMYKTKWTWYGWACCRGGAKTAYRMKYWWARKKKYMCAATKAKYRCATCYKAM
42                   RRWSYKGRMYKYKTKWYKWRYYGKAGKYWTAMKYTWRRGKKYMYWRKKKRKYRMMYYYKCM
                     ****** **** * .** *      ..*    *** * * ****   .* ***    **.*

38                   KRGSCRKYRRWKKCGTACTCAYCMRWCRGRGAKYWAKCGGGKMRCAACTRGWRAKTRTCK
42                   KASGYRKYRRWKKTRYGYWYRYSMRWSRRARMTYWGKSTAKKMRYGTYWGRAGWGCAKAT
                     * .. ********    .    *.***.*    .**.*. . *** .:          ...

38                   CRKMARMRACWRSWYYTACRWTGTYKTGRRRTCYKCYTKRKYYARRKKMWRMYMKACCCG
42                   YRGMWRMRRYWRGWYYWMYAAWRYYKWKRRRWGCGTCGKAKYYGAGKKCAGMCMTRTTAA
                      * * ***  **.***        **  ***        * ***.  **   * *.    ..

38                   TTACCACYCGGCCKCTYKRGTACAGGACCGRYYMGRWMSACRSGCCAAGGWGRMYTGRKR
42                   CCCATWAYMKATYKACCGGRYWYTRRWYYRRYCCRRTCSMMRSTTTRGTRWKRACKAGGR
                       ..  .*  .  *.         :      **   *  *  **     .  * *  ..  *

38                   WKMSCKCCRRYRAAKCYSYAMKARRKGMRTMYCTCAWTGRRRMRCMGKACKTKGYCGYAT
42                   AGCSYTYAAGYRRTGTTSYWAGWRAKTCGCACMGMMTWARRRCRYARKRYKKKACYRCRW
                        * . .  ** :   **    * *             .*** *   *  *.*.

38                   CYKKMMGCTYGWWYRYTKGMGCGGCKKRTTCAKMGTCCRRCRYARCGAKRYYAGCCTCMA
42                   SYKGMCATAYATWYACYKRMKSRRMKKRWKTWKAKYMYAGTGCCRYRRKGYCTAYMCMMR
                     .** * . :*. **    * * .   *** .  *          .*   * * :.    *

38                   GTATKRSGAWKRKRKCGKSKWYGTKAKGKKTKYTWYTKTYYKYWKKGTGGACRWMKWKTR
42                   RWCKGGGACWTGKRKYRKCKWTRWKMGATKKKYCWYCTKYTKYWKKRCRRWYRWMGAGCG
                       ..  ...*. ***  *.**   *   ..*.** ** ..* *****      ***

38                   RCACRCWTYRCRCCTCTSWYKYWWYGTAGAACCTGMCCRMGYCCTWCYKRKAYCRKKKKK
42                   RMMMGYWWYRYAMYYYCSTYKYWWCKKTKCGTMACMMYRMRCTTATGCKRKCYYGKTGKK
                     *      * **        * *****  .: ..  : *  **     :    ***.*  *. **

38                   RWKYCYKYMACCTMTCYGRSWGAGGGCTKGAYYYKRKRRYCTKWYGCGAGCRKGRKMCYA
42                   RWGTYYKTMGMTYMWYCSGSWRRKRRYYKKTCTYKRKRGCSYKWYRYRRRYGKTRKCYTR
                     **    ** *.    *    . **        * :  *****  . ***        * **

38                   ARMCKGTGRYWYTRCKKYRRKGTRYTKRTKAGGKAKYYCRWRTAYKAYAYRCGRYYYRGR
42                   MAMMKRWKRTWYKRTKGYRRKAYRYWKAWGWRRKRKYYGGTRCTYKRYRTGYARYYYRRG
                       * *    * **.* * ****. ** *       * ***    * :** *     .*****

38                   KRYKRYYRKGYKCKGAACKAYAACMGRYAACRCRWAAGGCTKTRRAYRMRYYKKCCCGAK
42                   KRYKRYYRKKYKYKRMGYKWYRRTMSRYRWYRMRWWRKRMYGCGARYRMRYYKKTTMRRG
                     ********* ** *   . * *   *.**   * **          ********
```

```
38          TCKRAWTGKSGKKRRKYKCGYRRYYYKYYMMYKKAWYAKRYAGGSTWRRAKKSKRAWYKY
42          KYKRWWKRKSRKKRRKYGYRYRRYYYGYYCCTKKRWYRGAYRKASWTGGMGKSKAWACTT
            . ** *. ** ******   ****** **   ** **   *  .*      ***    .

38          YMKCTKYTTTGAGCKAKKATTKTATTRACKKMWATAGYRKMYKKGGACKTCGAKMWATRC
42          YCKYCKYYGCCRAYKRKKRYYKYCYKGRYKGCWTACKYRGAYTKACGTKCTAWKMWRWRY
            * *  **     . * **   * . .   *  *::. **  *.*. . *  . ***  *

38          GTKAKRYMCATAAWCKTAYKYRKMYWWYAGGTYRGACYMMKGRKCMKYYKRRTCYGAKKS
42          KCKMGGCATGCGGTYKGRYKYRKMYWWYRAAYYRTRYCCCGRGKMMGYYKRRKMCATKKG
              *      . ..  *  ********** .. **         * * *****.  .:**.

38          WYRCTMCYYRKYCTTTTKGCTTWWRRKRKMTTCCTRKRAAYYARYYYTAAAMAGTYKTAA
42          AYRMWCYYYRKCMYCYGGATCCWAARKRKMGGTYWRKRWRCYRGCCCWGTGAWRWYKGRR
             **    ****       .   *  *****     ***   *      .:.    **

38          RRCCTAYRYRARGAMCCAKTGYAAGARGYKSCMMRWRAKYTRGCYGCRGWMCGRTTKRKY
42          GRYTYGYGTAMAKGCATTKATYRMRGARYKSTMMRWRGKCGAAYCKMGKTMMARYKKAKY
             *   .*      . . :*: *   .  *** *****.*   .       * .* .* **

38          RYKRMTAWKYYRTGAAATTGMGCRTTGAKRKTCRKGARTKWMKASGCCSTYTGAGTSRAC
42          RCKGCARTKYYRYRMRCCAKMRYRWWKGTGKCTAGRRRYKWATGGRMMGKCARTCYGGCY
            * *  :  ****    . : *  *   .. *      * ** ...   .. : :  . .

38          CTTTYACGKSTACYRMAATMKMRMRKRTRYYCCRCTWYRGRYGWKTTTSYGTTKKKKTAW
42          SYGCTMTRKCWWYCACCTAMKCGCRKRKRYYYARAAWYRRACRWKKYWCYRCGGKKKCGW
            .       *.      .::**   ***.*** .*.:***    **.  .*    *** .*

38          CYMWTCCAAGYKMCGATCGMTRTYCYYTCYRTGGTTTCMGGTACGWGGTTYRTRRRKRKT
42          YYATYYYGRRCKMYKMWYRCAACTMYYYYYGCRACCKYMRRCWTAATAYKYRKRRRKRKA
             *     .   **       :    **  *   .  . *     .  . .**.******:

38          TKKACTCKGCATRRAGCAYKYKYKTRCAKGGKAKRKTYARCWCWSAGTAKSYTCAKKTCR
42          AKKWYCTKSYWWRRCKYWYKYKTGCGTRKTAGWGAKWTGGYTMWCGKCGGSYYYTKKYYG
            :**    *.   **.   ****      * .    *  .    *..  . **  :**

38          CWAKGTACKKYWKWCCCAKCAAGTCTCYYGAATGACAGKYRYYSRKMCKKMAMWYKGRGY
42          YAGGTCGTKKYTGAYTGCKAGRAGYYYCCKWRGAGTMRGYRTCGGGCMKKMRMTYKCARC
              .   . ***      .*.. .          ..    **  .    *** * **

38          KMARKCCTACWMTTGTATGAYGGCYATTCTTATMYYRCGKKTKMWWWTKRGMGKRKYGRW
42          KCRGTTTGTYTMYWKAWYRMYAAMYGWWTKYTACTCGAKKKWKMWWWWKRRMKKRKCRRT
            *   .   :  *   :    *.. *.   . ::    . ** ***** ** * ***  *

38          ACGYYGYYRMKCAARTYKTCCAGAYYATCTCYYKWYKWMMACCTRCTKKTYGRRCAAATR
42          RMKYTRYYRCGYRRRYYKCTTRSMYYRYYKYYYKWYKWMMWMYKGMGKKCYRGAYMWMYR
               *  ***     * **    . **   . *********   .   ** *        *

38          KCATGTRYTGKAAYCGGKGYKGSKCCRKRYCTYKAGAWAWWRYCGGMMSCACCGYKAKRW
42          GMRYKKRYYRKRRYYRRKRTKKGKTYRTGCYGTKMRWWRWWRYSRRMMSMRMYRYKGKRT
                 .**  *  *   *  * .*  *.     *   * ****.  ***     **.**
```

```
38      YYKRWRYGYKCTACRGRRARMKMRSTYGYRRCACWKYGYYKGSMGKKRCMYRRRYGGRYR
42      TYGRWRYRYKYCCMGARAGGMGCRSAYRCGAYRYTKYRTYGRSMRTKRTATARGTARGYR
         * **** **  .  .* . *  **:*        **  *  ** .**    *  .  **

38      GRYWKKYYYATRARAWRMGAAYGTTKSRAMMYKACKGGGGRAKKTRYTRCARRMMMKAYW
42      ARYWKKYTCMCRCRMWRCKGTTRWWKGARMMYKCMTKKKRRRKKCRCCGYMRAAMCKCTW
        .******    *.* **  .:    *.  ****. .    * ** *     *  * *. *

38      RYKKKCCTCMKYKYGYTCGMKKKSYCCWMTCRCTRGCKMRRAATYYGTYWARCAGCRKTA
42      GTKGKTTCACKYKYKTYMRMKTTGTATAMCTAYKRRSKMRRWWYYYRKCTWGYMRYRKYR
          * *   . ****     **... .  *    .* .****   ** .        **

38      GTGATARGGGCAKTGTYKGGKCCYKYGTKTSGTTTWCYTTCGGYKKTTACKMSKAWWSTT
42      AYKRKRRKKRTMKYSKYKKSKYMTKYACKKSKKKWTYTGYTCRYTKYWWMKMSTMWWSYK
        .   . *     * ..** .*   **. *.* ..         *.*    ***. *** .

38      SYRCGYCMYKATAATAATYMRWTWYTGAATTYCWYWWTTATATGAGGMTAMKTTTTGCGG
42      SYRYRYYMYKRWRWGCCYCARAATTGAGRYACATTTWWGCCCKRRRKMCGMGCYYYKYRR
        ***  * ***     ..   * :   ..  : .   *  . ..    * .*

38      CATTKGGARYRYTGACCKKKMYWGKTTMKGCMKMCRRYRYGGTRTAKGGYTAGYKRKACR
42      MTACGKRRGCRCKRMYMGKKCCTRKYWMKATCGCYRRYRTRTWRAWKAAYWRRCKRKRSR
         ::       * .     **    *  **.     ****    *: *..*    *** .*

38      AGTGYGAAMKKCGGATCAYRSRKWYATACGKTGCYRTRRAKMCTWRYYCMCAATGCYWGA
42      MKGRYKRRMKKYRRGWMWYACRKTCTYWMKKAAYCGWGRRKMACWRYYTCYRRKTYYWKW
            *   ***   .   * .**  :    *:.     * **. ****     .  **

38      TCCCYKTRGCYYGKKYCKTYGKTGGCTARTKKMRRYKWKKCTCTGGAAMCCGACTTKKAA
42      WYMYYKGRRMCYRKKYMKYYKKCARYYTRYKKMRGYGAKKTCTYSSRRMATATAACGKGC
            ** *   * *** * * * .   :* **** *  **    ..  *. .:.:  *..

38      KCGMACGRGTTATMKTKCTGTYYYYRKTAYSKRAGGWTKKTTCRTSYWYYYYGYYSRRYK
42      KAKAWYARRKCRCAKKKMYAYYYYCGTYRTGGRGRRTAKKWCMAASYWYYYYRYYSRRTT
        *.    .* .    *.*  . ***  .   . *.   :**    :******* ***** .

38      KYRRAMYTARWYGRWKKAYMGGYKCTCKWRMTWKKRTCWCGTRMCCGAAAKYKWCYCTKC
42      KTGGMMCKGATTKGTKKGCAARCKTWYTTRCYWKKRYYTMAYRAMYATMWGYKTTTTCKY
        *    * ..      **.  .  *   . *  ****    . *   .:   **     *

38      YKWKGYKKGGMTTAYRKCMKKRWRMKYYMYCKYKTKRCTTTKGYTTTTKRRAGCAWCYYG
42      YKWKATGKKKMYYRYRGMMGKGAACGYTMCYKCGYKRYKKYKKYAAAAKRRTTYRTMTCK
        ****.  *  *   **  * *     * *  *   ** .. * *::::***:

38      KTTASKKARWAGTTMRKMKTWTKGCCTGTTKCGTYCKTCRYYKAKAYAGKTCAAGKATTT
42      GKYRSKKWAACAGGCRKMKKWCKKATCAYYKYKCTYKYYGCYKRKCCCAKYSMGAKCAYC
         .  ***   ..   ****.* * .  .  *     *    ** *. ..* . ..*.:

38      RYYACCTATYWYCCGKRYYRRYAYYTTCGTRWTGMWKMSKKCRRWAWWKRYYCTKGYCRR
42      RYCGAYYWWYWYYTKKRYYRRYGCCCYYKYRAAACWKMSKKTRRWRWTKGCCAGKRYSRR
        ** ..    ***   *******.       * :. ****** *** * *   . * *.**

38      GGGMCAATRKAARKMMKTKACAAKRCYKCCCYMRMKKRGAYTKRYCAAKMKARWKWYWRW
```

```
42            RRRMTGMWAGRWGKCMKAKTYGGKRYYKYYYYMACKKRAMYWKRYMMRKMGRRAGTYTAW
                 * .       * **:*: ..** **   **  ***. * ***   **  *   *  *

38            KRRKGAKATTTGGWYMCKKAAAAATKMATSYKGCKACKTMAWAYWRYYYKSRYTGKGCTA
42            KARKTRKGGKYTRACMYKKMWWRRKKMMKSTKRTKRSKYMRWWCWGYYYKGACYRKRSYW
              * **  *. .     * **     .** .* *  * .* * *  * ****.    * .

38            AGKKGTMYKYGAYTGCWMMYGYYKGTCWARCGTCAGTCCSMYRGGKRKYAMGTTMTARCC
42            WRKKKYCTKYRMYYRYWMMYRYYGRWYWWRYRYYWRKYMSATGSAKRKYGMKAAMKRRYY
                **    **  *   **** **    * *      .  *   ..****.* ::*. *

38            ACCARTGYYRMYSCACCARTAKKKAWCRKYRYRTTYRKGTYKRKSYRARKKKARKCRYKG
42            CTTTRYTTCRMYCMRYSRRKRKKKCWYAKCRCRYYYAGKKYKRKSCGRRGKKRRKTGCGT
              . :*    ***.   . *. ***.*  * * *  *   .*****   * ** **

38            CYWRCRARYRRRCYYCRYKKSYKKAKYKWGARKGCTYRYTYCKAAWYTMRYKTYCMWMRT
42            ACWRYRWRYGRAYCYMRYKKGYGKWGCKAKRRKKSKYRYWYMKMGTYAARCKWYYMWMRW
              . ** * ** *   * ****.* *   *   ** ..*** * * . *: * * * ****

38            KMGTGCYRATKTYRYWKRTGYGGMKRCATGRWRRKCAAKTTGTGYTCACYYKKCARSTTT
42            KCRYATCACCTGCGCAGAYKYRRMKRMRKRRWRRKYWWGAWAAKYGMGMCYKKYMGSGAG
              *   .   . .         *  ***  . *****    : .: *  .  ***   * :

38            GAYTARYMTRKKSKKWKRRYAATTYWKRTTGKKRWKMCTGGMMWACTCKYMKKKAMRKYC
42            ACTYMRYMYRKKSKKWKGGTRMYYYWKRYWRGKATTMMWRSACTTYAMKYMKKKCARKYA
              ..   *** ********       ****    *  .*   .   : : ******. ***.

38            ACKYYMCKKKYKWKTACKKCATGGCMYAWKYRKWRKYCRWTSYGYYTKKAAAGWRWKGAW
42            GYKYYMMKKKCKWGYRSKKTTCTATAYRTKYGKWGKYTRWKSCKCTKKKRRWRTRWGRTT
              . **** *** **   .** :  .  *  ** ** ** **.*    .**     **  :

38            CGRRGATARMYYKYYKAYGAGKYGGYAMYYMTKRKTTKATTCGYRYYWYKSYKTSCTKRT
42            YRRRRWYRRMYYKTCKMCTGRTCRRYMMYYAKKRKGGKMYYYRYRYYWCKSYTYGYYKGK
                **    *****  *   . .   * *** .***  *     ***** ***. .  * .

38            MGCARTMYYRAAACSYTTTYKGTTKYYYMYWTACCACRKGKCTCRACMRTYRTYKKKGKA
42            CKYRRYCTTAMTGTSCYYYYTSWAKYYYMYWKWYYMYAGRKTATRTYMRKYGWTKKKKGW
                  *      :. *    *.. :*******.        * : *: **.*   ***

38            ARRKGKTAKTWKKGRRKKKYTYTTKMRSWAKCRRKKRWAGARRRYYRAGGRYRWRCTTYK
42            RRRKSKAGKGWTKRRRKKKCWYYWKCASWWTYRRKKAWWRTGGGTTGRRSRYRARYWYYK
               ***.*:.* *.* *****  *  ** . **** *  :        .*** *   **

38            YTCCTYKGYYYCYWGKYKRYYKATTARRKKKKMATGGYWRWYWACCCAAKRYKYMACKY
42            YYYAGTGSYCCTYWAKCGAYYKRYWMRRKKGGGMGATKYWRTTWRTMMCGKGYKYMMMKT
              *  .   .*   **.*   ***     ****   *.:  ***  *     ..* ****  *

38            KRKTKYAYRYTKAKMYAYAGAKRRYCRAYYYCMYATKKTRCKCTAAGGKGMMRRRCAGTT
42            KAGCKYGYRTKKGKCCGYWKRKRRCYGRYYCMMYGWKKWGYGYYGGTATAMCARGYGSKC
              *   **.** .*.*  .*   ***    **  **. **      .. ...*  *  ...

38            YYCGARKRRCGTCKSYMAMAYSAGKYYYYAGGKKKACCCYWTGGGKKYYYTTCMKRKGTT
42            CCMKMRKGGATGAKCCMTMWYSWAGYYYYMRRGKGRYMYCWYKRKKKTYCYKMMKGKKYY
```

```
                     **  .  .*. *:* ** . ****    *      *    ** *  . ** *

38                RYRRRGWMAYGKRKYYCTWSWTRARYYAAWRTYRSSRMCRYKKMRYWYMKKYTTWYRSRW
42                RYRRRTWMGCTKRKYYYYTSWGRRRTCMWTGAYRSSAMMRYKKMRYWYMKGCYWTYGGRW
                  ***** **.  *****   ** * *      :**** * ***********     * .**

38                YYKMRKKCKKCACTSTGRWCTTCKYKGKYYMMYKTGKCRCRYTRTRCRWYKMGGGAAASR
42                YYKCRKKMKKTGTWCGRRTTCYMTYKRKCTMMYTYKKTGTRYWRKRYGAYKMACATMCGA
                  *** *** ** .  .  *     .** *  ***.  *   ** *.*   ***. .: ..

38                MKYYKKRRGYMRTTYYRKGKRRGYTTGRYYATGGWCKGYRACWKRKRSAARGATTMGGCK
42                MKTTKKRARYMGGCYYAKKKRRRYYCRRYYTCKRWYTACGTYWKRGRSMGARWYYMRATK
                  **  ***  **   ** * *** *   ***:   * ..  : *** ** .     * . *

38                RKKARTRMCKTKSACCGYYYATWTKAGGCKTKTAAGCKKTATCRWRKWYAKRGAKKKYKY
42                RKKWRYRMMGCGCWYMKYCCWYTKKWKRMKYTWMGATKKYRYMRWGKTTGKGRRGKKYKY
                  *** * **    .    *     .*    * .  .. **    ** *  .*    *****

38                RY
42                RY
                  **
```

The pairwise alignments of two accessions 38 and 43

```
38              YCCGRWRRTCCYAACYKAGYWRYARKWWWYTGKWMRTRCCKKKWRYRRRRRKYYAYRKKW
43              CYYRRWRRCYMTTCAYKMRYWRYRGKAAWYGTGWMRCRYYKGKAACAGRRRKCCMTATKA
                    ****    :..**  ****  *  **   *** *  * *     ****     .*

38              KYYYTRRTYMMMWRCTYRWRMYRMKWTRMWACCTKYKRAYYRYRTATTSRWTRKWKGKTR
43              KYYYYAGGCCCMWGAYYRWRAYAMKTYRAWWTTYGTTRTYYRYRAGCAGGWWRKTGKKKG
                ****       ** . **** * **  * *      .*:*****:. :. * **   *.

38              YCYYRRCKGWAYYAWYATWKRYKCTAYYYYYWYKRKRYSCSTRYRSMMRWYRYGYMKTKC
43              CAYYRRSKRTWCCWTYRAWKRYGYCRYCYYYWYKRGRCCYCWRCGGCCAWTGYRYMKYGA
                 .****.*       * :****    * ******* * . . *  .   *  * ***  .

38              GKSRWWYARYRMTAGGYCYMMYAMWKKMYCWWMYWWSMRKKGCGRKRYRWCRYWKAKRYY
43              TKCGTTYRAYRCYRRSCTCCCYRMWKTCYMWAAYTTCCRGKCTRGKRYRWAAYWKGGGCT
                 *.   *  **    .     * ***. * *  *  . * *    *****. ***.

38              YSMGRMAWMATCATWYKRGTKWMTKRWARSWRRKGTWYTKRKYYYRRMKKAWGKCMKYMK
43              CSMARMRACWYTGYWTGGRYKWAKGRWGAGTRRKAATTYGRKYYTRRMKGGTRTYATCCT
                 **.**      . *     ** . **. . ***.:    **** **** .  .  .  .

38              SYYYRWMKSRKKMYCTWMYKMYYYYKGGYAMRRYMRGAMGGGGMYRTGWCTYTRRWWCYR
43              STYYRWAKSRGKMYMAWMYKMYYYYGRKYRCRATCGRMCRKKKMCRYRWSYCKGRWWTYG
                * **** *** *** :*********   *  *           * *  *.  . *** *

38              YYYCGTRYWKWWSYRCTGTRCCTCMACYAGGCYKWAGATMCGWMRTRKMTTYKGCRYYAT
43              YYCMAKGCWKAAGCGYCAYRMAAAMGTCMRRMYKWTKGGCARACGKRKMAGYGAARTYMW
                **  ..  **  .    . * .:.*.      ***: .  .    .***: * ..* *

38              KGMKMRWMCAACGCYYSCYYKGWRCAGCCWRTCTATCKRCYKYTGAKGKCCKGMCTKTTR
43              GTMKMRAMAMWTRSYTSMTCGRTGYMRYMTRWTKRYTKGYYKYCAGGSKTAKKMYYGKYR
                  **** *.    .* *             *  .   *  *** .. .* .* *   . *

38              YYWGATCTCTGAAKAMTKKCRTTGAGARYTKKYMCCARCCASRYTTTKKRTCTKWTKKCR
43              TCWRRYYYMYRRTKMAKKKARYYTGCMGYYKKYCTTGAMYMGGYYYYKKRWYYKWKKKTR
                  *         :*  .**.*   .   * ***   .    . *   ***   **.** *

38              TTKYRKGYKAKCGTMCKKGKRKKKACSGTKKWCRRKKAYCGRKWCCTGKYWGCATTRMYA
43              KWKCAGRYKGGAKGATKGAGRKKKGTSSKKKWYRRKKGCMRRKWYTWKKCATTGCYRMYM
                . *    **. .    * . ****. *..*** ****.   ***    *    .  ***

38              KYMRTCAYKYTCKYYCGCCTYKCYTRYYRCARKKKYKTCTTAAYSRYGTTRKMGGCTCYW
43              KYMGWTTYKCCTKYCYTASYCKTYYRYYGYGGKGKYKGMKYRRTCRYKYKRKAKKTCACA
                ***   :**   **   ..  * * ***  . * ***  .    .**  .**     .

38              TYYTYTTTKYKWRCRYKKKCRTGCYKTYACYYGGCWYWACCKRRRKTAATAYRMRWRKTG
43              YCTATCCYKYKWAMACKKTYAWRMCKYYWYYYRATWYWRYYKGGRKYWMYWTAMGTRKAT
                   :    ****    **.      * *  ** . ***   *  **       *  **:

38              RYKRKKKYCRWMMYKYMKRWAGMKYYTATGAYRRWKKYATCAAMTTCMRWCACTCMMKYA
43              RYKRKKKTMGTCMYKTAKGARRAGCCCCAACYRRWKKYGWTRRMCCGMRWSRYKYCCKCR
```

```
        *******     ***  *         .:..*******.    *   ***.  .   *

38      AATGMTYGKRAAKTTRARAAMKAATTCWSKKKSACCCWGCRMMWRWYRTTKRTYWGKYYK
43      MGGRCWCAKAGTKCCGRGGGAGWTCGSAGKKTSRTTMTRTRMMWRACGCWKRACARGTCG
         .     .* .:*     ..   :  . .**.*       *****     **:

38      TCYKATWYTTATGTTYAAAMKYTAKYATGYKRYTTARTYKYAAYWCGGSTRCKTYKGKKG
43      WYYKGATTYKMKAGWYTTTAKYCTKYMWKTGGCCYMRCYTCRRTTMACCKGYKKYKKTTK
          **.:   . ..  *::: ** :**          * *.      . ..  *.** ..

38      GGCAGTYSCCTCGKGYKTCMWWGGACKGRRWRKKTYYAACCYKGTARKKGGGGAWAWKKT
43      RAYGRYYSMYCYKTRYTCAAATTAWYTKRRWRKKCYCRMYYCKRYCAKKAARKRWRWKKG
         . .  **     . *. .    .  . ****** *      *  . **..   * ***

38      TGATGAWMCRMKKGYYYYRKAKKRYKRSGCTAAAATRYKTRWKKTYCAAKCCKWRMAATW
43      GACYKWWAYRAKKRYYYYRKRKGRYKRCRAATCTRKRTTWGTKKYCMRCKTTKTRCCGCW
         ..   *  * ** ****** * ****. .::.: .* .   **    .*  * * .. *

38      KRYKKKRTWCGRWRGATKYYGAKRRKSAMMMRYWAAGKKYYYKRAAKTRTMTCGKKSWGR
43      KRTKKGRYTMCRAGTTWKYCKMKGGKCWMMCGTWWMAKKYYCKATGGYRCCKYSKKSWTR
        ** ** *    *   : **   *  *. **   *  .**** * :.  *  . .**** *

38      RGMYKWTCRTTCCTWGTYAAYRTGKARTTSSRTYYRWMYYKWRTRWRAWKRYWYRAGTTM
43      GTMYTWATRCAAAAWTCYMWYRCKTRRYYSSAYYCRWMYYGTRYGWRMWKRYWYGGTCCC
          **.*: * :..:*  *  **  . *  **  * *****  *  ** ****** .

38      YYYKMCMKYYGKYWCRCGKGCMGRWKKKKMRCTMAKRMKKCTATCRKRRRGKKRTTRWMC
43      CCYKCGMTTYRGCAYRASKKSCRGWGTTKARACATKGCGKACTAAGKRRGATKGAGRWMT
          **  *. *     *..* .   * ..* *.  :*   *. ::. *** ..* : ***

38      STGWKTATCYKYKKTCAACMYGCTAMYWWYCWKCRKRYAYYYYRATKKGYKKMWWKGMCK
43      GCRWKCCYYCGYKKYSGGTCCAYYWMYWWYYTKYGGGYGCCCCRWKKKACKKMWWGRMYG
        .  ** .    *** ...   .   *****  *    *.    * .**. *****  *

38      RGWATGACYKRMCWMTKWRTAGASYAMYYKKWYCKKGAACGGRCGCTTKKKARCATYTYY
43      RAWGARRMTKRCSWAYGWGCRSMSYRCTTTKWCTKKKTGGATGMRYCGGKKMGMRGCGYY
        *.*.:    ** .*   *   . **    .** ** :. .        **       **

38      KAACCCRAGAACGKGCTGTKTGKAGATRWCMMRAYRWRYRYGYKTRTRRWYKMMRTRKSR
43      KRMYAGGGRRRYRKKMCKWTGAKMRWYAWAMMRRTAAGCRYRYKCRYRRWYGMMGWRGGR
        *   .  .     *     . .*     *.***      ** ** * **** **  * .*

38      RMRRYGRRCKAGCCMAWKKYKCSRAGCKMMACGRATKCRYAGYYYRYWMRRTGCGRCCKT
43      RMRGCRGAYKGAYYCWTKKYKTCRRSYTMMRMRGWYKTGYRRYYYAYACRRAATAGTAKW
        ***      *..     **** .* . .**      *  *  *** *  **:. .  .*

38      ACCAMGACAMCYTWCSCTRMGAATAAGACRYAATCGCWYRMRRYWYSTACTACKYKRGAA
43      WMYRMTWYTAACYTTGYGAAARTYRGTWTACRGGYRMACRCGACTTGWGTCGYKYKRTGW
            *   : .    .    . :  .      .      *      . .  . **** .

38      TGCYGWCGTMRTYGCTCKRKACRTTGGCGTCRKGGTKTKMMKTGGMGKYRAARCRTACAK
43      ARMYRAMSYCGCCTTCTKRGRMRYYRRYTCTRGRRYKYGCATCTAARGCRRRGTRCRYMK
        :  *   .         **   *        *    *    .  .    *    *    *
```

```
38     YRMGCMRMMAKCAKCYGCAWKGRWWMKRKMASTRWCYYTTCCRCGKTYKCTCAATTYRRG
43     CACTTCGCCCKTTKAYRYCWTRATTCKRKARGGRWYYYCCATAYRKGYKTYYGGGATAGR
                .* :*.*  .*.     ***  . ** **  .    * **   .. :

38     TTTTRRKRAAKRYCGTTAACKCGYYYYRKCCCYTSTYTGCAYRMMKCWYKWKRCTARRMR
43     KACGRRKRRRKRYTTGCRGTGYRTYYYRTGTGCCSYYCTTGTAMMKTWTTTKATCGRRMR
       .:  ****  ***     .     ****.     * *   .  *** * . *   .****

38     KRYMWYACKWMCGCRYWYTCCSGARKGAKCRYYRCRKATRSYWARCRRRCMYMCGKKKKT
43     TRYMWYWMKTMTKSGCWYWMMCSGAKACKTRTYGMAKGCRSYWWRMARGMACCYKKKKKG
       .*****  * *  .  **   ... *..* * *   *. **** *  *       ****

38     KRKCRSRKRWGGYTWKCRGAKCTYWWSKKGKAGKRTKYACYRMGSGCTRYCCKKAWTKMC
43     GGKYGSRGAWRRCKWKTGAGTTCTATCGGRKGAGGYKYWYTRMRSRTARCYMTGRTWKAS
         *  **  *   .**  ...     .   *..   **   ** *  :*   .    * .

38     TMRWKWWWGYCKWGTAGRTKGCTCGATCTMYATTYACMARMCKKTYRATGAGYTWYGCTW
43     YAAAGTATAYYKAKCGKRWKRTWTTRYMYAYCYKCTYMGGAYGTYYRTCARTCWWYKYYT
               .* *   . * *          *. . : *.    . **: .    **

38     KSWKMAMYRTYTACGTTMTWRMKCCGGGYWRCGTTYKTAYRKGKYGRRRSYKYCAYRARR
43     GGTKMGMTRWCYRMTAYMGARMKMTRRRTWRTACKCKYGYAKRKYAAGAGYKYAGYRRRR
        . **.* *      : *  ***      ** . . * .* * **.   .***..** **

38     CYRYYYTGTKYTKRAATGGRAACAMMYCKKWTMRYRRCCWTTTCRRYAYYSGRAYKAKGK
43     YTGYCCCRWGYKKRRWWRTGRWYCCACMKKWCMRYAGATWYWCYRGCGCYGRRRYGTGRK
          *      *.**         .    *** ***  . *    *  . *. * * :   *

38     GGRRYMRYYYKKKYKMYGMTWRRCMTGMKKKGYCCWRTGGRTAYCTTTCTYMMYYKGWMG
43     RRGRYMRTYYKGGCGAYCMKWRRYAYRMKKKRYTGTAAARRYRYTAAAYYYMMYYKKTMR
          **** ***     * *.***    **** *    :. *  * :::  ******  *

38     TTWAYKWRGRSRTACYWAATCKRYKKCAGGGYKYACYWWKCTACCGTCAGGWWWGGAWTT
43     AGARCKARSRGAKRMYWTTGMKGCGKYGRKRTKTGMCATKYKRYSKYTWKKTTWRATWYY
       :    * *.*. .  **::  *   * .    * .    * .  .        * .:*

38     TGATYKMGATTTRKCKKCRYYYKSCMGGTACCACTKTATCWCYKKSRRYYYGTWSYTGTT
43     CAGACKMRRCCCRGMKKMRCCTKCAMKRYRYMGMCKCGCATACKKGAATCCKWTCYGACG
        ..: **     *  ** *   *..*      .  * . . . **.        .* .

38     AKCYCCTCWWSCTRRWKMYATCYYMRAWMYWYAYRTSATRKWTCTRKGAWARCRGCRTMA
43     GGYYTTAMWWSMCRRWKMYGAMYCAGGAMTTYMCAKSRYRKWWAWGGAGTTGTARMGKMM
       .  *  : ***  ******.: *   . *  *   .*  *** .   .. :      .*

38     RWYGACGAGCKRAACTTKCCWKTTCAMTTGAKACMKKYMRRCRYKTAKGGGGRGTAGYKK
43     RWCACAKMTMGARRAWYKYSTGYWMWMGGTTKGTCGTTCAAYRTKWWGRSRSRRWRRCKK
       ** ...        .  * .      *   :*.   .     * *    . .*     **

38     AGGACACRTACKGTCKTKCCTATTKTTAAAGKCTYRMYCCRACTKTCWRKRKKKKTGKTT
43     GATTGGTAKRYKTGAKKKMMYRCAKYWGRRSKACTGCCATGGYYTYTARKRKKKKKKKAC
       .. :  .  .  *  .*.*     :*  .  .*.     .  .  .   *******.  *:
```

```
38          TTCTCYGYAKAYTCAGATGTTWYGCYAAGTGRKACARACAWYRWMYCRRTRMACCTCAKK
43          CCTGTYRYGGCCCTGACCACWWYRMCWWKGAGKRYTGMYTWTGAATGAAGGMWYMCYGKK
                 * *. .   ... .  **       . *  :   :*          *     .**

38          CTTRAKYYTRWAGGCWMYRCTRAATWKYRSKYYYYYKATCTMTARGCKRARKWCRGATTC
43          ACCGGTYCCGWGACTTATATWRGGGWKYRSGYCTCCKRKYWMKRGKYGGRRKWYARWYCM
            .   ..*   *..        *.. ***** *    * .  *.       ***

38          WYRYRYCRYRYAAYKSGKYRKRSWYRKGRKKKTTGRGSCCYGRCRYRRCCGGACRYYWRT
43          TCGCRYMRYAYGGTKSKKCGGRSWCGKAGTGGYAARKGMYTRRYRYGRYTTRGMRYTWGK
                ** ** *.. ** *   ***  *.  .   :.* .   * ** *   . **  * .

38          GTYYRYRRCKKTTKKMWRKTMTYRYWKRMKAYYKAACACAGGMGRYCATGMRATYKKKYR
43          AKYCRYRGYKTCWKKMTGKCCGCGCAGAMKTCTKGGGGTGRRAKRYAGGAARCYYKKKYR
            ..* ***  *.  ***  *         **:  *.. . .    **.. . *. ******

38          TCTAYMMKKWYARCKWWWGGMSWAKRYKCTRMYKSRYYYRATTARCCKMMKRYYKYYTCG
43          WYYRCMMKKWYRRYKWWWTACSTTKACKYCRMYKSRYYYGGCCWAYMGCCKRYCKCYCTA
                 ****** * **** . * :*  *  ********* .         *** * *  .

38          GTTRKKKCCGCGKCAYKYRWGGYRAMTKWWCKKKGRTAGMYKRRTGYKKKATKAWGKMGA
43          KYYAKKKMYTYAKYRYTYRWKRTGRACGTAYKKKRRYWRCCGAAKCCKKKGAKMTAKART
                ***    .*  *.***           *** *        .  ***.:*  .*  :

38          CCTYYCRGCTWKATGARMCACRSYRRYRRTKMYTRGGYRYYCCWWMYTCTGRKRKRKGRS
43          TYKYYTAATATTGCRGGCGMYGSTRRYRRKKATYRARTGTCMTATCTGYATRTGKAKRRG
              .**  . : ..  .      * *****.*   *.             : *. * * *.

38          YSKWYRYRRATGCYTCTAYAGTTSCYMWKCAKSYCRCCTRATAARKYRCCTCSWYKYWWM
43          YCTTYRYRRRWKSCCMAGCGTYYGMTCTKTGGSYTRTTWGCKCMGTYRMYYYSTYGYWWA
            *.. *****   .   :. .   .    * . ** *    ...  .**    * * ***

38          ARTMKTMTKRRKRGCTCACATATGKYAAYKTARAYGGRRTGCRKRWCTGGKKACKYYYKM
43          GGCCKYAGKRRKGKMGMGMRYMYRKTWMTTYMARYRRRRKRTGKAWAAASKKCATYYCKM
            .   *   ****     .      *    .    *  **.   * *.:..**...** **

38          YKATGGCGAWKRYTGMAKSYTKACKKTKKRYRRRMYYKKRRKRWKAKRWWKKKRRTRKYT
43          CGTCTATRRTKGYAAMMKSCKKMMKKKKKRCGRRMYYKKGAGATKRKGAAKKGGGGAGYC
              :  .    * *:.* ** .*  **.***  *******     * *   **      *

38          GCTCAACKKKKAAAKRKKRKGTGKGKGARKWCKGMRAKAYCCCWWRWWCCGTATKRTTGC
43          ATATRWYKKTKWRMGAKKATAGAKTKAGRKAMKKMGRKRCYMMTTAWWAYTKGGKAAGCA
            . :    **.*     **  .. .* *..**  * *  *        **.  .. * :  .

38          KKMWKAWRKKGRYYYKSRYTGWYSYAGKRKKYCAGTGYGCRMYKYYAWRACYWRTYTTAT
43          KTAWKRAAKKRGTCTKGACYRWYCCWRKGKKYGRRYRYRYRACKTYGWGTYCWGCCYYRW
            *. **   **     *.    **.   * ***     *  *  * *.* :  *

38          GTKYYMGGYRRKWSAYYCAKRRYKGAAYTKGGKTTMAGTKMYRMCKGTRKKKKWKKCCGT
43          RWKYTASRTRGKWSMCTYTKRGCGTCGYKKATGCCMMKKKCTAMMKRWGKGKKWKKTTAY
              **  .  * ***    :**    ..*.*.    *  .*   * *   * *****  .

38          RKKKKRYYGRYYRRRKYKKAYCKATRCCAYRAACRKYMYAAGGKTRRGTRMRYYCGARKS
```

```
43                RTKTKRYYARCTARRKYKKWCTKCKGMYRCRRWYGGCMYTGTAGCRACCRMRYYMRRRKS
                  *.*.****.*   ******   *..     *      **:. .  *   *****   ***

38                KKRR
43                GGAG
```

The pairwise alignments of two accessions 38 and 44

```
38              YWKKYYTKYCKSMYCMACYYSYRGWWMSRRTRRRGRGRRYTMYGMSYKRRGCRCYAAMWM
44              YWKGCTCKTYKGMYYMRYTYSYGRATMSRRYRRAKARRRYWCTRASTGGAATATTCCCWM
                ***    *  *.** *   ***    **** **    ***     *    .    .. **

38              MMRKYWMMRKKRRMYYRYKMYRAGTATGYMKYRACTAWRWTCYRWRYRYGWSWRWWRKAY
44              MCRKYWMMRGKRACTCGYKMCATKYWWRYMKYRTTGTTRTGYYRAACAYSWSAAAWRKWT
                * ******* **     ***  :     *****:  : *   **    *.**   ***

38              RWYAACYWATCTYYTTRAYWRWWWGYKWRSTRWYYMKWRRWRRKAAKCRKGAGYKGYYRK
44              RTCRMACTMYYYCTCARWCWRWTTKYGWRCWRWCYMKWGGARAKGGGTAGRTATKSCYRK
                *    .         :*  ***   * **. ** ****   * *..     :. *. ***

38              KMYYYRKMKTRKWMCYWWWSMMWYMWRCGYGKCRCGCGWTRYMRTYTKGGGWYCTGRYMW
44              TCCCTGKCTCRKAAYYWWTCCMTYMWRYRYRKYRTTMKWKATCGACWTTRAWYYCTRTCW
                .     * . **   *** . * ****  * * *    *.    :  .  .**   *  *

38              WRRWYWRRKYKYYRRAAGWYMYMYYTRSRRMAAWKKKMRYRMTYCGWGKYRAYTCWKRCM
44              WRRWTWRRKTTYYRRWRRWCMYMYYKGCRRMMWAKKKCRCGCKYMTTCTCGCYAAAKRMC
                **** **** .****   * *****. .***   *** *   .*    .  .*:.  **

38              RYYWWMYKTYYWMRRKRRRYYRYRGAYMSRMMYKYGGYWTGGGTAYTKTKYYMCYGCYCY
44              RYYWWACKWCYWMRAGRRRCCRYGAGCCSGMACKYAAYTATRRWRYYKYKYYAMYTMCTY
                *****  *  ****  ***  ** ..  * *  **..* :     * * ***  *    *

38              MMRYRWMRYGWRCRRYYRTAGKYCWKYAKSRMTSYMYWRRYGTRYRRKRYWYMRYYYWSM
44              ACATGWCRYKAGMGAYYRATATTGAKCGGCRMWCYCYWRRYAYRYRRKRYWYMRYYCACM
                     * **      ***::..   * . .** .* *****. *************  .*

38              YMRMRWRSGYYRKTYRYKKMRWWMSMWYYWWKSGCRRATYGKTAARKYYACGYWRYYYRS
44              YMRMAARSRYTGKYCAYKKMRWWASMTYYTWGSRYRGWKYRKWWMAGYYCYRTWGYYYAG
                ****  ** *  *   ******* ** ** * *  *  .* *     **.   * *** .

38              TYKWGCAYKACCATYRRRCKGCTKKYYAARRRKKRMYYMKKAYMWYACWKCCCGCYYYRR
44              YYGTATRCKTYSWWYGRRAKATWKKTCGMRGGKKRMTTCKKTYMWYWMTKYYYKSCCTGG
                 *  .   *: .  * **.*.  **  . *  ****   **:****   *    .

38              RRYRWGKTATYGCATCCASRTYYSKYGARKGACMYRWYYCCTCRGRGSRCRKYMRGCRYS
44              RRYRWTKYWYYAYRWMMWCAACCCKYSRAKACYMYRWYCATGTRKGTCGAAKTAGAMACS
                ***** *   *.      . :  .**.  *.. ***** .   *   . . *   .   *

38              YWRRRYKKARWKKTKARRYGRGYYWKRMRWSYYGAGAAWKGCYAYMWYMKWSKKRYKAGR
44              YWRAGYKGTRWKKGGGRRYRGTCCTGAMRWSYCRGAGGWKRTCWTCACCKTCGKRYKGAG
                ***  ** :****  .***        *****  ....**         * . ****..

38              CCKGWYCTKTKYKGRTKACKKAACRYYYYMKKWGRMTCTTYGCRKRKKRKGMRCTCYYMY
44              YTTKACYCKYKYKARWTRSKGWWSRCYYTMKKWRACGMWACAARKRKKGKRCGTGTTCCY
                  .     * ***.* . .*   .* ** ****      : ..***** *         *

38              TGCTMAAKAAWAAYRCYYGYRKTAKKCARGATGTYTTGTYTCTTKYGCKTCCMTAWRTAR
44              WTMYMWRKMRTGRCGYCCRCGKKRGGMGGACCACYYKRWCCTCCTYRMKWMMAKRTRCTA
```

```
                    *  *   .          *.    . .. . * .       .*  *    .  * :

38                  CGTTAAWYACRAWKYYRRMKGTCATRMATAYCYRGGAAGCACAKWRGRYKSKYGKTYYTR
44                  TRACGTWYWYRTWKCCAACGTGYGGGMWCGTMYGATTRCTMTTKWGRGYKSKYKKYCCCR
                      : .:**  *:**         .  *  .  *  . :      :**   ***** *    *

38                  YYWMATYCGWYRYATKAYYYGYRKYWGAGTTYWRGAYAGCRKTTCACRRCTYCGYGMKYR
44                  YYTARYTMCWYGYRCTGYTCACRTYWRTACGYWRAMCCAYRKWYYMTGGTCYTRCRMKYA
                    **       ** *   ..*  . *.** :.  ***.   .. **         *    ***

38                  KKMRKYCKGCKKCTWASYWRCKRATYKYTAKMCAKSKTKTGTKYARRMYRKSKRYAWYRR
44                  KKCAKCAKKMKKYWTMSYWRYKGGKCGTYWKAYGTCKKKYACKYGRRMTAGGGRYRTYRR
                    **  *  .*  **    **** *  ..      *   ...*.* . **.***    . **  ***

38                  RRCAACTGTKTRTRSYYYYYKRCYRRGMCGWGRYMWKKKKCKWTWRTAWKKWCYKKRTWC
44                  RRMRRSKRKKCRARCYYYYTKRMTGGRMTTTRGYCTKTKGYKTYWRYMWKKTTCKKRYTM
                    **    .. .* *:*.**** **      *     *  *.*  *  **  ***   ***

38                  GWRYKKWRKTCGTKWKYACTATRKKACTYKCGCAGYRGCTAYGKYWKRGRRSMAACAYKY
44                  CWRCKKWRGYYRYTWKYMSGCAGGKCMYYKGATGRYRRYYWYAKYWKAKGAGMGGTGYKY
                     ** ****      .*** . .:  *.   **  . . **     *.****     .*.. .***

38                  ARKGYRARRRTTWKYKMRYYKWMRYKTCRTGKKYRTAAAYCTCWKKKMSGCYRSKGAMMK
44                  RRTRYRRRRGAAWKTKMRCYKWMRYKGTRKKKKTRCTMTCMKYWGKKMGAYTGGKKGAAK
                     *. ** ** ::** *** *******  *. ** * : :   . * ***..    .*  .  *

38                  RKMKYRCSWYKKCCAKTTYTCRRRYTWTACMYYYSACWAMCRCRKRYTGKCATKCKKYKK
44                  GKMKTAYCTTTTATMKYWYCYRRGCKWWRSACYCGGTWRMYAMAKRYWRKTTGKTKKYKK
                     ***    .   ...   *   *   **   .*   .   *  ..  * *     ***   *  :  * *****

38                  TKGAARMWYGTCTAGMKKGYMMARMAWRYKCTCKRYGRTRKCKAKMYSRCWMKKYAYKTR
44                  YKRMGGATYCKYYRKMTTACMCRRACARTTTAATRYRRWRKATWKMYSRATCGKCGCGYG
                     *   .    *  .     *...  *   *  .  *  .  :..** * **..  *****.    *  .

38                  AKTYWTAATACAKCTCGCCMYRKYCTRSWCAWGKRRYTRGWSAGARRRKYRACGATTGYC
44                  RKKYTYRRCCTCKTATTTTCYRKYTCGGTYCWKKARYWAKWSWKRGGAKYAGGACYKAYY
                     *.*      .  .* :      ****     .   .* * **    **        **  .  ..  ..*

38                  CTCMCKRYYATAYRYTMRAATTGKYCGKYGTYSRKKAGGWTTTCCKKAMSRTTGGCCMGY
44                  SGYMTTATTGGGCRCGCGTGCYRGCMRKTTCCGRGGCTCWKKWMYKKTMGGKCRKYYMRT
                    .   *  .    .  .  *     :.           *      .*   .   *..    **:*.  .       *

38                  RYAYCYYAAMTGCTGCTCWWTMCGAAYGCGCYYYCAKAYGKMGMCKTTGRMTTKKGCTAC
44                  RYTYMCCGGCYRYWRTGTAWYMTRMRYRYSYCYCMRGCCKTCAMAKCKAGCAYKGSTGTT
                    **:*    ..             * *     *    .    *        .   .  . .*.*  ..   :  *  .   :

38                  ACRGTGARGKATGYMRKRRKRKTCRKGMTWTMMGKYYATRKTGTTRKYMCTKTKACACAS
44                  WMRKYRMRRKGKRTCGKRRTGKCTAGRMYWCCARGCYWKRKWSKGRKYCTCTCTRYCTMC
                      *     * *..     ***. *       * *        *  .**  .. ***      .   .    .    .

38                  KYRARKRKYYAKYCMCSKCGRTTGKWYYRAKKGARAKKKAGTGGARTACARYYMYYTCCT
44                  KYGGATAKCYRGYYMAGGYRRKAKKWYYRRKKSWRGKTKGTWRRGRCCTTGCTMYYYYYY
                    **  .  . * *   * *..     *.: ***** **. *.*.*.      .*  .  :     ***
```

```
38    AYRYARMCGAKCMCMKMMRYWYRYTCAATTYKTGAGRKYRTGTTTAAKKKYRKTAKYRRY
44    GYRTMRMYRGGMCMAKCCATWYGYYYRRKWTTKKMRATCGARCGKRRKKKYRKYRGCRRY
      .**  **  .     *    ** *    .  ..    .  :   .  ******    ***

38    MMKKWYRGGACKMWWWGRGRRSCATKTSKRYAMSRGTYMKCMTATKRMYASRYYWYRKRA
44    MMKGWTRTRRTGMTTTRATGGSTMAKGSKACMCSGRACCKYMCTKKRMYGSRYYWYGKAG
      *** * *     *        *  :* **    *  :  * * :.****.****** * .

38    TTKATYRMATAKKCCGCTKMTGGYTCTMRCAYGAKKKCRYTCATKYKTCGAAAMYKMWKR
44    CCKMGYRCTWTKKYMSGGKCACRYGTCCRMWYAGKKKMRYKYRYKYKCTARWRMYTMWKA
        *  ** : :**  .  * :  *    *  *..*** **.   ***  .   **.***

38    KTCTKGCRMKRTGKTGMRWKTTRYWCKYRYWAKYRKKGWWCYMMTMYRMCCGCRCTRRYK
44    TWTCKTMGCGRYRKCACATKKKATTAKYRTWRKTGGGTTAYCMMKMYRCMTRYGTAAATK
      .   *     *  * .   *..   .*** * *         **.***       :   *

38    TRYKGGWAACCTMKYCKTTYKYYKSWKYYSWGYAKKRTYYGYGRRAYKKKAKKRWYCKYY
44    YGCKRRAWTMSKCGYYKKWYKCYKGAKYYSWKCWKKRYYCRCKAACYGTGGKKGATYGCY
         *    : ..  * *. ** **. *****   *** *      .* . .**      *

38    KYMYGYGRKSWGKWSTCCRYKYYCKRTKRYGGCRWRCGAGYCCKKMYRMCKKGTCGRYKY
44    KTACRYCGGCARKWSYYTGTKCTYKRYGACAATRTRYRRRYYYKKCTRMMKKRWYKGTKY
      *    *   .  ***     *   **    .. * *    *  **  ** **      **

38    KKKCKKTWKCCGTTAGRRRYRAKSKAKCRAACTTAMRKAAAYRCRAYYRKRKRRRTRYSY
44    GKKMKKKWKYYAWYRSRARYRWKSKRTSRGCMWWWCAKGGRTGYRWYYRKRKRRGCGCGC
       ** **.**  .   .* *** *** ..*..      *..    * ********    .

38    MGTTYKYWKGCYYAAKGYSTKYKTKCRCYMYAACRKYYCTTGAAGGCKTTRMRYYKCCCG
44    AKYCYKYWKATCCGTKSCCCKYKKTTGMCCCRWYRKYTYCACTGTAMKWWRMRYYKTYMR
          *****.   .:*. . ***..         ***   : :. . *  ******

38    ARKTAYSKKRCTTKKCAGGRAMYWRMYYYCRSYRYAMYAAMGKGARGKKKACCYYYTYYC
44    RRKWRYSKKRYCCKKYCCRRMAYWAACYCMRCTATRMCWRAAKRMARKKGRYMTCTYYCA
       **  *****   ** .  *  **   *  *.    *    .*    **        * .

38    TCYRKTRAGTRWAGGWMAYYKWGRWYAWAGCYRKYAYRWSSSCRRYRMCCRYRGRGWCCC
44    KYYRGYRGRCGTWRRWMGYYKWKRWCGWMATTGKYMYRWGSSYRRYRMYMRCRKRATYYT
      . **  *.       **.**** ** .* .   ** ***.** *****  * * *.

38    AMATGGTGSKKYAYTRGCWYTTTRWRAWKTKRWGCCGTAKMMRKMACGGAGYAGKWKCCT
44    TMRYRSCKSKGCTCCASTTYWCYGWRMWKKKRARTMRCCKCMRKMGMSAWKTWKGAKYTC
      :*   .  **  :   .  *    ** **.**      .* ****. ..       *

38    TTKRWAYKSMKKKYRKMMRWAMRGKRTYRGTTTTKRRTCGAYMCYYGKKKRRWATTMKYR
44    CCKGWGTKSAKGKTGKCCRTCCAKGRCYGKAAAAKRRCATTYMMTCKKKKRAAMKKCKTG
        * *. ** * *  *  * .    * *  ::::*** . :**    ****   .. *

38    ARMWGKSTTKWYCGTYKRKWAAAWYRTTCYKKYAYKAKYAGKKRRYGTCTACCCTRMTTS
44    GRMWRKSCCKWCTTCTKRTAGMRTTGGYYCKKCGYKMKYGAGKRRCAKYYWYYYYACYWS
      .*** **  **     **. .         ** .** **.. *** ..           *
```

```
38  GGAATATYCCTGCTKATAYYRYAYYKGTYRRTCYTYATWKMSKKGYKCKCCKAGTCYYGK
44  RAWRKRYYYMWAYWKWWMTYRYGCCKCCCRGCTCYCWKWKMSKKACKYKTAKGTCYTCRG
     .  .  *   .  *    ***.  *   *       .******. * * .*.

38  KKMGRYMTKKGKTKKTMKKRMCTRRKWCATGAKAAYTCSTTTCKKYYRKATTYTAYWKKG
44  KKMSRYAKKKKKKKKYMKKRCAGRRKWYRCTCKWRYATGYYYATKCCGKTAYCCCYWTKR
    ***.** .** *.** **** . ****    .*  *: .   ..*   *::   .**.*

38  TYAACRTYKYMCWWGGGTKCTTGKKAGCAAGCKTCWCAGRMYKCGCSKGYRKKKTACCAY
44  WYRRYGYTKTCYTATSKKKTCGTKKTATCCRYGGYWTRARMTKYKMCKKYRKKKYGTMRC
     *      *      . .*    **:. ..     *  .** *   .* ***** .

38  GTAGKWKKWCKRRKKYKYMRKKTTKMKWACCGGGRKGYKTRWKTGATRYRWGTARRTYYR
44  CAGKKWKKWYTRRKKCKCMRKKYYKAKWWTTAASRKRYKWGWGCATCRYRAKWRRGWYYR
     :. ***** .**** * ****  * **   ...** **  *  .: ***    *  ***

38  KYTCCRKATGCACTYKKYTSRRRKMWKRKCWAKAWKTRSARYYTGCRKKKYKYGCGCGCG
44  KYYYMRKRACTRMWCKKCWSRRRKMWKRKYTRKTWKKGCRRYYKRTGKKKTKYKYTMAMR
    **   ** :      **  **********   *:**. . ***.   *** **    .

38  KAGRRYKGRTRKWKWSWRGACGGAYMGGSWTARRYTTYWRRRYTKCRAATYKKRRGYTKT
44  KGRARYKARKRKAKWSWRAGYRRGYMKKSAWMGRCYYYWRAACYTYAMCAYKGARRTCKY
    *.  ***.*.** *****..   .**  *    *   ***    .   .:**  *   *

38  TCWTWYRWACRRKTATAYRTYWCWCCWTYGMKCGRRWMMAATKAKKYRYCAMKYYYKYTA
44  CGTKWYRWRYGRKGGGWTACCTYTMMWAYKCKMKRGTCMRRKKRKKYRCACAGCCCGCAC
       .****   ** .           *:*  *  *   *  .* **** ..       :.

38  MKGTCRGATGYGAYRYCMRWKCCAGYWMMYKGTRYRRYAMRYRCYWKSRKKAKAAKTTAT
44  MKSWTGAMAAYRRYRYGMRWKSMWRYWMCYKKWATGACTMAYGMYAKSAGTTGWMGGAGC
    **.   . :.*  *** ****.   *** **       :* *  * **  .:     :.

38  MWRSWYYSRWGAATYKMWWYRRRRRRKCYTKARKYYMKRKTKMKRGCCGAATRKTTGYYK
44  MWRGWYYSAATGCCYKMTWYRRRRRRGYCGKRAGTCCKGKCKMKGKYTAGRGRKYYRTCK
    ***.****   .. *** ********    *      * * ***    ..  **     *

38  RYRMYKCRCTMCRMKRKYMKKSRCRKCMTRRKSKGKKYRKRKCYYSGTYRGKRCKMKKTY
44  GCRMYGTGYKCYRAKRKYMGKCRYAKACYAAKSKKKTCGGRGTCCSRYCGKGAYTMGGYY
      ***    .  * ***** *.*  *.    *** *.   *    *        .*   *

38  RCKRGTGKGAAWYWRRRYRRCRCKTTYCRYKKYCKWRRTAGGTGCAYYRYRTYTYKKRRY
44  GMGRKCTKKMMWCWRARYGRYAYGYWCMRYKGTGKWRGKGKRKAYRYYRYRWYCYKKGGC
       *   *   * ** ** *        ***   *** ..  ..  ***** * ***

38  TATATTAKTGKMWRKTWGCYRATATYRYWKRMRTCGGTGGMGCRRKWTAKKCARWYMRKT
44  CGCWAWTKWAKMWAKCWATCACCMKCGCAGAMGCTRKYRRMTYRRKTKWKKTGAACARKY
     .  : :* .*** * *.   .  .      *        *  *** . ** .    **

38  TYYMRTSTTTWCYTTTAKKWYCKMMWATTWKKATYYRWWSCWSSYWTTTARRWKRRCMYK
44  WYYMRKSKKWTYYGYWGGKWYYGCATWKCTKTGACYGWWSMWGSYTCYKWRRWKGRTMYK
     ****.*..   *   . ***      .  *..: * *** *.**   . **** * ***

38  ATRWACGTGYTKGAGTYYSTKCYATTGTYRKSCYATTKKWWTRKYRKGAWATATMYACMK
```

```
44      GAGTWTCCAYAKSGTCCTGYKYYCACKYCAKSMTRYWKKTWWATTGGAGTTWRKMCGTCK
        .:      .*:*..    . * *.:     **     ** *  .   .. :  .* .  *

38      TRRCAGCWGTYGMAYKTKKMYWRMTCATCTTGAKACMKMYRYTYCACAYAGRTKRYYKCR
44      YRRYRRMAKGCRMMYGKKKCCTAMYMMYSKKKWGGTCGMYRCGYYGYTTGKRYGGTYGYR
         **         * * .**    *    ...   .   ***  * . : . *    *  *

38      GRYRAGYYKSAGGCYRACAAARYCTCGAACCMYKACCARYGYCKWYKGKKYARRARRAAT
44      RGYRRRCTGCGATYYRTMWRTAYMCTKRRYYCCGGYYGATRTYKTCKTKKCWGRGRGRWC
          **     ...  **:   : *           .  .     *  * **   *.*

38      GCAKKSRRMYRMTGTYWWKKWWSYCYYRARKTRAWASGTAACYRYARTTTAYKRRAGCGT
44      RTWKKGGACTGCKKYYAWKKAAGCTTCGMRKYRMWWGACGGTCGYTAGWAGTGAARRMTC
           **.      .  * ***  .      ** * * .. ..   *:   :.

38      MYTKGYKKCKKGGGRTCMKGMGCKKRYYRYGGYTWTYCTKMYMKKKARCAAYRRAGAYMW
44      CCYGKCKKYGTKKRGCMAGRMSYKKRTYRYRACYAWTAKKMTMKKKGATTRCAGTCGCMW
              **  .         *. *** *** .     ..** ****.  :    : . **

38      STYKYYTGRCYYYRCAMSKTYCRTAKKGCTARWACYGCTTTRTYWAGWKGKGYKYYKGYG
44      CWCKYTKRGMCTCGYMAGKKYTAWRKKKAGRGTGYCKYGWWRGYWWKTTRKKYKYYKKYS
        .  ** .          .*.*    ** .    .       * **   . * ***** *.

38      TTTCCTACATWAGTATKRMATGACYRWKKTGGMWWKRCTGKRRGGGGCYRWGTTCMYKKT
44      CYYTAYMYWAWMKKCGGACGGATTYRWKKYRRMWWGRYCRKRARRRKMTGTAGGSCYKKY
            .    :*  ..    . .: *****   *** *   **         .  . ***

38      GGCWAGAYAGAMTCTCYYKKWWYKCKCTRKGATGWMCCTAACAAWSRWAYKGTYRYKTGK
44      RRYWRSMYWKRCYYKYTTTKWACGTGTKRKATCRTASMYTMMWMWSRWRYKTGYRYGYRK
           * . *      .   .**      .**.:    .  :    **** **  ***   *

38      CGAACGGAGKGCSRRKRYKCRYTGAWWTRYMMKMSCCGAKYSMTWRRYYKTRWGWAAGRY
44      YRRRYRRRRKKSGRRKRYKYRYGAWWACGCMMKMSMYRRKYCCCAAATTGYRWCTWGARY
                 * ..****** ** . *    *****    **.         **   ..**

38      GKMRMRRRKWTKCCCCYAGMTTSSRGSARYTTTAKRCYRRCYWACKKYATYCSMTYAKMM
44      RKMRMGRRKWYKYYYMTMKMKWSSRRSCRYAGARKGMCGAYYTRYKKYMAYYCCGYRKMM
         **** **** *       *. *** *.**: : *      *   *** :* .  * ***

38      CRATCRYYYYYCRKRRKGTYATGTYKAYKYAKYKTARCYMATGCWGAGSKACSKKWKRRY
44      YGWYTGYCYYYYAKRRGRYCRKKKYKCCKTMGTKCCAMYMRYRYARWRGTGYSKKWKRRY
              * ***  ***     . .**. *    * .  **        ... ********

38      WRGACCKKRWCGMACTCKRYYMKKAAMMRKAKRCMRMKGKKKTYRMKCYKWMMKKATTKA
44      TGATATKKATYSCTTAMKAYCCKKWRMMRKCTGAARMTKKKKAYRMKMCGWMMKGRKYGT
          .:. **   . : : * *  **  ****.. . **. ***:****   ****  .  :

38      GCATGGCTKWKYRAKRTTTCMYWRKYRTYSYYYTKKYGKAAAGARRWYKRKKYKYYKRRK
44      ATRCTATYKWKYGCKGCACTMYWRGYRGYGCCTKKGCKTRRWRGRRWCKRKKYKYYKRRG
        .    .  **** .*  :  **** ** *.   .*   .    .*** ***********

38      RTKGAMYATGGYRWYTTYAAKMRRYYGCMKAAAMATCAKRYYACMYYKMAYRKMKTATGY
44      RWTSMAYGATKYRTTYWYCGGMGGYYRYMKRWWAGGTMGGCYRSMYYKMGYRKMKYMWKY
```

```
                * ..  *.:  **    *.. *  **  **    .      * .*****.*****    *

38              MYRRWMRKAKTTAYYRRCAAAACSYAGKATYCKTRYTTCKAAAGGTGKKAKRWKTGAAYY
44              ACRRWCAKRKCGRCTRGMCCRGMCYWKKGCTTKKGYCAAKGGTTACAKGMKATKWAGGCC
                  ***  * *     *  .. . .*  *.   *. * :.*..: . .*  *  * ...

38              WKGWWYKTYYKGYCRKKKTCRRTKYGATMTCAARRRYKMMKTGRAAKKAARWKRKAYMAR
44              TGTAACGYYCKRYSGTGKWSRAKTCRGWCWYRWGRRYKCMKWTRTTTKRMRTGAKWYMRG
                        * * *. . * .* ..  .       **** **  *::.*  *   * **

38              YMRATKGTGTTGRGTGCCRCRKGWCCMCYKMCTAKARSKYCCCWRWRGKACATRKKTGCT
44              YMAWYTRKRWKKRAAAYSRYRKKAMAASCKMYWRKRRCKCYMMTAWATKGYGGAKTKSMG
                **   . .  . *.:. .* **   . . **   * *.*      *  *. .  *...

38              GGRRRKMCGACWRKKARYAWGYWKKSRYTGGCYSYCAGKKRKRKKYTGGCKYKGCWMYYY
44              CKRGGTAYKRYAAKKGRYRWRCWKKGACYRRYYSYYWRKKGKRKKYWRRSKYKRYWMCYT
                  *  .       **.** *  ***.      ***   ** *****   .***  ** *

38              YKYYRRACYKCAMYKRGTWACCMCGKRCRCAGRTGCGRTTCAMTACCAKRGTCYATTTGM
44              YKYCAGTYCGYCATKGSYWMTTAATKRYRYRRRAATTRCYYMMAWMYRGGRYYYRYGCCM
                ***   :    .  * . *    . ** *   *:.  *    *:         *     *

38              YYGKKRWGYATRWMGCATCTTGAAGGTAAAGCYMMKTYYRCSRACAGRWMYKRRMCRMKW
44              YYAKKGTRCWYGTCATCGMACARRRKKCRGAMYCAKCYYGYGGRYGTRWACKRGCAAMGA
                **.**         . .  : .    .. .. *  * **  .   . **  **  . *

38              ATCGYCCWKCRYYYWCCKAACMAKACTGCAMAYAMYCCKGAATGMGMTRTMYCKTCAAWK
44              TWMRCMYWKYRYYTWMMKWGGMGKRMYSMRCRYGAYSYKRRMATMRCCRCACYKGMWWAG
                :      ** *** *  * . *.*   .    *. *. *   : *   *    *

38              KCACTSTWGGYRWYGYYYMMYGRTTKGKRWMYKAACKGTTRCKWWYRRWKWKGCSRTACC
44              KTGTACGWRACRTCRYCTMMYKGAGGRKRWMCKTTAKRAWRYKATCAGAKAKRYGAKRMM
                * . :. * . *   *  ***  :   **** *::.* : * *      * *  . .

38              KAAKMKYGTRGYGCCGKKRRWKCRMKKAAAGKKCGAMATAGTAWAWRCTTKKYCAGYRAC
44              KRCKMKYKWARYAYYRKKAAAKYRMKKCGCAKKYRWARYWAKWTRTGMYYGTCYRRCGTY
                * .****    *.   **   * ****....**       ..         .      :

38              WTGTKWYRKGATTMACKRMKGRKYTRTTRRRMKKKSCYMCACGSGKWKYYCAKMGGGKRT
44              WCAGKACRTAWCCMMTGGAGRRKCYGYYGGRMKKKSMTMYMYKSKKTKYTYWKMKKRTRW
                * . *  *..   *       **       ******  *    * * **   **   .*

38              AKKTATCRYRTWRKYKGCRTKSRRYRKATAAYRYRAWGGGGKKRCMCTGCTKYKKMGCTC
44              MKKCGCAGYRKTGKCKATGCGSRRYRKGCRRCRYAMTAATAKGAYASKRYKKYKGCATAT
                 ** . . **.  * *.    ******.    **   .. .*    ..  .***  . :

38              TRKWYRAYCWGWRYYMKCKRKYMYMYRRYAARRYKGYMYMRYYKKWAAMKRYWAATAKYA
44              CGKWYGRYMWRTAYYCKGKRKYATAYAATTGRGTGATCCAGTCKKAGCCKGYWRRARKYC
                  ***  * *   ** * ****   *   :.*   .       **  .. * **  : **.

38              RRTMWGRKGCGGRWKRKGTYSKYSKGCGCGAYKRKKCKATRMKCYYKSACWYSRYKYAWA
44              GRWCTTATRSRRGTGRKRKYCKTCGKYRYSRYKRKKSKRYGAKYYCTSMTACGGTKYGWM
                 *     . .      ** .*.* .     . *****.*    * * .*    .  **.*
```

```
38                  KKCSKWRKKCRTYKWKRCAKKWKCCTCCATGKWCYWTAWKYARKMG
44                  KKYSKWRKGMGCCGAGRMMKKTKMYYYTWCTKATCTGRWKCGGGCK
                    ** *****        *  ** *        *      ** .
```

The pairwise alignments of two accessions 38 and 45

```
38              YRMKTRKYWMYGRWCTKYYWYRKWRYRRYCGRYMKCTWACKCRGYRCCAKRCYMRYYAYR
45              YRMKYRKYAAYKGATGKYYWYAGWRYGRYTRRYMKYWWCYGGRTYRTTRKRAYMRYYRYA
                **** ***  *     *****  *** **  ****  *.   * **   **.***** *

38              YCKKMWKMMRGAKWKYWRCYGAYRWMMYMKKRKYRYKMTTTTYRRRRRGAKRTYGCKYWR
45              YYGKMWTMMRRCKWKTARMTTGYRWMMYMGKAKYACKMYCCWCAAGGAKRGACCAGTYWR
                *  ***.*** .***  *   .******* * **  **                . .***

38              SRWWRWYMMRWRRYWYRYYKSYKKCYGSSCATYRSYKYYYRWATAMKSYSYRKAYYSYGK
45              CGTTRTYMMRWAAYTYRYTKSYKKYCRGSMMYYGGTKYYTGWRCMATGTSYRGGTYCYCT
                .   * *****  * *** *****   .*   * . ***  *    .. *** . *.* .

38              YRAYRRRCKRRRSGAYMYYYRMKTRYYRRYWRKTKGYSGSYWRRRGGSTRGKGTCWRSYA
45              CGCTGARYTRAASARYMYYCGCKWGYYRRYWAGYKTTCKCYWRRRRTCWRAGRAAARSCG
                  .   * .*  *. ****   *  ******   *  . .*****  . *.  :. ** .

38              CAKSYCAAYKWYTGRWSRGRRGATSWYKRYAKTTGYGTARYCRMYSKRRYKYYYMRWYSY
45              ATKSCTTGYKWCKRRWSGARAAGYGTTGGTRGWKTYRCGRCYGMTCKRRYKYYYCRWYGY
                .:**  :.*** . *** .* .. .        . *  .*   * .******** ***.*

38              WWRWRCAKMRMMKSRKAYWWYCGGTWSMKCCARCTTGKYYYYMYYRRCACWMAYMRRYMM
45              WWRWGAGKMRMMKSRKRYWWYTAACTCMGTMWAMYWTKYYYYCYYRRTGGWCCCCRRYAA
                **** ..********* **** ..  .*         ***** **** . * .  ***

38              RWRWTCKWMRRRYYYYWRKRKWYTYCKAYWGYWTAGRYRWKYRYYYWSYYWKYYYCKMYY
45              GAAWCTKWMRAGYYYYWRKRKTCYTYKTCARYWYGKATGTKYRTTTTGCYWKCTTYKMYT
                   *  ****  *********     *:   ** .     ***    . ***    ***

38              GMTMYMMYGYYKYYWYMCCMTMGYYRKYMRYGAYRRYRWYYWTCAARARCCCRYYRRCKK
45              RMWCTMATRTTTCTTCCYYMYCRCYRKTMGYRGCRRYRWYYWYYRRRMRAGYRCCAASKK
                 *   *     .       *    *** * * . ********    * *.  *    .**

38              YYRTRAWMYTKMRKWRYRRRRKYYRAYRYRWKYMCMAMRYRKGYYTTYRWWKGKTTKKGT
45              YYGKGWAACWKMRGAACARRRGCCRGYACGAKCAAMTCRCRGCYYYYCRWWKTKKYKGAC
                ** .      ***     ***   *.*    *  .*: * *  **   **** *. * .

38              GTACTRTRKYRTGAKKRTTGCTRTTGGACGYYMRRYKCCCKRGAKTCRKGTGGYCMGGGR
45              AKWYYRWRKYAYRMKKRYYKMGRWWKAGYKYTCGRCGMYYTGSRKCTAGRYAKYAMTTTR
                ..   * ***    ***     *   ..  *   *     . . *      . *.*   *

38              TRYYMMATAATGRKTRYACCKRKYYTCKAGCTCSKWSGGCYRTGGYCYMCTWMSTRCCMK
45              CATYMMWCRGKSRGGRYGYMGAKYCCTKTTTATCGAGAATYRKRKCYYMAWAMSAGTYMK
                   ***   ...*  **.    **   *:  : .  ... **.    **.  **:   **

38              CAYGAKKTCGKYRRSYWTMYWMSTCMWYACYRYAAMCGMYRRRWYGTGCTGGATCAWGGG
45              TGYSWTTKYRTYRRCYACMTTMSCYATYMTYRYWWMYRMYRGGWTRWSYGCTCYYGTKKR
                 .*. ...  .***.*  *  **    *  ***  *  ***  *   .    .  .

38              ARTYCTAACTWWGGCTTTKRRRKKMGYWRWYCCACAAKYCRYWATTCKMYYKTTGGMKTC
45              MGCYTYWRTWWWRRACGKKAAATGMKYWRTTAAGACGKYAGYWGACTGMYYKGCTCMKCY
```

```
                   *      **   .  .*   . * ***  ......**. **.:   ****    **

38                 CKARKRYGKYKYRYKTGATAGMYACRKAKRYCRTTTAYYCKRTARAYTTGCRSMKTCKYY
45                 TTMRGGCTGYKYRYKARCYRKACMMRGRKRCYGYCARCTYKRYRGGCWYTMACMKCTKYY
                    . *      ******: .         *  **       :     **    .       .**  ***

38                 RRGKKWRYMGRCAYKKCAYKYACYCTYGKWGAGKRGCCCATKRKAGCTAKKCYATTCKRT
45                 GARGKWGYCKAATCKKTTCKYRSYTYTRKATRRKRKTTMTCKRKCCMYRKKYYMWWYKRC
                       ** *    .: ** : ** .*     *     **     : ***.     ** *    **

38                 TTMGAMRCTCWKAGCSKYTGCTKRKYYCRWRGCATKKMSTSKKMAGGTCGYTTMYCTYCT
45                 YYAKGCGTGTWKTAACKYWRMYKRKYYTRTRKAGKKGCCCGKKAWAKCYAYYKCCTWYMC
                       .     **:...**    ***** * * ...*  . .**  .    .* .     *

38                 GCAAACKTTAATATCRRCKAAAGCGGGGGCCGTCAWYCKGRYGTRRGAAMGAAKACCGCC
45                 RMCWWMGWWGRCMCTRATGTGWTTACAKTYYTGTCTCYGRGTAAAAKWRCKGCKWSYRYS
                     .      .     *    :.    . .       .       .:        ..* .   .

38                 GRYTATTAATMWKRKYKSWMRRCRGYYYRGKAWMYMGWTKGYKKRYGRGCGGYRKGRTTR
45                 ARTKGKWGRYMWKGTYKSWMRRMRRCCYRTGWTATMAAYGRCTKRYARCSAACRKTAYYG
                   .* ... .  *** .******* *    **      *.      .***.* ... **

38                 YKAACCCCMRCGTGCGWYAGGCATTTACCATATKRGTCTCCCAYACAGCTCCCATCYMRT
45                 CTTTTATTAGMRAATTACMAAAWKCWTATGAGAGGRYAKYTYGCTTTRAYYYMRYMCCGG
                    .:: .      :.      ... .  :. .:.:     ..    . : : .

38                 WCYYTTCARWMKTTGTSTGYKCGRKYMAYRCATTACWKMYRGTYYGATCYRYWSGAWKMR
45                 TACCYWMWRTCKACACGGATGMRRKYCGYGYRYYGYWKACRKKCCRGYYYACTGATAKCA
                    .      *  *: . . .    *** .*      . **  * .   .  *    ..: *

38                 YTACGGCWKCRTRKYGTYKKYGRMAKACACTGGGACYRKKYAAGTRKCACTWWWRKKGGY
45                 YCGTTTYWKAAARKYKYTKKTRRMWKGYRMYSRRGTTRKKCRMTGGGTWMWATAAKKRAC
                   * .     **. :***    **  ** *.     .  .  ***                **  .

38                 CCWSRKYWCCYYMMYYGYRTTCTRWMYKTGKCAGTTKTRRWWRWKWRCSRTACRKCGCKG
45                 YGTGAKYWMYCTMMYYKCGWYYGRTCCKAAKYWAAAGGRGATGAKARYGAKRMGGTTGKR
                      . ***    ****           *    *:.*   .::  *        * *  . .        *

38                 CKKMRYGTGRGCMCGRRWCAKRMKRWMYKCMKKAAAGCCGCGARMWKRYKARWWCTTKGG
45                 AKTARYKWRARMCARGRWYGKAAKRWCYKYMKKMRMRYSKYRWRMWKRYKWGTTMYYTRA
                   .*. **         .   ** .*  *** ** ***        .      *******         . .

38                 YGTTRYRACCWKRKMAYYWAATGTTAGGKARRRMGTCKAMRYGYYWKKTCATMRACGTAR
45                 CCCCRCGTYAWKRKCWTCARMWAWWMCRTWGGRMKYMKMMRYTCCWKKKTWWMRGMRWMG
                       *   : .****            .         .    **    * ***    ***.    **.

38                 YRTATCYRYRWCYRSRRYSWKYRYAAKMMWWRKKRGMAGRCKTAATGKRYRKAKCWARKK
45                 TACGCACAYRTACRSRRYGAKCRYCGKMMWWAGKRRMWRRTKATTGRKGTRKCGSWWRKK
                       . .   **  . *****. * **..*****   ** *   * *:::   *   **. .* ***

38                 RTGCWCAGACYTAGTAMYYKWTWWCYYRRKKAYYMRCGAAGCTKKYRAACRCTWSRACAG
45                 RYRYWTRSMGYCGCARCTTTWWWAYCYRRKKRCCMRYKTRKTWKKTAMMTGYYACGWYTT
                   *   *   .   *  . :      .* *   *****    **   :     **            .    :
```

```
38      TKTGKCRGAATKYYTAWCGKYKGASTYKKTGTRAYAWYRTCAWGTRYMKKYSWYWWRKTR
45      KKYRKYRRRRCGYCYMAMRKYKKWSCYKTGAYARYWWYRWYRAACGCMKTYCTTAAAGYR
        .*  * *     *      ***  * **. .   * ***    .   **.*.       *

38      WWRYKWGWAACARYGKTMRRYRATSCKKSRWATRRTKYGCAYCGTSCASWRCYTYGMRRY
45      WAGTGTCTGTGGRYRKCCGRYRRKCYKKSRWMGAGYKYCAGTTKWSYWSWRYYCCTCGAT
        *       .: .** *   *** .. *****     ** ..    *  *** *

38      CKACWKKTCGYKCGSMTTCKGKGMMCACRATKCYRYAYYYKARCRYCTRTAKRGRTCKKY
45      YKRYTKKATAYGYRCCCGSGKTKMMTRYGWYKTYRCRYYYKCGMRTAYRKWTRRRGAKKY
         *   **: .*   .   .  . **      * **  ****.  * . *. .* * .***

38      MCWWKAWKCKYCACGTMGAGMAYRKYACTKTARSCWCACKRGMGAGGCCACSAGKAGRWK
45      AMTTGGTKTGYTGAAACTTTCTYAKTGYKKCWGGYTTGYKRRCKGAAGYWYSWRKTTGAK
             . *  * ...:  :  :* * . .*   .   . **   ...    *  *:   *

38      SGTACGGWATTCCYTMTMKCCCYRMYTTGGARYYGCCYRRKCYTAAKYWMCYSRKMGGKK
45      GTCGTRAWCYKYSYCMYCTTATTGCYCGRRGRYYRAACRRKTCYTGTTWMMCGRGCAAKK
        .  .  .*. . .* *  . .    *    .*** .. ***   :.. **  .*  ..**

38      MWRAGCTKMGYRYCYTRYKMGRRKKGTGYCAGTTACWKGGAGCGCWATTKCCAKARCMCG
45      MWRGRYCKMRTRYMCGACKCSRGTTRKKCTGRCYRMWKTRMRYSYWGCGGAMCKGRTMAA
        ***.   **  **     * .* .. .   .     **     . *.   . .*.* *..

38      CMCTRTGTYGYGGTMRKYWGYAYKKKKYRTCMKYKRYRWRMKYMRYYKRKRYRTTTTKRR
45      YAMKGCTAYRCKRKMRKYWACGTKKKKYRKMMKTKGTGWRCGTCACYKRGAYGAAAAKRR
           .   :*    .*****. . ******. ** *   **      ***  * ::::***

38      WMKKWTKKCAKSKYYYRTTAGTTRKMRMKAGKSYTKAYCGYAAKTCKYYTKKYAKAGKSC
45      TMKKTAGTYTKSKYYYGWKMRKKGGMRAKRRKSCCGRYYKTMRKYYGYCKKGYMKMRKCY
         *** : . :******  .  ..  ** *  **    *     *   * .* * *  *.

38      YAGKWCRTTCTASGKGYCWTKATAMRYYRYKYRRTCTAGYTYRMKMKKYKGCTTCKGGYS
45      YGAKWMRCAYYWSRKRYYWWKGGGMGTYRYKYGGYYWWKCYCRCKAKKYKAYKYMKRRCS
        *..** * :   * * * * *. .*  *****          * * ****. .  *   *

38      TTWSAYTCTGTGCKWCYTRRKTTCCTYGCKMWMRWWSTCWRCCTTMRWKCGATRMWKYGT
45      KWTCMTGMGRAAGKWYYKRRKACTAGCAYGCTMGWWSGMWRAACCARWKTAGARMWKYCC
        .  .      :. ** *.***:  .  .    * ***  **..   *** ..:*****

38      AGTCYMRAWKCTCYATTAYRGCATSATRWWWRKYRCARCYCMMMTRRKRWTCTCRRYKAC
45      GTCYCARRAKTAYYCACMCAKYMWSRYRWTWRKYRYWRYCYAMCCGAKRWKSCMAGCKGY
        .     *  * : *.:        *  ** *****  *    *    ***..     *.

38      CCACTWTCATKWRKKMYWRKACMKCGCMGWSRRTTYRRACKKGGCRGAAATAGYWWRKRW
45      SYMMKWAATCGAGKKCCTAKGTCGAKTMATCRRKWTRGCTGGSSYRRGTCWRRCAWRKGW
        .   .*:.:    **    *.   .  *. .**.  * .   .. * .:.     *** *

38      AGGCYRAKGGKMTTMYCKWGYKKYRYTARRAGYWRRKTACKKTTTCGCYRKKCASRRYCT
45      RRKYYRRGKRKMKCCTYKTKCKKCATWRGRGKYWGAGCRMKKCCAYRMYGKTTMGGAYAC
            **    **.    *   **      *. **      **  :   * *.  .  *.
```

```
38    RWYAWGKKRRCMGSACCATGTKYYATKAAMGCTRGGCAACTRWYYRYYRGWRARRGGKMC
45    RWYMTAKKRAYARSTTYCAAKKYYRWGWRCATARKKSGGYYGWYCRYYARTAGGATATCG
      ***  .***    *:  .:..***      . :*  ...   ** ***   .    ..

38    ATTRCKTYYWRKGMYMTRRRYAKCAMYKMARYYYKWRWTMCCMAYKRATYTARYMWAGRW
45    RKKGYKACCTRKKATAYRRRYGTACCCGARGTCYGAGAKCMTMMYKGCCYKCGYCTRKAT
       ..  *:   **     ****....        *    .   * ** . *.. *

38    CAWKTKRKARWRKGRSKSKCTAACYRCKGAKTRGKGGTMRKMYATRKYAWKYCATKTTTG
45    TTATATGKWRTAKRRGGCKYYRRYYRYGSRKYGKKRCACRGMYTCGKYGWKYATWKCCCC
       : .:. * *  * *. .*     **  . *   *  : * **:  **.***.: *

38    YKKRMSTCAYARRMGATKKYRATTTGSWYKWYWWYGTMWCKRCTMMAKRRMGTGWYKCYT
45    YKGGCCWYRCMRRMRCATKYRRYCCKSTYKAYWWCKKATATGTGCCRKRRMRARTCKMYG
      **   .     *** .:.***     * ** ***  .  ..      **** :   * *

38    ATTRKGKAAKKRGGRYKTCTAKSMWYKYAWRKKGAGAGRRRKKYGCMRYYTRGTRYWGMC
45    MYYRTKKCCKKRRRGYKKYYMKGMWYKYGWRTKRGRTARRRKKTKMMRCYYRRKRYWKCT
         *. *..***   **.   *.*****.**.* . :.*****   ** * * .***

38    KTYRKTGTYKAMMKTCWKTCMRYMTTTTCGYRSKRKTRRCMKYRCSGYKKCCATTYCWRR
45    KAYRKYACTKCCAGKYWKYYMACMGCGYYRYRGKGGYAGSMKTAYCAYKKATMYWYYARG
      *:*** . *.    . **  *  *      **.*     .**   ..***.    *  *

38    YMTAYTCYAYYRYYSRYMCRRKKMRTRKKGCCKCGGCATCWKKGTTWTCRTTKTYYKGYY
45    CAWRYWGCTCYRCCGATMYAAKKCAWRGKRYMKTTRTTGTAKKACCAWMAGAKCTYKKCY
          *   : **  .  *   **   * *   *    :   **.       :*  **  *

38    GTKMMWMMYCCTACGCTMKMTTACCCTYKTRRKRATWTAYCTCGTTAKYYYCTAGTCGYW
45    ACKAATAAYTTWTYCYYMKAKKMYYYYTTARRKGWKWCCYMYTTGCRGYYYYYTAAYRTW
      . *     *   :    ** ..      .:***  .* .*        *** :.:    *

38    YMTCYYYWTTRYAATGGYTGKKYKKWGTWGYRYYYKCYCATAMCACKWYCWCCTARRWRR
45    YAKYYYYAGCGYRMYSRYWKKKYKKWRGTTYRYYCKYTSWWRCSMTKWTYTTTCGRRARR
      * . ***    *   . *  ******    **** *  .    .  **      .** **

38    MRWKKMRAATYMWYACWKKYCGCYYRRRYRCGYKYTCCRTMYSTMCCMCKGTACARKMYY
45    MGWKGARGTAYMWYWMTKKYYKSCTGAATGYTYKYWMMAAATCYAMYATKSKRYRAKATY
      * **  *.::****   ***  .         ***    :  .      *..    *  *

38    CCTRRCCWRCRKYMRGCRCYSYWRACKRYGGRTAKRGCGCAWYMYGWKGRCCTMAKATWG
45    MTGAATGAGMAKTAGKMAYYSYWGRYKRYRRGACGAAATTGWCMYKWKRRYYYAWTGAAR
                 *       ****   ***   :.  ..  .* ** ** *     ..:

38    RRRMYMACTAWTRRYWSKKRYKAGRCTCTARYGYYTTCKKRCYYGCKRYMKACKAYYRKA
45    AAGCTCWSGGAYRRYTCGKRYKGAAYATCGGYACYACSKKRSYYATKRTMTRSKRYYRGW
             . .  *** . ****..  :  . *. *: .***.**. ** *. .* ***

38    ASYRMRTTCGYKRCCMAARGACTTYMYCCARMYTCRKRGWKSWYTCCYRRGKCYRYRTTR
45    WSYRMRKAYRCGRMYMGGGTGMGYYCYTGCRMYKSGKRKWGCWYCTTCGATGYCRCRYGA
       *****.:    *  *..  .   * *  .***.. ** * .**          * *

38    YKKGYRRYYSCGMKGTRGCRAWGCYARKKRRKRWGMRRYKCAGCYARCRYACACRYKKKY
```

```
45          YKKKCGRYCSTKMKRYGATRTAATTCAGGRRKRTKMRRYKTGTACWAMRYWTWMRYGGKT
            ***   ** *  **   . *: .  .   ****  ***** . .    **    **  *

38          TTSKKKYYYRGARTAAYYRYCTYWRYTATKGYGKYMWGKTKWYRYWKRTMTCTCACAGGM
45          WWGGKGCCCRKRGKGRYTRYGCYTRCCGCKATTKYMWRKCKACGCAGAYMCTCGGTGRRM
              . *    *   .. * **  * *  . *.  **** * *        *    . .  *

38          KGKKRYWYCATGMKYGRKKYRYKCTCCKTAKYYCTMMCKKWYARAGRKKYRYGGATSKSR
45          GRKKRYACAGGAAGYRGKKYRTKMAYTGYRGCYSYMMMKKWYRGWCRKKTAYTATKSKSA
              ****  .. .  *  **** * :       *. ** ****    ***  * .:.***

38          TYYMSMKCCKAKRCYKSRYKCRAWKKATRMTKCTMKGWCMRMRYMRKAWGKKCMKYKTCG
45          YCCCSCKYTGTKRGYGGGTKYRRTKKMKRMGKGGCKKAYARMRYMRKGARKKMMKYKCTA
                * *   :** * .  * *  ** .** *   *    *******.  ** ****  .

38          AAAKAMYKMCWATGGKTTTRKTRRYGKRYCGKTGCGRWRYKACTCMKYRRYYWARKYRGG
45          GTGTWMYTMYWRWKTGCCCAKCRRYTGGCYTKATYTATGYKCYKYATYRRYTWRRGTGKT
            .:.. **.* *         * ***      *:      **. .  .**** * *

38          TYMMCYYRCCTKRRYKRAAKCCMKYACGGAKKTMYYKGAYWSYYSRKWMTYKKRKYAYAG
45          YCCCYTCRATAKAATKRRMKMSCGYWMRRRKKKMCYTRWTWGYYSRKWAYCKKRKYCYWK
                   *. :*   **  * .  *     **.* *.   *.******   *****.*

38          KKKSCRWRYYCKYYYGKYMMTGRSWTYKTGWSYCGYYCKRTKRTGTCTYTWYGARRCGAG
45          GTGCSRARYTYGYCYKKTAMKCGCAKYKKRWSTSACTYKRYKRWRCTGCYWYKMARYRRR
             . ..* **   * * *  *.  . .**. **  ..   ** **       **   *

38          KKCRKGGKTTCKKYMRKYGRYKGKKKWGKMYRTYRRRAKTKAKRAGATKMRTCCCCGRYY
45          KKYGKATKCCYKKTMRKTKGTKAGKKAATCYRYTAAAWKWKRTRGKCGKCAWYGYTARYY
            **  *. *   ** ***    *. ** .. **      * * .*. . *       .***

38          RKRRRRYSYMCAKKTMRAAGGAYSAATGKYTRKRTATRCTYACTGGAACRMYRWKAAGGK
45          RKGRRGYSYMYGKKYMRMGSKWCCWRWRKYCRGGKCAGMWCCYCRRRWYRMYGAGTGTAG
            ** ** **** .** ** ..   .    ** *  ..:    .       ***   :. .

38          RGARTYTGRMRYYKRCTKTAYCTKSYYKRRCATCGKMYCSMKTCGKRRTGKRKCGMYYKA
45          RRRAWYYRRMRYYKRYYKWRYYWKSCYKAGACWTKGCYYSMKKYRKARCAKRKTAATYKC
            *    *  *******  *  *  ** **  ..     * ***.  * * .*** .  **.

38          ACGTTCGARYKSKTKMRRRKRCAAKCGTSYTTTKRGGTKAAKYCTAYYRYYMCWWTYKCG
45          GTAYYYKWRCKGKYKCRRRKRYRMKSRAGYYYYGGTRKKWMKYYYRYYGTTCYWWYYGMR
            . .     * *.* * *****   *. :.*       .*  **   **     ** *

38          ACTKCKYCCTGCGRACKTTWAMYRMTYKAKKRARACGKWKKGRRKKRYYMRMRKMAKMGC
45          WTWKYTYTACAAAGTYGGGTGMYRMWTKRKGRTGGTSKWKTRRRKKRCCMRCAKAWTAST
               * .* . ... :     .****  * * *: . .***. *****  **  *  . .

38          GGSRKTWMTCGGYKKRWRTTRTAARYRWRTGRYTCYRKYTCCCATKYSRRKMWWRKCAWK
45          AASGGKTAYMRRYKKGWGWWRWGCRYRAGWKRYGATRKYYYYMWGKYSRRKMTTRKTGTK
            ..*  .      *** *   * ..***    ** . ***      *******  ** . *

38          AKACTACTKWRKAKRKYKKYKYGKGKATRYSCRKKCTCATCTRKKTRGSTYWTAKGGTYR
45          TKCACGMKKWGGRKGKYGKTKYKKAKGKATCYRKKYCYCYYKRKKGRKSACAWMGATACG
```

```
      :*.. . .**   * ** * ** *.*..  . ***   .  .*** * *:     . :

38    KRAYRKWRTTGGRYTKCRATGCMYKKRGTGYGTTTTCWGKTWYWACCKRRRKAYRMWYYY
45    KRGYRKWAYYRKRCCTYACARMACKGARKRYKCYYKMTKKKWYWRYYKGGRKWTAMTYYY
      **.****     *  .  .:    *   . *    .   *.***   *  **   * ***

38    KYCGTRWMYMAKAYYYCAAGYKYTAMKRGKYRARAWCKCCRKTRAYRYCMRWRTGYWYYM
45    KTMKYGTCYMRKRTYCACRRCGCACMKASKTARATTTTTTGKWRGYRYGMRWGYRYWYCC
      *       ** *  * ..     :.** .*    :  .   * *.*** ***   ***

38    MKYTYKWRWYRRYAMTRRGCGRWGCRKRMKAKTTATATWRSYKWYYGWCRWKMWYCTYRG
45    CKCWCGTATTGACTMWRGRTRGARTATGAGWKGAGCGAARGYTWYYKWTAAKMTYYYYRR
       *           :* *         .    * :. .: *.*.*** *   ** *  **

38    RTYGRACTKYYCRRWRKAAYWGYKTMKAGKRGYCYAGGYTTGKCAKKAAAKTYCGYYMTT
45    RATKARYYGTCYARWRTTTYWKYKCCKCTKRACMYCARCAKRKMTKTWGCKCTMRCCMGA
      *:           ***.::** **  *. **.  *..  :. * :*. ..*      * :

38    CKMWKKSYRKKKKYTGGKCMCGGGSYWRCRMYTTKAAMKTKTAACRRMYYYYRKYSAAWM
45    TKMWKKCCAKKKKCYSSKAMYRKTSYWRTGACKYKWGAKKKYRWMRRMYYYYRKYGMRTM
       *****.  ****  ..*.*    ****    . * . *.*    **********.   *

38    SRCTRYYYYYYYSRKRKRMARWYRYWWKKKCCAAGAWKMTWRTWKKYGWTTYKKCTAARK
45    CAYAGTYYYYYYSRKRKRMWATTGCATKKKTTMRRMTKCYWRYWKKYRTWKCKKTKGCRK
      .  :  *************        ***       *  ** ****   . ** ...**

38    CWCGRCKKGCCYYYRKSCAYAAMGKYYYAMYKKKYYKYTCMTCKGRGKYCYGTTRGWMAY
45    TTMARYKKATMYYCRKCTWTWGMAKYYYMMYKKKCCKTYAMKYKKRRGTTTRYWGTWMGT
         .* **.  ** **.    .*.**** *****  *  .*. * *          **.

38    YKRYMWAATGYWWGCYKRYCAYGYRYCSGYTRCMCWYMRSYWKKYATYTRWRRKYCCCTC
45    YKRYAWGCCTCWWATCGGYMTCAYRYSGAYYRYMYWCMRSYWKGCTCCCATRRKYTATCT
      **** *..   **.    * : .***...* * * * ******  :     ****  .

38    YKRKMMCGKMRYWWKKWWSYKYYRSMTGMTTAGYCGATMMCGKCTRCGTTTYRKKKGGGW
45    CTGKCMYRKMRYAWKKAAGTKYYRSMWRMWAGRYYKMCMMTAKMYRTTCCCCRKKTKKRA
       . * *  **** ***  . ******  * :. *    ** .*  *      ***.

38    GMTRKTGGTAACGMRRGTYRGRKKWRYKYCRWMWYMTGTARARAYYWCKTAGRSYGCYYA
45    RCYGKCTACWRTAMRRACCRKAKKARYKYTATMWTCCACGRRRTCTASGWRAACCRSYTW
          *  .    .***.  *  ** ****   **   . .* *:   .   . .  .*

38    GRWKKMYCYYYWKRGTGSCTKGAKGGCCYRKCGCWGAARGRGWYMAYKYKGGKTTGCCYC
45    RRTTKMYMCTCAKGKYSGMWGRMGKSATYRKAKYTRWGAKRKWYMGYKYKKSKYYKTATA
       * .***     *   ..       .. ***.     .  * ***.**** .*    . .

38    TCTGGGCRYRRCGTTTGYKGCATCKGRYATYGWGTYRYACKCACACAAYTARCYMTRYWS
45    CTARRRYGCGAMKKYYKYKTTTWMKAGCRKYKWKKYRYWMKYGMRYMRTYMAMYMCGYAC
        :          .   **  :  *.   .* * .***  *  .          **  * .

38    KSKKWSKYYYWRGWKCCRWMGGTTGMMRWKRYKGAGGKKAKRTCCCCTRCRAAKYMRYCK
45    KSKKWSKYTYWGKTKYYATCATAYSACRTKAYKTGARKKCKRGTSTTYRMGWRKYMGYMK
      ******** **   *     . : .  * * ** .. **.**  .   *    *** * *
```

```
38              KKYKKWMMKKACRKCRAYCKAGAACRTGGCMKWKYRKRCGCKCYTGYKKTYYSYYYTARM
45              KKCGGWACGKRMGKYGCYYKTAWGYRCTATMKWKYGKGYRYGMYAAYTKGTTGCCYKMGC
                **   *   *   *  .* *:. . *  . ***** *      *:.*.*   .  *.

38              KKKKRYGRKKTRMYYKGKWGGWWGRRWKCACGTMWWTTATYYYRKGYCMTCRYKKKMRRK
45              KKKTRCRRKKKRMYYKAKASAWWAGATKYRYTACAAYCWYCCYGKRYSMWSRYGKKCRGT
                ***.*  ***.*****.* ..**.   *    :         * * *.* .** ** * .

38              YGRTCAARKKMMKGAAKYYWKRYCKRYWWGAAKCCGYMRKRATMGTGRMKGKRGYRKRCA
45              CRRWYRWGKKCMKTTTTYYWKGYYGACTTTWWKTTTYMRGAWYCRKRGAKTKRAYRKRYG
                  *     ** ** ::.**** *         *   ***      .   * **.**** .

38              ATGWCCMCMMRGMKGCTYAARAKYGCCCWRKRWAYRYYATKKKKKAAAKMACCKCGWRKR
45              GCKAMMASMMRKMKATGYRRRMKCKYMMTAKAWGYAYYRAKGTKKWRRTARMYKYKWRKR
                .      .*** **.  *  * *       * *.* ** :* .**   .    *  ****

38              KAYACYTAKMWGYKSRGYGCYKKWWGTACAGTRKYKKCGGAYYYAYGWTRTAAYKAMYWR
45              KGCTTCATKMWRCKGARYRYYKGAAACCYWRCGKTKTSSRMYTYRTTTWGWRTCKCATWG
                *. :  ::***  *.  *  **   . .     * *...  * *        : *.  *

38              CCK
45              YYK
                  *
```

The pairwise alignments of two accessions 38 and 46

```
38                  YMKTTRKWMYRKWWWSMWYMRRKYWRYRYGTKRGYRCRCCGYRKYYYYYRRGGAYMWSRM
46                  YMKYYRKAAYRKWWTCCWYAAGGYWRTGYCCKGRYRYRTTTYRKYYYYYAGTAGCCWSGM
                    ***  **  ***** . **    ***  *  *  ** *   ********   ..  ** *

38                  MRMWRKYRRRYMKWRRAYTRMKKYYSRRMAMYMYYMKRKKRSGSYYRKWWRMYMRMRRST
46                  CRAAAGYRRRYMKTRGRYYRMKGTCSRRAWCTMYYMKAGKRCKCYTGKTWRACAAAARCW
                     *    ******* *  * ***   ***    *****  **. .*  * **      *.

38                  RKRRACARSMMRRGCYRKSYCTRYKTRYMRSYTKGGGKAAMYMYWYRRYRWYYWYRAYCK
46                  RGGGRTTRSMMRGRYTGKSYMWGYKKATCGGCWKTRAKGMMYCCAYRRYRWYYWYRTTYK
                    *     :*****     ***   **.    .  *  .*. **   ***********:  *

38                  MYCCTAACRMYTKWMRRRWRYRCMAAYYRMKSYMKSYRKRYYSYTCYGAKYRAYGRRKGR
46                  MCMMYMRGRACWKWMRRRAACAMMRRCCRATGTMKSYRGGYYSTCAYCRTCGCTKGATCR
                    *       *   ******     *    * .. *****  ***  .*  .  .    . *

38                  RRSYWAYKTGYRMRTRSRWWMMRWTRRTGTYMCAYYRMYCGTTATCTTCYYSRYYYWTYR
46                  RRSYACYKCAYGCGWRCGTTMMRWWRAAACYMYWYYRMTMKAGRWMWAACYGRYYYWCYR
                    **** .** .*    *.   **** * :. **  ****   :     :. *.***** **

38                  RKRSYYWKYYYYMYCMACYKRMRGCRCYAARRSYKRTWMWTRYRWWAMWTCRCGSYWWGR
46                  RKRSYYWKCTCTMYYMRYTTRMRKMRTCGMARSTGGYACWKGYRWWGATWSRMRSYWWTA
                    ********    ** *   .***  *  .  **      *. ****.   .*  ****

38                  YRWKYRCWRRWYARWSRKYYTWRYYYMKRRYGYRKCGMRYMMRCRYSMSWTCRYYRMRGK
46                  TGTKYGYWGAWCCRWSRKYTAWRYYYMGARYKYRKAACATCCRSRYCMSWCMRCTRMRKG
                       **  *  * .****** :******  ** ***..     *.**.***  *  ***

38                  ARRRYWGYCGAYKTYRCTMRRYCTCYCGRKYCRAWTYAAGYWYMRYMKWRRRWYYYRCSY
46                  RAGGCWKCYRMYGWYRSYMRRYACTYYKRKCYRWWCTTCRYWYMRYMKWRRRTCCYGTSY
                         *     *  **. ****.  *  **  * *  :. ************   *  **

38                  RTCMRYRYKMYRYMSCYYRRCYKYGYASATYTKYKKKTYGGYYKYTGRKRKTGTTAKTAC
46                  RKYAGYACKMCAYMSYYYRRSCKYKCGGCCYGGTTKKYCRKYYKYGTGGAGCAWYTGKWY
                    *.   *  **  *** ****. **  ... *   .**    ****       .  : .

38                  TCARKKKTTAGMCRTTGAYMRKRAKTCRKGATRTMGYSGKYMATRRTKYTARYSRTAGCT
46                  GTTGKKGGCGAAMRWWKGTCGKRRKCTAGRRYRCCTYSRGCMWKRRWKYGWRYGGKTTTW
                      : **   ..  *   .   ** *       *   **   * .** **  **. .:

38                  CCCYCTACACSWSGKYRKRATGWAYGRMRKRMGYYGGYCYRTATMKCTGGCGSTTATYGC
46                  YYATMCCTMYCAGAKYATGRAATRTRACRKRMTYYRKCYCAATCMKAWAAYASWGCYYRY
                      .   .   . ..** .  :.      **** **      :: **. .. .*  . *

38                  RGYRKKGRAKKTKTKACWTTTMWCRTTWCYYTACTAAAMWCTAYCKMCCGYRKRRCKCGT
46                  GRYRKTRGTKTKKWTMYAYYCMWYRKKWGYTWGYCRRRATTAGYAGMYYRYRKGGYKTTC
                      ***.  :*..* .      ** *..* *  .        :.*. *   ***   *

38                  RMRWAYYMYRTCAATTTCMYYGARYRCTWWGYTRRRKGGGAAYKYGMGAMYYRKAWKRYK
46                  AAATRYYMYRYYRRKWCTMCTRMRYRTWWWRYKAAATTAAGRYKYKAACAYCRKGWKRYK
```

```
             *****    .   *    ***  ** *.   . ... ***  .. * **.*****

38      YSRRYRKKCTRRKWWYMKWYTGKSWYTGRKRYYWRRTRRGKYGAKTKCRAATKRTTYYWY
46      TCGRYRKKYKRRKAWTMKWYCKKSAYAKGKRYYWRAYARKKCRRKYTYAMCAGACYYYTC
         . ***** .*** * ****  ** *:  ******   * *   * .   .:    **

38      GTKWYGGGACAWACCATARRAYRMWGRWYKYCRWMMAATRKAYAKYCARRAGYKKYTAMK
46      RKGWYKTTWMWWRYYGCWGRWTAMTKGTCKTMGTCMRRKAKRTWKCACRRRRCKGCACMK
         . **      *   .   *   *     *     *  . *   * ..**   *  :.**

38      RGSRRWKCAGTCTKKTARYRYCMRWCCYWYYTRMMKGAWATRRRWYRRYAKAMKRRGSRR
46      ASGAAWTYWRYTYKKWRGTRYGMRWSYYWYCAGCCKKGWCWRRATTGACTTWCKAGRSRR
         ..  *.      **    ** ***. *** :   * .*. **      :.  *   ***

38      GRTTRTKRMGKMKTTATRRSYRWGAATYRKMWTAWYRCCAYKYRTRRGCYKCYKARKYYC
46      RACKRCTGAAGMGGAGCRRGYAATGCCYRKMTCGTTRYYRTTYRARRKSYGYCKRAGTCT
           .* .  . *  :. **.*   .. ****  .  *    .**:** .*   *

38      AKKKMCKKRGMAGKRWMRYGAARCGKRCKYRWYAGTTRMMTARYKRTATGGAGAAMTCTC
46      TKKKMYGKRRMWRKRTAACSWRRYRKGTGTRATGAGGRCAYTRYKRCWYRRRSMRCYYKY
        :**** ** * **    .  *  *    *  ..  *   :****      .     .

38      YYKTYKKTRKWMKMAGRAAGRKWSRYRKGGGGTGYRYYYKTGCRGAACCYTAWCKYWRYK
46      TTTWCGGKRKWMKMTRRMMAGTWSRCGGTRRRGAYRYYYKYRYRRRRSMYYMAMKTAAYK
          .    .******: *  . .***        .******   *   . *    *   **

38      YTGKAAGWGAASWAWTCAWWTRYMMWAYMYSMWYYKTRWRYGYYMRTCAACTTSWKWTRT
46      YGAKGWKARTGSWRWATGWACRCMMWRYMYCCTTTGYRWRYRYYARCYTGYYKCWGWGAA
        * .*.    :.** *: .*  * *** ***.      **** ** *  :.  ..* *  :

38      RKRWTKGMAATKWKSMMYGARCYSGRYRCYRRRCKACWAKKWYKSMCWKMMRATCYRYYY
46      RKRTYKCMWRWTTTSCAYRRRYYSKRYGMCGARYKRYTRKKWYGCCSWKMMGWYTYRCYY
        ***  * *   . .*  *  * ** **     * *    **** . .****    ** **

38      YAYTCCKYKKKTAYYKYCYMMKKWYAYRKRYRYAGCGGRGRRSAKKKSKRCRYMSAGTMK
46      YTTYYYKTKKGYRYYGCYYMMKKWYRYRKRTAYRKTARATGGSMKKKSKAYACCSRRACK
        *:    * **   **   ******* ****  *   .     * *****     *  : *

38      CMACKATKRYMYSRYYKAYCRAWTAKKKGGGTTRTRTTCTMWRRMKRMRYMRKAKCGYTC
46      YMWTGTKKRCMYSRYYKCCYGRTKRKKKRRRKYAKRCAGGCWRRMKRMRYMRKCKMKYCT
         *   :.** *******.     . ***   .  .* :   ************.*  *

38      GMYKYKRKTCGTRRYKAGMRKRKKATAARKAGMKYRRYMKKCYRCYCMMRKCTRRYMGRM
46      AMYTYGRTWYKCRRYGGRAAKAKKGCGGGGRRATYRRTAGKYTGYYYMMRKTAAATMRRC
        .**.* *.    *** .   * **. ..     .***   *    * **** :   * *

38      KCYMYKRRCYKMATTYYKKKSWYSRKWGCKYYGCTGKKMGRRWYMCKGMYYKATTTMWKY
46      KSYCYKRRYTKMCCCCYKKKGAYSRKWRYKTCRMYRKGMSRRWYATKAATTKCYYKMWKY
        *.* ****  **.   ****. *****  *      * *.****  *.   *.  .****

38      WKSTRMGRRYMAAAAYCGSYTTTYCKTCKTYWGGCTTCTYKYMTGYYMYWWWTKCKRCKR
46      WKGYGCKRRYCRMWRYSRGYYYYCATYYKKYWSSYKWYYCKTCCACYMYWWWKKTKRYTA
        **.    ***     *. .*    ..  *.**.. .    *   . ******.* ** .
```

```
38              KGAGCARCACMRMYKKRKTACAYAGRGKTAKKWKTTWKAYKYMRKKKMACYMRGKAARYR
46              GRTATCGATYMRMTTKGKYGTRCMSRRKAGKKWGKKAKGCKCMRKKKAWTYMGRGRMRYR
                  :.  . .: ***  .* *  .     .* *:.***  .. *. * *****    **      ***

38              WTRTCRTCYTCRMKYYGGGCKWKWWRCACWKRAKAKAKRSCWTCACKARKKKRKCGCCAC
46              AGGWYGGAYYYRMKYCKAAAKWKTTRTGMTKRRKWKGGGCMWYYMTTMGKKKAKMKTTGT
                      .*  ****  ...***  * .  ** * *.  . *     .  *** *    .

38              TSTWGGYWSTCKKYYYWCYKTRTTWRRCTGCYTYRCRCYKWKCSRYWCGCMTKKYCCCGC
46              ACGWRACTGYMKKYCTTMCTYGWYTRRYCAYYWCRYGTCKAKYGAYWTTGMYKKCYYYRA
                :. * .  .  ***       .      **  . *  *     * * . **   * **      .

38              RGYMTCCCCRGGRKKKRMKRWGGRYKKTMKTARCGTGGYCKKKKMAARGGWWCTTKCAGG
46              ARYMGTTTARATRKKKAAKRWKKRYKKWAKCGGYKCRTTYKTGGMWWGKKTTMYYTYRRA
                  **     .*. ****  ***  ****  * .         *.  *            .   .

38              TRACWAKKRMKWYRKMAGRASGKYYGKRCTTTCARRTKKSMYCMSGMYRYYWYKACTMMA
46              WGTYWCKKRMKACAKMMKRGSTKCCRGGYYCCTGGGYKKSMYMMSTMYRCCWYKTYWMMG
                  : *.*****   **  *.* *           .   ***** ** ***   ***:  **.

38              CGTAMRYRYKKTATCWYCRGYYMYRWRCYRKGGAKSRRYKSKYRYMKAAKAAATRYKRYC
46              MRWMMRYRYGKCGCATCAAACTAYRTGACRKKKWGSRRYKSKCRYCGGGKGGGYGCTRYT
                    ***** * . .  . .    **   . **     ******* **   ..*...    .**

38              CATCTMTACTYYRMTGCGARSTKMTTYKTATKRGTYACTCCAKGCKTCCCCAGTTTYRWM
46              TGCTCCKMMCYYRMAATRMRCCKCKAYKAGAGGRYCRYYTARKRATCAMTMRRYWGCGTC
                 .    .    ****:.    *. * .:**:.:          . * .. .

38              GTGSGYKCCMCYRYRSRACGWKGRMYAYCAKRCMRRYWYSRKKTCGRCAARKSTTCWKCY
46              KYRGATGMMCTYRYGGGRYTWTKRACGCYGKRSCGACTYGGGGWMRRTGMAGCCGYWKAC
                   ..      ***  .     *. *  .  .**.      *.       *  .    .   **.

38              YAWYTRAYWARMMYAKKYYCYWRAGAACGAGGAAGKYTRMTCRWYATWCRGTRTYGGCGT
46              TGTYWGRYWWRMMCWKGCCYYWRRKRGTAGATRMTGCKRMGYRACTWWTRRYGKTCCTTG
                 . *   ** ***   *     ***     . ...       .**   *   : * *     .

38              TGKKACKYCGYTGAGCYYYYTGTYTKWCTMGAGCTAYYARCTYMMCARCKWYWYGRCTAR
46              CKKKRYGYYRTCRCAYYYYYWKYYKKWYKCRWKYYRYYTRYYTMMSMRTKWTTTAATCGR
                  **    *      .. ****    *.** .         **:*    **. * **     .    .*

38              RRRRKRMYMRKMASYMWYWTCCKCGCYYYRRYRRYRWGRRYKGATAATCCRATSGKMKGR
46              RRGGKRMTCRGAGSYMWYTAYYKYKSCCTGGCRRYRATRRYKTGYWWWMMRWACAKCKSA
                **  ***   *   .***** :   *   .       ****  ****  .        * :..* *.

38              KMYGCCTCRTCCGRCRKAYMTRTTRATGGTRMCASYWRRRAACGKKGKRWAGTYKGRYGG
46              KMYKMTGTRCTGKGMAKWTAGGCAGWGAAGAMTTSYWRAGTGAAKKSKRWTSYYKRRYRK
                ***      *       *      :    ..  * :****   :...**.***:.  ** **

38              KRGCGCWSYWKGCGGYMAYGRRRMYYKYRWRWSKRMKKRYGCAGRCTAYKTTKCYYRKATA
46              GAAATTWSYWKATARYATYRAAGCYTGCGARTCGRMKRYAYGAAYATYGYYKSYYRKMGR
                  ..  *****. . * :*      *      * . *****. ..   ::*    *.****
```

```
38              CKYGCYWGWWCTGKRCCTRWACACMGYYWKKWWKGYCRKKKGTCAGAKKMRRRKTKYTCK
46              SKYTMYAAWWYWKKGTTAATWYRYMKYCTGKAAKRYSGTGKRWSRKRGKCGGAGKTCWYK
                .**  * .**   *   :       * *   *  * *. . *  .    *     ..  *

38              AAYRGKMKCCAAMKAAKKYYWRRWYCAGYAYMCCTRRATYKGGAGATYGRCGGWGCMMRG
46              RWTGKKCKTTCWMKTTTKYYWRRWYYMRYWYMYYYARWYTTKTGACCCARMAAATSMMRK
                     * *  . **::.********   * **     *    .  ...  .* ..  .***

38              MAKKAARKYCCCWWRKRWYGAKATGGKRGKMACGRMWRCCKCCKAWCTMGCAWYYGYWCK
46              MRKKRRRKCYMMTTAKAWYTGKGGTRTRKTAMYKGMWRYTKATKGWGYMRMRWYTRCWYK
                * **  **        * ** .*.    .* .     ***  *. *.*  *    **   * *

38              SRYTGGCGYSYCACKGAYRKMYKYTGGCKAAGAGCWRYYRAWGWRKACYSRYYYYMYRGG
46              GACYRRYKYGTYMYKRGTGKCTKYWRRSGWRRWRYWGTYRWWKWGKTYCSRYYYYATGSS
                .       *.     * .  *  **    .        *  ** * * *:  ******   ..

38              AKRKTTCAKTCKWRRKMYSRKCWMKYAKTRCAWACMRCWKWRSAACWCTRRGGCMWAYAA
46              WKRKKYYWKWMTWRRKCCCAGTTCGTGGCGTRTWTMAAAGAGGTGYTYKRRRATCWGYGG
                 ***.    *   .****   .         .         * .     .:.   .** .   *.*..

38              GCCACSAGGACCYYGGGGCMKRARKGTACKCCGYCCMRMTKMKYRCTGCYKYGAARRWGA
46              ATYWYSWKKRYSYYTKKRYMGGGGGTCGTGSMAYYSCRMCKMTTGYYRTCKCKTGRAWAG
                .    *     .**     *   .     .  . .* . ** **.        *   :.* *..

38              TGGAKYACKYTMYRYRYWKWRRYWYACAAGAGCCMGWYRTYKYCGYTCRKMGTRRKCCTA
46              ACRGKYRYKTKMTRYRYWKTAGCWTTTGCTGKYYMRTTRWCKYMTTAAGKCSYRGTTTGT
                :   .**  * .* ******     * : .. .    *    *  **    :. * . * .    :

38              KGGGGGYCCTWAMKGACCCMYAAYCYATATGAMCAWGRKAKGGCTKCATAYTCKKCWRAY
46              KRRRRKCMYKWRMKRMSTACYGRCMCWWCCRCMAMTARKMTKRYKKYRWWYYMKGYAGCC
                *         .* **   . . *.        .  .*.  .** .     .*     *   *     .

38              WAAGCTRGMKWYATYRYYRCWGWRARGGKYRMCTCKGAYRRYKTYGAGCTRAMRRCSKRY
46              TWRATAAAMKWYRWCRYYAMWRTAGRKRTYRCGWYKTRCRRTKYTAGKYKRWMGGYCKGT
                   . : .****   ***   *    .*   .**      *    ** *    ..   .* *    .*

38              YTAAKWRWGAAMYAYARMATATWAGKGGRWKRKKCGTRSYSKWKCYRKKRTTTTGCKTCA
46              CCGTGAGASMRCYMYCGCMYRYTRKTRRGTKRKKTRKRGYCKWKYYRGGRWKKGTMKYYR
                  .:     .    * *.         .    ****   .*.*.*** **   * ..    *

38              TRMYYMWTKASAWTGTRYSMTRTATCKKYWKKCYACCMKRGTTTTGYWWSGCATTYRCRC
46              YRMYYMTCTGSMWYSYGTGCKAKCGTKKYWKKYYRTTCKRACCYCCYWWSKMMWWYRTAM
                 *****   ..* *  .     . . ..  ****** *     **.       ****      **

38              CTCATGWYRRRCYWYWWGGACCGKYGTGKKARRMTMTTSTGTTTWCGSYTCTKTKKCYKM
46              YYYRYKTCGGGTYAYWWAKGTYSKCKKRKKRRRMYAAKSWKKKWTYRCYGMYKGTKYYKM
                            * ***. .   .*   . ** ***  :.*    ..     .*    *  .* ***

38              SYKGATGTAWWSWTGTTRRWKCMYKRMWKCTGCYSRWYTKYCAKYMWKYCRKTCCTMYWY
46              SCGRMCAATWWSWWKCYRRWKTMYKRMWKTCAMYSRWYAGYTTKCMAKTYGKAYYGATTY
                *      .::****     **** *******   . *****: * :* *  *    *:       *

38              ATTKCKSATRTKGWWTRKYTAAARKWMKMTRCRRSSGGCTRCTYTGTCYTWMAWRKRAGK
```

```
46      CACKTKSRYRWKRTWWRKYYWRRGGTMGCCGSARSSRRMCGTGCWRKYCYWMMWRGRGRK
        .: * **  * *  * ***       *    . ***          .   ** ** *. *

38      KRMKACMKRYMYRWRRYRYCAGGCGCCYWSRTTYCTCCMRKGTYGACKTKKCRCRYTAGY
46      KAMGGTCGRYMYRWRRYRYTTRATATGTWSRKWTAAGYCRKRYYRRYGCKGYRYGYWRRC
        * * .   *********** : . .   ***.  .:   **  *     *  *  *

38      YKRRKYRAAARKYCGTCGKMKAMYRCWYRGKKKRRRRAAACTTTKYTSKSRRAMCTWTGT
46      TGRRKYRMTCRKYMTGAKKMKGCTRYTCGRKKKARGRRGRMCCAKCGGTGGAMCACWKKY
          ***** :.***   . ***.  *     *** * * .    :*  ...    . *.

38      RYYYSWYMYYYYKGARYGKYWYKAGTWCRTGAGCYKCAWKAAGTTCATTYCYYAMTATTG
46      RYTYSWYMYYYCGTGATKKCWYTCKGWGRKACAYYKMTWKWGCWYYMCCTMCCWAAGCCK
        ** ********   .   * **..  * *.... ** :** .             :.

38      MKMYWKRKKRKWKKKCTGTTTGGGKCGACTGTRCTSYWRMCYTGGCKTKTKCYKYGACKT
46      MKMYWGAKKGGAKKKAWRYYWRKKTTATACRYGYWSYWRAYCCTTGKKKKKACGYAWTGY
        *****  **   ***.        . .:.      ****       *.*.*.  *.

38      KKMRAAWMGGRKKTYGTCCTTYRRMAKRYWCMKTRSCTYYSRKYKRRKYRRMTRRMTCAW
46      KKMRWGWARARKKYTACYYYWCRRMRGRYTYMKGACYWYYSRKTTAAKTGGCAARCCTWT
        **** .*  .***  .      ***  **  **  .  ***** .  *    : *

38      CAGRWKKKCCGRTGCTTKAWKMYWRTWKKKKCCRTWKCGGRRGTRYWKKWWSYRYKWMAY
46      TWKGTKGKTTKAATGCGTMTKCCWRCWKGKKTGRCTKMACRRSWGCWKKAAGYGTKAAWT
             * *    :    .  *  ** ** **  *  * . **.   ***  .*  *

38      YRTATTGATAWTCGTYSMAYKGMRTASTYKRGCGTMTRMMMYGRGAYCTGTGYKKKKGGA
46      CGYMCAAGCMWATCATGMRCKKMRWMGAYKRRYTCMWRMMCCRGRGYTYRCKCKKGTTTR
             :..  *:  : .*  * **  .:***    * ***     .*      ** .

38      GRYGTKCGRCAYRWKRSWKYYWYRRYKYRKRWYMKTGTAKRMKCTGCCRRGGSTGGCGKA
46      RGTACKYRRYMCRWKASWKTYAYRRYKYRKAWTMKYRYGKRMGYKSYYRARRCWASYKKW
           . *  *   *** *** * ******** * **   .***  ..  *   . ..  *

38      SRMCTAWKKMAAMTGCYYYWKGRCAMGSTTGWKCKWGYYKCTKTKARWTGTACTRAGGMK
46      CAAMKWTTKMRGCATMCTCAGRGYMAKSGATWKTGACYTKAGKKKRRWYRYMSCRWKRMK
        .   .  .** . :             * : **    * *. *.* **    . *   **

38      WYMRTAAGAYKYKGCCCAGYRGTCAYTTTGGGYGWYKRYRYRSYMYTCRRCGRMKGYATG
46      WYMACGRKGYKYKKMTARTCGKKMWYAAATAKYKTTKGYRYRSYMYAMGRTAGMKTTMAT
        ***  .  .****   .     .  *::: . *   * ********:  * . **   :

38      KGCKRRMYYYKYRKYWRCTTTMGRRKYWMRKAKAYYTRTARCSMGCAYYAKYKYYKGAAA
46      GTSKGRMYYYTYRKYWRTCKWCKRRKYWCAGRKRCTYGYRGMCMSYGTYWKYKYYKATRR
          .* *****.******  .   *****    *         .*. . * ******.:

38      AGGRTGMRACYWWYCYAAWRKYAATTKTAAKATARYKGCGCAGYYGTRKRRRWGYSKMWT
46      WKRGWAMAWYCWWCMYMRWRGYWMYYTACCKCGGAYKKGATGRYYRCGKRRRAAYGKMWY
             .*    **  *  ** *    .:..*. . **  . . **   **** .*.***

38      GAAACTRKCGYRRRRTTTYRMRYYCATGSAGARGTRYWYTKCYRGKYCATCAGMMKWSKM
46      KWRWYYRTYRYRRRGACATRMRCYMRYKGGKTRRKRYWYCGTYRRKTYMYYCACAKWSKC
```

```
              *.  **** : : *** *    .. :* .****   ** *     ..  ****

38      CKTCTCACKMTCAMSKKMMKRRCMTYRRKCSGWYKKCCWATTYYCRRRYMTKYRKYYTYS
46      TGYYAAGYKMCTGMGKKAAKAGSMKTARKYCAWYKKATWMYWYYYRRGCAWKYRKCYKCG
            :.. **  .*.**  *  .*.  ** ..****. *   ** **    **** *. .

38      RSRMYYKRMCRCRKRYKKCATCYKKTRTTAGCACGTTAKKAMMWMMTYATGCTAAMAMRY
46      GGRCYYKRMYAMAKRYGTATGTTGGGAGAGAYGAAYYWKKMAMTMMYYRKCYYRRMWAGC
         .* *****    ***  ..:        :.. ...   **  * ** * .     *

38      CCTYKGGTYYYGAKWYGATYTWGGCCKYCACCKAKMRCTCYRRTRCMMYYAYRYYYRKCT
46      YYYTTARWCYTRRKACKGCYWWRKYMGTYWMYKTKCGTGTCRRWRMMMYYGCGCCCRKYY
            ..   *   *   . * *          *:*      ** * ****.     **

38      TTAKMAGATGTGTTTKWACKTTRMATGTRMWARRCAGTCAGYKYARRGAAWAMMTRMTTA
46      GGGKAWACCACKYKWGACMGWWRMWKRKRCARRATTRWTCRCKCCAAKGGARMCGGAKKM
          .*  .. .   .   .    ** . .*   *  :   .  * .   ..  *    ..

38      AYGYKRAKWKKAKWMMRWRKCCGKTCGYYCYGAAAWMYWAYKKYYKYCGKCYCYRRRWRS
46      RYAYKRMGWKTRKWMMRWRKMYRGCTAYCGCTRTWTATAMCTKYCKYYRKYYYCGRRWRC
         *.***  **. ********      .*     :       .** **  * *   ****.

38      CRGMTAYSAGRGAMYKMYGKYKYTWCTCGKYGGYGTTRKGWMAYKTTGAAWKKRKYKYKA
46      TRAAKWYSRARRMMYKCCRKCTTYWAKYSKYKRTRYWGKTWMGYKYYSRRWKKRKCGYKM
         *. . ** .*  ***   * .  *.. .**       * **.**  .  *****  **

38      ASGKRTTCSSACMCCYRCMYMRSYWYSCKYWRYRWSRAKKYTAYKMAKCMTAKRGTRTTY
46      WSKKRYAYSSGYMAYCRTMYARSYWYSYGCTRYGWSRMKKYCCYKCRKMMKRKGTKGCYY
         * ** : **. *.  * ** ******    ** *** *** .**  * *. *  .   *

38      KRYRMYKCYRRCRKGKTATKWYCRKKATTCTYARYGYMWKTKYRCKAGTMKKGYYRCACR
46      KGYRMYKTCGRYRKRGCGAGWYYAKKWYYMCCGACRTCAKYKYAAKGTCAKKKCCGGGTG
        * *****   * **   .: **  **      .      * ** .*.   **     .

38      KRKYSCWKYRRAAKKKTKRTTRRGGMRRYKYWKRMCAYCCRYKYCKKKYYYKGAGCGRMT
46      GRKCSTAKCGGGGGGGYKRCCRARKMRRYKCWKAMYWYYMRYGTGGKGCCCKKRATAGMK
         ** *  *   ..    **  *   ***** ** *  *  **    *    *  . . *.

38      GAYRYRRCYWMTAKTKWRWRKKTMTGCYRATYWKRMRTCCTYAAAMGCKRRWYCWYCATG
46      ARYRYRRSYTCWWKWKWRTGKKCCYATCACCYWKAMGCTACTGRGMKYKRRWYYACAGGA
        . *****.*    * ***  **   .   . *** *   .  . .*  *****   .. .

38      MCCCYYYGACMRKRRMKKCKATMYWTCACYGTYKGKYTTAAKSKYKTARYMTAGSKRRSK
46      AYTTYYYRRSMRKRRMKKSKRKMYWKAGACKKYKKKTKYRWKSKTKYMAYMYWRSKRRSK
            ***  .********.* .***....  .** * .   *** *   **   ******

38      YAYWKRYKRWSMCRGMWAKRYGRKKAAMRKATKGGRYRCACYRCCKYCKYKCMKWKGCAT
46      YMYWKGYKATSMYRSCTTKAYRRKKWRMRKCCTSCRYRMRYYGYMKYMKCGTAKWKATRC
        * *** **  ** *.  :* * ***  ***. .. ***   *   ** *    ***.

38      GGCKWKYRKRTTCGTGCKTKAYKGGTRSYYYRYYTTGTKAMKKKWYRYRYKRKKGSCRTR
46      TATKWKYGKGYWYRKRYKCGWYTKRGRSCYYRYTCCAKKCCKKKTYRYRCKRKKRSMAKR
         . **** *     .  *   *.   ** ****   ..*. *** **** **** *  .*
```

```
38    MYYKYYKRRMCRTWYGAGAGYGTKCKRKKRWKGAYKKKRRKKYYYMCMKKYRYKRWRTKM
46    MYYKTCKRRAMGCWYATTTRCRKKYKRKKGWKAGTKKKRRGKTCYCMMKKYGTKGAAYGC
      ****  ***    **.: :   .* **** **.. ***** *  *  ****  *

38    RWMRRGYCKYKGAAKRWYWWTKCGRGWMCYYGKKCSKASKYYYRTTAGTTMRRWMMAKAA
46    RTCAGKCYKCKTTRGRWYWWAKMAGKTMMTCKKKMSKRSKYYYGWKMRKKCGRWCMRKRG
      *       * * :  *****:* .   *    ** ** *****  .  ..  ** * * .

38    AGKSYKYCGTYTYRAAAGKRYGYAKAGKTCYKGRYYGKRKCTASGKWYCGTATYCCTKGT
46    CTKSYKCTKCTKTGRMRSGRCRYMKMRKAGYKARCTATRKYYWSRKTYTRCCAYYMWKKK
      . ****     .     . *  * *  *: **.*  ..**   * * *   .:*   * .

38    KYYRRYKKTCGYTYGTMCKKYKGYCKGGYYCWTYMKKRTKYTGTATYACKKKGCKWCKYY
46    KTYRRYKGYYKCYCKGCAKKYGAYMKRRCCTTYCMKKRYKYYRYCYYWYGTGAMKAYGCY
      * *****          .*** .* *        **** **   . *   . . *    *

38    KYMMKGGRKRWSWGKKWGSGKYYCGYYCYMRTKRRGYCKTACGGAACGAGYCKKGGKTTY
46    KTAMGRCGKRWSWRKKWKSKKTTGACTYCCRYKRGKCAKYRMKKMRYRRRYMKKATKCCY
      *  *    ***** *** * *   .     * **   .*           * **. *  *

38    GGGGCKMYRCKARKMRKKKAKKACWWGKMKRKKRATKAKTACRAGATCCRWTGRCYYRKG
46    TARSYKCTRMKGGKMRGKGRKKRYAWATCKRKKAWWKRTWRYRGKCGYGGTKARYYYRGR
       . . *  * *. *** *  **   *.. ****   * .   *. .     ..* ***

38    RKRCRRYSYMYYRKYKGGKYTMGWWTGCKKTCTKAYSKCAMCTKTTGKTRCATYGAAACR
46    GKRYRGCGCAYYRKYKRKGTGAAATATGKKYSYKWCCKTGATCGGCRKKGMGWCKMRWYR
       ** *  .  ******      .  :  ** . *  .* .       *.  .       *

38    TMYWGKYAAGGTKRARRMRYYKCGARKTAYCTYKRRG
46    KMYWAGCTGTAWGGRARMRYYKMRRRKWRYYWYKARK
      .***.  :. .     ******   **  *  ** *
```

The pairwise alignments of two accessions 38 and 47

```
38      RWYMWRMMRKKTYRRRYMYRWRYMYMKTKYYSRRMKYYWWMYMRYYRRKAKYSKKRRMMW
47      GWCCWGMCRGKYYRRRYMYRTGYMTMKYKYYSRRAKYYWAMYMRYYRRKGGTCKKRRMMW
         *  * * * * ********  ** ** ****** **** *********.  .*******

38      KYRRCCTAKYRYWKAKYYTYCKYSRMYMGYGKSYKMRRWMKRRRRYTGYKRYRTMWTTMY
47      TYRRMYWGKYRYWKRGCTCTYKYGRMYMRTKKSYKMRRWMKRRRRYWRTGGCAKCWYKAC
        .***   .******       **.****   ***************       . * .

38      KWTYWMRYMCATCYGRWRYRGTTRAAAAWWRYYRKSYKYRKRCKWKWRAKGYRWYTYWWG
47      KWWYWMRYMAGWYCRRAACATCARRWRRWWRCCRKSYKYRKGSGATWRGKKTATCYTWAR
        ** ******..    *      :*    ***  ******** .  .**.*       *

38      YWRTYCRYATCYTRWWWYKYRYACYWWYGKYRAMMTMYGGRRRYAGGGYGYRYKRYYYWW
47      YWRCYTRYTGTCYGTTTYGYRYRYYWTYCTCGCACWMYKKRRRYRATTCRCAYGGYYYWW
        *** * **:        * ***  ** * .  .   **  **** .      *  *****

38      AWARMYYRWTYRCTATSYWRWKRWYWYMYMRMRWTRMYWWCRKKTMCATYRYCYGYYRWY
47      WTMRMYYRTGYRYAGASCARWKGTYTYMYMRMRWWAMCWWTRKKKCMMYYGTYYKYTGWY
           *****  ** :.:*  ***  * ********  * ** ***.    *   * *  **

38      RAKKMRYYYRRCKRMKCCYRYMRRRGYMKRWARWRGCCCTKGYMYRGYRYYRYKMWWYRR
47      RWKKCRCYYRRSKGCKAGYATCGRRKYCKGWRGWRAYATCKAYMYRRYATCGCKMWWYAG
        * ** * ****.*  *. *    ** * * *  **. .  *.**** *     *****

38      RAWKRKGCYCKMMRYYKMKRWRMRYWSMWMYRGACKKRRMSCCYYYYRGYRKYYYRMKTS
47      GRWGAKTMYYKMCATTTMKRWRCRCTCCAATRRRYGTRGMSTSYCYYRTCRKCYTGMKYS
          *  *  * **    .***** *  .    *    .* ** .* ***  ** *  ** *

38      MYRRYMCWGRYCCWGTRYRWWMGGGRACCRRYMWRYRRRYCGSWCMKMSRKSSTRGRSRW
47      MTGRYMYWTGTYATKKGYRWWMATKGRYYGAYCWRTRRRYYRGTSMKMCAGCCWRARSRW
        *  *** *    .  . *****.        * ** ****  . .***.  .. *.****

38      YTGGRTTYYRWYMMKSRYYGYCTTAMTYYYSRKKCGTRKMKWYGYCMWTAYWGGGRYTTY
47      CATTGKYCYRWCMAKSRYYKYYYKCAKYCTCAKGYAKRKMKWYACMCAWMCWRRKGCYYY
         :   .  *** * ***** *  .. .*  . *  ..******.       *       *

38      WKKKWRTCTGGRTCRKKYTKCRAGCMYTKRMGRYYKWTCWYWACCRKKAYRRMGRKKKYC
47      WKKKWAYMYRKACGRTKCYTYACRMACKGAMRAYCTTKTWYWRYYRKKWTAGMKRKKKTM
        *****         *.*  .  .    .  *  * . . ***   ***    * ****

38      GYRWMMAMWATKAYMYCARMKYYYTMKRGYRRWKCRAYYTAKMTYRYCMRWCCAGCYWYK
47      KYRTCMRATRKGRTACACRAGCCCAMKASYRRWKTGGTTWRGMYYRYGMRWSMWKYYWCK
         **  *    .     ..*     :** .*****  .     * *** ***.    ** *

38      MMTAKYGTRWYRRYAMTTKRWKGSRMKMYRKTTATWRKSYKCWYYRWYRKMWKTYRKTRC
47      CCKMGCKWATTGACTMAAKRWKRSRAGMCRKGAGCWRKGYKAWYYAAYRKMTKAYRKYRY
          .           :*::**** **  * ** :. ***.**.***  **** *:*** *

38      TRRCYKCYTKARKYYKATGGGYWRWYKCTTATAAKATARKMYKMGACYAKRYAKYKYRKG
47      ARRGCGTCGKWAGTCGGATAKYWRTTKGYKMKCGKGCGGKMYKMRMMTMGGCRKYTYRKK
```

```
                   :**      *      .: . ***  *  . ...*. . *****         **.***

38                 MYYYKWKAKAYYTTARRGGSMMTCYAKKYKYYAGRRRMMGGKTRCRAGRKRATWGTCGTT
47                 MTCCTWKRKRCTCYRGAACCMMWYYWKKYKYYCRRRGMMKRKWRMRGCRKGWKWCWTTGC
                   *   .** *        . .**  * ******. ** **  * * *. **  .*

38                 RCTAKAKCGYWRYYYRSYKAYKACTYYCACKWKWKTTYRCTARRTRRKMYAYMWYACWTC
47                 RTCTGRGYRTWGYYYGSYTRCKTYYTYSMTKWKTKWYTATCGRRKGRGAYGYMWYWMTKY
                   *  :      * *** **.  *:   *.  *** *      .**. *  *.****   .

38                 GCKYYRRYRWGRKTATATCCATKGTACARKCMYSRYRMYCCTCRGCCRCRKRATGGRGCY
47                 KSKYYRRYRWTRKYWYWWMMWCKSKRYRAKYMYGRCRMYATGTRKTGGMAKGTKRRATTY
                    .******** **         *..    * **.* ***.   *      * :.     *

38                 SYWRRRYKGAKGTKRYGKRMRGCGCWSYYMYAWKMYCAGRRRMYWRYARMKWSKKRYGKA
47                 SYWRAGYKRWKSYKRYRGAMRAATTWSYCMYCWKMYSWRAAGCTAGCRACKTCGKRYRKG
                   ****  **  *. ***   **..  *** **.****.             * . *** *.

38                 GCTAYYRTKYYYYYKYTTGMGYRKCYYYKMGTTMTKKKCKTTMGGCWCRYSYKSKRYKTK
47                 AYATYYRYKTYYYYKYWWRMRYRKYTCYKMSYYAKKKKTKYYMSRYWYRCSYTGKGTKYG
                   . ::*** * ******   * ***   ***.   .*** *  *.  * * **..*  *

38                 KTACCMKMRSRRCAKCYCGTSYTTTYCRWKYYYWKKYTCTKYRYMCTKKCTKCAGCAKCA
47                 KYWYYCKCASRRYRKTYSRAGYYYYCYAAKYYYWTKYWYYKYRTCYKKKTKKYTATCKMT
                   *     *  ***  * *. :.*       *****.**    ***   .** .* :. .* :

38                 CMRKTRMYKGAARTACCAYCGRKKWKGRRKKYYTKTTMYAKCGRWCWGGKRYRWARTRWW
47                 YMRTKRMTGRWRGYGTMRCYSRKKATRRRKKCCYKYYACWKTSRTYAKAKRYRAGRAGWW
                    **..**       .     .*** . ****   *     * .*    .**** .*: **

38                 TYKCKRYRYCYKWWRKCWKGYMKCTRSKGARKKYKRWYKKAAWRTTRKAYKKAAYKKGGR
47                 WYKMKRCGCTYKTTRKYTKRCAKMKGSKRGGKKYKRTCKTMRWRAYRTCCKKCCYKKRRG
                    ** **    **  **  *   * . ** . *****  *.  **: *.. **..***

38                 YTWTRSMYKYARKYRKRRRRYYYRYMRYYWGGGRGTRYGKKKYATCAGMMWKATCKCTYMA
47                 YCAARGMYKYGRTYRKRRRGTYGYMRCYARTRRRKRYAGKKTMYYCACAWKTYYKTYCAM
                   *  :*.****.*.******  * *** *    * .**. **    ..  **:  *

38                 CSKKMRYGRCRTCKMKYRRCSYKKCCATTYTMCRRYKMAYMCCYYWSGGMCRCRKKRYKY
47                 YGKKMRYKRYGKSKMKTARYCTTTATMYWYAMYRGCKARYMMTCYWGRTMYAMAKKRYGY
                    .***** *  ..***  * . ...    *:* *  *  **   **.  *    **** *

38                 KKYYRMYKKCATCRRTTYKKCKYKMYGTMCMTTACCCTYKMCGRKYGTCRAKYKKRWAYK
47                 GKTCRACKKTTGTGAGAYKKYKYKAYCKMYAKKMMYYYTTAMSRGCRKTGGKYKKGWGTK
                    *  *  ** :     :*** *** * .*  ..      .  .*   .  .**** *. *

38                 KKTTKYKRYRWRYKMRKKRTWTYCKTATRKRRYGRKKSKYYAGCTTCRCCGRWMRMRCTG
47                 KGGGKTKGTGWRTGCAKGRCACYMKCCCGKRRYKGKKSKYTCTTWKAGAAAAACGMGYWT
                   *   * *   **    * *   * * .  ****  *****  .   .. ...    *

38                 AYMKWKSYMCCTGTKGTYTRTAAATCAYMKGKKAGYKCAYYCRCKCTATCSGYYKWAYAC
47                 CYMKWKSCCATCACKKCTYRCGMRYYGCMKRGKMRYKGCTCARAKYYWYYSRYYKTMYMY
                   .******  .  . *    * .    . **  *  ** .  .*.*     * ***  *
```

```
38          YCTKGYYRRYYWTAMTWKMSKKYKKYWRKGYTKCRRKGYCYAGCYTKKTTCATKCYYKGY
47          YAAKKTYRRYCWGWCKWKMSKKYKKCWRKKYGKSRGKACSYCAYCWKKWYYMCGMCCKRY
            *.:*  **** *   .********* *** * *.* *. .*..   **         * *

38          KTWCSYRKKRKKKYTGGACKGGCTSYWRTYYATKTTMAGRCGKMCYCKTYYWCAARRMAK
47          KYWYCCAKKGGKKCYSSRAKRKYWSYWRYYCGKKGYMWTRMSKAYCYKKYYWYCCRRMRG
            * * .  **  **  .. .*    **** * ..*  *  * .*    *.*** ..***

38          RYAWKKRRSYYYYYYSRRRWRMWKRRARWKCKWGCAKKTKWRTWKKYCTKCARTWWCGGR
47          RYRTTKARCYYYYYYSRRGWRMWKGGGGTKYKTKYWKKYKWRYWKKYTAKYCRYATMACR
            **  .* *.********* *****  .  * *    ** *** **** :* .*    . *

38          YMRTYARGACCMMTACYTGCGYWWKSAKMCMTCKRGTGCTKMKACACYYYMGTTYATSTG
47          YCAYCCRSRTTACYMMYAATKYWWKCWKAAMAYGGAYATCTMTTTTAYTCCRYWCWYGCT
            *    .*.        *:.  ****. * .*:   . .  .*.: :.*         .

38          CGCAGTCRCATCAYYRCSRACWKYGRTKRYWYSWKCCCWKASRYYMAKCWRMKSAYKCAC
47          TAMCTATRMGCYRYTGYGGRYWKCRRYGACTYGAGYTYWKRSAYYARKYTRMGSWTKYRM
             . . : * .   *   .   **  *     *.     ** * **  *  ** *  *

38          TGYACAATTTKGAGGTCMGGWTKAYYCGRKKTTYRKGARRKSMAYSGTRGAKYGGKTKWK
47          YSYTTWTCACKKMTTGYMRATAGCCCTAGKKWYYRKKMRGKCMWYSAGRAMKYARGYKWK
             .*:  : : *      * . : .   . **  ***  * *.* **. *. **.   ***

38          CTTGYRTYYKYYGYCTCGMTYGGKTRKWMTGYGRYKWKYYWKRRYYYYRSTTCSSAACRY
47          YCYTTGGCTKYCAYAKYSCWYKRGYGGWMYRYRRYKWKCYWKGRTCYYRSYWYSSMGAAY
                     ** .*.. .  *      **  * ***** *** *  ****   ** .. *

38          YMCCRYCCCGWCAKTMGTGWSTKKMWYRGRAKKYCGTRAGACCTTTAKMMGAKMYRYKTC
47          YMYMRCYMTATYTKWCRYRWSKKGMTYGKRMKKYGAAGTATATYYCCKCMRRKMYRTKAY
            **  *    .  :*     **.* * *  * *** .: :.:.    .* *  **** *:

38          CTAYYYYGKGMMKKAGAGRGRRSMYKKKSRYMSMKCTMAKRMRRWKYRRAKYKATYRCTK
47          TYRYYCYSGKMMKKRKGRATGGSATKKKSACCSCKYAMMGRMRRWKCGRRKCKMGYRGGG
               ** *.  ****  .     *  ****   * * :*  ******  * * *  **

38          MRMKKKRKKTCGAYMYKMWYKKRKKRRKCRRYMRMRTKKKYRACKRYWAGMKKTYRKKGW
47          CRMKKKAKKCTAWYMYTMWYKGRKKRRGMAGCAAMRGKKKYRRSKRYWRRAGGKTGGGTT
             ***** **  . ***.**** *****       ** ***** .****     .

38          WCKTYMMRKACGACKKRSWAKGMKYYKRRKMKYAYKCSWYRYGKWTYYKTTYKCCKWRRM
47          AYGYYMMRKGYTTGKKAGACKKCGCYKRRKAKCWTKYGTCAYRKWWCTKYGCKMMKWAAC
                *****.  : ** . .*    ***** *   * .   * **   *   *  **

38          ACTKKRKRGAKACTGYGKRTTKGGAGTGTACGTATAAYYTAKRTTKMGATGTTTRMYTYG
47          TTGTKRKRARKWMKKYTKRAATTTGAGCCGTKWCKCGYYKGKGCCKAACCACYKRCCWYR
            :  .****. *  . * **::.  ..   .   ....**..*   * .. .  .*   *

38          CTAAGYTRCAWAWRATTCTGTGYAATAMCAWWYKKRRKGAKKMRMAMGGTTGAKAATKWK
47          MCGRCTCATWATWGRKWMWKGACGGWWMTCWWYKKAAKKRKKMGMWAAACCTTGGMYKWK
              .        :*  .     . ..  * .*****  *  *** *  ..   : .  ***
```

```
38      YKSWYGAMWKCKKWASYGWMCRYTTGWKYKKAGTATYGRYRTTWRMKGARTACGATTGKR
47      YKSWYATMWKMKKWGGCRTAMRCKYSWKCTGTKWTCYARCGAAWRMKKRGCCTAGGCAKR
        *****.:*** ***..     * . .** . :  : *.*  ::****    . ..  .**

38      MYCRAMWKAYCRKKKAGAMKKYRYYKKGKYGACRKKYYCKKGTYRKGGCRGKMKKMRKRG
47      MYTGRAWGMYYAKKKWSRCKKYAYYKKKTCAGTGKKYCTKKRYCGKARYGKTMGGMRKRK
        **    *  *  *** .  *** **** . ..  ***  **    *.    .*  ****

38      MRKYGGAATYGCRRYRRKWRGCRYKKYKTKAKWRGARRTAYRYRYRYYKTKMMRKTTYRY
47      MRKYRKMMWCRYARYGRKWGAMRYKGTGCKWKWRKRGGKRYRYRYRYCKWKMARKCGCGC
        ****         ** *** . ***    * ***    . ******* * ** **

38      WKRTMTCYAMYWYKRMTRWYMKYAAYRKGWGGYWCTGYMMYYRWCYKGYRGTKTRWMKKT
47      AGAYMCTCGMCWTGGACRACAGYWTYAKTWRACTSWRYMMYYRTMCKKCGKYKWRTMKKW
            *   .* *     *    * :* * * .  .  ******   *     * * ***

38      TCCKWYRRWKSRTACMCRYYRYTKGTCRGYMCGGCRRKCWTCYKARMKRWMRYKKCRMKR
47      GMYKTCAGAKGAKRMMMRYYGCYKKWYARYMYTTARRKAWKMYKCAAKRWCATKKYRMKR
           *     *. .  * ***   *     **   .***.*. **.  ***   ** ****

38      AAAGTCWCTGAKRKAGAYGGWRWKCAGGTRACCWTAYKRMAAGYCYGTGGAMCKKSCCMK
47      MRMRCYWTARWKRKWAWYKKTRTTYRRAWGTYYWYCYKRMMWKCTCAAKTRMMKKSMMMK
              * :  *** . *   * .   .  :  * .****      .:   * ***  **

38      GMYRYYWYKKCATMGGRACGTARYRYKWGYRYYYMYRWRCYRKSRRYWTGKYRYRKKKKK
47      TMYRCCWYKKYWWMKRRGMSWMRYRYKTRCATCTAYRTGACRKGAATAYKKCRYAKGKKK
         ***  ****   *  *. .  *****        **  . **.      * ** * ***

38      KGCGKKKTRTTGRARAKRTCMKRKYTGGGGTKGKMSAKTGGARYSMRYAACASAAATGYR
47      GCARKKKGRWWKRGRRTGCTAGAGCYRTTTCKAKMGWKARCWRYGCGYMCTMCRWRYSYG
          . *** *   *.* .              *.**. *:   **.  * .  .    .*

38      MMTRKRRYGYCRTACMCTCYYRWTAKYYRGARTGRAKTTYRSYAKKKTKGMWGTAAKAYA
47      CMKRKRRYKCYAATSMAWYYYRWGCKYYRSWGYAGTTKWTASCGKKKCKKATRKGGKWYW
         *.*****    ::.*.  **** .****.   . :..   * .*** *    ...* *

38      MCGCMYRKAYTCAATTTGMCYASYYMMGWKCTGTTTTRGRRWCACKGWWAWYKYYKKGTC
47      MYRSMYRKRYYYRRKWCRCYCWSYTMMSWKTWRKYKKRAGATYRYKTAAMWYKCCTGRWS
        *  .**** *    .       ** **.**   . ..*.      *    ***  .   .

38      RKKRRKRKTYGRTCAARGTAGKMATAKKGAAAMKKAARWCYKAGCCRKMKKKKGRKATCC
47      RGKGGKAGKCRRWYRWGKYCKKCGGRKTTTTMCKTRMRTYCKWSYYAKCTKKKARKGCMM
        * *  *  .  *       . * .  *. ::  *.  *   * .   * .***.**.

38      MCKGMAKKRKYWRKRCWGAATWYRKAKYKKKKMSKKCGCWRKCCKATKAWRWRCWYKCSR
47      ASKKMRKKRKCTAKAYWTGGGTYAGGGCTGKTASKKYKYWRKATKGCKRWRWRTACTYGA
         .* * ****   *  * ..  *  .  . *. ***   ***. *. * ****   . .

38      YTGGCYKSYAAAGACKGGGKRKMYRMKRYCYCGCMYYKKAAGRATYWSRMYKGRCTAKRG
47      CYRRYYKSYGGWRMYKRAAKGKCTACKRYSYMRYMTYGTGWKGTYCASRATKKGYYRKRC
             ****..    * ..* *    ***.* *  * * ..   :   **  *     **

38      TTWWGMKKWWSYTAATYTARTMKGARTRTTKGGASTAGCKGGTYRKMMAMCYYAAKAKKM
```

```
47              AYAWRAKKAAGYAGGATCMRCMKRWRKRWKKTAGGAGRMKRKCYRKCAGCYCCRWKCGTC
                :  *  **  .*:..:   * **  *.* .* ...:.  *   ***  .       *. .

38              GRYGAAKGMYRAMRYRCKKYYWKYKRWMWYMKKKKSYARCYYRKWMMGGSTKGCASRACK
47              RGTAWRKRMYRMMRCRMKKTYAGYKATMWTMKKTTGTGATYTAKAMARRCWKRSWCAGMK
                   .  * *** ** * ** *  **  ** ***... .  *  * *   . * . . . *

38              MYGYYRCAMKYAGTAMKCMAMAATRRTCGCTKTWGARKAGATGGYCKATARGYKYATCKW
47              MCKTCGYMAKYRRWRMKTMRMRRWRRGAKAGKKTRWRKWKRYRACTKCGGGRYKYWWMKW
                *        **    ** * *   ** . . *.   **     .  *.  .  ***   **

38              TTTCCCRGGGCTGTCTTGYYCCTWTTKTTKRYYAKKKACCASCCRARGRYCKYYKYMMKK
47              GYYSTAAAARYWKKMYYKCCYYATCWKYYKRYYWGTGTGAGCYMGWGRRTYGCYKTAMKG
                   . . ..    .        :   *  ****  . : ...      *    **  **

38              AYGRSWKKTGKTTCTKYCRYYKCGYYCRRTRGYTTATGAARYCGAGKCKTTKKMYRMRAK
47              MYCGCAKKKRKKCMYKTTGTTKGACTYARYGKCACCKKMRAYYRRRTMKCCKKCTAMRWK
                 *  . **. *.   *     * .    *    : ..    *    . * **    ** *

38              KGTRCGRYWYRKYMATAKWWKGGYTRRRYRAKKAKKRAAKMRRWAYRCGRAYYRKRRKRY
47              GRWRYKGTTTGKYMTYWKAWKAKYYRARYRWKKRTKRGMKCAGTRTGYARTYYRKGRKRY
                   *       ***:  * **. * * *** ** .**. *        .*:**** ****

38              KRRYSYMYYRKMKYWWKKYMRRAAAAGYSACKGMRAAYGAAACRMYWYWAAGGGTTARMR
47              KRGCGCAYYRKMKYWWKKYCGRMGWWSCCRTKRMGRGCKMRWYRMYWYWTGTASYYRRMR
                **  .  ************  * .  . .  * *  .      ******:. ..   ***

38              YYCCRKTAYSCTKSYKKRRGKKMGCTKKRGMKRMRYWGGRYYKYWTTAATYMRMRYKKCA
47              YYTTRKWRYSYWKSCKKAGKKKMRTWGKRRMKRAACTTRGTYKTAGGGRWCCRCRYKKYM
                **  **  **  ** **   ***    ** ***        **    .    * ****

38              TRTYGAGAYAAMTCTCYYKTWWCYYRRCKKMKARKTTAWSRWKGGTGGGTGYRYSTGCTK
47              YGCYRRSMYWRCYYKYTTTWWAYCYRRTKKMKTGGGGMACGWKATYRRRGAYRYSYRSWK
                   *  . *     .   . *   *** ****:      . **.      .****  . *

38              CAAYWCGTSRSYKGTCCTTRTKYKKRRYTGKAGATRAWYRTCWWTRYATWCCGYAKYSWY
47              YRRCAYRYSRSYKKCYYCARCKYKTRRYGAKMRWYAWWYRWYWACGCTAWMYRYRKYCTT
                        *****     :* ***.*** .*      ***  *    ::*   * **.

38              TRWRYGYRSKTRRYRRATTCAGACTCYGTKSRWTGWMGGATRGYGKSRCYRKRCYRRYCK
47              CRWRYRYRSKCRGTGRRKCYTRGYYYYSKKSRTYCTCATWWRRYRKSRYYRGGMCGATYK
                 **** **** *   * .  : .   *..***     .   * * *** **        *

38              ACWKKMMYKSMAWATTCKYGGMRATCCGCAYYYKRWYRRRRWTAKKKGKRRKCTMGAATK
47              RYTKKMMYGCCRWTAAAKYKKMGWYTTATGYYYKRWCAGRRAYWKTKRGRRKYKMRRWYK
                   ***** .  *::::.** *     . .******   **   *.*  *** .*    *

38              YCCAKWSRAKKYKWCTRSCWCRGMWGACYKACCSAGGRGAKYYCWGGYYYWKWRMCTKMK
47              YMARKTSRWKKYKWYKGGYTYRRCWKMMYKWYYSWRRRKRKTYGWRAYYTTGARMYCKMT
                * . * ** ***** . .   *  *   **   *   *  * * * .**    **  **.

38              TTRYYGCKYACRTYKYATWGCWGWAKMWYYYATAKCGYMTRRKGACTGGCRAMCGACCCG
47              YKRYYRAKCTTRACKTWYWAYTTWGKMWYTYTCTKMKCCCRGTRRYWRRYRRMYRMTAAA
```

```
         .*** .* : *: *   *.   *.**** *: :*     * .       * *    ...

38       CGYWWKCAYMKMMCKKGMYAYATTCACYYATYKYARSKYKKRCYMTTAGSSKKWRRSKYY
47       YATWWKMCCMTCMAKKSMCRCRKGMGMCYCCCKTMRSGTKKAMYMGCWRSSKKWRRSKYT
          . *** . *. *.**.*    .  .  *.  *  **  **  **    **********

38       YWRYKRWSGMMCWKRYRMMRKKAKRMRKRGKYRKYMRYCKCYWMKWKCRKGCCAKCTGGC
47       YWGYKATSSACYTKAYRMMRKKCTGARTRKKYRKYMGCMKMCWMKWKYGGRYYTKYCTAT
         ** **  *.    * *******..  *.* ******   *  *****      :*   .

38       TMYWKYRRAYYAYGCMGWRKTTKTKTRRSYTAKKTKKTARGKRWYKRKTRGSRMYYKYYK
47       KAYWKYGGCTCGCASCTTGKYCKKTGRRSCKMKKKKKATAATRWCKRKWGRSRMYYKTCK
         . ****  .  . ..    *  *.. *** . **.**:: ..** ***   ******  *

38       RRKAWGSYRRKRCMKKGACKACCATGTAATYKRAWMGCTRCTRWYARYGYRCKGWRMCRK
47       AGGMTACCRRKRYATKARATTTYCAAKRRWYKGTTCATAAYYGWYRRYTYAMKRTACGRK
              .. ****  .*. ..:  .:..   ** :  . :    ** ** *  *     **

38       KYGRTKTRRRYRAMGYRYCMAAAARTTMWAGRKRKGRRYRYSYYKGAYKATRMYYTKMYK
47       KCRRKKYRRRYRGMRYGTMCMRRCGGACTRTAKRKRAAYAYCTYKSRYGRYRMYCCKMYT
         *  *.* *****.* *       .  :     ***   * *. **. *   ***  ***.

38       GRYCRATCYARYWCKKKAYCCCKGTTTGGCACWYRKCCTCATGSWYRRRGKCYWTAWKYG
47       CGTMACGTYGGCTYGKGRTMTTGTCYCCKMMYWYRKMYYYRYKSTCGGGAKTCTGGWKCK
              .  *.     *                ****       *     .*    .**

38       TMRMTGMTGTSTGTTWCCGTSCCCTCKYRRKRYSKMTYAKTTKCTYAWWSCCCCTTGTGR
47       KCRMYRAARKSWKKWTYYRYSYMYKYKYRRKRYSKMYTWGYYKAGCMWWSMAAACYACRR
         . **   : .*  .      *   . **********      *.   *** ...  .  *

38       MRWKCKMKYGGCTYCKCYATTSATYWTKKWTRKYRKAWATATAAATRTYGCTRTYTMAYG
47       ARWKTKMKYSAYYCTKYYCACSRYYWWKKWWRKYGGRTWGCCCTGCRYYRMAGACCMMCR
          *** ****..    * *.: *  ** *** ***      . .:. * *  : :  *

38       ACMKKKMYWRMTTGAKACKMKCSMYCGYWSYYRRTAKGGYGRYAGYGAGGCYRMAAGGTC
47       MAMGKKCCTAMGKTTKGTKCGAGCYTAYWSYTRRWWGRRTRGYRRCARRKYYRMWRKTGA
          .* **    * . :*. *  .. * .**** **        *   .    ***     .

38       GAMTTGRAKTYRSRCKYAKKYRRRARCKTAAAGAKKAARCSRRKMTM
47       KWAKAKGGKYCGCRYKCTKKCAGRGAMGCGCRSRKKCCGTGGAKCCM
            .:   .*   .* * :**   *.    .. . **..  .  *  *
```

The pairwise alignments of two accessions 38 and 48

```
38              MYWAMKKWMAGRKYWWSMWYMRKWRYRCGYCSWGCRYMYGTYRCWRMYRYRWYRMGMYWR
48              CCATMKKAAMRRKYTTCCTYMAGWRTGYAYTSWCTRYMYTCYRYWRMYRYRTYRMTMYAA
                   :***    ***  .  **  **   .* **  ****  ** ******* *** **

38              YRKCARGYTYTGCGGKRKKWTGAKCYSSYYRTWRCCGYRCRSKCKRRRSMMRRWYRKYTA
48              CAKGRARYACWRTKAKAGKWKKRTYTCCYTRAWRTATCATRCGYGGRRSMMRGWTGKYWG
                  *    *:     .*  **.  .  ..* *:** .    *.    ****** *  ** .

38              YKRGCRRYYRAMTRKYSRKYMCWRTKAKSRRRRYRYSRWKWWRYCYWRGRKGGRAARTAC
48              YKGRYGAYYRRAWRGTCRTTAGARKGRKSRAGGYRCCGTGAAGCYYTRAAKRRRWWGYGM
                **     ***   *  .*.    *.  ***   ** .        * *. *  *    .

38              AYRWRYWKYMTYYRYATWYYWKYMRYKYYYRRMKKMAKCKSYKWRYYMMYWGRKYKRSCA
48              GYRWRYWKYAYCYAYWAACCWTYMRYKYYYRRMGKARTYTGCKWRCYACYTKRKTKGSMM
                .********   * * :   *.*********** *  . .. *** *  *  ** * *

38              TYRYSRYGYRYWWSYKRYYSYGKYRAWRKCCRYMKRRSGAWGTYKRMRAAGMYATMYAYC
48              YYGYGRTRYRYWWSYGGYYSYCTCGCTRGYYGYMKRRSARAAATKGCRGCAACCWCYGTM
                 * *.*  *******  **** .  . *    ******. .:  *  *...  .  *.

38              MRYMYGAKCCGYWRYYCMACYAGGGSWYYTYYKYYRRTYRGYTCKMYRYMRKYYRYTRGW
48              CRCMYTGKYYRYWRYCYMRYTGATTGTCTYTTTYYRRAYARCWSKMYRTCGKYYGYKGKW
                 * ** .*   ****  *    ..  .      .****:*    .****   *** *.  *

38              YMKMWYGMRYGYMRYCGMRCYACRKWMRRRYYYYMRWWAMWGARMCSYWWRRKMRAKYCK
48              YMKCWCAAAYKYCATYRCRTYGAGKAAGAGYYYYMRWWGATRGRMYSYWWRRKARRGTYK
                *** * .  * *      * *.. *     ********.   .** ******* *    *

38              SRRATRCATMYWYKRRMMYMKMRKYMRCMCGYTTCKMYYYMKRRRYRKYTAYSYRYRCCY
48              SRRGCRYGWCTWTGGGCCCCGCRGYMGYMTRCCCTGATTYMKGRRYRKCAGYSCAYRAGY
                ***. * .   *          * **  *          *** ***** :.**  **. *

38              ATAWYWRWSKYWRRRCMKRYCYAYWWYWYGKCTTKGRGTACRGTGAKKTASKTTTTGAGY
48              TCGTCWRWSKYWGAGYMKRTTCWCAWYWCRKTCAKAAAGTTRAYRMKKATCKGCWWKGAT
                : .  *******    ***      ***  *  :*. . : *.   **::.*     ..

38              MRRRKTCGRKAKGGGMGYMGKYATSRKYRYSKGTCAKWSGGCRKRGYCYWKCTAYYRWTY
48              CGRRTCTCAGWGTTTCTCAAGCWKSRKYRYGTRYTMGAGAATATAKCYTWKAWGYYRWYY
                  **.              .   .******..      ...  .     **. .**** *

38              TAYMKGRGAAYTRYRWMKKTSCWGKKKAWCRTCMWARYMCGCKGYRRCYWYYYTCAATTY
48              KMYAKRGRMCYYRCATCKTASYAKKKKGWYAGTATWAYMYRSKACAGYTTCTYYYRRKWT
                . * *    .* *    *.:*   ***.*        **  .*.        *    .

38              TRGRTCAAYKYKKYYWKYYTKGTGYAGCYAKYGRCATTCAGCRTCYYMKAMKRKCKKKGG
48              YRRRKYTCYKCGKYYWKCYCKCWACCAYCGKTTGYWACTCCTGCYCCMKMMKAKMKKKSR
                 * *. :.**  ***** * *  . ..  .*     :  .       ** ** * ***.

38              GCTRCAKAGACKTMRTTGCGMCRTKTKTYAYCCTTRRSTAGRRMYYYYRKAAKRYWMKTT
48              KYWGMGKMAWYGYMRACTARAYRKKKKYCTCYYYWRRSKGSAAAYYYYRKRRGRYTMKGC
```

```
             .* .    **:  .   *.*.*  :     ***...   ******   ** **

38      RCRSCYYYYYYSRRKKYRRWRCYRWKKCCACKRWRMCTTYWKKKAACTYKYKKRKWCGGR
48      AYRCAYYYYYYSRGKKTGGWRMYGTKKTTRYTRTRCTCCYWKKGGGMAYKTKKRKTMACR
          *..******** **   ** *  **    .* *    **** .. :** ****  . *

38      KKKMCKKRGMAGKRKGGGGGKYRMKYYRKGTTTTKWAATYYRMARRKAGGKGCTGGATRK
48      KKKMAGKRRMWRKRGRKAARKYGAGTYRKAGGKCGTGRWCYRCTRRKGRTGAMYRKWKRK
        ****. ** *  **   .. **    ***.  .   .   ** :***.   .     .**

38      AGAYAMYYKKWAWWYKKTKMKGAGKKYRRAYMRRMRGACATGGGTGYRYYYKYKTGCTKC
48      RSMYRCTTTKWRWACGGGKMKKTRKKYRAMYMRRMRTCTRYRRRGAYRYYYKCGYRSWKY
         . *    .** *     *** : ****  ****** .       .******    . *

38      GAATAWCCGTSRGGGSKYKKGCTGYAGATRSWAWARGGWWGTRCYAMKGTYTYSTWRRWY
48      RRRYMAMYRASRSAKSKYKTAYGAYMRWYASWRWCRAKWARAGYCRMKRYYAYCCAAATY
                 :**.. ****..  .*     ** *.*. *  :    **  *:*.     *

38      YYWKTRWYGYKYTRGRRATTCAAWTRTKAAATRRTCGTRRRMRRGYRRYCKYWKKYKGWR
48      YTAKCRWYRCKYCGKRRRKCYTGWARYKCWWGARYMKKRRRMRRACGATYKYTKKYGRAR
        *  * ***  **   ** .  :.*:* *.    *   .******.     ** ***   *

38      MMTCRATCYRYYYYRKGWTYRRRAGSTTWSYCKTKKGWYYTKKYKYRTKTYYCWWSCCAA
48      MMGTGWYTYRYCYYRKAAKYRRRTTSKKTSYMKGKKKWYTKKTTGCAYKACCYWWSMYRR
        **      *** ****. .****: *.. ** * ** ** .*.     *:   ***

38      WGARAMCCKTRMWKACGTGSRWYKTKAGYMAWKGGCYATTAYRSATCTKKWWTRKYRKGA
48      TKRRGATTKCGATGGTCCASRWYKYKRSCARAKRAYYCACMCASRYMWKKWWWRKYGGAR
           *.   *     .   .***** * .    * . *.:    *    **** ***  .

38      WARCARTAMACRYMKTRGTCTCKYMKTTCAWRRCMAYGACKKKMYWRACKACMKRYYKGY
48      TWRYWRYRAGTRYCKKRRKSKMKYMKCCYWWRRMMMCACAGKKCCTAMMKGTCGRYTKKY
          *  *   . ** *.* ... ****    *** *  ... **      *.   ** * *

38      MYGYAGCYGWYRGARKGAAKCRGRYKRTAGYKRCGRRWTAGGCYRAAWARKKGCCYKTGA
48      CYRTRRYYAWTRTGRKSMRTYRRGYTRWRRCGRYRRGTYRRKYYRMRATRKKKMYCKKKR
         *     *.* * .**.  . *  *.*     *  *       **   :***    *.

38      GTATTTTTAYRTYRSRCKKKYRRAAAAATTYKTTTTYYRTSGKSRRMCTYRMKRRKAGAR
48      KYWKKCCWTTAYCGCRYKKKCGRGMRWRACCGCCAYTCAGGSKGGACACTGCGRRKGAMA
           ..   :     .* ***  *.    :     :     ..*.   .     ***..

38      MWCGTKCCYATTGTYKYKYCARSGCTGYKWKGYAGAACGRRSKYYMYRMWWTCTGRWKCT
48      CAMAAKSMCRKKKKYKCKTTMRSATAKTKWKKYWRGRMTRRSKYTMYGCTTCAAAATKMW
           .:*.   .. .** *   **. :  *** *  .   ***** **     .:.  *

38      GAGMMWKRYYMKGCKKKAKRCRKRYYRGCTRKGYCCYYMRYSKWWKKCRTKACMCRTGGC
48      RMSACTKAYCCKRYKKKCTGMRTRYYRRYYRKKYMYCCMGYCGWTTKYGKKTYCYRCTAT
          .   * *  *  ***..  *.****   ** *    * *. * .*  .*:    *  .

38      KWKYRKRGCKYKRYCKTWKRYKYTRMKKWKRWYRKKGASRMYYYYKRRKRKGYWGWYAAK
48      KWKYGKGRYGYTRYYGGWKRYKCKGCKKTKRWCRKKRGSACTYTCKRRGRTKYWRTTCGK
        **** *    *.**   ***** .  ** *** *** .*   *  *** *. **   ..*
```

```
38          RCKYRRATGAGTMYYYAKRYCYYCTGKTGGKYRAGAYRRYMCAKRRWMRKAKKAKAYTAR
48          RTGYGRGCAMRYMYYTMGGCTYYMWKGYKRKYRRKGYRRCCTMKRRWCRKTKKRKRCYRG
            *  * *. .   ***      **       ***  .***    **** **:** *

38          CSMACCYKYRYKYYTTAKYWTKRWRMGGRGMYGATTKTKCCMTMKACGCCGYRYTGCYMT
48          MCMMYYYKYATKYYKKCKYWAKRWRCKRAKCYAWCAKYTTTAKCTCAAYGATRYAATYMW
             .*   ***  ***...***:****      *.  :* .   . ....  . **:. **

38          KRGTAKGCCYTCYMGTTTTRWMGTGSGCRMYMMACYYRATTKWYRAWYTCGWRTGCCYGK
48          GGAYWKRYYTYMCCRYWGYGTCTCAGTTGCYCARTCYGRYYKWYRRACWTTWRCRYSCRG
              .  *                  ..    *     *    ****      **   .

38          TCARRMRGTAKRMRKCAKSAGCWKCCCTRYGYTRYWGAWYRMTGYATKYYTCCCYKWGAY
48          CTGRGCGRWRGAMGGTGGCGRYWKAYTWRTKYWGTTRWWYRMYSYWKGCCGTTYYKWKTT
              .*        *   . ..  **.   *  *      **** .* .       *** :

38          KTAGKGRTCKGGKTTKMMYGCRGRTWKGAGGGTTKGTTKYRAKYTTTTWYRRKWRSRTAC
48          KKRTKTRGYKKRTYYKMMYKYAKGWTKRRRRRKCKRWWKYRGKYCCGKTCAGKARGAKRM
            *.  * *  *  .  ****       *     . *   ***.**   .    * *. .

38          AGCKKKGRGTTAACCCKYRMRYKMCRMKGTYGAAKGGWWKKKWGGYRCCWMKAGCRKYYT
48          WRAKKGKARCCGGYYMKYACATKCYRMGKYCATWKKKTTKTKTRACGYYWCKGTMRKCCC
              .**      ..   **    *  **    .: *    *.*  .    * *.  **

38          KRRRTKKSCMKSGKYYWKYKAMGGRTATATCYRKCTRYYMTRWCYKYRKSRRYKYRYYRY
48          GGGGYKKSYMKSKKCCWKYKWAKRRWMCGCACRKYYAYTAYRTACKTGGSRRYKCRYYRY
                 *** *** *  ****    *   . . **   *   * . *   ***** *****

38          WWKKWWSYGGCRGCTTCGCYYRTATRAWGYSTCTTAYCYRKTGGMMMTYYCKGACGWRYG
48          AWKKAAGYAAMGRMACMSYTCGYMYRMWSTGWYWAGTYYRKKKSMMCKCCMKSRYRWGTR
             ***  .*..    :  .       * *. .   :.  ***. .** .   *.   *

38          AAKCCCRYRARWCAKGTATAARCCTKYWWGRYTRRRKRYMAKKSYTGWCYRRGGSTYGCA
48          WRKYMYGCAMRWSRKRKRYWRAAMCKYAWKRYAAGRKGTMGTTGTCTAMCAGRRCWCASG
              *       **. * .     .  ** * **:  **  *....          .  ...

38          GCYYAMYACYKYYWRCAMGKRCTAGCRCGCTKYTACAAGATGGKWRAGKTTCCCATGWGT
48          RSYTWACGMCKTCTGYMAKGGGCTKTRAKAGKYKRMRWKRYRTKWGGKKYYSTAGAAWKK
             .*    .  *            :  *. . **.         ** . *  . ..:.* .

38          YTYCKRYAYAKKKGTGCTCKRWCKYYYKYMMYGKGRSWKKWGSYRKYYCGYYCRTKRGYK
48          CYCYKRYCYWGTGKCAYGSGRAYGYCCKTAMYRKCGCAKKTKCTRKTTSACTYRYKRKCG
                ***.*  .   .  . *   *  *  ** *  . **  . **  ..   * **

38          GKTGYYKGGKTTAAYKMYRCRAKKGTYKRYKMKKWWGTRCGCKKRRYRTKAKRATMRCYG
48          KKARYTKATKCCMWYKCTAMRWKGRWYKGTKMGKAWKYRYSYKKARYRAKRTRGKCAGYK
             *: * *. *    **    * *   **  ** * *  * . ** ***:* .*..   *

38          GARYYYRYACRRKRYKRYYRKKWYWWKKMRTAAAGYSKCMGCAAKTAGGRCAYATAACKM
48          RTRYYYRYGYGRKRYKRYYRKKWTWWKKCGYMGWSCCKTAAMMMKKWRAGMGCMCRWYKM
             :******.  ************ ****    . . .*  .   *.  .  .      **
```

```
38              YAAGGKTMKRMRYYKKYCGARTACTGKSYYTRRAWKGRAKRCMGKACCATGTKKYSRTWW
48              YTGTAKWMKRMRYYKKTMRRRARYWAGSCYYAGMTKARMKRYARKTTYCAAKKKYSRWWW
                *:. .* *********    *:   . * *     *.* **   *:  .:..***** **

38              ATAKYWGCTRWWYRKRYARGWRKARCRRYAKMATRRCRYAGGCAATAAKARRYKCATWMT
48              RWTKCWATAAWTTRKRYRARTAKGRGGRYTKAWYRRYRYMKAYRGCGGKGGGTGMMAACC
                  :* *. : *  ****     *.*  **:*    ** **   .  . ..*.      :

38              RYYAGCCGMYYKRGSCGRAYARTTMGSGGKRKMRSKWSCCTTCMKKYYGCCRRCRTTKKM
48              GTCTAYMKCYYKGRGTAGGYCGGACRSRRKRKMRGKWSYTGCYAKGYYAYARRARWWKKM
                   :.     ***  . . .*.  :  *  *****.***       * **. .**.*  ***

38              GWTYRKMRYKKYARYWWTATKKCCSCAWCCKRSKKKTTGCYTTTARRMCATTCTWTGWYR
48              ATKYRKMGTKKYGGCTWGGKKGYTSATWTTKRCGGTGCAYCYYWMRRMMMCGTGWAAAYR
                . .****  ***.    * ..*   *.:*  **.  .  .      ***      *:. **

38              CTAWKCGCCTCATGSWYKYWWKYYGTMWWYRGKMRRMGMTGRWKCACMGYTYCWKRACTT
48              TWTTKTAMYYYRYKSTYKYWWKCCKKATWYRKKCRRMRMWRATKYRYMKYYCYTGRGACY
                  : * .       * ******    .  *** * *** *     *    * *      *..

38              TWYKARKGYCGWTCYKKMRRRKKYARTCAARAGKTRTYKMKGRAAKAYRRAAARTGKRYK
48              YWCKWGKRYSRTWSYGKCGGAGTCGRWYRWGWAKCGWCKCKTRTTTMYRRGGGRAKGACK
                 * *  * *.   .* *     . .*       .*    * * *::. ***...*:    *

38              KYYCCCRATMKKTTGRGRRAGGTTCCMCAYMRMAKARKYCCCWWRWKRYMAKTATRAAGR
48              KCYYYSAWYCTKWKKRARRTRRYWMMAYGCMRMGRRKCYMMWTAWKAYMGKKGGRRWTR
                * *   .    .* . *.**:         . ***   **     *  ** **.*.. *   *

38              YKKWKRRRGCACTKMWKCGYKAYCTAWRYGRCKSRYTGGCYKKWWMACAGKTTTTGGKRK
48              CKTAKRGGKYTYKTAWKYKYKGTSYRWRYAGYKGACYRRYYKGAACRYWRKCYCCRRKGK
                 *. **    : .. **  **. .  ***.  *.       **         *       * *

38              MMKYTGGCKAGGKAAGAGCTMYYYYYRCYGTTATAWRACYKKGACTMYKAGCGRGKGAGC
48              MMKYWRRSKRRRGWRRWRYYMYTYYCATTTWWRWWTGTYCKKRCYCATKRRYRGSKAGAT
                ****   .*              ** **             :  ** .     *      .*...

38              AARATWWTYCGTTTRGTAGKAKYYAAACGYAWYMRCYWYTGSTYTKWYGARAYRAYRYKA
48              RGGWKWWKYTTGCKRCCGAGRGYYRCCYRTRWYAGYYTYWKCYYGGTYATGGYRRCGCKT
                 .  .**.*     .*  ..    ** ..     **    * *   . *    *.: .**     *:

38              KRCYMMGCRRCCYKWKTTYCRRMRRRAKAACAAYMWYACWRKKCGCYYWGRYKATATCCA
48              KRYTCCATRRTTTKTKWYTTRRCRGGMKRGMMTYMWYWMTRKKYKSCTWTRYKGYWWMMW
                **     . **   * *     ** *    *  .  :****    ***    .   * ***.

38              GSYCGGTACARKMYCCTCKWGCKYMRTGGCMSYWARRRGRYKKAGTKTKRGYSRKRGMRW
48              KCCMSSKRYRAKMYATGTKWKAKTAGKRRYMSYWRRRAKGYKKWSYKGKGKYSRGAKMRW
                 .  ...     ***.    ** .*     .    **** **    *** . * *  ***    ***

38              GTWKGAGCYKRRRMYWRYACRMCCKGRKAWSKGAGKTGTRCCKWTAKRMRYMYTTKTKCY
48              RYWKARCYCKAAGCTAGCGTACATKKRKTTCGAGAKGACAYYKWATKGAGCCYYYKYKST
                  **.     *          .    . * **: . ...* .     **::*      *   * *.

38              ARTTGGMACKKYYYKYGYGAMYYKAYSRYCGTRKKGKCGWYGWTCGTTRKYCCTGTTYWY
```

```
48    GRWWRKCRSKGYYYKYKYACAYCKGTCRYYAKRKKKKTTACKTCTTCCGKTMMCAYKYWY
      .*      .* ***** *.. * *. .** ..*** *            *    . .***

38    RKKRTTRRKYRRGAKCRAATGCKKRAYGGTTCWGKYSGCTWYWACKRRAYRMKWRYYKYC
48    RKKAYYRRKCRRRRTYAMCARMKGAGYRKYKMTKKCCRTKWYWRYKGRWTAMTTRCYKTM
      ***   *** **  .   .:  *  .*   .   * .  .***  * *   *. * **

38    GRRWMMKKAKYCARTAGYASAYRRWKCTCGTCCRAGYRYCMRWCMRMKCACYWMMKYGKM
48    KAGTCMKKRKCACRCCACMGRYRRWKTYYRYYYGGRYRYGMRWSAACTMWYYWCCGCKTC
           *** * ..* ..  . *****        . *** ***.   .   **     .

38    TRWYRRYARRCYWGKKMAYCCTKTTATCCRKSYKTWGAATYTAWYRYKKTYGRTTRRYKA
48    WATTGACTRRMYWRTGMWCMTAGGAGCATRKGYKYATGCCYCGTTRYKKWCRRKARRCKR
             :** ** . *    :  :. . **.**   .. * .  ****   *.:** *

38    RKYYTRMTATRGYGGYYAKRYRMYSKCRCKRKYMWKYTSKYCRKWYRYYKYRATMKKGKY
48    AGTCARAKRKGTCKRTCGKGYRMYCKTGYKRKYMWGCCCKYYAKTCRCYKYAGYAGKAKC
          :* . .       .* ****.*   ******   .**  *  * *** .   *.*

38    RKAACTGAYYSCKGYRKRKRKRKKTAKRKRGMRYWGRCYGGRMRRMYRRWARCRYKSYKK
48    GGWGTCTWCCSTKKCGGRGAKAGGYGKRKRKMRCWRRYCARAMARMYGRWRRMRYKGTGK
         .      * *    *  *    .**** ** * *  .  * *** ** * ***.  *

38    KKCTARSWTMYRYRCYGWAYTATTAKRKKYWRCCTCKTMTYRYWKRTMKTCAAAMKKCRK
48    KKAWMGGWKMYRYGSYKWGYCGCWWKRKKYTGTTGTKCMGCGCAGACMKCTGRGMKKARK
      **.   .*.**** .* *.* .   *****      * *        **  . .***.**

38    WYCKTCAYRTGWYCRGAGAKGYGTRWWYKCAMKYKWGAYKMRKKMAYYYMTCMKKRYRWR
48    ACTGKYWYGCTWYTAATTTGRCRKAWWCKYRMKYKWAGTKARKGCMTCCCGMMKTGTGAA
          .  *   **  .: :    . ** *  *****.. * **         **.

38    KMRWTMRRYKYKCKRKYATCCRTTTTCYYGKKKKSKACRKRWMRKKMRKAWKSAYCCTGT
48    GCATYCGACKCKAGRKYTAAMGAAAAMYYKKKKKSKWMGGRWMGKKMGKRWKSWCATCAC
               * *. ***::.  :::: ** ******    *** *** * ***  .  .

38    KAGGTYRAAKMKRCTGRTYKAKGTKTYYKKTCYKCWYACCTASKYCTKRCWYYRRKGGYT
48    KGRTYTRCGKMGGTCAGYCKMKAYGCTYKKAGYKYACGAYYWSKYYKKGATTYRRKRKCY
      *.    *..**    .   * *.    ***: **   ..   *** .* .  ****

38    YRWMTKYKGCKGGYCGCACRACKKAAAYKAARKTYTCGYWYYYGAGKSAKYAAAGAMGGR
48    CRACKKYGAAGAACTTAGYRTTTKWRWYGRCRTYYWYRCWYCYKMRKCTTTWRMAGMARR
       *  .** .. ..   .. *: .*   *  .*. *    ** *   *.:.    ..*. *

38    RRWKYYTSYYYMWCATMTCSWMRGRRGGCCKGWMAYYWGRWYGTTTRRYYYYAYRTTCSS
48    RRWKCCYCYYYMWARCCKYSWAGKRRRRMMKTWMGYYWKRWCKAWKGRTCYCMYRYAYSS
      ****   .*****.   . **   **    * **.*** **  : . *  *  ** : **

38    CCCCRYCYTCCKGRGWAMRSYWWSCYWRYKRYRAKCKRTAYKMMYGGTKGYMAAATAAYM
48    YAYMRCTTKYAGRGATTMRSYWWSYCTRYKGYRMKMKRCCYKCCYAKGKKCARWTATTCC
       .  *   . .   . :*******   *** ** * ** .**  *.  *     ::::

38    AMMYKYRKCGCAYKTRCAKTGWKKSRATCCGCTMRKYCTARSWRTRGMKGAAGCCAATKC
48    GCCTKYAKAAYGTGYGYRKGRTKKSRWWYTASCCAKTTCGGGTRGRRCGKGGATGGGAKY
```

```
                    .   ** *.. .      *   ****    ..   *   . . * *    ...  ..:*

38                  RWKGSAKCYGACCKYGKRWAKKGGYGYCCMCMMKRTTRYYGRKAGMCRAAKYCRGGKYCC
48                  RWKASWKTYKRYSKTAGGWGTKRRYRCTGMYAMTRYKRYYRRKMKCYGTGTTYGRAKTGT
                    ***.* * *   .* .  *..*  *    *  *.* .*** **     :..    .*

38                  AMWRYYKKAMWYYGRKRRCYKMGTRKGYWTGKAWTTGKMGCTAMTMCGGTGCCMMGCYYK
48                  CMTGCYKKGMWTYRGGARMCKCSCGTRYTAAKGTGYTKMCAACCGMTRAWRMMCMATYTK
                    .*   ***.** *    *  *  .  . * :.*.    ** .:.  *  .     *. * *

38                  KTTTYKKWGWKGKRKMMKKACCTGKRTTKMRCTCSTTTMCRKKCGRWRTARYYYKYRGTT
48                  KCYGCGKWRWKRKRKAAKKWMYKKKRAATCGTGTCYKWMYAKGAARTRKGGCCCKYRAYK
                    *      ** ** ***  **    . **::.      . . *   *  ..* *..     ***. .

38                  TYKYRRARCTMACGGTYGYARYRRKGTAYGTTKWKKYKSWMRKTCGYYTYAYWMRWATAC
48                  WCKCAGRATACGARRWCRYRGTAAKTYRTACCTWKTYKSWAAKCTAYCCYWYTARWMCGT
                      *      : ..      *      *     .  .**.****   *   .*   * *   **   .

38                  GATYKKTACCTYKYRTMMMWKAAWRKTTCGTKKGTMYRKAAYKCGCAGGAARTRGRCYST
48                  TGCCTKYRATCYKCRAAACTKMRWRGYYYRYTKKAAYRKCCYKSRYRRRWTGCAKGACGY
                     .  .*  .  ** *:     *   **       .* : ***..**.        :       . .

38                  KGAAAYKYARKGYRRWRKRYRRMRYATWKSYAATCKTYYCKTCTAMTTGTSKMMRKRCRK
48                  GTTGTYKYRRTRYRAWGKGTRGMRCRYWKCYRWWYKYTCYGYYCGMGCRGGKAARKRYGK
                      :.:*** *. ** * *   * **    **.*     *         .*    .*  ***  *

38                  MYRRCSGAYCCACATTYTGCTYCRRMTARCYWRYYGYYSMCRCRKKYKKKCKRGCATCKK
48                  MTARYCAMCATMAMYWYYKAWYYRRAWRGMCWRCYKCCGCYAAAKKYGKTAKRRTTGTKK
                    *  * ..  .  .    *  . * **       ** *    .   . *** *..**  :  **

38                  RTTAGGAYYYKKKMMWMMYTATGCGTCTMGYMTGTMTTACCTYKKCMKKKRACTGYYMTR
48                  AGARASRTYYKKKAMTMMYKGGKYCKYYMRYAYRYCKKMYYYTTGMCKKKARMYRYYAKG
                      : ..  ***** * ***..     .  * *     ..      .   ***     ** .

38                  KRGMYYKTTYRGAGTKASKRYKTKKCGAMMGGGRRCAAATWGYGRKTCRSTTTYKKYCTY
48                  KRAATYKYYTRKRSKKRGKGTTYKKYKRCCAAAARYRRRYWKTKGKAYRGYYYCKKCYYY
                    **.  **    *   ..* .*  . **      ... *     *     *: *.     **   *

38                  RYKYMCWWTKCAKYRKGYAGCAKACKTTARYTTTTRMYKKTACCSGCRRYKTAKWYKRYK
48                  GTKTCYWWKKTTKYAGRYTATCKTYGWGRRTCCACRMTTKYGTMSSYGAYKAGKWCKRCK
                      *    **.* :**    *:. .*:      *    : ** .* .  *.    **:.** ** *

38                  YMRMAKACKCGGKKYRKWTCRYRWRTYKKKYRCYTCGYCYKSRRKMWKRKAWKAGYMKKG
48                  CMRARKWYKTAAKKYAKWYYRYRAGWYKKKTATYYYSYTCGSRRKMWKRKGTKRRCAKGK
                     **  *  *  ..*** **  ***    ****    *  .*    *********. *    *

38                  AKRCWAKCACRKAGYTRAAKYKYRYKKTCCTATAGYYKYYTYMMCKKKWYAGCCRGGGKG
48                  RKGMWGKTCTGKTTTGGGGKYKTATKKAYTYRGMCYYGCYAYCCYKKKWYRTYYRRCRGK
                     *  *.* .  *:     ..***    **:       **  *:*    *****     *

38                  MRYAGWWWSMYKKSRKYMSMKCMTGCKARMRYKYRKAKKATAGCTMMWGRMMYKKKKKTC
48                  MAYGTWWWSATKKSAKCCSCKYMCRTGTRARYKYGKRKKMGMRGGCMWRRMMYKKKKKCT
                    * *. ****  *** *  * * *     :* **** * **       ** *********
```

```
38                  GMYKMWKRKTKRKCRRYSRTKKCAGAMKYRRYWAKKCYRGCYCMMGKCCTAGAKRRYKKC
48                  AMYTMWKRTCKRGMAGCSGGKKYRRCAKYRRYWRKKYTGKYYYMMRGATAGTTKAATKKT
                    .**.****. **     *  **    .****** **     * **  . :. :*   **

38                  GKCMKYYKRRKGTYYKGKCKSWKWT
48                  CKMCGYYKRRKRKCYTRKMKGAKWY
                     *   ****** . *. * *. **
```

The pairwise alignments of two accessions 38 and 49

```
38              MYMTKGWKCYGRWKRTGCRKSWCRKYWYKRRKWMTGYAKYYCAMWKYRWSMGYWMYYSYR
49              MYMWTRAKGYRRWKAWSMAKGTYATTATGGAGWMAATRGYYGGAWGYRTCCRYAAYTSYR
                *** .  * * ***  .  *.   .       **:.   ** . * ** .  *  * ***

38              KGYYSYYMMYAKRYRTRMYMKRRSGGCRYGCRMWRKYMRYGRYAYYRYYKKYGRRMMKTT
49              GRTCCYYMMYTTRYRWGCCCTRRSSAMRYRYRMWRKYMRYRRYRYCRTYKKYRAACMTWY
                    .*****:.***      .***.. **  ********* ** * * ****    *.

38              YYSRRWWRWRMYMRMRYKSSRSTYCYGYYRWWTTRSYRWWMRKMYRRYWWMCCCKMCTMR
49              TCCGGTTRTRACAAAAYKSGRSYTYYKYTGWAYKGSYRWWMRKMYRRYWWMYYTKAMKMR
                  .    * *      ***.**   * *  *  . ***************   *  .**

38              RRRGRMMRYSMRRCAATMRKMYYRYRRYKWMARYKRRRYMMRMMRMKKYGRRRRYRWWRK
49              GARARMMRYSAGGMCCCMAKMYTRCARYKWMTRYKRAGCCMGMCRAGKYKRRRACGATGK
                  *.******    .. * *** *  *****:****    * * *  ** ***      *

38              YRKYYSRRMYKYYWYGCYAYSKYTWRWRRYMGKRRSGRSYYMCSWYTMKTKYTRCCCGRY
49              TRKYYSRRCTKYYWYRYYMYCKYWWRAGACARGGRSRGSYYMYSWYCMKYKYWRMMYRRY
                 *******  *****  * *.** **        **  **** *** ** ** *    **

38              GAYYRRYRYYWTWYCKMCGGCGGRWRYWYGYWRKYTRMKYRRKYMWYMRYAWRGTCCCYG
49              ACYYAATATCTYWTYKMYRRYKSRWRYTCRYWGKCARCGYRRKCMTYMATRWRRKYTTYC
                ..**         *  **     .****   ** * :*  **** * **   ** .   *

38              TYKMYYRRYCMRYMRMRCYAMWRWKRYYYKYYRRCKMKYMYKAAMTKSRYWWTMYAMRYG
49              AYKAYYRRYTCATCRCRSYGMWRWKGCTTTYYRRSKMKYMYGTRAWKSRCWACACRCRCK
                :** *****     * *.*.*****    .****.****** :    *** *      *

38              MYYCCMYARRCCKWRYRYGGYGRYWKTTYYYTGCGCARRTYYCTYSAYYYTATCTCTKAY
49              MYYTTMTWRRTMKAACATSRCRGYWKAYCCCAAATYGRRGYCYYTCWYYYGTYTGGAKWY
                ***  *  **  *      .    ***:    :..  .** *    . *** :    :* *

38              KMKKRAGYGWRCTKKCTWKWWWRCRRSKGCMCAWGAGACGACAARAAGYGCCSATYGAKM
49              KMKGGRATTTGTAKKYCAGTATRYRRSKATCTRWKGKRMRGTCTACGAYRAYSWWYKRKC
                ***    .     :**        * ****.     * .     . .: ...* . *  *  *

38              CKGGYYCCCCRMCKKRTTYTGCYYCGGTYKYRYRKYCAGCYSRKMCGTWAGCKYWRYKAM
49              TKRRYCYSYMRMYKTRYKYKKMCCACTACTTRYRKTGGRTCGGGCMCYAMRYKTTGYTGM
                 *  *  .  ** *.* .*.      .  : . ****   .   .          *   *..*

38              WYYKRKCTACYYTCRTGCRWKTMATCCYYAYCMMCGMCKWGYYCATMMYCKGRTTKRWYK
49              WYTKRKMYRMCCYMRYRMRTGYMCGTMYTCCMMCTAMAKWRCYSWYMMYMKKGWYKRTCK
                ** ***        *    *    *.    *  .  *  .*.**  *.  *** *    **  *

38              KWWGTTTYCGWWYRRKKGSRTACYWAAARYYKKYTAGKKGTCYRGGCCCRKGRGGRYMAA
49              KWWRWWKCYRWTCAGKKKGAKRMYWTTCRYYKKYYRRKKKWYYARRTTARKRATTGCCRT
                ***    .    *     **  . .  **::.******    **    *      .**         :

38              RRTCTTAAWKTRMAGAGGWWKYCAGYGTRGACWMKAGARGATTMCTGGTYYGTRAYKCAR
49              RRCTYKRMWKAAAWKTKKTTYYRRTAWGKTYWMGRATAATCCMGCKRWCCAARWTGMGG
```

```
                         **     .   **:      :      .*      .    :  **   .:  .:   *          .:*      .

38                       TTTTGGRRKKSMTGAKYYWYTKTTTGGTAMKKTATCKRWRKRCGTAATAAKYGRKKRGKA
49                       CCCCAAGGKKSMCKWTCCWYWKKWCKRWMMKTCGCAGRTGGRMCCGGCGGKCARKGRRTM
                              ..   ****    .   ** *.       **. . . *    *    .. ..* .** * .

38                       ATGGYWYKTCCKKRRCTAAAYYKAGTAKKRYCYYRKYYYKGKMAGWMRMWSWWAKAYYRT
49                       GATKYTTGYYTKKGRYCTTTYYKRACMKGGCSYYRKYCYKKKAMAWCACAGAATTRCTRY
                         .:   *       ** *  :::*** .   *     .***** ** *   .*      .   :.    *

38                       ARCSRMMAKRKRRKTTAYMKTGKKRRYKGKTRYKTGCCMTRTGCGRYTCTACCATATACW
49                       RGMCRMMTKGKGRKYYWYMKCKKKARYKAGYGCTASTTAGRAATTGYCTWWMYRAGAWYA
                             .***:* * **    ***   ** ***.      .:.      *:.    *         :.:

38                       MACATTYTCAYMSRMCRRYMCYRCTATTAGARMYRMRRWMKYYWRCAARKSCTTATTTWW
49                       MWYTYYYYATCMGRMMRRYCTYGYWRYYRTWRACGCGAAMGCCTGTGCAGCTWCGAAAAW
                         *   :   *  .:  *.** ***   *           *         *        ..   .    .::: *

38                       KKGACRMGAKATAKMYYAKMKTGCMCSYKWAYGACCRRKMRTYCRGCARTGGCTCGAYKA
49                       KKRGTRMKRKRYGKACYWKMKATTCTCYKWRYKRGTGAGARGCMGRYGRYACTCTRTYKG
                         ** . **   *   .*  * ***:      .*** *          *        .* .      :**.

38                       RACCACGYWKAGATRWWTYKYKKKMRRKYGAARRKTCRGKMTAAKAAAKWYCAKRKKCCY
49                       GGAYRYTCTKGRRGGAAYCKCTGKCGGTCTRWGAKCTRKKCATTTGRMKTYYGGAKGYYY
                          ..        *.          * . *    .         *   * * :::..  * * .  *   *

38                       MKRTMYKKKAGATTRRTAGGKCTYMMRMKKARSKCCCWRWARWYCTAKGAATRTRYYATA
49                       MKAYCTTKKRACCARRATAAKMGYAAGMKKRRCKYMMTAWMRWYYCCGTGGGRCAYYGCR
                         **      .** .. :**::..*  *    *** *.*      * ***    .   .. *   **.

38                       TATASAAMKRGGRYAAKCGKYYKTAWRWTAWCAGRKSRYTGGCKSYKMATAGACKGKRKM
49                       CRCWGRWMTGRKGYRRTYKKYYKYRWRWWMWTGRGKGACYRRYKGTGCRKWRMYKRKGKC
                              .  *.     *  .  ****  ***  * .  *.       *.     .     * * *

38                       YRMKKYYYTAAGCMKRYYYKYYRAWACYKASYYYYKGRGRKYCRTYYTCTTKMATWTYWT
49                       TGAKKYTYYRMRYMGRCTYGYCATWTYCKCSYYYYKKGSRKYTRCYTCTYGKCGYAYTWY
                            *** *      * *  * *  :*:  *.******  .*** * *      * .     *

38                       RWKAGWGGTATGARRKKACCTGYKRKYMRTCYAYTYMTARRGWTGARWAYRYYYRTCTMG
49                       AWKRRWTRGGYCMRRKKWMTKKYKRTYAGWTYGTWCCWWGARAAATRTGCGCCCGYTCMK
                          **  *    .    ****    . ***.*     *.           :.:* .          *

38                       CMGATGTTATTTYTYTCKTTRRAKRCRARAGCWRTCTGTMRWWCCTWMTTSTRRMMATMT
49                       YAAMCACGGGYKYWYYMGWWRRRGATRGGTAAWGWTCAACRWWSYYAAYCSAAACAWYAY
                           .  .  .  .* *      **     *. :..*     .: ***.       *:

38                       ATCGCGTTARGATWKKKCYKSYKRTATRAKYTCGWYRYGAAWMCAGRSYGKRYGTTKYKA
49                       RCTAYACCTRKMKWTKKAYKSYTACGCRGKYCTAWYRCTMWTAMCRASYKTGYAYAKYKC
                            . .  :*   .*.**.****.  . *.**  .***         .  ** . *. :***.

38                       MYYKAGCRYRYCTRKYRKWKWASWMKWYGCGWTAYRKRCYWRKYKWRCRKYKCRAGCYKR
49                       AYCKGTMRYRYYKRKYRKTTWTSWCKATAMTAWMYGKRYYWRKYKWAMATCTYAMRMYGA
                          * *.  **** .***** .*:** *   .      * ** *******    . .      *
```

```
38    YTTYTTCWGTWYWACCRRAYRMWKCRKKYYRWMMYCCRKAYMKTYCAYYKAMKGKKCRKR
49    YCYTYKMTKKWYWRYYGRWTAMTKYRKKYYGTCMYYYAKRTAGKCACYCKCMKSKKTGKA
      *    .   .***    *   * * *****   **   *    . ..* *.**.**  *

38    RSTKTTTTGTKYCCCACYWYYAMMMGKYGTTTTGRTARAKTTTRWRRATTCRTKGGCKMA
49    GGGGYCCGRWKTGSMWTCAYCRMCCKKCKWWWGRRWGRTTAAYRTGGTCYMRWTSKSGMW
       .        *  .     *  *   *       * .*:.:: *   :   * .. . *

38    YWRKSKCCWCTWGAATKTWTRRTTRKARKYYTKKKMKRGMTKRMGRYGTAAGYYRKRMMT
49    CWRKGKTATTCATGCCKWTARRWCATRAGTCKKKKMGRRMYKRMRRYSATTRYCGKRCAY
       ***.* .     .. *  :**   .     .**** * * *** **.::: *  **

38    TYKTYYYTTTTAKATTTKACCGAYYMTCTCYYKWGGWYKKAAGTKKTRRAAWYKTTGTGK
49    ATGAYTYWYKYMGGKKCGGTTSMYYCYYKYTTTTRKWCGKGTRGKKCGAMWWCGCGTGAK
      :  :* *  .   ...  .  . **   .   .   *  *.:  **     *      .*

38    KKACRGAAYCWCGAGCKYKKRYTGKCAGAGWASWAWMCCGAKWYYWRWTACMRRYGYKYM
49    KKGYRRRRCSAYRTKSKYKTRYGAKYMRWKATSWRWMMYRRTTTYARWAGTMGRYRYKYA
      **. *    .   : .***.** .*      :** **    .  * **:. * ** ***

38    RYRRRYKYWYRRKKTYMTSRMSCSGSAYGYRRYCACKKYGYCSMGTTWKKYMKMMRATCC
49    RTGGRYKYWYRRKKYYCWSRMSMSRSMYACGATYRYKKYRYYCCRCGWKKYMKMMGWYTY
      *   ********** *  **** * * *.       *** * .    ********

38    CGGTAGGMYYYYRRRKKYRKTAYMMKWYRRYARSAKKSWKYYMCMTTKRMKYGSRYYWYR
49    ATTCTCACYYYYRRRKKYRKYRCMMKWYAAYTGSMKKSTGYCCYCCKKRMTYRGGCTWYG
      .   : . ************   *****  *: * ***  *     .***.* .   **

38    AKATYARCKCTMKKRYKKTCGAAAAYMYKMWYRKCRTMRATAATRTKACAKYRWGKKTYR
49    RKMGTWAAKGGCKKRCKKCTAGTGWCMYTMWCRGMAKAAGCGGYGKKRYGKYRWRKGGTG
       *     .*   *** **  ..:.  **.** *   .  . ..  .*  .**** *

38    KKWWYCMMYRRYKRYKYKCMYKRRKTYYYKCAKKSAYRMYSKWRWYKKWRKAATCGTTGM
49    GGTAYYMMYAATKGCKYKMCYKRRKKYCYKTGKKGGCACTCKWRTCKTWRGWMYYRYGTC
          * ***   *  ***  *****.* ** .**..    .***  *.**

38    KTYTKKYKGAGTGGRKACSMTYKYMRKGYRATATGKRYKYKMRYGWGGTAGCKKGTYWTM
49    TKCKKKYKRRKWRKGKCGGMYYKYMRTAYRGYTAAKGYKTKMRYRWAACGATKKACTTGC
      .. .****       *. .* *****..**. ::.* ** **** *.. .. **.

38    CATCTCAKYTCACCAGSKMMKRCGRCKMKYRSWCCATTYCRRYTCRYWCTGGYYYYSGYT
49    TMYYAAGKCCTGATTRGKAAKRYKGSKMKTACAATMYWYYRRYGTGCWSWTRCYCCGRYA
          :..*   .. : .*  **   .***  . .    * ***    *.    *  . *:

38    MCKKAYKKATAATTCTTKKKTTYKGGKYACTKMYGTCTACKYYMTTARCCTYKCWYMTYA
49    MYKKRYGKTCGTGKTCAKKKGAYKRKKYMTCKAYCKYYRYGYYAKKMGYYYTTYATAWCW
      * ** * *: .: .  :*** :**  **   * * .     ** ..      .

38    RMCACKKTGMRYCCTWRGTTGGAYMRRKAKTCRKYCRRYYWSMMYTTGGGMGKYAYSRTT
49    GCTCTKGYKMRYMMGWRRWWKAGTCGRKRKCTAGYTGRYYWSMMYYATTTARGCTYSRWG
         . *   ***   **    ..   ** *    *  ******** :       :***
```

```
38    AGRYYRTRKAGCTSKRKRTAWMMKRRKRYCYKCTCCCTCTTTAACKGAKRGRAYARWWMW
49    WRRYYRYGKTTTWCKRKRYRTMMKRATAYYCKAWYYMKMACKCGAKCTKGRGWCRATTCT
        ****  *:   .****   **** . *  *.    . : ....* :*

38    KTKYRRKKWTGWCTYYMWACMTCMYRYCRCRGTGAARYTCAATTWRWGACYGTGTCTATA
49    TGTYRRKKWCAWYCYYATMYMWSMYRYMGYGRWSGTGYYYRRKWTGWRMMTKCACTKTCG
      . .****** .*  **    * .****      ..: *    .   *      .  .:  .

38    CGTTTRAAAYGKRMGRCMGCTKARATYSWWRKYYAAMGCTGAACTRKCRYGAYRCGWRRR
49    TRYKKRRRMCAKRCKAYARYKKRRRWYCTTRKYCWRCATAKGTYYGKARYKCYAMRTARR
         ..*    .**       .* *  *.  ***    . : .:   *.** .*     **

38    MCAAACRKRTRTKMYRMYTCGKMYKMRYYAGCCKWARWAMKCRKKTARAAKRCWCRKGRS
49    CGTMRYRKRKRAKATRAYYYRKCCGAGTCTATMGARRWMCTTGKGYRGCRKRTTMAKRRG
        :   ***.*:*  * *   *       :.     **  .  *    . **    * *.

38    KGYSKYGCCKYRAAGKYTKMGKRMKYYATKYCTGTRAKYGARYWWKGGCCCTCCCGAGWC
49    GKYCKTRMMKYRRMRGYKGMKTGAKYCWCTCAYSYAMKYKGGCTWKAAYMYKTYTRWKAT
        *.*    ***    *. * .  **   . . .   ** .   **..   .

38    CMKTGTTGKYYCARCGTTGWWYRKCCTCGSWGKWYWWKGTTYGTKMRWYRRMATYCGWYM
49    TCGCRYCCKYYMMGMKAAAATYRKMYYYKSTRKAYWWGAAACKKGCRATRRMGGCTRTCM
              ***     ::.  ***     *  * *** .::  .  *  ***.      *

38    TWKRTKYYKKKKKKRRRGYCKYYKKYMGGKMGRSWKTGYRYYCGYYCKRTKRYKKWYTTG
49    KAKRYKTYKGTGKGGGRCTYGYCKKTASRGMKGCAKKRTGTTGACTYKRYKRCGKWYYKK
      . ** * ** . *   *    * **  .  *  . *.      .   ** **  *** .

38    AARCGAGYMMKKMYRMCKKYGRYKKKKCWGGRYKRKAKAKRAAYKGTGRAYYRKRKRRRY
49    MRRYRRRYMMKKCTAMMKKYKGTKGKKTWATRYTAKWKRTRGMYGAKARWYYRKRKRRGC
        *    *****   *  ***   * ** *. **. * * .*. * ...* ********

38    SYMYYRKGRKWWKKAAAGCYSACTTRCAYAAACRKAAGGKRKTRTATTAGTYRMRYYKCC
49    GCAYYRKRGKWWKKMGRATCCRTCKGMGCMRWYRGTGTAKRGCRWWYYGCCYRMRYYKAM
      .  ****  ***** . .  .    . .     * :. .**  *    .  *******.

38    GARKKTASCTKTKSTTWSGCAYTTTKKWCYRYSMSCTYYCKAWWSCCCCTMRWKCGGCTK
49    RRRKKWRSYWKYKSKKTCKYMYGYGGKWYYATCMSYGCCYTMWWSMAAAYARWKTRRTCK
        ***  *  * **.. .   *    ** *  .**     . *** ... ***      *

38    CGTGYCAGYMRAWCKCGCYATTYRSATTKWTRKYRKWTTATACYKMTARRGTRAGYGTCT
49    TCCRYYRRCAGGAMKTTYYCACCASRYWKWWRKYGGTYWRCGGYGCCRGARYRMRYRKSK
          *      .  *   *.:   *   ** ***       . *        *  * ...

38    CWRYKMAYKKTKKMYWRKACKMKKYAGWSYYRRYMKKGGCRGRRTAGYAARKAGYCYRGA
49    MAGCKCCYGKCKKCCTAKGTTCGGCWAWSYTRGTCGGSSYRRGRWRRCRRAKRKYYYRKT
          * .* * **    *. .     .*** *     .. *  *       * * **  :

38    TCCYGTCRTTTCYRKCKAKYRRRATKMACSRRMCTYRYRKRYWMKKWWSYRCKATRWGRT
49    KYTYTGARCGGTTAKAGTKCRGRGAKACTGGACACTGYRKRYWAKKAAGTGYTMYRWKRK
      .  *  .*      *. :* * *.:* . .   .   ****** **  .   .  ** *.

38    GGRMRTAATYRKTKAYYKACKKGGGRYACAAKRGTMYWMWRCMAKARKYWRYKCGRYKGR
```

```
49          KRRMRWRRKYRKWKTYCKWYGTKKRGTRAWRKRACMCWMWRYMGKGRKYARYKYRRCKAG
              ***   .*** *:* *   .      .  **. * **** *.*.*** *** * *.

38          AWRARWYKYKYAMWYMKKKSYCYRRYASGGCYYASRCKYCRCYYRCARGKACTCGGWKYY
49          GTARATYGCKYRMWTMKTTGTTCARYRSKRSYTWCAMKYMAYTCGYMRRGMTKYATAGTT
            .     *  ** ** **...    ** *  .*  .  **        *    . .

38          YCCTKGGTTAGAGATYMWYMYAYKYKGGKCCTGKTTATWRWRMYSKGYWTYGTGYAYWAY
49          TAMGKRRYYRKWKRYYAWYMYGYKYKKSKTAATGACGAWRWRMYSKKCWKYKCACMTTTC
             .  *          * ****.**** .* .:  : .:********  *.* .     :

38          TTTCTKGKCGRTAYKYAACCKYWKSYRTCKKAGGRRCTRKTKTAMCRTATKTRRSARRMY
49          ACWAKTRKYSGCRTKCGCTTKYWKCCAWYKKRRKRRYWRKGKYWAYRCGCKKRRSTRRMC
            :  ... * .    * ..  ****.    **   **  ** *    * . *.***:***

38          CRKTKRYGWMSCMTGTKTCRSRCRKRARWRWKAKKCCRKKGWRTRTWKKWCYKKYRTTWK
49          TRTCGRYRTMGSCKRKKCYRCRYAKGGATGTGGKKTTRTKKTRYRYWKKTTCKKYRYCTK
             *.  **  *.. . .*  *.*  * .     .**  *.*  * * ***   ****   *

38          RRRGTAGKKTGTKKCYATWCAAATTYKKSAYAYRRARSKACATAYSKKWRRKRKKRWMGM
49          RRRSCGAKGAAAKKMCRKWYWMMYKYKKSMTRYRAGASKGTMWWCSKKWRRKGKKATMSA
            ***. ..* :.:**   .*     .****   ** . **.    ******* **   *.

38          RWRYKAAKMRKAKRKKRKYYYRCTKTMRYKKYKWWKCRYWGCCRKTGGCKWKYRKRWGCW
49          RTAYGWRKMRKCTGKKRKYYYRYYKAMGTKKCGTWKYGYWRYYGKCTATKWKYGKGWRYW
            *  *   ****.. ********  *:*  **   **  **    *  . **** * *  *

38          TYKARTRCYCCCSYYYATKAKATKKAYRRKGKRMYKYYKKCRTCYYGAKSMYAAARKKYC
49          CYTWRGAYYYYYSCCTMKKMKGAKKGYRRKRKRMYKYCKKMGWYYCKMKCMCWRMAKKYS
             *. *   *   *    .* *.:**.**** ****** **    *   *.*     ***.

38          YYKYYTYWCCCTCCYGRGKTRGWMAYSYWWYGYKGWMWKAARTTCTWTGYYRYAAYRSAK
49          YTKCTYYWAYAGTYYKRAGYGKWMGYSYWWYKYKKTAWKGCRCAAAWCRYYGYMWYRSWK
            * *   **. .   * *.    **.****** **   **..* :.:*  ** *  *** *

38          YACMCRKYKRMYRTATGGYWSCKATATRWRYRTRACRRWYTTAYKMMRYCTTGMRRRGCG
49          YRYMMRGCKGMYRWRYRSYWSYGTCMCATRYGCRMMRGWYKCCYKCCGCTGGAMRRGRTT
            *  * *  * ***    .***  :     **  *  * **. .**       .***

38          TTTATGRKAYKCGYWCYYRGWGWTCYTGSKKWAYYCMMYRWRCCKWTAKWAKCTARRTRK
49          GCYRAAGKRYKYRTWYYYAAAAAGYCATGKKWRYYYMMTAARATKWCTTTTKTCGRRKRG
                :. * **   * ** . .    : .*** ** **   *. ** :. :*  .**.*

38          RKRMKKMGGTARRACYMWYACWKKYGCGYTTCCAGYRRSGRGACMYSRTGYCCTCRYTGC
49          RGRMKGARKYRGAGMYMWYWMTKKYKSKYYWMMWKCARCSAACYMYGGKRCATGTRTCKA
            * ***        .****    *** . *        *.. .. **. .  .   *   .

38          KYMRATKSYWAGRGRGYMKRYGKKKTAGGGSCRKRWRAACCMSKRAGRRRMCYWAARRYW
49          KTAGWYTSYWWRGKARYMKGYSKKKCWRRRGYAGAWRRCTYASKRWRAAGCYTAGGGGCT
            *     .***      *** *.***     .    ** .   ***         ..

38          SKRYAGCCTGCGGWCKKWYCRAMTGACYRCTGKAASYRWYKGTGMRKKCMKGYKRWGAAY
49          CGRYGAYGCRYRKWMKKTYTRGMWRRSYRYCTGWWSYRWYGRKKAAKKYMKKCKGWAMGT
```

```
                 . **..       * ** * *.*   .**      *****  .   ** **  * *. .

38               KKKSKKKTMKKYKRRWRKMRWKCMRYKYKARTWYWWTYATGCYRAGCMYKRCTKASKYKT
49               GKKSKKGGMKKTKGGWRGCATGACATKCGMGKTCTTCYGARMYGRRYMYGATGKRSKYTW
                  *****  *** *  **      .   *    .     *.:  *    **    * ***.

38               TRGRTMRKRKYMTAGKWGKTWGCYYKAAWYRYTKKCGKCGTYMKAGTCAGYKTCCTACSR
49               KRAAKCGKGKYMATKKWAKCTRMYTKMRTTGCKKKSRGSTWTAKMRAGGAYKKAYYWYSG
                 .*. .  * ***:: **.*    * *      .**.  .    *  : ..**..    *

38               TAWKGACGCCYCKKAWYRRYAYYKGKTGYTYRAMGGKGYTKRRWTRRGACGWGKYATKRY
49               YRTKRWMKMMYYKKTWYRRYRCYKRKWKCYCRWCRKKACAKRRWKGGTCTATAKTRYKGC
                    *      * **:*****  ** *     *    *. :****.   . . .*   *

38               RMYCRCKCTKRKYMKAYCRCGKYKTCAGKAYRTYKYRGTATWSAGKYRACGCCTYYSYRA
49               RMYTGYKYWKRKYMGCYYAYRTYKAATAKGCRYTKYAAGTAWSCKKCGGYCYSCCCSCGM
                 ***   *  ***** .*    .**:.:.*. *  ** . ::**. *  .   .   *

38               KSKMKKTGRKCRGKTTWGRGYRRYRRCRKYWRCRYKKTYKKKYYWGARTKYRYYKRGGKT
49               GCTAGGYKAGYRKKYYWRRKCAAYGRYRKYWGMRYKGCTKKGCYWKRGKKYRYYKATTKW
                  ..        * *  * *    * * ****  ***    **  **   .******   *

38               YRTRKTRKMWKTGWGCYRATTYRYWKRTMGYAAACACAAAWYTAGGKRKWYMCKGYCAMG
49               CGAGKWRKMWGCRWATCACCGCGCAGAYMACGRGGGTGWRWYWGRRKRKACAYTAYYRCK
                   : * ****   *.   .         *. . . . .  ** .  ***    ..*

38               GCYGGGRAMYMTKCKGMYYKTTMWYAAGAGTKSKRYKTCTTKKYMYWCAYTRYTAAAAYS
49               RYCRRRRRMYAKKTKAATTKYYMWTRWKRSKKGKGTTWYYWGKYCYWYRCCACCMTWRYG
                       * ** .* *.   *  **     ..*.*  .     ** **        :  *.

38               YTTTYYYWCTAAACWWTKCRKRKGRAKCMKYRMACCGKGGKRKACCGRKGKKWKGRRKKT
49               YYYYCYYWYYMRRYWWKKTRKAGRARKYMKYRMTGTAKKTKRKGTYSRKRKKWTRRRKKC
                 *    ***      **.* **     * *****:  .*  ***.  .** ***. ****

38               YMAGSTWACCGRWKKWKRYRWCRARTCCTTYRRYTCMKGCYKAGKSRRKMKWWAKSRYYR
49               CAGAGAWWGTARWKKAKRYRAARGGYYYWYYRRYYYMKCYYGWAKGRRKMKTTRKSRYYG
                   ...:*   .**** **** .*.      ****  **  *   .*.*****   *****

38               SGAAASCARKKYKK
49               CTTCTCMGGKKYKK
                 . :.:. . *****
```

The pairwise alignments of two accessions 38 and 50

```
38              AKYWWTAWMGYMRKKRGCMRYKMGCYRYYAGKRCKATYGCWWYWCTGKKRWSYKWRKTGW
50              RKTAWYRWMRTMRGKRAYMACKMTMCACYRTKRYTWYTKAAATAYCCKKRWSYGWRGWSA
                 *  *  **  ** **. *  **     *  ** .    .       ****** **  .

38              RSRWWYMRWMGWYYRMKSYKYWTASGTTRRYSRYCKAWAKGYYRYTGSMKYRYMYWRYCR
50              GCGTTYMRWCATTCRMKSYKCAYCSAGCARYGRTYKRWWKKYTGTCAGMGYRYACWRTMR
                 .   **** .   ******   .*.   **.*  * * * *    ..* ***  **   *

38              MRYMYKRRYRRMRTGGKAAARYAGGTYGGYYCACYYMRYYYYRRYMMMWRMRYRWRKKYR
50              CRCMYKRRYRRMRYRRGGCGRYRARCTRRYYYRYTYMRYYYYAGCCCMWGCRCATGGKYR
                 * **********    ...** .     **    *******     **  *     ***

38              RRYKWTYRRKYYSRRMMYCYYWMYSTRRYYKCYKRYYKSYWWRYCTRKMYYRYMKTTMRY
50              RRYGTKTRRKYYSRRACTAYYWMYCARRYYKAYGGYYKSYWWRTTGRTCCYRYMKYYARC
                ***  . ********   .*****.:*****.*  ********   *.  *****   *

38              GWWWSMMRARWRYRGYYGGCCGTCAGAYRCWKAGGCCYCGGCRKCWRYRRKSGAWYWRRR
50              RWWTCCMARGWRTGAYYRCTYRYMWATYRYWKRKRTAYTAATRKYAACAGKGKWWCARRA
                 ** . *   **  .**        .:** **    .* .. **      *.  *  **

38              YMRKWRRGWYRYKCTTYKRYRKSSKRRCSTRRYRSTACKARYWYSWYRKKSWRYWKTTYR
50              TCGTAGGKWYRTKYWYTGGYAGCCTRATCWRRYRSAGTKMRYWCGTTGGKSWGYWKWYCG
                   .    *** *      *  ...*  . *****:. * *** .    *** ***

38              MYYYRCWWCWMRRKRKTARKRRMYMRYYKMRYGGYRRTRYYYYRRRCKYMYTRGTCMKRS
50              MYCCGTWWYWMRRGRTGCRKAACYCATTTCRTTKTGACGCTYYRRRSKYMYKGACTMGAS
                **    ** **** *. .**   *    . *          *****.****. .  *  *

38              YMKTRRRAGGRACCMSTTKYYRRKKSRYRKRWWKRMYYYACAKTAACCYTYGATRWGKRG
50              YMTCRRRWTKGRYYMSYCKYYRRTKSRYRKRWWGGCCYTRYRKYWRYMYYTRWYAARGRR
                **. ***       **  *****.*********    *    *     *         *

38              CTGKRATCMTGKCYCGMCGRKWACKYTAMAGATGTRAATGCTARRCTGCGCTGTCAYRYR
50              YKKKRGGTMWTKTCMSCMKGGARTGTWGAWACCACRWWWRMKRRATARSRYCAATCCRYG
                 . **.  *  *   .           .  .. . *     . *  : .   .: . **

38              YSAGRYRTAKRMAYYKARGAYKSRKGGYTRCYGWMTTAAKKYARTYACRYRWKAAWRKTT
50              TCTARYRCGKGMWTYKTRARYKSRKRRCCGSCTTAYCGGKTYTGCYRTRCRTKMRWRGYY
                 .:.*** .* *  **:*. *****     .      ..*.*:  *  * * *  **

38              CGTTGMCWKYTARRGKKGCCKKYKYGYGYSYGWRGRRYSMTACRKAARRRRYRMRYYKGT
50              YRYGTCTWKYKCRRKKKAYYKKYKCRTRYCYAWAKGACGCYWYRTMRRRGGTGMRYYGSW
                       ***..** **.  ****    *.*.*     .    *.  **    **** .

38              GTRTRMKTMRCATTCKAKMTAKRCRMYRCSCCWYTTYCRCARYMTRYYSRYSWWMCKAYK
50              ACRKRMGGCGTRKYYKGTMCMKRYGMTAYCATWYYWYYRTGGCAWRCYGRYSWWMYKRYG
                . *.**      .  *..*  **  *   .. **  * * .    * *.****** * *

38              KYAGRCATCYKARRMRRTTCGYGYKRKKMYRGTCTGMTTACCTYKMRCTATAACKMWACC
50              KCTCRTTGTYKGGRARAGAAATAYKRKKAYRCKYYCAKKMTTCTTAAMGGYTCGKCARTT
```

```
                    * : * :  **. * *  :.. .***** ** .    ..     .    . :. *

38                  MWMYCRGTGCTCMAKAYTTYTRMSGGGTGCAAGGMYRWSWKTCGWKGCKYKAMAGCWCAC
50                  AWMYMRAAATYYMWKRTYWYYRMGRAAYTYRGTAACAACAKAAAWKRYKYKRMGRMAYRM
                     *** *.:.    * *     * **. ..     . .     . *:..**  *** *.

38                  TGATRYRAYGRTYRYWWKKKWWSRYRWCATGWAKTRYTYKRAGGYRKMTRGCYKRCTYKG
50                  YSRKRYRMCTRGYRYAWKKKAAGGYRTYMCAWRKYRYWYKRRRRYRKMWRKYYTRTCCKR
                     . .***    * *** ****   . **     .* * ** ***    **** *   *.*    *

38                  RCCGCTAAKATTMRYRWGCMCACKYWRYKRWYYKYMWYKRCCCCATAWCKGGCYYWWYTK
50                  GTAAYCWRKGCCAGCRWAGCACYKYARYKATYYKYMWTKRATTGTYTASKKRSYTATCCK
                      ..     *.      **.   .. ** ***  ****** **.    : : .*   .*      *

38                  CYCKRYYKWYRCARYCMGCTTYATGAGATYRCMKWYMGYKYYKGGKTAKCCWCATGKKYG
50                  GCMKRTCGTYGYMRCTMKAGCCRCAWKRYYRMAKTTCKYKYYKKSKGWKTAAAWGCKKYK
                       **     *    *   * .      .    **  *     ***** .*   * . .    ***

38                  YGCCAYKWKRYKCGTRRKGYGMGSWTATKAGGGRWRKKWRTTTRKAYRKCRAGCRMRRWS
50                  YRYYCYKWKRYKYAKRRKACRCKSAWMYTWATAGWRKKWACCARTRCRTYAMRMRMRRWS
                    *    .******* ..***.     *     . . . *****    :*.   *.      ******

38                  YKKTYWTWYRWACKRRRKKTAYRMTYGWCWCKKYCGATMGKAYMYCARTYKKCAMMCKTC
50                  YKKYYTKTTGWRYKGGRKKYWTAMCCRTYTMKKTMKWCMAKRTACACRACKTYCMAYKCT
                    *** *  .    *  *  ***     *        **      *.*     ..*: *. .*  *

38                  CRTTRCMRWCCAYWCMMTKYKWRRWYRRYAGTARRKYKAKTTATWRSKWYYCWCWYYKTG
50                  TGYWRGCGTSMWCAMCCWGCGTRATTGACTRWWRGKYTWGKCTCARGTWCYMAMAYYKWK
                        *    .            *      :    * **.  . :  *..* *     ***

38                  KYRGYTRRTKYTRKKACMKYAACKKKKRWKKKYRKKTKMRYRWRKMRYCKRWKTWYCYRT
50                  KYRRYARRWGCCATKRYMTYGGYKKKGGWKKKYRGKKKMGTGWRGCACYKGWGCACMYGW
                    *** *:**      .*  *.*.. ***  ****** *.**     **      * *      *

38                  SKRKSKGCYRYRMKTWTKYGTYRGATCKRATYKYKAGKTCAGKYYCYGCGGCTAASRCSW
50                  SKRKSKKSYGTGMKYWCKCTCTGRGYYGGTCCKYKMRKYSGRKTCACAMAAYYWWSRYSW
                    ****** .*    ** * *        .     :  ***   * .. *   . . ..     ** **

38                  KWYCYKYRRYAYYKAGYTYRAMWKCCKGYRMRAGKKAACRAWMTAGKTCGATCGRTRYMG
50                  KTYYYKYRRYGCCKWKCYCRWCWKTSKACRMRMAGKCGMRRWMWWKKCTRRYTRRCATAR
                    * * ******.  *      *  ** .*. *** . *.. * **    *        *

38                  KYATSRKYTGAGRYMRTCYTACCSYCGYMMCRKRGCACMGYCYCTRKTAMYYRWMKYTTA
50                  GCWGSRKCARTCRYMGCATCCTYCYTSYMMMGKRTTGTMTCYCAWRKYRMYCATCTYCCG
                        *** : : ***   .   .   .* .***   **   . *      . **   **     .*  .

38                  WCRYYMWTARAGAAMCGGCGTTTCAYKGCRKYRGGGTTRRRKAAWRWRTMTKGKYTMWWG
50                  WYRYYATKGGRKWWMYRRMKKKYYGTKRMATCGRTTCGAAATRRWRAGWAYKKTYGAWWA
                    * ***   ..      *       .. . *    .           .  **      * .*  **.

38                  TYKKTTCATCYAACCKSYRKKKKCWGGYWRKTKMKKATKTKAMCRKKKKKYYRRMTTCYRK
50                  KCKTWYYMCTCRMYYKCCAKKKKAWRKYWRKYGAKTGKKYKWAYRKKKKYCRRMYKMCRT
                    . *.            *.  ****.*  ****    *...* *   ****** ***  .   *.
```

```
38                  TAKTTKTKWMKTTRCTKRYYYYRKYRRRRRGRWKGYKKCCCGRGKKWMRTWKKKKKRTKW
50                  CRKWKGYKTMKGCAYAKRYYYYRKTGGRGGKGTKRYKKTTYKRRTGTCRYWKKKKKRYKT
                      * .  * **    :********   *     * ***    * .   * ******* *

38                  RKTAKYRYKKYRKGRYAGGRGRRSAGCWTKKCTSRYYKMCARMKKYSRYYSCGKGAKKTR
50                  RKYRGCRYKKCRGKAYGTRATGGSMSATYKKYYSAYCKCYRRMKTYSRYYGATKRGKKGR
                    **    **** *   *.      * ..  **  * * *   ***.*****.. * .** *

38                  CTMMTRMRYMRKKKCKTCGAKMYKKRKKRKGAKCRRYMRWRGGTKTYKKWKGMGKRYRCT
50                  GGCMCRMRYMRKKTAKCTAWKMYTKRKKRKKMGMAGCAATGTTKKGYTKWKRAKGGTGYK
                       * ********..*  . ***.******             .* *.***        .

38                  AYMYGTWKKAGGARMYKRRKTKYYKSGKKKKMGGMWWTKRGMRRWGACYYKGTTRMRYYY
50                  MCCCAKTKKRAACRCYKRRKKKCYKGRKKKKMRRMWWWGRRMRRWRRMYTGAGGRCRYYY
                        .. ** ...* *****.* **. *****  ***  * ****   *  .  * ****

38                  KWKAAYAMTCTCYYKWWYKCKKWMMGTAGRTTKKTYGRRCARGRTTGKGGGTTGKCYTAW
50                  KWKRMYRCYYKYTTTTWCGTGKWMMKCTRRCGKKCCRGAYMRTRGYRKRRRYWSKYCYMA
                    ***  *    .   . *    ****  : *  **       * *   *     .*

38                  CAGCSKYKKCTATATRWTRYMMRAYAWYYCYTRWRARTRYGKYMTACTYRRYWRKWGKGC
50                  MWKSSKYKTYWRWWCAACGCMMRMYRTTYYTKAAGGGCRYRKYACCYYTRGYTGKWRKCA
                       .****.           *** *   *  .   .  ** **  .   * *  ** * .

38                  WMGMTSMTCSRYRRCKACWKTGGYYKGSMAAMMRATRMCAYYYYYCRTAKYYTWTGCKKC
50                  TCKMWSMYTSRCGAYKRYWGYSRYYGRCCMRMMGWYRMYGCCYYYYRYMTTCCTKRMTKA
                       * **  **    *  *  . **  .   **   ** .  *** *  .    .  .*.

38                  WAYWTGMRWKWWWTGGKGGYTCGKWSAKKCYYKKYCCCRRMKYCKYYGGTRKYRKMKKAM
50                  WRYTKRCAAGTATKRKKKRYGYKKWSWKKYYYKGYYSYRACTYAKCCCTKRGCRKMKKGM
                    * * .        .  *  *   *** ** *** * . *  .*.*    .*  *****.*

38                  WYRCGCYCTACKMGTCTGGKWGGTGAYWWRCYCMCCAACKRRGWKTGYATTCACGTYAYT
50                  WTRMRMTMYRMKCSYYWRRKWKCWRMYWWRMCTMTYTTYKRRTWGYKCRKGMGMKKYCCA
                    * *        * .     **     ****   *  :: *** *     .  .  .*. :

38                  YACAKYYRGRYSYSKKWRRYRKRWMMAWKRYMKKKACRCAKCKYMRYCKCYKCRGCCATG
50                  TWTMGTYAAGYGCSKKWRRYGKATACRTGAYCKKKCARMRGMKCMGYMKACKYGAYYTCT
                          * . *. ******* *        * ***..*    * * * *. *  .  :

38                  GCKWKYRKRGMYWRTKGYRYSYTTGACTTAKTKKAAAGTRCCAGRWCYKGSYKKRRKKMK
50                  ATKWKYGKGKMYWRKGAYRYSCCGRCAAKMKKKKRRWRGRMMTKGWYCTRCYKKGAKKMK
                    . **** *  ****. .****    ..:. *.**     *  :  *  . .***  ****

38                  ARTTGCTGGYKYYMTMYYKTTMWYRYGAGSKRYKTYMGCAKKGYRSYKTGTGTAACRAYK
50                  RRYYRMYRRYGYYAKATTKYYMWTRTKRSGKGTTYYCKYRKKAYRGCKWAAAARRYGRTK
                     *       * ** .   *  ** *   ..*  . *    **.**. * .:.:      *

38                  WWTKCAKKCRKGTGCAGCKATTCAKRYKGYKKRRGCTRSAGRGKWCYGRRKKATARYTWA
50                  WWKKTTKKYAGRKTAMSYGGGYYRTRTTRCKKARAACGSMSRRKWYCRRRKKTCGRCYWW
                    **.* :**    . . .  .    .* .  ** *..  * .* **   ****: .*  *
```

```
38              CGGCRTWKRYKSRTYMRMYTCKGCYWKRKYKCKRYYRSGTACACRKKYKKMYYWYYMAAK
50              TAAMAYAKRYKSGWYMRMYYYKCTYWKRKCKMKRYYGSCCGYGYGKKYGGAYYTTYMCGT
                 ..    *****  *****  *  ***** * **** *  . .  ***   **  **...

38              MYRKGTTMYKGTAKYYRKMYYCRYWRKCCAGKWAYTAWCSAAKTRKAAAGGRTMTWYKWR
50              MYRGAYAMTKAYMKYYRKCTTARYWRGYTGKKWRTYRTMCMTKKGKGGTTARWCYWCKWR
                *** . :* *.  *****   .****   . **      . :*. *..: .*   * ***

38              MKARGCTCCAKRMYTGRCRRKYMWKKWCRKGKRTYRGTATWSKKYRKARGMGARKMKTRK
50              MKRGKYCTGCGRMYGRGYRRKYMWGKWYATAKRYYAAGTAWSKKCGGRGKMATRTMGYAG
                **       . ***    ****** **  ..** * . ::****      *.:*.*

38              RAGSYGRCGYRRCYCRYTKKYKCRYTTRYKKRCTYKTCARYWCATCTAKKMTTTKTMTYR
50              RRKSYARYKYGRTYMRYGKGTKYGYKKRYKKGSWYKCSGRCAMGCMWWKKMWAYKCMGCG
                *  **.*  * * * ** *  *  *..**** . ** ..*   .    *** : * *

38              YWKRMGGGKACCCCAWCATGMCKGKTTRGYAKKKCGTSKKCAYCKYYKYMGGYKRKSWKK
50              CAGAMKRRKTTYYYRAAGGAAYGKKYYRRYWGTGYRKCKKYMTYGYCKTASRTKGKCAKK
                    *   *:      .. .    *  * *  .   ..**     * *  .  * *. **

38              WGSYAYWYYYCRTYKYGARCGAGMKCRCMYRMCKGRYWYKKMKTKKMGCATRYRAKAKKA
50              TKCTMTACCTYRYCKYKMAYRRRMGTGYCTAMMKKGTWYKKMKKTTCRMRYRYAWKRTKC
                  .        *  **       *       * *   ******...     **  * .*.

38              TYAYCGGRYAYYRGKRYKRKYKKTYYRCCAYSACRTGRATYAAACKTMYYAAGGRKTRRY
50              GYTTTKARCWYYRRKRYKRKYKKYCYRATWCCRTRKRGGWCMRWYKKMYTTGTARGCRAY
                 *:   .*  *** *********  **.   .  *.  .      *.** :. .*  * *

38              ARMRYYCCCGAKTASKKKGKTRCCRRATYCCGTTGTAKYCGYWRKYYYSKTKKWTACATA
50              RRMRYYTTMRRKWRSKKKRKWRMTRGWKCYTTGCCCRGYYRTWGKYCYGKYKKWCGYWYW
                 *****     *  **** * *  *  .          *   * ** *.* *** .

38              AYKCMMCMCCRGKTTRCTARRRKRKATTACGKMACWMKKGCRRYRGGTATCCSMKGRGAK
50              RCKYMMAMTYRRKWYATCGRRRKRKGAGGYKGAWMTMKKKSRRYRRTCTWMMCAKSAACK
                  * **.*  * *     .******.: .       *** .****   :   .*.  ..*

38              RKGYCCRGKCKYMATGGCTSYWGKGGTKKARGSRKRMRWGCRGYYWTGAGWSKKWTGWKT
50              GKRYMTRTKMKTAWKRKMKSYWSKKRGGGGGRSRGAMRWKTGAYTACRMRTCGKWYRWKY
                 * *  * * *   .   .***.*     .  **  ***   .*       .**   **

38              TKKYYYGYYRMRAWGKCCTTRCMAAGGKACTKKMRSGYWKGCTWMCTRWTKAWYARYRGA
50              CKGYYYRYYRMRMTAKYTGCRYARMRRTTTKKGMGCACTGATAAAYYGTGTGWYRRYARG
                 * *** *****  .*    *      .: .* * ..   . :       ..** **  .

38              RMCAACRKTRYKYKYMGYKGKARTRRRYRYRYYACCMSGATTMATATWKRWKRGSKKYKR
50              RCGMRYGGKRYKCKTARTGAKTGYRRRYRYGTCTYMCGACGACMYRYTTGTKRRCKKCGR
                *       .*** *     .*:  ******   :   ... :      . ** .**   *

38              RGAGAAKGTTSYSARYGCRATCKWRKCKYCCKWATTTYYCAGTKYRCCTCYKKAYWWYYG
50              GAGTCWTRWYSCGRGTSMACGYKWRKYKYTTGWMGCCYYMMKAGYRMYYYYKKGYWWYCT
                 .. . .   * .   .  .  **** **   *    **   : **    ***.****

38              AAACAMRRMTTTMTWSYTKTWCYTKMKKWWSTCWTWYRRWKCMYKRMWKASRWYCYATTT
```

```
50          RCGYCCRRMCKCCWTSYGKYAYYKGMGKWWSGMWYWYRRWKTMYKGATGGSRWYYYCACY
             .. . *** .    ** *   *. * ****  * ****** ***    .**** *.:

38          SYATWTKWRKGAWAGTSRCARAMMKRYRGTGCCTTKWMWYMTKKKMYWRMKACKMKYKYM
50          STRYWWKWRKRRWWAASRYWRWMCKAYRRCTATAAKAMWYMAKGKCCTAMGGTKCGTTTC
            *   * ****  * .:**  * * * **   . ::* ****:* *    * . *     .

38          CGWSYRRTRYYYCRGYKTRTAGYSKRWGRCTCRTTYKGMKYTYKAKYYYWTRRGYRAACK
50          YAWSTRRYGYYYYRRYTAAWRRTGGAWARAAAGYGYKKCKYKTKGGYCCTCGATCAGRYK
             .** **  *** * *.:     .  *.*.:.   **  **. *. *         .  *

38          KSRRMCTKCACKGWWWYKGGRRRKTKYGAAKAGGAAMMKAAKAKRWKKAGGWMRKGTKKY
50          TGGACACKYRYKTAATCKRAGGAGGTCRRWKCRRWRCMKTTTMKRTKKCATTCATRKKKC
            ..   . *   *     * .     .    *.     **::. ** **..    . .**

38          YKGTRWACKMMRGMAKSKYCCCKWRGAKRGYKKTGCKMMKKTCKKAYYTKAGYRYYYKSR
50          YKACRAWMKMMRTMRKCKCYMMKTATGKAKCTGGCATAATKKTKGGYYKKWRCGTCTKGA
            **. *   **** * *.*    *   .*   .   ..  .*. * .**.*       *.

38          YTGTGCWYSYKWAGKRKKYCKYGCYYYYRARATGGTTCYRGAGCKRKCKWCYKMMYYRCK
50          CYRYRYWYSYKAWRKGKKYSKYRYYTYCAWGTWRAWAYCGSGATGAKMKWSYKMMYYAMK
                  *****   * ***.**  * *    :  . :   ...   * **.******  *

38          GRTMKKCCRTGGWYRRWGYWMKKGTRCRRYRRYKCGRMKYTATTCCGTCAGGWWRKTRAW
50          KGAMKKTYGGATTCAGACYWMKKKWRARGCRRCKYRRMKYKRCYYSKYYWKKTTGTWGTW
              :***    .       *****  *.*  ** *  ****.    .         .  :*

38          TCCKMRRSGARKGATTGWRAYYTRCKGKSRCGMYRYYCTGMMTACCACGGATTAKKTATC
50          YMMKCRRSAGATRWYYKWRWCCYGMKAKSRMKMYRCCYWRCMYRYMGMRACWWMGTCGCA
               * ***.. .     **      *.***  ***      *    .  ..    . . .

38          TGAYRWKKKKTATTCKYRYYKGAYCGAGKRKSTAYSYYYAYYYRRRRKYYYKYTGRAAKY
50          CATYRTKKGKYGWCTKCRYYKKCCMKMTKGKCGWYSYYYMYCYARRRGTCCTTYRATGGY
             .:** ** * .   * **** .     * *.  ***** * * ***    .    :. *

38          CRTCYGRTYRGMAYSYWWYWGWGSWWACRRYYYAAYRSCATRMCAKKYCWYRAWWYYMKK
50          ARKYYKGYTGTMGCSYWWYWATKGWTWTGRTCTMWYRGYTKRMYTKGCYTYRMWWYTCKK
            .*. *      *. ******.  .*    *     **. :.** :*    ** ***  **

38          M
50          M
            *
```

The pairwise alignments of two accessions 38 and 51

```
38          CYWYMGRRAYYWWYYRKYWCSCTYSRWKYYRYRAMRKMRKGAYKARAAAMCKRWYCTGWR
51          MCTYMSAARYYWWCTRKCAYGTGYGRTKYTGYGRARTMRKTRYGGACGGMYKRWCYYTTR
               **.   ****  **    . *.* **  *   *.***  * . ...* ***     *

38          YKKTYRYRKYTTYGAAYMYWKRWYCKMTAGCRKCAMYYCWACTGMYMRYMYRRGYSTCRT
51          CGKYYRYRKCWWYRTCYCYTKRWTYKACTCTRKYWACYYWMYYRMTCRCMYRRCCSYYGC
              * *****   * :.* * ***  *  :  **    * *    *  * ****  *

38          CCYAACAYGRKKMYWRTCRYWMCACYYMRYKYKYRMGWRKCATRAYRMYGWRTAGTRGMC
51          YMTTCARYTATKMTAAATRYWMYRYTYMRYKYKYACRWRKTTWRRCRMYKWRCGRYRCAT
               :.. *  .**   : ****    ********   *** : *  *** ** .  *

38          YKTKTRWKCAYMKWYWTRYAGAYRYGYRRWTWYMRCARCWRWRRSYMCRKACKSSRRCRS
51          YKKKYGTKGRYMKWCTWACRKRYRYAYRRWWWCCRTGGAARWAGSYMYAGRYKCCRAYRC
            **.*   *  ****        ***.**** *  * . .  **  ***     *..*  *.

38          TRKCKRMMRGRKKSYMKTYYMWMKYRRRYKRAYRSRRMMYKYYWCYSGCRYYYRRATASW
51          WRGYGGMMRRGKKSTMKYYCCWMGYRRRCGGWYRSRRACTKYYWAYSRYRCYYRRTCGGT
             *    ***  *** ** *  ** ****    *****   ****.**  * ****: ..

38          RYYYKYGCTYCKCYRYYTRKMYWSSYYWKAKRACARRSYKRRCTTYKWRYRKWRYRRWWA
51          GCTTTYAGGYAKTYGYYGGTCYWSSYYWKRGRGGGRRSTGAGYCWYKAACAKWRCRRWWT
                .*.  *.* * **  . ********  *. .***       **    *** ****:

38          YGSYWWMKYRTGGRTTRYYKGRRYWYKMYRYAAGRYWMYYSRKRWRCTCWMWSMYMTRTT
51          YRGTATATYRCAARCARTCKARRYWYKMCAYCTARYWMYYCRKGARYYMAATCCYMKGCY
            * .    .** ..* :*  *.*******  *.:.******.**  *      . **.

38          YCRTGCKTCTAYMGYRTGCMTATMMYMARWAAMSGRCTGGACAGTGKMYWAYAAKRKKAT
51          CSAYAYGYTCRYCTTRAATMAGAMMCCMGTWRCGKRMCKRWTGTAAGACARCGGKAGGGC
             .  .      *   *:. *:.:**        . *      . :.      ..*   .

38          KSCCRCAGKKTRGTAAKAGCCCCAKMRMYACTRMCCYATSGTTWSYTTCGKWCYTKMSYW
51          GCYAAYRRKGKRKARGKRTYYYMMKCGACTMWAMYYCCKSKKWTCYGATRKAYTKKMSYT
             . .    * .* : .*       *    :   *   ..* .  .* :  *   .****

38          WWSGCCGTYRMRWKGMYAACATRWKCGTGCTYYGTTTTTCYYRAGSRMTWTGTYMRRKRY
51          WWSKYYKYYRARWKTMYTTTGAGTGTCCAAKYCRYWWWKYYCAMTSRMWWWRCYCRAKRY
            ***     ** *** **:: .:      ...*      . *    *** *   * * ***

38          RCKMCKATAMARYGACKTTKMYWKMTCATCKACKGMGWSRTCGTYAAARYRYYGRTAGYY
51          RSKMMKTATMMRCACAGWYKCCTKMYMMYSKGTKRCAWSRKMRGCGRRRCRYYRGWRRCT
            *.** *:::* * ...   *   **    .*. *  .***.    .  * ***

38          KWWGGAGGCYRAAYKGKTTTACYCACKTYGRATTTYGYATKMCTRWYRMYGRYYMKYKTA
51          GWWAKRRKYYRWRYKKKCYWTMTYRYKYCRGGCCAYRYGCTCACGTTGCYKGTCAKTKYR
             **.     **  ** *   :     *    .  :* *. . .      *     * *

38          KYYYMMKWYAAGCCCGCRAGRGRRCASAKSRAWCYMCTGTKRMTSRYYWKYRRAYRKACT
51          GCYYMMKWYRMCTTTTMGRKATGGYGSMKSATTYCCYCRKKRMCSRYYWKYGRRYRGGGG
```

```
              *******                .* ** :       .*** ******* * ** .

38          KMTMMKYMGCRKTKKKMYKRKCGMGCRYSGATAAWKTMKYRKWAGYRRKKGWWCYCMMMY
51          KCWCCKYMKARGKKKKMYTRKYRMTMGCSTGCGGTKGATYRKWRRTGAGGTTAYYYMMMY
            *    *** .* .*****.**  *    * . .. *  .****           * ****

38          KKRRYKKGGWAKTAGYMYKRRKTAGTTYYTKCTCYWRYWKKSATGTRCRTCRGSARTYTR
51          KKAATKKRRATKYWRYCYKRRKCTTKCCYGKYATYWRYWKKGMCAYRARAARRCTAGYAA
            **   **   :*   * ***** : .  * * : *******.  . *.*:.* .:  *:

38          YKRGCMCYTKYSYKKMCKRAAKKGGTYRKGRYKMKRWYWRYGTKKMKKKKSYWWMKTYTG
51          YKRRMCYYYKYSCKKCYGGTGKGRACCAGKRYKAKRWYARYKCKKMKKTTGTWTATWCAT
            ***    * *** **    :.*  .     *** **** **  *****... *  .  :

38          CCTGCYGGCYYSAKCCMCYRYYRCAGCRYCYGTAWRMRKGCAGCACYCKTAAAYKYYKGG
51          TTGCTYARSYTCGKGYCMCGTCGYMRYRYTTKKRWAMRKSMWKYWYYTKCGTGYKYYKKS
                 *. .* ..*             **   . * ***.      * * .:.***** .

38          YTCCCKCKMGYWWYTATKKGTYRCYAKKYMKKTYCGAYYKYKKRTMMYATAGTCKKYKGR
51          YGTTAKAGMKYWTTKMKKKAGYGYYGGGYMGGCTTTCTTKYKKGCMMCMYRAYYKKYKTR
            *   .*. * **  . .**. *  *.  **      .  ****  **    .  **** *

38          YCRTAYYTARYWCSMAKTRKRKKAGGGCWKKWRTCGTRKKAAGGYAKTWTRGRRCGCCSY
51          CYGARCTYRGYTMCMTKKGKGGGGARKYTKTWRYYRYRTKCWRRYWKYWAAKGAYRYYGY
               :      *  .*:*. *   ..    *.**    *.*.   * * *:        .*

38          KYYRMATCRGARKCTGYYRMRYYWCCYKWRKYRTKTTCWKKRYKGTTGCYRTGGAMMKRM
51          KYYRMRAAATCRTYYKYTGCGCTAYYCKWRGYRKKKKYWKKRCTCGCATTGCRRMAAGGM
            ***** :.  .*.   *          *** **.*.. **** .   .           *

38          KYRCCCWGATCYTRRYTYCCRYGYKRMCRRKKRYRAYKGCATCYKKGGRCCAYYYKKKAK
51          KTAYATWCGAMYYRGCWCSSRCRYKRMYAAKKATRRYKRTTGTYKKATGYTGTYYKTKMK
            *   . * .: * *    ..*  ****   **  * **  :  ***.    . ***.* *

38          MYGYCTGMTWGKCMAGGCTKYKKRSTCWMRWYCRYAGWRKAYRMCTCTAACRYRCKTRRR
51          AYCYYYRACTAKYAMRRYKKYKKGCAMTCAWYMRYRRTAKGYRCGWYWMRYGYRYKYAAR
             * *      .*      .**** .:    ** **    *.**         ** *   *

38          ATTRYAGKMYAMATATWYAGRKRWKRKKGTRATCGRRSYSYRYCWCCGACGTKRMWCMKY
51          RYCGTTRTCTCCMYRYTYRTRTGTKRKKRYAGYMKGRGYCTACSWATRMSCATRMTTAKT
                 : .  .      *  *.  ****   .    *.*.   .*.   . :.**   *

38          TCKRYWACWACGYCCTTTGWKCWYRKCCTCSWKYWWYGTKGGMGKKKMWWTATKKYRGMW
51          GAKGCTGYWMYAYTTCCCCWKYTYRKMYYYSTGCTWCTGKRRARKKKMWWYRWGKYRRMW
             .*   . *  .*      **  ***    *    *   *    ******    *** **

38          YRWRWKRTAAACWACCACYKGTTRMCCCGCYKTKTTRTTGAGAYTAMYYKKWWYKCKCTK
51          YRWRWGRATTMTACATRMYKAGGRCSMTTTYKCKCCRCCARSMYGRCTTTGWWCGYGTKK
            ***** *:::   ..   **.  * .    ** *  *  . . *     . **     .*

38          KKAAGRKKARWSRTGTYRYTGCTKRGAATCKCTAMYWCGGAKGCYKGYTAAGAGWAWRRW
51          KKTRKRKKMRACGATGYRYYRSWKGRRRWYTAWCACAYRRTKKSCTTCYMCAMRAWARRA
            **:  *** * . :  ***  . *      .. .      :* . .    ..     **
```

```
38     TRMMRYAKGCKSMYWRWRYGKTTRGCKGTWKCMRSGKASCRGYRRCKWKKCKYSMWRMKM
51     CGMMAYRKKYKCCTAAARYRKCKAKMKSKWKMMGSRGGSYRACGAYKTKKYKYCCWRMKM
         ** * *  *.     ** * .   *..** * *  .* *.    * ** **. *****

38     MWRATCRYYYRYWMYKYTTTAMTYYKARYCTRKWAMTKWASWWTATTAYWTYKRTCTRAY
51     MWGWYTRYYYRYACYKYGYKCAKYCKGRYYKRKACCCGWTSWAWMGCGYWKYKAYMGRGC
       **    ******  ***  .. .* *.** .** .   *:**     .**.**    *.

38     RKKCRAGCKYKYGTCGAGTYTTCWGRTTWYWACCKRKKAYRMWRYKYCGGGRWMTMATRK
51     RKTYAMRMKYKTKYYARKYYYKMTKRCKWYWRYYKRKGWTAMTRYKTMKKKGTCWMRKAK
       **.     ***    .   * .   * .***   ***    * ***         * . *

38     AKYKYKYYKAYYRRWKKCRTGMTYAYRYCMRWCAMMTGYWAATTRRACRAKRMKATTRSK
51     RKCKYGCCTCYCGGTGKTGWKCGYRYRYGMRWMWCCWRYTGTAAGRTTAGTGAGWKCGGK
        * **   ..*     *      * *** ***      * .::: *: ..     .  .*

38     KRMMRTARWYTARYKYRGTRRKKYYYYRTGRAWYCYGARKRRKSMWAGRRWWKACCWYKA
51     KRMMRYRAAYCGRYKYRRARRKGTCYYRCRGRWYYCKMRKGGKCMWWRAATWGGTAWCKT
       *****    * .***** :***   ***    **    **  *.**     * . .* *:

38     AKYGTRCATMTCGYYGTRGWAYMGTTSCYKGWMWYRMRGSWYRRYYYYRSRGKYSSYMCR
51     GGYRYRAGCCKYCCYKYGTWGCMKCAGTYKKTAWYRMRKGWTGRTCYYRSRKKYSSYMAG
       . *  *..  .   *    *. *  :. **   ***** .*  *  ***** ******.

38     KTMYYTWSKKYATRCTRRATTAWYYTAYKMGKMMRTYGTTTKRTKTGTATYAKKRYYKYM
51     GWMYYAWSKGCTCAMYGRMYCGWYYCCYKCAKAAGCCRKYYKRYKYRYCYYGKKGTYKTA
         ***:***  :     *   .*** .** .*      .  ** *   . *.**  **

38     AGRKCTWSWKWSYAYYRTRGYTYTACGGRCGCKGGAMKMYMCKGTAARRRYWKTAGRRAK
51     GCGKTCACAKAGTCCTRYGKCKYKRMKRAYRYKATCMKCTMMKRWRRGRRYATYRSRAWK
       .  *   . * . .  *    .*.        *. .**  * *     *** .  .*  *

38     KRAAYGCRTAYYRYGRRKRYKRRYSYMYYRKYKYWWAAKAAGSACTCARKRTTRCYAAAC
51     KRGCYATRYTYYRYAGRKRYKRGYSYMYYRKYKYATMWKMGSCRTCYCRGGKKGMCMRWY
       **..*. * :****. ****** ***********    * ...    .*  ..

38     GMAKTTTTAAGGGRTYRMRYYCCGAKAYKKTRRYYCGTTAKAWTCAYRRSKAAKRCCMRC
51     KARGCCAGTGTARRWYRMRYYTMRRKRYKKARGTCTTGCRGRWKYMTGGCKRRKRYYCRT
             : :. . * ******    * ***:*          *.     .*  **   *

38     TCYKKTCTARRRKKWKMWMRAYMWYACWMYCGCRRYRRRYTTTCCAGSCAMMKGKGAMYY
51     YYTKKCTCGRRRKKWGAAAGRYMWYWMTMYYKSRRYRRRCCYWMMWKCTCACKSKACMYY
          **   .******      ****   **  .******        . .  *.*..***

38     CCTCRYCACYRRKRTGGRRRRYKGGTGGGTKTKARGRYGSRKRMRKATCGYSKRRRRMYW
51     ATGTRTTRMYGAKGGAAARAGYKSAAAASYKGGGGRRYRSRGAAAKCCTACSKRAAGCTA
       .   *    *  *  .. *  **..:... *  .  ** **    *.  . ***

38     RYRMRRCTTYGKYGKRAACKKYYYKYGMRCWTWAKAASYKMKMKGCKACTGYKYARWYYM
51     GCACGGYWWCAKYAKAMWYKTYYTKTACGTACTTKWWSYKMKMKSYKRMYRYKCMRWYYA
                 .**.*    *.** * .      :*  *******. *    **  ****
```

```
38          TKKCKTTGYYCYWKRYKTKTYMGRRCAYYGRAKYRSWYTTTYCAGKTCTAKTCKYMYWWW
51          KKKYKYYKTYYYWKGTKYKWYCTRGYRTCKGMKYRGAYYYYCACRTCTWWKWYKYMYWWW
            .** *    * ***  * * *  *        ***. *    .. .    *  *******

38          TKCYYRTKCRKGAGCAAACMKYTRMRCAGYARRSGAGTGKWCKRRKKWRYMRMKWAKCGS
51          KKTCCRKKYAGRTATCGTYMTTARMAMWKTWRGSRMRYSKWYKRRKKAACCAAKWWKTRS
            .*   *.*    :. ..: *. :**      * *    .** *****      ** *  *

38          CATTYRYRSRTAMMKGYCYSRWKWWRKCRYYRGTACARKKKTKCCTRWKCACTTCGAYRY
51          MTCYCRYRSGWCMMKCYTYGAWKTTRKARYYGCCRYMGKKKKKTTAATKYRYWYYAGYGY
             :   ****  .*** * *. **  **.***       ***.*  :  *      ..* *

38          WWYGTCRRRTAARYRGYMKAAMKKKRRRRKYKKYYAWGGWYATACCKWRMKKKRGAGYRR
51          WTCRWSRGGGRWGCRRYMKTTCKTKRRAGKCKKCYMTAKTCTGWYYGTACTKKRATATRR
            *    .*       * ***:: *.***  * ** *  .   :        .***.:. **

38          CKGMTTARSYWRWKYKCATKRTAYKKAKMYCCMKKKGKAWRWYTCAYGRKYYYKSRYTGW
51          MKKMCCRRCCTAWKYKYGGTAAGCTGWTMYMYATKKKKRWAACWTWCKGTTCTKGACYRA
             * *   *.   **** . . :. .  .**   .** * *         .   *.

38          CAGCKGKRKMMYGYRMKYYCKAYKRARACTTCYTTCWTTGASMYRGGCWRCYWYYSGYGW
51          YWRYKRKGKCCTKYACKYTSKMTKAWGTSKYYCYYYAACAGGATGSSTWRYTWYYSKYKT
                * * *    *  ** .*  *   :..       : ...   .. **  **** *

38          CGAGYTYGGRKKGKATKCYYKASKYARKRYKWKARGGCGAAGKAKTACYCKYKYCCTSGA
51          SACACWCARGKKKTWCKYCCKCSKYMAKGYGAKRGRKYRTWRKGKYWYCYKTKYYAGGAG
            ....   .  ** .  *   *.***  * *  *      :  *.*     * ** . ...

38          RRMYYYYAKKKKYWKRSAGTGTTCRWRYRWKKYYKCCACATCKGGWRCGMWKKWCYKKCC
51          RRMYYYYRKKGKYTKRCCAGAKCYGWRYGTKKYYKTTMYRCYTKRTRAACWKKWTCKKYA
            ******* ** ** **... ..   ***  *****       .   *.. ****  ** .

38          GARTWCGYACWAATKTTYRKRKRKRTGYKRGAGKTCKSWKTWCCWCMGYYKYARGRTYMG
51          KCRYTMCCRTATGYKKYTAKRKRKRKKYKRKTAKGTGCWKWWYTTTMKYYTYGGARWYMR
             .*        :. *.   ******. *** :.*   .** *    * **.*. .* **

38          CATGKAACARRCTRTGKCAAGYTGAGTCCAWRYRRGATRMAYYKKAGWKRYACCCTATYA
51          MRWAGGRYTGGTAGGRKYGRRYCAMRAMTCWGTAAKRGGMWTYKKTAWKRYWYYTCRYTM
               . .  :   :   * .  * .  :  .*        *  ***:.****

38          YAWKYRSKGGTYGRYRKKGRKKKYKRGAYKKGRKKYKMRRYKMKMRYKKKRTCRKRRYGM
51          YWTKCACKTCCYARCRKGARKTKYKGAGTGKKRGKTKMRGTKCGCACKKGRCMGKRRYRM
            *  *  .*   *.* ** .**.***  ..  * * * ***  *     ** *   **** *

38          CYYGKKKYAGCYRWMKKRWGGKSTKGCGYYMWWYRTKTCYRKTYKKAGKTTCAGKKRYCT
51          MTCKKKKYCTTYAWCKKGWTCKGCKAYKCTMWTTGCKYYYGKCCKKMRKYAGGAKKRCYY
                ****.  * * ** *  *. *.    **    *  * *  **  * : ..***

38          ASGAWYCCTWCYMYWYYRYAYYTAKYYRAMKCCYGYYYKKTRYKKAYKTRCAKWTARSGY
51          WSTGTYYMWAMYMYWTYRYGCCWWKCCRTCKTGYACCCKKARYKKGTGCGYRKTKGGGRT
             * . *     **** ***.    *  *: *  *.   **:****.      * .. .

38          KYCTRMWSYWAAWGATMGMGKWYCSTKYRRMYTTCTAWYGAGYCCRMCCKCYGCCAAYCT
```

```
51      TCYKRCWCCAWWWKGWCAMKKWYYSWKTGGCYKGAYGTCTTAYTGRMYTTACRYMTGYTC
        .  .* *.    * .  .* *** * *    *. . .   :.*  **  ..    :.*

38      WMKYACYYGSCKMAWKKCSAMYRCTGCRRTACKMGKGYMKYTCGAATMCKYCYRGKTARR
51      WMTTRYTCRGYGCCWKKYGGATRYYRYARYRMKCSTKYMKCWTRRWYMAKCMYRRKKRGG
        **.      .   .*** ..  *     *   * .. ***       *.*  ** *.

38      GKCRKCAACRKCRKYATTCAKKWCRAKRYRGWSGMKKKGCTTKRRSKKCGCSCKRKKTTS
51      RGTGKYRMYRKTGTCTCGTTGKWYAMKRYAAWSCAKKGKMKYTGRSKKASMSTKRGGYYS
            *    **  . :   : **   *** .**  **   . . ****.. * **    *

38      GYRGMRGGRYKRYYSYYWGRMRYRRTRTACCRYKCTKRYKKWRYYRYKRYKRATAKTKMY
51      RYRKMRRRATTRYYSYCWRAMRYGRCACGTMRYGYKGACGKWRTYRYKRYKRGWWKWKMT
         ** **    .***** *  *** *   .  **  .    *** ********.  * **

38      GCCCTTKTYYWKRMKAACACAAGGMACRWYCATGMCCTYKRMAGWKKARRKKKTTWTCRK
51      KMTMAGKCCCAGAMKGGGGTGGRRMTTRACAGGAMYTCCGGCGAWKGGGRKKKGCACTAG
            : *      **.. . ..  *: *  .. .*       ..** . ****

38      TWTMGKYYTKSAASRRYSRCSAYATKYGCTAGAGTRRCYYRARTYRWWMKYRKKKYYMWA
51      YWCCRGCCCKGWTSRRYGGTCTYCCKYKAWTSGTCRGYYYRRGKCATTCTTAKKKYYATW
         *       *. :****.  .:*. ** . :..  *  ***  .     .  *****

38      TCRCATCGKRGCTRCKTCAATTYRKYRGTTTYCKAARKCKWGGTWTCYYYMMYRCKRKRR
51      KMRYMWTAKGKTCGYKYYRRKWTATCGRCGKTYGRRAKMKARAWTCSCTYMMYAMKGKRR
        . *    .*      *    .   .     .      * *  .   .  ****  * ***

38      GGKCTATTYCKCTYTGWWWGRSRCKRKGTCRGGACMCRCCCCRRKRRRKRKTMTATCGAA
51      ATKYWWKGCYKAGCKAWTASRGAMKRKKWYARRGACMRTGTARRKRRRKAKAMKRYYRWW
        . *   .   *.  ..*  .*.  ***      ..  *   .******* *:*.

38      GGWWKCAGTRACWAKTKRMKKTMKGRRRCKKKSGTYYCMKYYWYACMGGTCCTKTATCYR
51      KKWTKYRRWATTWCKWKRMKKKMKTRGGMGKKSAYCCMMKCCWYWMMKRYYMYKCGCACR
          * *     : *.* *****.** *    ***.    **  **  *      * . . *

38      WRKCYRKKSRRYKYRYMKATCGKGTKTGTKATKTAATAYKCCTSATRCSCAYKCYMSKKW
51      TGKACRKGSRRYKCRYMGGAYKKRKGAAAKRKKGGRYMYKATCGMYAYGYRTKMYMSKKW
          *. ** ***** *** .:  * . :.:* .* .   **.  .    .   * ******

38      RTKCARWWRAWKMGTACGMWRGGKKKKCGRGGRKYYCKMKKKRKAGGCYTWKRYYGRYTK
51      GGKTRGTTGWAKCTAMYSCTARRKKKKARRKRRKYYYKMKKKGKTTATCKWKGYYKRCKG
          *        *  :  .     ****. *  **** ***** *: .  .** ** * .

38      KCTKKAAAGRWCYCRKMKGATCGRMYKYYKRRK
51      KMKKKRRWRRWYCYRTMKRWGTTRMYKTCKRRG
        * .**    **   *.**     ****  ***
```

The pairwise alignments of two accessions 40 and 42

```
40          MWRMKKGCSRRATGYYWRTYWGGSYKMYRRKKGARTYKWKTAKYWGYYWYWKYKRRASWK
42          CWRMGKRTCAAGGACTWGWCWAKSYTMYGRGKKCRWYKAKGGKYWKYYAYWKYKRRGSWT
             *** *  .  . .  *   *. **.** * * .* ** * .*** ** *******.**.

40          YRYAYGSMMKYYRRYKACCRAWMRRMMAGWKWRRRRYWMMTRYRWRRRYYWKYAYMMCSW
42          TRYRCTSMMTYYRRYKGTTRRTMRRMMMKAKWGRRRYWMCYRYRWRGAYYWKYGCAMMSW
             **   ***.******.  *  *****   ** ******  *****  *****.  * **

40          MKYRSCMMKWRTRKYYRWWRTYYCMYRYYWRYKMYMYGSYGCRKAAGYYYKTWYYYTKYY
42          MKCRGMMAKWGYRKYYRWWAYYYMCCRTTARYTAYCYRGTRMRKRWRYYYKCWYYTKKTY
            ** *. * **  *******  **   *   **. * * .   **   **** *** .* *

40          YKRKAKMMSGAGMCCRATRCWYKGTCYMWRYYRRYYKWWKKWMRWKATKYKRKYYRSRRR
42          YGGKWKAAGAGAATTRWGGAWYGRYMYMWACYRRTCTWWKKTCAWKWKKCGRKYYRSRRR
            *  * *  ....   *   .**    ***  ***  .****   ** .*  *********

40          GWRRCKWMYRCRKYRAGMYMRCSCYYCMWRKYMRRTYRKWWRYYRAGGCAYRSWYAYRGY
42          AWRAYTWCYRYRKYACTMYMRYSYYYMMTGGYMRAAYRKWARYYRCATMCYRSWYWTRKY
            .**  .* ** *** . **** * ** *   *** :**** ****..  .*****  * *

40          GYRYCRGATTRTTKCYWMKRYKKYRATYYRYYWRYYGRYYRYMWKYMRYCKKWTMWWYYK
42          AYRYYRAGCWAACGYTWAGGYKKYRGYYYRYYWRYYTRYYRYMWKYMRYYKKTAMWWCTT
            .*** *..   :    *   *****. ********* ************ ** :***  .

40          WKACGGWMMYWMYKKYCACKARWTTAYRYGWYRTYKKYCATTCATAAATAGCGKCYGTGR
42          AKMATTWMMCWCYKKTYMTGMGAKCGCAYRTYACYGTTYWCGYRCGCTGGRYRGTYAAAG
             * .  *** * ***        . .  *  *  * .        ..: .     *.:.

40          ACGCTSYGATGGCCGTYWTAACRYRWAAAAYAACAYKAACGYYKACMKAWSTCCGMGCCA
42          MYKTYSYKRWTKMMKYYTGGRTRYRWTRTGCGWSRYGMRMKTCKWMMGRWSGGTRMKMYR
                 **         *  .  ****: :. . . *       *  *  **    *

40          GATTKGCMTGKYRMKRCTCTAGGKAATGRGWRGKMKCCRMKKWYTWKSYMRTAATGATTA
42          RWYYKRSAASTYGMKGYATAWAAKTTGCRRTGRTATTYRCGKTCCTKCYMRKCGYRRKKR
                * . :..* **  : : ..*::  *    . .  *  *    *.***...   ..

40          RSYKATAAKYGTTGMARTKCYGCRKYCWYMYGAAYWYKCYCKGGKAYWRYATAKKATARK
42          GGYKTATCKCKAYSMGGKKTCATRKYMTTATRWRYWYKTCAKRRKWYWRYRYMKKRYMGK
             .**:::.*  : .*. .*  . ***        ****  .*  * ****   **    *

40          WMRKTTAKAASAMCCTRKAKKWGRKCYSKMMTTTGTKCWWWRWGRWTRAWKAGAAKYAGC
42          AMRKCARGGGSRAYYYRGRKKWARKYCSGCCYYYRYKAAATRTARWGRMAKRRMRKYRRT
             *** :  ..*     *  ***.**  *        *.   * .** *  *    **

40          SAAGWGAKRRMYKTAAAWAARTTACMMKRRRCAMCKWMKCGTGRTMKATRRGSKWYYYWK
42          GCCKWRRKGRMYKGMMCTTCRWCCYMMGGGRYRMYKTAKTTYRRCMKRYRRRSKWYCCWK
            ... *  * ****   . :.*  . **   *  * *  *    * **  ** ****  **

40          GTTCTRKKYKMMAWKGTKAAYCKCRRWYYTTCTTGAGKYRRYRYMYKWAGTTACGARRYT
42          KGATARKKYKAMWWTKCGRGTYKYRRTYYWWTKYRMRGTGACRYMYKWGKACCYSWRRYY
```

```
                      : :***** * *.     .  * ** **    .         ******. : . . ***

40                    CKTKYKGKARCTCWWGYMKKCGGGCTKYRKSTAKRTTYYGRYYTCRRRATKYYRMGKTKR
42                    MKWKYKRGGRYYYWWAYMKKTAAKAAKCAGSYWTAYWYYAACTATRRRWYKCTRMRKYGA
                       * ***   .*   **.**** .. .:*   *  .    **.   : ***  *  ** *

40                    TYRKGMCTTTGARWYWYYGTATTCRGRTYYKSYRTRAKYKGCKCTKATTCKTKRCTCAGG
42                    YYRGTMYYKCRMAWYTCTKYWKAAAKRWTCKGCRYACKTKSYGTGKCCCTGKKRGCMMKS
                       **  *  .    **      .:.  *   *. *  .* *.     *.    .**     .

40                    TKRRMAYWKMTGAGKCGTATGYCYAGTGTYGGCMCTTGGYATCKTGGTRCRYWKKTTWTC
42                    CKAGMRTWKCYRRKKYKKWYAYYYGRWKWCRRYCYGWRACRKYKCRRAATGYWKKKWWYS
                       *  *  **      *  .  .* *.              .  . *   :   ****. * .

40                    YAKYCTCCYRYTACRATAWCCCGSYYKCKYYRATKARCRTGTRGAAAAGTAMYAGYMMYT
42                    YRKYYWMSYRYCGGRGAGAMYTKCYYKTKYYRRGGTGYAAAAGATGWGAWWMYWKTMMYW
                      * **   .*** . *.:.      .*** ****    :    :.: .:. ..  **   ***

40                    AGCYCCRYKRKCMARYMRWAGCCTTAAGGCRTYRCCMMCWTATYCCCCCGRGGCCGACRY
42                    WKMYTYRYKRKYCWGCMRWRATGAACGTTAAYYRYMMAMTCGATYYMTGARAATAKCARY
                         *  *****     *** .  ::..  .  **  *     .:       .*.. . ..**

40                    YTCCACKAYYCWMSRYRAYWMRCKRCTTTTRYSWKMSWKKKKKKYRAAMKYWRCGKMKKY
42                    YYYYRTKRYYMWMSRTGGCWMRYKRYKKYGRCSWKMSWKKKKTKCGRGCGYWRMRGAKKY
                      *     * ** ****  . *** **  ..  * **********.*    .  ***    ***

40                    TYRCATATTKRRTKAYRKTYYYTYKACRWKKKYTKKRTKRYGCAKRACRCTATTTRCTCT
42                    YYRTGYGACKRRYKWYRKCYCTWYKGTRWKKKYWKTRWKRYAYRKRGARYWRYYYGMYAC
                       ** . .: *** * *** *    **. ****** *.* ***.  **..*          .

40                    RYYKKRKGGKCSWARYYGYYYRYRKWCWWRKMWTGKRKKGCATTMSKKKKKGGKYSRTAA
42                    RYYKKRKRRKYCWRRYTRYYYGCGKWYWWRKATWRKGKKAAWYWAGGKKKKKRKYSACWR
                      *******   *  .* **   ***   ** ****     * **..     . ****  ***

40                    KGKWRTMMYTASTKRWYRWKMMRKYRRTKTGRGAYKCTACKYRKYYRMMMTACYYMCCCC
42                    KKKTAGMMCYRCYTGTYRTKMMAKYRRCKYRRTGYKTCRMKYRKYYRCCMWRYCYAMYMY
                      * *   **    . .  ** *** **** *  * .**    *******  *     *

40                    GWRGYRYGCACMCYWTKTYCCGTCYGWYKKGATCYMMYRARCCCTCTKCRGCGTYKTGKG
42                    RWGKCRCRYWMCSYWYKCYMMAAAYRWCKKRMYYCMMYAMRSYMYYAKYAKYACYKYRKA
                       *    *       .** * *  .:.* * **      ***  *.     :*     . **  *.

40                    KTTTACGTSSACRKKARWYAAYARWCKKGTTTAASYWCCKMTKMGMYYYCGMATAAWCTK
42                    KGCKMMRKCGCYAKKGRTTCGTRAAMKGRKYWWRSYWSSGCYKMRMYCTATMCYGCWSYK
                      *  .    ....  **.*  ..      *  .    ***..    ** **   . *. ..*. *

40                    KAATSKWWSACTGKKWYRMWRYYYKKMMKCACAGAKRCTCKYTRTGYRTYAYTCTRAWYY
42                    KRRKGTWWSTTAKGTAYRMWRYYYKKMMKTTAWRGKGYYMKCWGYRYRYTGYCTYAGWYY
                      *  ...***: :  . ************ :.  .*     *      **  .*     .***

40                    CCRSACGGYYTCGWRCCTAAAKTGGYAYAAKRYMKWMTAATRRKMKYRTTKTCCGTYMTG
42                    TTGGWMSKYYGTTAGATCWMTTAAACCYCTGGCCKWCYWMYRGGAKTRCAKWYMTWYMCA
                         .  . **      .     :.:.. .*.:     **      *    * * :*     ** .
```

```
40              GWAKYWMYRKWTCKGGCGYYKMKKWKAYYATGGATGYYYATTCKYCCTWCTTAWSKRACG
42              AWGKTWMYAGTWYKKATAYYKCKKWKWYYWYARGCRCTCGCYMKYSMWWAYAGTGTRGSR
                .*.* ***     * . .*** **** **  . .     .   **.  *. :. ..*..

40              CATYAWRWKTWKTSKAGYGTAGKAKWYRKCTGAATRCKKSAYKRAGATRGCCCMYYYKCA
42              TRYCRWGWKKWKKSKMKYRYRAGMKWTGKAWKTCCRYKKSRTKRRKMYAKTTMCYYYKAC
                     * **.**.**  *   .  **  *.  :. * ***  **          ****..

40              AYSRSWGWYYTTAGCAYRMYWYSYRCGTGATMTCMARATWMGATAYYRKGKCGCKCRACT
42              GYSRSTRWTTKYRTMMTRMYTCSYRYACARWACYCWGTYWMKGYRCCGGKKYKYKMRCAG
                .****  *  .      ***  *** . .        : **  .       *   * *..

40              GRKYMACMKTCAAKSCCYAWGRRKAWGKKCCWKAKGTKRYGWYMRAKCGRRCGMMTWYRR
42              AATCAWYCKCYTTGGYSYMWTRAKRWRKTYYWKTKRKGACRAYMRRTYRGRYRMMCWCRG
                . .     *  :: . .* * * * * *.  **:* .     *** .   *  ** * *

40              KTAGCKKAWRMYRRGAKTTGWKCATAKYYYMYYMSCCGGCGARYMMYRCCTWKATCGRRY
42              KKWKGKKCWRMYRATGKCYAAKYRWMKYYYCCYCCYMATTRWGCMACRYYKWKWYYRRRY
                *.   **.*****  .*  . *    ****  * .  .      *  *  .**    ***

40              CRMWRYKYRYRMGGYRTCCRWGMCRRGACACTCGYTCGTYAYRRGYYAYAWTKAYWYTMR
42              YRMWAYKYGYRMSRTAKSMAWRMYAGAGMGTCTTCCTKWYRTRRRYYGYWWYKGTTCGMR
                 *** *** ***.   ..  * *   .. .         *  ** **.* * *.    **

40              YYGKCAAAYYGYTAAAACARMWMRYRSWYRRYTMGYKGCKAKRYTAGKCTCCRYSKGKRK
42              YYKKYRMMCCRTGTCMTSRAMWMGYAGACGGTKARYKRYKWTRCYRKKMCYYRTGTATRK
                ** *         :. :.  *** * .     .  **  * .*    *    * ....**

40              CTCKKTRKMKYRRKRTGCSYYSRGYKKAGTGTTTTGRAKWRMYSGWRCATCRKKMTTYYK
42              MYYGKYRKMKYGAKRWKYSYYSRTTGKRRYKYYGCCRRKWRMYSRWRYWGMRKTMCYYYK
                    * *****  **   *****   *         * ****** **    **.*  ***

40              MYRMAKCCCRRRRGMWMCCCTCAKTWWAGWCAMRKTCCGCGMKYYATWRRTARACAAKAT
42              CTAATKATSRRRGRCWMTGYYYWKGTAWRTAMMRTATYRAKMKTYWYAAAYWRGYRGTWK
                    :*. .***   **      *      . **.:   . ** *       *.  .. .

40              WCTYTGKMMYMRAYYGKYKTAYYYRTCRCTSCKATACGGAWTAMTRRRTTMTAKRCGTGG
42              WMCYYRKAACMGGYYRKCKKGYYCGCYRTCCTKCYWYCARAAWMYRAGWYCKTKGYRKKR
                *  *  *   * .** * *..**    *  . *.    .  : * *     .:*   .

40              WGCYKAYKRRCRRTYATCGGYMRTGAGAYRGAGGTMMCWGCYGYYTYTCTCYKAMGRKKG
42              WRYYKTYKRRYRRGCTCTRKCMRCACRMCAKGASKMMTAAACRYCYTWYYTCGGCAAKGR
                *  **:**** **  :     ** ..     ....**  ..  *         . . *

40              GTGTRMGRCTCYTGTGCCKYRCKKWYYATGAGMRKYYRTGCRYGMKRRKKKRYMKGGMWY
42              SKKWRMKRGKSYKACATYKYRYKKWYYRCRGKCRGCCGCATGCKMKRRKKTGYCTRACTT
                ..  ** * ..*.. .  *** *****   .  *     .    ******. * . .

40              MYYMYKKKTACTRMRRRKYMYWCWAGCRATWRAMCTYYRKKKMYCWKGCCRACKGCGKKT
42              ATYCYKKKYWMKGMRRGGYMYWTTTAAGGWTGRMTATYRKKKAYTWKRYTGGYTTMRKGA
                  * ****   . ***  ****  :.. .    * : ***** * **    . .   * :
```

```
40          ATTTGTCKKGKRSKYYCWKYTTARTGKCKMGACRCTYSYWYKAYKAMCTAYKAYYKAYMG
42          GCGWRATGGTTRSKTYYWKYCYTRGRGYTCCGTATKYSYWYKGYKMCMWCCGMYYKGYMK
            .    :    .*** * ***  :*    .  .   .******.**    .   ***.**

40          AKKYRYGAYWRRWWKWGYMCGSTCACKTGRYTGMMCAATCAAKAAAKTCYWYRTYCMRAW
42          MKKYRYKWYTGAWWKTKYMMRGCMGAKACRYYRMMGTGCYRRKGRMKGGYWCRWTTARWW
             *****  *   ***  **  .  ..*: **  ** :.    *.  *  ** *    * *

40          GGTRKRRRYMATYMYGTARYACTMAWKCCRRKASRRYCRCTCRCWTGTAWGGTKWYYGTK
42          KRGGGGRRCCMGTACRKCGYMTGMRAKMYAAKWGGRTTGSGAGAACACCTKRYKWCCRKK
                  **        .. *   *  *    * . *   . . .  . .    **   .*

40          GCYCAMGYMYYATATMRAMTTYYKKATCWAKTTWYMYYYRGWWCTMRGTYRRRRACRRKA
42          KMTYRARTCYCRYCYMGGAWYYCKKWYYWTTAATYATYYRRTAMCARAGYGRRARYRRKR
                     *   . * .   * **   *:.:: *  ***      *. * **   ***

40          MAWRYMKCGCCCACCYRWYTTTKKYYCGAYYKKARRRKGKRYYCCKTAYKKTAAKGTRAR
42          ARARYMKTRAMYGMYYAATGCGKKYYMRRYTKKWGRGGAKRYYYMKWCCGKCCGKKYRMG
               ****  .  .  *      ****   *  **  *  .****  * .  * ..*  *

40          RYRWYRYMKWGWYGGGKGKKYRYKAKWCGRYKTWCKAKWTAAAACTKGWYYYCTTTGKKK
42          ACGTYRYMKAKWYKRRKKKKYRYTRGWGARYTCWMKCKTYWMRRYYGRTCYYACCYRKKK
                *****  **   * *****.  * .**. * *.*            **.    ***

40          AMYRGWKRGTGSWTYYRMRMYSWRWKAMWYAAGKRGYTYAMKKKRKTYSRGGYWCRKRTG
42          WCTGKTKRRWKSWYYYRMRATGTAWTRMTTWRRKRRYCYCATGGRKYYSRRACAAGKRCA
                  **   ** *****  .  *. *     ** * *. .  ** ***  .  . **  .

40          RRRMCAYWYRRCYKYRYKWATACCKTWRGAGGGKKKKMACGCTTCCATSCRGAGYRCRSK
42          RRGMMRYTYAAMTKYRYGTTAGTGGGWRRRKRRKGKKMWYRYKYMYGCGTRRRACRYRCK
            ** *  * *    ****  ::.    **     * ***    .   . . *  . * *.*

40          GCKYKKKAKARGRKWKYTYGTGKTCCAKTRTMYKKYSMKKAYTGGKGWYYKYWWGTYMKR
42          AAKYKKKRKWRRGKTGYYYRARKWSTTKYRCCTKKTGCKKGCYRKKAWYYKYWWRYYMKR
            ..***** * *  *  * * : * . :* *   ** . **.    *.*******  ****

40          YKGGGYCTMGGCYRKWAGTTRMRRGYKSACGGYKKGTMYTTKKCMKCGAAGTAKYMRMWK
42          YKRRKYYWMKRGYRTTGRYKACRRRYTCRYRKCKKRWCCCGKTYMGMTCWACMKTMGATG
            **   *  *   **. .  .  ** *..     **      *. *   . .  * *

40          RKKCKSRCGRYYYKGTTCRGTCTWRCWYWKYRYMCYGWKTGACMTKKWYKKWKKCTCKKT
42          RGKTGGAYKGYTTTACAARTCTATRYWCTKYRYMMTSTKWRRYAYKKWYKKWGKTATKKG
            * *  .    *  .. :.*   : * *  *****  . *      ******* * : **

40          CTGYRKCARCKRKTKRCYRGGMYYYRRYKKYTGAWAYMTRCTKKCAAACGYTRGCTRKMC
42          TCKYRTGTGMGAKYGAGYRAAMYCTRRYKTYYRWTRCAYRMYKTACGCTKYKGRTWGGCY
               **. :    *    **..**  ****.*        *  *.....  *.

40          TKCAMKYRRRTGRYSKACAKKYCACYTCTYYWGTTCTCAACAAGCYCCKKGCGGMTGGYR
42          WGYWCTYRRRWRGCGKCYMKKYSRYTYYWCTWRYYYYTCWTGGATYYMKGSMRKMCTTCG
                 .****    .*.  ***.        *      .  ... *  * .   *

40          ASYKGGYSTTCTKAYYCAATATMMMCATKCCWYTRYYKTGGGKGAKGATCTCTMYYYAMY
```

```
42                  GSYKRRYSGGMCGCTYYWWGRKAMAYMKKYAATCRTYKYKSRKSWKRRYYKYYMYCCGCC
                    .***  **     . *     . *   .* .   * **  .*.  *    .  **  .

40                  GKKAMRSSSRRYMMMYRTKTKYATGGTAAAKYYTTKKTAMCGCYARKKYYARRRGMATMR
42                  TKGGMRGGGAGYCMMYRKKYGYRYRRKTRWKTYCCKKYMATAYYMAKKTYWAAGKARYMR
                     * .**...  * ****.*  *    .:  * *  **     .*  ** *        **

40                  RRGAKCKMCKCCCTRCATKWGARARWTMTYGCTKCKAYWYRKRRRRKYRCSMRYMYTRKT
42                  RRKWKMKAYGTYMARTGYKWRTGGGTKAKCRSWGYKMTACRKRGGRKTRMSMGTCCGRKK
                    **  * *      :* . ** : .  . .  .   *    ***  ** * **     **.

40                  KRYWTTKYKAYSYRKTYYRRWMYYYRYKKTGGRKTKYCATKYGCYAYRWKKWKWKYYAKM
42                  KRYWKYKTKWTGTRKYYYRRWMYYYGCKKWRRGTYKYAMYKYAYYMYRWKKWKTGCYMKC
                    ****. * *  . ** *********  **    . **.  **. * *******   * *

40                  RCCKAKYYYAKWGYRATYWTCAWTWRCGKKCAGAWRATAARKKTCGGTKYAATKRYMSWG
42                  RYMKRKYYYMTTKYRGCCACYGTCTAYRKKTWRRAGGGTGAGKAMAAAKYGWWKRYMCTR
                    *  * **** .  **.     .      **      .:.   *: ..:**.  ****.

40                  RRWYCTRWGGYRKGGAAKYGRGRCRGCTYGKCAGCCGKKCKWMRRYYSCTGTAYYRWTTG
42                  RRWCYGATRCTRKCACRKTKGAGTRTAATKTTTCTTKKGTKTCRRYTCYCACWCYRACYR
                    ***        ** .. *   .  * .:  . :    *  *  *** .  .   **

40                  RYKYYMRYRWYGYCRRRYMARGACKGMRRCCKAKRYMTCRRAGYCATCSARYMATKYRCR
42                  AYKYTMRYAWCRCARRRYMWRRWYKACGGMMKWKRYMGTAARRYYRKTGCRCCRYKCRMG
                     *** *** *   .***** *   *.     * ****      * .  ..*    * *

40                  MKRYTKAGKKYRYGAWKTKCAYRAAACYWYCTRYGKKAATTRGKKGGMMKRCCRYAWRYR
42                  CKRTCTWATKYRYRMAGCKYRTRGGTSYWYTAGCRKKCTCYRSKKRAMMTGYTGYWWRYR
                     **  . ..****     *   *..:.*** :   **.:  *.** .**.    * ****

40                  YMMGTAKRYRKSRSRKKAKKYKCAATTGGGWKYRTCKGTKGGWRRKRCYMKYRYKRRKMR
42                  YMMKWMGRCRKCGSRKKMKKTGYRRWWRKRWKCGYSGKKKRRWGGGGYYMKYRCKARTCR
                    ***    * **. **** **          **   .  .*  *     ***** * *. *

40                  KGTAYRKGRKKCYCGKRYSGSWKTTRYKRYMYTKCCARYKKRGYCYYKCKMSRYRMRYKY
42                  KRKMCRKKRKKYYYAKRYSASWKAARYKGYMYCKTATRYKKRKYMYYKYKMSRTGMRYKY
                    * .  ** *** * .****.***::*** *** * .:***** * *** ****  *****

40                  TSGRSKYGTGWGCTKCGRMCTATKKMYRYYYRWGGKGAWRSKRGGGRRGTGAYYWGGGCM
42                  GGRAGKCKCTAATCKSRGCMWWWKKMTGCYYRWRRKARWRSKRRRRRRKYRRCYWRKRAM
                     .  .*     .  *.       ***   ****  *. *****   **     **   .*

40                  RYTMWMATAKYKYMKACCWCAGRRASKTGTRGRRGKMTTMWKRYYRKRKKYTATTTCRMM
42                  ACKCWCTGGKCKCMKGATTTCARGGGKKSWRRGRTGMYYCTGRYYATRGKYYWKAKGGMA
                      . * : .* * **..   ..* ..*.. *  *  *     *** .* **  .:.  *

40                  CKTGCASYKYWTRGKGAGGCWWWWRRTYRWMACGCYYTTKWYRTSRRACTRGTRSMRTKY
42                  MKCTAWGTTTAGGAKAGKAYWWWWRRGYRWMWMRYTYCCKWYRWSRRGYGRRWACAAKGY
                     *  . . .    .*.. . ****** ****     *  **** ***.  *   .  . *

40                  YTRYRKKYRSCCAKMYYMAAWKTAKRYMMYCYTYRKKRTRYCMGKTMYAACCGCAYWKKC
42                  YYRYGKKYRCYMRKMCYMRRWGCTKRCMMYYCAYRKKAKRCYMRGWMCGTYTRYTYWGKM
```

```
      * ** ****.   ** **  *  :** ***  :**** .*  *   * .:    :** *

40    KAKAGTRGCCCYCGGKTRKARACRCAGWATRRKCMTAYKCTGMYYWKRRKGKGRKGAYKT
42    KWKTRYRAYTGYYRSKKGGRRRTRMRRTMKGGKTMGGYGYKTMCYWKGRGRKKAKKGTKK
      * *:  *.   *  .*.   *  *     .  * * .*  . * *** *  *  * . *.

40    CMKRKARGGTASRTGRTCYWAARTARMGAYKRYYRYAKTCKWRYGTGRRYAYCCCWMCGA
42    TCKRGWRRRWRSRYRRCYTTRRRYWRMKRYTATTRCWKYYGTACRKRRRCWCAYMWMYRR
        **  *    **  *      *  **  *.   *  *       . **   .  **

40    GKAGMCWRAYKYAATMGARKCAATGGGAACWMSCTRKYRRACRYCMRKKYAGAKKAACTG
42    RKWRCYWARYKCRGGMRWGKMCGKRKRCGMWCCYYGKTRRMYACSAGKKCTRWTGRGYGA
       *    *  **  . *   * ...   .. * .   * **    .  ** :  .  .  .

40    RYWGAKTSYKGTCKCAKAYATKKGTTRYKWYYGARTACGCCMAGRYAKYYY
42    RYWRRKCSYKRYMKMMKWCRYKKTGYGCGWCYRWRYWYRYYCRKRCMKYYY
      ***  * ***   *  *    **      * *  *         *  ****
```

The pairwise alignments of two accessions 40 and 43

```
40                RYTGRTGCMKYYRRWYACCKYAMWRWRYCCCKAGWWMKACCTTCAGTRYYTGRYKTAKRG
43                GCCAAYAMMTYYRRWYGTTKTRMWGWRTATTKMSTTCGTYMCCYGRKGCCWRAYTGWKRT
                     .  . *.******.  *  ** ** .  * .    :     . .      *.  **

40                TATWMRGCRARCTRCYWYYKWMCGATYGMGGTKYRRKRKCGCYRGAKMYKAYWGAWYRAG
43                GWKWMRTTRWATAATYWYCTWCSATYCTMKRKGYRRGRTAAYTRATKACKRCWKCTTRTR
                    .***  *   :  *** .* ..:   *  . *** *...  *.:*  *  * .  *:

40                RRYRSKWWRYMKKCWKKGKGCYYGAKWYKWRYYGTAGGTYWCRTGYTAARKYKRKMAWYC
43                RRYGGGTWRYMKKYWKKAGRTCYKWKWYKWRCTAATATGCWYACACKTTGGYKRTCWAYY
                  *** .  ****** ***.    *  ******  .::.   *   . .::  ***.   *

40                WRYKWASTATRMYYYWKYWRRCGWKCRYRYCYKYMYKRYYMYWRYCGRYYRWWYCAGGKY
43                WGTKATSWWKRMYYCAKYWRRTTWTTACATATGTMTGGYYMYWGYMCRYYRATYMRRRTT
                  *  * :*  .****  *****  *.     .   *   ***** *  ****  *    .

40                MRYTCYSYAWSYMMKMKKYTTARCRRYKRRKAKCGTMYYRYKRMRGGYYWMYTYAYGTTG
43                CGCYMCSYMWSCAMKAKKCYWWRYGRYTGRGWKMRGAYYRYKRMRRTYTAMYGCRYACCR
                        ** **  ** **    *  **. *  *    ********  *  **   *.

40                RWRRGSYMWYMYYKRRRRMCYWRRCTGTYGSGRGAYGMMMWKRYRYGCYGGKRAGCTGTG
43                GWRRKCYMWCATCTGGAACYYWRRAARAYRGKGKGCAMCAAGRYRYKMCRKKGTATKSKR
                   *** .***    .      ****.: :* .   . .*    ****     * :. ...

40                CMRAMCARKRYWTATYYYKYKTTTATKGYTCCAWWYYGTKATGTTRTRTAWCSYARRRYK
43                TCRMMYWRGGCAYRKYYYKYGWYKMKKKYWYYRTATTKYKGYSKCGARWWWYGYRAGRCK
                    * *  *      .*****   . .* *          *. ..  :*  * .*   * *

40                TTWGGKYKAMMAGGYKYRWYCKGGMTRCTGGTYYCCTGRYYKCCCRTATCACTMRSAKYT
43                GKTKRGTGGMMWRAYGYRWYYKRRCCRYCRRYYYTAARACCGYTTGKTAYRYWARGRKCG
                   .      .**  .* **** *    *     ** .:         .::     *. *

40                AYTTCAGWAWRYKAGYTTAMWGYKYKAMWGAWYYGTYRKGCTKYTAYKRRRMTRKATKRC
43                RTCCTRSAMAAYKGRTWYRMTAYKYGRMWACTCTKYTAKRTWKYKMYTRGRCYAKWYGAM
                        .    **.     * .***  **..       *   **. *.* *   *

40                TGRYMTKTCYCTWTCWGMRKCCTWYMYCYTCGTSRWRCRYRMTMTGKAGYRRACCRGKTS
43                CARYCCGCTYAAACTACMRGTTGWYCTYCWYRKGATGTRYRMCMCAGRKTAGRSMAAKGG
                   .**     *.:     **    **       ..    **** * .       .  .* .

40                RACAKRTTYWKATTCYWCGTRYRCYARRKYWYGCRYYWATKYYRYKRKTAWYKYCYTTYK
43                RRYWGAYYYWKRWAATWYRKRCGTTWRRKYTCRMRYYWRKKCCRYKRKWRAYKYTYCCCG
                  *       ***  :. *  .*     ****    **** .*  *****   *** *

40                GAKATWYATAGCTTMCGKGGRKMKKRTGRKKACTGATKSCGRYRCRSKCWAWAAKATTYT
43                RCTMKATRYRKMCCMYTKKCRKMKKRCAGTKRAARRYKSTARYGYRCGMAGWTGKRCYTY
                   .. .         *  *  ****** . .* .:   ** .**  *.   .*:.*

40                CKKACTRATAKTAKYMKYKWRMTKYSMTTGKGTAGKWYAWWTCCTWKYACGAMGGYMYMC
43                SKKGSKRGCTKYTGYMTYKTRMCKTGCKKAKAKMKKWYGTWYMAGTTYCTAGCKRCCCMT
```

```
                  .**...*. :* : **.** ** * . ...*..  ***. *  .  .*. ..      *

40                KRWTKRKYRKWYRMWRTGTTTTKCGGKRKMYSKKTRCCAKTGACKCAKSRGGAYYGKTCT
43                TATCKRKYATTCACARKRWKYCKYRKKRKCCSKKAGYMMKWRRTKTTKGATTGTTRKCTA
                  .   **** .     *.  .  *   ***  ***:    *     * :*.    .   *  :

40                CKKKKWACCKKGYKWWKYCGTYSYKYWYKYAYRWKWRGWWRRWMACYGCKAGKWYAWAWY
43                YKKTKTRYYGKKYKWAKCYKCYGTTTATKYWCGATTRRWWRRAMWMYRYGGRGTCWTGWY
                   **.*     * *** *    *. .   **    . * **** *  *    .      .**

40                RARRRAACGTRSMRACCTGYKTWYCRYRAKKYRTKKKRYYCGCGTACRTRKGMYTYCWKR
43                RMRRGGGTRWACAATTYCTTGKTYYRCGTKKYRWKKTACYYKYRWRYAKRTRATWCMTKG
                  * **  ..    .  :      . * *  :**** **.  *        .*.       *

40                TACKCKTYRKRTRWKWKKMKWYYCKKTTCRCMRYGTYCMYCRGWRTCTMYRAGCRRATYY
43                GGMKAKYCAGRGGTKWKKAGACCYKKKGGRYMRYRWTTATMRAWRYYYMYRGTTGRMGYY
                   . *.*    *   ****      **.  * ***       *.**   ***.   *  **

40                RYCCCGGCTMTCWRYYRGRWSRWGYCTAATKYRMTTCKGCTAGYRKGGMTYGYKWMRTYY
43                RYYMMRRYYMGYTRYYRRRWGGTRYTKRRWKYAACYYTKMKGAYRKRSAWTKTKTARGCT
                  **        *   **** **.   * .   **      .  ...*** .     *  *

40                RGRRYRAGCKKRYCAKCGCCGTRYCAYRMYKKAYMRTMRYRKKRRKKGTKRYKCAKYMKA
43                ASARYRWRMKGRYYRKTAAAKCGCATYRCYKTRCCGYMGYRKKRRKGKWTGYKSMKCCKM
                   . ***    * **  * ...    .:** **.     * *******   . **. *  *

40                GCAGGGRKGRMRRKGCKTGTGKKKKGMYCGAATAGGYYTGKKMYKRSKACYAYCWCYRTT
43                TYRACTAKRRMAGKRYGYRYRKKKKRAYTRRWWRTRYYAAGKMYTRSGCMTMYYWYYRAY
                     .    * **  *       ****  *        **:. ***.** .    * * **:

40                GGGRKTAGARGKSRSYWYCCASYWACAMTKYYKYYRRKGTTMGAGCGMRGAAACCAGGKR
43                KARRKCMTRRTKCASYWYAYGSCACMRMYTYYGCTRRTKGYCKMRYKMGRRRMYYGATKA
                   . **    * *. ****. .*  .  * .**   **.         *        .. *

40                WWKCTYGGAGRYMCGYAYYWTKWRYYWKWTTWGWRMCYCAAAWSCTTGCKYTYTGAGAGG
43                AAKMWYCACARYMMRYGCYWYKTGCYWKACYTSWRMYCAGTGTCAAATAKTYYCKWAGAC
                     *  * ...***  *. ** *   ***    .***  ..:. ..:: .*  *   ...

40                CKRTYTTRAYSWTYGACTYYWCMTTKYRTTGYKGAGYCTRGARWKTKKKTCMGTTKRMMA
43                TKGGYYYGWCGWATATYKCTWAMYCKYRGGRCGRMKCYWARGRWKAKKKCYCKYWKRMMG
                   *  *      .*: .: .  *.*  ***                .***:***      ****.

40                TKKTMCKTCTWRCGYYARWCKWGCCKWYKSYWTKMRRMCATGCTMKTMYRKRKYWGGCCK
43                AKKYMYKGAAWRYRTTRAAMKWAMTGTCKSYWYGCRRMYRWKSYCKCCCAGAKTAKRMAK
                  :** * * .:**         **.     ****   ***     .  *       *     .*

40                WWSRKKTTYMYCAWYKCKRYYKMYAYCYWRRKKMKKRKARKAAGCKTAKWRYRYMRTCRG
43                WWSRGTKYTCCSGTTKAGRYYGMYRYMCAGGKGMKKGGRGKRMAYKCMKTGYRYMAWTGA
                  **** ..     ..  *. *** ** *      * ***     *  . *  *  ****     .

40                RGCAKKMYYWWYGGKYAKSGRYKKTKGYRARRWTTGCAGAATCTGAKWKMTTRGKSRATG
43                GRMRKKCCYAWYKKKYRKSRRYTTKKRYGWRGWYYAATACGAYYARKTKACCATKGGWKK
                      **  * **  ** ** **...* *  * *   ..:...:  . * *     *.  .
```

```
40                WRYRRRWKWYAAAKAAGGACCYMKKKKCKKAWTKYKGGGRKRTTWYYRRWTCYTCKCSYR
43                TRCARAWTTTGGTGTGTAGTYYATGGKSKKCWCKYKRRARGRYGTTCRGTCATKYKYSYR
                   *  * *.  ..: :. ..  * .  *.**.* ***  .* *     *   . . * ***

40                YKRYKKRKTTAAATAGCYYTTWCWKKSRSSSCRYMMMYTYYTGGTKYAAKAYYCKAYTRC
43                YGRTTGGGCCGGRKMCMTCYYTYAKTSGGGGYGTCMMYYYYCAAKKYCWTRCYYKRYCGT
                  * * .     .. .           *.* ...    *** **  ...**. .   * * *

40                TCGYRRRRGMGAKYMMCKCATGKRKWYWRKRCGARAKCCTRYWKMYSYTYRMYYYTTYGK
43                WMSTAAAGKAKWKYMAYGTGCRGRKWYWRKAMRMGRKMYKRYWKMTGTYCGMYYYAGYSK
                    .         ***     .   *******       *   .***** .     ****: *.*

40                CTYCKGYTRYRKTKGWYYYRWWKSGRGATCGAKWWYCTTAKMKTKARKGTGCCATYYAGG
43                TACAKAYKRCGGKKRTCCTGTTTCRRRWWMKRKWWCMKWCKCGAKGAKKAATTRYYYTAR
                   : .*.*.*    .*          .. *       ***   . .*   :*. * :.     **:.

40                TRGTCRCYRTTRYRYCAGGCAYRKKKGRKGTAAMRYYGAYRKTTGYMRWTTYSGYARTKK
43                CRRCYGMTRYYAYGYMRRRMRCRTTKSRKTCTRMRCCRCYRGGGAYMRWYAYSTYMAKKK
                   *      *    * *        *..*.**   : **    .**    .****  :** *   .**

40                MCCCSTAKATARYYYGCWWCCCGTRKGYYYRKACAYYRRTKKAGSTTRYKKCCMMYKRKC
43                MTTTCYGKWGTATTYRTTTMTGAGGKKYYYRKRTRTTAAYGKGASYCRYKKMYCMYGRTM
                  *    . .*  :    *         .  * *****          *..*  ****    ** *.

40                TKKMYKGYMATYYYCKTKRYRMRRWCKGGMGGRTTAKWKRSAMYKKRCAYRKMYCAYKKK
43                WKKMYGRTCRYYCYYKCKRYRMGGATKRKCKKAKWGGWTASRATTKAYMYRGACYWCKKK
                   ****        * * * *****     *        . . *. *    .*    **       ***

40                MMKGTTCTTGSWKAGAAKYGKGWYRYYYYYYMWSYRRYYKKTATYYAGYAKYRYMKYGYY
43                MMTRYGAGYRSAKRRMRKTRKRAYRCYYCCCMWCCGATYKKGRCYYGAYGKYRYMTYRCC
                  **.    .    * *     *  *  ** **    **.     ***    **..*.*****.*

40                CTYYTGRYAARGRTAAGMYYKWGTATGARRRYTCCGTTYYGCWKGGKYKYRYGTGKSMRW
43                AATTKCATMRGAGCGTTCYCTTCCCCCTAGGCWMTAGAYTKTAGRAGYKYRYACRKSMRW
                  .:  .       .  .:  * .    .   :         . :*        . *****.  *****

40                MRARRCYMGYSMKCAWYRTRKMRTRGGKKTRYMWRTWSRTKKKKMKRYRSRGCTCCWCKK
43                MGRGGTTCRYSCKYWACGCAKCGYGKKTGGRYCWRKWSGCKKKTAKRYRGGRYWYSAMKK
                  *        ** *        *         .  ** **.**   ***. ****.      .  **

40                KWRYKTRRYRAYKKMCMKYCRYRYMATTTAKMYYKSRTKGTWAWWKGTTKCGAATTKCKA
43                TTGCGYRGCRRYTKMTMTYYRYRYMMYKWMGMYCKSGCGKGAMWWKRWYKYARRWWTSGR
                  .      *  * *.** *.* *****   .    ** **        ***    * .    ..

40                WKCACCAACGGKRTAYKMRRKMGRKTRKYRWCATGMYKRGGRKWRYYMKYATAKCYYGCY
43                AKYCTYGGTTAGACGYTCRRKMARGYRKTGAMWWRMYKGARRKARYYMKCGYWTMYCTAY
                   * .  ..  .    .*. ****.*  **        *** . ** *****  .   .  *   .*

40                YAGTTTGGTTYTYKKYAWGKWTRYRAKTRCTSCTTGWCYKRRAATRRTGTGAMYGYGCGY
43                YMRYWCRRCAYKYGKCCWCKAWACGWKYGYACTACRWYCGAARWWGGGKCRCACKCRYSY
                  *        :*.* * .* *       *    :. :  *                .      .*
```

```
40              RYRRGKRMGRCTWKCTKRYRYCKYYYYTKGRACCCCKKRKKRGYYYYYYYRYKWRKWMTY
43              RTAGRKAMKRSGWKTAKRYRCMKYYYYYKRGRMMYMKKRKTAKYCCTYYYGCKARGAMWY
                *    * * *. ** :****  ***** *       ****.  *   ***  * *  * *

40              KRWGMKMWWYKRSKKKWRYRKSRGRAKTTKWRAACCSTKRCTKYRWKAMMRRCRKTKTKY
43              KRWRMKMATTKAGGKKWRYRKSRKAWKYAKTGRWYTCYTGMKKYRTKWMMRRYAKCKKKY
                *** ***   * . *********   * :*      . .  .*** * ****  * *.**

40              KCTGCRMRGCCYWKYRWYGTYCYWCYTCATTRKRWKGSYWTTACRKGTTRMKYRYKKTWY
43              KYYTYAMGRYYYTGYRTYKCCACTMCCTRWWGGGAKASYWACGAGKRYYAMKCRCKKYWY
                *     *    *  ** *   .             *.***: .. *    ** * ** **

40              GMYKYTSMSKYTGCACCTKCKMRYRYAYAWKRMGMRYKSWRYMMKYMYARWGGGRCGTSY
43              RMTTYGGCGKCCATCGSKKMGCATGCGCGWKAMACRTKGAACMCKCCYGGTAKKRTAWGY
                 * .* . .*  . . ..*       . .** *. * *.   * *  *.  .  * . .*

40              MKKRRGRRTGKRYWRRKTTTGYTGRCCATCYRKWAYKCKRCGTRCMTYKYYGGTAKTACA
43              AKKGAAGAWTKRYWARGGGAAYGTGMYRWYTGKTTTKTGRTTCGTMYTKYYAACTKKRYT
                 **  .    **** *   :.*          * : *  *     *  ***.. :*.  :

40              ATAGAATYGCAAAAGRTRAMCGTKMTTWTRCTRWMCKWARMMTGRTYKGCCKGRRRCRMY
43              CYRSGWYYRYTTGTARKRWAMKCGMCKTAGAYAACTGWGACMWKGKYKRTMTKAAGTRCC
                .  ..  *  ::.:.*.*      * . : .      *.  *   .**   .     *

40              GKRSTKYWCCRWATACACCCMATYGKARMYYYSRRTAKCGGYMRRCKCTKMCKYRRGTAK
43              KGRSYKCAATGAGYGARTMYMWKYAKTRMYCTSRRWTTASKYAAAYKMYTMTKCGRSYGG
                  ** *  .   . ..    * .*.*:***  *** :... *    *  .* *  *. .

40              ARRRRRACKRACGKYTTCTYYRCYCCRKKRTRSYMSWACKKWYTGTTTTWYRRKKCATMA
43              RRRRGGGYKRMYRKYCYYWCTRTTTARKGGGASYMGWRTKKWYAKCYGCWYGGKKTGCCT
                 ***  . **   **      *   .**    ***.*  ****:     **  **  .  :

40              AAYCKKKKRGGCTCTCKKYCCRYKCATSGTKWYRYRRGCAKYMWYKGKRARMRRWGYYCA
43              WTCYKGKKGAKTGSCYTGCSMGCTYMWSRYGATGYRRTTGKYMWYKKKRMAMRRTRTTAC
                 :  * ** .   .  .  .   .   *      ***  .****** **  ***    ..

40              YCGTGMTGRATRWTKKGCCWRKAWKKCCWGRKATGTKWARAATTAAMMKCATAAYCCRRG
43              TTACAAKKARYAAYKKRTSTRGRWGKTYWRRKTCAKKARRGGGAGGMMKTGYWCYYYART
                  . . .       **  . *  * *  * **: ..*  *.. :..*** .  .*   *

40              AGWKWRCTYTYAMYGAAAAAYCKGGCGGGCAGCTMRRTARATCCAWCACAAGAACTCACM
43              GKAKTRYCYYTMMTAGRMRWYTTATTTTATMKYWCARYRGRYAMMTYGTWGKRMMYTWMM
                .  * *  *   * ..    * ..    .       *     .    .  .        *

40              KRCAGTTYRAWTTRYRAGTMTAGGGGMWGWRKGTAWKCGTARCGYYKGTWACCCTRTRRK
43              GRYTAGCCGGTYYRYGWRGCWWRSRSCTATGTAGGTTTAKRGYSCYKRYTRAYMYGYGRK
                 * :.    .   **        . .  .  .. . . ..   . **    .      **

40              KKTTCKKAYTKRACRRRKWRRYSRATCRYKAACKCTAGAYKRYYTGAKWTAGRRYRCKTG
43              TKCATKKGTWGRWYRRRTWRACSRTCTRTKRRYKTAGARTTATTYRMGTYRRAGCRYKAR
                .* : **.   *  ***.**  **:  * *   * :..  .              * *:

40              CKTAACWYYRGWRTTYAWCRCRKACTTRYRYYMRKKGYAAGWCARTGKKGGCKYKKGATC
```

```
43              TKWRGYTYYGSAGYKYCWYGAAKRAWYGTRYCAGKKRYWRATTRGCSKKRRMKTTKKRKS
                 *  .  ** .   .*.*  . * .    **   ** *  .     .**   * .*  ..

40              AAAYCKWYTTTTAACGMYRYSKGCATRMAYGAKYYYAAMA
43              RTTCYGACYKCAGWYSCCGCGKRYGGAMWCACKYYYTWMW
                 ::      . :.  .    .*  . *  ..****: *
```

The pairwise alignments of two accessions 40 and 44

```
40              YMRRASMMKYYRRWYRGCARYKCGGYWMRMMKYKWRRRRYCCCKCCCCCCRRRYCACWKK
44              TMGAMCCMTYYRRWYGRTTRYKYKKTWMRMMKCKWGAGRTATTKTTYYYMRAGTYRTWKT
                 *   . *.******   :***    ****** **   * .  *      *      **.

40              YYMTGYKKYYCWKMMKAYTYKYRTWKRKKYMWYRYMKRYRKKRRKAKCCGYMYYKAGMRR
44              YYMCTYKKYYYWKAAKGYCCTYAWAKGGKYMWYGYMGGTGTKGRGWKYYRCATCTWRAGR
                ***  ***** **  *.*  .*   *  ***** **    .* *  *       .    *

40              ARGRYYWRYYWWCYRMYMKRYYRYMWRTYYKKAKYCRAWAMKYKRYYSYKKKWCTTYMYY
44              GACATYWRYYWWYYRMYMKRYYRYMWRWCYKKRGYYAMAMCKTGGYTCTTKKTMWWCMYT
                .    ******* **************  ***  *      *   * . .**     **

40              YWMCTCCTKCYKYYRWKAGGGYYGSYYRRRAATGRAKTYRYYYYKKMGGKMKGTTRYKYY
44              TAMMYYYWGYCKTTAAKRRRTTYRGYYGRGGTCAGRGGYGYCCYKKMKRKMGAKAGCKCC
                  *        *    *     * .** * .: .    * *  ****  ** ..:  *

40              YKYYRKYYRGGTWYRRMYYYKRRRKMYMRYTCMWCSSYWWYYYWRTCYYYTAYRSRTRKK
44              CKYCGKTYRRRGTYARMCTTGGRRKMYMRYWSMWYSSYWWYYYARGYTYYYWYRSRKGKK
                 **  * **    * **     ******** .** ******** *   **  ****. **

40              YTGYYRRYKGWATWMRTRCAYYMKYKYGYKYWCYWWYKAYKRKYYGRYKMWRRWRSTAMT
44              YYSYCRRYKCWWKWMRCATCYCCGYGTKYKCASCATYKRCGRGYYKRCTCWGGAASWWMY
                * .* **** * .***   .*   *   **  .   **   * ** * . *    *  *

40              CAYRWRYMYSTGRCAKGGRWWSTGGWGSYGGMYCYAGYMCYKYMMRCMYYRGYKCCWRRW
44              YMYRWRCAYSWSRYGKRKAWWSWAAAKSCRRMYYTWATCYYKYCAGYCYYRRYKTGTART
                  ****  ** .* .*   *** ..  *   **   .   ***     *** **    *

40              RAYKRKKYAWYWRCRYKRYYWWGGGTARSRASAYYMKYCWRGYYRTMMYSTARTATAGGC
44              RCTGGTKTMWYWRTRYGACTAACTTCTRSGGGWTYCKTMTRKYYAACCCCYRRWTCCKAA
                *.   .*  **** **          :** ..  * *   * ** :   .  * : . ..

40              GGTCCTCCRKCYCRRCAWWYYAGGCTAGKARKTTKRRRMRSAGGCWKYRGSCYCRYWKCC
44              AAYYYYYMGKYTTRRYRTATTWKATYWKKGRKKCKGRRMGGRAAYWKYGRGTYSRCAKYY
                ..       *   **        .    *.**. * *** . .. ***  . *.*  *

40              CTCGKWMCGCACTTACCYMWTCRRMRYYGKKYRRYSRYKCKGKCTGCWTCCCGCTSRRYT
44              YYTAKWCYAAWYCCRTTTCACYGRMRYYAGGTGACGACGYTRKTCRYWYATTKYYSRRYY
                   .**  ..             *****.      .    . *    * .     ****

40              CCTTTMGKATKGAYCGTTCRAGWAWRYRGAYMKKYCTKYRGKRTCTYGRRACCAMTYYKG
44              YMYYWMRKTYKTTYTTYYYGRSAMAATARTTMKKTAAKCAKTAWMCYARRWMYRMACTKR
                     * *: * :*       .       : *** .:*   .    *.**    *:  *

40              MRYGYKYCAKRYRGGCTKTRKTGAWYYGTYCRKTKAAKTRYKYRRMRAATKKKKYYYTKR
44              MRYRYKYSMKAYRRRTCTYGKCACTCTKYYAAKWKWRGWGCKYRRCAGGAKGKKCCYWGA
                *** ***. * **    .  * ..     *. * *      ****  ..:* **  *

40              CAGGTGYKRYWAYCRSYKYWKYKGGTAYKRWKWSAAGWRRYRWMACYYYWWCGCKGCCTA
44              MMKSCAYKGCARCTAGTTYAKYGAACWCKGATTSTWRWRRYRWMWMYYTWWTRYKKTGAW
```

```
                   . .**       . .* ** ..   *  . *:  *******  ** **   *   :

40     TYYATCTYCRRRWYRSRRRMYRYGTRSMRKRYYYYRYRTKKYRKMCKKGCGSTCGSKYYR
44     CTYWCACYYGAAWYRCRRGMYRYRWACAAGRYYYYRYGCKKYRGMYKGAYACGMRCKYYG
          *  . *    ***.** ****    .   *******  **** * * . ..   .***

40     KRYAGKMYKRYGRATRYCMMSYMKMCYMGYTGGTAAKKRAKCAKRGTTTYCGGTTARWRK
44     KRCGAKMYKRYSGTKRCYMMGCAKMYYCRCKKKWWTKKGRKMWKRRWCYCTACWKWRWRG
       ** ..******. :.*  **.  ** *    .     :**  *  **       .  . ***

40     ARAMMRTGYTCRWWKWKCTYWTKCGAWYYTGRTGTGRWKYKGYRWCWGCRCMYWCTTTRY
44     RRRMMRGRCGTGTTKWKYGYWYGARRACCKARYRYAGTKTGRYGAYWKMRMACWTYYYGY
        * ***        ***  **  .     ..*   .  *   *   *  *   *     *

40     KRRMKCGRKAWAWCGWGWAYKKTYYRAGAAKATWRCTTYAMTYYTTKKCGYRWRKTAYAA
44     GAGCKAAGKRTWATATRAWCKTGYCGGRGGTWKWGMYYCRAWYYYWKGARCRWRKKGCGR
           *.. *      .     *. *  . ... .*         **  * .  ****.. .

40     GRAAYGYTAAMKYRTYYAGGAWMARYCRAYTRCCTGATTGRCACGTYKARRTGYATCYYY
44     KRGGYRCCGCCKYGYCYCCKWACRRYYGWCGGYYYRWKKRAMWYTAYKWRRGKYWYYCTC
        *..*   .. **   *.      **             ..      :** **   *

40     RYAYKSCMCTRRKGMGTYGMTRKRTATYMKAMGGTKYRRGTTCAYCRTYKACYTATKYYK
44     ACMYKSMAGCGAKRMAWCRCKRGRCGYCMGGCAAKGYRRRAAMWCYAYCKTSYKWAKCYK
          ***       * *.    .* *  .  *  .  ... *** ::       *:.*. :* **

40     GGWAKRRCTGAKKWKYRSTRKKGAYTTTGKKRMGGTARYGRYWTKAACCKAYKRYYCAWK
44     AKTGKGGYCRGKKWKYRSCAKKTGTACCAKKRCRRWRRYRRTTKKRRYYKRTTATTMGTK
       .  .*     .*******  ** . :  .***      ** *  .*     *   .    . *

40     YKWTAACGRRYYWCGAGWATRWWCYYMYMSKAAYYWYMYYGARWTYYTCRAAAASCWSCY
44     YGTYRGTRAGCYWYRRRATGRWWYYYCTASTGMYCWYCYYRWRACCTYYRCGCGSAWCYT
       *     .     **      : *** **   *.. * ** **  *       *....*.*.

40     TRAYKRCYTYYACAGATRYGCMRKKKAGYAACKWCYRCWARTCYKKARYCKYTCKWWRCA
44     CGTYKRTTCCYMMRRRKACRYAGKKKWKYRGYGWYYRTWRGCSYKKRRTMKTYMKAAGMM
          :***     *      .       ***  * .  * ** *    .*** *  *    *

40     TKYYMAAAKMYKRSYYRAKWWYYGYRACRCMKKTCYRARRRRYSSATTTTGACGAYYYYK
44     WKYCCCGGKACGGSCYRMGAACCATRWYAYCKKYYYRTRRRAYCGTCACCAGTACYYYYK
        **   ...*    * **       . *      **  **:*** *..: :   .. ..*****

40     WTYKKCYTTCTKYTCKACCATCTATTYYYYTAKKYCRYYKRKRKYKRYKAYCYYYTRYRK
44     WYYKKSYWAAAKCKTKGASMYYYMYYYYYYYTKKYMRYYKGKRKYKRTKRYMYYYYGCGG
       * ***.* :.:* . *...        **** :*** **** ****** * * ***

40     WKWWKRYKWGRTTYWCMTKRWGMKYWCKGYRSGKTKKWRYRKSRGKSRKACAKKTKTAKT
44     AKAATRYKARGWCYWMMCKRWRMKTTGKSCAGAKYKKWRYRKSRKKSAKWMRKKYGGTKW
        *  .***      ** * *** **    *.   ..* ********* ** *    **    :*

40     KWAGAYAMAYCYRYASTKRWYRWKGKYARGAMMRRRYRKCYKCTYRKKRMMMTACWKKRK
44     KTTTRYWCWYMTACRCYTGWYRTKRGCGGAWMMRRAYRKMYKYYYAKKGCCMWRYTGKRK
       * :   *   *      . . *** *    . . **** *** **  * **    *      ***
```

```
40              WAACGWARRAGRWRKKCAWGTAMYYYYYCTMWGYYSRRYRACWTYTCYGYRRRCARCTYY
44              TGRYRWGGARKAWRGKYWTRCGMYYYYYTAMWRYYSRRYRGYAYTCYTRYRRRSGRACYC
                 .   *.     ** *     .****** :** *******.       ****..*. *

40              RTAYYMRWGYKYRYMAYSWRRYYYWGTGSYYCGYYTKYMSRSRTTAKWRRRKRYYCYCCS
44              GGTYYAGTAYKYRYMRYSWRRTYYWKYASYCTRCTKKYMCRSRYCRTTRRRKRYCYYYYC
                  :**   .****** ***** ***   .**     .***.***    . ******  *  .

40              TYCCARMKTGRAGKTYYKYMRYSMRRTMYGKRRRYMKRGAGCWTRTRTCGCGMRRYKKKK
44              WTTTCRMKCAARRGYCYKYMRYSMRGAMCRTRGGTCKGATATAKGYRKMSMRMRRYKKKK
                    .*** .      ********* :*  .*    * .:.  .  *. .  ********

40              MYTMRYACYCWYGACWGTYGYYRRKYMYYRCGGRYKTKCTCYRGKSYYWKKACATGYSGR
44              MTCCGYRMCYWTACMTRWYSTTGGGCCYTGYRRGYGYGYKYCARKSYYAKKGMRYRYSKG
                *    *    * ..    *.       *      *     .     **** **.    **

40              SSSCRYMKKAMMYKTGRYMGTKYYRTGGYGAYTCTAGTTAGTGRYRRAAACTTAYMYCAM
44              GGGYGTCKKRMMYKYKRTAKYGYYRCAACTCYYYCGCWKRRCRACAAGWTTYYRYMYYMM
                ...     ** ****  *     *** ..  .*    .  .         . :    ***  *

40              YRGYYARRRRATMMRMRKACAKCKMKKRKWARARWRYYRMMTGYCTCATCAKCTCKYWKR
44              YASTYWAAAGMYAMRMAKTTCKMKAKKRKWTGRGTRYYAMMKRYSWAMWMWKYYMKTWKA
                * . *        *** *: .* * *****:    *** **. *. .     *    * **

40              RRTKYTCGYYTYWTYKMRKYYSYRKRGAYRKCAARGRWKMYYYYRCTRYTWWKTMCGKTK
44              RGCKTWMRYYKYWCYKMRKYTGTRKGKTCGKMRCGRRAKMYYYCGYGGYAWWKKAAKKYT
                *  *     **.** ****** . **  :  *   .  * *****      *:***. . * .

40              KGYRMCYRTCGGKWKRRWGCAWKKWWKSRKKRYGKAYYMKYYYGMKYCCMGGCGKCCTWY
44              KAYRMSCGAYRKKTTAAWAMWWKGTTTCGKKRYKKRYYCKCCTRCKYTTMATTKGYYKWT
                *.***.  :   *  .  *.  **    .. **** * ** *      **   *.      .*

40              KKMYRMYYRRCGRYCRGTYYGKYRYRMGGTGKCTYRAKGCCRWGWRTSMGRRSYRTGGAC
44              KKCYACTYGRYRGYTRKCYCKKYGYRMCACAKTKTARKSSMAWKARGGAKRRSYRWRRRY
                ** *    * *    * *   *   ** *** . .* .    *..   *   *  .   *****

40              AKAWWRKRRYKAKKGTTCACGYGYRKATGTYRYWGKYWRYGCMRYYKYCGYYGTAMWMTA
44              WGGTTAKRRYKRKTTAAYMYKYRTRKCCRKCGTWRKYTACRMMRYYGYTCCCAAWATCWR
                  .    ***** *. ::     *   **.   .    * **      **** *      .:

40              WRRTRSKKYGWRYCGAKYKCRGGAAKAKRKWSTAGAKCTRYKGGRRKKRKCKTCRKMKKT
44              ARGKAGKKYAAGYARRKCGMGRAGGTTTAKWSYRKRKMKRTTKCRRKKAKSTWYGGAKTW
                 * . .***.  *.  *       ....:. ***     * .* .  **** *..      *.

40              AGCCCCGYYRYKTYKKTKRYYGYKCKYCWCYAAYKSRCRYRCGMKWSKRCGWWKWAKSCA
44              RRYYMYKYYRTGYCTKWKGYYACKTKCYWSYWGTGCAMRYGTACKTGTGGATTTACKCAC
                       ***      .* * **. * *   *.* .   .  **   . * ..   .   . .*...

40              TGGTKTCKAWYRYRTKSMCMGARAYYKAACCTCYCKGYRGKRKAAYYCSWGWTYYTTARM
44              CKRYKWSGMATGYRWKSMMMRMRRYYKGGMAWYTYKRYRKGAKRMYYSSTRTYTTKYRRM
                    * .     ** *** *   * ***.. .     * **    *   **.*        .  **
```

```
40          YAWYYSYTRMTRYRACKKYYMKKGRCCAWKKWKAWKCKYKKWMMTWCRWWRRWYATMGTA
44          YTACTSYKRAYACGRYGKYTCKKRRYSWTTTTKRWKYKYTKWMMKTTAWWRRWTTCMAKG
            *:   **.*        **  ** * .  .. * ** **.****.   ***** : *...

40          ASWAYGWACRACTGCTRMAATTRKTATTTRKMCTGRCCWMYRKRRGATGCKWRKGYGAKG
44          RSWMYRARYRCYYRYYGMRRKWRKYCCWAGKCGCTRYYTCCATARTGKTTKWRKRCRRKK
             ** *    *.      *  . ** .  : *    *      . * ..  ****    *

40          AATTGTAYYYYYTAYAYMRMRRCKGCYSYRKRRKKRYMKAKYKKKYMMMMKYWRKGTCRR
44          CTGYAWMCYTTYWRYGYCGMGRAGATCSCGKRRKTGYCKMKYKKKCCMMMKYAAKKWYRR
            .:  .   *  *  *.*  * *. .  *  ****. * * *****  *****  *   **

40          RMRRAARGCGAKMMKYMYWKGAAWAMTTWRAKAAKYKKKGGKCKGAATARGWYKKTCKGA
44          RMAGTGGRYRRGAAGYMYWTKRWTGMGATRMKWWGCTKKRKTTKRRWWRGTWYKKATKTC
            **  :.         ****.    .* : * *    .**  . *       ****: * .

40          AAGAATTCGTAKRSTAAKYKTATSTRKGKMRSRCYKYSYWYKCASKTAYYWAGCTYATYY
44          GCKMMWCTKAGGRSCCTKYKWMYGCRKTKMRCATYKYSYWYKTGSGCGYYWRSMAYCKYY
            ..       :. ** .:***   . ** ***.  ******** .*  .*** . :*..**

40          TTTAKYRCGKYKMKGRYGYAGGKGRKWGCRAGGMTTGWWKWYMCTKTGGGAGRYTGCAAT
44          CCCTGCAMKKYGMKKRYKYCKATARGTRYGGATACGAWWKTCAAYKWKCACARYYRSWGC
               :     ** ** ** *. ...*     ..    .***   . *   ...**  . .

40          KATARRMKCATRCCGYWMTRRTGWKYGACKSWAMRGTRGKCTTAYMRKKYTAYKMMYYSC
44          KCYWRRCGAGAGYYRYWCGRRYAWKTTTTKGAWCRACATTMWWWYARKKYKRCKMAYYSY
            *.  **  ..:    **  ** .**  : *.   *.   .    * ****.  ** ***

40          CSKCACTYACTRCTCKKRTYCYCGATCWGTTGYGYCAGCCRWRSTAACAKCCMWKTYYCY
44          SGKYTAGYMYYRTAATKRCCYTMRGYYTAKCATACYGRYTATAGWMRYMKMTMWKYTYTY
            ..* :. *   * :..**      .   .. . .  .      .     *  ***  * *

40          ACCCRGWCMCTAKMKCCCCKYCCTCYCRRYRGATTWRYWGCRTYWGATTKRTRTYGMKTT
44          WMYTRRWSMMGWGCGYAATGYTTGACYGRYRAWCCAATTAGACCAKTGCKRGRGCRMKCY
                * *.*       ..  *   .   ***.       .      :  ** *   **

40          MTCWKAGTAWCYGMGGATTCYKGGTSYTYKWTKTAATYTYRGKARTKKKRRMKMGGGCGM
44          MATTKRCCGTMYAMRRGAGAYKKACSYAYKWYKYMGGCYYAAKCRAKKKRRMKCARKTRA
            *:  *   .  *.*  .: .** . **:*** *  .   * .*.*:******* .

40          MMYKCTAKKGTTAGCCCKWRYARWCKKKGGYTRKYSYWRTKKMCKTGWGTYRMKTMYYAM
44          MMYKYKTKKKYYCATTYKTGTRAAMGKKKRCYRKYSYWRYKGCYKARTAACRCKWCCYGC
            **** .:**   ..   *        **    ******* *   *:  .: * *   *.

40          MYTRKRCGCAKKYWWSRAKKRKTTYWYRWCYKWYARRKKTTAMYRTTRGYTYGRYGMYTT
44          CCARKRYRMTKTTWWSRGGTRKKYTATGTYTKWCTGRKGYYGCCGYYGRYKCAGTKMYYY
              :***   :*. ****. .**.        ** : **   .       *. .   **

40          AYKACAKRCTKGCAKYTWYWATTTKRGAACRRSYTGYAGRMGCATAMRAAKGGATTKRGA
44          GTKGTGKRTCGRTGKYAATTTKYGKRRTCYRGGYYKYCRRMKMMCGMRCTKRRRCCKRKT
            . *. .**     .**:   :.  ** :. * .*  *. **    .**.:*     ** :

40          TGCCTYGGTAAWTMCARWACYGTTKYGATWARWTYWAMTAAARKKMWTAAWMTTRYRYMK
```

```
44          CRYMCCATYRMWAAYGATCTCRYYKYRRWWRRWKYWGMYTRWATGCTWMRWMCWRCRYCK
                  .    *:  .  .     **   * **.**.* :   .      **  * ** *

40          KKTGYRASRRRGMRTRYWMRWTTTTCKAKRTAGKAGTGTGGAGMYRYRYSRRTKCAATAY
44          KKCKYGRGGARACGKATTAGTYYCWTKCKRAGAKWSYKYRRCKMYRYRYSRRWKMTCCGY
            **  *  .  *.  .           *.**:..* .     . ********* * :. .*

40          CRAYCGMRCRKKCCTGKMCCGYYRRGRKSMKKTKCGARGYATGARRKRYSWTMMKAAKGA
44          MRWYMKAAGAKKMACRTMTYKYCGGRGGGCKKWGYATATTRCTGAGKGCGTAMCKWCKRM
             * *      ** .  .*   *      . **   .:      .  *  . :* * .*

40          YGAYCTCTYYKKYCCRKKGYMYRGCRRYKSYAMKKKYRWAMYMACKWYTKCAKKCAWKAC
44          TAGYSYYWCTKKYTARGKKCMYGRYGATKSYRMKGKYRWRMYAMTKWYCKYRKKTRWKGM
             ..*.     *** .* *  **      *** ** **** **   *** *  **  **.

40          CTTCRKTYGAAAYTYKGYKKRMAGRTTWKKGGRWAAWRRMYRKSAYYYRKKCCCMATCWT
44          MKGGGKYCRGGTCWCKAYGGRCGARYYWKTRARWMRTARMYRKSCYYYGKGYMMMGYYAG
             .   *   ..:   *.*  *  ..*  **. .**    ******.*** *    *.

40          GAKRCRMYKSWWGSWTCATKMMRWYTRYWWCAGCGGKGCAMYGMARTRGAMYATMGGRTR
44          RWKAMRMYKGWWKSAAYWAKACAWTYATWWMRRTRRTTMWATKMCRWGAGMCMAMKAGCG
              *  ****.** * :  :*   *    **      .      *.*  ..*  :* .

40          RYYRATCCYKGYCTRKWGRYKYYRATGAWWGAGCATWYCTCCGTMRTRCACARAAKACCY
44          ATTGGCTTYKTTTKGKTARTKYYRCAATTAATTYRYACACTTRCAAYRYRTWACTKGYYY
                .   **   . * .* ****.:.:  .:      .        *     .:*.  *

40          YTRKWMRAGCYTCYKRYACAKRTWWGGTKKRYCAGRKCTWTARKKGMKWKMGKKTCWKYK
44          YCRGWMRRSYYYYTKRYRYRKRAWARKCKKGCATAAKTYWGRRKKRMKWKCRKKYMAKYK
            * * *** . *   ***   **:*    **  .:. *  *  *** ****  **   ***

40          YTYATKTKAKGCGWYWSKCYYTKWMMSWTRYRRRCCMCKAAGTCRATAAGRWWGRRYRAC
44          CWYRKKWKMKKYAWYWGKYCYKKWMASWCATRARTYATGGTRYGRGAGRAGATAGRTRMT
              * .* * *  .***.*  *.*** **   * *     .:   *.:. .   . * *

40          CRAAACWTGRTYWRKRYWYTCCGGMTCTGACGKKATTGTGMMYYRTGATRTCAAATRYYY
44          GGGGGYWCTRKYWRKATTYWYYRRMATYRRSKGKTCGRYRMMYYAYKMYAWYCWGCRCCT
              ... *  *.****   *     *:    .  *:     ****        . . *

40          SAGKCAKTRATMGGCRTKKGMMCATACAAATTYYYTCCCCGRKGCCKKTCCARAAKCYAY
44          CGKKTMKCRWKMKKMAYKKKMAYRKMYMRMYKTTYCTATGAGTKTTGKYYYWRRRKTTGT
            .. *  * * .*     ** *   .      .  *  .  . .    *    *  *  .

40          RRATKKTYMCCAGGRRAKTTTCRYKRYMRKYKKGRWKYYYMYKKCMCCGAAYRMGYGACT
44          AAGYKKCCATAGARGATKCYATRYGGCMGTYKKCRWTYYYMYKKAAYTAWRYAMRCKRYY
              . **    ...   :*  : **   * .*** **.*******.   .  * *

40          TKSYRYKAYTCATARYCCWMKRTTCAKWKRACCYSAMTGGARKKRCWKGRYCYGGRTKCA
44          WKGYRYKGYCMRYMGYYYACKRKCTGGWTAWTSTSRAGTRRGGKAYWGRRYMCTRGCKMR
             *.****.*      *    **.  . *.   . *        *  *  **      *

40          YCRGAYKMCMTCKYTYWYKTCTRKWAGAAKKGRYKKYRRMYWRSRAYYKKYRYYYKCTGG
44          CYGTWCKMYMYMKYWYACKGTCAKARRMRKKRGYKKTGRCCARSGRCYGKYRYCTKYWRR
```

```
                  ** *  ** *  *    *     **  ***  *   **   * ****  *

40          MKRATRYYRYRACCKYKTMCKYACRKTKCTRWRSKTWCRKKKWGKMWYARTAAKCYGGAK
44          MKRCAGYYGYGRYMKTGCMAKCGTGKCKTAAWGGKYWTAKKKAKTMWYTGWMGKTYTRRT
            ***.: ** *    *   *.* .  * * : * .* *  ***  .***:   .* *   .

40          GKTKAYRYKKCKKRSKAWAGYKTWRKAARGTGCTWMGGMRSRRRWSCGRCKAYWTYYGWY
44          AKAKCYRYTKYKKRGTMWRKYTCTRKCGGRAAYYTMARARSRRAAGMTGMKWYWKYYKTY
            .*:*.***.* ***.. *  *.  **..  :.   *.  ****  .    * **.**  *

40          KRKGGKCCWYMRMRRWKWYMAAAKAAGGMKAKRKACYRKYTAATRKGRCTCRYRCYYRAM
44          KRKRRGYMACMAMRAWTTTMGGTGTGTAMGTGAKGTYRKTCTGARGARYYTRCRMYYRRM
            ***       * ** *.  *..: :. .* :  *. ***  :.:* .*   * * *** *

40          WGGRCYYSYGRYKRKWYYYCCKTWRKRKKYTCCGTGKYRWMRRYSYGSGRYYYRMATMYM
44          TARAAYYSYKRYGRTTTCTGYGGWGKGKKYCTTAYSKCGAAAATSCASKRYYYGCRKMYM
             .  .**** ** *.        * * ***   . .*       * .* ****   .***

40          YKGGTRYYTYCYRKSRKMCYMRYKATCCCRSKYYAGTTYKGTGWAMMCARYRKMYRYKYY
44          YKTACAYYACAYAKSGKMMCMRTTMYTMGAGKCYMKWCYKTCTAWMMYWRYAGCTGCKYY
            ** .  **: .* ** **  ** .      .* *    **     **  **      ***

40          WKYRGMRRYCCATGYKSAKKCCGGKCRYTMAWWWSRKTTATTAYSRYRYKYMRYSAMRWK
44          WKYARCRRCTTCCAYKGRKKYYKRTYACKCGWATCGKCCGCCTYGGCRCKCCGTCGCGTG
            ***   **   . .**. **    .   . .*  . *  .  :*.  * *    ..

40          CMTMTAKWYARRYASKKGTRRRTAGKGRRRKRRYWGKKMWRYWGWYWCTAGYRKRKYWKK
44          MCWAYTKWTGRRYGGKGCAAGAWGRGRAAAKGRCWTKKCTRYWKWTAAGGRYATRGTWKK
                 :** .***..*  :    .      * * * **  *** *  . . * .*  ***

40          WYYMGGGGAYRGMCKKKKYWYMGKKYYYYWWRSMYKAYGYGTKYGYTGGKRTYKRYRKAA
44          WYYMRRKSRTARAMTKKKTAYMKKKCCYYWWRSMYKCTATAWKYAYCARKGGYKRYRKMT
            ****   .      .***  ** **  *********. . . **.* . *  ****** :

40          WYCAYYYYTYARYRKKCGAATCRRARSKYYCKKMRAYKCWCATWKKAMSCRWYWGRSWGCG
44          WTYTYCTWYGRCAKKTARWYYRGRRCKYCGKKMRGYKMWMRYWGKWMSYRWYAKRGAATA
            *  :*   *.*  ** .    *  *.**  ****.** *   * * ** ***  *. . .

40          RMTATCATWCTCTKYSMGRRRKMCKGTKKTRYYYYAYKTCMGGTGGYGGYKRAGMKTKGC
44          RMCTCTGCTACMCKTGMRGGAKMYTAKKKYRYCYTGTKWYCKRCRRCRRCGACAMKYKRY
            ** :  .  .   * .*    ** ...** ** * . *              ..** *

40          KYRKWWTCRRMWMRYYGTTTSCGGTKMYSMKTTTKTTGAKCMCAYKMWRGCTGGKKCAKT
44          TTRTTWCYAACACRYYRWKYCYRKKGCCCATGCGKCCTTGYMMMYKATGKAYRRGKTTGY
            . *. *       ***  . .   .   . .   *   :  *  **    .    * :

40          SRYYYKCCYCYATYYWKKKRRRMWKYAKCMKGKYWWKKWRWCTWRKMTSCGYGYYRYRSW
44          GAYTTKMYCYTRYYCWKKTRRRCTKCRKYAKTKYWWKGWRWYGWRGMCGTKYRCTGYRSW
            . *  *       * ***.***  *  *  * ***** ***  ** * .  *    ****

40          KARKKYMKAAARCYWYRWCYKKYWYYWRYYGTKGWRGTTAATTAATRARTMGMKKKMCMA
44          KRGTKTMKGGTRGYWYRAMCKKCWCTWGCCRCTCAGRKYTWCYTWARRGWASCKKKMTMT
            *  .* **..:* ****   ** *  *     .    . :   : :*    . **** *:
```

```
40              GRYKGCCKYRYMKKWRTAKRYKGTAACAKKRSRYRKGYGTTYKTCAATTGKWKKRMCKAY
44              TGYKAYTTYRYMKKWRWMGRCKSCGTMCKKGCGYRKKYRWYTKYYWRWWRKWKKGCSGRC
                  **.  .********   * *. .: .** . *** *    *       ****  .

40              WYGAATYMKYACKKTCGGYRYYKRKRAKMRYKGTGGCYAKKYKKRKKAAC
44              ATTWMYYMKYCYKKKYKRCATCGAKRWTCGTKSGAATCRKTCKKRKKWCY
                      ****. **.         ** .   *. ..   *. ***** .
```

The pairwise alignments of two accessions 40 and 45

```
40                  WRRKAKTYMYYKRRRRRCCYWRKWRRYCYRCMMMYMKRWRKYRWTRRYKTACAYKMRTTY
45                  WGRGWKYCATCTRRGRRYYYWRGWRRYTTAMMMMYMKRTGGYGTKAGYTWWYWYTMRWYT
                    * *  *     .** **  *** ****    *******   *  .  *.    *.**

40                  KWKWSKMYYRYYWKYKRYRYMYRYKKWRYKKGYKMGGRKCMGCTMMYAWKGYTMRRCRKR
45                  GAKTSKAYYRYYWKYKRCGYAYRYKKTGYKKKYKMKRAKMMAGKCMYCWGACAMAAMAKG
                      * ** **********  * *****  *** ***   * *. . **.* . :*    *

40                  YYYYKYGMMYYTTTTKYYWYWMRKYMGKGMCYYRSWYYWAYRCYGKWKACAGGGCKKCAC
45                  CCCCKYRAAYYWKCKKYYWTWARKYMKTRCMTTRSWYYWTTAYYAKWKWYWAARYKGATY
                        **   **  . .**** * **** .     ******:   *.***   ..  * .:

40                  WKYGWRCWMRKKKRMMMRSWRMMRCYWMYTRRAGKWATTRGRRWRYYMMMRRYMYCAKAY
45                  WKTAARYWMAKKKRMMMRSWRMAGYYWCYKGRMAKWGYCGAARTRYYMMMRRYMYYWKMY
                    ** . * ** ************   ** *. * .**.   . * ***********  * *

40                  YGCYRWRGCAGTRRRRKGATTRTYKYWKYMRRYMWYKRSMMGCTGRRCMRYGYWYGKMRY
45                  YRMYRWRSMRKCGRGRKAGCAAYCKYWGTMRACAATGGSMAAAKARAMCGCTCAYRTCRY
                    *  ****.     * **.. :   ***  **       ** ....*        * . **

40                  MRKWYWTYWGSKCYYKRYYYRYWCCRMYRTRWRYRYYCRSWKYKWYYRWMRYWKGKYATKW
45                  MAGTYWWCAKSGYCCGGYCRYTTYAAYRKRWRYACCMAGTKYKWCYRWMRYTKRKCWKKT
                    *   **    *      * **     **.****     . **** ****** * *  .*

40                  RYKRKKYRYMRKYYRRWYCAATGWYRMMRSRKRYYRRKMGMRGASYWWYWTWYRWAGGYM
45                  ATGRGTTGTMGTCCAAWYYWRYSWYGMMGSRKACTAAKMKCRRWGYWWYWAWYRARRRYM
                       * .   * .    **    .** ** ***     **  *  .*****:***    **

40                  RYYWYRWYGGCYGYTMGYRAKRTWAMYKRYRRSWTWMYCRTGTTYYSACWWRCAAKGRTR
45                  RYYWYRAYKRYTTYYCKCGRKRGAMCCKGTRASTWWACTRKAGCTYCCYWTRSTRKAGGG
                    ****** *     *      **     *  * *  *   *..   *.. * *.: *.

40                  WRWCGCKYRSTATGKWCTAKCCYMYRMAGTKYCTCARMKGMATTKAWAMKCKMKYYYKAT
45                  AGAYKMKCRSWMYRKWYYRKTYYCTAATAAKYSWTTGCKKCGGCKRTWMKTKMKTYCTWK
                          * **    **   *  *    :.:**.  :  *  .  *   ** *** * . .

40                  WCTTTYCMRTYTGTRMRAYYAGGTTACTKATCCGYTAGYWTYCSGRTYYRYTTCAARTKS
45                  WYWWWTYAGYYCTYRMGGYYCASYWWYYKWGAARCWTRCWCTYGCRAYCGTCWYRGRCKS
                    *         *   ** .**...     *  ..   :  *   . *:*       .* **

40                  CWYACAYTYRCYAAARGGKRCTYAGCAAGRYARTTGGRRGYTYARATTTYRYYATKSYRA
45                  YWYMACCYCATCTGCGRRKGYKYRRMWMRRYWAWARARRTCKTWACCGGCATCMYKSCAG
                     ** ..      :..   *  .*      **   : .**  .   .         **  .

40                  GKKMYKGCMCCGTRAYMKCGMGKYRYGWMKYRRMRRGKCATYRMGTGTTKRCMTKWCCCW
45                  AKKACKAYMYARWRCCMGSRCAGCGYATMGYRRCRAKGYRYYAMSKKWKKAYCYGTGATT
                    .**  *. * .  *. * .  .   *. * *** *      * *..  .*       .

40                  GKCKGSGYMRTAAGYATGKRYRYTCCCGAYARGTCYKYYCKKGMYRRWKGGYAYYMYKKG
45                  TKMKRCRYMRKCGATWCKKRYRYAGYMTGYMRRCTCGCCMKKCACARWTRRYWYYMYKKR
```

```
               * * . ***....    *****:     .* *        **     **.  * ******

40             CGYYKCYMGYRRGMGGGAYGGWYYCWRYKAKKCYATARRTKGGGWMRTAKSAMRYKKCCT
45             TAYYKTCMRCGRACTKRRYRRWCYYWRYKCKKYYRYMGGATRCCACRCGGSRARYKKYYY
                .***  *    *.      *  * * ****.** *      :.      * . *  ****

40             YRCWKKYGCGGGYKCYAYSKKMCMTTCKKKSRKCKMAKKKGAGWKYRGYYSGRKMYYKCG
45             CAYWGGYRYRRRYKYCWCCKKMYMYWMKKKCRKMGCTKKKRMKAKTGRCYSRRTMYTKTK
                 *   *     **     .*** *   ***.**    :***     *     ** *.** *

40             KYYYYGAACKTGMWGYWYSYTWTYTCRAATAYKWRRCTTSRTYACCAYRYYGTYYCCYWK
45             KYYYYRRMYKWACARYWYSYCWYTCYRTRCCYKWRRYKWSRYCMYSGYRYYAWYYYAYWK
               *****    * .   ***** *    *:  .***** . **    ..****. ** .***

40             KYRYGMAMSWWYATGYCYYYWGRYYYSYGRWRCAGTMMATWKKGAGCCTYMMYKWSSRCM
45             KYRYCCMMSWATGYRYMTYCWTGYYYSYKRWATRSWCCTGWKKRWKATGYMMYTWCGAMC
               ****   ***   .  *  * *  ***** **    .    : ***    .  ****.*..

40             MYMYTTASKWYYCYCCAATTCACYCTACMCRRKYKYAATTCMRYTKCYKYYAYRMAAKRY
45             ATACYCGCTTYCYYYYWRGWYGMTYCGGMAAGGYKYMWWWYARYCKSCKYYRYRACGKRY
                       ... *  *       .     . *.    ***      ** *. *** ** ..***

40             RYKTKKYKACACTTKRYCATWCAAGYYRRRKYYTGGKTCTKKMRCRTTTTRKYAAGKWRR
45             RYKWKKYKWMMYYYKGCYGGATTGRYYRRRKYTYSAGGGCGGMRARWWCGGTYMWATTAG
               *** ****       *     .    :. *******  ..       **.*      .*  ..

40             RRRAYTTYMCYTTCCAGGCCGCCCMKWGAAKCCCRRWSACWGRAGWYTGTMKKMMTRATT
45             RRRMTGGTAYYACTARRKYYKYMMMKWRWMTMYTARWGRYTRGGAAYCACCKKACAAMWW
               ***        *:  .         ***    .     **.      .. *  .  **  :

40             CRMMAYYGGCCTYTAYTTATYGMRRAGGTTWARRRYMYYRAGRRKTCTYTATGGACYKGC
45             YAMMMCCTRYYKYKGTACTAYKARRGATWKWCGRRTAYTAMKGRKYYWTCGCTTGTTKRY
                 **        .*.. : ::*  **..  .*. **  *       **       .    .  *

40             CGYRYWMTGRYKYYRWWYACAWYTMKRGRAYYTGYYKCAAKRYGAKKAYTWWMGKKCYTR
45             MRYGTTAGTRTKYYRTAYCGWACCAKRRAGYCCRCCGYRRKRYKGKKRTAWAMRKKYYCG
                 *       * ****  *.       **   .*          *** .**   :* * ** *

40             WRMWTRYRMCTCYKTYSYKYWYSCSCAATYYAYCGRKGGACGYAYCCRWGKYSTWRRMAC
45             AACWGRYRAMWYYKYYGTTTATGTGTTGWTTRYYRRGAAWTCCCTYTGARGTCAWRRMWM
                   * ***     ** *. .     . . :.     *   * ..      .          .:****

40             CTMKYRRCRYWYRTGGASRATRTRSMRCCCKAWRKYYYYATCTYYRRWRRKKMRAKKYRT
45             TGAKTRAYRYWYRCACGCRTAGKASMRYYYGGTGKYYYYMYYYYYRGAGAKTMRRKKYRC
                   * *  *****  . ..*:: . ***     .  *****     ***     *.** ****

40             CCCMYYYTSCGSMYCKRYTKCMMRTGKCGCYYGKYGRKATRRRMGMKMYWKMGYKRRKYC
45             TTAMCTCYSMRCMCTKRCCGMCMRCTGTTTCYRKYSAKTKRRRMKAKMYWKCRCKAAKCT
                 .*     *  .*  **      **         * **. *:.****  *****    *  *

40             KRTAKKRKRGMYKCGGYGTRCRWRKRAMRGTWYCTMACCYCKWYYTCTGKTAAWAMRYTT
45             KGGGKKAKRRCCKTACTAKATGAAGRRMRTGTYYGMRYYTMGACCKMCAKYMRTTMRYYY
               *   .** **   *  .  ..         * **    *  *           .  .*     :***
```

```
40          GRRKCARCRTTRWRMGKTYCATAWYTYATRCGAWKATTTKYYKGARYCCGATGMYYYYCA
45          KRRTMRGYGYAATRMRGKYYRWCTYYTGWRYAGWKTKYGKYYKATRCMYSRYKCTYYYAM
             **.      :  **  .*   . *  . * ..**:.  ****.:*   .     ***.

40          RTWAYAGKGTWTRGTACTGRYMGGWCCRWTCTCTTTRTTTGAKTTYATMCAYCTKYAYKM
45          GCWMYRRTKCWKRRATMYSGCCATWMYGACMKYWCCGYYGAGGCGTRKCMGYTWKYMYGM
              * *  .  *.* ::  .   . *      .        ..     .  .*  ** * *

40          TTYMTRYCTGGWARTMTTRYCYTWCTKTKAGRYGMCATTGAKRYMATARTYWWYRYRAYY
45          YGCCAGTTYRRWGGYMKYGYTTCWAAKWKCTRYRMACAGATKRCMCACGCYWWYGYGRCY
                :      *.  *.  *   *.:* *. ** *..: .:** *.:.  **** *   *

40          CKKKKCKRKRKCRTAYCTRYRWCCKTCARRGRGGCCAKKWGWYKRTAGGGTKGGKKRKAR
45          YKKKKAKRKGKMGCMYTAACGTATKYYRRATAKRMYRKKAKWYTGWMKKRYKRRKKRKCR
             ****.*** *    * :    . *   *        **  **.       *  ****.*

40          YKRMKKKAAGAAWTRKRAATAGCRWCTGYAGYCTCCGKKYRYTGTSYKRKGGSWYRRWKM
45          YTRAGKTMWAWWWYATRRGACAGGATCTCGTYMAYTTKKTACKKYSYKRKRRSACAAWTA
            *.*  *.  .  *  .* .:..       . * :   **   .  *****  *    *.

40          WYMKAAAKAAGGKACKMGCRYTYTGGKYYKKGTTASRRKGRRGTCRMYWTRCYCYYAGGR
45          TTAKGGTGTGTAKGTKCAGRYGCCRTKYCKKKKYRSRRGARRAATGMYTYAACGTYCTAG
               *..: :. .*. * . **     ** ** .  *** .**.:  **   .   *. .

40          CRYKRGATCWRTCCWRKRGKAGGYYTGRGTYWYRRSYYYWKKYKKGAAKGTRYRRAARGR
45          YRYGGKCGTWRAYGWRKGRKGRRYYYTRACYWTGASYYCWKKYGKRMRGKGRCRARMRRR
             **   .  **:  ***  *.  **  *. **   *** **** *      * *   * *

40          RKRYTKTYMKKRRMTTTKTKGMWWTCGTCGKMMCKYGYMYRYASGTRKYTCTYARWGCKR
45          RKRYAKCCCKKRRMYKWKWKRMWWCTACYAKMMYKYAYCCATCCRWAKYKMATRAAKMKR
            ****:*   ***** . * * ***  .  .*** **.*    ..   **. :      **

40          KWMTTTYTKRSYWGKMMCKRRMMTYRKRKKYWGCGCAGCYWWSACRKWYWYKKWMRYCGW
45          GTMGCCCYKASYWKGMCMKRRMCWCRKRTKYWKYRMTSMTWWSGYGKATTYKKACATMKW
              *     * ***  *  ****   ***.***    :.  ***.  *   ***      *

40          TTYYMTRCAWKGYTKKACARCRYYRKWMWWRMYKKTKYGTTYTGTKRRTAGRTKWWYAYG
45          KYYCMARTGWKKYKKKTTWGMRYYRKWCWWRMYKKWKYACYYYRWKGGGWRRYKWACTTS
            . * *:* .** *.**:    ****** ******* **.  *   *     * **  : .

40          GCTAAMGATYCKKRGWYCRGAKCTGAATYAGWATWKYYRKRMKSTGMAGCKYMKWAAGGC
45          RTGCTARCYCSKKGRATYRAGKYYSWTGCRRWTYWKTYRKRMKGCKMCKYKYAKWWRRKY
               .:  .  .**     *..*  . :    *: ** ******.  *.  ** **

40          CCGMGTCTCTCRKWCGRKMMYYRYYGKGCATACGSAYKWWCACTTARSRKKKWTACMWMM
45          YSKMCCAGYWMRGTAAAGMMYYGCTRKKMRWWMKSRCKWTYGTCAWAGGKKKAYWMMWCA
             . *  .    *  ..  ****    *       *  **  .  :  . ***    **

40          RTYAAYRWSCCTCKGAGAGCGKGCKKKAGGYRWATRYCAYKRMKGGGTCTTKYWWYKMWA
45          GCTWWCAWSTTGAKAGTTTTTTAYKTKTTACGTCGRCAWYGRMKRRAATKKKCTWCKMWR
                   **   .*.. :   .. *.*: .   . * . * ***  .: ..*  * ***
```

```
40      YRAYKCTKRKMKYTKKKMRTGRRRCCRCTGTYCGRYASRRWYWWRTKCKGCAYRKMYGCT
45      TGMYKTGKRGCKCYKTGCRCKAAGTGGAAKCTYKGYTGGAATAARCKMKRYTYAKMYRYY
           **  **  *  *.  *        .:      *:.      * * *  :* ***

40      TKGACYTGRTAGGAGTGTCCYRGKMYRYRYSRRCMAKYKCCCRTAYCAAMRKCRCKGCKA
45      CGSTMYKARAGAKWSYKYMYCGKKMYATATSRRYAWKTKYMSGCWYGGTAATGAYKRMKT
          .: *..*:..  .        ***    ***   * *  .   * .:  .   *  *:

40      KGYKYGGMRRWAKRGRRKSMGKTCTKCMYKYMAWTKRGCGCYSWKAWTKGAGATGGAYCC
45      TKYKCTRARRWCGGRAGGGCKKCTWGYATKYMRTWKGRTCTCSAKCWWGRMTCAAAGYST
        . **    ***.      .  *       ***   *      * *.*     .:...*.

40      YATTCTYYMKMWTTCTCRKCCTAKGCYYRCGYTCMSYAYMTYCGSCKTKKRATRCMCKGY
45      YTCYYWCTMKMWYYYYYRKTAYTKKYYYGTAYKTCSYGYMYTYKGMKCTKRRCGYATKRC
        *:      ****     ** . :*  **  .*.  **.**    . * .**      *

40      CAACYKRYCTSWTRRGTCGTKKCGGGRYYKATGAWKGCAKGKYRGWAGKGKTKKRKAKAY
45      GMTTYKRYYCCACGGACAAATKMAACGTCKGCAGWKKTGKKKYRRACAKRKAKKGKWGTT
          : ****  .    . ..:.* ..    *. ..**  .* ***  ..* *:** *  :

40      TTWYYGKGACKYCWGCMAKMRCKGWSKCYRCGMWTRWYKWKKTATGGTATCGKTTTATWY
45      CCAYYKTKGYKCYWTSAGKMGATKTGTYYGGAMTKRTYTAKKYCYKRYRAGRGACCGGAT
           ** . . *  * . .** ..  .. *  .* .* *. ** .     :   :  .

40      RYRCGGRARYYACYCGYRYRMRGKRAKAMYKCKYSCGRSWGWYYYYKCARMYSYRTRMTT
45      GYRGTTRGRYYRYTYRYRYRMRKKRRKGCTKSTCGMKGSTRTCCTTKTCRMYCCRKRAWK
         **   *.***     ****** ** *.  *.. .   *       * .***. *.*  .

40      TYGRATRTCCKTKCTRYRRKYMTCRKCCWYTGKGTYATCRAWKKYKKCWWRSWWAATYYS
45      YYKARYAYMYKKKTGGYARKYMWYRKYSTCCTTAYCCCGRRWKKYKTYTTRSWWMTCYYS
         *        *.*   * ****  ** .    ..  .  * *****.   **** : ***

40      WKGWACGWRTGAGTCYWCTGAAKRCKTGACTYRWWYAKKGATRRYYWCYCCWRCMYRKTG
45      WKRARYKWRCACRWMYWYYRCGTRYGCRCMKYRWWYWKKKGCGRYYWAYYYWRMMYRKYT
        **     ** ..   **   ...*    . .***** ** .  ****.*  ** ****

40      AAWKRTCGWATKYGKAATAKYYRTRGCGKACAWWYYKAKGYCCCKRRKMWYYGRKSGYCR
45      GRWKRCYKWRKKYKTMGWMKYYRWRTTAKRYRTATTKGKTYATTKGRKMWYYRGKGAYSR
        . ***   * .** . .  **** *  .*       *.* *.  * ******  *..*.*

40      YYKCTAGKAMMKATCGTKCYKKKCCRCGCGSRYYRKKYRRYRYMYTRCCSRRYCTTTTWW
45      CCKSWRAKRAMKWGARKKTTKGKYYRTAYAGRYYRGGTGACACCCYGMTSRRYMYYGWWW
          *.  .*  **  . .*  * *  * . ..****              ****     **

40      AKYCAATKAGWCAGWAWRYRATTMGGGAATKTYYCTKCKRTTGAAKAGMTAACCAATMGY
45      TKYTGGAKTRAYRSAMAATATCCMRAATGWKKTTAATMTAYWKTRKRKACGTATGRYMRY
        :** ..:*:    .      :  * ..:. *.  .:. .    : *    .:. .  * *

40      YKAACYYRYMRKYKYKRYRKMTMCTKRKGAWYYGTYCKKKTYTAYMKKTKKAYYKTRAAC
45      TKRWSYYRYCGKYKYGAYRGMYMTCTGGACTCTKYYAKKKCYGMCCGKAKTWYYTYAGGT
         *  .****  ****  ** * *  .  ..     *.*** *     *:*. **.  ..

40      TKAKGCKCTKTTGATTCKYYRTYKRCAGGWTGKYYMSAGGAGGCWYKMYRMCCYRGGRYC
```

```
45      AGGGCTGTGKWWKCCCTKCCGCYGAMMKSWCAKCCCSRSKWRKYWYKCYACYTTGRRGYT
        : .      *   .   *    *     .* .*   * .     *** *         *

40      RRYYAWCGKYRARKYRATMCCAAGGAGGKGRAGTAKRCTCWGTSMRRSYRKKKRKRYWYA
45      RAYYTATTKYGGRKCRGWCMTGGRRRKRGRATKYRKAACGAKGGARACTAKKGAKRYWTR
        * **:   ** .** *.    ..        :   * .     . * .  **  ****

40      RKYRKRTRTYRCKGRKTYRYWRRGGCCKMYYYMRYYWKKYYWGYTATAKWRYRSKAGWRY
45      GKTAGAWRCTRYKRRGACGTWRRRKYGKMYYCMRYYWKTCCTRTGTWRKAGYAGKMAAGY
         *     *  * * * :   ***    **** ******.      :  *  * .* .  *

40      YRGCGGTKYKCRGMGACCAAKYRCTTARYKGGMATCAKKCMTKKTTAYKRRRKMKAKTAY
45      YRKYRRYKCGMRRMAWTATTTCRMYYRRTTKCMWWTGKKYMWKTACGYKGRAKAKTTWRY
        **     *   * *.  .::. *    * .  *   .** * *.: .** * * *:.  *

40      YSRYKKYKATCTCGTTRKYMKAGRRSSSCRYMMYCKRRRKTYMGTKYYRTGGSYKWKAYT
45      YSRTGKCTGYYKYKYYRKCMKGKAAGGGYGTMMYSGRGAKGTAKYGYTACAACYKWKCYY
        ***  * ..  .    ** **.   ...   ***. *  *      *   ...****.*

40      CTTTAGTKRYRRWKAYMRKCGTTYCRKYTRCTRGAGYKYRRRRMRRMKMRRGAKKCMKWK
45      YCAKGRCKACAATKRCCAGTAACYYRKYCGTWASRKTKYAAAGARGATMRGKWKKYAKWK
          :..  *     *     .:  * ***     .   **     *  .**   **  ***

40      ARRWRYYYRKMMSYCWRKTTGYCAGCTCMMAKARRRRYKYAATYKMRRMTTCCRRTYYRW
45      TGGTRYYCRKMMSYYTGKAKRYSWKYYMMMMKWARGGTKTMRKCKMRACCCTYRGWYCGW
        :   *** ******   *:. *.     ** *  *   *   . ***      *  *  *

40      CTCYRCAYTYRKKCTGKKGYYRCAYKKWGGKRRWKWKKYGGRWWKSMTTTGKRACAYYYW
45      YWYYGYMYWYRKKAYRTKAYYRSRTKKTRKTAAWGWKGYKAGTTTCCAWKSKRGTRTCTT
           *   * ****.  .*.***.  **   .  * ** * .   .. : ..**.

40      ASCMCGRRATSCRKRYSGCKYACCGCTCATCYACYKRRACTAACWWKYCWGKRKRTCKTW
45      MSTCTARGGYGTRKRYCTAKCTAYTAAATCYCGSYKGGRSKGWSWTGYYARKGGRKTKYW
         *   .* . . ****. .* :.  .:.:   ..**   ... .*  *   *  *. * *

40      RMTMYATTCAMCKTKYSMMWKCGAGKRYTYMKRYYKWYKACGGCTCMKATTKRAGKKTCK
45      RMCCTTKCTGMMTCKTGCCWKMRWAKRYYTMTRYCKWYKCTRKYWYCKGCKGACAKKCGK
        **   :.  .* . * .  **   .***  *.** ****.       *. .  ..**  *

40      KRMRKWTCMRYGTTTTSSCGGRMAAGKGCKKMYMKTTKCMACAYKYMRMMWGKCKTRYKT
45      KGMRTTCTCRYAAAGYCCYRKAMRWRGRMKKCCATCGKYMWMMYKTMGMATKKTGARYGY
        * **.    **.::  ..    *      **   .  * *   ** * *   *  :**

40      SRCGGRTTYYYMWKKYAYKWYYKCTACKGWYWWAWRYKCKYYKRWKACGRARRRRMKKWR
45      GAYKKRYKYTTCWKTYRCGATCTYWTMKTWYWWMWATGYKCYKGTKTTAGGGRRRMKKWR
        .    * .*   **.*      .  : * **** *    * **  *: . . ********

40      WTYTRKAGAYRKTTATATACKARKRACTCTKKRRYRKCYWAAGRCKAACAYKRYYWYKWC
45      WCYKAKMTGTAKWARCRGRMKWRKRMYKAGKKRRYRKYTTRRKAYKWGARTTATYTYGTY
        * *. *  .  * :      * ***  .. *******        * ..  .  * *

40      TGRRYTCCGAGAAMCKYTMYMKRSKKCKKRKRGYWYYTGAMYYRCGKRAAAATAGWKMSC
45      KRAGCATYRRRWWCYKYYCTAKRSGKYKGGGGRCWYCYRWCYYGYRKRCGCGCGAWKCCY
```

```
        .    :          **    *** * *      **      **    **.... ..** .

40      KTKRRTYGAARYGCMRKRKKTYKGGYAKRCTACAATGGGRTAYRCCWARTCRTCYGTTWK
45      KCKRGCYKMCACRYAGKRKKKCKCRYWGAGAGTRMYSRKGYRTRYTWGGWYRCMYSYYAK
        * **  *  .      ****. *  *    :.    .      *  *.   *  *.   *

40      KKACCCTTCTKKKSARKAKGGGTKAGGYTGWYYYRCACYRTGTYRRTRTSCTCYGAKYYY
45      KKGMYYWYMCTKKGRRKRKATAYKTRRYYRACCTGMWYCRWKKCAGCAKCYCSCACKYYY
        **.       .**. ** *. . *:  *           *  .     ..  . ..****

40      AYACAGAKCYYWKYMMRKMCRKKYYTTAAYKRWRRKKGYACSYRTGCRYGMKTAKRYMKY
45      WYGYWTWKMYYWTYAARKAYRKKYYYYRRTKGTGRKGRCMYSCGCATGCKMGWWTGYCKY
         *.    * ***.*  **  *****     *   **     *   .    *   . * **

40      WYMYKRRMYMYWRKRRMRRGGCYGRGCKGAAGRCCAYYRKKKTTCTKGKRMYCKTGRAAT
45      TTATKGGMCCYWRKRRMAGAATYKRRYGKRWRGATMTYRKKKCCTWKRTRAYTKCRARWW
            *  *  *******  .. * *        .   *****    * .* * *

40      AYYGGGTACKGGGACKKGKRSKYCKYACTKSGMMRKCRGKMYRSRCYKKSYWYKCAKSMC
45      RYCKSRYWYKRRRRMKTAGRSKYTKYMAYKCRMCAGYRTKMTGCATYKKSYWYKTGKSCM
         *  .    *     *.. **** ** . *. *    * **  .  ******** .**

40      TGAYKYYARRAGCYKMKGGYGRMRKKWGRYAGGRMKSYMWKYMCTKACTGAGTRKCTARY
45      ARRYKYYGRRGTTYKMKACCKAARGKTRGCRRKAAKGTCTKCAMYKTTACMAWRKTGCAT
        :  ****.**.  ****.     * *         *.   *    *: :  . **  .

40      CAATYCACTGKGRMWRKARKGYKKYYKCRGYRSKAYKYRGCRCWCYCGACKKKTYYYWRY
45      GTGCYMMYKRGRAMTGKTRKACTKYYGMGRYRSKRYKTGRYRGAMCYTWAKTKCCCTTGC
         :. *   .    *  *:**. .***    **** **    *       .*.*

40      KYYGTCTKKGAATYTTRATGGTTTKKMMKRWAGRACTCKWCRCTTYRYMMKMYYKRKRSR
45      KCYRCYKTGRCGCCYARRWRTWWKTGMMTGWWKGMYWYKTYRAYWYRYMCGMYCTRKGCG
        * *   ..  ..   :*      .. **. *       *  *.  ****  ** .** .

40      CYKAMKGTTYKTCRATTGWKGTAWYYMKACRCRGACYYRKRATGTCYRKMGKYMRKCYGA
45      AYKMCKRWYTKYYRRWWRWKKKRACYMKCYRAGACGCTAGATGTCACATCRKYMRGYCRG
        .**  *    *  *    ** .   ***. *. ..      :   .  .  ****    .

40      TGKYKKKAATCKWGKYTRTCTGGKGATTRKYMAACGTTYYYTCGKCKKCYCCCKRKAAAS
45      KKTCKKKWMYYKWTKYWRYMWKKGATCCRKCAMGARYYYYYYTAKYKKMYYMMKRKWRRS
        . . ***    ** ** *      .:  **   ..   ***  .* ** *   ***   *

40      YYYYTRYKRWKWWRAMYYRKKRWGCYRKSKKKWRYRKKCGCTKYSRKTAKKATTARYKMA
45      TYYYYGCKGAKAARRMYYRKKRWRYCAKGKKKWRYRKKARYYKTSAKAWKGCYGTGCKAR
         ***   *  *  * ********    *.*********.   * * *: * .  :  *

40      TWAGTKMWMMYRASTKRKWYRWYRAMMRRCGCMKKCYKCTGYGRKKRYMWYRYTACCYYK
45      GTRCGKCAMMTARCYTGKTYRTYGWMMARTAACGKMYKYYSYRAKKGCCWYRYWRYYYTG
             *  **   . . * ** *  ** * ..  * **  .*  **   ****    *

40      RWKMMMTWTTAWRAGRWRCACWKRKGTGAGMRCTRYCTWKMKAARGAWYKYACGMRYCKT
45      RTGAMMCTYCRWGRKAWRYWTWKRKKYRMAMGYYGCMKWKCKRMGTWWYKYGYKCRCTGW
        *   **     *    **   ****    .*      .** *     ****.   *
```

```
40      WGAGSKKTGMTGMSWGAARKKTATYTGYWWRAAGCTWKRRMYKCKRGMCRKAYMGTAATY
45      WTTTGKKKRMKKASWKGCGKKCGCCAATTWRGTRYCWTRRMTKTKRTMMRKGCCRCGGCY
        * : .**. *.  ** .. ** .  :.  **.:   *.*** * ** * **.    .. *

40      RTCTGACWKKYCATCCTWYCCTRTTCTATMRWTAYAKGAACTTAAWYGAMYGGCRTTKTA
45      RATYRRSATKTMTCGAWWYYYYAKWYATCMRWGRTRKAGCGAACGTCARCCKKMAYYKYW
        *:    . .*  :  . **    .  :: ***    *... ::..  .         *

40      RRKCKGMTKTTCYAWCTCCCCYRGGGCGCRGKGCGACYCGKTGGKMRRYYGKGSYWKRYK
45      RRGYKTMAKGYYYGWYCTATMTGKSRYRYRRKCGTWYYYAKYSAKAAAYCKKASYWKRYK
        **  * *:*   *.*   .     .    * *     * .* ..*   *  *.*******

40      RMATMMYKGTTRKYKCMRYCCYYKYKMSRCGGTYMRYCCAYRYGTTKGTGWCYCRCTKKM
45      GCRKMATKRYCAKYKYARCYAYCKYTMSGMKKWCMRTTGRCGYKWCKKYKAYYTAMYKKC
           .*  *    ***  *  .* **.**      **      *   *     *    **

40      CRTYRYYYRWYKYYTWTYKRWYYARMCRYSYKYYKYCRRRAKSWGCTGATCCGGCCRYKT
45      YAGTGCCTGAYTYYAWKYKAWYYTRCSRYSCKCCTYYRRRMKGWKYGATCMTTATMACKK
                  *.**:*.** ***:* .*** *  .* *** *.*   .:    .    *.

40      MMKRRGTSAMATAGTCYKRRMCYYMYKRYSKAKKGRWKGGKWKYGKMYRCRCRYKSKGTM
45      CMKGATYGMCWKRCATCKRRMTCCCYKGTCKGKKAGTKRKKWKTAKMYRARYRYKGGSWM
         **    .   .  :  ****    **  .*.**.  *  *** .****.* ***. . *

40      KRRTAGRRKAGGRTGKMWRYWYWMRKRKTTYTTGCKWKKGKAGACMCCCKKCYAMRR
45      GGAWGRGRKRRSAWTKCTRYWTAAATAGKKYCGRGKWKKRKMSRYAMAYTKATMMRR
            .  **  .   *  ***    .  ..*    **** * .    . .*.  ***
```

The pairwise alignments of two accessions 40 and 46

```
40              KYMWCCRGYSCGYMRWRRKAKGRGYMGSRRRYRWYWRRRYWCCMMYYMKRWRKGRRRYRR
46              KCCTYYGTYSARCARWGRGWKARTCAKSGRATRTYARRRTWTMMMYYMKRTGGKRAGYRR
                *       **.   ** *  *.*    * *  * * *** *  *******    *  ***

40              MTWRYWGSRKSRRYYWSWAGRCYTGMKWWMRCYKYRSRTRKKYYTGRYYRRYGATYWKCW
46              CYAGYWKSRKSRRCTWSWRARMYYKAKTTMGYYKYRSRKGKKYCWRACTRRYAWKYWKYW
                    ** ******  *** .* *   *  *  ******. ***      ***. .*** *

40              MRYRKRYYSYWWYGKYRYRYMKAKTGYYRRMYYCYYMYKAYWGGGRRSRRRSWYRKRRYA
46              MRYRGACCSYWWCRTYRYRYMKRKWKYYRRCCTYTYACKRCWKKRRGSRRRSTYRGGGCR
                ****    ****  .******* *  ****     *  *  *   * ***** **

40              TYKGTGKYRAYYYYWRTRCTGGCRYYYRYWMRYKTKKRRYGWWMMRCGATKRAGRARAGR
46              CYTCYRKYRGYYYYWRCAYCATTRYCYAYWCACKCKKRRYTATMMRTRRWGRCRRCGTKR
                 *.   ***.******     . ** * **   * *****   ***     *. *. : *

40              RMCRWCTTYYYKYGYYYYYYCKTWAYYMGCAWAYCCKKWRYKRKYGRYKYCYRGGATWKY
46              RMARWMYKCYCGTKYYTYYYYKYWTCYMRAGTTCTTKKWRYGRGYKGTTYYYRAAGCWKY
                **.**  . *    ** *** * *: ** .. :   ***** * *   .* **... ***

40              MMRCMYKYKYYRAYTYAGKRAYATCAWGYMYKRAYAYYMKMWYRRTCKWGWYMKMKRGKG
46              MMRYCTKCKCTRTCKYRTGRTYGCATWTCMTGGGYTYYATCWTRRWMKARTYMKMGATKR
                ***   * *  *: .*   *:*. .:*  *   .*:** . * **  *   ****   *

40              TMKRRTTCCTAKAYWYMRTMMWYRYYWGSYMRYYRWYWYMMMAGGKYAGAAWCWKWSRRR
46              WMGAACYTYYGKRYWCAGWAAWYRYCWKSYMRYYRTYWYMMMRRRTTTTGWAYWKWSGRR
                 *        .* **      **** * ******* ******   . : .   **** **

40              SRWTRGYYRWCWGGWWMKYMTAACYKWYWRYMYMYYSYYYWYMSKWKTYKYRRAMYKRRK
46              SGWKRTYYRWYAKCTTCGTMWCWYYKWTWRYMYMYYSYYYAYASGTKWCGCRRMCYKRAT
                * *.* ****         * .  *** ************ * *  *    **  *** .

40              YYRWKKCAAYKCGGRKYRRMYSKTCGRYRWYMYGCKRKAACCYYMYRMWRMRKSRGMRCM
46              CYRWKKSWGCKMRAGKTRRMYSKYSRGYGACMCKMKRKRRTTYYCTAAAGAAKSGACRYC
                 *****. . *  . * ****** .  *   *   ***    **        ** . *

40              YRRKAWCTAWYMACYKYCAATAYSRRAMTMKYTCYRCKKATRCYCCGTCRYTMRAYYTTK
46              YRRKRTTGWTYAMSCKCTTTGCYCGGGCGMKTASCGYKTWKGMTYYKYYRCYMGGYYYWK
                ****      *  . *  :: .*.  .  ** :.   *. .        *  * .**  *

40              KTYCGYRWAASCYYTGCTKYCMCTCGTMAYTTTCAAYKSCAAKTYAGCTATGGWTCRAYK
46              KYYARCRWWCCYYYYKSYKYYMYCYKYMRYCWYYRGCTSYCGKCYCCYYWYCAAWTRWCT
                * *.  ** .. **  . ** *     * *     . .* ..* *.      .   *  .

40              YKKRCKTGRGYKYKRRRATYGRRTKTAGYATCYACTRYRYYKYACYAGCMTCACGGYCWR
46              CKKGYKKRGRYKYKRRAWAYARRYKYGTCTCTCRMYRCATCKYMMCGAMACTGYRKCTAG
                 **  *.   ******  :*.** * .  :      *    **   ..    .

40              KCACRCYTAYKAMGKYYYAYRRGKKCCRYGSTKTGATMARRRKWKMAKKTAAKGCKMMRY
46              KYMTAAYYRYGGCAGCCYRYRRRKGYYACKSKKYRMWMRGGGKWKCMKKYRMKRYKMCRT
```

```
            *    .*  * . .   * *** *      *.*    *    ***  **   *  ** *

40          YATRMRRCGKRYCTCAMCTWAKTGCCCCGCRCRTSSKYGATAAMAMSWRKTRRTGCGAGK
46          CRKRMRRTRKRTAYMMMTCWTKWTMYYMATRGRGGSKYRRYGWMMASWRKCRRGAAATTG
              .****  ** .   *  *:*      . * * .***   . *  **** ** ...:

40          RAAGTAACGRYRGKRRTAYRCGSAGMMGGMRGTCGASYGCCTGACCGKKKTGKYCTGMMY
46          GGTRKGTGRGTRTKRRCRCRTACGCCMRSMRAWYKWSYKGTYRRSYRGKTYRKTMKRMAC
             .: ..:    * ***   * ...  * .**.    **      .   *.  *  . *

40          WYMRKGTTCMRWAYGAGKCTTAARTCTKSYCRTMYYATGYRWTYRTCTTCCCYYRKGACC
46          WYMRKRYAYMRWMTCRKKGAACGAYYYKSYYRCMTYGAKTRACTGYYWCTATYCGTKMMM
            *****  : ***     * ::..    *** * * *.:  *         . *  .

40          RGWWMKRCCGCTTTAYMKYMYYMMSTKTCMGGCRKCCTWKMYGRYYTSRWRARYKCRYRR
46          RRTAAKRYYTTKGYRYCKCCYYMMCYKYYMRKYGGYYKWKCYRRYYWGATGRGTKTRCRA
            *    **    .   * *  ****. *  *       .** * *** .      * * *

40          YYYRMGKYAYRAAARAAGGAARWKRTSMKRRRCRSYRTGGGRGKKRYWYAYRKRTRRTKR
46          YYYGMAGYRTAMRRGRGATCGAWKRGGAKRRRYRSYRWRRKRSGKRYWTRTAGAWRGCKR
            *** *. *        .. .. *** . **** ****    *. ****       *  **

40          RYRCKATGKTGCCCGCCGWYRYAWRRGGCKYMYYWRYACTMRYYKWKKYYGGYTAAYWKR
46          RTRYKCCRGAATTYRYYRWCGTWWRRRKYKYCCCTRCGYGMRYYKWKTCCKRTGTWTAKG
            * * *.   :.       *    ***   **    * .  *******.      :   *

40          MGRSTGWYRYYRMACRMGCACTTAKACKRWYCAWYWSTAGKKCTATARYKYGGKMKRMKC
46          MRAGYAACGYYRCRYAAAYTYWYMKTTTAATYGACWSYRKKKYKRYRRTTCKCKMGAAGY
            *  . .   ***     . :    *: .    .  **   ** .   * .   **

40          KACKRTRKMMKYKTATATCTTCCRCYYSRYKTYGKRMCGCRRYRTSKKRYGWCTCATCKY
46          TWMKGCAKMAKTTWRYCWYYCYMRYYYSRTGYCAKRCTAYRRYGKCGKRCTTTCTTYSKT
            .  *   ** *  .   .      * ****    .**  . *** .. **      : .*

40          KAKRCKYKMSACCWARRKRGRTMYATTCACRTKAGYTKKTRYYKGKWKATCMTYYCAKTC
46          KWKRSKYKAGTTYARAGGGKRCCTTKCTGMRCKTACKKKYRYCKKKWKGWYCCCCTGTYG
            * **.*** .:         *   :.  . * *:. .** ** * ***.        ..

40          KYGRGKWWKTCMRYYGTTTYTSCGGRCGCMYSMKTTGCATCMKTGAAGCGYAYKYMRKKK
46          KTRRATTWKCTCRYYRWWKYYSYRKATKSCCCATGCRYRCYMKAAGTCATYMYKTMGKGK
            *  *.. **   ***   .* *      .  . .       **:..: . * ** * * *

40          CAKTSRRATAYYMKWWWYAKAAKYCRRCYWATYYCTWKGKKRYMKKWKYTGACAKGKYWW
46          TTGYGAGGAGTTCKTWWYCGGGTYTRGYYWTWTTTYWKRTGRCCKKWKCWRRYRKKKYWW
             :  .  .:.   * ***. ...* *  **:     ** . *  ****      * ****

40          WKKKCKKKMKYKAATTGYRRRYRMASRRRMGARGAGWMRWTTAGMKGGMYRRRRTGMYCC
46          WGKKYKKGCKCKMMYCKYAAGYRMRGGARMRTRARRTAGWYCRRCKRRMYRRRRWRAYYM
            * ** **  * *     *   *** .  ** :*.     *     *  ******   *

40          CAYKMRCRCKGCCWTGGAMWCGYCRYKCYRGYWRRKSMKGATKCKAKATRRRARRRYRYK
46          SWYKAAGAYKRMYWCRSWMWTKYYRCGTCGRTARGGGCKKMWGYKTKRKRRRMGAACGCK
            . **     *   *  . **  * *        *  . *     *:* .***       *
```

```
40      KAGACKAYCYCGKYCTTCTYYKTTCTKCCAGGTTTAGRMGTYTRGTWGSYGAMTYCSCTT
46      KWRMYTGTTYGRKYSYYYWCTGYYYYKTAMKRYYCWKRMKYCWGRGWSSYARMYTYGMCK
        *    ..  *  **.           * .        **       *.**. *   .  .

40      AKKKYRWACTYMWWKWGGCKWYTKRYYKWYWKWATRACYAARKGYCARCYRCTRGYMYYA
46      WTKKYRWRMWYATTGTKRTKWYCKGTCKWYWGTRWGTMYRWRGRCMRRYCGYYGRYCCTG
         .*****   *        *** *   ****     : *  *     *       *   .

40      GCKYTAKKYRGCCKCWACTCKMYCTGCGAWGAAKCAYYAAKTTGKRYMGGKWKGRMRRCR
46      RYKYGMKKYRKATGTTCTYYKCTYKRYRRARMMKMMYYCTKCWAKGCCATKWKKRARRYG
          **  **** .    .   *   .        *  **.:*  .*   . *** * **

40      RWGTCRGTTTGAKYCRARWTRRYSCTYYYRTRCWRWAKKRCCATCYTAKAAKTAAMKKTK
46      GASKTRCYYGAGKCTRRRWKGGYSTWYTYRKRATRWGKKRYYWKYTGGKMRKWWTCTKAK
          .. *    ..*  * **.  **  * **.*. **.***   .  .*  *   :  .*:*

40      RAYYSKCYGAYKSRCRYKGSKRCGWRWYKSGTKTCGKAWYRYRSCKARYRKCTKCKACYM
46      GWYYSKYCTGTGCAMRTKAGTGGATRTYTSRYKWSRGMATGYRSAKGRYRKMKKYKRYYM
          ****   .  .  * *...  . * *.*  * .      ***.*.**** .* *  **

40      WYKYGYRGKRTAGARGYATYYCKRRGWGWTYYGGYAYRMYAWYYSYTRCGTGTKGRKKAW
46      WYKTRYRKKRYRKMATYMYYYSKRRKTRTYTTRKYCYRMYWACTSYKRTACAYKKAKKWW
        ***  ** **      *  **.***         *.****    **.* . . *  ** *

40      KAGCTTRYCCYYWYYWRAWKKYKKGAACKYKKCWRWRSWWKATTSWKTGWACWTCTGRAA
46      GWRTKGGTYSYTTCTTRRWTTCGKRCGYKYTKYTRTACTWKTCKSWTWRARYWGYYRRRR
            .    .*     * *..  * .. **.*  *  . **: .**.     *    *

40      TTGTARCKTTGTYRMMMTYRKMCAACYKCCWMYRKKRRGATTKRWYTATGGATCKRTMWK
46      KWKGCRYKYWKWYRMMMCYGGCTGCYYKYYTCCAKKARTGKWGRWYWMCTAMGGGGGCWK
        .   .* *    ***** *    .. **      ** * ..  ***     .       **

40      KMTRYCMGRTTGGTTKCYTRAAGRTCTGGCAWRRMYRARRRAYYMTGYCCCMKWGAAGKC
46      KMWRCTCAGYCAKCGTMCWRMGAGYYYKTAGTARMYRCGRRMTCMKKCYMMMKWRWMRKM
        ** *   .   .   .   * ..      ..  ****. **   *.     ***    *

40      WCRYKRGTRSCRSTARWTGTMKKMMRGMYYGMCCGKYYKRRRATMAYAYTRRYRMTYTTK
46      AYRYKRRYRGYGSCCGAYRYCKKACAKACCKMYTRTYTKRRRGCMMTWYWRRYGAWTYYK
          ****  *.  * .      **        *   .* ****. *   * ***      *

40      AATGYRYWMYRYYYRGGWWCWYTATAAKKCARRKAYRYYYKWMRYAWRYAKCYATAWWRG
46      CCCRCGTTAYRTYYRTRTAMACCGAGTTKYRRRKGYRTCCGWMRYRARYRKYCTGCWAGR
        ..       ** ***        .:.:.*  ***.**    ****  ** *  : .*

40      CYTGRYCAYRMYTRMWCTCKGGMTTKKGKGCCSYGTYKMYKKMKRYYRCCRTKCKKCKRR
46      YYCRGCATYRCCGRMWMWYKARMWYKKKKAMMSYAAYKACKGCGRCYAYMRAKYKKYKRR
         *    .:**   ***   *. *  ** *.  **.:**  *   * *   *:* ** ***

40      MTTKTMGKGMMWGYTAYKKRGCACSRCAWGYYYARWKWTTTGYTKRSYWKKMKRRMTCTY
46      MKWKWCRKRMMAKCWTYKKRRYCYSRAGTRTYTRAAKTGCCRCYKASYWKGCKRRMWMYY
        *. *   * **    :****  . **..   *    *       * ****  ****   *
```

```
40      TMTYYAYRKRYWGKGAARSYWWSRACTAYWYMRKRWYCCYYKTTAYKACRKKMCTYRKKY
46      YCWCYGYRKRYWTKAGGRGTWWSRWTAGYATMRKGTCGYTTKKYWCKCYATKAYCTRKKY
            *.****** *...*. ****  :.*  ***       *.    *.  .*    ****

40      YMRTMRRWRRYYKTRYGKRRKKKRYKKRAKKMGKYMRYWRKTGRRMTRRKRYKMMRKGYM
46      YCGYMGGTGRYYKCGCKKRRKKTGYKKRMKKCRKCCRYWRKKRRMKAGKRYGAAGGKYM
        *   *    ****    *****. **** **  *  *****. ***.  ***      **

40      YWTWGAARAKYKYYRGKKACACGKKMKAYCKGWGAKKTYTGRKSYKGAAAKSYMKRSKYC
46      YWYTKRWRMKYTTYRTKKRYGYKKKAKRYTKRWAGKGCYAARKSYKTCGCTSYAKRSKYT
        **     * **. ** **  .  ** * * * *..*  *:.***** ....** *****

40      WYKTSTTGMCRKTCCACGKMYRKSRCYKYSYWYKSMRCTGAYYKMKGKYYYARRTYKRYG
46      WYKYSGARMGAGCTYCATKMYRKCATYKYSYWYKSMGMARRYCKCKKGCYYGRRKYKRYK
        *** * : *      .. *****.  **********  :  * * *   **.**.****

40      MRKWRRCKWYMCCTACTYKGTRYTGYCAATKCACTGGMYKYYYTCCKTMWTTYTSTGCCR
46      MRGTGATKTCAMMWGMAYKCWRYYRCGTGCKMMYKRRMYKCCCKMYKWCAYACKCAAYTR
        **     *      . :**  **    :. *   .  ***   .  *    : ..:.  *

40      RAYAWTGMTYCATYGWWATRRAWTCRYGAKYYRKRRYTWYRYTYRRWYYTAKCCYYYYRW
46      RRYGACRCYTYGATRWTMGRRGTWSRYRRKCYRTRGYAWYRYKTRRWTYYGTMTYYYCRW
        * *.       .:  *   **.  .**  * **.* *:****. *** * ..  *** **

40      RACRMATKWRYTRGRGARGGGGYKTAAKWMSRGSRMMYYKWARRRYYCYCCCYGWRCKCT
46      AWYRMTGKWRTKGAGRMGARTTTKCGTTTMCRKSRCMTCTTWRRRYYTTMTTTRWRGGMC
           **: *** . .    .    * .:. *.* ** *  .  *****       **

40      GRAMRYGSMRACWTTYGMYTMAMMYRRCCGYRYKCKMYARGTCKATWTYGTGKCGKRWMR
46      AAMARYASMRCGWCKCRMCYCWMCCGRYMRYRYKTKMTRGSYYKRKACYSYSKTAKRAAA
        .   **.***. * .  *    *   *   **** **   .  * .  *. .* .**

40      RYGKGSYWKCTCCRYKKKMATMMYKGTKKRTRWKMGYCRYRKYCTTWMSKMCYMRYKCCC
46      ATKKASYWKTYTYRYKKKCRKMMYKRYTKRCRWKAKCARYGGYTCCTMSKMMCMRTTTMG
           *.****    *****  .****  .** ***   .**  *    ****  ** .

40      YSKYYRYTYRGMCAYGGGCRCKKMRYRYYYRWGYWKCYRWCYYRTCSKRMRGKGATYYKY
46      YGKCCGYCYRKMYMYRRRYAMKGCRTGCYYRWAYWKYYATMTCGYSSKRCRKTRGWCCKY
        *.*   * ** *  *      *  *     ****.*** *     .*** * . .   **

40      WGKYCGGCCMRYRTMMWWSTRSMKYKYMCKRYSGYRGYRWKMMGGKWGTTTCRYSCKCGT
46      WKTTYKRYAAACRKCMATCAGGCKCKCCYKGTCRYRRCGTKMMRKKWAGWKTRYGYGMSW
        * .     .   *. *  .: . * *   *  . **   ***   **.  . **.   .

40      RMRMRTAGYYGRRKRTWGGGKMRWMKWYWWRKRRCKWYYMKKTCGGTWMMCCTACYCTKK
46      RMGAAWGRTYRGRKAWARKTKARWCKWTAWATRAYGWYYMKKGATTKAAAMAATTYYKTG
        **    .  *  **       * ** **  * .*   ****** .  .    .:: * ..

40      CYMRKTCTCAACYRTKMYMAAGSSSRCRYMKACKRRTKTGTTYYTGGTWYTTKRYRKCWA
46      ATMRKYYKYTGGCRYKMYMGGKGGGGYGTCKRSGRGKKYKYGYYCAAKATCCKACAKYTR
        . ***  . :.  * ****.. ...     * . * .*    ** ...    *   *

40      MYCAMYRGKYYGRRRRMRTTMRMTARGAKCMKCKCTRCAKKWRWRYYMATYGCCTCCMAG
```

```
46                MYYMMYASGTYKAAAGAAYYMRMYRAKWKMAKYGGARTGGKWRWRYYMRKCRSYYMMMRR
                  **  ** .  *         ***     *  *   :* . ******** .  .    *

40                CKAYWWRYARKGACKCACKKYRKRGSWTRYWYKMGKATRKYYRARMYYYKKYRCKYTGGK
46                MKMTWWRTWRGACAKYMTKKTRTAAGTKRYWCKMRKWYRKCCGTRMYYYKKYGYKYARRK
                   *  ***  * ...*   ** *. .. .*** ** *  **   :********  **:  *

40                MYCTKKGYMCKYRKWGGKKRRWKWKYYAYGSTMWWKSMKKTRTKKAYKCGWYCRWMWYWK
46                CCAYTKAYMSKCGKTRKKTAAWGWKCCRTKGGMTTTCCKKKRYKKRYKMKWTYGWMWYWK
                    . .*.**.*  *   *.  * **     . *  .. **.* ** **  *   ******

40                WYYWRYRKWRMRYWRGRGTAWRMYTCCTGMYTKTKYYGYKRTKRRTYRKWCCYTYACCYY
46                WYYTAYGKTGARYARARRCMWRMYAATCRAYWKGGTYAYKRCKGGGCRKWGGCYTTATTY
                  ***  * *   ** *.*   ****:.    * *   *.*** *    ***     :.  *

40                RTWYGAKKKRRTKGCAWWYYAACYGGKAKAKSSMTTGRTGKMWGGAYKRMKRATGGSYSR
46                GCATKMKKKGRWKKYRTATTMRACKKKGKGKSSMYYSRAAKMWKRWYKRMKGGKKRGYSR
                        *** * *         .   *.*.****  .*:.***   ***** ..  .***

40                YRYKWGKYKTATAACRCCYWYCASRKGGCMTCAGMRAWTATKKYRRYSRRYMYKTCRCCC
46                CRCKWAKYKYRYRWYRTTTWYYGGRKRATCCAGCARTWGGAGGTGACGARYMYKYYGMTT
                   * **.***      *   ** ..** .   ..  *:* .:      . *****

40                ASRRYTCCCTTTTMSWTTYCTGYKTTRYACRTTCRYCAGWAWTAARYRGAYMKKYCTTYR
46                RSRRYYMYMYYGWMSWGGYMYRCKAGGYRMGYYYGCTRSAMAATCATARTYMKKYAAYCA
                   ****        ***  *    *:  *          .   ::.    :*****.:

40                KGKRCCAKACTGTTAAMYYGAACMRYKYKRYRTTACKMKRKTGAWYYGTYCRTACTKRTY
46                GKTAYYRKTTGCCCGRMCTRRWSMRYKYGAYRCWWAGMTGKKACTCTKYYAAWRYWGACT
                    .    *:     . *     .*****  **   . *. *...     *.

40                KRYYTSRYRCKCTRCCCTTGAAACYKGGGKTTGTAAYYRYKRRRRGCACAGGTGKRYAWK
46                GGTCAGAYRYTMYAYYYYYSRGGTCKATAKWWKYRWCCGYGAGGARYRMMKSCAKRYMWK
                      :. ** .        . ..  *. .*         *           . .*** **

40                CAGCACTKKRYATRTYGRRMKRYYYCCKKYKRYCKKRKTGGCTRRARWKCGGCCARAWYG
46                ARAACYAGKRCCAGCYRGGCGRYYYYMKKYKRYAGKGKCAATAGAWGTKYAAATWARWYK
                  . ... : ** .:  *     ****  ******. * * .. :     * ...    **

40                RKGRKTTGGKYRYKMCCTTKAGWRWMKACGAKYKKTWCKAKAGATAGAAACGWTWYGGCG
46                AKRRTCGTRKYRYTAYYYWKMAWGWMKRYKRKYTTYTMKCGGRGACRMRRYRAYTYAATR
                   * *.    ****.     * .* ***    **..   *. . .:.         *..

40                MCTTTRGKKCYWGWTARSYKRKCCSMWYRMRRWKMWYMAAAKAAGGTACKGCYGCMWRRK
46                AACCYRTKKMYWKTAGGSYKRKYMSMACAMRAWTATTMGGTGTGTACGTGTTYRSATART
                   .   * ** ** :.  *****  **   ** *.   *..: :. . .    * .   *.

40                GKGGCYRRRSRGYWKGRRRRRYWGRCYRYKWATARTGGKRGKKGKKYGCTMTYMCGATTG
46                RKAAATAGASRRCAGARRRRGCTRAAYRYTWTAGRCTTKGRKKKKKTTTGMCCCYRRYGA
                   *...    **    .****     .***.*::.*   *  ** **    *        .

40                WYCCASRWTTGKCGYMTTAYYGGCAATACCTWYRCCACACTYYKGGRCACGTTMRRGCGS
46                WTYMMGGAWYTTYTYAGGGYYRKYGTGGYSCTTRYYCYWAGYYKARRYMYKYWMRRKMAG
```

```
                    *    .     .  *   .**   .: . .    *  .  . ***. *      ***  ..

40                  KMATATKKCGTCWACATYTRTWRYRSRKRKGTKCMWTYGKKYRRCMWTACKTWSYCWAGR
46                  GARWRWGKMKYYTMYGGYAACTRYAGGKAKRKTTMWYYRKKTRGSMWYRTKCWSYYTTAR
                          *      . *:   ** . * * .. ** * ** * .**   * ***  :.*

40                  WRYWGGCKKRMWRATKCTGTTRMKCTTTAGGKYRKMWATAGTGYCGYYKACGRARMCRWK
46                  AATTAASKKRMTRCGKMYSYGRMKMGAATAAKCRKMWRCMTGAYTACYKTYRGRGMYRWK
                        ...**** *. *   .  ***  ::::..* ****      .* . **:     * ***

40                  TKGAYKTWMKYYKRAYYRCRYWTAARKRKRMYRAYKRYAGYGGRYYKWCGTGRRYGYWCG
46                  CGTRTKAWCKCCKRWYYRYGTTKRRRKAKRMYRRTTATGRTRKRCYGTYRKRAGCATWYR
                        *:* *  ** ***    .  ** *****  .  .    * *     .    . *

40                  AGKTCGAGCRKKMCWRRSKTCCCCYAMYYGWYRYCSKAACSCMSRWTATGRRTKAGAAKY
46                  RRKCYKRAYRKKCYWRASTATYYTCGCYYSAYGYYSKCGMSACCGTATGTRRCKMAGGKC
                     *    . ***  ** *.:      . **. * * **.. *. .  ::  ** * ...*

40                  CGGCMRKTYKGAGGYARTYWAYCCAGKTGAACKGRGCWARTSYWMKAGCGCKYKWWRCAY
46                  GKRYAGKKCTSWKRYWACCTCCYTRRGYTWGYGKGKTWRGCSYAMKRRYRMKTKWWRMMY
                        *. ..   *      .         .      *   ** **     * ****  *

40                  KMYARCCAAKMKYRWYYTTGYRCACTYARRKAYGGAYYYYKYCRKTWWGCKGSGTYMTAG
46                  KACRRMGCGKAKCRACCYKATGSWYYYWRAKWCRACYYYYKTYAKYWWKMKRCRYYMKGA
                    *   *   ..* * *     ..  .    * * *   ..*****   * **   *  .  **...

40                  TTARATKRKYYMKCYKAYKRYMRKYKGAWYRKYAYYMYKYRRYKCYMGYGACTKGTYKRG
46                  KKRGWCKRKYYMKYCGGCGGTMGTYKCRTCRKYWYYMYKYRRYKTCMRCKRYYKKWYKGR
                    ..    *******    .    * .**    *** **********  *       *  **

40                  GAYMGSKYWRKTGAATAKGARRYCMKMRTTKYAKKRAMRYYMCCTYKRCWKRRGWAAKKC
46                  RGCMKSKCWRKGRGGGGTRRGGYMCTCRCCKCGGTARARYYCYYYCKAYWGRRRWTGGKM
                      . * ** ***   .. ..    *   . *  *  . .   ***      *   * ** *:. *

40                  AACAYKMCMTGTKKWYKRYRRKWAKKKGAKWCTCCTMGKYRRYSMYWRYKKTRYWGYRCY
46                  RMYWCKMYMYRKKKACKRYRAKATKKKRMKAYKMYWCTKTGRYSMCWRYKKYGCATCATT
                         ** *  .**  **** * :***  *   .      *  **** *****

40                  GGWYRCAAGKTRRMYRGCKYYYYGGGWYRRWSRRRSMRCCCKRYYYYATCRYYGYRTCCA
46                  RKWCGATTRKWRRMYRRYKTYCYRKKWYRGWCRGACAAYAYGGYYYYMYYRYYRYGAYSR
                      *   .:: * *****  * * *   *** *.*   .    .    ****   *** * :  .

40                  AWKYCRYRGCSTCCCMWMYWYYKSSMRMATTTKRRKKKRGYAGTKMMYYCKYYCGKGKRA
46                  WWKYMRYRRYCCTTAMWMCWTCKSCMRCGCCKKGGGKKRKCGAAGMAYYYKTYTAKSTAW
                     *** ***   .    .*** *  **.**  .   .*    ***   ..: * ** * *  .*..

40                  TRRYCMMGKKSMYCYMGYTWKRRWRAKKKAKRYKCGGYTRKRATMRGWKWCTYWGGAMRR
46                  KRRCYMARKKSMYYYCRCKWTAAWGGGKKWKRCKTACYKAGRRCMRRTKWYGYWRSMARR
                    .**   *  ***** *    .*.  *  .  ** ** *  .  *.   *   **    **    **   .   **

40                  MKWYYKWRGTGGRYACGTSYCKAYKYYCYRGSKACGAYYRKAAARCYWWCYKTKWRYKTK
46                  MGACCKARTARARYWTSCGCMTTYTYYMYRRSKRYRRYTGKGGTRSYWAMCTGKTGCKCG
                    *     * *  :  .**    .  .   .:*.** ** **     *   *..:*.**     .  *    *
```

```
40      RGKYRGTTTRAGMTKKMTYRGWRRYKRWATCTGYMGWRMKAGAGKAGAAARSRRKATGMM
46      ARTCRRKYARRSCGKKMKTGKWGRYKRTCGYCSYMKWRCGRKRSGRCRWRGCGAKCCACC
          . * . :* .  ***.   * **** .   .** **     .       .  *. .

40      WCKGTTYMKCRRAACMATTGKWKKCKGTAYWKRMWYYMKACYTRKRRRKMRKAATGKWKK
46      TYKRWYTCKYRRRRTCRWWRKWKKSGKKRCAKRCAYYMKCTYKRGARATCRKRRYRGTKT
          *     * **        ****.  .   **  ****. *.*  * . **      *.

40      YYRRKAKCWKKCTTCTKGTCYKKRYWMKGACCTATTKYYYWRYYTAKKKYCCCKRKTGGC
46      CYRRKWTYWKKSWAAAKRYMCKTRCWMKAGATWMYYKYYYWRYYYTKKKYMMMKRKWSSY
         **** . ***. :.:*    *.* ***...     ******** :****   *** ..

40      YSYRYYYYTRYKKWCMGRKTTKYRKSKTKKWRYRKMKSKRKKYAKTKTKRYKYKWAKCKA
46      YSYRTYYYYGCKKAMMRGGKWKCAKGGYKKWRYRKMKSGAKKYWKYKGKGCGTKTRKSKW
        **** ***   **  *   . *  *.  **********  *** * * *    *  *.*

40      MMMYASTKRCAACWYRWYYRAMMRTRYRRRMKKCGGCTGRKMRCCCGGMMMTAYKMKYWK
46      MCMCRCYTGYWWSWYRTCYGWMMRYAYRRRMKKMRRYYRAKMGMYYSKCCMWRTGMKYTG
        * *  . .    .***  *  ***  *******       **    .   *    ***

40      MYMYRWKTTACGWRAGYYRKCACRY
46      AYMYRTTYCRYRWGRKYYRKYWTAY
         **** .     *   ****    *
```

The pairwise alignments of two accessions 40 and 47

```
40                      KYMYRRRKAKYMCGRRRRRYRWCCYRRMGWSCMMTMYMKRWRKRWYYGTTRAYWMGYYAK
47                      KCAYRGRGWKCATAGGRRATATYYYRRMKTSACMYMYMKRTGGRTYCSYWGRCTAKYYMT
                        *  ** *  *    . **      ****  *. * *****   * * .        ** .

40                      MRTCTRYMTRKACRGYYYYMTCKWGGRTYGSKYYYKMYRWKYRWKYWYTGKYRRSYRYKT
47                      CRCACRYMKAGMTATYYCTMYYKWAARWCKSGCTYGMYRWKYAWKYWTYAKYRRSYRTKY
                         * . ***.      **  *  **..*   *   * ****** ****  .******* *

40                      RAYYMWRRTYKARRWAMYKYKRGYYKCRRRKKKWRGGYRKRAYKARMRRYYRTKCYWRYT
47                      RWTYMTRGYYKRRRAMCYTTGGAYYKMRRATKKTRAAYRGGRYTMRAAGTYRYGYYWRYC
                        *  ** *  ** **   *.   .*** ** .** *..**   *. *    **   ****

40                      RCARYYYRYWMRYMYYYRWKCYGCMGYYCAYGKWRRRRYYGCKYGSWYRKWMRWKKWAYY
47                      AYRRYYYGYWCACMYTTAAKMYASMKYYYRYRGARRRRTYRMGTKGTYRKWMRTKKAWCY
                           **** **   **    * *..* **  *   **** *     . ****** **   *

40                      RATACMYKAYKRKRGGTRRMMGWYYCSYTWKMSRRTCWCKRYMRYRWMRRRGYKSWGYKS
47                      RWKTTCYKRCGRGRAAYGGMMCATCACTWTKAGRACYWAKRYMRYRAMRRGAYKCARYKG
                        * .:  **   * *..   **    ..   * .*   *.******* *** .**.  **.

40                      MYTMRCYKYRSRGTRTGYRYGYTYRRYGWYWKRYYWWWTKTTCTMKAAKCGCGWYKRAWC
47                      MTWMGYYKYRCGRKGWRTACRTKYRRYAWYWKRCYWWWATCCMYMKGCKYRARATKRMWY
                        *  *  ****.  .        .****.***** ****:.    **..*  .   ** *

40                      ARRMYRRKGKGGAYMKKRCWKMKMKGCYYRRYTMGYCYMYRWWGRRYYYKYRTWYKRMMA
47                      RRRMYRRKKTAATYMKKRYWKAKMKTTYYRRYYCKCYTMYRWWRRGCCCTYRGWTGGMMR
                         ******* ...:***** ** ***  *****      ***** *    .** *   **

40                      GGKYWSRWYMYCYYRWCSRRRRRRCGWRKKYKMYYMYKYRYYCGCYRTTAKGRWYTRYAR
47                      RRTTWSGAYMYYTYRWYGGRGRAATAWRGKYKMYTMTKYRTYYRSCRGCGGAGTTGGYTG
                          . **  ***  *** . * *   .** ***** * *** *  . *  . .     *:

40                      WTGGTCCRYTMTTYMYRMWRMRKRGSTMTAGCWYRRKAWAWARSMYACTYCRMAMTYKKT
47                      ACARCYMRYAMCYYCTAAAGAAKRACYCKRKMAYGRKRTWAWRSMYTTAYYRMGCGCKTK
                          .    **:*  *        **..  .    * **     ****: :* **.   *..

40                      ATRKYCTYTMYGKAYYTTKTCGYTYWRSYTGCTTMAAKCRAAAYYCRYTWKCWAWKAGTG
47                      WKGKTYYYYACKKGYYYWKCARCWYTRCYYKSYKMRMGYARGGYCYGCCWKTACWKCCCC
                         . *   *    *.**  * .   * *.*  . .*      ..*     **  .**.

40                      TAYWTRAKKYYRRTTMTRCTGSWGYKAYKRRRRKGYYYYAGMRKTYTTTMAGGTYKWKYR
47                      YWYAWGWKTCYAGWYCKGYKRSWRYKWYKRRRRGTYCCCGAAGKKYWKYMMAAKYGTGYR
                          *    *. *     .  . ** ** ******  *   ..  *.* . * ...*   **

40                      RGKRGKCAKYKRCRTTGRRRMGAMKAGAYCCAKSRYMRTTKKYCCGCCKYCCACTYYTGR
47                      RRKRRKMWKYKRSRKYKAGAMAGAGGATTYMMKGRYARCGGTTYMKTYKYYSWAGYYAAG
                        * ** *  ****.*.     *..  ..:    *.** *   .      ** . .  **:.

40                      AGRRMRRKTYGTATTWWRRCATAKAACMWKTCACACCMWTAWCCMRYWSGYWRYWKCWSK
47                      MKRAAGRGYTKKRWYTWRRYGGWKRMMMWGYTWMWMSMWGWTATAGTWSAYAATTKSWSK
```

```
              *   *    .    *** .  *   **       .**   .    **.*    *.***

40          RRRWATRTTYGRMMKTAGGWGYRYRWRYTCKYCMCRCRRRTTTTMMRKYYAAKYRWTYCW
47          GGRTCGRWWYCRAMKWWRRTAYGYRWRYGGKYYCYGMRAAWCGKACGGCCMWKTRWATGT
              * . *  * * **     .* *****  **     *     .        * **:

40          RRMYRKCTYYKKAAATTGMWGAYCTATCGRWSKCRWKKGWTRAGKSGKGYKMYCACMRYM
47          ARMYRKTCYCGKCCCAATMARWCMCGAYRRWGKYGAKKRWCARAGCRTTTKACTGGMRTC
             *****  *  *...:: *      .:  **.*   ** *   . . .  *   . **

40          AAGRKYMTYTGTYKTKGWWMGSYYKYYYRWWWYTMATAAACKMTTARKRTARYYYKAWRY
47          MRKRKYAWTYRCTKYKRTTAAGTCKTYYRTAACCAGAGTWAGCYWRAKRKGRTCCGRARY
               ***       * *    ..  * ***      .:.: .      **..*      **

40          AYGKKGRKGGTWGMWKGYYKWAKRRCTKTYRGAYRKTRKAYYYCKKRYMKRRYRTGCYWK
47          RYRKKAAKAACARAAKKCYKTRKGRTCKCYATGTRKARKWCYCGKTRYMTRRYRGRYTTK
             * **. *..     *  **   * *  * *  . **:**  *  * *.***.****   *

40          AARCKTYYRAYKRYAAKYWKGYKWCGGRRYCWCGAGAMCTTWKSKKCYMAMCYTGAMRCC
47          RRRYKYYYRRTTATGRKCTKRTGTYRRAGCAWYRRRWCYKKWKSKTTYCRAYCYRWCGYY
              * * ***  .  . *  *          .*       ..****. *

40          RAAGAAWMSCACKAATRTCTKAGAAKYAMRKKTYGAGGYAYRRKAWMKYRWYRCWAKGTK
47          RCGKCGWCCYTTKGTCRCSYKMAGGKCRAGKKKCSWKRYWYRRGGWMGCGWYRTWRKRCK
            *.. ..* . : *.: * . * ...*    **. .   * *** .**   *** * *  *

40          WKKGYGCGCKYWWRCAKGYMGCGYTMCKCRKKGGGTMYAYRAKWYYYGTYRTACRYKTAC
47          AKKACAYRMKTWWRMMKRYMKTTCKCMKSRKKATACACTCRMGACCTAYTGYWYRYKGWT
             **. .   * ***  * **    .  *.***. .   : *      .      ***

40          KTRRRGTTWYGAKYYYTGKKKTGYCWMYRCYRSRRTCGCARKRKGTKTKKYCMYRYRRRG
47          KARRASKWTCACKYYYCCKKGCKCMTCYGMYRGGACMRYTGGAKACTAKKYMMYRYRRRR
            *:** ..   ..****  **       *  **.      :   *. ..:*** *******

40          CMKYAYMRTRCKCKCGKWYGYKYRGCTTYRKSMTTKTKCMYRYATRRYGRYKRAGGACGKG
47          YATYWYAAYAYKMKCTTYKYGCGRYWCCGGGCYYKWGYATRYRCGGCRRYKRWARMYAKA
              .* *     * * . * *          .   *     **      **** .   .*.

40          AATGYGGCYCTTTTCTYYMKKTTCTCYRYCCKGMMTTRGAACAGRMSYKTYCGATKCKRW
47          MCAATAATYSWYYYYWCTMKKYYYYYYRTTAKKMAYWGRTGGRRAMSYKYTYKWWKYKRW
             .:. .. *.        ***     **   .* *     :.    ****      * ***

40          KACMCATCYWYTKKRYYKRWCCAAAAKKGYKWKYTCMRYRTSACTAGGAWKMSTWCKKKW
47          KRYAYMKTCATCKKGTCKRWGYRGGWKKRTGWGYGYMGCGCCGTYTAAGWKMSKAGKKKT
            *     .     **   ***   .. **   * *  *    ..  :...****.  ***

40          RYYCTYACGRMSWWKKSWRYTCCTCYKYYKACTGRKWRYAYACCTCCTAGTCGAGTTYYR
47          RYYTYCCATRCSWWTGSWRYAATCTYKYYKCYCTRKWRCWCTATCTTCCAWTARRAGCTA
            ***   .. * ***. ****:.   *****.   ****   :.     ..  .  :

40          WRKGMAYTKYRRTKCSTGYKGCGAYKCYRYGASWKTAGGAYKYRAAARGCWCYKKKWWRY
47          ARTAMWYCKYARCTYGYRYKAAARYKYYRYKMSWKCTTARYKTGGGTRRSAMCTKKWTGC
             *..* * ** * . .  **... ** ***  *** : . **  ..:* .   .***
```

```
40          TKKKGAATTTTTRYACTMMKKMKKGRWMRKTKCWRYKKMYRYKRAARSRSRKAKKGTTYM
47          KKTGRCGAGWYYRCTYAMMKKMTGTGWMRKYKSWRYKGMYRCGRRWGCGSRKMKKRWYTC
            .*.  ..:    * : :*****.   **** *.**** ***  *   . *** **

40          CGRKKMRAATTGKCKGTYWACGRCTCYYMKYYRRACYRRCGRCSKRRGYRKMRAATGKWK
47          YRRKKMRRRWWRKSGKKCAMYRRYYYYYMKYYRRCTCRGTAAAGGARTYATCRRRYRGTK
              *****     *.  .     *   ********.  *  . ..  * * . *      *

40          KYTAAKGRKKAKYCKKWRGKACGGYCKYGARKRWSKCCGSYRGTWTRWYWKKSKCGTATC
47          TCCGCKARKKWGYYTKWGAKWAKKYYKCTGGKRTGTYGACCRACTKRTYAKKSKARYRWS
            .  ..*.***  * .** .* .  * *  . **  ..  .. *.  .* * ****.    .

40          KAWYRYRRWAYYRYKGACYMWYYCYRGKRTAGACCACYYKGSSCWGTYYYYCGRMYAWYY
47          GMATGYRRTGYYRYKRRYYMWYTYYRKKRYRKMMYRTYYKKGCGTRYCCTTTARMYTACT
                 *** .*****   ****  ** **        ***  ..         .***:

40          ASYRRMKGRATRWTKKKYMWKYYCCWYTGKYYTAWRKAWKGKKKKACKWRWRKATGTCGS
47          MSYRRAKKARYAAYGKKYMWKYTYSTCCTTCCCCARKRWGRKKTKGTKTAWRKTCAKATS
             **** *        *******  .    .   . ** *  **.*. *  ***: ... *

40          WKGWACGGWRTACTGATARAAATTTARCKTWCTACTYRMTYRKKCKCCCWRCMYRRTGAT
47          WKRARYKKWRCCYYRGATACRRKWGCRYKYAYGGYCYRMCCRKKYKMYYWRMMYRAYTGW
            **      ** .   .:: .  .  .* *    .  ***  *** *   ** ***   .

40          RKRYKRYGAATTTTAKYYYCGTWTTKYMRRYKGSYWRYRMATMYYMYKGTRAKKTKMRGY
47          RKRYKAYACCGYYWMKYYYYAGTGAGCAAATKASYWRYGCRKMYYMYKRYRMKKCKMRKC
            ***** *...     ****  .   :      *.*****   .******  * ** ***

40          CCMMYYKKMSRYRRKCYMTRYKYATCCCRSKYYYGTTYKWGCCGMCKYGGGCKKKMYRYY
47          AACCYCKKMSRTGGKMCCYGTTYMYTMGAGKCYYKWCYKWAATAMYKYRRRMKGGCTGCY
            ..  * *****   *      .*      .* **   ***.. .* **    *      *

40          YRWAWAKWTWKYKRKRKCCYAYKYTGKSWGCGCCMRYTMMWKSMRKYKGYMYRKRYSAKR
47          YRWGWRKTAWKYKAKRKTTCGCKYYRGGWKYKYMAACKCMWKGCRKCKKCCYRKGTCGKR
            ***.* * :**** ***   . **   .*        . ***. ** *   ***  ..**

40          WKMMGGTYGAGMYRTRYYSKGTMKKRRAGKGRAATRRTGKYKYRRWYMWRYSYWYWMRKR
47          TKMMRKCTRGKMYRWRYYGGSWMKKGAGKGRGRMKGAWTKYKYRGWYCTRYSYWTAAATG
             ***     . *** ***. . ***  .      .    ***** **  *****    .

40          KYGGAWKKWYYMGKGACCKKYAAGCRWMTGAGTAGMARRWWWKMKGAAKWKTKMYKYTAT
47          GTRSRWKKWYYMRKSRMATKTCCCYRWMYRMAWRAMRRRWWWKCKKRMGAKAKMYKCWRK
               . ******* *.  ..* ..  ***   .  .* ****** *     *:****   .

40          CATTCKMTATAGKYTGYGKMSWTATTACRRAKRYRAAGTGMRRYMRYKRGAAMTTRKSKG
47          TGYWTGAAGWTKKYKRCCKMSWCMWKMYRRWGGCGGTRCKCGGTMGYKGTGGACGRKSKA
             .     :. : **.   ****   .  **     .:       * **  ..   ****.

40          GCKAMCCYRTGRYWGYCTCTGTGKAKKRWMWAYGKTYTCMRWKTYYSCACTTAARTCTCG
47          SYKRMYTYRARRCAKTAWTYRCTGCKKAWMTGTKKYYWYMRWGGYTSAGGAACGAYYYYK
            . * *  **: *    .       .** ** .  * *  ***  * *.. ::..
```

```
40              RMRMWKAYYRSRYRYYCWTCCCWYRKRCCGYYGYACGTKRGGAAAAAYMTKTCCCMMGGC
47              RMRATKGYTRSAYRYYYWCTATWYRKRTGAYYKCMMRYKGSKWRRGGYCYKYYTTCMRKY
                ***  *.* ** **** *  . *****  .**      * .    ..*  *     *

40              WRMMYRCCTKRAYRYYCGSRWRYAAAGCKARCTTYYKRYRMGATATRAKYAYRAAAKKRW
47              WGMACRYYKKATTRYYYRGATGYRRWRYGRRMCKYYKGYRMRMKRCGGGYRTARRRKKAW
                * *  *  .* : ***  .   *       *  .*** ***  .   . *      ** *

40              GMTSMKRYGKGKAWRWGRYYWAKRRTWYRCATGTCGWYRYYACATCAKWRYGCMRYYKYY
47              RMGGAKRYKKSGGTGTRRYYWRKRGCWTRYCCRATRTCGYTWYGCTGKTRCRMMRYYKCC
                 * . *** *. .    **** **  * * .  :     *   .  .* *   *****

40              GYTAAAAWRYGRSGWYRRYYRMGAYYKCACTGAARAKAATCATKRRYTAGKCKACCAATT
47              RTGTCMTAGTKAGAACGGYYRCRRYYKTTTAKRMAMTMTKYMYTARCYRKKMKMYYGTKY
                   :. :     ..    ***   *** : :     . :.   . *    * *   .:.

40              ATATRYGGTTCGMKRKAKRRYKKYKTAGTACGKTGYYSRYKAGTYWAYKRYRTCGKMMTT
47              RYMYRTKKCCTAMGAKGTAGYKKTTWRKWRYKGYAYYSRTGGTYCTTYKRYRYMTGMMYY
                    *      .*  *..  *** .         .****  .    :*****    **

40              GARRCARRRRGKRATTATYKRTGCKCKAGAACRYATTAKMYYYSGARWRKKMGYCAYCWA
47              KTRRMRRGRGRKRGACGCYKRGRYKYGTRTCYAYWYCRKCTYYSRMRAGGTCSCMMYMTC
                 :**  * *  **.: . ***   *  : :.  *    *  ***  *   . .   *  .

40              AGGKRRYMGGWRRRRGTTTRAKTTYRTGTYSCTYYKRMWAYKMGATAAGAMKCTRCRSYK
47              TKAKRGCCATWRRRGCYYCRRKCKTRCRKYSTWYTKGAWGTKMRMWMRKTCKYYATAGTT
                : .**   . ****     * * . *  .**  * *  *. **      : *     . .

40              YWYTKYYRRGAGGCAAYKRWKWGWYYYSTWRRRRMACCMGCTTAGCKTKYKWYAWTYGAA
47              TATWKYYGRAGKAYTWCGGATTKWYTTCWWRRAAMWMTARYYGGCAGGKTGTCWTYYTTG
                    *** *.. . :     .  **  . ***  *        . .  *       * :.

40              TCTYSRRGTGRSMRCKYYYATCTRTYYKRRYRCCAAKYRKSWMWTTKMYRYGSTSKCCRK
47              CACCCRGRWRACAATGYYYMYYAGYYYGRRYGYSRWKYRGSWMWGKKMCRYRSKCKTTGG
                 .  .*     .    ***   :  ** ***  .  *** **** .** ** *..*

40              KRYMRTGKCGYYGATGYYRKRGGTRMCMSYKMYWRYMGYTRRWKKCKGKRMYKCGGTRRR
47              KRCMRCTGTKCYAGATYYRKAACKRMYMGCGMYWRCCRCKAAWKKMKRKRMCKTACKAGA
                ** **      *..: **** . .** *.  ****    .  *** * *** * . .

40              KMRAWMRGTGWKTYWYKWYYKRCGCGACYYRRSWWKWAKKRKKRACTARRTCKCTGGYCW
47              GARRWMRTGRTKGYWTGACCKRTAYKRYYYRRCWAKWGKKGKKRMSKRRGCAKMYRTYYA
                  * ***    * **     ** .    ****.* **.** *** .. *  .*    *

40              GRMTMYATCKAATYMTGGAMCKGKKCAGCCTGTRYYYKCCMGGTMYCCMMKCCGKRMRGK
47              KRMCCTTKMKTGGTMKRRWMYTAGKTCRTGARYRYCTKYYCKRCCYYYMMKGSKKRMRAT
                 **   :. *:.  *.   * .. * .   :  **  *       *  ***  . ****..

40              WWTCARGTTTAYCTTGATKCGGWMRRKKMYSKTKCMKCTAGAKYAACRWMRMKTKCAKTS
47              TWCTWRRWWKTTGCGTCYKYRKWMRRKKCCSKGKYMKMYRRMKTTGTRWMGMGAKTTGCG
                 *   *   .:     . *   ******  ** * **     * :. *** * :* :  .

40              RTYYYYAKKCRCWKKRYYGWWYAKKGWYCKWYCYRATCTCYKSYYKSGRRSSSMKAMMYR
```

```
47          ACYTTCCGTYGYWKTRYCRATCRKGKWYSGWYYYRWYYKYCTSYCGSKRGGGGCGRMMYR
              *   . .   **.**      *  **. ** **   .  .**  * * ...   ****

40          RTTYGTYYWATCTTTAGTTGTRYRRYGAYMKYCATACGYRRRRMYAAMRMTARGAKKMYC
47          GKYYKYYYACYYCAGGACCACACAACRRYMKYYMCGMSTAAAGAYGGMRMYRAKWKKAYT
             . *  ** .   : ..  .        ****   . .      *..***     ** *

40          AKKWYARRWRYYTRTMAGYKMMYKCAAYYKKRKYRYGSWTTYKMRGGKYSYAAATGRCRA
47          GGKWYTGGWRYYYRYMRACKMMYGTCMYTKKRKTGYSGTKCYKMRRKKTGTRWRYSRYGT
            . ***:  **** * * . ****  . * ****  *.. . ****  * .     .*  :

40          YRYWTTCRWMYYYGYCTKKRYKMCYYCYTKKGYYRWMCKWKRRGWKGGKYYYKYYCMWWK
47          CGYWAGGRWMYYYRCYAGGGCKCACYAYYTKAYYRWMMKTTAARWGRSKCCTGCTYMTTT
              **:  ******   :    * . *.* .*.***** * .   *  .*       *  .

40          SRKCRRKAYTKCTGGKRKGGRMATRYRRAYCCKCKKRTRAYRYYRWCTTYMCCCWRRGCA
47          CRKMRRGRYYKYWRRKGKAKRMCWGYGGRYYMKAKKGCRMYACCGTTYCTAYATWRARMR
            .** **  * *    * *. **.  *   *  *.**  * *           . **

40          KKMWYKRTAKKYGMRYGGTCYRYKKKKKAWTACGARKWKAARTYGCCCTTTWMRSRRKKW
47          KTMWYKGWMKKYRAGTRRKMYRYTGKTTMWYRYKRRGTKCGRACRYAYYAYTARSRRTKA
            *.****   ***      . ***.*..  *     *  *..*:   .  :   ****.*

40          SRAYWGWYKRTGGYCCTASWGYRMRRWKMWYMKAAGGACCYGCKKMAKYKGGCSRRATKG
47          GGWYWKTYKRWKRCTACGSAKCAMRAWTATTMKTGTAGTYYTSKKMMKYKAAASRRRYGA
            .  **  ***     . .*    ** *.   **:. ..  * .*** ***...***   .

40          TGYRYKWTYRRCRSYARYRAATWATAKWRKRGACAYGKTGTCCYKMYKGKCATTGYYRRW
47          CACGCKTYYRAARSYRRYARCAWTAGKWRKGRWSWYAKCAAMAYKMCKAKYGCYCCYARW
             .   *  ** .*** **  .:*::.****   . *.* .: .*** *.* .    * **

40          KTKTKTCKRRMTKMGGKGGMMKYYKKCGGMYRYAKWRTCTWRWYYYARWKKYTTTGAARS
47          KGKATCYKRRMYKCARKRRMMKYCKKTTACCATCKWRKMAWRTTYTRAAKKCKWWRWRRS
            * *:.  **** * . *  **** **  .    .***. :**  *    ** .     **

40          YWRTWKMRAACTMTAYTMAMYTAGMKYWGCGCKKYWWSTTYWYKKRCKTKKTAAGTTRGC
47          YWRCTGCRGGTACWRCCCWCCATTCKYWRMSMKKTWWSKYTATKKAYKYKKKCGACYRAT
            ***    *.. :         ::  ***  . ** ***.    **  * **....  *.

40          YRRYMYKGWYMYRRKYACGAAYCYMGYGKGGYYWRYKAKTRRWMATKAKTSAMCCTYRCW
47          YAGTMYKCTCACARTYWMRWRYTCMRCKKRRCCWRYKGTCGGACWKKGGKSRAYYYCAYW
            *   ***      *.*     *  *   *    ****..      .*. .*        *

40          KKMGAGGGCCACYCAYGSKMCMTTYKWYRYRKWAKGARKWKYRRGYSMYWRYKKWGTKCT
47          GKMRWKKAYMGYCYWCTCKMYMWWYKACRYAKATKKRGKATTGRKYSMYWRYKKWTCKSW
             **    .  .      .** *  **  ** * :*   * .  * **********  *.

40          KKYYKRYKWAKGACYYYTTGYCKKKCKKCKRKKKRSYYYYYTYAAGGKWKWWRAYWCMCK
47          GKYYKRCKWRKKGAYYYYKTTGKTKMKKMKRKKKRSYTYYYYCWGTTKAKAARCYWMMYK
             ***** ** * ..*** .   *.* ** ******** ***   .  * *  *.** * *

40          RWGRTMYCTTKTKGYRKSTKKWRYRKGTCTCKSKTARKKACAKRTARYKMACTWAKKWGT
47          RWRGCMCYWATGKSCAKGYKKWRYRKKYMWYKSGWWAKKWMRKGGTGCKARYGTRKKTSG
```

```
            **   *   :. *.  *. *******     **    **   *  :  *      ** .

40          AMMMYRCASKTGATCCTKRCTCAACWYWAMMMRYRMKKYKCTGYGRKMRMGMMTATKWKK
47          WCMCTAYRCKCAGWYTYTGMKYWWSWYTWMMMAYRMGKYKYYRYKAKMGCRCMWRYKTGK
              *     .* ..    .  .   .** *** *** ***   *  **    *   *  *

40          MYRKACGWAAGRWYRAYGMAGAYCRCYRCAGKRAAGWRTATAGCGCGKGYCAWWYYGGRK
47          MYRKRYRWRWARWYGCCTCRKWCYGYCRYWRKRMMRTRWTCCAAAMAKKYYRTATTKKRK
            ****   *  .*** .           *   **    * : .... .* *        **

40          AKAKTSKRRTWWKGYGTRGSYYCWKGTRWGYRGGKKATCYMTTKACCYKYCTCSKYYTAK
47          GKGKKSKGRATWKRYRCGRGCCYTKKGRTRCGAKKKWGAYMCCKGTTTKYYCTGGYYGGG
            .*.*.** *: ** *    .    *  *    . **  .**  *.   **   . ** .

40          KYRRYSRRYKKSTCRCCCTSRRYTCCTTKWKYKTGGCAGWAWRYRGAYKCTKYRKSYKRW
47          GTGACGAACKKSYYGATTYGAATYMYCWKWGYKACTTRSAMAATARTTKAAKCAGSCTAW
                 .   ***   .   .        ** **:    .       : *.:*   * . *

40          TTCTYYYRYMTATYKYKRYRRAAGMKKWKTCAKKRCTGAWYYGTYCCGTKTYYKRTRCAA
47          YWYWCTYRYMCWYYKYGAYRRRWRMTTWGGAGKTGMCACTCTKYYAMAWGCTCKRYATRM
                  ****   ***  ***   *..*  ..*.   ..     *. .     **

40          KKCCTTTGATTCKYYRKRGCACAGGCTGMYYYYYYACKRGYYSYARTCYCRYGYKWCTCG
47          KKTAGWWKCCCTKCCRGARYRMMKSYCAMYYYYYYMTTRRYYSYMRCTTYRTRYTAAYYA
            ** .    .   *  *        .  .*******  .* **** *    *  *.  .  .

40          TCRKRATYYRYYCMRWYAAKRYRYGYSWWRRYTAYYYWMYYYGAMMAKYMMYTTAAKWSK
47          WSGKARGCYRYYAMRWYTGKRYRYTCSWARRTYMTYYWMYYCRGCCRKYMMTKCGTTWCK
             . *    ****.****:.*****  ** **    ******  .   **** . .:.*.*

40          SRMMYMYTKWMRRRGGTRYYSGYYCCATMKATGRYKYMRYMMYMRRRYMGAMCYRYRYKR
47          SRCATACKTTMRRGTTCRYTCKTTTCCMGRCAAYKYARYMMCMGGGTCKRAMYRYRYKR
            **     .. ***    ** .     . *   . *** **** *         *******

40          CACTTARGTGYCCAGATWATGTTCTTGYCGKKYYYRMARRYYKCYSYRRYGMKRRKYKRY
47          MMYYYRGSYRCYYTAGGWCATYWMKYSCYKKTTYYGMMGRYYGTCSCGGCKMKRRKYTGY
                   .     :.. *.:    . .   *. ** *  ***   *     ******. *

40          AKKWYMYYMKWRKRRRMRRKGRGCGAKMMAKTWGAARTATYYRATCMKKKKGKCGWGKCC
47          MKKTTATCCKWRKRRGMAGKKRRYRRGAAGGCTKRWRCTATYRGCTCKKKKRTTRWRKYT
             **      ****** *  * *       .      * :: **.   **** .  * *

40          TWCRGTTCSYGTTCKGAAKRSKYCWKAGTGSGMRKTAYRSRCYKYSYWYKKSCWMKMCTA
47          CTYGCAATSYRWWYKKMMTRSKYTWKMRYRSRMRKYMYRCATYKYSYWYKKSTWMKCMAR
                 :: **    *   .**** **    * ***  **.  ********** ***  :

40          YMYYKYAKRKYGAKGGCGAGKGGRKGWRYCAGGKRAMKWYMCTTGTRYACAATCACTG
47          YCYYGCGKAKYKMKKRYKCATAKRGKTGCARRKKAMAKTCAMYWCWRYRGTGCMMYKR
            * **  .* **  *    .... *     .   *   *        **  :.    .
```

The pairwise alignments of two accessions 40 and 48

```
40              CYRYMRWRYMYYKCRYRRMWRWACCMMYMKWRKGRYGTYYKYRGYSWKKKGKYRWMRWGK
48              AYGCARWRCATCTYGYRRMWRARYACMYMKTGGKRYAWYTGTGKTGTKKKKKYRAAGTAK
                .*   ***    .  ******   . ****    **. *      . *** ***    .*

40              AYRMMTKAYKWRYKRYATKYYRGRYTAKSRWCAACWRACCCSWWYRYYWGSRYKYMRYGC
48              WYRMCGKRCTTATGRYRYGYYRKRCATTGRAMRRAWAGATTSWWYRYCWKSACKYMRYCT
                 ***  *  .    **   *** * ::..*    .* ..  ****** * *  *****

40              YMYRYYYWYKAARRRRRKTGRRKCMKKCMRYMYSRKMRYMKARMGCKRGSRRGAATRGYY
48              YMYRTTYWYKGCGRGGRKAAAAKYMKKACGCAYSRTCRYMGRAMKATATSRRRGGGAACT
                ****  ****.. *  **:.  * ***.    ***. ***   * ..  *** ..  .

40              WWRMRTYWCGTAKCYAYMWRYWGSRKARRAMKYKRYKRRWRKYYYCCMYRYMWMKKAGGY
48              WWGMRWCAYKWGGYCTYMWRYWRSAKRGRMCKYKRYKRRTATYCYYYMYGTMWMKGRRRT
                ** **      .   :****** * *  *  ********  .* *  **  ****

40              YYGYKRKTKTCTYYCTCWSWYRCYWMKTRRARYKWYRMRRWRYRYYRRKMYGYCYKWYTT
48              YYRYKGGAKCGAYYTAAASWYGYYWCGKGRMRTKWYRMRRTRYRYYRRKCCRTYTTWYYY
                ** **  :*  :** :. ***  **  . * * ******* ********      .**

40              AKTRYCARSKKKRMRKTTSWYYYYRYMTTRRYRGRKMYKMMYKSRTGYRYYATTRTRRYW
48              RKWGYSCRSKTTRARKKGSWCYYYRTCGYGAYRRRKMTKMAYKCGWRTACTGCAAARRYW
                 *  *..***..* **. ** ****      ** *** ** **.       . : :****

40              GCRRCYYSYWWKYRMTYYAGAGCKWATTTSYTCRYYKYGYWYWRWCGSKYRRYYMKKMCW
48              TTRAYCCSYWWTYRMKYYRAWKTGARWWGSYAGGCYKYAYWYWRAMKCKYAGTYMKKMYW
                  *    ****.***.** .         **:   ***.*****   .**   ***** *

40              KYGGWMRGKYRYMAYWKYRWYMMYRCYKRKRGTGTGRTGGCMCAAYKMGRKYYWRWYGCA
48              KTKRWMRTTYAYMTYATTGWYMMYRYTGGGGRYACCGWKRSMSTGCGMAGKTCWGACAYR
                *   *** .* **:* .  ******        .      .*.:.  *. *  *   .

40              TCGRKWYMYRMWRAGKTCYCMKMYTWATTWGTCYWAACYCKRGKATYMKCGCYWRYRMRA
48              CTKRKWYCTAAWRTAKWYCMCKCYGAGGCTTATTAWMSYTTGRGGGYMKTAACWRTGMRW
                   ****    **:.*     * *  .    :     .* .   . *** .. **  **

40              TWCYTTTGTTGAKRRMRRYYKTTKYMYWARYYYRGTGCTCTMARAATYYYRTRCAMYCYK
48              KWMTYCCTYWRWTARAGRYYKYWKCACWWRYYYGTYKSYTKMRGGGCYYCGCRTCCYYYK
                .*          . *  ****  *   * ****    .  .*  .. **   * . * **

40              KKTWYRYAGTAGGAMTRRYKYKKAGRCGAAWGCYYKRRAATRRYRCYACAACTKRYGMAY
48              KGWTCGCCCYWCAWCAGACTCKKTRGYRTGWRYYYKRRRWWRRTATCGTRGAAKRYKMRT
                *      .    .  :   . **:    :.*  *****   **    .  ..:*** *

40              RYRYYYKSKYRAGYRKMKRRMTCKCRRAMKAMGKRWYRRGCAGKCARMRTKKKTKRAYYG
48              RCATCYKSTCAGAYRKMKRRACGKGRRWCGGCAGRTTAGRYMRKMWAMRKKKKYKGWYYR
                *    ***.  ..*******   * **   . . *        *   **.*** *  **

40              GCKYCWCCCTYAKSCARYRGAWSKRCGSYGATATWTRYKTCGKSATAGKAWYTRYRMRYT
48              KYKCYWSAMCYGGCMRAYGKCTGTGGACCARCGYTKRYTKMKKSGGRRGMATKGYRCRTW
```

```
          *  *..  *. .   *  . ..  .. .  .  .**..  **.       . ** *

40      GCKACYMWYKYGYRGKRAGKAGAYAYYAKGYSTCRASWGWYYYYGYRMYSYTCGTGMWWG
48      SYKRYYMWYKTRYRKKRGTKMTGTRYYTKKYGATGTGTRTCCTTKYRMYSYGTACAAWWA
        . *  *****  ** **. *  .  **:* *.:  :.        ******  . . **.

40      TCCATGGRGYKCAARKCCWKRGRKAWKGCKKCWWATTGSWKGWAKTCTTGAGTCYWCTGA
48      ATTTYKKAKCKAGGRKYSTKRTRKRWGRYGGYTWTCKASWGRARKWYYCACRWMYWYYRC
        :  :      *...** . ** ** *       *: ..**    *    ..  **    .

40      MAATTKCGCTACKTACMMWWYAMKCRGAKKCCWRCMYRGACGRKWGKYTAYTCCGTGGGA
48      MRRKWKMSYKMYKYCMCCWWYRMKTGRCGKYYWRMMYRTGMKGGWRKYWMYCTYAYSRRW
        *  . * . .  * .   *** **   . *  ** *** .    * **  *   . .

40      RKYRWRWYGSWRYTKRCTWAMMMYKTRMAKRMAMGYCYRYKYKMSRYRKCCYMRYKCCCS
48      RKCRARWCASWRYKKGTGTGCMMYKCRCCKRMRAKCMYACKYTMSRTGKYMCMRTTTAGG
        ** * ** .****.*    . **** * .***     *  **.***  *   ** . . .

40      KYYYRYYGTTKWKCAYGGGCRCCKMYRYYYRWKYAGTTTWKKRWYYCTGKRRYGAYAYTG
48      KCYCGYYKWCKAKYMYRRRYAMYGCTRCYYRWKYCKAGCWKKATYYSWRKRRCRGCGYYR
        * *  **   * *  *          * ******. :  ***  **.  ***  . .*

40      SWGKYCGGCKRYYRMWRTMGCYMCYKKYMKRYSRWGGKWGTAKKRSKGTRKCRTAGKRRG
48      GWKKYYKRYKACYRMWRKCRTCCTCKKCCKGTCGTRKKWACGGKRGKSWRGYGWGRKGRT
        .* **    *  *****.       **  *  .    **. . **.*. *    . * *

40      KKRWKMKGWYWRKRKAWKKYYMTKTCCCMCCTAYKKKWYRRYYTKKRYMATTTYYRYRAR
48      KKRWKCKKWTAATRGMWKKCTCAKGAYYAMAATTTKKWTRRYCYKKRCMCAKCYYGYGRR
        ***** * *   .*  ***   :* .    .:: .*** ***  ****.:.  ** *  *

40      YYCCYWRRKKAACKRYRTAKRTGRYCTYRKTSTKTWCRRRYRGCKKWGWYKRGGRRKYGG
48      YYYACAGGKKCCATGCGKMKRCRRYTACGKWGCKYWTAARTRRMKKAKWYKGKKARKYKR
        ** .    **....   . **  ** :  * . * *   * *  **  ***    ***

40      KGTCRYKKCCTTSAKWRTRCGRAGRYKWKATTGGCTCTATWKYMCTTTRSRRKKKKWSRA
48      KRKMRYTKYYYWGGTWGYRYKRGARYTTKCACAAGYYYTYTKCAMYYYRSRRTKKKAGGW
        * . **.*    ...*  *  *..**. *.: ..    :  *      ****.*** .

40      GWSYKRGCYSMWYRMRRWKWYKAAGGACKCGYRTGRRMWRAAGTRYKARKARGGATGRRR
48      KTSYKRRYYSMACAMRAWTTTGTGTAGTKAATGYTRGATAWRRAGYKTGGTAATRCARRG
          ****  ***   ** *.   :. .. *..    *       : **:  : .   .**

40      TMYWTCKRYYRYRWTWKRGKKYKYRMWKYRYRSYSKYKWWWGCRMARYYWGRWGWRYGCT
48      GMYTYAGRYYRYRWKWKGRKKYKYRMWKCRCRSYSKTKTATCTGAGRCCTARTRWRYAYA
         **  . *******.**  ********* * ***** *       .*   .*  ***. :

40      CCTKYCGTGYYTGRYRKGTGRTKGKKWCCRYACYGTCGYYYYKTCTCAACAGYWMTTKWY
48      ATCKYMTGAYYCARYRKRARRYGRKKWGTRCTMYAWTACYCYKYTAYRTYGACWCKYKWT
        .  **   .** .**** : *   ***  * : *.  . * **  :  : .. * . **

40      CAYKSTRWGMAACYMKKKYWYCCCGGYCCGYYCKCTRGYSKMYKATGCWYYATCCRSRAA
48      YMYKGYRWKCWTTYAKKKYTCTGYRKYTTACYYKYWRAYGGAYKRWKYTYYGYTYAGGRM
          **. **   : * ****       *  . * *  *.*.  **     **.    .
```

```
40      MAACGRCYTMWTKYYTYCMWGMKKWSWCRWRYWGGRTWGCKGRTCTTTYRGTYMWTMWRY
48      ATGAAAATCMWYKTYYCSMWACGKWSWYRAATTAAACATTTAGGMGCCCRRGYMTKCTRT
         :... .  ** * *  .**.  **** *    ..     ..       *  ** .  *

40      RATCATCKYRWRYAKYYRKRRKWKMWTRKKGAYRKTWYKRRMRYSRRTGYYRRYRTRYWA
48      GMGYGYMTTGWGTGKCYGKGRKWKCACAKKTGTAKWWYKRRCRYSRRYRYYRRYRKGTTR
            .  .  *  .* * * ****    ** .  * ***** *****  ******.

40      ARCKYWRAYKRYAACRTYWYKWRYGTATACGRRYGYWKTYGRAAKCWYMRGRTKKCKYCY
48      RRYKYWRGTTATGGARKCWYGTACRYRKGTRAGCATWKWTRGWWKYWTARSGWKTYKTYC
         * ****. .  ...*. **       ..     . **      * *  *.  *. *

40      TCWGTTGAMYYGRKSAAGAAWMSCRCGRKCYRARYCGMRYKTKAGGAAGCKGRYRCWART
48      GGTAYYRWCYYSGKSCGRCGWCCYGAKRKTTGMACSKAGYKKTWKRWGRSGKGYATWRGC
           .     **. **.. ..* .  . **      .   **..    . .   *  *

40      TWMKARTCWCAKKARCKKGGGTMRKKAAWYYRAKACWWRYTARRRRAYGACWYWKTGKGK
48      YAMKRRYMWMMKKRRSKKATACCRKKTMACYRTTWYTAAYGWRAGAWCACYWYWKWRKKK
          ** *  *  ** *.**. .  ***:   **:.     *  *     .. ****  * *

40      GRTGYYRRGWKYWKTTYYYYTCACKAAYCKRRCKKCCTKRRSYYKYYYTRYKRKWKWWRK
48      RAWRYYGGRAKCWKYYCYYYYTGYGGGYMKGRMKKTAAKRRSYTKYYYYGCKGKAKAARK
            **    *  **   ***  .  ..* * * ** .:***** ****   * * *  **

40      MWAGMYWCMTKRWGRGGTMSGAKKKTWRYRKGKSWRACCAGCGCCKTRTAKRYKWAYKKM
48      ATTCMYWMMYKRWRGAACMSKRTKKYWRYRKKKSWAWMAGTTATMKYGGTKGCKTGYKKC
          : *** * ***  .. **  .** ***** ***   ..  .  *   :*  * .***

40      CMMYRRASTWYRWKAMMARRGKKWYKGGCTYGYRKMRYRYMMMTACYYKMYWKMMCGWMR
48      YMMTAGRCYAYRTKWMMWARKKKWYKRRTCYRYAKMGYRYCCMWRYYTGAYTGAMYRWMG
         **    .  ** * **  * *****    * * ** ***  *   *   *   *  **

40      WYRRAGRYYYCAKRCWKRYGYTTSRRTARTRRMGKTYMGGYTAYRGTTAYKAGCRCTKYY
48      WCGARKAYYYYWKRMTKGYRYYYSRRYWGYRRMRGKYAKTYKGCATYCGTKGRTRTCKCY
        *      ***  **  * * *  ***    ***  .*   *..      .*.  *  * *

40      WYWGKGYYCGCWRSTMYTCRGAAAAARMCWSAYAAKAKKRYWRMGRWCGGCRYTRGKTAK
48      ATTRKAYYTRTWGSYCTCTATTGTMMRAAWSMYCTKRGKRCWRATRWYKSMAYCRCKGGK
            *.**   * *       :.:  * .** *.:*  ** **  **  .  * * * .*

40      RTGGRWTATTGAYASCTTARGWMACTAATTGGACTARYMSKMCGARSKGGTYKRTTYAKK
48      ACACRWKGYCKCYGCAKKGRKWMGTWMRCWRRRTAGRYCSKCTATGCKAAATKGGYYRKK
          . **..   .*......* **.          :.** **  .: .*..: *   * **

40      KCTAGTTARKWKCWGKRATKRCTGRTAATCCKAGTYSMMWGYYWYKGACKMCGKGMKCAC
48      KSKGACCWRKWTYARKRRYGRMKKRCTTCYMKTRGTGMCTAYYWYKRWYKMYAKTMKYCT
        *....   ***.   **   * . * ::   *:   .*  .*****   ** .* ** .

40      RMWGTYGYKCACYGGCTMGGTTACCCGTTGYGGYKRAGKCWKRAYRGKWTCAMRYGTTTK
48      RCWRYCKYKYCTTRKYWCKRCCCYYYRCGRCRRCGACAKGWKRRTRATTCTTCRYRWWKK
        * *    ** .           .             ..* ***  *..   : **   .*
```

```
40      KATASSCGGRMYSMKTTTKMCCTYAKYKRWMRMKKKCAKTSRYYYMTYTAYTYYWKKARY
48      KRYGSCYRKACCSAKGCGGMYMYCMKTKRWMGCGKKTTGCGAYTTCWYWWCYYYWKTMRY
        *  .*.      * *    *     * ****   ** :  . *    *    ****. **

40      YYAWYTGAKKKKKKACKKCRTKWKCKGSGTAAAGCCTCTGYRCARKGGTCYRKKCCYRAK
48      YCMTCWRRKKKGKGRYKGYAYGWKAGACAKCWGATGATCTYRTCGKAAGGCGGKYYYRTG
        *       *** *   *     **. ..... ..  :   ** . *..     *  **:

40      RMYKGWYYRRCKTYAYYMYKCGYKCGYGACTGAGRCKKGGWYYWRYRRYKMKARAAKRSA
48      GMYKCTCCARYTKYWYYMYKYRYKTRCKRYYKGTATKKRRWCYWRYGGYKCKWRGMTASR
         ***     * ..* *****  **        .   **  * ****  ** * *. . *

40      MRYARKCCTKRCWKWRAAGCGTKYACTYKYKMMCKAGWYCKSRKWKAGAAKYKGGKYWKK
48      ARYGGGYYYKAYWGWRTGAYAAKCRYKCKYGCCMKRRACTKSAKAKRRMRKTKRKKTAKK
         **.     *  * **:.. .:*   . **    *     ** * *    * *  *  **

40      YRRYSWRKKKCMAWWWKMKTCKYYMAYCGYMRRTTCKGMCATCTTTTCTGWGAYTTGGTM
48      TGRYSWRGKKAMRWWWKCKYMKYYCTCAACMRRWCTKKAATATGAAAAYKWTCCCKRACM
          ***** **.* **** *  *** : .. ***   *  .::  :::.  * .  . . *

40      MSWTRARTYRRRRGGARKYRGTACCCCGRMCGYKKCRKSKCRWAMRYRGYRYKWTMMMYM
48      ASWCATGCTRRRRTAGGGTGRYTAATTTRMYKTKKTRKSKGRARMGYRKCRYKWACMMYM
         **  :   **** ..      :..   **   ** **** *  * **  ****: ****

40      CTGACCKKMYMGCATKYYMRAGYTTCAAATCMRWYAAAGKARASGGCRCTKCRKWMKRTM
48      TYRRSYGTCCCRTGCKCYMRRAYKWYCWGCAMRWTRWRRKGRTCKKMAYYKYATACKRAM
            .  .     . * *** .*.  . . .***     *.*:.      *  .  **:*

40      AAMMWGAYRCCGYYKRKGACGGTRKAARAYCKAAAMKYYMYYMMTKTCATCTMKGGCWYK
48      GTMCTRMTRTGAYYKGTKCARRYRKWRGGYYTRGGCKYCCYYMMYKYYTCYYMKRKYWYK
        .:*     *  .*** . ..   **   .* . .. **  **** *  :   **   ***

40      CMYYRYYCGSRWRYCRTYYCCRYAWRKMYAYRTTAKRWKGTSMRRTRKRSYRTGGTGTAG
48      YCYYRYYYRGATGYTRKYTYMRYWAGGMYRTAYARKAWKAGGAAGGAKRSYRWRRARAGS
          *****  .   * *.*   **    **    : * **. .     *****   : :..

40      MKRYWYTAAYRCKATKGCGYRYGGGCKYYGACTAYKGMRYYGRGKKYYCAWMTWRRSYYR
48      AGRYWTYRWTRYKCCKAGRCGTKRKYKYCKGYGWCGRMRYYAATKTCCMWTCWAGAGTYR
          ***     * *. *.         **  .      ****.  *.          . **

40      KRMKAAKYMGRYKWSTAGKKCTWATTCTAKRWRKKRYKCCYGKKRMKGTCGGTTRMKTCT
48      GRMKMTTCMRRYKWSYRKKGMAWRGKYYRKGWRGKRTTMTCSKGAAKSWMATCCGAKATC
         *** :. * *****   *  :*  .   * ** ** .   .*   *.  .     *:

40      RCAATAKKTKTAGSCCRCYYSRKAGMYYYYCKTMWGYYSRRARATYKACRGTCTTGGKAG
48      GATRYRKKGTWRSSYMRYYYSRKGTMYYYYYKWMWRYYSRRMRGCTKCSAAKYKGCTKRR
         .:   ** .  .*  * *****. ***** * ** ***** *.  *.. .. .   *

40      ARYYAGCYAYRYTYTGMRRWYYCTYCWYSYYGAMATAWYYMMTTAAGCTKWGCTAACRTR
48      RAYYGAMYGYRYACWTARRWTYMKYTTTSYCRGCTGRATYMMKCGTKMYTTKMCCGTRYR
          **.. *.***:    *** * .*   **  . :    ***. .:   .    .. * *

40      YGTYGYRTGMKCRYKYRYSMRCWMYMGARGACAKKRRYKKKAGTCACTTARGTGGYCCWT
```

```
48          YRYYRTGKKMGAAYKYRYSMRSWMCMKRRRWYCTKRRYKKKWRYMMYYYRGSYARCTYWC
            *  *   . * . ********.** *  *   ..*******          . .    *

40          TCAGGYGAWRKYRTKSMYKKYRYASKKTAGGARYAYRCRWCYKKWRYKGKKKRYRMMGKA
48          YTTRAYSMAGKCGCKGMYTKYRYMSKKCTTARATMYGYRAACKTTGCGKKKTRCAMCKKG
              : .*.   *   *.**.**** *** : .    *  * . *.     **.*  *  *.

40          MMRACTTGTWYMCRYWRKWMRKYKRSRSRAACKAMMMYKAGTTYKTCKAACMATTKCKGT
48          MMGTYGYAAWYMSGCWRKWCRGCKGCGSRRRYGMCCCTKTRWYTGYYKRGYCRWWKSGKK
            ** :   .:***.  **** *  * . **         *:       * .     *.  .

40          AYGCWWYKCYCGGCYTRKKCYRKRRKMRKKYMRKACCTCRSYKYWYAYYTTCCGRGRYYW
48          GCRYAWYKYCARATCWRKKYYRGARTCRGTCMRKWYYYTAGTTTATRYYAATYAACRYCW
            .    ***  . .   *** **  *. * . ***      . .    **::  .  ** *

40          YRYCCCRRWGKWKRRRRAYRWMACCGCKYKKWWYGTTWGGGWYRSMTRGTRSMRARYKRK
48          YRCTATGGAAGWKRRAAGTGAMWMTRYKTTGTTYACKAATTWYRCMWGRWACAACRYGGT
            **  .    . ****  .   *     * .   *. . .  ***.*     .  .**  .

40          YYYYKGRYRWRRKAKYGKYRTCCCGMYCRWCGMTYCSSKRKKRGYMTYGCGRKYSKKKRG
48          CYYYGRRYGWRRKWKYKKYRYYYMRMCSRWYAACYMSCKGGKRKCMYTSYKRKYCGKTAA
             ***  ** **** ** ***     * .** .  * *.*  **  *  .  ***. *. .

40          GRTRYMCMSYGKMCYGTARMGYGAKWKAYTTKKKKKKRYCGGTRRWRKMRAMRWACACTY
48          CRKRCMYMGCRKMYYRGGRCRCRMKTKMCAGGKKKKKRCTACKARWRGMRRMRTRYWYGY
             *.* * *.  ** *  .*     * *  :  ******  . . *** ** **      *

40          WGGAMCCYCKACAKMTTCWWGKMRMYTYCMYCTAGKKKGKWRRMYRTKCRAYTTKCCCTC
48          WRSMAYYTMGGAMKMKGGWARGMRMCWTTATGYMSGTTAKTARMYRCKYAMCGKGYMMGT
            * .       .. **.  *   ***         . ...*  **** *     .

40          MKGAKARCKSTGRRSWTKTGTYGKKMMTRWGWGTTGMYYYGKGCKGRMCGYYCKACKGTK
48          MKATKCAMKGCRRGSAYKYRYYAGKACWAAKWRYWRMTCCRTRTKKRMTACYTGGGKRGG
            **.:*.  *.  * *  *   *. *      *    *    .  * ** . *  . *

40          YATYRAGRKTCYYTKGTYRYWKMRRTYYKRYYRWWTWYWTTRRYCYAAATGAKYYYRTMA
48          CMYTGGTRKYYTTCKACTGTTGARRYTTKRYYRTACACWCGRACATGGTCSGKYTCRKAR
                 . **     *.       **   *****     *  *  . ..: ..**  *.

40          CAWRYGCAWKGACAARYATWTTSSCYCYKRKMCKYKKTYAMRRWGACRYYKRYGMKRRKK
48          YRARYKYRWKACATGGTCGWGCGGYTGYTGKASKYKKCCGMGGTRRYRYYKGCKMGRRKG
               **   **...:.  . *  ..   *. * .****  .*      ****   * ***

40          TKRYMKGGAKGCYAMGKYMMGMGRMKYKKTRRMRRKGRGCRYKTTGTACCWATTWAYKYY
48          WTGYCTTAMKAYCRCRKCCCRCCACKYKKKRRMAGKKRRYRYGYYSYRYTTGGATWCGTY
             . * . . *.     *        ****.***  * *  **   .     . :     *

40          RKKKKGMAYCKKCYAATAATGWYRKAKYKATCKGGTAKKGMAYKRSKAAKYCYRKTSYGM
48          RKKKKRARYTKKMYGWWRGGTTTRGWKYGGATKAACGKTRAGTTRSKRMKYTYRKYSCRM
            *****   * ** *.   .    *  ** .: *.. .*.  . .***  ** *** *  *

40          RKCSRCYKKKSYWYKCAKTWSMCTGAYYTKSRTTATARKYGAKGGCCARYGARKWRAGGY
48          RKYCATYKKKSYWYKYRKGWSMMARRYCKKSRGCTCGAKYKMKARYMWRYKTRGTGRRKY
```

```
      ** .  ********* * *** :  * .***  : . **  *.    ** :*      *

40    KRMWWKWYMCCGTRYKCAATCACTGGTATAGCGGMMMGRRRCAAWWYYATTTGGAATGTR
48    KAAWWKTCAMACWRYKGTGCMMYKRKWTCCRMRRMMMKRRRYRRTATTRGCGKKGGYSKR
      *  ***    .  *** :.    .   : .    *** ***             .. ..*

40    SKRRACTTGWYGGSYCYKTRWYGKKKMMGCATGTGRYKCTKRKMYARKGKYRTCCCKCKS
48    SKGRRYYAAWYRRGCSCKGGTYAKKKMMAAWWRGSRTKYYKRGMYCRGRGTGYATTGYKS
      ** *   :.**  . . *   *.*****..    .* *  ** **.*      .    **

40    RRYTCCTAKYYKTTCAGWAWTAARYRGAGRYKCTYRSKRTTGGAGMYRRRMYYKGRYMTT
48    RRYYMYWTKYCKYYYRSAMAATCATARTTGTKAACASTAYWRRRKACRRRMYTKARYMAC
      ***    :** *    .   ::.    :   *.:  *.         ***** *.***:

40    ATGKYKYKATKRYRMKKTGAWYYGTYRTRTYKRKYCYMKGCWRKTRYTTGCTKYYRKRGC
48    TCRGYKYKMYGAYRMTTWACTCTKYYAWACTGGKTMCCGRTTRTYATWWKTCKCCGGAAT
      :   ****    ***.. ..     *       *        *.        *     .

40    AMCTGYRYCWYRSYYYYKASGYCRYMKMMYKRRRAATYMGTKYYTGGTWYAYTCGCGGTT
48    GAMCAYRTYWCRSYCCCKGGTYYGTCKMMYGRGARMYTAKYGYYCAAKATCYYYATATCW
      .   .**  * ***   *.. *    **** *          ** ...  .*  . .

40    TGTKRYRRYTTAYYCKAWRGAAACGYARRRRMMRMRAKKWKKCAKWRAKKTRRWMKTYGC
48    KRCKACAAYCYRYYYKMWARRMRMSTWAAAGAMRMACKKWKGTGKWGWGGWRRWMTKCRS
      .  *    *   ** * *      .        ***.****  .**     ****..  .

40    KWRYGKCTCMMKYKRGGRRRKKYTAKRTRYWGGRKAYSYRRYYRAGRKMYYYRCKWWYKK
48    KARCAKYYMMMKTKAAAGGRKKTWRTAKRYWRRRKWTGTRRCCGRRRKMYYYGYKWWYKG
      * * .*   *** * ..  ***   . .***  **  . **    ******  *****

40    YMCGTGKKGYYRMYYRATTKGTWKRRGWGCAKYYYTTTGRTTGGWWKSMRRKKAKYCCTK
48    YAAKYRTKACYRMTCGMCWKRYTTAARWAMGGCCTKCCAGCAAATTTCCRRKGRKCTYKK
      * .   .*. ***      *   .   *. .    .  .  :..  .. ***  *   .*

40    KTGRRRCCWRCKCYASRRRMYRTMTGAMYKRARKATAGMTAYYMYRYRRCMAKAYRCKCA
48    TCKARGTGAGMTMYRGGARCTRCMKRTCCGACRKWKRACCCYYMYRYRRYAWTWYAYKMT
      .   *      . * .  *  * *. :    .** . .  .********   . *  * :

40    TKMGYYKYRKMKCTKCKARATRRRRRKKRRRYRAYSKAACWGMATGGAYGCTTTTCTYYM
48    YKMKYCGCGGCKYWGYKTRRKGAAGRKKGAACGGCSKWCTWTMCAAAGYKSWYYYYWCTC
       ** *      *    *:* .    ***     . ** . * *.:...* .

40    KTTCTYKKCCRKRYKGCCAGMTGYGYAGRTYMRTASYKAYTYCSKKRWKATRCMAGCWYT
48    KYYYYYKKTARKGCKAYYTKMCTYKCTCGYCAGKMSYKRYYTYGKKRWKRGGYARRTATC
      *    *** .**  *.  : *  *  :      . *** *   .*****

40    GCRRKCGGGRYYCAAAYCKGGKTCKGTRRCKRKGAYYRWRCKKTKKTTGTYMSRRMKMGM
48    SAGGKMAACGTCYRGWCYKRRKCYKACRGTGGKRMCYAWRTKKAKKCGTCCCGRRMKCRM
      ..  * ..      .   *  *  *. *    *   * ** **:**      .****  *

40    KGAKMMYMYRYACKTRWRYYYARWKKKGYTSYWTYKKMRRKRKTTMYCCGTKYWGCKWYW
48    KAGKMMYCCATCYKYRWRTYTRAAKKKRCYSYWCYKGCRRKRKWCCCTTAAKYWRMKATW
      *..****    . * *** *    ***   *** **  *****      .:***  *  *
```

```
40               WSKKGTWYGCGWYKKMM
48               WSGTRCATSAATTKKMM
                 ** .    ...  ****
```

The pairwise alignments of two accessions 40 and 49

```
40              KYGYSWYYKYWKYKKAMYKAYKWRYKRTYRRYYARGTRTAWTKYKRRYYRRYSGYWKCRY
49              GTKTGTYYKYTKYKKMMYKRCTTATGRYYRRYYWRRKGWRTAKYKRRCTRRYGCCATTRC
                    . **** **** ***  .    * ***** * .    :*****  ***.   . *

40              YSYWWKYRYMKMRYAAGRRRYCWKWYMKKRWGCWKKAMGAYYKGGRYCKCKSRWTTCGMS
49              CSYWWTCACCGMRYGTTRAGTYWTWYMKKAWTYWKKMMSWCTTRRAYAGYKSRAAATAMS
                 ****.     ***.: *    *.***** *  *** *.   .   *.  *** :: .**

40              WYRCYWMRAWRYMWCRWRGMRWYRYWMYCYYRKMWRYMMRWMKRKKRKWSRRYYTYWAGS
49              WYGYYWCRMTRYMWTRTRKMRWCGCTAYYCCRTCWACAMAWMGGGTGTTSRRCTWCAGKS
                **  ** *  **** * * ***     *   *. *   * **   . . ***     . *

40              KCYAATYCGYTRGYKKYKKKGRWMCAYATTWMYKAKYWCYGYYYMRYYYRGGTGYKYYRR
49              GYCTWYTMRYYRKCKKYKKGTAAMYRCRYWAMYKRKYWMYRCCCMRYYYRRRYRYKYYRR
                   :     * *  *****    *       *** *** *    ******    ******

40              CCMYYKCYKYRWWRWMRRSKRWMMYMKYMRRKAKYMMRRWKRMYMKRWRKYRRRMMRRWY
49              YYCCTKYTKYRTWRWMGRSKRACAYCTCAGRGWKCAAGAWKRMYMKRTGGYAGRAARRWY
                     *  *** **** ****   * .   *  *     ********   *  *  ****

40              KRCCRWAMYKRYWYRCYSYRRWKYGYCYMYYMTYRKAGGYYRRRKMGRGTWCRTYWYRRG
49              KRYYAAMCCKRYTTAMTCTRRATYRCTCMYTMWYRKRRRTYGGGKMARKYWAGYYWYRRK
                **       ***     . ** .*    ** * ***    *   **.*  *.  *****

40              YYGTTGGYRYMGSCKMKYKCWKTCASKYMRWAKRYAMCGKYRGYYGTTRRYRKRRYYRWY
49              YYRAARKYATMKCYKMGCTTAGGYGSKYMRAWGGCTATAGYRRCYACAAGTAGRACYRWT
                ** ::  *  * . **  .     .*****     :  . **  *. :     *  ***

40              KRMMKYMYMRYYKWWAAACAACYYMWMWYTWTYCWYMAGWYCWYRYRRWGTAGGTYRAKK
49              GGMMTTMYMRYCKAWCGCARRYTTCWMWYCTKYYTYAGAATTWYGYRATAATATGCRTKK
                  **. ***** * *....      ****  .*  *  ..   ** **  .::.   *:**

40              TKRTATGTTTTKKYCCMARWSKCRCGSMRYRGTAWKKSKCGTTCKAWYRYRKYSMCGCGG
49              AKGCGGTGWYKKKCYSAGGTGTYGGACMRCGACRTTKSKARYWSGMATGYRGCCMARSTK
                :*  .     .**  . .  ..   ..**  .   .***.   .     **  .*. .

40              ARCKACYMWYKYGRYRMRKRKAKKRYYTTYYCKGWGWKTYYRMYAWYYMTTAGACGRYAC
49              GRYKRYYMWYKTRRYRMRKRKRKGAYYYYYYSKKTRTGYTTRMYTACTAKYMRRYKACRY
                .* *  *****  ******** *  **  **.*        ***:    .

40              KCCKYYMKYTARGGGYCCTTGTWYGKTTRKAWKKKWWKRWTATSWGWAKRTATTCTGRAA
49              GMYKYYMKTYRAKRRTYSAGTGTCTTYYRKRWKKKTTTAWKTCCTRARKRCGGWYYRRMR
                   *****         .:      .  ** ****  . *.: .    ** .     *

40              ATTRKTAYRMMSRSRKMTACACGCCGATYKGRKRKCAGTAKYYYSGTKTCTWMGCGMYKM
49              RKWRKYCYRMMSRSRKCKWTGYKYYTGWTTKGKRKYRAWMKYYYCKYKYYGAMSYRMYKM
                 . ** .********* .  .     .  .  ***  .  ****.  *    *.  ****

40              KCCTWYKKRYCGRRYKCARRYAKYRYMMYAYRTTAGACACCRKTSMRRRKKGAAWCRKRK
49              KYYKWYKKAYYRRGTGTRRAYGKYGYMMYRTAYYRRRYRSMAKGGAGAGKGTRWTYATAK
```

```
              *  .**** *  *     * *.** ****          .  * .    *       . *

40            YTAGKWCGKYRKATCRRGYRYTAYAWRRGGCKYYYWRYAMRYYAAATYYTAKRMRSAGWY
49            TYRRGWTSTTRKCCTARRCGCAGTWWRRRKYKYYYTRCRMRYYWMMYCCWRGGMAGMAAC
                   * .. **.   *    :.  ***   **** *  ****          * . .

40            RYYRYKGCGCCTRGAAAAAKYCTRAKYWSTAGAKKCTCTATARYGKGGATKRKCKKTCTR
49            GYYRYKRMSYYWRKRMMMTTCTWGCKYWSYRKGGKMKYKRYRRTRTKCMKKAKSTGATCG
               *****  .   *     :.    .****   . * . .   *  .   .* *.. :

40            TTKKYRCKTACGSYYKTYKTATGSYKYWYTRCAGTAYACRWRKYSWRRMACYCWYGYRTT
49            KGKKCGTTWRYSSYTGYCKYRYRGTTTATCGYGCCWCMTGARKTCWRRMWMTYTYRCGAC
              . **   .   .**    *    . .      .        ** .****     *   :

40            ARYMRKKYYYTYYYRACKAAWYKKYRATKGSTSRKAAARRKKRYCAAAMYGKYKKGKRKT
49            RRYMGKGYYYYYYYGTTKWRATKKYRRYKRSKCRKWRWRGGKRCTGGCMTKKTKKSKAKK
               *** * *** *** : *    ****  * *..**   *  **  ...*  * **.* *.

40            RYCMSYATGKMKYGMGYTCGWKKKCYCAKAKRYKCGGGKRKTGTGAWRAMRGWTYCTYWK
49            RCYMGCCARKAKYTCRCKYAWKKKMCMCKWKRCKTACAGGGCKKAWWGRMRRTWYYGYWK
              *  *. .: * **    . .****   .* ** * . .     .. *  **   *  ***

40            MCGAWYYKWCRCGAKYRRMGKKGRACAAGRCTYCRTRRGGTGYCWYGCKATCKCTKRTRT
49            AARRACCKTYGMKTKYRGMTKGRRWMRMRAYWTYGYGGACCRYTWCTTGGYYKTCGRGRC
               .     *     :*** * *  *              .   * *    .  *   * *

40            GAACRRRGGCCWATATYARMGKCTYAAACTYKCTGKWRYMGGTRACARWAAGCCKKMKTG
49            RTCYAGGSKSYWRCGYYMRMTKMWYCTRYYYKMCAKWGCCATCRRTRRWMGAAAKGAKGA
               :.    . . *  . * ** *  *.:   **  .**   .  *   ** ....*  * .

40            AKKTCMRWTCYRKWGMRKKMWTATMGTAAAGACRYMTYGTCGAYYMKGATGSGTKKYCCA
49            GKGYYMRWKTTAKWAMATGCTCCCMRWMRCTGSRYCYYKAMTWYYCKKMCSSRYKKTYMM
              .*   ***.   **.* .    . *    . ..**  * :   ** *   .*  **

40            AKKSAWTYCATACCSKKRARTMRARGWKYGMATYTGTSYKYACCTRSRACRKTYYMCMKC
49            GKKGWWYYTTGGYSGKGGMRYMRMGTAKYKMRWCYAYSYKYWTTCAGGRYAGYTYMACGY
              .**. * * : . ..*   * **    ** *    . ****     .       **.

40            CYKAGAACGARGATTRKWRYWGCACARTWKYRRWKATTATYYGTKTTMMKCAYRMKTCGG
49            ACKRKGGYKGRATCCRKAATTASTTTACAKYGRTKCWYRGYCAWKYYAMKATYGMKATSS
              . *  ..  .*.:  **    ..: :   ** * *.    * . *   **.:* **: ..

40            GYWATCGGTGTGTAWYRCYRWRYTSYYYMWSAYARMYTAYYKGTTGTRYYYRYYCCKYRA
49            KCTRATTRYRATCMTYGTYRWRYYSCCCCASRYMRCTCCTYKSACCCGTCYRYYMAKYGM
                  :     :    *  ***** *     * * *   . **.:      **** .**

40            YRYMSWRRYGTYSYCAGCMMAKWYAYYMMTCTAAKWMSRSRMMYCAKWCRYYWCYCCKYC
49            YRYMSWRRTAGYSYMWRYMCRKWTRYYMMKYCGTTAMCRSRCMCGGTTYRYCTTTTTKTY
              ******** . ***    *  **  ****.  .:. *.*** *  ..  **      *

40            CAKTRARKTCRYKYRYWSMRGTYAGTRWARRYMCSAMRGACKCCKGCRYRYKKRKAMMYA
49            YMKCRWRKCAAYKYRYWSMRRCCTCAGACRGTCTSCMRRWYKYMTRYRYRYKGRKWMCTG
                * * ** . *********   : :  .*    *.**   *  .  ***** ** *  .
```

```
40          YAKATWYAYYRCRASKKCGCYRRSKKTCKKKCTRRKCWRKGCAKCATGYTAMYTTCACKK
49          YRKGGWTCYCGSAMSKKTAYYGRCKKYSTKKSKRRKYAAKKTTKYRYRYCMCTKCTGMKK
            * *. * .*  .  *** . * *.** ..**..***   *  :*    *    .  . **

40          RYKYRGKKRMWYYKYWWTRYYYKGWMCYTCMGGTATTTACKKCGCYSMAARKWRMRGTTT
49          RYKYRAKKRMWYYKYWWYRYCYTKACAYWYCKRACKYYRYKKYKMTGCTGRTTACRRWWK
            *****.*********** ** *.   .*     :..    **    . :.*.   *   .

40          YYCGGRYCKRKMYTKGCTCAYKYMRCCTKKATKCASRGGRCCYYYRWWAGRCYTYKKKRR
49          YTYRKACYGGKCCGKRYYMMYKTAGYSWKGMAKTTGAKKGMYYTTGAARTRYYKCKKTRR
            *         *   *     **    . *  :* :.      *       * *. **.**

40          AAAYRYKWRRGAWKCTAYCAKGGAWYKWAWKWCRRKYRCCGTCGTRWGCRAMKTTYYKMY
49          WRWYRCTAGGKWTKYWWYMRKTKWWYKWGAKWYRGKTRASTCGRGGATMGMMKCYYYKCC
               ** .      *   *  *   ****. ** * * *..           **  ***

40          RAMCKRGRMRMATTRCMRKTAWWWYKKCCKRACAACTYMKYKKYCTTKMRCGYYTTKYTC
49          GTCTKRTRCACGGYRYMRKKRTATYTGTMKRMAGCTGYMKCKTTYCCKAGYAYYYWKCKS
             :  ** *   .  * ***.    *.   ** ...  *** *.    *   .**  * ..

40          TCGYARYKTYGYRTGRAWKGYGYAWTAGGMGGRYRACTKTKYKRCKMCTTATRGGAAWCY
49          KARCWGYKKCRYGCKAGWKSTAYCWYWSRCRKGYGWACKCTCKGYKCYWKMKRRCCWWYY
            ..    **.  *    .**. .*.*  .     *  . * . *  *   . .*  . * *

40          AYKRRRRGYMYRAYAGGAAYCMTCAARYKRCTMKAMGTKYRAYRRGGRGKCARKSRMGCT
49          TYKRRRRTCMCAMYGATTWYMACTGWRCGRMYMGRCRYGYRGYRRRKRRKMWAKSGMKSK
            :******  *   *.. : *    . *  *  *      **.***  * *   ** * ..

40          YYTCRRRTWKKWYRSTRCKGAYTTRAKRRYTKKYRTTCTYWAATTATRKWCRCAYKRYAK
49          CYGAGGRCWKKWYRSCAYGTGTWCRWKRRYAKKYRKWYWTTRRACGARKWARARTTATGK
             * .  * *******     .   * ****:****.       : .:***.*.  .  .*

40          WKCAGGGTAKGRRYWCGAGKYKMCWYMSKKYYKYGTTGAMRWSSKAACWMSCYCRCTRAC
49          TKTCATRYRKRAGCWYRRRKYKCYWTASKTYCKYAYYRWCGASGKCGAWCCYYMRYCGMM
             * ..    *    *    ***  *  **.* **.       *.*...* . * *

40          KARMRKKTKKGAGGYAKKAKYCWKRYGTTWKKKAWGCKYTCKCAYTAYAMCKYRAKWYYK
49          TMAAGKKGKTSWKRYWTGGGYTWKRYSYYAKKKRARMKTYMKMMCWWCRCMKCRMGACYK
            .    ** *..   * . . * ****.   ***    *   *         * *    **

40          RGTRTACTTKKRKGTTAYGAKYYYKWAGCKYTTGGGRYGGCKACGKYYYYMCYTCKKCCC
49          RAYGYWYYKKKAKSKWWCACKYYYKWTTTKYWARKKGCKRYKGASGYYYCMTYYYTKMTA
            *.      .** *..   ..******:  ** :        *... *** * *  .*  .

40          RCKRGCTYAAASYAAATAYYYYRYRWKWAWRAKMTGGTMKMWWYKRSKKKKSRYKSARKT
49          GMGGKTCCTGGSYGCGCCTYYYGCGAKAMARRKCGAACMKAATTKAGKKKKSRYKSWAKC
                    :..**... . ***    *   * *  .. **    * .*********  *

40          YYACARKTRYKWGYKCKMMMYRASKTKRYRWKAMMRRRKKKCGCYKMGYRKRCAMRMMTA
49          TCWMRGKYGCKTKYKTKCMMTARCGYKGYRTKWMMRRAKGKTAMYKCRYAKGMMCRCMWR
                  *   *  ** * **   .  * ** * **** * * . **  * *    * *
```

```
40          YKMYWKMMACGWRWYGRRAGRCGRKRYKATGCRMRCKKKKWWRYYWGRWKWSWRKMYKYG
49          TGAYTGAMRYRWRTYRGARKATRRGRYKGKKYGMRTKKKGTARYYTARTKWGWRKAYKYA
               *   *   ** *        * ***..   ** ***   *** .* **.*** ***.

40          TGAAKRTYYRWWYAGAYYYYKYYKRCCARKKAYTRRRACYRTKCSGKKGKYRSKTCTWAG
49          CRCGKGGYYRWACTKMCYCTKYYKRYYRRTKWCCAAGMMTGCGMGKKKAKYRSKYMYARK
              ..*  ****  :   *  *****   *.*             . **.*****

40          KYRYCYYRMATWRYYKKKRCAYRAMTGYKCKKKMCMKTAGRCRYKCMRKMYYGTRAARYS
49          KTGYSCYRARYTGCYKTKRTGCRRMGKYKTKKKMTMTWWKGYATKSCGGMYCSYRMMGYC
            *  *. **      **.** . * *  ** **** *.       *.   ** . *   *.

40          RKYAACKAYYGYYWWKKGTWKGTTYMKYRCACGKKMRATTGTWACYGTCKGTRWWYYRYW
49          GGTRRYKMYYTTYWWKTKATKRWYTCKYAYWYKKKMRRWWRYWCGCKCSGKKRWWTCRCA
                  * **  ****. : *     **     ****     *.    .  .***  *

40          YYYYWYGGYKYRRKRRSKRKRGYRKMRRYKGCRKYKYRMRKKCKKTKKKSAAGGAAGCGY
49          YYYYWYRKTKCRGKGAGGAKRKYRTCRGTGRARTCGYRMRKKYKKYKKKCCTKCMGATAY
            ******   * * *  .  ** **. *    .*.  ****** ** ***..:   .. .*

40          CYYCKRKYCKCGGYKMGYYAGRGGYWRYTKRRWKKMARAKKRSMCCTGRCRWKKGKKWKA
49          YTCMKRTYMKTARYKMRCTWKRRRCTRYCKGGAKKCWRGGTASAYYYAATRWGKRKKWKG
                **.* * . ***     *    ** *   **  *. . *    .  ** * ****.

40          WYKMAGCAYKMCMTTCKKKRYCTTACSRKCCCRWKAGAAKYGKGWAGGCYYRRSMYKKKK
49          WYKMMKYWCKMYMYWMTKKRYSCCGTSAKGTTAAKRRMRKTRKRARARTCTGRSCCKGTG
            ****     ** *   .****.  . * *     *    *  *   .     **  * .

40          GMCCKCYCGASRRRKMGKRKRTAKCTATGACGCMATCCGTAGGAAYYGTMYYYSRRRYTA
49          KMYSTYYYKGGGARKMRGAGGGGGTYMYRWMKMMWATYKAGAKMRYYRGMYYYSRRRYKW
             * .. *  ..   ***      .          * :   :..   **  *********.

40          YCWAYRCRCKCTKGAMGYRYRYARKSMKCCTKCARRRRRRRYRYTYYTKRAGGACRTKYC
49          YYWWYAGAYKMYTSWMKYACRYRGGGCKYTWGYRAGGGGAACGCKCTCKRWCRMYAAKYS
            * * *    *  .. * *  **   . *                .   **      :**.

40          TTCTYYMTTCTCCRCRKTGCGMRAMSYAYMATYCGSTKKRAMCYMKKCYWYACCGTYGTT
49          YYYWCTMYYYYTARMGKYSMKMGRMSYRYMTYTYKGCKKRRMTYAKKTYWYGYTACYTWC
                  *     .*  *  .  *  *** **:    . *** * * ** ***.  . *

40          RRKRYYRWCCAKGAYKGTCTCYYRTKYMGKMCRSSSRYKKAMMYRRTTRGTKYYYRTGGC
49          GGKGTCRWGYRKKWTKCYYKYTCRYKCMKKMYGGGGGTKKGMMYRGKYRKYGYYYRCAAM
              *   **   *   *   .   * * * **  ...  **.****  . *   ****  ..

40          SYTWKATCGYTTRMYMCGYCAYRCGYYARRRRRMRMRKCKMCKTAGTKKKWRRWMAMGSC
49          CCKAKCYYRYCCACCMYAYYMYAMSTYWAAAGRMRMAKMKAYGATTAGKKWRRWMRACSM
            . .*.    *     *  .*  *  . *     ****  * *   ::  : *******   *

40          RWYRYGACACCCKMMYKGAYYWKRRKKYRCKRGGCTYTRYWYKMRRKYSYRAAKYRRWMY
49          RACRCACMMTTTKMMYGAMYTWKRGKKTGMTAAATCTKRYWYKMRRTTGTRGGKCGRWMY
            *  * ..     **** . * *** **   . ..   .********. . *..*   ****

40          YYRCGKKKCKKTGKGYYRKRKMCMYYYRGCCTWKRRGWKWKKYMTTGGWWKSMYAKRRKK
```

```
49          YYGYRKGKAKKYRTAYYRKRKCYATTCGATTATTAARWGWKGCMCAAATTTCCYRKRRKK
            **   * *.**  ..******       .  : .   * **  * :..  .. * *****

40          GGTCGKCTGYRKTKCKKKTTTMMKGAMMMGMYRMCGCAKKCAKCRSRKKGWRWCGCYYCK
49          AAAARKYGRCAKAKYKKKYWKCMKRRMMMKMYGMTTTTKKTCKYRSRKKAWRTYRYTTMK
            ..:. *     *:* ***  . **  *** ** *   :** .* *****.**       *

40          WAGAGYTSYWRTKTTMRRCKRMTCRCGTMYARCKWWSRKKKTTYWYRWWMRMMKKMAGGW
49          TGAGACYSYWRYGWACRRYKACWSRSTWCCGRYKWWSRGTKKYTATGTWMRMMKKAWTSW
             ....  ****   : ** *   .*.    .* ***** .*.      *******   .*

40          TKMTKKKGAYKGKMYKYCRMRRTCCTGMMCYYSKYTGYGMSWCTRRRGKAAGKCRAKKKC
49          WKCKKKGTTYKRKAYKCMRMRRWTYATACGTYSKYKRCRMSWMCARRKGGTRTTRRKTKT
             * .**  :** * **  ****   :     ****.   ***   **  .: . * *.*

40          KAGGRAYCTKSKKKYKKAGARYKWSCMGTCASYCTGACRKYAGMCWTTCYCTCTTMRWYY
49          KGTARCYAAKSKKKYKKGRRRYKWCYMAWYWSYTYRRSRKTTAMATGGTCTYAAYCGTCY
            *. .*.*.:********.  ****. *.   **    .** :.*.       .:     *

40          YSAKKGGCTRTSYCTAMMSTGKTGATTKTTYCCCCGYKTTTTCGAGYWKAGGATAAYAAA
49          TSGKKKKMKAYSYYAWMACAAKGAGYWKACTYMTGAYGWCWWTRGKYWKGTTWWMRYMCM
             *.**   .  ** : * .:.* ..  *:      .*       . ***.      * .

40          TKGCGCGMGMMKKCAWWYYGGTKATRRASWGYYGRTSACTACKTTTKGTATYKCKACAGA
49          WKAAAMAMRMMGKYRTATTKKYKGKRGRGWRYYRGCGGMWGYKCACKAGTAYKGKWSWAG
             *... .* ** *         *..*  .* **   ..  . * : *. ::** * . ..

40          YWAKTCTTKCCAGSKYRMYAYWTAGKGKYRRYYWTCCCCMGCGGGTCGTTTKYYKAGTTT
49          TTTGWYWKKYTCAGGYRMYRYTGGAGRGTGACYWYYATTARARAKYMCGGWKYCKMAAYY
              :    .*  ... **** *  ..       **  .    . .       ** * .:

40          CAGWAWRYRGTTYKKYCTYRKSGKRWTAMYACCATYYKGAACRYYKYKAKRTYRWMKGGA
49          YRSAMAATARCCTKKTAACAGSKTAWWTAYWMYRACTGRRWSRYYKYKMGAYYRTMKRAC
              .          ** .:   * . * : *    :      .******    ** ** ..

40          YWYYGTCRKTKKWWYKSWWRRRKCTAKYYTTGKYYRTKRGCACAGGRTGYRKKWRRYTYR
49          TTCTKYAAKWKKWWCKGTTAARTAYCKYYWWKKCCGAGAATGMMKSRCAYRKKTGRCCYG
                  . * **** *.    *.. .***   *   :  . .   .* .****  *  *

40          RACKMCGCAGKCTKTRRTKTRAKGYRRWCCKCKRGCAGACKWGKKMWYTGGMCAGGKKGG
49          GRYKMMAYCSKACKKGGCKCGGGKYAGTATKTKARMMRWMKAKKTAWYKRKMYRKKKKKR
               ** . ..*. *.   *  .  *   . * *       *  *. **.  *    **

40          GKRGKTYRYKKKSCWRAWMKCATRKKRAATYAAACATWMMYYYRTKKWSGRAYWMYRGWR
49          RKGRKAYRYTKKGYWGGWMGYRWRGKRCGATMRRYMYTCACYYAWKKAGTGWYWMYRKTR
             *  *:***.**. * .**    * **..:           **  **  .   *****  *

40          TGGCCSMWYYRMRRWKWYAAGGTKACKGYGCKKAKKYAGMYRRRKRRYYRCRKWKRCRCA
49          WKRYMSMACYAMRAWTTTTGTACKGTKKYTSKKWKKYTKCTAGAGRRYYRTRKTKATAAC
                 **  * ** *.  :. . *. * * .** ***:       ***** ** *   ..

40          TTCGSYTGRYKRWGCTAGWRKRGCATCKTYGTAKKYCYYYWKCRWKYYKMRTMRRYYKRY
49          AATRSYYTRYGRWTTARTWRGGRTGAAKWYTAGGKTGYTTWKYRAKYYKCGYMGRYYKGC
```

```
    ::  **  ** **  :  **    .:.* * :. *  *  ** * ****   * ****

40  GKTAKKRYMKCAYKRRMYMKKCAGGGKRMRRRGCRYCKMMGKWCGRGRYKKYYRKKKGWK
49  KKYWGTGYCKYMTKGGMCCKKYRACTKRMAGRRYRYYGAASGTYRGKRCGKTYRKKKRWT
     *   . * *   *  *  **  .  ***  *  **    .      *  * ***** *.

40  CKKGGCYKAKCCTCKKGKRSKYCWKATGAKGMRKGRSRCYKYSYWYKCWAKSYKTTKYAK
49  TKKRTYCKTKYTATKKTTRSKYTWKMYAMTRMRKTRCATYKYSYWYGYARKSCKCCGYGK
     **    *:*  : ** .**** **  . . *** *.  *******    ** *   *.*

40  YKGAKWWGCRYGRKGWRRWWKSYMWYMCTATKTCACAATGGAGTRYTGYCAATACTGKRT
49  TTKMKWWRYRYKRGKTGAWWKGTCTCAMYRYGWAWTGRKCACAWRYYRCGTGCMYKRKGW
     .  ***  ** *     ***.           .  . . ... **    :.   . *

40  TTAGGKAKKRWRMYRCRRTGAARAKAATGCCCAWKRCWRYYCGWKSTCRYTGTKGMMRGK
49  CGMTRGRGKRWRMYRYRRYATCGMTCGYRYMMGAKAMTRYYTKAKGKYGYWKWGKACAKT
            ******* ** .:.  ...     . *   ***   *..  *         .

40  ATCWGKGYAKMGAGTRTKRYAARGRGAARYTAYCTYYKKWMYYYRWWWYTMRRGTAYRRK
49  CCATRTKTRKMTGAARYKRCRMGKRAGGRYWWTTYTYGKTATYYRTAACCARGAGGYGRK
    . .  .   ** ..:* **     *...**      * *   ***      * . .* **

40  AYYYKTCWRYCAWWAAACKKCYTRWGYCTCTCYCGTGGKYRWRMRRYKYKGSKKRAMATM
49  GTCCGYYARYYRTATTGTKKYYGAARTTAAATCYAYSRKCRAGAAATKCKASKKGMCRKM
    .       **    ::. ** *      :.:   . . * *      * *.***    .*

40  GYMYKGTKYKKCYCYYYKKMSACYKYATRSKYGTTGCGCMCCTKKMYRYYYRWYTYWRWA
49  AYMYKRYKYKKYCAYYCKTMSRYTTYRYAGKCKWCSYRYMYTCKGCTGCYYRWTGYTATT
    .****  ****  .** *.**   .*   .*    .   *   *     ****  *   :

40  SKRGMRRYKYCRRRAKSWMRYRTMKRRGTYSMRKMRCYKTRYMKRYSAKRWKWGYAMYRG
49  SKRRCRRCKYYRRRMKGWAACRKCKGAKWYGMGGCGSCKCRCCKGTCGKGTKWKTGMYRA
    ***  ** ** *** *.*   *. *    *.*    . * *  *  ..*  **  .***.

40  GCCGTSRAGMRRTAGKKRTRAATTSMRTGWGKKKWYWMRKRKYGKTACKKAMCKKKKYR
49  TAMTWGRMSMGAWGRKKRCGRMKWSMAWRWTGKKWTAAATRGYRKCGTKKMAMTKKKTR
     .   .* .*   . ***     . **   *  ***    .* * * . **   .*** *
```

The pairwise alignments of two accessions 40 and 50

```
40              RAYTTGGRTMKYYRRWCGGKACCKKGGRTMRMMWKWRRRYCCCKWGWWMKKCCKCGWRRK
50              GRCYKRTGAMTYYRRWTRRKGTTKGKAAKMRMMWKWGAGYATTKWCTTCGKAAKYTWRRK
                    .   :*.*****   *.  *  . .*******   *.  **     *..*  ****

40              MYACAKGAYYYYRRSRYYWCCCKTTAYRTGYMYKRYCYRWYYRCARRRGCCKACRWRCMY
50              MYRYRKACYCTYRRSGCTTYSYTCCTYRYAYMYKRYYTGTTYRARRAACTAKRMRWRMCT
                **   *..*  ****     . .  :** .******     **. *    .*  ***

40              TGAYTRCAKRYRWYKYYMYKRTTYTYTRKKYCYTGYRARRKKWWMRMRWSYWWMYKWMTT
50              GAMYYRMWKRYRWCGTTCTGGWWCYYKGKKTMCWRTARRRTKWWMRMRTSYWWMYTACCC
                 . * *  *****            *. **        **.******* ******.

40              CTYKRRRWYYMRRRAGGKCWKYMRYWKSWYWRRGGARYRYMKKMWYAGKKWRCYMYTGTY
50              ACCGRRRTYYMRRRRRRKYWKYMRYWKSWYARRKRMRTRTAGTCWTWKKKAATTMYWRWY
                .   *** ******   * *********** **   * *   . *   **    **   *

40              KYRWCCYRAYWCGYRYKRWYRYYGSKRTAGAMACYSCWRCMCYYRRKRMYYKRRRYKKCT
50              KYRWYYYRRCAARYRYGRWYRYCKSKRACTCCCYTCYAGYCYYYRRKRMTCTRAGTTKYY
                ****  **   . *** *****  ***:. . .  .      *******  .*   .*

40              CARKMYRRKAKMRYGYYKRYRMRYYMYKRMYKCTGTKRAATTGRKAYRCMKWTMGYWGRS
50              YRATMYGRGWKMRYKCYKRYRMRCCAYGACYKAARAKRGTGCKGKRCGYMGTKCKYWKRS
                   .** *  ****  *******   *   **.: :**.:    *    *  .  ** **

40              RRYMGMKSMCMKYKGMRMRRKCYMYGSGCAYGYWWRKGAGYYYKRWKWRYRYGSTATAKC
50              RRCAAMKSMYAKYKRCRMGRGYCCYAGRYGCAYWTGGTRKYYYGGAKWRYAYKCWTCCKY
                **  .****  ***  ** *    *..  . .**      ***   **** * . : .*

40              YRRRCAWWYYGGKKWMKSKRGRKMYKWYCARKKKWAGGGTATYKCAAYKWYTKCSRRWAA
50              TRRRYRTATTKKKKTAKSKGKGKMYKWYSRAGKKWRRRAGTAYKSWGTKWYCGYGRRTTC
                 ***        **  ***   ******.   ***   . ::**. . ***   .** :.

40              GKRTAKKYRRYCCTRTKYARYTTTCWSAGWAWRYRAYKKCTKRWTATMRRRCYYKYKYKR
50              KKGGGGGTGACMTCRWKYRGCYYYYWSRSAMAATATTKKAATAWWGCCRRRACTKYKYGA
                 *  .         * **       ** .      : **.:. * .  ***.  ****

40              YRMKGARWYYGTCYRKTYKRYKWSTGYRATRAACKAKKCKMTRRTTMRKRAKWRRMYRKC
50              YRMKACRTCTKYACAKCTGGYKWGYRYRWYAGGTKWGKMKMWRAWWCGTRMGWAGACAKY
                ****..*     .  *    ***.  **   .. *  * ** *     .*  *     *

40              TTRCCCCMCGCWARCKARRCMWRAGCCMMRWGTYCTRACGKGCYKCACTCKACGRRACRT
50              GKGYYMMMMAYTWRYTGGRYCARTAYMAMAWRKTYYACYRTKMTKYMYKYGGGRRRMGRW
                 .     * .   * .. *   *:.   * * .    .  .   *   .  .  **  *

40              YTGSYYTKWGMRYYYRGWGRKWKTMKGYRKGRCAYYYKWMRWRYKGMKRKTGGKRMAYYR
50              TYRSTTCKTRARTYYRTTSRKAKCAKRYRKSAYTTCCGAAAARYKAAKRKWRRKRMCYYG
                   *   *   * ***  .** *  * ***.  :        ***. ***   ***.**

40              RAKKYKRKCKTAATRMRKRKATARTYRYMRGCAKWGKWYRGGKTRKKGKRYYKKKWKAAA
50              GRGKYKRKAGKRWKRARKGKCCMGYYRYMARMRKAKTWYGKKKGRKTRKRYYTKGWKMGG
```

```
               *****. .  .* ** *.    ****    *  .**   * **. ****.* ** ..

40          GAWTRKWRMKATKGTAAAACGKMYYYRACATAKKYRRRYAGWTRRSRGCCMWYRRWKWYM
50          ATWCRTTRMKCAKKCTMRRYRKACYYAGYWYWKKTGARCWKTGGGSRAYMMACAAWTTTM
            .:* *. ***.:*  :     *  ** .    **   *       **.  *    *.  *

40          AAGGACKYYRAYKWCGYWCKRGRYWRCMCRYRYRKRKRYWRKTRYWWKWRWKAAAKYRYK
50          TGTAGTKTYRWYKWTACAAGRAGYTAAMTRYRYRGRTRYWRKCGYWWKWRWKCCTKYGYK
            :. .. * ** *** .  . *. *  .* ***** *.*****  ********..:** **

40          WWRYYWRWKAGGCWRRYCAAAMKYTCAGTCKRYTRTCKYMSKWCCYAGTCGWWYYTYKWR
50          TARYYWRTKGRRMWRRYAGTGAKYCYWAYYKRYYRYYGCCGGWGGCTAWTAWACTWYKWG
              ***** *.   ****..:. **   .  *** *     . *   :.  .*    ***

40          GTGTAYGTCCYKTRTYRWRRWYYTTGCTKCTCMKTKYKMGKMMGKSGATCKTWRYCKKKY
50          RCACGYAAMAYKGAKYAWRRWYYYKRYAKMCYCGYKYKCRKMMKKCRCKYKKWRYMKKKC
              . .*.: .**  .* ****** .  :*      ***  *** *. .. *.*** ***

40          TTSYWTWRKMRRMACTKTGGTRMTMTAMYGRKRATAAKYWGCAKGSAKWWSWYRMWCCYK
50          YKSYWYWRGMRRMGTAKAAAARCWMCRCCTRKRCCGCKYWRMTKKGCKWWSATAMTYTTK
             .*** ** ****. :*:..:*  *     ***. ..***  :* ..****   *    *

40          MKKYYMGGCGWYMCAMKYYCRRSRWRYCRTYWKYRMTYRAGGAWAGCCRGRTRRKGRYWY
50          CKTYYMAAMTWYCSMCKYTYRGGATGYTRCYAKYGMYTATAAGARSSMATRGAGKCAYWT
             *.***..  ** .  **  * .   * * * ** *   :...  ..   *   *  **

40          ATCRCTTWYRCYRYYAWGGCKYWAGYGGCRCGMRYYKYYTATAKWRRSKKYYYKTGAAKR
50          RWTGAACWTRYCGYTWWRKYKYTTRCTRMRATMRYYKCCAWWRKAGAGKKYYCGKAMMKR
                .: * *   *  *   ** :     *. *****  :   *   .****  ..  **

40          MYWSTAGKCCTTTATYGGKKCKCKRKKRRRMTKTAAAGGSCCRYYSRYTCCYYRGGKSYW
50          MCWSYRKKMACAYRKTKSKKGGSTGKKGGGCWTWGGCASSYMRYYSRCYYYTCATAKSYT
            * **   * . :  . .**  .. **     . .....*  *****        .***

40          WYYMKRGSSSKMMYTGTKYATGGTTCTAGRYRRACACAYRGAMMRRGAGKTYMCKMWKKG
50          AYYMKARGGGKMMYYKYGYMCAAKYYCRSACARTYGYMYASWAMRAKWKKGCAMKAWKGA
             ****  ...****    *  ...    .   *: .  * .  **    *    * ** .

40          MWCAACATKTMGMCKRYKMTAMMMCYWRKRRRKYRTKCKRRYMYRSWTKRYWGMYSYTRR
50          CAYCCTGCGCACCAKRYTAAGAMMYTWRKARGKTRWKMTAGTCCRSWKKRYWKMTGTWRR
               .. .      .***. :. **  *** * * * * .     ***.**** * .  **

40          YRCAARMYYYMRRRCTTTGMKCKMYMCTKGYRCKYKWKRRGWWKYYYWWKSRKRCTTKMK
50          CGMRCRMYYYMRGRYYAGKCKTKMCAAYTAYRYKTKTTAAAWTGCCTTTTCRGRMWKKMK
                .******* *  :   * **  . ..** * * .  .*       ..* *  .***

40          AMCKYMRTMRRYSYRKRRKYKRYKAKYMKYKKRMRRGKKRGCAKRAAAKKGMRAYKCKGG
50          RAYKYCGYMGRCSCGKRRKYTGYTMGCCKYGKRMAGKKKRRYGGRWWWKKRARRYTTKRT
               **   * * *  *****. *.    ** ***   ***  . *   **  * *. *

40          GYCRCCTTCSYKSYKTGTTRSKTYCWCRAGTSKMTGRKCRYRKSRCYSYWYKAKTMTRMC
50          RYTGYYAATSYKSYKCRAARSGKTTWYRMRYSKMCKRKYRYRTCATYSYWYKGKYAARCA
             *    :: ******  ::** .  * *   ***  ** ***..  ******.*  :* .
```

```
40          TYKAAKYYKKYGAYRMYGAKGCCRYGGGRKKWRACGAMWWKWYMCCTACAATGGAGTKYC
50          AYGGCKYYGGYKGYRACKMKRYMRYKKKRGKTGTTAGAWWKTCAMMWTTGRKCACAWKCG
            :* ..***  * .**    *   **   * *  : .. ***      : . . ... *

40          AATWTCGTGGKRMRRYKGCGGKYRRKKGWRYRAGTGGGYMYKKKGKMTAAGYYYKMSRYR
50          TGCWCTACCAKGAAATKKATKGYRRKKAWRYGGACAARYMYGKKAKCCTRKCYCKMSRTG
            :. *  .  .*     * .   *****.*** .. .. *** **.*  :   * ****

40          KYKYATRRSKYYGTTTAGRAKGGWKTGCYRCKMYRYGATGYYRWYYAMKRGGTTTGAYYK
50          KTTYRWRAGKCYKWCCTRRRKKTAGKKYYAMGCTGCAGWKYYATTCTMKRKTYKCARCYK
            * .*  * .* *    : * *    .  *       ..  **    :***   . .  **

40          SRYRRMWKGSMMKCKCCYYMKRYSAKKGARTACARYSKCGTRRAATTRMYAGKKKWKWYW
50          GACRRMWKSGCMKTGMTCCCKGTCGKKKGRYRMRRYGGASWRGRMKWRMCRTKGGTGWTA
            .  *****.. **       *  ..** .*    **. .. *   . **   *    *

40          WGTRRKWGWKWYYMGGRRCCATCKKYKGKTAAAWTARRYKGYRRRWYATRYKWWAKYYKM
50          TTCARGWKTGWYYMATRGMYRWYTKTKKKCCGGTCTRGYGRYRRGTYWYRYGATTKYYKM
                * *   ****. *      .* * * ...  :* *  ***  *  **   :*****

40          GMGAACRYACYAAGAYGCAAAWYCRWWTRGTGARYARRWYKMAGYYRKKMWKKTCTKGGM
50          SMRTCYRYCAYTTRMYAMMCTWCTRWTWGCGAGGCWGGTYGMMRYYATGCTKKWMYKRKM
            .* :. **..*::  *.  .:*  **     ..      * *  ** .   **   *  *

40          TKSYKYWYKCGYRAWYKACCTRTRAAAGKKTWWRRYRWMACRGCKGYYGAAMATTWYRKR
50          YKGTTYATKARCRWWCKCYTYGWGTWWRKKYWWRRYRWMWMRRYGATYSRRMRCCWYRKR
             *. .*  *.  * * *.      :   ** ********  *   . *.  *   *****

40          TRKRYYYTGAYYRKKMRAYRSWKMCTRKKYMYKCCYYRGRTRRMSYGMKCYTYYYTTWRR
50          AGGGYYYARRYYGKKMRRYRSWKMTCRKKCMTKYMYYRSAKRRMGCRAGYYCYYCKCTAA
            :   ***:  ** **** ******  *** * *  ***. .***.     * ** .

40          AAKCKAKKRGCCTGYKCGGRKCMAWTRAWWTKYKWYYTTKKTTGCRTCSCATRYCCCTAA
50          WWKMKWKKRRSYCCCKTACAGMARWGRGTTWKTGACCKYKKKCATAYTGTTARYSMTGCT
              * * ***  .    * .      * *.   *     . **. .    . ::**.   .:

40          YMMRKYKRKYACKYKMGYRTCRGGACTYGKGGWYWRYTKATARRWKMRAKWTSAMRYCCT
50          TMMGTYKRTYWMKTKMRCGKYGRKRYYYKKRRWCWRYCKRYMGGAKCRGGWYSRARYTTC
             ** .***.*  * **   .       * *  * *** *      * *. * *  **

40          GKRCKKCAYKMCMTKGWYKAYSRKWCKMMKKYGKGAGAYRSGYYGWKRRMCWKKTTYTCC
50          AKAYGGYWCGCTCCKAACKGTSAKAMGCMKKTAKRCKMCRSACYATKGGMYWKKCACAMM
            .*            *.  *. * *    *** .* .   **. *. *  * *** : :

40          TRCGAYATAATTKRYYYRMGGTARYSRKTKYRTYWKKCKRMGAYKRYYGGKWGTGTATGR
50          CAYTGTMWWGAGKRCYCRCRRWRRYSRKWKYRKTTKKYKRMARTTATCATGTRCKYRKRA
                .    .: ** * *     ***** ***.  ** ***.  .   .        .

40          RYYWCGAGKYRAACYYSKKCYCTMYYRRAAKWAASWMSCYKYRYGKKTKYAYRMRKYKRY
50          GCYWYRRRKYRWWYYYSTKYCMGCTYRGCGGWCGSWCCYYKTRCKGKGTTRYRMRGCGRC
              **    ***   ***.*      ** .. *..** . ** *   * .  ****   *
```

```
40             GRRATCTRYWKKGYCKYTCKWCAYARMGRACTCAYYAKWYACKYYKKRRMRTCSYGAYRK
50             KGARWYCRYAKKRYMKTYMKWMMCGRCCRGACYRCYMGACTTKYCKKRAMACTCCACYAK
                      ** ** * *   **    .*  *..     *     : ** *** *    . ..* *

40             CGRYRRSWKGKTTKKRMCTAKRKKCTGMRCWGYRKTRTMYTTCACKKYSMKGACKGTTKW
50             TARYGRCAKTTCYKKRMSKWKRKGMYRCRYARYRKKRCCTKCTGMKKTGCGATYTAKCKW
                .** *. * .  ****.. ***      *    ***.*    .  . ** .   .: ... **

40             YCKWGTYMKRYYGGKCACTMYMYCCTGCCYTCKRKWRMWTAGTTRTCGGRAKTKKCKCTA
50             YYKWAYTATRYCKRGYCTWCCCCYYKATAYYGKRTTACAWGAGCRYYRKAWKGKTYKMYT
               * **.    .**      .          .. .*  **.       ..  *       * *. *  :

40             AKKKCSRGGRYYYTCGTCCTKKRYGKKYWTGAKAGCGAAKGAYKKKCAMAAGTKMKWKMC
50             MKGKTGAKKGYTTAARCYYCKTRCKKKYTWRRKGRYSWWKTWYKKGYRMRWSWKCKAKMT
                * * .     *  :.      *.*  ***     *.  .  *  ***    *  . * * **

40             KGYRMRRCGKKTRCACAGTCAARRWTSKSYGMSWTTARYATYRRRKGCACRATAWRRCTA
50             TACRMRRTAKKWRYCACACTGRAGAAGKSYRASWACTACWCYRRRGRTGGRGAGARGYYW
               .. **** .** * ....  .     :.***  **: :     ****    . *.:. *

40             CGKCKAGGGCCCKSKKGCMAGMTSYTGRKKKYYMYYCCMRTMRWAKKYCRTAACWMMKCG
50             TKKTKRRKRTAMTCTKSYMMAMASYYRRGKKTYMCCYTMAGCGTGKKYGAYRRYACMTAR
                 * *      . ...*. * .*:**  * ** **     *      .***         *..

40             KATCCCCCGYGKYYKAYAAGCGYWYMKKTKGGKRYWWKCAYYYYTGKCCCYYCGSYYYRY
50             KWCTATTGAYKKYYKCTRTRMAYWYCKKWKRKKRCWAGYMYYYYYTKMMMYYMASYYYGC
               *   .   .* ****.  :  .*** ** *  ** *     ****  *    ** .****

40             KWKWWAMYYRKCMAKRWGRGGTTAKRSTKKWRYRKSRGMKGCSKTACRACAKTRKWARGW
50             KAKAARMYYRKMMWKRWRGAACYCKAGYKKWRYRKSRKMKTYSGWWTAWAGKYGKTTGTW
               * *   ***** * ***   ..  .* . ********* **   *        ..*  * :  *

40             GRTAMYRYRSWYWKAMMRRRKKYKATYRKRMMMTACCYAWGGMRRKGWCTTGCGRKYGRW
50             SRKWCTACRCWYTKWMMRRAKKYKGAYAKGCCMWRTYYRWAKMGRKRTMGCCSAGGTRGA
               .*.       *.** * **** ****.:* *   *     * *. * **         ..

40             CGCRGWTGTATRYKCMYRCRWKCKAMKYWRKATWYTYAYYTTKKCGYGCRTSTACYYAYS
50             YKMRKWWRYMYGYGYCYRTRTKTTGMKCWGTWKWTCYGYYYWKGARCATRWSKRTYYGCS
                  * *       *    ** * * ..** * . .*  *.**  * .  . * *.  **. *

40             CMYRYATATAGGWTRYKYKKRCTWWCYKYKRRCRRYYCKYTCYTRWWYWMRYRAAGMYTM
50             YMYGTCKRYWSRAWGCTCKKGYWWWYYKYKRRYGGYYAKYYYCYRWTTTCAYRMGAACYM
                **  ..   .     . **   ** ******   **.**    **     ** ..   *

40             KAGKCAKCRYGTGTCMYYYYMWGYSRRYTGRYRSRGAMYYYYKKYRYYGMMRRWYCYSYC
50             GWRTMWGYACSKKWSMYYYYCARYSGATCARYRSRRRMYYYYKKYRYCAMMRRWTMTGTA
                  .       ..  .*****    **     .*****  ********** .*****    . .

40             WWWMGYTTTYWGKKYRGRMMTAASRCMMYYKMRCTTKWCKATTYKMKCRYMRYSMRWMKAR
50             TTTAACKGTTAKKTGAGMMCGTSRMCMTCGARTYCTTAKRYGTKMGAAYMRYSMRWMKWG
                    . .   .**  . ** .:**  *     *    . .*     ** . *********

40             GARCTCTRTGCGYKRRYKKKYAYCCTAMRYCGCAAAGCACAAYYRRKTGKKGTCCKYASR
```

```
50              RWRYCSCGCRMRYKARYKKKYTTTYCTMRYAAMGGTRTCTTGTTGGGYSKTKCTSGYGGG
                  *  .      ** ******:    :***.. ..:  . :.      .*.   . *..

40              RRTCKSTKTTYYMYSRTTAKCCAAYMRRKTWKKMYKRGCTACRKSMGTCYTRRRARYSKA
50              AGYYKSYKYCYYMYSRCKWKTAMWYAAAKYTTTMYKGRYWWMGGGCKWYYARRRMGCSKC
                    ** *  ****** . * .  *   *  ..***        .    *:***   **.

40              ARKYCTTTCTYYTTTCTCYCCTKGMTGGCRRKKSYAYMTYCKRYWRAMYMWWKKCWYTTR
50              WAKYSWYYYWCTYYYYYAYTAWKKMCTTYRGKGSYRYMYTYKRYWRRMYAWWGKTWYCCG
                  **.            .* . * *    * * ** **   ***** ** ** * **

40              KRYYCAYGKRGMASTTAKYKCYYKCWYRCGWKSKTGTWYRYRRCACKYRGAGAKATTKGW
50              GGTCYRTRKGAMWGWCGKCKSYYGMTYGGATKSGYRYATGYRRYRYKYRTGTMKMYYKKT
                        * .* .  .* *.**   *  . **       ***   *** .  *   *

40              GWTYYYYRMYAWYYSYCGTGMRTTTGRKKCTGYRKYMKCCWYRKAWKKKRWKAATGTAKG
50              RTYCCTTRMYTACTSYTACAARWKYKAKKGGAYATCAKYSTYRKRWKKTRWGMTCAKWKR
                       ***:   ** . . * .   **  .* .  * . *** ***.**  : .. *

40              WACGWWGAAACKMMWWTKKAYTCCRCGGATTKRWKCATAYKMYAATMTGKYMKAWMYMYR
50              ARYRAWRGGGYKMMWWGKKCYYYYAYKTGKWGRWKYRWMYKMCRMYAGAKTMKMWMYMYR
                     * ... ***** **.*       ..  ***    ***      .* ** ******

40              YCKCCYYYCTRAATGCGCWCAAACTTYTACCMWSCGATTWRYWGGRMKARMKTYTAGGWG
50              YTKYSYYTGYRWMYKMKYTTRRMTYCTCWMSMWSYRWYYAATTAARCTGAMKCYAWAATR
                * * .**   *                   .***         ..* .. ** *: ..

40              AMTGTACYRWRYATCCARKATATTTYRTATTKKYSWCAYKYRKAAACKWCYKKWRYCTKK
50              MAATCMYTGTGTGKTTGRTTATCCKCGCMWKTKYSWYRYKTGKGGTSKAMCKKTGCTGTK
                  :         ..  .*.:::  .     ..****  **  *..:.*   **     .*

40              MAYRGRCAGGKKKMYMKYRWWCYRTAKMYTYTKRSRYRKKMMTWTGGTTYTCAATTKKCK
50              MCCRKGTGASTKKMYMKTGWWYYRWMGMYKCKKGCGYRGKCCATASRWYTYYRRWWKTSG
                *. *   ....******  ** **   **. .* . ** *  : :.          *..

40              KAYGCWWKYYKRRRKMRKKRKKYKRKKA
50              KGCRYAWKYYGARATCRKKRKTCKRKKW
                *.    ****  * . *****. ****
```

The pairwise alignments of two accessions 40 and 51

```
40              KGYYWSYWRWCARSMYCRRGRYRKWRGTTRRACGWMKRCYAMYRYCRAMKAGGRYYRWRW
51              GKYYWSYWRWTTRCMTAGGRACGKWRRACAGTTCTCKRMTRMYRTMRGCKRRRRTYRWRW
                  ******** :*.* .      *** :   :    **   ***  *. *   * *****

40              AMYGSYCGGWRRRGKKYYRRRRMCYKWMRKAYWCCTYKARWMRWYWKWGSWKYYRKWACC
51              CCYAGYTTATGRGTKGYYRRRRMYTKTMGKRCWYTCYTMRWMGWYWKTKGTTTYGTWGTT
                . *..*  .  *  * *******  * * *  *   *. *** ****  . . * .*.

40              KRCRAWMMKWTARCCCKMWTYRYMGKKTAYKRCCGYWYYYSRGTRYYKKYYYYRYYYKRG
51              KRTRRWMMKWCTRATTKMWWCRCAKKGYWYKRYMRYTYYYCGRGGYCKKYYCYRCTYKAK
                ** * ***** :*.  ***  *   *   ***   * ***.    * **** **  **

40              CYRYYSYWWYRCRRCRCCYWAMMARRKRYRYGSRYKYMRYTGCCCWRKKRRRRKAKRYMY
51              YYRCCGTATYRYRRTAYYYTTCAMAAGRCGCKSACKYMRYYKMMAWGGKGRGRGWKRCAY
                 **  .   ** **    * :      *    *  *****    .*  * * *  **  *

40              YKRYRWWWYWSMMYMKRAMMGMCYYKYMYKYRGGTYCCYMKGCWWKCMCRSMYWYARRGY
51              YKRYRWAWYWSMMYMKARAARMYTCGTMTGYRRRKYMMYMTKAAWKYMMRSMYAYMARRT
                ****** *********     *     *  **  .*  **. . ** * **** *  *

40              MWYTWKMSTGCRMMWKYRWGYCTRRYWGCTRKKKKKYAGCKYYRARCKYYCTACRYRGGG
51              ATTCATCSCRTRCCWKYRTRCMWAGTWAMWGKKKKKYRRTKCTGTRYKCYYYWYGCRRAA
                     . *   *  ****        *.   ******   *   :* * *      * ..

40              RCRYYKMKAWCMYKYKKYWYYARGTKRMWKTSRRTYKCWRKATGRTKSCKKMKGTYATYR
51              GYRYYKMGRTAMYKYKKTTYYWARAGGAATKSRRCYGSARGTCAGCKGTKTMKTGTTYYR
                  *****   .******  **   :    ..*** * . * : .  *. *.**   : **

40              YSWKAYKYRKWCKKWCTYARYKKKARYRAMMKKMMYRATACTYRYMWKWRTAYKKSRKYY
51              YSWGRYKTGKAMTTTYCCTGCGTKCRCRRMCGKMMYGMWWYYYRYMWKWRWMCTKCGGTY
                ***  **  *  ..    :   .*.* * *  ****      ********   .*.   *

40              RKAGWYGTTYKKCAATTGWKKKGTKGYWWYMKACYCATYKRRRRMRKAYRAYKYKWRKKY
51              RKMKTCRWYTKGYRRWWRWKTGTGKTCAWYMKCTYACGYGARRRCRKWTGGCKYKWRKKY
                **        *       **.   *   ****. *.. *  *** **   . ********

40              RWKKMYRRYCRWKYWRYAAYCTYCTCATTWCKKRRTAYMSMAGCGYMYYYAGCARWKYTK
51              RATKAYGACTRTKCWRYRRYTCYYCYWCCWYKGRRYWCCGMTATAYMCTYGTTGRWKYAK
                * .* *    * * ***  *  *      * * **    .*:. .**  *.  .****:*

40              CYGCGKRGRYGKACCTYYYYTAAKKYACMYYYWKKRKWCTGKRYYYYRYARKWKWWYKWY
51              YCRMKKARRCRKGATWYYYYYTTKTYMMMYYYWKKRKATCCKRCYYYGCGGKAKAACKAY
                     *  *  *..  **** ::*.*  *********    ** ***  . * *   * *

40              WCMYKRWGRGAYKMWWYSKSKKWRYRKSRGCYYSKRRACAKTKYSRYKYKWARGAKAMRT
51              WMMYKRWRGRCYKAATTSKGKKWRYRKSRKMCTSGRAWMRKYKYSGCGTKTWGKGKWCGY
                * *****   .**    **.*********    * *    * ***    *    .*

40              CASTWYRWKAMMGGARKYRRTGGYTKYKRKYYRRYRYWYRYTACYTACCYYKMRMYWTGC
51              TRCYWYRTGWMMAAGAKYRRCRKYYKYKAKYYGRYRYWYRYWRYCYWYYYTGMRAYWCAA
```

```
        . ***   **... ****   * *** *** ********        *  ** ** ..

40    RWYYMKRKRAWSKTTTKCGRSWWYCGTTCKKRAAAKWKTYGCWRKTWRMTMYTTCAKTWC
51    ATYYAKRKRMTSKCCCGAARCAACGRWYSKKRGWMKWKAYKYAAKKWRMCCTKCTGGYWM
        ** ****  **    ..*.       .***.  ***:*    *.***   .  .  *

40    TCKKYSMMGGACKGYTKRWYAGGYTYACGTCMGGYYKTCWKTWTAYMRKWWKRAARKGYT
51    YMKKTGCMRRWYTACKKRWYGRRCYYCTCWYCKRCCKYGTKCACTYMRTTWTGGRRKRCK
        ** . *    .. .****.    *.         *   *   :***. *. . **  .

40    TKTAGAATCMCATGKYRWMRKKCAKTSRTGCYYYYCYAKKCKKYKKRYKRYTKWTGAYCC
51    GKCRRGRCYMMTATKTRWMGGKTTGYGAAATYYTTTCCGTYKTYGKRYKRCKKTAARCTA
       *   .   * :: * ***  * :  . :. **    . . *.* ***** .* :.   .

40    KKGAYKWGKCAATWRTATAGCGCGTATKRRRWWYYGGGTAAYARATCAWMKTKRMRRGGR
51    TKTWYKARKYMMWTRWTCCAAAMAKMKKGGATATTKKRCTTTGGRYMWACKKTRCRGTRG
      .*  **  *     * : .... .. .*           :: .       *..* *

40    TCGRKMCTKRMKSGTTARYYYAKKRMGATRWKGYKACKKWTKCRYCMYRMMAGCMYAYWT
51    CYRRKMTAKRMKGRYYRACYCWKKRMTGGGTKAYKWYKKACKTRCTMYRMMCKTMYRYTG
         *** :****.      *  **** .   *.**  **  * *  *****.  ** *

40    ATKGKYRKRYWTYCCCRATAACSRRYTWKAYYGTTTKTAGWAWTAAYRGAYMKYATYCTY
51    GWGRGTGKACWYYATTRTCRTYSRRYWWKTYCTAKKKARSAMAATCTARTTMKYCCYAAC
      .      *  * *.  *:  : **** **:*  :..*: .   ::.   : ***. *.:

40    RSKRTTTAMKRYYKYKRCKKTGAWYYGTYCRRTCTAGYKRYMKCTTRRYRRRMTRTTKCK
51    ASTAYWYRMKRYYKYGAMKTCACTCTKYYAARWYCRRYGGCCGYYARRYRRRMYAGAKTK
       *.     *******   *. ..     *. *     *       :*******   :* *

40    YKRRKTKWKGYRSTYTKAYRKTRKYYRMGGTAWRRYRCYKWTKCTTYCAYKRYWKWCGTA
51    YKGGKCKWKSYRSCYAKGTAKWRKCCRCRRWRARRYRYTKTKKGYYYARTTATWGTYRYR
      **  * ***.*** *:*.  * **  *      ****  * .*   *.  .  *

40    TGRRYGYWCRCKKAYTCYYMYYCYYAACWSCYTAGRKTYTACATGCYGKKYKYATAYRMR
51    KRAGCRYWYRYTKRCGAYCCYYYYYCGAWCYYCWARKYCKMMRKSMCTKTTKYRKGYRMR
      .     ** * .*   .*  ** **...*. *  .**  .   ..   *. ** ..****

40    RYRCWRTYKKRGCKYYKAYRYRYSYCYACCKAATAGGTRKWYGYGYRTAKACKYKGCYRM
51    RTATWGCYKKRRMKTYKRTRCGYSYMCRMSGCGWGAACGGACKTATGAGKWYKCKRYCAA
      *   *  ****  * **  *  ***    . .. ...       .  :.*  * *

40    YGAKAAAKYYYCTAMYYMMYSMYKMKWYKMYRTKCCTTTYCGSRWRYKCRATTTYYGTCG
51    CACKTWWKYYCYYRCYYMMYCMYKMKWYKCYAKKMYYYWYYRGATGYGTRMKYWYTKYYA
       ..*:  ***     *****.******** * .*     *  .   *  * .  *    .

40    AGKYRMGATMYRAARATTAAACCGGAACCRGSMRRTRKACAKYWYAYRKATTYRTYGAWG
51    RKKYGMRMKMTARRGCKYRGMSYATCGSMAAGAAGGRKRYWKYWTRTRKCCKCGWTTWWR
        ** *  .*     ..  . . . ...  ..    **   ***   **. .      *

40    GCKYWCYGCKAMRYYKYYCATAKWYRSWYRYMTYKGGAMAGTKAKYWSTAGAKKMCGCAC
51    KYKYTYCRMKGMRYYKCCAWWRKAYAGACGYCKCGRSWMTKYKMKYWSYRKRKKMMRTGY
        **     *.*****  .   * * .   * .   . *:  * ****    ***   .
```

```
40          TATAYKGKCCGYKCKRMKCKRKKTTKGTGSCCCYYSRYKYTCCGTKRWRMRRYCYKGSWK
51          KRYRTTATTTCCKGGAAGSTGKKYWTRWSSYMYTTGGTGCCTTAGKRARAAATYYKASWK
            .    ...    *     .. **  .  .*     .       . ** *     **.***

40          RYKGCGTRMYRKKYCRYRYKKMSRYRKMGGTYKCCRSKYCTYRYCGTTKKKCYKKYRYKA
51          RYKTTKGGMYRKKCAGYACKKMSRTGKMSKWTTTGAGKCYYCGCMKWCKKKYTKKTGCKG
            ***     ***** . *  *****  **.   .   .*          ***  **   *.

40          TGKYYWWTGYKRYYMSKRMYRYSKTKRYYRMWRTAKTTAMRACACYKYMTYRACKAKYYR
51          YRKYYWTWKYKAYYMSKRCYRCGKWKACYRMWRCCKCCGCRWYMYCKCCWYRRYKGKCYG
              ****   ** ****** ** .* *  ***** .*  . *     *   **  *.* *

40          WMMGGRKTARRSCAMRTAGKGRATTSMKRRWGKKAMWWYWMRKYRTWRKWKKWGCCKKKK
51          TMMRKRKCGAGGYRMGWGRKRGMKWSMKAGWTKKCCTWTAAATTGATRGWKKWKYMTKKK
             **  ** .  .  *  . *   . ***  * **.  *    .  : * ****   .***

40          CMRGRMYGTCTRWGCATGRAGKWAYYKMYRMKRMKMRKCRSWKACGTCKCKRTMKYKKWC
51          AMRCGMTRYSGGAAYRCRGWRKWWYYKCTAAKRCKMRKYGCWKCGSGMKTKRGMKCKTWM
            .**  *   .   .       ** ***    ** ****  .**. .  * ** ** *.*

40          YTMMYWYYCGYAGRTAYCRAGRATGATGAAGGWTAKYKCRCTGRGATATWCAGWCYYKRR
51          TYAACAYYARCRARKRCYGGSRTCCRCSRGCAAAWTCKAGYATGATCGYWYWKWYYYKRR
                  **.   .*.    ..*:    . . . : . *.  :  .: . *   * *****

40          RRTCAYTGYGYRYRYYAGGCMKRKMGKYRRGCACATKRGTGTSYKYTCGGKYTMTYRYKK
51          RRGMMCGTTKCRCATCGAAMMKRGCRGYRRRMWYRWKASKKWSCGYKYRRKYGCGCRYKK
            **         *    ... ***    ***      *  ..  *  *.   **    ****

40          YYRWRYRRRWAGTGCTKKKRRMCCTKMCGMKGRKMMWYGMYRYACKRRYYRWCRKTGYKT
51          CYAWRYRRAWRRKRYAKKKRRMMTYKCMRMKARKMMAYACYRYCYKRAYTAAMRKCRCKY
             * ***** *  .  :******   *   **.**** *. ***. ** *    **   *

40          KAGKMTMRTMYACGCKCWWSKKRKWYWCKGWCCYMYYAGTWWGKKAYCRSSSCRYMMYTR
51          GWRKCWMRCCCGYRMKYWWSGTRKATTTGSTYYYMCCGRCTAKKGGYGGGGGYGTMMYKA
               *  **   .   * *** .**     .   **  .     * .*  ...   ***.

40          TYMAKTYYYCYTAMWTGGKYTAGTRYRKAMATTAYMRMATRRCGYYARRRRMMRMTARKC
51          YTAWGGYYYSCWWCTCAAKTCRSCACAKGCTCYRCCAMMYRAMSTYWAAAGAMRMYRAKM
                  ***.      ..*   .    *. :      *  *  . *      ***   *

40          KMWKTCATWCKKWRYYMKMMMYRWKKRRRKTKYYTYTKRYWYKMRYSYRYRRWMYYYSGR
51          KAWKKTGKWAKKWRYYMKAMMTRWKKAGGKCKTYCCKKRYWYKMRTGTRCGRWMYYYSKG
            * **. ..*.******** ** ****   * * *  .******** . *  *******

40          YTKKKKCRKKYCTGKGYRAYYRYAKGGKRRGWKWKKYYMTRKAYRTWMKYYRTTMRRKGK
51          YAGKKKTRKKYAYRTAYRMTCGYGKATTAARWGWKGCTMYGGRYRGAMKYYGYCMGRKRG
            *: *** ****.  ..**    *.*. .   * **   *    **  ****   * **

40          YKTAKRYKAKYKWYMYYMKRMRRKGRGCRYGKYMYWGAARYRACKTGKCKGAATAGKATT
51          YGWWTGYTMTYKTTATCCKRMAGKKRRYRYAGYMYWKRWRYRRMKARKTKRRWWRTKWCA
            *   . *. .**      ***  * *  **. ****   ***  *: * *      *  :
```

```
40          CKGSYKRSKTKCCSRKSRCKCYSYWYCAAKSRWMAGCTYKYYYAYKYGAKGRKWRAGGKT
51          TKTSYKRSKKKTYSRKCATKTCSYWYYRMKSRWMRSMAYKYYYGYKYKMKKRGTGRRKKY
             * ******.*  ***.  *  ****   ***** . :*****.***  * *      *

40          CKWKYMGCATCAKTGGAGKCAATCCTGRCARSYKYWYYAYRWKCKWWRRMACKRYKTCTY
51          TKTKCARGRCMGKWCACAKGTGCMYKRRYRAGTTTATCWCGAGYKWWRRMWMKRYKCACC
             * *       .*  ...* :.   . *   . .          ******  **** .

40          WYRRTWTRRGTWKYYYRYRKKYRKSTSRKRRRRKRYYTKCYMKRTRMKMMYKTMGYTKTR
51          WYRGCAWRGRWWGYYYGYGKKTAKCGCRKGAGGKRCYYKYYMGAKRMKAMYKYCRCKKCA
            ***    *   * *** * **  *. .**    ** * * **  .*** ***    .*

40          WKCAKGGTYCGGTRRWRMRCGTTTWKTKACYRCYCMGRCTCAGATYCMYKWYRMYRKRRT
51          WGMWKRAYCTACKAGWRCGMKGYGTKAKMAYGTCTCAGAAMMRGWTTATGWYRMYRKRRC
            *   * .   . .  **        *:* .*     . .:   .      *********

40          AYMRYKATCTGMGGCWAATTGAACRCRRWKSRRCCWGGRWKYWTGTTTGAAKTTATKMGG
51          MTAGCTCATATMKAYTWWGGRCTARYARWKGGRMYWRRRAGYWYRYAARWRKCCRYGARK
                 ..: : * .       .:.*  ***. *  *  *  **   ::   *

40          CGKGAGYKRKYTRRATCCYMTYTKTAYGWMTGRYYRWWGWYTAMGRRKKACWRYCAWWGR
51          MTKKGAYKRGCYGRRYYYYAWTYTAGTRTAGTGYYRTAKACCRACARKGRYARYYRWAKG
              * ..***    *    *    .:.       ***          **    **  *

40          KKCYWRKCASGTKRYYGRCARGTMTATTATYAKCGGAACWWRSYARAGYKYMGGWCKCCA
51          KKYYWRKTTCRWKGYYKRMRGRCACGACGCYGGYARTCYWWGGYMRMKYTCCATWMTYMR
            ** **** :.  * ** *       .: . *.  . :. ** .* *  *.  . * .

40          RTGAKKTGTTCYRWRWCKAGTRKKMWATAATCAAGCCGMYYYMWGSARTGYTYARYYYCC
51          GGAGKGYKKYACGTRTTKGKCATGCTCWMRGMGGAYMTMYYYMWRSCGCRTCCCRYYYTA
              ..*   . .   *  *.   .   .     ...   ****** *.      .**** .

40          YKRYRYSWRRRYCGGCYYMYGTRYWAWYRGGRCCGYAKTAAMSRMKWTAARRRYYTRMKC
51          YKRYRYSWRRRTYRAMYYMCKGATTRWTGARGMAAYTKCGTMCRCTTACGRRRYTCRMGA
            ***********   . ***       *  .   ..*:* .:*.* . :..****  ** .

40          RYKYYMRYGRKWMKRCMYMYMRKTRWTWTTCKRYKAMCAYACYGYYRRTKKRGRCKKGTS
51          AYKYYARYKATACKRGMCMYMRKCAACAGAYKRYKWMMRTGACCYYRRKKKGAGYKGCAS
             **** **  .  ** * *****      : **** *   ..  ****.** .  *  :*

40          KRYKYKGRYMMMAKRKWSKYACCCGSWCGRSKTGTATCKAWYRYRTKSCMGGYRRYRCAC
51          KRYKCGKRYACCGKGKAGTYGYMGACTMKRSKCRYRWSGMATGYRKGCACRKYRRYRYRY
            ****   **   .* * ..*.   ..   ***     .     **. ..   *****

40          KYCGYRGAGAYYSTCYYCKGWGWKYYKYRYYATMTAGAGRTACKCCYRKYMCYWKWTGYK
51          KTYRYRKRKMTYSYYYYSTKTRTGTTKCGCCRKAWTKCKAYRYKYYYRKYMSTTTTCSCK
            *   **     **  **..       *     .  : .     *  *****.  .  . *

40          AWCKCWATTKKGGWAKCGRATGWCTGRAATTRCWYRGCCAGYKCAKACACCWMYRRTKCA
51          RWYKYWTYKKKARARKYKRCCRWYYRRRRKWRYAYRKYTGRCKTGKCYWYYTCCAAWKTG
             * * *: .**.   *  *.  *   *  . *  **   .  *.*.          * .

40          RYRKGWRMYAYCKKYKRMYCKKRKGTATAKRAWCRKAWGTYRYMRGCKWGWYARGGKYGK
```

```
51          RYRGAWRMYRYTKKYKRMYAKKGKTACCMKAGTSGKRWRAYRYMARMKAKWYRGKKKYRK
            *** .**** * *******.** * :.  * . . * * :****   * **     ** *

40          AGGTKTTCTGRYKKKSAWRKKCKARRWTGGTAAACTYCRCTMTCKKAKGWYKRCCMWCRR
51          RKRAKCAAGCRYTKKGGWRGTMKCGGWARAYMRRYYCMGTKAAYKKWKKTYKRYMMAYAA
               :* :.  **.**..** . *.  *: .          . : ** *  ***  *

40          WKWYMAAGGKACKYRKMKKAKGCRGTCRWRCCRCTGGTYYKWWRKKKKKKTGKYRMCAMR
51          WTTTMTGTAKGTKYRKMKKTGAMRAATGTAAYGYATAYYYGWWRKKKKKKCKTCGMATCA
            *.  *:. .*. *******: . *.:    .   : . ** *********  .  *.:

40          MYYMRWTAGTKGKTAGAKYYMYYSRRTTGKAKKYGYMRCKMCGYKYRRYRKSMTTTCAMK
51          MYYAGWYGKWKKKWRRTKYYMYYSRRWYRKWTKYRYMAMTMTKYGCGRCGGGCYYWYWCT
            ***  * .  * *   :*********   * .** **  .*  *   *   .       .

40          AWARKAACGTGAYCTCTYYKYYKCCCRKGMARYRGTGYWMSYYMSTKRWARYMWYTGTTT
51          RWMGKCMYRWAGYSYYWCTKYTKTAMRKKMTRCGRGRYWCSYYMGCKRWRGYAATATCGC
             *  *.    ..*.     ** * . ** *:*     ** ****. ***  *   :

40          YRRACTATGAKAKKMKCATRAGTWKYKSAATTRCCGTCCACTCTCGYGTGKGTSYKTACC
51          YGGRYYGCAGKWKKCGAGAGGAYWKTKGCGGWRMYAKTGMMYTAAKCAAATKYSYKYRYT
            *     . ..* **  ..: .. ** *...  *  ..      :.  .:..  ***

40          ATWCWCATTCTMWACMCWWSYTMKTCYRMRTWGCCCYMKMWACKWTWKMKKTKAAAAGGA
51          CATYTYGYKYYMWWMMMWWSTCMKYSYRAACATTTACMKATWYGWWWKCKKWKCGGRKTT
            .:    . .  **  * ***  ** .**       . **     * ** ** *...   :

40          YKARGMYRTATRMRRCKTRCWKTCCMSKYGMSWCTRYRAAGTAAAWCCCRTTCKKRKSKG
51          YKMARCCAWRKRMRRTKAATWKWYMCSKYRASWYCAYRGTAWGGGAYTAAACAKKRTCTS
            **        .**** *:  **    ***  **   **.:. ...   . : .***....

40          CGAARTRTGKRKMTAACTGMRWYYSCGGCRTCMACATATAKKYYTCCCCCGYGYWWGTYK
51          YKCTRARYRGRKCYCGYCRMRWYTSGKKMAYYMGMMWGCGKKYYCTATMGAYKYWWRYYT
              .:*:*   **  ..   **** *       *.   . .****  .   .* ***  *.

40          YRAYAGMGTKKTYKYRCKYAMMYKRKTYCKKAAYKMYCCTCCKYYWRYTTRRCYYCCWMR
51          YRRYCAATYGKKYKCAYKYTMMYKRTYYMKGWRYKACSYWMYKCYWRYCYGGYCTTTACR
            ** *..    *.**   **:*****. * *   **  .    * ****           *

40          AKWSAKRYYYKKRCWRGYYCAYMMCATKTYSRRKRKKMMAGAAKCYGKGGRSYWKY
51          GGWSRKACTYKKAYWRRYCYWCAAYTAKWYSRRKAKGCMRRMRKMTAKRKRSCAKY
            . ** *   ***  ** *       ::* ***** *  *    *  .*  **  **
```

The pairwise alignments of two accessions 42 and 43

```
43              RRRMRRKRCRTGGRWRKYRGYWYWKMCWMCGRGKKRKWYRWWMKRTYWKWKGCCYRYYYW
42              RRRMRGKRMGYACRWRKYRAYWTATCYACYSRRKKRKWCAWWMKRAYWKWGKYTYRYYYW
                ***** **   . ******.**  .     .* *****  *****:****    ******

43              SWWCYYWTCTGYKYRRYRRMGYKKRWRYWMYWYTCCWYKCTRYWKSYYKRGWKRWYYRWW
42              SWWMYYAYMWTYKYRRTRRMACGKRWGCWMCWYKMTAYKSYRYWKSTYKARWKRWYYRWW
                *** **     ***** ***.  ***  ** **.   **. ***** **  *********

43              RRRKYRKMWRWKKYYRKYWRYMRCCTTYCRRTAKYTYRKRMMGMSCCCKTATYSRYYYRM
42              RRRKYRKMWRWKKCCRTTAGYMRYYYWCARGYRKTYYRKRMMAMCMTYKGTAYSRYYYRM
                *************  *.   ***     .*   *  ******.*.   * ::********

43              WAAWRMRRKMCYRRMSKMYMYRKWCYARMRYYTMKMYYRKWRWWARAWMRRYSRWMMMGW
42              WGGTRAARKMYYRGACKMYMYRKWYYTRMGYYGMKCCYRKWRWWWRRTCARYSRWMMMAA
                *.. *  *** **  .******** *:** ** **  ******* *    ********.

43              YRYRYYRRKTWKMCRGACAYKACKTCGKKYTKACKCTAKWYWAKRWRCRTGWRRMCWRYK
42              CRCRYYRRKYWKCYGAGTGYKMYKYMRKKYKTWYGYKRKWYTRKRWRYRYAARRMYTRYK
                 * ****** **   .. .**  *   ***..    . ***  **** * . ***  ***

43              TRMMRWYGAMMYYGYTTAKRGMCKRRMRYKARYYYYRKCCCKRYAGASMMKWYKRRKCAM
42              WRMMRWCKRMMCTRYYKRKRRCTKRRMRYKWRTYYYRKYYYKRYTAGSMMKWYKRRKMCC
                 *****   **   * . **   ******* * *****   ***:..********** .

43              CYYGMARSKKGTWKWGRWWMYRARAKGYYGRCKYARYYGKKRMMKKRCKYTCWWYYGYCW
42              YCTTCWRSKTCWWKWKRWWMYRGRWKRYYKGYKTWRYTKKKRAMKKRYKYYMWWYYRYYW
                      ***.  *** ******.* * **   *  **  *** **** **  **** * *

43              KRGRRRKKYGTKTCARKAAGARKRRRRYTGYMMYAATTATKRMKRRGCRKTCCKGKTKSK
42              TRRRRRKKYSKKCTRGKRRRRRKRGGGCGAYMMYRRYYWYKRMKRRRYRKKSMKRKWKSK
                .* ******..*    *    ***     .****      ******  **.. * * ***

43              KCAAKRCGACARKMATGAGRATTMTCRYRYRMKCRCMTMGKYRKKYKKYKYTMRRYKKCA
42              KTGGKGTAWGRRKMMCAGRGTKKMWYGCRYGMKYGYMAMRKYRGKYKGYKYWMRRYKGMG
                * ..*   .  ***  ..  :..*    ** **   *:* *** *** *** *****  .

43              AYYGCRMRYKYYKKKKKRYKKSSCYGGWYWMKCGRSYGSYMMMKKYYRWYACRYKRAMYK
42              CTYRAACAYKYYKKGKKRYKKSSYYRKWYWMKTARSYACYMMMKKYYRWYGYRYKRMMCK
                . * .   ****** ******** *  ***** .***..***********. **** * *

43              KRYGATYMGYKWARCMKYKAGMMKCYTKRSRKCMCAYGGRARYMTRAACCKATTAWTTAT
42              KRYTGAYMRYKTRGYMKTKTKCMKMYKKRSRKYMYRYKRRWGTCCAMRYMKCCCGWCCCC
                *** .:** **    ** *:  ** *.***** *  *  *          *.  .*  .

43              RRWMYAYMKTTAAYRKKKTAAGWKGTRRKKRKKYCKKMYKCYCRRMRYRMKCTCKTCCCT
42              GRWMCRYMKKWRRYRKKKCRWRWKRYRRKTRKKYTKKMYKYTMRRMRTGCKYCYGYMAYA
                 ***  ***.   *****    **  ***.**** *****   ****   *      . :

43              RRCGGAKSRKCGGGKYKYGGCTAAKAGKRRWRCGGRTKMGRKKRYWCYCRKKYYWKYTKR
42              RRSRRRGSRGYKAAKTKYRSSGGCKGKKRRWRYTKGYKMRRKKRYAYYYRKGYYWKYYKR
```

```
                    **.    **   ..* ** .. ..*. *****     ** *****  * ** ***** **

43                  YTSAKKTKKYATKACKKMKKRWMAGGKKGTMYRKYKKAAGCCYYSYWKKATGSRKARKKY
42                  YYSMKKYKKCGAKMMKKMKKRWMWTKKKKKMYRKYKKMRRYYYYSYWKKWWRSRKRRKKY
                    * * ** ** .:*  ********   ** .*******     *******   *** ****

43                  YKKKYTRCYKKKTRYGTTKYWTTAYMCAACKCRRGWKYKKYRYYTRRYRMMYYATWKRKG
42                  YKKKYYRTYKGKGAYRYYTCWYKMCCSRMYKYRRKATYKKYRYYWRRYRMMYYRYWKGGR
                    ***** * ** *  *   . * .   .   * **  .******* ********  **

43                  WRCTTKRYYYYYAGGTKAKCTKYYYRKYCTCYYKAAKYYWYSRSYWYAAYRRRWYMAMYG
42                  WRACGTACYYYYWRRWGRKMYTYYYGKCAYMCYKRRGYYWYCRSYWYMWYRRAATMMMYR
                    **.  .  ****      *  .*** * .   **   ****.*****  ***   * **

43                  AKGWTTMRSCRWRYCWRWRYWCAGMRRYRYMARYGKYWKRTGTGAAKYWATAMYYKRCTT
42                  WKRWYKMRSMGWRYYWRWRCWYRRMRRYRYMRGCRKYWKRKKYRMRKYWCWMMYYKRMGC
                     * * .***  *** **** *   *******    *****.     ***.  *****

43                  AARMRRTYTKGGGKAKGKKGGTCYTYCRYGYGATCGCTMYRCGAGCTTTAARRGKYRRKR
42                  CWRMRGCYKKRSAKRGRKKACAATCCGGYRYAGGYAYYCYRYRRRYKKYGMRGRKYGGKR
                    . ***  *.* ..*   **. :.     * *..  .   **     .. . *   **  **

43                  KKRYYCMTWTYKKAAAAKGWYCTRRYMATRCYGWKYRMTGAYTRRKYKMKKWRTKKYAAG
42                  GTGYYYCCWKYKKMRRWKKWYACRGTCCYGTTAAKYRMYAGCWAGKYKMKKWGAKKYCGT
                     . **   *.***    * **. *   .    . **** ..    ******* :***..

43                  AAACGMYYYAASRRGAYSYYRGAGTGTTASWYRAATYRYRKMMWYYRRMKYCYYKYKMYK
42                  TGRMSMYYYRWSGARRCCCCGRWRGCWCTSWYRWMYTATAKMMWCCRRMKYYTTGTKMYK
                    :.  .****  *     .          :****       ****   *****    ****

43                  GAMKTKTYGYYKMRGAYYGKSCRSMGYSYKKKKKSCCRWRRRKRMMRYMKCCTKKGCCWR
42                  KCMKCKCYRYYKCGCCYCAKSSRSMSYSTKGKKKCMYGAGRRKRMMRTCKYYKKKKMYWR
                     .** * * ***   .* .**.***.** * ***.     *******  *  .**   **

43                  MCKWKWKCGCAKYYYTGGWYKMGYTKYGKKRKYSSCKYTKYKKTGTYGYSRYTGCAKKMY
42                  MYTWKWKYRYMKYYYKKRWYKMAYYKYRKKRGYSSMKYATYTKYRCYSYSRYYRMCKKMY
                    * .****    ****.  ****.* ** *** *** **:.*.*   *.****   .****

43                  WTCAARKACTRRRSAACKGRYARMYMKCACAKAAYYYAYWTMCRKRYCKRMYGRYCRMRA
42                  WWMWMRKRMWRRRSGGAKRGCCRMYMKMTTTKRRYYYWYWGCYGGRCTKRMYRRYYRCGM
                    *    **   ****...*   .***** : :*  *** **     *  **** ** *

43                  KYWTRGAKMMYTYTKGYMKKTTGAWGYYRMGGMKRGWKKRYRYYYYMWYWKKKCTGWMYG
42                  GYWYRRRKCMYWCWKKYMKKYYRWWKYYRCRRMKRKWGKRYRYYYCAATWTTKTCAWMYK
                     ** *  * **   * ****    * ***   *** * *******   *..*  .***

43                  GATTYYYTRYYKMKYAMYYYTYYMRKYKKMRGWMKKRKRTATTMRCWYKYMMKYYTMYRY
42                  TGAAYYYGACYKMKYRMYYYYYTMRKYKKMGTTMKKRKRYMYCMRYWYKYMMKYTYAYRY
                     .::***   ***** **** * *******   ******    ** ********   ***

43                  MYTCCARKRYWKMYYYWGKWKRATTKKKRKTWMGGRKWRYRKKGYRYKARMRAYTYKKCC
42                  MYWMMRRKRYAKCYYYTAKWKAMWYKKKRGCWMSRRKWRYRKKSYRTKWGARMYCCTKYY
                    **    **** * *** .***    ****  **. ********.** *   * *  .*
```

```
43                    TKGCGWGGYGCKKRGGMGWRSKWMGATMYCCYRTTYRYRKKRCTARRRGGRRCYCCYYYK
42                    YKKYRWSRCRTKKAKAAAWRSKWCRGGCCTTYRAKYRYRKKRTCGRRRRKGRGCYTYYYK
                       *   *.    **  . .***** .      **:.*******  .***   *    ****

43                    RRWYRKAGKCRCTYGTGACKRMAARWRMCRRAKYWGYYRYMCGTRCAWKACTCWYRRCAY
42                    RRWYRKRRKYRYYTAYRMMKRMWRRWRMTAATKYWKYYRYMYRYRYWTGTAYYWYRRGRY
                      ******  * *   .    ***  ****   :*** *****   *    :.  ****  *

43                    TGKKTAKYKTCGCRCAGTTAMMGATKYCTAYSGTAAAAKRWKYTWCCGYKRAAAYCCAAR
42                    YRKTWRKTKYMRYRMMRYYRMMKWYKYYGMYSCGCTCRGRWKYCATTAYKRMGWYYYRRR
                        *.  * *    *      **   **   **  .:.  ****    .*** . *    *

43                    YSYYAYKAGGAYYRKYKKCYYMMKRYCTTTKGAYYYASRYKYGKMYRGYKYRMCRWAYAR
42                    YSYYMTTMRKRYYRKYKGTYCMMKRYYYYWKSRYCCGGRYKYRKMYRKYKYRMSRWRYWR
                      ****  .    ******  * *****    *. *  ..**** **** *****.** * *

43                    RYKKRCTCACSGCKRTWWTKMTATAKAATRKTCTTGCWRYMMTKMRRWGCGCGWYMCGRW
42                    RYKKGYYYWSGCTKRYAWCKMWWKWGCTYRKYYYYKYWRYMCKKMRRWKMRTAWTAGRRW
                      ****     ..  **  * **  .  .: **      **** .*****    .*    **

43                    KCWMKAARCMMKKKMWMRTKMRGKGWRGKGAATKYTKYRMRKCRMRMRRWYKRKCGMATY
42                    KYWMKRRRYMMGKKMWMRYGMRRKRARRKRMWKKYWKYAMGKYRMRMRRWYKRGAAAMYC
                      * ***  * ** ******  ** *  * *   .** ** * * ********** ..

43                    TTKAGYYKYKTWKKKYKWRMKYYWMMWWARRYAGMKTRTGKTRYWYMTCKCRRRMYCGAA
42                    CYKTKYYKYKKWKKKYKWRMKTYWMMWWTRRYCTAKKRYKKKRYWYMWYKMRRRMYYKRR
                        *: *****.********** ******:***.  *.*  *.*****  * *****

43                    KARKKYYYYRRKMKWGKRCRAKRCMYCAKWAKCGTKRKGWRGGWRSYGKYKRKKTTGAYY
42                    KGRKKCYCYRRKMKTRGGGRGKRYMYTMKWCKYTGKRKKWGKRWRGTRKYKRKGYYRRTT
                      *.*** * ******     *.** **  **.*   *** *   **.  *****

43                    YCTKARKAYRGKTYGKTMKKYKRAAAAWRYTYKYKTCACACYCAMMMRTYYCRYCTGYWG
42                    YYYKCGTTCGKKYYKKCCTKYKRRRRRWRYKYKYKCGGYCMYSRCAMRYYYMRCYKRYWK
                      *  *. .:   * * *  .****    ***.****  . . *.   ** ** *  . **

43                    TGYRAKAWKATYMYWRTKRYTWMCRMYSYMTKCTAKMTYMYGYWRMSRCKKCTAWKCGRM
42                    YRTRMKTWKRYCMCWRKKRYAWMYRMYSYMYKYYRKCGYMCKYTRCGAAKKYYMWKYRRM
                         * *:**   * **.***:** ****** *   *  **  * * . .**   **  **

43                    TAGAATAYYRWYWYTYKWGCGATYTAYMYKSMRAAYRKWCTCAGYKKGTRYKRKGTTMAT
42                    KRSMCYRTYRWYACYCKAAGRRKYKRYMYKSMRRMYGGAYYYRKYTGKYRYKRKRKYCRY
                      . . .   ****    * .   .*. *******  *        *.   ***** .

43                    TGKAWWCMMYATSTKYMKMGGKMCCKCYMRYYTAATYYAGACCYYCYRGRMGMRAYYYRY
42                    YKKWWWMAMYCKCKKYMKMKKKMYMKMCMRYYKMMKYYRRCYYYYYCRRRMKMRRYYYRY
                        * **  **....*****  **  *  ****.  .**  .  **  * ** ** *****

43                    TMMKYCGGACSKAWRYRGGWWACRCYKMKWWMKAAATKYMRWRRKATTTKYGTKKKCGRW
42                    KMMKCMSRRYSKMWRYRRAWWGYRYTTATWWMKGMRYKYMRWRRKRWWYKYACKKKMRRW
                      .***  .   ** **** .**. *  . .****.   ********    **. ***  **
```

```
43          MRGGSSMKAKRAATTAMYWGCCACKRGGGAWCCTRKGCGTKAKMMGKYKCYYKKTCRRKY
42          CRRRSSMKMKRRMYYRMYWAMYGYTGAATGWMYKGKKAKAKRKCCRKYGMYYKGKYRRTY
             *  **** **     ***.  . . .. .*  . * . :* *   **  *** . **.*

43          GCRYCYKMYGAKGWRGRYKKRKTAYYAGKYMRRRCYTRYTRTCMRKKKYKTTYRRRRSWR
42          KYGYYYKMYRRKRAGRRCKKRKKRYYRKKYMRRRMCWRYKRYMMRGKKYKYKYRRRRSTR
               * ****  *    * ****. **  ******   **.*  ** **** .****** *

43          YARTKKYKYTWYMGGYRYCAGYWMTGCRAWYGTKMKRKKATCCKTYTRMYYWKKKATSYR
42          YTRWKKYGYKTYMRRYGCYRKTTCYRMRMWYRYKMKRKKWWYSKYCWRMYYWKKKRWSYG
            *:* *** *. **  *           * **  ******   .*   ********  **

43          CKKRKKGYKYRC
42          MKKRKKKYKYRY
             ***** ****
```

The pairwise alignments of two accessions 42 and 44

```
42                AMMYYCCAWYRYWGCYWKKYSYKYKTMSYAYWKCRRMRMYKRYKMMKRMWTRRWWKYMRK
44                MMAYYTTGWYRYWTMYWKKCSCGTKWMSYMYWKYRRMRMYKRYTAATGATYGRATKCAAG
                   * **  .*****  **** *   * *** *** *********.  .     *  *

42                GGGWYRYKKSRKYRKKSRRRRMYWKYYWWWYRYSKWWCMMKYKYKYYRRMKYKSYMRYKK
44                RARTYGTKKSGTTGKKGGAGRMYTKYYWAWYRYSKTTYCCTCKCKCTRRCTYKSYMRYKK
                   .  *  *** .  **.    *** **** ******     . * *  ** .*********

42                RWYTKRYYKRMRCARYYYYRRRRTRSTYRYWYRCRRRYYYYYGYGMMKRYYYYMYKYGRM
44                RWCAKAYTKAMRMRRYYYYRRARKRGWYRYWYRYRRRYYCCYRYRCMKRYYYYMYKYKAC
                  ** :* * * **  ******* *.*. ****** *****  * *  ***********

42                RYKWGYMMWRMGYRKMYWRCGYYRKYKCTKYKRYRWRYWRKWYKKRGWRTYYRYKRYSSG
44                RYKWRYMMWRMRCRKCTWRAKCCRKTTSWKYKRYATGCAAKWYGKRRAGKYYRYKRYSST
                  **** ******  **  **.   **  .. *****      *** **   .*********

42                YYWWRYCAYYYWYWRKMMRCWRMKYMYTTYMYRAKATTGRRRMCRGMYMRWKKWRYRWKC
44                YYTWATGGCTCWYWRKMMRMWRCKYMYCGYMYRWKTCCRRRAMTRKMYMRWKKWRYRTKT
                  ** *    .   ******** ** ****  **** *:    ** * * *********** *

42                YGCTTYYKRWWWYKWRMRRRRWWMYYTRCKRMKRRKYKRCMRWWWWRRMYMMCGYTYWYC
44                CTAKYYYYKRWWWYGTRMRRRGAWMYYWRMKGAGRRKYKRMCAWWWWGGMYMMTTCKYWTY
                    .. ********  *****  **** * *   ******   ****  ****   .**

42                MRGWYGYYWAACCKRCYGGTWKRWTMGGWGTRRRMSYMRRYYMYMGCGATTRKRCKKRRR
44                MRRWYSYYATMYTKRMYKRCAKRWGARRWKCGRRMSYMARYYMYMATARWYGGRAGGRRG
                  ** **.** :   ** *    ***    *   ****** ******. .     *.  **

42                RKYRRYYRKYMYWYYGGAAACRYKAMSRWYCCCCMCCWYYGKMCRARMYYAKMYRYRYRW
44                AKYGRYYGKYMYWYCTTTGRMRCKRMSAWTTYATMATTYTATMSGTGCTCGGMYRCRYRW
                   ** *** ******   :.  * * ** *   . *.  * ..*. :    . *** ****

42                MMWWWKMKRYCWKWKKKWMTTGKYMKYGSRMKYCGCWTAGRRAYYKYYRRKRRKTWGYKM
44                MMWWTKAKRYYWKWKKKWCGCKKTCKYASRMKYTATWYWKGRCTYGCTRAKRRKKTKYKC
                  **** * *** *******    *  **.***** . *    *. *   * ****.  **

42                GTKKWKYRRRWKYRYKRKAYGGCCGGGGYMSKYYRRAYWTGGYYTAKSRKCYMRKYKYCT
44                ACKTWKCAAGWKYRYKRKMYKRYTTARKYCSKYTARWTTCAAYYKRKSRTMYAATYKYGY
                  . *.**    ******** *      .  * ***  *     ..**. ***. *  .***

42                YCCYRMYGYSYKMRCGCMCGCCKTARMRRRMYYCRCCTYGGCMKGYYWYRKTYRYMRKRG
44                CMYYGCCKYSTKCRYRSAYRTAKAGRMGRRCCCTRTTCTRRMMKRYYAYAKCYRTCGKRK
                     *    ** * *   .    .*:.** **    *       ** ** * * **   **

42                TAKRRTKYKYYRKWYYWTYRKRCTCMGKKRKYYGKYRGRGGTYRMKSRCTYRAYYRWYTG
44                CGKRRWKYKYYRKWYYWYYRKATGTCKKKRKYYKKYRCGTTAYRMKSRAATAWCTGWYAK
                   .*** *********** ***       ****** ***    :******.:      **:

42                MAYYYGYYKCRMGRTKTRRRKKYKMKMYYRKYKMMYWKAATKRYWGGYYWKRRKMGRTKY
44                MGTCCKYCKYRAKRAKCGRRKKYKCKMYYRGCKMCYWKWTGKGYTRRYYWKRRGCCAWKC
```

```
                 *.    * * *  *:*  ****** *****  ** *** : * *   ******     *

42               YMARRTYRMRYGRRCAAMAGTGTTARTRYRYRWYYGCTYKSRKTYTCKRMMWYYWMAAWY
44               YCRGGCYRMRCAGGMWRMWSYKYWRRWGYRYRWYYRMCCKSRKCYKYTRMMWYYWMCGAY
                 *     **** .     * .     *  *******   **** *. .********..  *

42               KMMYRWYWMRYYRAACWWMGSKGKRWMRKYWWYYKTAYKKAYTKKYRYYYKRYKKYYMWG
44               KMMCAAYWMRYYRTTAWWMAGTKGRWCRKYWWYYKWCYKKWTWKGYRYYYKRCTTCCCWA
                 ***   *******::.***...  ** ******** .***   * ******* ..   *.

42               ATRGAMKRTGWKMYWYYCMKYTGAAWYCRYTKWSWYRYKKWWKRKYMTCTRRKWRKCRCC
44               RGRAWMKRAATKAYWYYYMKTGACTTYSRCYKWSWYRYKKTTKRKYMYYYRRKTAKTGYA
                   *. ***:. * **** **  ..: *.*  *********  *****   ***  *   .

42               ACCTGCCCGTWRWKWRKYWYRAKKRSYKKWKRKRRMKKCYYYKTCWKKKRRKYWWKYART
44               RYYYKYMMKWWRWKWRKYTCRGKKRGTGKWKRKRGCGKYYYTKYYWKKKRRKYAAGTWGW
                           ********  *.***.  ******   * ** *  ********

42               CCCKYWYRYMRRCMYRRRRKWRSCRRWRKTGRTTGCCRWYRRYMGKRYTCGRYRYKYCRK
44               SMYKYWCGCAGAYCCRRRRTWRCSAGTRKWRGKCCAMRWYRRYMRKRCKATRYRYKYYGG
                 .  ***         ****.**..   **   .  . ******* ** .. ******

42               CCTGTYCYKYGRWGKMKMKAATAKRTTRKKYMYRMRRKYKRSYKWRYYRKKGKYYAGWTY
44               AMYAAYTYKYRRTKTMKCKGCCMTAYWRTTYMYGCGAKYKRSYKTACYRTKKKYCGRACY
                 .  .:* *** *  .** *..  .   *..***    *******   **.* ** .   *

42               YRARYMKAKMKMYRWKYRRWRCGMKMKRYWWMRCAKKCATTRAYGAKRGRMRKMTYTCCM
44               YRRRYMKWKMKMYRWKTRRWAYKCKMKRYWTARYRTGTCCARWCRWTAARMAKMYTCYTC
                 ** **** ******** ***    ******  *  .  . :*    . .** **

42               YYRCKKKWYWYRKWKYKKRRAKYKMRYYRWYYSCYKMWRCYYKRYYYMMRWYRRWWRYKK
44               YYRYKKGACACRKWKYKKGGRKYGCRYYRWYTGSCTCWRYTYKRYYYMMRWYRRWWRCKK
                 *** **     *******   **  ****** .. . **  *************** **

42               KKCSYRYRRRTYGATRMRKKYYGYYMTKCAAYMRKYKYKCMKMYYYRMKTTACYTTRWKW
44               GKYSYRYRRRACKRYGMGKKYYRYCMCKSRRCAGKCKCKTMKCYYTAMKAYMSYKARWKW
                  * *******:     * **** * * *.     * * * ** **  **:  .*.:****

42               MCAGKWRKYMACWTTARATTYTYAKAKRKCSYCATTRAGGKGYWACWKKARYKMYKKYGW
44               MTRRKAGTCCGTWACRAGYKYKCMKGKGKTGYYRYYGCKATACTGTAKKCRYKCCTKYKW
                 *   *  .  . *:   . .*.  *.* * .*     . ...  .  **.*** .**  *

42               WYKKWCRSKYRYMRYYYRSKTYMRKCKKYCYRKWRKKWKRYRYWYKRMYMKTKKKRYYWK
44               WYKKAARSGTGCMGYYYRSKKCMRKTKKCYYRKTRKKTGRYRYACKRMCCKCKGKRYYWK
                 **** .**    * ******. *** **  *** ***  ****  ***  * * ******

42               SMYYRCKRKGTCMKGCCCKGKRCMRRKRRYSTTARTKTKKKTGACRGGGGTRTGMCSWMR
44               GMYYAYKAKKWYMKTATAKRGAYAGAGGGYSKYRACGAGKGGRGTGRRRKYAWRMMSTMR
                 .***  * *   ** . .*          **.     : *   .          * * **

42               YACYRWYCGAGTTKWYYYRKWGATTTYRMYRRAKKKRAAKYSTWKKMMGATTTRRRYKYA
44               YRYYRWYMKWAKYKWYTCATAKRYCYYRCTGGMKKKRGGKYSKWTKMMKCCCGRRRYKCW
                 *  ****   .. ***   .      **     ****..***.*.*** .   *****
```

```
42    GRMYYRTKGTRYKKMATTTWCTKAAYKYRRRYWWAATYRWKRWGMGYYKTGTRGKGARYC
44    RAATTRWTKYATGGCTGWCTTCGGTCKCAARYWACCYYRTGAWKCRYTKWACAAKRRRYT
           * .       :       .: *   *** .. **   *  * *  .  .*  **

42    RTYYRYCAKWSKTAYWSRRYRRRKMWCCRYRKYAYMKRCTRRRRWSGYKRTCAATTKKYR
44    GYYTRYYGKWSKGTYWCGGCGRGGMTATRYGKTWYMKGTCGRRRWSCYKRWTTMCCTGYG
        * ** .**** :**.    *  * . ** *  ***    ***** ***   :  . *

42    YMCTWYTGRRYWWGATTRTRCTARWATCTTKRYYGGKGGCYMGTCKRATRKYRTRACGWY
44    YMTCTTARRRYWWAGCAAGRTACRWRWYYKKAYYRAKRRYYMKYMGGRKGGTRKAWMRWY
      **    : *****.. :  * :.**    .* ** .*   **      .   *.    **

42    YRKCKYYYRKMWYWGYWYRSYMYYRYKWARMMKTRWWRWRMKGGCKKMWWWYKYYMRYYK
44    YRKSGYYYRKMWYWAYWYRCCCTTGYKTMRACKGRWWRWRMKKKYKKMWWWYKYYMRYYK
      ***. *********.****.     **  *  * ********   **************

42    RRWRTCGKKKKMWYRMRGRYYRKRCRKYWGKSYWCYWRRYYMMYKCRGKRKKMYMAKTSK
44    RRWRGYKKKKKAWYRARKRYCRKRMRKYAKTCYTTYTRRYYMMYKYRRKRKKMYMCKWSK
      ****   **** *** * ** *** ***  ..*  * ******** * *******.* **

42    KWRWMWTWRRRKTTTTWYSYRYKWYRKSRKRKYRGCYKRRYRKKKRRRYKMGYWTCCTRT
44    KWAACAWWRRRKYYYGTYSTRCGTYRKSRKRKYRKYYKRRYRGKKAAGYGMKYWYGTARC
      **     *****     ** *   **********  ****** **   * * **   :*

42    YYSKRCKYAWKKYRMRSGYYKMWYYKKYYGRTAARYKYMRYKKSRATKKYKYKRYRYYKK
44    YYSKAYKTMTGKYRMRSRYYKCAYYKKTCTRATWRYKYMACKGGGMKKKYKYKRYRYCKK
      ****  *    ****** ***  ****   *:: *****  * .  .********** **

42    RRRYWYWKTYMTRKGAGGRCYYKRRYMYTWRCKRGYKMKYWKSKKRYWCKCTRKGYYKYK
44    AARYWYTGCYMYRKKMKRAYYYKARTAYKTAYKRSYKMKYAKSKKRCWGKGGGKKYYKYG
        ****   ** **      *** *  *.   **.***** ***** * *   * ****

42    RYRMKKKWRRRKKYKATGWKWWYMGKKYSRKMTTYYYWYKWMKKRYRWMKWSSMRRKWMS
44    RYGMTKTTAARKTYKRWKWKAWTAKKKYGRKMCAYYYWCKWMKKRYRWCKWCCMRRKWMS
      ** *.*.   **.**   ** *   ***.*** :**** ********* **..*******

42    STTRRWYMRYKACKAARKYMKRYKRGYTRYYKRYYRRWYCTYCMYYKTKKCGYKYWMKTR
44    SCWARTYMATKTTKRRRKCCKRYKGKCARYYKGCYGGWYMWYMMYYKWTKGTTKYTCGGR
      *   * **  *: *  **  ****   :****  *  **  * **** .*   **    *

42    KYYKTKRKYRRCGCGTCRYYWWKYWYMKKYRYYTCKKYMRKKKRYCAYYATKWAYMKRRM
44    KYYKYGRKYRRYATRCTRTYWAKYWTMKGYRYYYYKTYMGTKKRYMCCCRCKACCCGRRM
      ****  ***** .    * ** *** ** ****  *.** .**** .    * .   ***

42    CKRWRKKRWSYKTTCYWWWSYKRMYRWRYTTCYGMTSAGYRYKWKWYRRYMSYKSMRKAG
44    MKRWRKKRWCTGCCTYWWWSCKRMYRWGCWWSYSAYSCACRYKWKWCAGYMSYKCCRKCK
       ********.     ***** ******    .*.  *.. ******   *****. **.

42    RRYRGWAWMYYYGAGTGCTRMKRKWCRCTTSAAMCCTTMKAYYWKTKRYMYMWMWKWYGK
44    RRYRKWRWMYYYCWAAAYCRMTRKTYGYYGGRWCYTCCMKWYYWKKKAYMCCTCATWYRK
      **** * *****  .:.  **.**      .       ** ****.* **     .** *
```

```
42              KKKTTAWYKKCRYWRYYWYSKSRMYRYRMYYAMYKCRAGCRYYTCCWYRTCKRAWYYKGY
44              KKKYYWWYKKTAYWRCTWYSKGAMYRYRMYYMMCTYRTTARYYCTTWYRCYKGWAYYKKY
                ***   ****  ***  ****. ******** * . *: .***   ***  *   *** *

42              KRTGTCCGATYWCCCGYSKAASKRKSKKSMRKKKSYYRKRRGGYKSMRMRYKTKYRSMMY
44              KGCACYAAGGYWATTACGKGGSKRKGGKSMRKKKSYYGKRRRRYKSMRARYKWKYRCAAY
                *  .  ... **.  . .*..****. ********** ***  ***** *** ***.  *

42              KYWKKWMKMKYYRKWACCGTRKWKMRYKGKKRRKYKWYGACKMMMYYKKRYWCRRRGMWW
44              KYAKKWCKMKYYRKWWMYSYGTWTAACKKKKGAKYKWYATTKAMATYKKACTYGGRKCAA
                ** *** ********    . .*.   * **  *****.: * *  ***      *

42              KSAGGRYYYMRMYYYGGGGRARSWKYYKMKWSGGYRRYKKRGCRKYRYKYMRCAYCRMWK
44              GCRRRRYYYMGACTCAAAARRRSWKYYKMKACTKYRRYKKRKYRKYRYKCCAACCTGMWK
                 .   *****     ....* *********  . *******  ******   ..   ***

42              YKKRWKRMSYWYYMKGCKKYKKYWWKSAKKYYKMCSWRWYYRKRWYKYRMYKKRSCCYRW
44              CGKAWKRMSYWYYMTRYTKYKKYTTKSCKGYYKAYCAAWYYRKRWYKYRMTKKASMMCGA
                  * **********.  .*****  **.* ***  .  ************ ** *

42              TGAARYRSYWKRGTYKYYYMKMMGYWRRKMKGYGKKWRAGGGRRGCCYYYAMGTYKRKYT
44              KKRWGYRSYWGGCWYKCTTMKAMATARGKMKKYAKKTGMKRKRRKMMYCTWMRWYKRKYY
                .    *****    **   ** *.  * *** *.**      **   *   *  *****

42              ATYMKKRKKKWYKRMRRYTCAMWMTMMWYRRRRGATRGWYRKWRYKKRRKWKKGRYYKRY
44              RWYMKKGKKKWYKRMRRYWMRATCKMMWTRGGGKGWGCWYRKWRYKKRRKWGGRRCCGRY
                  **** ***********      .*** *    .   *************   *   **

42              KMKGYYKAYYKKKMRKRTYCRKRKYYMKYTYRWRRKYYAYGMYKYRMYWKRRKKKKKRKC
44              KMKCYCKWYYKKKMGGGCTTRKGKYYMKYWCRARRKCTGYKAYGYRCCAGRRKKKKKRKM
                *** * * ******      ** ******  * ***  .*  * **    *********

42              TYGTGARRKWYKGYRYYYYRKKWCCYMMKTKTYYRRGRKYYWAGGGYAYMTWWYYAKRYK
44              YYAAATRRKWYKKYGCCTTGKKWMYYMCKKKCYYRGRRTTYWGATACTYMWWACCMKRYK
                 *.:.:****** *      ***  ** *.* ***  *. **.. . :** *    ****

42              YYKWYYRMKKYKKRSWCKARYTGTKRRWWYMYRWRRKKYRSKYYMSKYGKMYKRARWKMS
44              YYKWYYRMKKYTTAGTMKCRYYRYKRGWATMTGTAGKKTACKYCMSKCTKMYKRWGWKMS
                ***********.. .  *.**   ** *  *     **  .** ***  *****  ****

42              TSRAKWKYCYKWAKRRKKCATRMMYMYKYYMYTGCGMGAATKACRMKYRMRYYRRAGYGC
44              ACRMKWKTMTKTWKRRKKTTCRMMYMYKCYCCAATAMCGGCKWSRMKYRMRYYRRGKTTT
                :.* ***   *  ***** : ******* *  :. .* .. * .***********.

42              YYACGCRGAAGMCRMMYYYKMYRWACGTKYRKWGRYYAAGMYRYGRSKYYYYARYAKRMG
44              YTWMRTRRMRKMYRMMYYYKMYRATTTCKYAKWRGCYWRRMYRYAGSGYYYCCGCGKGCK
                *     *    * ********** :   ** **   *   ****. * ***  . .*

42              YTGWRRRYWMCGKGKMGKGASATKKKKMKKMKGRWYTMGRMRRKYTKRMYKRGYKYKWMY
44              YKAWRRGYWCTRGKKMSKCRCTGKKKKCTKMKKRWYYAKRMRGTYAKAAYKRRYKYKWCY
                *..*** **     **.*  .: **** .*** ***   *** .*:*  *** ***** *

42              GCMYAKMRRRMMWMKMWMTGRSYMRRYWKTMTKWKWKKKKGGKGRRRRYYKRKRRRSMCM
```

```
44          RYAYGKCRRRMMWMKATCAARSYARGTAKYCATWKAGKKKARKRARRRYCKRGGARSMYA
               *.* ********   :.*** *   *  :.**  ***. *  **** **   ***

42          TTYKKMYYRAYKRTTRRMRKRYYGRCTYYKAMKKMKMYCWGMMARYYRRRKRGYKKRYRY
44          CYYKKAYCGCYKRWWRRAGGACCRGYWYYKCMKKMKAYYWKMMWAYYGRRKRKYTKRYRY
              *** *  .***  **          ***.***** * * **  ** **** *.*****

42          YYYRRKTMRKKKKKRWAMYKGWARYRMYYRKWKKKKYTRYYTWKRKKRRKCARRWMRYGW
44          YYYRRKYARKGKKKRWMMYKATGGTAMYYRKTTKKKCCRCCYWKRKKRGKMRRGAMRYKW
            ******  ** ***** ***. .   ***** .***  *   ****** *  *  *** *

42          RRKKYYYCRGKRYKKRRRRRYRRRYRGGCYCYTCCACAKTTRRWMRRTKTSKYKRRWARM
44          RRKKYYTMRKKRCKKRRRRRTAGRCGTRYCYYGYMCYWTKYARAMGGKTWSKTTRAAGRM
            ******  * ** *******   *       *   .  ..  * *  .. ** .*  .**

42          RKKTACGKKTTTYAWGCYAYRRRRWAWRRWYRACGKRRRMWYCCTYRMYRKRYMRRWCCY
44          RKGAGYRKKGACTGACTCWCAGRRWRWRRWYRGAAKAAGMWYTTCYRCYGTGYMRRTYYY
            ** :.  ** :  .        *** ******...*   ***   ** * . ****   *

42          YKGKCGCRYMRW
44          YKRKTRYRYMAT
            ** *   ***
```

The pairwise alignments of two accessions 42 and 45

```
42                  MWRYRYWSYRSYSMKWRYAMSWWKKCTYRRGKTTTKYRKRYRKKYRMRRWRRWMMKYYKS
45                  MTAYRYWSYRSYSMGWRYTMGTWKKACYARKGGACKTATRYRKKYRMAAWRRWCMTYYKS
                    *  *********** ***:*. ***. * *   : *  .*********  **** *.****

42                  YYYMRYMKTAYRWKYRCGYYYCRCAGAGGSRYSRWMWMYKYYMMRYRSAATARMKYYYWW
45                  YYYMRYAKCRYRWKCATRYTCYAMCKRRRSRYSRTMWCYKYYCAAYRSTRCCGMKCCTTW
                    ****** *  ****     *     .    ***** ** ****   ***:  . **    *

42                  RYYRYMYYWRRKYWRWWYKYMYGGYGTTKRWMMMYWKRYWTYCWYRKYKMYWKYWKYWKR
45                  RYYRYMYYWRRKYWRWAYKYCYTSYACCKRWAMAYWKRYWWYTTCAKYKMYWKYWKYWKR
                    **************** *** * .*.  *** * ****** *    **************

42                  RMRMGRYMYRKYRKMRYWYYRMKYRMYCRYYMWRKYMWMKTRKKYMYYWYMMMMYRYYYY
45                  RMRMRATATRKYRKMRYTYYRMKYACTTRYYMWRKYMWMKWRKKYMYYWYMMMMYRYYYY
                    ****     ******** ******    ************ *******************

42                  YYRWRYKWYRRYKMYYYRRYRSYKKRMYYMYKMCYRYWRYWWWMKWSRRKRWMRRWKRWC
45                  CYAARYKWYRRYTAYCYRRTGGTKKRMYYMYKMYYRYWRYTATMKWSRRKGACAATGRWA
                     *   ********. * ***   . ********** ******   *******      **.

42                  RRMKRMYSRRRYSAGAYAYYMKYGAWWMCTKTGWRYRYYRWWMRMKRYKMMKRGKSKRWR
45                  RGAGRMYSRRGYSTAGYTYYMKCRTAACYKKCRWRYRYTRWWMRAKRYTAATRTKSKRWR
                    *   ****** **:..*:****   :    .*   ***** ***** ***.   .* ******

42                  MRTTARRTKRRYKTKTWMRGWRMRSMMKTKCCCCKKATWRCCYRMGGMRMRKRKKKRMYR
45                  MGKKMARCKRRYKYGCTCASWRMRSMAGYKYYMTKKRYTRMYCGMKKMRAGGAKKKRMYR
                    * ..  * *****       .******   *    **    *    *  **    *******

42                  TWCMRKYRMWKGCTKRCMRYYKRTYTTCCWYRMYWWGKACKRYKRYYYYYRRKYWMKMKC
45                  AATCRKCRMWKRYYKAMMGYYKGCYACTAATGATAWKTRMKRYKRYYYYYRRKCWMKMKY
                    :   ** ****   *  * ***  *:   .       * .  ************* *****

42                  AWMRKYKRCMAGYYRMKMKKKRKMYYTWKYKKMRYRRMYWYRYYSCWKMRKGCGTYYYKT
45                  WTMGKTKAYMCKYYRMKAKKKAKMCCYWTYTKCRYRRMYWYRYTGAWTCRKRYTCYYYKK
                      * * *  *. ***** *** **    *.*.* **********  ..*. **     ****.

42                  MAGCCAMRWYRRWMWRRWKKSYKRARRKCATSGATRKGRTKYMCCCCAATARATRYMRKY
45                  MMRYMRMRWYRRWAAARTKKSYKARRAKYTGSKRYGGAGWKCMYTMSRRKRRGCAYMRKC
                    *     *******   * *****  * * : *      .  * *    .  . *.  ****

42                  KKYKMKCYGYYWRRYMMKKGGKWYTKWMKYMRYMMMGGCMYWWTTARARCMKKRRRATGA
45                  KKCKMKYTTCCTAATMMGKRCKWYATTAGCMAYMMMRRMMCWWACGAGRTMKKRRRRYAW
                    ** ***          ** *  ***:.     * ****   * **: . .* ******   .

42                  KKYYRKCYSRYMRWRWRYYWKWRTTRKRTAAWMRYWRAGTGTKTMWYTGKWRRWYYSYMT
45                  KKYYGKTYGGYARWRWRYTTKTRWYRKAYTTACRCAGWSYKYKAAAYYKGARRWYCSTAY
                    **** * *. * ******   * *   **   ::  *     .    *:  *     **** *

42                  TGYACWRYRKMARCGGTTCTACCYYYYMMWYYRWMKWTGRCMTYGCGCMAAYKCWRKKYM
45                  CRYCTAGYRKMCAMRKCAMCWTSCYYYMMWYTRWAKWAARTMCTKYRYMRRTTTWRKKTC
```

```
                  *.   ****.     :    . ******* ** **:.* *      *   . ****

42                AYWKWCMWKRRMRTRYRWMCRWRYWYYWAAGYKKKYKYAKYYKGKYYMWYWKMYKKWKYW
45                TTTTTAAWKGGCGCRYRWMYRWRTTYYWWGAYKKGTKYGKYCTKTCCCWYWKCCTKWKYW
                  :  . . **     ***** ***  ***  ..***  **.** . .    ****   .*****

42                KYTKYWCKCRWYYRSRTCTYRKWRKKMRKYRWWTWYMYKKCCWAYATKRGMSMARTKRYW
45                KCYKTWAGYRTYYRSGAGYCRTWRKKCRKYRTACTYATKKTGAWCWYKRKCCCTACKRYW
                  *  * *.  * **** :    *.**** ****     *  **        **   . :  ****

42                TKYKYCGGCAGCYCGGATGTRKYRKKRYAYTRWWWWWMRGWWKKKGAGTWCRRYYKCYRR
45                KKTGCASRYCRYYMKKRYSCRGCRKKRYTYKRWWWWWMRRAWKKTRWSYTMRAYTKYYRR
                  .*    ..  .  *      . *  *****:*.********  ***.   .    * * * ***

42                RWKAYARCAYTRYRKYRSMYYMMARMYYWRKRTWYYCYKKRRMMYMYMKKSTYKYTTKMR
45                GWKGTGGMCCCRTGKTACCYCMCWRMYCTAKRWWCTMTKKRRMMYMYMKKCCTKYKWKMR
                   **. .  .  *  *  . * *  ***   ** *    ************.  **. ***

42                YYRKAGCKRYYTWRAGAACCRMMYWYYYWYYKYGKYWAKRRAKKCTACYRMSYMYYYRGC
45                YYRKGCTKRYYGAARRMRAYRMMYWYYYAYYKCCGTWWKGGGKGYYRYYRMCCCTYTGRM
                  ****.  ****        . ******** ***    * *  .*      ***.    *

42                CMKWKYMKRRRWRMKSMRACKMKKMMKRWYWRMAKRKKGAYYGRRGRGGYRYMRWKKCKW
45                AMKWKYMKRRRWRMKSMRRYKCKKMMKRWYWRARKRKKKRCYKRRTRRTYRYMRWKKSKW
                  .****************  * **********  ****    * ** *   ********.**

42                RKYWKSCWKRKTWGKYMMKRKKMYCYTAKSKKWKMKRRRKTTTTRKTTCGRYCRKMRWKK
45                RKYATCTTGRKKTKKYMMKRKKMYYCYRKSKKWKMKRRRKYWYYRKAGTTGTMRKMRTKK
                  *** ..   **. **********    ***********     **:       **** **

42                RKKKCYKMRRYRKYRRRYKRYRYWGRGWWKRGGAYMMRKKMCMMKRMRKYWRMAGYKYGR
45                RKKKYYKARRYRGYAAGYGRYRYWRAKTAKRRRMYMMRKKMYCCTGCRTYWRAWKTGYAR
                  **** ** **** *    * *****     **   *******    .  *.***      *.*

42                GWYRYYGASKWMRCTWGRKRWWWKCCWKMTCCYWYMRKKGAGYKRYYRGYCTAKRTAGGA
45                RWTRTYSRSKWMGACARRKRWWWKMYWKAWAYYWYMAKKKGRYKRYYRKTGKWKACRART
                   * * *. **** .    *******  **   . **** ** . ******    . *    . :

42                GAGRYYRATTKYYTATRRYRMMSRYYRCGGGWKYMWKTYRWCGTYGAMRMYYMTYRGCWY
45                ATTRYTGRKYKYYWMYGGCRMMSAYYRATTAWKYMWKKTGAYRKTACMRCTYMYYRRTAT
                  .: **    . ***       **** ***.   .******.       . ..**   ** **

42                WSKTWRYTTWGCGYYMYKRTYMKRCTRRRRWSGRKMYKTKYMKTTMRYKMRWRWKWYGAT
45                WSKCTGTKAATARYTCCKGKTAGGTCGRRRWSRGKCCKKKCATYKMRYGCAARWTWYAGC
                  ***     .:  . *    * .       *****   *   *.*   . .***     **.**..

42                RGCRGKKWARCSRYKYWWRWWRRTAYCAYGCYRMTYKRKCTMYYYSRGTACTGRAWKRTA
45                RTTGAGKWWAMCRYKCWWRWWARKGCYRYTTYRACYKRKYYCYYYSRRATMYSRMWKRCT
                  *    . **    .*** ***** *..    *  **  ****    ***** ::   .* *** :

42                AGWKKCGRTKKTRRTKKAKGTTACAKKKSYYKKKRRTYKKKRKRKYGRKYTYKSSYRKRY
45                CKWGGYRRYKKGRRKKKRKRKWRTTKKGCYYKKGGRYCKKKRKRKYKAKTCYTCGYRKRY
                  . *     * ** **.** *  .    :** .****  *   ********   *   *...*****
```

```
42    CWGTYARYYYYWWMKCTGKGAGMCMGKRRKKKYCYMRYTWTCAGYYYRRKWYTKMYARKT
45    GTAWTMRYYYTTACKYYKKATTATMKGRRKKKYYYCRYAACTGRCYCRRKWYKKMYWRKC
        .   ****    *   *.:   *  ****** * **:   .  * *****.*** **

42    RTGTYYRKWYTKYRMYRRKMKKYYAGGKKKKKWGCKKYKWKSYYKKAGTTRACTWGWKKY
45    GKKCTTGKWCKKYRMCGGKMTKTYRRKKTKKKWRYKGTKWGCTYKKCACCARYKAKWKKY
       .     ** .****   **.* *   *.****  *  ** . ***..     .  ****

42    YRYSYSSRMYKRMYYWRMYKRKSWYYKRYRTATKYRRRRWYMYKKTRKRKYKTGYMATYW
45    YRCCCSGAATKRMYYWRCCTRKCWYYKRYRYGCKYRGAGAYATKKYRKGKYKYAYMGGYW
      ** . *.   *******  .**.******* . ***    *  ** ** *** .**. **

42    YCCACCCGYSYRRSRSKKKMRKKGWRSKYKRMCYSAYKRKRRMYGKSMRYRARYKYRKSG
45    YTSRATTAYGTGASRGKKKMRKKAWRSKYKGCYCGWYKGKRRATKKSMRTGGRYKYRKCT
      * . .  .*.   **.*******.******    . ** ***   ****  .******.

42    KYYRTMKWYKAGSGAYRRRYKYCRTKYRKCCCCTAKYGKMKKKKKRKRRMRRRYRSYWWK
45    KYCRCMKWYKWRCKMYRGRYKYMGCTYRKYYMYWRKYRKMKKKKKRKRRCGARYRSYTTK
      ** * *****  .  ** ****   .***      ** ***********   *****  *

42    RYKKGYRKYYYKATCYGRYMRKKTTAARRMKMRYKKRRWMMYYKKRRYRRYTKATTRGKR
45    ACKTKYRKYYCKWGAYKACCRKKKYWRRRMKMRYKKRRWCCCYKKRRYRRYGTCCAGAKA
        *. ***** *  .*    ***.   ************   ********* .. : .*

42    YKAKKYRMTKAMTTYACKKKKKWYTCGYWKKWYYYKRRKKRKRRRYYTKMAWKAWKGRRC
45    YKTGKCRMKKRMYCTTGKKKGTACWYRCTKKWYYYKARKKAKGAATTYKMTAKRWKSRRY
      **: * **.* *   : *** .       ******* *** *      **: * **.**

42    MKRMKGGTKKGKGTGGKRAYTGCGMRYYRYRKTYMYMTCAGTTKWYRYTRCKTTCTYYYW
45    CGGAKCRAKGKTAKRRTAWYCTYKMRYYRYRKYYMYAYYWAKYGWYGYYGYKCCTYYYYW
          *  :*  ...  .  *    ******** ***    ..  ** *   *    ****

42    RMRYMMGCTRTGYSCYYRTSRCRAAMRRRYRKRRGKWWGGMYGKWYYRRCWYYMYGKATG
45    GMGYMMKYWRYKYSSTYRWSRYRGRCRRRYRGARKTTTAAMYTKWCCAATATYMTKKCYC
       * ***   *  **. ** ** *.  *****  * .  ..** **       **  *.

42    TKKYRKYRMMYYWYYTYKYRSRMWYRYTCAAMGRCYSKKYKRMSKWWRWWMSWKRYWKKA
45    AKKTAKTGCMYYWYTWYKYGCGMWYRYCACGMAGTYSGKYGRMGKTTGAWMSWKGYWKKR
      :**  *   *****  *** . ***** ...*.  ** ** **.*    ***** ****

42    RAATKACKCKKMKGKRMACKCTYTKCYMCGCKKRMYKRKGGRKKWGTCACGMYKAKRMMK
45    GMTGKCTKYKKCKKGRMTYKSWTCKYYAYCSKKAAYKRGARAGGWRYYGTCMYKGKRMMT
        : *. * ** *  **: *.   * *   .**  *** .    *   .  ***.****.

42    KMKKMWMYMRSMYRTWRKSYGRWKKYRMRWATKKKMKRRKYYGYCARKACYMRYRKMKRK
45    KMKKATMYCRSMCRGWGKGTRRAKKYRMGWRYKKKMKRRKYYRCMRGKGTYMRYRGAKRK
      ****  ** *** * * *.  * ***** *  **********     *. *****  ***

42    KYKKKYYYYYKTWGRYRYGYCKRCRRTKRRRGCYKRKKCTRTWARRSKRKKYRGKRRRKK
45    KYKKKYYYYYGKASRCRTRCYGAYRRYKAGRRYCKRKKMKAKAWRRSKGTTTGAGAAGKK
      ********** . .* *       ** *  *   **** . .  **** ..  .    **
```

```
42    MKYRRMWRRKWRYRTKRRRSKYYYAYKMYRAWYYTTACCGKYRKKRYKMRKKRWCYKRRR
45    MKTGRMWRRGWRYRYKAAGSKYYYMYKMCRWWYYKKGTTRKYRKTGYKMRKKATMYKRRR
      **  ***** **** *   ***** *** * ***...   ****. ******   *****

42    MKKYKRRTCKRRYYRRRMRKWMWYRKAKKRRRTRKWYMTWMYYKRYRRGWCKRMYYRRYY
45    CKKCKGGWTKRRYYRRRMRKWMWYGTWKKGRAYRKWYAWWATYKRCGRKTGKGMYTRGCY
       ** *    *************** . ** *  ****  *  ***  *   * ** *  *

42    RSKKYTCTWKKRRKRYKCKTSKYMKCRRWRKYYRYTYYRRRATAYCARRMRYKKRKRRRS
45    RSKKYYTCAKKRRKRYKMKWSGTMGTRRAGKTYRYAYCARARWWCYRGAMRYGKAKRGRS
      *****    ******** * *  *  **  * ***:*  *         *** * ** **

42    YWYRKKKWYTGAAMWYMRRRCGRWWRYKCCCCRYRWKAMCCTTRYWCKKWYKKWYRRRAG
45    YWYRKGKWYGATTMWYMRRRYKGWWRYKMYYYRTGAKTCYTGWGCWGKKWYKKWCGARGS
      ***** *** .::*******   *****    *   *:       * *******   *..

42    YYRKYWYYKKRKSYKRKWYKRKRKKKKKKKCYMYKGWKKYYWRWWTKYACYWYRYMRRMM
45    CCGGYTTCKKRKGTKRKWCKRKRKKKGGGKYYMYKKWKKYYWGWTCTTRYYWCGCAGACM
          *   ****. **** *******   * **** ****** *  .   **       *

42    KMYYRRRMRTKAWWRKRRWRRKTGCCYRYMRKCKARKMRGKRRRYCYYYTKWRRMCKYWR
45    KMCYRRRMRATCWWRKAGTRRKCCAMYRYMGTYKRRKMRRKRRRYMYCCCKTRRMMKTAA
      ** ******:..****   ***  . **** . * **** ***** *   * *** *

42    KRRWRWMKTTTCGCKWWSGYCWGWMRKRYYCYRYYRSKMRYKKYRRKKYMSYGSMKGCRT
45    KRRWRAMKCCGTAYKWWSKTMWKWMRKRYYYTGCCRSKARYKKCAGKKYMSYRCCGKYGW
      ***** **    . ****   * *******     *** ****   ****** .

42    KYKRYRWYKWWGWMRMMYRKKKRYWMSYMGMYATKWKRYRRWMWKWYRYKKKKRYATKCT
45    KYKRYRATKWWCAMRMCTATKKRYWMSYMKMYGCKWKAYRRTCATWYAYKKKKRCGCGAC
      ******  ***  ***   .********* **. *** ***   .** ****** .  .

42    AYYRCYTRYRWMMWWYSSWKKMCMAAAKKGTCKKWKRKKTMTYMKYYRTTGMKYGCCKAK
45    WTYRYCKATAWMMWWYSSTKKCYCWTGKKRKYKKWKRKKAMKTCKYYRYYKMKYKYYKWK
        **  .   ******** **    :.** . *******:*.  ****   ***   * *

42    YACSRCTGAGCKYKYRKAAKTWYTWRKMGKKWTKYRRKRWKRYKRKRRRWRRWGYMTRRY
45    CWMGRYAATRTKCKYRKGGKKWYWWGKMAKTWYKCAAKGWKRYKRKRRRTRRWKTCYART
         .* :.:  * ****..*.** * **.*.* *   * ********** ***      *

42    YGRGCKWTARKKYSRKMRYYKYCYYCMYGGRRMYSYGCYMRTGGGKKYGCAGRRRKKGYG
45    CRGKYKAKRATTCSRTAAYYKYGYYSCCKKRRMYSTRMYCGGTACKKTRYCTGRRKKKYR
           * .  .. **.  **** **.    *****   *    . **   .  **** *

42    AACCMKGTAGCTKWRAATGKKRMKKKYYRKWYYMCGARRYWMRKYYYRKYYTKAAARKGC
45    RMTAMKRCCTYYKAARCAAKKRCKKKYYRKWYYAYKMRRTTMAKCYYRKYYGKMWWRKRT
         .**  .   *   .:.*** **********    **  * * ****** *   **

42    RYTRTTTTKTYRMKSTRKYYYCCARCAGATMCTAAYKGRKRYMRGWTATMTYRTRCAGGT
45    GCGRACAKKKYRMKSGGKTCTTARGYRRGWCYWMRYKKRKRYMRKWATCAGYGARTGAAC
         *: :.*.*****  *    .     .      ** ****** *::   * :* ...

42    YYWRRMKSKYKKAYYKYATGATKWTCAKYSRSWMMRSSRYGYRAMCYYKRWTKRWSARGA
```

```
45                YYWRRCKSKYKKMYYKCCGTCCKWCTGKYSRSWMMRCGAYSTAMMYYTKAAKKRTSMGRT
                  ***** ****** *** .  . **  .*********.. *.   * * * .**  *   :

42                YKRGKMKYRRMMRYTGAATYGYRYYRYMRRGKGMARRWYYYYKKKWMKTCRKYKRMRKCC
45                YKRCKMKYRRCMRYKRTTGTKCRYYRCAAGTGTCGGGAYYTYKTKTCGGSRKTGRCRKYM
                  *** ****** ***. ::    ****        .   ** **.*    .**  * **

42                YKYWWYMYWKMYCKKYKRYCKWGCTAYCTSMYGMYSMRKRYMKWKYKYSRRTAGGAYKKR
45                YKCWAYMYWKMTAGKYKRYYKAKMKWYSYSMYKMTGMGGGCAGWKYKYSRRCRRRGCKKA
                  ** * ****** . ***** *   . *. *** * .*      ********    . **

42                KWRAKYKTAGYMRGAYYRKWRYGYYWMSKRCGRRYWRWCAKCTYRAGGGKRRMKGRKTRA
45                GWGMKTGCWAYMRAWYYAGTACSYYAMSKRTARRCWRWGRKGGYGRRSKGRGMKRRKCRT
                   *  *    .***. **       .** **** .** ***  *  *   .  * ** ** *:

42                TKCGMTYKKKYWYRGKGGYMGCKWWKKGKKGYATYKCKYKAAGCCYWYSKYAAYSKKKWK
45                GGARAWTTKKYTCAKKTTTCKYKWWKKKKKKCGCYTYKYKTGRMGYWYSKTRCYCKKGWK
                    .    .***    *      ***** **   . *. ***:.   *****   .*.** **

42                MCGGATSWRWYRYRSKTCTRWWYYARKYRMYSWRGAATGAYRWSYWKKYYYCMKMGWWRR
45                AYCAGGSWRWYRYRSKYYWGAWYYGRKYRMTSWRKRWKKMYRASYWKKCTTMMKMAAARG
                     .. **********     ***.***** ***    .  ** *****    ***.  *

42                KYYYKKGTGYCCWRYTGKKKWRGGYGRRWRWGCCYMTRKWAGARWATTCATKKWMRRKRY
45                KTTTTKRYKCATWGCYKKKKTGKRTKRGARAKMMYMKRKAGCRGTCCGTGKKKWMRRKRT
                  *   .*    . *    ***       *  *    **.** .     .    ..********

42                MWMMWYYRRRKWYRKR
45                MWMMWTYRGGKWYRKG
                  ***** **  *****
```

The pairwise alignments of two accessions 42 and 46

```
42          RYWGGYRRYGRRKTTTWWKRYYYRMCYWCKRRRYRRAAYYWRRWMMKKYMRYRWKYYMRC
46          RYWRKYRGYTRGKCGCWWKRTCCRAYTTGKRRRYRRWRCTWRRWCMTKYMRYRWKTCMAY
            ***  ** * * *   ****   *     *******    **** *.********  *

42          RYYRSMYYYYRRWCGGMRMKYKYYWKRYRGRYMYYMGRCCTCKWRYRYRYKWYRYSYWKR
46          RTYASMYYYYRRWMKAMRMTCKYYWKGCAKGYMTYMRRMYCMKWRYRTATKWCAYCYWKR
            * * *********  .***. *****     ** ** *    *****   **  *.****

42          RKYRWWWYYRKWYSRGMRCYTMMCYRKCYKMMRARARYRYRWKSWWSYSCYMAKRKRYKM
46          RKTRWWWYYRKTYSRAMRTTWACYCRKYCKMMRWRRAYRYGATGAWSYSSYMRGRKRYTA
            ** ******** ***.**       **  **** *  ***  .. ****.**  ****.

42          MKYRKYMWKMWRMYSWYKSRKKYMYYTRKRMCWKMYYWRMRMYMGCYTYYRYWKYRRCTT
46          ATYRKCAWGMWRMYSTYKSGTGYMYYWRKRATWKMYYWRMRATMRYCYTCRYWKYRRTCC
             .***  * ****** *** . **** ***  *********  *      *******

42          YMKMRRMRRRMKYRYWYRRRRWMKYRRMMRYWWWMCCCMATCRYTGMRAACRTYYSCKWC
46          YMKMGRMAAGAGYRYWYRRRRAMKYRRMMAYWWWMTGACWYYRYCTMRMWTGKTCCYTAT
            **** **     ********* ******* *****  .    **  **    .  . .

42          KMYWKMWRCRYRYYYRSGKMCMGYRGTGRKKRAMYYKRGTGYRAMKRRWKKYKYMKYRMY
46          KMYWKMWRYRYRYYYRSKKMAMRCRACARKKRWMYYKRRYRYGGMKRRWKKCKYMKYACY
            ******** ******** **.*  *. .**** *****   * .******* *****  *

42          WRGGTTCYAYRRYYGTRWWKSYSYYYYRYYKMWMRKMRYRYRRMYACKKWYYSCTYKMWR
46          WRAKWYTTGTRRYYAGGWWKGYSYYYYGYYKMWCAGCRYRTGRMYRYKTWYTSSKYTCWR
            **.     . ****.  ***.****** *****    ***  ***  *.** *..*. **

42          WYYYYKGMMRWMMWRMMRWRYKKAASYCRTTATGTRTRKKRCSARKRKYYMYMTYRCAAR
46          WYYYYKKMMRWCMWRMMRWRCKKGGSYTAWATCTWRYGKKRASGGKRGYYACMACRSRRR
            ****** **** ******** **..**   ::   *  ***.*. ** **  *: *.  *

42          RKCARYMRYKGKMKMRTCMRRYMMKYYATWYYRYKGRMAGGWTTARARYTRRRYRYWMCR
46          RKTGRYMRCKKKMKCRWYMAATMMKYYCGTCCAYKKRMWKATACGAGRTYRRRYAYWCYG
            ** .**** * *** *  *   *****.     ** **  . : . .*  **** **

42          CAWKKTACRTKWTTGCGKRYMWMRWMGMYYYWYWGYARTMWAGRKRRWYYTCATMGRMCK
46          TRWKTWCYRWGTYWKTKGAYCAMRWMKMCTCACAKYRRYAAGRRKRRWYCKYGYMRRMYK
              **. . *          *  **** *       * *   . ****** . . * ** *

42          RRMGTRRKMCWYYRGWGRRMMRYAYYWMWRMRYRKYRRYKAWRATAAGAYRKGCWMSKKC
46          RRMTWRAKMTWYYARTTARMMGCCTTWMWRMRYRKCRRYTRWRWGGGAGYRKKMWMGTGY
            ***  * ** ***     ***  .  ********* ***. **  ....***  **..

42          RWWWKYWWYYKRYKKKYWTYKRYYCAKMSTAYWKYTTTWWCKTRKYGTYKACKMMMGGGG
46          RWWWKYWWYYKRYKKGYWWYKGCTYGKCGYTYWKYWWKAAYKYRKYKGYKGTKMMMKTKK
            *************** ** **    .* . :****   .  * ***  **. ****

42          ATRWWWWWMWYWKKKYKKSWCRYRYRRYRRRKYRKKGYRSKCYGAGMSYKYYWRKRACGT
46          WCRWWWWWMTYWKKKYTKGTMRYRYRRYRRGKTGKGKTACKMCAMRMSCKYCTAKRRMKW
```

```
              ******* ******.*.  ********* *  *    .*  .  ** **   **

42          SRKWYMWRTMYMYKRKYRTAWSYMRYYSRRKCGGGKYCTMWRRKRMRGKYYYWRRYYYKR
46          CAKWCCWRCMYMCGRKYRCRTGYMRYYSRRKTRRRKYAGCARRKRMRRKYYYARRYYTKA
            . **  ** ***  ****   .*********    **.   ****** **** **** *

42          RYMRKATKKRRYCAGYAWYMKWTTTTTRRMCAASYRKGAWRSYKKKKRWKRKWWSYYKCC
46          AYMGTGWKKRRYMGTCTACCKWYCAYKRRMMGWSYRKKRWRCTKKKKRTKGKWWSYYKMT
             ** .. ***** .  :   **  : .*** . ****  **. ***** * *******

42          CTAWMYWCTTAKKWAKRYYWKRYKAACAGWYYRRYMKGATGSYSMRKKYKKRARYRGYAT
46          TAGWMYWACCGKKTGKRYYWGRYKGWTTCACCARYMKAGWASYCCRKKYKKRCRYRKCRK
             :.****.  .** .***** ***.  :     ****.. .**. *******.***   .

42          SCWYMYYYRMWYMKKKAAYKWKYARGCCCCTKRYWYYWWGAKKRRKTWMWSKRRKKRRTR
46          CYATMYYYRMACCKKKCCCKWKTGGATYYYYKRYWYYWWCTKKARGWTCASTRAKKRRCR
            .   ******   ***.. ***  . .     ******** :** *    *.*  **** *

42          RYCKRRSMMKTCCYMGMRMRKRCAKKYRWMKRRRRYRCYYRWYRMWWWGTRRKYMCKRKK
46          RYYKRRSMAKYAYYMTMRAGGAYCKKYGAMKARRRYGAYYRWTGAATWRKARTYMMKRKK
            ** ***** * . ** **     .***  ** **** .****     * . *.** ****

42          RMCGAYYYYRTATWRRYYYRRKWKRKRYKKMYKWKYKRYKCMTCMGRMMKYKMYYAKKMR
46          RAYRRCCCTGCCCWRRYYYRRKTTRKRYKTMYKTKTKATTYMYMCKRMAKCKMCTCKKMR
            *          . ********* .*****.*** * *  . *    ** * **  .****

42          RGYMTCAGYCMWYKKMGKKRYCMKWKWKKRTRAAYRYMKYMCSKYKGRKYRTARKKRYYK
46          RKTMATRKYYMTYKKCCKKRYYAKWKWKTRCACTTRTCKYCMGKYGTRKCGGGRKKRYYK
            *  *:   * * ***  ****  *****.*  .: *  **  .**  **   .*******

42          KYKKYRAYGAKYMAYKRRKKCMTWTWYKKKKYRRRWKCRYWCYYGTMKYYRRYYRTCKRY
46          GYKKCRCTATGCMWYKRRKKTMGWYWYKKTKCAAGWKAGYTYCTKKCKYTARYYRGTKRY
             *** *. .:  * ****** * * ****.*    **. *     . **  ****  ***

42          YYRARTRRWWCKKYKMRYCMYGMKKKKSKKWCKYKKAGKGATTKACACWSCTYCTRKYYK
46          TYGGRCGRWAMKKCKAAYSCCKMTKKKGKKWYGTKKTAGTRYWKCTRYWSTWYTCRKYYK
             * .*  **  ** *  *.   *.***.***  **:.     *.   **  *   *****

42          WYMAGASRMYYRMYYYMYKRYYAYGAYWYTYKGARGTCYRRRRAWKKYYWYKYYRKRKYT
46          WYCRRRGAMYYRMYYYMCTRYYCYRWYWCCYKTGGRYAYRGAGWAKKYYTTKYCRKGKYY
            **    . ********* .***.*  **  **  .   .**     ****  ** ** **

42          YWCCCCCGYSYRRSYKRRSGCRKKSKKSTCTRKRAKWRWSYYYYRCCCMYYRKRKYYRTC
46          YWMSATTAYGTGASYKRRCTYRKKGKKSYYWRKRMKWRWSYYYTGYYMAYTAKRKYYRWA
            ** ..  .*.   *****.  ***.***   *** ********      *  ******  .

42          RCTKYRAKGCTYKRYCMTYRGTMWSKRYTATYGYYKKRMAYRKKRCTRRWSYTGYKKMKR
46          ATYKYRRKKTYYKATYMKTRCAMWSKGTKCCCRYCGKGCGTRKKGTCRRWSTYSYKKMKG
               *** *   **   *. * :****  ..   *  *  . ***   ****  .*****

42          CTRGCKYKSMMYYWTRKTCYGATRCGTWYGRYRYTRRWMRYTCGGCCKRGRYAMKKGMGG
46          TCGTTKYKSMMYTTGRKWYTAGCRYRYWTTGCRYYRAWMRYATTCATGAKAYRMKGAARK
                 *******   **   .. *   *    ** * ****:   .     * ** .
```

```
42      KRKRMWYCGCCGRKSAMYYYSRWGTCRMYWYRYMKKMRYMKYKKYAAYTCKTKKKTKKKS
46      KRKRAWYTATYARKSGCYYYSRWACTGCYWCAYMTKCGCMKYKKYRCCWSKCKKGKKKKS
        **** ** .  .***. ******.    **  **.*   ****** .  .* ** .****

42      WRWYYRKRTTTARYYRGYSWGAATGAAACYRWSYWKGATTKYYWKYKKYYYMKYYWWKRR
46      WRWYYRKRAGAMGYCARTSWKRWKKMWRYYRWSYWGKRKYKYCWKYKKCTTMKYTAWKRG
        ********: :  *    **   .     ******   . ** *****   ***  ***

42      KMYYKKWYYRYKWRYRRWGCCYRYYTMGRCCYMGGGCRKKKKMWKRMRRKRYGAYTCAKM
46      KMTTTKWYCGCKWRYRRAKMMYRCTCMRRTTYMRRRYRKKKKMWKRMRRKRYKWTWMRGM
        **  .***   ******    **   * *  **    ***************       *

42      WMMKKYWYGYRRRWYRKRMYKTTYWYKWAYWCRKAYYYRSWYMKRYRTTWWYWMYKKYYW
46      WMMKKCWTKYRGGWYRKRCCTWYYWYKWGTWYRGGYYYRSWCMKRYRYYWWCAATKKTYW
        ***** *  **  *****  .  *****. * * .****** *****  **    ** **

42      TYRAWRKRKMAYTGYSMKTKMMRYWKGYYKYMGGACARRWYAYYRSYMYYAWMKRMRWKG
46      WYRRARKRGCGCAAYCCKCKMMRYWKATTGCMRRMSRRRWYRYYRCCCTTMWMKRMRWKR
         **  ***  . :.*. * *******.    *   . **** ***.     ********

42      TYKGKKWYWWWYKKYMRYKRYWYWRMMRKYWSYYGTRGWKRRGTAWTAAKRKKYWGKSYA
46      ACKKKKWYTTTTKKYMRYKRTWYWRAMRKCACYYKKRTWKAGKYGTCCGKRGKYAKTCYR
        : * ****    ******** **** ***  .** .* **    .  ..** **  ..*

42      WYMMCCKCGKMYKTTTYCKMTWYRRRRKAGRRRWRKTWYKRKKGYRRYRKKYRRRRYKMR
46      TYMMTTGTAKMYKKCACYKMKAYRRRRKMRRRRTRKKTYKRKKKYRRYRGKYRAAGYGMG
         ***    .****. :  **. ******  *** **. ***** ***** ***   * *

42      YRGYYWTCCTRRTTTGYGRGWWKRKMGGAKYRMMRRMRMMTTYMWMYKYRWRYYKYKWYA
46      CGKYYWCGTARRYYWTTRARWWKRKMRRMKYRMMGGMRMMKYYCWMTTYRWRYTGCKWTM
           ***   :**        ******   *****  ****. * ** .*****   **

42      TAGCAGGKYSTRGYACRKKRWWWKMWWYCTCKYSKRRRMRKYAWMRYRKSYRKRRRRRAG
46      ACTYGAKKYSCGKCTARGKRWWWKMWWYACTKYSKARRMRKYRWMRYRKGYAKRRRARTT
        :.  .. ***    :.* **********.  **** ****** ******.* **** *:

42      MRYYRKKGYRRYYKYRWKGMYGAYYRWCGTRTMAMRRKMTTYAYYRSGRRTCGYKWAMYY
46      MRYYGKKRYRRYYKYRWKKMYRRYYRWYAKRKMGAGRGCKYYWTYGSRRRATKYGWMMYY
        **** ** ********** **  **** ..*.*.  *  . *  * * **:  * * ***

42      KYMRKKSRKKWYYKYSRRTAYRTYKRRYWRKYGYWYGCRKKKRWKYWYKRYRRYYMKWYR
46      KTMGKGGGKGWYYKYSRRCGYRYCKARYWRKCRTTCAMRKKKRAGYTYKRYRRYYMKAYR
        * * * . * ******** .**  * *****     . *****  * ********** **

42      TWKYMSKTRKRKYWCGAYKKCTKYYKYKKRYRRTGCRKKKKRTGYGKKGAAAKRYYMRCK
46      KATYMSKAAKRKCWYRMYKKGGKYYKYKGRYGRWATRTKKKACCYKKKKRWRKRYYMAYK
        . .****: *** *   ***  ****** ** * . *.***   * **    *****  *

42      KARTCTKAYKYRRCMMYRKKYRYGTYCKGRGGRYRYYRYRCWYCYMYMGKKYWYTRCRYY
46      KTGATCKWYKCAAYCAYRKKYAYACCYGRGRRAYRYYRYRYAYYYMYMKKKYWYGRTGYY
        *: :  * **      ***** *.         *******  * **** ***** *  **
```

```
42          YATYYYWGKYSKYRCKKSMGRRRYRRKWRWGGRYKKRWYYCTKRRMYRYRRRYWWYYAGR
46          TWYYYYWKKYSKYRYTKSMKRRRYRRTTRTAAATGGGWYTYWKRRMYRCAARYWATYGKR
               **** ****** .*** ******. * ..     **   ******   ***  *. *

42          MKWCTTMRAYYYYYRMWMKMKTYYYKRYCCWMKKRRMTTMRRWMSSRYGYKKTMRKMGTW
46          MTWSYYCAGCTCYTRMWMKMKCYYYKRCYYWMKKRRMKYMRRWMSSAYSYKKWCAKCSAT
            *.*.    .   * ******* *****   *******. ******* *.***   * .:

42          MMKRRKYRYMMMAKKRRRYMYYGYRYRWRRWYYTKKYWMKTKRKKRGYRRYYKWWKYSKM
46          MMKGRKYRCMMMGKKRRRYMTTKCRYRWGGWYYWTKYTMGGKRKKRRYRRCTKWAKYSKM
            *** **** ***.*******    ****  *** .** *  ***** ***  ** *****

42          YKYKTCMWKRYWRRKWKKKMKKMKYYKKRKGTATGRRKRRKKGCRWKRRWYSRYWWMKKC
46          TGYKKYAWKRYWRRKWGGKCKKMKTTGKRKAAGWARGGGRKKCYGTKRRWYSRTTACKKT
              **.  *********  * ****   ***.:. .*   ***    *******    **

42          YYTYSKRKYRRTKTCYGCKRRTCYYYRKWYKMYGYGAGYYRYYYYRKKCCYCYKTKMRMR
46          YYGCSKRKYRRWKWYYRTKRRAAYYCRKWYKMYRCRMKCYRYYTTGKKATYYCKKKMRMR
            **  ******* *  *  ***:.** *******      ****   **. *  *.*****

42          YYYRGAYRMKGYYYRKCKWYAYWATGRRMWWYRYTRWCRYWRKTARTCGRYYRAYAAWYW
46          TYYRKGCGMTKYTYRGYKWYGTACCARRMWWYRYCRWYRYTAKATGGYRGYCAGTTMWYW
             *** .  *. * **  ***.  . .******** ** **  *::     *  . : ***

42          KKKRSYKRKWYRCKRMRCYYYWWKATGGKKAKTYAKAGRWWKMARTCTKGCYRYMGMKMY
46          KKKRGTKRKWCAYKRCRYYYTWWKGCAAKKMKYCRGGRGWWKMTGAGGKRYCGCAKCKCC
            ****. ****   ** * ** ***. ..** *    .  ****: :  *        *

42          YRRRMRTKSKGGCGRRWKKRTGCCYRMKRKCAACWMRYKWRMRARKAKWMGYYSWYYYTR
46          YRRRMRATSKTAAAAGTKKACCAMYRMKRKMRGTAMRYKWRMRGRKMKWMKYYSTYYYKR
            ******:.** ...   **   . ******  .  ********.** *** *** ***.*

42          YRKYRKKRYRMCAGAACAKKTRRWKTYGRWKGKKYRRKRRGYMRKKCMACYRRMMRCRKR
46          CRKCAKKRCRMGCRRMYMKKYARATWYRGWKKKTTGAGARKYMRKKACGTTGAACRYAKR
             **  *** ** .     **  * . *  ** *.     * *****. .      *  **

42          KMGWRYRRRGKTTCTSKYKRWYYYYWAYRYYYRRYRRYTYMRWYKMMRTGRCGCATKKYW
46          KCKWRYRAARKGGTASKYKAAYYYYAMYRYYYRGYRRYKYMATTKMCGGTRYRMWAKKYW
            *  ****   *   :****  ****  ****** ****.**   **    *    :****

42          RWGCAGRGYMTTAGCKWRWTGAYYRAGAGCTWYMRRRRKWYYGTTYYRRYWKTARCTCMY
46          GWRTCTRKCMCACTYGARWCCGYYRCRWKYYWTCGGRRKWYYRYYTYRRYWKYRAYKYMY
             *  . *  * :.    **  .***.     *    ******    ******    . **

42          TCGAGCYYAMKRYRKATYTRMKSRYYRYMMYYYTWKKKKCMGTMMWCKRRRYYRKMYYRK
46          CYKRAYYYRAKRYRKGAYKRMKSRYTAYMMTCCATKKKKYMKCMCAGKRRRYYRKMYYRK
                . **  *****.:*.******  ***   : **** *  *   *************

42          RKMWTGKMMYKMYRRWYYWKRGGAGMWKYWYAYYATAYRRYKCRWKYTCGKWMKATCKKK
46          RKMWKKTACCKAYGRWYYWKRRRMKCWKYWYRYCWYWYGRYKYGWKYWYRKWMKMCTKKK
            ****. .   * * *******     ***** *    * ***  ***   ****   ***

42          RTGAKKRYKKYGRMRRYTRYRSYTKKWRYWKSCKTYRRYMWKGAMKMYRRWMYKKKKRCA
```

```
46              RCTCKKRCTTTRRCGAYKRYRSYYKKTAYTTSGKKYRAYMWKRCMKMYRRWCCKKKKRYR
                *  .*** ..  *   *.***** **  * .* *.** **** .*******  *****

42              TKKRYYKAKCTYTKYRCAKKKTRMWMWTGKKWYKKKGRKAKRMAYRYGRSKKKYTTAKGR
46              KTGRYYKTKMYCYKYRYRKKKARMTMWYAKKWYKKKRRKWKGMWYRYAGSGKKYGAGKKG
                .. ****:*    ***  ***:** ** .******* ** * * ***. * *** :.*

42              MKGYYRCTWRKRWRYWKKRTGTCCTAKKAKKGYRRAGCAATMRRRKGTKKCAAAGMKMAK
46              CKKYYRMKWRKGTGYWGKGCRYSTYGKKMKTSYRRRCARTGMRRAKRYKKTTTTAMTMRK
                 * *** .***   ** *    .  .** *..***  . : *** *  ** :::.*.* *

42              YRRRKYMCTRMRYKKWMYAKCGYWMMMKYWMMWMYRSMMRRGWKKYRMKYRMKWKYMWWK
46              YRRGTCCYAAARYKGWMYGKYTYWCMMKCWMATCYRSMARGKAKKYRCKCAATWGYATAG
                *** .   :  *** ***.*  ** *** **   **** *   **** *   .* *

42              GRKRTYWYWMTRKKRTRGRKWKMWYRKKYYKRRRTRAYMTGWMYRRWCYYYAKAAYYGGG
46              ARKRYTTTACGAKKGYRKGKWKMWYGTTCCKGRAYRRCAARAATGRTGTCCMGWMTYAAA
                .***        ** *  ****** .. * *  *   :     *          *...

42              RSMWKMCTGRWGYRRKRYCKTWSKYAACRYYKRWMRATAYRTRKMWRYTAYKKRWKRMKS
46              RSMWKMGCAGWKYRRKRYMKWWSGTMGYRYYKRWCARWTCGCAKMWRYYRCGKAWKRMKS
                ******  . * ****** * **   . ******    :    *****    * ******

42              YWY
46              YWY
                ***
```

The pairwise alignments of two accessions 42 and 47

```
42              YYWKYYMYWKYMYRRACWKGKKWRYRYSWKKCRWKYYYRMCYTKRRRYRMYYRYYKMMRR
47              YYWKYYMYWKYMYRRCYWKKTTTRYRYGTKKAAWKYYYRAYTWTRRRYRCYYRYYKMMAA
                ***************. ** .. ****. **. ******    .***** ********

42              YMYRWYAYWKYRYGGCCCKMMAYGTRYKAYRMKYRRRMRRWYYRRRGWWRYWRYRYWTYY
47              TATRTTMYWKYRYRKYYYKMMTYTCGYKMYRMKYRAACRRWYYRRRTWWRYWRYRTWCYY
                   *   ******     ***:*   ** ******   ******** ******** * **

42              YRKGYRWWYYRWWRYSRMCYRYYYYSRRKRMRYMRYRRWKKKKKKCKWRWMKYTKRCRSR
47              YRKKTRWWYYRTTRYSRMYCRYYYYSAAKRCAYMRYRRWKKKKKGTKWRWCKYAKRTASR
                ***  ******  *****  ******  **  ************  **** **:**  **

42              MWMRRRWRYRYSTTYYKYWWWMRSRRCCYTKKSRKMYKMRKRYRYMKCRYMMRYMWMWKY
47              MWMRGRWRYRYSGYYYKTTATMGGAAYYYCKKSRKMCKMGTRYGYCKTRYMMATAWMAKC
                **** *******  ***    * .    * ****** ** .** * * ****   ** *

42              RRMYRYWCYTWMYTRKMRCCMYYGRYMKYKWKYRRYKSCTKYYKMKRMKMMKKMYRCAYK
47              RRMYRYAATWWMYWRKARYTMYYTRCAKYKAKTRRYKGTYGYYKMKRMTAATKMYGYTYK
                ****** .  *** ** *  *** *  *** * ****.   *******.  .***  :**

42              SYWRKRMKRMYRYRRRYWYRWKYYYWRKYRRKKRYKGYYATTWKMKKRKYKKKRSWCRTG
47              SYWAGGAGAMYRCRRGCAYRWKYYYWRGCRRTKRYKKYYTGYWKMKKAKYKTTAGTMRAK
                ***      *** **   *********  **.**** **:  ***** ***.. .  *:

42              RYKYRRRWYYRKGYRSKYYMWAMARATAATKMTTYWRRWKMSSKWKYYYKRWRRKKRMMY
47              RYKYRRGWTTGKKTACKYCMWRCWRRCCGCKMWYCTARWKMSCKWGCTYKRWRRKKRMMY
                ****** *   *   .** **   *  .. **     *****.**   ************

42              KTMYMKMMRMTGYKYRWMRYYRRACRWRRWYYRGGAAKKKRTMWYYWKMYRWRYACGTYK
47              KCMYMKMMRMGKCKYRTMRYYRRWTRWRRWYYRRRMRKKKRKMATTAKMYRAGYTTTCYT
                * ********   *** ******  ********    ****.*    ****  *:   *.

42              YSWSKMRYRMYYYSCKMWRYKYYYKGMMRWTAGYRWWYKAAASYARKRRRWTAAGRYSRK
47              CSWCGCRYRMYYTSSTCWRYKYYYKKMMRWYRRYRATCKWGGSYMGKRRRWKMRKRYSGK
                 **.  ****** *.. ******** ****   **   *  ..**  *****.   *** *

42              MTYYMGKKCAAMYRRGAYMYYKGGMAMMMRRWMACYYTRKAGGATTTMRAAYWKMTRRRY
47              MWYCMKKGSRRMYRRATYCCCKTTMMCCMAAWMCGYYARKTTTWCGKMARMCWKMYRRRY
                * * * * .  ****.:*   *  *   *  **. **:**:     .*    *** ****

42              RCWYYTKYYRKAWRYMRWGAAGKGKRYCAATWMYKAKYYWYKTTKSKYKMYYYCCKTCYM
47              GYWYYGKYYGKCWRYMGWSMCATAKRCYMWYWCCTWKYYWYKYYGCGTKMYYYGTKKYCM
                  *** *** *.**** *. ....**     *  . ******    .  *****  *.  *

42              RKKKMYRWWMYKYKRRYRYAYYCKRKAKMKMMKYRYYWSKYKYMWMRRYMRMWSKRYYGK
47              RTKKCYRTAATKYKRRYRYRYCSKRKGTCKMMKYRYYWSKTGCMAMRRYMRMWSKGCYCK
                *.** **    ******** * .***.. ***********   * **********  * *

42              RKTMRCTRMRRWSACYKMYKYMYWTRYKGGATYGTTTRYYARWYRYYAMRYMYTWMKWRK
47              GKYMGTCRMRRWSWYTTMCKYMTWGRYKKAGCCRACGRYCCRWYAYYGMRYMCGWMTWAK
```

```
             * *   ******   .* *** * *** ..   :  ** .*** **.****  **.* *

42          YKRMKRRRAYRCGCRRKSYYYSRKYWTKKKKMWKKYGAKKGKMKKKRRGKYKMMRYKRKR
47          YGRAKRRGGYATATRRKSYYYSRKYWGGKKKCWGGYRRKKKKMKKKRRRKYKCMGTKGGG
            * * *** .*  . ************  *** *  *  ** ******* *** *  *

42          RYKKKYKATCYWYYSGWRKCWYKTGRGYWWMKRWRGMYYKRRKKAKMRKYATTRRWKYAR
47          RCKKKYKMGTYTYCGAARKYWYKCCRTTTACKGAGKAYYKRRKKMKCRKYTAWRRWKYWR
            * *****   * * .. ** ***  *     *     ******* * ***:: ***** *

42          YYYRKKWYKKMRMRYYRWYKMKKYYKWAGGYMKRAYYATCWTWTTRRMKKRRWRWSYWRM
47          YTTGGKWCKKMRMRYYRTCKMTKYTKWRRKYMKRCCCRCMTCWYKRAAKKRRWRACTWRM
            *    ** *********  **.** **   ****.       * .*  ****** . ***

42          GTTTCYKATTWWWSYKCYYGTGKCWRYWKMRKWCTRYYRKYTMYTTMKKRYRYRRKYSGY
47          KCCGTYKRKYAWWSCKYYCAARKMWRTWGMRTAAGRYYRGTGCCWYAKKRYRCAGKYSKY
                 ** .  *** * * .: * ** * **. . ****        *****   *** *

42          SMRMSYKRTAARYRGMMWYWMYYYGGTWKAYRMYKWKYRGCKRYWAKMTATCTTMYWWRR
47          CCRMSYKRKGGRYRRMMATWMYYYAAKAKWYAMCTWKTGAYKRYWRKMAGAAYYMYWWAR
            . ******...*** **  *****... * * * .**  . **** **:.:.  **** *

42          YMWMWKYYTAGRKKRWWYRYMRGCYSYKYGTKYYYKRKMWKRRWRYWKKTKCKKCKASKR
47          YMTCATYCKRKAKKGWAYRYAGRYYSYGCTCKYYTKRKAWKRGTGYWGKGKATKMKRCKR
            **   .* .   ** * ***    ***    *** *** ***   ** * *..* * .**

42          RKKKMGGKCKMKYKRKRMYKTMMRTKYKYKMYAKTYRRYMMCMRSMGCRRRWKKYTGAAY
47          AKKKMRRGYKMKTKRTAAYKAACRCKYKYGMYGKCYRRYMMYCRSMRYRGRAKKYCKRRY
             ****    *** **.  **:  * **** **.* ******  ***  * * ***    *

42          MMKTWKWRTRCGKCYACMWRWMGRGWTYRTATRYRWYYRYMAGTAYWRYWYYRMKWYACC
47          MCTKWKARWRTKGGYGMCAGTARRRAKYRYRYRYRWYCACARKWCYWRYWYYAMKWYWTS
            * ..** * *    *.       *  .**   *****       .******* ****  .

42          YYYMMRYGWRWMKGYMAAWRKRAAGRCGRYCKKKMRGGATGYRWRRKYKCKYKYYMWMYW
47          YYCCARYRTRWMKRTMRRWRKGTGARYAATTTTKMRKAGATYRARRKYKAKYKYYCWMYW
            **   **  ****  *  *** :..* .   ..*** ..: ** *****.***** ****

42          YYMRCKRTWKAYKRSAYYKTGCMTKRMRRKRRYKTTTRMKRKGRGGRSMYYRYCMWRKRY
47          YYMAYKAYWKWYKAGGTYKKAYCKGGAGAGGGYGYWYRMKAGAGRRASMYYRYMMWRKRY
            ***  *  ** ** .. **..  .        *    ***  .    ****** ******

42          YWRYAYYWRMKGKWKMKSMMGRRRYGRKRKKCTAYCTGRRSRRKYRRRYGAWYYRWGGTC
47          YWRYRTTTRMKKKWKMKSMMKRRRYKRGATKYACYYWTRRSRRKCAARYRMWTYRWKRAT
            ****    *** ******** **** *  .* :.*   ******   **  * ***  :

42          YKYAKYMYRWKMGAGAYYYYWWRSGYMMYRYYAMYYKKYRKSRKRMYKSRYMWKWYKKYS
47          CGCTKTMYAWKCKWRGCTCYWWGSRYMMCRYYGMYYKGTRKSRKRMCGGGCAWKTYKKYS
               :* ** **    .   *** * *** ***.****  *******  .   ** *****

42          RRYRYKKKRYYWKTATGGRRYYAKKTAKYKKTRYRYMTCKYWYKMSKAKRYWMWKCTKYK
47          RRYRCKKKGYTTGKGKAKRRYYRKKCWTYKGYACAYCYYGYAYKMSKWKRCWMWKGGKYG
            **** *** *   .... **** **  .**     *    * ***** ** ****  **
```

```
42          YRKCYGRKSGKWRRKKTRKKRYKYYRATGYGCCWTYMTKRAKRKMWYKYYYKYRAYWMKK
47          YGKYYRRKSRTTAAKKCRKKRYTYTARATYKYMWYTACKRRKRKMWYKYYYKYGCCWMKK
            * * * *** .   ** *****.*   : *   *    ** ************ . ****

42          KKTAYWMRSSRYGYTKWYRAKKRKTCWRKYRRSRRCYMMRMRYMYKYYGGYRYYWRRYYG
47          KKATYWMRSSAYSTWKAYGCKKRKGYWRKYRRSRGYYMMRCRYATGTYRKCRYYWGGYYA
            **::****** *.  * * .****  ********  **** **    *   ****  **.

42          YYKKKKYWKTCKKKRGYRRYKWWYAWYKMYMGGATCKKGRYRKWGTYWWYCYRSYMYYKW
47          CYTKKKYTGGSKKKRRYRRCKWAYGAYGMTMKRGGMKGKRYRKWKKYWWCMYRCCCTTKW
             *.****   .**** *** ** *. * * *  .  *  ***** .***  **.    **

42          KYRRWKRMKSMAGYKGKKMWWWWYYMMKRSYCYYKCTYRKKGRYKWWYRKRRKGGRAKCR
47          KYRRWKRMGSMGAYKKKKMWTTTTYMMKRSTGCCKMGCRKKKRYKWWYRKRRKTRRRGAR
            ******** **..** ****    ******    *   *** ***********  *  .*

42          KWKSYACYYACWKKYKRKKMYYYKRKAACRMKWTAKKRRKKRKKSRKRKYRKYRGRTRRR
47          KATCCRTCYRYTKKYKRKKMYCYKRGWWTAMKAYMKKRRKKRKKSRKRKYRGYRKRCAAG
            * ..    *   ********* ***     **   **************** ** *

42          KMWGRWWYRGGAKYRKSKWRYKYKWKAGKGATTACRACWYCTGWKMKCRCWSYRSYSRYM
47          GMWRAATYGRRMKTRTSKWRYGTKWKWRKTRYWCTARYWYYKKWKMKMATWCCRSYGAYM
             **    *    * *.*****  ***  *    .    ** . ****   *. ***. **

42          YRRMYYGMYKRSTAACWYYKRYWTKWRTTYKRRRRAWKKYKYRWKRKGYMTWCCCGYSYR
47          YRRMYYRMCTRCATGTWYYKRYTCKWRYYYKRGAGWAKKYKYRAKGKRYCYWATTAYGTG
            ****** * .*.::. ******  ***  ***     ****** * * *  *.  .*.

42          RRKSYKRRGKTRKTAATKKSKKACYRRSMTCTMKGKWRKWSYTKKYYYRKRRMYYRRKKC
47          ARKSYKRRKKYRKCCCCKKGKKCACAGSMYYWMKTKAGKWSYCKKYYYGKRRAYTRAKKT
             ******* * ** .. **.**..   **   ** *  **** ***** *** * * **

42          SMYYRYKRACYYTTTGYTGTYWYKKRYGKKAYKGTRRYKTRYAYRKKKYWKMCTGKKTTT
47          SMYYRYKRCYYCYAARYYKYYWYKKRCKKKCTTKGGRYKKGYGYRKKKYWKMTCKKKYWC
            ********. *  :: *   ******  **. .   ***. *.*********   **

42          GAKRYKKYGRMRRYRKYWYKWRYYKKRGTTYRKTTATYTYMGCGCRRKARRYMKMYRWKY
47          TCKRCTTTRRCGAYRKYWYKTACYGTRKWYYRKWYMYYCYCRGCGRRKWRRYMKMYRWKT
             .** ..  *   *******   * .*   ***    * *     *** **********

42          RRWMRWKCCRATATCRYYKKYATTYYKACACTGTRMKTYKKYKTKKRWYRGWWATAGCYK
47          RRWMRWKTTRGCCCMRYYTGYMYAYYKWYGTCTGRMKYYKKYKYKKRACRKWWGGRAAYK
            *******  *. .  ***. *  :***  .    *** ***** ***  * **.  ..**

42          KKYKWKRKYAWCTMKAWAACCWKRYKGRYYYRYRWTKRKYKMRRRYWYKWMMMYYKRGWY
47          KKYKWKRKTRTYCCKWWRRTTWKRYKKGYYYRYRWGGGTYTARGAYWYKWAMATYKGKTY
            ********      * *    *****  *******   .*. *  ***** *  **   *

42          KKRMYYYTRSMKMYKMYRRKRYKKKWSKYCARKMRTYCTKKRWKMRARTMWYYKKRWKRK
47          KGGMYTYKRSMKMYKCYRRKRYKKKWSGTYRRKMRCYTWKKRWKCAGGKMWYCGKAWKRK
            *  ** *.******* ***********    **** *  *****  . .***  * ****
```

```
42      GSYWYTYGRYWAGRKCYYMGYYKWWRRWYYRRRGATGRKRRKKYYRKWYYYWMTRMTYWK
47      KSYWYKTKRYWCTGKMCCMRYYKAWARWYYRGRRCAAAKRRKKYYRKWYYCWMYRMKYWG
         ****.  ***.  *   * *** * ***** * .:. ************ ** **.**

42      RKGRGKKAYARKTTTTGGRGGCTKRKMRMKSYYWGGWMRTKKGGKRGRTMMWSAAYCRCA
47      AKRRKKTTTRRKCGCATSGKKYKKRKMRMKSTCAKRWMRWKKKRKRRGYMMWSGCCYRSR
         * * *.:  **   : .    .********     *** **  **   ****..  *.

42      RWRRKKYYKKMYYRKKRRYMYRRRYYWKRKRMGATRGARKYAYTTCRRYMWYAGGMRRRW
47      GWRRKKYYKKMYYRKGRRYAYGGRYYWKRKRCARGRAWAKYGYKCTRRYMWYGRRMRRRW
         ************** *** *  ******** .  *.  **.*.  ******.  *****

42      GYRYKKCRYWGKCCCTRKMSGRWGKWRRRAYRKSYKRKKWKKRKKKCYMYKGARRWKKKR
47      KYRYKKMRYWAKATTCRKMSRGATKWGARGCRKGTKRKKWKKRKGGYYMYKKTGAWKKKR
         ***** ***.*.   ****    **  *. **. *********   **** :  *****

42      RKMRGGWWKRTCKTGGWYWYWRRMMKMYYRRRMRTKKWCWRKGCGRRWRKKWRKTGCCYT
47      RKCGRRWWKRGYKATAWYWYWRRMMKMCYRRRMRWTKWYWRKRMRAGTRKKWRKCCAMYK
        **    ****  *: .*********** ****** .** ***      ******  . *.

42      TARTRRKYYKYRKKKWRRMRRSMMRKCCCGCYRAACMRRGTKKKRKRMYKRGGYMRKMMM
47      KMGKRRKCCKYRKTKWRRMRRSMARKAYYKYCGWWSMRRRYKKKRKGMYKRAAYCRKMMM
        .  .***  ****.********* **.        .***  ***** ****..* *****

42      WMKYYGRRRKRKKWKYKRYYYKYRRKMKRMRKKRWMYRKRYKMYYRMMKCKKKKYYCTRW
47      WMKCYKGRGKRTKWKYKRYYYKYRRKAKRMRKGATMYGKATTMYYRMAKTKKKKCYYYAW
        *** *  * **.************** *****   ** *  .***** * **** *   *

42      YYAKTMKARRYKYGYGCAYRWMRYRMWARKKCYKYTYYSYYYKKTGRYKRGYRGRRKKRG
47      YYRKKMKMRRYKYRCKMRCRAMRYRAWCRKKTCKCWYYSYYYKKAKRYKRACAARRKKRR
        ** *.** *****      * **** *.***  *  ********: ****.  .*****

42      CYRMKKATRKWMRRSGKYTYGKYRTGRKKKTYRKRWRRMKKWKYRYRRKRKYSKYAMYTC
47      YCGAKKMYAKAMGGSKKYCYTTYAGARKKKCTRKGWRRCGKWKYRYAAKRKYSKYRMYYM
            **   * *  * ** * .*  .****  ** ***  ******  ******* **

42      TATWYAAYRRKMYYKRYKRYKRWMKGRGCMMYARKYKAKWRKKCAGCKKKYKGMCTWWYR
47      CCCAYTTCRRKCYYTGYKRYKATMTARRYMMYMGKYKRKWRKGYWRYKKGTKAAYGWWYR
         .  *:: *** **. *****  *..*  ***  *** ****     **  *.   ****

42      WKYYYRYRMYKSCTACWWSGAATGAAKRYYYRSYWKTGGTKYKYYYMKYYWKRKKAYYKK
47      WKCTTAYRMTKSTWWYWTSKRWKKMRKRYYYRSYWGYKKYKYKCTTMKYTAKGKGWTTTK
        **    *** **    * *   .    *********     ***   ***  * *    .*

42      RRYCCRYGAAWRRRWRGCCYYYMRKCRCKMARYWMMRKRRKGTCATKWYKTCCGAAYRRR
47      RRCATGCCRWWRRRWRKMMYCTMRGTGTKMGGYWAMRKRRKKWMRWKWTGYYMTGGYRGG
        ** .      ******   *  **    **. ** ******     **      ..**

42      RRRMRYATKWYMYRYMRKRYRWMYSSWYMKKKKCKWKWMKKKKYMYKYCGGRYYWYRKGK
47      RRRMRCGCGATMYRCMRKRYRWMYSSTYAKKGGYKWKWMTKKKTCYKYMRKRCCWYRKKK
        ***** .    *** *********** * **   *****.***  ***   *  **** *

42      WMTACYGAGKYWAYACRRKAYTKWWYKKKKRWKRYKRKRCYYRGYMKGRYRRCYYRKYYA
```

```
47              WCKGAYTTKGCWMCCMRRKCCWKWWYKKTKGWKRYKRKRYCTRKYCKKGYARYYYRKTYR
                * ...* :   *  . ***.  ******.* ********   * * *  * * **** *

42              RWKCRKKYKYYGYYSRRRMYKMGYMRYRYWRAAYKRWYYARWAMRSTYRRCGCWKRWRWW
47              RAKYRKKYKYYKYYSRGGMYKMKYMRYRYWRMTTTRWTYRGATMRSCYGGTKYWKRWRWW
                * * ******* ****  **** ******** : .** *   :*** *     *******

42              WKCGGYCMKRRSKYGMMRTASYATRKKKMYGCKRTRRKGRKAGMRYYRKTTKYRRYRYKR
47              WKAATTTAKRRSKTACMRGGGYWCAKKKMYAMGRWRRKRAKWKMRYYGKKYKYRRYRYKA
                **..    ***** . ** ..*   *****.  * ***  *  **** *. ********

42              TATYYRMTYKYYRWYMMWKMMRKMYYR
47              YMYYYRMKYKYYRWYMMWKMMRKCTYR
                   ****.***************  **
```

The pairwise alignments of two accessions 42 and 48

```
42                 YWKWCRRYMKKYMYGTGTRCRYRAYTGSCKWRYRCGYRYKWYRYSKWYRWWWYYRTWYYG
48                 YWKWMRRYMKKYMYRYSKRMRTRRYYRSMKWRYRTTTATKTTRYCKTTRWWWYYRYTCYA
                   **** *********  ..* * * *  * *****     *  **.*  *******   *.

42                 CMYCTTYSMKARRKMRMKCMRRRYCACTCCYRGYRRYMMMYRWKYYYYMYYRMMKKWRWR
48                 MMCTCCYSATGAAGAGAGTCRARYARMYYYCAKYRRYMMMYRWKYYYYMYCAAAKKWRWA
                    *    ** ..         * **.        *****************    *****

42                 MGWRCYRRYYKKWKMMWRKMYYMYTRRRWMSARYKSYRYWYYYWGTRRWCCCATWYKRTG
48                 CATGTTRRYYKKWKMMWRTAYYMCWRRRTMGRAYKSYRTWTCCWTGARAMMTWWAYKACT
                    .    ************. ***  *** *.  ***** *   *   *       **

42                 RKCSRRKWYMRYMRYRKAGCATRWKCWMAKKYRMRKRYMKCYRMKMYYKMATTCKMKYRR
48                 RKYSAAKWYCAYMRYRKTCTCGATGYTMTKGYRAGTRYMKMYRMKMCYKCRWWMKMKYRA
                   ** *  ***  ******:  .      *:* **  .**** ***** **     *****

42                 RWYYRRYKKMMKYGCWWYWWMWRTMRTTKYKSKYKYRYGGRKYRRCRSWGKKRWKYKRRY
48                 ATYYRRYKTAATYKYATCWWMWGYCGACKYKSKYKCAYRRRKYRRMAGTRKKAWKTTAAT
                     ******.  .*     ****    : *******  *  *****  .  ** ** .

42                 RRYRWYMTKCTWRMYRWSYMWMYYRTRKMYTRRMRWRCYRYWTTMSMYWKGRYKTCMTKA
48                 RRYRWYMGKYWWRMYRWSYMWMTYRWRKAYARRMRWRTYRYWCCMGMYWKAGCKYTCYTT
                   ******* *  *********** ** ** *:****** ****  *.****.  *    .:

42                 TAMYYYYRYMMKGTTAGTAWRCGKRGCCKRRWYRYYMRRCRKRYCYYMSGYWTRKAKRYT
48                 GCCCTCYRYMMKTCWRRYRWGYAKRRSYKRRWYRYYCRASGTRYSCCASTYWCRKRKRYC
                    .    ******       *  .** . ******** * . .**.   * ** ** ***

42                 RWWWWWMKWRTKYKKKSWTRKRYYRRRYYRKYSAAAKYYYMMGRYTAATKAMTTGCTYWR
48                 RWWWWWMKWRCKYKTKGTWRKRYYRRGTTGKTCRWMKYCTCMKGYCCGCKRMKARYCCTA
                   ********** ***.*.  *******    * .   **   *  * .. * *.:

42                 KRRKYRWRWKYYYKRWYRMMYRTMYGKKYRMRYYYYSRRRYGCRYCTMWARMRKYYYKWK
48                 KRRKCATGWGCTYKRWYRMMYRCMYRKKYRMRYYYYSRRRTTTRYAGCAGRMRKYYYKAK
                   ****    *   ********** ** **************   **.   .******** *

42                 GYWRSWRKKKRRRYRTGGCKWGRCKCTTGARRMCWKRRWSYTTCYWWKRKWWSYYYYKKA
48                 KTWRSWGTKKRRRYAWKRSKTTGYKAAWRWRRMMWKARWCTCGTYWWKRTWWSCCCCKKW
                     **** .******    .*    *.:   *** ** **.    *****.***    **

42                 GYRCTTAKCCYYKYAGWTKRKYYYYKWRYKYYMYRRKYMTSRWKAYKKRARYRKMMWYKK
48                 KTRMYYRKMYYYKYGTTGGRGTCTTKTRYKYYMCAGGYMASRWKCCKKRGRYRKMMATKK
                     *    *  ****.    *     * ******    **:****. ***.******  **

42                 WWMYYYWKKRYKATGMMSKTGGGGKRRTYRCYYMWMWAAWCYCSRKGMKRRKYYYYRACW
48                 WWMYYYAKKRCTWYRCMSKCRKKKKAGGCAAYCCTCATRWYCYCAKAMKGRKYYCCRGTA
                   ****** *** .    ***     *     .*     : *   . *.** ****  *.

42                 MRYYKAKRKKAAWYMCGYKKMYYAGWYYYKGRYRGGCCRGKRRRGCYRMKCKCKTKRWMR
48                 MRYYKCKRKKCCACCYTCKKMYYMRTYYYKKRYRARYMAAKAGRRYCGAKYTTKCKAAMG
```

```
                *****.****..      *****   **** ***.     .*  *      * . * *  *

42              RGRSSYKYRRKRTTTARKTCGKKKYYRYSAWKRWRRKMGRYRKRRKRMKYYYKYATYWYC
48              GRGSSYTTGAGGCACTRKATAKKKYTRTSGTKGWRRKCKGYGKAAKRMKYYYKYMKCAYG
                   ***.      : :**: .**** * *. * ****    * *  *********  .  *

42              MTYMTMCCAKYTGTYYRACRYKRKYYYMRWYCYYWYRRYSKRYAWYKYRRYKGTTGRTKY
48              CCCAWCMYWKYKRWYYRRYGYKRKYYYMATCAYYWYRRYCKRCGTTTCRRYKRYWRGGKY
                         **.  ***   ********   .*******.**  .  . ****      **

42              KCAYKRKKRYKKYYMGRCRMRRTRYRSYKWRYCAKKYRYYGTCYCYKRGRCAKWYYMRYR
48              KTTYKRKKRCTTTYMRRTRCGAKGYRSYKTACYRTKYRYCRWYYYYKRSRYWTATYMRYR
                * :****** .. ** * *   . *****     .****    * ***.*  .  *****

42              WYGRRWTMKMYRKKRGCRKRWYMGYGKRTKKRGARYYYGMYKKYKKKWYMWACKKGSWYK
48              WTARRWYCKCCRKKRKYGKRACATYKKRKTGRRWAYYYSCYKKYKKTACMWGTKKKSWYT
                * .***  *  ****   **     * **.. *   ***. ******.  **. ** ***.

42              KKMYWRRWYRTMSYMYYKWCTRACARWWRWKRMKCKSMRKTCAKMRTCCMKRWMRYRKYG
48              KKCYWRRWYRKMCCCTTKWYKRRYWRWWRWKRMKTKSMRKWYRKMRWYTMKRWMRYRKYK
                ** *******.*.     ** .*   ********* *****   ***   **********

42              KRKKMYMKGYYGSKWYCKGWARRKRCKWMKKYWYSCCYWYMMKRKKMYWYRCCTTCAYCR
48              KGKKAYMKKYYKSKWCTGATGRRKRMKWMKKYWYCTAYTYMMKRKKMYACGTYAATTTTA
                * ** *** ** ***     . .**** ********. .* *********     :: :

42              MKWTAKCCYKRRRKKKRKWKRKRKKKTYRWKKTYMYAGWYGKKRMAYYGAGMKKYRATTT
48              MKACCKYYTGRRRKKKRKTKRKRKGKAYRWKGKYMYMKWYRKKRAGCCRMKCKKYRGAAC
                **   .*    ******** ***** *:**** .***  ** ***  .      ****.::

42              GRACRGCYGGYMWRYGRYSGAAYKKYCTTGRMTGGCYCTTTAWKRRGRYWMCGCCMRTKM
48              TRWYGATCKRTMWRYAGYSAGGCGKYYGARRMYRRYYYCWCGWKRRKGYWCTAYAARCKA
                 *    .     ****. **...  **  : **     *     .****  **   . . * *

42              KGGKYRRCGATTKKGAKKMKRKRMKKWYRRWMKMGRRKKRKRMYKKYKKTKKWCGMYMKM
48              KKSKYAGMCTGKKKAGKKCTRKRMKKWTRRWMKMCRGTKRKAAYKKYKKKKGWYSMYCKM
                * .**     : .**..** .******* ****** * .***  ******.* * .** **

42              MKWMKMWMAYMTGYRSMYRKYMTRWYRCAACAGAARMRKKRKGKKMRYYYTWYYTGAKWY
48              MKWMKATMWYCAAYRSMYGKYCCRATGGCWYGKRRRCGKGRKKKKMRYYYGWYCGTCKWY
                *****  * * :.***** **  *     .  .    *  * ** ******* **    .***

42              TCAKRKKWKTCMRRWMSSRYYTRKWYYKRKRYYTTYAYMRRRKRYKRGYRYYWWKRGKWY
48              CTGKRKKWGCYMRRWMSSATTAAKAYTKGTGYYKCYTYMRRRKRYKGKCRYYWWKGKKWY
                  .*****   *******    : * * * . **. *:*********   ******  ***

42              YYYYRKMKKKYGGAWGGKRGYRATAYYYWWMYWKMYKYYGMRAAAKTCKAWRKKKYRKYY
48              YYYYRKMTKKYRTTTTAKRRYRGAGCCYWAMYWKMTKYYKCRWTGKYYKWWRKGGYGKCC
                *******.***   :   .** **.:.  ** ***** ***   * :.*   * ***   * *

42              KKTYAKRRGYKKKMMRYKTGYKYAACRCRKKTYRKRYKRYYYYWWMKTASMYKTCAYRRK
48              KKACMKRRRYKGKCAGTKGACKYMWYGTGKGWCRKRYKRYYYTTACKATCAYGKATTRRK
                **:  *** ** *    * . **      *   *********    *::. * ..: ***
```

```
42    CGMCRCTYGKTCYKCCYKMYRYYRKWYYGYYYSTKKYGTGYYRCRTTGATKKKTWCAYAC
48    YKMTRACCAKAAYKTYCGAYRTTGKWYCAYCYSKKKYRYRYYRYAACRWKTTKCTAMTRY
        * *.  .*:.**     **   *** .* **.***   ***  :   ...*  .

42    AGGKKKGYYRYRRYRMSAMGCKKKKRYCYWKWKRRMYMGYKYKKRWYWKSRYASYYYRKG
48    RRKKKGKTYRCRRYRASCATMKKKGRYYCWKWKRAMTCKYKYKKRWTWKSRCGGYCYAKK
         **   ** **** *.   *** **  ***** *   ******* **** ..* * *

42    KARMKWGWYKKMYRRWRWKYRYKRKRRCYKMYYYYGRRAYYRYYKRARTRCKYSRTGGYY
48    KCGMKWKWYKTMCAAWGWKYRYKRKRRYCKCYYYTRARWYYRTYKGGRCGYKCSRWRSYY
      *. *** ***.*   * **********  * ***   * *** ** .*   * **  .**

42    KYYYRMCYGYSYKKYWWMGWKCRYGRKYRRYWCCRKKRMRKKRMMRAKWYMKMYYKRGWY
48    KYYYGCMCKYSYKKTTACKWKGRYKGKYRRYWTMGTTGAAKKGAAARGWYAKATYKRKTY
      ****     *****     ** **  ******   ..   **      ** *  ***  *

42    YYMKAYYRSMWKYKKGGWSKYRRKRYKKKYRKKSRYYYRYRKATCAYGRMWRYWYRYKKR
48    YYMKMTCRSMAKYKKRKACKYRRKRYKKGTRKKCGTYCACRKRWMWCSGMWRYWYRCGKA
      ****   *** ****   .*********  ***.  *   **     . *******  *

42    WKRKSYWYRYRRRYTKCGYMYYTCYCKMRKKSKAMCACCYRWMTAKYKKKSYRTTKRSRY
48    WKGKSYWYRYRRATKKAATAYTYSYSTCRGKSKCMYMYTYRWMKGKCKKGSYAYYGRSRT
      ** *********  .*..  *  .*.. * ***.*    ****..* ** **    ***

42    YTYGYMKWRRYYYKYKYMYYRRRKWKYGKKWRKMAYARYKYKWTWMTTGRARGWMTKMRR
48    CWYKCMGAGRYYCKCKYMCTAAAKAKYAKKWRKMGCRAYKCKWCTCAYAARGAWCYKMAA
        *  *   *** * ***     * **.******.   **  **  : .   .*  **

42    RYWGRKYCTRWRRRYKKWKMKRTTAGKCRKYTCCRRRCMMRRKGGRAMGYYYRYRKYMYK
48    GYWAGKCMKRWRGATKKWKMKRCKTAKTRKYYTTGAAYMAARGRRARMRYYYRYRKYMYK
       **. *  .***   ******* .:.* ***       *  *     * ***********

42    WTGWYRYYKTGYTRCAMRTKCKYWYKSGMRRAARYKRRKWKCTRYKKYACMTGGCTWKRR
48    AYRWYRYTTYRYYRYWCGKKYKYWYTSRARRRRGYKRRTTKYYRYKKYRYMKRTTCWKRR
         **** .  * *    .* ****.*  **   ****. *  *****  *.    ****

42    AKYRRCCCRWRGTCYKTYAMKAMYKRWMRAYYYTYCTGTCYYMTAYATGGRKSKKGMWKY
48    CKCAAYMYRWRRATCGYCTAKMMYTAWCAGYTCYCYYKYMYTMKGYWGRAGKSGKKAWKY
      .*      *** :     : * **. *  .*         * *..*   . ** *  ***

42    WKSYKGCTKRACKGATWKMKYYWYWSKSRMYYMYYRMYKRSTWYYKRAGACCYKYRRRRW
48    WTGCKAMAKARYKRGCWKMKYYWYASKGAMYYMYYRMCTRCAWYYKRMRRYYTTYRGAGA
      *.. *. :*   * . ******** **. ******** .*.:*****      .**

42    KKYAWKRGYWTCCCCCGYSKRKSYKRRSKRKSKKMYYRGSKAKKKKWSYSKYRSRYYRKR
48    KKYMAKGRYWCTAATTAYGKRKSYKRRCKRKGKKATTAKSKMKKKKWSYSKCAGAYYGKR
      ***  *  **  ..  .*.********.***.**     ** *********  . ** **

42    RCCCCMWRYYRKSMKYYRYKYRSMAKGATGTAMMMRKMMWRYKYRYWRYKKGRGWAYYTW
48    RYYMMAWRTYAKSMKYYRYKYRCARKKCYRWWMMMRTMCWRTTYRYWRTGKRRRWMTYWA
      *     ** * ***********.  * .    ****.* ** .*****  * * *  *
```

```
42      AAACSRRWTRKRWWWKMYTTMWWKCRRCAYMATMKKYSMCTYTRKTKRGAGARRKRYKTT
48      GTMYSRRAYRKRWWWKAYYKMWWKTRRTTTCWYMKKTSMTGYCAKKKACGARRRKATKCY
        .:  ***  ******* * .**** ** :    *** **  *  *.*  .. ***  *

42      MGRYYRKYSYATTKAGGYMARYRGGWYGACAMYYYRWYKRMAARKMYRYRCSGYYGGKWA
48      ARRYTGKTCCRKYKCAAYMRRYRTAWTRMYRMCYYRWYKRMWGRKCTRYGSSRYYAKGWG
          **  * .  . *...** *** .*     * ******** .**  ** .* **.  *.

42      KATCTCTYYKRYSRWAKMKSRYMKWYYKYRTCAYRKRRYWRKYYWKYMRARKRMKWKKKK
48      KTGAKYKYYKRTSRWWKMGGGCAKWYYKYRYTGYRKARYWRKYTTGYMRRRKGAKTTKKG
        *: .. .**** *** ** .   *******  .*** ******   *** **  * .**

42      WRYYYMKWKCTKAMSTKKRAGTATKRYWWRKKCTCAASYRMKKKWRRKRKARKKYKYRYM
48      TRYYCMKAKYATWMSYKKARKYWKKRCWWRKKGGTMRSYGMKKTTAAKRKWGKKTTYRYM
         *** ** * :. ** **     .** *****     ** ***.   ***  ** .****

42      KMATGYMWGWKKGMKRKTRYYRTKYKRTCTWSKGGCARRKMWKRKYCCTRRRWSCYKKCR
48      KMGATTCWKWKKKMKRKYATTRYKYKRCYCWSKKAMMGGGATKGKTTTCRRRWSYTTKSG
        **.:   * *** ****    * ****   *** .       * *    *****  .*.

42      KGKKKYRACWRWYRMYGATYGMAKWAAKSCGARYKYRACSRARATYGKYRTRAAMYRWYW
48      KKKKKYRRYWRWYRMTAGCCTMTKATTTGMAWGCGYATASRRRGGYKGTRCRTTATAWYA
        * *****  ****** ..   *:* ::.. .    * :.** *. *   * *::   **

42      YYSRGATTTYMKKGCKYRKYYKRCCGCTKYWYSYAKKKTYKKMKSWRWYRMYKSRGKYRM
48      YYSRKGCWWYMTKRYTYRKYYKGGTTTAKYWYSYRKKGKYKKAKSWRWYRMYKSRRKYRM
        **** .   **.*  .******     :****** ** .*** ************ ****

42      YACGCRWKRWCCAYRWSYWKYYTKKKMYGRYWWCRGRKYMRKMYRYCTRRTTKARKCRCM
48      TGTAAAWKRAYYRYRWSYWGYCCKKKMYKGTATMRRGKTMRKCCGCAYRRKYKWRGTGTM
         . .. ***    ****** *  *****      *  * ***    . **. * *    *

42      YRCWKKRRYKWKRMRKRKYKAGTCAMWMGKWKYYRWYRKYWRYMRMMRRKWGGYKRKKCM
48      YRYWKKGRYKWKRMRKRKTTGAAAGATCKKWKTYRWYRKYWRTMRCCGGKAKKTKRKGYM
        ** *** *********** ...:..    *** ********* **    *    ***  *

42      RMKKYYWWRRWARGATGYMRRKRRMKKKYYRKTKMRYWMKRAYRMRGYRYAKYRKTYRKC
48      RMKTYYAWARWRGRCAATCGAKRRCKKKYYRKYKARYWMKARYGMRKTRYRKYRKKTRKT
        ***.** * **   .:.    *** ******* * *****  * **  ** ****. **

42      RWKYKYRRRYWWYAAAMRYYYKYKCATMKAYKARRYYCTGKYYRGAKRRKGMMYKRRTYY
48      GWKYKYRRACAWYRMMMRTCCGYKYRCMKCCKGRRYCACAKYYRRWKRRKAMMYKGGYYY
         *******   **   **    **   **. *.*** . .****  ****.****   **

42      WRTGGATRGAYCCCCKYKWYMKYWRGCCRYYGRRYKRRWKYRYYMTARAYRRKWRKCCYR
48      WRCKARGRAWYTTMTKYKWYMKYAGKTARYYKRRYKRRWKYRYYMGCRGYRRKWGGYYTG
        **  .  *. *    ********    .*** ************* .*.*****

42      WKTWRKTATWRTGMKCGRWGGYKRMGYKKRSYRKKWMRKTMKCYGWYARWKARMWKKKRT
48      AKGTAKWRKTGGKAKYRGATKYKRMKYKKRGTRKKWMRKGCGYYRAYRRAKGAMWKKKGC
         *   *  .     *      **** ****. *******    *  * * *. *****

42      RRKMRWWTKYCRTCTKCGGYWYRYMYWYWMKMYMRTKKRWRSKGGRRWKKTGCCRTTARR
```

```
48          RRKCGTTCTTYGAGGKTACYWCGCAYWYWMKMCARATKRWRCKKRAGTKKCCAMRKKMAR
            ***     .   :  * . **    *******  *:.****.*     **  . *..  *

42          RKKMRGTKWRSMTKKKTMTRCYRMRKRYAMKMSYRKKYRMSRYKRWMRKAYSWRMAWCKC
48          GKKARKYKTRSMYKKGKMYGYYRAGGAYTMKMSYRKKYRMSAYKRWCRKMTGTGMTWYGY
             ** *  * *** ** .*   **    *:************ **** **  .  *:*

42          TYKYYYYKRYKCCATCTGACKRYKRYYYYRTATKMRMRSRRYKRMRKKRWMYKKRKYKYY
48          YCKYTYYKRYTYAMWYYKRMKRYKRYYYTGCCCKMRARSRRCKRMRKKGAMYKKGKTKYT
              ** *****. .       ********   . *** **** ******  **** * **

42          KMYYRMKMKKKKKYKGTWCMKKTAACGCYRYMWTAKAYRKYCRYRWYTGSMAKSRCCTGW
48          TMYYRAKAGKKKKCKRKWYCKKWGGYRTCAYCAYRGRYRKYMRYRWYKRSAMKSRTYCRT
            .**** *  **** * .*  ** ..     *      **** *****. *  ***

42          TYTMYYMMYMRWMWTMMAACWRGGAKYRAYRRYTAKMRRYGGGTGGGYRYRKGCSKTWKY
48          YCWMYYMMTARWMWCCMRRYWRAAGKCARYRRYCTKMRRCARTCAASTRYRKATGTYAKY
               *****  ****  *   **...*   **** :**** .   ... ****. ..  **

42          WWYYTYKKGCKCCCWGKKTRTYKGKYYMYRWKWMYKKYWYWKAKGWKKMYKRSWYMRYRK
48          WWYYWYKGKTKYMYWKKKYRKYKKKYYCYRAKWCCTKYWYWKWGAWGKMCKRSWCMRYRK
            **** **   *   * ** *.** *** ** **  .******  .* ** **** *****

42          MRYTYRYRWWMYKYKRYRAWAKRSMTKRYTTTWTTCKRMRKMYW
48          CGCGTAYRTAATKYKRYRRARKRCCCKRYCCKTCGYKRMRKMYA
                  **    ******   **.  ***  .    *******
```

The pairwise alignments of two accessions 42 and 49

```
42              KYKSYMRYKMYRWKYKRYRYYKRTSMRYYYRRRCMCGGRMYYYYRMYYRMKWRWTAGRMK
49              TYKSYMRYGACRWKCKAYRTCKAYSCRYYYRRAMMTKARMYYYYRMYCAAKWRWKMRACG
                .*******   *** * **  *  * ******  *  .*********   ****.

42              YRYKWKKGRRYMMWRWRWYYRRYMSTYYRGKKYRYKMKKYYWKWWASMKYKYKRGCCKMK
49              TRYKWKTRGATMMWRWRACYRRYMGATTGRKKCGYTATKYYWKAWCSMKYKYKRRSYKMK
                 *****.    ******  *****.:    **  *. .***** *.******** . ***

42              ARYKRSWTRWYMCGCYTYYRMTMYWKRYKCRWWGRYYYGRRYRKWKYYYKCYRMRMKWWM
49              RGCKAGTKGWYMYRYCYTCRMKCYWKRYKYGWWKRYYYKRRYAKWKYYYKTYRMGAGWWM
                   * . . ***       **. ******  ** **** *** ******* ***   ***

42              YWRYTGAMRMYRYYMGMKRYGGRKMYRRYASYGTTGYYWKCGAMWMYYRTRKMYMYYRYR
49              YTRCCAGMAMYRYYMKMKRCRRGKMYGRYWSYRACKCYTKAKRMWMYYRWRKATMYYRYR
                * *  ..* ****** ***    *** ** ** :   * *.  ****** **  ******

42              RMRWYMWWWMWYMMWRCRYMWYMKWRRKRMRRWKAKMGCRWCSYYWKRWWYRWACYCCAC
49              RMRWYMWWWMAYMMWRYRYAAYMKWRRKRCARWKTKCKMRWYCYCATRWWCATCYYYTGT
                ********** ***** **  ********  ***:*   ** .*  .***   . *  .

42              MMRRRKRYMKWMMACTYYRGYWMWWYKSRRRCYRYKCRYWSGKAGMMMKCACCWGSKYRW
49              MARAGTRYMKAMMMMYCYRKYWMWWYKSGRRYYRYKYRYWSTTGAMMMKYWYYWRSKYRW
                * *  .**** **    ** ******** ** **** **** ...****    * *****

42              TKATKYMRRRTWMRGYYYWRAMYTAARTKRWYRYRGTYYWKSRYCTATGYMMWYWYMYRK
49              WKGAKYCGGRCTAGRCTCARWMYWRGRWKRWYRYRRCYYWKSRYTGCCRCMMWYWYCTGG
                 *.:**   *         * **  .* *******  *******  .   ******

42              RYRMWMAMAAYKWRYGYMGGARATGRATTYRTAAGYRYRKWWKRKKYYKTCCGAMARRRW
49              RYRAWMTCGGTTWRYKYMAATAWCKRMKWYRGGGAYRYRKWWKRGGYTKYYYKMMCRRRW
                *** **: .. .*** **..:    * . ** ...*********  * *     *.****

42              RCGWYRRYGKRYWCYGATAYRRYKRTTATYKWAYWCKMKKRKKKRKRYKKYYMRSRYGKT
49              RYRWYRRYKKRYWTCKTCTYGRYTGACGAYKWMYWMKMKKRKKKRKRCTTTYMRSRYSGC
                *  ***** ****   : :* **. : .:*** ** *********** .. ******.

42              WKYYACAAYRRKKKCGYRKTYYARYKKMMRKKAAARRGCACAYMKMYKYRRWRYKMYMRK
49              TKCYGYRCCGRGTKGRYRKKYYWRYKGACAKKGTGRRKYGACYMKMCKTRRWAYKCCCRK
                 * *.  .  * .*  ***.** ***    **.:.**  ...**** * *** **   **

42              KRRKRTAKKRRRMSMTYKYRRKKKKKKRTAWWYYKWTMYKMCCYASARKYMRYRAMWYAG
49              KRRKRKMKTRRGMGMYYKYRRKKKKKKRCGTACCKWYCYKAMATRGCAKTCGYRMMWTCT
                *****. *.** *.* ************ .    **  **  .  .. *   ** ** .

42              RGTCARYRYMYYWRAYRYKYRYMRWSRWKRRCGKKKKYYRKKYAAGKMRGRYYKKRRGRG
49              GSKYWGCRYMYYAARYRYKYRTCGWSRAKRRTTGKKKYYRKTYMMKKAAKRYYKTRRRGS
                 ..    *****   ******   *** ***   *******.*   *   ****.**  .

42              CKRMKSYRTYWWMYYYKWYKWKKSKGCGRGKRRYYWATCKMYYRTGACYMWRRYMAYRRY
49              YKRMKSTAKCAWMTCCGWYKWKKCKKYAGAKRRYYARATKMYYRCRCTYMWGAYAWYRGY
```

```
         *****  .  **    ******.*  . .*****  : *****  . ***  *  ** *

42      YWRRMRCACCKCAATMYKYYWYKGMYYYYKYGYKSKMYYRSRKYMRKKRKYRWYKWYKKR
49      YWRRCRSGYMKYMWYCCTTYWYGRMYCYTKYRYGCKMYYRSRKCMRKKRKYRTTGACTGA
        **** *..  *      . ***  ** * ** * .******** ********     .

42      ARYRYRKMSMTKGMYRYYYKYCKRYKYYMCACTYYYKGTCKYWACTATKWSGMSRRGKYK
49      TRTGCRKMCCCKACCRYYYKYYKRTGCYAGCYYYYYKAAYKYAMYAGCKWSKMCGGRKYK
        :*   ***.  *.  ****** **   *  .  ****.: **   :. *** *.   ***

42      ACKRCCTKAMKKTGTKKGRCKYMGATYAACMYKRRKRGRWRTRYGCRYRCCAMWYYWACW
49      RYKGYTCKMMGKCRYKKRRYKYMAGCCWWACYKGRKRRRWRCGCSSAYAYYGATTYWGAA
          *    * * *   ** * ***..    . ** *** ***   .. *   .   **..

42      MKRKRRRYWATYCRKKMRYRRMYYYAYMWKMRKRGMYRGWMWCYKSYAKRRKWMCSYYMY
49      MKRGRRGTTGYYYRKGCRTGRMYYTMYMWTCRKRKMYRTWMWACKSYCKRRKWAGSYCMC
        *** **   . * **  *  ****  ***. *** *** ***. ***.*****  ** *

42      RCWCAAWAGRYMRKYKYKMMYMRRKYYYKAGRWYAGRYYCRKCMWRKYMAWMYRTGKTCA
49      RYASRRWTKRYMRKCTCKMMYMAAKYYYKMTRWYGRGCYYAKYMARTCCCWMTGYRKWSC
        *  .  *: ***** . *****  *****  ***.   *  * * *.  .**    * ..

42      AKYRTRKAKYYRYRWRRATRYTGCKRRKYYRWMRCGCYYKWTKKKKKYKGMYYGKKKYRW
49      RKYRKRKCKYYGTRWRGCWRYYSYKRRKCCRAMRTATYYKWCKKKKKYKKMYYRKKKCRW
         ***.**.***  *** . ** . ****  * ** . **** ******* *** *** **

42      RRACYKGRRRGCYRMKKCKATCCTCTRAWMRRYRSRKSRKMTGRKKKTWYAATRWAKRWR
49      RRMMCGAAGRRYCGAKTTKCGTYWMYAMAMGGYRSGKSRTMAAATKKGWTRWAGWRKGWR
        **    .  *     *. *.         *  *** ***.*:. .** *   : * * **

42      MWRMWKRYRRRRKAMYKYYAGAMTYRRCAWATAATKYYYRYKRWYRRRKRKKKWKKKMRM
49      MTRCWTRYRAAGTRMYKYYWKCCCCRRSRWGCGGKKYYYGYKRWYAAAKAKKKWKKKMRA
        * * *.***   . *****  .   **. *. ...**** *****   * *********

42      RRKRRRKRCGGCYTGGKRCRYRYRAYRTCTYMGKYYKYWGAYAGYRYGTKGKYYWTMMMM
49      GAGGGRKRMKRYYKRRTAMRYRYRMYRWYWYMKKYYKYWTTYGAYGYRYKRKYYWYMMMM
             ***    *.  .  ***** **   ** ****** :*..* *  * **** ****

42      CTGKGCCRYSAAATGTGCMRRYRKRTCAGKKWACCAAYGCTCGYRRRYCCAGWYKCMTTC
49      GAAKRATRYSWMMYTCRMMRRYRKRWTWAKTTWMYRWTRYKYRCAARYYYRRATTAMGGT
         :.* . ***        *******   .*.         .     **      ..*

42      KRGMKMRACGGGTCTCAAYRMGWWRKGYRWAMRTKKKYTKKTYKKKKACRACCCWRWKRG
49      KRKAGMAWMRRRWTYYRGTGMKWWRKATGATAAWKKGTCKKKYKKKGCTARYTTWAWKGR
        **   *           .  * ****.   :   **   **.**** .      * **

42      KMRYMSKSRYMYYMYYYMYKRKSACWTTYRKKYKKRTRTGYARMTWCCGYSYRRSYKRRK
49      KMRYCSKGAYMYYMYYCMCTRKCTMWYYYRKKYKKGYRYAYGGAGTTTAYGTGASYKRRK
        **** **. ******* * .**.: *  *******  * .*.      .*.   ******

42      KSKKSGKKWRGKWSKYRKRRRMWYRSMYYRYKMMTTYYTTAYWRRRSYMYYYGYMKSRAW
49      KGKKSKKKWRKKWSKYGKRRGAWTASMYYRYKACCGYYCWCYWGGRCCCTTYKYMKSRWW
        *.*** **** ***** ***  *  *******    **  .**  *.    * ***** *
```

```
42          WRWRMSMGKMWKKMKRWMRKKSKKTMKMKCCCYMRCGKWYKTTTTGCWRKYWKYYYWYYM
49          WRWRMSMKKMWKKMKRWMRKGSKKKMKAKGTTYMRAKKWCKKKCATTWRKYAKYCYTYYM
            ******* ************ ***.** *   ***. ** *.. :  **** ** * ***

42          MYARMRYWYKCKRRRRKTTGGGKKMWKYRCYKKKKRGTRRRKMGYWCRAYCSKGRWWYKY
49          MYRAMRYAYKTGGRRRKYYKTKKKMTKYRYYKGKKRKCAAGGMRYWYRCYTSKRATAYGC
            **  *** **   ****     *** *** ** ***      * ** *.* **    *

42          KYRATRKWRRARTTRSRWAGMYKRYMGMRYSKYTYRRRCTYGGCGKGCWTWKRRYWKGKR
49          KYRTYAKARRMAWYGCGWMCMYKRYMAAGYSGYCYRRRMYYATATKATAGWKRGCAKRKG
            ***:  * **     . *  ******.  ** * ****  *. . *.   ***   * *

42          GCSKKKTAATKKGGTMKRWMACGKRMSRRKRRMYKWRYKRKKKMYYAKKMRRKMMKMWMM
49          CMCKKKYRWYTKAACMKRWMRTRTRACGGTRAAYKWRYKRGKKMYYGKKCRRGMMKATCM
              .***    .*.. *****   .* .  .*  ******* *****.** ** ***   *

42          AYMRRSWKSMTRKWGTCKKKRGRYWRYAKCGCCAYKWAMRRMCSYWSYWRKMKGCCWWST
49          MYARGSAKCCAGTWKWSKGKRAATARYCKMTTGMYKACCRRMMSYWCTWRGMKAGTWWSK
             * * * *. : .*  .* **.   **.*     ** . *** ***. ** **.  ***.

42          YTTKATGTMWKCTATTKMMKAYKYWRKYMTGAYWKYRKYMMGCYSMRKMYRKMMYSWYCK
49          CYAKTCTCMWKACGWCTMMKRCKYTRGTAYSWCTKCAKYMMRMYCCRGAYRKMMYSATYK
              :*:   ***. .  .***  ** *    .   *  ****  *. *  *******   *

42          MYYYGCWMKKRKKYRKRYWYYMYGCKKGGGKRAGCCYMKWMWSKRRTKKKMRKMKYYYRG
49          MYYYAAACKKRKKTGKRYWYTACTTKKKKKKACASYYMKTCASTRAKKKKMRKMKYYYRT
            ****..  *****  *****     **   * ... ***   *.* .************

42          TAYRKWWMMYKKKRRWRMRRWMKKRWMMMRKSRSRMYRCYGCAGMKARMRYMYKYGYRYR
49          GCCGKWWMMYKKKRRWRMRRWMKKAACMMAGSRSGMTRYYRATTMKCRCRYATGTRCRYG
             .  ********************   **  *** * * * .: **.* **      **

42          YRYRRACKRWYYKTTAKKKTYYTRKYYKMGYYYYYWYAWMYMKKCMRRGYWYRYWMKMKK
49          CRCGRTAKGWYYKYWRKTKYCYKRKYYGCRYYYTYWYRWMTMKKYMRRKTWYRCWMKAKK
             *  *:.* ****   *.*  *.****   *** *** ** *** ***  *** *** **

42          MGCKWWAARMTKYMKTGYKKWRYWCGCAKRYAAYYKTARRRWWAGCKKYYKKKTYRRTWK
49          MKYKWWWWRCGKTCKYCYKKTRYWTAYMKRYRMYCKWMARRWAWRYKKYYKKTKCAAAWK
            *  ***  *  *  *  *** *** .  ***  * *   ***    ******..   :**

42          WKRRYKRKRKKMSTYKYRYARYYGRGCYYWYYSRKMRKYKYYSRSGYTKYYKYAARKKSR
49          WKRRYKRKRKKCSACKTRYMATCRGKYTYAYCSRTAAKYKYYSGSRYYKYYKTTWRKKSR
            *********** *: * **         * * **.  ****** * * **** : *****

42          WAKKMTKSRKKKWKCKKYKRYYKKKRARYWRYKYGYWKYMRTRKWYMKKYKWKRYWKTMS
49          WWKKMKGGGKKGWKYKKYKRCCKKKARRYWRYKCATTKYMRWRKATATKYKTKAYAKKMS
            * ***. . ** ** *****  ***  ****** .  **** **   .*** * * *.**

42          RKKCGTRRYWWRWKAKKKCTYKYRKRKKKYYMYMKKAKGKWATYMKAAAKYKKYMKYRKY
49          RKKTAGRRCWWRWKRKKTGGYGYGKRKTKYTCYMKKRKKKWWYTAKRWRKYTKYMKYRKY
            *** . ** ***** **.  * * ***.**  **** * **    *   **.********
```

```
42    YKRCKYKYKKKKKCKTKGGGYCMSWRWYCGWYTKYRKYMYKRWTGYGAKRSWWCTGAAYR
49    YKGYKYKTKKKKGAKKGRKRTMASWRWYAKWYKKYRKYMTKRWCATATKRSTAYKKMRYR
      **  *** **** .*.       *****. **.****** *** . .:***   .   **

42    WSYWWWRYKYYYKYMYWKRKYKYKYRYKYKWRCGGRRYYTAMTKRYMWKKTRGKKYKRMR
49    WSYWWWRYKCTTKYMTAKRKYKYKCGCKYKTGTKKRRCTCRMCKRYMAKKCGRKKYKRMR
      *********   ***  *******   ***     **    * **** **   *******

42    YAYTCAKMWYYRRRMKKRYGCTMKMYYKRRYYCRRRWWKYRYGKRGRTWKRKKCRTTTWC
49    YMYWMRKMWTYRGGMKKAYSGAMKAYYKRRYYSRRRWTTYRYKKAARKWKRKTMGKKCWA
      * *   *** **  *** *. :** *******.**** .*** * .*.****.  .. *.

42    KKYWMCRARYGRCKYRRAYWYRMRSYKRKKWCKYKCYYAKRRMWGKKKGKYYWWKYYYYR
49    KKYWAYAMGYRRYKCGAGYWYRAAGTGRKKWYKYGYYYRKRAMWTKKKRKYYWWKTYYCG
      ****     * * *   .****  .  **** **  ** ** ** *** ****** **

42    YMKRRMMKMYYRRRRTKGAKWGKRRWYRWYWRKKMAYRYMWASWAGYGKMRGWYYKWTCR
49    CAKRRMMKMCYRRRRATRCKWSKAGTYRWYWATKMWCGYCWMSWWKTGKARAWTYKACMR
        ******* *****:. .**.*   ***** .**   * * **      *.* **   *

42    YRRWGRWWWWKMTCAARRRYMRRRRYKTKGCTARKRRMRYYRTTRYKYRMYRGGWYYYKM
49    YRGARRWWWWKAYMMWARRYAARRRYKAKAGGTRKARMRYTGKYRYKYGMYRTAWYYYKM
      **   ******      ***  *****:*.  :** ****  . **** *** .******

42    RGCYKYYRWKTGTTTMARKGMYYGCGRYYKYWWKYTMWKWMRGRRYRKWCAGYMWYRKKK
49    RTAYKYYRWKCAWYKMMRKKCTYAGTRYYKYWAKCCMWKTCGKRRYRKWYRRYAATGTKK
      * .*******  . .* **   *.  ****** *  ***    ******   *    .**

42    YKRRTTCCCCKKGCMMWMYYKTRGWYYYMYYRSMWKYKGYKKYCYRRKKYYRMRYRKATA
49    CKGAYYMYSYKKRYAMWATYKCRKTYYYMTYRSMWKYKKYKGTTYRRKKYYRCACATRWW
       *      . **   **  ** *  **** ******** **   ********    .

42    GYRMRYKYKKRYSYWYWKRWRRKWKKRRYYKKKRRAYWKMYGKKYKCRMRKRYKKWKYKA
49    KCGCRYKCGGAYSYWYWKRWAATAGGRRCCKKGRRRTTKMCRKKYKYRMGGGCKKAKYKT
          ***    *********  .   **  ** **   **  **** **    ** ***:

42    WKWKRWYSWYRKAMCYKKYMMKTYKRKWWCKTRTTCRGWKYRKMYARYYYRWYYYKYYSK
49    TKAKRWYSWYRKRATYKKYCAKGCKRKWTYKGRACARATKCRKAYWRYTTGWYYYGCYSK
       * *********   ****  *  ****  * *: .*. * ** * **   ****  ***

42    GKKYYYRWRRTKKKYKYWTRRKGGKYYYKAYYTKRSMTKMTYRYRRMRKRRYAAKMKKYK
49    CKKTYYRWGGCTKKYKYWKARKRKKYCCKMCCGTRSMYKMYCGYRRAGGARYCGKMKKYK
       ** ****   .******. **  **  *    .*** **   ***    **..******

42    MYTTCWCKGMMRCKCTRYRRRYRRRTGWYKATTGKAKRKYYYYRTATYRRKMYKWTAMRK
49    MYCKTWYKKMMRYGYCAYAGGYGRRCAWYTGCCTKWKRKCCCTGYCCYRRKMCKAGWMRK
      ** . * * ***      *  * ** .**..   * ***      . ***** *   ***

42    RWMYKRKYRYYKMKKKYRYRYRWKYKRKYAATTTARWWRKYRGKRATYYKRKRWWWWWMW
49    RWMYKGKTAYTTMKKKTRCATAWTYKRKYTGCWWWRWAAKYRTKRTYCYKRKRWWWWWMW
      ***** *  * .**** *    *.*****:.    **  *** **:  ************

42    WWKKYKKKRSWYRYKGYRRRWGYSYRTSKGYRSKYYGMWSRCTGRATKMYYRKRKRWRWY
```

```
49            AWKKYTKTAGTYRYKKYRRGWKTSTGGSKKTACKYCKMWSGTWAACAKMYCAKRKRWRWY
               ****.*. . **** *** *  *   **   .**  ***   . .:***  ********

42            YYCYKKRYRMMYTMMKKMKYKYRMRYYRRACYCYCARMMTRYYYKMYRMYWYGGCC
49            CTMTKKRYRMMYCMMKKMKYKYRMRYYRRWMYATSRRMMYGYYYKMYRMTWYAKYT
                  ******** ****************  *. . ***  ******** **.
```

The pairwise alignments of two accessions 42 and 50

```
42                  YRMRMYYRRYCATRRTTYMMWRTWRKKRMMKRGAKRWRYMYYMCRYAYYYKKMYYYYYMY
50                  YAMACYYRRYYWARGWCYMMWRGWRGKRMMKRRRKRWRYMYYMMRYGYYYKKMYYYYYMY
                    * *  *****  :*   ***** ** ******  ********* **.*************

42                  YRMMGCTYKKWRKMMYMRYRRCKRMGMSYCRYKSYWYRRYWYYYWYWYRRRMWKMTAYKK
50                  CAAARTWYKKWRKMMYMRCRRMKRMRMGTTRYKSYWYRRYTTCCWYWYRRRAAKCYWTKK
                           *********** ** *** *.  **********    *******  *    **

42                  WRASYYMWWGKKWYRRCGYRSWKKYRWMWKRRYRMKTAWMMRWWRRMRRRMKRYWRRRYM
50                  TRWGYYMWWRKKWYGRATCRGTKKYAWMWKRRYRMKWRWMARWWGRMRRGAGRYWRRATA
                     * .***** **** *.  *. *** **********  ** *** ****   *****

42                  YCCTRKWTYWKGRYYSMKYYYRYWRWRSKRCTYAMYWWWRWSYRKMMAWMYRRMRMYYRS
50                  TYYKRKTKYWKARYYCMKTYCACWRAACKRTATCAYWWWRWGCAKMMWWMYRRMRMYTRS
                       .** .***.***.** *   **  .** : . ******.  *** ********* **

42                  RRYYKSRACRMTKYYMRYKKWRCRATARRYWSWRYRKYCYYYWYYRTWTTYSRYSWRKRM
50                  RRCCKSRRYGCATYYMRYKKWRMRTCGRRYTCWRYRKCAYYTWYYRCWAAYSRYSWAGRM
                    **  ***    :.********* *: .*** .***** .** **** *::******  **

42                  YKMKGMYMMKRRRRWYKMMKKSKRKYYMGYKMRRWKRRKRYACYWKRRKRKKMYKYKMKR
50                  YKMKRMCMMKGGRRWYTAATKSTRKYCMAYKCARWKAGTRYCMCWGRRGRKKMYKYKMKR
                    **** * ***  ****.  .**.*** *.**  ***  .**.  * ** ***********

42                  RKKYGMKRRAKRKMMGYKKRKRKRAYCRAAKYMKRRAWYWRRKWTGAAYWYRMYWWMKYY
50                  RKKYKCKRRCKRKMMACKKRKRKRGCMGRCKYMKRRMTYARRGTAATTYWYRATTACKCT
                    ****  ***.*****. *******.    .******  * **  :.::****     *

42                  MYMMKYARYRRKKKWRRKTTCYRRWYKYYYRWYYMAYYKKKYTWWYAAGRKKTRYYYYKC
50                  AYCAKYWRYRRKKKWRRKKAATRRWYKYTTGWYYMCCYKKKTAWTCGRKRKTKRYYTYKY
                     *  ** ***********.:. ******   ****. **** :*  .  **..*** **

42                  KKKRRRYYYKRCMKRRRKRRYYKKRGGRTYRGYRTATAGWRTCKTTRYSRGRRARRYTAA
50                  KKKRAATTYKAYMGGGAGGGYYKKGRRACYRRYRATATAWGCTKYCGYSRRRRRGRYGTG
                    ****    **  *       ****     ** **::::.*   *   *** **  ** :.

42                  RKWMRRYKKCWSKYGYWWYYYTRGRWYRRAYYYCYTGRWWKWYRRYGACTAYYCRYKRKK
50                  RTTMRRYKKYWSKCKYWATTTYRKAWTGAGCTCYYWTRWWKWYRRYRRYYGYCYRYKGKK
                    *. ****** ***  **     *  *   .    *  *********    .*  *** **

42                  KCCACGYYKYWWKRYSWWKWRATAAYKRKKYKRSYMMRYRSYWYKYRKYTMMMRKWKACA
50                  GTAGMKTYKCWTTRYSWWGTRCATGCKGTTTTAGYMCRYRSYTCTYRKYCMMCRKWKGAC
                      ..   ** * .*****  *.::. * .. . .** *****  .**** ** ****...

42                  YMKWKKKRMKRWRYMKMYRKRRYAKATTYKYWWYGYRKRKMCYKKKKKWYKKKYKKMRKA
50                  YMKWKKTGCKRWRYCKCCRKRRYRTMCACKYWWYTCAGRKMTYKKKKGACKKKYKKCGKC
                    ******.  ***** *  ***** .  : *****    *** *****   ******  *.

42                  SYMYYKARKRWWRWSMYKCATKMWYYYRYKRRWRKYMRYKGKYKRKWKRRMRKYKWYCGG
50                  CCCTTKWRKRWWAASMYKTCGKMWYYYRYKRRWRKTAGYKKKCKRKWKAGMRKYTWYTCA
```

```
            .    * *****  **** . **************   ** * *****  ****.**  .

42          YACMKYMMCKKKMYCKTCSKKWRMKWAWMWRRKAWRGMMRTWGKYRYKKCCYYYATYYRG
50          CRACKYMMYKKKMYYKWYSKKWAMGWGACARRKWAGRAMGWTKKYRYGKYMCYYTCYYAK
              . **** ***** *  **** * *.   ***     *    **** *   **: **

42          WWCKYRCGWKYRTTAGCCTRRKRKMRRYGYYMKCTGRGYRKWRRSMTKMGWAMRCYRRMR
50          WWYTYRTRWKYRKKMTTTARRKGKMRRYRCCMKYGRAKYRKWARSMYKMKWMMRYYRRAG
            ** .**  ****..    :*** *****   **     **** *** ** * ** ***

42          KRYRKKKKYRYKRTGMCKGCTKRKRRYYKRKAGTATKAYYYRRKTYKKMKRYTKWRWMYK
50          GAYRKKKKYRYKRCAMYGAYCKGKRRYYKRTCTGTWGTYYYRRKAYKKAKRYYKWRWMYK
              *********** .*  .  * *******..  :  :******:*** *** *******

42          GWRKYRYYKMYYRKMKKRKKYKKMRYCTACYCTYYSKCAGGYKMCGARCRGCCKKTGMYR
50          ATGKTAYTTMYYRGAKKRKKCKGCRYMYRYYACCTSKSWKRYTCTKWRYRRTTKKYKMYR
            .  *  * .****  ***** *  **    *.   **.   *.    * *   **  ***

42          RTTYWTYKKKSYRRRKGATRRMTYYGGYAWAACGCTYKMKYYRMRRKGKYKCAWCMKKWR
50          RYYYWKCKKKSYRRRKKRYGGMKYYKRCMAWGTATKCKMKCYAMAAKKKYKYGWYMKKWR
            *  **. *********     *.**      . . . *** * *  * *** .* *****

42          KWKKWKCWMTTCMYWKYYRWCRWRRRKYMKYMTAATYKKYYAKKKKKKSWRWYRMWKYWT
50          TWKKWKYWMWCTMYWKYYGWYRWRGRGYMTYMCCGYYKKYCRKKKGKKSWRWYAMTKCWY
            .***** **   ****** * *** * **.** .. ****  *** ******* * * *

42          RYRMYKSYKSCCYRRWTGARSYWKGYWRKYMYMRYWRRKMCCYKKRYKRWRRRWYYTKKR
50          GYRMCKSYKSMMCGAAKKMRSYWGKYWRKTMYMRTARGKMYTTKKGCKRWRRRWCTCKKR
             *** *****      .  ****  **** ****  * **   **  *******   ***

42          RKMTRRYKYYYWMMRYKRYYGMMYWYWYYRWYRMKKYRKMRYAKARRSRRYKRKKACTRR
50          RKMKGRYKTCTWAMRYKRTCKMMYWYWTYRWYRMKKYAKMRYGKGGGCGGYKGKKGTCRR
            ***. ***   * *****   ****** ********* ****.*.  .  ** **.  **

42          RWSRKKTAACTKTWMGMYTRCRWGCKACWRYYYMRMYYRAMRRKRRYRCGCRRKCCTMYY
50          RWSGKKKRTTAKKWMKCCGGARWRYGWYWRYCTMRMYYAGAGRGRRYRYRMRRKSYYCYY
            *** **. : :*.**     .**     ***  ***** .  * ****   ***.   **

42          SRGTCMRAGMYKKYRYMYWRYKCKRKMYRYRKMWRKKWYRYMYRWWMYRKKWKYRYRAKW
50          SRACTCRTACCTKTRYMYWRTKAGRKMCGYRKAWRKKWYRYMYRTAATRGKTKTRYRGKA
            **.   *:.  .* ****** *. ***  *** ***********    * * * ***.*

42          WRSMGKRYMWRWRCAWAYKYMRMGKMWKYAARYYYMMKWKKGRWMYRRWRWGKRRSTKRK
50          TRCCTKRYATATRMTTTTGCMRMRKMWTTRMRTTYAMKAKGRGWMYGRWRWKKRRSYKRK
             *.  ***    * : :   *** ***.   *  * ** *   *** **** **** ***

42          YYMYYRKRRKKYRMRYYRKKYRRYKMYKYRYMACASKMRRMGYKMYCRCKKYRTYMYKSY
50          YYMYYRKRAKKTGMRYTGKKYRRYKMYKYRYAWMRSKCGRMAYKCTTGTTKTGKYMYKSY
            ******** **  ***  *************    **  **.**     .*  .******

42          CYWTATCCTAYYKYGCTTRSMKSRYMKWYYKYMRYGYKKKYKRRYWRYCYYYWTAGYMRK
50          TYWYTGAYKMYYKTRYYCRGMGGGCAGWYYKYMRCKCGKKYKARYAGYMCTYTCTAYMRK
             ** : . . ***     *.* .    *******    **** **  *   *  :.****
```

```
42             KYYKRYYKTWRYKGWKAAAMSRKKTWKKCTYKRYWWYYKYKGSCRMKKKKWRRRKKKGKT
50             KYYKRTYKGTRYKRAKTWWMSRGKYWKKGGYKRYWWYYKYKRSMGMKKKTTAARKKTKTY
               ***** **  ***  *:  *** * ***  *********** *  ****.   ***. .

42             TYRYRYMWYRGGMWWWKYMMTAKKRAAWYGAMRWMWAWYMKTGCGKRRWYTYKRMYAMRT
50             KCAYRYMATAKKCWAWKTACACKKRTTWYRGCRWMWCWYMKYRYKKRRWYGTGRMYGMRA
               .  ****      * **   :.***::** . ****.****    *****   ***.**:

42             YYYYMTRKYGRAMAAWRMYRCRWMRYKAATAAGGRKTGKRTKRYYYWKKTCTYRKWAYMT
50             YYYTAYRKTRRTMRRWRMCATRWMRYKRMGGGASGKCAKGKGAYYCWKKATCYRKACCCY
               ***   **  *:*  ***   ******   .... * .* .  ** ***:  *** .

42             TRRMCSYRSYMKATGCWWSYYYWRYCGAAKCCMCMYYTYYAGRYWRTGTKYMSACSMRYK
50             KRRMMSYRCTMKRKATWWSCTYWRTYRRMKTMCYAYCWYTCARYWRYAYKYMSMSCCRYK
               .*** ***. ** .. ***  ***     *     *  * ..**** . **** .. ***

42             TACARYRKMMWYWWMYYYKCAKKWCCTKRYWMATGKWKRTTTYMKWMWGKKKKKRRKGAC
50             YRMRRYRKMMATWWMYYYKYGKGTYYYKGTAMGCRKWKRWWYYMGTCATGKKKKRGKTGT
                   ******  ******* .*     *   *.  ****   **      ***** * .

42             WMRYYKWKKKKMAYYWGKWTMACGKKRMRYRWRKRRRYRGGCYRMGTWGGRKKRWMRRAT
50             AMRYYKWGKKKMWYYTRKTKCRTAKKRMRCRWRKAGRCGKRYCRMKGTCKRKKAAMGGWK
                ****** **** **  * .   .***** ****  *      **     ***  *   .

42             TSKRKKRKMRKKKYTCYGGGARGRWRKMWKRYRRRKMYYMTYRGTRYWWYYMKCTWKRKY
50             WSKRTGATMRKKKYCTTTRKGRRGWRKCWKRYRAAKMYYCWYRRYGYWWYYMTMATKGKY
                ***.  .******      .*  *** *****  ****  **   ******. : * **

42             AWWTYWKRRYYWCGKRRYWYRYYWRKRTCTKRYRWKKTTTRKMMKCGRWGYRRAYWRYSY
50             GAACYAKRRYYWSKKRRYWYRYCWRKRWYGKRTGAKKYAGGKCAKYRGWKYARGYWRYGT
               .   * ******. ******** ****   **   ** :  *  *   * * *.****.

42             RTCACRKWGRRKYKYGKTKKCYMKWKKRRKWWKYRTKYWYRYMWYYRRRMTKAKCGSKRR
50             RKYWMRKWKARTYKCRKYKKYYMKWKKRRKAAGTRGKYWCGCAWCYRRRMGKCKYRCKAG
               *.   ***  *.**  * ** *********    * ***    * ***** *.*  .*

42             WKKKTGCYRYRYMYWYYYCRWTWGAYAKRGCRYGRYYYRGCAGAACMKYYWCSTRARRYM
50             TKKKCCAYGTRYMYATCCMGWYWKMTWKRSYGCKRCCTRKYMRRMTMKYYAYCGARGATC
                ***  .*  ****      * *    **.    *   *       ****  .

42             RWSKKYAGKYYRKTRRYWRGYRYKYRKRRKCGAKYYWRCMYRMKSKYWYYWYMRYKKMYT
50             GWSKKYCTKYYRKYGAYWAKYRYKYRKRRKSRWKYYWRSMCRMKSKCATYWYMRYKKMTC
                *****. *****   **  **********.  *****.* *****   *********

42             MMTCKRRRYYAKMYYRRKKMMKRYRRYGCYYWRGGMYKTCGTKKGMWGAKYKKWGGYKAG
50             MCAGKRRRYYTKMYYRAKKMMKGYRGYKYYYWRRRCYTGGRAKKKAWATGTKKWRCYKWR
               * : ******:***** ***** ** *  ****   *.   :**  *.:  ***  **

42             KTKKKACACGASWCCRRWKRGKMRRYSKKKSRYMYYMYYYKWYYKKYYGYKAYTYRRRRA
50             KWKKGCTRYAGGWTTAAWKGKGMRRYSKKKGAYMYYMYYCKWYYKKYYRTTGTCYRGAGW
               * ** .   ...*    **   ********. ******* ********  ..  **
```

```
42              WKKYKKRGGAACRKYCCGYSYRRRSYKRRKRKKKMYYRACGSMRGRGKWRKWKYKRKRRG
50              AKKYKKRRRWRYGKYTTACGTGARSYKRRKRKKKATTACATSARARKKWRKWKYKGKRRR
                 ******      **  . .   ***********    .. * *.* ******** ***

42              GGYKGRKSMYTGYYRYKYRGTGTYAWRSMCYKRYTMRYSYKYMRYRWKRRWRYWMCKMKK
50              AAYKKAKSMYWAYYRYKYRKYSKTGTGCMYYKRYAMGYSYGYCRYRWKRGTGCAMAKMKK
                ..**  **** .*******  .. .  .* ****:* *** * ******     *.****

42              YKGCRSKKKKMKAWCKMKKCAATKCCTMKTRRCTGKRMYKRTKKKKWMYGYARKRRTMMW
50              YKKYRCKKKKMTRWTKMKGSMMWKYYWAKYRGACAKAAYKRYKKKGWMYKYGRKRGAMMT
                **  *.*****. * *** .   *    * * . .*  *** *** *** *.*** :**

42              YMRRRKWWRWMMMKKWGKKKKCSKRMMRYYYWYRWMYKKRYWKRKWRSKMRRWMRYGYRK
50              YMRRGKWWRAAMCKTWKKGKKTGKRAMGYYYWYRWMYKKRYWKRKWRCKMRRWMAYSTAK
                **** ****  * *.* * ** .** * *******************.****** *.  *

42              RYYYWMMRKRCCRRTTTKAAGCCAGMRMYMYYYYRYRWTGKRCCRWYYYMYKKKYWATSK
50              AYYYAMMGTGMMGAKCYGTGTTATTMRCYMYYYCGCRWATKGTTGWYYYMYTKKYATACK
                 *** ** .     .   :.  .: ** *****   **: *    ******.*** ::.*

42              WMKTKKKYRRKYYWMYWMYKTCYMWMTTTCGATCGCRKYRKKRYKWKYYRMGAKGYCKKR
50              TCGKKKKYRRKCTWMYWMYKYYYMWMAYYTTTYYATGGCRKKRYKWKYYRMRCKAYYTTA
                   .*******  *******  ****:    :  .    ************ .*.* ..

42              KKYRRYRRRATYYRKTWRRSKGYRSKYRMWGKMYKRWKMRRWKWYCYKRMMYTMYYMKYK
50              KGYRRYRRGRKTTGKCTAGGKKTACKCRMWKKMYKRWKMRRWKWTMTKRMMYCMYYMKYK
                * ******  .   *    .*   .* *** *************   ***** *******

42              MRYYRRCRYSYTKMMYYYYRKGMRRYYRYYKWMMWWWMMCKKKCWKWWKKWYYMCGYSYM
50              MRYYRGTRYGYYKMAYYYYRKKMRRTYRCYKWMMWWTMAAKGGYAKWWKKWYTCMRYSYM
                *****  **.* ** ****** *** ** ******* * .*    *******    ****

42              KRKWRYWYCYGRCGTKKSRKYASRRYKTKYKRKKWKWKYYMKYKMYRRWRWKKRCWTTTM
50              KRKWRYWTTYRGMKWKKSRKCRGRRTTWKCKAKKWKTKYYMKCKMCAAWGWKKGMTGCCC
                *******  *     *****  .** . * * **** ***** **   * ***

42              TMAYTTTYGKCAYYYGCRYRRYYRKYYRRWRKKAKCYTYCCYMYSYKYKKWKRKYRRTWK
50              YCGYAAAYRKYWYYTRYRYARYYRKTYRRAATTGTGTYCMYYMYSYGCKKWKRKYRRKAK
                  .*:::* *  **   ** ***** ***  ....      *****  *********. *

42              CGRKKKKTRRRKRMMMYYKRGWYKGCTRSWYKTMGWASKKRKRYCKYWRKYMYYRWKRYR
50              TAGTKKKKGARKRAMATYKRKTYKKYKRSWYKCCAAGCKKRKRYMGTTRKCMYCRWGACA
                 . .***.  *** *  ***  **  .*****  . ..******    ** ** **

42              KRRMRYKKKKRKWKMSYWY
50              TGAARYKKGKAKWKMSYWY
                .   **** * ********
```

The pairwise alignments of two accessions 42 and 51

```
42          MKYRRRMRWYYRRYRRWRSWRYRYCGYWCYCYRYCCYTKWYRWWWYYRTWYRYYSRMRMM
51          MKYAAACRWYYRRYRRWRSWRYRYTTTTTCAYRYGTYKKWTRWWWYYRYTYRYYSRMRMM
            ***    *****************      .***  *.** *******  **********

42          RKRWMYYYYRRRKWKMRYMRYRSKYRATRWRWKKTYRKKTTYYYMYKAYGRYRYWSYSYY
51          RTGACTTCTRRRKWKCAYMRYRSKYGRCAWRWKKWYRKKGACYYMYKRCARYRYWGYSYC
            *.       ******  ********    ***** **** : *****  .*****.***

42          RWYMKRRYWYRATMWKMCAYMCATGRASRYKKWKYYWKRYWMKCKKYYKARYWRYSKKMW
51          RWYMKRRYTYRRYAAKCYWYMMMCTGWGGYTTWKYYWKRYAMTYKKTCKGACAACCKTCA
            ******** **    *   **      . *..******** *. **  *.     .*.

42          GRSTAYRYWMMRWRMKRYWYRRRRTTRMMYYMKRYYGTRKRRYWYYWSYGYRMMTWRMKY
51          KRSAGYRYWMMGWGAGRYWYRRRRYWRMMYCAKGTYAWGKRRYWYYWGTTYRMCAWRCKY
             **:.****** *   ********  ****  *  *.  ********.  *** :** **

42          TKYTCRRAMRRRYRMMKRAWRYRYYYYYRKYTAACCRWRCYYYYRYYMYYAGTWKYRGYR
51          AKCYMRATARRRYAMMKAWWRYRYYYYYRKYWTCTYRWRMYYTCAYYATYWCKWKYRKYR
            :*   * : **** ***  ************ :.  *** **   **  *  .**** **

42          RMYYYWWKRYKSRYYMWYRKMGKRRMRMMRWKTYMCGRMKRYRRWKRWRRKWKKYRYYRK
51          GMYCYTWKGCKSRYYMWYRKASKRRMRMMRWKWYMTKRMKRCRRWGRAAATAGGYRYYRK
             ** * **  ********** .********** **  **** *** *   .   ******

42          YYKCACYGCKYYYYKYKWYKRKYKKKMRWTKMKYKKKYTRYMRRTTTACCGRCRWYSWRK
51          CCKYMYYAAKTTCYKYKWYKRKTKTKMRTGKMKYKKKTGRYMRGYWACMTAGYGTYGARK
              *   *..*   ********* *.***  *******  ****   :.   .    *. **

42          WYRYYYKYYMYYAKMMGGSKRCGRRKWRRCYKMGGTYCRCRRKYARRGGATYYRKWYYMS
51          WYRYYYKYYAYYRKCAKKSKRTCRRKWRRYYKCAACTTAYRRKYTGRRRWCTTGKWYYMS
            ********* ** *    ***  ****** ** ..     ****: *       ******

42          KYYSKKYTYYRKKKWGYYKTATCTYCWAGGYMYRKRKACYKWYMKMTKARRKRWRWSYYY
51          KYYSKKYAYYRTTKTKYTKKMKYYCMWRRKYMYRKRKCAYKACCGCYGWRAKRWRWCTYY
            *******:***..*  * *. .    *   *******..**        * *****. **

42          RKKWWSYYAGKKRWCTTAWKYYWRKMYMRYCCATRYWKSGYSMAMRYKRYSCMYYYWKRW
51          RKKWWSCYRKKKRAMYYRWKYYTRKACARYTACGRYWKSKYCCGMRYKRYSYMYYYAKRA
            ****** *  ***     **** **   ** .. ***** *. .******* **** **

42          WKKMKTTGCRYWAATMGKGTYCCTAWKRMKTYGYCTAMKWMWSKTCRYKKRYMYRYMRCG
51          TKKMKAGCTGTATGCARKACCAMKCWKRMKWYACGCRCKTCASTCSAYKKGCMYRYMGYR
             ****:      :.   *.   . ..***** *.     *   *. . ***  *****

42          SYKWYYYCCRAYKYGGWMWRKRRMCGRKMRSRSATKKKWKKKYKRWMTCCTMCCTRRKRM
51          SYGWCYCYYGGTGYATWAWRKRGMMRGTMGCGCTGKKTWGKKYKRWMYYYWATTGRGKAA
            ** * *    .  *. * **** *   .* . .: **.* *******        * *

42          KYKTMMRYKMYKYMRMMKMMWYMYMRRWKMKKWKKKRRRKMRYKYYKKYYWRRRSMATTK
51          KYKWACRYKMYKYCRMMTMATYMYAGRAKCKTWKGKRRRKMRYKCCKKCCWRARSMMYCK
```

```
            ***   ******* ***.*  ***  * * *.** *********  **  ** ***   *

42          RCMCAGMMRWRRKMRKRCTYKKKMSKKRAYKMYKRAAAAYWCYWYGMMMMWSKYYGAYRK
51          GYMYWRMMRWRRKAGGAYWYKKKCGKKGGTTMYKRTTTTYWTCTCRMMMMWSKYYTGYGK
              *   *******      **** .** . .****::::**     ********* .* *

42          YYRMYWGAYKGCACKTATRYWYKRYYYYRRKMYKYMRKMYKRYYKRKKKKYRYKKKYRGM
51          YTGATATTYTTTTYKGGCRYWYKRYYYYRRKACKYMRTMYKGTTTRKKKKCRYKTKCAAM
            *      :*.  : * . *************  ****.***   .***** ***.*  .*

42          KKRRMRGMRWKSMRKTKMWKTGYCRRWRRYYMYKYRYRYMRCKWKKCYAMCKWKWRRKKM
51          GKRRMRRMAAKSMRKKKMWKCACTRRWRRTCATKCRYRYMRMKWKKTCGCYKWKWRRKKM
             ***** *  *****.**** .  *****    * ****** ****  .  *********

42          MYYRKCYSKKWRWTAMWKYRRRKRCRWMMWKKRKCGCCRYRKKTRRRKYRKWWGRWWAKY
51          CYYGKTCSKKWAAYMCAGCARRKRTGTAMTKKRKYAYMRYRGKCAAGGYRKWWRAATRKY
             ** *  ****         ****    * **** .  *** *     *****     **

42          RGGAWWKYRYRYMYATYYYCCKWYRTKAACYGCMKYYKAYTGYYRRRYMRRWSRAYYYRK
51          RRRMWWKYRTRYMYGCCTCMTKWTGYKRMTYAMMKYYGRYYRYYRGRTCGRWSRGYYYRK
            *   ***** ****.      **   *   *. ****  *  *** *   ****.*****

42          YRMTARGRKYRKKRGRGCKYYWKYRRRYWYWMYYYKKKKCGAACARRAGTRRKYYRGRYK
51          TRMYRARRKYRKTRRGSYKYYWKYRGACATWMTCCGKKKYAGGTMRRRRKRRKYYRKRYK
             **    *****.*  . *******     **    *** ...  **  .****** ***

42          KTGAAMMKMRKRRYYWRYMRKMYKKRYMCTYWYKWYKWKCRKMYWKTYMRWRCRYRYMAT
51          GCRGGMMKAGKGGYYWRYCRGCCTKRYMSYYWYKWYKAGYRKMYWKYCMRAATRYRYMGA
               ..***  *  ***** *   .****. *******   ******  **   *****.:

42          TYRGRKKWYWTYRKRAYKMYTGWSMTCRYATATAAGYYYTRWKRWCTYAYKYMKMRGKMG
51          GTAKRKKTTTAYRKRRYKMCAATCCCYRYWYRWRWKYYYGAWKGTYACGTGCMKMRKTAK
                ***   :**** *** :. .   **       ***  **   : .   **** .

42          TTYKYWWKKKKTCAKKKGKKMWYRCATYRWYKSGYRGRTYRMKRAYRGAAMYRWKYYMRY
51          YCYKYTWKKKKKYMKKGKKKAWYGTRCCRWYKSKYRRAYYRMKRGCRKRWMYRWKTYMRT
              *** *****.  **  ** **     ***** **   *****. *   ***** ***

42          YWWRYKKYGTRTRRWRGCTCYAMYKARKRKYYYKKMWKRMRRMTKWYKGGRRRRGATRGW
51          YAWGTKKCAYRKRRWRTACATWMCGTRKGKTCTKKMWKRMRRCYKWTGAARGGGKGWGCW
            * *  ** . *.**** . .  *  :** *   *********  **  ..*    .   *

42          YRMKCKYWRRYKRCCRYTATRYCRYWWKAKMTRTTYKKRMRRRRSYKWYYCACYAWKKYR
51          YRMKMKYWRRYKRYMRCKGYRYYGYWWKWKMKRYWCTTGCGARRSYKTCYYRYCRATKYR
            **** ********  * .. **  **** **.*   ..    *****  *      .***

42          CTTYMKKTTGAATAYKWRYRWMKTTRTRWGMKMRCRRRMYKYKKRAKRCKYRRCMAMKMC
51          YCAYMKKCCCTMWWYKWRYRWMTWCRKRWKCKMRYGRRMYTCTTGTGGYGYRRYCRCKMY
              :****   :   *********. *.**  ***  ****. .. :    ***    **

42          YKKWMKWYWKKKYKRYKAAAKMGRYYMCWMMGGCWYRYMRWRRWYCTTRKYRYMYYWKMK
51          YKKWMKACTKKKYKGYKGCGKCKAYYCTTAMKKMWYGYARARRWYSCGRKYRYMYYWKMK
            ******   ***** **...*   **    *   ** * * ****.  ************
```

```
42          KRWAARMWRCTCYRYKRRKYYKMRYKYRKWTTTKKYWWYKTYRKMWCAYMYKRKYGRYAK
51          TRWGTAMWRAGYYRYKGAGTTKAAYKYRKWCACKKYWWYGAYGKMWACYMYKRGYTRYMK
            .**.: ***.  ****     *  ****** : ****** :* ***..***** * ** *

42          GKKKYYRMKKKRSWCTRARYKKKRRYWRYRSKYRSKYGGYGMGYKKMYRAACGRMKWYYK
51          KKKKYYRMKKTAGTMWRTRYKKKRGTWRTGGKTACGCATYAMRYKKMYRRRMKRAKATYT
             *********. .   *:******  **  .*  .  . *.* ******    * *  *.

42          CTATRRKKRMMYMMKKMYKRMKYRRYYMRYYRRGCSRGRKKMMKTWKWAACGTRMKKCRK
51          YWWKRRKKRMMYMMKKMYKRMKYRAYYMRYYRRTTCRTRKKMMKKAKWWTTTCRMGKYGK
               .******************** ********  .* ******. ** :   ** *  *

42          YYGYCARRYYWCATKAAKKKYGAYYSGKAGKKWRRRKRRRGCYRMWKKCRRWMRRSGYKK
51          YYRCMRRRYYWTGYKCCKKKCRCYYCRKGKKKWRRRKAGRRYCRMWKKMRAAMGGSKYTK
            **    ***** . *..***  .**. *. *******  *   ***** *  *  * *.*

42          RRGCAKMRKKKTMYKTTGYRKRWRRMWKRTYRYRCRRRYKYCGMYYRRYRRWYYYRKRYK
51          RAKTTGMRKKKWCYKCAATRKGWRRCWTRYYRYGGAAGYTCTRMYYGGYRRWTYYRTGYK
            *   : *****  ** :. ** *** *.* ***     *.   ***  **** ***. **

42          RTYYWRWTTMAWCWCKGWSKGAAGSRRKRTYYMCTYCYKKGTKYYYKKSKYKMYYCMYTC
51          RCYYAATAWCWWTATGAWSKKMGKCGGKGWYYMTCCYTTKKKKYYYKKSKTKMTCACCGA
            * **   :   *    .***  . .  *  ***     .* .******** **  .   .

42          RGRKWYYGRYTTYYMRWCTAWRYYSRRMARCKRKKRRYYRKRRMYYYKGRRSKTGTCYRR
51          RKRKTYYRAYYCCTCAAYYGTATTGGRMRAYKAKKAATTRKAAACCYKRGASKCAKMYRG
            * ** **  *         .    .  **   * **    **     **   ** .. **

42          YMGAYYTRCKSGGTAKGKSMRRYAYYRKKTYGCRRKKCYCKYRWGGCMRTCYKYAKMRMG
51          YMKTYYWGYKSRRATKRTSMRRYTCYRTKYYRMRRKKTTTKCRWRRYCRATCGCTKMAMR
            ** :**   **  ::* .*****: **.* *  ****   * **    *:    :** *

42          YYYWACRSGYMMYKAASKYYKMYMRAKYKSMKSRKWKKYMRYKTATTKRRYWRYYKRRYY
51          YYTATTGSRYMMYKMTCKYYKMTMRMKYKGMGGGKWKKYMRCKCTCGKARYWGYTKRRYY
            **  :  * ***** :.***** ** ***.* . ******* * :  * *** * *****

42          RRYMKYWRYMKYWKRMSRKKRWWKKKCTYAAKYYKYSYTTRYRKKTRKTYKKKGYMYMWY
51          RGTATYTRYMKYAKRMSRKKRWWKKTGGYMRKYYKYSYAAGYRKKCRKWYKKKKYMYMTC
            *   .* ***** ************.  *  *******:: **** ** **** ****

42          KAAMYMWWKKKRMRRKCAYYRCCYGYWMMMWWWMKMRYKATTWRMKTKYMKYTYCTTGGA
51          GGGMYCAWKKKRMRRGYGTYRTYCTTWMMMWWTAKMRYKTAAWGMTYKTCKYYYTGKTCC
             ..**  ********  . **     ******  *****:::* *. *  ** *  .  .

42          YYSYMMKWRCRTCGAAGTKSRTYASYYGKYKYCRYKKWGYWCTYKKYYKMYRRWRWKRYK
51          YYSYMMKWRARAARCCTAKSRACGGYYKGCKYYGTKKWKYWSKYKKYYKMCAAWGWKRYK
            *********.*:. .. :***: ..**   **   *** **..*******   * *****

42          RKKKYTTSYGRRYYGRGCKYWRKKYSRKKYKYYYSKKSGMWCYKYTTKKWGTGSYAGYTK
51          RKKKYKYSYRRATCRGKYKTAATTCSRTKYKYYYSGKSKAWYYGTCCKKWAAACYWRCAK
            *****. ** *       *  ..  **.******* **  * *    ***.:..*   :*
```

```
42    KACRAWCCTWWYMKKWSKYMYYYYMYWYYKCCRGKAKKACCYRYRRAWKKYKRKYRYWC
51    KCTARWYTCAWYMKKWSKYMYYYYCMCWYYKYYRKKCKKRYYTRYGAWAKKYKGGYRYTA
      *.   *    ************** * ****  * *.**    **    ****  *** .

42    CCGSYRRRSYKRRSKCKKKSKKKRGKKWSYYRKRRCCCMGGYRKSMGCGARKYYMTRWRM
51    TTAGTGARSYKRRCKYKKKGTGKRRKKWSYYGKRRYYMARRTAKSMKTTGGKYYAKGWRM
        ..   ******.* ***.. ** ****** ***        ***   . *** . ***

42    WGRRRYWYMRTMTGYYMRYRYRRGWRAWMYGRRMYGRKAAYRAYYRCRAKTTGAKYRRYC
51    WKGGRYWTMRGCGAYYARYRYGRKWRWWAYKRRMYRAKTWCAWYYRYAWKWYRRGYRRTS
*   *** **   .** **** * ** * * ****  *:    ***   *     *** .

42    WKMRYKYMYWTKATRYYKMYWYKWATGYYYMMCMTATMWKMMTMYYGKRMKYMWKYKYWC
51    WKMRYKYMYWYKRKGYYKMYWYKWGGAYYCMMTACTCMWKMAYCTYRKRCKCMTKYKYWT
      ********** * . *********. .** **   : ****    * ** * * *****

42    AGRKKKRRKGACMMMYRRYKKRYKKRMYYYSRSMWKYRMRCWGCSCKYRRKRYKKCKTKY
51    GAGTKKGAKRTTAMATRRYKKRYKGGMYTYSRSMWKYGMRMATMCTKYRRKRYKKMKAGT
      .. .**  * :  *  ********  ** ******** **    . ********* *:

42    WCRKYKRYGRWKKMRRMWKYKKRWKRGKSYWYKMAMKYYYWYRYRWMYKKYRAGRKKWGK
51    WYRKYTATTRWKKCAGMWKYGKAWKGRKSYWYKMMMKYYYWYRYRWMYGKYRWRRKKWRK
      * ***.   ****   **** * **  ******* ************* ***  **** *

42    MRRMSSMTGYCAYRKCTARWMKRKYRCCRMMAYAGYCMKRRRKRYGYYRYKWYTGMRRWY
51    MRRMSSMGAYTCTAKTCTAACGRKYRMMRMMGYTCYYMKRRRKRYACYGCKWCATAGGWY
      ******* .* .  *  :    ****  ***.*: * ********. *  ** :    **

42    YCMKKKWMKTCYKTCGKRGRYGTYKSYWYKMYYKKAYRKTCKAKRYRMYYKMCWRAMYKK
51    YYMTKKTCGGSYKAYAKRRRYRYCKSYWYKMTYGGWYRKYYTWKRYRMYYTCAWRWMYKK
      * *.**    .**: .** **   ******* *   ***  . *******. .** ****

42    KGMTYRRWTTMWRYKAGTSYTKRMYCRKCAAGCCRKMMRRATGYACTKTACYGAAGKRGY
51    KKMYYRRWWKAAACKRRYSYWKRMYYGKSRRAYMGKCMAARARYTGGTYMSYAGGCKGRT
      * * **** .    *   ** ****  *.  .   * *   : *:  .  .*... *

42    KARKGCYWMYRTRCSWYRWRRRYKTSRTCGYYYCWMYYCGWYRYKAMKWCCKKMCWYKYY
51    GRATRYCWMTGCGTGTTRWRGRYGACGAMAYYTTWMYYSKWYRYKRAKWYYKTAYWYKCT
         .   **     .  *** ** :. : .**  ****. *****  **  *.  ***

42    AAWYWRSYRYKWTRMMKKRKWACYYGKARWKTKKKTGWWKYKYYRYMRRMYRWRMWRSRR
51    TGTYWRGTRCGACRMMKKRKTMYYYAKGAWKAKKKCTAAGTKYCGCARRMCRWRMWRCAG
      :. ***. *    *******   **.*. **:***      **    *** ******.

42    WKKYR
51    TKKYG
       ***
```

The pairwise alignments of two accessions 43 and 44

```
43                  KRYYRRKWWYWKGWYGTGARMYCTWCYRGYRGWRGARRKKRKWTACGCMRRKYYKYWWRR
44                  KRYYRRKWAYWKRACRGAGAMYYCTTYRRYRRWRRRRRGKRKWCWMRTMRRKYTGYWWRR
                    ******** ***     .. **    ** ** **  ** ****     *****  *****

43                  RMRYRATTRKGTTYRWRYRRWRYWMYYWMWYTTYAATYTKCCYTKMTAYKRYWWWCKGGR
44                  RMRYRRYKRKAAAYRTRYRRWRYWATYWMWYYCYWWYYYGTTYWKMKCYKRYWAAYGARR
                    *****  .**.::** ********  *****  *   *    * **..*****     . *

43                  RRYCYRRAYKRYYCYTGRWRMKYRYAYCGCACAKSTYKMRRWYKMKSARWWTCRYYKYYR
44                  RRYTYARGYTRYYYTCAGWRMTTGTTCGATGTWKSGYKMRRWYKMKSWRWWYMGYYKYYR
                    *** * *.*.***   . ***.   :  . .  ** *********** ***   ******

43                  YMWYYMAWKKYYRRARGMWRWRRSWYAYYWYRWKYYWYAMRRRKYRCMWWRYMYMSKKYM
44                  YMWTTMGWKTYYRRTRKMWRWRRCWTTTYWYRWKYYWYGAAAGKYRMMWWRYMYMSKKYM
                    ***  *.**.****:* ******.* : **********.    *** *************

43                  YYCRYWRWWRYKYKYWTSMACYRYYCYCTTRYRKKMKRMMWKKRWKKKCRWYGRYWSSMY
44                  YYARYWRWWRYKYKYAYSMCAYRYYYYMYWRYRKKMKRMMWKTRWKKKYRWYKRYWSSMY
                    **.************  **..**** *   ************.***** *** *******

43                  KRRWKGMARYKRKTACRKKYAWSMKWYMYSKRTTCATYYYYKYYYYTYCYRYCYYTSWWA
44                  KRRWKKMWRYKGKYRYRKKCGWSMKWYMYSKAGAGRKYYCYTCCTCCCYYRYTCCASWWW
                    ***** * *** *   *** .**********  :  .** *.       ***   :***

43                  RWKKMYYGGAKTAGATRYKKRWKYAAGGAKYGKAATAAAYAAKYWACGYKYRYTAGYGST
44                  RTKKMYCTAGKCGKCYGCKKRWKYTGRTTKYCGRWWMWMYGRGCAWTRYKYRYYWKCRSC
                    * ****  ..* . .   ******:.  :**        *.       *****     *

43                  RMKYRRMYYTTRYWRYSKAWAWKWRATWKRTAKYGATWKAKKKRKCYKYKGAGAKRYRGC
44                  RMKYRRMYYCAGYWGCSKGWWWKWRTAWKGWGKTKTWWKTKKKGKTTKYKKRKRKRYGRT
                    ********* : **  **.* ****::**  .*  : **:*** *  ***    ***

43                  ACYGYMKRRGGGRGMMYGRYYYMYRYGCMRRKKTRYRCGRKACTGTKKRYSYKKARKYRR
44                  TATTYMKRGAARRKAMYKRTCTMYRYKYMRRKKKGCATARKTTYAKKKRTGTKKMRKYRR
                    :.  **** .. *  ** *   ****  *****.    .**:  ..*** . ** *****

43                  KKWYMMYAAYKKMYSKRYKRRKGYKGKKATAYKRKACGRWYKMTKYGRYKRWRYRYWKYA
44                  KKAYMMYWGYKGATSTRYKRRKTYKKKTGAGYKRKMTAAACKMKKYAGYKRWRYRYWKYC
                    ** **** .**   *.****** ** *..:.****  .   **.**. ***********.

43                  YKKRWKKTAMTKKCCCCCRYACTGYKKYGMKWYKKMRWRWRRWMMRYAYRRMWCYTYKG
44                  YKKRWKKKMMGKKATGGGTRMYSYTKYKMKWYGKARWRTRWRRWMMRTGCRRMTYYWCKK
                    *******. * **.        .*.** **** * *** ******** . ***  *  *

43                  AYRKMRWKKGMYRYWTRRGCYATWRRYKMRRRRGRWRYYRTYTKATTGTCRAYCSGTKKS
44                  WYRTMRWKKRMTRYWCRRRYCRCAGRYKMRRRRARWRYYRWCYKTYYRYAARYMSKYKTC
                     **.***** * *** **       ********.******   *:    .  * *  *..

43                  RKKKCKKMGKKYYYYKYSYWTGTYGTAGGARRKCCGRAAYYKCATYKGRGYYKKKKRTYY
44                  AKKKTKKMKKKYYYYKYSYWYKWCAYRRRRRRKYYRRWWYYKTRKYKKRRYYKKKKRYYC
```

```
         *** *** ***********    .     ***   *  ***  .** * ******* *

43      KTTGGYYWCKRRGYRYRYCTAAMWYKYGTCTKRMYYYMYKKYWYRKYYMTAKYTYYRSGC
44      KAAARYCWMKRRSYRYGYYYRMMWYKYRATCGRCYYCMCKKYWYRKTYMKRKYWTYRGKY
        *::. * * ***.*** *    ***** :   * ** * ******* **. **  **.

43      YTYMWARTYCATTAWYMYKSRRYRGTYTTCCSWAYYTKRKTKGMRYKRKATKKATGKKYA
44      CYCMWRRYYTRCGRWCCYKSRRYRRCTYYTYSWRCYGTGGCKKMRYKRKCCKKCGKKKTR
           ** * *     *  *******       **  * .   * ******. **.  **

43      YAGYRTTGKACKYMRRYYYCRMWWKYMKMKRRRMCCCRYGAGACTAGAAYTGTTRKWAGK
44      YMKTRYYKKTTKTMAGCYCTRCWAKYMKMKGRGCTMMRYRRRMYCCRGRYWRWYRKWCTG
        *   *   *: * *   *  * * ****** *     **      . . *    ***.

43      YYMCAAGCWTTCTGWWGCWYMKCRKGAMWYAYRCKTTMRRWGYYKMKWWGWYMKKKRYAG
44      CTCMWGRYWWCTCKWWRYTCCKMGKRRMWYGYRYKCCAARWRYTTATWWKWTMKKKGYTK
             .  *     **     *  *  ***.** *    ** * . .** * **** *:

43      MCMKRMKWRRKKAGKKRGYRKMYSKRTRAKRYMCCTYYYYYKKYKKAGTCKKYKRYGKKK
44      CYMKRMKTGRKKRAKKRRYRKMYSKRYARKGTCTTCYYYYYKKYKKGRWYKKYKRYRKTK
          *****  *** .*** ********   *      **********.   ****** *.*

43      RGYYAKCCATMKYYWAATYRSYWYKYRGWYGMTCYKYWMRRWRYRYWRCKSGGWMYYKRY
44      RCCTRKASWWMKYYWRTCYRSYWYKYRAATKMAMCKCWMRRWRYRYWRAGGRSWCYYKRY
        *    *..  ***** : *********.   *:  * ***********. . .* *****

43      YKYWYKRYYRYWGYCWSKKWYKRGRRRWWWYKRCMWYCMYYRWRWAWRAYRKYRYMCRTY
44      YKYAYKGTYGYATCYTSKKWYGGTARRWTATKAYMWYYMYYRWATWTRCYRKYRYAAAKY
        *** **  * *     *****    ***   *  *** *****    *.****** . .*

43      RYSYKKRGWRRRRRCAAGCGTWKTKRMYYMWRKKTGYRYGRCGYKMARRCWMCKYYYYRS
44      RYSYKKRKWRRRRRYRRRYRYAKYKRMYYMWRKKYTYRYKRYAYKMCRRTWCYKYYYYRS
        ******* ******        * **********  *** * .***.** *  *******

43      MKRAARYYRTRRMCCYCCCCYGMKKKRGTGSGKKTYGKKKSGKKRKWKGCMYTYYKKMKY
44      MKRMMRYYRCAAATTYYYYMYKMKKKRAGAGRKKKYKTKTCKTKRKWKTYMTAYYKKMGC
        *** ****      *    * *****. .. **.* .*.. .*****  * :*****

43      GRKKRWWSCCSAKSCMTAGKKGGCGWRCKKRYKKKTGTMRMWTAGCRYGTYYTRRYMRTA
44      KGKKRWWSYTSRKSYMCTKKKAATAWRTKKRYKKKWRKMAATKTTAGCSGYYKRRCMGWR
          ******  * ** * : **.. .** *******  .*   .: .  . **.** *

43      MGASCCRMRTCKATGWKKYMGTTRYMKRACYYRKMRRKTGKKKGYRTGGKCWTYRATRCK
44      CRTGYTGCGAYKCWKWKKYMACARYMKRRMYYAGMRRKKKKTKRCGAAKGGAKYARWGTK
          :.     : *.  *****. :*****  **  ****. *.*   :.    .*     *

43      KATTGMCRRKKKYMRKKYTAATAKYRRRYMKKYYGMAAYTKKKAKTAGYYCKGARYKGYY
44      KCCKCCAGGGTGYMRKKYWCCATKYAGRYMKKYYKCMMCWKKKRKACRCCYKRRRCKKYY
        *. .  .   . ****** ..::**  *******      *** *:.    *  * * **

43      RYYYWMCTRKKTGKWTKKWGGAKWRRSWMTTTAYYTGSKMKRYKCWAMRYKRYMYKRWYK
44      RYYYWMMKRTKCKKWYTKACARKWRGCWMYYKRYYYKSKMKRYKMARAGYKRYMYKRWYK
        ****** .*.*  ** .*  . *** .**  . **  *******     ***********
```

```
43      ARWKKRTRGGTRCCSGRKYTCKKKMKWYRASTYYAAARMRWMTMTKGMCYGGGGRKYMYK
44      RRWKKGYGRAYRYSCTRKYYYKKKMKWYRMSCYYMWGGMRWMYMKKAMTYAAAARKYMYK
         ****    . * .. ***  ******** * **  . **** *.*.* *....******

43      CWCRGCYYGYGYWTKWKKWMRKRMMTKCAGTATWMAKTTKRYCGYTRKMATRAKTGCKKY
44      YATARTCTRTRCWAGWKKWMGKRCMYKYWACTCWMWGGYKGCMRYAATATCRMKKRAKTT
                    *: ***** ** * *  . : **    *    *: . : * *. .*.

43      YKKGYCGAMMAAKACMRWMKGMYCARYWYWTYYRMMRMKRYWMAMMWWARWWYGCCAMTK
44      YKKAYGTMCMGGKGTMATMKKMCTGGYACWCCYRACRMKRYWCWMMWWTATACAYACAKK
        ***.*    *..*. *  ** *  . *  *  **  ******  ****:    . .. .*

43      GYTKMTTKRYMRTAKWRCYRMRWRKMARYMKCGRKTGTMKWKKKGAAAKCCCYMYYRSRR
44      RCYKCWKKATCGKRKWRYCAAGWRKMRRYMKMKRKCRYAKWKKKKMWRKTMYCCCCACRR
           *  .*    . ***     **** ****  **    *****    *        .**

43      KCKRKKKMMKWYYATTRYAAGATGRMTSRGKYKKGGGWACKTYGKRRWRMYKTGRMGMTR
44      KYGGTKKAMKATCRYKRTTTTRYKRMCSRRKTKTRATWTTGKYKKRRWRMYKWKRMRCKA
        *   .** **     .* ::    ** ** * *. . *: . * ********  **  .

43      RCAKTRYWKRRYMRKTYYWRKSYCCGCTKMRRYKKKGYTYGWSYCARGAYKKWKKKYKAR
44      RAGKYRYWGARYMRGAYYWRKSYYYKMWKMRRYKGGKTWYRWSCTTGTRTKKWKKKYKGR
        *..* ***  **** :*******     ******     * **  :    ********.*

43      KKTSSGMARYYGTGMKKTTTSYSTMKYRYYRKWRRYKMYRYRKWSMKWKTGGTMYMGCGG
44      KKKSSKMWGCTAWSMKKAACSYCYMKYRYYRKWRRYKMYRYRKWSMKTKYTAYACCKTAT
        **.** *    . .***:: **. *********************** *  .      .

43      MKGKRSKSMKWWKRYMRCYTKWKMWKGYKKRRRRTTAKRTRMRGYKTKKKRATSWRYRRC
44      MKAKRSKSMKWWGGTCRMYKKWKMWKRYKKRRRAAWGKGWRMRKYKYKKKRWKSWRYRRY
        **.*********    * *.****** ****** : .*  *** ** **** .******

43      ARCTYKYRMMYRMRYTMYYCYWMRKWAKRMKRKKGTTCTACCRGGGYYWMKGTRKTYRKK
44      WRYAYKYRCMYRMRYKMYYMTTCRKWMKGMKRKKACKYKGYGGRKRYYWMGKKGKKYRKK
         * :**** ******.***    *** * *****. . ..      ****  . *.****

43      KYCTCTGRGAKCKYKTCWYYRGACKRMRSYYRMYGAYKYKKKMSKAGGTGGAYRCKCTWY
44      KYSCTCRAAGTTKYKCTWYYRKWYKGMGSTYRMYAGCGYKTKMGKGKCYCRRYRSGAGWY
        **.     ... ***  ****   * * * ****..  **.**.*.      **. . **

43      RGKGYMAAGKKASTTAMTKYSWKAYGYRRKYMMYWYRTKMTWRMCWKKYGYYRYRGSTCM
44      RKKKYARWKTKCGYKRCCTTSWKMYRYRAKTMMYWYRAKMGWRMYTKKYKYYRYGSSKYM
        * * *    .*.. .   . *** * ** * ******:** ***  *** **** .*. *

43      GWYRKKYGGYKSRKKKAGYKKKSWMMRGCGGCGRKYKRYYAKYMKKRKKYYYYMWYWKAS
44      TWYRTKTRKYKSRKKKGAYKKKSWMMRTYRRTAAKYKRYYRKYMKKRKKYYYYMWYAKGG
         ***.*   *******..*********     . ****** *************** *..

43      CWWTCAAMCAWCYYKKYYYTKKRKMMYYYYRKYGRKKYKRCYYMMMYYYYYKYKGKAATA
44      TWWYTCWAYWWTYYKKYYYYKKRKMMYYYCRKYKRKKYKRYYYMMMYYYCYKYKKKWCGC
         **  .    * ******* ********* *** ****** ******** **** * . .
```

```
43                  RKKKRRKRYYYWMMWGCGGKAGKKKRWMWRYRGYGRTTYYYMRKTGYKCRCGGYRRRYRW
44                  RKKKRRTRYYYTMMWRSRRKTTKKKRWMWRYRSYAACAYYYMAKWKYKYATRRYGRRYRW
                    ******.**** *** .  *: **********.*.  :**** *  **     * *****

43                  MGKGMYRTGRCRKKRTAAKKTAGYRWKRRRMGRWRCKKMKYAAYKKWCKKYGRCCRTRGG
44                  MKKKMYRGCRYRKKRCGGKKYRRYRWKRRRCARWRYKKMKYTRYKKWTKKYKRMARKRRR
                    * * ***  * **** ..**   ******* .*** *****: **** *** * .*.*

43                  GAWKKKRRYYGMRWRYYRYRGAGCTKYKKRTGAYRRRYWAYRWCMYGWKMWYKAKAGKRK
44                  KWWKKKGRCYRMRWRTTRYGAWAYWKYKKRYRMYGRRYWCYRWYMYKWKMWYKRKWKKRG
                      **** * * ****  ** . .  *****   * ****.*** ** ****** *  **

43                  KYTCCCCGYYCRWGKCYKGTGKTTCGCKKKMYMKWWR
44                  KYCTMAYTTCGRTRGYYKRGTKKWYRYKKKMCCKWWR
                    **   .     *    **   *.    ****  ****
```

The pairwise alignments of two accessions 43 and 45

```
43                 YTCGRRAAYYRWWCKRTYYRWWWMRMSWMRRYYYCCKYYRRYWKGTTAACGYYCYRRARM
45                 YKTRRRWWYYRWWTKRKTYGWWWMRMSWMRRYYCYYKCCAAYWKRCGGGMSTCMYRRRRM
                   *.  **  ***** **. * *************   *    ***   .. .   *** **

43                 MARCKKGGTYYYTRMKYYGWAWKMRRRYKKRMMAYYMGWGAYAMTRRYRKYRWYRAYKGS
45                 MRRTKKRKYYYYWRMKYYAWRAKAAAGYKKRMCRYTARWRTYRMWRRYRKYRWYRTYKTC
                   * * **   *** *****.*  *    *****  *   * :* * **********:** .

43                 WRYRRKSRYWGYYSWYMGKATWWRWKMRRMRKKGRYGCAGRRYAMKAGTGTRKCMYYCAG
45                 WRYRRKCRYWRCYSWTMRGTCWAATGMRRMRKKRRYRTGTRRYTMKWKAAWRKYMYYYRA
                   ******.***  *** *  : *    ******* **  . ***:**  :. ** ***  .

43                 KAMWKRYWMYKWGCGMRYWMKSYYKWKMSRWMKYTGGKGSKKRMYYMMRMSYMKYRMRKG
45                 KCMTTGYAATKWRAKCRYWMKSYYKWKMSRWMKYGAAGTSKKRMYCCMRMSYMKYRMAGA
                   *.* . *   ** .  ****************** ..  ******  **********  .

43                 SMKYRTAGCCTGKGRYKYRTTCKYRMCWRYRGTGMGYMCYYRRKRYKRYWYRCTCGGMKK
45                 SMKYGYGCTTAAGRRYKYRAYYKYRMYWRTGTAAMSYMYYYRRKRYKAYWYRAGMKTMKK
                   ****   .  :.  *****:  **** **   :.*.** ******** ****.    ***

43                 YYSYYKGCKYKKYRTGSRKRAKRGTYYKYWYSYACTRKTKRMRATACWKYKKRYRYKRYY
45                 YYSYYGKYKYKKYRAASGGRMKRKWYYKYACCCGTARKKKRAGGWGTWKYKKRTRYKRYY
                   *****   ******:.*  * **  ****  . . :**.**  . . ****** ******

43                 CGKRSCACWKWTTKGCKRYMKTYAKSWMTCRTKCGKRKMYWKKAACARRMTTAAAAAAAA
45                 TKKRSYRYWKWCCKKMTRYMKWYCKSWMGSRYGYAKATATWKKWWTWARMKWRRRRWMRR
                     ***   ***  *  .**** *.**** .*   .* .  ***     **.

43                 YMCYTTTGCYKKAAAKKWSGKYRGAACMMCCCCARTWMMWWGWKYYKYGCMTKGGARKCC
45                 YCMTCWYSTYKKRRWKKWSRKYRRRMTACTMMMRGKWMMWWTWKYCKYRYCKKATTRKYY
                   *      . ***   **** ***            .***** *** **   .*. :**

43                 RWYMWYWKKRWKCGAAKMWYAMRWWRGARAYKRMGKWMMGYSTKKKKYYRYYWCKCYTKT
45                 RWYMWYWKKRWKYRGCKMTTWMRWWGRRRMYKRMRKWMMRYSKKKKKYYRYYWYTTCCTW
                   ************  ..**   ****   * **** **** **.********** .   .

43                 CTGCYRYMRSKYMYKTAKWRWGRYYKMGGMKGRYYYRKKRMKKKCGSYWCTWWYMYYYRM
45                 YWKYYRYMRSKYMYKYWKAGAAGCYKMRRMKKRYYYRKTRMKKKSRSYWSYWWTMYYYRM
                       ***********  *   .  ***  ** ******.*****. ***. ** ******

43                 ATYYKYKMKCTKRKYKMGKMMKKKAARTYRRGGWYYMMKTKYRGTAKKKYYMATGARYYW
45                 GAYYKYKMKYYKRKYKMKKMMKKKGCRYCARRRWYYMMKWKYRRYRKKKYCCRCATRYYT
                   .:*******  ****** ******..*   *  ****** ***   ****    .:***

43                 MMWKKRRKRWMWRYRYGYKRKCKWWSSKSYAWRYKMTAYGKRKMYSRKYRKKMKKGAATA
45                 MMWKKRRKRWMWRYRYSYKRGAGWWSSGSCRTGTTMCTYKGGKAYSRKYRKKMKKSRRYR
                   ****************.*** . **** *     .* :*   * ***********.

43                 ACKYYRYTGTMTTTCKCRYYYYKKRYMRKMSKACGRMYRMRKAGWKTTATAMYTCCCCRK
45                 RMKCCGCAAGMKCATKYGCYYYKKRCMGKMSKGAARMYRMAKCKWKWYTARMYCYTYTRK
```

```
                      *    :. *. : *   ****** * ****...***** *. **  :: **     **

43                  KYRTRKYKYGTTTYGCRATYCCTWKCCCTGRKTTTWGRTKRKCWMYYGRTTWAMYTKYRY
45                  KYRKAGYKYRYYWYKTGMYYYYYWKSAMWRGTWWKARAYTAKYWMYYRRAGWCMTKKYRC
                    ***.  ***     *     *   **..    .  .    . * **** *: *.* .***

43                  KKWTTKKSCGCTRRRKAAAKKWYTKCRMAYYKGYTGAGCRYRYGRAGGRGTCCGYKTARR
45                  KKTYYKKSMKYYRRRKRWMGKWYYKYRMWYYKACGATATGTAYRRMKKRKKMTKYGAMRR
                    **   ***    ****    *** * ** ***.  .:.    * *   * .   * : **

43                  YTRYMRKSYYMCAGAASRSMCRCAYYRACAYCTCYYRSKKTGGCRGRCMSKKKGKKCRTR
45                  YYRYMRKSYYAYWRGGSRSMTRTGYYRRYRTMWYYYRSKGCTAMRKRYCCTKKKKKYRWG
                    * ********    ..**** * .***       *****    . * *  ..** ** *

43                  GTYCWKRWKKYAWCCATGYYGCTKKYKRYKSWMKCTAMRRRKAKWMTAWGYSKRKKMATG
45                  RCYMWKGWKGYTASATWKYTKYWKKYKRYKSWMKGCCMRRGKWKWCCGWRCSKRKKMRYR
                      * ** ** *: ..:  *    ***********  .*** * **  .*  ******

43                  KWRRKCGATRYYKKTCGYRKKKRYSYRYMYYKKKKYKKRTTGAKCTTGGRYYYRWRMRWR
45                  KWRGKMATARYYGKCTAYGKKTACGYRYMYYKKKKYKKRCYRMKACCRRRTYYGWRMRWG
                    *** * .::*** *  .* **.  .*************    *.    * ** *****

43                  YYGYKATCCYGKTAASKKTKAAATYCYYYMCRTTRGWKGCRRKYYACYTGMYGKYCMRKK
45                  YYATKTYYTCKGCCCGKKYKWCWYCYYCTCTAGARRTGRYRRKYYCYCGRAYRKYMMRKK
                    **. *:        ...** * .    *      :*     *****.     * ** ****

43                  YKGTAKRRRYKRWRRRWYYKYKYKTTKKYASWRYRTKRAGGKCWGMWYRMRYTMYYYMKT
45                  YTAGGKRRRYKRWGRRWYCKYKYGYYKKYWCTRYRAKARRSKATAMWYRMRYKMYYTCKC
                    *.. .******** **** ****   *** . ***:*   .*. .*******.***  *

43                  WWTTATMTRAWKMKRKKMTCCCCRTRARYYWYMTGKKATRGWKGYKTRAGCYKKYGGCRY
45                  AAGWCYCYRMWKMKRKKMAYTGTRWRGGYYWCCYRKKWAGRWKKTTWGWRTYKKYRRMRY
                        .   * ********:    * *. ***    ** :  **   .     ****   **

43                  MRKRYAYTGKTWRMKKYWGYYMYMMKYWAKRWRRCCGKAAGGGKTKMGYRYYRRKYRRRW
45                  MRKRYRYKKKYARMKKYWKYYCCAAKYWMTRARRMTAKWGRTKKYKMAYRYYRRKCGRRA
                    ***** *. *  ****** **    *** .* **  .* .   * **.*******  **

43                  MMYAYWRWYCKRYMKKTYAKKCCCCTRWYKCCKYCTKYYRRKYKAYRATKYAAAWYMYKR
45                  MMYMYTGACAKRYMKKCYGKKATMTAAACKSMKYYYKYYRRKYGMYGMGKYTCTWYAYGG
                    *** *     .****** *.**.   :   *. **  *******  *   **:.:** *

43                  TCTGTRKYKWYRYYRMTAYRYYYCGRWKAYGTTKMRRKTRYRCSRYMYKSMCRKRRYKYT
45                  YAYACRKYKWYRTYRMKRYRCYYYARWKRCTYAGAGATKRYRTSRYMYKSMYRKAACKYG
                     . . ******* ***. ** ** .***     :     ..*** ******** **   **

43                  WRYKRKAWKRCRYWAKYKYCGTTKTTGTRMMRYMYKTTTYTYTKRRYRMCGYYYYKKKYY
45                  WRYKRKGTTRYRYARKYGYMRCAKAYAYRAMRYMYKYCGCYYKKRAYGMTAYYYYKTKCC
                    ******. .* **  ** *   :*: . * ******      *.** * * .*****.*

43                  CYRKTSKKRRCYKAKYATSYWYYRTRGYRYRGWYGMYKRYTWGTTMRWGKRYWTTRYRTA
45                  SYRKCSKKRRACKRKYTCSYWYYRKRRYRYAAATKMYKGTYWKCGMRWAGRYWACRCRCT
                    .*** *****. * **: ******.* *** .    ***    *    ***. ***: * * :
```

```
43          GGRRMAYRMYWRYRRCTRMYYSYTAYYCCTCTACCRAAGGYWKWMMKGCKRYWKRRRKYR
45          RSRRMWTGATARYRGYCACYYSYKRYYTMAMCWTSRCWRRYWKWMMKKMKAYWKRRRKYR
             .***      ***     ****. **  :    .*.   *******  * *********

43          TAKYRKKTTYKKSGCKCKMCCCTMKWRCTAMYRSTYYATGWATMAYAKMGTYKYRARSCG
45          AGKYRKKGCTTKSSYKMKMYYMYCKWGGKWAYRSCYYCCTWGWCCTWKMSKYKYGCRCYR
            :.*****   .**. * **     **  .  *** **.  *.  .  **..*** .*.

43          KTCGYTGTAYWRKCYMYKKRKGCYRYRYYGYYMMYRWYYYGKCTAGCTKMWKTGTYYMMW
45          KKTAYACAGCWRKYYMYKKRKKYCRYRYYRYYMMCRWYYYKGMCCTYYKCWKKKCTCCMW
            *. .*: :. *** *******   ***** **** *****    .   * **.     **

43          YRYWGARGCGWATCKKWRYRRTTMRYARRWWGAYKMKAWYRKWYMWWKYMKWKMAKMRRY
45          YATTRGRCTRARKYKKWRYRGCCAAYWGRWWRGTTATMWYRKWYMWWKYMKWTMRKMRGC
            *    .*     . ******     *  *** . . . **************.* ***

43          GYAKGGAGKCTKGTGMMKARYTCKWYGMYKKGGYGWYKKKMCTCRKYKYGMKMGTAAMRT
45          AYRKKAWAKTCKKCKMCKRRYYYKWYKMYKGRRYRWYKKKMYWGRKYKTACKMKWRWAAA
            .* * . .*  *   * * **  *** ***   * ******   **** . **      :

43          RRTTYYTKYKYRWMYTCCTRRMYYYKAKTYWWSYYKTTCCGGGMRWSCAYRRYCAAKKWK
45          GAYKYYYKYKYRWMYCATARRMCCYKWKKYWWSYYKWWTYRAACGWSYGCGGTAGRKKWK
               .** ******** . :***  ** *.*******     ..  ** .    .. ****

43          TKKYKARTRSSGMKTTTRSYKTTYRKKKYKMGYAGGKRWACCTMYMYGTAGMAKRSMKWC
45          YKKYKGGARCGKMKAACRSYKCKYRKKTYKMCYWAAKATRYYCACCYKKRRCWKRSMKWY
             ****. :*.. **:: **** .****.*** * ..*         * .    ******

43          RTKKWWTTTMYKKKGRKYYYMTWCATTCYRRRWMKGRYSTKKYYWYRKWKRRCYYRAKRK
45          GWTKWACCWMYKKKRRKYYYMCWYWWYMYRRRTMKKRYSCKKYCWYRKWKRRYYYATKRK
              .**    ***** ****** *     **** ** *** *** ******** ** :***

43          MGTYKMKRMKRKTWKGTWTYMRWRKWRRYRMRKCTMGKGAKYMYRKKACAWCTAKSMTKK
45          MKCYKMKRMKRKKWKSWWACMRWRKWARYRMRKYKMKKKCTCMYRKKCGCWYGMKSMYKK
            *  *********.**. *: ****** ****** .* * .. *****. .*   *** **

43          AACCGRRGCWCAGGWWKRYKYRRMRYYRMKTKCGGATTKARKYRMKSRCWTGKCGGTRKC
45          GGTTARGTAWYWRSTWKRYKYRRMRYYGCKYKSKTGKAKGRGYRMKSRYWKRKTAAAGGA
            ..  .*  .*   . ************  * *.  ..:*.* ****** *. * ..:  .

43          CCCWTTARKCWTKKWGTMCRCGKRRCKRYYRACMRWRKCYKGAAGYCTCKKGRAWCRYCR
45          YMYWCCTGGYAKKKWKWMYRTAKRATKRYYRRYCRWRKMYGRCMKYYYYKKKRMWYGCTG
               *  :    .***  * * .**  *****   **** *  .  *   ** * *

43          TGWRYCYRAGGARWCCGMKMTCKKYARKYTCAAAKKKKYGMYKKKGYYRYMRRKKKGYCY
45          AAWRYGYRWSRGRWTAKMKMCMKKYGAKYATTGCKKKKYACCKKKRYYRCMRRKKKKYMY
            :.*** ** . .** . ***  ***. **: :..*****.  *** *** ****** * *

43          KKWKRKKYYYWKAYYKRKKSRMARKRRKKMKMMYRYYYYKGKTTCCYGASYWKTAGARKK
45          KKTKRKKYYYATTYYKRKKSRAMRKRRKKCKCCTAYYYYKKKKATTCAWSYWKYRRRRKK
            ** ******* .:********  ****** *    ***** *.:   . ****    ***
```

```
43                  KYTCGYKKYKYKTWRCYGCTAKYWARRATGYKYACRYRRKRWTTKAWKKRRGGACTGYKY
45                  KYYMRYGKYGTGKAASYTYYGKYWTRRRKSCKYWYGCRRKGWKCKMWKKRGRRRMYSYKY
                    **   * **   .  .*   .***:** .. **    *** *. * ****      .***

43                  RGTGARKCCCCGRWMMRSTRRTYTCMYKMRWKYMTYGAWRYCKASMMKYRACMGRRKMTY
45                  RKKTRRKTYYTTRWMMGCCGRKYYSMYTMRWKCMCCAMAGYYKGSMMKCRRYCRARGMYY
                    * .  **     **** .  *.* .**.**** *  .   * *.**** *     * * *

43                  YRRRGRCYRCCC
45                  YRRRARMYRYYY
                    ****.* **
```

The pairwise alignments of two accessions 43 and 46

```
43                SRRWRAAYRRSWRWKSRSCMYMYYYRKKKYRWYMRCCCYCAYMKYRGYRARRRYWKRMGC
46                SARWRRWYRRCAGATGRGYCYMYTYRKKKYRWYMRATTYSMYMKYRRYRCRRRYWKRMAT
                  * ***  ***.   ..*.  *** ***********.  *. ***** **.********.

43                YWRYYMYRKRARKYYGGWMYSRYWMYRKKTWRYRKYACWCKARRWYMWRWKSKRYYYWMY
46                YWGYYMYRKRWRGYTRRWCYSRYWMYRKKATAYRKYMMAMKWARWYMWRWKSKRYYYWMY
                  ** ******* * *   * **********:  ****    *  *****************

43                MWMYCRWYKRWGTYKKRYYWYRCTYCRCKRMKCYKSYKKCGYSRWWTRKKGMYYYGTRYW
46                AAATYAWYKRWRWCTKRYYAYRTATMAAKRMKYYKSYKKMRYSRWWWRKKAMYYYTKRYW
                        *****   .**** **  :  .**** ******  ***** ***.**** .***

43                WMRSMSKMYKMYWKRGCYYRRSGWYMTWMAYWMKRYWKKWGWYGTCACMATYWRKYYYYM
46                WMRCMSKMYKMYWKRCTYYRRSAACMAWMGYWMKRYWKKWRATKYTMYARYYARGYYYYM
                  ***.***********  *****.  *:**.**********           * * *****

43                WYKCKTYGTAGCGCCWCYYYGCKMMYYYGCCRYRYYMWRMKKTGYRTYSKRYATCGKYKY
46                WYKYKKYACGKSATATTYYYATKMMYYYRYTGYRYYMTRMKKYRYRYYSKRYWCTRKYKY
                  *** *.*. . .. .  ***. ******    ***** ****  ** *****    ****

43                MMGCATCKCGAGCRKKYYYYWKKKKWRYRYTGYRKRMKYGTCGSCGTYGGGAGAYYYCAA
46                MMRTTATKYKRAYRKTCYYYTKGKGWRYRYCSYRKRMKYKKMRSYTYYKAARRRYYYMGC
                  **   ::*   . **. *** * * ***** .*******  . *   * ..   *** ..

43                GAYMTMRAAAYYYKRSMCRYTRRMCCACCYGRYGRAWMYRKRYWKMYGKTYYKKSKAGAY
46                SMYMYCAGTMYYYKRSMTGYCAAATTWYYYKRYAGRWMYRKRYWKMYKTKYYKTCKGRGT
                  . **   .: *******  *         * **.  *********** .. ***..*. .

43                YRRAGAGCTKKTAKWKYKYWTYGTTTSAYYTTRMRAGRSYWTRRSMRTMCAAMYYGRWYR
46                YARCRWKYYKKKRKWKYKYWKYKKKYGTYYKKAAGRRRSYWYRASMRCMYMMMYYRRWYG
                  * *.     **. *******.* .. .:**..     **** * *** *   *** ***

43                CWMATYCCGYRYTCGMMRWRYMYYRRTYTYGKKACWRRRRRKGWWKRYYYTRRWTKCMYY
46                YWARGCTTKYRTKYKMMRWRYMYYARYYYYRKKWYWRRRRRKRAWKRYYYKRRWYKYMYY
                   *       **  . ********* * * * **  *******  ******.*** * ***

43                RTCCARWTRGTYGKWARYKKKGGYRKYKRWYWGKTYRWKRAYKWMMAARCRMGGRYMYYC
46                RGYYCRWKRRYYKKWMRYKKKRRYRKYKRWYWKKCYRWKRTTKWMMWRRYRMATRYMYTY
                  *   .**.*  * ** *****  ********* * *****: ****  * **. ****

43                YKYKYRMRWYGKMCMMYYGYYWKRTRKTKAKRRRGWYRRRKWTGATTTMYSKYMKMYYRR
46                YGYKYRMRWTRKMGMMCTRYYWKRWGKYKRKRRRRTCRRRTAARRYWWMYSKYMKMYYRR
                  * *******  ** **   *****  * * ****   ***. :     ************

43                RKRYTTWRYMRGTGYWYRAYTYYSSYYKKKKAYKGYRGGARGCACGKKYKTRKAAATCTC
46                RKRYCWWRCCRKWRYWYRMCCTYSSYYKKKGTTKRYRTTCGATTYKKKYKKRKRWRYTGG
                  ****  **  *   ****    ******** : * **  . . :  ****.**

43                TMMCRKKTTRYKKKRTYYRATKRYYKRYKYKKMSMKMRCRYRKKKTAYARWSRACTTCWW
46                AMMYRKGGGGYKKKRKYYRMKKRYYKRYKCKKASMKMRTGYRKKKCRCCGWSRGTAAYWW
```

```
                    :** **    *****.*** .******** ** *****  *****    . ***. :: **

43                  GTYKWYRKKRGYGYWACAKKKCTKKKYWGKYKRAAAARWYRYMTWRGKYMRRWMYYYKMR
46                  RYYKWYRKKRRTKCWTMRKKKYGKKKYWKKYKRMRRWRWYRYMKWGKKYMRRTAYYYKMR
                      ********     *:  ***  ***** ****     ******.*  *****  ******

43                  SYCKYKTKCWYCRKMAGGAAGRAAGGKSCTTAYGYMWRRWWYSRKYSWKYKWYMMGTWRY
46                  GTYKYKKGYAYYRKMRKTTGRGRWKRKSYYKRYCYMWRRWWYSRTTSWKYKWYMMRYWAY
                    .   ***.    * ***    :.       **   . * **********. *********  * *

43                  YYRWKKYMGYAGTGRKCTRWRWRWSKMKYYRRGYYKKAGASGTKKRWGKGGGKCTCYTYR
46                  CCGWGKYMTYGTWARKMYRWRWRWSKMKYYAAKYTKKWKMSRKKKGTRKRAKGYWMTYCR
                       * *** *.   .**  ************    * **    * .**    * .       *

43                  KRRATTCGYTAGMMCYWTTTWATTATACRRYGMAGRRKWCKKYTYAGYWKRYKSWWWKYM
46                  KRRCACTAYYWKAAMYWACGWRKYGWRYGGCRCGRRRKWYKKYYYWRYWKRYKSWATGYM
                    ***.:   .*      **:   *  .  .        . **** *** *   ********    **

43                  GRKCYAYYYKGGGCCYKSYWKRCRKKRYRKKYMTCCTGGRKYYKYKWYTYYGSCKAWYAG
46                  TRKTCGYYYKKKRYYYKSYWGRYRKKRCAKKYMYAMWKKRKYYGTGWYYYCAGTKMWCCC
                     **   .****      ***** * ****  **** .     ****    ** *  .. * * .

43                  RCAAMRRGGYRYRRKYMCAKRRCARTYTRGTMRYYRMYARYGACTCKRKYWTCGYAGTWC
46                  RARMARRKSYGCRRKYMMMKRGYRGYTYGRAMRYYRMCMRTKWTAYKRKYTYYTYWRCWY
                    *.    ** .*  *****  **           :******  *     : ****     *   *

43                  GAYGCMRTYKRMTMMAKGWTWRCYKKWRYRRMRWGAGCRYKMKKWAWWWYKMKGYAYGMK
46                  RMYRYCGCCKRMKCMCKRAKAAYTKKWRYRGAAWRGSYRTTATKWMWWWYTMKKYWYTMK
                      *      ***. *.*   .     ******    *  .. *  .  .** ****.** * * **

43                  RMMRKKMYRAKTYTAYRRRYYRMKRRCGKWATKGARRYYRMRCGYKAKKYKYAATSWRYG
46                  RMMRKKMYRRKWYCMYRRRYYRCKRRMCKWRKKRWRRYYRARYKYKTTKYKCTWKSWRYR
                    ********* * *  ******* ***   ** .*  ***** *   **:.*** :  .****

43                  RCKKCWGWMWKRYMYKRYYYMTRWMYKRKKKYRRGGGYYWYYKKSGKRYWKKKATKKKRR
46                  RYGKATAWMWKRYMYKGYCTCARWMYKRKKKCRRRRRYYWTYKKSKTGYWKKGGKKKKRR
                    *  *. .********* *    :********* **    *** ****  .  ****  ..*****

43                  WSTYKRTKRYRKMMRYCRCAMYWCTWGTKRRYYYYYKKKKKTYWYKTYWKKYCRYYAYKK
46                  WSCTTGCKRYRKMMRYTGTMMYWMKWRYKRAYYYYYKKKKKYYWYKYYWTKYTGYCRYKK
                    **   .  *********     *** .*  ** ********** **** **.**   *   ***

43                  YYWAATASGGKRSYWYAKRWMAAAGRYKWMRSTGTTACYWSRCSRGRGGMGCGRMYRMWW
46                  YYWRTCTCAKTRSYWYRKRWMTGTARYKWMRSYRYYRYYWCRYGGRGRSMSYKGCTGAAT
                    *** : :.. .***** ****:.:.*******        **.*  .      .*.

43                  AGATCKYAGAMYRCKCRYGGAYKKKMRYRRMKKYWWYYMYMMKKKGGYTKYCCYRCYKWM
46                  WAGCTKYRKRATGTKSRYKRRYKKKMRYRRMKGCATYYCCAAKKKKRYKGTMTCGGYKWM
                     ..  **        *.**   ***********     **     ***   *.         ****

43                  KRYRRRTATYCGACTCGTRYKRKMRRKRRKCTAYKKMYKRYYYRKTYATRWYTTRRRYMY
46                  KRYRRGGCCCTATTYMAWGCKRKMRRKRRKMKRYKKATTRYCYRKKCCGAACKKRRRYMY
                    *****    .    .:    .    ********** . ***   .** ***. .     ..******
```

```
43          ARKKRYYMRRMGMTAAKYGARRATCYWKKTTAWMKCKRGRYMKTATCCTMATMRGGWRGY
46          RGKKRYYMRRAACCGTKYKWRRMKSYWKKYKMWMKMTRKRYMKATASTGMGAMRARARRC
              ******** .  .:**  ** ..**** . *** .* ****:::.  *.:**.  *

43          AYCTRKMAWKKKYKGGRAAMRYKKRWCACGCAGATAATTRYYRKWMKWYYRWKWRMMKYT
46          MCYAATAMWKKKYKRAACCMRYKKRTARTATMRWYWWGKRYYGKWMKAYYGTKWRACKTA
               : .  ****** . ..****** .  .        .*** **** **  ***  * :

43          CCARWMKKYYYMTKRACGAARSYYYYMTRRKKKGRTGCCCKMRRYWSYCGAAKGRSTAAR
46          MMRRWMKKYYYCKKRCTATRRSYYYYMWARKKKTAATSYYKMRRYWSYYKRRKCRSYGGR
               ******** .**. .: *******  ****  : .  ********    * ** ..*

43          RYYRWYRKRWRMGTCRWRCMKAGYGTMTGAGYRRKYYRRTGGGAWKKWKGTMWMRWRYCR
46          AYYRWYRKRWRCAATRWRYMKGKYKYACSCCYRRKYYRRKRRKWWKKWKKKMTARWRTYG
             ********** .: *** **. *    .. ********.    ***** .*  ***

43          CKYCKRTCTKYWWSMRRKCWKSWAAATCKGYKCTAAKSSKGMKKRSYMMTAGAKKTYYRK
46          YKYYGGCMWKYWWSMRRKMWKSWRCGCTKKYKTCGTKSSKTCKKRSYMMCTCTKKCYYRK
             **      ********* **** ..  * **  .:****  ******* : :** ****

43          MKCRYKYCYRMMMKYYKKMRKGYAATMYMYTAGGMKKKKRSSMKWKARYMRCYTYKWKMC
46          ATYRYKYAYRMCAKCCTKMAKRYRMYACCYCGARMTKKKRSSMKWKWGTCRMYKYTWKMY
             . ****.***  *  .** * *      *  .. *.**********    * *.*.***

43          GGYKRKKCRWYAKWYWKKRRWKGAWRKMSWGMWRTACATTKKWMMGYKKKKTGKMGATGG
46          RKYKRKKYRWYTKWYWKKRRWKRCWRGACWKMTRCGTGYYKKWMMRYKKKKKKKCAGACR
              ***** ***:********** .**  .* * * . .  ***** *****. * ..:

43          ACYRYAWMRYTYTTTWKKRTGCGAYKGCTARYYYTKYRWGCAWKGAKYSCTGKYYKRTGC
46          GYYRYRWMRYKCWYYTTKRYATTTYKKMCRGCYYKKYRTRTGTGRCTYSYWRKTTKRKRY
            . *** ****.     .** .  :**      **.***   .   ..**   *  **.

43          CWCACTKKYGWYGRTKATMRKKKYMCKKYCAAATAGTTYRYTYRRYRWWYWKWKKTTTAY
46          TAYTTCKTCSAYRAYKWCMGGTKYMMKKYYTCCATACCYRYWTGRYRWWYWKWKKCAWWY
               :  *. . *   *  *  .*** *** :..::.  ***   *********** :  *

43          KKMYKKRRYCTGKYAATYYCKTCAKYTCMACAKRRTTKMKKYRKKRKKTYWWSCGCGSCS
46          KGMYKKRRTACCKCGGCYYSGCTRTYYMAGMGKRRYYKMKKYRKGRKKCCWWSYAAKGYS
            * ****** .  * .. **.    .*   . .***  ******* ***  *** .. . *

43          MRYMGKMKRYSRYYKKAACYTGTCCTAYTGTYACAAYYMKRCYMRAKCKGRMYKATGWKK
46          MRYMKKAKRYSRYYKKRRMYCRYSYYGYAAGYRMWWYYCKRTCMGGKYKARMYKMWKWKK
            **** * *********   *   .  .*:. *    ** **  * .* *.****   ***

43          MCYKYGCRAYWWMKRYKYASYRYMYWKRRRWKYKAGACKCCRYRRRRAGAAKCRKKYKTC
46          MYYKYKTRCYWWMKRYKCGGCRYMYWKRRRWKYKRATTGYYRYARRRGAGRTYRKKYKYY
            * ***  *.******** .. ************* .:    ** ***... . *****

43          TYMMWYRSYYGTCMRRRMCCTCAGGGGRRSKCGYMYYYWYYYTCTARMWYRMTKRRWYRW
46          WCMMWYRSYYACTMRRRMYMKMRAAAAGRCKTAYMCCTACCYGYYRGMWYRMGGRRWYRW
              ********.  *****  .  .... *.* .**      *     *****  ******
```

```
43                  RKKYKYRGYKYMGYCGGGYRWKYTAMACATGYMGRYRKRYWTMTWRKKWARGRAGKTTGA
46                  RKKYKYRKYKYMRYYRKRYRWKYKRCWYWKAYMRRYGGRYTCMCWRKKAMGKRRATYYKR
                    ******* **** *    *****.     ..** **  **  * ****    * ..

43                  GAWKKAWMYRMRMAGGGRWAYMKTTGGMYRKYTYKAAARYKYMYCGCAGGKMYKTRRMRY
46                  AGWKKWWACAAGCMRAARWRYMKYYKKMYRKYCYKMWRRYKCCCYKYWRSKMYKKRRMGT
                    ..*** *        ..** ***    ***** **   ***        .****.***

43                  YATRMKTTCCGGGAKGGKRTMSRATARGARSRKGWMGRGWRAKTRRYWKCGKCAGRCMYM
46                  TRAGCKCWYYKKKRKKAGGWMSRMAMRRCRSRGKWMKRRARWKYRRYWKYRGMWARYMYC
                      :  *        * .   *** : * .***  ** *  * * *****     .* **

43                  RKCGTY
46                  RGYKYY
                    *    *
```

The pairwise alignments of two accessions 43 and 47

```
43                 RRRMKRYRYGRATYRRMKKTTRTTKRYYYRYWWRTRRYYYKRMWWRRYYTAGMSKWRYSR
47                 AARMKRYRYSRWAYRRMKKCARWWKRYYTGYWWRKRRYYYKRMWWRRYYYGRMSKWRYSG
                     *******.* :****** :*  ****  ****.************** . *******

43                 RYWRMTYRKKMYRKYMRRKRMYSKKSKCWMRACGRWKSMTCYYCYYWRYRRSRYWWYRRG
47                 RYWRMWYRKKMYRKYMAGKRMYSKKSKTWMGCYARWKSMYYTCAYYWRYRRCRYWTYRRK
                   ***** **********  ********* ** . .*****    .*******.*** ***

43                 YYKKMWMWYKMRMRYWGGWYYCCRKWKRYWGRMYRRMYRWYKKCKRYWKYKMRMYAYMYY
47                 CYKKMWMWYKMGMGCAKAACYTTGKWTRYWKRMYRRMYGATKKMKRYTGTKCGCTMCMYY
                    ********** *    .  *   **.*** *******   ** ***   *      ***

43                 TCKRWYWTRYKRYAGWGRYTKWYYMWMRRYYRRSKYWKYMKCMWYMRYRRRWKSKYCYSR
47                 YYKRWYTGRYKATWTWTRYWKAYYMWMRRYYRRSKYWKYMKTMWYMRYRRRWKSKCYYSR
                     ****  ***    * ** * ******************* *************  ***

43                 WWRWKSKWKATATATYRRRMMYMYRWGGWKRMRYYYRWYYYKYKYKTKKKATSWRYRRCK
47                 WWRWKSKWKTCGCMKYRRRMMYMYRWKKWKRARYYYRWCCYKYKYGYKKKWKSWRYRRYK
                   *********: .  .***********  *** ******  *****  *** .****** *

43                 YWGWMYRMRYMYCMRYYTWWTTATMTRAWSYTKKYMKRKKMTCCKTYGCGATGCKCYYWR
47                 YTRWCYRMRYMYTCRYYCAAGWCYCYRMWSYWKKYMKRKKMATGKKTAAACWCYGTYYWA
                   *  * *******  ***     .   * *** *********:  *. ....     ***

43                 YMYRKKAKCSYYKWRKKKKCYTYRGAYMKRGKKKYYYYMWYAKRMWCGSTWWGCWYYYRK
47                 TMYRKKWKMSTYKWRKKKKAYACARTYMKRKGKKYYYYMWYMKRMWSRSKATKTWYYYRK
                    ***** * * ********.*:   :****  ********* ****. *.    ******

43                 GYKKMMYYYKYRYMRYYMKYYATWRGKMKKRKRRRTRGKKRGGTYKMKYRKGYKMYYYWM
47                 RYKKMMYYYKCRTMRYYMKYYMKTGRKMKKRKRRRCAKKTRRRKYKAKYRKKYKCYYYTM
                    ********* * ********  .   *********   *.*  .** **** ** *** *

43                 MWGYACKKKWMWRYRYAKGYRCKMKMYYRTRYRRYYGKRRKRYMYMRKWAYGAAATGKYM
47                 MWTYGTKKKWMWRYRYMKSYRYKMKMYYRWRYRRYYRKRRKRYMCMRKWWTKRWRYSKYM
                   ** *. ********** *.** ******* ****** ******* ****       .***

43                 GCYMKGYYYRYTGAMWYGGCCKKWRYRTMWGAYKMKWYARCCATMKMMGRYYKAWWRRMA
47                 RYCCKACTTACKAGMWYCAYSKKWRYRKMWRGYKMKWTGATTCKMKMMKRYYKGWWRRMW
                       *.     ...*** . .******.** .***** .   ..**** ****.*****

43                 AKWRMTYMKRSAMRTGWKGKMARWWYGRKYKMMKKAKKRAAASYMMMAKCKACYGKRYKK
47                 RKWRMWYMKRSTARKAWGRKMWRAATRRKYKMMKKMKKRRWRSCMMCTGYKGAYKKAYKK
                    **** *****: *..*  ** *    ******** ***   * ** :  *..* * ***

43                 RYYRRGKGWCTAYGMATMYKKRCTCWYYRRKRMRSYMGYRMGWKYKYYCATGKTKMYARM
47                 RYYGGKKKTMCRCRMWYCYKKRMCTWYYRGKRMRSYAKYRMRWKYKYCTRWKKYKMYRRM
                   ***   *       *   ****   **** ******  *** *****     * *** **

43                 CKKRGGATCKYAATYKATTTTARRAYCMCTTSYKRYRWKMTWYYYYYMRKKGMRYRWGWS
47                 SKKRRCGKMKYRWKCTTGCYKRRRCYMCYGCSYKATAWKMYWCCYYYMAKKKMRYRWRAS
```

```
            .***  .. **  . .:   . **.*     ***   *** *  **** ** *****  *

43          YCGAAKMYRRASKYTYRRWKRRAKRWRARMYYAGKYRKMKYYTTTRRGGGAYYWWKGRCM
47          YYKRRKMYRRRSKYWYRAWKRRTKRWRGRMYYRKKYRKMKYYKKGRRRRKWYYWWKKRTM
            *    ***** *** ** ****:****.****  ********.. **    ***** * *

43          WRAARMKYCKKMYKRYTRRCTKYKCAYYKMWCKRSMRCAYYRKTRRMCCKSKYKKTTGTK
47          WRGGRMKYTGKMCGAYCRRMYKYKTGYYKCWYKRSMRYGYYRKCAAATTKSKYKKCARWK
            **..****  **   * **  *** .*** * ***** .****      ****** :  *

43          KGCGTASAWRRRYMRCCRKKSKKGKKCRYYWKWWYRKYKWYTGAARATCKRKKSRKGMKM
47          KRTRYGGCWRARYMGYTRKTCKKKKKYRYYWKWWYGKYTWYYRGCGCYYKRKKSRKKAKM
            *    ...** ***   **..** ** ******** **.**  .. .  *******  **

43          KRAKYYYYRYYYMYRKKCCYAGCKSYWTAGKAKRKRGYYRKKYTCRGYKKKKWYTYRYYW
47          KRCKYYYYRTCTMYRKTTTYGCTKSYWYRRKRGRKRKYYRKKYKARRYKKKKWYYYGYYW
            **.******   ****.  *.  ****   *  *** ******..* ******* * ***

43          YSRRTKGTKKKYGRYRGTKKKCTAWYMYCGGTYRMGRGRKKYWYRGCMTAGGAWARGWAY
47          YSRRKKSYKGKYARYRKWKKKYYRWYMYYRSYYGCRRKRKKYWYRKYMKRARMWCACWGC
            ****.*. * **.***  ***   ****   . *   * *******  *. .  *.  *.

43          YAATRYRARYTYMYKSKRKYRCYRKMRKYMRKKYTWRWAKAYRRTTGKWWKCYTCCCGAG
47          YCGKRYRRRTCYMYKSKRKYRMYGGAGTYMRKKYATRARKCTRRYYKKWWKTCCYMYSRA
            *...*** *  ********** *    .******: *  *. **   ****      . .

43          RGKCCGTMCKKYRTKKRYWWSTCACGCGCGSCKSYAKKMMYKKRGKMKYSRRYGTMWRKT
47          RKKTAAAMMKTYGKKKRYWWSCMRGRYAAKGYKSYRKKMMYKGGKKMGYSRAYTCAWRKY
            * * ..:* *.* .*******      .. . *** ******   ** *** *   ***

43          TAKRACKYMMYMTTAACAAYYYTTYWTKKCRMYKWKKYMYKTKKYWARKMRYMYYRKTCY
47          YWKRRMKYMMYMKATGTTTYYYKGCWGKKYRMYKWKKYMYGCKKYWTRKMRYMYYRKCYY
              **  ******.::. ::***.  * ** **********  ****:**********  *

43          ACTKYKGAAAAKAWYMYYAKKCKKKGTKRRYYAAAKTGGATTTRGKRYGRKGTTGMMMRS
47          WYAKCKKMWRMKTWCCCYTKKYKKKKKKRGTTRRWGAKTGKAGRAGGYARKAKARMMCRS
              :* *     *:*   *:** *** .**       :  ..: *.  *.**..: ** **

43          GWMTMYCGRYARYTKRGRYWRRKRYWKRRYMRYGTYKGTKCTRTAYRMKYYGRSWRRKWK
47          KWMAMTTAGTRRYWKRRAYARRKRYWKARYMRYKYYKRCKYCGCGCAAKTCAGSWAGKAK
             **:*  .   ** **  * ******* *****  **  *    .   *  . **  * *

43          RYMCWGRTTWMYYTATMGMYYRYYRWRMTGKKRWRRKAGCGKKCWKYRCCYKRCRRGGRT
47          AYMYWRRAGWCCTYCCAKCCYRCYRAAMYRKKRWRRKRRYRGTTWKYRGYYKRYRGRKRK
             ** * *: *    .     ** **  *  *******     . ****  *** *   *.

43          RYRYKAAYKRYTGGARSMTCGYCKTTTCKRRYRRYRMRKTAGKYYTCRWKKTTACTTRWK
47          GTAYKGGTKRYCTATGSMCTKCYKACAATGGYRRYRCGTYTCKYYKYGWKKCARTAAAWK
               **.. ***  .: **     *: :..  *****  . : ***.  *** :  :: **

43          WRGCKWYKKRMYACTGCAKKKKKKKRRKYAYAGARYYRKKARRMKAKCYKKYWWGYYYGY
47          WRTYKWYKKRACGTCTAGKKGKKKKRRKYWTRKRRYTRKKWRRMKGGYTKKYWWKYYYKY
            **  ******  .   ..** ********     ** *** ****.   ***** *** *
```

```
43  KKMKRYRRRKRRTKGTTKTAYRRKRRYYMMYATGTTYGYGGATAGCRRKKRYMKRRKYKK
47  KKMTRTRRRKRRKKSKWKYMYRRKRGYYMMYMYRWYCAYKRWWGATAAKKRYMKRRKYKK
    ***.* ******.*.. *  ***** *****      .*    ..   ************

43  CYYKYTTRRWYKRKRRRRKRYGMRRRWRMMKYRYMKKCCRWYWKYWAMKMTTKRWKWRTT
47  MYYKYYKRAACKRKRRRRGRYKMGRRWRMMKYRYCKKYYRWYAKYWWAGCYWKRWKWRGG
     **** .*   ******* ** * ********** **  *** ***      ******

43  RMTARYRKKKKMKYMMYSRTKKKYYRGYYWKCYRYYTKYKGWRGCARYKRYGYYRRKKRY
47  RMKWRCGKKKKMKYMMYSAKKKKYYRKYYWKYYRYYYTYKRAGTAMRCKRYRCYRRKKRY
    **. *  *********** .****** **** **** .**    . * ***  *******

43  AYWGKSKWMMKKKRRAYKRYYAMRRRWCYYYRRYRRWKGACYWYAAYCSKSTTCTAMKYG
47  CYWKKSKWMMKKKRGGCKRYYTMRRRWYYYYRRYRRWKATTYWYGGTYGKSKYYWRMKCS
    .** ********** . ****:***** **********.: ***..  .**.    ** .

43  GAYSYYRKWKMCAATMAGYYCARYWYMYGRKKRWKMYKATKGGKTACTGYAKYGRKAWKA
47  RRYSYYRKWKMTCGKMCAYYGGRYWYMCRRKKGWKMYKMKKRRTARMYSYRKYKRKRTKR
      ********* ...*..** .*****  *** ***** .*  .:   .* ** **  *

43  RWKCCCCGRKAWRWMMRSRRMGRYSYCYKYKYYRYRWKMRWKGKCGTGYGGGAGRRCRAA
47  AWKTYTTTRTRTRWMMGCRRMRRYSYYYKCTYYRYATTMRWKRKTAGTYRRKRARRTRRR
     **     *.  **** .*** **** ** .****  .**** * .  *    .** *

43  CKKYWRRRYRKRYARYYMTTKKKAYAGCGKRWRRYGWWYSKMKKYYYRWMRWCYRRYRGA
47  YKKYWRRRYRTACRRYTACCKKKRYGTMAKRTRRYKWWYSKMGKYYYRWMGWMYAAYRKW
     *********.   **    *** *.  .** *** ****** ******* * *  **

43  GTAKCGKRWKMWYCYKTMGATKKMGRYGMAGKTRTCSKYCKKRGTCTRYWWSKYKKYTAY
47  KKCKSKKRTGMWYTTTGMRRKKKMKRYRCGRKYRKMCKYYKGGTCAARYWWSKYKKTWMY
     ..*. **  ***  . *  .*** **  . * *. .** *    .:*********   *

43  KYKKKKKKKYKTKSSMMSWKYYRWCYMMYGTKYRYKRMGKTYMKCYRSKKMYWWKRYMRY
47  KYKKKKKKKYKKKSSMCSWKYYRWYYMMYRCKYRCTRMKKYCMKYCRSKKMYWWKGTCRY
    ***********.**** ******* ****  *** .** *  **  *********   **

43  WTKRKGRCRWKGAMYYKTTAGCKKTYYKGKKRYKKCYRRWMMRKRRTAGTTCYCAAATGY
47  WWKRKRGAAWTAGMYYTCCCCGKKKTTKKKKRYTGSYRRWMMGTRRYRRYCMYTTCCAKY
    * ***  . *...***.  .  **.  * ****. .****** .**      * :..: *

43  TWYRRYTKRKKGAMGMAAYTKKYYGAYWKKYYWCKKKRTYRWGTKAACTKMWKMRGRAMM
47  WWTGRYKKRKKRRMKCMMCWKKYYRRCAKKYYWMKKKRYTRATGTTRYCKMWKMRKRMMM
     *  **.****  *      ****    ***** ****  *   .:   ****** * **

43  TAKKGAMRTGWRGRKMKKKARMYKKKRWGAAAAYGGTYKWMKMYRYWCKGRMMRMKYWMK
47  YWKKTTMRYAARRATAKKKWRMYKKKGTKWMRRCAAWYKWMKCYRYTMKKRACRMKYWMK
      ** :** . *  . *** ******        .. ***** ***  * *  *******

43  CYAGKYKMWGKTKAWAARYRMMRYMYKATWACRYKRYRYYYYYKKTMKGTKCKKKARYKR
47  TCCTKYKCWKKCTWWRRRYRMMRYMYKRYWRYACKGTRYYYYYKKGAKAAKYKGKRGCKR
      . *** * * . *  **********  *    *  ********  *.:* * *   **
```

```
43          KRKKKKWAYWAATRSYWYGWKYRRYRGWYMRMYKKWTTRKWYRRYWKRRYYYRRRGGGCA
47          KAKKKKTGYWRTCRSYWYTATYRRYAAATMRMYKKWYKRKWYGRYWKRRCTYGRGRSSYW
            * **** .** : *****  .**** .  ******* .**** ******  * *  ..

43          GAYRWMRRMYYK
47          RRCRACRRMYYK
               *  ******
```

The pairwise alignments of two accessions 43 and 48

```
43              GWCGKCRMCWMRMKCCGSYKYYYKYYRRATMAYYRRGYMRMKYKYYWYWKMKATARRWWK
48              KTYRGMRMYWMRMKYYRCYKYYYKTYRRRYMRTYAAKYMAMTTKYYWYTGAKGCTRRWWG
                      ** *****   .****** ***  *  *   ** *. *****   *. :****

43              CKRYYYRWWWYWRYYWRRWWKRYWWWWWRWMRKGMSRRKYGCCYMKYRRRMRKRRGYRKR
48              SKRYYTGWWAYWRTCARRWWKRTWAWWWRAAGTKMGAGGCRYYYMKYRRRMRKRRAYRKR
                .****  ** ***   ****** * ****   . *.       ************.****

43              STYMKRRAGKWYSMARYSSKCYTMYRTCYRAWRYYYRKCTYRKYRKGRYWRWKSRYRYSR
48              GAYMKRRRKKWCCATRTCSKTYYMYRWTYRRWRYYYRKAAYRKYGGTRYAATGCRYAYSR
                .:*****  ** . :* .** * ***  ** *******.:****   **    .** ***

43              WYYWYYRRYWKYGKWYGGRCGAGATCGYKCMYYSRKRMWCGAWKRCCGAAKCRYRRYKWW
48              WYCTYYRRYWKYKKTYKKRSAWRWCGTTTYMYYSRKRMWAATWKGTSAWWGTRYRGYKWW
                **  ******** * *  *..       . *********..:**  ..    *** ****

43              KMAAAGMMSYGWMRWRRMMYRKMCAYCARWKKTTTKYRYWKSKRWYTCYYMRYKRMMYRK
48              KMTGRKCMSTRWMRTGRMMYRKMTWTTTATGKAAKKYRYWKSKRWYWYCYMRCKRMMYRK
                **:.   **  ***  *******    :   *::.***********   *** *******

43              KCAAKYYYKGRYCTKAYCRTTTGYKTMKKRMRRKKYCAAATARWRWKYTAWAYRRYYYAA
48              KARGKTTYKKRYTCKCCAGKAYKYKWMKKRMRRKKYTTCCATATGTKYWRTRYGRYYYRM
                *. .*  ** **  *. . .:  ** ********** :..::    **    * ****

43              YRYYYKKTSYYYSRYYYGKGAYKWAWGGCYKWGKCKKRCTKWMYRGKWRYRMRKMRGGAY
48              YGYTYKKCGYYYSRYYYKKAGCKWCTKAYCKWRKTKKRAAKWMYRKKWRCAAGKMRRKCY
                * * *** .******** *.. **.  .  ** * ***.:***** ***    ***  .*

43              RRKYYYKYKKYKSMKRKYRRKRRKKWKMYKGRRMRYYAKGGWGGWKARRSRGYRSKWRYC
48              RAKCYYKCKGCKSMGRKYRRKRRKKTKMYGKRRMGTTRKRKATAWKRRRSRRCRSKWGCT
                * * *** *  *** ********** ***  ***    *    .** ****  ****

43              GCKCTGWTKKGTYRMWRKRYKKCCKCAGYRCWMRKGTYYYKRYRSYSMRGKYRRKKKKMA
48              AAGMKKWKKKKWYRCARKRYKKYAGMWATAMAARKKYYYYKRYGSYSMGRKYRRKKKKCG
                ..  . *.**  **  ****** .   .     **  ****** ****  ******** .

43              GTYRYWSKCRWGCKGWYRGKATKRKCKKKKYMMKRWYMRYCTGMCCKKRRYYTTKYAYRW
48              KACRYWSKYGWTYKKWYRKKGKKRKYKKKKYMMKRWYMRTTAACTAKKRRYYKYKYMCRW
                 : *****  *  * *** *..*** *************  :.  .******. **  **

43              AYTGTWGCKYWWRATTGKGARKMWYAWRYRCTMRRWWYKMKGCWGWYMMMWYWMYYMMTK
48              WTYRYWSYKTWWRWKCAKRRRKMWYRWRYRACAARWWTTATATWKWYMMCTTWMYYCCWT
                     *. * *** . .*  ***** ****.   *** . .. * ****   ****   .

43              RRMRRGGCRYKMTKMYWKKWMKMMRYMTAKACCKAGMMKKTYGWRGYTTRMYRMRKKRTA
48              AACRGRATRYKMCKMYWKKWMKACRYMYWKMMMKGCMMGKWYRARRCCCAAYRMRKKRAG
                   *  . **** *********  ***  *   *. ** * *  *      *******:.

43              MRYKKRWACGAKCCMYAAKGKKKGMYRYWKKRGMKYCARYWGKCAYRYGYKYMTRRYMAG
48              MRYKKRWRTATKYYCYGGKTKKKTMYRYWKKRKMKYMRRYWKKSRYRYRCKYCKRATCTA
```

```
                    *******   .:*    *..* *** ******** ***   *** *. ***   ** .*    :.

43                  RRGGWKWKCARMMKAGARRTYRKYWKWYKKMWCCCMRRRKRKWMMRRGYYATAKTAMYKK
48                  RGATAKWKSWAMCKGCGRGKCRKYAKWYGGATYTTAGRRTRKWMMRRKYCCCGKKWATKK
                    * .  ***.  * *. .* . *** ***          **.******* * . .*.   **

43                  TAYYGKMMKRYWKRYRRKMWCAKYWGMARRRYRYCCTYYKYRTAAGGYWMKRYKGCKWGT
48                  CGTCTKMMKRYWKRCRRKMWYMKYWRMWGRRYRCGTWYYKYRWCWARYWMKRYKKMKWRY
                     .    ********* *****  *** *  ****    ***** . . *******  **

43                  RYMYSRRRYKYRRTTGKYRRWKRRRYYRRAGAYAKMAGGKMMKATWRTATYRSTYYYYWW
48                  RYMYSRGAYKYRRCCRKYRRWGRAGTTAAGAGTRTMTKKKMMKGCWGKWGYRSCYYYYWW
                    ******  *****   ***** *       ...  .*:  ****. * .  *** ******

43                  KMGCCTKRKYMCGYYGGYRKCCYGSMKKRKYMWYWKRMKKGCGYGYCATCGGTTCCWCYY
48                  KMKYMKKGTCMTACTRKCRKYYCASMKKRGYMWYWKRMKGRSRYKYTGCSTTACTMTTYY
                    **    .* . * .     **    .***** *********  . * * . .  :     **

43                  YYGCYMRYKKYMYMYYYYTRYKGRMMKKYRYSRRGYTRMYMMGKYYAYGKCYCTRYKRYR
48                  YYRYYMRYKKYMYMYYYYWRYKKRCAKKYRYSRRKYCAMYMMAKYYTCTKYCTYRYKRYR
                    **  ************* *** *  ******** *  ****.***:  *    ******

43                  RRYYYYWTAWGCGGKKKTWYMCTWRYRYYSYRKMYYRRRYRYRGCCGCGWTAKMRGRRYY
48                  RGYYYYTKMWRSRRKKKYWCMTAWRYRYYSYRKMYYRRRCRYRRMTRYKWWRKCGRRRYY
                    * ****  . *  .  *** * * :*************** ***      *  *   ****

43                  YRAGCWGYKKKTYKKATGASWRYACRCAWWMAYRMRYYYYRYYTWAWMTGRTGAWKKYMC
48                  YRWSTWKYKKKKYKKWKKMSWRYMYATRTWMMYRMRYYYCRYYCATACYRRWRWWKKYMA
                    ** . * ****.*** .  ****       ** ******* ***   :     *   *****.

43                  CTKRKMKTAGGCYTYYWKKYGKYRKKKYYRCYRRKRMKYGTCGCKMGKYYKGGGAMTKRY
48                  MCKRKMKCWAKYCWYYWKGTRKYRKKKTYRYYRRKRMKYKKMRTKMTKCYKRRSMCYKRY
                      *****   .    ****   ****** ** ********  .    ** * **   .   ***

43                  MRTYGMYYYKRKGTRSMKRRRYYRTRRMCCYKYRKKKKRGTGSRKMYARCKKKSKKKKRG
48                  MRCYRCYYYKRKAKRSMKRGRYYRCAAATTYKYRKKKKRAGAGAKMYRRYKKTCTKTKRR
                    ** *  ******..***** ****      *********. .. *** * **...*.**

43                  GCYWKTWKCATAWYYAWKKCACRWYGYYWKSSMKKRKGWRKKMCAKRMCSRWYCAKKMYK
48                  ATYWKKWKTCAGWTYGWKKYMMRWYACYWKSSMKKRKSWRTKCTGKRMGSRWYTGKKMYK
                    . ***.** .:.* *.***   ***. **********.**.*   .*** ****  .*****

43                  TACMKRKYYSKMKKGRGKYRRCCAMKCYRYKCAGCMYYYYKKGCRYWRGKKRGKYMWGAK
48                  YRYMKRKYYSKMKKKRKKYAAATWCKYYRYKMTASMYYYYKGRMGCAGAKKRKKYMWAGK
                       *********** * **   .    * **** :..******        .*** ****..*

43                  YRGRYKCCGYARYGYCKAMRRMKKYWWRYYYMYRYKSCRYRRCCRKTRRYRCGWKYYTKT
48                  YRKRYKTMATCGCTYYKTMRRMKGCATRTCTMYRYTSTRYRRMYRKYAGCATATKCCYKY
                    ** ***   . .    * *:*****     *    ****.* ****  **        . *    *

43                  AKGKYATTWMMYKYWRWYMKRYGKWRKKSKYGRKYRWYRRRYKGATCRRYMMGATYRGCG
48                  MKKKCRGAAMMYKYTGACMKRYKKWRKKSKYRRKYAACRRRYKSTKYGRYMMRRYYRRMR
                     * *    : *****     **** ******** ***    *****.:.  ****    **
```

```
43      AYYRYMRKKKTYWWSSKKSGWCYKMARYMTARKKYYYWGRKRRKGAATAACKYTGTKYMG
48      WYYRYMRKKKCYWWSSKKSTATCKMMRYMCTGKKTYYWTRKARKSRRYRRMKYAAGKYMR
         ********* ********    ** *** : ** *** ** **.      **:. ***

43      RCARAYCMTTTGATCAKYYYTYMWRYARWMYKKATGTGRCKGKKMYYYRKTKTGAKKKWY
48      ATCAWTYMWYARTWTTKYYYGCAWRCRRAMYKKCAKWKRSKAKKMYYYRGKKGAGKKKWY
          .    *  : :  :****   **  * ****.:   *.*.******* .* ..*****

43      RKYWGYKCCCMRKYYAKGAWYWMYATRKMAGSRYYSCWRWMYRRKRRMKRWRCKKMYYKY
48      RKYWKYKSYYMRKYYRKKGTTAMYGGRKCCASAYYSMTRWAYRGKRRCKRWRYKKMYYKY
        **** **.  ***** * .   **. ** ..* ***  ** ** *** **** *******

43      RKKRCWWGRTGGGATWKWKMGGTKGSYKCKKMRRYKKKTTYRGKGRMRWKSYKYKKKYKS
48      RKKRYWWRRKRRKWYWKWKMARYKASYKYKKMRRYKGKWWYRKKAACGTKCYKYKGKYKS
        **** ** *.     *****.  *.*** ******* *  ** *.    *.**** ****

43      SMACCYCGKGRSYSWMMMKKTTYRYRWKYMKYRMGGKKRCMTMYMAMAKRGAKRSKSMYW
48      SMRTATTRKKRSYSWCCMKKACYRYRWTYMKYRMARKTAYACACCWMWKRKRKRSKSMYW
        **  .   * *****  ***: *****.******. *.        * **  ********

43      WWKAKWKWATTMWYKAGWKRKCGGWGKTGGCRAGYTWYGGTKKSRKKRWWCCAGYKKMKK
48      WWGWKWKAGCWMWYKWKWKRKYKRWKKYRKAAGTYKATAAWKKSRKKRWWYYWTYKKMKK
        **  *** .  ****  ****   * *   . . *.  .. *********    ******

43      RYKYYYSYWAAYKKRCTAACTCRKKKMTWWYRKYTCGTRGYYKYTAAYAKKYKAGAAAMT
48      RYKYYYSYWTTYKKAGCTWMKYRKKGMKWWYAKYKMAKRRCTKYATCCTKGTKMRRMRMY
        *********::***   :  . *** *.*** **. ..*   **::. :*  *     *

43      KYWMYYYCTCATKRRGWKRGYRYYKTRGCTAYMYCGTYRMGYCTGAGYKRRTGCKKYCAA
48      KCWMYCTAYTGCKRRKWTRKYRYYKCRKYYRYACTCCTGCAYMYRGRTGGRAKYKKYYWW
        * ***  .  . *** *.* ***** *    *        .*   .    *:  ***

43      YRYKYYRRYMTATGCTATAAGYWRSMYCARYTRYRRMMYMYKSRRYRMRWYKWTRMRYKR
48      YRYKTYRRYMKRGKYCTYCRACTRGMYTGRYKRYRGAMYMYKSRRYRMRAYTTCGARYKR
        **** *****.     : . .  *.** .**.***  ************ *.    ****

43      KRCCAGKTYCWGAKWAATWCRYTKGTCCKKTTGCCGTRYRYRTRYYYCGYTKKKYYKGKR
48      KRATCKKGYTTTRKWGTAWTRCCGKCTTKTACATAAAATGTGKRYYYTAYAGKKYYKRKR
        **. . * *    **.::* *       *.: . ..:     .**** .*: ***** **

43      TGAKRWYWRRYSKRYYKRWKWRGKGTTTAGAKKGKRWKYRRYMYAGGATGATKYTATKKA
48      YRMKGWCWGRYSKGYYKRWKWAKKKYKWWKRKKKKGTGYRRYMCWKRWKRRKKTCCAKKG
           * * * **** *******  *  .    ** *   *****     .  .*  .:**.

43      RGTYMAKWKKRYRMCGGARTMYKWRKRRYKRYYYYYKKKKYTCWTYSKMKRYYYTYWYAA
48      GACCCGKWKKRYRAATATAYMYKWRTAACKGYYYYYKKKKYYSWCYSTCKRCCYACTTMG
         .   .******* . .:  *****.   * ********** .* **. **  *:    .

43      KKKTASKRSYWYKKYRRYYRWYMKKWTTRSGCRWTYATKGASRRGGKAYMRMRRRARMKM
48      KKKKRSKRSYWYKKYRRYYAATMKKWYKRSAYGWKCCWGKRGRGRSKGCCRMRRGCACTM
        ***. **************   **** .**.  *. .    .*  .*.  ****  .  .*
```

```
43          RYTGRRKYRRRRMRYTTWCYRCCTWKYYACRSMRMTCWYGRMKKMKKYYYTYRGGGAATK
48          RTCARRKYRRRGAGYKKAACRYYYWKYYRTASMRACYWYRRCKKMKGCYTKYRRKATGYK
            *  .*******   *.. . *   ****   ***   ** * ****  * .**  .:. *

43          KYRYAAAWKCGCYYYKKRAKCRGYATYRGKMWRMKYYRCTTWYRMKKKKATCGCYTRAA
48          KYRYRWMAKTATYYYKTRWKMGRCMYYRKKMWRMKYYRYYKWYRMTKKGKRYMRYYKGGG
            ****    * . ****.* *      ** *********  .****.** *     *. ..

43          GRCKRGGKKKKKKMMRYAAAGGTWARWRYMCTGWACKKRKYYKRYYCCRCCTCAAAMRMM
48          TRTKARKKKKKKKMMRYTTGTKYTGRTGCCYKRAMMKKAKYYKRYYMYRTTCAGCGCGCA
             * *   **********::.    .*     .    ** *******  *   ....

43          KKYKRKWYAGCWGKYKYKMCYRYCGTTTARGKTRKKKTTAGRWWYSKTSKYWKTCSYYRR
48          KKYGGKWYRKYWKKYKYKMYYRYSKGGATRAKYRKKKYKRKRWWYSKCSKYWKCTGTCRR
            ***  ***   * ****** ***.   ::*.* **** .  ****** *****  .  **

43          RKKYMMTWRCYY
48          RKKYMMYWAYCC
            ****** *
```

The pairwise alignments of two accessions 43 and 49

```
43              YARRTYYWRWYRRWRRRYRWTYYYWYRSAKTGACRWWGRTYRWWWMRYRWYRGRRRYKYY
49              YWRRCYYWRWYRRWRRRYGWKTYYWYRGRKAKRMRWWKRKYGWWWMRYRWTRAAGATKTY
                * ** ************* *. *****. *:   *** *.* ******** *.    * *

43              RKWMMWCYRWKKASYMRRMRMKWYRYMRMKARRYYWRAMTCAWWMRYRTRKKGGSKCYSY
49              RKWMMWTYRWKKMSYMRRMRMKWYRYMRMKGRRYYTATMGATWWMRCRYAKGKSSKMYSY
                ****** ***** *****************.****  :* .:**** *  *  .** ***

43              KRYWAAYTMRMRMYYKWYRKYKRWRRWGTTTGYGWYYAWMYRKWTRTMYMYYYRKCRWYY
49              KRYWWRYYMRMGCTTKWYRKYKRWGRWRAAARTAACYMWATAKWCGGMYMYCCRGMRWYY
                ****  * ***    ********* ** :::  .  * *   **   ****  *  ****

43              RRWYWKYYWYRRRWYMKYKYYTRKYRWYRMKYYYKCMMWKKRKYWRRKMRMYKWMWYRYS
49              RRWYWKYCTYRRGATMKYKCYCRKYRWYRMKCYYKYCAWKKRKYARRGMRMTKWMAYRYS
                *******  ***   **** * ********* ***   ****** ** *** *** ****

43              YSKAYACYRAYYRKYRKWRWKCYWRRYCAGKWMKKMCTRRWMKSKWKRMTMCYWRMSYKK
49              YSKRTCTTRCYYRKYRKTRWKAYWRRYYRRKWMKKMAARRWMKSKWKRMAMYYWRMSYKK
                ***   . *.******* ***.*****   ******.:***********:* ********

43              SGKKATTTYCYWRTKKKYKYRGGKKGKYYTAAASKGCATAAKRRACYKCTWCGKRKYYRR
49              SRKKRWKGYACTGCKKKYKYRTAKGSKCCKRWRSKAMMGGTGGRRYYKYYAYAKRKYYRR
                * **  . *.    ******* .* .*  .   **.   .:  *  **    .*******

43              TATAKRYYRGRGTCGCKMGTGCYYTKMRRYKATACCRTTKYYYKYYYYWCRSMRRTCCAA
49              YMKMKRYYRKRKKMRYKMKKAYCYKKMRRYKGYCYTGAYKYYCKYYYCWTRSMRGKGAGG
                  . ***** * .   ** ..  *.******. .   : *** **** * ****  . ...

43              YKYKGKWMYRACYYMYGARCKSKKGTKRKYWKTYAATMGKKWKRKYCKAWKWTMACARYM
49              YKYKKKWMYRWTYYMYKRRYTCKKKGKRTYWKKCRMWCRKKWKGGYTKRWKWYMWTRRYM
                **** *****  ****  * ..**  **.***.      ****  * * *** *   ***

43              MAGCCKYYRKYKMTGTRTYGWRGKRKMTKMRYKRWWTAYAAAAKRMRYKGCKRKKKKGGC
49              MMRMMGYYRKYKMGAYRCTRWGRKATACGMRYKRWTARYTCGGKRCRYKRYTGGKKKRKY
                *     ******* . *   *  * .   ****** : *:...** ***  .  ***

43              TMGAYRYWGKYRTRMGTATTATKKRAACTYWMMKKYTYGGMKRKRKGCAYRYKKSMMCRC
49              ACRTYRYWKKYRCRMACTACGAKKRGGTAYWMMKKYYYKKCKRKGKKTGYRYKGSCMMGT
                :  :**** *** **. :: .:***.. :******* *   *** *  .**** * *

43              GCKRRYKRYYGKMKGTTMRWSRAWRARGRWWYWRKRTKRMMWACAKKSCGMTTYGAWCAK
49              TAKRRYKRYYRKCGAGCMRWSRRWRGRRRWWYWRKGKTRAMWTMRKKSMACCACAMATGK
                 .******** *  .  ***** **.* ******* ..* **:  *** .  : .   .*

43              YTKRYCKRYRTYAGGYKKRTTWRGKRYYRGWRRRKRMYRYYWKMCYTKAYKYYTTKYYAY
49              YCKACAKGYRKYGTAYKKRWYWRKKATYRKWGRRKGMYGYYWKMTYGKMYKYTCCKTCWY
                * *  .* **.*. .****  ** *  ** * *** ** ***** * * ***   *   *

43              AGKKYTKRRRMYMWYYMKGWMKGRKGMCYRKYSGGAYCYGRGKYARYKARKTWRGYRWKK
49              WKKKYYKRRRCYMWYYAKRWMKKGKRCTYGKYSATGYTTRGKKCRRYKGGKKWRKYRWKK
```

```
            *** **** ***** * ***  *   * ***. .*     *  ***. *.** *****

43        RMTTRRRGTGKKKYMRRCCKKYTGGTYTAGRYACYRRYWKKRKCTCACAAAGTYAMWYMY
49        RCWWGGRRWRTKKYMRRTMKKYKTACYWWARYGYTGRYWKKRKYCGGYCWRKKYCAAYMY
          *     *   .******  ***. . *  .**.   *******   . .   .*.  ***

43        TGTRAYYRGTKWYKCAYKYCAAKKRWMMRWYRMYYTGYGSMRKRRYRKYMWYWKRMKWCG
49        YRKRCCYGKYKWYKTRCTCAGGKKRWMMRTTGMTYCACASMRGARTAKYMWYWKRMKWSR
            .*. *   ****   . ...*******   * * . .***  *  ***********.

43        RYKKWTMWYYYKYYKMRYMYYYKRYKKRMWGRKKKTGYRKRRGGMATACYKYMMYKTARK
49        RYTKWWATYYYKYYKMGYMYYYKRYKKRMWKGKKKCAYRKRRRRMGCCYYKYMMYKGCRK
          **.**   ******** *************  *** .*****  *. . ******* .**

43        KKYYYWKYKWKKKKGRWMTKGYKCTGGKTTYKYKRAAARTCKWYYRGWYMTKKGGYGGCG
49        KKYYYTKTGWKKKKKRWMCKSYKTCRKKGYYKYKRMRRRCTKWYYRAAYCKTKAATARYR
          ***** *  ***** *** *.**    *  *****   *  *****. * ..*.. .

43        YRMKWKYRRYKYKMYRKKCKKYAAKKTTAYYTGCASKYRATKGKWMMWYYYRKCTAMYYY
49        YRMGWKYAGCGYKMYGKKMKKYRWKKYKRYYKKMCSKYGRCKRKWMMWCCYGKYYRAYYY
          *** ***    **** ** ***  ** . **.  .***   * *****  * *    ***

43        KYKGCCKKYKYRYYGAARMAGTKKCGGRWRACTCAAAMCTGGTYMYKSRRRRYYCAYKKR
49        KYKKYAKKYKYRYYCGCRMTCCGTTKRRWRGYKYMWMCMKAKCTAYKSRRRAYYYCYKTG
          ***  .******** ..**:   .   ***. .      ..    ****** ** .**.

43        KGMRYKRKATARCRSKKAAMGTTGKWKYTKATTCGTCACRTRCRWYGYWSSMKKRRWKAC
49        KKMRYKRKCCTRYACKKMRCRYYKKWGCCKCAATRCTTGRYAYRWYRYWSSMKKRRWKGA
          * ******. :*  .**       **   *.::    : *   *** ***********..

43        ARKSWKWRTARAAYKTACMAKKYMMMKYSYRMKMKKKKRYYWYCYYRKRKCYRYYKKKRM
49        CRKCWTTRCWRGRYKYRYMGKKYMMMKYSYRMKMKKKKRYYWYYTCRKGTYYRYYKKKRM
          .**.*. *  *. **   *.***********************   ** . *********

43        YKKWGCKATAAKYCCGKRYAGTKTYGACTCYTTYKAYKYTYRRRYAAAWWWRRRCGRKMY
49        YKKWKMKGCTGKCTTTKRYGCCGKYRRYGYTCKTKCYKYGYRRGYMRGWAWRARYARKAY
          ****  *. :.*    ***.   .*       . *.*** *** *  .* ** * .** *

43        RYRGYKKRMRRRGKKKYYKKRCRMYYKKKCTKYYYGMKYKKRYWWSSTAGGRSKAYAGRY
49        RYRRYKKGMRRRRGKKYYKKRYRMYYKKKSYKCCCRMKCKKAYWWSSYRKRRSGRCRRGC
          *** *** ****  ******* *******. *    ** ** *****    **

43        CCMRYMTAYKRKMKAYCGYKTTKKRYRYYRTGTKYMACAKRYATKWKTSRRYKGGRMYKA
49        AYMRYMYWYGGKCKCYTAYGYKKKRYRYYRWRKTYMTTTKRCMGKWGKSRRCKCARMYKC
          . ****  *  * *.* .*  .********  ..**: :**   ** .*** * .****.

43        TGWGMAKKCMMMYYKTGAMKKKKTYWRMCRKWWKTYGYRRKTKYTKSMAAAGGKYMCCYG
49        AKWKARKKMACCYYKGAGMKKKKKYWRAYRKAAKYYTYRRKKKCYKSMMWRTTKCCYYCR
          : *   **    *** ..*****.***  **  * * ****.*  ***     *

43        TMKYRKRAKRKCKWKMYTAATKYWKRYKRRMGRYYAACKTGGGTTTACMYMSRRTCKMRS
49        KCGYGTRWKRGYKWKMYYRWYKYWKRYKRRMRGYTTCTGCKKKWCACYMYMSRGYYKCRS
          .  * .* **  *****    **********  * :.        :. *****   * **
```

```
43    RKKWMRKGGTYCGAKRMATRKRYYKKGTCYKYKSKYCTCYMRRYKYGYKYKKKKKKYRGA
49    RKKTMRKKKWYYRWKRMWWRKRYYKGTCMYKYKSKYYYMYMRRYKTKYKYKKKKKKYRKM
      *** ***   *   ***  ******    *******   ******  ***********

43    YWATCGCYRGCMCYYMGKSYMMTGTKKYRYRKWYKKKKRYMYRYRWSMRKKTWKCMYMGR
49    CTTATTAYGRMMACCCAKSYMMAACKKYRYRKWYKKKTRYMYRYRWSMATKYWKYMYMKR
        ::  .*   *.   .*****:. ************.********** .* ** *** *

43    MKKRSKKSMKWWKRMAWKAAGGCTTSRYKGRKYYYMAKYKKMRGCTYTRRRWMGSYCGAA
49    MKGRSKKSMKWWKGMRWKWGCTTACSRYKKRKYYYMRKYGGMRRYKYCRRRWMKSYYKRR
      ** ********** * ** .   : **** ****** **  **  .* ***** **

43    KGMYRRKYYCRWYRRWGKRWKRGRTCMARWRRRACGRKMYAGAKYKKYMRKYCCAAARRG
49    KKMYRRKCCYRWYRAARKRTGGRRWYMTRWRRGGYRRKMYRKRKYKKCMAKCTTGCCRAR
      * *****   ****   **    *  *:**** .  ****   **** * *   ...*

43    GGATAMWWYYGWKWRAAGAGMRSYKTCTWKMYKKKKAGRRGGKKWWRGRWRWWMMRYSWY
49    RKWCGMWWYYKATTRRRRWRMRGTKATCWKMYKKKKGCRGRRKKWWGKRWRWWMMATCTY
          .*****  . *     **. *:  ********. *   ****  *******  . *

43    CGKARWKWCGTGMKYKYYGATAAWMYRRRYYYAKRRRGRRRYYRCRRYRRGTTGGYMRRW
49    YRKRRWTATAGTMKTKCTARYRMWMYRRGYYYMKRRRARRRYYAYGGYRRRAARKYMGGT
        * **.  .  ** *  .    ***** *** ****.*****    *** ::  **

43    TKAMGARRKRYCATYGGRRYYYRGMYCRTGTTRWKGKWATSYRMKAGCWYMRATGGCGCM
49    GTCCKGRRGRYYRYYRARRYYYRKAYYGWRWYRWKKKWWCSYRMGCAYTCCRCWATATYC
       ..  .** **   * .******  *      *** **  **** ..    *. . .

43    KKYKSGKYRKACRRWWGGYKMKWMKKRGMKYRMGRRWRRYYKTKYRCRYRYAKYKKCAKK
49    KKYKCRKYRGTTGRWWAKTTATTMKKGTMKYGMKAAWRRYYKWGTRYRYAYRKYKKYRTK
      ****. *** :  ***.  . . ***  *** *   ******   * ** * ****  .*

43    YRRMYYYYRMTCMKRKRRMRMTCKRGTTATCGKKTYYKKWARRKYRRKRYRRTGYWYMMY
49    YRRMYYYCGCATMKRKRRMGMKYKRSKWTATKKKCYYKKTGGRKYRRKGYRRGACAYMMY
      *******   : ******* *. **.. ::  ** **** . ****** *** .  ****

43    AYKRYMKRRYKKRYYKYTCCCRRRTMYWRRYMKRTKTGYYYACRGTRGYCKACTRAGSKM
49    GYKRYMKRRYKKGYYKYGTMYRRRYMYWGRYMKGGKYKYYYWSAACRKTYTGTWACKSKM
      .*********** ****    *** *** ****  *  *** . . *   ..   . ***

43    MRYWYRCYCRRYMYKCCTYRKKWRCARYRRYYYKCRTTAKAGWTCKMMKYRSCTYYKSKW
49    MRYWYRMYYARYMYKTYWYRKKTAYGRYARTTYKYRKKMKGKTCTKMMKYRSMYYYKSKW
      ****** *  *****   ****   .** *  ** *.. *.    *******  ******

43    RMKKKYRKRMTKRYMYYYYKKKRRRYMAMRGYAKGGTCWATKRMMRAWKYYYKTGKTKYW
49    RMKKKYRKRMKKAYMCTYYKGKRRRYMMMRRYRKRKCYWGYKRMARRWGCYYKYATKKYW
      **********.* **  *** ****** ** * *    *. *** * *  *** ...***

43    KSWRYGRRCKTCCYMWRYTMYYKYCCCCGRWGTCYKRCCGTGCKRMKKGGTCKGAGATAW
49    KSWRYRRRYKATAYMWRYGMYYKYYYYYKRWKATTGRTTTATAKGMKKARGGKAGAMYTA
      ***** ** *: .***** *****     ** :   *   : .* ***.   *...  :
```

```
43             YGYYWKTKWSWGRKYRTTGCYYYCSKKKSRWMRRRKKMRYYRACGYKKKKKCYSYWYGKR
49             CAYYWKAKASWKRKYRGCRAYTCYCKKKSRTCGRRKKMRYYRCTKYKKKKKYYSYWTKKR
                .****:* ** ****   .*   .*****   *********.  ****** ****  **

43             KKKRWAKRGYCMRKCKYYYYWCKRRKYRYCCARYRMRYSKKACGCAYYYYYKKAGRYGKA
49             KKKRWWKRKYYMRKMKYYYYWYKRRKYRYYYRRYRAACSKKRYRTGYYYYYKKCRAYKKR
               ***** ** * *** ****** *******   ***   ***    .*******.  * *

43             YWRYMWATTYGWRRRKKCYRYWYRRYYYAKYAKYARARSYWYKYRYYWRYRWYKMGYTWM
49             YWGYMWGCCYRAAGRTKTTRYWYRRYCYRGYTKYGARRSYWYKYRYCWRYAATKMKYYWM
               ** ***.  *    *.*  ******* *  *:**.  ********* ***   ** * **

43             RSGWRRYWKTRYAYRWKRMCYYKKARAAATA
49             RSKWRRYWKGRYRYRWKGCSYYKKTRWTTGG
               ** ****** ** ****  .****:* :: .
```

The pairwise alignments of two accessions 43 and 50

```
43              GGGYYRMYCWWYWRWCGCMMRYKYRMYSKYYRMYYYRRTGTMYRRCGMYGKMYYYWGRWW
50              RRRYYRMYYWWYWRWTAMMAAYKYRMCSKYYRMYYYRRCTCMYRRMKCYTKMYYYWRRWW
                   ***** ****** . *  ***** ***********   ****   * ****** ***

43              KRRWWWRMWMRRYCCYYKRRKWRYMYYWRWMYWMYYMYRYRSKWWMYRAARWKSMRCYKK
50              KRGWWWRMWMRRYYYCYKRRKWRYMYYWRWMYWMYYMYRYRSKWWMYRRMRWKSMAAYKK
                ** **********   ********************************  ***** .***

43              MMRYWYSYRGCTAYTTCRKWGYMYMYCKRMYWYRKGTAAARYRYWKSYKMCCRWYYMWSA
50              MMRYWYSYRRYGGYCCYAKWKYMCMYYTACYWYRGKWWWWGYGTAKSYKMTARWYYMWSW
                *********   .*    ** ** ** .  ****       *   ***** .*******

43              KRRKYRARMKGYMWKRYYRRYWKRYYRKGATYWMGRMYRKYKRWMRKKAGKWRWKMRMWW
50              KRRKYRRRMKRYMWKRYYRRYWKATCRKTCCCAMRRMYRKTGRWMRKGTCKWRTGARMWW
                ****** *** ************   ** .   * *****  ***** : ***   ****

43              RYRWKRRMRMKRYAYWRCSMKRMRYCTKCKYYWGWYRYYYKKTCGKRMTYWWSRCAASGY
50              RYRWKRRMRMKRYWYWRGCMKRMRYAATYKYYWAWYRCTYKKYYKKRCYYWWSATRCCRC
                ************* *** .******.:. ****.***  ***   **  ****   ..

43              CAYKKMKYYKYSRKTMCSAWRKYRRYAACYKMYMRRKKKYYKTCAGRRTTSRKRYCTRTM
50              MCCKKMKYYTYSRKCMYSRWRKYARYGGAYKMYMRRKKKCYKKYGAAAACSRGRCYWGWM
                 . ******.**** * * **** **...********** **. ..  : ** *     *

43              YKRAGWKYMYGATKKKKKYRTWGMRYMYKRAKTGGYRKRKKTTMRKSMAAAKCGCTYRRK
50              YKRCKWKYAYRRWKGKKKYRKWRARYMYKRGKYKKYRGAKKCWARKSMMWRKTTTCYRRK
                ***. *** *   * *****.*  ******.*   **  **   ****   *    ****

43              TKRKKKTKKMKYYWKRYKRRMYKKTGGMKSRYRMRRMWRKMRSCKGGRKCTKGRRWRGTM
50              KGGTKKYKKAKYYWKRYKRRMTKGYTKMKSRYRMRAMTGKCRSYTRTGGMKKKRRWRKWM
                .  .** ** *********** *    ******** *  * ** .     .* ****  *

43              YCGRYAKCGTTRKCATRRYWCYRKYTKWKKKWMTTCKMMTAKKMTRKGRKWYGRGKKRMM
50              TTAGTRKYRCGRKSMWRRYWYYRKYWKWKKKWMYYMKMMYWKKMKRGKGKAYRRRKKAAM
                  .   *    **.  **** **** *******   ***  ***.*   * * * **  *

43              YKCTCYGGYKAMKRWKKTRWTAAAAKMGTRYYATGMKYYRYWSWYRGAACMMMGKKYRAA
50              YKTCSTRRYKWMKRWKKKRTAWMRRKCAARYYGCAMKYYRYWSWYRRRMTACMKKKYRRR
                **  .   ** ******.* :    * .:***. .***********      * ****

43              CYWMMWWGARTCYGAYGRAGYMRRYGGTAKRCTRASKTTGRKKGCKRYYRWMGYKKTKWM
50              YYWMMWWKTRCYCSCYRRMRCCRRYKACGKRTWGWSKGCCRTKKYKRTTGWMKYKKYKWA
                 ****** :*   ..* *    *** . .**    **   *.*  **   ** *** **

43              TGYKKSRKARKMRMYMRRWAYKWKYYMCCRGRYKKKKWKYTKKRYAAKKYRYYRYYYMKG
50              YCYKKSRKGRKMRMYMRRWWYKWKYYMAARARYKKGKWKYYKTRYRMKKYRYTRYYYMKR
                  ******.********** *******..*.**** **** *.**  ***** ******

43              YKKKRRYRWMGAKAWYWGYKRARRRRYRKWRKMKKRRKGMRWRWRKMKYCCKGKKWRKYC
50              YKKKRRYRWMKRKRTTAKYKRWRARRYRKWRKMKKRRKRCRWRWRKMKYYATKKKWRKYM
```

```
           ********** *    *** * *************  ********* .. ******

43         RRRRRKWKYCMRYYYRRYWMMGMRKKYCAWYMYTTSGTKWGRRYMYWRWARYGACRCATY
50         GGRRRKWKYACRYYCRRYWMMKCGKKYYMWYACGYCKCKWKRRYMCTRWGRYTGTATGKY
             *******. *** ******   ***  **     .  ** ****  **.** .   ..*

43         KSRRCYRARCKMAAARTTGKWAMGTKTKRYWYWRMKYWYYRAYKTGGWYRMKKKGCTTAA
50         KSRRSYGGGMTMCGRRYYKKWGMAAKAKRYWYWRMKYWTYRRTKWRKWYRAKGKKYCYWT
           ****.* .  .*.. *   **.*.:*:*********** **  *   *** * *      :

43         WKYTCGCTRRKCTKKKKATKTRKSYCWMMAAKRKGYRKYKRCTYKRYYRMYSACKGWTAG
50         AKTYMRYKRGGAYKKKKRYKKRKSCTWMMGTKRKTYRKYKRTCYKRYYRMYSRYKRAYRA
            *      .*  . ****  *.***  ***.:*** ******  *********  *     .

43         TYYYYKKWRTKYWRRKRRYYKYWGTSYWYYRRRYCGRRYYTTMRSGTTKTTRRYWKGYKG
50         CYYYYKKAGYKYWRRKRRYCKCTACSYWYYRRRYYKRGCTYKMRSKWKGWYRRYWKACKA
            ******   ********* *  . *********  *     .***  .   *****. *.

43         AYWRATARKYYKYYSWKRCGTMCCKYWKRMKCGYWKCAAYYWRWYYYYKKWKMRYMYYYC
50         GTWRCYRRKYYKYYSWKRTACCTAKYWKRMKSRYWTSCTYYWRWYYYYKKWKMRYMYYYY
           . **.  *********** .    .*******. **...:*******************

43         KRYATYKMTRMKKRGSRKKRYKTMYKYMMKYRYMWKYRYMYKRRYRRKYYYYWKGYWCKG
50         KRTWKYKMYGMKKRKCAKKRYKCMYKYMMKYRYMWKYRYMYKRRYRRTYYYYWKRYWYKK
           **  .***  **** . ***** **********************.****** ** *

43         WMKWRYRYGYKRWYYWSSMKRWKGACAWRKRMSWYKRYKWYGTACKKWSKRATKYKYRMK
50         WMKWRYRYSYKRWYYWSSMKRWKRGACWRKRMSWYKRYKWCRYRYKKWSKRMYKYKYRMK
           ********.*************  ...*************      ******  *******

43         RKYYKKKYYCYYKCKGMKCYRYMRKYGKCYKRYKWRYYKYKAGGCCMAAAKWYYRATMWC
50         AKYYKKTCCTTCGMKKCKYYRYMRKYRTYYKRYKAGYCKYKRKRTSMMRRKWYYRTGMWY
            *****.       *  * ******* . *****  * ***     .*   *****: **

43         KKRATMTYRMYYKMRSKYYRGKMGGAAGRKKYSYYRWWYSAYSYGGCMMKYKCRYTMRKR
50         KKRTCAYYRMYYKMRGKYYRAKMTTTGRGKTYSYYRWWYSCTSYKRMMMKYKSRYKMGGG
           ***:   ********.****.**  :.  *.*********. **   *****.**.*

43         YYKTRMRYYMKGYYGYKKYWGYRRCCRKWKKTACAKACCRRRYRRTAGTMMYKRRRKKKR
50         CYKYRMRYYMKKYYKYKKYWTYRRMYRKWKKCRYRKMYMRRGYRRCCCCMMYKRRRKKKG
            ** ******* ** ***** ***  *****    *   ** ***  .  *********

43         CGCYAKWCKATGKRAYRTWRYRRYMATTKYGKRAKYKRYKSMMTGCGKWYRYYKRKKMYY
50         MKYYGKWYKGCAKRWYRWWRYRRYMGCAKYKTAGKYKRYKSMMCKYKKWYRYYKRKTMYY
              *.** *. .** ** *******. :** . .*********    *********.***

43         MWSKRWWKWYKMKWYRKKRRAGTWKKARWKKYKYRYRYMYYGTKYYYKKTKGGGGYWWTT
50         MWSKRWWKWYKMKWCGKGRRGAAWKKTRWKKYKYRYRYMYYRWKYYYKKYKRAKKTWWKK
           *************  * **..:***:*************  ****** *  .   **..

43         YWMYYRWKMMARYKWTWYGAGAGGTWKKGKTCYTYCMYYYATGAGKKTMKWGCKMARTKT
50         YWMYYRWKMMRRYKTYTYKWKCRAATKKKKWMTYCGMYYTCCKTATKWAKAKYKMCGAKC
           ********** ***    *   . .: ** *      ***  .  :..*  *   **. :*
```

```
43    RYMKWGWKKWKCRGACAKMYKACCCGGACTGYRTYMWYRWRKRWMMRRMWYRMRWYCYRY
50    RCCKTTTKKWKYGAGTGKMYKMYSSRRRMYSYRKTMWYGWRKRWMMRRMTYRCGTTTTGC
      *  *   ****  .. .****  ..     .**. *** ********** **

43    KGKWMWKKKKRKMWYYKRGRYYTKCCCKYYWRYGCKKKYMYWRRRGKYRRRCYRRYKKGC
50    KKKWMWKKKKRKMWYYKRKRYYKKYYYKYYWRYKMKKKYMYWRRRKKYRRRMYRRYKKSY
      * **************** ***.*   ******  ********** ***** ******.

43    KAMKYWYRCAGGYYMYCGCMRRWYMGYRKCCTMRRWKKYYRKRTYWKKRRRYRRWKRMMR
50    KCMKYWYRAGATYYMYTAAMGAWYMKYRKTAGMRRWKKYCRKRKYWKKRRRYRAWKRMAR
      *.******... **** ..*  *** *** . ******* ***.********* **** *

43    KWCKTKYWKRRRATSWRYRCYKKRKYYRMRTYGMWTYGCCCARKWSYKRKMCRKAGRRGY
50    KWTKYTYWKRRRWKSWRYRYYKGAKYYRMRKCTAAAYATYTWRTWSYKRKMGRKRRAGRY
      ** * .****** .***** **  ******.    :*.    *.******* **     *

43    YWKYKKYRWKTGRKYYRKYRKKYCRYMRRYKKTTGWKSKKKKCRKKRYKGMYGGGGYKWK
50    YWKYKKYGWKKKRKYYRKCRKKYYRYMRRYKKWWRTKCKKKGMGKKRYKAMYARAAYKWK
      ******* **. ****** **** ********    *.***   *****.**. ..****

43    ASWMMKKKYYYRKGMYRWSMYCAKRWKKTMYMKTRSKKWWTTGKKWKRKKYWKGGRRYSW
50    RCTMMKKKYYYRTKMYRWSMYTRKRWKKYMYMKKRSKKWWKKKKKWKRKKYWKKAGAYSW
       . *********. *******  ***** ****.******.. ********** .  ***

43    WRKYKYYRRYRMRSKGWKTYYWKRMGRWRWWTKYRYMYRASMYYRKKRWRRRRRCGKRRR
50    WRKYKYYRRYAAGSTKWKKCCWKAMRRWATWKKYRYMYGMSMYCRKKRTGARRRYRTARR
      **********   *. **.  ** * **  *.******  *** ****   ***  . **

43    WRYKGYCCAATYRKRGTCGCCKKMGRYKAKWYYYYYRSMCCRRAGYYRRRMKYYKSKMWR
50    WRYKKTTSMRKYRKRKKMRYYGKMKRYKGKWYYYYYRSMATRGTTYYRRRMKYYTSKMWR
      ****   .  .**** .     ** ***.**********. * : *********.*****

43    WKRMGKTKKSKKGYKCWCRRAKKSWKYWMKWKYCAAAGYWCKYMMYYMGTMCCCKRRYYG
50    WKRMKKKKKSKKKTKYTAGGMKKSWKCWMKWKYYWRMKYWAKYMMYYMKCCTAAKRRYYR
      **** *.*****  *  .   ***** ******     **.*******    ..*****

43    MATCTYRKRMCMWTMYWWCYMRGCACKWRYRATCACTTACGTRMWWGGKYKMKKKAGMKY
50    MGWYCYRKRMTMWYMYWWYCCRATGYKWRYRTCTGTWYCYRGRMWWKRKYKMKKKTKMKY
      *.   ***** ** ****   *. . *****:  .   .   ****  *******: ***

43    MKTGYMGMKKRRMCTAMGMAMYRRKYYGRRYKKRYRTRRRWYYKRKGMRKKKKRMRTGKK
50    MKWAYMKAKKRRMTCMMAMGMYRRKTTKRRYKGGCGWRRRWYYKRKRMGGTKKRMRCRKK
      ** .** *****   *.*.*****   ****     ********* *  .*****  **

43    YAAATARYWYRYYYKKGKYKKGGATRGTGTKRYYCYKARYWKTYKYSYMGRCA
50    YTCCATRYWTGYYYKKKKYKKKTRKRSWSYKRCYAYKRRCWKYTTYSYMARTT
      *:..::***  ***** ****   .*. . ** *.** * **  .****.* :
```

The pairwise alignments of two accessions 43 and 51

```
43              RYRYRYWRRYYGWCKWRRMKKGTMYKGRTYTCGTCTMKRYYSRYSTRMYYWRRWYWYAKK
51              RYRYRYWRGYYKTYKWRRMKKTYMYKSGGCAYAAYYMKRYYSRYGWRMYYTRGATWYRKK
                ******** **   *******  ***.   : .:  ********. **** *   ** **

43              RWRTRYRRSGRYWRWKSAWYSYRRMRWYYRRRTYWKWYWWRYRKMYYKRWMYWMRRYGKC
51              RWRYGYRRCKRYWRWKSMTYSYRRMRWYYRRAKYWKWYWWATRKMYYKRAATWMGGYRKM
                ***  ***. *******  ************ .*******  *******   **  * *

43              SMWYYYMCCRGKRWYRRRYWRMYRKCRWKKYYRKTCTGTRKTTATRRKYYYYYMMYTKKA
51              SMWYYYMTTRRKRWYRAGTARMYRKTRWKKYTGKYSWKCRKYWWKRRKYYCCTMCYAKKR
                *******  * *****    ***** *****  * .   **   .*****   * *:**

43              RWKSTTARRRMRKSRMYYYWRWWYRWRWWRRMRRMSRYRYYWGKRSSKMYRYWMYWRTCY
51              RWKSCCGRRRMRKSRMYYYWRWWTRWRWWRRMRRMSRYRYYTKTGCCTMYRYAMYWRWST
                ****  .**************** *****************  . ...**** **** .

43              SYYKCRKSKYRRYKRTWSRYYKYKTAYRTYYGKGRKKKGKKWRRRRWYRWKKMRRKKYMR
51              SYYKSRKSKYRRYKRCWGRYYKYKKMYRKTTTKKRKGKATKWRRRRWYRWKKAGGTKYMR
                ****.********** *.******. **.   * ** *..************   .****

43              RAKKKKYCAAATAYWMKYRYYWWYWGRKAAGTYKTMYSTKGRYGKYWYRKWKRRYRYYKY
51              RWKKKKYYTCCATYWMGYGYTATTTRRKWRKKYKAMYSYKRRYKKYAYRKWKRRCRYYKY
                * ***** :..::*** * *      **   .**:*** * ** ** ******* *****

43              MWKYGKRRRRMRMMRCRYAKRRWKGTATKKKRRSWRYGRCYYWMMWYRMRMYAYRYGYWM
51              MWKTKKRRRRMAMMAYRYRKRRWGKWMYKKKRRSWRYKRYYYWCMWYRMAMYCYGYRTTC
                ***  ****** **  ** ****     ********* * *** ***** **.* *

43              WWMKTACRWSARWKRKMKRKKKTYCGRRWYYWYAYKTKKRKAYRCGKKGYRRKKKKYYRY
51              WWMGYRYRWGCAWKRKMKRKKKCTTTRGWYYWYTYKKKKRKWTGYKKKKYRRKKKKYYRY
                ***    **.. **********    * *****:**.****     ** ***********

43              RKYRMYRKRYMRYRCTYKMKAGTRMKKYWKYYRYYYMYRKKKYKKKRYRRGAGKYTKARK
51              RKYRMYRKRYMGTATGYKMKTAYRMKKYWKYYRTCTMYRKKKYKKTRYRRRTAKCYKMRK
                ***********     ****:. **********   *********.**** :.*  * **

43              RYRRKYRRRMMYGKKYYMKRRYYRTRKKSKKCGCYRCCCCCCMRYYRKARYRYWGAGARY
51              RYRRKCGRRMMYSKGYYMKRRCCGYRKKSKKMTTYRATTYGTMGCYRKWRYRYWATKWGY
                *****  *****.* ******    ******   **.     *  *** *****.:   *

43              KYMKKRRTCAKRKKYRWWSSCAYAKMYYKKAKRACTCWRGKKMKRRRGAATKKAAAMGYG
51              KCMKKRRCTGGGKKYRWWSSMCCMKMYYTKTKACMCTWARKKMKARRSRRYKKMGTMRYK
                * *****  .  ******** .  ****.*:* .   *  **** **.   ** .:* *

43              RRTRYMWCKKRYYYYCTYWTGRTRTKRYAGCMYKMYKATGWKKYYMYYYKTGTAMKKTWW
51              GAWATATYKKRYYYYYGYWGAAARCGRCRRMMYKMYKCWKWKKYYMYYYKGAYGMKKKAT
                        *******  ** . :*  *    ******.  ********** . .***.

43              WRRMKYGTCCKMYKTRRCYKRCAYYKYKKGRSMCKCGAWKKYCCKTTRTMKKRSWRRMGR
51              ARRMKYKKMTKMYKCRRTYKGYGYYKYKKARSMAGTRGAKKYMYKAWRCMKKRSWRRMKG
```

```
                     ***** .  **** ** **  .******.***.   . ***  *: * *********

43                  KKCAAAWKRRKKYKWKKMKWYYTCYKRKRWKSMKRWKWRKYKTMGRAWYKTACWSKRKTW
51                  TKTCGTWKRRKKYKWKKCKWYYKACKRKRWKSMKRWKWRTYKKMKRCWCKYRYWSKRKCT
                    .* ..:*********** ****.. **************.**.* *.* *   *****

43                  CATYMMKYSYRMYWRKMKKSKYYYYWKYYKCYRYTCGGYYKKYYKKKWRKKKRKYGYAYY
51                  SRYYMMKYSYRMYWRKMKKSTYYYYWTYCKYYRYYYRKYYKKYYKKKAGGKKRKYRCMYY
                    .  *****************.*****.* * ***    *********   *****   **

43                  GKKRMCTTGTYGGRGSKTKSRKKKKKRRKKMKRRYCKKMMYRKWKTAGKASRRKRKYMRY
51                  KKKRMACGAAYKAGASKCKSRTTKKKRRKKMKRGYTKKMMYRKWKYRRKGSRRKRKYMRY
                     ****.  .:* . .** ***..********** * *********   *.**********

43                  YKKKKKTCRKATKYWYTMTCAKTATRYRYTTGKYKYTCAGYTTRYTRMKKKWKYCRKWWR
51                  TKKKKKYTRKMYTYWCYCYTGKYRKRYRYCCKKYKYYYMKCCYGYWAATKKWKCMGKAWA
                     *****  **  .**     .*  .****   ****        *   .****   * *

43                  KYYWYWYRSKWKRTCTARWGWKTYRCYYMYRMYRKKYRWRRRKRCGKYRYYYYKRCRMYY
51                  KYYWTTTGSKWKAYYYWATKTKKYRYYYMYGMYRKKYGTGARKRYRKYRYYYYKATRMYY
                    ****    ****         *.** **** ******    ***  ********  ****

43                  KCCAYCACRTCKYRMRKKKYKRAAAKWYYRKKYTYRMWGCYYYKMGKRGTCTKYRMKYAY
51                  KTYMCTRYGYTKYRMRTGKYGRMRRKWYYRTKYYYRMWKYCYYKMKKRTWYGKYRMKYRY
                    *          *****. ** *   *****.** ****   **** **    ****** *

43                  RARGWWYSKMKYKYRTSYRRWMYYYYKWTTTWYRMRARYMKYTGKKKSKYKYAWKSMKAT
51                  RWGRWWYSKMTTKYRCGTRRAMYCCCTTGAGWYRMRRRTCKYYKKKKSKYKYMWKSMKRC
                    *   ******. *** . ** **   .  : ***** *  **  ******** *****

43                  CCYMCYGRCCTKKKKWKMYKYAGKMSRKCAYRSYCGYKGKRRWRWKTRMRMGMGRWRYKR
51                  YTCCYCRRYYWKGKKWKMYKTRRKMSRKYGYRSYYRYGKKRRWRWKWGMRMRCKAARYKR
                           *   * *******   ***** .****  *  *******  ***     ****

43                  RYWKKRCCKCAGCGWKKGTYYRYMYRYMKYYTKKYYTYTRKWRTKRKMYYYRGRMAAMAT
51                  RYWTKAYAGMWAYRWKGKYYYRYMYRYMKYYKKKYYYCCRKWRYKRKMYYYRTRMGGMTA
                    ***.*  .   .  **   ************.****   **** ******** **..*::

43                  ATKYRGMYASKAGGTKYRWKYRYCKRYRGYYGAKYYCRTCGCARYMAKKWKYCWKKKWMK
51                  TCKYRTCYCGKCRAAKYGWKCGCYKGYRRYCKTTYYYGKYRYWGCCGKKWKYYWKKKWMK
                    : ***  *..*. .:** **    * ** *  :.**  .       .***** *******

43                  MMRYMMKKKMTCKKTTAAMYACTKKYMRAAKKGATRRYKKYRGTTTRYMKMYYRYWCRYG
51                  ACRYMMKKKMGSKKGKWMMYWMYKKTMRGMKKARWRRYKKYRKYYCRYMKMYYRYWMRYK
                      ******** .** .  **   ** **. **.  *******    ********** **

43                  KYRAACMKKYRYWMMWWRYKTCCYAYYMRTASYYYYTTRYKGSCKMMKRYKRCKMYRYKY
51                  KYRRMTMKKYRYWMMWWRYKCYTCCYCCRKRSYYYYCWRYKRSYKMMKRYKRMKMYRYKY
                    ***   **************    .*  *. *****  *** * ******** *******

43                  RKRRWKKGRYTRYYCYKCSSRKKKMMKWTASYARWYMYRGTWGKKKYYYRYMCYYYKYRK
51                  RKRRWGGRRYYATTYYKYSSRKKKMMKWKWSYRGWYMYRAKWAKKKYCCRYMGYCYKCGK
                    *****   **     ** **********. **  *****..*.****  *** * **  *
```

```
43    KYKCTKYMRRYKKYTYKYYAAGYTKKKKKRYKSSKRSYCTGKKYRYRKCGTAKKRYMWYR
51    KYKMWKYMRRYKKTAYKYCTMKTYKKGKKRYKSSKRSYYYAKKYRTAKYAGGKTRYMWYR
      ***  ******** :*** :    ** ***********  .****  * . .*.******

43    YMRYGRTYGTCMRMKRSRKMYCRYMTKWAKTWTKMCKYGGRMWRGCMSKKKYYRRCTTTG
51    YMRYKAYCKCYMRMKRSRKMYYGTCWTARKKWYKMYKYKKRMWRSYMSKKKYYRGSCGAA
      ****       **********     .  *.* ** **  ****. ******** .  :.

43    KWYRATWYRAKAKCCCTATCGCRAKRGTTKGYKKTTKKKMGTARGACCKRRKYRYKRKKA
51    GTCGGCWYRTTGKMYYYWYMRYRRKRAAATRCKKACKKKMKYRRATTTTRAKYRYKAKKG
          . ***:..*         * **.::.  **: ****   *.:  .* ***** **.

43    TYRYYKKKSMYYKRYGYYRYWWWYKGCWRAAKRWRYKKKKAAYCRYRMMYRKWYRCTTYK
51    GYRYCKKKSMYYKRYRCYAYWWWYKKYWGRGKRWRYKKKKTTTMRYRMMYGGWYRMCGYT
       *** **********  * ******  *  .*********::  ******  ***   *.

43    RRYRRYYYYYKKKWARYAYKYKYYYYWATSYWYMRRWYMKSYRSKWRRYWWGTRMYRMYW
51    AACRGTTCTCTKKAGGCRYKYKYYYCTTCSYWYMRAATMKSYRSTWRRYWTAWGCTGATA
         *      .** .   *******  : ******   ******.***** .

43    RKKRYKMYYRYYMKTGYRRAYMWKCGCMYWWGGCWYMKTGARYYYRMYYCKCAWRTTTMA
51    RKTRYKMYYRYYMGCTCRRWYMWKYTYMYWWKRYTCCKCAMRCYTRCYYSKMRWRCYCAT
      **.**********    ** ****   ****      * . * * * **.*  **    :

43    RGRCRWWWTGYKMKKWRRWYMKWWKMKKCRMTTRCGTKKKCTAYKYTYYCAKCKKWYRYY
51    AAGTRAWWYKYKMKKTRRWYMKWWKMKGTRMCWAYRYKKKYYRYGYCCCYRKAKKWYRYY
       .  * **  ***** ***********  **      ***   * *     *.*******

43    KRYRTMGCKTCTGGAWWACAYAKMRRKTRYRYMRKYRKWKGCYKKKMATTARGAGARTTG
51    KRYRKMATKCAAARMWARTGYGGAGATKRYRYMRKYGGAKKMCKKTMMYGWGKGKRRYYK
      ****.*. * .:.  *   .*.    ..********   *   **.*      .  *

43    KRGGCWKCCCCAGKMKCGTCRGACAMKKKAGARRAYTKYCWGYMARKRWRKKKRWMMYRS
51    KRAATWKAYMYRAKMKTAAYGAGTGMKKKGCGRRMYKKYTTRCAGGKGWRKKKRWMMCRC
      **.. **.    .*** .:  .. .****. .** *.**     . * ********* *.

43    TRRRMRRRGYCTTRCGGGAGTYYRWKGWKMMMYTCGGKRRRYYWYWCAMAKRRRYYYRWY
51    CGRRMGRAKYYCGAYRKTMAWYYATKRTKMMMYKTAAKRRRYYWYWYTCTKRRRYYYRWY
        *** *  *         . **  *  *****. ..********* : :**********

43    TTCAYYYYATYKMKRMRRYRMYCGAAKGMYGTTRRKRCCKYRRWKMKRRRWRMKYGKYYK
51    KWYMYYYYRAYKMKGMRRYRMYYKRRKKMYRAWRRKRTTKYRRWKMKRRRWRMGYKKYYK
      .   **** :**** *******    * ** : ****  ************** * ****

43    TKRTGGGAKWWKWRTKAMRYWKKTCTYYMGTGMRMSMKKRTWKRMKMRGTAWMYYYYMRY
51    AKRKRRKWKWWKWRKTMMRYWKKYTCTYAKWRMGCSMKKRAWKRMKMGAYRWMYYYYMGY
      :**.    ******.. ******    *    *  *****:****** .  ******* *

43    MYYYKRYYMRKYKMRKKYRYSKKTKGYKGYRYMMYRYMYTARRRMYYYWAACWGYCTKKY
51    MYYYKRYYMRKYKMRKKYRYSKKCKKYKRYRYMMYRYMYGCRRRCYYYWMRYWATTCKKY
      *********************** * ** ********** .*** ****   *.   ***
```

```
43              WMRKMWRYRGY
51              WMRKMWRYRSY
                *********.*
```

The pairwise alignments of two accessions 44 and 45

```
44                 KMYKSYYYYRRRYMYTCMMYWWRRWRCYWYYMRRGCKRYSRWCKARSWWGAYRRWYGMRY
45                 KMYKSTCYYRRAYCYGAMMYWWRRWRYYTCYMRRASKRYSAWYKRRSWWRRYRRWYRMRT
                   *****  **** * * .********* *  ****..**** * * ****  ***** **

44                 RKCKKCGGYMGAMYYWMYYWCAKTRYMGCYGKRMKMKCYRSRKYCCKKMMKYMKKWYYKT
45                 RTYKKGRKYCRMMYYWMYYAGMGCGYASMYAKRMKAKTYRGRKYYYKKCAKYMGKWCYKG
                   *. **   *   *******      * . *.**** * **.***  **  *** ** **

44                 TCAARKKYRKYRYRSYRYRYRCGGCRYYARRCMRGACCYYRWYYMRRYCYYAAGYWSYYR
45                 CYGGRKKYGGTACASCACRYRYKRMRCYMRRYCAACTTTTAWCCMARYSYYWWKYWSYYR
                     ..****      *   ***    * * **   ..     *  * **.**   ******

44                 CSYKWRKYWYYRKTATAAWRCYKRTKCKWYYYWWYSMWTGCGGMKMRKKYYMRYYRMWKR
45                 YGYTWRKYWYYRTAWCGGTRAYKGKGYKWYCCATYSMWCATAAMKMAKKYYMRYTAMWKR
                    .*.********.:  .. *.** .  ***    **** . ..*** *******  ****

44                 KCYMGSKMYRRRKWYMRGCARYWRMKTRMMYMWRRKAAYTTRYRWKYRWKWGTCRYKYRK
45                 KYYMKCKMYRRRKWYMRRTRRYWRAKWRMMYMWAGGWWCYCGCRWKYRWKWAWMATGCAK
                   * ** .***********   **** * ******          ********.       *

44                 KYWRYRCMGGMRKMKTWMMYWRRKRRKRYRRWWYYYYKWCYRKWYRRYRYYYYAATAKYR
45                 KYWRYGGMKRMRKMKYWMMYWRRKRRKRYARWWYYYYKWTYRKWYRRYRYYYYTRCCKYR
                   *****  *  ***** ************* ********* *************:  .***

44                 YRKYRYSRRTTKAKRYYGYYCTYGWAYYRYYYRKRRCRMCGTWMRWYRYWKWYRMRYWWR
45                 YRKYRTSRRCYTRKRYYTYYAATRWRCYRCYYRKRATRCMKAWMRACRYWKWYRMRYWWR
                   ***** ***   . **** **.:  *  ** *****  *   :***  *************

44                 TYYYYKRYSAWSWYKYYRRWKSYMMMYWKRSATAYYWYRRYKMWKKKSRYKTTCKRWRYR
45                 CTYYYKGCCMWSWYKCCAAWKSTAMMYWKRSWYGCTTYRGYKMWKKGSRYKYYYKRWRYR
                     ****  . *****    ***  *******  .    ** ****** ****   ******

44                 KMMYYYRTTAGGTGYKYYKWYAKRGTAYGKYKKKKYRYKRKYCKKYRRRRMMRTGGRRKW
45                 KMMYYTAAGCSRYKYKYYKAYCTATATCAKYKKKKYRYKRKYTKKYRRRRAMRCAARRKW
                   *****  : ..   ***** *..  :: .************* ******* ** ..****

44                 CYYMYKYAKCYKSYWYKMMRGYYYRSSWWKCCYCKTKYMAMRYGYKYYKRKKYTWGCYKK
45                 GTCAYKTRKYYKSYWYKMMAKYYYRSSWWKYMTYKGKYMRCGYRCKYCKRKKYWTKYYKK
                       **  * *********  *********    * ***   *  ** *****    ***

44                 SRYCRYRMKGSRGRMSKKCTRRKKRRKKYSRKRGGRKWWYWMRATSRYKAGCTTGGGWAC
45                 SRYARCAMGKSRRRMSKKYKGRKKRRKKYSRGRKRGKWWYTMAMYCRYKCRACCRRKWGM
                   ***.*  *  ** *****  .********** *   **** *   .***. .     *.

44                 GWWKMYWTRWRRSYCGKRWRKTMWRKTKGRYARRKRKYCGCGSRRMMYMWSYKGKSRRWM
45                 RWWKMYWCRTRRSYTTKRWRKCMWRKYKTGTWGGKGKYAATTGGRMMYMWSYKKKSRRWM
                    ****** * ****  ***** **** *      * **..  . ********* ******

44                 MKKMWWYYKKGRCCCARSKGRRRCTKTCWKKCCCAYATMKSYKMWCKRMRKKKYYRWSKK
45                 MKKMWWYYKKRGYYYGRSKAARGYGKKAWKKSMYGYMYMKSYKMWAKRMRKKKYYRASKK
```

```
            **********     .***. *   *..***.  .*  *******.********** ***

44          YYRGRYYWKMKAKKTKAWSACGTKSYMYKYWSKRMYYKCAMYKTTYRYKRKYKKMCWYKA
45          YYRRRYYWKMKRKKWGGASMGKWKSTATKYWSKRMYYKMCMYKKWTRYKRKYKKMYTCKM
            *** ******* **  . *    **   ********** .***.  *********   *

44          KSYTYKRGATCKACWYAYKWCTRATKMRRKKYGWCYKACTTGCYYWKGATKRRTSCCTTS
45          KSYYYGARWGAKTGWYRYKATCGTYTAGRKKYKAYYKRTYYKMCCAKAGKKRRCGTTYAG
            *** *     .*: ** **    : .  ****   **         *...*** .   :.

44          CYGWKKRWMMMRRRKRRYATTGMYTMCKRRKTWKRYMMRKKYRMSRYATMCKGATTCCYR
45          SYRAKKRWMMMGRRKAATRYYRAYYMYKRGKWAKRYAMAKKYRMCRYGWCMKATCCYYCR
            .*  ******* ***        * * ** *  *** * *****.**.   *.:     *

44          YKKWYYMAGRGYKGRCYKGGYRRGYACKSKYRRWKKKYYYYRYRGACWYRRCYYKKRKRW
45          YTTWYYCGAAAYKKGTTKRKYRRAYGYKSGYRRWKKKYYYYRTRATAWYRGTTYKKRKRW
            *..*** .. .**    *  ***.*. ** ************ *.:.***   *******

44          CACYAKRKKAYTRTTYWYCRMYKYKKKYRRYYYWMKKYYKYYYSYYKKGKKCKKGGTYGY
45          YGTCCKRKKGYAGYYYWYYRCYTYKTKYGRYCTAMKKYYKYYYSCTKGKKKYKGKKCCKC
             .  .****.*:   *** * *.**.** **   **********  *  ** *

44          ARTRYCWTTCTYYKRKMAKWYYWCYYRYYKRTCGKWYCAAARAATCGTTRYYGCAATKKY
45          RACRYATKGTWYYKRKAWTWYYWACCATTKRCTAKWYTTGCRMMYMRAARYYAATTCKKY
               **. .   *****  .****.     **  .*** :..*     ::***..:: ***

44          KMAAACCYTTYYRMRWYYSTKYWKGTARGWMYWRRTYWYKRWKCGKKKSRWAYKWWYCRM
45          KMRRRYYYACYCAMRATYSWKYWKTCGRKWMYTARYTWYKGTKMAKKKSRWCYKWWYYRC
            **     *: *  **  ** ****  .* ***  *  ***  * .******.***** *

44          MYRRTTTYRCYKYKTCGKKKKYYKRYKKCWMYRYRKYGWKMYYRKTRYYSGYCYYMSRKY
45          ACGRACAYRYYKYKAAATKKKYTKRYKKTTAYATAKYRWKMYYRKARYYGKCYTYMSRKY
               *: :** ****:...**** *****   *   ** *******:***.    ******

44          MYTCWSRYCTRKGGYYCCKAGSSATARCGATKYKMYRWWYCYMSGARYYYYWTKKYTCAT
45          ATYYWSRYMWRKRRCTYYGRSCSRWRRYSWWGYKMCRWWCMYMCRGGYYYYWGKKYCMRK
                ****  **        ..*   * .   *** ***  **. . ***** ***   .

44          TKRWYATCTRGGGCAAGCRWGKRKYAWYRMMCCYYGRWRRGGCYYTARKYKWRCACKSGK
45          YKRWYMYMYRRRRYCCTAGTTKRKCMWYRMMTTCCKAWRGARMYCCGRKYTWRYGTKSKK
             ****    *    .. .   ***  *****      ** .  *  .***.** . ** *

44          MYKRYYYMYRRKKKRYWAMKTYACAYYKKRKYKMKRRAGAMMRGAKCGWTSYWRGTRRMR
45          MYKRYYYMYGAKKKRYWWMKCCGARYYKGRKYKMKRGTAGMMRKRKMRWWSYWRACRRAR
            *********  ****** **  .. *** ******* :..***  *  * ****. ** *

44          RMKKMRKRCAKMMYKMAGTGKSYGAYKRKTTKKKMCRTKRCGKMCTSYKKKKCMKYCCWK
45          RCKKMRKRYRKMMYKMTTATKSYAGYKRKAAKKKATGATGTKKMYYGTTKKKTMKYYYWK
            * ******  ******: : ***..****::***   :.   **  . .*** ***  **

44          YRCGWKRKKWTRWYRWMYRTYWKKKKRRRTRMRGSTAAKKGYKTWYWTYMKRWGTKKYRR
45          YRTAWKRTKWKRWYRTCYRWYWKGKKRRAAAMRKSYWMTKAYKYTYAWYCKGAAAKKYRR
            ** .***.**.****  ** *** **** : ** *   .*.**  *  * *  .:*****
```

```
44                  RKMMYMMYWGCYYKARKKKCGTAWTKYCYKKCAAGRKRTRGYGCTKYKRYKKGYCRYYRW
45                  RKMMYMMYWKAYYKRRKKKYRATWKKYYTKTYRMRRKRKRTYAAKKYKGYKKRYYGTCAT
                    ********* .*** ****  ::*.**  *.    ***.* *...*** *** *

44                  RGRKTGKKKKKTTYWYKYYKWKMYYKWRRTTKYYKYWMKKKRRRRYMYMMWKAGKYWWSK
45                  RAGKYKKKKKKKKYWYKYYKWKATTKWRRWKKYYKYTMKKKRGRRYMYMMWKWRKYWWSK
                    *. *  *****..*********   **** .***** ***** *********  ******

44                  KGKKRKYKYKKSGRRYYYRRRKKYCRKRKKYAMRKGTYRYRRRYKKSWMAKRKGARRYWC
45                  KKKKRKYKYKKSKRRYYYRRRKKYMRKRKKYGARKAACGYRRRCKGSWMTKRKAGGGCAY
                    * ********** *********** ******. **.:  **** * ***:***..

44                  CGAKMWMRYKTGWCMGWKYYYARRAMSYRYYRACKGYKGAKWYYWKYYTTMWWWKRKGTA
45                  MRRKAWCAYKWKWYCRWKYYYRRRTCCYRYYGRYKSYKATKWYTAKYYAKCTTAKRKRKM
                       * *  **  *   ***** **: .****   *.**.:***  ***:.    ***  .

44                  KCGKRMKYRYMYYYRKRRKRMKRTCAKTSGRRWTCKRYRYTMTTCCCYKRKYGKMRWKYC
45                  GYKKRMKTRYMYYYRKRRKRMKRKSRKKSAGGTCTKRYRYKMCKYSYYKRKYTKMGWTTM
                       **** ***************.. *.*.     *****.* . . ***** ** *.

44                  MKRWRRYTRRWKRYYMACGMRKMKWGKYYYRYKSMYKWRWMGYWRAKCCCKKRRRSYYYW
45                  CTGTRGCYAGWKRYYMRSKMRKMGWRKYYYRYKSMYKWRTMRCWRWGTSMKKRRRSYYYA
                     .  *     ****** . **** * ************* *  **   . *********

44                  MYRMMRYRYMKCMRACTRKKKTKMRYYWRYYGKKRTRRCGYRWMGSKTMMWCCTKWRYRK
45                  CYRMMRYGTMKTAGMAGRGKKKKMRCTAATTRKKGWGGMRYRWMACKWMMAYYKKWRYRK
                     ******  **    . * **.***       **      ****..* **   .******

44                  KKRRKKYYYYKRKKWCKTGKKTGYYSGYRWKKYKCTTGTTKWRRGGKRGGRMYYGKCGCC
45                  KKRRKKYYYYKRKKWYKWKTKYKYYSTTRWKGCGMAKKWYKWAGRKGRAARMCYRKMRTT
                    *************** *  .*  ***  ***    :.   **     *..** * *

44                  CWTAKRYRYMKYYSKSKKWYACTAGKKKKYSYWYRRKCGWYYMYRRRMRAMYMYKWTWCY
45                  TTKMKRYRYMKYYSKSKKWYWYYTKKKKKYSYWYRRKYRWYYMYRRRMRCMYMYKAWAYC
                      . ****************    : ************  **********.*****

44                  RYMWYRYTMYKWRRRKCWAYKRWKYTMWYKMATCTTAGMMRWWGKCCCKMYYGTYGWKYK
45                  ACMWYRYCMCKWRRGKMWMYKRWKTCATYKMCAGAATRCCRWWTGMMGKCYYKYYKWKYK
                      ***** * **** * * *****    ***.: :::   ***     * **  * ****

44                  RYYGYMGTKRRYKYKAKCCRYYRKRWMYKRYYRKKGATKYMCKAGGAMRYMRKYWGKCRA
45                  RYYKYMKYGARYKYKMGYYGYYRKRWMYKGYYRKKKRYKYMSKGTTWMRYMRKTTATAAC
                    *** **    *****     ********* *****   ***.*.   ******  ... .

44                  KTAKYGYKWYRKYGKMKRKYYKCGCAKAMKRYCWYRTATSYKWYCKRKCYYWYRGYRKKY
45                  KYRKTRCKWYRKCKKMKRTCCKATTTKWMKRYAWYRYWYGCKWYYKGKYCCACGRYRKKY
                    *  *   *****  ****.  *.  :* ****.***   . *** * *       *****

44                  RCYAYMMRRKKKGRMYYKWMYYKKYMMMKRRKKYKYRYKWGMGWRKKKYYAKCRYRKMRR
45                  RYYWYMMRRKKKKRMYYKWMYYKKYMMMKRAKKYGYRYKWAMCWGKKKYYRKARYRKMAG
                    * * ******** ***************** *** *****.* * ***** *.*****
```

```
44          GWKRMYYKRRKRSKYCCAMKRYRYMCKKCAGGATKWRWWMCARYRAYRYGKRSMKRTYCA
45          RWKRMYYKRRKRSKYYYRMKRYRTAATKTGARRKKAGTTMTGRYRGCACATRSMKRCYTG
             **************   *****  ..* ..  .*    * .***.   ..***** * .

44          GRRMGTKKCKYKWKRYGCWMRRMTTGRAYYYKSAWKYRKTGYRGAGRKAATGKRRMKRKM
45          AARCKYGGYKYKWGRYAMWMRRMAGTRGYYYKSRWTYGKKKYRKRKGKWWYKKRRMKRKM
            . *      **** **. *****:  *.***** *.* *. **    *    ********

44          MMKYMKGTRYRYSKRTTTWKWAYRKRRMWRWTTAWMMKKWMKYKWYYYWCCTYYRTRYTW
45          MMKYCTKYRYGTSKAAGCWGTMYRKRRMWRWYWWWMMKKWMKYKWYTTWMYYYYRKRCAW
            **** .  **  ** :  *   *********   ************  *   ***.* :*

44          YRKRAYRSRRYRKGKRTYCGCAMGACRMYYWYMAKYKRRRATKYRYSKWYYWYRKKKAGR
45          YRKRRYRSRRYRKKGRKYMRTTAAGARMYYWYMWKYKRRRRWKCGYSKWTCWYRKKKRRR
            **** ********  *.*   : ...******* ******  *  ****  ******  *

44          RTKCKTCRARYYSRWRYYYRYRKTTTKKTYCGYWKCCTKCCAKWCMYYMCWTYRWKGMRW
45          RKKMGCMRWRYYSRWRYYYRYRKCAAKKKYYRYWKMSWKGTTKWYMYYAYACYRWKKMRA
            *.*    * ************** ::**.*  *** . *  :** ***   **** **

44          YCTWKKRYYCGGGYYRYWRCWKYYKYTRTGCYRRGMKAKYYMYRKYRMYKKCMYCRKKKM
45          TTAAKKRYYAAAAYYRYWRMAKYYKYCGAATYAAKMTMKYTMCAKTRCYKKMMYARKKKA
              : *****....******  *****  :. *   *. ** *  * * *** **.****

44          TWRAKWRKWCYWYRGRCCRRAMKMYKSYRKYKSWCRYYCARMWWAMKRWMKTTTKCKYTA
45          YTRGKAAKWYYWYRRAMTRRCMKMYKSYRKYKSWYRYYTGRMWWMAKRWMKCAAGAKYYR
              *.*  ** ****    **.************* *** .****  ***** :: .**

44          YKRGWRRYYWYYCATGAWRCAAYGRCTRRWRRYCCTRRMRMRYGCTAYRRCYKYRSKAMY
45          TKRKWRRYTATTTTATRWRYGCCCAMGRRTARTYAGRGCGCRYCYYRYRAMYKYRCGGCT
             ** ****     ::  ** ..     **  *  . *    **    **  ****. .

44          YWCCGKKYRKYATTRWWAMWMCKKYAAKKGKKAYMWCTRYRAWYYSTATTSCGWYYKYKA
45          CAATAKKYRKYTAARWWRCTMYKKCGCKKAGTRYMWTGRYRMWCTGWWCGSTTTYYKYGR
              . .******:::***   * **  ..**. . ***  *** *  .    *   ****

44          ARKRMATTCYKGRGMCRGKTKKCTTYWYTCWAAYMYKRYRKKKRYYTARYWTCCCTTYRM
45          GRKRMMWWYYKTGAMMRKKCKTTKYCWCYYWWRTMYKRYRKKKRYYYRRTAATAAAWCAC
            .****    **  .* * * *. .  *   *   ************  *  : ..:

44          TMRKMKGRGTYGYRKWGRKKGRKMTTGMMGTYRRYMMYRKWAGGCKKGTWWGATKWKRAA
45          AMGTMKRGRWYRYRKARRKKKRKCGATMMRYYRRTAMYRKWRCAMKKTATTAGCKTGRTC
            :* .**    * ***  *** **  : **  ***  *****  . ** :  .. *  *:.

44          ATCKMRSWYYAAGCRRRKRRYMRMYWTRGAKMGCGCTRACYCYWKGKKTWYRAATKRRMK
45          RGAKMRSWTTMRATGRRGRRYCGMTAGARRKMKYRYWRWTCTYAKKKTAWYRGMCKRRMK
              .*****    .  ** ***  *      **     *    * * *.:***.  *****

44          ARKRCRYMRYGKKYYMRWMYAKWYYKYTGTCGRCYYTRYMKRYRRKACYWGYAGMSWTYG
45          WRKRSRTMRYRKKYYMRTAYWGWYYKYYKCTARYYYYRYMKRYRRKMYTWRYMAMSWCTR
             ***.* *** ******  *  *****    .* ** *********   * * .***

44          MKYYKARYRMMTTKRRRYKTACACTRYRKKKKSRRYYYKKKTRKKKRYTKKRCTMWMCKA
```

```
45         MTYTTRGYRMMAYTRRRYKWWYWTYRYRKKKKGGGYYYKKKWRKKKRYKKKGYYMWMYKM
           *.* .  ****: .*****      *******.  ****** ******.**   *** *

44         SRAYRKMYKSRRRYRCKWGCGCYRYKRAGKGTGYYKRMMKWAAKSMWWYKKRKGGTGYKK
45         GGRYRKMYGSRAATGYKWAMKYYRYKRTKKACATCTRMMKTCCKSMWWYKKGKKSYKYKK
           .  ***** **     **.   *****: *. .  .**** ..******** * .  ***

44         GCACTKAARRAKRCCRWRKYRCCARGRTTCCTCAMRWMTGRTTAYYKRKYRMYRMKYMMG
45         KMMYWKWTRRMKRMSRWRKYRYYRRKRAAGAATWMRWMYRRWWRYYKRKYRMYRAKYMMK
                * :** ** .******   * *:: .:  ****  *   ********** ****

44         GKTKGTCATCAKKCKYKKYYRYMRRWCWMRRRCATCACGCYRKYCWTKMTRKCCCCWYYK
45         KKCKKWTMKSRKKMKYTKYYRYCRRWMACAGATWWTTTTYTGKYTAGKAGRGTTATWYYK
            * *    .. ** **.***** ***          :     **   *  *   . ****

44         KRYKMRWKACCGGCARGYYMWMKTGYAAGGKRGRCARWRRRGATYRRMRKRKWKSKYGWY
45         KRYKMRAKRYTKTMGGATYMWMKWSTTGKKKRTGTGAWRRGSGAYRRMRKRKAKSKYKWY
           ****** *      . . ***** . :.  **   . *** ..:******** **** **

44         YKKMYMSKCTSRKKKWTTTRAKRRRYWYRYYGTYKYMRRYRYTCYRCKAYKKMGAYAYCK
45         YKKMYMSKTKSRKKGWYWYRWTRRRYWYRYYKYYKYARRYRYYMYGTGRYGKMRRYRCMK
           ******** .**** *   * .*********  *** *****  *    * **  *   *

44         KGKATAAWKTAGAGGCYWTTAWKYYYYMYCCGWKKSMMATWSYRAMYYYRRKKKYCKKYY
45         KKKTCTTWKCTAGRRMYAAACWKTYYYMCTTATKKSMMRWWGCGMMYYYRRKKKCMKKCC
           * *: ::** :..   * ::.** ****   . *****  *.   *********  **

44         RCAGTRGAGYYKKAMKKKCGKKAWYYYGTAAAYYRKYGTRKYTTRSKWRKYRAATGKKKK
45         GTTRCRRWSYYKKRMKKKTRKKRWYYTKWTCCCYRKYKGRKYWGRSKWRKYARGCRKKKK
             :  *  .**** ****  ** ***   :.. ****  ***  *******  .  ****

44         KYRYGKYYRKKKKWKARKTGRKKYYSYWYSKKRKYCRYKMMRWYKCYTWKKCKGYRKGWW
45         KYRYCGTTRGKKKWKMRKYRRKKYYSYWYSKKRKYGRYKMMRAYKAYKWKKMKKYRGKTW
           ****    * ***** **  *************** ****** **.*.*** * **   *

44         GYRKGKYRRKKRRKTKYAARWKRWCGTKKKKRMTRYKYYSKGCATYGRYKKYKMYYKKSR
45         KTAKAKYRRKKRRKYKCMRRWKRWMTCKKKKRMCGCKYYSKTAGAYAGYKKTKMYYKKSR
              *.********* *   *****   ******   ***** ..:*. *** ********

44         KMTKRYSYSKYSYKKKYWRRRYGKMMKKCAYAYSKGSKMCGKKRKKKYKGTYYYAYRTAT
45         KMKKRYSYSKYSYKGKYTRRRYTKMMKKYRYRYSGKSKMYCKKRKKKTKKACCCTTGCMW
           **.*********** ** **** *****  * **  ***  ****** * :   :

44         CYKYKWKYYYYMRCTMRYSKGACGKRYKSAKYYWCAGCTMWMYGKGGGTWKYKCGARWTT
45         MYKTKWKYYYYMRYWMRYSKRGTTKRYKSRTYYWYMRYYMWCYKKRRRYWKYKYSGRWGG
            ** *********  ***** .  ***** .***     ** * *    **** ..**

44         CTGTTTYCKRYKKRKWMTWRRYWMYRCKTGGYKYGMRYYKRGYKYWMWYYAAKATRWSGM
45         TWRWWKYYKRYKKRKWMKWGGCWMYRMKGTKYKYSMRYYKRRCKYWMWYYGMKRCATGAA
                .* *********.*   **** *   ***.******  *******. *    ..

44         GWKKWRRRAAC
45         AWKKTRRGGWT
```

.*** ** .

The pairwise alignments of two accessions 44 and 46

```
44                  RYGAYGATGYWRSWYWKSYYMYRWKSMRRYKACGWYRMSARKYYYKMMKYKMCTGGMKMY
46                  RYTRCRRWSYARSWYWKSTYMYGATGMRRYKGAATYRAGGRKYYYKMMKYKMMWRSMKMY
                    **      .* ******* ***  ..*****... ** ..************   .****

44                  SMMYWRRRRWKWAMMWRYYYYYGYRRWWKKYYYYTCYRAKCAKAYRRRKTYYRWRYMMWW
46                  SMMYWRRRRWKWRMMWRYYYYYRYRRWWKGYYYYWTYRWKYWGRTRARKWYYRWRYMMWW
                    ************ ********* ****** ****  ** *     * ** **********

44                  WYRRCRCYYMRRGATRCMYMRMWYGYCCCMYYTYMRKMYRRWYYRYMWKTYRYKRKKYRS
46                  WYRRGATYYMRRAGCRYACAAAAYKCTTTMYYAYMRKMYRRWYYRYMWKYYRYTRKKYRS
                    ****   *****.. *       *     ***:**************** ***.******

44                  AKRCKWWMRYWKKGKKWYRYRYAACMYYYYWKRSYRYYWWWRTKKRYRRTSRYRKYYWAR
46                  RKRAKWWMRYWKKAKKWYRYRYWRYMYYYYTAGRSYRCCATAAWKKRYRRASRYGKYYWWR
                     **.*********.********   ****   ****       ******:*** **** *

44                  GKYYMKCRYWWRRWYYTATKRYYRWYYKRYRCMGSWKYKGMMTWYWARRYRRRKWYGRKY
46                  RKYYMKYRYWWRRWYYWWWKRYYRWYYKRTGYCRSWGYGSAAYTCWRRRYRRRKWYTRTY
                     ***** *********   **********     ** * .     * ********* *.*

44                  MWYMTWTWKWKRMRTYYWCKTYYGYKWWSYWWRWYRKYMYYWKWYYYYYKWGGYKYRKCY
46                  MWYMYWGATATRMRCYCWTTKYYKYKWWSYWWRWYRKYMYYWKAYYYYYTWRRTKYRKMY
                    **** *  . .*** * * ..** ******************* *****.*   **** *

44                  TAKKKSMMYSRRRRYSRMKYCAAWYSYKYKRRTACRRYKRYCWRSSWMYYKKYRYRWYMY
46                  WWTKKSMMYSRRRRYSRMKYAGGWYSYGYKARAGAGRTTRTYWRSSWCYYKTYGTRWYMY
                      .*****************...**** ** *:.. * .*  ***** ***.*  *****

44                  YKYKGCRKRYKRRYRYMYWKWRRRYMRWMTKTTYKKKMWYKRKRMRYMMWGKKKTKKAKW
46                  YKYKRARKRTTARYRYMYWGTGRGYMRWMCGACYKKKAWYKRKGCGYMMAKKKKKKTRGT
                    **** .*** . *******   * *****   :**** *****   ***  ***.*.

44                  KRYYRYGTYKYMRWGCGKMRRRKTTYRRKGATASRTTGARRYKKRYKAGYAMRKYMKRYW
46                  TATYRYRCYKYMRTCAAKCAGATWYYRRKACCWSRYGRWRGYKGRYKTCCCMRKYMKGYW
                    .  ***  *****  ..*    .  ****..  **    * ** ***:  .****** **

44                  MKYMTMYRAWTYYKRACKKWRCRRYRTRRRKKSYKKMYWKRKKMWWYCYGSWKRKYKTKT
46                  MTCCGMYRRTKYYTRRMKKWRYRRYRKGGRKKSYKKMYWKRKKMWWYTCASWKRKCKYKA
                    *.   ***  .**.*  **** ****.  ******************  .***** * *:

44                  TKGMRCKKMWGTYATWRYSTYKKKRRARCTYWYYCKRGYMACKCMGRRGTKKCGKGRGRT
46                  KGKMRMKKMTKCCCCWRYSCCKKKAGGRTYYWYYMKRAYCTYTYARGRTAKKTKKKRKRC
                    .  ** ***    . ****  ***  .*  **** **.* : .    * :**  * * *

44                  WKARMMKKRKRRMKRRWATTGMYMRTWCGTTKTKWMKRYKATGTCCCTAYYYYCSKKKRR
46                  WKGRMCGGRKRRMKRRWRYYRMYMRCTTKWYKKKWMGRYKRCRYSTYYGYYYYSCKKKRR
                    **.**   *********    ****      *.*** ***    .   .****..*****

44                  RYYWATKYSYWTMMKKCTTKGTGYGGAGWKYKYSYWCRATTYRRKKYRRRARRWWYMYRY
46                  RYYWRWKYSYTYMMKKYKKKKKRYARTTWKYKYSYWMATAWYRRTKYRRRCRRAACMYRY
```

```
        ****  ****  **** ..* . *.  : ********  :: ***.*****.**   ****

44      KMWYRYCARRRTMTGYGCTTTTKTATGRCRGCTRWYRRYWAYTGCYKRRYKGTTTKYKYY
46      KMATGYACRRGCMGTCATWCCGKCCCTAARKYWRWYRRYARYCAAYKRRYKTCWKKYKYY
        **   *..**  *   .     * .   .*   ******  * ..******   .*****

44      KRRCCGTSTATWRGGWGKWKGCYYWYTTCTRWYYMGTKRKKKRYRYKRWRCYWMMKYYRT
46      KRRMTACSATGARRATKKWKKAYYTTACMCRWCYMKYKRKKKRYRYKRWRYYTMMKYTAG
        ***  . *::  * .  *** .**  :   ** **  ************* * ****

44      AGYKKYYRKMWCGAKKMKYRYKRKTYWYKKYRRRRMMRRRRWTMKKYAYGKKSYWYKMYR
46      CKYKKYYRTMAMRMKKMKYRYKRKKYWYKKYRRRRAAARRAWKMKKTRYKKGSYWYKMYA
        . ******.*    **********.**********   ** *.***  * * *******

44      YMYYYYRYMKYSKRRKAAWKWYRKMKMKYYRKKKRYSYYKRYRKKMRRKSRKSKRAKRRK
46      YMYYYYRYMKYSGARKRRAKWYRKMKCKYYRKKGRYSYYKRYRKKMRRKSRKSKGMKRRK
        ************  **   ******* ****** ********************  ****

44      TMRYTAGACWWYWAMRKRYAKKRYSGTTGRGYYKTATKYACARMKRYKKYRWRWYRYCGT
46      KMGCACTRYWWYTRMRKRYMKKRYSRWARRRYTKYRCTTRSRRATGCKKYRWRWYRYMRY
        .*  :.   ***  ***** *****  : * * *   .  . * .  **********

44      GCCRSGKKGWYMCCYRYRTTTMMTGRYYRAKKGTTTRWYTKYRMKTTYTYWAWKYMWMYK
46      AATAGAKKKWTCYYCACGWKCCCKRRYYRRKKAWAARWYKKYRMTCWTCYTCTKTCTMCK
        ..  ..** *         .   . **** **. ::***.****.    * . *   * *

44      YRCCAGCCYYRKGCWRYKTAYATCCAKKKYYMYAMYYCTTMGMKYRRGTYSCGTGYRYKW
46      YGTYGATYYYRGRYWRYKCTCWCAATKKTCYMYWCCYTYCCRMKYRGTGYSTTCATACGW
        *   ..  ***   **** :   ..:**. ***   *     ****   **   .    *

44      AYMARTTGCWRRGKWAGGKCKRGKMRAGGWYYKMYKRYTKGKRYYCTGARYRYGRYAKRR
46      RYMRACCAAWRRTKWWKKKTKRATMRGAAWYTKMYKRCYKRKRYYACAGAYRYRGCRKRR
         **    ..*** **   * **..**...** *****  * ****. .. ***    ***

44      RWSRKRKMYRYTRKKKATTWKYMYTTGMMYKKRMKAYRWTATWRKTTAYYYYWTGCCGRK
46      RWSRTRKMCRYCRKGKCGAWKYMCGATMMYKKAMKWYRWWRCWRKCAGCYYYWCRYYRAG
        ****.*** ** ** *. :****  : ***** ** ***   *** :. ****

44      YTWGARTRKYSWWYYYYTRGYTRRGKYSRYYMRMRRKYMKATTKMMRRMKGCMRKTKWKK
46      CAWTRRWRKTSWWYYYYKRTYGGGKKYSRYYMRMRRKYMTRYWKMMRRMKKAMRKKKWKK
         :*  * ** *******.* *    **************.   ******* .***.****

44      KWRWRTGCYKRRWKRRKRYYRYMRKATKYRRYYKRRYRKYKRYCCKYKTGRYKYWWTKKC
46      KWRWRATGCKAGWTRRKRYYRYMRGRYKYRRYYKRRYRKTKRYTYGCKKSRYKYWWKKKT
        *****:   *  *.**********   ************ ***    *..******.**

44      GCKRWWTTAWWYYYCCCMYWKMMGKMRGCSWSYRMMWYTKCTYSCWTRKKRCACAWYGAC
46      KGGGWAWACAAYYYATAMYWGMMSKMRTGSWGCGMMWYGKYYYGYTGRKKGMMTWTCRMY
            *  :.  ***. .*** **.***  **.  **** *  *.   ***

44      WYRTWRYKTMKGGYMKYKKRTAYKMYRYTGKRRGYTKWMRKRGSGKWYRMWKRRKCWTAK
46      WYRKWRYGWAKRRYMKYKKRAGYKMYRYWTKRRSCWKWMRKRRGKKWYRMWKRRGTACCK
        ***.***   *  *******:.******  ***.  ****** . *********    .*
```

```
44    YYRTMRYATTAKKSYRRWWMKGKKRRRWKYYYAGKRRYCAGCMKRYRRKTMYYWGRRWMW
46    CYRYMRYCCCCKKSCRRATAGKKTRRRWKCCTRSKRRYAGCAMKRYRRKGCTYWKGAWMW
       ** ***.  .*** **     *.*****    .****.. .*******   **   ***

44    YMGYGGGARKRYRYKRYRWMRSRRKKAYGKRYKRKKKSRRKKKYKCAKTWATAKYKKYKR
46    YMKCRSRRGKRYGYKRCGWMRSRRKKWYKKRYKRKKKGGRKKKCKYRKWWTAGKYKKCGR
      **   .   *** ***  ******** * ********. **** *  * *::.****  *

44    YKGKRMWMSTYKYRMYAKGARRYRWATCYRYRRRKKMKTTAKMKGATGGASSKYCMMKRK
46    YKRKRMWMSCYKYRMYRGRRRRYRWWYYYRYRRRKTMKYKMGMKAGACRGSSKYTCAKRK
      ** ****** ******    *****   *******.** .  **..:  .****   ***

44    SYYMYRKYMRYKKKYRGWWRRKRCTAMKKAYCYSGYKWAWTCAGMRRKWYKRWRAGRMCR
46    CTYMYRKYMRYKKKYARAWRRKRTGGMKKCCYYSKYTATTCMRCARRGTCGRARWRRMMR
      . *************   *****  .***.  ** *. :      **    * *  ** *

44    YKATCRRYYKYWKYRRWKYMYYYYRKKMRYSKKKGMGRKKYYWMMCTMWYGGGTWKYKYR
46    YKMWMRAYYKYWKYRAWKYMCCCTRKKMRYSKKKKCAGKKYYWMMYYMWYRRRYWKYGCG
      **   * ******** ****    **********  . *******  ***    ***

44    YKRWRKGYKWMWYWMYCTGAMKTYKMMYKMRYYKYYKKGYYWMWCCYRRWKWWKKRRAKK
46    CKRWRKKYGWMWYWMYMKKGCKCYKMMYKMRYYKYYKKRCYWMWSTYRRWKWWKKRRMKK
       ***** * ******* . . * ***************  ****. *********** **

44    YWRWMAYGGTGCTMGAARMMGMRRYTTYKTTGTRRCTKYYRWMMMYCTWKGYYWRMYRYY
46    YWRWMCTRAYKSYMCTGRMCTMRACCGYKCCAARGGAGTCRWCCMTYCWKAYYWRCYRYY
      *****.  .  . * :.**  **    **  .:*  :   **  *   **.**** ****

44    GKKGCTRWYRYRYMGYWCCCCKTGYKCCTTKRTSMRRAYAYRTGMKGAAYMRKRRYKYTG
46    RKKRMYRWYRYRYMCYWTTTTKWRYKTTGCGAASMARMYRYRKKMKTCWYCAKGGCGYKK
       **   ******** **    *  **      :** * * **. ** . *  *    *.

44    MRCCRRGGYCTTYTCKRYRYRYRKKYKTRKKKCTYTRATGGMKYMYKKYRKAGAKTTYMC
46    MGYTGGKKYTCCYCYKRYRYRYRKKYKARKKKYKYKRRKRRMKYMYKKYRKTTGKYYYMY
      *       *   *  ************:**** .*.* .  **********: .*  **

44    YYRYYKMKMWWKRRTTKYTYAKKCGATTAYKAYRYWYRTTWKTGWKKRRTRGKKGCKKYR
46    TYRYYKMKMWWGRRKYKYCYTTKGRRAGCYGRYRCWYRWWWTKSWKKAGKRRKKSTKKYR
       ********** **. ** *:.*   : .*  ** ***  *...***  .* **. ****

44    TTGWCGCARKYYGTYACMMRKKRWRKKCKKTYWMKYYCMGKKKKMGGTCSTMYRTKKKMY
46    CAAWMRMRRKYYAAYTSMMRKKRWRKGYKKKTWMKYCYCKKKKKMKKWMCKMYRCKKKMY
       :.*    ****.:*:.*********  **. ****    *****    ..*** *****

44    WRKRMWRGKRWRKKKCCTCGMKWTRWMKKRYSKKTMKKCGCGSRKMYMWKMWSYKRKSWR
46    WRKRMTRRGRWGKKKTTCMRCKWYRWATKRYSKKWMKKAATTSRKMYMWKMWSYKRKSWR
      ***** *  ** ***      ** ** .****** ***..  ******************

44    WMKYSKCMWWYYRKKTGMGGGCGRSKGTAYMKSACGGYACMWKRMRKKYACYRSKTTSGY
46    WMKYSTTMWAYYGKGAAMKRRYKRSGTGCYMKSRMKRYGTATKRMRKKCTYYRSKGKSRY
      *****. ** ** * :.*     **   .****    *.   ****** : **** .* *
```

```
44              YRTYYYWKMYARWCGKKKYRRRYGSYKKRWRSMYKKYKYYWRASKRMYRRGKYKWKRYRK
46              YRWYYYWKMYGAAYAKKKYRRRYKSYKKRWRSMYKKYKYYWRCSKRMYRRKKYKWKRYRK
                ** *******.   .******** ******************.******* *********

44              YWKGMYYYGCKKCAGAYRKRRMCTSRARYGMKMRGWSRYKMYKTGSMYSYAGYYRMAKYY
46              YWKRCTCCAYKKATTGYRKRRMYWSRGRYRATARSWSRYKMTGYKSMYSYRSYYRMRKYY
                ***      . **.: .******  **.**  . *.******    ***** .**** ***

44              KYKYYRMTGKYKAYYMRSRKGGKAYARMMYRKRRATTKWYWRGRWACAKRKMKCCCATGT
46              KYKCCRMWKTYKGYYMRSRGAAKGYGRMCYRKRRRKYKWYWRARACYMKRKMGAMYWSY
                ***  **   .**.****** ..*.*.** *****  . *****.* .  **** .    .

44              KATCTRYTGMRKKKWRRWMTGCKCYKRGYKKKKGYCYYGGGGYCRYGYRYRKKKKWRMAA
46              GGYSYRYAACRKKTWRATCCAMTTYKRRTKKGKRTYYYKRRRCYGTSTGTRKTKKWRMMM
                 . . **:. ***.**     . . ***   ** *    **          .    **.*****

44              KYRTRKCSYWYRKRRKRYRKAYARYYYCKMKMRWRWRKKYYKKKRYKYRKWWRKYYRKKY
46              KYRWGKMSYWYRKAGKRYGGGYRRYYYYKMKMGARWRKKYYKKKRYKYRKTWRKTTRTGY
                ***  * ******  ***   .* **** ****  *************** ***   *. *

44              YYWYRWYRKKRRYRTTATKTGTRGGAGKRKKKRKATYYTSTGCCKARYKGRRYKKYKYYK
46              YYWYRWYRKKRRYRYKRYKAACRATTAKRKKKRKGATCKCAAYTKRRYTRRRYGKTKYYK
                ************** .   *:. *. :.*******.:   ..:.   * **. *** * ****

44              SMRKMWYCYAGYRYKYGCTCGRYMCCRTTKKKRYWAMRYKYYKARKKTKWMRYCAYTWCR
46              SMRKMWYYYRKCGTKYAGATTRYMMMGYAKKKRYWWMRYKYYKGAKKKTWMRYMGTKTTA
                ******* *      **. :  ***     :****** *******. **..**** . .

44              TTYKRRGAAGMRRKRWSYWRGTYARAGMRKRKYTTYRRWKMRATRCRKAYKMMYWMRMKR
46              CYYKRRAWGAMRRKRWSYWRACYGRTTMRKRKYCCYRRWKMGTAAYRKRCKMMYWMRMKR
                  ****. ..**********. *.*: ******  ****** ::   **   **********

44              RSYMAKKRKRRRKKAKKKYKKWYYYKWKRMYKKMYYYWRWCCGRKSKMKKRRMRYWSKMM
46              RSYMRKKRKRRRKKMKKKTKKWYYYKWKRMYKKMYYYWRWMYRRKSKMKKRRMRYWSKMM
                **** ********* *** ********************   ****************

44              GYMRKKKRWKTCARRGMRKKKYAACATARYWRCKYWYGYKWYKKYKTTRKKKAYYKMGGC
46              KYARKKKRWKAATRRKMRKKKYGRTTCTRYWRMKYTYAYKWYKGYKAKRGKKMYYKMAAT
                 * *******:.:** ******.   : :**** ** *.***** **:.* ** ****..

44              CKKKKRMYRMCRKAWMYGRKKYKKKKMWKRYWWRMYRMYRYKYKWMYCRRRRMYKKRKCT
46              TKKKKRAYRMYRKGWMYTRKKCKKKKMWKRYWAAMYRMYRYKYKWMYYRRRRMYKKRKMK
                 ***** *** **.*** *** **********  ************* ********** .

44              CSYRKGARCCTRRYGRKRSWKAYKYYKWKKRRRWTWAMKMSYYYKRWKYMMMYRYRGYYS
46              YGTRKRTATMWRRCARKRSWKCYKYYKWGKRAGTYWRMKMSYYTKRWKYMMMYRYRKYYS
                 . ** :    ** .******.****** **     * ****** *********** ***

44              GGGKRYACYTTCAKKKKYWMSMRKWGCKWMMGTTARKKTCTKWYCTAMWRKYYRRRGTRY
46              ARAKAYMYYWKAGKKKKYWMSMRKWAGKWMMSYYRRKKYTCKWYMWGCWAKYYRRRRART
                . .* *  * ...***********. ****.    ***    ***   . * ****** :*

44              YRCCRAKWRRSRRYRKRRARCCCKRCMYMKYRYRRRRRKKKYYKWYKRGCTRRYRYRMCY
```

```
46          YRYYRMKWRASRRYRKRRMGTSAKRGMCAGYRYRRRRRKKKTCKWYKRATYAAYRYRMMY
            **  * *** ********   ..** *   ***********  *****.    ***** *

44          TRYYKKSYKYGCGGYMTTRWTGTTKKWMYYRRTCYKYRYYRYYYKMRMKKWYYYRKSGKY
46          CRYYKKSYKYRMRRYMWKRAYRYYTTTCCCRRAMYKYRYYRYYYKMRMGKWYYYRKSRKY
             *********    ** .*     ..    **: ************** ******** **

44          WRWACRGCATAKACCYAKRKKKGMRKRSYWRMKMRYYMKWRYKARKMYYYWKKAAATATT
46          WATRYRRYRYWKGTTCGKRKKKRCGKRSYWRMKMRYTMKWRYKRRKMYYYAKKMMRWRAC
            *    *     *.   .*****   *********** ****** ****** **     :

44          ARTCGRGAYRTMKMKYCCKWRYRGKKKRKYYYKKWRCWWRKMYKYWKRSGMKRATWRYMR
46          TAATAGARYRCMKMKYYYKWRYRKKKKRKYYYKKWRMWWRKMYKYWGGGSMGRGAARYMR
            : : . . ** *****  ***** ************ *********  ..* *.: ****

44          MSRYMRRKKTYTTTGGKWKWYRYYKYRRKYRCKTSYRRKKKKRRRRWWCYYYRWYRTRWR
46          MSRYMRRGKWCACCARKWTWYRYYKYRGKYRTGASYRRKKKKRAAGTTYYYYRWYRWGWR
            ******* *  :  . **.******** ***  :*********      *******  **

44          WRYAKWRACYAKRKAAYWYMRRRMYKKKKYRMCGAGRGKKKYYKKYCCWKRRAYRRWGYK
46          WRYRKWRGTCCKGKRMYWYMRRRMYKGKKYRAARWARRKKKYYKGYSAAKRGGTGATSYT
            *** ***.  .* *  ********** **** .  .* ****** *.. ** .    .*.

44          RRYWATTARGYYTRKKKKCRCAYRYMYYMYKYTCRRYMYYKMSMKYMYRRYKCYYGGAGT
46          ARYWRCYRGATYWRKKKKYRMWYRYMYYMYKCAYRRYMYYGMSMKYMYRAYKATYATTRW
             ***     . * ***** *  ********* : ****** ******** **. *. :

44          TCRKKWRWWRSMRKRCCGCRKKKYKTCWGTYKWRKCKCATRTTWTRAKGAMKTGATRYWK
46          YMRKGWRWWRSMRKRYMTYRKKKYKYYWSWYKWRKYKTTWRWKWKGMKCGMKYATCRYWK
              ** **********    ******  *. ***** * : * .*.  * .** .: ****

44          RYRRATGGSGACAYYKWMYKWKKKRRRKMKMKKWYMYWKKYKYYKSKTWMWKKYCCYYKK
46          RYRRMYSSSKRYMYYKWMYGAKKKRRRKMKCKKWYMYWKKYKYYKSKCWMWKKYYYYYKK
            ****  ..*    ******  ********* **************** ******  ****

44          KRKRRTCTYGGCCMTTKCAAWMYRYWYKKSRTKGYKKKKYKYRYKKACKCTMYYYKSCRK
46          TRKRGYYWCKAAYMYKKSWRWMYRYWYKKGRKKRCKKTKYKYRYKKCTKYWATCTKSYRK
            .***      .. * .*.  *********.*.*  **.********. *      ** **

44          YYKYWMKKKRRWRRKMYMWKWWSRTAGKKYKYKACTRRAATGYRKKYKSYKCRYRRRYKA
46          YYKYTMGKKRRWRRKMYMWKWWSGCRKKKYKYKWYWGRMRKKYRKKYKSYKYGYRRRCTC
            **** * ****************    ******    *  . *********  **** ..

44          GAGCTYYKWKGMKRWYTCGRYCKYYYKRWGAAKTCTGYKYRKRCRGGCTRYCYYGTGCCA
46          RWKYYYYKWKKMKRTCWYKRYAKYYYKRWSMRGAMGKCKYRKRMGKAMARYYYCTCTAAT
                 ***** ***     **.*******.   :    *****   . :** *    ..:

44          RGMACACGYYRMMKYSCKKYWKATGTTAMWWYRWKKWYRWRAGCAAKYMRTCTRRTGMKK
46          RAMRYRYKYYRMMKYSYKKYWKRWKKWCCTWTRWKKWYRWRTRTGTKYMRAAGGGCSMKK
            *.*     ******** *****   . .  * *********:  .:****:.    .***

44          KKGGGGWYKR
46          KKRRRCWYKR
```

**        ****

The pairwise alignments of two accessions 44 and 47

```
44              YMWYMAYMATCRTAWRRKMAKRRCYMMRYRYKMMWRRRMRWYYGRYMYWKARWRWWMSRM
47              YMWYCWYMRATRGRWAGKMGKRRYCMMRYAYKMMWRRRMRWYCKATAYWKRRWRWWMSRM
                ****  ** : *  *  **.***  **** ************     *** *********

44              YCTTRYMYYSRGRCRRKWYKKYSSRWGTKCRYWYYRYKGTCMYMRMRYKYRTYKCWRYRY
47              YTCYAYMYYCRSRYRRKWYKKYSSRWRWKMRYWYYRYKAAAMYMRMAYKYRKYKMWRYRY
                *    ****.*.* ************  * ********.:.***** ****.** *****

44              KSYYRSYYRRRYMRYRYGTYRMRSAAYRYWKSTRYYRMMRYRSKYRRRYYMRYSRYYYMY
47              KSYYRSYYRRRYMRYRCTCYRMRSGWCAYWKSYRCCRMMRYRSKYRRRYYMRYSRYYYAY
                ****************   *****.   **** *  ********************** *

44              YWRYRWWYYRYSRRKRYMTGMYYRYYRMKMMRWCYTGTGGYCRKGWYYKYWWKYRKWKYG
47              YWRYRWWYYRYSRRKRYMWRMYYRYYRMKMMATMCKRWKRYYRKRACYKYWWKYRKWKYR
                ******************  ***********    .    * **   ************ 

44              YKKYGCYWMGCGKYRRKYKMTYMMKMSMWRRMMRWRYRWWKYYYYKYRYYYKKWKYRRTG
47              YKKYATYWMKYAKYGATYKMKCMMKMSMWRRMMRWRYRWWKYYYYKYRYYYKKWKCRRAA
                ****. ***  .**  .***. ********************************* **:.

44              YWYRKWRCRKKYWYGRMYSRKYAWRSAYAKYKYAYYRRYTCTRRYYRWTKWYYYWWGRRY
47              TTYRTWATRKTYWYARMYSRKYWWRSGYTKYKYRYYRRYCYCGGYCATWKWYCCATTAAY
                  **.*  **.***.******* ***.*:**** *****     *    ***       *

44              WSCYYYARRWYYRWYGYMGYWKWYKRMYWYYWMRTTRKYYRYMKRRYRWKMKRYYWKRWY
47              TCYYYYWRRWYYRWYTYMRYWKWYKRMYTTYWMRCARTCYRYMKRRYRWKMGRYTAGRWY
                 . *** ******** ** *********  **** :*. ************ **   ***

44              RRYTYKYYWYYRYKSWTMRKKYYCGGKMWTRYGYCTWYKWTCMYASKRWKYGACTSYYGR
47              RRYYYKYYWYYRYKSWAMRKKCCYARKATYRTAYTCWYKWYSMYRSKRWKYRWMASYYRR
                *** ************:****   . *   * .*  **** .** ******   :*** *

44              CKATKKYYTMWWWRCAKRWKMKRYKYGGCRYRKRRAMKRWRGTCATCATCKWYKTYAYWG
47              MKRYKKYYKCTTARYRKRWGMKRTTYRKMRYRKRRGATGARKKSRGYWWTKWYKCYTYWT
                 *  ****.    *  *** *** .*   ******. .  * ..      **** *:** 

44              GYYKMYRKKYWRRKRRRWWMMKYTTYCYSSYKGTAWMGKWYKKRKGYRMMKKKTYYAKYK
47              AYYKATGKKYWRRKRRRWWMMKCKWCYYSSYKRGGTCRGTCKKRKRYRMMTKGACCTKYK
                .***   *************** .   *****  .      **** ****.* :  :***

44              YWYYRRCYWKMYYYYRKKACWYRTYYMRRWSKKRYYYWMWYKRKMWKKYKYRYKRWKRYT
47              YWYYRAYCWKMCCCTRKKTTTCRCYYMRRWSKKRYYYWMWYKRKAWKKYGCGCKRWKRYK
                *****   ***    ***:   * ******************** ****    ******.

44              GYKRWCWCAYCGGGGRYKWRKRCYKKTKMRYYKYYYKKGTAYKWMAACYRRWKTAWKKRR
47              RCKRWMATTTYARAARYKWRKRMYKKAKMRYYKYYYKKRYCCTWMWWYYRRWKYRWKKRR
                  ***   :  . ..******* ***:***********  . .**   *****  *****

44              MAGYKKYWMTGCTCTGGTGMRKKWMYYKRKRMMRKMWYKCGAACTGTYCARRYRRTYRYY
47              CGRYKKYWMYKSYYYTKAAMRTTTCCYKRKRCMRKMWCTMAGGTCTGTTCRGCAGYYRYY
```

```
                 . ******  .     :.**..   ***** ***** . ...      .*     ****

44               YKACGMRWKKKWAYYYRKKMGTGCCCATRWYRYWRRKRRKRMRYYKRRRWRMRKRYMRKK
47               YKRSKCGTKKGWMYYYRKKMTYKYYAGKGWTRYWRRKGGKRMRYYKRRRWRMRKGTARKK
                 ** .     ** * *******     ... * ******  **************   ***

44               TRKWWMKYMRAYWGYGGTRGKTRYRAGKMMKTGMMKWWRYRKTGKKKARRKYYYKYRKGK
47               CRKTTMKYCRCYARTRRWRRKWGYRTTKMMKWKMMKWWRYRKCKKTKRRRKYYYKYRKAG
                  **  *** *.*      * *  **: ****  *********  *.* **********.

44               KASRWRWKMGGTGKKKGKYKRRMYSCTATYRKARMYRCAGCMRWMTGRYKKCARKRKYGG
47               KRSRWRWKMKSYKKTKRKYKRRMYSMWWYYRGRRMYRYWRYMRWMYRRYKKYRRKRKYRK
                 * ******* .  *.* ********    **  ****    ****  ****  *****

44               KTKKYKKGKKYWYKRMRRCCCWMRRRKYAKRGYRRKCYGSRWATTTCWCCTGYRATWYKK
47               KKKKYKKRKKYTYTRMRRTAAACAGATTGGRTYRRKTYKSRWGKAAMTMAKTYGGGAYKK
                 *.***** *** *.**** ..      . . * **** * ***..::    .. * .  ***

44               RYYMRKSYAGMWKKCYTMWMKTKTGYGKTKTTTGRARKWKGMRRRYRRYWRRYSKMWYKY
47               RYYMRKSYCTMWKKYYKMWMTGKWSCTKKKKWGAGGAKWKKMRRRYRRYWRRYSKMTYKY
                 ********. **** *.***. * .   *.*.  . . *** ************** ***

44               GMRMSMYGTTKWYWGTGKYWWGTYWYKKYYMKYMYKKRRKKMRRKCWCTMYYRRMSYKRR
47               AMRMGCCRYYKWTWACSKYWWAYYWYKKYYMKCACGKRRKKMRRGMTTCMYYRRMSYKRR
                 .***.      ** *. .****. *********    ********     ***********

44               WKCKRGAMYMTATKYCRWMRKRGATRSMKRKGAGRTTTMKAYKKRKYTTKWGTGGMMKTR
47               WGGKRKMMYMYMYKYARWMRKRRRKRSMKRKTTCRKKYCGTYKKRKCYYKWRAARCMKWR
                 *  **  ***   **.******  .****** : *..   :*****   ** :.  ** *

44               KCWMRRRRRRATCYKRAMGRKMYRYMYRSYGARRRRWRYTAKTAKYKYCTAWKGMRCTYR
47               TMWMRRRRRRGAYYGGCCRRTMTRYCYRSYKRGGAGWRYWWKYRKYKYMYRWKRAGYWYR
                 . ********.: *   .   *.* ** ****      ***   *  ****    **     **

44               RRWYGYYYRWRTGRRGMWTYKRYYRYCCGRYTYRAAAYMYKWYRARRRKKMYYWKKRRYM
47               RRWYRTTTRWRGARRKMWGYKACTGTTYRRYYYRGTRYMYKTTAGGRRGKCTCAKKRRYM
                 ****    *** .** ** **        ** **.: ****    . ** *    ******

44               GYWTYKKKMACATYRRKCKWRCTKMWYMTGGWYRYCSYCGGRCYYKYMAAGTGTCARKRY
47               KYTACKKKMGTGCYRRGYKWRYYGATCCWRAWYRYYSYTTKAYCYKYCRRRYSKYRAKRY
                  * : ****. . ***  ***          .**** **       ***      ..   ***

44               RKKYYRKYRYMYKMKYRKAGAYCKYYMKKGRWYGKCRARSKYYCGKCAKTCTATCYYSKM
47               RKKYTGKYRYMYTMKYRKTTGCTKYYMKKKRWYRKMRMRCGTCYRKGGKWMYWYYYYSKM
                 ****  ******.*****: .  ****** *** * * *.      * .*       *****

44               KWWARRRCYRYWARYKWMYMTAYATRRWKGWCTKKTRRKTKYKRKKKRRKKYRWCGCAKK
47               TATGRRRYCRYWMRYKWATMACCGARRWTSWYYKKKAAKWKYKRKKKRRKGYRWMRMRKK
                 .  .***  *** ****  *:. .:***..*  **.  * ********** ***     **

44               YYTGCKKRRRTWRYRKKRRTRWKYYMKYMCGRKYRTKCKARYYYMYWRKTTRYWYYRKKY
47               YYCRYKKRGGGWRYRKKRRWRWKYYMKYMYTRKYRWGMKWRYYYMYWRKKYRYWYYRKKY
                 **   ***    ******** *********  ****    * *********. *********
```

```
44                YTCGGKKKKYSKWCMRAYKYYKKRSRRKGKTMRGAWSYWRGTMRKRYRRCKMKRRWMATA
47                YWATAKKKKYSKWAMRGYKYYKKRSRRKRKKMRKRWSYWRACMGKRTGGTTMGRRWMCAG
                  * . .********.**.*********** *.**  *****. * **    .* ****.:.

44                YKCRGMYKKRRSYTRKTTAYKRKKYRKYKKMTTTKYKKYKWYKWKMYYYWGRMRYRATAT
47                CKTGACCKKRRSYYRKWYMYKRKKYRKYKKMCGWKYKKTKWYKWKMYYYWKRMRYRGCGW
                   *  .  ****** **    ************   **** *********** *****. .

44                RYWRKKKKSRRRYKAWRTAARGTTKKMRKTMKWGAARRRKGATKCYCKWYYGGKKYSYWC
47                RYWRKKKKSRRRYKWWRYWMRTACKKMRKCMKWRWRRRRTRGWKYCYTATCTCKGYSYWM
                  ************** **   * : ***** ***   ***. . *   .     * ****

44                TRATTYRRKAKYRRACARRMYKYRYKKMYYRMMMWYRCAKRRCGMCGYKWRGWYWYKRWY
47                AATAWYRRKRKYRRMYCRRMYKYRYKKMYYRMMMATGSCKRGAAMSRCKWRKWYWYKRWY
                  : :: **** ****  .*****************   ..** ..*.  *** ********

44                RRYWYKWWATCTTAAAMRRYGKMAWCAAYYKKKWSCRKYKKRKKRRTAYYKRKYYMMMKT
47                RRYAYKWWMWSWWWMRAARYRKCGWYCRYYKKTWSYRKYKKAKKRRYMYYKRKYYMMMKC
                  *** ****  .       ** * .* . ****.** ***** ****  ***********

44                RRRYRYYAKAYYKKRKYWRRMKYKWMMKKWMMYRKWTKYYMRKKKGATGTAYYYRKWYRR
47                GARYRTYRKMYYKKRKYWRRMKCKWMMKKWMMYRKWCGTTCAKKKRWKAYRTYYRKWYRR
                    *** * * ************ *************      ***  ..   ********

44                KATRSRRYRKRGRCYKRMMRRKKYKRYWWCYKYSTMKRAKWYRKKKYRYMRKKMYKRKRR
47                KMWRGAATRKRCATYGGMMRRKGYGRYWWYYKYSCCGGMKWYRKKKYRYCRKKMYKRTGR
                  *  *.   ***   *  ***** * **** ****     ********** *******. *

44                YKYSYYYRCTGSYYKARKMCCGTKKKRMCYYYKGKYRYKKKKKKWRMGGKRYRKMKRRYK
47                YKYSYYYRYASSYYKCRKMYYAAKKKRMMYYYKSKYRYGKKKKKWRMKKGRYRKMKRRYK
                  ******** :.****.***  .:***** ****.**** ********   **********

44                KYSYWYRKKRKKWYRYKRKTYMMRWRKACTKKGCKYYMYRRRYRWYRKKRRYRYYKYKYA
47                KYSYWYRKKRKKWYRYKRGCYMMRWRKCAWKKAMKYYMYRRRYRWYRKKRRYRYYKTKYM
                  ******************  *******.. **. ********************** **

44                WKKYKYYSKRYTKMRGCGCCYGCTKKCYGKMYYMYRSMRKMWYYRRYYSRYKYYMYMCAR
47                WKKYKYYSKRYYKMRKYKYYYRMYKKMTRKMYYMYRSMRKMTYYRRYYSRYKYYCYMYRR
                  *********** ***     *   **   ************ ************ **  *

44                YSKKKKMRKWRKYSTYKKRWYYYYRYYKKMRRGKKKMATCRKMKRKWKKMKYMKKKRRWA
47                CSKKKKMRKWRKYSYYKKRWYYYYRYYTGAGRRKKKMCGYRGCKRTWKKMKYMKKKRRWR
                   ************* ************.   * ****.  *  **.*************

44                TTGMYMMAGYWKRYRRYKWCWMYYKKMYRKCAGYYGAGMRRCYRMKMRWSRYATCTSACS
47                YYRMYMMWKYWKRYRRYKWYTMYYKKMYRKATTYYRTAMRRAYRATARWSRYMYYKSGYS
                     ****  **********  *********.: ** :.***.** . *****   .*. *

44                TKYYKYKRRAWYRKRYCMMKAYGKGATTKRWYCCRKRTAKRKKGYWYRYRAACARYGAGG
47                YKYYKYKRRMWYRKRCMMCKGYKGKRKYKAWYYYGTACCKRKKKCWTGTGGRTMGCKGAT
                   ******** *****  * *.*    . *  **   .  .****  *    .      ..
```

```
44              AGRYYRWKKCMYRRRTMYYKYKYTCGRAWYRWGGGCCKAGTCRRRMRMYRYMMRYKKYYT
47              TTRTTRAKKTMYRRRYMYYKYKCWMKRRWYRWRRRMTGWRWYRRRMRMYRYMMRYKKYTY
                : *  * ** ***** ******    * ****          ****************

44              SAAKKGTCGRAWTKWYYKRMKRYRRKGWMMYRYMKYKYRMKKCTWYYKGGCTTGGAGCYT
47              CGTKGACAARWAYKWYYKRMKRYRRKKWAMYRYMKTTYRMKKTCWYYKATYGYSSTTTYK
                ..:* . ..*   ************* * ****** .*****  ****.    ..:  *.

44              KATRKKKAMTRYKRKKYMRKRATAYKGGRYKKRRRYKKYRKMYKSRRRYRWYRYKGGKRM
47              GWYRKKGGAYRYKAKKYMRKRTAGYKTARYKKRRRYKKYRKMYGSRRRYRWYRYKKRKRM
                   *** .  *** *******::.** .*************** ***********  ***

44              MSYKYKSAYKYKKWWKTRKKRMGRRYGGRWKKTRWRMRYMKRGATKKMMKKACSRMMRSW
47              MCCKYKSMYKYKKWWKYGKKRMRRRYKRRWKKYRWRAGTAKRKWKKKAMKKTYSRMMRSW
                *. **** ********  **** ***  **** ***    **  .** ***: *******

44              SYKRKSRYRKMYWYKYKYGRAGRWCCMRSKTCAGWKKTAYMKSMRSMYAYGCAYKRKMRY
47              SYKRKSAYRKMYWYTYKYAGMAGTTTMRSKAAGAWKKGCYMKSMRSCTTTTGTYKRKMRY
                ****** *******.***.  .    ****:...*** .*******  :   :*******

44              RRSKKYYRKTYRKYYWKMYTKKRRRYKSRATAYAKSKMYKRYKKYYWRSKGRMYRKKYKM
47              RRSKKYYRKWYRTYYWKMCGKKRRRYKSGMWWYWGSKMYKRYKKYYWRSKKRMYRKKYKM
                ********* **.*****  ********    *  *************** *********

44              YYWYKKKRRRKMMYMKKYRKKTKYCCKKWCTMCWCKYKKRKRTKGCMKRKKYYKRYKYKW
47              YCWYKKKRRRKMMYMKKYRKKKKYTTKKWGAMYWYKYKTRKRKKAAMKGKKYYKAYKYKW
                * *******************.**  *** :* * ***.***.*..** ***** *****

44              CCMRKGKKWYMYYKKKRWKYCTSWRKRCGYYKYRCCTMKKRRRTRRMWKRYKMYWWSRTK
47              YMMRKKKKWYMYYKKKRWKYYWSWRKRYAYYKYAATCMKKGRRKRRMWKRYKMYWWSRYK
                  *** **************  ***** .**** .  *** **.************** *

44              YTGKKYKYYKCCGYRTGGTYRRKWCATTWMYWWYMKRMSYKYRRRCKYYYMAKGKRWKRG
47              YWTKKYKYYKAAAYRYKKYCGRKAYTYGAATWWYMKRMSYKYRRRAKYYYAMKTKRWKRK
                *  *******...**      **  :     **************.****  * *****

44              MRWCYWRYYCGGGKCMRYRYWRCKACAYKYMYYKTGCYMKKKKSKMYTGTKWGRMYARKK
47              MRAYTARYYAAAAKYCGYRYWRMGTYMYKYMYYKKKAYMKKKKSKCCATCKWARMYGRKK
                **    ***....*   *****  :  *******. .********  :  **.***.***

44              KKKWRWAWYRMYRKKYWACRWRRKYTAWGYTMYKCTCSYRKYYGKSWKKRMYYWRKRRKC
47              KKKWRWGWYRMYRKKYWMYRWRRKTCGWRCCMYGMKYSYRKYCRTSWKKRMYYWRKRRKY
                ******.**********  *****  .*   **  . *****  .**************

44              YKKKKSWRYYKKRWCRYGYKWSYYKRKAMKKATGCTKYRYKRKYWYYYRRRRMAGRRRCG
47              CKKKKSWRYYKKRWTGYKYKWSYYKRKGMKKRKKMWKYRYKRKYWYYYRRRRAWRRRRGA
                 *************  * *********.*** .   ****************   *** .

44              AGCMYYMRYRKGCCRGTKYRSYWYMMYYYYYRKYRKKAAAKRMKKKYMKKYRKRKKKYKS
47              RRTCTCAGYAKATYRCCKYRSYWYMMYYYYYRKYRKGRWRKRMKKKYMKKYRKRGKKYKS
                        * *.  *  *******************    ************** *****

44              AYTKRRGRTKSRKKSKTGKRYWRMRCWRRKYRYKAWYWWSYYKRWKRYMMMKYRMRYMAA
```

```
47          RTCKRRRAAKSRKKSKYRKRYWRMGMTRRKYRYKMWTAWCYCKRWKRYMMMKYRMRYMRW
               ***  :*******  ******   ******* *  *.* ****************

44          RYWWYYKRKKRRYGTGRRACWSRYRKKAATGKYYYYRRYRMRRRRWRYWYYMKKKRKWYY
47          RYWWYYGRKKRRYACRRRTYWSRYRKKRRYSKYYYYRRTAAGRRRWRYWYYMKKKRKWYY
            ****** ******.  **: *******   .*******    ******************

44          RYKRKRWKTYWYKTGKRYRKYMRCMRTCARRKKKYYRYYYSAKCKKMSTCGMRRKKWKKW
47          RYGRTATKYYWYKWKKGYAKYMGYARATRGATKKYYRYYYSGKMKKMSGTRMRRKKWKGW
            ** *.  * ****  * * ***   *:    .*********.* ****   ******* *

44          CRYWRKRRRKRYRYRGKAWRYYCMMWRYKYKKKAKYKKKAYKRTTRYATCTMCCYWKKKC
47          YRYWRKRRRKRYRYGKKRWRYCMAATGCKYKKKGKCKKKTTGRAGGTRWYKCTAYWKKKY
             *************  * ***       *****.* ***:  *:      .  .*****

44          CYWTWGWYYYMWASWSYRMWYARTMWYKMKKYYYRTAKGWYKTTGTGYTAGGTKYMKYWK
47          TYTGAKAYYYMWWSWGCGMWYGRYCACGCKKYYYRWRKAWYKWYRKKCCMRRKKYMKYWK
             *     ***** **.  ***.*      ******  *.***   .      .*******

44          YYKYKYRCYYKKRKTTWMRMTCRKRTMKTRCTCGYMRKSKTCAAAAYMYKRGYKYAGKKR
47          YYKYKYRYYCKKRKKWWMRMAARKGYCKGRAATKYMRKSTYYTTWRYMYKRRYKYMTKKA
            ******* * ****. ****:.**   * *.:  *****.  ::  ***** ***  **

44          YTYWTRYTYRMRYAKKCRYGYRKRKCAWRKYRYGTGYKYWMKYKMKYWWAGGCKYAYTKW
47          YCCAAGCGCACGCRKKYGTAYRKRKGGWRKYGCACKYGCWMKYGMKYWWGCAAKTWYKKW
            *   :         **   .***** .****  .  *  **** *****. ..*  *.**

44          KTTAMKMMRKTRSWAYYKRAKACYTRGMRRKAGYYKCGGGKATTGAKMWKGKCKYWWKKK
47          KCKRMKMMRKKRSWMYYKRGGTTCCGTMRRTTTYYKYTATTRYWRRKMWKTKYKYWWKKK
            * . ******.*** ****. :     ***.: ***  . .     **** * *******

44          RRSCYKRYKRYYKSKTAAKGRYRYYKKRRYGYYAMMTYMACACKKYYYMRMMKKKYYSTK
47          RRSYYKRYKRYYKGKCWTKRRCATTKKRRYTYYMMCCCMRYRYGKYYYARMMKGKTYSKK
            *** *********.*  :* *    ***** ** *   *     **** **** * **.*

44          YYWKYTYYKRRRYWYYYKKKSRWKRYYRKRKYYYKKMGAWKCCYYR
47          YYWKCCYYKRRRTWTTCKKKSRWKRYYRKRKYYYKKAACAKYYYYR
            ****  ****** *   ******************* .. *  ***
```

The pairwise alignments of two accessions 44 and 48

```
44              RTRRGKMMYCCAGCYSRRYTRMGMKKKYCCMMCRAWWKWKACTCCYMWYYGYYRSGAMCG
48              GKRRKKMMYYYWKYYSRRYYRMRAKKKYYYCCYRMWATATRMYYYYMWYYACCRSAWMGA
                 .** ****     ***** **  ****     * * . .     *****.  **. * .

44              WKGGGTAMYRTGGMKMYSMYCTMYWRKRKTRGMMWRYKYRWWKKARYYYYYKYRGKYMGK
48              WKKRKWMMYRWRSMKMYSMCAAMYARKRGGAAMMWRYKYRWWKGGGYYYYYKYRAKYCRK
                **     ***  .****** .:** ***   .*********** . ********.**  *

44              AKWRRRRYWYWRRKWYYAMKYCCRYYRWCMMKYGWRYKGCYSWYKYYRRGSAYMRYGRTG
48              RKWRRGRYWYWRRKWYYWMKYYYRYYRWTMMKYKARYKRYYSWYKYYRRTCGYMRYRAWR
                 **** *********** ***  ***** ****  ***  *********  ..****

44              MRAARWAWKSRYMRKMYRRATAGWRRRYYYAWGRRYRRCGTRMRYGYYYYWKWMCYYSWR
48              MRGWATRWKSRYMRKMYRRGGTTTRRRYYYWWRRRYRRYAWRMGYRCTTYWKWMYCCSWR
                **.    ************. :  ****** * ***** . ** *    *****   ***

44              RKWRYRWYKTYKKWMRAYAYMRRKYSSYCCTRYMYYWKYYRGYKYWKRRYYWWYYYRGGA
48              RKWRYRWYKAYGKAAGCCGYMRGGTSSYMTWRYMYYWKYYRRYGTAGRRCCATYTYRRCG
                *********:* *   . .***   ***   ********** *    **    * **  .

44              CCAATKYYWCRGKYMAACKRYYYYKSRYYRCRYWWCRCYMRMWMKYMKAGMKRMMWTRRK
48              TTWWYKYYWYRRKYMTGTKRCYYYKSRTYRYRYWWGATYMRMWMGYAKCKMKRAMWWAGG
                     **** * ***:. ** ****** ** ****   ****** * *. *** **

44              RKKRRWYYCRCWKTYYGGYKGRKYRWYWWKYGKKYYKRMTYKKAMCMCGMRMKYRCYMWY
48              RKKRATCTYGMWKACCRKYKTAKCAWYWWKCKKKYYKRMAYKKGMTCATCGCKYAMCMWT
                ****       **:    **  *  *****  *******:***.*  .    **   **

44              YSYYRGWAWKWTAAKMYWRWGTCKKYRTGTYASWYKKWYYYKWRSGTCCCAGGRWYYYAA
48              YSYYRATGWKWCGTKMYWRWAAAKKYGAAYYRSWCKKTYCCTWRSRWYYYGARRWYYYWR
                *****. .*** .:******.:.*** :. * ** ** *  .***     .. *****

44              ARWRKKYCKCKKYKKRKKRKMTAMRYRKMRKRRGYKMYKYKCGKCKMKYGGGGRYGGKKY
48              WRWRKKYYKMKKYKKRKKRKAWWMRYRKCRKRRRTGMYGYKMRGYKMGTKRRRRYKSKKT
                 ****** * **********   ***** ****   ** **    **      ** .**

44              KKKTMRCKYWRKKSYWYYYKRKKTGYYRKYYRYKKCRTKYMWKMRRKMWKKRKKYMRYWR
48              TGKCCGTKCWRGKSYWYYYKRKKGAYCGGYYRYKKYGKKYMWKMRRKCWGKRKKYARYWR
                . *    * ** *********** .*   ******  .********* * ***** ****

44              KKCYATYRYYMYRKKRRYRTTATYYKKYKKMRWWTAGKKKRYYYSTTGKRYYYRYRKGGR
48              KTYYRWCATCMYRKKRRYRYKRYYYKKYGKMRWWGMRKKKRYYYSYCRKRYYYRYGKAAR
                *. *      ********* .  ***** *****   ********   ******* *..*

44              MKKCTKMYYSMRKMRRYYYRYYRGYKYCSAGAKCGKKKWWYRMRKYYYYTRAWYAKRRGA
48              MKKYKKMYYSMRKMRRYYYRYCARTKYTSGAWGTKKKKWWYRMRKCCTCGAGAYCKRGRR
                *** .****************    ** *..    **********      . *.**

44              GRRAAMYWGTGKWCATKKWYYMYSYWGATYYRRGCAKYCATRKRYMGYKYRYKKRCGTCW
48              RRRGGCYWAATKTMTYTKATCMYSYWRRWYTAGRMWKYYWYRTRYMKYKYRYKKRTTCTA
```

```
                 **.. **.: *  : .*   *****   *      **   *.*** ********

44              TSMKWRTATATGMRCYKTYRYRYTGKKRGACYYWYRARTCGYAGTTCCCYKMGCAAAYYS
48              CCMKWRCRCCCTMRTYKKCGTRCCTKKGKRMTYWYGCRCGAYRRYCTTYYKCCMWCWYYS
                 .****   .  ** **.   *   **     *** .*  .*       **    . ***

44              MGRRCYMMWGWKYYKKYCGYMGRKRKKKYKKRKCCCGKYRKMYMYRMKKAGAKKATTCKT
48              CRRRYYMMWKWKYYKKYTACCKRKGKKKYKKGKAYYRGYRKMYCYGMKKTTGKKMKKAKA
                  ** **** ******* .   ** ****** *.    ***** * ***: .** ...*:

44              RWYGKCRAMTKGGTYGMKRYMRRKYMRYWWMYTYRRWCCATRGTCARKKTTAAKSWYWKK
48              ATTRKMRMCYKATGCKMKRYMRRKYMRYWTATAYRGWATTCGAYMWAKTCWGGKSWYAKK
                    * *   *.    *************   :** *. :  .    *.  ..**** **

44              KKKRKKRRRKYRWCGCARWYKRKYWYGMMMRKKRYRYWMGGYAAWKWYTWKKYYYTGCYK
48              KKKRKKRRRKYRWMRMRRWYKRTTWYRMMMRKKRYRTWMSAYWTAGWYWWGKYYYCTTYK
                *************    *****. ** ********* **..* :  ** * ****   **

44              YMMYTCKSTGRRMYRRYRKKAGCGAACWTAMACGCYAGGGRWKWYRKRRRMKRYRTRKTW
48              CMMYCMGCKKRRMYRAYRKKRRYATTATATCGSATYWKRKRWKWYRKRRRMKRYRYRKYT
                 ***   .. ***** ****   .::. :: ... *    *************** **

44              TRCAMCTYTAKSYTATMCARAYMRWYYKGCKRYYYAGAKRYWYRRKCTATMYWYGMAYMR
48              KRYTMMKYCCKSYYRWMYGRWYMRWYYKATKACCTGSWGRYTYRRKACTKCTWYKMRYMG
                .* :* .* .***   * .* *******. *    ..  ** ****. :.  ** * **

44              YKKYRMKRSRRKKTTCCYKRYRKKKSRRWAYRGMTMWYKKRYKKRMWMSKYRKMYKRRYR
48              YKKYRMTRSRRKKWCTAYKRYRKKKGGRWRTAKCWAWYKKRYKKRCTMSKYRKMYKAATG
                ******.******   .********. **       *********  *********

44              WYRYRRTKYWYMCRCYKMKKKTYKKGARRKMKRWRCTATYRMKSMRTRYYRYMRWWYKTT
48              WYRYRRKKYAYMAAYYKMKKKKYKKKGRRKCGRWRMWWYYRMKSMRCAYYGYMRWWYKYG
                ******.** **.  ******.*** .***  ***    *******  ** *******

44              KKGAKTATTAYKRKKYMMTWCRRGKKYTCRGTKRYYMRGCTAGCYCAYAGKRTTGYCCYY
48              KKAGGACCGCYKRKKYMMCTYARKKKYKTRAWKRYYMRAAGRTTTAGCGSKRYWAYYTYY
                **.. :.  .********    * ***. *. ******..     .. ..**  .*  **

44              CGCKKKTRMRYAMKKYYKMWMKYMKTMYMYMTWRRTYKWKGKSYKKATMGYWRKCAAGKR
48              YSYKGKKAMRTRMKKYYKMWMTCCKWCCAYCYWRRCYKWKKKSTKKMYAKYWRKTCMATA
                 . * *. **  *********.  *    *  *** **** ** **    **** . ..

44              YKRMYRKYMRYKYKYRWKRRKRGTRMMKKMYYYSSMRKWWGKMKKKTKKSSMCACKRRYY
48              CGRMYRKYMRYKCKYRAKRRKRRWAMMKKMYYTGSCATATCKAGKKYGKSSMYRYKRRYY
                  ********** *** *****   ******* .*  .   *  **  ****   *****

44              KKWKYYYMYYYYRMRRWYSKKKRYKSGKRKYYWMCTMWYKRGGTKWKYRWMCCATCTRYK
48              KKWKYYYMCCCTRCGRWYSKKKRYKSRKRKYYWMYYMWYKRAACKWKYRWMYAMWYYGYK
                ********    *  *********** *******  *****.. ******* .     **

44              KRMRKWMYRRCGRCYTKMKWMCAGMRYYTYYKKYAKWMCAATKRRRRKKKWSWKKKWYTG
48              KRMRKWMYRRYKGYYKKMKWMTTKMRYYKYYKKCCTWMSMMWKRRRRKKKWSWKKKWYCA
                **********    *.***** : ****.**** ..**.   **************** .
```

```
44      CRWMYAKCGAYKMGGTKKAKRRRMMYCYKKRKYKYYKCGWKKYTKKKYKRGYTTATTKCA
48      MRWMYMKYKRYKMATYKKMKRRRMMYYYGKRKYKYYKYRWKKYKKKTYKRTYYYRYKKSW
         **** *   ***.  ** ******* * ********  ****.**.*** *    .*.

44      AKRRKKKRYWYKKRGMWYRRKCTAKTKYWRWYKYACGYKKTMYYYKSGWRCRKKYRKKMT
48      RGRRKKKRYWYKKARCWYRRKTYRKYKYWGWYKYWYRTKKWATCTKSRWRYRGKYRKKAW
          ***********   *****   * *** ****    **     ** ** * *****

44      TKKKRRRRWRRMYMWKKMYWWSKCACTKYKSKKSRGYTAAKYWTAGCTWYYTMRMSYRKC
48      KKKKGGRRWAGMYMWKKMYWWSKTTTCKYKCKKSRKCYRMKCAKCAYCTCYCMRMSYRKY
        .***  ***  ************ :  ***.****     *  ...    * *******

44      RWWYSKYMRGTKYGTRKKKWWYTRRKRYYYKKWYYYGYKCCTGKARYKKTCMGATAARTA
48      GWWYSKYMRRYGYRYRKKKWWYWRRKRYYYKKWYYYKYKGTAAKWRYKKWMCKTCTTRCG
         ********   *  ******* ************* **  :.* ****    : ::* .

44      CAKRYYYRKWYYYYWSCGCYCMRYSCWMMWYKYAWRRTRYMWYKRYYYRKYKGTTCGTCA
48      YGKGYYYRKAYYYYTSYSMYTMRYSSWMMWYKYGWRGYRYMWCKRYYYRKYKSYCYRYYT
         .* ***** **** * . * ****.*******.**  **** *********.      :

44      YWKKGTYKSYRCRTGRCYTARSKKYRKKCRTAGTTRATGKKKRRMRTAWRYYKKRWRYMM
48      YWGKKWYKSYRYRKARMCWGACKGYRKTYRCWKYCRCYTKKKRRMRCRARYYTGAARYMM
        ** *  ***** *..*   . .* ***. *     *.  *******   ***.   ****

44      YMRTATGTYKTWKMCWAYKKKKKYRYKRKYWKYYYKYRRRRCAACGCGRRKYYMRYWRKK
48      YMRKCGAGYKKWKMTTTYKKKKKYRYKRKYWKYYYKYRRRRYWTYRMARRKTCAGYTRKG
        ***.. . **.***  :************************  :   .***    * **

44      AYKRSYWYKMKYYYYRYMSKRTGKWWKRYRWKKAKKWYMRCWCYKRKRKYKSKTAARRRK
48      RYKRSYWYKMKYCYYRYMSKRKRKWWKGTGAKKWKKATMAYATYKRKRKYKSKCGGRRRK
         *********** ********. ****    ** **  *    ********** ..****

44      MRRYRKYGSRGKSKKRKMYRKKYYSRGGMRKGRWWYWARCGKKSRYKWRYYYYMYCCGRT
48      MRRYRKYKSRRKSKKRKMYRKKYYSRTAMGKRRWWYTRRYAKKCRYKWRYYYYMYMMKGA
        ******* ** *************** .* * ****  * .**.***********    :

44      KRYRMWKCTYAAYYYYSGKWMRYAGCTYCGCYRSRRYKRAGGAKGYWRTTTRKMKRTYKR
48      KRYRMWKTCCGRYYYYSAKWAGCCRAKYTATYRSRRTKRCAKRKKYWRGYCRKAKRCYKR
        *******   . *****.**   . ..* . ***** **..  * ***   ** ** ***

44      RWKGRAKATRMMYMKRRSMKKRKKKYRRRKKKKKKYKKWYKCWKKMYYYWRRRGMGGRRY
48      RWKAARKTARMMYMKRRSMKKRKKKTAAGKKKKKKTKKWYKMWKKMYYYWRGGCMRKRAC
        ***.  *::****************    ****** ***** *********   *  *

44      AYRTTKYRKMYKKKMKWACMYMCTCATYRKRGKTYYMRMWMRKWMARRWKYRMGGRYYGR
48      GYRCCKYGKMYKKKMKAWMMTCTCTGGYRKAKTACYARCWMRTWMRARWKYGCRKRYYRR
        .**  ** ********   *     . ***  .: * * ***.**  ****    *** *

44      ATAKKRTGKTATCCTAKAYMKRTTYARYGYYRKRKRTKATCAKYCWATATWRRYKYCWGT
48      RATKKRWRKYTYYTCGKGYMKRKCYGRYKYYRKRKRWKRKSRKYYWGWGCWGGCGCMTRC
         ::***  * :    .*.****. *.** ******* * .. ** *. . *
```

```
44    TYCMRMMKMKAYYGTMYTRKYGRKGRGKKKKCCGGGKRCAGATWYCGCTYKKYTRYGKYM
48    CYYARAMTMKGCCACAYCGKYRRKKRKKKKGYAAAAKRMWRWWAYYSTWYKKYARYRKYM
       *  * *.**.  .  *  ** ** * ***  ....**      * .  ****:** ***

44    KKGKYMAMWMYTGKRKCKKMKRYWATRMYRYKWYWYRRYGCTCSYSKYRRGASWKKWKCA
48    KKKKYMWMWMYWTKRKYKTMKRYWWWRMYRYKWYWYRRYTMKYSYSGYRRRRSWKKWKAC
      ** *** ****  *** *.*****  *************  . *** ***  ******..

44    TGTRATTRCYYKYYRYRRWWKWYTTCMRTYMRSYKYRMRRTYMRKWWYKKKKYWYGAGRG
48    CAYRWAWAYYYKYYRYAGTWKWYYCYMRYYARSYKYRMRRKYMRKWWCKGKTTWYRGAAA
       . * :   *******   ****   ** * *********.****** * *. ** .. .

44    YYKRRKKWYRRYRMWSYRRKRRGYRWYRTGKRYCWRMAYKKKRKRWWYKKKKKYWYMRRT
48    YYKRGKKWYRRYRMWSYRRKGGAYGWYRWTKRYMWRCRYKKKRKRWWYGKKKKYWYMRRY
      **** ***************  .* ***  *** **  ********** **********

44    CGGAYYKKMRKYYYYYSYYWKKWWKMYTGCRRKRWATWWRRYKKRWYGKKRYYKRKKMRY
48    YRAGYYGKMRKYYYYYSTTAKKWWKMYCATGRKRTTWTWRRYKKRWYAKKRYYKRKKARY
        ..** **********   ******* .  *** :  *********.********* **

44    RKGKKRACCYGGSRCRAYTMMRSYKRSRRWSWTWYKKCTTYTCKYMTGRKRRRSGRKYMA
48    RKKKKRGAGYKRGGYRRYYMMRSYKRSRRWGTKWYKTAWYCYYKYMAARTRRRSKRKYMR
      ** ***.. *  .  * * ***********. .***..     ***:.*.**** ****

44    WKKYWYRTKKRAAAMRKKYYRSKKGKSYYRTYRKGGAKYYWKKKKYRRRYGSRKRSWYMY
48    WKKYAYRKTGAGTTMRKKCYRSKKRKSYYRWYRKACTKCTTKKKKYRRRYKSRKRSWTMY
      **** **..  .::**** ***** ***** ***. :*   ********* ****** **

44    KTYYWRRMYKCKRKYKYKYWKRSYKTWYYYWYRGGAGRWRYRYYKRWRKARTYYRRRKMM
48    KKYYWRRMYKGTRKYKYTYWKASYKCTYYYWYGRSMKRWGCATTKRWRKMGAYYAAGKMM
      *.******** .*****.*** ***  *****  .  **     *****  :**   ***

44    GCKYYGARMMKRKKYGCRMSKGKATRYMTCYYYRWRRWKYKMKSRWKWWTRYSMRATGGC
48    AMKYCRRRMMKGKKYSYRASKRKRWRYMCMCTTGWRRWKYKAKSRWKWWYAYSMRRYKKY
      . **   **** ***. * ** *  ***      ******* *******  ****

44    YKYKYTKKKKMSMWRRCRYTACKYCAGWTAAATAMAYRWTRYTYTYCKRARYRAYRRATR
48    YKYKTCKKKKMCAWRAYATYRYKTATAAACGTKGMTYRWKRYYTWCMTAWRYRMTRRGAG
      ****  *****. **       * .:. :..:..*:***.**     .  ***  **.:

44    TTMMRYYRTAAGAGKWYRYRWRKKYRTCGCCTTTMYYYRMMYYCKGWRRYMMWRTGRYYR
48    KCCCRYYRWRWRWKKWYACRWRGKYRYYAYYWWWMCCTGAAYYMKRWRRYMMTRWRRCCR
      .   ****      ***  *** ***  .     *      ** * ****** *  *  *

44    TSTRRYCSGCGRYWATGTAYAGTCCCARYKYRRKWYYYSKRRYWYRYRKKKKRWYKYMYK
48    WGCRRCMSATTACTGYSKWCMKKYYYRAYKYRRKWYTTSKRRYWYRYRKKGKRAYKYMYK
       . **  *.     . ..    .     ********  ************ ** ******

44    RYRRKRRYKYRWMYYRRMYGKKMKRYRYMMYGGAMRMRSACAWMRKRGATKKRYRRSMKG
48    RYRGKRRYKTGTMYYRRMYRKKAKRTATAAYATTCGMASRYWWMRKRRRKKKRYRRSMKT
      *** *****   ******* ** **     *. :  * *   *****  .*********

44    AGRYYMTKAYKTCWGTTKWYSWRKGRGYTRCARGCYYWYYRKMRRRYYARYTTYYKGGGT
```

```
48          TCAYYCCGGYKYYWSWYKWTGTGKRRRYYRYWRAATTACYRKMRRRYYMAYKGYYKAATC
            :  **   .**  *.  ** .  * * * *  *..    *********  *. ***..

44          YWRYRKYKWYKKCWGYGYMRGSRKARWSRYATCTCGSMYSYYSAAKYYYAKYKKCYMYSR
48          YWRYRGYKWYKKAWTYKYMARSRKWRWSRYMYYKASSMYSYYSRRKYYTCKYKKGTATCR
            ***** ******.* * **  *** *****   ...*******  *** .****    .*

44          AYWWYMYCRRKKYWMYYCSAKTATTRRTGCTGAGKRKMGYWGGGTRRKYRACGARYGTYT
48          TCWWYMCGRRTKYWCCTYGRKYRKYGRYAYCTCTGRKMKYAKCTCRRKYRGAAWAYKYCW
            : ****  **.***    . *  .  * .   .  *** *     *****...  *

44          YYRYWGCAAYGKWMYKRKRRGRKKYKYKMMMKYKRRGTRYRATAGKKMKYWTGCTWAGCG
48          TTGYWRTCGCTGAMYKGKRRKRKKYKYKMMMKYGRRKAACAWWMKKKMKCWYAGWAGKAT
               **  ..    *** *** ************ **  :       **** * .   . .

44          CWMMKKKMKKWYYCMRYYRAGYWYRRKAYRSRRYRKRGKMTMGRAKYYWKRRRWKKKWYA
48          MWMMKKKMKKWYYMCAYYRGACWYRRKMYRSRRYRKRRKACMARWKYYWKRRRWKKKTTW
             ************   ***.. ***** ********* *  *.* ************

44          AYKWYMARKRYAGAMYKRYYSYYYRKSYKYCGMMWRKKTRTWTYRKKSKMMRKAYYKRRT
48          CCKTTATRKRYRRMCYKRYYSYYYRKSCKYTAMMWRTKAAYTYYRKKSKMMRKRYYKRGA
            . *   :****    ************ ** .****.*:    ********** **** :

44          AYKYKRYYKCYRYMKYCGMTRGTRTKGKGCGGKKKCTWRKYCYMWMMGTYKKYTYRMKWW
48          GYKYKRYYKMYATATTARMCGTCACTAKRAATKKKAGWRKYSYMWMARYYKKTWYRAKWW
            .******** *   . . *      ..* .. ***. ****.****   ***  ** ***

44          TKYYKKGGAGTATCYGRKYTARACCTCAMCRRGGRCRCKSKTKYSWYACYRGMMYTRRAC
48          YKCTGTRSRKYWKYYRRKYYCGRTTAACMYAARRAYRMKSKKKYSWYRTTRKMMCCGGMT
             *   . .    . * *** .    :..*       * ***.*****   * **

44          GYMKYKRYCCTCRYWRCYKRKCCGMRMRGKWSRKKRKYRYKMRMCAKWYYYGYRSSYMRM
48          AYMKYKGTTACTGCAATYKRKMMRMRMGKKWSRTGRKYRYKMRMGMKWYTYKYRSSYCGM
            .*****   .       ****   ***  ****. *********  *** * *****  *

44          GARRWKCYWRRKKRCYCATKTGTRYYKGTGKRRSYYYWMMYKRYKKKMYRYYKRATKMKY
48          KRRRWKYCTRGKKRTCYGCKYTARYYGRWRGRRSYYCWMMTKRYKKKMYRYYKRCAKMKY
              ****   * ***   . *  :***     ***** *** *************.:****

44          YMYCCKWRYRKKKRYKKYKKWRRKRGKKYKSWK
48          YMYYYKWRYRKKKRTKKYKKWRRGGKKKYKSWK
            ***  ********* ********   *******
```

The pairwise alignments of two accessions 44 and 49

```
44              ATKMMYYGCAKMYCYKGCRMRYCYRMMYRTMWWRRWYYYMWRYYYYCGGRKYMMYYKYYR
49              RYTMMYYATWKMCTCTRMRMRTAYRMMYRAMWWRRWYYCMWRTYYYMRRRKYMMYCKYYR
                  .****.  **   .  *** .******:******** *** ***   ****** ****

44              YRYKYRRRYMRWTTSCRGKSYKYKRWWWWMRYWKKKRMRGYKWYRYMMTYYYYMRYWMKW
49              YATGCARRYMATKCSYGAKGTKYKRWWWWMRYWKKKRMRRYKWYRYAMWCCTTMRYWMKW
                *     ****  . *  .*. ****************** ****** *     *******

44              MYSMYWRRCMMYRYAKKKYYYYYRCTRYWYAWRRWYYRYYKKRYRWYMKYYWYRWYYTCK
49              MYSMYWRRTMMYRYCKKKYYYYYRTARYWYWWRRWYYRYYKKRYRWYMKYCWYRWTCCGK
                ******** *****.********* :**** ******************* ****    *

44              WSWKYKYYRRSYMRYGYSKGYYMWYRCGTASACWAWKSYMWGGCAWRGRWKKGYKYWYRY
49              WSWKYGYYRRSYMRYKCGTRYYMWYRYKYTSMYAWWKSYMWRATGTRAAAGKRYKTWYGY
                ***** *********  .. ******   :*    ****** . . *.   * ** ** *

44              YMAYYYRKMMWKYWWRKMMYTMWWRYRYKSRMRYYYMYSYWRKYRWWMTTYSRRWTTYCG
49              TMRYTYAGAAWKYWWRKCMYYMWWRYGYKSRMRYYYAYSYWRKYRWWMCGYSRRWCCCTK
                 * * *    ******* ** ***** ********* ***********  *****

44              AYYMAYRRYYKRTCMYRYTYYWKRYGCYYYYCRRRTRMSWRRWMSRRKWYGCYWRYKMYC
49              RYYMWYRRTTTRCAMYRCGTYWKRYTTTTYYTRRRWRMSWRRWMSRRKWCRYYWRYKMYY
                 *** ***  .* .***   *****    ** *** *************   *******

44              TRKWAYMYYWWMGTYMKSMRWYRYKGGYYMYYYWYATCAKKACRRKGCYWKRSRKRMMRK
49              WRKWMYMYYWWMRYCMKSMRWYATTRKYYMYYYWYRCYWKKGAGRKASTAGRSRKRMMRG
                 *** *******   *******  .  ********    **.. **..   ********

44              KKYYWYCTTTRKWYRGKCMWGGYGCAYRYMWYMGCWMMKARYMWKTAKWKYAYTRGCRYG
49              KKYYWYYCAARKWYRKGYMARRYTTCCRYMWYMAMWMMGTAYCTGYRKTKYWTCRTMRYA
                ******  ::*****   *   *  . ******. *** : *     * **   *  **.

44              CCRSGTYGWRKYRSTKKRGTTTKGCAYCAYRTTTMRMTYRCCAKKKTRYKACGYWWWMYK
49              ATAGACYKWATYRSKKKRKKYWKKAGCTGTGAGCCGCGYRMYWKKGARYKGYAYWTTMCK
                .  .. * * .***.*** .  * ..  .  :      **   ** :***. .**  * *

44              KAAMCCCCYRWKYYKRKKRTYRMYKTRTGGGRRARRSGKGCTRCYKYRRGYMTAAWRCCW
49              KCTCGTTTYRAGYYKGGKRGYRMCGKAWRAARARRASTKTYWATYKYGARTCKRGTATYW
                *.:     **  ***  ** ***  .   ..*  * * *     ***     . .    *

44              YSWTRKKMRWGKRSYYTGRYAARWYYYTAAAYRRYATKRRKKYCGTRKKWGKRMYCKRWK
49              YSWWRKKMRTKKRSCYARGYGGAWYYYKWRWYGRYTCKRAKKYTKYRKKWRKRCCAKRWK
                *** *****  *** *:  *.. ****.   * **: ** ***   **** **  .****

44              KMKMRKMRRWTYGAAGCKAYYWCKYTWCCYMRCKSKGCTCAAMYKRCYKYKKKRYYAYRY
49              KMKCRKMRRWYTATCTYGMYYWTGYYTYAYMGTKSKAAYYWRMYKRSYKYKKKRYYGTRY
                *** ******  .:.    ***  *   .**  ***..    ****.*********. **

44              RMTRYKYYMTGKKWWRRKYWMKYMKYRYTCTKWWKCGKWKCGYRYTGACTARKKWAYSWM
49              ACAGCKYYMYKKKTWGRKYAMTYMGYRTCAGGWWKARGWKATTACCRCTCGATKACYCTM
```

```
                    :  ****  ** * *** *.** **   .  ***.  **.       .  . .* .*. *

44                  WYYARYRYMYKYRMAKMGCGWCCKCYCACATACRWWYYKKRTYGYYKMARRAWTCYTSCT
49                  WYYCACGYMYTYRMRKMRYRATTGTCTGTCGGARWWYYKKRKYACCKMGAGGACACCCSG
                    ***.   ***.*** **         . . ..********.*.  **.  .  .  ..

44                  ARRRRCGMCGGKYSYAKKWYYYRKYGAAKYMWRSYWCTRATTKYCGYRRKWYYRRMKGKM
49                  CRRRRTACAACTTCTTKKACYYRKYARMKYMWRSYWMAATAWKCMKYRRKWYYRRMKKKM
                    .**** . .. . . :**  *****.  ******** : :: *   *********** **

44                  YYRMATMRYYYWRTRATTTATRYCYKWGYTMAACKRATYRYYRKAKMYYKCRRSMCKSRA
49                  YYRMCCMRYYCWRYGWYYCCCAYTYKASYCATCAKRCAYRYYRKMKCYYKTRRGAATSRM
                    ****. **** **      .  * ** .*  :..**.:****** * *** **. ..**

44                  WKYGRTCYCGGTKKTKCAYTTYKTTCGRWRWGGAGKKRRRMKMCRKYRKGRYKKYRKYMG
49                  TGCKRCYTAKKYKKYKATCAYYGYYMKRWRWCRWRKKRRRMKMYRKYRKRRYGKYRKYMR
                        *   .   ** *.: : *     ****    ******** ***** ** ******

44                  AGRKKKRKKKRYMKWYRRKWRCGYGMMWYRYKYMMTAGTGTGKWSYRRYRATAKKAMGKK
49                  MKRKGKGKGGRYMKTYRRKWAYRTAAMWYRYKYMMKTACKKRKWSCRRYRGGTKGTAAGK
                      ** * *  **** *****    . *********.:.  . *** ****. :* : . *

44                  KKWCKRMRTMTAYKRYKKACWRRYRYYKKYRMYSRRRYRWYRYKKKRRKYKTRAAAWKKK
49                  KKWYKRMRWAAGYKRYKKWYWGATRYYKKYRMYSRAATGWYRYKKKRRKYKYRMMWAKKK
                    *** ****  :.******  *   ***********    ************ *    ***

44                  GKTYKYTTKKGKKRRMAARWRYRTATYYRAMRWKCKRYMGCRAYWYRRYKAKKKRKYMMW
49                  RKYYGYAGKKKKGRRMMRRWRYRKGATTAGCGTKYGGCAKYRMYWYRRYKMKKKRTYMMW
                     * * *: ** * ***  *****..:   .   *       * ******* ****.****

44                  TCCGKTKKYKKAKGKYCRCYKYTAAMWAGRRWCCKMRRRKYRGCCYAYCAGYRRKYTCCC
49                  KMMKKCKKYKKRTKKYAASYTYKGRCARKRRATAKCAGATCRATTYRCTRATGGGYYTTA
                    .   * ***** . **. .*.*..     **  .*    . *.  *    .    *   .

44                  CRWYYKGKTYMRKSKYGMWKKCGAYMWMKYMKCGGCGYYTKGTTYWRRAGTYRRRRKTRK
49                  TRWYYGRKKTMRKGTYRCWKKYTGYMWMTCCGGAATCYYKKSKWTWRRGSAYRRRGKKRK
                     ****  *. ***..*  ***  .****.     ..  **.*..  ***..:**** *.**

44                  RYKKKMKKAAGYWAKMYRKWRYKSRYMRRKWWYGWYYKRKKWYRRTYAMWSKYRRKKWKY
49                  RYKKKMKKTRTYWRKMYRKWRYKSRYMRRKWWYRWYYKGKKWYRRAYGMWSKYRRKKWKC
                    ********:  ** ****************** **** ******:*.***********

44                  YRKRCYKKWRRWSWRYYKRWWYKRKYKYWYCTRMRRYYKTAYGCKYRKRYKKYYKGCTRC
49                  YAGGYYGKWRRWSWRYYKRWWYKRKYKTWYYYRMRRYYTARCTTGYRTRYKKYTKRTCAT
                    *    * ******************** **  ******.:      **.***** *

44                  RMWWMYYKRKMRKMTWTKCKTRTAYRYYAATCYMKMWKACKAGYYYRSSKMWYAAKARAC
49                  AMATCCYKRKMRKMCWATYKKRYMYRYCWCYYYCKMWKRYKWRYYYRSSKMWTGTKTRTT
                     *    ******** *:. *.*  ***   .  * ****  *  *********  .:*:*:

44                  RWMKRYYYKKRSYYRYWRMARCCMYMRYKRYKGRTTGYRKCGATTRTCTGRWTMMTCYMC
49                  RWMKRYYYKKRSYYRYWRMMRYYMCMGTKRCKRRCCTYRKYRWYCGWYWTRWWMMWYYMA
                    ****************** *  * *  ** * *   ***          ** **  **.
```

```
44      GGCWTTWKKKRAKYYYKYMKKKKKKYKYWKCMRMGACGCCCKKWYYWGCGTGYMGWATAA
49      RCMAGYWKKKRMKYYYKYMKKKKKTYKTWKYMRMRWMKMTAKKWTTWATACAYMSAGCGW
              ***** ************.** ** ***      .***  *. . .**. . .

44      GGMCKKYYRKKKYYYMKYMYKCGRRYKRRRYWAWTMSGMSGMMKKGKRRGRKRKYMRTTG
49      KKMYTKYYRKKKYYTCKYMYKAARGCKRRRYWWWWMSKMSRAMKTKKRRKRKRKYCAWWR
          * .*********  *****..*  ****** * ** **  **. *** *****

44      KSTKWRWWAGAYRYKRSMKKGAGRRMKYGTCCKKKWYKTRYWMRRYRTATCWRTYGTTAT
49      TSYGWRWWCAGCACKRSMKKKWCARCGYKAGTKKKWYKWRYTCGGYRCTCGWRYCAWGTA
        .*  ****...   ******    *  * :  ****** **    ** :  **  .  ::

44      AKRCYRKYYMAYKSYMYAATKKKWKGCTGKRYGGCKRMTCWTTCTTAWYYKKAYRYYKRW
49      MKRYYRTYCCMYKSYMYWWYKKKWKKTATKRYAATKRMKYTGGTYCRTYYKKRCATTKRT
         ** **.*   ******   *****  : ***.. ***.         ****     **

44      ARKMRTKKWRYRTWTAGTRKKWRMMKGATCYCMSCGCYCYWKRTACYWRWWYKKYCRGKK
49      CGTCRCKKWGYRYWATRYRKKWRCCKRTWTYSASTTACYYWKGCTMCWGWWYGKYMRAKK
        . . * *** ** *::  *****  * :  *. *  .  ***  :  * *** ** *.**

44      SRWKWWYCYRKRTYYKAKKKWAAYRKKTKKYKRCCMYRRRRKMMWKRTYYSKAKKKRCRM
49      SRWKWWYYYRKRCYYKMKKKWCWTRKKKKKYKRATMYRRRRKMMWKRKYYSKMGKKRTAM
        ******* **** *** ****.  ***.*****. ***********.****  ***  *

44      YKSSMTYYAYRRRRYYKYKYRTWKWWWMYYRCCTYCMRYWYCKSKRKYYWYMMCAMTTWK
49      YGSSMACCTYRRRRYYKTKYRKWKWTTCCTRYSWYYMRYTCMKSKRKYYWYMMYMMAGWK
        * ***:  :******** ***.***     * . * ***   ***********  *: **

44      TYKGGTRWKYKKRWKMRRYTAGYGGKRWKWMKCGWKKTMRYYYYKKGGTAMYWTKACTRW
49      AYKAACAWKYKGRWKMRRCGTKTRTKRWKWAKMKAKKWMRYYYYKKRKYCAYWKKRSYRT
        :**..  **** ******  :    ***** *   ** ********   . **.* . *

44      GCSMWKWRKMRRAKYWRMYYRYRYATCRWGGTKGCWGYTCTCRYRGSRCGRGGYWSCCYG
49      RYGMWKARKMRRGTCTRMYYRYRYGCARWTAYKKATTYATCYRTRKSRMSAAAYWSYAYA
          .*** *****..  ********. .** . * .  *:   * * ** . ..*** .*.

44      CWYYKAGSTTKYYCATMTCRWYKYYYMYTMKYGYYYSYAYWATTTTTKWYWRKCTAGGGC
49      TATTGRSGGYKYYYWKMYYRWYKCYYMYGMKCAYYYSYRYWRWYYKYKWYWRKYYWAAAT
              ..  *** .* **** **** ** .***** **    . ******   ...

44      TKKRKTWTGTWCCRMRATRYATCYYYYYRKKKWCAAARGACAAYKWWKRRWTGWARKMKR
49      GKKRKYAYSYAYSGCGRCRCMCMYYTTYRKKKTYCGGARGTCTYKWWKRRTWKTWRKMKR
         ****   .   .     *    **  *****  ...  . .:*******     *****

44      WKWRYMKCRKTTYSYRKGMKCRGRMTRSTTTGTSYKRKSAWYRRRKWYSKRMWKKKYMKK
49      WKWRYAKYRKCCYSYRKAMKMGRRMKRSGGCAKSYKRKSWWYRRRKTYSKRMWKKKYMKG
        ***** * **  *****.**   **.**   ..****** ****** ************

44      KMARSRKYMKKGRYYMKSYRCMRKYYRWSKACCGKGYYRRYYWKMCGGCTWGTKKYCKWS
49      KMGRSRKYMKKRRYYMKSYRMMRKCYRTSKMMARKAYYRRYYWKMYRKMWAAKKKYTGAS
        **.******** ******** *** ** **  . *.*********      ..***   *
```

```
44          GRWMYKYKKYYYWTGYRSKRMYWKKRRKRKCKKMMMYRYYYMSWCKCWATCGKAGYACYA
49          KRWMYKYKKTYYWCTYRSKRMYWKKRRKRKYKKMMMYRYYYMSAYGMWWWTTKMKYMYYM
             ******** ***  *************** ************    *    *  *  *

44          KYRKYWGRRKMKYWRRRYYKWMMYKKWWMKYKWYYRTWGKCYYRMGYWYRAWRRKRRGTA
49          KCGGYWKRRTMKYWRRRYCKWMMYKKWWMKYKWYYGYWRKTYYRCRCWYRMWRRKRRCYW
            *   ** **.******** ****************  * * ***   *** ******

44          AYRMKYMGWRRMAKRCAKRYYWRRRKRTYRKKRTYKRYGKMYKRKYSTTCYYYRKKSYMK
49          CYRMKCAAWRRARKRYRGRYYWRRRKGYCGKKRWTKRYRKCYKRKYSAYYYYYRGKSYMK
            .****  .***  **   ********    ***  *** * ******:  **** *****

44          YMKGYCCARYKCCKKCCATYKGACRYYMYRKYYKYYGAAWMRRYKMWSKRCCARAGRKYT
49          YMKRTTAGRYKMMKKAATAYKRRMRYYMYRKYYKYCAGRWMRRYKMWSKRYACRCARGTC
            ***   ..***  **..::**   *********** .. *********** ..*..*

44          CGGCCGWRGTYAGRKMKRRTRRWKSRKMKKRRTSKRKKKKKYKTKKKWYKCCRAGKMYYY
49          YTATYAWRACYGARKMKRACRRWKSRKMGKRRCSKRKKKKKYKCKKKWYKYTRWAKMYYY
              .  .**. *..*****  ******** *** ********** ******  * .*****

44          WRRTGTRYRSKYRRYWKYKKKSRMRGYYRGRSKCKKWGKKWYRTWWYYMYKRMAARYYAC
49          WRRGAWRYRSKYRRYWKYKKKSRMRCYCRSRSTMKGWKKKWYRYWWYYMYKGMWGGYYGT
            *** . ******************* * *.**. * * ***** ******* * . **.

44          KATYARYKKKYWAGWCWGRTTAKWWYYYYWYSYAWRMMWYYWGCCRTMMKYKKMGGACCG
49          KMWCWRYKKKYWMAWYWTRWACTAAYYYTAYSTMWGMMWCYWKSYRYMMGCGKMRRGYTA
            *    ******* .* * * :..  ***  **  * *** ** . * **   **  .  .

44          WAYYGAYKTTMRKCKYYWKYKMYRMYKRTYYTYRRKRMTRYMYTKATRRWKRGRMRKKKK
49          WRYYSWYKWKAGKYGYYWKYKMYRMYKGCYCWYRGKRMCGTMCCKRYRRWKRKRMRGKKK
            * **. **  .  *  ************  *  ** ***   *  *  ***** *** ***

44          KCYRCWMRYRYWRCWAKCYYWRKKYYMKKYYKKRMCCGGCKAWAKCWMYAARTATRKKKY
49          GYCRYACRYRYWRMTMGMYCWRKKYYMKKYCKKRMMTATTKGWTGATMYRGGCRKRKKKC
               *   ******     * ********** ****  .  *.*: . ** .   .****

44          RKKRWAAMKRMYRRYYKGYWATTYWYRRAAARRWMMYACTKCTCSYWSRKGYKMSWKWYY
49          GKKRWGWMKRMYRRYYKRCWMKKYWYRRRWRRRTMCYRMYKMKYGTASRKRYKMSWKWYY
             ****. **********  * ..*****   ** * *   * . .  *** *********

44          WAKKRTGTWRYKYRWKKYTKYYRRRTKMMMYMGMMKKKMYMWCCKCYKKSGRRRKKKWKG
49          TMKKRGRCWRYKYRWKTCKKYYRRRWKMMMYMKMMKKKMYMTTAKYYKKSKRGGKKKWKK
              ***   ********. .****** ****** ********  .* **** *  *****

44          KYKYRKKCAAGYKTGAGGTYCTCTKCAAKWMYRYCKWYKRSRYYRWRRRKGYKKKCKYWW
49          KYKYRKTYRMRYKYKRRTAYAGYKKSWRKWMYRYMKWYKAGGYCATRRRKACKKKATYWW
            ******.    **     :*.  .*.  ****** **** . *   ****. ***..***

44          KYYGKKKARKGGSKRKYYKYYRKMKKKRRRRYMYMKYWWSAKYTAGGRYKYKYKRRRKKC
49          KYYKKKKMRKRRSGRKYYKYYRKMKKKGGRRYMYMKYWWSTKYYRKRRYKYKYKGRRKKA
            *** *** **  * *************  ***********:**    ******* ****.

44          GTKAKCRYKRACAAMYGRMKGTYRRRCYKGYKKKYGGGGRYKKYRCKKKKAAYGRCWRYK
```

```
49          RWKMTTGTTARYGCMYKRMKAACRRRACKRCGKKYKRRRRTGTYRSKKKKTWCRGYWRYK
              * .   .   ..** ***.: ***. *   ***    *  .**.****:     ****

44          YSYWYTYKARCKRMKTCYRYYYYKTGTTAKYMGWKMRWRKACGWKKKKYYMKYRGKGKYR
49          CSCAYCYKCAMKRMKCGYRYYYYKWRWYWKYMRAKMRWRKGTRWKKKKCCCKYGRTKKYR
             *  * **.  ****  *******     ***  ******.  *****   **  . ***

44          RKYRKKTGTTAATRWRCCKCKAKKKKYYSKARYAKAATGKCACGGTGYKKYMKTAKKMYY
49          RKYRKKACCCWMYRWGTTKYKMKKTKYYSKRRYRTRTCAKYRYRRWRYKKYCKGCKKMYY
            ******:      **   * * **.***** ** . : .*       **** * .*****

44          WMYRSRKMKKRRYYYRYKYYSWACYRTCAGATRYRGTRRYCSTAYMWTKTARGMTWKMGWK
49          WMYRSRKMKKRRYYYRYKYYSWTAYRGYGKGYRYRKWGGCYSKRYATYGGMAKCYWKMKWK
            *******************:.**  . . ***      *. *          *** **

44          YRAYYRYGKKKCCTYCYRKYYGATAYCRSMKTYARCGTGCSRMKYTRRYYMYYRKRKRWA
49          YRTYCGYRKKKYYCYTYRKYYAGGRCTGSMKATRAASARTSGMGTCRRYYAYYRKRKRAG
            **:*  * ***   * *****..     ***:   ..:  * *   **** ******* .

44          RTTTCACGAYGCATTTMAMRRWCKWGKRMAYAKKKSKYKKRYKMYCYKYYKCRRMYRCCG
49          AYWKSRYRRTRYGGCCAWMRRWTKWKKRATCTKKKSKYKKRYKMYMYKYYKYGGMYRAYA
               ..       .     **** ** ** : :************* *****   ***. .

44          KYKKKKKYRYKYRRYYKYRRRRMRRAKAWKKSYWYMKYKYYYRYMSKKCCTCWMKCRGYK
49          KYKKKKKYAYKYRRYYKYRRRRMRAGKMWKGSYWYMKYKYYYRYMSKKTACYWMKYGRYK
            ******** ************** .* ** **************** .  ***   **

44          YRRKRKKGYYWYAAGCKRYYRWKAGCAMMKKKSKRASRWRYYARRMRTACGRKKSRYCAW
49          YRRKRKKRCCWYTCATKRYYAWKGAACMCKKKSKRWGRWRYYMAGMGYWTARKGSRYMMW
            *******   **:.. **** **....* ****** .*****   *    .** ***  *

44          AGYKRYRYTKKYKKYCYRRKKKKYYYWKMYMYKRRMKKAGAKKTYTTTKGRRWYYWGKCR
49          GACGRYRTYKGTKKYATAGKKKTYYYTKMYMYKRGMKKTTGKKGYYYYKKGATTTTATAA
            ..  ***  *  ***.   ***.*** ******* ***: .** *   *       ...

44          ARCAAAAKKYYAKMYRKWTRTMYRKWCGCSWAKKKAKRKAKCKRRRKCGCARYKTYACMM
49          CRYGGCGTGCTRGMCRKWCRCMYRKTYKMSWWKKKRKRKWGTKRGRKAAAGRYKYYCAMM
            .* .....      * *** * ****    ** *** ***   ** **....*** *..**

44          RRKKRKRYRMYAAGGWKWRYKRYYTGRYKKYMYMRYRKYTTT
49          RRKKRKRTGMYTTAAWKWATKRYYCKRYKGYMYMRYRKYGCC
            *******  **::..***  ****  *** *********
```

The pairwise alignments of two accessions 44 and 50

```
44                  KYMRYYAMYMGKYRRWKYMKGRKYKMMKYGKCCWMMGRRKGTYGRMWRCCCAYRRTKGGM
50                  KYMRYYTMYMRKYGRWKCCTARKYTMMKYRKSTWMMRRRKRWYTRMWRTAGWYRRYKKRM
                    ******:*** ** ***  ..***.**** *. *** ***  * **** .  *** *  *

44                  RWMKTATMMMYWRRRRWTTTMMSGKRYYYYYYRRRRWWKYYYYKYRKGMMTWRYWKRKYR
50                  RWMKYWYMMMYWRRRRWYYWMACTKRYYYYYYRGARWWKYYYYKYRGSAAYTRYWKRKYR
                    ****   **********   * . *********  *********** .    ********

44                  KYYYYWKRRTRWWMSYMYRRKTSRRYCGKTRMKMMYSRWKKYYGRWMMWYYAGYRSWMKY
50                  KYTTYWKRGKRWWMSYMYRRKWSRRTYRTKRMKMMYSRTKKYCARWMMWYCGAYRSWMKY
                    **  **** .*********** ***   ..******** *** .****** ..*******

44                  YYKRKYRYRSAYRWKWYKYRRGTCWRMWYWMRKKRMTTYYYYRCAKMRRKWYRRCYMASY
50                  YYKRKYACAGWCAWKAYKYRRAAAWRMACWMRKTACWYYYYYGYGGMRRKWYRRTYMRGY
                    ******   .   ** *****.:.***  ****.    ****  . ******** ** .*

44                  WCCTWKRWCYRWYRWMYYRSKRAYWYMYWWRKYYRMYAMYWYWWKRWYYKMGAKSRYYYY
50                  WYYWWKRWMYRWYGTAYYRSTAGYTYMYWWRKYYRMYWMYWYWWKRWTTKMRMKCRCTTC
                    *   **** ****   ****. .* ************ *********  **  *.*

44                  CGTCYYRRYKYYWKAYWKCTWRYYKSCGMYYRRRGGYKWKSGRYRYRYYYRGKWYYRGRK
50                  TRAMTTRRYKYYWKGYWKMYWRYYKSMSMYYRRRKRCKWKSTRYRYRYYYRRTAYYRKAK
                      :   ********.***  ****** .******   **** ********* . ***  *

44                  RYYKYYKRCTTAWYRWYWKYRRRRYWCCRRKRKKMAKYMYRKKYACCTTGCCGKKMRYKY
50                  RYYKYYKRMWKMWYRWYAGYRRRRYWMMRRKRKKMWKYMYRKKTRTAAYRMAKKKAGYKY
                    ********  . *****  *******  ******* *******   .:   . **  ***

44                  KKMWRAGCGCARYKWGSYWRYGMYTRCKAKWSRYKYMRRRTSYCGKRKKKYKMCRTKCKA
50                  KKMWRWKSAMGRYKTKSYWRYRMYGGAGGKWSRCKTMRRRASYTRKRKKKYKAYGATTKG
                    *****  .. .***  ***** **  . .**** * ****:**  *******   :. *.

44                  GCKKCKYKKTKWYKRKTSYKMYYYWRRAWWYRKGSKYKKKSRRAAYSYKKMYYYKMGCAW
50                  CYKKYKTKKWKWYKRKYSYTMYYYWGGWTTYRKASKYKKKSGRCRYSYKKMYCCKMSAGW
                      ** * ** ****** **.*****     ***.******* *. *******  **...*

44                  YYMTGGMMYCGCTCCKYKTACTYACRKRKKMKRKRRGCKKYYATCTTGCGYWCTGGMRKA
50                  YYCKRKMCYMSYAAMKYKWRYACGARKRKKMKRKRRTYGKYYTAAYCKAAYWMAAKCRKR
                    ** .  * * . :. ***   : ..***********   ***::.   ..** :.  **

44                  TTYTYTKGTTCCAAKCTKTKRMARGGAMMRCWKYWKTAACMRWMSYKYGCAMGKGTTRYW
50                  AWCACYKRCWYYGRGMGKGGACWRRRRMMRSWKYAKYTRYCAAACYKYRMGAAKKCYAYW
                    :  :  *     .    *     *   ***.*** * :      .***  . .*     **

44                  TSYCGRTGYKKRGTATKWKCTGYATYMRRATYRYGATCWYWWYYMRMRKKWAGGMKYKTT
50                  KSCYKGWRTKKRRGRWKWGYKRTTCYMRRRKYRYAGGYAYWWCTARARKKWGRRMKYKGK
                    .*       ***    **  .  : **** .***..   ***   * ****.  **** .

44                  AKRMKKAGRARRYYKGWAKKGKYSYWTRATTATKYRKGYYYTRWWMKWRTRWCYRRWWAM
50                  CKRMKKRRRCGGYYKKTTKKKKYSYWWAWAWMWKYRGKYYYKRWWMKWRCRAMYRGAARA
```

```
            .*****  *.  ***  :** *****   :   ***  ***.******* *  **

44          RKKYKWCARWKWKYKRYTRWYMKKMYRYMRYKKYRWKRRKMMKRYYYSSYKKCKMYMSAA
50          AKKYKWYWAAKWKYGRYCRTYMKKMYRYMRYKKYRWKRRKMMKAYYYSSCKKMGMYMSRW
             *****    **** ** * *********************** ***** **  ****

44          KSSMKTTYYAKRRYYKYKRWRYMRKRGTKYRKYYWMYMMYKKYWKYKRWKKCCWMYMACK
50          GSSMKGACCTKRRYYKYKRWRYARKRTCKYRKYYWMYMMYKKCWKYKAAKKTTWMYMRYK
             **** :  :************ ***  ************** ****  **  ****  *

44          CGKKWMRYYYYKKKYAAAKWCAATYRRAWKKAWKRRKMKYTACCYWMTKKKKSRYYGKKY
50          YKKKWMRYYYYKKKYWCTTWSMMWYRRGAKGRWKRRKMKYAGAATWMAKKGKSRYYSGKY
              ************* .:.*.   ***. *  ********:... **:** *****. **

44          CCKRMCACTKKRYTRYSWYRKYYCARYKKYKKMGWKRKTWKKCSCGKKWGYYKRYWWKKK
50          YGKRMYMYKTKRYWGYSAYRKYYMWRYKKYKKMKWKRKGWGKMCMKKKWKTTKRYTWKGG
              ***   ..***  ** *****  ******** **** * *.   ***   *** **

44          YYKYRRTKKKRMRWRWKRKAYTKTYYSRYTCYYRGAKTCYRYSWWKGYKMYYMWYKYKYR
50          TTKYRRKKKKAMRWRWKRKWYGKGYYSRYYYTTGARKYAYRYSWWKRYKMYYMWYKYKYR
              ****.*** ******** * * *****     . * .******* *************

44          YKKYTAWCCGYWKRYMMKKYRMRYMWWYKCWYRYARGYYKYRKRRWSRYRRKRGCRAMWG
50          YKKYGMWYTRYWKRYMMKKYRMRYMWWYTYWYRYRARYYKYRKRRWSRYRGKRRYARCTA
            ****  *   ******************. ****   ************* **      .

44          GGCGGYRWTYRYKWRYYYRRKKWYYRKCACTYARRWYKYRYRWYMRRCGAYKYKRTRYMC
50          ARTARTRWAYGYKWRYYYRRKKWYYRKYGYYCCRRWCGYGYRTTMRRYSCYTYKGYAYMY
            .  .  **:* **************** .   .***  * **  *** ..*.**   **

44          MRKKYKCYMRSKCCGTTCKRYYGTYRTRYRYRRRWSACCKKTTTWRARMKKYCYRKMMYR
50          ARKKYKAYMGSTATAAYYKRYYKKYRKRYRYRRRWSTTTKKGAYWACAMKGTYYRKAMYR
             *****.** *.. .:  **** .**.*********:  ** : * . **   *** ***

44          KWWGCTTTKAKAWAATCMCRGRKRYRYMMYKAGMKMYYCCTTRYKAARRCTCGKMRKKCM
50          KWWSYCYWTTTGAWMKMMYRRRKATRYMMTKRAMKMCCYTCCGYKTGRRMYTKKMRKKMC
            ***.    .:..   . * * **  **** * .***       **:.**    *****

44          GGKYRYWRCKYYMRKKMYMKKKYRKAKAMKRMYRKRKKRMYRMYRYYRRYMYMRKGKKSW
50          ARKYRYWRMTTYMRKGCYMKKKYRKRKGAKGMYRKRKKGACACYRYYRRYMYMRKRKKSW
            . ****** . ****  ******** *. * *******     ************ ****

44          GKWMYYMWWRYKRRWTGWGRKKWYKMKWKKKTYMARRRWRRKGMMYKKMYMKRKYYTYRR
50          KKWMYYMWWRYKRGTYKWKRKKWYKMKWKKKATATRRRWARKKMMYTKMYMKRKYYGYRG
             ************    * ************:  :**** ** ***.********* **

44          RKCSGRTCWKKKYKKCAAGRKRKYGKTTKRYTKWYKTACACYRYYRWATRYKYYKKKRKT
50          GKYCRRWYWKKKYKTYRMRRKRKYKTYKKATYKWYKYRMRGYGTCATCGRYKYYKKKRKW
             * . *  ******.    ***** . .*   ****     *      . **********

44          TYYKYYRKGMKKATAKRRRRRYRRYMMCTKYWWSYKMCAGAKYKYYMKSYYYGKRCTGKT
50          KYYKYYRKTMKKRKRKRGGRRYRRYMATCKYWWSYKMMWATKYKYYMKSYYYTTAGGAKA
            .******* *** . **  *******   ********  .:*********** .   .*:
```

```
44       CRMSYRRRCYAWWYSRRWRRKRAKWGRSKWKRWYRKKRYGKCGYAAARRKMKRMMWSYKR
50       TGMSYRRRYCMWWYSRRWRRKRWKARRSKWKRWYRKKRYCKSTYRRWAAKMKRMMWSYKR
           ******   *********** *  ************* *. *     ***********

44       YKATTGRYRRRWAWYAYKKKCTYYAYAMRAARRKYMGWKYYMKSRYYGMRYYRYKKYRKK
50       YKCWYSGYRRRWGWYRYKKTACYYRTGCRRGRRKYMKTGYYMKSRYYTMRCYRYKKCRKK
         **.   .*****.** ***.. **  . * .*****   ******** ** ***** ***

44       KRYKKSYKCCTTMYKKKYYWYRRMYKCKRKKWKKARYGGYRYYYTKRMRRWWMWMKRKKW
50       KRYKKSYKYAWCMYKKKYYWYRRMYKGTRKKWTKWRYAAYRYYYAKRMRAATAAAKRKKW
         ******** .  ************** .****.* **..*****:****      *****

44       RKKGYYMMMYGRRMMMYWRKYKRMWKRWTRTTYSYKYCKSRWKYCTARAAATGGYYYKKW
50       RKKRYCMMMYCARMMMYWRKYKRAWKGACRYACSYKYMKSRWKYTCTRWWRYKKYYYKKW
         *** * ****  *********** **   * : **** ******  :*      ******

44       KRGRTYKRKCTAGKMKWKCMWWKKARRYKTYTKWYMTTKWRYWRRKGRKTKYYTGTKAKY
50       KRRAYYKRKYCCRKMKWTMCATGKCRRYKGCYGWYMWYKWRYWRRGARKYKYYCACGGKY
         **   ****  . ****.     *.****    ***  ******* .** *** .  .**

44       YYMAATRRYMKRKKKMGRKMWYYWRKKKRRWYMYKKMMCKYACKWRYRKKRKRYYYKYKK
50       YYMTTARRYMKRKGGMRRKMAYTWRKGKRRWYMYKKMMYKYRTKWRYRKKGKRTYCKYKT
         ***:::*******  * *** * *** *********** **  ******* ** * ***.

44       MKYWMKKKKRWRYRAYMRWRKCYKKKYYRKKGYMMKKGGTCYGATKYYGRGYMAMRCKMT
50       MKYWMKKKKRWRYRRYCGTATMYKKKYYRTKKYMMKGAACYYRRKKYYTGRYMGCRAGCC
         ************** *    . *******.* **** ..  *  .***   **. *.

44       CCCCCWWCYYYTYMRKRKYKKYKKMWMKCGGCYKWRGARTCYMRKRCKKSKYMYWKRKMK
50       TTTATWAYYYYAYMRKRKYKKYKKMWMKGAATYKWRAGGYTYMGGGYKKSKYMYWKRKMK
            . *  ***:**************** .. ****..   **    *************

44       KRRRKWKYMMYRYYYMSRWRYWRRMGRWRYTKKKYKWMMKWMKMYRGKYMMYKWYYYRGG
50       KRRRKWKYMMYRYYTMSRWRYWRRMTAAGTWKKKYKWMMKWMKMYRRKTCMYKWYYYRRR
         ************** **********      *************** *  ********

44       WYRRARKRRRATRMRRRWRMRKKTRYYYGARKRYRRRWKKWYKRCCACTKCTRRYKKRGR
50       WYRRMRKRRRGAAMRRRWAARKGCATYYAMRKRYRRRWKKWYKRATGTCKAARRYKTGRR
         **** *****.: *****  **    **. **************. .  *.:****.  *

44       KKSTYTYYYRKSKYCKMWMTRYKTATTCTTMKRTAAACCRRYYWRGCCRSGRKWRYAAGG
50       TKSACGYYYRKSKYMKAWMCRYKYRGCAYYATGACGTYTRRYYWRAATAGAGTTGCRMRS
         .**:  ******** * ** ***    .   . :..:  ******..  .. .      .

44       ATTAGWRGYGAYRYKCYTCCRKRRYWYTCYRTYYYTTTYKYRKMYTACYYRGRRCGYKYA
50       GWWMATARTRGTRYGYYATYRKRRYWCYTCRGYYYCCKYGYRGCCGGTYYRKGRTTYKYR
         .   .     . **  *:  ******    * ***  .* **    . ***  *  ***

44       RKCTAGWRMSMYYGAMATGKATGKKWYYYTYMYKWKYRRKKMRRYRGMWWMYKRYYCTGG
50       AGACGCWRMSMYYATMCAKKRWKGKAYYCCCACGWKYRRKKMRRYGRAATMYKRCYMYRR
           . . *******.:*.: *    * **      ***********     **** *
```

```
44          RKTYRYTKRSMKGRRCWKKWGKKRKWMRRYYYRRRRYYGTATSTAAMRGCTTYYCSKTGC
50          RKGCACCKRSMKTARYTKKWRKKRKWMRRYYYRRRRYYAGTACGTGMRRMYWCYMCTAKG
            **     *****  *  *** *****************. ::. :.**     * ..:

44          CAYMYKKKKSAAATARYYKRWYKYYYRMRGYKKYYKKGRCMKCCGYKWKKKKYRYKYWKY
50          GGYMYKKKKSTGTGMRYYKRWYKYYTAMRRYKKYYKKKGYMKTTAYKWGKKKYRYKYWKY
             .********:.:  **********  ** *******   **  .*** ***********

44          YRRRRMAAGYYMRKYYKRSYWYKMMYYYRKYWTWCKRYRWTAGWKTCCKKWRTGKRKKKK
50          CAGAAAWWRTCARKTYKRSYWYKMMYYYRKYTYTYKRYRWCCRAKGMYKKWGWAKRKGKK
                        ** ****************    *****  . *   ***  .*** **

44          KKRGYRKMRRYWKRKKKYYRMKKKRRSRRYYWRATSRYCWYYCGRKWMRRRRMKAGWRRA
50          GKRKYRKMGAYWKRKKKYYRMKKKRRSRRYYTRMYSRYAWTYYKRKWMARRRMKMKAGRR
             ** ****  ********************* *  ***.* *  **** *****    *

44          MYRRMGTWCRYRCYTYKGARYMAKTGWYKCMKWYRRKKMYYWMYRKYRMGCGSWYYWSAC
50          MYRAMRYWYRYRMCWYKKGRYMGGCKACKAMKWYRRKKCTYWMYRKYRMKACCWCYWSMT
            *** *  * ***   ** .***.     *.********  ********* . .* ***

44          RKKSYAMTWGRTYKRYKKTSKRKMYKRRYRWYRYCKTCKCSKYMRKRKKKKKKWRRAGRT
50          RKKGYMMWWRRKYGRYKKACGRKMYKRRYRWYRYTTCYKTSKTMRGRKKKTGKWRRRRGY
            ***.* * * *.* ****:. ************* .  * ** ** ****. ****

44          YCYYACYCGGYGKKYAYGGRYKKCYWKKKGARKYKYYYMWYWMWTACYWRYKACAWKWRG
50          CYYYRYYTRRCRTKYMCKTRYKKYYWKKGRTGGCTYYYMTYWMWYWYYWRCKRYGTKWAA
              **  *     .**    **** ****  :   .**** ****   *** *  . ** .

44          RYTGTMYAGTAWYYCKYKSYRGRKRSRKYAKYARKYWKMMTYYTGSRAGGRYWSSYKTYK
50          RYYRKMYCRATWYYYKYKSYRKRKRSRGYTKYCGKYWTCCYYYASSRWRRRTWSCYKKCG
            **  .**. ::*** ****** ***** *:**. ***.   **:.**   * **.**.

44          AYYMYKMRWGKWYKWMYMWMRTYWSRKGAWAAMYRCKKYKKRRKKKGCYGARRCMMYKGY
50          GYYMYKMRACTWYKWMYMACGCYASRKTGWMMMYRYKKYKGAGKKKKYYSRRRYMMYTTC
            .*******  .*******    * *** .*  *** ****   ***  *. ** ***.

44          AMKYKKKRKKKKRMAGCKAMYKRWYRWWCWKRKKKKKKGKTCCTRRRYRCATKYRRYKRY
50          CCKYKKKRKKKKACTATKWMYTRWYRWWAWKRKKKKKKRGYYYWRRRYRTCCKYRAYKRY
            . **********  :. * **.******.*********      ***** . *** ****

44          MMRKRYRKYTWWWKYKMKKMSMYKK
50          MMRKRYRKYGAWWKYKMKKMSMYKK
            *********  **************
```

The pairwise alignments of two accessions 44 and 51

```
44          KMKMKWWRYRRGGWRCTKWKRCKRGCYWRRWSRYMWSAGAAAMAGCYCRTKRWWYCYWKC
51          KMKCKWWRTRGKKWRYWKWKRYTGKTYWRRAGRYMWGGACGGMGAGCTRYKRWWYYYWKT
            *** **** *   **  **** .   **** .****......*..   * ***** ***

44          AAKCMRRKMTYYYRYYYKRCAMWCYYKSKYRYMMWMMWWKYRRMRCYMRRWWMKYSRRKR
51          WCKAMRRKMWCCTRTCYGGMTMTYTYKSTYRYMCWMMWWGCRRCRYYMRGTTAKYSRRKR
             .*.*****    *  *   :*   ***.**** *****  ** * ***    *******

44          MRKKYRWYAYWCWTYACARYSYARYYYWRKYRYRYWCGYRYRYMRRYWYWWRRRYYKYYR
51          ARKKYRWYGCWYWWCMYTGCCTTRTYYWRKYAYRYATRYRYRYMRRYWYTWRRRYYGYYG
             *******. * *    :  . :* ****** ***   *********** ****** **

44          WCYRMCWWYYYRWSYYKSKWKKCRKMGGRRCAKRKYMRYMRWYMRRKRRTCWKYRYYCTY
51          TYCRMYWWYYTGWSYYKSKWKGYGGMRKRAYMKRKYMRYMRWCMGATGRYSWKYRYCAAC
               ** ****  *********    *  *   ********** *  . * .***** .:

44          YTYYRRRYCMRMWSKSYMAAWRYKWSSKWYWRGCSRKGMMTWTRMSAKRWMRMYSRRKWR
51          CKYYRRRYMMRMWSKSYMGGTRYKWSSKWYWRSSGAGSAAYTWRMSCKRWMRMYSRRKWR
             .****** *********.. ***********...  .     ***.*************

44          AYWMACYGGYRKYTRKRYYWWMYTWGKYKKYYRYYYYYYRKKYYWKTRKMYRKCRMKAAR
51          MYWMWTYKKCAKYWRKAYYWWMYATRKYKTYYRCCTTCTRKKYYWKYAKMYRGMRMKRMR
             ***  *    ** ** ******:  ***.***      *******  ****  ***  *

44          RKKYKMMKMSMWRRGTYKYRYRWYAYYYMRMYKCRMKGCCGYRYYMKYRWRWRTMRWKGM
51          RKKYKMMKMSMWRGAYYKYRYRWYMCYYMRMYKTRMKATYAYRYYMGYRWRWRCMRWKKM
            ************* . ********  ******* ***.  .***** ****** **** *

44          RRRYGMGGGKWRMYRRYWRYYTWKRKATMRKKYMYWKKKYAGRMYAMKMKMYGKYRKYKK
51          RRRYSCARRKWRMYRRYWRCYCWKAKGAMRTKYMCTGKKYRRGACRMKAKMYKTCRKYKK
            ****. .  ********** * ** *.:**.***   ***      ** *** . *****

44          KWRRMYRRYKWCYGMYKCTCRKYTTTSWKKRYYWYRCCACTMWWRMKYKRYWACWRKCKM
51          KAARMYRRYKTYYKMYKMKYRGYCCCSWKKRYYACGTYRYCMWWRAKYGRYWRMWRKYKM
            *  *******  * *** . * *   *******        **** ** ***  *** **

44          CGCTAKATKGRGAAYYKYCCYYMRKYTGYRMRYKYYMYMATCTACGKYWWKWMRMCGGGK
51          AAGCRKCGKARATGYYKYYMYCCGGCKRCACRYKYYMYMTGYCRARKCWWKWMRCTAACK
            ..   *. *.*.:.****  *     .    ********:    . * ******  .. *

44          YRYMYYYWYGWWYKTCKYAMMTRKMATAYYYYKRCYKYMYKMACKKKKKSRYYTKRWCGA
51          YRYMYYYWYRWTYKWYKYRMCGRKCRKTYYYYKRTYKYMTKMTMKKKGKSRYYYKRWYKW
            ********* * **  ** *  **  .:****** **** **: *** ***** ***

44          YKMMYYRTTAKKGMYKYKKGYYKTGGCKCGCKYKKKYMRCKKYRYKKRKYKYYYKYRRRR
51          YKMMYTAAGCGKKMYKYKKKYYTKRRYTTAAKYKKKYMGYKTCATKKRKYKYYYKYRRRR
            *****  : . * ****** **..   . ..*******  *.   ***************

44          GGMWRYKKTTYSYWYKKYKYRYYYYKKYRWKWKWKMRTKYACCRKKRYYKRKMRRTKKRK
51          RRCTRYKKCGYSYWYKKYTTAYYYYKKYRWKAKWKMRGKCGTTRKKRYYKRKMRRYKKRK
```

```
                        ****  ********.  ********** ***** * .  ************ ****

44      SKYYRMKKRRYYSRMRKWMRYSRYKATAWYYKKKKKGGGGRWYYKTSGAKKMRRMGYYWK
51      SKYYRMKKRRYYGAMGKTMRYSRYKWYMWYYKKKKKKRRRRWYYKGGTMKKCGRMRCCAT
        ************. * * *******   ********    ***** .  **  **    .

44      WRKWATARRYKSYCGAKRMKCGCYYRKACRYKKKKKYMKWKRCGYKWKKTSCCCKCTWRY
51      WRKWRYRRRYKSCTAGKRMKACYYCGGRYRYKKKGKYMKWKRAAYKWKKKCYYMKTATRT
        ****   *****  ..****.  *     ***** *******..*****..    * : *

44      YKKRKYRKKKYYYKYRWYKKRYRTATAATKARWKRYRYKYRYRTAGWGKYMKKKMWSMKY
51      YKKRKYRKKGTTCTYRWYKKRYRCTCGCYKWRWKRYGCKCRYRYRAWAKYAKKKMWSMTY
        *********    .********* : .. * *****  * ***  .*.** *******.*

44      YYYRYKKYCYYRMYKKWYKMKKATGCYRRAKYAYTTKKCCWKRYRRRRGTGMWMKYMRAA
51      CCYRYKKYYYCRMYKKWYKMKKRWKTCGGWKYGCCCTKTTWKRYRRRRKYKATMKYCRCT
          ****** * ***********        **.   .*  ********     *** *.:

44      WKAGRMYWCCKRYRMYRRGARGYCRYRRSMKCCKCWGTKKWKRGCKWRRYYGYMRGRKMK
51      WTMKRCTWTGKRYRAYRRRGRRTGRYRRSMKTAKTACAKKWKRSAGAGACYTYCRTGKMK
        *.  *  *  **** *** .*   ******* .*   :*****..     * * *  ***

44      GTSARMRRGYGGKYYMYMWRGGRWYRTGMATRYWMTRCYYKYRRYKKRYYKKMYRMMKYR
51      AACTGCARRCTTKYYMYCAGKKRWCRYRMWYRYWMKRMCYKCRRYKTACTGGMYRMMKYR
        .:.:   *    *****     ** *  *  ****.*  ** ****.     ********

44      RKMYGMRMKRYRMMKRRRGKCRKKTRKSYCMRKYKKKMSTYRKMYKRRYRWKWMYKKTYK
51      RKCTKMRATRYRMMTRRRAKMRKKWRKGYAMRKYKKKCGWYRKMYKRRYRWKAMYKKYYK
        **   ** .*****.***.* *** **.*.******* . ************ **** **

44      KKKYKRKKKRKYWMMYKYYKKRCCGAYKTCGYKARKYYWGKYRKGGTGKCYYRKWCTACR
51      KKKYKGKKKGGYTMMYKYYKKRMTRGYKYTRTKCGTTTTAKYAKAAARKYCYRKWYACYR
        ***** ***  * *********   .**    *. .   .** *..: *  **** :. *

44      WTRTTRCGTAGCWKYKKKKKRKKCCTKRRTGKKYRCRYKYYTCGYKKYRKMYKYWKMYCY
51      TCGYWAMSYRKMWKYKKKKKKRGKYYWKRRCKKKYRYRYKYYYTRCKKYRKCYKYWKMYMC
               .    ********* *   ***  **** *****    ***** *******

44      YKYCMGCYKMCGCTGCGWYYRKYTTTRYYYYMYYYKRKYKYRYMCYMKRGTAKKATGYWY
51      YKCAACYYKMYATCRMRAYYRKYWYYRYYYYMYYYKRKYKYRYMMYMKAAAGTKRKKYWY
        ** .   *** .      *****   ***************** ***  .:..* . ***

44      SYWMRAAKSKRGAKMKCGTMARRGATGGMKYAYKMMYMRRSYMKRKMKTKKKKKYKKYKK
51      SYWMRGCKSKRAWKMKSACMMRGRTCACMKYRCKCCCMRRSYMKRKMGGKKKKKTTKTGG
        *****..****. ***.. * *  : . ***  *    **********  ***** .*

44      KWYKKMYYYWRGAYRATAKKSKRYWKKKKRRKKTATCAKAYMSWYWYRRYGCMYTYGRYR
51      KWYKKMYYYARACYRGCGKKSGRYWKKKKRRKKYRAMRGGYCGWYTYARYAAACAYRRYR
        ********* *..**. .*** **********   :    .* .** * **..  :* ***

44      SGYAGYYMWACGYRYRARMYRRCCARKMGRWRRRKMYGWCGCCRRKWKYKKCCCYCTYAA
51      SATGKYTCTTTKTGTRWGCCARMYWGTCKGAGRRKMYATYRYYAATAKCKKTTTYMWYGT
        *. . *   :     *     *     .     *****.       . * **   *  *.:
```

```
44              CRKYYYWRKMGYARSTRKYKRCYYKYRWRGWKAKRRMYSKRWRYRYMYWTTCCTCARYKK
51              MGGTCTWRTMAYGRGCRKYKAYCYKYGTRRWKRKRRMYSKRWRYRYMYWAAGAATWRYKK
                      **.*.*.*. ****   ***  * ** ***************:: .:  ****

44              KYTRKGKYTWCRRGKKYKGYWYYAKRRWMRRRKGCKAYWATTWYYYWKRYKKKSYAYMKK
51              KYKRTRTYCTTGATGKYKKYWYYGKRRAMRGAKSMKMYWRKWWYYYWGRYKKKSYGCMKG
                **.*. .*       *** ****.*** **  *. * ** . ***** *******. **

44              YGGTYYYTWCAARRCRCKGYRMRKRAGYTRYTKYKMWYWKKKMWATGTTMCYKRKYTRTSA
51              YKSKYYKWTGGRGYASKKYRMRKRMTCARYWKYKMWYWKKTAWTYRYYMYYKRKYKRCGC
                * ..**.* ..*   .* ******   :** *********. *:    * *****.* ..

44              RMMRKMRKWKTRYTRRCATAYRYKMKMMYYYRSSKMWYAWRRGAKKCYAWMGRYYYRRSY
51              RCCGKMGKWKWRCCAGTGCMYRYKMKMMYTYRSSKMWCGTARAGKKTCGWMTRYYYRRSY
                *   ** *** *      .  ********* ******* .  *..**  .** ********

44              YMKRYMCGTGRYYRYTRMCMYTCKKYRWTCGTTSKGKWWRYRKKKRKYKKKWYKKRYSWK
51              YMKGTMTKCAGCYRYYRMTMYWMKKYRWATTCGSKKKWWRYRKKKRKYKKKAYKKRYSWK
                ***  *    .  *** ** **  *****:    ** ************** ********

44              MKYTMYRYMRYKKTKKKYYYWRRKAARWWWMMKRYAYSSMRTCAGKKRMSTKSMYGTYYY
51              MKTWMCAYMRYKKCGGKYYYWRRKCTGTTWMMKRYRTGSCAYYMRGKRMSCGSMYKACCC
                **  *  ******   ********.:   ******  .*      ****  *** :

44              AKYKGWYKYRTWKWMRRYMRRWYKKKYYWMMMWMYKKWKYKRWMKRKYGKRYWMYCRWKR
51              TKYKRTYKYRWWKWMRRYMRRWYKKKYYWMMMWCYTGWKYKAAMGGKYRKRYWMYTAWKG
                :***  **** ******************** *. ****  *  ** ******  **

44              MTMCRACGGKYKMRYYKYYKAKKYSYAKAAYMRYWKAAWTCTRMYAWKGYMYRYWCRCKT
51              CWMMGGYRRKYGMRYYKYYKTTKYSYRKMRYMRYWKRMAYYCRMYRWKKYMYRYWYGYTA
                  *  .   ** ********:.**** *  ******      *** ** ******   .:

44              ARSRWSMMKKWKYCGKCGTYKRYMMYSRYKYYWKYYTGGYKGMGSTGRSWWYGGKRYWRM
51              WGSRWSMMKKWKCAAKYRAYKRYMMYSRYKYYWKTYCARYGRMRSYKRGAWYKRKACAGC
                  ********** ..*  :************** * . *  * *  *. **  *

44              RTRACYTKMRACAYMKAGAYWWCAKRMRKYYMKYRMWSRYMKRKKCWYYKRRKKRRMRMW
51              GCRGYCCKMRRYRYMTMCGTTWGGGRMRKYTMKYRMWSRYMKRKGGWYYKRGKKRRMRMW
                  *.   ***   **.  .  * . ***** *************  ***** ********

44              SYRRKKKRMWYYTCSYMGWYRKMGWSWRKRKWCACYAKRYARWKKKYYWYMRRYRARMYK
51              SYRGKKKRMWYCKYSYAKWYRKMRWSWRKRKWYGTCCKRYCATKKKYYWYMRRCAGRMYK
                *** ******* . **  ***** ********  .  .***.  **********   .****

44              KGRMRYKKYYSKKKTTKRRWAGTWTARRKRYWYRYGYYYARYATYRRYRYAKKKYYARYK
51              KKRMRYKKCCGKKKWCKRRWRRYWWWRRKRYWYRYAYYYRRYWYYRRYRYRKKKYCRRYK
                * ******   .***  ****   *  *********.*** **  ****** ****  ***

44              RWTKWYRWWYYYGCCMCCGWCAACTWMKMWYYYYWSWTRRMWKCRKYYYRGTAKAACGWG
51              RWGGWYRAAYYYATAMYYRWYTTTAWMKMWYYTYWGTGRRCAKARKCCCAAWGKRWTKWR
                **  ***  ***. .*   * :: :******* **.  **  *.**     . .*    *
```

```
44              RWRKTTGTKCARTARYMYAWAYYKGYKMYWWRSRGKRTRRTTYACYSKRYCKKMRRWMKG
51              GAAGWYRKKYRRATGCMYTWMYYKRYKMYWWRSATKRYGRWYYRMYSKRYMTGAGAACGT
                       .*  *::  **:* *** ********  **  *  *  ***** .

44              TKYCKKKRKARWKKWCWMGCKGWMKYKYWKKRRKRYCTGYKRKRAKTRKTAKKCRYWYKR
51              AKYTKKKRKCRWKKWYWMTTGRWMKYKYWTKRAGRYAGKYKGKRMKYRKCWTKYGTWYKR
                :** *****.***** **    *******.**  **.  ** ** * **  .*   ****

44              KTCAGMKMYMYYMYRRRATRMMKCKYRGTYKYKCWYWAAAGRKYKKWKWWWTRYSYRYYG
51              KATGTMKCYMYYMYRGARYRMMKYKYRRCTGYKYTYWMRRRRKTKKWKWWACRYSTRYYT
                *: . ** *******    **** ***    **  **    ** ******  *** ***

44              YTTKKKWGCKAYRMGGTTKYMYKYYAAYSTAGRTYRYMGCCGKSKKWKKWYYKKYTRATY
51              TKYKKKWRMKCTRATTACGYMYKYYGGTCKMKRYYRYMKMYAKSKKWKKATCGKYAATAY
                 . ****  *. *    : ******.. ..  * ****   .*******    **: ::*

44              RRKKYKRMYRYKKRCRKMKWRAAWYGCTYRRYYMRGYMGKKGAAYWCWYGCYMKKKWTYK
51              RRKGYTRMYRYKKGYGKMKWRRWAYKTACGRTYCGAYMAKKSTCYWYTTKAYMKKKWWYK
                *** *.*******   *****   *  :  * *  .**.**.:.**    .****** **

44              WKKYKKYYWRRRKMMYCMKMYKRWKRKYYYRRRKKKRGWGKAARKCAAGYCCGCWMKTTK
51              WKKYKKYYWRRRKMMYYMKMYGGWKRKTYYAGGKTTGRWKKRGRKYRMRYYYAYTMKCGK
                **************** ****  **** **   *..  * * .**    *  .  **  *

44              RYKYKRYWYKSWTAGTRYYRWTRYKKAYYKACCTMYYYWRKRKYYKYYRCAKMKATAKTA
51              ATKYTRCWYKCWKTTGGTCATKRYKKMYYKCTTWATCTWRGRKYYKYYRYRKMKTCGKAG
                  **.* ***.*.:       .**** ***.       ** ********  ***: .*:.

44              TRRWTYYWMGMYWWSRGKAYTRWAGGAGGKKYKYAAKSARRTGYTTAKRTRYCYGTACAC
51              GRRAYTYWMKMYWWSGAKCCCGTGRAGKRKGYKYRGKSWRRYRYCCGKRAGTYYKCTARY
                 **   *** ***** .*.    . ..  * *** .** **  *  .**:   *  :.

44              GGMRKTWYASWGARRMRRKKMWGKKWCKMYYYYGRKTYMRTCMYTGCTKTGYYSRAGAYY
51              KSMRKWWCMSWKTRRARAKKATRGKWAKMYYYYRGKWYMGCYCTYTTAKAAYYSRTTCYY
                 .*** *  ** :** * **    **.******  * **        :*:.****: .**

44              AYRKRAWTTCYYMKRRRRRKYMMMKYRSYRTTTWAGYACGAKKWKWYYWYYKWMMKKWMK
51              GYRKRTWGCYYCAKRRRRRKCCMMGYRSYAWWKWMKYMYRMKTWKWYTACTKWMMKKWMK
                .****:*   *  *******  ** ****   .*  *    *.****    *********

44              KATKWYGAYCMWRKYYYRYRYRGACCAWYWRKKASRRYRKGKRMRRRWRKKRYRYRRRKK
51              KTAKWYRRCMCWAKYYYRYRYRRTTTRTCWRKKCSRRYRKRKRMRRRWRKGRYRYARRKK
                *::***     * ********* :     ****.****** ********* **** ****

44              RCKKCTACKCTRGMYRYSYYYRYYAMARRWRRRKKKGRWKYKKGYYGRRKKMKRMMSYKK
51              RMKKTYRTKAARRCYRYSYYYRYYWMGRRWRRRKKKRRWKYKGCCTKRRKKMKRMMSYKK
                * **    *.:*  ********** *.********* *****     *************

44              RKWYRYSRMWWWKGTRAGACWAGKCTRRRKWKKACGYTRSMYYKRMRYYRWYKYKYYRKR
51              RKWYRYSRMWWAKTCRGRGMWWRKMWAGGKWKKRMKYYRSMYYKRCGCYRTYKYKYYRKR
                *********** *  *. . *  *     ****   * *******   ** *********

44              KKMKYSGMKKRWMYRKCKYKYCYYRYKWKK
```

```
51                KKMKYSRMKKGTMYRKTGCGCYYYRYKWKG
                  ****** ***  ****      *******
```

The pairwise alignments of two accessions 45 and 46

```
45                  WGCWKWYGGTYYRWWAKYYYWRKRRMARMKMWWRTYYRCRRRYGWWYKWYWMRRMYSKRK
46                  TTYAKWCRAAYYRWWRGTCYWRKRRMGRMKMTWRKTYRSRRRYATWYKWYTAARMYSKGK
                        **  .:*****    *******.**** **. **.****. *****   ***** *

45                  RWCWKWCCYRWWATKRSSYWRGYWARMMMRRRCTRRRMRCMMYYRSYRYMRYTYGCYYGT
46                  RWTWKWMMTRAARKKRSSYWGRYWWACMMRRRTWRRRMRMAMYYRSYRCMRCAYSYYYTG
                    ** ***   *   .******  **   *****  *****  ******* ** :*. **

45                  YMYKYWKWRWWKWRYWRYYKRAYYYYKWMMYRTMRMYRKWGGWRRKRGKWYRSGCKRKWW
46                  YMYKYWKWRWWKWRYWRYYKRCYYYYKWMMYRGMRMYRKTTAAGGKRRTWTGCTAGAKWW
                    *********************.********** ******  .   ** .*  . .  ***

45                  WYRMYKRMRWRKYRWMYGRWTGYMRRMYCCWCTTGTAYYRYAYKKMKGGKKGRRRYYGYK
46                  WYRMYKRMRWRKCRWMYRATYRYCAGAYYTTTCCACCYYRYCTKKMKTAGKARRRYYAYK
                    ************ ****     *    *      . .****. **** . *.*****.**

45                  AYWWKYYMAKYMKCYRMRYRRYRKCCATRWGYKKKYSRYRAYWTRTKKRWCTMYYGWMMR
46                  WCWWTYYCCKYMKMTRMRYRRYRKTTRKGTKYKKKYSRYRGYWCRKTKRWTGMYYRTMMR
                      **.** .****  *********   .   *********.** *..***  ***  ***

45                  KKWKKMRRRYWKYMYWYKWTYMRYCYRCRMYKKGTRYRKYYSMKMWWYSYYKGGCKMYWS
46                  KKTGKMGRRYWKYMCWTKTYYMRYTTRARMYKKAARYRKYYSMKMWWYSYTKTRTKMYWS
                    **  ** ******* * *  ****  *.*****.:*************** *   *****

45                  YYYYMKYWKKRMYWCMYRRWWTRRYRRYRYTYMKKKYTYYARWKRRRYMYYKAMKKYYRR
46                  YYYYCKYWKKRMYWMMYRRWWGRAYRRYRYYYMTKKYYYYWRWGRRRYMYTTMMKKYYRR
                    **** ********* ****** * ****** **.*** ** ** ****** . *******

45                  MWYGSYKGYWWRRWYMMKACGRTAWKYRWYCYRKYRYKGRKRSKRYMWGMWKYTYTRYRK
46                  MWCRCTKSYWWARTYMMKRMRRWTWKTRWYMCRKYRCKTRKRSKRYMWRCATCCYYRCRK
                    **  . *.*** * ****   * :** ***  **** * *********   .  * * **

45                  MGGCRWRYRCTMSCRCGKKYYRAAGKRRCYMGTCTYMRAAAGTCKRRRYMKCYRYYGRCM
46                  MRAGGAGTGAKMSMRYKKKYYRTCCKRRYYMKCTGYMRRWRAGTTRRRYMKMYRYYKRYA
                    * .      ..** *  *****:. *** **    ***   .  .****** **** *

45                  RYYRARYRKKKAYKYRTRMKYMYGCYGCRYGKYRRTRYKRGWCMKRMKYRKKSWMYYWCW
46                  RCCRMRYRKKGCYTCGKRMKYMCAMTRTRYRGCRACRYKRATMCKRMKYRKGSWMYYATA
                    *  * ***** .*.  .***** .    **   *  ****.   ******* *****

45                  AKAAYRRSKCCAKCGTYGGYAACWGTGRWYKRMTYATKYCSMMKKYYYYRRKMCCAYKYY
46                  GKWWYGRSKTTGTMTAYRKYWGTTRKKRTYKGCKTGCGCTSMMKKYYYYRRKMSMWYKYY
                    .*  * ***  ..  :*  * .   . * **  . .    *************.  ****

45                  MYTGYWTYYKCARWRGCCCAWRMYSKRTSRRKKKTYKKRWSYRYRKKKRMMKKCWRWMRR
46                  MYWKCWYYYTTGGARAAGAWWRCYGKRKSRRKKKGTKKRWSYRYRKKKRAMKKAWRWMRR
                    **   * **. .  *.. . ** *.**.******  ************* ***.******

45                  KYKRMRRYKKGYKRSYRYMKRKKYRCRRKGKMRYRATAKACKYYYTWYMMYKYKWKYRMK
46                  KYKRMRRYKKRYKRSYRYCGRKGCGMRRKKKMRYRTYWTMYKYCYYWYMMYKYKWKTGMK
```

```
                **********  *******  **    *** *****:  .  ** * **********  **

45              MMWKKYWRSGYTYKYKKRKRMYRYTCTKWYKKMKSYWKRKKTAWKRRYRWRCKKYRYKKR
46              MMWTTCWRSTCCYKYKKRKGMYRYYYWKWYKKMKSYWKRKKYMWKRACATRTKKYRYKKR
                ***.. ***   ******* ****   **************  ***    * ********

45              KKYKKKKRKRYRYRKYMRWKYKKKRKMKYWRRRKYRKYKKGCGAGYRCCMWKRKYCKGAT
46              KTYKKKKRKRYRYRTTCRWKYKKKRGMKYWRRRKYRKYKGTYSGATGMMMWKRKYMKAGA
                *.************.  ******** *************   ...    ****** *..:

45              GGAWRRCTYRWSYRKARRCCARGKCRKAAMMKKRRMYWWKGKKYMMRKAKYYKKTGRGKK
46              CRGARRTGYGAGTGGRRRSMWAKTMRKRRAMKKRRMYWWKKKKYMMAKRKYYKKATRKKK
                  . **  *  .    **.    . **   ********** ***** * *****: * **

45              YKMYSRTRWWRRKKYWRMRGCKWYWWKKRRRSMKKKWYRKYRKCTTKMMKKKRWRKMRRR
46              CKMYGGCRTAGRTKYWRMRKAKWYWWKKRRRSMKKKWTRKYRKYYWKMMKKKRWRKMRRR
                 ***.  *   *.****** .**************** *****   **************

45              GCMSYRRARKYKMMCAKRWWRYTAKARMKYKKYKGAKYMWKMACSRRACTTACACCKRKM
46              KTASYRRRRKYKMMYWKRWWRCWRKWRMKYKKCGRWGCATKMGTGRRGYAYTTRYYKRKM
                   **** ******  *****   * ******        **. .**. : :    ****

45              RMKYGGAKRKMWKKKKRKRSWYTKYKMATRYKKTWRRMRRYYYYCATTWMRYGYYRKTTG
46              RMKYKKWKAKMWKKKKRKRSWYWKYKMRYRYKKWTRRMRRCYYYTRGAWMRYRCYGGKKK
                ****   * ************* ****  ****  ***** ***   :****  *  ..

45              YMYRGACTTKTYYTYRKMWGCAGATMWWWWYCCCWRYACGYYRYYKCTATYYWRSYKRWG
46              YMYRAGTKWKKYYYYRKMWKSRKGKMWWWWYYYMWRYGAAYYRCTKAGGATYWRSYKRWA
                ****.. . *.** *****  .  ..******   ***...***  *. .: ********.

45              YRKYRMYRKMRYMRMRMRKAKKYKYYSTTKKKWKKKKMYKRTGCGRKKKKKYMTRRYSAM
46              YRKYGCYGKMRYMRMRMRKCKKCKYYSCGKKKAKKKKMYKRGASRRKKKKKYMYRRYSRM
                ****  * ***********.** ****  *** ******** .. ******** **** *

45              TYMMCCKKRRRAYRKKYKYYTGYWKRYKKCKTCKKWKCYCYYYMMTTTRMMRKRMYAYRR
46              WYMMYMKGRRRTYRKKYKYYCACWKRYKKSKWYKKTKYCAYYYMMCCGRMMRKRMYRYRR
                 ***  * ***:******** . ******.*  ** *  .*****   ******** ***

45              AKMYYKWRYYKKMMRYYKRRRYSWKYRKRKKGRYRCYACARRRRRRYRYYYRYCCRTRYS
46              TKCCCKWRYYKKMMRYYKRRRYSWKCAKRKKTGTRTYMTCRRGGGAYRYTTRYYTGYRYS
                :*   ********************  ****   * *  .**    ***  **    ***

45              KKKRKTMMSRKRAAATYWKKYWGWWKMKMRTRAYRKYKGWRYRRWKRRWRRKYRYYKRKM
46              KKKRKCAMSRKRMMMYYWKKYWCTWKMKMRGRCYRKYKSARYRRWKRRWGRKCGTYKRKC
                *****  *****    ******  ****** *.*****. ********* **   ****

45              RWMMYWMCTYYYRYKYCCTTYMYYWKAYKRRKWRSYGKKATYYYAGTYYSRRTKRYYKAK
46              RWMMYTMSAYYCGYKYYYYWYMYYWKWYKRRKWRSYKKKWGYYYTTCYYSRRKKRYYKWK
                ***** *.:**  ***    ****** ********* **  ***:  *****.***** *

45              MMYWKTTAKYCAKKKAGWYRRWWKGYRGCRKKRYMRYKGYYRKSMRKCMTKKWTKSYWYK
46              MMYWTACTKYAGKKKRTWYRRWWKRYRSYRGKGCCRYTRYYRKSMRKMMGKKWKKSYWYK
                ****.: :**..***  ******* **. * *   **. ******** * ***.******
```

```
45                  ATAKYMKTKCYKSKRKKWYKKAAKRYKRKRYYKYWRRRKKKGSRKYKCTTCYWYTRWYYA
46                  TAGKYMKYGMCKSKRKTWYKKWWKRYGRKRYYKYWRRRKKGRSRKCKMKWYYWYYAWYYR
                    ::.****    *****.****  *** *************  *** * .  ***  *** 

45                  TTGTMKKYYKRKRRMYRRKRYKMWRRCYKKKCGWSKRYYKKKYMGYYRKRRYRRRGKYYY
46                  KYAWMKKYYKRKRRMYRRKRYKMWRAYYKKKATWSKRYCKKKYMKYYRKRRYRRRKKCCC
                    . . *********************  ****. ***** ***** ********** *   

45                  YRYKYWYYYGRCGCAAKYCKKKYSCKKMYKKKGMRYRRGYRWTAGATKYGRKRYMYKYWK
46                  YRYKYWYYYAAYRGTRKYYKKKYSYKKMYKKGRMRYRRKYRACCCGKKCARKRYMYKYWK
                    *********.    : ** ***** ******  ***** **  . ..* .**********

45                  AKMMTSMMSMYKCGYTYKCGYYCTRMRWYRSYTYRYAKKKKAGGTYWKSYYRYRYRRCCG
46                  WKMMYSMMSCYKYRTCYKATCYYYRMRWCRSYYTRYRKKKKRRKYYWGSYYGCGCAGSYT
                     *** **** **    **.  *  **** ***  ** ****    ** ***      .  

45                  CCRKKRKWCACGYKRWKYWARWMYRYRSMKMMRYYYARGMWMMWRGKRYRYRYRRGWYRW
46                  TARKKRTWYRYTYKRWKYWGRWAYRYRSMKMMRYYYTRAMAMMWRRKRYRYRYRRKWYRW
                     .****.*    *******.** *************:*.* **** ********* ****

45                  RWKMGYCKRMGTTRAWYWSWKRKGYKYYKMKKSKRKMYYYRRKAAARKCYKKRYYRRYAA
46                  RWKMKYTKRAKKWRRWYWSWKRKRYGTCKAKKSKRKMYYYARKWGCGKYYKKRYYRRYWR
                    **** * **  . * ******** *   * ********** **  .. * *********  

45                  RCRAGYKYKMWWYYRYKWAMMWGKKKYRWRCMTRWARAATKRTTYYTYTKTATYMKMKWW
46                  RYGWRYKYGATTCYRYKWRMMWKKKKYRWRMMWRTWARWWTGWCTYCCGKYRKYCKAKWW
                    *    ***     ***** *** ******* * *      .    *   *  .* * ***

45                  KKRMYRRRKKWWSYACAYKGYTAAAGYRRKKYMMWKATKKYSRAWYWWAGCKGSYWSKRT
46                  GKRMYRRRKKWWSYTMRYKRCYRRWAYRRKKYMMWKTAKGTSRRWYWWTAGKKSYWSKGC
                     *************:  **      .**********::*  ** ****:. * *****  

45                  TCCRYKYKKYRRMGYKYKKRKKAGAMKKWYMAGAKYKYWYYWKMKKRMYYRYRYKAYYYT
46                  YYSGYKCKKYRRMKYKYKGRKKWCTMKKAYMRRGKYKYWTYWKMGKRMYYRYRYKGCCYW
                      . ** ****** **** ***  :*** **  .***** **** **********.  * 

45                  KGRYTKGKKMYAKKKKYYCYMYYKSKRRTAAAKKAMKWYWRCKMYYSRTMRWYRKYRMYM
46                  KKRTWKRTKMYRKKKKYYYYMCCKGKRRATTGKKWMKWYWATGMYYSRYMRWYRKYAMYM
                    * *  * .*** ****** **  *.***:::.** *****   ***** ******* ***

45                  YYRTCTTTRCCMYYYKAKAGCTAAAKTWWKRKTCGGCTRTTTMYRKSRKMGGRYYKMCKY
46                  YYRWYWWAATAAYYTKGGCRYCGGTGCWTKAKWTRAYKAAAAMYRKSRKCRRRYYKMYGY
                    ***    :  . ** *. .   ..:  * * *   . . :::*******   *****  *

45                  WATAYKYRKRTYKKYSTYKMKMGGGAAKYCKYKKKKMRWKKWKMTATGTYRYYYYMYAKM
46                  TRYRYGYRKRYYKKYSWCGMKAAAAGRKYYKYKKKKMRWKKWKAKTATKYRYYYYMYRKM
                        * **** *****   ** .... ** *************  .:: .******** **

45                  MKYYMKMYSRKYRGYKKMRKMKRRRYKGCRKMYKWRMKRYMGWTATYRRRYMYWKWKKWK
46                  MTYYMKMYSRKYRKYKKMRKCTRRRYKAMRKMYKARMKRCCTWCTWYRRRYMYWKWKKWK
                    *.*********** ****** .*****. ***** ****   * : **************
```

```
45    KYKWYMYRYRAGYWRYRKKCCRTTGKWRTTTYRKGATAGYTGRMYGSRGKYATYGYRTYR
46    KYKWYMCGCACTYWRYRKKYTRGGAKWGYYCCRGKWGGACAAGCYRSRKKCWACRYRWYR
      ******    . *******  *  .**     *    .. :.  * ** *  :  ** **

45    KGYKCARMWRMKAGATKKKWMRGGTKWCYYKYKKKYAYRKWWKKYGTYRATAACYKATKR
46    GRYKYGRMWRMKRRRAKKKWMRRWKWAYYKYKKKYMYRKWWKKYRCYRMGGCYYKMKKR
        ** .******   :******   **.******** ********  **  .. ** .**

45    RYKKRWWSSSYCYMRWRKWKYTCCCKSKKMYKRRKWKKYSKGWWYTGMYGWGTKATKYYK
46    RYKKRWWSSSTYYMRWGKWKYYYYMKSKKMYKRRTWKGYSKRWWYACACTTKCKMYKYYK
      **********  **** ****    *********.** *** ***:       *  ****

45    KKKKWRKRRWWKRYAWMKSYKYWYYKKRYKYRYRKKKMRKYMYKKTGYRYWGYKRWSRYR
46    KKKKWRKRRWWGRTGWMKSYTYWTTKKRYKYRYRKKKMRGCMTKKYRYRYWRYKRWSRYR
      *********** * .*****.**  **************  * **  **** ********

45    YRYYRRRKKKMKKKKMGRAAWKCCGARRYRYYWYRRYRTRMYSKGWCRRMMRSKSKYRYR
46    YRYYRRRKKKMKKTKMKRGTAGTTTGRRYRYYWYRAYRKRMYSKSWTARMMRSKSKYRYR
      *************.** *.:     .********* **.*****.*  ************

45    RRCKKTGGTGCKRRGRRYKMYYRAYYSGKKWWRKYYRYRRKGSKKATTTGRARWKATYWT
46    RRAKKASTAAYKGGRAACGCYYRRYYSKKKWWRKYYRYRRTKSKKWYYYKRMRWKRGYWG
      **.**:. :. *        *** *** ************. ***     * ***  **

45    AKKCYCRYWKKYYAKKAATWRWMAMYYKMAYYMGYRGCRRKYKAAGAATWKCWWYWRRKK
46    CTKTTTRYWKKYYGKKCCCARWMWMYYKCRCYMAYRRMRRKYKWWSMWYWKTWWYWRRKG
      ..*   *******.**..  *** ****   **.**  *****  .   ** *******

45    YTRMYSWKYMCCARWYWRYAGAYYKYWAGRRRAKKMWMWCYYYWKRASRRYYKRRKMKWY
46    YKGCYSWKYATTTRWYWAYRKRYYKYTWAAAAMKGMWMWMYYYWKRMGARYYKRRKMKWY
      *.  *****   :**** *   ****  .    * **** ****** . ***********

45    RKKRRKWKKMKKCYYKRRKRRRKWMYMRAWGRRYRYMSWGWKKRYYAKWYYGRKRGKRWS
46    RGKRRKWKKMKKMCYKRAKGGRKWAYMRRWRRRTGTCCTCAKGRYCMKWYYRRKRKKRWS
      * **********  *** *  *** *** * **    .   * **  **** *** ****

45    GCCTCSMYYRRRKKKWKKYRKKAKWKKKRYACRYRGYRRKKASARAYKRYWYKRKKCCYC
46    AMAAYSMYYRRRKKKWKKYRKKRKWKKKACMTRYRTTARKKWGRGWTKRYTCGRTKTYYY
      . .: ***************** *****    ***   *** .    ***   *.*  *

45    ATATYYRYRARYKRYTTARGTKKWAGTTCRRWKGAACRMMMKAACMKCCKMKRCCKKYWY
46    GCGGYYRYRGRYKRYAAGRRCKGWWRWWYRRWKRRMYRMMMKGTAMKATKAGRYYKKCWY
      . . *****.*****::.*  * *     ****    *****.:.**. *  *  ** **

45    MMYYRCMRKRKRRYMWKRRMTATMYYRKMWKGRYKYMRYAYTYRKWKWSKKGGARMMKYG
46    MMYYGSARKRKRRYMWTRRMWRAMYYRKMWKKRYKYMRYRYKCGKWKWSKGRSWRMMKYA
      **** . *********.***  :******** ******* *.  ******  . *****.

45    KGTGYKYWYKKGGYRCKTYMRMYYRKACMAGRRRYACGRSYAGTCMTATTRCKRRYYYMK
46    KKCKYKYWYKKATCATGKYMGMYYRKGAMRKARRCCYAGSYWSYYMCCACGYGRRYYYMK
      *   *******.     .** *****..*   ** . . ** .  * .:    *******

45    KYCGTGGCCGRKKKCCATCYTKKSCYRCKKAACCRYYAGRRYRKRYKYMRKKSRSYWRRR
```

```
46      KYYRYRRTYTRKKGYYCWSYWKKSTTATKKRMYYRYCRRRRYRTRYKCCATKSRSYWRRR
        **        ***   . .* ***    **    **   ****.***   .*********

45      YKRMKYCWWMTGCYTYWRMRKWMMRMWTYRRYMYMCSKYMYWYMYTRWRGCKMYAGYYKY
46      YKRMKYMWWMATGYYYWRMRKWMMRMWKCRGYMTCTSKYMYWYMYARWRSMGMYWKYYKY
        ****** ***:  * ************. * **   *********:***.  **  ****

45      KMRGCYKYTMWWMRRYYKARYYRMMYTGTYWSKTTGTKSKCYMKKKYYWRWRMTCTACAY
46      KMRKYYKYKMWWMRRYYKRRYYRMMCYKYYWSKAAACGSKYCMKKKYYWRWRMWYYRYRY
        ***  ***.********* ******    ****::.  **  ***********      *

45      WWRAGKWYSKKYYKKRYYMRKYWARRATSKKWKKKTCTAKSRCMKWYKMKKKMKYYKYWY
46      TTAWRKWYSKKCCKKRYYARKYWGRRRKSKKWKKKYMWGKSRMMKTYGMKKKMKYTKYWY
             ******  ***** ****.** .*******   .*** ** * ******* ****

45      RYYYGMATMYYYKKKRWKKAYGRWRKYRRKKKRKRYMSKMRKKYGCATAYYATGYTRKWG
46      RTYYRAGCMYYYGKKRWGKWYRRWRKYRRKGKRKRYMSKMRKKYRYRYWTTRWKYKRKWT
        * **  . **** **** * * ******** *************          *.***

45      GGARKRMKKRWYYKMSYRMKMYSGMKYKRYKKYKYTYYKRKWYRRWGGCACYYWCMMYWA
46      AAGAKRMKKRWTTGMSYRMKMYCRMKYKRYKKYTCCYYKRKWYRRWKSYCYYYWMMMYWT
        ... *******   ********. *********.  ********** . . *** ****:

45      ACKWKWKKTMKRYKKKRCCAAKKCYWYYYMYGCCMYWMCAATKKKWKYYKKTAARYRKKY
46      TTTWKWKKYMKRYKKKRYYRRKKYTACCYMYRYSCYTMATCAKKGWKYYKKGTGRYRKKY
        : .***** ********    **     ***  . * *.:.:** ****** :.******

45      RGRYYRMMRYASRYCRYYGTRMRRYKKYWWYKMRKTGATGGWKKKYMYYGKKKTKRYWGA
46      ARAYYRMMRYMGGCMGCYTCRMRRYGKYATTKMRKKRGKKKWKGKYMYYKKKKYKRYATC
           ******* .     *  ***** **   ****. ..  ** ***** *** ***  .

45      YYWRMCTGTMRMRAYKYTAMATYAATCCYYYRRCRRRYRMYKTCRRKMKMWRCTKKWYKW
46      YYWRMYCACMRMRRYKCYGATGCTWCAAYYYRRYRRRYRCCKKAGRKMKMWRACKKTYKW
        *****  . **** **  . :  :  ..***** *****  *.. *******. ** ***

45      RKSYKYMGMYCKSRRYKWRMKRYYGYRYRYYYYWYRSACCCMKYRWKRWKMRRRKKWWGY
46      RGGTGYMRMYTKSRRYKWRMKGTCRYRYRCCTYWYRSRYMYMKYRWKRWKMRRRKKWWRY
        * .  ** ** **********    ****   *****    ***************** *

45      KYKMRWRRKRYMMMYYYRAKRMYRYYYRKYKCTGAKMKSRYYRRYRWKGAKMKKKRGYMS
46      KYKMRWAAKRYMMCYYYRWKRMYRYYYRKYKYYRWKMKSRYYRRYRWKSRKMKKKRKYMS
        ******  ***** **** ************    *************. ****** ***

45      WKWAATGYRRAYRRGMSKRRKRYKRYGYKKAYRMGTKTKKTYYTTGCCARKRRRKYCCYC
46      WKWRRYRCRRMYRRKMSKRRKRYKRYTYKGCTRMRKKKKKATCKAAYTRRKGRRKYTTYY
        ***     ** *** *********** ** . ** .*.**:  .:.   ** ****  *

45      YTGGGAMTTWTAAGCYKRKKRWTTAGAGCTYYTCGCGYRRRRKACRSMRKWWYKYKYKKY
46      YYKTARMYCTAGGAAYKRKKRWYWCRWKYYTYGTRARYRRRRKRYRSMRKWWYKCKYKKY
        *   . *   :....*******  .      *   . ******  ********* *****

45      SWRKMMWYRSKKWKKAARYMKAACGAGMKGRKTKKKYYARYSKKTKKRKWWYKCAWTAAA
46      SWRKMMWYRSKKWGKGRRYMKCCAARAMGAATWKKKTTGRYSKKAKTRKWWYKYWWCMTT
```

```
          ************* *. ****.... .* . . ***  .*****:*.******  *  ::

45        YAGCGWKWYRARGATTCYGGWKCGRRRGRRYAGTKCYKMRYYKWRWKAYSYSKGYGAKYS
46        CTRYRWKWCARRRRYYYYCAWKMSRRRRRRYGSYKTYKMRYYKWRWKGYSYSKRYRMKCS
           :   ***   *     * .** .*** ***.. * ***********.***** *  * *

45        YMTTYKRRAGCWKGAACWGYTCGMCKKTMYKTACCAGRRSRKYRRAKWWTTCGSYYC
46        YMYYYKRRRRTWKKGTGASCWYRAYGTCMYKWRYMMRRRSRKYRRMKWWYGTASYCT
          **  ****   ** .:  .       . ***      ******** ***   .**
```

The pairwise alignments of two accessions 45 and 47

```
45                  GCYRKRWYKWGYCMWAWMMKRYYYKMYYRYRRMMTKYRYRCYYWWCYGRRMSGMMYRCYR
47                  KMCAKRWYKWTCTMARWMMKRYYYKCYYRYRRMMKKYRYRTYYWWMYRRRMCKCMYRMCR
                        ******   *  ********* ********.***** **** * ***.  ***  *

45                  MRCYCRMYWRYGKMYYCKYMGKYWKKMYMMRWKRRWKRYKRYAYYRYYYKYYYYKRRKWR
47                  MAYCYGMYWRYRKMYYYKCMKGYWKKMYMMRWTAAWKGYKRYRYYGYYYKYYYYKRATAR
                    *     ***** **** * *  **********.  ** **** ** ********** . *

45                  YMRAYRSWMKKWRRYYYRYWWRYWYMSYTWYWWWWYYWYRWKYYWRKMYKYRYRRACCRT
47                  YMGMYRSWMKKWRRYYYRYWWRYWYMSYCWYWWWWYYWYRWKYYWRGMYKYRYRRWTTRW
                    **  ************************ ***************** ********   *

45                  RYRYWYRRYKKYWCGYCYYTTARRYMYYYCGMYKSCSKMYKMKYKSKRMRYWYMYRKRRW
47                  RYRYWYRRYKKYAYTYYYYYWRARCMYYYYKCYKGTGGMTKMKYKSKRMRYWYMYRKRRW
                    ************   * **    * ****   **. . * ********************

45                  WARAAMKMMRYMKYWKKAYWYWYWRWKRKYMKYGMYWKRWWKYRRYRGCKYYTRMKMKYR
47                  WRRTWMKMMRYMKYWKKWYWCWYWRWKRKYMKYKMYWKRWWTYRRYRTAGCYGGCKMKTR
                    * *: ************ ** ************ *******.***** .  *   *** *

45                  KYKTYWWWTKYCKYYKKRCRYKGGTCYRKYYKYWMKKYSRRYRKYYSYYMKKRRYWMWKM
47                  KYKYYWAAKGYTKYYKKRTRYKAKATYRKYYKYWMKKYSRRYRKYCCYCMKKARYWMAGA
                    *** **  . * ****** ***. : ******************* .* *** ****

45                  MRYRWYSRSMKMKWMRKRCTRTMCKCAGYMRMMYSWWYWYKYMRYMYKRKKKRWKACGKR
47                  MRYRWYSRSMKMGTAAKRMWACCAKTGAYMRACYSWACAYKYMRYMYKRKGKRWKTTAKR
                    ************    **     .* ..***  ***   *********** ****: .**

45                  YSKGAWYWYYRMKKYWRTYYYYWRYRYKSAYYRKAGRKRYCKKRRCMTRRKKWCTYTAGS
47                  YSKKRWYWYYRCTKYWRYYYYYWRYRYTGMYYRKRKRKGTTGKRRYMKGRKKWSYYYCKG
                    ***  ****** .**** *********.. ****  **    *** *. ****. * . .

45                  MARYKRGRYWTKGARSAGTRAYTAYYTWTKTCGKRYYKKYMRKYRMRSYWRRYRGCAKRM
47                  CGRYKATAYWYKATGGRRKRMTWGYCATAKYSSKRYYKKCCATYRMRSYWRRYRTTCKRM
                     .***   ** *.: .  .*   .* : :* ..******   .***********  .***

45                  YYKKWWMWMYRYRKGTAWMMMGRARMWMKSMCCGGYGMCARWRKYGTKMRAMGCKRMMRR
47                  TYKKWWMWATRYRKRAWWMMMRRCRAAMGSMTAARYKMGWRWRKYKKKMRRMKYTRMMRR
                     *******  **** : **** *.*  * ** .. * *  ***** .*** *  .*****

45                  YYRYYRMMYWYKRCTRKGKSRGGTGGATCWTTYTTYTTRWRASRYGRACAKYWKSMRKKY
47                  YYRYYRMMCWCKRAWRKAGSATRAAAGAMWYWYWCTCGRWRRCACRRRYRGYWKSMRKKY
                    ******** * **. **. *   :...: *  *     *** .   *    *********

45                  MRKKWYRYWKKMKRWRKKRYRYKKRKKYAKKYTKKKKYRYKCTTRKKKRKMMYWRKRGGW
47                  MRKKWYRYWKKMKATAKKRYRYKKRKTCGKKYKGKKKYRYKYCGRKKKRKMMYWRKRKAW
                    *************   **********. .***. *******   ************* .*

45                  KWRYRMWYRKYKWRRKYRRKRKKKAAKYYSKGRRGAWKTTKRMMRRKKSKKYYTKKKYMK
47                  KWRYAMWYRKYKARRKYRRKRGKKRRKYYSKKRRRWWKKYKRMMRRKKSKKYYKKKKCMK
```

```
            **** ******* ******** **  ***** **  **. *************.*** **

45          YRYRYKKYWRWYKMKWYKKRRRSMKRKYRCTTMKRKKKSTTCATCYMTAMRKYMMRKSKG
47          YRYRYKKYWAWYKMKWYKKRRRSMKRKCRYYWMKRKKKSWWMWKAYMYRMRKYMMRKSKK
            ********* ***************** *   *******    ..**  **********

45          WKKWYGCGSAGTTRYKKKKMKYYKYMWKYRYKYRKTTAGAWYYYYYKKRWWKMTAACYCT
47          WKKACRYSSRTCAGCKKKKMKYTKTATGYRYKCGGGYTCGTYYYYYKKRWWGMATGTCYG
            ***    .*   :  ******* *    ****     : . ********** *::.

45          YKGWRRTTKRYKRYGTTKYYKSRMKRKAGYGTYKRMKATGKYKWTKRRKRKCTYCKRWWY
47          YKRWRRCKKGYGRCTCCKYYKSRMKRKTAYRKTKRMTRWKKYKAAKRRKGKYCYYKRWWY
            ** *** .* * *    **********:.* . ***.   *** :**** *  * *****

45          RMKRKSGMAAYTKKTKRWGGGRYKKYYTKYKKWAGCYRKRWYWKRTYRKRGRYRMKWRRY
47          RMKRKSRMGTCCKKGKRWRRKRYKKYCCKYKKWMRMYRKRWTWKGYYRKRRRCGCKWRRY
            ****** *.:  ** ***   *****  *****   ***** **  **** *   *****

45          YYRYGTTKKWKMKKTKKKWYKSYWYKKYMGRKKKKTRGYWKAARCTTKGGRRYYWRRRRK
47          YYRYACWKKWGMKKGKKKWYKSYWYKKYMKRKKKKKRKYTKWWRYYYKKSRRYYWRRRRK
            ****.  *** *** ************** *****.* * *  *   * .**********

45          KSYRRKKYWAAAGTYYMKYWWKCKRKRKRMRCCTKKWRYKTSYKAYCGYYKKKCTRMAYY
47          KSYGRKKYWCGTCYCYCKYWWKYKRKRKRMAMTAKKWRYKGCYKCCYKTCGKKYKACCYY
            *** *****..:   * ***** *******   :****** .**.      ** .  .**

45          RKWRKTRRRYKYYYRYRYWYYYYWKACKKKRCAYRYYRRKKKYCSRYMGRWKWRRYAYWR
47          RKWRKYRRRYKYYYRYRYWYYYYWKMTKKKAYRYRCYRRKKKYMSRCMKRWKWRRYMYWR
            ***** *******************  ***   ** ******* ** * ******* ***

45          GAAAKKRMCTRCYYRYRYRSYKYRMKYYKWYGKMCRTTAARYKRYYRTCAGCAKCGWCKR
47          RRWRKKRMMCAYYYRYRYRSYKYRMKYYKWYRKMTACCGCRYKRYYRGTCRYGKMATMTR
                ****    ******************* **    ..*******  .  .* .   .*

45          KSYWKYYSWGGRTMYYKYRRYYGTMKRYWRMWRWYRKRCKMTMKTYWRKKKYTKCYMGAC
47          KSYWKYYSWTTGCMYYTYRRYTRAMKRYAAATGWYRKRTKMWMKGCWRKKKYWKYYMRGT
            *********    ***.****  :****     ***** ** **  ****** * ** .

45          AYGATMGMKCSRRYKYWRKYRKYRRSWSKRSCCTCKRWWWWKAKGRSYCKYKGYMKMRWA
47          CYAMWMKMKTGAATKYWRKYRKYRRSWSKRSTGKAGRWWWWKRKKGGYYKYKTYMKMRWC
            .*.  * ** .   *****************  .. ****** *  .* *** ******.

45          CCCKGCCAAKYYYRRMRYMYKYCTGAGYRGAMKKRYYMKRYRWKKCTGAMGTRCKKKRYY
47          TMTKATYTGKYYYRRMRYMYKYYYRWTCGSRMKKRYYMKRYRWKKTASRMAYRYKKKRYY
               *.  :.*************       . ************** :. *. * ******

45          YMSWRYMMYSRRKRYKSMKRYYKARKTMMAKRRYWYSMKYKKRWWSYKRRYKYKKWKYKY
47          YASWRCMMYSRRKRYKSMKRYYKCRKYMMMKRRYWYSMKYKTRWWSCKRRTKYKKWKYKY
            * *** *****************.** ** ***********.**** *** *********

45          GGGGGKCGRKYSSRRWKKSYWYMTGRYGKTATKYRYYKYYKWKRRTTWCGWMKSYYKRKW
47          SKRSRKYKGKYSSRRWKKSYWYMACACTKYMYKYRYYKYYKWKRRWYWYKWMKSYYKRKW
            .  . *   **************:    *   *************  *  **********
```

```
45              RYKKKYRYRKKKKKAGRKARYMCYKKTGRRYWYKYKWSKRYYRYYTAACRKKKMMKKACG
47              RYKKKYRYRKKKKKGARKRGYMMYKKYRRRYWYKYKWSKRYYRYYCTGYRKKKMMKTWMK
                **************..**  ** ***  *****************  :. *******.

45              MRWAAAACRYRTYRYRAKKYKATRRKSKKYKGCAARAYTSGCRKTKCCCAKKGGRRYYTG
47              MRARWTWTRYRCYRYRRKKYKCCRRKSGKCGRAWGAWYGGTTRKKKYYYRKKCARATCCR
                **   :  *** **** ****. **** *   . .  * .  **.*    ** .*

45              ASTKKRKYRRSGGCYTRWKSYWRKKRYYYYRMMKTWRKWYMTKYMRCKTYYTMACYRRYC
47              GSKTKRKYRRSKKSTYGTKSYWRKKRYYYYRMAKKARKWYCAKTARMTCCTCCGYYRRYG
                .*..*******  .    ************** *. **** :*  * .     . ****

45              WRRKRWKWGKSKKWRRYYWRSRRKRYSWCTTTASYMRWYWRYRYYRYYRRKMWMYWTKYM
47              ARAKRWKWRKSKKWRRYYWRSRRKRYSWTCCCRCYARWYWAYRYYRYYAAKMWMTWCKYM
                 * ***** *******************     .* **** *******  **** * ***

45              GMYSRRAARGYACYRYWRYRRKMKRKWYRGARKWKKMKKYYRYMKGYKWMYMRAWMMRYR
47              KCCGARMRARTGYYRTAAYRRKMKRKWYRCGRKWKKMKKYYAYMKKCKWACMRRWMMRYR
                   . *     . **   *********** .********** ***  **  ** ******

45              YYYYWYYRWTKTTKKKYKSMYYYRKRKRRRYKKATYKWKATGTKAYRAKRRKKKTGCTTK
47              YYYYWYYRACGYWKKKYKSMYYYRKRKRRRYKKGACKWKMWKKKWYRMTRRKGKCAMCCK
                ********     ********************.: ***   .* ** .*** * .   *

45              YMKKRYCGKGRKKYMMRKYYYSKKGWYKSKYKMYRGRKKMSRTMRKCGYSTYMMTYRRSY
47              YMKKACMRGKRKKYMMRKCYYSKKRWYKSKYKMYRARGKMSRCMRKMRYSKYMMCYRRSY
                ****      ******** ***** **********.* **** ***  **.*** *****

45              KKTYRRRKKGMMYTYYWKYAYKCKKKTGKYAMKKRYYYMMSYWMRKRTMYAYRRKKKMYY
47              KKKYRARKKKMMYYYCWKYGYKSKKKWRKYGCKKRYYYMMSYAMRKRKMYRYRRKKKMYY
                **.** *** *** * ***.**.***  **. ********** ****.** *********

45              KWTCTTYKKMRYMRYYMRSWRKTKWKYKMAYMYRYYRWTYAKRYKYKAKMTTAAKRRSMM
47              KWCTGGYKKMRYMRYYMRSWRKCKTKYKCCYMTRCYRWKYRKRYKYKWKMCAGMKRRSMM
                **    **************** * *** .** * ***.* ****** ** :. ******

45              MYKTMWCYCTRMRWYMKSRKKKAYWRRRCGGCRKACKRTGRAKRKWKAAKTATGRWRMMY
47              MYKCCAMTYYRMRWCCKSRKKKRYWGGRYSKYRKWYKRGRAWKRKWKGGGARWRGWRMMY
                ***       ****  ****** **  * .  **  **    *****.. :    *****

45              WYWRMAWRMYTYGRGAARRGYWWYRWYRGACARMWYKKKKSKRKYRATRATCCAAKKKSR
47              WYWRMWWRMTACAGTWRRRKYAWYRWCRRGTGRAWTKKKKSKRKYRWYGGATTGGKKTGG
                ***** *** : .    ** * **** * . .* * **********   .:  ..**..

45              YYKRMWKATATRYCTARYKKWKRWKKAGTTACAGMMMWMYGGGYKGRWKWCRYRMWYYKW
47              TYKRMAKRCGKRYTAGRCKKAKRWKKTAAARARAMMMWMCCTRYKKRWKWSRYRMWCYGT
                 **** *  ..** :.* ** *****:.:: . .*****    ** ****.***** *

45              YCATMMGYYYCTCACKKRRYTAAGAWMTKSKMYYRMKRYKYKAKKGYRRKWYWSYMGTRA
47              TTTCMMAYYYSAYRAKKRRYKCCACWMCKSKMYYRMKGCKYKRGGKYRRKWYWSYMAWRR
                  : **.***.:  .*****.....** *********  ***    **********. *
```

```
45      TTKGYMKYYWYRKRWGGCYYWKWRRRYYWMAYGKWMKRYYKKKKKGYYKMKYTGRKYKWR
47      CYGKYMKYYWYRKRWKSYYYWKWRRRCYWCRYRKWMKRYYKKKKKKYYKMKYWKRKYKWR
            *********** . ******** **  * ************ ******  ******

45      YKKTAMRYKKYRRYRMMKRWYYASRYCRYRRKKWYWWYKRKRKTKKKKKKKYMKYYKKKK
47      YKKYMMRYKKYAAYRMMKRWYYMGGCMGCRRKGWYATTKRKRKAKKKGGKKYAKYYKGKG
        ***  ******  ********* .     *** **   *****:***  *** **** *

45      YWCGAKARYYWRAYMTMRMRYAAGCWRKYYWKKYRRRRRRYCKMMYYRRKRKKYYYYRRK
47      YAMTCKRRYYWRWYMCMRMRYCCCYWRKYYWKTYRRAGAACMKMMYYGRTRKKYYYYRRK
        *   .* ***** ** *****..  *******.***      ***** *.**********

45      MGYYMYYAWRMAWRMCYTTSRRKKKKKKKRYRYYWSGYRYMKKKRKMRRWKGTCGWMRRS
47      MRCYMYYRARMWWGCYYKKSRRKKKKKKKRYRCYWSSYRYMKKKRKARRWKTYTRWMRRS
        *  ****  ** *   *..************* ***.********* ****    *****

45      RYSYTTKKKYSKKKYRYWYRRRYKYYRYKKYRKMCKKRKMRYYRAWKWYMTKKKYKYMGK
47      RYCTCYKKKYSKKKYRYWYRRRYKYCACGGCGKCYKKRKMRYYRRWKWYMKKKKYKYCTK
        **.   *******************       *  ********* *****.******  *

45      KWYRKMMYMYCRWRGTTSATKRYYKKKRKKKRAYYYTRKRWRYKKGGKASYWKRKKKWKK
47      KWYRKMCYMYYGWRSKWSRATRYYKKKRKKTGGYYYAGTRWRCKGKRKRSYWKRKKKWKK
        ****** ***  **.. * :.*********. .***: .*** *   * ***********

45      RRWRRKTGCYYYMYKMMYMKMSRRKYRYRRSKMRKKMRKRMAGAYYKYKTYGGTYTMYAR
47      RRWRRKCTTYYYMYKMMYMKMSRRKYRYRRSKCGKKMRKRCCATYYKYKYYAAATYMYCR
        ******   ***********************  ****** ..:***** *..:  **.*

45      YYRRRYWYMMWKKYWMTWYCKTATKYWWKCAGYKCATWRYRKKTRKKWYKKYRGTYRKYY
47      CYRRRCWYCCWKKYWMYWTYKKCWKCTWKGGKTKYMKWRYRKGWRKKTYKKYRRYYRGCY
         **** **  ****** *  *.. *  ** .  *  .*****  *** *****  **  *

45      KYCARGRMRKYYKKWMRGGWYYKKKKYRYYRTWWKKYARWAMRMYYRYKGRYKYTCCGRK
47      KTYRGRRMRKYYKKWMRRRWYYGKKKYRYYRKWWKKCMRTCMRMYYRYKTRYKYCTTARK
        *     ***********  *** ********.****  * .******** ****   .**

45      YAKYKYYMYMMYATGGARRGKKKYKWYAAKKKYSRKRYMYKYRMYYRTRCCGMAYRTTAA
47      YRKYKYYMYMMYWWRKRRRKKKKYTACRWGKGYSRGRYMYKYRMYYRAATARMRYRKYRG
        * **********     ** ****.     * *** ***********:  . * **.  .

45      RAMCMKKKTKMYRCRKYKAMKATAWSYWYKYRKRYKYSCCGYKYACATKWYYKYRWKKAT
47      RCCYMKKKYGMYRYATYKWMKRYRWCYATGYRKRYKYSYMKCGCMYMYKWYYKYRWKKRA
        *.  ****  ***  .** **   *.*   ********          ********** :

45      YRYCGTGRRYAGMYTYMWGTCTKTGRCYAKKCTWWWWKGYRYYRMCAWWYKRWKCGCGRR
47      YRYTTGTRRYGKMYWYCWACTYKKRGYYWKKYKWWWWGRYRCTAMMRWWYKRWKMSSKRR
        ***    ***. ** * *.   *.   * ** .****  **   *  ******* .. **

45      RRRYKYRYKKYKWYKKGYRKMYWYCCSYCGMAGTYRRTCRYRCCTAYMRCAKTCGTCKMC
47      RRRYKYRYKGTTWYKKRYGTCCAYTTSYSSMWKWCRRKTRCRYTATYMRMMKYMSWYGMT
        *********  .**** * .   *  **..*    **. * *  ::***  *  .   *

45      YKAATWMGWTCSKRRKACAAYRKYYCRYRKYRCATTAWRCAKRTKMSKYYCRWKGTRYWR
```

```
47                  TKGTAWMRWCTSGRRKTAMGYRKYYTRYRKYRYRAAGTRMRKRCKMSKYYYATKRKRYWR
                     *.::** *  * ***:. .***** ******  ::. *  ** ******   * .****

45                  WYKYWAAKKTAGKMYYYRYRYRRKKKTTGKKRYRMRWMMWYKYAYMYTGTYRYCCARYKR
47                  WYKYWGWGTAWRGMYYYRYRYRRKKKKWRKKGTRMRWMMWYKYWYMYYKYYRYYTTRCKR
                    *****.  .:   *************.  **  ********** ***   ***  :* **

45                  RKWRSTMYCTYKMYYYYYSRRYYKKWRYWWCKWWKYTTYKRRMTAGKAYCTKYYSGCARK
47                  RKWGCKAYSYYKAYYYYYSRACTKKWRYWWMKWWKYYGYKGGCKTSKGYTCKYYSATRRK
                    *** .. *. ** *******   ******* *****  **   .:.*.*  ****.  **

45                  RRKATTCMYWWYTMMRRMRKYKRKRCAGATTTKYMKYYKMYYSWGYWKWMYKYAGCCKKK
47                  RRGRCGTMYATYKAAARMRKYKRKRMWKTWGAKYMKYYKMYYSWKYWKWMYKCTTGAKKK
                    **     **  *.   *********   :  :************ ******* :  .***

45                  CKWRKYWKTRYTCGCTMKKKRKRMCAKKRYATATSYRKKKYRCAYRWRRKYYMMKKMYWS
47                  YKWRTCWKYRYCYATCMGTGRKAAYGKKRYMKTASYGKKKYRTWYRWRRKYYMMKKMYWS
                     ***. ** **  .  * . **   .**** .::** *****  ****************

45                  YMCKMRMKYKWWYYKTATTKKMKKTKRRSWCTCYKMRYGMRTCYYWMRYWATYMYGRKMY
47                  YMYKMRMKYKWWYYKGCCCTKMKKKKRRSWYYMYKMRYCMRWMYYWMRYWTYTCYKRKMY
                    ** ************ .  .****.*****   ***** **  *******:   * ****

45                  KKKMRRATKTGWMMWKKACGTWKYKYACGYGCKAWAARKMRYMRMRMRMKRAAATGTKKK
47                  KGKAGRRKKYKWMAWKKTMRCWKYKYGAAYRMKRTTGRKMRYMRMRMRMKRRWRYSGKKK
                    * *  * .*  ** ***:   *****...*  *  :.**************    . ***

45                  RYYTGTCRKCKRYYMRRYKGYMWWYRYKRMWAMMYWKWRWCTTYYYWTCRKKRWKRRRMY
47                  GTCAACGRKYKRYYARRYKRYATTCRYKRMWGCACTTWRWSYCYYYWKARKKRWKRRRMY
                       :.  ** **** **** *    ******.    .***.  ****..***********

45                  RRRWWWKRKYRRKWKATWKYCGAKRCCCCCAGCYKRYSWSKRTTCRKYRGMYYKKRKKMK
47                  RRRWWWKRGYRRKWKWAAGTSKRKRTTMYTTAGYKRYSWSKGCYYGKYRRMYYKKRKKMK
                    ******** ****** :   .  **     :. ********     *** **********

45                  KGACRKKKWKKC
47                  KKGTAKKKWKKY
                    * .  ******
```

The pairwise alignments of two accessions 45 and 48

```
48                    YRRYKRWMYKMMMYRYKWWRRYWSGCTYYYKYYTTYWRYRYRAAAGKMKYYRCWWYRWRY
45                    YRRYKRWMYKMMMYRYKWWRRTASAAAYYYKYYCWYWRYGYRWWRATMKYYRTWWYRWRY
                      *********************  *..:******  **** **   ..***** *******

48                    YKSRYYYMRYYRWRWKYRYYGGRYRYRRWMKRMRRKRSWWRCKKYMKRYYYYYYRMGYKM
45                    YKSRCTTARYYGAAAKYRCYSARYRYRRWMKRMRRKGGWWRTKKTAKRYYYYYYRMAYKM
                      ****    ***    *** *..************** .*** **  **********.***

48                    KGRRYKYAATTTAGSRRMMMWKKYRYYKRMYYYYKWWYRYKRYWMWRGYKYMRRWCWWRY
45                    KTRRYKYGMCKKCAGAAMMMAKGYRYYKRMYYYYKAWYRYKRYWMWRAYKCMRAWYAAAC
                      * *****.  .....  *** * ************ ***********.** ** *

48                    YMKYMRRMRWWKKWKRRMCAYMYTTWWWGYYYTYCRRCRKRTAMKRRKYRTRYMRMMWYY
45                    YMKYMRRMRWWKKWKRRMYTYMYCAWWWAYYYCCTGAMRKAWGMKRRKYRYRYCGCAWYY
                      ****************** :*** :***.***      **  .******* **    ***

48                    ARKWCYYRRWRWYGMYMRWCGWRYSYSRWYKGGYYYRGCTYGRRMRRYYYWMMMYYMRYY
45                    TGKATCCAAWRWYKMYMRWTAWRCGCSRWYKARYYYARYYCSRRMRRYYYWMMMYYMRYY
                      : *      **** ***** .** . *****. ***     .******************

48                    YRKYRYYRRYWWGGYMRYYRARCCSKGKYRYKRRGWRRKYKWYRYGWTRKYATRYWKKSK
45                    YRKYRYYRRYWWAAYMRYYRMGYYSKRKYRYKGRTWRRKYKWYRYSTWRKYTGAYWKKSK
                      ************..******    ** ***** * **********.  ***:  ******

48                    TYKSRCTCGRYYKRGCATTACTKCRYRMMRYMTWYMYMYKWRRRRMCAGGTTYWKYRYYY
45                    CYKSRTCAARYYKRRYGGKTAYGMGYGMMRYMAWYMYMTKWRRRRMACATACYWKYRYYC
                       ****  ..*****  . .:.    * *****:***** *******... : *******

48                    KKKYSKWTRSYRKAAMKYYATATKRYYSWMYYYKYSKGWRYRTKAKKYKRWASKKWKKKY
45                    KGKYSKTGRSYRKRRMKYYTCGKTRYYSWMYYYKYSKKWRYRKKRKKYKRWWSKKWKKGY
                      * ****  *****  ****: ...************* ****.* ****** ****** *

48                    RKKYKRYKMMKMGAARKGAGTRAAYYTSYKKAKAWCKRKYCTGCCTTYWATCRKKSYRRK
45                    RKKTTACKMAKMAGGRKRWSYRGGTYKSYKKTTGTMKRKYYYRMTAAYWMWMRKGSYRRK
                      *** .  ** **...**  . *.. *.****:..  ****     ::**   ** *****

48                    RKKWKRMMYKRYKGYGAYYTRYKRRRYKYTAKYKKRYAKTRKKMKKKKRYCYWYRWYKRW
45                    RKKWKGMMYGRYKACTTYYGRCKRRRYGYCGKYKKAYTKAGKKMKGKKRYYYWYRAYKRT
                      ***** *** ***.  :** * ***** * .**** *:*: **** **** **** ***

48                    GTARRRWGRKKATYMRYSRGSMRGGYMKWKGKMRRTGAYKRRYYKTTWYYKTRRMYGAKR
45                    TGCRRRWKRKKWAYCRCSRRSAAAAYMKWKTKMRRGTTCKARYYKYKATYKYRRMYKRTR
                        .**** *** :* * ** *  ..***** ****  : * **** .  ** ****  .*

48                    RWKMCMKKWRWGTGTCRTMYKMRRYRRRCACCGACTKMYMTYKRRSRYRTGCGRRAGRYY
45                    RWKMMCKKTGWKAAWYRYMYKMRACRAATWSMRRMYKMYMKYKRRSRYGCCGTRRTTRYY
                      ****  **  * :.  * *****  *    .     ****.*******     **: ***

48                    RTRYWWACRAARAAGKRWSKWKGWWCGATKKKCGCRMRYYYTKYMRMYKRKYRWAAYYAA
45                    RYAYWWWYGGWRWMRGRWSKWKKTWGTRYKKKTAAMRYYYKTYMRMCKRKYRWMGYYRM
```

```
            *  ***   . *    ******  *    *** .. *****..**** ****** .**

48          RWMAKSYYSRSATGCWMTWSYKYCCATACYYRKKYKGGAKRMARMYKGKSYRYKMKRMWY
45          RWMRKSYYSRSGGCTTMYWSYKYYTGKGSYYRKKYKCCTKRMMRMCKKKSYRYKMKRAWY
            *** *******.    * *****  ....*******  :*** ** * ********* **

48          KKKRRYKGMTKYWYYRSMGAAKATGRKKKKKRYWYYYRRTCKYRKKRKYKRYRRKWKKRW
45          KKKRRYKRMWKYWCTRSMTGCGTAARKKKKKRYWYCCGAYYKYRKKRKYKRYRRKAKKRA
            ******* * ***  *** .. ::.**********      ************** ***

48          KKWKWRGYKRKTKTYYYMKGRATCMGRYKACARKWAMSYKRKRKKWRYKRRKKAAAKYRY
45          KKWKWRAYKGKCKKYYYMKRACCTMKRYKRSMRKWWAGYKRKRKKWRYKRRKKTCTKYRY
            ******.** * *.*****  .  * *** . ***  .***************:.:****

48          TGTKTAAMYYYRWKRRYAYWGRGYKCGTCKKTCGCGCKSKYTTYYYYRYYKKWYKMKYYA
45          ARAGYMTMYYYRAKRRYRYWKRRYKTAAAKKGYKYRYKSKCCAYYYCACYKKWYKMKYYW
            : :   :***** **** ** * ** .:.**      ***  :***   **********

48          AGTKMKYRKWKAKSRSKKKKGTRKRKMKRSYRCCAYSWWMKWSYYCKRYKKWKATCTKYK
45          WRYKMTYRKWKMKSRSKKKKRYAKRKMKRSYAMMRYSWWMKWSYYMKRYKKWKMYYYKCK
               **.***** ********   ********    ********** *******    * *

48          KWMRRMMWRKWMWKKKTKCYYKWWYSTRWRKWAGCAYMWRRWWMRWYGMATYMYMRRYKW
45          KWMRRMMWRKWCATKGKKYTYKWWYSYGWRKWTTTRYMWRRWWMGWCCMGAYAYMRGTKW
            ***********  .* .*  ******  ****:   ******** *  *.:* ***  **

48          WACRWKSMRCACKWKSKYYKYMRAKSWRYKCCKWTYWYRKKMKRRKMMRGKRKCGAYYYG
45          WMYRWKSMRACTKAKSKYYKYMRGKSWRYTYTKWCYWYRKKMKRRKMMRKKRKATTYYYK
            *  ******.. * *********.*****.  ** ************** ***. :***

48          ACKKKTRYRTWRRMTGYYRCAGGKRKYYTRKCTWKTCRGTTMKMRWMRWKGGGKWKRAGK
45          GSKKKCRYRCWRRMYSTTAYWRSKRTCCAGGTGWKGSRKGCAKCRWMRAKAAAKWKRTRK
            ..*** *** **** .      .**.  :    ** .*    * **** *...****: *

48          KWWSKKRYKRWKYRKKRWWGCKRRKRYYTRYYRKCAAGKYKMYGRRMMYYYTWYYAASKR
45          KWWSKKRYKRWKYRKKRWWRYKRRKRYYGRYYRKYRMRKYKMYKRRMMYYYYWYYGRSKR
            *******************  ******* *****    ***** ******* ***. ***

48          KKWYCACCTRKKYMRGRCWTKYRWKCAAARKRYYWYAACTWTAGKKWRAYGCWYKKYYYC
45          KKWYTGATARKKYMRARTTYKYRWKACGGGKRYYWYCGAGWYRKKKWRWCKMWYKKYYYM
            **** .. :******.*   *****.... ******... *   ****    *******

48          YYKYSMMGWKMKYYYYYKYWMMYKRRRYRMKWRKKTCKAYYKAAKMRKRKTGYAKGRYCT
45          TYKYSMMRWKMKTYYYYKYWMMYKRRRYRMKWRKKYMKTYYKWWGMRKRKKKYGKTRYAA
             ****** **** **********************  *:***   *****. *.* **.:

48          CMYWGRYYAGRYARMYMYCCGTWKKMKGCAAYKRYTTTWWKRYRRRKKWRKKGCMRYTWW
45          TMCTKRYYRRRYRRMYMYYYRAWKKMKRYGCCKRYYYYAAKRYRRRKKWRKKTAMRCWWW
             *   ***  ** *****   :*****  .. ***     ************ .**  **

48          WRWRKKGMRWKWRAYYYYRKCTTRTKWMYRKRMTTRCCYATYTTGCWARKRMRKYRKYMS
45          WRWRKGKMRWKWRCYYYCGGYKYGYKWMYGKGMYYRTYYRYYYYKMWCAGRMRKYGGCMS
            *****  ******.***    .   **** * *  *  *  *    *.  *****   **
```

```
48                  RRYASCWMKGKCGMKGCTRARMRGCCGKMKKYRMWKAWTCTRRMRACMMGGRKKRWATYK
45                  RRYGGYTCKRKTRCTTMAGTRMRATYKKMGKYRMWKMWYYKGAAATAAATRRKKRWRYYK
                    ***..   * *   .  : :***.   ** ****** *  .    :.    *****  **

48                  RRKSYYGCGSMKMRRYRMCTCRYRRYYRKYWKKWRKYYYKKYKKKGYKRYYCRTTYMMYK
45                  AGKSYYTTKSAKMRACACYYMRYRRYYRKYWKKWRGYYYKKYKGKTCKRYTYRKYYMMYK
                      ****   * ***       ************** ******* *  ***  *. *****

48                  RRKYKTCKMKYGYRKCYGGCARKMKRKKKWYYMSYKKRYSKAAMYRMRKKACTAKYRRRK
45                  RRKYKKTTMKCTYRKTCTRTWRKMKRKTKWYYMSTKKRYSKRRMYRMRKKCACGGYRRRK
                    *****. .**  ***      ******.****** ******  *******.. . *****

48                  RRKGTRYWYKWYRYSYRKCACYYMYSKYRKYMMKRYGKRWKKRCCYWWSKYMAYYRWRYK
45                  RAGAGRYWYTAYACCYGGYWTYYMYSKYRKYMMKRCKKRWKKRMYYWWSKYMGYYRWRYK
                    *  . ****. *  .*     **************  ******  *******.*******

48                  YMYYKKKACGGAMWKTYYKKTCYMGRRYGRRRRYYRKRKKYAGMRYYCKYTYKYRWKKAY
45                  TMYYKKKRTTCRMWKWCYKKCTYCKRRYKRRRRYYAGGKKCGRMRYYMKYGYKYRAKKMY
                     ******     ***  ***  *  *** ******   ** . **** ** **** ** *

48                  WRARSRRYYRYGYAKKKKKKRYYYMMWRAYRMYMMCWSGRYYRWKRAYWKYRRMCGYMGY
45                  WRGRSRRTTACRTGKKKKKKRYYYMMWRCYRMYMMAWSKRYYRWKRGCWKYRRMMRYMKY
                    **.****      .**************.******.** *******. ******  ** *

48                  GYKYKKGTGYWGRKRKGTCAGKMCTGYGGCAARKRKKGKCMTRKYYYMMSATCTWKGKTA
45                  KYKYKKTCAYWAAKRKRGMGKKMYWRYKATGTRKRKKRKMMCAKYCCCCSGGTWWKKKYW
                     *****  .**. ***   . **   * . .:***** * *  **    *.   ** *

48                  YRMRKKKWGKYRYKRGTKRGKCWWYKKKRYKRYYSKYYYMKATCKRAKSKKCKAATYWMY
45                  YRMRKKGWKKYRYKRRYKRKKYWWYKKKRYKAYYSKYYYMKWCAGRWKSKKYKRRWYWMY
                    ****** * ******  ** * ********* *********  . * **** *   ****

48                  SKRKCAGRYYCGGCAGCCRKYSRRRWKCGYGKKKYWWMRKRKYKRKRRRYKYWMRMWYRK
45                  GKRKMGTRYYASRYCRYTGKYSAGGAKTTCRKKKYWACGKRKYKRKRRRYKYWMRMWYRK
                    .*** . ***..  .    ***    *    *****   *********************

48                  RTTARCYMRYYMWKGTCKRYWRKMYRKRKKGYMTAYMYYKYTYMTGYKGKTTKRAGKYYA
45                  RCAGGTYMRYTCAKRKTKRYWRKMYRKRKKKYMYRCCCCKYGYMCTYKKKKKKGRKGYYW
                    * :.  ****   * . ************* **      ** **  ** *..*    **

48                  KRKWTYGKGAKYYYRRRMMCTRTTGKWAMRTMAKWYKKKYKMWKYRKRRKKTWKRTCGAK
45                  KRKWYYKTKWGYYYRRRMCYYRKYRKWTMGYCMKWYKKKYKMWKCGKRRKKKTGAYYRRK
                    **** * .   *******   *.  **:*    ***********  *****.       *

48                  RYYCKYRRMKYACCCMKKKMYRRYKWTYKKMYWKTKAATRYRWMWKKYWYTWGWTCTYWW
45                  RYYYKYRRMKCRYYYMKKKCYRRYKWKYKKMYWKKKTRCRYRWMWTKYWYKWRTWAKCWW
                    *** ******     **** ******.*******.*:  ******.****.*   .. **

48                  RCGYGRCRMSKRRYKYKKMYYYRRMKKSRRKSRKRKRAYCKACTCGCKTWAYYRTCGYGA
45                  RMKYRRYRMSKRRYKYKKMYYYRRMKKSRRKSRKGKGRYMKGYGTATGYWCYYRYYKCAW
                    *  * * *************************** *  * *.   .   *.***    .
```

```
48          CYRRYRCTKTKRWATYKYKKWRYAACTAWYRSYTACCTYMWKKKAARMKRARRTCGCCCC
45          YYRRYGAWKYKRWGCYKYKKWRYRRYYRWYRSYKRMTCYMWKKKGRGMKRWARGYRYGSY
             **** . * ***. ********     *****.    ******.  ***  *     .

48          AKYTTCTKYKRKRMKRYATRKRWRSRYTMWYTRAKKYAAMMCKMRAYRGMMKKGCMMSTT
45          WKYCATCKYKATGCKRTRARKRWRSRYKAWTAAGKKYTRMMYKMRCTARCCTGRMCACKY
             ** :  *** .  **  :********. * : .***: ** ***.     .     ..

48          TTTTGCWKMRMRWKWAGYKACYKRCRYRYATRCKCRMSKRSRTGYGAYGASKWWWRWWKY
45          AAAYATWKMRMRWKWGAYTWYYKGTRYRYWKRTKYRMGKRSRYSYTMCTRSKWWARWWKY
            ::: . *********..*.  **  **** .* * **.**** .*     **** *****

48          TACGKYRYKRCKCTSTKGWKMKMYCACCGTRKSCTARYRRRGAAYRYSRWAKKWGYWMMM
45          YMMRKYRYTRYKYYSKKAAKMKMYMRSYKWRKSYYMAYRRRKRWYRYSRWRKKWKYWCMM
                ****.* *  *.*.*****  .   ***    ****   ****** *** ** **

48          RRMRMAMKGKYYWRGKGTTRKKTKAGCCCRMYRRAAATMCKRRWYYKYAARRGAGKYRRK
45          RRMRMGCKTKYYWRRKKWYGGKKKGRYSYRMYRRGGGWCYKRRWYYKCGGGGRGKKYRRK
            *****. * ***** *     *.*.  . *****...   ******* ..   . *****

48          KYYYYKYAGTRRRWMKKMGYGCRKGRMWRTYRRRKYRRYKWAACGCRGYGAYGRKKKKCA
45          KYYCTKCMRKRRRWMKKCRCRGGKKRMWRKYRRRKYRRYKWMMMKMARYRWTRGKGKKAC
            ***  *   .*******      * ****.***********       *     * **..

48          TYRWKRYMRYYYCRCGAAKRRRYRCACARRRRYYKKMRKKTKMMYKRRWKMYKYYYACCA
45          AYRWKRYMRYYYAGAAGRKRRRYGYMYMRRRRYYKKMRKKCKMMYKRRWKATKYCCWYYG
            :***********. ... *****     ************ *********  **     .

48          GTWMCWKRSMKYYYRMATKRRMRWTCCWRYKGKKYRCKKWKWWSMMRKMMKRRKRYWYYR
45          RAWAYWKRCAKYYYRMRYKRRMRWCTTWRYKKKKYRYKKWKWWSMMRKAMKRRKRYWYYR
             :*  ***. ******  ******   **** **** *********** ***********

48          KKTTTKYKKACKYWMYWKKWRMRYKKTTACKRCYKYKRTTKMWMYRRGKKWWTCAATSWK
45          KKWYYKCKKGGKYTMYWKKARMRYKKWYRYKRYTKYTRGGGMACCRAKKKWWGYTRYSWK
            **   * **. ** ***** ******    **  **.*   *   *  ****  :  ***

48          AKWRMGCYGMCCYCRGTKYCYYYKTCGYWGGGGYYRARGRTACMRMCCTCTATYYKGYKW
45          GKWRARMYKMYYYYRRYKYAYCYKKYRYWAAKTCCGRGRRAGAMGMAMKTCCCYYKAYKW
            .***   * *  * *  **.* **.  **..        *:..* *. .  . ***.***

48          GTSYKAMYKRKKYRYYYRRKKYRYCCSRRGCATYRCCMMRWRKGGACACATGMGKARMSY
45          KYSYKMMYKRKKYRYYYRRKKYRYYTSRRKYMWCGTGMCRAGKRRGTTTCGKMAGRRMSY
              *** ******************  ***        * *  *  . : .  *.  ****

48          WYKKCTAKYSYWKSWKRGCRGRRYAAGKYKRKKYWYKWRCWKRKAAGYRKGYYMMKKGTA
45          WYKKYKGKYSYWKSWKRRTGRRRTRWRKCKRKKYWYKWRYWKRKGCKYRKKYYMMKKAGR
            **** ..**********    **    * ********** ****.. *** ******.

48          YRKRCRYYRYRYYYRYMWKGKRYKGCCKKAAYRYYTRSRKKRMKKACGKKCTAKKYYSKM
45          TGKRYRYYGTRYTYRYCTTRKRYKKSMGKWRYRYYARSRKKRMKKRYSKKYYRGTYYSTA
              ** ***  ** ***  . **** .  *  ****:*********  .**    .***.

48          KCCCKKKWRRMYKKYYKKYYRCGRKYRWGKKWGRKRWYSYYAKKAKKKYKTCYKRGKKRK
```

```
45              KTATKKKWRRMYKKCTGGYTGATGKYRWKKTTARKRWTGTCRGKMKKKYKYYYKGRTKGK
                * . **********    *  .  **** *. .**** .    * *****  **  .* *

48              RWKKTAARKKKACCARCCCCTYRMRRWWAYWMRARKRWMSKCKKKMYYKKRYSYYYGGTT
45              RWKKWMRRKKKMMYWRYTTYKYGMRRWWGYWCATRKRWMSKYKKKMYYKKRYSYYYKACC
                ****   ****    *    .* *****.**  :******* ************** .

48              TYYYKTGWMYRYWKYTSMGKKKKKKYKYRRYWRMMYKKYKKKYRGKYRYMMYKKWRMYKT
45              YYYYKGAACCGYWKCGGCKKKKKKTYKYRRYWRMMYTKYKKKYRRKYRYCCYKKWRMYGA
                 **** .    ***  .  *****.***********.******* ****  ******* :

48              TGKKKKWMYKARSKYYKTYRYRRMWYMYRTTRGKRYGWYTGTTAWKMYTGCYKKMYCCGC
45              GTKKKKWMYKTRSKYYKWYRYRRMWYMYRYKRRKRYSTTATAATWKCTYRYYKKMYSYRY
                  ********:****** *********** .* ***.  : :::**     *****.

48              KKYKMRRKKYYRMGYGRYWGYKSMKKMYMYKTGKSRRCGCARKMYMYRKAYKRYYRKKRM
45              KKYKMRRTKYYRMKCARYTACKSMKKMYMTGCRKSRRAATGRKMYMYAKGYKRYYRKKRM
                *******.*****  .** . ********    ****.. .****** *.**********

48              RTYKKTRKRKGGRGSKGRCKYYRKRAWRYMMMKKTRKTAGCYRYYKCTMKYYYRRWRKWY
45              RYYKKYRKRKRRRASKKRTTYYRKRMWRYMMMKKKRKYRRYYRYCGTAMKYYYRRWRKWY
                * *** ****  *.** * .***** ********.**    ***   :************

48              YYYYRWCAYCWYRRKCRYYSWRYYKKMYKRWKCATATWRRWKYRMRKRYSKMYRWAACKM
45              YYCTGAAGCTWTRRKSGYYSTRYTGKMYKGWKATCGATGGWKYRMRGGYSKCYRWRRYKM
                **    ..  * ***. *** **  **** **.: .:   ******  *** ***   **

48              WSTAWKKKGKKYKCRYWWKYMYRWWYRKAGKCGYWRYKMKTTCWYMCYTTWYYRRMKRYM
45              AGCGWKKKRGKYKMRYWWKYMYRWWYRKMKKMCYWRYKCKWWYWYMYYYKWYYRRMKRYM
                 . .****  *** **************  *  ***** *   *** * .**********

48              CCCWKMKKKWTTYMARKGTTAMRSMWYRYYYKMAYYCAKRMAYWKRRRKKKMWYWTGKRY
45              GMTWKCKKKWYYYARRKRKWRMRGMWTAYCYKMMYYYRKRMTYWKRRRKKKMWYWYRKRY
                   ** ****  *  **  .  **.**  * *** **  ***:*************  ***

48              KYRYRAWYYMYRRWYYMYKYKYRYYKYYKRWGYCRYKKWRRYTTRKKCRKRGCCAKYRKR
45              KYRYRRWCTMYRRWCCATKYKCRYYKYYKRWRYTGYKGWRRYGKAKKYRKGKTAWKYRKG
                ***** *  *****    *** ********* *  ** **** . ** **   . ****

48              MAYGGM
45              CGTATM
                 . . *
```

The pairwise alignments of two accessions 45 and 49

```
45              AYYWYRWKYYYWRKYRGCTRYYMKWWRYRYKGGMKCKWMYGYKYYYRYGYYTAMWKRWGY
49              TYYWYRWKYYYWRKYRTTGRYYMTWWRTATKRKMKTKTMYRYKYYYRYSYYCRMWKRWSY
                :***************   ****.***   *  ** * ** *******.**  *****.*

45              RYRKYMYYWCMYKWRYWRYKMRMYWKKWMMYYRWMYKRRMYYWYKWYTRATMYSRRRRKA
49              RYGGYMYYWTMYKWRYWRYKMRMYWKKWMMYYRWMYKRRMYYWYKWCKGWWCTGRGRRKR
                **  ***** ************************************ .     .* ***

45              MYCCYCYWWCRSSYCTAWCYKCYWTTGMWWRMMYRKWYRRKKMKATCTYRRRWWRYRYRR
49              MYMMYYYWWTRSSYTWTAAYTTTTCYAATARMMYRGWYRRKKMKGGYYYRRRWWRYRYRR
                **  * *** ****  : .*.     .   ***** ********.   ************

45              RGYKYYRMGYRRYCATTRYKWGGKKTAGYAYMYRWYYKGTRRTKKWGAYWYMMRWKRRWY
49              RRYKYYRMAYRGYTRGWGYKTKKKKWMACGYMYRWYYKTCRRCGGTSWYWYMMRWGRRWY
                * ******.** *     **   **  . .********  **    . ******* ****

45              WRYMYMYYYKYYYYKYYSYRKWYRMRYWRWSWMKMCYWKKRMYKMKYWMRRYYYKGYGCG
49              WRCMYMYYYKYYYYKYYSYATAYRMRYWRWSWMKMYCATKRMYKMKYWMRRYYYKTTCTA
                ** **************** . *************   .****************    .

45              KYTRKWWCTKCRMKCAKSTAYGAYMWYRMYKRRWGYRYYYRKYRRYMMRRATYYYKTYKK
49              KYKRKWWYCKTRMKYRKSCCTTGTAWYRMYKRRWRCRYYYRKYRRYMMRRTCTCYKKTKK
                **.****  * ***  ** .  .  *********  **************:   **. **

45              KYGCTGMKKYMRYYTYRYKMSYKSACYRWYRRSMTRKYRCYCCKWGGKMYYWKACSKCAC
49              KYAMWKMKKYMRYYYYRYKMSYKSMYYRWYGASMARKYAYYYYGARKKMYYWKRYSKYGT
                **.   ******** *********  ****  **:***  *      ******  ** .

45              AYTWMWKYTRGAARRKKKCTWRRKMSAWCRYRYMKWYMKWYAAAKRYYRRYCTWRRYSRY
49              CCCWCWKCCRAGRRRGKKYYWRRKMSRTYRCRYMKWYMKWYRWRKAYYRRCTWWRRYSRY
                .  * **  *.. ** **  ******   * **********   * ****   *******

45              AGGCATCGKKCAGRTMWRKMCAAKRYWWYATGGMCGWCCCTTYRRRMKKSWGTYYRGCAT
49              RRRTCKTTKGYGKGYMWRKMTTRTRTWWYGAATAMATMGTYCTARRMKKSWRKYYRAMRY
                    ..  *  .   ***** : .* ***.:.   .        ******* .***.

45              CAGCAAAAAKAKTCTTRRMGYRWYTYCRAATGACSKAAKKYRYKKSRCTRMYKKCKWRRC
49              TRRYGMGGMTCKCTCARRMKCGAYCYYRRGYTGTSKCWKKYRYKTGGYARMCTGYKWGGM
                    . .. ..*   :***    * * * .  . **. ******..  :** .  **

45              CYRKWMYATACGWCGGRKKRAWYKCRKCATMWMAMMWMYCCKKRYYGAYRWYRKCAAAGT
49              MYRKWACGCGMRTYACRKKRRWYKTRKAGKMWMGMMWMYYMKKRYYSGYRWYRKMMMRKK
                 ****  . .    . **** *** **...***.*****  *****..******     .

45              WYYKWRYYRRKRMWWWWMKKCYMRRKKCTYCWYWSTWTAKKCGRKYGGTYRYKRGMGKRC
49              TYYGTGYYAGGGMAWWTCKKSYMRRKGYWTYWYWSAACCKKAAGKYATCYRYKGKMRKGT
                 **   **    * **  **.*****     ****:  .**.. **.  ****  * *

45              AKATRWAGKTAACRMYKWYSYCGAMYRTCKTTAMKSRMKWKMKKKKRTRKWKRRSWYMRW
49              TKMYRWRKKKTTTRMYGWYSYAATMYRKTKWYRMKSRMKWKMKKKKRKGTTKRRSWYMRW
```

```
        :*  **  *.:: *** ****..:***. *   **************. . *********

45      RYWAAMRYTYWTMRYAGYYYMMTTYYGGTAMYRKGYRSCGRKGKKKKSYYCKGYRKMWCT
49      RYWRRMRCKYWYMAYWRCCYCCYYYYAAATMYRKATACYRRKKGKKKSYTYKKYRKMWSW
        ***  ** .** * *    *    **..::****.  .  **  *****  * *****.

45      YAWTKMWWCYTRKYYCTGGAKACGTWYYKYSYYYYYTKAAKTWGKYKMRKYSKYYYYRRK
49      YRACGMWWYTWAGCTYGRRWKTMRCWYYKYSYYYCYYKGGKWAKKYKMRKYCKYYYYRRG
        *    ***            *:   ********* * *..*   *******.*******

45      MAGTGKKTYMYWTYAATRMWRMSRRKKKKKRKKWSGGTYRYKKKKRRGKWMRRSRYKKYK
49      MTTYRTKACMYTYYMMYRMWGCSRRKKKKKRKKWSAKWYRYKKKKRGSKWMRRSRYKKYK
        *:   .*: **  *   ***  *************.  ******** .************

45      KRYMRKKYMGMKKTTCCKKYRYKCKYMCGGKKYRGKAKMRYYRAWCYMKTKYKKWYRKYG
49      GRYMRKKYCRCGKKKYYKGCGCGMKYCYAKKKCGSGWKMATTARWYYMKKKYKKWYRKTK
         *******    *..  *      **  . **  .  **     * ***.********

45      RWYMKMATGWRGTTRKKGGYTGAGGTWYWRWKKMYKASYWGTKRKKWKKWWAGRYRRKMM
49      GWYMKMWCTWRSKWRKKRKYYKRRTAAYARWKKATKRSYWCKKRKKWKKWWWRRCRRKMM
         *****   **.. ***  *     : * ****  * *** .*********  * *****

45      RMACCGAKRYTKGGGRYYRRWYKKATRRAMYKKRYACYYKTRRCRRWYCRYYYKKKKWRY
49      RATTTTGGRYYTRKTACYRRWTKKTCRRMMCKKRYGGYYKCGGYARAYAGYYYKKTGWRY
        * :   . ** .     **** **: ** * ****. ***     * *. *****. ***

45      YRWGYRRYRMRGSKWAGGWYKRWYKKATYTAKKKCYCKKAAKATTWKYKKYMYRTWKYRY
49      YRTKYRRYRMRRSKWCAAAYKRWYKKRGYGCKKKTYYKKCCKRYWAKYKTYMYRKAKYRY
        **  ******* ***... *******  * .*** * **..*    ***.****. ****

45      MYCWRKKKKGRRGMYKYWSSSKKWRRYGWYSWGMYRYMWYWRYYYRMYYRRYKMWMKKYK
49      AYGAAKGKKRRRKACKYWSSSKKWRRYAWYSWTMCRYAWYWRYYCRMYYRRYKMWMKGYK
         *   * ** **   ************.**** * ** ****** ************ **

45      MMYCTCYRWAGGKCAWKRYYYKCTAYYRKYWKMKRRSMSMYKATCMMYCYKYSRATYRKK
49      MCYYCYYRAGAAKYGAKRYYYKMKWYYRKYWKMKRRCCSMCKTATMCCMTKCSRRYCRKK
        * *   ** ...* . ****** . ***********. ** *:: *    * **   ***

45      AYWGCTRRAKCKWKRKRCTWWYRYARMMYKMYYYRGRKYGYTAKTYKRYGAAYMWYRRYK
49      RCAKTAARWTTKWKAKRTGWWYRYCRCMYKMYYYRKRGCATATKYCGRYRWMYMACRRTK
             : * . *** **  *****.* ******** *  . ::*   **   **  ** *

45      RYRWYYRKYYWMKMCGATTKSACTKRKYATGKYKYYSKYKTTKKKKMYKRSGYYGGGRKY
49      RYGAYYGKYYWMKAYKTGCKSMMYTRKCRYRKCKYYSKYKCCKKKKMYKRSRTCATAGKY
        **  ** ******   :  **   .**    * *******  *********   . . **

45      MYSRMKAMMKARRSKYRYTCYKARWMKTCTYWKRYKSTCCKYACCRRKKKGKKKRYYYYM
49      MYSRMKRMMKCRRCTYRYKYCTRRWMKCMAYWKRYKGCAGKYCTTGGGKKTKKKRYYYYM
        ****** ***.**..***.  . ****  :******. . **.     ** *********

45      MSYCCTKKMAYRRKKYYKWKKMRMRYYKRRRRTWRAARRKCRRRMGCRSGKCGWKWYSRG
49      MSYTTAGKMRYRRKKYYKWKKMRMRYYKRRRRGWATGRGKYAGGCKYRSRKYAWKWYSRK
        ***  : ** ********************** * :.* *       ** * .******
```

```
45      RCKMMKMKKYATATCKAYWKCYKYYKRRWTMKYKGSRWRAKRKYKSRMSYKKKRMKKKYR
49      GYKMMKMKKYMCCATKWTTKYYKYYKRRWCMKYKRSRWRGKRKYKSRMSYKTKRMTKKYG
          ********  .: *   * ******** **** ****.***********.***.***

45      KKRRRKKCTGKTRWYRAGMYKSMTKKYKYKKYKKKRRYKKATKMWWSKYGTYWGYSYKKR
49      KKRRRKKAATKKRWTRMRMYKSAKKKYKYKKYKKKRAYGKRYKMWWSKYACYTSYSYKKR
        *******.: *.** *  **** .************ * *  *******. * .******

45      AYKKWKYGCKTRTKKTKKRRRWKKKKKTMRGYWYCMWYMKKKKTATKAGYRYYKKYKKKW
49      GYKKWKYAYKCRCKKAKKARRWKKKKKWMRRYWYTCWYMKKKKYMYGMKYRYYKKYKGKW
        .******. * * **:** ******** ** ***  *******      ******** **

45      KKMWKCRTCKAGGWKAWMKSYKWAACTTGKKYRYKRKKKRCYCKKRRYTYKKWSKRYWYR
49      KKMWKYRWYTRRSWKGWMKSYKWGCAAATKKYRYKRKKKRSYAKKRRYYYKKWSKRYTYR
        ***** *  .  .**.*******...:: ***********.*.***** ******** **

45      RKKMKKRWYKAKGRRYRRSRRYMRYGCGGYRATCKMTTYKKAKSYWYTTAKGYRWYMKTG
49      RKKMKKRWTKTGRRRYRASRGCCRYCTRRYRCCTKMCCYKKMKSYWYKAGTTYRWTAKYR
        ******** *:  **** **   **    **.  **  *** *****.:.. ***  *

45      AKSKRKKAKATACKYKRTYKRRRKKKRKTGCKCGYWWRRATTAGRKKYKKKRRMYRRKRY
49      MKSKRKKMKMWGMKYKRYCTRRRKKKRKYRYKTTYWWRRWGYRTAKKYKKKRRMYRRKRY
         ****** *  . ****  .********   *  *****      ***************

45      CYMYYTCGCWSKRAKMRYYRRKYYRRKCCRRKGMKAATYYSKGWRKWYYKYYYGYRYKMR
49      YYMCYCTATWSKRCKMRYCRRKYYRRKYMARKAMKGGGYYSKRWRKWYYKCCCSYRYKMR
         ** *  . ****.**** ********   **.**.. **** *******   .******

45      YWWKKCRCKKKMKRMATARRYMRYYMYTARWCYCCAYKKKKKSKYRKKRYKYCGWARGGT
49      YWWKKYAYKKKMKRMCGMRGYMRYYCCYCRTYTGATYKKKKKSKYRKKRYKYMAWCAAAC
        *****   *******.  * *****   .*    .:**************** .*. ..

45      AAAAYRYMYKYYWSMRKKYRKWWTMMYKKMCYYTSYRGAATKKAAAGRTKKGCKKRKRYY
49      RGTTCATMYKYCWSMRKKTRKWWYMMYKKMYYYCSYRRGWYKKCTGAGAKKKYKKRKRYY
         .::   **** ****** **** ****** ** ***  .  **.:.. :**  *******

45      RYYWMYRMWMMWWYYKRMWMMYWWKKRYRCGGRMTTAKYKAGYMKKKATWKGRRMYRRRR
49      RYYWMCACWMATTCYKRMWCACTWKKGYRGKRRMYYRKYKGCYMKKKMKWKARRMYRRRR
        *****   **    *****    *** **   **   ***. ***** .**.********

45      KSCRYGKYYRRKYMMWMYKKCRWYWWKSSGRRYYYWMMRKYYRCMRYYMKKKGRKTKACG
49      KSMGTRGYYRRKYMMTMYKKYRTCTTTCSKRRYYYWCCGKYYRYMGYCMGGGRRKWKWYK
        **     ******** **** *    ..* ******   **** * * *    ** *

45      AMRRSKCAKKTKYTKSMRYRRGYRYYSKRTCCWYTMKGCMSYYGKGTRGTCWGWYMKKYR
49      GMAGGKYTKKCKTAKSMRYRRKYRYYSKGYTYAYCAKTAMSYYAKRKACWTTAWYMKGYR
        .*  .* :** * :******* ******     *  *  .****.* .     .**** **

45      KYKTGTGCKKRYCGRKTTRWMMYWMTAKACTYYYRKYYYYMYTCYMKYGKYKKKMKKGYT
49      KYKGAARTKTRYYRRKWWRWMMYAMCWKGTAYYYRKYYYYMYATTMKYRKTKKKMKKACA
        *** .:  *.** **  ***** *  *. :***********:  *** * ******. :
```

```
45    YKRTYTTATYYWYKKWKYWRTWKATTTKKYKRKKKRYKKRKKKTCTGAGCRYRYKKRMKY
49    CKRGYACTCYYWTKKWKYTGKTTRYYKKKYKRKGKRYKTAKKKWYWTWTTRYRYKTRMKY
       ** *: : *** *****  . .   .****** ****. ***       *****.****

45    KRYMKAKYWRRAAWKRWRYRMKWYMWKRKYKWGTKRTYRWSRRKRKAAMMRRMACYRRKK
49    GRYMTGKYWRRGTWKRWRYRMKWYATGRKYKAKAKRKYGAGRRKRKRRACRRMMTYRRKK
       ***..*****.:***********   ****  :**.*  .*****    ***  *****

45    KKCRRMMRRKKKMYKGKYWRMRKMRMKYKMTTATKWYWWKRRSMWMTTTTCMYTRKRKCT
49    KKAGRMMRRKKKMTKRTCTRMRKMRMKYKACATGKWYWWKRRSMWMCCCGYMTARKRKYY
      **. ********* *  .  **********  :: ************     * :****

45    TKMKRKMRYRRTMKMRKYKMTATGWCMTMKRRYYKRKMTWYRRKKKCYYTMCYCGKCTMK
49    WKMKAKMRYGGAMKMRKYKMWMYRWYMAMKARYYKRKMCWYRRGGKYTCCMAYYKKTYMK
       *** ****  :********    * *:** ******* ****  *    *.*  *  **

45    KRWMRKAAYAWTRGGGGRAYYGMCYRSRKKTGKRWSYMTWTYWRRRRRKYKKYRATMRRC
49    KRWMRKRRYGWCGRKAARMYYRCYYRSGKKYRKRWSYMGACYWRGRRRKYKKYRMKMRRY
      ******  *.*    ..* **   *** **  ******   *** ********* .***

45    CKKKWYMKKKCKSKTTMWWKAAGRRMYYRYKKGTATYKYRKKGKKRKYYMSKMMYYYRYW
49    YKKKWTCKKKMKSKKYMTAGMGKRRMYYRYKKKAGCYKYRKKRTKRKYYMSKMMYYYRYW
       ****  *** ***. *    . ********* :. ****** .****************

45    RRKKKGYYWRKMYKYRMRYSRWWRRWYKRKMYYRMYYKKTAGWYGRATTTTAKWKMYRCW
49    RRKKKRYYWRGMYKYRMRYSRAWRRWYKRKMYYRMYTKKYRRWYAGGAGYWRKWKMYRYW
      ***** **** ********** ************** **   **. .:    ****** *

45    RAKMRKYATYATATTKAKKYKKYKCTGGGATYYKYKKKMAAWKKWYKWTGCYCKRKMKYR
49    RWKMGGYMCCRYRCCKGKKTKKCGYGCAACGYTKYKKKMWWWKKWYKWCTTYMKRKMKYR
      * **  *        *.** **     ... * ******  *******   * *******

45    YMKYKKKKKKSRWRRAYRKKTKYYAKYYYRRRYRYYWYCMMWAGWKKRYKRRRWRRSYWY
49    YAKYKKKKKKSRAARMYRKKCKYTRKYCYRRRYGCCWYTCCWRRWKKRYKRRRWRRSYWY
      * **********  * **** **  ** *****   **   *  ****************

45    MYKGYKKKRTTTTGAAAGTAYKKRYRRWRCAKYAKYYCARTMGAYAWKKTCGCRYYKCAA
49    MYKKYKKKGYAGAAMGGTATYKKRYRRWRYCKYGGCTTGGKMRRYCWKKYTSYGYYKYTG
      *** ****  : :. .. ::********* .**.    . .*  *.***  .  *** :.

45    RRYKKCCTYYWAYRSKWWKYSTRWYYWYGYKWKGGKWWKCRYKKKRCCAAKYYYMMMYKT
49    RRYKKYAGCYWMYRSTWWKYSKRWYYWYRYKWKRKKTTKYRYGKKRYYRRKYTYMCMYGW
      ***** .  ** ***.*****.****** ****  *  * ** ***    ** ** **

45    GKGCGTYKKMKWKKYYAKMKRRRRTYWKYGKYTCGMYWKGATCYRMMRKYCAGRRYKYYM
49    KKKTRKYKKAKWKKYYCKMKRRAAKYWKCRKYYATMYWKKRYAYRAAGKYTCTRRYKYYM
       *   .*** ******.*****  .***  ** . ****   .**   ** . *******

45    KKRKRKRKTKTGTSTYGKKKKMKYWYTKKWTGYCGAKWYSWRRTMRMRKCAYYCACGGCK
49    KKRGGGRKWKKTCCWCKKKGKMKYWYYKGTGRYMTCKWYGWRRYAAAGKYRYTTCTAAYG
      ***   ** *.  .   ** ****** *    *  .***.***     *  *  . ..

45    GYKYYYRRTARRCYYKYYRMRGKYWYRKASKKGKYMKGYYACYTGKYRKYRKKWTCRSMK
```

```
49          KYKYYYRRCGGRYYYKYYGMRRKYTYRKTSKKRKYMKKYYMYYATTYRKYRKKWAMRSMK
             ******* . * ***** ** ** ***:*** **** **  *: .********: ****

45          KKCGRYAAYRWAKWAYKWTWGWYRTGACYRWKYKCAAKYRYKRKSRKCCTGATKKCSMKK
49          GGTRRCMWYRWMKWRYKWGAATYGYAGAYRWKYKTTGKYRYKRKSRKYYYRWCKKMSMGK
                *   *** ** ***  . *  ...****** :.**********      ** ** *

45          YRYRWKMKKCGACGYYYSWTYRRKATAGSYSMGKSMRYYKRMGAKRTCCMYYRRRCKAYT
49          CRYRWKMKKYKRYKYYYCTACRRKCCTKSYSMRKSMRYYKRMKMKRKGMCYYRRGYGGTY
             ********     ***. : ***. : **** *********  **.   ****   .

45          KGRYYYYTYATARSRKYCGSGCTGTTCYWKKYRTTGTKTWGTYCKYRYRWRTMYKYMGTW
49          KKRYYYYKYCCTRSRKYYKCATAKYWSYWKKYGWATAKGWTWYMGYRYRWGKMYKYMKCW
            * *****.*. :*****  .. :   .*****  : :* *  *  ***** .*****  *

45          YRWTMWKYMYKAWYRWYTKRRWMMRTGMRYSCCTRGGAGKMMGGYKYRYKKYYKKRCRMK
49          YRWYAWKYMTKCWYRWYGKRRWMMRKKMATSYTAGRAMKKACAAYKCRYKKYYKKGYGCK
            ***  **** *.***** *******. *  *  :  .  *  ..** ********    *

45          KKYYSRKMKYYKRTKYMKRKACMTTWTRYKKKRKKYRRRKKYSRAAGCMRKATCAKKKSW
49          KKYYSRKCKYTKAYKYMKRKRAMYCTAAYKKKRKTYRRRKKYSAGGTAMRKWWTGKKKSW
            ******* ** *  ****** .*   : ******.******** .. .***   .*****

45          KWGAKWKCRYSMMKRKKKKRKKRAYSYYKTAKKKTAYRGKTRRMYYKMKCGYKGCTWYRK
49          TWTCKWKMRYSMMKRKKKKRKKRGYSYYKCCKKKYRYRKTCRGCYYKMKTATGASYWYRK
            .* .*** ***************.***** .***  ** . *  ***** .  .. ****

45          SGRYAGGWGTYRTAGCTAYRRYACCKRYKRKRCMYWCYASYTRRGRRRYGRCKMYYACMA
49          SAACTAKWKACAKWSAAGTRRYGMYKAYKRKRYCCAMCGSYYRRKRRGCKRYKMYTGTMM
            *.  :. * :  . ..:. ***.  * *****      .** ** **   * *** . *

45          TCAMTYCRYKYRGRTATGGTATRWRYYSWSK
49          CMMMWYTRYGYGRRAGGAAYRARWRYYSWSG
               * * ** *  *:. ..  :********
```

The pairwise alignments of two accessions 45 and 50

```
45                 YKRTTCYYKWCCRWWRMKRSSWTWWCYYACGTATMCYRWYYYYYYAWRMRKYCRKYRYRC
50                 CKRYYTYTKWMMRWARMKRSSWYWWYYYTATYWYCMYRWYYYYYYCWRARKYMRKYRYRG
                    **   * **  ** ******* ** **:.      *********.** *** ******

45                 TYWWGYWWYGWWYKMCGGWKWAKYYYYRMMRRYYMKMWWRYRMKRMYYRMMKYRRAMKRY
50                 AYWWKYWWTAWWYGCTTRTGWGKYYYYRMMRRYYMTMWWRTAMKRMYYRAAGCRRRMKRY
                   :*** *** .***       *.*************.****  *******    ** ****

45                 KKYSRYRKSWYWWRMMYGGYWRCKMYKYMYRWRCRWWWYRRRWKGAKKRYYKCGKMYWKK
50                 KKYSRYRKSWYWWRMMYKRTWRYKMYKYMYRWRYRWWWYRRRWKRRKKRYYKYTGMYWKK
                   *****************   ** ********** **********  ******   *****

45                 RWMMYYGWYRKRRTKKKWYWRYYYYYKKYYSYRKWYMWRWAGCCTRYYRRWGYTYTAYTG
50                 RWMMYYRACRTAACGTKTCTRYYYYYKTCTCTRTAYMWRWWTYYYRYYRRWRCGYCGYGT
                   ******   *.    .*   *******.  . *. *****     ******   * .*

45                 AYTACYKMKYGAGTWRRGSMMYKCKMKRYRRMGCTYRKMKYYRYRYYRYKGGTKCMYYKS
50                 GTCTTYKMKYAWRWWGASGMMYKTKMKRYGGMRTWCRGMKYYRYRCYGCKTKCKSMTYKS
                   .  : *****.   *  ..**** *****  *    * ******* *  *   *.* ***

45                 KRMKWAYWKSKKYYTRMMMRMTAKMYKKRYKYSYMYCWMYWRYKWYMRYYMYCAMRYWKR
50                 TACTWGTAGSKKYYGRMMMRMWMKMTTKRYGYSYMYMWMYWRYGWYMRYTCYAGMGTAKR
                   .  .*.   ***** ******  ** .*** ***** ****** *****  *..*   **

45                 WYMGGRMWSKYGWYYACTTAATTMMMCYYRSWTRKKYYYYKKYKWYMTYWYARMWRMYYT
50                 WYCRRRMWSKCRWYYGAGGTGAYMMCYYCAGWCRKKYYYYKKYKACMKYWYRRMWRCYTK
                   **   *****  ***..  :.: **  *  .* ***********  *.*** **** * .

45                 KTSRRKKKTYKKRRYYWSYRYMKTCTGCRMRRKRRSRYYKTGTCCKGTATKRYTRRMKKT
50                 TKSRRKKKGTKGRRCYWSYRYMGWYKSYGARRKARSRYYKGSAYMKRYMYKRYCRRMKKW
                   ..******  * ** *******   ..   *** ****** .:  *    *** *****

45                 TMMKKYRYRRKKRATKWKWKKWYRKYRMKYYYKKWRSKKYYKAGGAKKYRWWRKRKYKKS
50                 YCCGGCGCRRKKARGKWGWKKWYRKYRMKYYYKKWRSKKYYKCATTKTYRWTGTGKYGKS
                           ****   ** ************************.. :*.***  . ** **

45                 YWKKTRRRMRYYYKKKWYRKYKYYSKCTCAGGTTKYKYKKKCRTKTWTYKKWGTTYYRRR
50                 YWKKCRRRMRYYYKKKWYRKYKYYSKACTGRCCCTYKYTKKMACKYTWYKKWRYWYYRRR
                   **** *********************.  .    .***.**   *   ****   *****

45                 ACRWMRKKYYACYYKKGTAMMYRCWACAWMKMSRYMMKKKKATKKYRYKRYRKYMKTKYR
50                 RYRWMRKKYTTGTTGKRWCMMYRTWRYGWMKMSRYMMGTKKTCKKYRYKRYGKYMKCGYR
                     ******* :    *  .**** *  .********* .**: ******** ****  **

45                 ARAMRGCATAMWGGRYYYTMYRMRMYYWRYMYWARGAGKSYRMRRACCAKTACYACAGRR
50                 MRRMAKYCGMMAKRRYYCYMYRMRMYYWRYCCWWGTCAKSYRARRTTAGKYGTYRYRARG
                    * *   .  *   ***  ***********  *   ..**** **: ..* . *   .*

45                 WRKYRYGKYKTTTTCTAASKMRKYKKMCTRRYKCCCTWSRRKWRRYRGWWRYRKKKYYGR
50                 ARKCATRKYGCGYWMYTWCKMRKYKKMMYRRCGMMTYWSRRKWRRYRTWWRYAGKKYYAG
```

```
                 **    **       : .********  **      ********** ****  ****.

45               TRWRRRWSTAAMWRYKWYGAKRRRYRKKGWCRMAATMCKGKRYWRMWRMRKTRKMMRMKT
50               YRWRRRWSGTGMWRYKWTKTGARRYRKKATMRCWWYMMKRKRCAAMTRMRKAAGMMRMKA
                  ******* :.******  :  ******.  *    * * **   * ****:  *****:

45               TTGACAGTYRKYMYRRKRSSYWKKMAAKRRCMYKMKYRAMRWRSYKKYWSYRRKRKRGTY
50               YGRTYCAYYRKCMYRRKRSSTTGKMGCKRRTACKMKYRTCGTRSYKGCASYRRTRKGKWC
                    : .. *** ********   **..***   *****:   ****   ****.**

45               WYRYRMMYYYMYAKGMRYGYWRMRGGYTGGWRWKRYKKYKYWKMYYKKSYKMRTKSMKSK
50               TTACRMMYYYMTTGKMGCKYWRMRARTATAWRWKRYKGCKTTTMTCKKSYKMRAKSMKSK
                    ******* :  *   *****.  : .*******  *   .*  *******:******

45               KKYYCAKWGRRRYKYKKGYYKAAARRKWKYYRCYRMRGWSTCGTTKRWKGGRYWKYYYRM
50               KGYYAGGARRRRYKYKTACTGGRRRRKWKCYAYYRMRAWCCMKYWGRWKRRRYWKYYYRM
                 * **..   *******..   .  ***** *  ****.*.      ***  *********

45               KKAAWARYWWYGYYYKYMYKKGYCARAGYKKYWAGSKKWWWWYWTRRAATTGRGYSGMAY
50               KKCCWWRYWWCRTTYKYMYKKCCTTAGRCKKCWGRSKKWWWWYWKRRWWYARRAYSTMRT
                 **..* ****    *******    : .   ** *. *********.**    : *.** *

45               YWYWRYRYYYRRYMWMYRYMYYRKYMWMYKRWRKMTARRMGYGTAGCTTKYWWYYKMKKK
50               YWYWRYRYYYRRYMWMYRCMYYRGCMTMTKRWRKMCCRRMRYRWWTTYKKYWWYYKMKKG
                 ***************** ****   * * ****** .*** *       .**********

45               RRKWKRRSKWMRYWMRARYYYMGGRYAYCCYKRATKMAATGTGGKCGGACCKTSYYGCKM
50               RGTTKRASKWMRYWMRRRYYCCSRRYRYYYYTGCGKCGCACWRAKYARGTMKKCTYATKA
                 * . ** ********* ***   . ** *  *. . * ..:   .* .  .  *.. *. *

45               RRKTCKWAWGGTWATWWRKGACGTWMTYKYACGKYKARWKCGYMRYMRKMRRRKSKWKMW
50               GRKCTTWGWKTCWTCTWAGKWMRYWCCYKYGAAKYKGRWKARYMRYMRKMRRGKSKWKMW
                  **   .*.*   *:  *        *  ***...***.***. ********** *******

45               WKATGRKYMMKYTATCKTCGGTGYMWKYRYYYKTTRKRWYKRWRYAAMTCKKTAKATGKK
50               WKTAARKYMMKYYCATKWYTRWRTATKYRTTCKCYAKRWYKRWGYGWMAMKTACTRWKKK
                 **::.*******  .: *         ***   *   ******* *. *: *.:..   **

45               TKRRRYYKMYRMKYKGKMYTKYKRRKWYRTGKYGYRKATTRGAGAGRYRRMYCYWKTKKK
50               WKRRGTTGMYRMKYKKKMYYKYKRRKWYRGTGCAYRKTGARKGAGARCRRMYTYWKCGKK
                  ***    ******* *** *********     .***: :* ....* **** ***  **

45               YYKRRKKRRWRKYTGRYKRMRWMMYAGGYYTMYYYMKRYWKGYKYKKRYYSRRRYYYRMM
50               YYKRRTGRRWRKYKKGTKRMRWMMYRRRYYCMYTCMKRYWKCYKYKKRYYSRRACTCRCA
                 *****. ******.   ********   ** **  ****** ***********    *

45               YYYKKWYRTRWYYWAYCYKWYYYKKKTMKRYKKCCAAKKGYGMMWMKKRGKKKKYYKYMR
50               TCCKKWYRKAAYYWRYYYKWYYYKKGCMKRYGKYYRRKKSYRCMWMKKGRKKGKYYKYMR
                    *****.  *** * ********  **** *    **.*  *****  ** *******

45               TYMYWRYRRYKWKWWYRKKRKKKKKMMYRWKKMYYKKKTYWGKYCYRMKKYRYSWWYYKY
50               CYMYWRYRRYGWKATTRGTRKKKKGMCTAAKKMYYGKKGYATKYATRMKKYRYSWWYYKY
                  ********* **   * .***** *    ***** ** *  **. *************
```

```
45    RYYACKKYRMGRGTKRKKGTACKWTTMKMAMMYTGYRKYRRWKYKWKKKWRKWYMWMYWK
50    GYCGTKKYGMRRAYGRKKRWRYKTCKAKMCCMYATYRKYRRWKYKWGKKTRTAYCAACWK
       * . *** * *.  ***    *  . **. **: *********** ** *. *    **

45    KRSRYCKKRKGAKYYRYTRKMYKKKCCCYKRWKMKKAKKRKMRYYYKAYKAKSKGCGRKR
50    KRSRYTKKRKCWKYYRYGRKMYKKKATAYKRWKMKKWKKRKMGYYYKWCKWKSTKYRRKR
      ***** ****  ***** *******. .******** ***** ****  * **.   ***

45    YKKYYKKRKMMYRRTWYTKTGKKTTCTWCGMAGTTYRCGAGTMRMYKSTYARCTGRARSY
50    YKKYYKKRKMCYGAGWYKKYRTTYCSYWTAMRRKKCRGRGSYMRMYKCCYGGYCRGGRSY
      ********** *   **.*  ..  . * .*  .. *  .. *****. *.     .***

45    TKRRYTATAWGTTGCGCAKYCAATRGYYRWRCTACWTGTWRMYAYRYYKKWYRKMYYYWK
50    ATRATYRWGAKCAAAKYGKCMMCKGSTYRWRMYRYWYRYWRMTMYAYYKKWYRKMCTYWK
      :.*     .   :..  .*   .. . ****    *   ***  * *********  ***

45    YTACYCAGKYWMRTGGYYWYYAKKKKYRKRRRYKKYKWKKKYRRRAKKKKWACCCKKKMC
50    CAGYYYGAKYWCRYAACCWCYCKGKKYRKRGRYKKYKWKKKYRRRWKKKKTGTTTKKKMS
       :. * ..*** * ..  * *.* ****** ************** **** .   ****.

45    RAKYRTAMKKAYYKRKKKAKKKRRKKYCKCARKYMKKKKKRRMKYWRWRKKKWRYKWYWT
50    RRKYRACMKKCYYKRKKKWKKTARKKYMTTCRTTMGKKKKRRMKYWRWRKKGWRYKWYAK
      * ***:.***.******* **. **** . .*. * *************** ****** .

45    TGTMKARRRRRMYYRRKKKTTGAKKKRWKYRYRYRRKWYWYKKKKRRSMKWMRKRKKMRY
50    CCGMKRRRRRRMYYRRKKKGCCRKKKRAKCRCGTRRKTTTCKKKGRRSMKWMRKRKKMRY
         ** *************    **** * *   ***    *** ***************

45    KMMRKYKCMMYYMYCYRYKMAGMYWRYRTYYMYRMRKAGKKRKAAYTKTYRRRYWWWTGC
50    KMMRKYKGMMCCCTMYRYKMCTMTARCGCCYMYRMRGGAKKRKTWYAKKYRRRYWWWYRY
      ******* **     *****. *  *    ****** ..****: *:*.********

45    KRYAKRSYKMWRRMYSRWRYRKKCARCYSKRYRRKMKYAMWRTKKKWMRYYKYYMRRYKA
50    KRYRKRSYKMWRRMYSRWRYRGGYRGTYSKRYRRKMGCTMWRWKKKAMRYYKYYCRRYKR
      *** *****************      *********  :*** *** ******* ****

45    GGCAATARKCTAYTCCTYYSKKACYMMAASTKMMRMYYRYKKYYCKMKKAKYRKYMKMMY
50    KRTTTCGRKYYTTCTAGCTSKKGTTCCGCCWKMMRMYYRYKGYTYKCKKRKYRKYMKMMY
         :: .**  :   .   ***.    ... ********** *  * ** **********

45    GYWATWRRKYRKMKYSRCYCKYGGMYKYWWYSYKCRGWYRGKRKKMRKKAGKTTMWKTYY
50    TYAWYTRRKYRKMKYSRTTTKYAAMYKYWWYSTKAAAWYGRKRKKMGGKRKKWKMWKCYY
       *    ***********   **..******** *. .**  *****  *  * .*** **

45    RCKYRRKYKTAACYYKYKKKKRWKKYGTCRTYYRYRYKRYYRKTGTYRYRKMYRCWWSMM
50    RYKYRRKCGCTGAYYKYGKKGRWKKYRYYRYYTATGYKRYYRKCCGCAYAKMYRMWWSMA
      * *****   :..**** ** *****   * *    *******     * **** ****

45    KKKYKKYGCRCMYKRKGMRRWWKKSMWYMKTATKYAYRYMKTAAKWKWYRGRTYCKMWKK
50    KKKYKKYAMGTMCKRKKARRWWKKSCWYMKYMYKYGYRYMKYGRGWKWCGRRWTYGMWKK
      *******.   * ***  ******* ****   **.***** .  ***   *    ****
```

```
45              RWMKSYKKMKRYKKRKRRRYSWTMKYKCYYKKRYWGWRYKKRRWSKRYRYRRRKKMKKKK
50              RWMKSTKTMKRYKKRKRRRYSWWMKYKSYYKKRYWATAYKKRRWSKRYRYARRGKMKKKK
                ***** *.************** ****.*******.  ************ ** ******

45              CKAGRYAYMRRGTYCRGYKSGCTGCCTKYCCCYGTYRGTSRYRMMGCYWRRKRKYKRKMA
50              GKGTRYMYMRRTCYMRRYKSSTCTATAGYYYTCACYRRYGGYRMCKYYWRRKRKYKRKMR
                 *. ** ****  * * ***.   . : *    . **  . ***   ************

45              WMMRTYRRKATYMYRRGRKKACCYACGSCCCTCRCYCKKMYKKMAGRWCGGRYKKRAYRG
50              WMMRKCRGGRCYMCGGCGKKTAMYTGTGATAYYGAYAKKMCTGMWRATYACRYKKRMYGA
                ****. *    **     **:. *:  .. .   .*.*** . *     . ***** * .

45              TTKYRTRYGAKKKWMYGGGCTKKRYYKWKTAGTWTMRRYRMKYYKAKSMWMKWYRYCKYG
50              WWGTRCGCKGKKKWMCCTRTAKKRYYKWKGGTGWYMRRYRMTCTGMTCMTCKWYRYMGYK
                    *    .*****     :******** .  * ******.    ..*  *****  *

45              CGYYRWYKCTMTYRSYAYKTMKYYSWYYKKKRWGWCCTAKMYYMRSRRYMRYYRKMYKKK
50              SACYRWYTTYMKYRSYWYKGMKYYCWYTKKGRWKWYGCGKCCYARSGGCCRYYRKMYKKK
                .. ****.  *.**** ** ****.** ** ** *   .*  * **    **********

45              KACYKSYWYKKKYKWKCKKRKYWTTRRRKRWKYYRYYRRYKYYWKKRRRWRKRYRCTYRG
50              KRMCKSYWYKKKYKWKMKKRKYWYARRRKRWKYYRYYRRYGCYWKKRRRWRKRYATAYRK
                *   ************ ****** :***************  ************  :**

45              YYRKKCCRRKMKRAYGMKWRWYYYYYRYKYWKKKTKKRYYGTCAKCYKMGTKRYYGAAYK
50              YCRKKTMAGKMKRCYRCKWRAYYYYYRYKYWKKKCKKGTCACGTKYYKMCYKRYYSRRYK
                * ***    ****.*  *** ************* **   .  :* ***  ****.  **

45              RRYKRMWMMYWKRGCCGKKYTYRGKRMRKAACKMKWKYYRRRTCKSKRKKAYRRKYMMKT
50              RRYKRMWMMYWGGAATKKKYWYRKKRMRKTTTKAKWKYYRRRYAKSTAKKCYRRKYMMKA
                ***********  ..  *** ** *****:: * ******** .**. **.********:

45              RKGRCGCCCRKSSKRYTTYKYRGMKRKKKMKKWYTRRSAATTTCAKKKKKYYKAYKYYYS
50              GTKGMTTTTRKSSKATCYCKYRRMKRKKKMKKATYAGGGTGGYYGGKKKKYTKRYKCYYS
                 .       *****     *** *********     ..:    . ***** * ** ***

45              KKTCGRTTKKKTTGCTMYRKRYKRMRKYSKTMMYRRSRAARGGCTTKGKYRWRCSKYKAW
50              KKCGARCCKKKCGRAAMYGKRYKRMRKYSGKMMYGGSGMGRACAWKKKKYRWGYGKYKMW
                **  .*  ***   .:** ********** .***  *  .*. . .* ****  .*** *

45              YKMMKMMRKRMRRKWRYYKKMMRYYYAGGKMR
50              YKMMKMCRKRMRRKWRYYKKMMRYYYWSRKMR
                ****** ******************* . ***
```

The pairwise alignments of two accessions 45 and 51

```
45        AAARYGRKYYYKYRRGYRRGGYAMYKKRMYYYWKYYWYMRYWTYGCAGCTATYKAACYMW
51        CGGRYRRKYYYKYRRACRGRRTCMYKKRMYCYWKYYWYCRCWYYATRRTYRYYGWMYYMA
          ...** *********. *    .******* ******* * * *.       *    **

45        KRGYRYYRYCYYTCGWYWWRMYKYWYAYKKWMTASKYYRCRGYRWWRYWYKWYRYGCYKY
51        GRKYRTYAYTYYCTTWTTWRMYKTWYRYKKWMCGSKYYRTRAYRWWRYWTKWYRYRYYGC
           * ** * * **   *  ***** ** ***** .***** *.******* *****  *

45        ACRGYCRMYGAYKGCTYTMYKMKYKSTARRRKRWTTGWCTWYRWSRGYYMKRYSTRYRMY
51        RTRKYMAMYKRCKCYACGMYKMKYKSWMRAAKRWWWRTAAWYRWSRKTYMKRCCYGYGMT
            * *  **   *  :  ********  *  ***    .:******  **** .  * *

45        STKGKWRSKKKTWYYKYMCMRYMRRWRKMYKRYWKTTCYYMKRWYRYYWMTRYMKTWCWW
51        GKGSKWGCTGKCWTTGCMTMRYMRRWAKMYKRYWKACAYYMKRWYRYTAAWRYMKYAYWW
          .. .** .. * *    * ******* ********: .*********    ****   **

45        YRACACTGKWRYRYRRWYYRKWRRRMKRRKYYRRRRTCYTTKKRRSWACAASGYRTRTAM
51        YRRYMYYRKWRYRYRRWYYRKWRRRMKRRKTCRRRRGTYYYKKGRSWWTTGSKCAYGKMM
          **      **********************  ****  *  ** ***  :.*     . *

45        ATWMMMGRYGGMMRKCAYAKKYRRCKCWMYWYYRYCYWKMMRKKKGYYYYYYRYYCYKRY
51        TAWMMMRRYKKMMRKTGYRKKYGGYKYWMYAYYRYYYWKMMRKKKSTTTTCTRYTYTKGT
          ::**** **  **** .* ***   * *** **** *********.      **   *

45        WAAMWWYTYKKWWYKYMRYGWRWWCTAKCGTRKRSSRCTKWRRYKKMCYKKKSYWYAAGT
51        ATWMWWYYYKTATTKYMRYKTAWWGCGKMRKAKRSSRMWKWRRYKKMTYKGKSYWYWRKK
           : **** **.   *****   **  .*  . *****  ******** ** *****   .

45        TGKRYMKGAGGKKGAAYKRKYWRRRRKKGTGSYRYKKYRAAGWRYRKWKCKRRRWKRKGG
51        YKGAYMKKCATKKTWWYKRKYWRRRAKKRKRSYRCKKYRGTCWRYGKWKMKRRRWKAKRR
              *** .. **   ********* ** . *** ****.: *** *** ****** *

45        CRYAMWRYYGRCYYYAWTGCSKRCWYKRYYRRAAKKMKYSKWRYYYYYYRRYYWWRYWYW
51        YRYCAWRCCAAYYYYTTAAYSKRTWYKRCCARRRKKMKTSKWRYYYYYYRRYYWWRYWYW
           **. **  .  ***: :. *** ****   *  **** *********************

45        KYCKKRCYKKSRYTKRKTTGAYWKWGGRWCGGTRKRWWRTYKCAMYWRRWMKRMAKMWMY
51        KYYKKGYYKKGGTKKRKWYRRTWKWKRATYACKRKRWWRCTKMRMCWGAAAKRMGKMWMC
          ** **  ***.  .***     ***     . .******  *  * *    ***.****

45        GGRKYAKYTRGCAMTRCTYYCYTTKRRKMAKWKKSKMGKKYAWYWYCWTSKYYWTTAKYY
51        TRRKYGTYAGTTTMGGGCYYSYWYGRRKMCTWTTCKMKKKCRAYWYTAYCKYYWYGWKYY
            ***..*:   :*    **.*   ****..*...** **   ***   .****   ***

45        KWYGWYYKYKCMRAYYMMYYRYRRAKKARRRGMKSKKKRYGCAARYTKKGGRCMCKKWRK
51        KWYKWTTKYKTMRGYYMMYYRCAGRKKMARRKMKSKKKGCAATRAYKKKARRMMTKTWRK
          *** *  *** **.*******    **  ** ******   ..: *.**. * * *.***

45        YRMRSYTWKTRWYCCKKYRMMKGAGCAGWKYKTYKMYGKWRYWARKKRKAGSKKWRRAAY
51        YRMRSYCWKYGTYTAKKYRMAKRCRMCAWKYKACKACRTAGYATRKKRKGRSKKWRGTWY
```

```
                    ****** **   * .***** * .  ..****: *   .  * :*****. ***** : *

45                  TYSKWAGCCGMTYYRWKYWRYYYWYYYRRYKKRMTAWRKTGAAKKGGTMKGTYKYSRRYK
51                  KTGKWGAYMTMYYYRWKYWRYYYWYYYRRYKKRMCCWRKCAGCKKRACMKKYYGYSRRYK
                    . .**..    * ********************* .*** ...** . **   * ******

45                  CYRYRKRWSWCCTCWKWRTWYYYKTAWMGWKKKGKGCCACAYYKYYMMMRWAGCCAAYRY
51                  TYRYRKRWSTTGKAWKWRKWYYYKYWAAKWGGKRKSYYWYGYYKYYMMMRWMRYYWRYRY
                     ********    ..****.*****     *  * *.     .**********       ***

45                  MMYGTTMRYMYKKKRYYKRKYYRWKTGMKKRRYTCTGKRYYGASYSWRRKRYKKRYYKRM
51                  MMYKYYMATMYKKKRYYKRKYYRWKYRMKKRRTATKRGRYYKWSYSWRRKRYKKRYYKRM
                    ***   *  ***************  ***** : .  ***  ****************

45                  GRCTAAKKRYYGTCKKGCKMYKYRTKRMMMYWWCKTRRTCCTYRSYWYYKCYWWRMYRRR
51                  ARYCMMKKGTCACGKKKYKMYKYGYKRMCACTAYKWRRKYTCYRSTWYYKAYWWRMYRRR
                    .*     **    .  **  *****  ***       * **.    *** ****.*********

45                  WRSYRRKAYMYYYYWWCCTKTCCGRRCCCCCSYYRKRSKRYSYKRYYMYRKKMRGKRRSA
51                  WRSYRRKGYMYYYYWATAYKATYRRRTTMYTSYYRKRSKRCSYKATCMYRKKMRKKAGGG
                    *******.*******   . *:    **      ********* ***   ******* *  ..

45                  RKTCCKAAYYKSMMYRYTKTKGWWRGTYYRAGAYRYRYKYWKCGRAGTKKYKRWKKASYG
51                  GKAGTKCGYYKSMCYRYYGYGRWWRRKCTGGRRTRCGYKYWKARRGSYKGTTRWKKGSYK
                     *:  *..***** ***     ***  .    .    *  *****. *.. *   .****.**

45                  MYAKGCYGGTRRAWCRTTGAMKKTMYGGMAKCRWRYACSYYKYRGCRKKGKTYRAGAGRY
51                  MCCTKTYARCRRGAYGCAAGAGTYCCKKMRGTRWRCTASYCKYRKTRKGAKATRGAGARC
                    *  ..   *.  **.     :..   .      *    *** :.** ***   ** .*: *....*

45                  RAKRKKGTTYKTKKYRKYKRYRWCATCKACAAKCYAYGYGTKRTMRWACGYYCACTYCYY
51                  RRKRKKRKYYKCTKYRKYKRYRWAGATKRTGGGYCRYATAAKRACRWRMRTYSRYWYTYY
                    * ****  . **  .**********..: *   ..     *. .:**: **     *.    * **

45                  YYTMYRCACKTRYYSRKWKATTYYMKKYCYRYARRYGYKKTYYWGMRMMAAGCYMRWKKR
51                  YYCMYRYRYTKRTTGGKAKMWKYYMKKYACAYGRRCKYKKAYYWRCGAMTCCAYMRTKKR
                    ** ***     ..*   . * *  .******.   *.**   ***:***      *:. .*** ***

45                  ATGGKTSRKKAWMKYKWYGGAGTYGACTARGWCRRMYRRWKSWTAARMSWGTACYRWTTT
51                  TCACGYCRTGCAMKYGATAKTKATARYARGATMRRCYGRWKSWAGTRMSWRYMTYRWCWW
                    : .    .*. . ***    . : : .   :   .   ** * *****:.:****     ***

45                  MMKWRMWRGSAMTTCAWYRKYKKRKTYGTCRMSYRRRMTTWTKTYKWYMYRRRMKSWMRT
51                  AATAAATGKSGMKWYMWYRKCKKRKCCKCMRMSYRRRCCCTAKAYKWYMYARRMTSWCGC
                      .      *.*.   **** ****      *******      :*:****** ***.**

45                  TAKMRYYKKWKMMWYRRKKAGRYMKKTKKRYSYKYMKKYRTCMMKMRTYRKRKMAAAMKR
51                  CCKCRYCGKWKMMWYRGKKRKRYMKKCKKRYSYKYMKKYRAMMMKMRYYRKRKMGMGMKR
                     .* **  ******** **  ***** *************: ***** ******. .***

45                  RGCCSYKMASRCCMRWCRSYGGGKRKYCCKCKKGRRRCACYKRYWKCTGTYGWYRYTRRM
51                  RKATGYKMMGAYYMRWYRSYKRKKRKCYTKTKTCRARYRYYKRYWKMWRWCKWYRYYRRM
                    *  . .*** .    *** ***    ***    * *. * *    ******        **** ***
```

```
45    CGRCKMCRARTGRWRYAARYGRWARWTWWWRYKGKSKRKMRYCMSYYYRAGGKRRYKCRS
51    YRGMTCMGTRCARWRTGWGCKRWRGAAWAAGYKKKSKRKMRYGACYYYRRKRKRRCGYRS
          .   :* .*** .    **   :*   ** ********  .****   ***   **

45    RKGCKAAWRRKRRMRRKTTAYRMYYRRRWKYSRGCWMAARWWMKRMWMMTMWSMAYRGMR
51    RTAAKGWTARKRRMRGKCCWYRMYCGAGTGCSRKMWMMWRTWMKRAWMMKMWSACCGKAG
      *...*.   ****** *   ****       **  **  * **** ***.*** .

45    MRRYSYYMRYGRYRYYMMYTMRMWSCCCCWKYWRKACAYRYKKYKWKSMRKKMRKRYKKM
51    CARYGYYMRYKRYRYYMMCYMRMWSYMYYWKYWAKGTGYRYKKYKWKSMRKKMRKRYKKM
        **.***** *******  *****    **** *. .**********************

45    RWYYWYYGKGMMYGWRWWAATTWRYCCAAKKYGAKKYGCYYMKGYTAGKTCAGKKWKKTA
51    AAYYWYYRTRCCCKWRWTMCWGTRYYYRRKKTAGTGCAYCYMKACCCRKWYRKKGWKKAG
        ***** .     ***  .   **    ** ...  .  ***.  . *    * ***:.

45    TAACYWYMRGKKKMKYRYRKKKATRTTRMKKKYMYYKKKKTKRTTWYMCKYAKKCAYCAG
51    GTGYYWCAGKKGKMKYRCRGKKRYRCYRCKKKYAYYKTKKYKRWWTTMYKYRKKTGCTCC
       :. **    * ***** * **  *  * **** ***.** **    * ** ** .  .

45    TGWTYYWKYRRRRRYRYKRGRTKKYKKRKKKRGKGYGRYRYCCKCARKYRYKTKKKKKRM
51    CRTGYYWKYRRRRRYRYKGRAYKKYKKRKKGGKKRTRGYRYYYKTCRKTATTKKKGGGRM
          **************    ********   *    ***  * .**   ..**   **

45    ATARKWKKWRMWKKKWRRMMRYYSCYAWTRRKKYTMMMRRKAKYKKYKCCKTAGCCGKMW
51    WCGRKWKKWRMWKKKWRRMMRCCGYYGAGRRKKTWCMMRRKRKYKKYKYYKCGRYMAKAW
        .******************  . *.  ****   ***** ******  * .   .* *

45    YCYGKKRRSMKWMRWCYYTAARKRKCTTKTCACCRKKKMYGCTTAMTAMMTTWSKKRWRR
51    YYCKKKRRSMKWMRTMCTARTRKRKYYWTYYMMAGKKKCYKYCGGAYRMMYGAGKKRWRR
      *   **********    : :****   .    . *** *    .   **   .******

45    KWTATAAKMKYTMWWYRRRKCAKWWYYKKKTCATCKKRKWKKRRSWAGCAMTRYKTWMRY
51    KWYRWRWKMKYYMWWTRRRKAGKWWYYKKKYMMYSKGRTTKKRRSWGTTGMCRYGWTMRY
      **     **** *** ****..********    .* *. ******.  .* **   ***

45    WCYTTGWMRYMYARTKKAKKRYWRRYAATAGCAWTKRGTACCKKACGTWYKYACGAGCKK
51    WTCYYRTMRYMYMGKKKMKKRYWRAYGMYGASGWCGAACMYYKKTAACWYKYGAARRMKK
      *      *****  .** ****** *.  ....*   .    **:.. ****...   **

45    ARAWKRYCMRMRKSKSWKMWWYYKWYSGACRKKKYTATCKKGCCACYYCAGCKCTRYKRW
51    GAGAKRYYMRMRKSKSWKMWWTTKWYSRRMRKKKYCCATKKKMTGAYYMRSMKTARCTRW
      . . *** *************  ****   ***** .: **   ..**  . * :* .**

45    RYKYTKACAGCKAKRYSRMSCMRGYAKKKYRWTKRRCGRKTRYTGTCRWAYKMKRWKTMK
51    RYKYCKGSCTYKRKRYSRMSMMRRYCTKKYRAAKAAYTRKYRYATWSGTRTGMKRWKGMK
      **** *...  * ******* ** *..**** :*    ** **:  .     ***** **

45    KKKYKKKTKYKWWKYRMWARYKRKAWYKRKRKKMTYYYMYKAKWMRAWMMRAGSCTYKKK
51    GKKYKKKYKYKWWKYRMWMAYKRKRWYKRKRKKMYCTYMYKWKWMRRWMMRCKGYCTKKK
       ****** **********  **** *********   **** **** ****. .   ***
```

```
45      YTYRTKWKAGMYRRRKKKKWKKKCTRCAYKTCGCKRGGGTKGGTAGGMRRYKKRKCKWWS
51      YGYRAKWKGACTARRKKKKWKTGTGRYRCKYMRMKGTRRCKATACKRMRRYGKRKMKWWS
        * **:***..   ********.   *   *    *     *. :.  **** *** ****

45      YMKKYKKYAMYGGAGTKKKRRKKCYWYWMTTTYMAAGCCAKYAYRYCKKKKYKCKWYRYK
51      TAKKYKKYRMCRCWRAKKKRRGKTYWYWCCCGYMGCAYTTKYGYRYYKKKKYTMKWCGTK
          ****** *     :***** * ****    **...  :**.*** *****. **   *

45      RMWKKAWKKKKRYKRKKCRRTKCCYWKKKTGRRRYCYKYKWSRYGRRGKKKMKKRKYYSM
51      RMWKKGWKKGKRYKRKKMRAWKYSYWKKKCARRRYSYKYKWSRYARRKTKKMKTGKYYSM
        *****.*** ******* *  * .***** .****.********.** .****. *****

45      KKGTTTCGKKRKRYYCTYRSGWCYKRRSYKAKYCYRRGKAAKMMWWKRYKSKYKKCCTCK
51      KKKCCYAKTKRKRYYTCYRSRWTYKRRCTKMGYTYRGRKGCKMMWWKGYKGKYKKYYWYK
        **    . .******  *** * ****. *  * **  *..****** **.****    *

45      YKKRYMMTWMRRKKWKKAWYKKMMRYYMKRRRWRKYRTTGCYYYMYKMYATKYGATGSKK
51      YKKRYMMYWMRRKKWKKMAYKKMMRYYAKRRRWRKCRCCTTYYYMYKMYCCTYKGCACGK
        ******* *********  ******** ******* *    ********. .* . .. *

45      KSRCKKKRKYKMKYYYMWKRYYWMRSKKKWWYKGWRYRKYCGYYRKRRGRWKKYCAMRGG
51      KSRSKTKRKCKMKYYYMWKRYYWMRSKKKTWYKTWRYRKYYRYTRKRRKRWKGCYGMGRR
        ***.*.*** ******************* *** ******  * **** ***   .*

45      KTMRCCYYKYYMKKKKACCTYYYRGKKYMRKYYYRRKMWGCTMGYRMWRMGYTTSRRKKK
51      KYMRTAYYKYCCKKKKMTAGCTYRKKKYMRKYYYRRKMAATGMAYRMWGCKYKKSRRKKK
        * ** .****  ****  .   ** ************* .  *.****   *..******

45      KKWSYRYYWMKKRKRKRSYWKWKKKAKYMWKATTAGCGKKRKTTMRYYRWKWKYYTKYKK
51      KKWSYRYYWMKKRKRKRSCAKWKKKTKYMWGWYYMRARKKRKCAMATTAWKWKYYYKYKK
        ******************  *****:****      . **** :*    ****** ****

45      KWYRKMGTYKKRARWRASTYYCGRGYYKKKWYYRCMAKKKAGKKKKKRYRWRTMMYKYKY
51      KWYRKMKYYKKAGAWRTCCTYYRAKYYKTKWYYRMMRKGKRSKKKKKACATRYMMYKYKY
        ******  *** . **:.  *    ***.***** * * * .*****    * *******

45      RYCTTYAKKKAKYRKMMMYTWWKKWYYKAYKYSRAWAYCKKYGGMYTTGTYRYGMKKKRT
51      GCYKWYCKKKRKYRKMMMYAATTTACYKWYKYSRGAGTYKKYRRMYAGACYRYKMGKKAA
           . *.*** ********:  ..  ** *****. .  ***  **: . *** * ** :

45      TTMRKWRRARRAKYWAKKKTRRYYKCKYMRWKWYK
51      AAMGGARGMRGRGYTGKKKWRGTYKTKYMRWKWYK
        ::*   *  *   * .*** *  ** *********
```

The pairwise alignments of two accessions 46 and 47

```
46                RYMKRWYWYYSKWYWYRYMMRKMWRYMYWMRWKYYTRMMYYYYRKYKYRAYWGWWRKRWW
47                RYMKRWYWYYSKWYWYRYMMAKMWRYMYWMRWKYYWGCMYYYYRKYKYRTTWRWWRKRWW
                  ******************** **************   ***********: * *******

46                WWKWWRMYRMYKKAAGGRCRRYRWRMRYKMRTGKRYARKRWYSMGCGRRKKWRYMTCKYY
47                WWKAWRMYRMYKKWGKTGTRRYRWRMRYKMAARKGCCAKRWYSMKYAAAKKWRYCYYGYY
                  *** ********* .    *********** : *  . ******  .  *****    **

46                YSRRCYYRRWRKKACYGKKYRKWKMYCTRKTTWTYRYYMRRSKSMKTWYKRRMGGYYYRR
47                YSRRMCYRRWRTGTMYRKKYAKWKCTTARKKWWKYRYYMRRSKSMKGWYKRRMTTYYYRR
                  ****  *****. : * *** ***   :**. *.************ ******  *****

46                YMYMRWMMRRMMYYYRYKRWRRRYWRRWMYYYWKMKMGTRWRWKRMRKAYGWYKWYGYRM
47                YMYMRWMMRRMMYYYRYKRWRRRYWRRWMYYYWKMKMAYGTRWKRMRKWYTWYKWYRYRM
                  *************************************.   ******* * ***** ***

46                MKGMKWYTGYWWRWWYYYYKMMKYMTCKRATYYRRRRKRKRYYWYKMYMRYRWYRYYYRM
47                MTRMTACACCWTATAYYYYKMMKYMYYKGRWTYRRRAGAKRYYAYKMYMRYRWYRYYYRM
                  *. *.  :  *    **********  *    ****   **** ****************

46                RAYRWKKKYYRWRRYYRKMYTRYWWYRYRYRMRMKRWKRYRRWKWSKRRKWRKYSRWWYW
47                GTYGWKKKYYRWRRYYRKMYYRYWWYRCAYRMRMKRTKRYRATKWSKRRKWRKYSRWTYW
                   :* **************** ******  ******* ****  ************** **

46                WSMRYTCMKYGKYYWMGATGTKYYRTTYSYRRYRYWKYTKSAMRRKWTYAAACRKTYKRA
47                WSMRYYYMKYRKYYWMRMWKKKYYRWAYSYRRYRYWKCYKSTMRRKWACGCGYRKYYKRT
                  *****  *** *****    .**** :**********  **:*****: ... ** ***:

46                MWYTKKMRYYKMYKRWYKSYRWWRKYWMRYYMYYRYMTTAGMRYARRAASYWWMYYYYY
47                MWYWKKMRYYKMYKRWYKSYRWWRGYWMRYYMYYRYMCATRMRYTRRTRSYWWMYYYYY
                  *** ******************** ************ :: ***:**: ***********

46                RKWKKRMRYMYKKWAYCKKMYYRCTCTGTGCKMKRGKCTKSRKTKYWKCRWKMMRMYGRS
47                RKWKKRMRYMYKKWGYTKKMYYRYWMCRWRTKAKRRTYWKCRGCKYWGARWKMMRMYARS
                  **************.* ******        * ** .  *.*  *** .********.**

46                KYMWRGYKRRRRYRRYKKKRYYGAWTWKKKRRSAYARRRYGTRWYRYGRRMYTYRYRKKT
47                KTCTAAYKRRRRYRRCKKKRYYATACWKKKRRSRYTRRRYTCRWYRYRRRMYYYRYRKKC
                  *    .********* ******.:  ******* *:****  ***** **** ******

46                KKYTGRRKCCTKWKRACGTRKCGKKTTKMYACGAMSRKGYTCCAYMRRYRKKYSYYTRYY
47                KGYARRRKTTCGWKRMYRYRKYTKKWYKMYMAAGMSRKKYYYYWYMRRYRKKYSCCKRYY
                  * *: ***    ***    **  **  *** ...**** *    **********  .***

46                RRWKYKSYMYKKRYMGKWRGRYYKATRMGGAAYTYSYYKYYWATTCSYRWWRMYYKRYRA
47                RRWKYKSYMYKKRYMTKWRRRYTKWYRMKAMRCCTSYYKYYWWWKYSYRWWRMYYKRTAW
                  *************** *** ** *  ** .     *******  . ***********

46                KAMGYRAYMKAAAGTRYKWKYTTWCSWGRYRSWGAAYYMMYYWAACYKKARTGRYYRRGT
47                KCCKYGRCMKMRRKCRYTWTTCGAASWKRYGCAATWYYMMYYARWYCKKWRCAGCCRRKW
```

```
            *.  *   **     **.*.     .** ** . .: ******     ** * .   **

46          AYAMYYKRGTYTGKRMYYMSYWRTKWMGKRSWYRRMYYCATRTRYTWYRYGGKKKMMKYK
47          RYWMYYKGTGYACKRMCYCSYWRKKWMSKRSWYGAACCYTGRGRYGTTGYRRTKKMMKYK
             * ****   *: *** * ****.***.*****      : * **    *  .*******

46          GRMAYMRWRKYKTTKRMRAKYMKYKKYKRKCTYRYYYRKGRSCRSMGTTMYYWMYKTTCW
47          RRMGYMRWRKYTCAKRMRRKYMTTKGYKRKTAYRYYYRKKRSYRSMSKWMYYWMYKWCAW
             **.*******. :**** ***. * **** :******* ** ***.. *******  .*

46          KCWGGKWKGGTGRYRRKMKGWRKKRMKKTRKMKMMYYGRWMKMTYYAKKRKYKTRKTWRK
47          KTWAKKWKKSYKAYRRKCKKWRKKRMKGYRKMKCMYYARWMKMCYYMKKRKYKKRKYTRK
            * *. *** .   **** * *******  **** ***.***** ** ******.**  **

46          YRRSYYKWTTGKKGYRWYGTKCKRYYKWKCAKYYYTGAYMKYKKYKWKRKMYMRKWRYKM
47          YRRSYTGWGCTKKRYRWYAATAKRYYKWKTMKYYYKRTYCTYKKTKWKRKMYMRKWRYTM
            *****  *   ** ****.:..*******  ****. :* .*** *************.*

46          TYCYWYYYKKYRYWSYRYYWRKTTKRKKGRTGTYKWRYRYKKAYYRKYKRRKKYKAYKCY
47          CYTYWYYYKKYRYWSYRYYWRKCYGRGKRGYSKYKWRYRYKKWYYRGYKRGKKYKMYKYY
             * *******************   * *   ..********* *** *** **** ** *

46          RKKKKMRAYGYYYRRKRYYKYRKYARTRGTTTKKYKYMKCCKYWKATCRARWKYSRMYYC
47          AGKGKCRGTKYYYRRKRYYKYRKYRRKRKCCGGKYKYMKTTKYWKTATGTRWKYSRMYCY
              * * *.  ************** *.*     ******  ****::  :********

46          RKKKKKYRGATRGAKKMRYWWYYWYYMYRWTWKKKRSYKRWYKWMKKMYKYCRWYRKRRK
47          RKKKKKYRARGRAWKKMRYWWYYWYYMYRWKWKKKRSYKRWYKWMGTMYKYARWYRKRRK
            ********.  *. ****************.************** .****.********

46          KKKTKKSMKYCTAKMMKRCKRMGKRKKKYGRWRYCRMKKYWRYRKYWRMYRWWSYKKGSK
47          KKKYKKSMKYACWKMMKRYKRMKKRKKKYKRWRYYRAKTYWRYRKYWRMYRWWSYKKRSK
            *** ******.  ***** *** ****** **** * *.****************** **

46          KKYRKKYRWAAGACKTARRYWCRYYKKKKRRYRMYKTYYMGMGKMTKMYATGGAWWSMRW
47          KKYRKKYRWTTAGMKWMRRYWMRYYKKTKRACRMYKKYTCACAKMAKMYWWRKRWWSMRW
            *********::.. *  **** *****.**  ****.*  . .**:***     ******

46          RKSCKKKMRYWRYAKKTCYSYWYTKYYYRSMSWYRKMGGKYRRWYWRMKRKKMKKYRYRT
47          RKSMKGGMRYWRYWKKCTCSYWYCKYCCRSMSWYRKMKKKYRRWYWRMKRGKMKGYRYRY
            *** *  ****** **   **** **  *********  *********** *** ****

46          GMWRYRYMRYYRYSKRWTSWKKKTRATGGRYYYKKMYKWYMCWCWGTSRWYKKRTATYMR
47          AMWRTAYMRYYRYGTGAYSWKKKKAWKSRRTTCKKMYKWYMYWYWTCSRWYKKRYMYYMG
            .***  *******..   *****.  .. *   ******** * *  *******   **

46          YCCRMAYKYYCMKRTCTYRYGAYWTYRSYYRRRWRYSTWYKMWMMYGWWWAAACCCRKYY
47          TGTAARYKYYMMKAYTWTGTTGYWGCRGCCGRRWRYSYWYKMTAMTSWWWTCTTGYRKYY
                  **** **        .**  *.   ****** ****  * .***:.:   ****

46          KSSRKYWYRKYYKTARYKRWYMKRRWRWRKRYKCKYYKKKMKRRYRKRYAKKMYRYRKAK
47          KSSRKYWYRTYYKAWRYKRWYMKRRAAARKGCKTKYYKKGMKRRYRKRYMKKMYRYRKRK
            *********.***: **********   **  * ***** ********* ******** *
```

```
46              KRKYRKKKKWRRKWKRMYSKRKCSKRRWMGYKRATMYKKTAKTTWYRRYYTAATTTRCSK
47              KRKYRKKKKWRRKWKRMYSKRKMSKRRWMRYKRWWMYKKYGKYYWYRRYYWTTCGGGAGK
                ********************** ****** ***  **** .*  ****** ::    ..*

46              RSYKKMRWKKGTACGAYTMKYSRYSKYAWRWWRMRKWSMRRWYWKMWRRWKYKKRYSYWW
47              RCTKKMRWKKRWRYRWYCMKYSRYSKYGWRWWRMRKWSMRRWYWKMWRRWKYKKRYSYWW
                *. *******      * *********.********************************

46              RGAATTTAAKRWKMMAGGKRWSYRRKTYMKRYWWWKYKKTTKRYMCKGRYYGYYKWRYYA
47              RACCGYYRWKRWKMMGAAGRASYRRKYYMKRYWWWKYKKCCKRYASKKRYYKYTKWRCYC
                *...     ******... * ***** ************  *** .* *** * *** *.

46              KRSRRKYYRKRYYWYKKRWKRYTKSAKWKAARRAKKYRRRKWMMMWKKRCARRKYRRTMY
47              KRSRRKYYRKRYYWYKGRWKRYYKSWKWKMMRRGKKYRRRKTMMMWKKRYRARKYGRWCY
                **************** ***** ** ***  **.******* *******   *** *  *

46              RYWRYGCAYYRRYRRWRYCYYMKSYRSSYGMYRYYYRYRMKARCWMWKKYCWWMKCKKGY
47              RYWRYATMYYGRYRRWAYTYCMKSYRGGYRMYRYYYRYRMGTRSAATKKYYTWMKAKKRC
                *****.  ** ***** * * *****..* ********** :*.   ***  ***.**

46              KKRMRRKTYMRSMYKTMYKGRMRMMWKMYYRTCASYTTKKYRRAGCRYKWSYRTKKRRTM
47              GKRCGRKKCMRSMYKAMYKTRMRMMWTMYYRAGRGCYYKKYRGMATRYKWSYAYKKRACA
                 **  **. ******:*** ******.****:  .   ****  . ******  ***

46              MTCKKKMYYKYRYKRKWWYKRYYYRYKCRYRKMKAYWAMRKGAKKKKYSMRYMRKKKYRK
47              AKTKKKMYYKYAYKRKWWYKRYYYRYKYRYRKMKMYWGARKSMKKKKYSMRYMRKTGYRK
                 . ******** *************** ****** **. **. ************. ***

46              YTYTCARSYYKRWMKSSARMKRMTRYRKKYRRKYKYKYGYKKRYRRYYRKMRMRKRRKRR
47              YKYAMMRSYYKRWMKSSMRMKRMGRYRKKYRRKYKYKYTYGKRYRRYYRGMRMRKRRKRR
                *.*:  *********** ***** ************** * ******** **********

46              KKMYCKMRKCYSKRYGCYKRYYKRYYYYKRKRKGAATTMRARMGKWRYRKTGTSRRRKRY
47              KKCYGKMRKTCCKRYCACKRYYKRYYYYKRKRKATCWAMRCRMRKWRYRKAAGSRRRKRY
                ** * ****  .*** . *************** .:. :**.** ******:. *******

46              KRYKRRWYGRKKKKYRRRKYYYKRRGYWYMYYYMKYRYWYRRRKYKRRWYRMYMYYYATR
47              KRYKRRWYARKKKKYRRRKYYYKRGRYACAYYYMKYRYWYRRRKYKRRWYRMYMYYYTWA
                ********.***************  *   ***************************:

46              WKKKSRRKMRKRKKRRWYRRYKCYKRAYTKYMYKRWYYYRRRWKKKMYWRKMRWRRKWWR
47              WKTKSRRKMRKRKKRRWYRRYKTCGAGCYKYMYKRWYYYRRRWKKKMYWRKAGARRKWWR
                **.*******************    .  **********************   ******

46              MKWAAMRYYRATKSGAKYKSGTCRRKSYRSCMWRYYTGAGTSWTKATRRRRRKRYSYAMY
47              MKWMGMRYYRCKKSAGKTKSKYYARKSYRSYMTACYCKMACSWCGTGRGRRRKRYSCGMY
                *** .*****..**..* **    ****** *   *   . **  : * ******* .**

46              RYYKYKWKCGCTWKWTWGCRKMTKYWKRTRRRWYMYRYAWRYYYYKYYKKKYYRWKYRKK
47              RYYKYGWTAATAWKWGWRYRKMGKYWKRKRRRWYMYRYMAACYYYKYYKKKYYRWKYRKK
                ***** *... :*** *  *** *****.*********    ******************
```

```
46             KYWMAAGTTAWKKGKKYYRYYMKMYKRKKKMKWYMWYWCKTGRYRYKMKKRMYTRGCRKK
47             KYWMTRRGWRWKKAKKYYRYYMKMTKRKKKMKWYMTYWMTGTRYRYKMKKRMYCRRTRTK
               ****:     ***.********** ********** ** .  *********** *  *.*

46             RYRKWRRYYYYRYCRRKKYKCTRCMTRKKKMKKWKWWKKRYSKYYMWMTKKKYASYWKWR
47             RYRKTAACCYYRYARRKKYKAARTAYRKKTMTKWKWWKKRYSKYCMTMGKKKYMSYWKWR
               ****     ****.******.:*   ***.*.************ * * **** ******

46             RKRRKYWRWATYKMRKMRKYKSYCYKKMRKRSGACRKYYKKMYKKSYKWMRWYRRKWYYK
47             RKRRKYWRWTGYKMGKMRKYKCTYYKKMRKRSRMMRKYYKKMYKKSYKWMRWYRRKWYYK
               *********: *** ******.  ********   *************************

46             KWMWRKWYRRKKYWTKKKYYKKRWCATCRWSKYMTRKRRSMRWATAAYYKYRWYRKRKRK
47             TWMWRKWCRAKKYAYKKKYYKKRWYTCTRWSKYMKRKRRSMRTWWRRYYKYRWYRKRKRK
               .****** * ***  ********* :  ******.*******     *************

46             YYYRYRAGCWRRAWKGKKSYRRYYSCTMACKKKTACRYGYYCWRKYKKSKYRKKRYMKRR
47             YYYRYRCRTAGAGATKKKSYRACYSTAMGTKKKKMTRYTYYYWRKYGKSKYRKGRYMKRG
               ******.     . . *****  ** :*. ***.  ** ** **** ****** *****

46             KWMYRRWKYYMRTYYKKYYSYCYYGKRACTWKYYKYRRTGACAYYAKTRSYGAMYRKWCG
47             GWMYRRWKTYMGACYKKYYSTYYYKTRRYKWKYYGCRRARRYGYYRKCRSYATMYRKWAT
                ******* ** : ******  ** .*  .****  **:   .** * ***.:*****.

46             TTKKRYYARYTCAWAKRKYGCYCCRKRKRKKKRKMRRKGWYMYSMRKWWKYMKRYKRKYC
47             CGTGGYTGGTWACWRGRKCAMYYARKRKRKKKRKMRRKAWYMYSAGKWWKYMTRYTRKYT
                 .  * .   ..*  ** . * .**************.*****  ******.**.***

46             TCGGCWWKWKKMRMKKARKRRKKMRYWYSKKAKKRYYRMRYKRMKSWKKWKYKRKYRKWR
47             WSRRYWWKWKKMRMKKRRKRRKKMRYWYSKKMKKRYYRMRCKRMKSWKKWKYTRKCRKWR
                .   *********** ************** ******** ***********.** ****

46             KKKYYYRRKKWTKYSSRMYRRMGYGRMKSYKRKYYKAYMRMSCTTASKAKSRYRRKKKRT
47             KKKYYYRRKGWCKYSSRMYRRMAYKRMKSYKRKYYKMYCGASTCCCSKMTSRYRRKKKRK
               ********* * **********.* *********** *   *   .** .*********.

46             ATYYKRAGKRYMRGSAAYTKRKKYRRGGRAGYYRRMKRRWKRASYYYMYKGYCYRYACTT
47             RYYYKRWRKRYMRKGMMTWGRKKYRRRRAMRCCRGAKRRWKGMSYYCAYKRTTYRYGYYW
                 ****  ***** .     ******       *  *****  ***  **   ***.

46             RMKYCYYMYAKTYKKRWMMKYRWWYRGAWCRTGTYKKYTGATRKKRAWRKMRMWYWWYRA
47             RMKYYYYMYGGCYKKRWMMKYRAAYRRWWMRCTYYKKCARGCGKKGRWRGMGAAYWWTAG
               **** ****.  **********  **  * *   *** : .  **  ** *   ***  .

46             WGGAGASWKKYYRATYRYWAYKRRYRGCKYKKKKAMWWMYKMCYCCAKYWRRKRYRKMYR
47             TTRMTWSWKKYYRTKYRYWGYTGRYRRAKYKGKKRMWWMYKMGCYTGKYTAAKRCRGMYR
                     *******:.****.*. *** .*** ** *******    .**   ** * ***

46             MWKMKYKRYCCTCKMMRYWKRWRCYSWRYKWKKRKYWMKKMKMKGTMKKWYRRYYMWYKK
47             MWGMTYKRYTGKAKMMRYAKRWRTYSWRTTWKGRKYWMKKMKMKKKMKKWYRRYYMWYKK
               ** *.****  ..***** **** **** .** *********** .**************

46             YKMRGTATRYAYCTMKMYRTAYAYYSKMAAWKCRWKYYYWKYWKKRRGGCTYWRMYYYRW
```

```
47          YKMRCCGCGCTCAAMKATGWWYWYYSKMRWWKAGAGCTYWKYWGKGGATTGYWRMYYYRW
            ****  .   : .:**     * *****  **.     ***** *  .   *********

46          KTGATKKYSMRKRKRYRRMRRKWKKKYKRYRYYCAKYTGKKKATRGYYYKSYWWMMKRYM
47          KWKRKTKYSMRKRKRYRRCRRKWKGKYKRYRYYTMKYWKKKKMAGKYYTKSYWWMMKACM
            *   ..************ ***** ********  **  *** :  ** ********  *

46          RKRKKRYRKMWMRMWMRKYRMSYRCYMRYYMKTRRKWWGYYYRKRRKKRTTYTCRKMKSK
47          RKRKKRCRKMWAAAAMGTYRMSYRYYMRYYMGYRRKWWRCYYRKRRKKRGYTAYRKMKSG
            ****** ****    * .****** ******  *****  *********   : *****

46          RYYRRRKTKKGKMRKAKRMKMYYCCCRGRYYSRRKYKKYKKRRYKCKTCYRKKRKRKWYW
47          RCYRRRKYKKAKMRKMKAMKMYYTTYRKRYYSRRKYKKYKKRRYKYKCTYRKKRKRTWYW
            * ***** **.**** * *****   * ***************** *  *******.***

46          YWMKKKRYKWMKYMTKYKTWSRRMYWYYKWRRYKKCKKTATYRWKYYCRCYYRKRKKYMK
47          YWMKGKRYKTMKYACKYGYWSRRMYWYYKWRRYKGYKKACKCRWKYYYGYYYRKRKKCMK
            **** **** ***  **  ***************  **:.. *****   ******* **

46          KMKMMKTTKKMYKRAMYMRKMRKSSMKSKKKKRYTCTAWWRKKKTMYRKMRRKKRRKYWK
47          KMKMMKCWKKMYKRCMTMRKMRKSSMKSKKKKRYATWRWWRKKTKMYRKMRRKKRRKYWK
            ******  ******.* *****************:   *****..***************

46          KKKKKYMRRGCMKSMRYRKKKMKWKKWTACKRMRKYRKKWRCRMMKKKKKAKKRRKKYRR
47          KKKKKYMRRKTMKSMRYRTKKMKWKKWYTYKRCAGCAGKWRTRMMKKKKKRTKRRKKYRR
            *********  *******.******** : **      *** ******** .********

46          KGARYKYCYRMRYYKKRYKRKAMRKKRRKYTYKYYTAGRRRKWKKKYWMMKKKRCTGGKK
47          KAWRYKYMTRMRYYGKRYKRKRMRKKRRKYYTKYYYRKRRRKWKKKYWMCKTKRMCAKKK
            *. ****  ***** ****** ********  ***   *********** *.**  . **

46          YMRKYKSYWWMYRYRRYRAGRRYYKGRSKSMYCCTKYKAAYTKKKYRKKKKKSMRTKKAC
47          YMRGYKSYWWMYRYRRYRTARRYYKSRSKSMYYSWGYTTGCCKKKYRKTTKKSMRGKTCY
            *** **************:.*****.****** .  *.:.  ******..***** *..

46          SMYRYYWKYYSAYRGAACRTTCRWTGATKRGKKWKKCTYTCTGATGTKTMYCCKTAGCRY
47          SMYRYYWKYYGRYRKRRYRWYYRWYARCKGRKKWKGMYCGMCAGATYTWMYAYKCGAGRC
            **********. **    *   ** .  *  ****       ..:  . **. * .. *

46          RRMYYMRRSWRCYWKYTRWYGAACGAKWWKYMKMRWYRMRWTGKKRKRWWSARMTTMAGR
47          RRMYYMRRSWRYYWKYKRWYRGCMCCKWWKYMGMRWCACGACKKKRGRWWSRRMWGMTAR
            *********** ****.*** ..  .****** ***       *** **** **  *:.*

46          MYSSTTAMAMRMYRWWGKKRRRYTGRRY
47          MYSSKKRAWCRMYRTTAKGRRAYYRRRY
            ****..    ****  .* ** *  ***
```

The pairwise alignments of two accessions 46 and 48

```
46              YMRYRYYMKYWAMYWKYRMCTWYSKWMRYRYWRMKRYKTMKRRSKWRWRKMKMYYYYKCC
48              YMRYRYYMKYWGMYWKTACAKWYSKAAGYRYWRMKRYKYAGAASKWRWRKMKMYYYYKTA
                ***********.****   ..****   **********     *************** .

46              GTYRWRRKWKMYYRRYKTCTYKCKGWMWYRRKWYCAARYYYYWRYYYKYYYYRRRRGKMW
48              AAYGARRKWKAYYRRYKYYKYKTGRWMWYRGGWYGRTAYCYYWRYYYTYYYYRRRRKKMW
                .:*  ***** ******  .**   *****  **  : * *******.******** ***

46              RKWYYKRSMYYYYKWRRYMKWRYRRYKMTRRYMMRRMWRMCCYYWKYMYRSYYWKRKYWY
48              RKAYYKRSMYYYYKWRRYMTTATRRYKMYGRYMMRRMWRMYYTYWKTMYASYYWKGTTAC
                ** ****************.   *****  **********   *** ** ***** .

46              YYRWYRRTYYYCTYRMRYMRWYWYYRRMWKSRRYMKWWRRYCRRTARGKYKKWYKWWYRW
48              TTRWYRGKCYYAYYRMAYMRWYWYYRRMTKCAACMKWWRRYYAGAWRRKYGGTTKWWYRW
                  **** . **. *** *********** *.   *******   : * **    ******

46              YYRRWWYWRSYWWYGKRWGMYRRCATCCKYWWSSGWWTKYYKKYYTWMRWYKMRYCCGGK
48              YYRRWWYWRSYWWYRKRWAMYRAMTAYTKYWWSSRWWAKYYKKYTCWMRWYKMRYTTTTK
                ************** ***.***  ::  ****** **:******  *********    *

46              KYYGGTMYKRKSWKKSGRYTYACYYMRMWYMKGWWRMRSYYRWWWYYYKRRMMCMWYWMW
48              KYCTKAMYKRKSWKKSRAYWCTYTYMRMWYMKKWWRCGGCTATWATCYKGRMMTMAYAMW
                **   :**********  *  :  ******** ***  .    *   ** *** * * **

46              YRMSRRYCGAYKWYKWYMRWKSWAKTWAARRMTRMKYYMYRMRYYSKMRKYMYKYYWKRS
48              CAMSRRCYKRYKWYKWYMRWKSWCKWWWRRRMKGCKYYMYRMRYYSKMRKYMYKYYWKRS
                  ****    *************.* *  ***.  *************************

46              WRWWYMKAKWMCYWRKRWRKYYSRRWCTGYMMWKYRKYYKRRRKKRSYMRMYRRRYTGRA
48              WRWWYMGTKWMMYWRGAARKYYSRRAYYKYMMWKCGGYYKGRRKKGSYMGMYRRGTARAM
                ****** :*** ***   *******    *****   *** **** *** ****  :

46              TKTRMRRWKWKRCGYYRMYKMWRYMKYMRKMGYKRRYYRRRWCGAKKRTMMKYRKYRTGC
48              AKKGARRWKAKGYKTCGAYKMWRYMKYCRKMACGRRTCRRRWYKRGKRCMMKYRKCRKKT
                :*.  **** *       ********* ***.  **  ****    ** ****** *.

46              TTTAYKKRYYGARKKGWWRRSMYCTYKYKWRYYYYWRWSGCYCRYRKYYTCKWRKMRRCM
48              CGAGYKKRYTKRRKGKWWRRSMYTATKTKWRYYYYWRWSTTYTATGKYYWMKWRGMRRYM
                  :.*****   **  ******* : * ***********  *    ***  *** *** *

46              GWWMSYRYMYAKCARRYTKRWRRYYKRRRKKYKKMMRRYYYWKCCKRWTCGSWRKMRKYW
48              RWWMSYRYMYMKYCGRYYKRWGATYKGGGKKYKKMMRRYYYATYYKRWCATSWRGMRKYW
                 ********* * . ** ***   **   ************  .  *** . *** *****

46              MMKMKATKMKKKAGTWYTMRYKKYYMWKKYRYYRACTKCYRYKYRKRKKKTMKKYYKTGA
48              MMKMKTCKMGKKRKYWYKCRCGKYYMWKKYRYYRTTAGYCGCKYRGRKKKWCGKYYKACC
                *****: ** **   **. *  ************: :     *** ****   ****: .

46              CKKKKTYYCRWKYYYWKYWRRRKRSYYWRYMSKYSMRKTCARRMMATRKRMKAKRRKKKK
48              YTKKTGYYAGAGCTYWKYWARRGRSYYWRTMSKTGMRKWYWRRCCTCRKRCGRKRRKKKK
```

```
       .**. **.     ***** ** ****** *** .***   **  : ***   *******

46     KYCGTKWYTTTRMWRYWKTMRTRYCSYAGAYKMKCYCGTRCYRYRRTRKWRRYYYRWYYR
48     KTYRYGWYCCCRMWRYWTKMRAGYYSYRKMYKMKMYYSYRYCGYRRCAKTAACCCRWYYR
       *     **   ******..**: * **   **** * . *   ***  *      *****

46     YYKKARAGCGGKYWRYRRKKYRATTCRKYKKCRYRYAMRYYYKRYKRYYTKYKCAMMMKK
48     YYKKTRTKYRCKYWRYRRKKYRRWWMRKYKGMRYRYWMRYYYKRYKRYYAKYKAGMMMKK
       ****:*:    ***********    ****  **** ************:***..*****

46     SWKKTRRKYYTAGRWRCACYYYKMYKKKTCCKYWMAGATGARKYKKYKYKKRCGCSWYWR
48     SWKKKRRTYYYRKRWRTGYYYYKMYKKKAYMKTACMATGCCRKYTKYKYKKRAASSWYAR
       ****.**.**   *** . *********:  *    .:  .***.*******...*** *

46     CGGARRYRRTRYKKKRTRSKSMYKKKKTKRSYRMKKYMAMYRYYWGGCKKKKKMSYTRRW
48     YRRWAATGRKRYKKKRGRSKSMYKKKKWKRSYRMKKYMRMYRYYWRTTKKKKKMSYYRRW
               *.****** ********** ********** ******   ******** ***

46     GCYWWRRYRKYKKWGGSYWWKTRKYYSMYRSYRRKAGYMTRAKWYWKWWKKKKAGYMGYT
48     ATYWWRRYATTKKWRKSYWWKWRKYYSMYRGCRRKRKCMARGKWYWKWWKKKKRKCCTCY
       . ****** . ***  ***** ********. ***   *:*.***********

46     KKRKYGKKYYYGCRYRYYWKRSWRWYYCAYTKRSKMKGYGKYYWYKKWWMYGTCAAKAAW
48     KKRKCRKKYYYRYRYRYYWKACWRWYYMCYWKRSKMKAYKKYYWYKKWWMYSYYMRGRGW
       ****  *****  ******* .***** .* ******.* ***********.      .*

46     KYGGRWKKYRMYRTYYYWYKTYWRRMMYTRYRKMAYMKRRKWRCARRKYKACMWYWMTAT
48     KYKKRWKKYRMYRKYYYWYKCYWRRCMYKRYRKCCTMKRRKARYRARKYKTTMTYWCYGC
       **  *********.****** **** **.**** . ***** *   ****: * **  .

46     YTKSYWYYWRRCGCKKKYRSMSMWYKWKRMCTMWYWRMKKKCWWCGCRYRYYSTYYYYYC
48     CCTSYWYYWRRYSYKKKYRSMSATTKWKAMYYMWYWRMKKKYWTYRMAYRYYSCYCYYYY
         .******** . ********   *** *  ********* *     ***** * ***

46     YKTCAACKSWMRKWYYKKKGATATGYSRAYWAGCCGSKRWYKRGTRMTCCACCSRMKWGM
48     YKYTCGTTSWMRKTTYKKKKWWTGCYSRRYTMAMYRSKRWYKRKCRMWTYCYYSRMKWKA
       **  .. .*****  ****   :  *** *  .   *******  **   .  *****

46     KYYAYYYTYRRWKWMYKYCWWKWWWKKYRRKRKYTAWWMYTCAATRYCGWMKRRKYGRCG
48     KYYWYCCKCGRWKWMYKYGWWKWWWKKYRRKRKYWCWWMYKSMMWRYTTTCGARKYKRTT
       *** *  .  ******** *************** .****..   **      *** *

46     RYYRRCTTCCSYMTRWCRKKMYTCAGGGTKTTTRYTRKMKYWKTYKAATTYYSYTYKYRY
48     AYCGAMCCTASYMCRWYRKKAYCMTTAKGTCWWRYWRGMKYTKKYKMRWYYYSYAYGCGT
        *       .*** ** *** *  : .  .   ** * *** *.**    ****:*

46     RARKMYSCYWYYWYYYGRGCGTYACRWYYRMKYMTTMKRARKCGAKWCCYYYKWSYYWRW
48     AGRKMYSYTATTTTCYKRRMRACMYAAYYRMKYAACMKRWRGSTCTTAMYYYKWSYTTRW
        .*****        * *   :     ****** : *** * . .. . *******  **

46     WRAGYRGAAMCGATGGGKATKRWYWWARKGAGWKMTRYSMKTRMYTRYCTWSKAYCRRWK
48     WRTAYRKRRMGTGCTTAKGCKRTCTTGRKTTAWKMKRYSMKGRMYKRTYCWSKCCARRWK
       **:.**   *  .   .*. **    .** :.***.***** ***.*   ***. .****
```

```
46     ACARYAYWCYRYCTGARAATMAAMGTRWWWKRMCGGGGRGKKTRKKRYGWRRWSAMCRYA
48     TACGYMTWAYRTYYTTARWWMMGCCYRAWWKRMYAARRGAKKKKRKKRCRTGATGGCMRYR
       :.. *  *.**    :    * .   * ***** ..   .**.****      ..  **

46     ACTCCWSMTTRRKWMYRKMSWAMYYCYRYRWRARYYRKYTGRGKRKWKCKYSMYAKAKWC
48     MMYYYWSMWGGRKAMYRGMSWGCCTYYRYRAGTRYYRKCCARTGAKWGAKYSMYWKMKTY
            ***   ** *** ***.    ****  :*****  .*   ** .***** * *

46     SGCAYYRWKRKKKYGTGGKRKWRRYKRWKKCAWKMYRMRRTKMYTCWYASYKYCYCAAYY
48     CRTRYTRWKRKKGYTGAKKRGARRYKRWGTARWKMYRMGRCKMYYTACTSYKYSTYTCYY
       .   * ****** *  . **  ****** .. ****** * ***    :****.  :.**

46     YKKYRRCKGRMRYYYKKRARYYATAMWYYYTTMGKRKWWKSKYKGMRYYRCRMKCCCGGS
48     YGTCAAMKKRARYYTKTACRYYGCGMWYTYCCMAKRKWWKSKYKACRYYGTGMGATTTAS
       * .    * * *** *. .***. .*** *  *.**********. ***   * .   .*

46     YWRRWKYRWMKMRWRKYKSYMKMRKRSRMRKYCKTWYKKATGGAWYYRKKKRWRYWRYWK
48     YWRRWKCATMKMGARKYGCTMKAGKRSRMRKYTTKWYKKTWSRMTTYRKKKAAAYWRYTK
       ******   ***  *** . **  ******** ..****: .    *****   **** *

46     WMTRRWWRKKRWKRYSRRKYRARKTTTTYYYCRSKYYKKKGGGTRYGMWSRKMKTRSMRC
48     WAKRRAWRKKAWKRCSRRKYRMGKGAYCYYCGGGKYYKKKRAAWRYKCWSRKMKCRSMRT
       * .** **** *** ******  * :  **   .****** .. **  ****** ****

46     YRYRKGCKYYCAGGYRRTKYRMTAYACGWAGATKKGMYYYKKYRYRWYKARYYYRYKWKG
48     YRTGKTTKYYACAAYRRYKYAAYWCGTAACRWKKKTMYYYKKYAYRWYKRRYYYRYGWKA
       **  *  ***....*** **     . . .  .** ******* ***** ****** **.

46     GTARYYKRRRKMKKRKKKKRRRRKWMMYSKRKRKYYSYYKCAMSSARKRMTYYRSSMYRK
48     ACGRYYTGGRKMKKRKKKKRRRRKWMMTGGRKRKYYSYYKTMMSSMRKRMGYYRSSCTRK
       . .***.  ****************** . **********  *** **** *****  **

46     KYRRYKYYKMMRRRCKKWGATGTYRKCGARKYKRKKRYRGRRYKRRRCCCGMWKMGYKCY
48     KTAGYGYYKMMRAGMGGAARGRYYGKGSTRKYKRKKRYAKGAYKRRRAYYKMWGMAYKTY
       *   * ******      .    * * .:*********    *****.   ** *.** *

46     KRYTTGGCATCYYMRGYKTACKRKRKMRKKCYMRYRRYRWMMKRGAGRMRWKRKKWWYKY
48     KRYWWKKACCYYYMRKYKCWYKRKRKCAKKYYMRYRATGTMMKRRWRAMRWKRKGAWYKY
       ***    ..  **** **   *****  ** *****    ****    ******  ****

46     TGKRMWKKRARYRRRWSWTKRKTKYRYRYYRRYKRKMTWMYKWTKKMYKWKCCRTRRKKK
48     KKKRMWKKRGRYRRRWSWGKRKWKYRTRYCGRTKRKMCWMYKWYKTMYKWKAYRKRRKKK
       . *******.******** *** *** **   * **** ***** *.*****. *.*****

46     MRMKRWGSSMTRTMYYKYWKRKRGCKKRKKSWYRCRKMRRKKACKSRMYKKWKWYKKGYR
48     MRMKAWRSSMYAWMYYKYTKRKGCTKTRKKSWYRYRKMRRKKRYKSRMTGGWKWYKKAYR
       **** * ***   ***** ***   *.******* *******  ****   ******.**

46     YRYYWKKYYYYGCYKYKYRKWATGKTGRCKMYMRRKKWRYTGAYKWRGMYYYYSKMATCA
48     YRYYWKKYYYTAYYKYGYRKWRWRKYRRMKMYMRRKKWRYCKTYTWGRMCTYYGKCWYTT
       ********** . *** ****   *  * ***********  :*.*  *  **.*    :
```

```
46          AGMTRYWSKRKCRKRKTYKWATTRAKWGYWYRYKYYRWSKTCCYRGCWRYKWRYRRTKYY
48          WRCCRYWSKRKMRGAGGYKWRYWRMKWSYWYRYKYYRWSKKYSYRRMWRCKWRYRAAKYY
                ******* *    ***   * **.************. .**  ** ***** :***

46          YRRRYKRKWKYKRYKGKKWYYKTRKKKYSRARYYWAMGGCCAKKYMKCWTYSRMCTTRTK
48          YRRRYKRKAKYKRYKKKKATCKKRKKKYSRRRYYWGAAAYGGKKYMKTTCYSRMYACRCK
            ******** ****** **   *.******* ****. ..  .*****   **** : * *

46          KYTKYKKKGATRGAKYTGGKYMKMTRWYYYRYWRYKTGGKARAKRKTGTYRYRSYKKRWY
48          KYKGCKKKARGRAWKTCACGYMKMCRWYYYRYWATKWRSKMAGKRKCCGCAYASYKKRWY
            **.  ***.  *. *  .  **** ********  *  .*  .***     * *******

46          KMMWMKKKMYKYRWMYRGTKRYKKKKGWYMYGCAMKKRYKYRYRMGGATTTKSCCGCTRW
48          KCCWMKKKMYKYRWMYRAWKRYKKKKKWYMYAATCKKRYKCGTRMTRMCAAKSTTATKRW
            *  **************. ******* ****..: *****   **    ::**  . .**

46          RYYRSRRKWRYRMYWWWSKYAKSKYRKATKKRKRMRCWWYWKRYTTYTKYRMKYYMAKKK
48          RYYRSRRKWRYRMYAWWSKYCKSKYRKGYKKGKRARATWYWKRYACYYKYRAKCTCGTKG
            ************** *****.******. ** ** *. ******: * *** *   ..*

46          WWSMYCRMKGWKKKCGMGGYKWKKYKKKCAKGSRGGYKYWYMRMMYGYMWRGYKRRRSRK
48          WWSMCGRMKKWKKKAAMRKYKWKKYKKKAWKTCGRRYKYWYMRMMYAYCTRAYKRRRSRK
            ****  *** ****..*  *********. * .   **********.*  *.********

46          KRYKRYYKKASRYYRRAKRRRWYWWGTYGCRWYRRMYKTYRYKKKKRYYMRYSRYKYGKK
48          KRYKRYYKKWSGYCRAGKRRRWYWWTCCCTRWYRRMYKYYRYKKKKRYYMRYSRYKYRKK
            ********* * * * .********     ******** ****************** **

46          RYMRGSMCTGCKYKKKKATYMRKYRCRKKYSYYGYYCYYTRARKYKSYKCYKKARYMRRW
48          RYMGKGCYARYKYKKKKRKYMRKYRYGKKYSCCRYYTYCCGTRKYKSYGYYKKMRYMRRW
            ***  .  :  ****** .******  ****   ** *   :******  *** ******

46          TRKGWMRYRYKYKKCYATGKWKSYWKGWMMMRKRYMRKTRKYKKRCMCMWMYKRMMWMRR
48          YRKRWMRYRYKYKGYYWYKKWKSYWKTWMMMRGAYMRGCRKYTKRMMACTMCGRMATCRR
             ** *********  *   ******* *****  ***  ***.** *.  *  **   **

46          KYMMKYYRWWTCRYRYKRRKGYGTCCTGACTYTRMKKKRRYKTKWKRMRKCRKWKMYYYY
48          KYMMKYYRWWGARYRYKRRKKCRCTYYTGTCTARMKKKRRYKYKWKRMRKYRTWKMYYYT
            ********** .********        .   :********* ******* *.******

46          YYSRRAWWYSKTTMMYKKRYYCGYGKKKKKSRRYYYGMRYYTRRMRRKKWRWTTYKKMSY
48          YYSRRWWWYSKGKMMYKKRYCMTCTKKKKTSRRYYYAARYYKRRMRRKTTGTYYYKKMSY
            ***** ***** .*******     ****.******. ***.******.     ******

46          YMMYRSMMMCMGTGGCRRKRYSKGATYYAKWKWWKKMGGRCKCTYWWWYRMRRRMRRRCR
48          YMAYRSMMMTCACKKMRRKRYSKAGCYTTGWKWWKKMAARMKTCYWWWYRMRRRCRRRTR
            ** ******  .    *******.. * : *******..* *  ********** *** *

46          YRTKGKYYRWRKKYMWWRWKYCTKRKKRGAYWKKKKKRAAAYYKKKSKGKYWTKRAYYYR
48          YRWKRKYYRWRTKYMWWRWKYYYKRKKGAGTWKKKKKRRWMYYKKKSKRKYWKKRGYYYR
            ** * ******.*********  **** .. *******   ******* ***.**.****

46          KYKRMKYWYMRYYTWYRRRWAAARKCYRKKKKMKKRMRYRYTCRKKKMGYRKKKKRWYTA
```

```
48              KYKGMKCTTMRYCAWYRRRWGGGAKTTAKKKGMKKRMRYRYGYRKKTMKYGTKKKRWYKM
                *** **   *** :******... *   *** *********  ***.* * .******.

46              KYGKWRRKKYMKAKRKSKRRRWMYKYYYRSYYMRRYRYMKRSYKKMMGTACGATKMKYRM
48              KYRKWRRKKYMKRKRKSKRRRWCYKYYYRSYYMRRYRYMKRCTKKMMRWRYRWCKMKYRM
                ** ********* ********* ******************. ****       ******

46              WSRKKRWWAMTRKWYTCCKKKRAKRKCYMRRCSKKKYKKYRRYKWKWWKKMMKRACTGWR
48              WSRKKRWWMMYGKWYCAYKKKRMKRKYYMRRMSKKKTKTYRGYKWKTWKKMMKRMSWRWR
                ******** *  *** . **** *** **** **** *.** **** ******* .  **

46              RYMKYSKKRRRKKRYMKKRYRYMMKKKYYKSKWKKYKRKRWKRKKRKKYGTKMKMSGRKY
48              RYMKTGGKRRRKKRYMKKRCGCMMKKKYYKSKWKKYTRKGWKRKGRKKCTYKMKAGKRKY
                **** . ************   **************.** **** ***   *** . ***

46              KKKYYYKKKKMRSTCTTASKSRKYMRWGYRGKRYRKKRKRYRKYYAKRKYKKYRRRKKYK
48              KKKYYYKGKKCGSYYYYMSTSRKYMRTKYRKKRYRKKRKRYRKYCCKRKYKKYRRRKKYG
                ******* **  *     *.******  ** ************* .*************

46              KTCYGYYYWKYRYAKYKYRRWYRMKMYYYTTGGWKCCRCTRKRRACRRKMRKRKRRRWYK
48              KCYYKYYYWKYRYTKYGYRRWYGMKMYYYGAAAWKGAGTARKRRWAARKCGKRKRRRWYK
                *  * ********:** ***** ****** :..** .  :**** . **  *********

46              CYKRAYKKKYTAWYMMMYCRRTRYWMYKWKKMRKMCTTRRKYWRSWKKMKAGMGKRYWYW
48              TCGAGCKKKYYWWYMMMYMRRYRYWMYKWKKMRKATCARAKCAGGTKKMKGTMRKRYWYW
                    . ****  ****** ** ************   :* *   . ****. * ******

46              WYRRTRRKCARYYKKGAAKYYKWRRMYWGACAYYKWKYKRKKGRWAGTYKYCYKRRCYYR
48              WYRRARRKYWRYYKKRRRKYYKWAACTWKWYRTTKWKYKRKKRGWGACYKYTYKAAYYYR
                ****:***  *****   *****    *      ********  *.. *** **   ***

46              YMKKMWKYARYRGRMYMRGCRRSKKSKKKRYYWRKKATGKRKRCACKYWKKYRKWTTMSM
48              YMKKCTKYGRYRSRMYMRKSRRSKKSKKKRYCWRKKMGTKRKRYRYKYWKKYRKWWKMSM
                ****  **.***.***** .*********** ****   ****   ********* .***

46              RYRKKAAGGKWKTTKGWAKGYTRKMKRKRGYRKTAKRYYKYSGAGCKATCGGKYASATGG
48              RYRKKMMKKKWKGYKKWWKRYWRGAKRKGRCGKCGKRYYKYSACRTGGGTRAKYWSRYKK
                *****    ***  * * * * *  ***    * .*******..   .   .** *

46              RYYWYRKYWAKRYMAYKRRGTRRAATAAMYYYYSYKKGSYKAACKRKYRWMMRCGYYKRR
48              RTCTCGGYWWKRTMTCTGGKCGRTRYWRCYYYYCYKKTSYGCTGKGKYRWMMGAKYYKRR
                *      ** ** *: .     *:     ****.*** ** .: * ****** . *****

46              WRGAYYMMKGGCCWAYRSYTMYKYRGKRYWRATKCMTRRAYSRRARYYYTWYMRWCYSRY
48              TRATYYMMKAKAAARCRSCWMYKTRCGRYWRTYKAMKRAWYSRRGRYYYKTCCGTYYSGY
                 *.:*****. ..   **  *** *  ****: *.*.*  ****.****.      ** *

46              MMYRMRT
48              MMYRMRY
                ******
```

The pairwise alignments of two accessions 46 and 49

```
49                  WRWCSMRKMYWKRYWRSYKSRRYMWCRTKCAWSCCKTYWKKYWWMRMCYRRCTMYKRYYM
46                  WRWMSMRKMYWKRYWRSYKSRRYMWMRYKMTWSYAKKYWKKYWWMRMMYRRMKMYKRYYM
                    *** ********************* * * :** .*.********** *** .*******

49                  WKYWMRAYKARSMYRAAYYWKRYYRWMMWSRYKGASKYKKRYAKWMWRTKWRYSRGYYRK
46                  AKYTAGWYGRAGMYRWWYYWKRYYRWMMWCRYKARSKYKKRYRTWMWRCGWRYSRTYYRK
                     **    *   .***  ************.***. ******* .****  ***** ****

49                  WKRGTTAYRYMKKKMKMSYWWMMRTGWCMAYRWYYYYYSKYWRMMRTACWYKRSYYMMRY
46                  WKRAGCCYRYMKKKMKMSYWWMMRACAAMTYRWYYYYYSKYWRMMRYWYACKGSYYMMGC
                    ***.  .*****************:  .*:****************     * *****

49                  WRYWWYYAGGRRRKWYRMKYGGYRRYYCRWAWAYRYMYRKYWKWRWWKCYMKCYSYYSRK
46                  AAYTTYYTARRRRKWYRMKYKAYRRTTSGATACYRYMYRKYAKWRAWKMYMTMYSYYSRK
                      *  **:. ********** .***  .  : .******** *** ** **. *******

49                  TTCWATKSTYKRRMMYYARGTYGYRWYKMKTKTRKYWYKWWYYRYKYRYRTKWRWWMWKW
46                  CCTAMWKSKYKRRMMYYRRRATRYRWTKAGAKARKYWYKWWYYRYKYRYRCKWRWWMWKW
                          **.******** * :  *** *  :*:***************** *********

49                  YRKRCYYTTAMKYYWRWRRRSWRYWRWYYRRRSKYYYYYGWRRWYYKRYAKMYKMKWYRK
46                  YRKRMYYCWCMKYYWRWRRASWRYWRWYYRRRSKTCCYYRWRRWYYKRCRKMYKMKWYRK
                    **** **  .********* **************   ** ********  **********

49                  RRKYYMWRYKRRYWRWYMRYYYYMKRYMYGCKYRKYYWRMYCTGCWWYRMMKTKYAKKMK
46                  RRGTCMWRYKRRYWRWYMRYYYYMKRYMYKYKYRKYYWRMYMWRAWAYRMMKYKYRKKCK
                    **   ************************  **********   .* ***** ** ** *

49                  KRKYTCRKRWAWMRKKKKKRYAMKCYKKKKSYWKKSWYRYRKKRKKMMRRKKKAYKKKYC
46                  GRKYYMRKRAGWMRKKKKKRCCCKYCKKKKSYWKKSWYRYRKTAGKMMGGGKKMYKKKYY
                     ***  *** .********* . *  ****************.  ***   ** *****

49                  YYYRKAKMRYRRYCKYAMKKKRKWYMRYYYMTYRWMKMRRRKRRKMYYARGWRYYYAGYY
46                  YYYRKWKMGYRRYTKYRMKKKRKWYMRYYYMKYRWMKMRRRKRRKMYCTAAWRYYYTAYY
                    ***** ** **** ** **************.*************** : .*****:.**

49                  YTTACWATCYYMWKTCCKATCYGYAGRGGMYYKYWKYRYTTTRYAMKAKWMYCAGGGYYR
46                  YAATGATATYYCAKKTYKWYYYAYRKRRKMYCGYWKYRCCYWRYRMGRKWMYYRRATCTR
                    *:::  :: **  *.  *   *.*  *  **  *****    ** *  ****   .   *

49                  WMKAKYRYYWYWGGSSGKYRKKYSKYMCRMYYYKKKGYATTASYKYAAMYYKACTATKWW
46                  WMKRKCRYYWYTARGCKKYRTKYSKYMGAMYYYKKTKYRYYWSCTYRRMYYGWMYWYKWW
                    *** * *****  . .. ***.******  ******. *    * .*  ***      ***

49                  KKKYRYGTATRYYRKGYTAAAAGGTATRMTKRRRMKRWRYMWRWWKYRRKRKKYAKRRKR
46                  KKKYRYCACCRYYRKRCYRRWRACWRCRMWKRRRMKRWRYMWRWWKYRRKRKKYTKRRKR
                    ****** :. *****       .    ** ************************:*****

49                  MRKWYWMMRKSKKKRTGRWTATKGRYKKYKMKSRYMRRKKRYWYYRTYTAKCWRKGGRRY
46                  AGKWYWMMRKSKKKRCKRWYRYTKRYKKYGMKSRYCRRKTRYTYYGWYACKYWRKTTRRY
```

```
                  *************  **    . ***** ***** ***.** **  *:.* ***   ***

49              RYKKKKGCTGGATKGCYYRRKYMCYKYKWATKYCTRKKKKYSYYYRYKRKGARYRYYKKW
46              RYKKKKATYAARWKAGYYRRKYMYTTYKTRCKTMWGKKKKYSYCYRYKRKACRYRYYKKW
                ******.  ..  *. *******  .**   *    ******* ******..********

49              KWARGWSGTKMTTKWTYYYRYSRKRKGRRRAAKKKCTGRCKYAYRKRATGKMRMWRACTC
46              KWRGRASRKGMYYKWKYYYRYSRKRKRRRRTGGKKGCRGTKYWYRKRRYRKMRMARTAAA
                **    * . *  **.********** ***:. **     ** ****   **** *:.:.

49              YRKKMKKYRWSKCMKKKRKRYKTGYRMYRTMMKKTCRAGRKTRYSMKSMRKGMSWRYKYK
46              YRKKMKKYRWSKYMKKKRKRYGCAYRMYRWMMKGWYRTRRKKATGMKSMRKKMSWRYTYK
                ************ ********  .***** ***   *: **.  .****** *****.**

49              MWTCYGTYGAMYKKGTTGRYKRRSKWYRRKYCKMCTYYCGGKRMGGGMMRYKRWGSYGYK
46              MWKYYRYYRRCYKKAAYRRYKARGTTYRRKYTKMTKYYYCRKRMKSRMMRYKRWASTKYK
                **. *  *   ***.:  *** *.. ***** ** .**   *** . *******.*  **

49              RYYKAATGMRYGKWCGRWYWYYMKWYRKTAKKRRYRRYRYCWYKACKRRWSMRYRGMRRK
46              RYYKRRKKMRYKKTTKGWYWYYMKWTRKYRKKRRTRGYRCYATKTAKRRWSMRYRKMRRK
                ****   .*** *    ******** **  **** * **    *:.********* ****

49              WYYTYRYYMKKYRMWYYKMRKTASWRRYWCTAYRGAWCCAMACTCTKKCYYRGYRRRYRY
46              WYYCYRYYMKKYRAWYYKMRKYMSWRRYWYCGCRSMWATTAGYWSKGGYYYRRYRRRCGT
                *** ********* *******  ******  . *. *. : .  ..   *** ****

49              KAGYRKYRMRYRYKRYYRYKRKGYWAGMCKKKRKYRYRYRKYKKRKMRMKMYTAWKKMKK
46              KGAYRKCRMRYRYKRYYRYKRKRYWMRCMKKKRKYRYRYRKYKKRKMRMKMYGCWKGAKK
                *..*** *************** **    *********************** .**  **

49              TGSATCRKRYTKRYMRWKMRRKAAKRGGCGWWYKYRMGYGCTYGKYRKRCTKYYRCAYRT
46              CACTWYRKRYYKRYMGAGCRGKWWKRTAMCWWYKYRMTCATCYTGYRKRGCKYYRYMYRW
                 ..:  **** ****    * *  ** .  *******  .  *  ****  ****  **

49              WYYWMRTTYRTTGTTRGARRRYSYMMYKRKMCTKWKYWRKMRKTRKKYYKRKRAMKYWRM
46              TCTTMRAGTGKCAACRTCRGRYSYMMYKRKCMWKAKYWRKCAGWRKKYYKRKGRAKYWRM
                    **:   . .: * .* **********   * *****    ********   *****

49              KKMSMRRYRKRYKKRYKAARARYCWKWATGWMYYRMYGATYYYMKKASKKRYMRCCCATT
46              KKMSMRRYATGTKKRYKWMRRGCMWKWRKKWMCYRMYKWACYYMKKWGGKRTMRYYYWKC
                ******** .  *****  *    *** . ** ****  : ***** . ** **     .

49              ACTAKKYSYWYYRMRKTRKWYRRWYWKKRMTGKRTCTCKMMMKKKRTCRMRTYRWTCTKM
46              CTYTGKYSYWYCAAGKKGKWYGGTTAKKRMYRKRYSYYKCAMKKKRATAMAYYRWCACKC
                .  : ******    *. ***     ****  ** .  *  *****:  *  *** . *

49              CRWMRRWWWRTCYAGYARRKMWYAAGRYKWMKSYMSATYRRMYAKRYYKYCTYTKYRKKA
46              MRWMRRWWWRYMTRKCTRRKMWYMRKRYKAMGCYMSWACGRMYCGAYYGYSCCWKYRKKW
                 *********      :******   *** * .*** :  ***.  ** *.   *****

49              CCYKTKRRKRKYTKRKAAYTKRWRKTMYYYKKTMRTRKYRMRRSGKRYYTYKKGCSSWTK
46              TYYKYKRRKRKYCKRKGTTCKRWRKKMYYYKKYMRGATYRMRRSKKRYYACKKTGSSWWG
                  ** ******* ***.:  *****.****** **  .****** ****: **  ***
```

```
49    MGGTTYYMMYKYKKKYYYKRRRYKKMMAMYRKKYWMWMKKRCTKWWYMRYYAGYTTYKKT
46    MSRGCYYMMYKYKTKYYYKRRRYKKMMRMYRKKYWMWMKKRYCKTTTMRYYGRYYKYTKC
      *.   ********.************* *************  *   ****. * .*.*

49    CACTRAARKKKKTTKKCGAWGAMYRRWKRSATKKRKMWRKKRYKYMAGSYCTRAKRKAGT
46    TGMWGGTRKKKKKWKKGRRARMMYRRWKGSGCKKRKMWRKGRCKYMRTCYMWRGKRKWSY
       .   .:*****. **      ****** *. ******** * ***  .*  *.*** .

49    CCAAGRWYKCWWSTCYCMRSKWYRKYGTYTGTTTATKKRKWYKKTKTKKCKRRRRKRKKY
46    YTTCARWYKTWWSKYYYMRSKWCRKYRCYAACKWRYGKRTWYGKATWKKYKRRRRKRKKY
        :..**** ***. * ***** ***  *:. .    **.** *:. ** **********

49    RTARKKKKWKCYCMWCMKWGWTCTAYWYWMYYMKTRGCYMKKSMSYRTTCCYMWYCYGYM
46    RWWGGKKKWKYYAMTAMKWRWYYCTCWYWMTCMKCATYYCKKSMSYRWYYSYMWYMYAYM
      *    ***** *.* .*** *   : ****  **    * *******   .**** *.**

49    KKWYYRCGAKWRWRYWKYKACKWAMRKKAKKKKTTTWWKYAAGTATRKWGATGGAGRMRM
46    KKWYYRMRMKWRWRYWKYKRTKWRCRKGRKKTKAACWWKYGTACGARKWAGACRGRRMRM
      ******   **********  **  **  **.*:: ****.:. .:***..:  . ****

49    KMYYRCYYARTGAKKTKKRCGACYCCYYKKYAMRAKYKYAGWTKAAGGRWTKTWTRRTMK
46    KMYYATCTGRWAGKKYKKRMAMMCMMYYKKYRMRRKYKYTATAKMMKRRWWKKWWRRCMK
      ****    .* ..** ***  .     ***** ** ****:. :*    ** *.* ** **

49    KKMMKYRYWCATTKCTYSRYYKMMKKKWCKKYYWRKYMKTMRYYTKYSYYGGYSKKMGKK
46    KTMMKYRYWYWKYTYYCSRYCKMMKKGWSKKYYWRKCMTKCRYYYKYSCTRKYSKKMKGK
      *.*******  . .   *** ***** *.******* *.. *** ***    *****  *

49    YKKWRTTTYKRKYRTGYYRRMRRCYWWKKRRRGYTGCYARMWYKYWKYCRCGAKRWSAKY
46    YKKWRWCYCKRKTGCRCCRRMRRTYWWKKGRRRCYRYYGAAWYKYWKCMRTRGKRWSTKY
      *****    ***      ***** ***** **     *.   ******  *  .****:**

49    YKKGCWYRTWKKCAKRRRRYKKKKWWYYYKKWYCWKKKKRRCAGCKYYYYMARKTTTWTK
46    YKKTYWYGYWKKYWKRRRRYKKKKWWYYYKKWCMWKKKKRRMGRYKYYYYMTRKCWGWKK
      ***  **  ***  ******************  ******* .  ******:**   *.*

49    RKYRKRKYATRRYKWWSMYYKKKWKYRYKKAKKKRRKYCTTKKRMYGCYCKYWGGYKWRW
46    RKYRKRKYTKRRCGWWCMYYKGKWKYRYKKGKKKRRKYSAWKKRMYTACTKYWRKYKWRW
      ********:.**  **.**** ********.*******.: ***** .  ***  *****

49    RRGRKYKRKMMRWRWSWKAYMRYYKTRYSYYRYMYRYAARKCKGKMKSYGGGYTACKYSK
46    RRKRKYKRKMMRWRWSWKCYMRYYKGRYSYTRYMYRYGGRKSKKKAKSYRRRYWRYKTSK
      ** ***************.****** **** ******..**.* * ***   *   * **

49    YRYYYACGGCSKRCAKYRYYAKWYWMYYCTCARGYTYKKKKAACGGKWKKKCAYRWKASK
46    YRYYYTSACTCKRYMKYRCYRKWYWMYYAAYCRKYYTKKKKWRYKKKWKKKTTYRWKCSK
      *****:..  .**  *** * *******.: .* *  ****     ***** :****.**

49    YKRRGGTKMGYRRRSRWGAKYAYWMYRWYYRRCCGACWKKYGTGGCCGKRRAMWSKMRKK
46    YKRRATYKMKYRRRSRWKMKYRYWMYRWTCRRATTRTWKKYRKATTATGRGTCWSKARKK
      ****.  ** *******  ** ******  **.    ****  ..  .  * : *** ***
```

```
49    RKYRWRKRRYYKAAYAATTARWKCATTKGRYRYRMKKMRYYYMRMKWRYRKYKKKKATYY
46    RKYGAGGRRYYKGGCCWACMGWKTTGCGAGYRYRAKKMAYYYMRAKWRYRKYKKKKTAYT
      ***    *****.. . :   ** :   . **** *** ***** ***********::* 

49    MYKKYYYYRYYWYSWRTYRKRYRATCRGMYYWSKAMRKYKYYWWWRWKRKGCSYWYGYTG
46    MYKKTYYYRYYWYSWRYCRKRYRRYSRKMYYWSKWCRKYKYYWWWRWKRKRYSYWYTYCK
      **** ***********  *****  .* ******  **************  **** *  

49    GWRMYTRRMWMYWYSWYGTKKKKKKRYRRSMWRRKMKKKYSWWRYYKSRKGKRKWWKTRM
46    KWRMYYRRAACYWYSWYKYKKKKKKGTGRSMWRRKMKKKYSWWRYCKSRKAKRGWWKGRA
       **** **   ******  ******   ***************** ****.** *** * 

49    YKRMKGRRTKYGAYWAYTKCYRKKKGTTCCKKRYGTWTTGMKGGKRTRRWMYYKRKKKYG
46    YKRMKKRRCKYKMYWRYYKYYRKKKTAGGTKKRYRKWCAAAKAAKRCRRWMYYKRKKKYK
      ***** ** **  ** * * ***** :   **** .* :. *..** ************ 

49    AGWRKMRKKKRRRYYWMAWYSRKKYYMWKRRRKWSMKKRMYYCATACAKRRKGGMMYYRT
46    RRWRKMRKKKRRRYYWACWYSRKKYYMWKRRRGWGAKGRMYYTRCRYMKAAGKRMMYYRK
        ************** .************** *. * ****      *     *****.

49    YKMKRKWRWSKKRWAKKMKWKKYRAYKYKTKYKACATAWKRTKKMKKKGRYGYKGYKKCK
46    YKCKRKWRWSKKRWCKKMKAKKTRRYKYKKKYKRAGCWWGRCKKMKKTTGTKCGRTKKMK
      ** ***********.**** ** * ****.*** ..  * * *****.        ** *

49    RCCYSMKKKKKYYGCCGTTYCKYYCYYWKMTAWCGKKYYYRKKYCTRTYYYSGKMYKTTY
46    RMYTSMKKKKKYYTYGKYKYMKYYMYYWKMCGTYKKKTCCRTKYYYAWYYYSKKATGYGY
      *   *********     .* *** *****  .   **   *.**    **** *     *

49    GTRGRKKMYWRYRYKMYYRMRRCGMYRCGAWCTRKMAGTTYSKGRWGWCGKCCRGGKGKM
46    AARKRKKMYWRYRYKMYYRMGGTAMYRTAWWMKAKMRRAAYSKRRARTTRGATAKKTAKA
      .:* ****************   .*** . * . **  ::*** *      .    ..* 

49    KKAGGYRKYTATAMRYMYYKMRMYMMYRRYYCAYYTMYKRYKWMKMKWRRKSRKRKRKMY
46    KKRRRYRGCKTATAACMYTTMRMYMMTAGTCAGYYCMYKRYKWMKMKWRRKSRKRKRKMY
      **   **  .:::   ** .******     ..** ************************

49    WKSWCRGKKKRRKMKYRYCTGCGTKWGRKKKMTKKGYRRRYMWYMYACGKKKMRRGKKMY
46    WKSWSRKKKKRRKMKYRYYKRYRYKWKRKKKMWKKKYRRRYAWYMYWMRKKKMRRKKKMY
      ****.* *********** .    ** ***** ** ***** ****   ****** ****

49    YTTSTRYMMYWSKRRKKKKWGWGYYTRYMGMKKYTAGSRCAAATAYKRYYWKYRSSRMYK
46    YCCCGATCMYWSKRRKKKKWRAAYYKRYMRMKKYYRKCRYTCCATYKRYYWKYRSSRMYK
      *  .    ************  .**.*** ****   .* :..::***************

49    KCGAYKTASKGRTYTMMMWRMRYCYKYWRYRRRKYGKCKKRRWKKAGTYYKMKKRYRKRW
46    KYRMYKGRSKRACTCMMMWRMRYYCKYWAYRRRKYAGYKKRRWKKWCCTYKMKKGYGKRW
      *   **  **     ********  *** ******.  *******    ***** * ***

49    SRKKTRWMRKRWKRKKWYYYYKYKRSRKMKMYRMKMGRRKYAAACARRKMRRYMRKYKCR
46    SRKKCRTARKRWTRKKWCYYYKYKRSRKMKMYRMKCRRRGYMWRYGRRKMRRYMRKYKYR
      **** *  ****.**** *****************  ** *    .************ *

49    TGCAGRAYYSRYYTRRKTACYKRRKWYGRRTTYTMTKTWGTWCTTWMKYMCGYRMRWKMK
```

```
46              CRTRKGGYYSRYTGAGGGRYYKRRKWYARRYKYYACKKWKKWYGCWMKYMTRCACGAKMK
                      .*****        *******.** .*   *.* .*   *****       ***

49              TCAGKATYRCYGRTRYYKAKTCYYYWAATGKMMYMYGRSACYKYSKYTYKYYMCMRKWKY
46              GTWRGTYYRYTTGWRYYKRTKMYYYWRRYRGAAYMYARSGYYKYSTCYYKYYMYMRKWKY
                     : **     **** .. ****       ***.**. ****.  ***** ******

49              KYWWYKKMRYAKKGCSYYYSRKKYWRYRYMSWRRSKMWRMRKAYWMRRWMKRCTKMRMKR
46              KYWTTTKARYWKKRYSTYYSRKKYWRYRYMSWRRSKMWRMRKTYWMRRWMKRSWKMRMKR
                ***  .* ** **  * *************************:*********. ******

49              RTTAATATKCRTRRWKYYKKGRCCKKSMYRYTAGWGWAAKYMRTTCMMKRWYCRGSKKSY
46              RAWRRCRCGMRYRAWTCCKKRGMMKKSMYATYRKARWRRKYMGCATCMKRWYTRKSGKSY
                *:        * * *.  **    *****       *  ***  :  ***** * * ***

49              RYTCKKKCTKSKMMTCGAYACRMYMWYCYYGYCGSRWKMKYRGYYYYKKRCWKGKYRAMC
46              RYCTKGKTAKSKMMGTTRYTTAMCAAYYYCKCTRSRWKMGYRRYYYYKGAAAKKKYRRMY
                **  * * :*****    *:  *   * *     ***** ** *****  . * *** *

49              KRYMYYKYYRRKCYSKSAYWWYYYRWRKMRYYMYGCCTMKTRTTCCRCG
46              KATMYYKYYRGKGYSKSCYAATYYRWRGMRYYAYKYMKCGCRCATTRTA
                *  ******* * ****.*   ***** **** *   .   * :  * .
```

The pairwise alignments of two accessions 46 and 50

```
46              YWKRKMRWGRKYRRYCTGARGCKYMYMKWYKWYGCCRRRYRYKGGRTKYRRWKRSKYRTY
50              YWKRKMRWKRGYRRCYWRWRTMKYMYMKACGAYAYGRRAYRCKSARGKYRRWKRSKYRWY
                ******** * ***     *  ******    *.  ** ** *..* *********** *

46              RMTSYRYKACRRYMRYYMRCKWRGCYRRKWKMRAARMYRYRMSYWWMYYRYMTRKCCTYY
50              RMWCYRCTRMRGYARYYMRYGWRKYYRRKWKMRTGRMYRYRMSYWWMYYRYMYRKYYWYY
                ** .** .  * * *****  **  ********:.***************** **   **

46              YYYKMRYYRRGACMRTWGATYKYRRKWYTGCMGATGCYMKYRRMWAKGKKSRRYWKWMKA
50              YYYKMRYYRRTGGMRGWRGYTKYRGGWYCTMMAMAATYMKYGGATWGRKKSRRTWKWMKT
                ********** . ** * .  ***  **   *. :. ****       ***** *****:

46              CCCKYTGGAKRKRYRRRMRRRMMMYMYKYTYAACKAMWYYWYRYMGKYSMRKRKRRYYTG
50              YTATYACTRTGGGCRRGMGGAMMMYCYKYACGGYKWMWYTWYRTMRTYSMRKRKRRYYCR
                  ..*:     .  ** *   **** ***: .. * *** *** * .***********

46              CCKYSRRGATYRCRMRWYYCYAGYYRGYYWWRMWWWWRWYKKYYMRYRRTATTKMYYRWM
50              YTKYSRRAGATRTRMRWYYYYWAYYRRTYWWRMWWWWRAYKKYYMRYRRYWCKGMYTAWM
                  *****..: * ****** * .***  ********** **********    .**  **

46              RRYWCYKGCRRYRKYCYMMYRYTYYWRKYMTGAGRTYMRWMMRWRYYYRYWKSRMKRYWT
50              RGTAAYKTTRRYAKYMYMMYRYGYYWRKYMARTCRKYMRWMMRWRYYYRYTKCRMKRYWY
                *   .**  *** ** ****** *******: : *.************** *.******

46              TKMMWRYYRSYYYYYRWWKSWKYSKYAYKYRKKMKACYMKKCGTKSYWYKAYAMTAYYMS
50              KKMMWRYYRSYYYYTRWWKSWKYSTYRTGCRKGMKWMYMKKYKYTSYWYKTCGCKWTTMS
                .************* *********.*    ** **  ****   .*****: . .   **

46              WRMYYCTKYWKRRWKWWRSYKWYWRKYTSWMKAYYMYKGKAAMGAAYYYKWMYMWWKKMK
50              TRMYYGCGYWKRGTKWWRSYGAYWRGYCSWMKRYYMYKRGGGASRRYCTKWMYMWTGGCK
                 ****   ****  ******  *** * **** *****  .. .  *  ******    *

46              GGYMKMWKKYTGCYTYSRKYMYSYRKWWSRAYRKWWKKYKKRRYRRMTYCTWKKAGCRYK
50              KTYMGMWKKYGKSYGCGRKYMYCYRKWWGARTGGWWKKYKKRRYRRMGCTAWKKMATRYK
                  ** *****  .*  .*****.*****.     *************   :*** . ***

46              YKTWYKKRYYYRYKKCGGGRTWCRRYRKKMYRYKKTAKRRRRKMMGRKKRKMYRKRAWAR
50              YKWWYKKRYYYRYKKYRRKRWWMGGYRTKMYGYTKCRKARRRKMMKGKKRKMYRKGTWMR
                ** ************    * *   **.*** *.*  * ******  ******** :* *

46              MRMGTRYRSSYRYYYRYYGWWGKKRYGGRKTRCYRRRKTCTTRCTKKYYGKMRKGMRTRW
50              MRMRGGYRSSYRYYYRYYKATAKGRYKKRGKGYTAGAKYYKYRGAKKYYTGMRKRCRWGW
                ***   ************   .* **  *  .     *   . * :****  ***  *  *

46              RKRAYYYAMYKRRKTMWKKRKYGRTCRCMKCYWRRKWMKKAYCCRWAKTRMKMMRKYWWT
50              RKRRYYYMMYKRAKYCWKKRGYRGWYRTCGAYWRRKWMKKCYTYGARKCRMKMMRKYATC
                *** *** **** *  **** *    *   .*********.*     * ********

46              ARKKMRGTRRYMSCYTKRKTASAMKWYRKKKRWRGRWKMRGSAWWKKRRKTACAYRKKKY
50              TRKKAGRYRRYMCAYKKRGCTSGMKWYRKTKAAACRATMGACGWAKKRRKYRMRYRKKKY
```

```
                    :***    ****..*.**  :*.******.*    * .* ...* *****    ******

46                  YKRWTACATRRYRWSMRKRRSMRSAYYYCMRKRKMKYYKKKYWSWKWKRKKKYWRKATYK
50                  YKRWKWYWAARYRWSMRKRRSMRGRYYYTMRKRKMKYYKKKYWSWKWKRKKKCWRKRYYG
                    ****.   : *************. *** *********************** ***  * 

46                  YYCGTKARKMRRKWCYYRCAKRKYKKMGMKYRYKRKKRRRKMRRWKRYSWWRRYWSAARK
50                  YYTTAKGRKMRRKWYYYGGCKRKCKKARMKYRYKRKKRRRKMRRTKRYSWWRRYWSWWRK
                    **  :*.******* **  .*** **  **************** ***********  **

46                  KKKRAYWRKYRRKKKKWYMGMAACCMMKKWTAKMKWKWTACKRKYWRKMKKAKRAYRCRR
50                  KKKRRYWRKYRRKKKKWYMRMCCGTMMKKTCGKMKWKTYTYKRKYWGKMKKGKRRYRYRR
                    **** ************** *..  ****  .*****  : ***** ****.** ** **

46                  KMWMSYMKYWACRCCRGKRRKKMRYAMATTRTGTKAKRRRRKYKSKMMRYRKWKRWCGTG
50                  KMWMSYMKYATYGAYGKKRRKKMRYTARAAGCAAGRKGGGRKYKSKMMRYGKAKRWAKWR
                    ********* :  .   ********:  ::  .:  *   ********** * ***.   

46                  YSWRKRMKKKYYACGMCRMWCMYCCGRYGCRCAGKRYKGKYRKKKKTGMMWKYYKRYMYK
50                  YSWRGRMKKKYYWTKMYRMWYMYTATRYRMRMTAKRYGTKYGKKKKARCMWKYYKRYMYG
                    **** *******   * *** ** . **  * :.***  ** ****:  ********** 

46                  WKKKCRWKYYGRKYARKKSYRMYRWKAGGYWMYRMRKGKRKRRKSYRSKMYRGCRRKRYR
50                  WKKKMGAGCTRGKYRRKKSYRMYRWKRKRYWMYRMRKTKRKRRGSYRSTMYGTARRKRYG
                    ****        ** ***********   ******** ***** ****.**  .***** 

46                  RMYTWYKYYCKKWKMTTRWKKAYWWWYRRRRRRRRWYARRWSRYWRRYYRTKGKKKYRKR
50                  RCTYWYKYTMKKWKMCCRWKKRYWAWYRGRRRRRRWYMRRTSRYAARCYGAGAKKKYRKR
                    *   ****  *****  **** ** *** ******** ** ***  * * : .*******

46                  WKKKYCKYWMWKKRKYRKKGYKYKCTYCKGYKMRYRYYYKKKWKSYWWMMMRWSTCAMRK
50                  WKKKYAKYWMWKKGKYRKKKYKYTTWTTKACKMRYRYYYKKKTKSYWWMMCAWSGSMMRK
                    *****.******* ***** ***.    *. *********** *******  ** . ***

46                  YKGKMGRYATRKMYTCGTYRRRYRKRYYWWWYYRTYKGWYYRKMTCYRRRKKKKWMRRWK
50                  YKKTMRATWYGTMYAARCTRRRYRKRYYWWAYYRCYKKTCYRKMYAYRRRKKKKWMRGWK
                    ** .*      .**:.   *********** *** **   **** .*********** **

46                  MYYKYYSRKAAGCYWWYSYMMMAGCRYYKRKRKWYYWMRYMCKKKWRWYAKKGAMAKGAT
50                  MYCKYYSRTCWTTCWWYSYMMMGTARYYKRGRTWYYWMRYMTKKKWRWYMKKTGMCKTRK
                    ** *****..    ********. .***** *.******** ******* ** .*.*  .

46                  CKYCCSWTMYKCYYRATGCYCWCGKCSGAGRRYYWRYYWKRKKKYGCMRRWKMTGGKCCC
50                  AKYTTSACMYGAYYRMAAACYTYRKTCCGKGGCCTRYYWKRKKGYRAMRGTKMWKAKYMM
                    .**  *  ** .*** :..     * . .      ******** * .**  **  .*   

46                  YACCYYSRCTKRGGTGCKKRRYYKRWYCAAARYMTCRYKKRKKRGTGSMYRRYYRWKGRM
50                  CTYYYYSGMYGRKTCKAGKRRYYKRWYMRRGGTCCAGYKKRKKRKCRCMYRRTYATTRRM
                     :  ***    *    . *********   .    . *******   .**** *  . **

46                  MKKRYGKRYMAACGRKRYGRKKKYRRRKYKCTGGRRRMWYYYYKYRTGTKAYMKWYRMMY
50                  MKGRYKKRYMRRYKRKRYARKKKYRRRKYKAAACARAMWYTTCKCRCACGGYMKWYRMMY
                    ** ** ****    ****.***********.:.  * ***   * * .  .*********
```

```
46          YYWWKWKTCKYRMRRKRKRKKWYRKRKYKWYKWKMMYYWKSKWKWCGSKMKYYRATTRAR
50          YYWWKWKAGKCGMRRKRKRKKWYRKRKCKWYKWKMMYYAKCKWKATRCKMKYYRCKKGMR
            *******: *  *************** ********** *.***   .******...  *

46          YSYKWMYSKKYYYKRCSCCAAMKCWCCTRYTYSWATGGYWYYKYYWAGCMYGRYMYKRYR
50          YGYKAMYSKKYYYKGTGAGWWMGYTTTCACCTSTRYKKCTYYKTCTRKTMTRRTACTGYG
            *.** *********  ..   *          *       ***      *  *   . *

46          RGATATYKKAWMYYAAAYRMKCGRMYYMMYYRTGTWTTRWMYRRMSYWRGSARAYKYYWC
50          RKTYWWYKKGWMYYRWMYRCTYRRMCCCMYYRYRYTGYRWMYRRMSYWRSSMRRTTTCTT
            * :   ***.****   ** .  **   ****      ***********.* *  .

46          RRCYCATRYMRYCCAYYTAKYMMCMRRYGAACCKATKMKRSYTTSARKYYTAKWRMGMKT
50          GGYCYWGAYMRYTMWCCCCKYMMYMRRCRWRYMKRYKMKRSYWYSCRKYYWGGWRMRAKY
                    ****      .**** ***      *  ******  *.**** . ***  *

46          CRKYRMYKCAGAMGTGYSRRMMRMMMYRRWCWCGTGKYGCTKARYYKGTAKCYWRRMYYK
50          MGTCGAYKTTTTACAAYGGRMMRMAAYRRWYWAACKKYKSWKWRYYKAAGGTYWRRMYTK
              .   ** : :  :.*. *****  **** *..  ** . * ****.:.  ****** *

46          WYYSRTRYWCCYTWARRWKWRRKYCCCKKSSYWKWKYKKAMYRRRKWKWWSYKKKKKGYT
50          ATYSRGATTTSYYTGRRWGWRRKYYYYKKSSYWGWKYKKWMYRRRKWKWWSYGTTKKAYG
              ***     .*  .*** *****   ****** ***** ************ ..**.*

46          YTKKTGYRTCAWGRYYGCKTWWSMKSRAGTTACSRKRGYYYGRRKRKATKYYWYKKRWRY
50          CYKKWKCGWYWARRYYRYKKTACMKSRWRGAGASRTRRCYCRARKRKGCKYYWYKKRWRY
              **         ***  *.  .****   :..**.*  *   ****. ***********

46          GTKKTKWAGRKRKAKAAMYYTWKAGYRKMKKKYTWRTAGAYKRRYWRYMKRRYKRYKKMK
50          SYKKYKWTTRKRKMKRRCYTCWKGKYRGCGKKYKTRKMAWYKARYWRYMKRAYKRYKKMK
            . ** **: **** *   *  **. **   ***. *. . ** ******** ********

46          RYRYWKKCKYRKKRWYKKKRCGGMYRKTGKSWYWGYWMYKYYRAAYYKRKMKKYKKMKGT
50          RYRYWGTYTYGTKRWYKKKRTRAMYRKCAKSWYWATWMYKYYRTGYYKRKMTKYKKMKRW
            ***** . .* .********  .**** .*****. *******:.******.******

46          ACGAYTMCASRWKYRMWKKGYAWRWYYKMKSWYRMMRWRRRAYYARRTTKCKTWYKKTRA
50          RYRWYCCMGGATKYRMWKKRTGWATCCKMKSTTGACGTGGAGCCGAACCGTKCWYKKKRT
                *   ..  *******  .*    ****          .  .      * ****.*:

46          GRRWSCKGYKYRCCYYRGKKYAGKKKWKRTRKRMRRKMGTGGYCYWMKKRCCACTGMRGK
50          KRRWSGKSYKYRTAYCRKKKYGTKKTWKGGRGRMRRKAKWRSCTYTMKKRGARTCACAAK
             **** *.**** .* * ***. **.**  * *****    .  * **** .   .  .*

46          KKRRTACKKMKRCTYCGTKKWRCYSTYYGCKKRRYKMYRTTYYKYRYRKKMYKRGMWRCA
50          KKRRYWYKKMKGYCCYRYKKWRTTCCCTAMTKGRYKMYRYYYYKYRYRKKMCKRKMWRSM
            ****   ****       ****  .   . .* ******  ********** ** ***.

46          ATYRYMKTTARYCGTWTYYGWTGCYSYRRCKMYRKSGTTCAAGKMKYWTMKWWYMYRCTA
50          MWYRYCGGGGRYAAGWGYCKWAAMYSYARTKMCRKCAWWYRMRGMKYTAMKWWTCYATCG
              ***    .**.. * *  *:. *** * ** **..       *** :****  *   .
```

```
46              CTCRYMRYGYMRMRYYRAYRRYRRYKRAGCGSCRRWMKMKAYYCYAAYSRMYRKKKKKKR
50              TAMGTMRYKYARMRYYRTYRGTGAYKGMATTGGRRWAKMGWCYTCTTYSRMYRKKKKKKA
                 :   *** * ******:**    **  .  . *** **   *  ::************

46              WWWYGMKARCYRWRKYWKGRAACGTGTKRSYKMMWMKMYKRTACCKMYRKKRKMKKYMYK
50              ATTTCMKGRAYRWRKYWKAARCYRCCGTASYGMMWMGMYKRCTTTKMYRKGRKMKKYMYK
                     **.*.********.  .     . ** **** **** :  ***** *********

46              KYRMWRKKAGTWRAGCYWKWRCAYWRYRKYAMYRSWWKRKRTAYAAWRYGYSMYKKKTYG
50              KYRMWRKKGRGWRGATYTGTRTCYWRYRKYRMYRSWWKRKRKRCRRWRYRYCCCKKTCYR
                ********.  **.. *   * .******* **********.    *** *.  **. *

46              CGMYWMKYMKMMWWSYRWRYRRKYYAYYKRYWGMRYYKRRRKRRYCTKKKYKYAYKRKYT
50              MRMYWCKCMGCCWWSYRWRYRRKYYGYYGGYWRMRYYGRGRKAGTTYKKKTKYGTKRKYY
                  *** * *   *************.**  ** **** * **     *** **. ****

46              AGRWKKWYKYWMARRKWYKKARKYYKCCGCYRCGGRRYRMRYRYRRWAGRSTAKAAYTKM
50              RKRWKKTYKTACMRRKAYTKCRKCTKTAAGYRYRRRRYRMRYGTAAAGSRSKWTWRCYKM
                  **** **    *** *.*.**  * .. **   *******     ..**. .    **

46              RKWAYRMKKYMSMRKMYRYYWKAKYKKWRWKKRKKTTGMRRSKKKAWYRYMGGGTTGKGC
50              RKWRYRMKKYMSMRTMYRYCAKWGYKKWRWKKRKKKKKMRRSKKKTWYRYMRRRYWSKRM
                *** **********.****  *  ***********.. *******:*****     .*

46              RRKGKCYKWWWKMRMKMMKRACTGWMRRMRYWYSKWYKKKKYKYRMKKRYRGCAWKKTAC
50              RRKCKTTKWWWKMRMKMMKRMSWRWMRRCGTAYSKWYKKKKYKYRMKKRYGRTWTKKYCT
                *** *  ************* .  ****    *****************      ** .

46              CKSTCKWWKYKKYKKKRKKTSSRMCRTACSKRKMKSYKKYKYKKCCAGAGMRMCKSYYCC
50              GKSYYKWWKCKKYKKKRKKCSSRMMGYGTGKRKMKSYKKYKYGKYATKTRCRAMKSCCYT
                 **  **** ********* ****   . .************ * .: :  *  **

46              TTRSKKRYKKKKWAKCKKKKWMMMYYKRRRSRRYWKKKRYYWKKKAGRARRRYGRMYKYW
50              CCGSKKRYKKGKWTKAKKKKWMMMYYKRRRSRRYWKKKRYYTGKKWRRWRRRYKRMYKYA
                   ******* **:*.*************************  **  * **** *****

46              KRYKKKCRYGRKKWRRRKMYRKGRTAAYCKWYCGRKGKMARKSMWYKKCRRWCYGCGTMR
50              KRYKKKYRYRAKKAGRRKCYRKKRKCWYYKWYMRGKCKMTRKSMTTKKYRRWGYRYKYMR
                ****** **  **  *** *** *.. * ***   * **:****  ** *** *    **

46              RKKWYKGYRWKWYRGRKKTAGGRYRKRTTYYKWKYYYYKRKWKMYMRRMKKYWYWRMRYW
50              RKKWYKKYRWGWYRKGKKYWSKRYRGRKYYYKWKYYYYKRKWKMYMRRMKKYWYWGMRYW
                ****** *** ***  **  . *** *. ************************** ****

46              YYYTWYRYWSKRYKYMRRAKWSYYRKKYKAWYKYRRKAKWKRYKYYYRWYMRTYKKKYRY
50              YYYCAYRYWSKRYKYMRRMKWSYYRKKYKMTCKYRRKMKWKRYTYYYRTTCACCGKKYRY
                ***  ************* **********   ***** *****.****       *****

46              KRRYKWYYARKKYKTRYRCWTGRTKKKRMYRRCRGYGKMYRYYRMCYARTTWTARACKCG
50              KARYKATCRRGKYTWRYGYWWRRCKKKAMYRRTRKCRTMYRYYGAYYGGAGWAGGWTKYR
                * ***    * **. **  *  * *** **** *   .*****   *. : *:.   *

46              CCATMACRGTTATYAACGACGKATTRTWTTTKKYWRCKGKCCARMGTATWYSSCTMWYYM
```

```
50            YYWYMCGGKYKMATGCMCCMTGGGCRWTCKYKKYWRMGRGYMMAAAARGWYSSMCAACCM
                  *.    . : ..  .   .  *   . *****         .:  ****      *

46            CRMMRWRYWRAMGYY
50            YRMMRWRYWRGCRCT
               *********.
```

The pairwise alignments of two accessions 46 and 51

```
46              AKYYSKKYWYMRATMYSYRYRTMWATCMYYWATACYYRWWGKTGYSYWWYWRYYMYMYSR
51              GGYYSKKCWTAGTACTGYRYRWMWGCTMYYWGCTYTYRWWTGKSYGTATYWRYYMYCYSR
                . ***** *   ::  .**** **.  ****. :  ****  ..*.   ******* ***

46              RGGGWWATTGKYGGCGWRRYYYWKTRWYYYWYRRYMKSWKTAYWMKKWMWCGCCGGYCRK
51              GRRRWWGGCTKYKKMRWGGYTYWKYGTTYYWYRRYMKSWGWRYWMKKTAWTATMATTTRK
                    **.   **    *  * ***    ***********   *****  * .  .   **

46              KTTYYWRMRWYMRGKSAMKWKWMWRYWYKYYRRRYRSTTYRYKRKTRYYRMRCWAYCRKR
51              GGACYWRMRTCAGKKSRMKWKWMWRYWYGYYRRGYRCKWYGYTRKWRCYRMRYTGTYAGA
                  : *****     ** *********** **** **.. * *.** * ****  .

46              YTYCCMYMRYCCCYRYWRTTYTRKYWGWAGWWRWYKRWRKTCMCRYYSRKMAGTYRRRKW
51              YKYTYMYMRYYYTYRYWRYAYGRKYWRWMRTTRWTKRWRKYYMYRYYSRKMGAYYRRRKW
                *.*  *****   ***** :* **** *    ** *****  * *******.. ******

46              KRAGCYKKRRYYYYSKKYWWRMTGYMKWYRYMKRATYYYYRYRMKYRMGARYRYYTGRYW
51              KACAGYKKRRYYYYGKTTWWRMWTYMTACGYMKGRWTYYYAYRMKYRMKTAYRYYAAGTA
                * .. *********.*. ****  **.   ***    *** ******* : ****:.

46              RMMKKAYKYRMYRCYWYTTWRKMYKRWWSMSRRGYCCWKSYCYYMRKYRRGWSYRMYKRK
51              RMMKKWYKYAMYRAYWYWAWRKMYKRWWSMSRRTYMTAGGTYCCCRKCARRASCRMYKRK
                ***** *** ***.*** :************** *    .     **  *  * ******

46              YMKRRMKYYWAYTTCYYWMMTYMSWRTYRTCYWRMRTGKWYCWSSYYCCYCGRKKKGAGR
51              YMKRRMKYYWGYGAYYYWMMCYMCWRATAWAYWRMRKRKWYTWSSYYTTCTAGKKKRRRG
                **********.* : ***** **.**:   .*****. *** *****    . ***

46              AKTYMKKKRRWYMRMMRKYGYKRRSRGRAGTYKYWKYYKKSYRRRYGTTYYRKRYRKKKR
51              RKCYMKKKRRWYMGMMRKYKYKRRSRKRTAAYKYWKYYKKSYRRRYTCACCAKRYRTGKR
                 * ********** ***** ****** *:.:***************  :   ****. **

46              KRCKRYKKWRWKKRRRMYRAKKMACARARWRMMACYRYMCYRCKKTTAYRRWKSYKSYKR
51              KRYKRYKKAGWKKRRRMYRRKKMTTTRWRWRMMGTYRYMYYRGKKYCGCRRWKSTGCTKR
                ** *****  ********* ***: :* *****. **** ** **  . *****  . **

46              YKMKTRRRYYMSRWWCMMAYRACACWYYAKARTYWRACKGTAAAYRMATAGCTYCRGYST
51              YKMKCRRRYYAGATAYACMYRWYMYWYYGKCRCCARWMKTCCWGCRMMCGAGAYTRAYSW
                **** *****  .       **    ***.*.*   *  *  . . **  .. :* *.**

46              CKMRTGTAYCARGGKGGTTRRWCMKKKRYWSKYRCKRYYAMRMMRWRYRAMGYYRAGYGT
51              TKMAGACTCACGKKKRTCGAAWAATKKRCTGTCATKAYYCCRAAATACAGCRCTRRACKW
                 **  . : ..   *      *. .***  ..   * **. *       .    * .

46              SYWYKWRYCACMSYMGWRMKYCWYRMKATTYYRYRYRYYGGYSRGKKRAKWRYYYSTCYS
51              SYWYKWRCGCTMSYMAWRMKTTTCRMKGWKYCGCGYRYYTTCSGRKKRMKWRYYYGCAYS
                *******  . ****.****    ***. .*    ****   *  *** ******. .**

46              WMKAYRAATGYYMKKTRMRMRYCCCRRMKYATATGRYTYSARYRYKMYSTTAWWKSRAYR
51              WMKGYRMTGCTYMKKKAMRMGCTTTRGAKYMYMYRRYGCGGRYRYKMYSGCRWWKGARTG
```

```
            ***.** :   ****. ***     *  **     **  ..********   ***.

46          WWWTRMRRKCCKYRRYKWARMYMYWWWWKYRMSKKKRYYKWKTYYYYYKKRWRKGCMMKK
51          WWWYRMRRKYMKYRRYKTWRMYMYWWWWKYRMSKKGRYYKWKWYYYYYGKRWRKAAMMKK
            *** *****  ******  **************** ****** ***** *****..****

46          AAYKWRRYRRMKYYYGRYWSKTTCCTYRKRKKWATTYRKWYRKYKAWYRTCARRKCTCTA
51          GTCKWRRYRGMTYYYKRYWSKCYTACYRGAGTWRYWYRKWYRKTKMTCRATCRRKYGTGR
            .: ****** *.*** *****   . **   .*   ******* *   *: .***

46          AGTTYYYYRKRYTGTRARKTTTYRTTAKGKYRGRYKKMKYCGTKCGTMWWRYCRYSTYYG
51          GTCCTYYYRKRYYAGAMRKGCCYRCGGGKKYRRRYKGMKYYRYKAACCAWRYTGTCCCTA
            .    ******* .   **   **  .  *** *** ***   *..   ***   .   .

46          GWKKMRTRYRKRWTCMYRGYWRYRKKRWYYYKCGRKKRWAKKYWSRMYMKWYKRYRKKKW
51          RTTKMRWRYRKRTCAMYRRYWRYRGKRWYYYKMARKKRWWKKYWSRMYAKWYKRYRKKKW
              .*** *****  .*** ***** ******* .***** ******** ***********

46          AGCKYYYRTGCKRRMRRCYYRKKKKMMWKMKCKRMRRMYRRSYKKKKKRAYWWRRYYATT
51          TCYKYYYGCAAKRRMRRYYYGKKKKCMTKMKSGRMRRMYRRSYKKKKKRGYWWRRYYWYY
            :  ****  ..****** ** **** * ***. ****************.*******

46          AGMRGRKWMKKYYRKKMKMKRYKRSRYRKKKWKTTKKRYTRWRRKKKRKYSTCRRWWWRY
51          MRARKRKWMKKYYRTKMKMKRYKRSRYRKKKWKGYGKRYWAWRRKKKRKYSYYRRWWWRY
               * *********.******************   ***  **********  *******

46          RYKSSWYKRAAAKKRKRYWWKKTKKYYGCMWKYYSRCAMYYKYRYYGTKRSCAMKKAGRK
51          ATKSSWYKRGGGGKRKRYWWKKWKKCCCTAWKYYSRMTMYYKYRYYRKKRSMMMKKRKAK
              *******... ********* **     ****** :******** .***  ***   *

46          YRKRKATYYWAYKKRWKAAKKRKTCAKRGYKSRMMYRYRKKMWKTRRMRRWGRYYKKMRR
51          TRKRKGCYYWTYKKRWKMRKKGKKMTKGRYKSRCMYGTGKKMWKCRRMRRWARYYKKMRR
             ****. ***:******  ** *. :*  **** **   ***** ******.********

46          RSKKWYRYMWKGWRYAGAWTTKWWSRWTTCCGYSWACTGWMWYMYSKKKRYYRKKRMTCC
51          RSKKWCRYMWTKWRYMACWCCKWWSRWWKYYKYSWMGAATMWYMTGKKKRYYRKKRMWAY
            ***** ****. *** ..*  ******  .  ***  :. **** .***********  .

46          KSKWKWKYKKGAYYRYRKYYRKMKRKKTGYRRGMAKKYRRMRSYRYAACWYKYKMRMKCT
51          GSKWKAKYTKRCYYRYGKYYRKCKRKKCKCAARMTKKCGAMACYRYCCATYKYKMRMKYC
             **** **.* .**** ***** ****      *:**   *  .***... ********

46          TAAKSRYRRCMYRKYMMKMKYKGCKKRRYKGRRRYKMMTRYKRWCRKKRKTCTKMKKWGY
51          CCMTSRYRRACYGKYMMKMKYKSYKKRRYKRGGGYTMMCRYKRAMRKGRGCTYKCKKWRC
             . .*****. * *********. ******    *.** ****  ** *    * ***

46          KYKYYMWKKYGCGRRKTAMMYRCYYKGMWKKKKMYCRWKYYCGRKYRKRSYRGRYWRMSR
51          GYKYYMWKTYRYRRRKAWMMCAMCYKRMWKKKKMYMGAGCTTKGKYRKRSYRRGTAGMSR
             *******.*   ***: **    ** ********         ********     ***

46          WKKMRKRRGMMARKRRYRRRMKRKKATCGRYYWRCSCCRYYYWYKYGRTGSYYGAYTKYR
51          WKKMRKRRTCCWRKGRYRRRMKRKKGGTKRCYTGTGAGRYTCTYKYKACACCCKTYKKTG
            ********    ** **********.   * *  ..  **   ***   ..   :*.*
```

```
46                  MWMRCYSYYMMYYRWYKKRCYGRMKGYRMYGMYYRGYSKMRCMAGTMRYSKYWKRKATCR
51                  CAMGAYSYYMMYYRWYTTGTCRRMKRYRAYSMYYRRYSKMGTAWRCCGYSKYAKRKTATR
                      * .***********..    *** ** *.**** ****        **** ***:: *

46                  AYRRYKKYARYRWYMKRRKKKKMKMRYYWYGGMKKCAKMRYRRKKMRKAKKWRKKCGKKK
51                  GYRRYKTYWRYRTYMKRAKKKKMKMRYYWCAAMKKYRKARYRRKKMRKMKKWRKKTAKKK
                    .*****.* *** **** *********** ..***  * ********* ****** .***

46                  SKWMGYGMYKKKYYRYYYYTKKTKKMMGTACGARTWCKAMTYRMWSWRWYCRYRKMKYSR
51                  SKWMATKMYKKGYYRYCYCGGTWKKMMRWRYRWRCTSKWAAYRMWSWRWYMGYRKAGYSR
                    ****.  **** **** *   . ****      *  .*  :*********  ***  ***

46                  SYCRMYYYWWRRKCATGCTRRMKWKKGCARAYMYKTMTCYCTTKTTGTRYCTYRRWYATT
51                  SYTRMYYYWWRRKMRKSMAARMKWKKKTGGWCAYKWCYYTAGYKWWRWRYMCTRAATRWW
                    ** **********  .. : ******  .    **     .  *    **   *

46                  RRMMKRWWYRTWRYKYYRKWWRRRRWGSMYRKSWRYYRYWRYYTRYWYAGKYKKACMACG
51                  RRMMKGAAYGWAGYKCYRGWAARGAARCMTAKCWGYYRYWGYYGATTYGTKYKKGTMGTA
                    *****   *    ** ** *  *    .*  *.* ***** **    *. ****. *. .

46                  WAKKKYYYKAYKYYKTWYKRYYYYKTCKKKAAKKAGKKKRKACMMRKKRKGKYRYYCRKS
51                  ACTKKYYYKRYKYYGAWYGRYYYYKYYKKKCGKKWRKKKRKGSMMRGKRTRGYRYYARTG
                     ..****** ****  :** ******  ***..**  *****..*** **.  ****.*..

46                  YYCMRSSRMYRSSTCGYAYTGYKYYMMRYRCCYYCRYSKRYGAKRRYRGRRKTGCTTRRC
51                  CCYMRSSRMYRSSYYRYGYYSYKYYMMRYRMYYYTRYSKRCRCKRGTAKGAKYKYKYRRG
                       **********   *.* .*********  ** *****  .**      *   . **

46                  TKKYYMRRTAAAGYKRCKMKRKRKKKKRCAKRATRGATMMMMRWRRRYYARRTTKAKKYRT
51                  AKKYYMRRWGCTTYKRTKMKRKRKKKKGMRKGTYGCGAMMMRWGRACCGAACCGGGKYRY
                    :*******  ..: *** **********   * :   .:***** *   .     . ***

46                  AKYYCKKGCKYYRRYYWAAKMTWMYTTCMRCWRAKKKMWKMRRRRKYKKAGTRKYGKWRC
51                  GTYYGKKKTKYCGRTTATCKMYWMYACAMRTWRRKKKMWKMRRGRKYKKRRYAKYRGWRY
                    ..** **  **  *   :.** ***: .** ** ********* *****    **  **

46                  GRCSKMRTYAASTTKMGMYGRKKRKKGCKRGMYTKMMMYRRGTRYMRKKYRSWKRCAAGK
51                  AGTSKMACTTGCYGGMKCYKRKKRKKRYGRKCYGTAAATRRRYRYMRKKYGSWKRYRMRG
                    .  ***   :..   *  * ******   *  * .    **  ******* ****

46                  MMWTGCRKMAWYYRKYRYRKATYYRMWAAYKTTGKAYARSKWSYAGCYKYKKWYACYYGR
51                  MMWCATRKMWAYYAKYRYRGMKYYRMWMRYTCCAKTYMRSKWSYCRYCGYKTAYMACYKG
                    *** . ***  ** *****  .*****  *.  .*:* ******.    **. * . *

46                  RYYWCCCWKRKKKRKYKWTRMRTCKYYTWCRGKRYYRYYGGKARSYWGGTRKMRMTGGAR
51                  GCCTMMMWKRKGKRKYKWARMGWMGYYKWMRKGRYYRYYAAKGACYWSAYAGMRMYRCTR
                           **** ******:**    **.* *  ******..*. .**..   ***   :*

46                  SYRSMWRYAGCCGSMMATRRKRYRAGYKYAAKKYTWWYKKGTMRMYKMKYWWYRRMKRTC
51                  SYRSMTACGAYMTGAMTGRRKRYRRRYKYTTKKYCWWYKKKYMRMYKMKYWWYRRMKRYM
                    *****   ..   . *: ******  ***::*** *****  ***************
```

```
46          TAGRWWRYYYKYYRMTWKKYRWRWKYKYYRKTGCKMYKKYRRKKGKCYGYYRWMRYYKRY
51          YRRRTAAYYYKYCAMGWGKYRWRWKYTYYRKCAATMTKKYRRKGRKYYKYYRWMRYYKRY
               *   *****  * * ********.**** ...* ******  * * ***********

46          KRYMKRYKKRCCKKKYYSKYRMYWRKMWYYWMGATGGRKKYKYYKCGSWTGYTTRWGRYC
51          TRYMKRYKKRTYKKKTCSKYRMYWRKMWYYWMRWKSARKKYKYYKAASWWRCYARWKRCT
            .*********  ***  ***************  ...********..**    :** *

46          GGYRRYRYTRYAKRRSKSYATKCAAYTKYGRKKKMYRYYRYTYRYYKKYYKWMMCCKYAC
51          AAYRRYRYKRYRKRRSKSYTYTSTGCCKYARKKKMYRYYAYYYRYYKKYYKWMMTTKCGT
            ..******.** *******: ..:.  **.********* * ************  * .

46          CMRRKYMCCTRKTATGYKCRMGMWMRRKYKYGWWGCWYYRRGTTKYRKYYRKTKWKRKMR
51          TMRRKYMTGGRKGCGAYKTGMKMWMGRKYKYTWWCYAYYRRKCYKYRKCYAKYKWGAKCG
             ******   ** . .**  * *** ***** **   ****   **** * * **  *

46          RKMYYYMRCYYSRKRWWYSYMMKKKKKTCACRYYYGKKYRKRGCKGTSRRYYGATAYYWM
51          GKMYYYARYTTGGKRWWYSYMMKKKKKYYMMRYCCTKKTRKRTTTKGSRRYYRTCRCYWM
             ***** *   . **************    **   ** ***  .  ***** :   ***

46          KRAYKYGMKKMTAGWKYRYMSRRKGRWRGCGRCAWTMRAKMGRKYMRRAKKKRYCYYTYR
51          KRRYGYKMKKMYWAWGYGCMSRRKRRWGRAARYRAAMGRKMKRKYARRTKKKRYTYYCYA
            ** * * ****  .* *  ***** **  ..*   :*  ** *** **:***** ** *

46          MKMRRWRKKCKKWMRSWWRKMYAGGRKGWWSRYKAGTCKKYYKRWTACACKRYSWSAKTY
51          MGCGRWGKKYKKWMRSWTRKMCWKSAGRATGRYKMRCYKKYYKRWKWYCYKRYSWSTKCY
            *   ** ** ******* ***   .     .***    *******.  . ******:* *

46          RSMRSKWAYGRWKMYKYTRYRGTTCACRKRMKKMRKYYKYMCYRKRKRKRKKYRRRKKYW
51          RSMRSKATYRRWKMYKYWRTAKKWYMYRKRCKKARKYYKYMYYRKRKRKRKKTGGGKKYW
            ****** :* ******* *   .    *** ** ******* **********    ****

46          YKYRYKRRKYWYRMMYYYRTWRKKRRAKKMRKRWKRRRRYCYKRAYKWYYYRYRKKKMRK
51          YKYRYKRRKYWYRMMYYYRYWRKKRRWKKMRKRWKRRRRYTCGAGCKWYYYRYRKKKMRK
            ******************* ****** *************    . **************

46          YATKYWGTTASKWKWCSKKAWRRKRTTSYCCKMKCAAKYSRMYRKKYKKYRRKYYMRRKK
51          YGCKYWACCCCKWKATSKKMWRRKRKKSYYYKMKTTTKYSRMYRGKTKKYRRKYYMGGKG
            *. ***.  ..***  *** *****..**  *** ::******* * *********  *

46          RWWYMCYRWKKCGWAAKRSYYKYCKGMYYCAWTACCKMYRCRYKWKMKKKMMKRTTMKKS
51          ATTTMMYRWGKYRAGRKASYYKCMKKMYCTGWYWYYKMYRTRYKWKMKKKMMKRKKMKGS
                * *** *   . * ****  * **  .*    **** *************..** *

46          WRTYWRYRKTACRSWWCKYKRKKYRYKAARKRMAYWKARYSMYKKYKYTYYMAGMGKYMR
51          WRKYWAYRKCGYRSWWTGTKRKKYRYKTTRKRAMCATGRYCCYTKTKYKYTCMACAKYMR
            **.** *** . ****   ********::***    ..**. *.* **.*   . .****

46          MYKYGWWSYW
51          MYGYAWWSYW
            ** *.*****
```

The pairwise alignments of two accessions 47 and 48

```
47              RRGGKWYTMWKKCARWKRRKAGWYMYKCRYYRARWCYRYRRATYYRYWKMYKRYKMYSRT
48              RRRAKWYACATGYRGWKRRKCATCMYKYGYCRTGTYYRYRRGCYYRYWKMYKRCKMYSGY
                ** .***:  .    *****..  ***  * *:   *****. ********** ****

47              YYGCWAMKYRKTTAGRKRWYYYYKRYWMWRYWKYWKGCTAYARMWYYGRWYKYGTCCMYW
48              YYKYATMKYRKYWWRRGAAYYYYKRYWMWRYWKYWKATWMTRRMWYYRAWTKYAAATAYW
                **   :*****    *   *****************.     *****  * **.:.  **

47              WMKRAYRKTMCAARRYRASTRKWRYSRRYYMRYMRWKRRCCWWCGCRWYKKRYSMYRMYR
48              WMTGCYRGYCGRTARYRMSWRGWAYSRRYYMRYMATGRRYYWTYRMAWYKKRYSCCRMYR
                **. .**     : *** * * * **********   **  *     *******  ****

47              WRRMYRYSYYYRRYYRRWMRYYYMGGCYYRGRWKYRRCACYMRARASSCGMMMRKRMMMT
48              WRRMYRYSYYYRRCYRRWMRYYYMTRMYCRRRWKYRRYWTYMRWARSSTTMMMRKRMMMC
                ************* **********   * * ******   ***   **  *********

47              GYRCKWYYRYYMRCWWRRSRKWYKWYYKWMKSYWYKRWRRRWRWTRRYMYKRKYYWYAMR
48              RYRMKWYYRYYMAAWWRRSRKWYKWTTKWAKSYWYKRWRRRWRAWARYAYKRKYYWYGMA
                 ** ******** .***********  ** *************   ** ********.*

47              RRYYRRYMKKWWYRKYRWWTYARWYRRYWGGSYYYYYWTRMKYMYWYRRKMYMGRYMKRW
48              RRYCRRYMKKWWYRKYRWWWYRRWYRRYTKTCYYYYYWWRMKYMYWYRAKMYMSRYMKRW
                *** *************** * ******   .****** ********* ****.******

47              RRYRRGWKYYGKKTACCMKKRKKRYYYYSWRRAYYYKYYRYWYWKYRTGTGGRYKKWKTY
48              RATAGKWKYYAGKWWAYMKKRKKRYYYYSWRARYYYKYYRYWYWKYRKAYSTGYKKTKYY
                *     ****. *  . **************  **************.. .  *** * *

47              KKYATAKRMYTKMKKMKMRYMKYACYYKRGKYKKTRMYKRSYKRCACCATTYWTTRMGWK
48              KKYMAMKRMYYKMKKMKMRYCKYRYYYKRKKYKKKRMYKRSYKRYWSYMAAYWWARMTWK
                *** : **** ********* **  **** ****.*********  .  ::** :** **

47              KRYYKKYYAKRRKYGCAKSYWKKMKRYYWAMRYYRRRCYKYYSKWYYKYMKAATTWYWTA
48              KRYYKKCYGKARKYRYWKSYWGKMKRYYAGMRYYRRRSYKYYSKWYYKYMKGTGKWYWYR
                ****** *.* ***   **** ****** .*******.*************.: .***

47              TYYCYRYRTTKRAYGWYMRWRRMTYKYKMYGTRYMYYSMRWYYCCYRKWYWKMYSYRRMR
48              KCTAYRYRYYKRCCKTYARWAAAWYKYKMYRYRYMYYSMRWYYTTYRKWYWKMYGCRRMR
                .  .****  **.   * **    ******  ***********  *********. ****

47              RRRYCYARYYSYTTKCCYKTGARSKKMRGSTRMYWKRKRKTRTRRKKYAARSWYGRYSYY
48              RRRYYYGGCYSYAGKYMYKACGACKGCGKSKGACWKRKRKYACRRKKYRGRSWYARYSYY
                **** *.  ***: *  **: . .*    *.   ******   ***** .****.*****

47              RYRKCYGKRCATRCTWYKGKYRYKKYWRKKGWYYYCRRRKKKMRYKTGYMKRMKCCRRRY
48              RYRGYTRKRYGGGTKWYKKKYRYKKYWRKKKWYYYMGAGKKTMGYTCTYMKRMGMTRRRY
                ***    ** .   .*** *********** ****    **.* *.  *****   ****

47              YRRYTTWRKYRSKYKKAKKRKKMMWGYTKWMRKYKMTYRKWKRRMKKYRWKMSRKWKRYW
48              YRRYWYWRKYRSKYKKRKKRKKMMWRCCKWMRKYKMKYRKWKGRMKKYRWKMSRTWKRTW
```

```
       ****  ********** ********   ********.***** ***********.*** *

47     WRYKGYKGGAGKRRKYRKYRSYYYKYRKKRYSKWRYRRWAMMRKRKKMTMMYYKRGTYAG
48     WRYKRCKRSMKKRRKYRKYRSYYYKYRGKAYSKWRYRRWRMMRKRKKMKMMYYKRRWYRR
       ****  * .  **************** * ********* ********.******  *

47     KMMMMMWKWRYKKKMRRGMKMWYYRRMRGRTTTGTKMRYKCWGGRYYYRCWMSKGRRGRY
48     KMMMCAWKWRYKKKMRRKMKCTYYRRMRKRGAYACTMRYKMWKARYYYRYWMGKKRRKRY
       ****  *********** **  ****** * : . .**** * .***** **.* ** **

47     YYKYRRKCWTAKKYWWACTCMMMTTACYYGCRWRMMMRKYTGTCGYYMWAWYYRSRKTYR
48     YYKYRRKTWKTKKCTWGGWYCMMCAGAYYRYRARMMMRKYYSYYRYYMWTTYYRSRKYYR
       ******* *.:**  *.    ** :..**  * ******* .   ****: ****** **

47     TRYYGRGRWRKRRRYYYKTYKKKKKRRKCSWYTCGMYYKMKRGKMRWYMMATYRRYWRMY
48     YRTYKRKAAAKRRAYYYKYYKKKKKRRKMSWYWMRCYYKMKRAKMRWYMMTWCRRYWRMY
        * * *    *** **** ********* ***    ******.*******:  *******

47     KMYYYYSKRYSYKSMAMRYYWYTTKKMKCKRYKYRWKRYRKKKYRYRMKSYSYWYWRTGG
48     KMYYYYSTGTGYKSMMCRYYWYCAKKMTTKRYKYRWKRYRKKKCGTAMKSYSYWYWRCTT
       *******.  .****  ***** :***. **************    **********

47     WGRMKYYKKKYMRKSYMKSRKKMRMRKWKRMRYRYKRWYYKKKRRWRWKMSMKWKYKKYW
48     AAAAKYYKKKYMRKSYMKSRKKMRAGKWGRMRYRYKRWYYKKKRRWRWKMSMKWTTGGTW
        .  ********************  ** ************************* .    *

47     MYYKRWWRYAKMCTGMYKKYKRYYWRKKKWWKARSWKMKRTRKACCKYRRKAAWTRCTGW
48     MYCKRWWRYTKAMCKCTTTTTRTYWRGKKWWKTRSWKMGRYGTTATKYRRKCCTGRYKKW
       ** ******:*      .. .* *** *****:***** *  .:. *****..  * . *

47     KRRSRRAWTCRRKYCCYKYKYKMKCGGRWKRRYRRMRKKKYARGYWYWKCKRYKTRYTTW
48     KRRSRRTTCTRRKYYACTYKYKMKYRRRWKRRYRRMRKKTCGGKYWYWKMKRYKKRYACA
       ******:   **** . .******   ************. .  ***** ****.**:

47     GMYRATGKRRGKYMMRYKMMYSYYSARYYKWKTGYKSWRMRYRKWRGTATTRKYYAGGGT
48     RAYRRGAKRGKKYMMRYKMCTSYYSMRYYKWKKRYKSWRMRYRKWRKAGAARKYYWARAA
         **  .**  ********  **** ******. ************ :.::**** . .:

47     KKRYYCAAMMYYYRYYKCGMRYSMKMRTRKKWKMTYKYKKRKRRGYKYYYMWTRKAKYKY
48     KKRYCYTRMMYYYRYYGYRMRYSMKMRKRKGWKMWYKYKKRKRGKYKCCTCAGRTTKYKY
       ****  : ********   ********.** *** ********  **      *.:****

47     WWWSCGRYRGTMMYRRKKRGCKYMRKKRYKKWKYYSRCKKYYMARAGKYGCRRKGRYYRG
48     WAWSTKGCGKWMCYRRKKRRMKCMRKKRYKGWKYYSRYKKYCMWRRRTTRYGAKKRYYRR
       * **       * ******  * ******* ****** *** * *  .     * ****

47     GYRMKTCKKKGRMKGRYKRWMMMWWRKRYCCTRMWYRCYCYYYGTYRKWMYRSRTRSMKT
48     KYRMKKYKKKCGMKKRYKRTAAMWWRKRTYMKAMWYRYYMYYTRWYRKWMYGSRWRSMTK
        ****. ***  ** ****   ******   . **** * **   ****** ** ***..

47     RYSMRKKRMRSKMAMWWYYRYYYRWTAYTRKKRYRYYRKKRRKSYKKRYKWYMYWRSRCA
48     RYSMRKKRMRSKMWMWWYYRYYYRWYGYYRKKRYRYTRGKRRKSYKKRYKWYAYWRSATT
       ************* *********** .* ******* * ************* ****  :
```

```
47          CMRWGKTKRKKKKWRSYWKGRCKWTCTRRKRYYGKWKCGKKKCCAKKKCAAWRRGAMRMY
48          TMRWKKCKRKKKKWRSYWKKRTKWCYGGRKAYYRKWKYSKKKMYRKKGTGCTRRRRMRMY
             *** * ************ * **    ** ** *** .***   **  .. **  ****

47          YKKKKTTYKWRKRGWTTTTRGRAKRYKRYYTKRGCKRSKCYKRWYWYKCCTARWWKWWKY
48          YKKKKAYCKWRKRRACCCCRAGWKRYKRYYYKRATKRSKYYKGWCWCTTTWTRWWKWWKY
            *****:  *****      *.  ******* **. **** ** * * .   :********

47          KRYRRMRYMRKWKKMWYRKRSKKWKRYRGYGYYMKTCGKTARMYCGMWKKWSRYKYKKWY
48          KRYRRMRYMAKWKKMWYRKRSKKWTRYRSYRYYMGGTRKCCAMYAAMWKKWGRYKYGKWY
            ********* **************.***.* ***    * . **..*****.**** ***

47          KYTMRTTAKWAKRKKSWYRKWCTTTKRWYGGYCWYRWKKWKYKRRRGYMKKKTCRRCCGA
48          KTKMRYYRKWGKRKKSWYRKATCKYKRWTAATTWYRWKKWKYKRRRRCMKKKGTRGGMKW
            * .**   **.*********   . *** ..  *************  ****  *

47          CGCCCSKKKYTYRRKRWKYWKWKYTKYWRYGCRRKMKRYRYRGCYGYSRMYTTCATYKYC
48          TRTTTSKKKYCYGRKAAKYWKTGCCKYWRCRYRRKMKRYRYRAYTKTGRMYCCMWWYKYT
                 ***** * **  ****    ****   **********.    .***     ***

47          GSWWKYYKRSRKKRCRRAKCYYYYCRGAGTKKYYRMRRKYKYRKRKRKYMKMRMAYSWMK
48          RSWWKYYKRSRKKRTRATKMYYYYYGTTTCKKYYGMRRKYKYAKRKRKYMKCGCRYSWMK
             ************* * :* ****   :  **** ******* ********    *****

47          MYSMWKMKKWMKYCWRRKRYTKRYYKWRRMMTTKRCCATCKMYCRYSYRKMYWYRTKWMS
48          MYSMWKMKKWMGCATRRKRYAGRYCKWRRMAKKGRYAMWYKMYMRYSYRKMYWYRYKWMS
            ***********  . *****: ** ***** .. * .   *** *********** ****

47          WYKYRYKTRAWTRRYYWYWYSKYYAGKATWWYWKYRKGGTGTKRMRKYCCTTAKWKTRAC
48          WYKYRYKGAGAARRYYWYWYCGCCCAKGAAWYWTYGKCAAACKRMRKYTTGGMTTKCRCY
            *******  . :********.   ..*.: ***.* * .:. ******     . * *.

47          CGRCRRWRYRRGSKRTCCAYYKYCKKGYAKGRCRWRTKTKSKMKWRKKGKKKKRYKKYYK
48          YRGYGGTRYRRKSKRWMYWYYKYTKKRYTKKRARARKKYKSKAKWRTKKKKKKRYKKYYK
                   **** ***    **** ** *:* *.* *.* *** ***.* ***********

47          KYYCKKKKRWYKRRRKRRYYKRRWSCRYTGAGGRKMRYRRWTMKRRYKMYRRCGKAMCGY
48          KYYTKKKKGTCKAGRKRATTKRRWSMRYACCAKRKMRYRRWKAKRRYGCTGAYKKWMGRY
            *** ****   *  ***   ***** **: .. ********. ****       * *  *

47          TCTMCCYKKKWRSKYMRRWRACRKCMTRRKWRKYTRKMAKRRYRKMACYWWKKKKCKKSR
48          GAGAAAYKKKTAGKYMGRWRWYGKTMYRRKWRKYYRKMRKRGYRKMGAYWWGKKKSKKSR
             .  ..****  .*** ***   * * ******* *** ** ****..*** ***.****

47          YRRKKYKKKKMRWYRYSMYRYTAKRKKYRKKRTCGKATRAMSGGGCWAGSKRKYYATYYY
48          YRRKKYKKKKMATYRYSMYRYWWKGTGYRKKRATAKGARRASRKRYTGASKRKCYGCYYY
            ***********  ********  * . *****: .*.:*  *     ..**** *. ***

47          KCYRMRRKKKKRRKKMWYYYMYMKWRYRWKYKRYTKYRYYKKYKTYYSYMRYKKYSYTKY
48          KYYRCARGKKKGGGKMWYYYMYMKWRYRWKYKRYCKYRYYKKYKAYYSTMRYKKYSYAKY
            * **  * ***   ******************** *********:*** ********:**
```

```
47              KKWMMKWSGCCMKRKYAGKKRYRWYYRMYKRYYYYYRWKWYRCYKWCMWKRKMRKKRKKK
48              KGWMMKWSAYAMKRKYTCGKRYRWYYRMYKRYYYYYRWKWYRTYKAYCWKRKMRKKRKKT
                * ******. .*****:  *********************** **   ***********.

47              MRKYRYYTAWYRTKYKRYKYKTTRKGCYKRKKMYGACAWYRRKYRWKCTCCRYMRWACKY
48              MRKTRYYYGWYGKKYKRYKYGYAGKKYYKRKKMYKWYRWYRRKYRWKTKYTRYMRWRYKY
                *** *** .** .*******  : *  *******    ********* .  *****  **

47              KYMYYKKRRKRCGCGRTTKGKKYRRKWSTYWWKKMMKKYRYYWKMRYRRCCTKCGKYRAT
48              KYMYYKGRRKRMSYAGYYGKKKYRRKWSCYWWGGAMKKYRYYWKMRYRRYYYKYAKCRGC
                ****** **** . .     ******** ***   **************   * .* *.

47              AKAWSMRKSRWRYWCTKKMYYRKKRMRRMAAMRMSMYKKRYYMRKKYWYTWAWAAYYYRW
48              GKCASMRKSRAGTAMCKKMYYRKKGCAGCWMMRMGMYKKRYYMRKGYWYCAGTTGYYYRT
                .*. ******      ********       ***.********** ***  . :.****

47              RCRMWMGKMRRYCCCYRWYWRMYMMTKWYYRKMMMMKWWKRMKYKYYGKWKYYYSYRYTG
48              GMGAWMKTMRRYYMTYRWYWRMTMMYKWYYRKMMMMTWWKRMKYKYYKKWKYYYSYRYCA
                    ** .****   ******* ** **********.********** ********** .

47              KKKKYRYKYYRWSRKRRKYYYRGCTGCTMWKRKWRRARWGCSYKKWRWRKRRRKKRKKYY
48              KKKKYRYKYYRWSRKRRGYYYRAAYRYWMWKRKWRRWRTRMSYKKWAWRKRRRKKRKKYY
                ***************** ****..    ******** *   ***** *************

47              KKRRKKYKKTKKYYCAAWWYRMYYMMRRWWSMAYMTTKYGACKKYKYKTRMYRYKKWYYY
48              KKRRKKYKKKKKCTYGGWWYRMYYMMRRWWSAGYMCCKYKGAKKYKYKGRMYRYKKWCCT
                *********.**   ..************** .**  ** ..****** ********

47              MRYRMKYAAAYYMRMRKYMWKAGWRMYRRKKWMRRGKTYYRKYYWGRRAKRMYYKWYWGA
48              CRYRMKYWRWYYMRMRTCCTKRRWACTRRKKWMRRAKKYTGTTCTRGRRKAMYYKWYWKR
                 ******   ******.   *  *   ********.*.*  .     * * *******

47              TRYGMYRYKKKKYRRWCRYYYYKKKMRYRYYKKWTCYRKKRWTKYAMYAYRYKRRRKWKK
48              KRYRMCGTKKKKYRRWAGYYYYKKKMRYRYYKKWYTYRKKRWWKYGMYWYRYKRRRKWKK
                .** *   ********. ****************  ****** **.** ***********

47              RYRTWMRRKRACKKMMMRKRYRRKKKYKYCKCCYWKMAAWMYKTRRKWKKKYGGGYYWCK
48              RYRWWCGRKGTSKKMMMRKRYRRKKGYKYYKYYYWKCRGTMYKWRRKWKKKCRATYYWMK
                *** *  ** :.************* *** *  ***  .  *** *******  . *** *

47              KATKYKKARRRTCCCRYYTYRYKWKKRKRRYKRWWWYKWWRKRKYYRRRKKRYYRRATCK
48              KCCKYKKRRRRYYYMRYYYYRCGWKKRKRRYKRWWWYKWWRKRKYYRRRKKRYYRRWWMK
                *. **** ***    *** **  *********************************   *

47              YRMYRWYKYYRKKRSKARCCRKYYYKRMKWMRKMYKWWKTCCYMYWYYKYRYRCTACYRK
48              YRMYRWYKYYRKKRSKRAMYRKCYYKRMKWMRKMYKWWKCTYTMYACCKYRYRAGTTYRK
                ****************    ** ****************    **   *****. : ***

47              RYRMRKRYYYYTKYYYARRTWAAGRWRYSYRRWRAYYRARWKYKARYTYAWTKSKYGAAT
48              AYRMGKRYYYYKKYYYWRGYTTWKAAATGYARWGWYYRGGTKYKRRYKCTACKSKYSRRY
                 *** ******.**** *   :      .* **  ***.  *** **. :  ****.

47              KWYYCGMKKKRRKRYKRKKKYKWMWGAAKKAWCKYYYYYTMKYKAWYYYGACCTSRRYRY
```

```
48          KWYYMRMKKKRRKRYKRKKKYKWMTSRRKKCTTKTYYYYKMTTKMWYYYKRATASRRYRY
            ****  ****************** .  **.  * ****.*. * ****  . :******

47          RKAMRKYRKKKWMAYRWTRWYKYRYRSMWRKKRKYRRRWWRRYKYKKMGAYRYWRYATCA
48          RKRAAKTRKKKWMTYRWCRTYKYRYRSMTRTKRKYRRRWARRYTYKKMKTYRYWRYMATW
            **   * ******:*** * ******** *.******** ***.**** :****** :

47          YRYKKKTCCWMKGTYWKYSMRSGKYYKTCAGCKRRMRWYTAGWTWKSYWRCACYRRSKWK
48          YRYKKGWTTWMKAYYWGTSMRGCGYYKCMTRTKRRMRTCCTATYTKSYAAGCYYARSKWT
            *****    ***.**  ***.  ***  :  *****   :.   ***   . * ****.

47          KMYAKGCARTSAGYGYRYYKGKYKRYRTCYKYMYAGAAAWRRYWWYRMYSKMRWMKCMYM
48          GMYMKTTTAGSGCCACRYYKAKYKRYRKYYKYCCMRRMRWRRYTTTRMYSKMRWMTYCTM
             ** *  :  *.  . ****.******. ***       ****   *********.   *

47          CGYTYKSKKKWKTRRYWRWMTYKKRGKCYRKYKSKWYAACCMYYYKRYKKKMKKTATRKK
48          ACYWYKSKKKWKWRRYWRWAAYKKGATTYRKYKSKWYCTYGMYYYKRYKKKMKKCRARKK
            . * ******** ****** :*** .. *********.:  *************  :***

47          TRAYKYYWYYYWKKYRRWRGAGYKRRAKGWRKWSWMMYRMRKKSYRRRKAWSKMRGRTGT
48          CATYGYYWYYYWGKTRGWRAMTYKRRMKRWRKWSWMMYRMRKKSYRRRKRWSKMRKRKAC
              :* ******* * * **.  **** * ******************** ***** *..

47          YACKGSMKKRMKKYKMYRKKTYWMYMYGYWYKWRKKRKKYW
48          YRYKRGMKKRMKKYKMYRKKKYWMYMYRYWYKWRKKRKKCW
            *  * .**************.****** *********** *
```

The pairwise alignments of two accessions 47 and 49

```
47          RTYYYWKMKRWTWMYRYRRKKKWYTWYRGTCYMAWAKWRARYRWYWKYRYRRRRMWTYWY
49          RAYYYWKMKRWKWMYATAGGKKWYCWYRKCTYAWTWTWRRRYRWYWKCAYRRRRMTACWY
            *:*********.***     **** ***   *    .** *******  ****** : **

47          MWRRYKRRSCWRYMKSWRRYYARWKGYWCAMKWMMKRWWKRYRRKYMWYYWRYGRWCGKW
49          MWRRYKRRSTTRYMKSWARYCTGTTKYWMCAKWMMKRWWKRYRRKYMWYYARYKGATTGT
            *********  ****** ** :  . ** . ******************* **

47          YYRKYYWYAKYYYYYYCARYRRASTRKWRCCARYYMRYRKKRWKYRRKTRRAYMYYYRYA
49          YYRKYYWCRKYYYYYYYRGYRGMSWRGWAYYWRYYMRYRKKRWKYRRKYRRTTATYYRYW
            *******  *******   **  * * *    **************** **:   ****

47          CYYKKKYRKCRYWRCCRKKRSGWGRWYKWYYKMYYSKMMKSYWRYKRMWKRKRMWYRCRK
49          YCYKKTCRKYRYWGYYRKKRSRWRRWYKWYYKMYYSKMMKSCAGCKRMWTAKRMWYRYRK
              ***. ** ***   ***** * *****************    ****. ****** **

47          RWRYRRYMKGMMWRWRYYMRSYYGCGRMYRKYWKRMMMYYWGRYYRMCTGGAAAYRKYMY
49          RWRTARYCTRMMWRWRYYMRSYYSYRRMYRKYWKRMCCYYWSRYYRMTCTAGGGYRKYMY
            ***  ** . *************.  **********  ***.*****   ....******

47          KRYYRRSMWAYKMKWRYYRMRWRRWYWRYMMWRYMAGWMYRMYWKYMYYWTCCWWKKGRK
49          KRYYRRSMWCYKMKWRYYAMRARRWYTRYMMWRYMGSAMYRMYWKCMYYWAATWWKKRGG
            *********.******** ** **** ********.. ******* ****:. ****

47          MYYRYMYKKRRWMMKYMMKYYYMRRRYRYRYYKYYWYRYKRRKKYGWMYYRKCGKWRMRY
49          CYYRYMYKKGRWMMKYMMKYYYCRARYRYAYYKYYWYRYKRRKKTKTCYYRKTAKWRMRY
             ******** ************ * **** **************    **** .******

47          YWRWCKKMMKYMKAWYRCGYGTTRKKRYRRYWMTKYACCYRKKCSMKSRMYTKMYYRKSK
49          YWRWMKGAMKYMGCTYRMKCAKCRKKRYRGYTCKKCRYYYAKKGSMKSRMTYKMYYRKSK
            **** *  **** . **   .. ****** *  .*    * ** ******  ********

47          RSRKYYTGTATKWKYKKYAKKYSYTWAMMMGRKKMWACTAGARGTGAYRMMTYCKAGTMM
49          RSATYYYAGTAKTTCKKCRKKYCYYWMMMMKRKKMAGTGRAGGAKCMYRCAWYYKMCCMM
            ** .** . ::* . **  ***.* * *** **** .   .. ..  **   * *   **

47          MMWACYRWWKWRMYMMTCSYKYYYWWKAKGRYCYAYSRWYCWRRRYWYWTKRTMGMRGRR
49          MMWRYYRWWKWRMYMMYTGTKYYYWWKMKTGCYCMYSRWYYTRGATTTTCKRCCTCRKRR
            ***  ***********  . ******* *      *****  *       **    * **

47          RKRKTMWSWWWSWMKRKWYKTYMKYKKRYYRKMMSKKTTKGTTTAWRRYCGYKARWKMYG
49          AKRGWCAGAATSWATAGWYKWYMKYKKRYTRKMMSKGGYGTAAGGWRRYTSYKRRWTCTK
             **    .   ** .  *** ******** ******     :: .**** .** **.

47          AGACYKSKMRTRKKMMKYYKKYTGTARCKRRRYRRRWRMGKYCWYWKGRCRCRSKYRRYY
49          MRRYYKSKMRKRKKMMKYYKKYCRYCATKARRYRRRWRASKYYWYWKRGYRYRSKYRRYY
                ******.***********   .  * ******** .** ****   * ********

47          MMYKRKRKGRCRYRKKYYGSMKRKKRYAKKKWGYAATGGCARYKRKRTCTRYWWYYRRRS
49          MMYTRGRTKGSRYRKKYYKSMKGKKGTRGKKWRYGWWRASWRYKRKRKSKRYWWYYRRRS
```

```
      ***.* *.  .******* *** **    *** *.   .. ******...**********

47    YRKRYYRKRYGYYCRKKKCRTKCKRMYRWTYKYAYRMMKSARCGYYYGKWMWMRRRRKYK
49    YRKGTCGKRYATYYRKKKGRYGMKRMYRWAYKYGYRMMKSRAMRYYYRTAMWMRRRRKYG
      ***    ***. * **** *   ******:***.******    *** . *********

47    WWKYMYTKKTWRKRYRCRYRMKKKRCKYGYYKKAATRYARWRYSTCRTWRMYYWYYRARW
49    WWKYMYGKKYWRKRYRAAYRMKKKGYKYKYYKKMRKRYWRWRYSYYGYWRMTYWYYRGGT
      ****** ** ******. ******  ** ****  .** *****    *** *****.

47    YYYRRYYYKGKKKWMRRKKKTCYKAYYKMKMRRWKYTRWRMYMYYYWKYSCKMWSCRACR
49    YYYRRYYYKKKKKWMRRKKKWYYGWYYKMKMRRWKYWRWRMYMYYYWTYSYKMWSYRRYR
      ********* **********  *  *********** **********.** **** *  *

47    YYKKKMKRYKYRWKMARYRKMSYSYWYKWKRYKKRYKCSYYMATCSRMRKYRYKRRMRKW
49    YYKKKMKRYTYRWKMTRYRKMSYSYWYKWTGTKKRYKASYYMRAGSRMRKYRYKRRMRKW
      *********.*****:*************.  *****.**** : ***************

47    RMRYKYRYRYWYKKWKTKKRRYWRWYRRCYTTCTYRKYKYYYKTYSKYKKWYMYYKRWRA
49    RMRYKYRYRYWYKKWKWKKRRYWRWYRRTYKGGCYRGYKYYYKYYSKYKKWYMYYKRWRW
      **************** *********** *.   ** ****** ***************

47    TYKMKKYYKYRTKYKYYTKYYWYMKTGCYYKYRRWRTGTYKRRTKMKCGGCTTGKAWSWA
49    YCTCKKYYKYAWKYKYTAKYYWYMTWRMYYKYRRWRCTYYKRRCTMKYRKMWCCTCWSWM
        . ******  **** :******.   ********   **** .**       ..***

47    MMMYMRKKKWWRRKWSSKRYRMTYMRARRMKKKRRMKTTKCKMYRYKKRMYMGCYWYSRR
49    CMMYMRKKGTTAAKWSSKRYGMGTCRRRAMKKKRAMKYGKAKMYRYKKRMYMRMYWYSRR
       *******     ******* *   * * ***** **  *.***********  ******

47    WRKKRKKAYKMCYTCGKWYWRYRSMWKKKKRRRMYRYKRKRYKKSYRRKKGRAYGKYCKK
49    WRKKRKKTYKMYYATAKWCWRYRCMWKKKKRRRMYRYKRKRYKKSYRRKKAAGYRKCYKK
      *******:*** *: .** ****.**************************. .* *  **

47    KASRWMRYRKRYWYRMYACTRWMYKKWKYRAMGMWRRRMKKKKWSRKTYYCCAYRRKKCW
49    KWSRWMRYRKRYWYRMYRYWRWAYKKWKCGMMACWRRRMKKKKWSRKYYYSYRTGAKKYA
      * ***************   ** *****   *. ************* **.     **

47    MKKYGKWRKKKTAATRYGYKARMMYYWYRYRSTYYAWRYWTGKGKAGTKRRRMRKCKWRY
49    CKKYKKWRKKKYRWYRYRYGRRMMYCAYRYRSACCCWRYWYTKRKRRAKAGRMRKYKWRT
       *** ******    ** *  ****  *****:  .****  * *  :*  **** ***

47    SSRCRTRKCGTYCCACGKYKYKRCRRYYRRKSSKGCKWRAKRRYKWMKRMYGKTWAGYRG
49    GGRSRWRKMRYYATTTRKYKYKRTAAYYRRKSSKKATWRGKRRYKWCKGCYKKCACTYRK
      ..*.* **   *. :  ******   ********.. **.****** *  * *  . **

47    YKYYGAYYRCYCKKRKRRRYYKRRWWYRKMWRYRWGCGAKRAYRKRYCAKKAKKCRKWMR
49    YKYYRMYTRTYTKKGKAGRYYKRRWWYRKMWRTGWRTCCKRGCGTRYSWKKWKKTRKTCG
      ****  * * * ** *  **************  *   .**.  .**. ** ** **

47    RGKYYRKCYRRYSMYGKYWGKTRMKYRYKMYKRRRYKYYCAAGKRWSGAACGKMMRKAYY
49    GKKYYRKYYRRYSMYSKYWKKKRMKCGTKMYKRRRYKYYYRMRKRWSTTRTTKMMRKWYT
        ***** *******.*** *.***   ***********    ****  :  ***** *
```

```
47    KMKYRYYYKKWRYRYSRYTTWMRKRRWKKKMYMKKKYRRKKKKKKRKTARMRKYWYWKAW
49    KMKYRYCTKKWRYRYSRYAAWMRKRRWKKKMYMKKKYRRKKKKKKRKYRRMRKYWYWKGW
      ******  **********::***************************  *********.*

47    KRRRRKWKKAKYRYRKKRMAAWKKKRRRRCWTGGMWAYWRYYMKYYTRKMMRCRRRYTTA
49    KRRRRKWKKGKYACRKKRMGTWKKKRRRRATATTCTWYWRYYMKYYYRKACRYRGATCGG
      *********.**  *****.:********. :     ********* **  * *     .

47    RSYAYGMRYWYMGGGGATATTCTYKRMRRYCCKKKWYYGGRMRKRWMKTRYWRYYYRYWY
49    ACYCTKCRYAYMKTAAMWWYYAYCKRMRRYMYGTKWYYAKRMGKRACTYRTWRYYYRYWY
       .*.   ** **  ..     .  ******   .****. ** **  . * *********

47    RYYKYTMRARKTGGTGTGCRYYYAYKYKSKATTCWMTMYMRYKMAAKKRRKYWMYKRGGG
49    RYCKYYMRGAGCKRGAARTRYYCTTKTKSKRWWYWCYMYCGYKMTGKKRRKYWAYKRASA
      ** ** **.      .:  *** : * ***    *  **  ***:.******* ***...

47    RCCKAATTMKKAYTYAAYKKRYRRKYYKWMYRWYKKRYGTGCRRYRSMKKAGYYYRKYKR
49    GYAKMRYYMKKMYCTGCYKKRYRATTYKWMYRWYKKRYRCCMGGYRSMKKTRCYYRKYKR
        .*    *** *  ..****** . ************      ******:  *******

47    RKKRCGGYWRRKYYRRYKYKMGYCGYYYRYKCWAGCYWMSRKYKKKWMMKMWKRAYTGAY
49    RKKRGCAYWRRKYYRRYKYGMATYAYYYRYGYACKMTTMSRKYKKKWMMKMWKRTYKSWY
      ****  .************ *.  .*****   .    ****************:*.. *

47    GGYWYGKRYYYWKYWMYRCARTCGWKWMRKKMSYRYKKAGAKMRKKRYKWYKKRYKTYKK
49    AKYWYKKRYYYWGTWMYRYGGAGAWKAMRKKMSYRYKKTACKMRKKRYKWYKKRYKAYKK
      . *** ******  **** . : .** ***********:..***************:***

47    YKRRRRKMYKKRWYKRWKTRYRKTYMYTYKARWYKKMRKRMWMMKTGGGKYYKYTTGCYC
49    YKRRRRKCYKKRWYKRWKKRYRTKYMYYYKRRWYKKMRKRMWAATAAAAKTTKYAGRSYT
      ******* **********.***..*** ** ***********  .:...*  **:  .*

47    YYGARKWCRGTAYKRSWGYYKATRRRKWKGWRWMMRGAWYWTYAYAYRMWACMCWGGTGK
49    TYRWAKWYRKKMYKRSWKYYKRYRRRKWKRARWMMRRWWYAWYMYMYRMWRMMGWSRYRK
       *   ** *  .***** ***  ******  *****  **  * * ****  * *.   *

47    RRRWCACAYSMRWKMRKYKWKAKKWTTSYWYKKYTKWRKKAAAYYGYRMRKKRKKKKRKR
49    RRGTTCTRYCMRWKMRGTKWKRKKTCCSYWYKKYYKWRKKGMGYYKYRMRKKRKKKKRKA
      **   .  *.******  *** **   ******* *****. .** *************

47    WMMWTMAYCGCGRTYYKRKYKWKCTCTKWTYKYARRMGGSRYKRKKRSCYGGWYKRCTWA
49    TMMAKMMTAAAARYCYKGKYKWKTGTKKWAYKTMRRCKKSRYKRTKRSTYARWYKRSYWR
       ** .*  ....*  ** *****   .**:**  **   *****.*** *. ****. *

47    TTGAWYRKRYRMAGYGSWKMKWKRRWKYYYKKYRKYKRMRRWWKRMRYYYRSRYYYKYRR
49    YWKRTYRKRYRMMKTKSWKMKWKRRWKYCTGGTRKYKRARRWWKRAACYYRSRYTYTYRR
           *******    ************     ***** ******   ****** *.***

47    YKRWWKWWWKYRRSKYATGYRYRRKKYKRKWYKRRTKTCKKGKKRRSRRMAKKRYKYKYK
49    YGGTTKWWWKYRRSKYMYKYRYRRKGYKAGWYKRRWKGYKKKKKRRSRRMRKKRCKYKYK
      *    ***********   ****** **  ***** *  ** ******** *** *****
```

```
47        KKKMRRWRKGGAGYKRWKKKYTCKYMWYKTAKKGTCCYGCRKKKKTKMKMKKRYKRMWWR
49        KKKMRRWRKRRRRCGRWKKKYYMKYMWYKKWKKAYYYTRYRKKKKYKMGMKKRYKRMWWR
          *********      ******  ******. **.      ***** ** ***********

47        RYMGCWCYKTGTYCYWRYYKWRSRTTKTKMKYTACRSKYMWMWWYYRYYYRWRKAKRKRK
49        RYMKTTAYTYAACACWRYYKWRSRWWKYKMKYAGTRSKYCWMWWYYRYYYRWRKMKGKRK
          ***   .*. .: . *********  * ****:. **** ************** * ***

47        KRGKYARKRRSKRYKKKCKRKWWKYMYRSCMKKKGMRTKRMKRSRCKACGTTCTMKGWAY
49        TATKYWAKRRSKRYKKKYGATWWKTMYRSTMKKKKMRKKAMKRSRSKGTAGWYCMKKWMY
          .  **  **********   .*** **** **** **.* *****.*. .    ** * *

47        KKYGWTTTRWMMKYRRMRKMRACKAWCSRWKCATKRYYKKKCTGGTKSRKGMRKYMYRYM
49        TKYRAGKCRWMMKYRRMRKMRGAKMATSRWKTTCKRYTKKKACCKGKSRKKAGGTMYRYA
          .**   . *************..*   **** : *** ***.    ****     ****

47        RKYRKKRTWGWRKGTASYAYMKYKKKKKKKKKWYCYCWYKRKKKYYCKKYGTYGYCYWSA
49        RKYRKKRWWKARKKWWSYRCMKYKKKKKKKKKWYMYYWYKRKKKYYYKKYRACKCYYWSW
          ******* *  **   **  ************** * ********* *** :    ***

47        KYYKKYRRYYYRRARRKKKKRKTYYWYCTRKGKRRKSKMKYYKYRYYRRRMWKRWRRKTM
49        KYYKKYGRYYYRGGRRKTKKRKYYYWYTAAKTKRRKSKMKYYTYRYYRRRMWKRWRRKYM
          ****** ***** .***.**** **** : * **********.*************** *

47        SYKGKWRYKYSKKKYGCAKKKKRKRKKKYKGKKKYTKWYWWSAAGAMYTCAAGKYKKKYKY
49        SYKKKWRYKTCKKKTRYWKKKKRKRKKKYKKKGKYGKWYWWSRTKWMYAYGGRKYKKKYKY
          *** ***** .***    ************ * ** ****** :  **: .. *******

47        KTYMMYRKYTKWTWRRRWAAKYYGAGKKMRMRKKMWRWWRMYSRRYSKTTCWTCKYRMYS
49        KKYMMYRKYKKWKTARRWWGKYYRWRKKAAAGKTCTGWWRMYSRRYSKKCYWYTKYRMYS
          *.*******.**.  *** .***   **    *.   ***********.  *  ******

47        WGWKRSYWSRKKRTKMRAAYCYYYYWKTYRKRYGKYKYRSKRYKRKKYRMRARYKYYRRR
49        WTWKRSYWSRKKRKKMRGTYMYYYYWKYYRKAYKKYKYRSKRYKRKKCGMRMACKTTRRR
          * ***********.***.:* ****** *** * *************  **   *  ***

47        MMYSWMMYWMWKKMWCWYCKKRYYYTRYRRKMKMYKYWYRKWWYKYRYKRGCGGYMYYAA
49        MMYSWMATWMWKKMWSWTTKTRYYYYRYRRKMKMYKYWYRKWWYKYRYKRRARAYACTGG
          ******  *******.*  *.**** ************************ . .*   ..

47        MRYYSYCYYCAACWAMYRTACRKYRYWTKYYYYRKMARYRYRWTGKMRGACAKMRWYYRK
49        MRYYSTATCYMWYWCMYRCTGRKYRYWYKYYYYRKMCAYRYRWAKKMAAGTRKCRWYYRK
          ***** .      *.*** : ****** ********. *****: ** ..  * ******

47        MYTMRRRRTAYYYKSYMYRSRTRKARKKYKYCGYRSYWYKWACGKRRKKKYGMRWWWRGR
49        MYYMRRRRCGYYYKSYACRSRKRKMAKKYKYTCCRSYWYKWWYTTRRKKKYKMRWTAGSG
          ** ***** .******  ***.**  *****   *******   .****** ***   .

47        GYYRRGKYYAKAMRYGKYKKKMYKKKKKTYYKYKYAKRMYRMYMAKKMRYMKKKRCYGGM
49        SYYGAKGCYTGGMRCAKYKKKMCKKKKKACYGCKYRKRMYGCTCGKKMRYCKKKRYYATM
          .**     *: .** .****** *****: *  ** ****    .***** **** *. *

47        RMYRSGYMRYYKRMKYKYTKWYKRKGGTSWWYKKAGKRYWTMRYYRYYYSKKYMGMCKKY
```

```
49                   RMYRSRYMRYYKRMKYKYGKWYKRKARKGWATKKMCGRYAYMRYYRYYYSGKYMRCYTKC
                     ***** ************ ******. ..*  **   **  ********* ***   .*

47                   YWGACGCYRRRSYRRYYKRKKRSKYMKKRKKMMWCKWMRKYKYKRKYWKRRMMMKMRGAG
49                   TTRWMKMYRRRSYRRYYGRKKRSKYMKKRKGAAWMKWMRKYKCKRKYTKRRMMMKMRRMR
                            ********** ************   * ******* **** *********

47                   RKWRSRSSKKKWKCMMMRYKGKTKKRRKKYYWRYYKYYKKYRRKRCGCKRCTCTGSKWYK
49                   AKWRSRSSKTKWKTMAMRYKRKKKKRRKKYYWRYYKYYKKYRRKRSKYKRYCYYSSKWYG
                      ********.*** * **** *.**********************.  **    .****

47                   KYRRRWCMYRRKRKKMMYYKRYMMM
49                   GYRRRWSMTRRKRKKMMYYKRYMMC
                      *****.* ***************
```

The pairwise alignments of two accessions 47 and 50

```
47              RTWRKRYRRWYRYWKCTYMWCCCTAYKWMKYRKKKKYRYGGGWMGAMYMRRRWMMRRKRM
50              RCWRKAYRRWYRYWKYKCMWMYYCWYGTMKYRKKKKYRYSARWMTWMYMRAAWMMRRTAC
                * *** ********* . **     *  ***********.. **  ****  *****.

47              RWRRYMYMWMRYWWKYYMKYYRYKKRRRRSRWMYKKRMTYWRRMWRYTGRWYYYYRYKYR
50              RWGGYMYMWMRYWWKYYMKYYRYKKRGRRGRTAYTKRCWYWARMWRYGTRWYYYYRYKYR
                **  ********************** **.*  *.**  ** *****  ***********

47              RWRRYWKYKWRYAGMKRWRWYYRYYYRRRMWRKRTYAARSRKWMTWYWWYYYYKMYMRCK
50              RWRGTAKYKWRYCRMKRWRWYYRYYYRRRMWRKRCCMRRSRKAMGAYWWYYYYGMYMRGK
                ***   ******. ********************    **** *  ******* **** *

47              RWSYAKYYWRYYMYKKYYMRYWYWKMWWKKYWRYYYMKGAKGKAYKMWMYWRRMYRRKMK
50              RWGYRGYYWRYTATKKYYMRYWYWKMWWKKYWRYYYMTRMKTKRTKMWMYWGGAYRRKMK
                **.*  *****   ***********************.  * *  ******   ******

47              RRWAWYRWYGRKCMRRYRASRKWRWYSRRWKWWRGWMKRWWRKYCCTYYYRKYYKYYYYY
50              RRWGAYRWYKAKAMGRYGMSRGWAWYSRRWKWWRKWMKRWWRKYAYWYYYRKYYKYYYYY
                ***. ****  *.* **  ** * ********** *********.  *************

47              RRRYGCKSWRTRYWKMGYRYYRRARYTYYYRRWRYYYYRYGKSWYKWMKRWGARYRTTRY
50              RRRYRYTSWRKAYWKMAYRYYRRCRYYCCTRRWRYYYYRYRKSACGAMKRWKMRYRYWRY
                ****  .***. ****.******.**    ********** **    ****  ***  **

47              RARYYKRWCAAKRWMKMTRYWYTYRGWYTYWKRRAMYCMRRRYYSGRYKRRWSMAAMKRW
50              RCAYYKRWARRKRWMKMGATTCYYRKATGYWKRGGMYYMRRRYYSRRYKRRWSMGGAKRW
                *. *****.  ******      **    **** .** ******* ********.. ***

47              KWRMSRKRTRRRMKKYWYWKWYKWKATYKATASKKKKWGKRYRKWTACWWRRKRKACKAM
50              KWRMSRKRCRRRAKGYWTTTWYGATGACKGGMSKKKKWKTRYRKWWRYAWAAGAKCAGGM
                ******** *** * **  .**  ..: *.  ****** .*****    *    *.. .*

47              YKCGGATRRRYKMRKKWRKMKYKKYWCCCRRRYYKYYKYYYMKYWWRYYYYMMYRKGRKR
50              YTMATCCRRRYKMRKKWRKMKYKKYWTAGGRRYYKYYKYYYMKYWWRYYYYMMYAGKGKR
                *. . . ******************* .  ************************    **

47              KTCCKYCRRRWKRWKYYSRKKYAGCKKAGKKGYKKRMKKMMKRWYYWWKTGTAYYAWMYR
50              KAMMKYSRRRWKRWKYYSRKKYWRAKKGAKKAYTKRMTKMMKRWYYWWTCRKCYYMWMYR
                *:  **.***************  .**..**.*.***.**********.  ..** ****

47              SRKKCRKKYRYKAGACRKGMRKGRYKRRKSYYKKKYAYRKWWKMRYWKRKKMCTMGRWRY
50              SRKTAAKKYRYKMRWYRKKMRKSRYKRRKSYTKKKTRCRKWTGMGYWKRKKMMYMTRWRT
                ***.. ******    ** ***.******** ***   ***  * *******  * ***

47              SSWCGMYWYYKRCGYRKMKYKYTAATTKYACSMKMYYRRMRRMRMSKYKRGTATRTGWSK
50              SSWYRMCWYYKGYRYRKCKYKYYCGKCKTGTSMKMYYRGCAGARMGKTKRAGTARWAACT
                ***  * ****   *** **** ... * . *******     **.* **. ::* . ..

47              YKTTAKWMRYYWRRAGMWYYGMMMMKYRMCRYWWYSKWKMYYRWRKKMMRRKGKCTCRRK
50              YKCGGKWMRYYWRRCCMTTYCMMMMKYRMYRYWWYSKTKMYYRWRKKMMRRKKKSCARRK
```

```
      **  .*********. *  * ******** ******* ************** *. .***

47    RKWMWCCYWYWWGAAKKMYRMWWYKKWYTTRRSKRKKRRYSWWYRWWRATGYYRSMKMKK
50    RKTATTMCWYWTSRRKKAYRMWACTKATCCRRSKRKKRRYSWWYRWTRRCCCYRSMKMKK
      **      *** .  ** ****  .*    **************** *    ********

47    YYRKRYYRKATTMRRKKYKKAGTAYYGCRYWRYRYWWYWRASMRKKWTRAKKKKSYWKYM
50    YYRKRYYRKMWKMRRKTYKKCRATYYTYRYWRYRYTACARCCMRGKWCRRKKKKSYWKYM
      *********  .****.***. ::**  *******    *..** ** * **********

47    KWYYRKYYTKKYYYKYWKRRKKSRRKKKYAKKKRWTCAGAYAKYKMCTCKWYRRKRKGKR
50    KWYYRKYYGKKYYYKYTKRGKKSRRKKKYGKKKRWWTCKRYGKCKMTKTGWYRRKRKKKR
      ******** ******* ** *********.*****  .  *.* ** .  ******* **

47    YRKKKRKKMRSYGKWYKKRGCTWWTWKMYGTWKKCATMTAWMKKWMMWSKCGCKCRYKYK
50    YRKKTRKKMRSYRKWYKKRKSYWWKWKMYAAWKKATCMGRWMKKWMMWSTMSYKYRYTYK
      ****.******* ****** . **.****.:***.: *  *********. . * **.**

47    RRKYRYYRMYKRYYYYYRTAYMKMWKRRKMRKKATATRKKRMRRWSYWRWKRYKAGATTC
50    RRKYRYYRMYKRYYYYYRYWCMKCWKRRKMRKKWYGCRKKRMRRWSYWRWKRYKRRCCWA
      ******************   ** *********  . *****************  .  .

47    MWYKMCRKRRRKRYMYKKKRRKARWGMYMRWKKCACYKKYATAMYYGAKRRYRYYWRKYC
50    MWYKMMGKRRRKRYAYKKKRRKMATKMYCGTKKYMYYKKYRKRMYYAGKRRYRYYWRKYT
      *****  ******* *******    **   **   ****  . ***..*********** 

47    GTMRYKYYRSKYKKRKKGMMTWCYYYKMKKRKGKRKRKWYTGSRKAKKWWYKKKYGRRKY
50    KYMRYKYYRSKYKKRKGSAAYTTCYYKMKKRKKKRKRKWYWTSRTTKKWWYKGKYRRRTY
        ************** .      ******** *******  **.:****** ** **.*

47    KYYAACKRATCKRYSKWYRRMYKRWKMRGRKARMKKRKMMRKMGGKRRYRRMKKYMTYYT
50    KYYWMMTRGYYKRYGTTYRRMYKRWKMRKRKRRMKKRKMMRKMRRKRRYRRMKKYAAYYK
      ***   .*.  ***.. *********** ** ***********  ********** :**.

47    KTTCGYRKYKYYYCACSKYCCYYYWTYYKKYYKYKKYYWYWKKRYKRRAKKMKKKTSWMM
50    KGATCYRGTTTCYYWYSKYTGYYYWCTYKKYYKYKKYYWYWKKAYKRRMKKMKKTCSWMM
      * :  **  .  *   ***  ****  **************** **** *****. ****

47    YRKKYRRRKWSKGTARKAMRYCMAMKKRWKMKCKKYRRAYKRWMMYWRYYYMKRAKKYRR
50    YRKKYRRRKWSKRWCGKWCRYYMGMKKRWGMTMKKYGAMYKRWMMYWRYYYMKRWKKYRR
      ************  . *  ** *.***** *. ***   *************** *****

47    WWRYMKWKKMRCAGTKWKKGGWCWRMWKMCMKYYRWMYRRTGRYGKRGTAKCRMRGWYWM
50    WWRYMKWKKMRTGRYKATKRRAYWRMWKCGMKYYRWMYRRAARYKKRAAGGTRMRRWCTM
      *********** .  * .*    *****  **********:.** **.:.  *** *  *

47    RRRYRRMMWGYMWYGKATCAMGRWRKMKWWSMYKYKKYTCTRRRWWYRKWYYARRWWWSW
50    RRRYRRMMTRYMWYRGCCTCMKAAAKMKWWSMYKYKKYWTCGRRWWYRKTYYCRRWWWSW
      ********  ****  .  .*    *************    ******* **.*******

47    WKKAYYRKRRSTCCYRYTGKMMKRKWRYWYRRYYKWRKKKSRRMAAARCKKMKKSACGGR
50    WKGGTYRTRRSAMTYRYWKKCMGGTWRYWCGACYKWGKKKSRRMGMRATKGMKKSWYRRR
      ** . **.***:  ***  * *  .****    *** *******.    * ****    *
```

```
47    WKSKRRWYRRRRKKKKMGRMKRGKRRRKMCMKWWKKRRRMKRRYYMRTTMTTGACCTRWK
50    WGGKRRWYRAAAKKKKMTGMKRKKRRRKMYMKWWKKRAAMKRRTYMRCGMGCCGAYAAWK
      * .******   *****  *** ****** *******  **** ***  *   .. : **

47    MKYCYKKKTMRRRYKCCARKRKYRKRRGGKWMYRKMMYRWSGWAGYSKYTAYMKYKKKKA
50    MKYTYKKKKMRRRYKATGRKRKYRKRAACKWMYRKMCTAASAWRRYSKYGGYMKCKKKGM
      *** ****.******. .******** . *******    *.*  **** .*** ***

47    KGKYTKGKWYWRYRWRYYKKCYYKYWASYYKKRRCMWKKKGKKMWYRMMYTMGRYMKRKR
50    KKKCATTKWYWRYRWRYYKKTYYKYWMSYYKKRATMTKGKKKKCTTGAMYYMRGYAKAKR
      * * :. ************* ***** ******  * * * **     ** *  * * **

47    YRMYRSKGYRYTYYCMYWCACGKRMMYYKATAKRKGAKCKRYTYMRYYRRYYSKKWYYKW
50    YRMYRSKRYRYKYTMMYWAGTAKRMCYYKRYMKAKARKTKRYGYMRYYRAYYSKKWYYKA
      ******* ***.*  ***.. .*** ***   * *. * *** ****** *********

47    YMYYGKKTTYCGTYRRMWKYYYCGYKTKKKMKTRRKKKAYYKMYYKMKKKMRWKYMMYRS
50    YMCCKKKWYYYRCYRRAAKYYYAAYKYKKKCKCRRTKKMYYKMYYKMKKKMRWKYMMYGS
      **   **  *   ***  ****..** *** * **.** ******************* *

47    YWRAKAMYYYKKRYSMTYGYTWRYYWYYAKKMYTGYRYWKMCYRKMKYSYWYWKYKKKSW
50    YWRRTCMYYYKKRYSMATRYGWRYYWYYCKKMYYSYRYWKATYRKMKYSYWYWKYKKGSW
      *** ..**********:  * *******.**** .*****  *************** **

47    KRYMSMRGYMRMRKKWRYKKRAYWYRMAYYTCKMRRMTWTCARYRYKRRWYYRRRAKKWW
50    KRYMSMRKYMRAGKKWRTKKRGYWYGCTCCCTKMRRCYTCYGGYRYKRRWYYRRRGKKWW
      ******* ***  **** ***.***  :    ****     . ************.****

47    RKRKKSYKRTTCCTRRMYRWYKYYRMKSAAKYYYMWMKWYYRMYMYWKKKRRYRMGYRKY
50    RKRKKSYKGCWYYKRGACAWYKYYRMKSRMKYYYMWMKWYYRMYMYTKTKRRYRMRCRKY
      ********     .*    *********  **************** *.******  ***

47    ATTGYYYYWMRRRAAAYRYKWKAMRSYAYAAKGAMMSRYWYTACCWMKSAGGATATMGCRY
50    RYWKYYYWMRRRWWWYGYKAKRMRSYRYWMKAGMMGGCWCYMTTWAKSCATTCGWMTARY
          ********   * ** * **** *  *..**.  *     * **.. : . * .**

47    KCCWMWRGYTATTYKACTYWYGYYMYTCTTMYMTTKTYKAGGCYYRMYSKRWYYKSAGAG
50    KSYWAWRAYATAGYKMTCTATAYCCYWMWAMCMKYGWCTWRKYYYRMYSKRWYYKSTCGA
      *. * **.*::: **      .*  *   :* *.    .    *************: ..

47    GTYYYRRKKRKKTKRRRTWMSKMKYYMWWTRRCWTKKKSYKCKTMKRWRKTTCGAATWKY
50    KWTTCRRKKGKGWKGGGCAMSKMKYYMWACAAYTGKGKSYKYKYMKRWRKGCAACGCWKY
           **** *  *     *********       * **** * ******  .... ***

47    RSKKKMWKRKKYKRYCGKKWYRMRYRWWSMTCACKKYKKYRYAKKGAKWRGGKYKWWYYM
50    RSKGKMWKRKKYKGYMKKGWYRMRYRWWSAWYTYKKYKGYACWKKTCTARKRKYKWWYYM
      *** ********* *  * **********   : **** *   ** .. *  ********

47    RYSYYYMRMRKKMWWTWYSYRKKYWKKKKRKACTARKMYYYYARTGTKRYAGMRKRKKRY
50    RYGYTYMRMATGCTWCWTSYRKKYWKKKKRKCMGGRKAYYYYGGYACKAYWRMRGRKKRY
      **.* **** .   * * *************.  .** ****.  . * *  ** *****
```

```
47          KRKYRKYRTACAYMMSMWMCKRWKRYRMYSKRRKRYRYKKKWRRKRMKMYACAYWKKWYY
50          KRKYRKYRYRMRYMMSMWMMKRWGRYRMCCKRRKRYRYGKKWRRKRMKMYRYRYWKKWYY
            ********    ******* *** **** .******** ***********   *******

47          RYYKMRYCWYYWYRMKTYYATATMWKYMRWGYSGGGYKRCKGAGMRKKRSRYYRTAMKKK
50          RYCKMRYMACYWYRMKYYYMYRYCTGYMRWKCSTRRTKAMKACAMRTKASRYCRWMMKKK
            ** ****   ****** **       ****  *    *  *...**.* *** *  ****

47          YYRKYYWCYTYMYKWYWTRRMYKKKYYRRKYYCCAAGKKKTCYKMKKYRYKKKKWRYKRY
50          CYRKTCTYTWCMYKWYWKRRMCKKKYYRRTYYYYRMRKKKCYYKMKKYRYKKKKWRYKRY
             ***       ******.*** *******.**     ***  ******************

47          RTCKWKRRWKRKMYMMGKCCGKRYRRKKKKYYWKCTWGMKKYRRKKYCYKRMYKKYKKKR
50          RYYKWKRRWKGKMYMMKGAMKKRYRRKKKKYYWKYAWRATKYRRKKCMYKRMYKKYKKKR
            *  ******* *****   .  ************* :*  .******  ************

47          RCRTWWWYWYRYRYYKRYKKMKCTYYRYCTTTRKWKTCARRYTGRKKTRYCSYKYKYKMR
50          RYRAAWWCAYRYRYYKRYKKMKAAYCGCTCCWRKWKCYGRRYYRRKKYRYTSYKYKYKMR
            * *: **  *************.:*       ****  .***  *** ** *********

47          YYTKRRKKKSCMWRRRKKKWAWAGKKRYKYKKTTGKKKWYRRRKKGWKRYYRWKMRYRKA
50          YYCTARKKKSYMWRRRKKKWGACTGKRYTYKKWCKKKGWYRRRKKKWKRYYRWKMRYRKW
            ** . ***** *********. .  ***.***   ** ******* *************

47          KKMWYRKRYKYKKWRGTAMYKKWKRYCWKKKKWKYKYRKYWKRRCAMYKTARWKKWYKYA
50          KKMWTRKRYKCKKWRKYMMYKKWKRYAWKKKKWKYKYRKYWKGRYRMYKWRAAKKWYKYG
            **** ***** ****   ********.*************** *  ***    ******.

47          GTKWWRACMKKYKTGAAKASRRWMKCAAGKKYWRKRYYRYYSM
50          AYKWWRGTMKKYKKRRTKMSRAACKTGTRKTCWRKRYYRYYSM
            . ****. *****.  :* **   * .: *. ***********
```

The pairwise alignments of two accessions 47 and 51

```
47              YYMYKRYSYRMYYRRYRRGGGCYMYYCTRCCRSASRASTRKWRWGTYRRCAKYWMTYRKY
51              YYMYKRYSYRMYYRRYGAKKATYMYYYAGTYGGWSGMGAAGTAWACYRGYWKYWMYYRKY
                ****************    . **** :    . *  .:    *. **   **** ****

47              SRYSWTYKKWRYRSMKSAYYWYWWMKCAAKRYRWKWCWRRKYRRCWAGWTYMYYWKRAAS
51              SRCCWWTKKWGYRSMKSGYCACAWMKTGCKRYRWKAGWRRKYRRYWMRAWTATYWKGCTG
                ** .*  *** ******.*    *** ..******  ******* *       *** .:.

47              GYGRWWKAYTWRTTYTGGYWWMKRGCCRKMRSSYCWMRWMMAYRRYMTRWMKYMKRWWYR
51              KTRATWKTTWWRWAYKSAYATMTAATTRKMRSSYTAMRWMMTCGGYMKRWMKYMKRWWYR
                     **:  ** :*...*  *. .  *******  *****:   **.************

47              RMKYRKMCCWRGGRWYYRKYYWYYYYRGMRCYTRWRRYWYYRRRYYKWYWRGRCCARKWT
51              RMGCRKMGYARKKRWYYRKTCWTTCTGAMAYYWGWRRYWYYRRRTTGWYWRKATYRGTWK
                **  ***   *  ******  *     .*  *  **********   ****      .*.

47              ATCYRMRRMYTRRKWKMCACMRWTYCWGYRKKWRAMMKYYYMKRYRRYKKKWRGMRYRWY
51              GCTTRMGRMYWRRKWKMYMYMRWYCMWRYRKKWRTMMKYYYMKRYRRYKKKWRRMRYRWY
                .   ** *** ******   ***   * ******:****************** ******

47              CMRMYGARYWRYRYMYRGRRYMRGGRWYWSRYRRWWWSRKCCRKYYWGAYRARYYYKYTA
51              YMRMYRCRYWRYRCMYRTRRYMRKARTYWSATAGWWWSRKYMGKYYWRMTRRAYYYKYGT
                 **** .****** *** ***** .* ***    ******   ****   *  ***** :

47              YRYWTGRMRYYWKKKCTYMRYYYCCKRWYCYWRRYYRAWSYTYRMKYYGTRMRSYWYKYY
51              YGTAWRRMRYYWGKKMCYMRYYYMTKRWYYYWAAYCRWWSYKCGMKYYRYRMRSYAYKYT
                *     ****** **  ******  **** **  * * ***.  ****  ***** ***

47              ATATGKARSYCGYCMRYKKRWYWCYKARYKRMKRRYWYTGMKYWKRKRTTYTKMWKRARG
51              MYRYTTGACYTKYYMRYKKRWYAMYKGGYKRMKRRYWYKKCKYWKRKRWWCKKMATGGAA
                     .. .*  * ********  **. **********.  *******   .** . . .

47              RYRAGGRTTTCAKGKMRKWRRKMKYYYGTTKRWMMKCCCRMKYKTARMRYWKKYCGCAAY
51              RYRGKRAACWTMKRKMRKWRRKMKYCYRAGKRWACKTAGGMKYKCGGCGYWKKTGATTGY
                ***.   :    * *********** * : ***  * .  **** .   ****  . :.*

47              WWSRYRKSKYMMYTTATRYGTYRKRYKKRRRTGCRRKKKKKKRYRRRWWGYYKKKKYSMW
51              WWSRYRGCTTMMYCCGGACRWYAGGYKKRRGCAARRKKKKKKRYRRRWWRYYKKKKCSMW
                ****** .. ***  .     *   *****  ..*************** ****** ***

47              KKKTCTAGRKRRYRMYYKMWKGGYWKRYKKTKYKRKRRKKKWYRKRYWWWTARWMKKWKY
51              KKKGTYMRAKRRYRMYYKMWKATYWKRYKKCKYKRKRGKKKWYRKRYWWAWRAACKKWKY
                ***      ************. ******* ****** ***********      *****

47              ATRMGMKKATWRCKKMYYYYWCRTMMCTSRCAYRRKKYKKYWRKRYRYYYKYWMGACTRR
51              GYGCACKKWWWRMKKMYYYYWYRCMMTGSRMCYGAGKYKTCWRKRYRYYYKCAMAGTGRR
                .   . **  ** ******** * **  ** .*   ***. **********  *..  **

47              YKKKMRRWKCATGCRCMMWYMWAAAMKYGGYGRKWMYAGRTACCRYKRKMGYYYKYYGAM
51              YKKKMRRWKMRKSMRAMAAYAAGWWMKYRKYKRKWMYMARYRYYGYKRKMSYYYKYCRTM
```

```
                                   *********  .. *.*  *  .  ***  * ***** .*     *****.*****  :*

47                                 MYRMMKGCTYKGWMWCWYMRSRRSMGRWRKMRYYSRKYWSYKAKYGCYKWWKRMKKGRTM
51                                 MTRMMKRTGYKKWMWAWYMAGRGCMKAAAGMRTTGGKYWSCGWKTRYYKWWKRMKKCGWM
                                   * ****   ** ***.*** .* .*     **  . ****   *   *********   *

47                                 RATCAMTWWAAKTRRYRRYKKTGTYYRYYYTACYRYWRRGYGKKMKYRSKYGCGCMAYRY
51                                 RWKYRMCATMWKWGRYRRYKTCAWCYRYYCGCTYRYWRRRCKKKMKYRCGYRMAYMWYRY
                                   * .  *     *  ******. .  ****  . ******   ******. *  . * ***

47                                 KGGTTTWYKKMGRKGYRRTKRRWKWCCGTAYWYYRKWWKMRSWGAATTRYTAMRRYYYKK
51                                 GAAGCCWYGKMRGKKCARCKRRWKWMTAAGYWYYRKWWKMRSWRCRWAAYYRCGRYCYKK
                                    ..   ** **  *   * ******  .:.************* .  : *    ** ***

47                                 KAWRYRRRKRRGRAYWYYAKKRRMKKSYRKKARRMYRWYKYKYRRYRKKTTYYYWAGMWM
51                                 KGWRYRRAKRRRGCYWCYGKGRRMKKSYRKTGRRMYRWYKYKYARYRKKYYYYYAGRCWM
                                   *.***** ***  .** *.* *********..*********** *****  *** .  **

47                                 YMYKTGCKKYWRKYRRYYRKKKYRWKSYYYYRGYYWGATARYCAYWRWMYYRARWRTMRY
51                                 YMYKCATKKYWRKYRRYYAKKKYRWKSTTYYGAYYWKGGCRYYRYWRWMYYRGGTRKMRY
                                   **** . *********** ********  ** .*** . .**  ********.  *.***

47                                 MKKKKRRKYYRKCTCKARYYRKKSRRRKGGCRRKKKKRWKMMTYSKRKKMMSWKWRYRKR
51                                 MKKKKRRKYYRKYYMKRAYYRKKSRGRKTATGRKTKKRWKMMYYGKRKKMMGWKARYGKR
                                   ************   *  ******* ** .  **.******* *.******.** ** **

47                                 RKMRSCWKYKAKWCWRYYRMYRWYWYRAKKYMYSMKKKYKKKKAKRMYACATGKRYKCYK
51                                 RTMRSYWKYKTKWMWGCYAMTGTTWYRRKTCMYSMKKKYKKKTRKRMYTGGCAKRYKMYK
                                   *.*** ****:** *  * *    *** *. ***********. ****: . .**** **

47                                 KYTTKKCKKSYKYKRMCRKKYYATYRKKKGCAYKKRYMRYYGRKWYKKWYYKYMTACATC
51                                 KYWYKKYKKSYKYKRMMRKKCYGGYAGKKRYWYKKRYMRYYKRKWYKKWYYKYMGTGCAT
                                   **  ** ********* *** *. *  **   ********* ************ : .:

47                                 TGCCMTCYWYKYRRRKYCRWTCTCTAAKYMRWMATMYYSKTRYWYRGGMRYYRRKYWTMY
51                                 YKMTMAYYWYKYRRRKYYRWWTAMARRKYMRWMMCMCCGKGATTYGCCMRYYRRKYWWMY
                                       *: ********** **  : :  ******  *  .*    *   ********* **

47                                 YMRRRYSYAKKKRRSMSYSKYSYYKRKRKRYKRMSAAAKWKYYYCAAATYKSMKYKKYWA
51                                 YMRRRYSYGKKKARSMSYSKYGYYKRKRGRYKRMSRCRKAKYYYYGGGACKGMGYGKYWT
                                   ********.*** ********.****** ****** . * **** ...: *.* * ***:

47                                 TGGMRGTCTAYGGATTGRYRWRTAKRKTATAAYMRARRMRRYKRRKKYRYMYRGGCKRRR
51                                 CACMRKYYWWYRRGGCTAYRWGCGGAKYRWRWYMRCRGMRRYKRRKKYRYMYRRAAKRRG
                                    . **     *  .    ***  .  *     ***.* *************** ..***

47                                 RWSMKYYRYRRRMWGKRRAKKSYKRRKRKYRKWRYWKKKRRKTRMYKKYWYYGKYYMKRW
51                                 RASMKYYRYRRRMWSKRRCKKSYGAAKRKTRKWRYWKKKRRGWRCCTKYWCYKKYYMTRW
                                   * ************.***.****   *** ***********  *  .*** * ****.**

47                                 WSTCKKKYKKKKKWYARSYKYASYYWKMGWMYRRYKRWYSKYYYYCWYWKYRYCYRRRKY
51                                 WSWYKKKYKKKKKWYRRGYKTGSYYWTCAWMYRRTKRWYSKYYYCTWYWKYRYTCGRRKY
                                   **  *********** *.** .****. .***** *********  *******   ****
```

```
47          CKKTRYSYRYRTKYCGYKYKGRCTARGKTMKYYKRTWRRRKKKGTCKKTYKRYYAAYYGW
51          YKKKRYSYRYRYKYTRYKCKKRTYWRRKYMKYYKGWWRRRKGKAWYKKCYKRYYRRYTKT
             **.******* **  ** * *   * * *****  ***** *.  ** *****  *

47          AKGKYKKRRMKRGKRKRYYTKWSYYRKWKKRYMYKSYKRCCSTRKCTRGRMKARWKWRRK
51          WKKKCKGRRMGAKKRKATTWKWSYYRKWKGRYMTGCTKRTTCGRKTCGKGMKWAWKWRRK
             * * * ***   ***    ********* ***  . **  . **     **  ******

47          WWTTRRRMYRKRYYMYYRWTSAMGRWGWKKYYRRKKGMRMKRWRYMKRKYRKKKWRMSWY
51          WAGGRGRMYRKRYYMYYRAYCRMAGWRAKKYYRRKKKMRMKRWRYATAGYRKGKWRMSWY
            *   * ************  . *. *  ******** ******** .  *** *******

47          SYYATKYGTAGGKKKKMWYYYMYMWRYRWKKYYKYSMYYRYSRKWCCTKRAWMYKARMAC
51          SYYGCKYRWWRRKKKKMWYYYMYAARYRWGKYYKCSMYYRYSRGWAGCKRRWMYKMRMWM
            ***. **     ***********  **** **** ******** *.  ** **** **

47          TKMWAKRTTMMAAKSSASRKRKTKWMYACRYSYYWTTKKRTTYKKYTKMSWRRCYYKCAW
51          YKCTCKRCYCCGGKSCTCRKRGAKWAYMARYSTYAGWKKAAAYKKYYKMSWRRTCYKYTW
             *  .**    ..**.:.*** :** * .*** *   ** ::**** ******  ** :*

47          GAYYTKSKRGKMWRTTTCAATCSGTRRYKKKMMCKRYYKCRRYRKCGRYAATKATYACKS
51          AGYYWKSKRKKMWACGGMRTAMSKCGRYKKKMMYTRYYKYRRYRKYRRYCGCGCATGTKS
            ..** **** ***      :: *   ******* .**** *****  **..  .: . **

47          MKKSRYKAGCYSKGMGMTRYCYKGKRWYKYTATTRRKMWKMRKRACTRRYCTTYRTMRAG
51          MKKSRYKTATYSKKARCYGCAYKAKRTYTYCGCARRKMWKCRKRGTGRRYTACYRCAGRR
            *******:. ***       .**.** *.* . :****** ***.  *** : **

47          TWYTYKKGCTTMMKYYRMKKYWTAKSYSWWYYCGKSARYRMRRRWYYYYCRGAKKRYSRK
51          YWYGTGKCYCCMMKYYRMKKYTCRKCYSWWYYMRGSWRYRMRRRWYYYYYRTTKTRYSRG
             **   *    **********   *.******   * ************ * :*.****

47          KKSRYKRKRMRKRRMMYSKMSKKMRRMACWYRKMTGRRKRYKRYYRRMKMYARKYKYKYR
51          KKCRYKRKRMRKRRMMYSKMGKKAARCGMWYRTMCKRRKRYTRYYRRMKMYRRKYKYKYR
            **.*****************.**  * . ***.*  *****.********* ********

47          YKYAAMAARYKKAKGAATMKKACMKGGTYKKWYMCKRRTGCYKYMRYRAYGGKYRMYCTG
51          YKYGGMRMRYTKRGSRRYMKKMYMGAAACKKWYMMKRRASMCKYMGCACYTTKYRMCGCK
            ***..*  **.*  .   ***  * ..: ***** ***:.  ***   .*  ****

47          MCYYYWWYGKKKYSAGCARWRYRWYKRYRSMKGRKKTCGTAGMKYRKRRWKRYMKKMYYY
51          CYYYYWACATKKTCWRYTRTRTAWYKRYRSMKSRKKKTRCGRMKYRKRRWKRYMTKMYYY
              ****  ..** .   :* *  *********.***.   . ************.*****

47          YKRYKWWACCYWWKKYYRRKYKYRKYSKKWMYTTCAYYWRKKWRKKYYKYKYYYWMWYWK
51          YKRYKAATTYCTWGKYCARKYKYRKYSKKWMYKWYMYYWRKKARKKYYKYKYYYWMWYWK
            *****  :    * **  **************.    ****** *****************

47          KYRRKKYKRRKKMWRKWWSWMMYRMRSYRRRKWSSKRKYYYRRMRGRMKGKKRWKKYKKM
51          KYRRKKYKRRGTMWRKWWSWMMYRCGSTGGGKWSSKRKYYYRRMRRRMKRKKRWKKCKKM
            ********** .************  *    ************** *** ****** ***
```

```
47          YRKYYRWGMYKKKYWYATCTWRTGATAYWKGTRRRRRYWWYKKYKKRKYKGMMTWAGYKA
51          YRKYYRWAMYKKKYWYRCSWWRYRRCTYWGKYRRRGRYAWYKKYKKRKCGSAAYTRKYKW
            *******.********  . **    :**   *** ** *********  .      **

47          KTWKRMMWKMSKKWWKSRGAKKWMMKKATTAGMYKACGRYYRKYWYYYRRCGTCKRKYWY
51          KGTKRMMWKMSKKWWTSRATKKWCCGKWYYMRAYKRAARYYRKYWYYYRRSKYYKRKYWY
            *  ************.**.:***   *      ** ..************.   ******

47          RYSKWRKMKRRKRRKKMMYYGAKRMMKKWMMMKTMTGAYKWMYYKKRRKWKRWGGWTYYK
51          RYGTTRKMKRRKARKKMMYYCWKRMCKKWMMMKWCYRRTGWMYYKKGRKWKRARRAYYTT
            **.. ******* *******  *** *******       ****** *****     * .

47          WRKKWWKKKYRKATATRRSKYKRKRYRYKRKRKWKRSRRMKYKYMKSRWMYWRRRRMWKY
51          WRGKWWKKGYRTTCGYRRSKCGRKRYRYKRKRKWKRSRRMKYKYMKSRWMYWRRRRAAKY
            ** ***** **.: . ****  **********************************  **

47          YYTCMKKKKKCRRAKYYYKMMRWYYRRSYWRRMYYKMYYYKKYSSKYMRYYKKKRYKYRY
51          YYWYMKKKKKYRRGKYYYKMMRWYYRRSYWRRMYYKMYYYTKTGSKYMAYYKKKRYKYRT
            **  ****** **.**************************.* .**** **********

47          WKAKYWRYRKKYKSYWYKWATGTRRGKACTGKRYKTYKYMRGSYYMKYRMRRRMRKWRYK
51          WKGGCTRYRGGYKSYWYKATAACRRAKRYYKKRYKCCKCAGKSYYMKTRMRRRMRKWRYK
            **.   ***  ******* ::. **.*    ****  *    ***** ************

47          KRWYMYKWMMWSWKKKGKYYKKYKYRMKRYYYYYRWYKMWMWKMRKKRKKCMRKKKKKMW
51          KRWYMYKWMMWSWKKKRGYYKKYKYRMKRYYYYYRAYGAWAAKMRTGRKKMMRKKKGKAW
            ****************  ***************** *  *  ***. *** ***** * *

47          YKMYYTRRRKRRRKMTYKCRYRKRRWRYRMRWKGKMYYKYKKYRMTRGKAGMRKYRKYYW
51          YKMCCWRRGKRRRKMKYKYRYRKRRWGYRMRWGKKMYYKYKKYRCYGRKCTMRKYRKTCT
            ***   ** ******.** ******* *****  **********    *. ******

47          YKMTMGMMKRKYRRYRTWTARYYYCAAGKTGCWTCAAKKMYKKWKYRWKAKYRYTKGYWR
51          CKMKMKCMKRKYRRYGYAYWRYYYYRMRKKKSACATGKKMYKKWKYRWKGKYRYYKRYWR
             **.*  ********    ****    *. .  .:.************.**** * ***

47          YKRYRTYCTTTMRKRWKKKMYMCCKYRRKKKKTGCKRRMKKCWYWKKYRRKRMYCYWSRC
51          YKRYRACYAAGMRKRWKKKMYMYAKYRRKKKKYRYGRRMTKYWYTKKYRRKRMCAYTCGY
            *****:  :: *********** .********    ***.* ** ******** .* .

47          KKRMAKYATTCKYKKKTARR
51          KKRMGGCCCYYKYKKKCTRR
            ****.  .   ***** :**
```

The pairwise alignments of two accessions 48 and 49

```
48              RMMKCWRWWRYKYKYYRRYWWYWRWYKRRWYKYCKYYMRSYYYRWYGYRYRYYYYTRYYA
49              RMMGYWRAAACKYKYYRRYWWYWRWYKRRWYKCYKYYMRSYYYRWYRYRYRYYYYWRYYW
                ***  **    *********************  ************ ******** ***

48              YMRYCYGKYYRWYRMKRMCYYSKRYRRWRRKSYARAWWGCKYRRYWKCYMTRGKCMKYWK
49              YMRYTCKTTYRWYRMGRMTYYSKRYRRWGRGSYRACAARTGYRRYWTMTMCATTYMGYWK
                ****   . ****** ** ********* * **  .     *****.  *   . * ***

48              RMMYMKWWYYRYGGKCRAGCYCYRYWYMCCWRKRYYMYKMKYMTWYYRMYMRRKWTRRWR
49              RMMYMKWWYYRYRKKYRTATTTYRYWTCYTWRKAYYMYKMKYMGWYYRMYMRRKWCGATA
                ************  * *:.   ****    *** ********* ***********

48              MKTTRYKGRSWWKMKYWRKYKRRWRMRRCWMWMYKYKRGAYRRSYYYKYRKYYCYACTCY
49              MTWWRYKARSWWKMKYTRKYKRRWRMRRTTAWMYKYKGAGYRRSYYYKYRKYYACTTCTY
                *.  ***.******** ***********   ******  ..*************. :   *

48              RWAYWKRRKYWWKSRGYCCYYKSRKKRCKYKTYRMRTYWRWWKKKMMCCRGWWTRYWYRM
49              RWWYWKRRKYWWKCRTYATYYKSRKKRYKYKKYRMRYYWRWWKKKMMTYGRWWAGCWYRC
                ** **********.* *. ******** ***.**** **********    **:  ***

48              WRYYRRMRKAGCYRYYWYRRKMKRYYYYWAYYYKSKKAYRYRKYRRTYRRRWKSYATTRY
49              WRYYRRMRKRATYRYYWYRRKMKRYYYYWWYYYKSKKMCAYRKYRRKYRRRWKSYTCCRY
                ********* . ***************** *******   ******.********:  **

48              YYYWWRSMYRMKKWRYKACCRRWYACYRSMMYYYRTTGCMRRYRWMYRYYRKYWAKCGMG
49              YYYWTGGMYRMKKARYTGATRRWYRAYRSAMYYYRCGTTMRRYRWCYRYYRKTTMKTTCR
                ****  .****** **... **** .*** *****    ****** ******   *

48              WYYRKKYAYWKKKYATGKGYYMKKYYRRKRACYMKKYRYKACGCYGCKMYKKRKMWYYWC
49              ACYRKKTTYWKKKCRWKKKYYMKKYTRRKAGTYMGKYRYKRMATCTYKMYTKRKMWYYWY
                  **** :*****    * ****** *** . ** *****  .    ***.********

48              YRYRRRKCGTAKWWRMYKGSRRGKKCWMYGCCRKKYRTMKKRKYYTMWYMYRWTTRRRYM
49              YRYRRRKTACGGWWRMTKRSRRKKGMWMYRYYGKGTRCAKKRKYYAMWCMYAAYGRRRYM
                ******* . . **** * *** *  ***    *  *  ******:** **    *****

48              WYRYKWMYRYWMKYKRKKKTKKKKRRYYRYGRYWYKSMKKKMYKTAMRYMKRKMYCGCTM
49              WYRYKWMCGCWMKYKRKKKKKGGKGRYYRYRRYWYKSMKKKMYKWCMRYMKRKMCYKMWA
                *******   *********.*  * ***** ************* .********

48              YGRMTGTGCAYYKRKMYYRKKRRWRRGRYCTSYGTWKKMGGAGGKWRYKRKKRCMWKKGG
49              CRRMGAARTMYYKRKMYYRKKGRWAATGYYKGCACWKGMRRWKKTWRYKRKKRTMWKKAT
                  ** .:   *********** **    * .. . ** *     .******** ****.

48              MYTRYWYGKRWWGMWWYACRYAAGARGGWWWTYAMSKAKYKMTMWRRRCTRKYKKKYKRR
49              CCCRYWYKKRWWKMWWYGARYRMRMGRKWWWCCGMSKGKYKMCMWRRRTGRKYKKKCGAR
                   **** **** ****..**       ***  .***.**** *****  ******   *

48              KTWYRWYCTRMRYKKRMYKYKKTGRKAWYYYMTWRARWKKKYMRWYTRRRKKKYTAGGKG
49              KYWYRACTGRMRYKKRMYKYKKYKRKMWYYYMYWRGAWKGKYMRWTYAGAKKKYAGRKTR
```

```
      * ***    ************* ** ***** **. ** *****     ****:.  .

48    TYYWRKTKKARKKYKYKATAAKGMYSMYKYMTYRYMKYKCMRYYARRKKYKKKKAGAGTT
49    YYYWRKKKKGGKKYTCKTCGTTRACSMYGCCYYRYMKYKYMRYTMRRKKYKKKKMATRYY
       *****.**. ***. *:  .:.  ***    ******* ***  ********* .:

48    CAKCCWKKKKKKRRRYKKYKCRYGKGWKMRTYRYRKGSYWYKKYKYKGYASYWKSWYRRR
49    YRKTAWKKKGTKRGRYKKTKYRYRKAAKMRWYRYRKKSYWYKKYKYKRYMSYWKCTYRRR
        * .**** .** **** * ** *. *** ***** ********** * ****. ****

48    RRWRRYMYAAKKKYMRKTMWRMYYSKYYRRAGCMKRRKWMWRAKYKRATGMGKASRKGKK
49    RRWRRYATTWKKKCMRKWAWRMYYSKYYRGGRYMKRGKWCTGTKYKRWYRCATGGRKAKK
      ******  : *** ***  ********** .  *** **   :****    ....**.**

48    RWCWWKYYMYRKWTRSRKMYKKKCYGKRCCYSKKRRRRKKKRMYRRYYKYTTTRYGGYCC
49    RTAWWKYYMYRKWWRSAGMYKKKYYKKAMMYSKKRRGGKKKRMYRRYCKCYCARCAACYM
      * .********** **  ***** * *   ******  ********* *   :* ..

48    KRRSMAGCTWGKYTKRWMKSMRKKYYWYGAAMRRRTSWRYTAYTCTTAKWWYKAAAKRCT
49    KAGGARRYCWKKYKKRWMKSMRKKYYWYKWTAAGACSWRYAGCGTCCCKTAYKMGGKGMC
      *  .     * **.**************  :     ****:.     .*  ** ..*

48    YKGAYKRATYYYTYKMKYRYKKRTTKRRMTGYYRKTRWKWWMWYRWMKKGYGKCYKRWWS
49    CTTGYKRGCYYYGYKMKYRYKKGACGARMYSYYRKKRWKWWMWTRWCKKRTRKYYKRWWS
       . .***. *** ********* :   ** .****.******* ** **   * ******

48    KKRYWKKRWKRYMKRYKAAGTYRYMRYYCAAGTTKYGTRYWYRMKRAKRGRTKYYRWRAM
49    KKRYWKKGWTRYMKRYKWRKKTATMRYYYRMRCAKYAGRYWYRMKAGKRKRKKYYRWRCC
      ******* *.*******   .   ****     :**. ******* .** *.******.

48    CAWTWRWKKRYMMYRRWRYGAMYKAYYKAKSSTYKKKTCYMRRYRMKWCRKTMRYYGTGY
49    TTTCARWKKRTCMYRRWATKRMYKMYYKMKSSKYKGKCTYCRRYAMKASGKKMRYYRYRY
       :   *****  *****    *** *** ***.** *  * *** ** . *.****   *

48    YKYTYRKKAWTKRYWMSYKWRRMKMKRYKRKRYMMRYKRGCYGWCYMYMMMSKKAATACC
49    YKYWYRKKGACKRYWAGYKAGRMKCTRYKRKRYMMRYKGRYYRWAYMYMMMSKKCGCCTY
      *** ****.  **** .**  *** .************   * *.*********.. .

48    WACRCWKCAMCGRTSYRAATYWKMKAYCCWKYRWYGCGCYTMMCAYKKKAAMMKWWYTMY
49    AGTRYWKYMMMRRCSYRGRYYWKCKGYYTAGCRWYRTRAYYCCTGTKKTTRMMKWWYGMY
       . * **  *  * ***.  *** *.*     ***   .*    . **.: ****** **

48    SCYKCAWYACTYYTKKKSKKKAYKKYCCYMMRASGGTCGWRWKSKAKKYYYKRWSYKRKY
49    SYYKMCTYGAAYYCKKKSKTKGYKKYYYYMCGGGTTYAAWRWKSKGKKYYYGAAGTKRKY
      * ** . *..:** *****.*.****  **  ..   ..******.*****   . ****

48    YRKYKRKYRYTYRYMKKKSYRRKYRKAKWKRRWKYKRTGCCKCTAAAKYRKKKRYKKMRK
49    YRKYTRKYRYGYRCCTKGSYRRKYRKWKWKRGWGYKRGRTYKTACCMKYRKKGGTKKMRK
      ****.***** **  .* ******** **** * ***    * :.. *****   *****

48    GRMCRYRKRRCATGGGYRRYAGGRWKTWTYGYGTRAAWYMWWRKYRRYSRATCACYCAGS
49    RGCMRYAKRRARYTTTYRRYCAARWKYWACKYSGRRRWYMWWRKTRGCSRRYYWYCYCAS
          ** ***.     ****...*** *:  *. *  ******* *  **       ..*
```

```
48          MMYMSYGRRSRTRYRYWWYKWRYYRKMKKCGAAKCRMYWRKMYARMKCCKMCCGGYKMKM
49          MMYMSYSRRGGKRYRYAWCKWRYYRKMKKTCGWKMRMYWRKACCAMKTMKMTTACYKMKM
            ******.**. .**** * **********  . * ******  . **  **  . *****

48          RYCGYSKMGATWYRMRATKKKTCTGGGGYYWRMKRRRTKKCKKTTWTMWYWTRYYRRGYM
49          RYYAYGKMRWWWYRMRMKKTKASYKTTRYYWRMKRRRYKKYKKCCTWMWYWYRYYGARTC
            ** .*.**   ***** .*.*:.     ********* ** **    **** ***

48          TRYYRKYMRGTMRKYKKTGACKRKGSYYRWKRYKSWKARKCRKKKGYRKGTCYTYCWSAC
49          KRYYRKYMRRYARKYKKWRMAKRKASYYRWKRYKSWKGGKTRKKKRCRKKATYGYYWSRM
            .********   *****   .***.************. * ****  ** : * * **

48          TMMKYKRRRRYGCGTSRTAGRTYGRKRCTTGYWCCTCTRCRWKKWYRTGCAACGTRGAKC
49          YMMKYKRRRRYKTAGCRAGKRYYRRGAMWWKYWTYCYGRYRWTKWYRYRYGGSKKAAGKS
             **********  . .*:. * * *      **     * **.****   ... . ..*.

48          KKKKMYYRKAKATCATGAARWMRYKKMACYCWMWSYTAAGGAGKKGSRRYKWMSKASTCA
49          KKKGCYYRKRKWCYCAACGAAMRYKKMTACYWMWSYGGCARWRGKKSRRYKWMSKGCCTW
            ***  **** *   .:...  ******:.  ***** ...    * *********..

48          RYRKYMRYYCTKRYKRSMKYRAGWRYAACTYWTSWSWRKCAWKKGTTYYRMYKTYGAAYW
49          GYRKYMRYYTAKRYKRSMKYRRRTRYCGAGTWGSWSWRKYWWKKSYYYYRMYKCYRWRYW
             ******** :**********   **...  * ******  ***.  ****** *   **

48          KSWKRKWMKATKYKKYMYWMMKYGCRRRAGAAMKWRKYKKKYYRYKRRWKYKRRYCARRR
49          KSWKRKWMKCCKYGKYMYWMMKYAYRRRMKRRMKWRKYKKKTCGYKRRWKYKGRYTCRRR
            *********. ** *********. ***    *********   ******** ** .***

48          RMRYATTGCTKRKYKYKYRTCWRRKMMYTKRRWKKMRRWYCTKYATAGTKRRWKKYRMRR
49          RMRYRWYRTCKAKYKYKYRATTAGKMMYYKRRWKKMRRWYTGGCGCTKWKRRWKKYRMRR
            ****      * *******:    **** ***********    . :  ***********

48          TAATWRYTAKGCCATTYRMTCKRSKTCTWMTGGGYRWTMYKTTKKKGGAKGACMMKRRYY
49          CTGYWRYYMKKTYGCGYRMKYKRSKWYWACYRRRTRWYMYKYYKKKKKWKRGTCCKRACY
             :. ***  *   .  ***. ****          ** ***  ***   * .   **  *

48          YWRWKKMTYMKKSYKMRYYCKKAYMKRCAMMRRKYYWRYRTGTRKKMCKYRRYGYWKRGW
49          YWRAKKMKYMKKSYKMRYYYKKGYMKRMWMMRRKYYWRYRCTYAKKMTGYRRYRYWGRRT
            *** ***.*********** **.****  ***********    ***  **** ** *

48          CRCACAKRYATGAKKRRYRKKWWKRWRKKKKWRKMTKKCYKMYRRMYYAYKCRWRWRCCA
49          YRYGACKRYMWKWKKRRYRKKWWKRWRKKKKWRKCCKKYYGCCRRMYYRCKTAARARMYR
             * ...***    *********************  ** *   *****  *   * *

48          AKWGYARTMTMYKYWKKTKCTYAMWYYWRYKTGYSWTKRYKYGKMWTRMYKYRTTWCMRM
49          MKWKYCAKMCCYKYWKKKKMGCCATCTWRYKWSYSWYKRYKYRKMWGRMYKYAGGWMMRA
             ** *. .*  ******.*   .    **** .*** ***** *** *****   * **

48          CKRCRRWWAMSYSWRKKRRGCATWRRYMTATGTKACTYGCYYYYYYKKTWGAYYYYKRMC
49          MKRMRRWWRCGYSWRKGRRCTTKWRRYACCCKCGWYAYAAYYTYYYKKCTKWYYYYTAMM
             ** ****  .***** **  :.****  .      :*..** *****    ****. *
```

```
48    CYYRKYTKYKMYKRKKYYYCTYKYRTTCWRRCRATTYGARCWRYSTYGGMYKRWRRRYGC
49    MYYRGYCTYKMYKRKKYYYYWYKYRGGTWGGAGRWWCRWGTTRYGYYRAMYKRWRRRYRY
       *** * .***********  ****   *  .          **. * .*********

48    AAGCAAMYRWMRKRKYWRKRKTYSYYYAAAAWWTKKWKYKRWKKATCTKTRYKACRCCTR
49    CWAYGGMTAWMRKRKYWRKRKWTGTCCRRWRWWKKKWKYKRWKTCKSYKKRYKGYRYYKG
      . . ..*  ************  .       **.*********.... *.***. *  .

48    MMWGGCYGCRWTCYMKMRKYYWMKWTCRYMYYGTTKAYYYGTTGYTYKYYKYKKMGAYRT
49    CMTRRTYRMRWYMYMKMRKYYWMKTKYRYMYYAKKKGYYYACCACACTYYKYKKMCGCRK
       *    *  **  *********** . *****...*.***.  . : .******* . *.

48    YTGKKCCTYWTTAKKYKKYRAGRRYWYYRYKTMYYYRGMKKRRAWKYAYWYKYKYKKKKY
49    CGKKKAAAYWGCGTKYKKYRGARGYWYYRYKYMYYYRKMKKRRMAKYRYTYKYKYKGKKY
         **..:**  ..******..* ******* ***** *****  ** * ****** ***

48    YYKRYMTAYGYKKMKYKMKYRWKRGGCRAWRKSRKRSSKGWWKKKKTGTARCKAGGGRRY
49    YYKRYMWMYKYKKMGCKMTYRWKGSKSGRTGKSRKRSSKRWWKKKKCRYCATKWRKRGRY
      ******  * ****  **.**** . .    ******** ******   .  *     **

48    MKKRKSYYYRYWKRSYWSYKRYWKTYKRMKWKRYTRRMWCKCTKKYGYAKWWYSRKKKGA
49    MKKGTSYYYRYWKRSYWSYKRYWTKTKRMKWTRTCGRMTTKTAKKTACRKWWYSRKKKRW
      *** .******************.. *****.*   **  * :** .  *********

48    KKYYMWRRWWYWRRSWRMTTTSAWWYKYKKWWGKKAYRRTAGCCSKWRYKKACCGKMKKY
49    KKYYMAGGATYWRRSWRCAAASTWWYKYKKWWTKGMYRACRRMMGKWRYKKRYTTKMKKY
      *****     ******* :::*:********* *  **      .******    *****

48    YACARKKTGMKRGKYYYMMSMKKSYACKRKYWYKKYYMWTARRSKKCATCWYMRKKGRYW
49    YGACRKKACMGRKKYYYMMSMKKSYWYKRKYWYTKYYMWAGRRSKKTTCMWYMRKKARYW
      *...***: * * ************  ******.*****:.***** :  ******.***

48    RKKYYWKKKSKKRKYMYWYRKYYYYKKSKWYTATTMACCAKRCARRTRGKYKKKKYRYRR
49    AKKYYWKKKSKKGKYMYWTGKCYYYKKSKWTYRYYCWMMRGASRGGKGRKYKKKKYRYRR
       *********** *****  * ********            .   .  ***********

48    KKTRKKYMRMSRYTYWMSCWKGKKRRYKKKYYRKYMGKRSMKKRMTAYGCCRKYWYYYRK
49    KKGRKGYCGMSRYYYWMSYWKCKKRRYKKKCCRKYMKKRSMGKRMAGYATTRGYWYYYRG
      ** ** *  **** **** ** ********  **** **** ***:.*.  * ******

48    KYRKKRRWKYYRYKMKCCYTGGRCCKKKWARGYKRKATGCMTWYMGKGTAATCCKAKCGT
49    KYRKKRRWTYYRYKMKYAYKAAGYYKKKTWRRYKGGCCRMMKWYMRKRGGGGYYKWGTRY
      ********.******* .*...   ***  * **  .   *.*** *  ..   *

48    GTYKKKKRKWRMTKKRGGCCGTKMWKKAYWSYYYTTYYRWTCWYYRRKYYWYSKKMKRYY
49    KCYKKKKRKWRMKKKRSKYYRCKMAGKCCACCYYYKYYRWCTWYYRRTYYWYSKKMKRYY
        **********.***.     **  *.  . ** .****  *****.************

48    MRKKCCACYGRRYCACKRKWKRRTWMSCCYCAKMCGMRYKWWKCYYCKASGTTWWYRKAK
49    MRKKTYGTYRGGCYWYKAGWKRRWWMSMACYGKMYRAGYKWWKMYYYKRSRYWWTYRGMK
      ****  . *       *  **** *** .  .**    ***** ** * *   * **  *

48    GGKRYWYWYYYATTCAMMYAGTKTRSYCGTKRYKRRTCTKGTKTSKYTTCTKKKCCGKSG
```

```
49              RKKRYWCACYYMYWTRMMYCKKKKRSCGAAKRTKRRCTGKTCGCSKTCCACKKKMYKKSK
                  ****   **     ***. .*.**  .:** ***   *    **   . ***   **

48              YRKYYMWYYRSRAGRGRCWAGGCYYYTACCKGKSMMRTRRGRKYYYWRCWTWKTKKARYK
49              YRTYYMWYYRSRMRRAGMWMRRYCCYWRYYKKKSMMGYRRARKYYYWRGTGWKAKKGRYK
                **.*********  *.  *      *    * ****  **.*******   **:**.***

48              WRYRRMYYRYYYRRYKKGTRRRTKTKRAAATKSGACTTRRWTYRMRMTWRKRRKYRMKMA
49              WRYRRMYYRYYYGGYGKRKRRRKTKKRWMMYKSRWYAARRWKYRMRMKWRKRRKYRMKMM
                ************  * * .***...**    **   ::***.*****.***********

48              MWTKYGSYGGAAAAKKWRYAAAKMRKWKKWYRMKRKRKACTAARRYKRMWYTKGKTRWMY
49              MWKKYKGTAKWMWTKKWRTGTTKMGTWKKWYRMKRKRKCTCTTRRYKRMWYKKKKYRWMC
                **.** . .    :**** .::** .************.  ::********.* * ***

48              MRTTKCCYKKRYWAGRTWYRKSRTKYGKKTKKKYMKTRYKMYMMACTKYTKKRMMRKKWK
49              ARWYKGMYKKRCAWKGWWYRKSRYKYKKKKKGKYMKGRYKMYMMMMYKYKKKRMMRKKWK
                 *  *  ****      ****** ** **.* **** *******   **.**********

48              YGYYSRYYYRKCKKKMKMWKRRKYKWWKKKRYYCCKYRRMKKTAKW
49              YKYYSRYTTGKYKTKMKMWKRRKYKWWKKKRYYTYKYRRMKKCGKW
                * *****   * *.*******************  ******* .**
```

The pairwise alignments of two accessions 48 and 50

```
48              WWKRYYGGGYYRKYRKRAYRKRKYYYRRCRSRCKKRSRRRYKTAGAAACSKKTKYMKTAK
50              WWGRYYKRKYYRKYRKRRYRKRKYYYRRTRSRTKKRSRRRYKKMKTGTTSKTWGTMKAGK
                ** ***   ******** ********** *** *********.  :.: **.   **:.*

48              YWKKKSWWRRWAARGKMKYYKAYAYWCRKRYTYKSRYYYMGKWMYRWRWKYAYAGYRRKM
50              YWKKKSWWRRWWWRTKMKYYKWYRTWTRKRYWYKSRTCCCAKWMYRWRAKYCYWRYRRKM
                ***********  * ****** *  * **** ****    .******* **.*  *****

48              YMMRMRWYATWYYYRYYYKWYCKYCWYWKRRRMSYWWRWRKYCRRSMTARRRRWRRKWWK
50              YMCGMRWYCYAYYYRYYYKWTYTTYWYWKRRRMSYWWRWRKYTRRCMYRRRRRWRRKWWK
                **  ****.  *********   .  ***************** **.*  ***********

48              MGRYRAGYYTCCWYYKYACGTGKKRMYYYYRYYYCAYWKAKWKWKGRCWWARMYMRYRWY
50              MRAYRWRYCWYYWYCTTCYRWRKGAMYYYYRYYYYRYTKRKWKWKCRYWWGRMYMRYRWY
                *  **  *    ** . .    *  *********  * * ***** * **.*********

48              YYYMRYYYYCRRGTYYSRYWMYWTSMRRRYRKYYMYRYRRCRMRWWMRKMKARKATTWRY
50              YYYMRYYYTYRRKCYYSRYWMCAYSMRRRYRKYYMYGYRRMGARWWMRKMKGGTRYYWRY
                ********  **  *******   ************ ***   ********. .   ***

48              CMMRMRYRYWSYKYKYYRRRRMMYKTYWRYYKRRMRTKCRSRRRYKRMCGGYYWRSWYMM
50              YCMRMRYRYWGCKYKYYRRRRMMYKYYWRYYKRRMRCKYASRRRYTACTATTYTGGWYMM
                  ********. ************* ********** *  *****.   .  *  .****

48              WKYYKMYMYYYRMRRMKKRSWKRGKGGATWWMCCGGTKYCAAYKRRYKRWWRGRYRGRKY
50              WKYYKMYMYYYRMRRAKKRSWKRAKKTRKAAMAARRWKYYGRYKRRYKATAGRACARRKY
                *************** *******.*    .  *..   ** . ******         ***

48              KKMMCYGYYCKRMYYKKKYCYWGKYAWYYKTYKRRRKWMRKGAATMYKAKCKRSYKRYMK
50              KKMMMYRCTYKACTYKKKCYCTTKTTWYYKYYKRRRTWMRKACGCMYKTKYKRSYKRYMK
                **** *    *   ****     * :**** *****.****... ***:* *********

48              AAKMRYSACMYYRYYYCKYRYSWWKRCWRYRYWRWYKAKMRKAWAKYWKKRYRKKKKKRK
50              RRKARYSMYMYYRYYYGKYRYSWWKRYWRYRYWRWYKTKARGGWWKYWKKRYRKKKGKRG
                  * ***  ******* ********* **********:* * .* *********** **

48              YRKYRYKRTTGRGCCSKKKCKKYRMRMKRKKSYKYRRRYTYTARSRCWKKKRRCGGAYMY
50              YGKYRYTRYKKGKYASKGGYKKYRMRMKRKKSYKYRRRYAYYWRSRMWKKKRRYRRRYMY
                * ****.* .    .**   *******************:*  *** ******    ***

48              MMRRWRMRYYWWKYGKKARYRTMKWYYKYGRKYYYTYWRTGTRRRKKKRKKTGTSAMKWR
50              MMRRWRMRYYWWKYRKKGRYRAMKWYYKYKRKYYYCYWRCRKRRRKTKRKKWRWCWMKWR
                ************** **.***:******* ***** ***  .****.****   . ****

48              WYTRRRRTTGTGYCYRWMMMYKWRRRWSRWYTGACATWRYRRKAGAGAMGCGMRAYMTYY
50              WYKRRRGKWRWAYYYRWMMMYKARRRWSRWTCAGACAWRYRRKGRGTGCRMRMRRYMWYT
                **.*** .   .* ******** *******  ....:******. . .    ** ** *

48              KYYRRMTRYKYRKRTWWMWRAAGGCGRAWMAKWGAAARCRRGAGKRYRYYMRAYKAGKRY
50              TTCRRMGRYKTGGACAWAWRWMRAYARRWMTKWTTGGASRGARRGRYRYYMRGCKRRKRY
```

```
            .  *** ***      * **   . .* **:** :.. .* .   *******. *  ***

48          RGATCYYSRACYSWRWMWSMRRKWWRMYKKYMYMMKRGKYKCRTKYKMYRRMWMCMYRKM
50          GKTGAYYSRGMYSWRWMWSCRRKWWRMYKKYMYMMKRKKYGTRGKYKMYRRMWMYMYRKM
              : .****. ******** ***************** **  * ********** *****

48          WKAYCTTYKWCMKYKKKWRKWTRGYWKSMKKKMKMKRYKSRYRYKRYKTMRYRMKRKKRK
50          WKGYMAAYKWAMKYKKKWGKWKRKYWKSMKKKMKMKRYKSRTGCKRYKYMRCGMKRKKRK
            **.* ::***.******* **.* *****************   **** **  *******

48          WKMYWARYKRKKRAAYKMKKYKMAKMRMKKSKMKTRMYWWKCCKYWKYKKKMTTWAYRYK
50          WKMYAMRYKRKKRWWYKCGKYKMRKMRMTKGTMKCRMYWWKYTKTTTTKKKCYCTWTGTK
            ****  *******  **  **** ****.*..** ******  *  . ***        *

48          KRYRYKYRCYGKKKTTKYRKYKYYKGKWKCRYRYYKRYKRKAGGTKCWAAGYKYRKKRKR
50          KRYRYKYRYCTKKKWAKCRKTKCYKAKWKYRYRYTKGYKRKCRAAKTTGTRCKTRKKRKR
            ********   *** :* ** * **.*** **** * ****. .:*  .:  * ******

48          KWKAKYKKWRATKKYKKKTTYKRWKKCTCKTRTTKYKKKRYCCRMMRWYWKWAWYRRYCK
50          KWKRKYKKWGGWGKYKTKAGTKRWKKSYAKKRKKKYKKKRCTTGCMRTCWKWWWYRRTYK
            *** ***** .  ***.*:  *****. .*.*..******     **  *** ****  *

48          WMKKYRYKMYYKYMMMKYWAYMSKKYKAKKKKYRGGYTKKKYKKYAYTRKRTRWYYYYGC
50          WMKKCRYKMYYKYMMMKYWMCCSTKYKCKKKKYGCAYWKKKYKKYWYCRKRYGTCYCYKY
            **** **************   *.***.*****  .* ******* * ***    * *

48          KTCKWARGKMATCTYAYWRKMGACATTYTKRYKRRYRKKMYRKRKCTTTTTAMKWRKYRA
50          KYYKTTRRKMGGGYYGYWRKMRGACAGYGKRYKRRYRKKMYRKRKYYWGGAGMKWRKYRR
            *  * :* **.   *.***** ...: * ****************     :.*******

48          YTTGMGKYYYWMYRAWAGTAKWKCTAKYYACRYWKMKRYCRKTYAWYYKSWMYYTGMWYW
50          CYCKAAKYYYWMYRWWRKKMKWKYWWTYYRYGTTKMKRYMRKKCCTCTKSWMYTCAMWYW
                 .******** *  . ***   .**     ***** **. .   *****  .****

48          YYSYMGAYWGTARKKACWKYWCATSRYWSMKKRRTKAMMKCKKCGRYWYKSCTGTTGTGA
50          YYSYCKCYTAAGRKKCTAGCWTGGGRYWSMKGRGCTTMMKTKGTTGTWYGSYACACCAAG
            ****  .* .:.***.    * . .****** *  .:*** *     ** * : :  :..

48          WRKYRYKYTTKGARRAYAAKCAATRMRTGKSGTACACRYGACAYKRYTMYTMTKRAASCW
50          AGGYGCKYAKKRGRRTYGTTYGRYRMRYSKSRATTTGRCRMYRYGRCCMYWMCKRTTCTW
               *  **:.* .**:*.:. .  *** .** :: : *     * *  ** * **::. *

48          KGTAYYTWYYMKRGWMYWYRYYYRRAMTKWWRKRYKTTKMWRCTRYRMYWMMRKRKGTGK
50          KTWCCCKWYYCKGRWCCWCGYYYRRRMWKWWRKRYKCCKMWRTGGCRATWMMRGRKACAK
            *  .  .*** *  *  *  ***** * ********  ****    *  **** **. .*

48          KYKKRTWKYWYMYMWARCACKKYKSMTKYYKSYKRKKCGACGTTAGTYMKGMYKWAWKYW
50          KYKKGYWKTWYCYMTRGMGYKKYKSMCKYYKSYKRKKTRGMKYWCACTAGRAYKWGTTYW
            ****  ** ** **    . ****** **********  .    ..      ***. .**

48          WWKRYKGKGCKGKKKSTRYWYRKKMKRMRYWKRRYACCRCWCYYWTYRMARMWKKYYKMR
50          WWKRYGRKKSKRGTGSYGYWTRKKMKGCRYWKRRYGTTRYWAYCWWCGCRRMWKKTYKMR
            *****  * .*  . *  ** *****  *******.  * *.* *     ***** ****
```

```
48      CYGRWMTKTATTCTMKRRYRWGRAKYTKKKRMYRYWKRYMKWWRKYWYRKCWYRYTRWRK
50      MCRRWMKKKGGGYAMTGRYRARATKYCKKKRMYRYWKRYMTWWRGYWYRKTWCGTYRWRT
           ***.*..   :*. ***   :** *************.*** ***** *    ***.

48      YYWRRYYYCYCMWKRYYMRYYYRATARCCYRYWARCKGTYRKRYRGRCCTYWAMMRKGSR
50      YYWRRYYYTYTMWKRYYMRYYYRRKRGAYYRYWGAYGRGYRGGYRKGYTCCWRAMRGRSR
        ******** * ************ .  . ****.     **  **      *  **  **

48      KYKYYYKKKRKGCGCGWYWCWYKRYKMYTGASGCARRYYKYYRKKRYWYKRRKSMYKYRK
50      GYTYYYKKKRKRYKMTWYWYACKRCGMYKAGSSTGRRYYKTYRKKRYWYKRAKSMYKYRK
         *.********     ***   **  **...*. .***** ********** ********

48      YKRRKKRYRWRKCWYKRKAASWKRKMAWWKRSKKGTTTAGGGYYWRRKRYSTYKYTTTCT
50      YKRRKKRYRWRTGWYKRKCCSWKRKMTWTGGCKKSYCARRRRYYWRRGATGGYKYCYWMY
        ***********. *****..******:*   .**.  :    *****   . ***

48      AASSMMWKMMKYKKKYYWMMRCGKRYRWRKACKKKKYKRTGTTKYWYMRYSTTKYKKRRY
50      TWCSMMWKMMKYKKKYYWMMRTTKRYRWRKWYKKKKYKRYSKWKYTYMRYSCCGTKKRRY
        : .******************  *******  ******* .. ** *****    *****

48      YRCKKRWKKYMRTRYRGGKATGTMMKYRAYYKYYWSSCGKKKRRTYRYMRSRRYKKMRYY
50      YRMKKATKKYMGCGYGRKKRYRYMMTYRWYYKYCASSAAKKKRRKYRYMRCGRYKKMRYY
        ** **  ****   *   *    **.** ****  **..*****.*****. ********

48      CGKYRKGAGTTAMAYYMCKKKKYAYMMRTRWKMMMSKRWKMRMYYRTTWKMYWWMKWYRK
50      YKKYRKRTAWKMMRYYAMGKKKYGYMMRCAWKMMMSKRWKMRMYYRYKWKMYWWMKWYRK
          **** :. . * **   ****.****  **************** .************

48      RCKYWKKRRYKRKKRARMWYRKKMCKTWRYWMKKKYCCYGCCYKKKYGTAKCTYTTGACT
50      RMKCTGKRRYKRGTAGAMWYRKKMYKCARCAMGGKYYYYKGTYKKKYAAGGTKYCGRCAA
        * *   ****** . . ******* *  *  *  **  *   *****.:.  .*   ..:

48      KYAYAYKRKKWTGAKYWGYGKWTYYRCYAKRMCRMKKMMKTKYKWKTRCAAYKRYYKATA
50      GYGTRCTRKKWYRWKYARTRTAWTYGYYGGRMYRCKKMMKKKYKTGYAYRGYKRYYKCCG
         *.   .****   **    .   *  *. ** * *****.***      .******. .

48      TMYWKKKKTKRYARRRRWRKKMRYCYKYKMMYKRRRKKMRRWYKAAGCKMYWWKRRWYYY
50      CMYWKKKKKKRYMRRRRWRKKMRYTYKYKMMYKRRRKGMRRWYGRGAYKCTAWKRRWYYY
         *******.*** *********** ************ *****  .. *   ********

48      RMRRWRYKKGKKCRYATWWRRRKSKKCAGYTGYCMRRYYKWYYRKTMYCACYKYKKMMYW
50      RMRRWRYKKKGGYAYRYWWRRRKSKKYWRYATCMAGAYCKWYYRKYMYYMYYKYGGMMYW
        *********     *  *********   *:      * ****** **   ***  ****

48      CRKKYYWYMSYKCWKMRYYKKRKRRRYKYRYYRMKRRYCGGWKYWYGYKYRYWRKKKKKY
50      MRKKYYWYMSYKYTGMRYYKKRKRRRYGYRYYRMKRRYTTRTKYTYKYKYRYWRKGKKKY
         ***********   ************ **********    ** * ******** ****

48      YYYYRTTCCYKKKMCYMKKKRAWRKRSRWKKRTKRKKMKSYYSYWRSYWSKRYKWKKACM
50      CYYYRYWYTYKTKCTYMKKTGRTGKRSGWKGRYKRKKMKSYYSYWRSYWSKRYKWTKRMM
         ****    **.* ****.    *** ** * **********************.*  *
```

```
48          KRMKWRMGWMMGWMWKCAWMKKKATWWYSRTSSYGAKKYMWRWMATSKYYYKWAKKSMRK
50          KGMKWRMKAMMRAMWKMGWMTKGGCWWYSRKGGCRWKKYMWRWCGASKYYTKWMKKSMRK
            * *****  **  *** .**.* . *****...   ******* .:**** ** ******

48          AGYYKSYRAAMYMRSSWRYAATGKWWYWYRRAKRCGAAGCCTKKYYYRAYRYKRRMKKRR
50          CAYYKSTGRMMYMRSSWRYRRYRKWWYWYRRCKRSRGGAATCKKYYYRMYRYKRRMKKRR
            ..****    *********    ********.**. ....  ****** ***********

48          KTCTWKTMGRWMWKKWGAYKRGWWSKKWKRGCCWKKYKAKRKRYRKRTACAAGKYKYRYY
50          KATGWKKARRWAWKKAKWYKRRWWSKKTGGTTMWKKYKMKRKRYAKRYRYRMRKTKYRYY
            *:  **.  ** ***   *** *****      ***** ***** **      * *****

48          RRTGGRMRKSKKMSCRYTYMKMKYYKKKRRGKMKYYYKRSMKATGCRKRYWYYKYRKRKR
50          RRGKRRMRKCKTCGMRYYYMKMKYYKKKRRRKMGYCTGRSMGRWKSRKRYWYCKYRKRTR
            **   ****.*. . ** ************ ** *   ***    .****** *****.*

48          YAKKKMCCYYRYKKKRTRCYYKTTGWGKKWMYRAKGCCKATKKRKSYWYKYYKKSYWRAK
50          YCKKKCMMTYRYGKKRCATYYKCGKWKKKWMYRGKRYAKWWKKRKSYWYKYYKKSYWRWK
            *.***    *** ***   ***   * ******.*  .*  ***************** *

48          TWWRATTCWMRWRYYKYKKKYRMGKWTGTWCTTCAKKKRKKRYKKKYMKKKYRKYTRKKM
50          GWWRMCATWMRWRYYKYKGKYRMKKWGACTAKYYWKKKRTKRYKKKYMKKGYRKYAAGKM
             ***  : ********** **** **  .  ..   ****.********** ****:  **

48          KYKKAYYSYTGAGKKMACRRCWRKRWGGAACKYRMYRYTACYRYYYRKKKWGKCCKWYKR
50          KYKTGYYSCARTCKTMTTGGAAGKRWRRWRYKYGACAYWRYYRCYYRKKKWKKMMKWYTR
            ***..*** : : *.*:   .  ***     **    *   ** ******* *  ***.*

48          KKTMKYKKRKRKGKYYKTAKKYYYMSGCCTMWWTKYRMRKRKCAKYKAATARRRWYRGTR
50          KKGMKYKKRKRKCKYYKKMKKYCCCSTAAACAAAKYRMRKRKTGGTGRRGRGGGACARYR
            ** ********* ****. ***   * ..:   :******** .               *

48          SYWKKWYCKKRYKWRKYCTCKKKRCGTKSKKKYYWYRYSAKKGYYRYCKSKWKYYKKKYY
50          SYWKKTTYKGRYKWRGYYYTKKKRTTATCTGKYYWYRYGGKKKCYRYMTGTTTCCKKKYY
            *****   * ***** *   ****  :... *******..**  *** ... .  *****

48          RKWKYKRYMYSYGATRMYYYRMRCCRWYRRYYRGYWAKYYYKYRKWKKATGKMYYMKART
50          RKWKYKAYMCSYAGCRMYYYRMRTARWYRRYYRAYTWKYYYKYRKWKKRWKKMYYMKWGY
            ****** ** **.. ******** .********.*  ***********   ******

48          RRKCGKMKMKRTKKYRYKRKKKKAKRKKAWMWWSRYYYAKCKYRKKKYR
50          RRKMAKATMKRKKTYRYKAKKKKRKRKKGWMWWSRYYTTTTKYRKKGYR
            *** .* .***.*.**** **** ****.******** :. ***** **
```

The pairwise alignments of two accessions 48 and 51

```
48                TTWCKKYMKYWGKTATYWKCWRRYYKRRYYYMKWCRWYKCKYRKYWCTWWCAKYRRYKRW
51                AWWYKGYMKYTAGCGAYWKTWRRYYKRRYCCCKWYGWYKMGCRKYTYWWWATKYRRYKRW
                  : * * **** .  .:*** *********   **  ***   ***   **.:********

48                MYKMRYWRYMRKYRRSAGGYYYKYRKYMYYRKGYKKGWYYYRYYRRMRYYRYWCARMMYK
51                MYKMGYWRYMRKYRRSWKKYYYKYRKYMYTRKKYKKATYYYRYYRRMRYYRYWTGRMMYG
                  **** ***********   ********** ** ***. *************** .****

48                YMYMRYYKCGGWATRWYRKRRWYTYYSKWRYMRKWYRACYGYKSRKKCTTTYRYRRYKRC
51                CMYMRYCTTCAWWYGTYRGRGTYWCYCTTRYMRKWYARMYRYKSRGKTCWWYRYRRYKRT
                   *****  . .*    ** *  *  *.. *******   * **** *    ********

48                YAYKRKRKYYMRYTGYAKWKWYMYYYYCRRRMYYKMKRYRYYRGYACCCGGYCYRYRRGG
51                CRYKRKRKYYMRYWKCWGWKWYMTTCTMGRAMYYKMKRYRYYRTYWYMTATTTYRYGRAT
                    ***********     *****      * ************ *    .   *** *.

48                YTRWRKYAAAARYYRRWSRTAGKCWAAKYWRKRMSYWWGCMARCTRRMRWWWKMRRKTYM
51                YGRWRKYGCGGRYYRRWSRAGTTYWMCKYWAKRMGTATRYCCRTGRRMRWWWKMRRKWYC
                  * *****....********:. . * .*** ***.      .*  ************ *

48                GRRKWAAYGCAKCRMYMRYWRWYCTCTTKYMKWRGKCYWKSAYYSTCYCKKWRRRRRRK
51                ARRKWTGTTGTKARMYMRYWRWYMWGYYKYMKWRCGYYWKSMYYSYMYYGKWRARRRRRK
                  .****:.   :*.**********     ******   **** ***  *  *** ******

48                YWKMRMRRCRYKYRYYYRWMKCYCWYCTGKRCTWCCKCGCKRWYASRWYRCCRMMKCRMY
51                YWKMRMRRMRYKTRYYYRWMKYYAWYTATKGMATTTKMRMKRWYGGAWYRYYGMMKGRMY
                  ******** *** ******** *.** : *  :   *   ****.. ***   *** ***

48                SCCCATKAMYRTYACCAKWGRYAGCACKRKYKRMAMYKYSMYGRWKKYYTAAYRGAKYCT
51                SMGTCKKGMCAWCWTYCKWKAYGKTTTGRKTTGMMAYKCGMYRRWKGYYYMCTAAGKTTY
                  *   ..*.*       .**  *.  :  ** . *  ** .** *** **  .  ..*

48                TCYYSCGKKTGKGTRSYKRYKTYWRRKKYMSRYYYRYYKYSKWGMRTRCKKACRMWKKMW
51                ATYCCYRKTWKKRYRSYKRYKCTARRKTYMSRYYYRYYKYSKARMRARMKKTYRMWKKAW
                  : * .  *.  *  *******   ***.**************  **:* **: ***** *

48                AWKTGRWKYKWRYKKKKCCKCTKMRGRKRRYRKKRCKYSYRRKRKWKRKRKYTCYARGAC
51                WWGCKRWKYKWRYKKKKTAGTAKMATRKGGYRKKRMTYSYRRKAKWKRKRKYYMYRRARY
                   *   ************ .  :**  **  ***** .****** ********  * *.

48                KRYKYRAAKKRYTYWWKKRYRYRWWCYRWGMKYRYGASRRYSMCKCGYKMKYYMRGKARG
51                GGYKYRWRKKRYKYWWKKRYRYRWWYYRWKMKYRYAGSRGYSMMGATCKMTYYMRRKMAK
                    ****  ****.************ *** *****..** ***  .  **.**** *

48                RWATYWMKRYWKMGMKKYWWKRRKAMKYRKRKGKTACKYYYYMMKKMYYRMAKRKKGCKK
51                RWWCYWMKRYWKMRCTKYWWKARKGMKYRKRKCKKMYTYCYYMMKKMYYRMWKRKKSYKG
                  **  *********  .***** **.******* *.  .* *********** ****. *

48                YATCAARKRRSGAKWYARYAYTRYMYKATCRKKKKYMCKYRKRYAGRYCAKCKWKKWMKC
51                TRYTTGRKRRSATKWYRRYMCKAYMYKWCARKGKKYATGYRKRYGTRYACKGTTKKAMGY
```

```
                        :.*****.:*** **  . ****  .** ***   *****. **..* . ** *

48                  SYKCYMMWKKRTRARWWTWWYWRMTKGSRRMGGKTKTKGKWYRRRTCKKRYKKRWWAYGA
51                  GYTACCMWKKRGGTRTWKWWYWGMWKKSRRMKKKWGWKKKWYRRRWSKKRYKKRTWRCSR
                    .*..   *****   :* *.**** * * ****   *    * ****** .******* *   .

48                  RMRYYYRYWRKKKMRMCACCCGKYCGMRMKKWWTCYCCGGGKWRMYYMKKMGCCGYRACYR
51                  GMRYYGYWRKGKMRMYGTATKKYYRMRATKWWYMYAATSKKWRMTCCGKMKYYKCGTYYR
                     **** **** ****  . .   **   ** .***   *.. . ****     **      : **

48                  CTCTYKTYWRRYYYYTGTKTWTTRWWCYYGGKAKMAMYMATARMMYCYYKRMYYMYKRKK
51                  AAMYCTYYWRRYYYYGAKGCTAWATAYYCRTKRKACACCWAWRMMYMYYKRMYYMYKRKK
                    .:    . ********  ..   :      *    * *  .     : **** ************

48                  KYWYRCYWMYGKRTTCAKAGRKCTCGAAYYRRKKYRTMRMRYKYSYTYGTKRMKRMASKK
51                  KTWCGTYACYKGGKWYMTWRRKSKTTGGYTRRKKYRGMRMRYKCSYYYKYKRMKRMMSKK
                    * *    *  *    .   .  **..  ..* ****** ****** ** *   ****** ***

48                  KKKKRRGKTKMYRKRSMRKKRTKKYWYYAAKCKRKCYTGTRTYWKKGMYYYWKKYMCCCY
51                  KKKKRRRKAGMYGKRSMRKKRYKKYWYYMMGYKRKTTCCCAAYWKKKMYYYWKKYCAYAY
                    ****** *: ** ******** ******    ***      :**** ********  . .*

48                  RMWMKYRYRYRCCCYWARYGYKTCCKYKKYMKWRYACYKKAKRKKRKTCCKTKAGARGCT
51                  RMWAKCRYGCAYYYYWWRYRCGAAAKYKKYMKWRYGAYKGWKRKKRKYYYGCGGTGAAMY
                    *** * **       ** **    :..**********..**   ******        . . .

48                  TGSRRYKRATRYRKRTCCTRGGAWKMGCRKKKTRSRGGCRYYMMCRCRCKRYKKKTCAKM
51                  WSSRRYKGGCRTGGAKYMWRAARWKCRSGGGGARSARKYRYYMMYRMGTGACKKTKYMKM
                     .***** . *     .    *.. **   .     :**     ***** *       **..  **

48                  TRKKYYSRWMYKTGTKMRKWYRKRRKYWKRKRKACTAAGKAAMWYKKRTMCKKRKKKSRK
51                  GGKKTTGGWMYKYRKKCRKWYRKRRKYWKGKRKCTCTTRKGTMWYKKGCMYKKRKKKSRK
                      **   . ****   .* ************ ***.   :: *.:*****   * *********

48                  YKTAAKKKCRWKKYKKGTYTCKWYKYRRMMYRMCRKKWYKWYSCMRRWKYKYKCMYRYKT
51                  YKYRTKKKMATTKYKKKCYWYKWYKTRRMMYGCYRKKWYKWYSYMRRWKYKYKMMYGYKY
                    **   :***    .****  *   **** *****   ********* ********* ** **

48                  ATCKKWWWRKYMYWTAGCCRYYGMKRMACYMARRWRGKWYYRWYKYYGRYWYSWWCTCMW
51                  MYSKGWWWRGCMYWYRRYMAYYKMKGMCSYCWRRWRSKWTCRWYKYYRGYWYSWWTCTMW
                       .* ****  ***       ** ** *..*   ****.**   ******   ******    **

48                  KWRAMRYAGAYRRCCYATTTYMCRWMRRRWCMWSTWYRYMSRCYMRRTYKATGTAKSYKK
51                  KWRMMRCWRGYGGYYCGYWYCAYRAAAGRWYMWSWWYRYMSRMYMRRCYKRYAAGKSCKK
                    *** **    .*       .       *     ** *** ******* **** **   .:.** **

48                  AYAWRRMYRWRWYMCYYMMSAGKGTKATYCTKRKYYYMYKRKRYMRGYRRWKKRKYMRGM
51                  TYWAGRCYRWRWCMYYYMMSTAKRKKGKYTCTRKYYYMYTRKRYMRKYRRWKKRKCAGKM
                    :*    * ***** * *****:.* .*..*   .*******.****** *******     *

48                  RGMYRYYKSSKCRYRMGRSRRYWAMRYYTKYGTKYTYKRKKKRAWRYKRCTMWGCYRATK
51                  GRMYGYYKSSKYRYGCKRSRRYATMRYYKKYCGKYWYKAKKKRMWRYKRTCATAAYRWKG
                      ** ****** **    ***** :****.**   ** ** **** *****     ..** .
```

```
48    KWRRYYRTYSTRWWRTMTYWTYYRMCGYRMRYRRKSGKKYKKRYKACTKKSYRRRGAAYA
51    KTRGCYAWYSCRWTGKACCAGYYRMMRYRMRYRRKSSKKYKKAYKRSWKGSTGGGKRWYR
      * *  *  ** **  .     ****  *********.***** ** . * *       *

48    GTYSRAGRCWMWRRKKRWCMMRRKKMRMRYYWRKGTTKYSKYRMGWYYKYKKGTRRRRKR
51    KKYSRMRRMWCAAAKGGAGMMRRGKMRMRYYWRKTACKYSKTAMKWYYKYKGAGRRRGKR
       .***  * *    *    **** ********** : ****  * ****** . *** **

48    KMKGACKATACCGWMTTWRYRYMKWRAKKKYYYSWRRCCCYCTGTKKWYWYRMGGCKGRR
51    KCGRTTKWAWMYTWMWYWRYRYMKWRGKKGYYYSWRRMYAYGARAKKWYWYRMKKTKKGA
      *   : * :    **  *********.** *******  .* : :********   *

48    KATKKYYYKSYRCYTTACRYRYKRRWRRCYKRKRWTCKKMRWMKKRRTAWYKYKRAGWWS
51    TRYKKYYYKSYGYYWACMRYRYKRRARRYTKRKRAGSKKARWMKKRRAWTTTYKRTRWWS
      .  ********  * :. ******* **  ****  .** *******:   .***: ***

48    KKKYRYRWKRGTKAGKRRKRYKTCCRRKTACAAGKYKRKKYKKWKAWYKWRCATRRTTRK
51    KKKYGYRTKRAGTMRKRRKRYKGAMRRKCGYRMRKYKRKKYKGWKRWYKWGTRCGGWGRG
      **** ** **. .  ******* . *** .    ******** ** ****        *

48    CKWRWKWRKYYKSYKTKRYKKAKACKYRMMSGWKWAWYRGYYYKGYKKRYKMKTYYKKYS
51    TGARWKWRKYTGCTKKKRYKGRKWSKCGCMGRWKWMWYRKYYYKKYKKRYKMKKTYKKYG
         *******  . *.****  * .*   *. *** *** **** ********. ****.

48    YYYAKTGWKYKCKYCRGWKKKYRKRMYKKKGWMYKYYRKRKKSKYYYGTCGAGRYRRMWR
51    CCYGKGAAKYKYGYYGAWTKKYRGRMYKKKRAMYKYCGGRKKSKYYTKYYARKRYRRMWR
        *.* . ***  *  .*.**** ******  ****   *******    .  *******

48    YAAYYMYRRKTGMWYKYKKKWRKKYYGYYRYYYSMKRYTCKSRKYAGTMRYCGACYKRWK
51    CGTTYMYRRKYKMWYGYKKKTRKKYYSYYRCYYSMKRYYYKSRKTRRYMRCMARMYKRWK
       .: ******  *** **** *****.*** *******  ****    **  .  *****

48    KMWKKKGKWARYKRKRCYSYWYKYYYYARMKYWYRKWKKRWMYWWMAAAARRYKYWYYRY
51    KMWKKTKKAGRYKRGGYYSYWYGYCTCRRMKYTYRKWKTRWMYWWMMRGCGRYKYWYYAT
      *****. * .****   ***** *    **** *****.******* ..  *******

48    RYKCMKRKKTCAKYYRCKKYKRRATYGTKCMKRYRYRYYTMKKYSKKMKCRTTGAKKYRR
51    GYTTMKRKGYYWKYYRMKKYKRGRYCKAGGMKRYRYRYYAMKKYSKTMKYRCCCTKKYGG
       *. ****    **** *****     :  *********:******.** *   :***

48    RRMYRRKKYRGGYKKKKRSMYWRRKYRYWKRMKRRYTYKYCYKKTSYYRTTTTYRYKYKM
51    ARMYRRGKYRRRYKKKKGGAYARRKYRYWKRMGRRYKYKYSYKKWSCCACCAYYRYKYKM
       ***** ***  ***** . * ********** ***.***.*** *     : *******

48    YAKKYWSRWYTCAGCCKKKKRTWRKKKKRRKSYYSYWRYYWSKRRYKWKWKKKMKTARMM
51    YMKKYWSRWYCTGRTYKGKKRYWRKKKKAGTSYYSYWRYYWSKRRYKWKWKKKMKCGRMM
      * ********  .   * *** ******  .************************ .***

48    WGKAACATKKWWYSRCWYMACTGRRWCTWWMKKKWWGTTKWKKKYYYKRCRRRRMRYTCW
51    WKKTGMRAKKWWYSRYWYMMYCTARWAKWWMTKKWWRGCKWKKKYYYKRYRRRRMRYATT
      * *:.  :******* ***     **..***.****   ********** *******:
```

```
48    KMMYRWTKAGCCGTAKMRRYGCGWMWKRRAYYKRYRMMKRRWRYKGTATRGWARKYCYCY
51    KMMYRWKKGAYMTWRKMRRYTGCWMWKRRMYYKAYRMCGAAWRYKKYGCARWWRKCTTAC
      ******.*..     *****   ****** *** ***    ****  .   * **   .

48    TGKSCACRRKWYYRTMYYKYKMMKRYYYWYMKKMRKKTTACRRKKAAWTCACMMGKKRWK
51    CAKSYWYRRKWYYRKMYYKYKMMKRYYYWYMKKMRKKWYRYRAKKRMAYYTAAATKKRWK
       .**   *******.**********************    * **     :.   *****

48    RRYMYGCYRTMYYYRTGRRYRTCTCTATYRRKKWRKKYWKATGKKYCYYWMYYMRRRYAG
51    RGYMTKYCRCMYYYRYRRGTGWYYMKMKYRRKKWRKKYWKRWKKKYYYYWMYYMRRRYTT
      * **    * *****  *       . .************   *** ***********:

48    KMMKKYYRYKKKCCACKKAKKMWWMKYRKKKYRKRTKYYYKKYKRKYYWKGTGTATGARK
51    KMMKKCYRYKKKTAMYKKRKKMWWMKYRKKKYRKRYKYYYKKYKRKYYWKRCTCRARGRK
      ***** ****** .  ** **************** **************     : .**

48    YYYWKRYKWKRKMWCTKKRRYRKKWACRWWAAMRKMKWRYSGGTKKYTYYACTYRRAGCW
51    YYYWKRCTWGRKMTYGKKRRYRKKWGTRATMWMRKMKTGYSAACKKYKYYGAGTGRMKMW
      ****** .* ***   *********. *    *****  **.. ***.**..   *   *

48    GYRMYGYYKMKRYKYWTYAMWYYRKSWKKKWKMGMRRGWTKKMKGCCGGYKMKMRTATGC
51    KYRMYKYYKMKRYKTTKCCATCTRKSWKGKTKMKMRRRWYKGMKRTTAKYKMKCGAGGRM
       **** ********  . .    ***** * ** *** * * **   . ****  :.

48    YKMRKKMGCACACCGAKKRMWMAGMWGCGGRYYRYRKYMCCYKKTAKRAKRKRGGCGYKY
51    YTMRKKMKARYGTTARKKRCTMTAMAATAARYYRYRKYMYSYKKWGKGCKRKRAAYKYKY
      *.***** .  .  . ***  *:.* . ..********* .*** .* .****..  ***

48    RKKWKRYRKKRKRKMAKKMRRRRYKKRKRKRYYKKRSSMGTTRWKKWRAAYKKKWSWKRM
51    RKKWKRYRKKRKGKCWKGMRRRRYKKRKRKAYYTKRSSMKYCGWKKWRGRCKKGWSWKRM
      ************ *  * ************ **.*****    *****.  ** ******

48    GACCYWWKRSKYWRRYKASMMKKYMKYGTCTTRYKMYWMMRRYRMKWRKYKRYKRR
51    RTGACWTGGCKYAAAYKRSMMKKYMKTSKYWWGCKMYWMMRRYRMKWRKYKRYKRR
       : . *   .**   ** ******** ..     **********************
```

The pairwise alignments of two accessions 49 and 50

```
49          YYSKWKRWYKYYGKYTTWMRYYMKYYRRMMKKSRKYKYMWRYYTYRKMYYGMWYKRYKYK
50          YYSKAGRWYKYYRKCYYWMRYYMKYYRRMMKKSRTYKYMWRYYGCRKMYYAMWYKRTTCK
            ****  ****** *   *****************.********  *****.***** . *

49          SYGRRRWWAMKMMYMKRMKYYWCARYYWRYYYYWRYWRYRWRSRRYKTWYWWYCYYTGYY
50          SYRRRRWWWMKMMYMTACKYYWYRRYTWRYYCYWRYWRYRWRSRRYGYWYWWYMYYCCYY
            ** ***** ******.  ****  ** **** **************  ***** **  **

49          MWYMARRRWTRMKMRCYYYKRRWMWRWWRMRRGTYYTAMKRCCYGMTARTCRYYYRKGAS
50          MAYMTGARTKGMGCAMYYYTRRWMWRWWRMRGRWYTWTMKRTTYTMYWAYMRYYYRKTWS
            * **:  * . *    ***.***********   *  :***  * *     ******  *

49          KRKYMTRKSYRYTMKWMCCRRMKRYKYYWKMRCWMRAKCYYKGYKYYGWYRKYKSRYRMY
50          KRKYMAAKSYRCYMKWMTARRMKRCKYYWKMRTAMGTGTYYTKYKYTRWYRKYKSRYGCY
            *****: ****  **** .***** *******  * :  **. ***  *********  *

49          YRYWAMYRRWYMRYYWRKKYKTTTCSKWRRKRYRKWRYTTRMWWSYCTTWYCTRMYTCWY
50          YGTAGAYRRWYMRYYWRGKYGYCCTSKWRRKATRKWRYYWRMWWSYYYWACTYRMYAYTY
            *   . *********** **     ******  *****  ******       ***:  *

49          YYKMTAGCRWWKYRWYYWWMYRMRRRAAKYRYGRYWAYKYWWRYMRKYWWWMRMWYSRYY
50          YYKMGGATAAAGTRWYCTWMYRCRARTCKYRYTRYWTYKYWWAYMRKYWWWMRMWYSRYY
            **** ..      ***  **** * *:.**** ***:***** *****************

49          YYWMTMGGYYWKAWRYRKRKYWYGYCCCMRYYRMKRAYRMGGTRMMWRKGKYMKCWTARK
50          YYWMYMRKYYAKRWRYRKRKYWYAYTTTMRYYRMTRGTRMSRYRAMWRKRKYMKMAYRGT
            **** *  ** * **********.*   ******.*. **.  * **** ****     .

49          RMYWRTYRRKKKTCWYKYRRYKTKRRWYWRKKGRTARKKCRCGGCTMSMMMYTMYRARAY
50          GMYWRKYRRKGKYYWYGCAGYKCTRATYWRKKAGYMGGKYGTSRYYMSMMMYWMYGGAGC
             ****.**** *  **    ** .*  *****.     *   .   ****** ** . .

49          AYYWMKGMKYKKMRKRAAKRGTAKCKMKKMKSRRYCWCWAKRKKKWKYYKRWWKYYRYRK
50          GCYWMKKMKYKKMRKRWWKRAAGGTKCTKMKSRACAWTTTKRKKKWKYYKGWWGYYRYRK
            . **** *********  **.:.  * .*****  .*  :********** ** ******

49          RTTKGGKYMRRKKKGAGGCSTTASAYRKKKKKYYRKTMKYKKACYKYAWWYSTCTYYYYT
50          RYWGKRKYMRRKKKKCTATCYCTSGYRKKGGTYYRKYMKYKKRYYKCRWWYSWYGYYYYC
            *     ******** . . .  :*.****  .**** *****  **  ****   ****

49          WKACAKKYKSGMTYWYRRWRKTKCKRWGTTYMWKWKYRSYYKYKKRMKAATYWWMRYKAT
50          ATTYRTKYKSKMCYWYRRWRKKKTGAWKYWYMWKWKYRSYYKCKKAAKTTGYWWMRYGRK
             .:  .**** * ********.*   *   ************ **  *:: ******  .

49          TGYCCTRWRSKWYRRKYKYTGKWCGGMWSTKYYRCAAYKKRMMAKYMMRYYRYRYKKKAM
50          CAYTAGRARGTTYRRKYKYCKKWYRRMTSAKYYRMTTYKKGMMTKYMMRYTRYRYKGKRM
             .* . * *.. *******  **   * *:**** ::*** **:****** ***** * *

49          YWMGTGRYYKRKYRCRRKKCYKSRYYYMKYRKKYRYRYKYRRKKMMYRMKMKRKRRRTKY
50          YWMSYKRYYKRKYRTRRKKTCGSRYYYMKYGGKYRYRYGYRRKKMAYRMGMKRKRRRCKY
```

```
            ***.  ******** ****   ********  ****** ****** *** ******* **

49          RKKYWMMKYAATKMTMMKAYTKARYSMKKYMRRKKMYYYRRKRYYMCRKYMWRMGRGYYW
50          RKKYWMMGYWMKKAKMMGMYCKWRYSMKKYMGGGKAYCYRRKATTCYGKYMWACKRKYYW
            ******* *  .* .**  * * ********   * * ****      ****   * ***

49          YYSWWRKKKATGMKTYYYGYKWYRKYSYGAATYKKTCTKKWYYYCYYKTTCAMKKRWRMK
50          YYSWWGKKKGGAMKKYYYKYTWYRKYSYKGTCYKKYYWKKWYYYYYYKCGAGCKKRWRMK
            ***** ***. .**.*** *.******* .: ***   ****** ***  .. *******

49          YYYYCYKYRYCYKYWCTACWCMYKWRWRRRGTYCKYYKKRMWSMRYYWKKYKKYRCAMTG
50          YYYYAYKTRCTYKYWTCRMTYAYKWAAARRSKCMKYYKKRMWSMRYYWKKYKKYRYGACA
            ****.** *  ****       ***   **..  ********************* .  .

49          YRYMKRYKWSKKKTRRMKKMKRYYKSMKAKGSAMTKWKKRKYWYYGWKYYRRKRKKTGTK
50          TRYMKRYKWSKKKKRGMKGMKRYYKSMKRKRSRCCKWKKRKYWTYKWKYYRRKRKKKKKK
             ************.* ** ********* * *   ******** * **********. .*

49          CYRWKKSCGYAYKWWKRMRYRYRRYWYCTAAYKRWMCTAMSYSWRRRKKGRKKYKKYRYR
50          YYRWKKSMKYRYKWWKRMRYRYRRYWYMCMRYKRAMTWTMSCCARRRKKTGKGYGKYRYR
             ******  * ****************    *** *  :** . *****  * * *****

49          KKMKKRRKKKARYGTKMKKRWGKKKYMYYYKMYYRKKRCGCKRYYSSWMMKKYKKGYRKM
50          KKMKKRRGKKWRYSKKMKKAAAKKKYMYYYKACCRKKRMSYKRYYSSWMATKYKKRYGKM
            ******* ** **..****  .*********   **** . ******** .**** * **

49          MRKYTMYWKWMKCWMTTAYCTKYKYRMKATAYRKWCTRRKKKYYYRRKKYKYYRKWKRYM
50          MRKCACTWKWMKMTCCAWYTCKYKYRMKTCWYRKWAWGRKKKYYYRRKKTGCTAKWKRYM
            *** :  *****    : *  *******:  ****.  ***********     ******

49          MMRSTRYWCRYMGATAYTCYRAYGGCKGGWYRKCTGRKGKMKTKWKCKKWWKKMKRWTKT
50          MMGGYACTMAYMAGYWYCAYRRCKRTKKTWYRTMKTRKKKMKKKAGAKKWWKKMKRWKKK
            ** .      **..  * .**     *  ***. . ** ***.*  .**********.*.

49          KMTYRYKKKRYRYKKYKMRKAGAYKMMRWTMYWTGCRAGKRRAYRYTTYRWKWYCGRRRR
50          KAKTAYKKKRYRYKKYKMRTWRWYKMMRWWMYWWKYRGTKRRGTRTKGYRWKWTTARRRR
            * .  **************.   ****** ***   *. ***. * . *****  .****

49          YCAYWGAACGWGCTGGYRKYYRCTRRKWYRRMYGTAMCMGTAGRYKAATGYWYRGTKYGC
50          YYRYWRWRYRWRMYRRCGKCYRYKRRKWCAGMYRCCAYAACGAAYKRRYRYWYRKYKYTM
            *  **     *       * ** .****   **  .   . .. **    ****  **

49          CWMYRMGTGRCTKWGYTYAMGASGWSTMARTRKYRATKAACACTWRWRYYYKCYYWYCKM
50          AWMYRMRWRRYWKWKYWYGAAGSSWSCARGYRKYRRWKWMYMYKWRWRYYYKTYYWYTKM
            .*****   *  ** * *. ..*.**     ****  *     .******** **** **

49          RRTKRRYWRWRRRYSYKYYRKYMYKTGCCTCKYMSWRRWATWKTCYYCMYYKGGCKRTCK
50          RRGKGRYWRWRRRCSYKYYRKYMYTCTAYAYKYMSWRRWMYWKCYCTYMYYGRKYKRCTK
            ** * ******** **********.  . : ********  **     ***    **  *

49          KKTKYWKMKKYGAYYKYGRYYMTCYMRYTKKRTYKKYGMMKGSKKTGYWMYKSCGKYTKM
50          KGKKYWKAKKYKTYYKYKRYYMWTYMRYKKKRCYKKTRMMGTSKKYKYWMYKSTTKYKKM
            * .**** *** :**** ****  ****.*** ***  **  ***  ******  **.**
```

```
49    RKYMKRWKKTKGAMARYRRAGYRSKKMSTTYKMKCRKKRWWSYRMGYRYYRKYKRKTWYK
50    RKYMKRWKKGKRMMWRYRRCTTRSKKMGYCYKMKAAKKRWWSYGMTTRYYRKYKAKCWYK
      ********* *  * ****.  *****.  ****. ******* *  ******* * ***

49    KTTRWTGCTTTGAYMKTMWYYRYKAATTYRYWRMKRRKWRSYKRCKYGACGAYRYMGKCA
50    GCCRACATYYKRGYMKKMWCCRTKCGCKYRTTAMKRRKWRSYKGGKCKTTATYRYMRKYR
         *  .   . .***.**  * *.. .**   **********  *  : .:**** *

49    WTGGRGRYRYRSKKMRWYGWYYGKKKMMYRKACTTMKYKAATTAARAKGWGKKYGYCYSY
50    WYKRGRGCRCRSKKMRWYKWYCCKTGCCCRKGMGCATCKGTAAGGRRKRWTKKYKYMYCC
      *       * ******** **  *.    **.    . *.:::..* * * *** * *.

49    RYRKKYKYYKRKKYWRMKRCKMRRRKRYRKKRYKRYCWKMKYWMYKACCCKRMYKMYRRK
50    RYRKKYKYYKRKKYWRMKRYKMRRATGTRKGAYKRYMWKMKYWMYKTYYAKRMYKMYRRK
      ******************* **** .  **  **** *********:  .**********

49    RKMWKRSAYKKYYKRYRRKKYGGGYWYYRMRGWRCRSYWWWYYRKWWKMRYMAATMMRKK
50    RKMWKRGTYKKYYKRYRRKKYATKYWYYRMRATGMRSYWWWYTRKWWKMRYMGTCMMRKK
      ******.:*************.  *******.   ******* *********.: *****

49    MCRCMKKKYGCCCTAKWKKRYRYSKRKWYWKGYWKKKWYSYWKKKKMTKKYRMKMKMRKR
50    CYRAMKKTYKAYMAWKWKKRYRYSKAKWYWTAYWKKKTYSYWKKKKMKKGYRMKMKAGGA
        *.***.* .  : ********** ****..***** *********.* ******

49    MCYYKTRMRYYKYRYYRARMYRRRYCYRYYKYRKRMRYSYWKYRYKRKAYKKKKRKYRYR
50    MTYCGWGMRYYKYRYYRGAAYRRRYYYRYYKYRKRMRYSYWKYRYKRKRYGKKKRKYRYR
      * *    **********.  ***** ********************** * *********

49    YYRKGYYKKRKKRYRWKGTCKKMKKRRYKYCCTRAAACCCKRYMRKKKKRWRYKYRKMGG
50    YYRKAYYKKRKKATGWKRYYKGAKKRRYKCYMYRRWRYMYKRYMRKKKKRWRYKYRKMTK
      ****.*******   **   *  ******    *      ******************

49    RWWKYKMWRYRWKCMTRYRYCKKKWYMRRYCYCAGRYMTKGCSKMWYSRKSKCKRTWAKC
50    RWWKYKMWRYRWKTCAAYRYAKKKTCMRGYACTCRRYMCKRYSKMWCCAKSKYKRCWTKT
      *************  : ***.***  ** *.  . *** *  **** . *** ** *:*

49    TKYMYATAKYYYMSKRKYKRMMWRMRKKKKRCRARAMKGGACKWSYWKRKMKKKRYRYKR
50    AKYMYRKRKYYCCCKGGYKRCAAGMRKKKKRYRRRMMKRRGTKWSYWKRKAKKKRYRYKR
      :**** . ***  .*  ***    ******* * * **  . ******** *********

49    KTASAMMMKKKYKYYRMYRRYMKTGCTAKKRRKWYMRYYYRTGRYKKRSMRRKRWRTYSWT
50    KWWGGCMMKKKYKYYRMYRRYMKYSMWRKKRKTTCRYYYRYRGYKKRSMRRKRWRCTGWW
      *  .. ***************** .   ****   *****   ************  .*

49    TKYMAWTKKRMYATYTYYYCCMYRYRYRKTKKKKACTGYRYGYCKMYYRYYYTTRAYYYY
50    YKYMGTCKGRMTWYTWYYYTAATRYRYRKAKKKKRSCCYRYATTKMYYRTYYCYAGYYYY
       ***.  * **     *** .  ******:**** .  ***.  ***** **   .****

49    ARKAMKYAYGGWWRCGTRRYRRYMMYRRYYTAMYYKTKYWYWCRMMRYWRKKMYYWWRSR
50    WATTAGYRCRRWWRMKWRRTRAYMMYRRYYCGMYYTKKCTYWARAMRYWRKKMYYWWRSR
        .:  *    ***   ** * ******** .***..*  **.* ***************
```

```
49              KKRRRSRGCRCKCRCRYCTKWKTKWRKYYWRWKGKWRKWKKRAWRWMGKGWKMKWWYMKS
50              GKRRRSRAMRGGSRYRYACKWKKTWAGCTWRWGRKWRGWKKRGAATMCKCWKMKWWYMKS
                 ******. *  .* **. ***..*    ***  *** ****.   * * **********

49              WYRYRKSMYRRTGKKKATTYWMYCTAYKKKYKKKKKWWRRRRGRTTTKGKRKKKKWYKWK
50              WYRYRKCMYRRYRKKTGCYYWACYKWYKKKYKKKKKAWRRRRRRCGGKRKRKKKKWYKWT
                ******.****  **..  **   . ********** ***** *   * **********.

49              YRWKMRRRWYYYMRYRRMKSWRMYRCKKRWMYYKKACACKRRMKWYKKKAMKRKCAAATA
50              TAWKARRRWYYYMRYRRMKGWRMYRTKKRWMYYKKTTGAKRRMKWYKKKWMKRTTTCCAT
                  ** **************.***** *********: ..********** ***. :..::

49              YRYYKWYRSSMYGRYWKKTRAAKRTTMRYYRYKYRRRYYRWRYRGTRRMYKKKRRYYKKY
50              YRYYKWYRSSMYKRTWKKAATGKAYYMRYYGCTCAGRTTGWRYRRYRRMYKKKRRYYKKY
                ************ * ***: :.*   ****  .   *   ****  **************

49              KMCAYYSRRYGCTTKSYKRWSRWRKMRWMKKRRCGKKYSGRKKKYYYYKYRSYKMRRKMK
50              KMSMYYSRRYRMYYKSYKRWSRWRKMRTAKKGRTRKKYSRRKKKYYYYKYRSYKMGGKMK
                **. ******    *************  ** *  **** ***************  ***

49              TCRRRGAYYKKRWSMMKKKYRYSRCCGGRRKTCKKKACRYKYRATTACKYAYYKRYYRMK
50              CARRRTGYYKKRWSMMKKKYRYSRTTKRRRKKYKKKGYRYKYRGCCCTKYRYYKRYYRMK
                 .*** .*****************    ***. ***. *****.  . ** *********

49              MRWACTCYRYSKKMKKRWKMKKKKKMYYWYWSRMYRMRKWKWKRKGGTCYGMKSMRCKKK
50              MAATAAAYRYSKKMKKRWKMKKKKKMYYWYWSGACAMRKWKWKRKAACYYRMKSMGYGKK
                *  :.:.*************************    *********..  * ****   **

49              GMKWRYRTKYKYYKYYSYYRRYGYGKAATKKKGMGRRYKYKCCKKYKRKKRRKYMRSYWY
50              TMKWRYGYKTTTCKYYSYYRAYRTTGTCGKKKRCRRRYKYKAATKYKRKKRRKYMRSYWY
                 *****  * .  ******* *    :. ***   ******...****************

49              YTGTCMRRMYKWRRWGYKYGTYRKKRMRTKYSWWRMKYRARKYRYMRRKGTYYAATAYAM
50              YWRYTCGRMYKARRWRYTYKYTAGKRMRKKYSWWRMKCRRRKYRYMRRKRYYYRRYRYMM
                *      **** *** *.*     ****.******** * *********  **    * *

49              KWMCYYKYRKKKWWCCCGKRYYGWMWK
50              KWMYYYKYRKKKWWYYYRGRYYRWMWK
                *** **********     *** ****
```

The pairwise alignments of two accessions 49 and 51

```
49              YRYYKKYWMRTYYYTWRYRWWKKGGCWKYYWRYRKCRYWCGGWMASRKRAWRRGYYYYYY
51              YRTYKGCWMAAYTTCWRYRWWKKRRYTKYYWRYRKYRTWTKKWCWGAKRWWRRAYTTTCT
                ** **  ** :*   ********    ******** * *   *  . ** ***.*

49              MCCCKYWAMYMGYRWYTSKYRWRWMKATTWYKSYGYGGAMMRYWMWMMCCTAWKWWKWRY
51              AATTKYWRMYMSYRWYYSKYRWRWMGWGCACKCTKYRKGMMRYWMWMMAMKGWKWWTWRY
                 .  *** ***.**** ********      *.  *  .*********. ..****.***

49              MKMYWRAWGATRRRAWWRKMYKMYKRKRYKKSWMYYWRMMYRYKARWWYYYMYKYWKWGM
51              MGMYWRWWAGCARGGWWRKMYKMYKRKRYKKSWMCYWRMCYAYKWRWWYTCMYTYWKAKM
                * **** *..  * .******************* **** * ** ****  **.***  *

49              KSMRCGRCKRWYKKWCTGKYKSRKYYYWKRYWAGATMKYMRYKWYKMKKTYYGYWGRGYR
51              KSMRMRAMGRWYKKWMYRKYKSRKYYYWKGTATTCCMTYMRYKWYKMKKCYYRCWRRAYR
                ****     ******   ***********   : . *.*********** **  * *.**

49              WKCYRRWWCATWYKAMWKYKYSRRKRTKRRKWGRYCYRKWYKWWCAYYRRAYYMWRRTYR
51              WKYYRRATTGCWYKTMWKYKYSRRGRCKRRKWTRYYYRKWYKWWMRYYRRWYYMWRRYTR
                ** ***   . ***:********* * ***** ** ********  **** ******  *

49              AKKYKYYMMYKWKMKRCYYYSRYGCKYRKRMKMMKKYYMYTYMWTRCRWRKYYRYRKYKW
51              RTKYKYYMMTKWKMKRTYYYSRYKYKYATGMKMMKKYYMTWYMWGGTRWRKYYRYRKYKW
                 .******* ****** ******  ** . *********  ***   *************

49              YYRYYWTYMRRTTCTTTTGCYCKTWSWMMKKYKKAKKSKWGGCMARRYRYGRCGCYRKKM
51              YYRYYWYCMRGACACAGCAYCTKWWSWMMKKCGGGKKGKWRRYCCRRYRYKGTKTTAKKM
                ******  ** : . :  .   * *******   .**.**    .*****       ***

49              KKKWRWYKKKRMKRKMGKKKRYYKRRKYSKKYKYMRKRKMKKMGRRKMRWKRMYKGYCGY
51              KKKWRWCKKKRMKGKMKTKKRYYKAAKYSKKCKYMRGRKMKKMRARKMRWKGMYKRTYTY
                ****** ****** ** .******  ***** **** ******  ****** ***    *

49              RRKSWTMYKKGTKKYMAKRYAYMMRYWKMRKMTMKRRRYYWRKYWWYYKRKYKKKRRRGS
51              RRKSWYMYKKTCKKYCCGRYGCCMRYWKMRKAYMKRRRYYWAKYWWYYKRKTKKKRRRRS
                ***** ****  *** . **.  ********  ******** ********* ****** *

49              RKKARMYRRYTGTKKTWKYYCYYKWCKRWAKMWMKWGYYWKKCCTAAKKKKKWWKKYTMS
51              RKKTRMYRRYCKYKKWWKYTGYCTWYKRWCTMWMKWKTTAKKTGGCRKKKGKWWKKYKMS
                ***:******   ** ***  * .* ***..*****    **   . *** ******.**

49              KWRRMWACYRYKKYWMRRYAAACKWSKGKKKKKWWCWRRKKRKKKKRYRKKGMKKKTWKY
51              KWARMATYYRCKKYWARRYGWGTGWSKRKKKKKWTYWRRKKRKKKKRYRKKKMKKKCWTY
                ** ** : ** **** ***. .  *** ******  *************** **** *.*

49              WKKGRRWYYYYACCMRKYRKMRYGRYRYAKMYRRSRWKMYYKSKYYCCKRMKWSKKWKMG
51              WKKKRRWYYYCGTTMRKTGKMRYRRCAYWGMYRRSRAKMYYKSGYTATKRMKWSKKWKMK
                *** ****** .  ***  **** *  *  ****** ****** * . ***********

49              AYYRYKRRRGMYYRYKYKYTKRRYSKYWGRYYRYGCKKRAGSGAGKYYRKYRKMMYKRKG
51              GYYRYKRGGRCYYRYKYKCYKRRYSKYWKRYTAYKTKKRWACTWRKYYRKYRKCMYKRKT
```

```
        .******    *******  ******** **  *  *** ..   ******** *****

49      KKYGCYKMKGSYYGGRKRYKYAMGRKRWRYRRYKYRAMYWYKMWYGACSAYWKYYAKYRY
51      KKYRMYGAKKCYYAAAKRYKYGMARKRWRYRRYKYGMMYWYKMWCACAGRCAKYYMKYRY
        ***  *  * .**.. *****.*.***********  ******* ....   *** ****

49      YMRYRMRGRAAYSKYKSKYSGRAKKASKYKRAKYMYYKYYMSYYMCAKGRRKGYTARKYW
51      YMRYRMRAGCGYSKYKSKCCRARKGRSKYKRMKYMYYKYCCSYYCYMGRARKACCGRKCA
        *******. ..******* .   *  ***** *******  ***      **.  .**

49      TRGCGKKKSKYKKYMRCTRKKACKATATKKKKRCWKKWYMYRKMWKRYCTKMKKKAATYM
51      YGRMRKKKSKYKKCCRYWRKKGTKTATCGKKGGMWKKATMYRKCTKRYMWKMKKKWRYYM
             ********  *  ***. *:::  **   ***  ****  ***  *****   **

49      YRRRYYRMKKRKKYYKKKKYRRAMRRKYMRWYRRTKTTCCYWKYYGYRKTMWKYYYMRRY
51      YRRRYYRMKKRKKYYKKGKYRGRMRRKTMRWCRGCGYWMTYWKTYAYRTKMWKYYYMRAY
        ***************** ***  **** *** *       *** *.**..******** *

49      GYCRRTTATGSYKKYYWGACTRYCAWYTRRSKCTRKWCTTRKKYSMKKKAKKAGCRKGWT
51      RTYARKAGWRGTKKYYWRMSGRYYRWYARRSKMYRKWGYYRKKCSMKTGWKKTTARGAWG
            *.:.  . *****  . **  **:****  ***   *** ***.  **: .* .*

49      GWASYGYSMGKYYKWRRKKYRKRYMMRRWYKAACTGGKMAMWCTRRTGRATRSKYKASCT
51      AWGGCACSAAKYYKWRRKKYRKRYMMGRWYKTCGWKTKMGCWTGAAAKGMWGSGYKWSSA
        .*.. . * .**************** ****:.    **. *    :     * ** *.:

49      CKYRWYRYKKYKYAWWMCKWRCAYKWWKAAGTATRWRKMMYYTSKGKAAYYKYAKMRKAW
51      MKYRWYRYKKYKCRWWMTKWGATCKWWKRWRYRWRWRKMAYYYSKKKWTYYKYCKMRTGW
         ***********  *** ** .: ****      ***** ** ** * :****.***..*

49      YRATCTKKWRRYYRAGRACCYTCGKYAGYYTTRRYYYAGWWAAMYRKRRAATCYRRMMYR
51      TRTKYWKGWRRYYRRKRGTSYATTKYGTCTAGGRYTYGRATMWMYRKRRWWCYTGGMMYR
         *:.  * ******  *. .*:  **.   :  ** *.     ******       ****

49      AKMWWAGTRRRKKGGYKMWCYYGYRTYRWKKAGYCMMYMYWYYYSSKWYRRKKMTYYGCA
51      GKCTWGTYAGAKGRKCGCTTCCRTGACAWKGGAYYAAYMYWYYYSGTTYRRKKCKYYRTW
        .*  *.     *             :  ** ..*   ********.. ***** .**

49      CGTYKGCGTTTGCAAAWYYSYGKRYWKKRGYARWKMRYWCRWRYRCMAAACKTKRWWMRG
51      MRKYKAAKYCCATGGWAYYSYKKRYWKKRKYRAWKMRYTYGWGCRYMGTWTKAKGAACAT
          .**..    . ..  **** ******* *  *****   *  * *.:  *:*

49      CGRTAKCWCYRRAGCTATACKMGMWYRYAKWYKGSYRWRWRGYAGTTCACGKAAAGKKKR
51      GCRYRKMTATAGWRMWGCWYKMKMWYRYRGWYKKSYRWRWRKYTACYYMMKGGWCRKGKR
          *  *  .       .   ** *****  *** ******* *:.       . . * **

49      YRRRKKMKMKKMKRKRRKKMMRKWWMMRKSKRKMKRYSMYKYRRWKYKKWCCAAYYMAYT
51      YRRRKKMKMKKMKRKRRKKAAGKWWMMRKSKRKCTRYSMYKYRRWKYKTTAYRGYYMWCC
        *******************   *********** .*************. .  .***

49      RYAYMWGRMCAWYYCGTAKYTYYWRRGKMRYATYTMGRKAGATKRWKKYYSYYYRKKYRY
51      GCRYMWAACYGWYYAAAWKYKYYWRRAGARYCKCYMKGGCACKKRWKKYYSYCYRKGYRY
           ***.   .***..: **.*****.  **..  *   ....********* *** ***
```

```
49      YYYYKKKRKMMRRKKYYRYKKRRYKWKMWWTAAKATARMYTRYRCGRCGKSRTMYMKYKS
51      YYYYKKKRKCMRRKKTYRYKKRRYKWKMWWGGGKRKMRMCKRYRYRRYSKCGAMYMKYKS
        ********* ***** ************** ..* . ** .***  * .*. :*******

49      WRWCTCGGTAAYMRYAYRWYYRMRMYKKKAGYRYRYRMTCKYYTTMGMMKKKRMAGCCGR
51      WGTTATCACCCYMRYMYRWYYRMRMYKKKWKCRYRYRMYYKYYAAMKMMKKKRMGAYMTA
        *   :  . ..**** *************   ******  ***::* *******..

49      GRYGTGCYTTMRRTWGAGTTWSRWRMRKKRWRKKATYCKRWRRYRYCKKKMYWYMRRTMK
51      KRYCCATYCACGRWWKRRYYWSRARMRKKRAGGGRWYMKRWRGTGYYGKKMYWYMRRGAK
         **  . * :  * *     *** ******      * ****   *  *********  *

49      KMRRRYYWYYKKYYRKYYMYRYRYGAWKRRKRWKKGRYRKWKKYYWMKCAARRTAAKYSC
51      KMRRRYYACYKKYYRKYYCCGTATTTWKRRTRTKKKRYRKWGKYTWMGMGRRRGGGKYCT
        *******  *********       :****.* ** ***** ** ** .  ** ..**.

49      CTSMRTMKYYRSTKKYKYCKWWYSAYYMWMYYGYYWWYKKTTAYCYWYRRWTKKYKRYRS
51      YYSMRCMKYYRSWKGYKCYKWWYSMYCAAACYTYYWWYKTACGCYYWYRRWGTKCKRYRS
          *** ****** * **  ***** *     * ******.: .  ****** .* *****

49      YYKWYYRKKTTTAGYTRRYWWAYAKKYSKRCTCACTSGACMCCYRMKKRMMRMKTTGSCA
51      YYKAYYRKKKYYMTCGRRYAAWYGGGYSKRACGTYKSSCTMTTYRMTKRMMRMKACAGTC
        *** *****.      ***   *.  ****.  : .*.. *  ***.*******: .. .

49      GMKKKKKMKAKKRKYKYRRYGRKCMRATWRRWYRYAYRRWKKMYSMMMYRYKKRWYMYYR
51      TMKKKKKMKRKKRKYKYRRYKRGYCRRYWRRWYRCGYRRWKKMYSMMMYRYKKGWYMYYR
         ******** ********** *   *  ****** .***************** ******

49      MKKKMRKRRKSKMKSRYRWKRCAGRKMYYWTYYWKRKKYYKKARMKYKKRTYYCYCCCKW
51      MKKKMRKRRKSKCTSRYGWKATGTRKMYYWKYYTKRTKYYKKWRMKYKKRWTYYYATAKW
        ************ .*** **  . ******.** **.***** *******  * *. .**

49      RWRRRRKYKRKKWSCMRYYWKKYKRKYRRYYRYMSKKKKKKSMKRYMYKKYRYYSMKAAK
51      AAAARAKYKRKKWSYMRYYWKKYKRKYRRYYRYMSKGKKKKSMKRYMTKKTRYYSMKMRK
            * ******** ********************* ********** ** ******  *

49      MMRWKRKKYYASYKRWSRMYKAYYWRKRKKKMKMTYRWKKRKSKACKWAMSWCAAATAYR
51      MMRWGRKKYYRSYKRWSRMYKMYYWAGRKKKMKMKYRWKKRKSKTTKWRCCWTTCCATYR
        **** ***** ********** ***  *******.*********: **  .* :..::**

49      YYWYRSSKARYWKKWYKRTYTKMGMYRYYRYKRRYRKRGCTCMYKKGYYKYWKTCCMYYK
51      YYWYRSSTMRYWKKWCKACTCKCKMTRYYRYKRRYGKRRMYMMYKKRYYKTAKATTCCCT
        *******. ****** *    *  * ********* **    **** ***  *:     .

49      MMCMRYGAKKRAGKWRRRRYMRKRACWCKYYKKYYKYKYMKKMTCGKRRRRGKKRYMRYY
51      MMYARYRRKKRRKKWRRRRYMRKRGMAYKYYTKYYKYTYMKGMCAKKGGRRRKKRYMRYY
        **  **  ***  ***********.   ***.*****.*** * . *  ** ********

49      YWYWKKRKYKTTRKCKKGKAYKKRKSYWYYTMRRMKWRMWWSWKKAGACRRSMKAGATKK
51      YAYWKKRKYKGARKYKKRGRYKKRKSYWYYYCGRMGARAAWSWKKTCGYGRSMKRRRKKT
        * ******** :** **   **********   **  *  *****: .  ****   .*.
```

```
49      RYSWRYYMGKKSRKKWAMMRYYKYKWYKKTCKKKKYMYWKKRYRSYWCWSMWKAYKYKAT
51      RTGAAYYMRKKSRKKWGMMRYYKYKWYKKWYKKKKYMYWKKRYRSYWTACMWKGYKCKWY
        * .  *** *******.************  ****************  .***.** *

49      TAGMRYRRKSKKCTMWRKGSRYKMGCSYRKCTWWSTCYCMCMYRSKWYRGKKKTKKRKTM
51      YMRARYRRKSKKYGATGKKGRYKMKMGCGKYYWWSKYYYMMMYRSKWYRKKKKYKKRKAC
            ********     * .****  .  *  ***. * * ******** *** ****:

49      AYRYMKYGCKRWCRGRGGRYRYKMKMRWCATGCMSRTMKYYYTCCATWYRYRKGTCYYYK
51      MYRYCKYATKAWYRTRRRRYRYKMKCGAMRKSMMSRKMKYYYCGTGCTYRYRKKCTCCYK
         *** **. * * * *  *******     .. ***.*****   .  *****     **

49      RKKRRKKRMYRWRKTYMYYYKKMYRRMKKRYYKKSSAKYYMMKYKKYKRKKMMRRKWRTK
51      RKKRRKKRMCAARKWYMYYYKKACRRMKKRYYKKSSWTYYMMKTKKYKRKKMMRRKWRWK
        *********   ** *******  ************ .***** ************** *

49      TGKRMKKRRYCTKMMYRMKYRKWRKATKKKYWWCKKRKYCKYGWKCRMGCGRYYYYYSAM
51      WRKGMKKRGYTCKMAYRMKYRKWRKGGKKGYWWTKKRKYAKYKTTYRMRYTGTCYYYCMC
          * **** *  ** **********. ** *** *****.**  . **      ***.

49      GRRKMYCYTRRRRKKTKYRYKYYRYYYYYWKAWRRRRRCKYMYRKGARKRTACTYYRRYY
51      TGRGAYYCYARRRKKYKYRYKTYRYYYTTWKRWGGGGRTGYMYRTARRKRATTCTYRRYY
          *  *    ***** ***** *****  ** *    *  ****.. ***::   *****

49      KKWKSKGKTKWYSRMRRWKKKCKKATGYYCCRYRKYAKKYYRYYRMKMACCYYRGCCKKK
51      KKWKSGTGYKWYSRMRRWGKKYKKRKRCYTYAYRKYRKKYYRYYRMKMTAMYYRRYMKKK
        *****    ********* ** ** .  *   **** ***********:. ***   ***

49      RWKMKKGGTAYKRMYYYWYKKYKRMYRMTATKWKARKYKKRKGMWRCWYYKYTKYMWYSY
51      RWKMKKKAARYKRMYYYWYKGCGRMYRMAGGKWKMRKYGKRKKMWRYWCYKYCKYMWYSY
        ****** .: **********   *****:. *** *** *** *** * *** *******

49      GTKYRMRYRMTWYRYYRRKRKKATCYKAKRKYYWYKKRGATMKRYGGGYYKYCKCKKKRY
51      RCKYRCRYRMWWYATTRRKRTGTATYKCKRKYYTCKKRAGCAKRYATRYCGYMGYKKKRY
          *** **** **   ****. :: **.*****  ***..  ***.  *  *   *****

49      AGCRRRYMMKYRWYRRYWKKAAAYCRAYKKTYWMSYGGTKRWRTAKS
51      GTTRRRYMMKYRAYRRCWKKTWTCYGCTKGWYWMSTSKWKGAGKGTS
        .  ********* *** ***: :   . *  **** .  *   ...*
```

The pairwise alignments of two accessions 50 and 51

```
50                SRRRRRYYMYYRTTWMKWKYRYKRWGYMRKKWYMRYYKYKSSYRYKRKRYYWKWYMYYMM
51                SRRRRRYYMYYRGGWMKAKYRYKGWKYMRKKWYMRYCGTKSSYRYKRKRCYTKWYMYYMM
                  ************  *** ***** * **********   ********** * ********

50                YRTTRRCGGYTYYYRRYYRKRRWWWRKAAWRYRMYKYWARCRCKYSYYYKRRRWWCAYKR
51                YGCARRTATYGYYYRRYYRKRRWTWRKGMWRCRMYKYWRRTAYKYSYYYKRRRWWYCCKR
                  *  :** . * ************ ***. ** ****** *   ************ . **

50                RYWYWKTWRWRCAYMGMMYWWRWKWWCMAYYAWYKYRWRRRCKRKRWRTYYYYKSKYAYK
51                RYWYWKAWRWRAGYMRMMTATRWKWWYMTTCRWYKYRWRRRSKRGGWGCTTCTKSKYTCK
                  ******:****..** **   ***** *:   *********.**  *      ****: *

50                CKYRYWCCAWRYRYWYMRCCGCTCKWMKRRRKMYSRARTYYRKMYGRRWYMWRYYYWRYR
51                YKTRYWMYWWRYRYWYMRAMKYWYKWMGARAKAYSRGAAYYRKMYRRRWYMWRYYYWRYR
                   * ***   *********.     ***  * * ***. :****** **************

50                KKKYCYYYYWRYYWKRKMKYTCWKGMWKYRYGYRYMMGMKTTCRKWYRYMMRCTARKRYY
51                KKKCYYYYYWRYYWKRTMKYWAWKRAWKYRYTYRYMMKMGWYYRGWYRYMCGYCMRKRYY
                  ***  ***********.*** .**  ***** ***** *    * *****     *****

50                KWTTTYYMKKRYRSKTYRMWKRRYYKYKYRKYKKYRYSKKYKMRYYKACYRCAKWKMKYR
51                KWWWKYYMKKRYRSKKYRMWKRRYYKYKYRKYKKYRYSKKCKMRYYKGMYRYWKWGMKYR
                  **  .**********.************************ ******. **  ** ****

50                KYSWGYRRRKWRTTTKKGRRYKRYWAKRYRRYRMRRYRRKMWRRYKMKKMYAKRKTTGYM
51                KYSWTYRRRKWRCWGKKKGRCKRYWTKRYRRYRCGATGAKMWRRYKMKKMCMKGKACKYA
                  **** *******   **  * ****:*******      ***********  * *:  *

50                WRRKYSYMKKMTAYRMRRRYRRTRRKMYMYYMRTTACKRYKYKAATTKYSSYKKGTTKMY
51                WRRTYSYMKKMGRYRMRRRYRGCRAKMYMYYMRWYRYKGTKYKCGGYKYSSTKKRYYKMY
                  ***.*******  ********  * ********    *  ***..  **** **   ***

50                CKYGCYYWGKYRRYTYGCARGTYRKRTSYTYYKCACCTCCTACKMKYMRYSWWKRMKWKY
51                TGCATTYACKYRRYYYTTCRTWYRKGCCCKYYKYGTYYYTACTKMGYMRYSWWKRMKWKC
                     .  *  ***** *  .*  ***  . .*** .     :. ** *************

50                GRRKKRKKKTCAYRCAAYRYYRKAYMSSGTMGRGGGYAYKTYYRRKRKYKKKMRYYYKKK
51                SGAKKRKKKYYMYRYGTTRYYRKRYMSSAWCARRAAYWTTGTYRRKRKYKKKCGYYYKKT
                  .  ******   ** .: ***** ****.  .* ..*  .  **********  *****.

50                KCYCKYGYKKKGKKKYWKMYRYWYYWTTAYRYKRRYYRKRYKKYRARKYWKKRCTYTWGK
51                KMCMKYAYKKKRGKKYWKMYRYWYYTWWGCGYKRRYYRKRYKKCARRKYAKKRTCTCAKT
                  *   **.****  ************   .  ************   *** ***      .

50                KKWRRRYMRAYKYYRKKAWGCTAGKCRYRYKKAAYYTRGYGCCTWCCGTGYCTYKKWRKY
51                GKWRRRYCRMCGYYRKKTTATGMRKTACAYKKRRTYYRAYATAAAMTCCRTYCCKKWGKC
                   ****** *   *****: .    *    ***   * *.*. .:          *** *

50                YKACCCTYCTYWTTYTTCKMYKKRKRRMKYYKRTMYMKKYWRKKRRWKRRKYRMRKYGAK
51                CTMMYYYCMGYWCCCCAYKMYKKRKRRMKYYKGYMYCKKYWRKKRAAKRRKYRMRGYKCT
```

```
            .        **    : **************  ** ********  ******** * ..

50          YYAGKKWKKKWYMRKSKCKGSMACKKMYKAYYWKMWRTATCCCKYRYSKKWYGYTMYSWY
51          TYMRKKWKGKWYMRKSKYKKSMTTKTMYKRYYWKMWRKGWYYMKYRYSKKWYAYAMYSWY
             *  **** ******** * **: *.*** *******..    *********.*:*****

50          YKKKYKKYTRKKMKRRRWKATRAKMRRSRTGKWTRYYKYRKKTYRRKKKWYSMMWKTKCK
51          YKKKYKKYWGKKMKRRRWKCGRGGMRRSRWAKWKRYYKYRKKYTRRKKGWYSMMWKCKMK
            ********  *********. *. ***** .**.********  **** ******* * *

50          KWRRYRKKYRRWRKSRTCMASRYYTWAKRSMCYMYRGKRTRKRKWKKKRMYYYRAKKKKM
51          KWRGYRKKYRRWRKSRAYCGSRYYKWMKRSMYYMYRKKRARGRKWKKKRMYYYRMKKKKM
            *** ************:  .****.* **** **** **:* ************ *****

50          KRYKKWYYYKKKKMMRGTKMKRWMAWGKGATATCRRKYSKKYRYARRCTGGYWMKKYRRK
51          KRYKGWYYYKKKKMARRKKMKRWCGTRKKGCGGSRRKTGKKYRYGGGAGRRYWMKKYRRG
            **** ********* * .*****  . * . . .*** .*****.  .   ******** 

50          RYKKYKTCYRAKKCGKKRWWKKKMYRKKRYYKYKKKKWRYKKKKKAYMYRRYRMKKYCAG
51          RYKKYKATCRMKKMRKKRWWKGKMYRKKRYYKYKKKKWRYKKKKKTYMYRRYRMKKYYWA
            ******:  * **  ****** ***********************:***********  .

50          KRMKYKMMRRRYYYKMYKRRAGYRKWMCARMYRYKRCTKMRYKKKWYKMAKKSRKYRWRM
51          KRMKYKAMRRRYYYKMYKRRMKYRKWMATRMYRYGGTGKMRYKKKWYKMRGKSRKYRWRM
            ****** *************  *****.:*****    ***********  *********

50          RKWWGWMYMMKYRGACYYWAMKWYWGYKRYRKKKTRKGYRMRRMCACSYKWRWKYKKYRR
51          RKWWKWMYMMKYGACACTWRMKWYWAYGRYRKKKCGKAYRMRRMYMYSYKWRWKYKKYRR
            **** ******* ...  * *****.* ******  *.******   *************

50          KCAGMMYYRWRWKKKKYKRGGARYKAGCCRYMKYYRCKTWRKYKRMRWKYKKYYCYYWTK
51          KYWKMMYYRWRWKKKKYKGKATGCKWKTMGYMKYYGYKKWRKYKRMRWKYKKYTTTYWKK
            *   **************  .:  *     *****  *.**************   **.*

50          YKSKTAWRYMYTKMRYYRWKYKRKGTTGRMSTKKSTYMWKGKKKRMKKKKKMKMWYKWSY
51          YTSKWMAACCCYKMRYYRWKYKRKACAARMSYKKSWYCAKRTKKGMKKKKKMKCWCTWSY
            *.**        ************. :.*** *** *  * .** ******** * .***

50          KCATCYWKRGYTARYYYYYRWKYWKMKRYYRKYWKYSYTRCRGKAKKKRKRKRGYCCKKK
51          KYWWMYWKRKTYGRCYCYYRWKYWKMKRYYRKYWKYCYKRYRTKWKKKRKRKRTYSMKGT
            *    ****   .* * *******************.*.* * * ******** *. * .

50          MMWKGWYCTYTCGRRYRRGCAKRKCCAYCCTTKKGRKKTYYRYKKRGKYRKWKSMYYMYY
51          MCWKTWCTCYWMRRRCGRSACKRKTTRCGTGCTKTRKKWYYRYKKARKYRKWKSMYYMYY
            * ** *   *   **  *...***        .* *** ******  *************

50          YYGYKSWAAYYRSRRRWKSYKKRKMKMSYWYKKYTACKMMKRYKARYTYYKKKMYMMKKM
51          YYKYKSWTTYCRSRRRWKSYKKRGMKMSYWYKKYCCMKMMTRYKGAYGYYKKKMYMMKKM
            ** ****::* ************ ********** . ***.***. * ************

50          RMKCKKKWKAYMMKYAYKWMSYYMRRMYRKYRKKYKWRYYRMKKMYWRYRWSRGAYYMRK
51          RMGMKKKWKTYMMKYGYKACGCYARAMYRKTRKKYGWRYYRMKKMYWRCATSRKMYYMRK
            **  *****:*****.**  . * * **** **** ************   **  *****
```

```
50      MYRMWYYRYWYYRMTACATRKMMYKWMMAYKKYKRYYRWYKKRYYMCYYMYAARMKRMCR
51      AYRMWYYRYWYYRMKCYTGRKMMYKAMMRYKKYKGYYRWYKKRYYCYYYMYGCRMKGCYR
         *************.. : ****** ** ***** **********  ****..***   *

50      RRWYRYKSYWKRMYRMWKWKSGGAYCKKRYYRCTRTMWKKRRWKRYYRKYYYKRRYSCTG
51      RRAYRYKCTWKRMYRMWKWKSRRMCYKKRYYRGYACAWKKRRWKRYYRKYYYKRRYSMYR
        ** ****. ************     ******     ********************

50      CCRYMMKCKKKYRYMKKYKYGMMKTCAWKKMKYYYKRMYRYCKKKKKWWWSYKRRKRYSK
51      MMRYMMKMKKKTAYMKKYKYKMMKKYMWKKMKCYTKRMYGCYKKKKKWWWSYKRRKRYSK
          ***** ***  ******* ***.  ***** * ****   ******************

50      RYMYAKYWYMMWATYKWKKKCGRYWMYTWCKTTGMGKWMMYYRWRKKYMKKAKYWTGKWK
51      RYMCRKYWYMMWGCYKWKTKTTGTTCTAAGKAAKARKWMMYYRWRKKYMKKRKYWWRKWK
        ***  *******. ****.*       :  *::   *************** ***  ***

50      RYMRWKRRKWKRYKRYRKYYKRRKCYTAMAKRWKWRYSSARMMYYSKMWSYKYRWYRGRS
51      RYMRWKRRKWKRYKACAGYTKRRKSCWWCTKRWKWRYSGCAMMYYGKMWSYKYRWYRKAC
        **************    * ****.    :********.. ****.***********  .

50      YYYRKYAKKKTKWYYKKWKTWRWRRARWYKAKRWKTCRWGTSMCYRSYAKKGYRKYYYYT
51      YYYRKYGKTKKTWYYKKWKWWRWRGRAWYKWKRWKATRWAYSMGYRSYMKKKYGKYYCYK
        ******.*.*..******* ****   *** ****: **. ** **** ** * *** *.

50      CKKKYKKWMWWMRYWRKMRYYGRRMWWMWGKGRYYKWYTKKYKYYYKKTRKGAARGKYKY
51      AKTKYKKWMWWMATWRKMRYYKRRMWWMWKKKRYYKWYKKTYKYYYKKKRKTMRRRKYKY
        .*.*********  ******* ******* * ******.*.*******.**   * ****

50      KRWWWYRYMWRRKWTKTMRKGTRSTYKKYYWYRRKKKKWCAYYGKAYKKTAYYKYYMCAY
51      KRAWWYRYMWRRKWKKWMRKRKRSCYKKCYWCRRKKKKWYCYTAGTYKKYRYYKYYMACY
        ** ***********.* *** .** *** ** ******* .* . :***  ******..*

50      CCRRYYWYYWRSKKRMKTMRCRKCKCTTMRRTWAWMWMTKYMWRMYRWRYMYMMSMMWRK
51      TGRRYYWTCTASKKRMKKMRGRKYKSKYMRRGTRTMWMWKCATRMYGTRYMYMMSMAWRK
          *****    ******.** ** *.. ***    *** *   ***  ******** ***

50      MKRYKGMKKRGYYWTAGYGYRKRWKRKYYRKRKKWYMKMKMYRSYKRRRMKKWWMKRYGA
51      MKAYKKMKKRTYYWCTTYKYRKRWGRKYYRKRKKWYMKMKMYRSYKRRRMKKWWMKRYRR
        ** ** **** *** : * ***** *********************************

50      CMGKAGRCYKGYRRYWMKKYWWCCGAAKYRKRSRSYMKYRKKRKRMRKGTKMKKYKRWKY
51      SMRKRKRMYKRCRRYWMKKYWWYTARGKYRKRSRSYMKYRKKRKRMRKACKMKKYKRWKC
        .* *  * **  **********  . .*********************. *********

50      KKRYYYWMWYTYKWWKYMMCRYKYTTKCGTATCYRGTRYSYSRRCMGKMKWMMCAGMKGK
51      GKACTYWMWYCYKWWKYMMSRYKYAKKYRGTYTYRAWGYSYSRRYMRKMKWMMMMTMKSK
         *   ***** ********.****:.*    :  **.  ****** * ******   **.*

50      YRYYCGMMSRKRKKWKRKKRTGMYYYKWRKYKYWYRMKWMKYKWMMRRKAAKKSRWKWRR
51      YRYYMRMMSRKRKKWKRKKRWRCYYYKTGTYKYWYRMKWMKYKWMMRRKGGKKSRTKWRR
        ****  **************   ****  .*******************..**** ****
```

```
50              KKYKARAGWRRMTCATKKWKKWWYSRYGCKKYMYGGTWKWTWYYRYRKKYKGGWTKKMKK
51              KKYKGRTRAAAAYYMCKKTKKWWYSRCRAKTCMYTSYATTWTYYRYRKKYKAAAAKKMKK
                ****.*:         ** *******  .*. ** .  .   *********.. :*****

50              AYKYKATRTMWKRWMRWWKRMCMWKKACCKTKCWKGGTYRTGCCTRYYWCKTKKYYRRMM
51              RYGYKRWRKMWKRWMRAAKRMYMWKKWMYKAGGWGAAATGATTYWRYYWYKKKKYYRRMM
                 * **  *.*******  *** ****   *:  * ..:  :    **** *.********

50              ATGAAGKRYMYYYYMKRRYYRAMTAARMRMKMYAGTAYRRKKYRYYYMKRRRKRYTTCMC
51              RCSRGRKRYMYYYYMKRRYYRGCGGGAMRMKMYRRYRYRRKGYRYYYMKRRRGRYKYTAY
                  . . ***************.  .. ******    **** ********** **.

50              GRRKKYRKYYWCGKRYGRRKYAWGMRYRRAATTWYRTYSMWR
51              ARRKKCRKYYWYKKRYKGGKYTWRMRYRRCGGWWYRKTGCTA
                .**** *****  ***   **:* *****..  ***. .
```
